# Perturbations shift the composition of bacterial DNA carried by virus-like particles in the murine gut microbiome

**DOI:** 10.64898/2026.07.08.737213

**Authors:** Jessie L. Maier, Benjamin Callahan, Breck A. Duerkop, Manuel Kleiner

## Abstract

Horizontal gene transfer (HGT) is a driving force in microbial evolution that allows community members to rapidly evolve to cope with environmental stressors and competition. Despite the importance of HGT for the generation of genetic diversity, little is known about the specific mechanisms or dynamics of transfer in complex communities. Transductomics is a sequencing based technique which identifies potential HGT by bacteriophages (transduction) through sequencing of the transductome - the DNA carried by bacteriophages and other virus-like particles in a sample. We analyzed the murine gut transductome before and after perturbations with antibiotics and *Clostridioides difficile* infection (CDI). We found that several bacterial families - the Oscillospiraceae, Butyricoccaceae, and Turicibactericeae - disproportionally contributed to the transductome. Some families, like the Butyricicoccaceae, were frequent transducers in both the baseline and perturbed murine gut microbiome while other taxa displayed condition-specific transduction indicating that there may be specific transducing subpopulations or regulatory mechanisms controlling transduction frequency. Additionally, we found a diversity of highly abundant and enriched mobile genetic elements (MGEs) in the transductome including plasmids, integrative conjugative elements, phage satellites and transposons. The detection of MGEs containing conjugative elements suggest that some MGEs may spread through both transduction and conjugation. Overall, our work reveals a complex network of gene exchange occurring through transduction in the gut microbiome.

## Introduction

The gut microbiome is a critical component of human and animal health^1,2^. A major unexplored aspect of the gut microbiome is the gene exchange network formed via horizontal gene transfer (HGT). HGT allows for the rapid dissemination of genes associated with microbial fitness through transduction, transformation and conjugation transfer mechanisms^3–5^. Many transferred genes provide a fitness benefit to the microorganism but also impact human or animal health by encoding, for example, antibiotic resistance, toxins, biofilm formation^3,6,7^, and adaptations to environmental disturbances^8,9^. HGT is a pivotal driver of microbial evolution, however, most of our knowledge regarding HGT is based on historical data. Most HGT events are inferred from signatures of past HGT in microbial genomes and methods that detect active HGT are currently lacking. Observing ongoing HGT would provide insights into the routes by which microorganisms in communities adapt to change and acquire new traits, however few untargeted methods exist to detect HGT in complex microbial communities, and none, to our knowledge, link HGT events to the specific mechanism of transfer.

The recently developed transductomics approach is a sequencing based method that can detect potential HGT events associated with transduction by detecting bacterial DNA actively carried by virus-like particles (VLPs)^10^. VLPs include phage and other virus-like entities like gene transfer agents (GTAs) and vesicles capable of carrying nucleic acid^11^. For transductomics analyses, sequencing reads from a sample’s ultra-purified VLP-fraction are mapped to bacterial contigs assembled from the same sample’s whole-community (WC) metagenome thus creating read coverage patterns that can both delineate the boundaries of VLP-packaged DNA and indicate the specific transduction mechanism used (i.e. specialized, generalized, lateral, GTA-mediated, etc.). While transductomics can not determine if transduced DNA is ultimately delivered to a new host cell, the method’s advantage lies in its ability to detect ongoing rather than historical transduction. Currently, our understanding of the microbial species involved in transduction, transduction frequencies, and transduction triggers in microbiomes is very limited therefore restricting our overall understanding of the role of transduction in microbiome evolution. We recently automated the transductomics analysis with the TrIdent R package^10,12^ which allows us to address these fundamental questions related to the role of transduction in microbiomes by enabling the detection of VLP-packaged DNA in larger sample numbers.

We used transductomics and the TrIdent R package to explore the murine gut transductome (i.e. the bacterial DNA carried by VLPs) under baseline and perturbed conditions^10,12^. The detection of diverse MGEs in the VLP-fractions of microbiome samples suggests that they are being transduced, however the exact boundaries, gene content, bacterial hosts, and transduction mechanisms of the MGEs is often unknown^13–16^. Additionally, abiotic and biotic perturbations of microbial environments can lead to induction of prophage and other MGEs like phage satellites, plasmids, and GTAs leading to their increased abundance in the VLP-fractions^17–21^. The gut microbiome contains a diversity of MGEs^22–24^ as well as large numbers of phages and VLPs^25,26^ which provide ample opportunities for transduction. Furthermore, the environment of the gut microbiome can rapidly fluctuate such as during periods of inflammation or antibiotics exposure^17,21^. Since gastrointestinal infection rate remains high in the United States^27^ and global antibiotic usage continues to increase^14^, it is important to understand how HGT in the gut microbiome responds to inflammatory perturbations. Our goal was to better understand the types, abundances and microbial community members involved with transduction under baseline and perturbed gut microbiome conditions. Additionally, we were interested to see if there was clear evidence for MGE carriage by VLPs.

We analyzed changes to the murine gut transductome after two different perturbations - antibiotics and *Clostridioides difficile* infection (CDI). We identified an abundance of coverage patterns associated with VLP-packaged DNA, many of which were associated with MGEs like conjugative plasmids, integrative conjugative elements (ICEs), phage satellites, and transposons that were both unique and common amongst bacterial taxa in the baseline and perturbed transductome conditions. In the context of this study, ‘VLP-packaged DNA’ is associated with VLP-fraction read coverage of bacterial contigs and indicates that bacterial DNA is carried in VLPs. However, complete transduction includes not only the packaging of bacterial DNA into a VLP, but also the delivery and utilization of that DNA in a new host, thus VLP-packaged DNA indicates only that the first steps of transduction were completed. Many MGEs were differentially enriched in the transductomes between conditions indicating that their packaging into VLPs is influenced by environmental factors. Finally, we found that there were several bacterial families including the Oscillospiraceae, Butyricicoccaceae, and Turicibactericeae among others that were enriched in the baseline and perturbed transductomes suggesting they are frequent sources for transducing VLPs.

## Results

### Specific bacterial families are enriched in the transductome relative to their abundances in the whole community metagenomes

To explore the effects of strong perturbations to the intestinal microbiota on transduction, we designed an experiment that followed a well-established disease model in which mice with a conventional microbiota are treated with antibiotics prior to challenge with *Clostridioides difficile*^29^. We generated transductomics datasets consisting of paired WC and VLP metagenomes from fecal pellets collected prior to (Pre-ABX) and post administration of the antibiotic cefoperazone (Post-ABX), and on day four of a *C. difficile* infection (CDI) when toxin mediated inflammation is severe^30^ (Fig. 1A). We validated the presence and absence of both CDI and inflammation in Pre-ABX, Post-ABX and CDI samples by quantifying whole-community (WC) reads that mapped to the *C. difficile* strain 630 and host mouse reference genomes, respectively (Fig. S1, Fig. S2)^31^. There was an average of 2.47% of WC reads that mapped unambiguously to the *C. difficile* 630 genome in the CDI condition compared to 0% in the Pre-ABX and Post-ABX conditions. There was an average of 30.6% and 24.7% of WC reads that mapped to the mouse genome in the Post-ABX and CDI conditions, respectively, compared to an average of 0.58% in the Pre-ABX condition. The elevated shedding of host DNA in the feces is expected to accompany intestinal inflammation and drive up the fraction of host reads in the Post-ABX and CDI conditions^41,42^.

**Figure 1.**
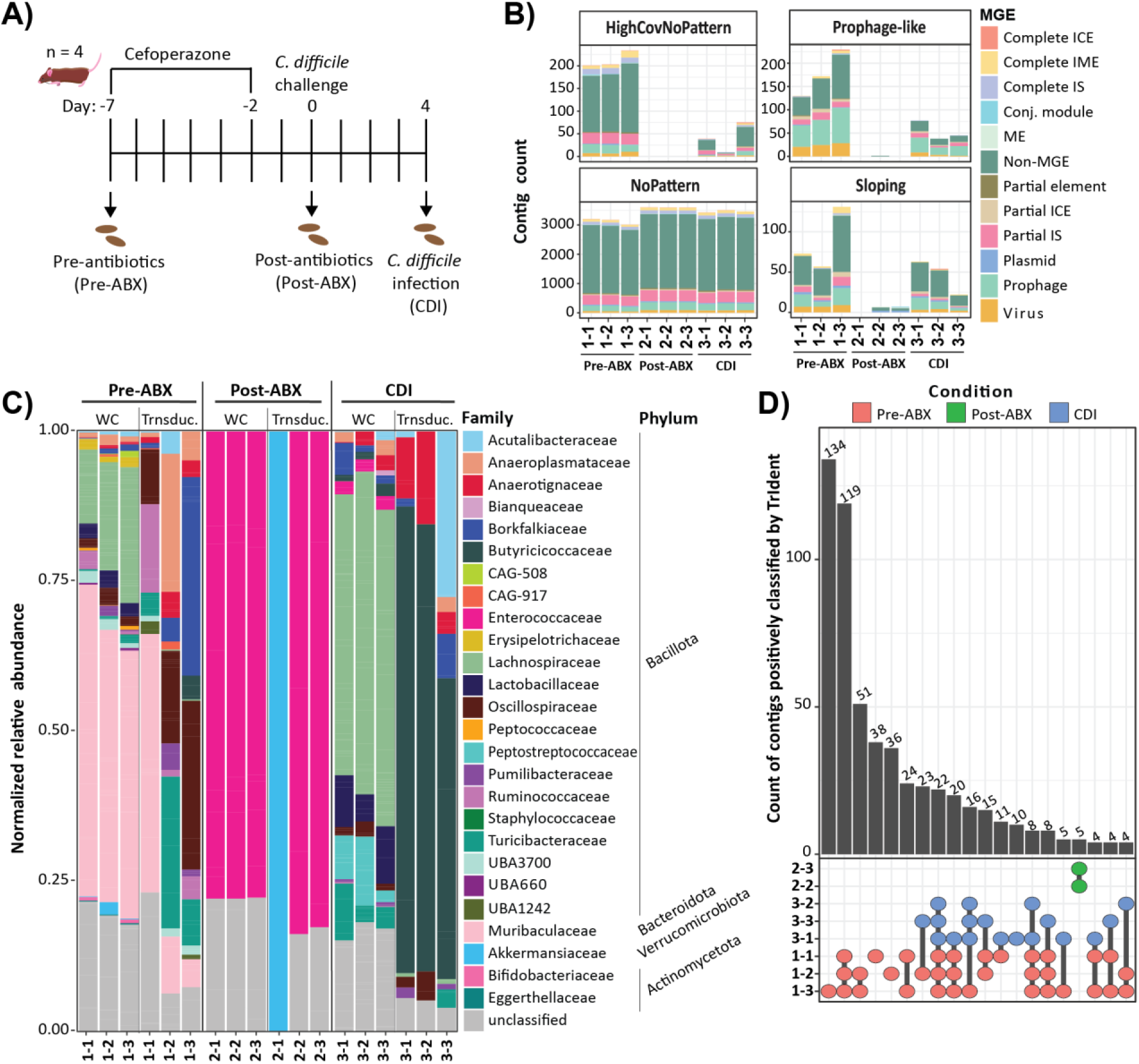
Gut microbiota and transductome compositions pre- and post- perturbation. Sample names start by specifying the sampling condition (1=Pre-ABX, 2=Post-ABX, and 3=CDI) and then specify the replicate (mouse) number identity. A) Sampling timeline for mouse model of primary CDI. The Pre-ABX fecal pellets were collected from all four mouse replicates (n=4) on day -7. The mice were treated with Cefoperazone antibiotics during days -7 to -2. After a two day wash out period, the Post-ABX fecal pellets were collected prior to infecting the mice with *Clostridioides difficile*. The CDI fecal pellets were collected on day 4 of the infection. B) Count of TrIdent-classified contigs in Pre-ABX, Post-ABX and CDI samples with associated mobile genetic element (MGE) classifications. ‘No pattern’ classification counts include contigs which had been filtered out prior to pattern-matching for low VLP-fraction read coverage. There were no positive TrIdent classifications in sample 2-1. C) The relative abundances of bacterial families in the whole-community (WC) metagenomes compared to the relative abundances of bacterial families in the transductomes (the bacterial genetic material carried by VLPs) after viral and prophage-containing contigs were removed. Bacterial families in the WC and transductome were quantified by mapping WC and VLP-fraction reads from each sample to the taxonomically-characterized non-redundant contig set and dividing the number of mapped reads by the number of trimmed and decontaminated reads in each respective sample (Methods). D) The intersection of contigs positively classified (‘Prophage-like’, ‘Sloping’, or ‘HighCovNoPattern) by TrIdent in each of the replicates and sampling conditions after viral and prophage-containing contigs were removed. Positive TrIdent classifications are contigs that TrIdent identified as having VLP-fraction coverage indicative of DNA packaging into VLPs.

In order to directly compare transduction across replicates and conditions, we combined the individual WC assemblies from each replicate and removed redundant contigs to generate a non-redundant contig set which served as the common read-mapping reference for all samples. We removed viral and prophage-containing contigs from the non-redundant contig set prior to comparing the taxonomic composition of the WC metagenomes and the transductomes so as to focus our quantification on bacterial DNA. At the phylum level, the Pre-ABX WCs were composed of an average of 32% Bacillota and 47% Bacteroidota which is expected for male C57BL/6J mice gut microbiota^33,34^. The Bacteroidota were completely lost and the Bacillota expanded to nearly 100% abundance in the Post-ABX and CDI conditions which is an indicator of inflammation in the murine gut^30,35^ (Fig. 1C). While richness and Shannon diversity did not significantly decrease between Pre-ABX and CDI WCs, both richness and Shannon diversity were significantly lower in the Post-ABX WCs (Fig. S3, Fig. S4). Additionally, the WC composition was very distinct between sampling conditions based on the Bray-Curtis community dissimilarity metric (Fig. S5).

The taxonomic compositions of the Pre-ABX and Post-ABX transductomes were clearly distinct from each other and their corresponding WC compositions (Fig. S6). While the taxonomic compositions of the CDI transductomes were distinct from the Pre-ABX and Post-ABX transductomes, they were very similar to the CDI WCs (Fig. S6). Additionally, while the Shannon diversity and richness did not change between the Pre-ABX and CDI WCs (Fig. S3, Fig. S4), the Shannon diversity and richness of the CDI transductomes were significantly lower than the Pre-ABX transductomes (Fig. S7, Fig. S8). The Pre-ABX transductomes were composed largely of the Muribaculaceae (avg. 19%), Oscillospiraceae (avg. 18%), Borkfalkiaceae (avg. 13%), Turicibactericeae (avg. 12%), and Anaeroplasmataceae (avg. 9.5%) bacterial families, whereas the Post-ABX transductomes were composed primarily of Enterococcaceae (avg. 55%) and the CDI transductomes were dominated by the Butyricicoccaceae (avg. 67%) followed by the Anaerotignaceae (avg. 10%) and Acutalibactericeae (avg. 10%) (Fig. 1C). There were large enrichments of several bacterial taxa in the transductomes with respect to their relative abundance in the WCs. In the Pre-ABX transductomes, the Anaerotignaceae, Butyricicoccaceae, Borkfalciceae, Oscillospiraceae, Ruminococcaceae, and Turicibactericeae families were consistently enriched across replicates whereas in the CDI transductomes, the Butyricicoccaceae was the only family consistently enriched (Fig. S9). The Post-ABX transductome was composed primarily of Enterococcaceae, matching the WC composition in that condition (Fig. 1C), however, it was not enriched relative to the WC (Fig. S9).

### VLP-packaged DNA in the Pre-ABX and CDI transductomes was often associated with mobile genetic elements

We used TrIdent to identify and classify read coverage patterns associated with VLP-packaged DNA from transductomics datasets generated from each sample. Positive TrIdent classifications (i.e., ‘Prophage-like’, ‘Sloping’ and ‘HighCovNoPattern’) are associated with VLP-fraction read coverage patterns indicative of different mechanisms of DNA packaging into VLPs (Fig. S10). In contrast, negative TrIdent classifications (i.e. ‘NoPattern’) are similar to HighCovNoPattern classifications albeit the VLP-fraction coverage is not enriched relative to the WC (Fig. S10, Suppl. Text). Samples contained between 7 (1%) and 594 (14%) contigs with positive TrIdent classifications (Fig. 1C). We screened the non-redundant contigs for mobile genetic elements (MGEs) including plasmids, prophages, viruses, ICEs and integrative mobile elements (IMEs), and insertion sequences (ISs) (Methods, Fig. 1B) and found there was a significant enrichment of contigs containing MGEs amongst the positive TrIdent classifications for Pre-ABX and CDI samples (Fig. S11). Prophage-containing and viral contigs made up an average of 28% of the positive classifications and 21% of negative classifications in the Pre-ABX and CDI conditions (Fig. 1B). As expected, prophages and viruses are very abundant in the VLP-fraction and are often detected by TrIdent (Fig. 1B), however these contigs (identified with geNomad) were excluded from the remainder of the analysis to focus on coverage patterns associated with VLP packaging of bacterial DNA.

We compared the positively classified contigs in each condition and found that there were 467 contigs unique to the Pre-ABX condition, whereas there were only 5 and 10 contigs unique to the Post-ABX and CDI conditions, respectively (Fig. 1D). There were few positive classifications in the Post-ABX transductome. This may be driven by the very low bacterial biomass in this condition, which is reflected in the small proportion of reads in the Post-ABX transductomes that mapped to the non-redundant contig set which represents only bacterial DNA (Fig. S12). The total number of Post-ABX VLP-fraction reads was similar to the Pre-ABX and CDI VLP-fractions (Fig. S12), suggesting that the Post-ABX VLP-fraction reads were viral or host DNA, or at least originated from unassembled genome regions not contained within the non-redundant contig set.

### Bacterial families associated with the most abundant and enriched VLP-packaged DNA differs between the Pre-ABX and CDI conditions

The positive TrIdent classifications that contributed the most DNA to the transductomes (the most abundant classifications) and the positive classifications that were the most enriched in the transductomes were of interest due to their potential impact on the microbiome as a whole. We used the maximum TrIdent pattern-match value as a proxy for abundance of positively classified VLP-fraction coverage patterns. To determine the enrichment of positive classifications in the transductome in respect to the WC, we developed a score termed the ‘relative transduction enrichment value’ (RTEV) which takes into account both WC and VLP-fraction coverage values (Methods). There was considerable but not complete overlap between the most abundant and enriched positive classifications (bolded contig names in Fig. 2A and Fig. 2B). The most abundant positive classifications in the Pre-ABX condition were associated with the Ruminococcaceae, Anaeroplasmataceae, Oscillospiraceae and Muribaculaceae families whereas the most enriched positive classifications were associated with the Butyricicoccaceae, Oscillospiraceae, Anaeroplasmataceae, and Ruminococcaceae. The most abundant positive classification in the CDI condition were associated with the Lachnospiraceae, Butyricicoccaceae and Pumilibactericeae families while the most enriched were associated with the Butyricicoccaceae, Oscillospiraceae, Anearotignaceae, and Pumilibactericeae (Fig. 2A, Fig. 2B). Next, we were interested in positive classifications that showed the biggest change in enrichment between Pre-ABX and CDI conditions (Fig. 2C). The Lachnospiraceae, CAG-917 and Oscillospiraceae positive classifications were all more enriched in the Pre-ABX transductome with the exception of one Oscillospiraceae classification. The Anaerotignaceae and Pumilibactericeae positive classifications were more enriched in the CDI transductome. The Butyricicoccaceae positive classifications showed mixed enrichments between conditions. We analyzed several highly abundant and/or enriched positively classified contigs from different bacterial families or MGEs in-depth below. We included the positively classified Turicibactericeae contigs in the following analysis of enriched and abundant positive classifications as they made up a relatively large proportion of the Pre-ABX transductome (Fig. 1C).

**Figure 2.**
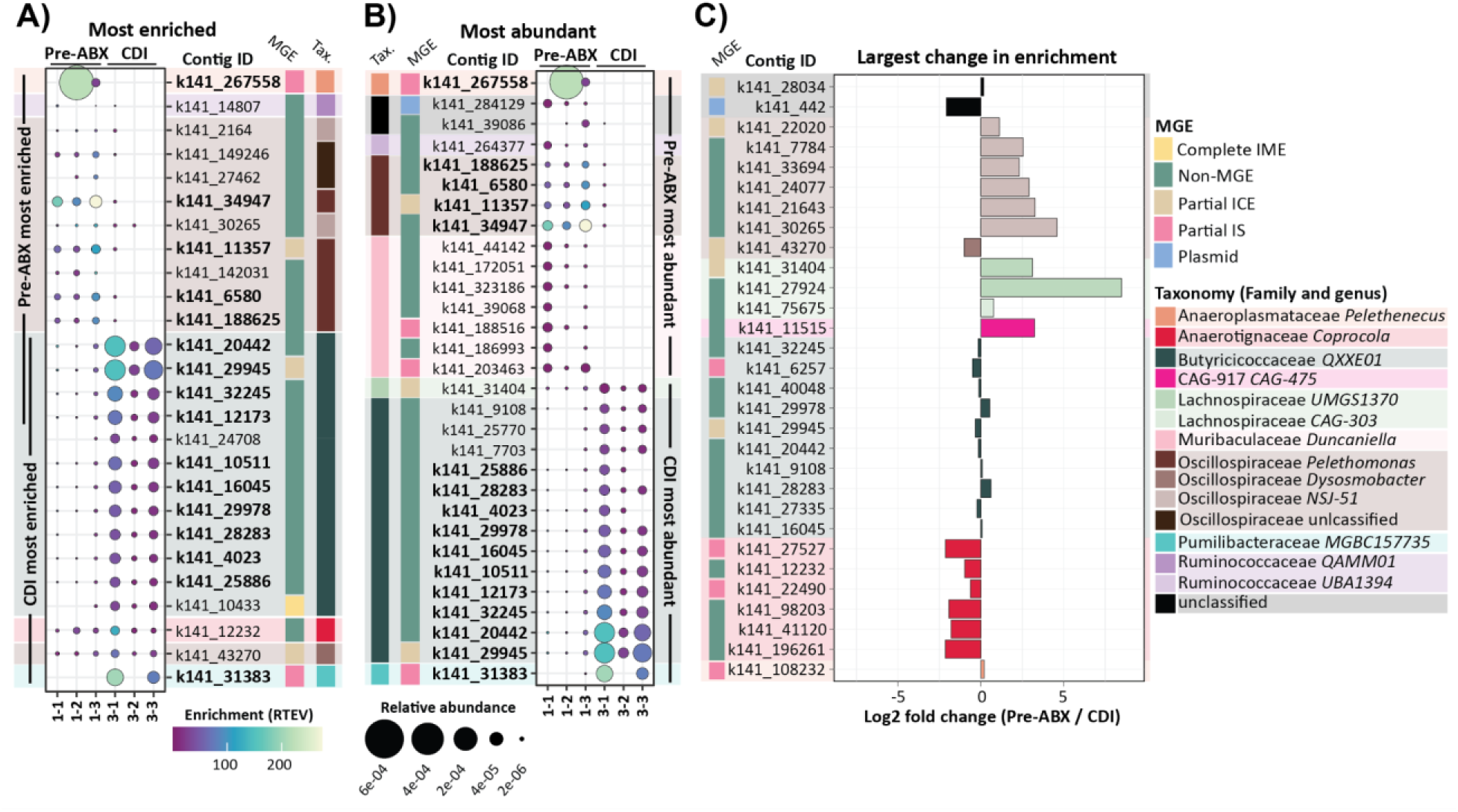
The most abundant and enriched positive TrIdent classifications in the Pre-ABX and CDI transductomes. Positive TrIdent classifications are contigs that TrIdent identified as having VLP-fraction coverage indicative of DNA packaging into VLPs via different packaging mechanisms. The sample naming conventions are as follows- the first value specifies the sampling day (1=Pre-ABX, and 3=CDI) and the second value specifies the replicate (mouse) number. The taxonomy and MGE (mobile genetic element) color palettes are shared between plots A, B, and C. ICE = Integrative conjugative element, IME= Integrative mobile element, IS= Insertion sequence. A) The 15 most enriched positive TrIdent classifications per condition. Enrichment was determined using the relative transduction enrichment ratio (RTEV) which was calculated by dividing the relative maximum pattern-match value provided by TrIdent by the relative whole-community median read coverage value. B) The 15 most abundant positive TrIdent classifications per condition. Abundance was calculated using the relative maximum pattern-match value provided by TrIdent. Relative values were calculated using the number of trimmed and decontaminated sequencing reads for the respective WC or VLP-fraction. Bolded contig IDs indicate that the associated contig was positively classified in both the most abundant and enriched plots. C) The top 30 log2 fold differences were determined by first filtering for contigs with positive classifications in at least two replicates in both Pre-ABX and CDI conditions, then averaging the RTEVs and taking the log2 of Pre-ABX avg. RTEV/CDI avg. RTEV.

### Specific MAGs were preferentially packaged into VLPs

#### *Turicibacter* sp. found in Pre-ABX transductome but not in CDI

Every Turicibactericeae contig (all of which could be further assigned to the *Turicibacter* genus) was positively classified by TrIdent in all of the Pre-ABX replicates, while there were no positive classifications in the CDI transductomes. The *Turicibacter* positive classifications were characterized by uniform (i.e., full contig coverage without major inconsistencies/abnormalities) contig coverage in both the WC and the VLP-fraction metagenomes of the Pre-ABX replicates (Suppl. files 1-8), however the coverage was higher in the VLP-fraction relative to the WC. Many (12 out of 30) of the *Turicibacter* contigs contained partial or complete transposons, some of which were associated with sharp spikes (3-6x the baseline coverage) in read coverage at the transposase gene in both the WC and VLP-fraction metagenomes (Fig. 3A, Suppl. files 1-8).

**Figure 3.**
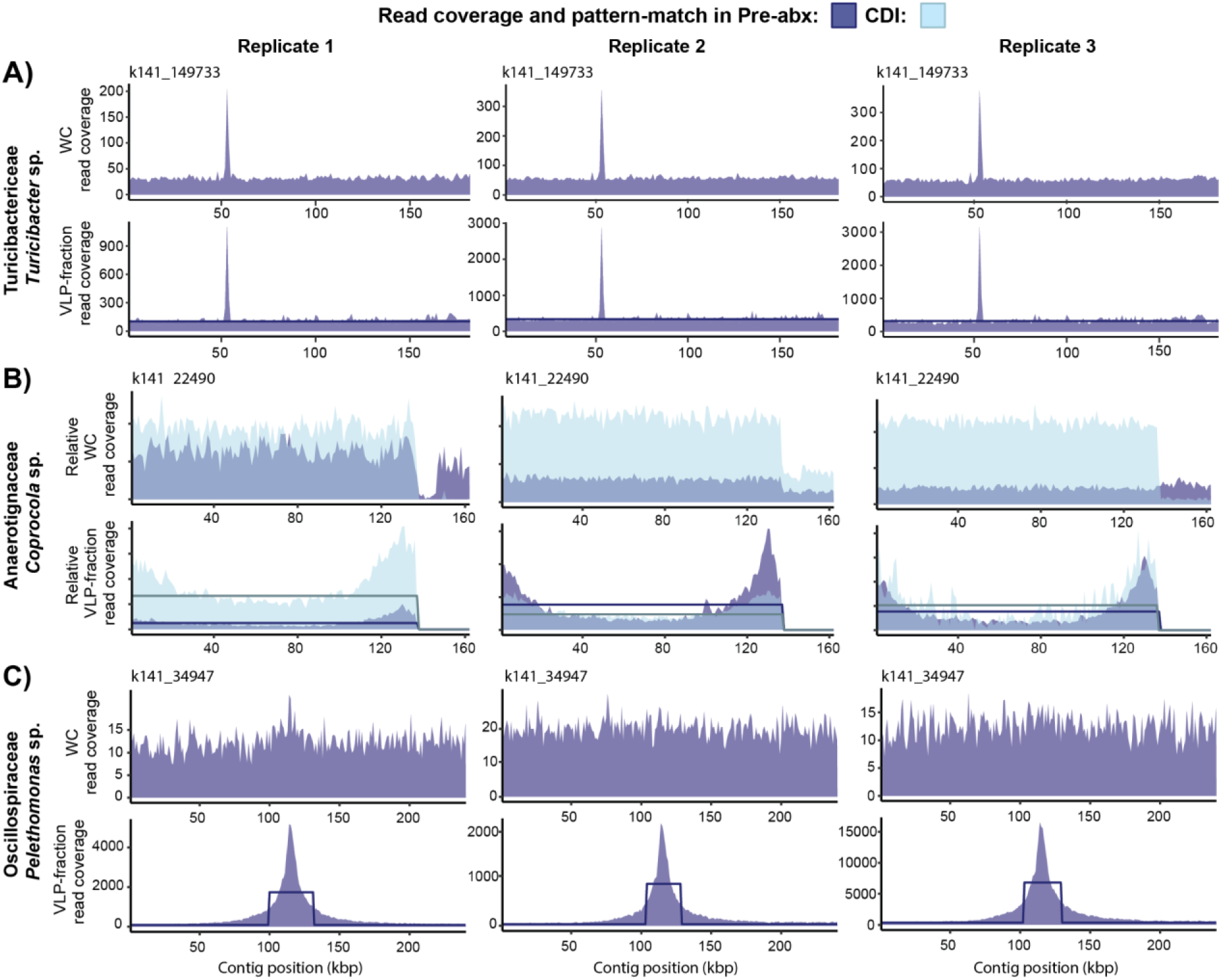
Read coverage patterns associated with contigs from organisms that had large amounts of their DNA carried by VLPs. The read coverage from the pre-antibiotic (Pre-ABX) condition is displayed in dark blue while the read coverage from the corresponding replicate in the *C. difficile* infection (CDI) condition is overlaid in light blue. The sample’s associated TrIdent pattern-match used for classification is overlaid as a dark (Pre-ABX) or light (CDI) blue line. A) The relative read coverage for contig k141_22490’s (originates from CDI replicate 2 WC assembly) whole-community (WC) and viral-like particle (VLP) fractions was plotted to allow the coverage between Pre-ABX and CDI conditions to be visualized on the same scale. Relative coverages were calculated by dividing the coverage values by the number of trimmed and decontaminated sequencing reads obtained for the respective WC or VLP-fraction sample. B) Contig k141_34947 (originates from CDI replicate 1 WC assembly). C) Contig k141_149733 (originates from Pre-ABX replicate 4 WC assembly).

#### Strong enrichment of VLP-packaged DNA associated with specific bacterial families in the Pre-ABX and CDI conditions

The Anaerotignaceae *Coprocola* positive classifications were strongly enriched during the CDI condition in comparison to the Pre-ABX condition (Fig. 2C) despite the fact that the general VLP-fraction coverage patterns and WC contig coverage was consistent between conditions. Each of the 25 *Coprocola* contigs in the non-redundant contig set were positively classified by TrIdent in at least one of the Pre-ABX or CDI replicates and most have VLP-fraction coverage patterns characterized by uniform or sloping (i.e., full contig coverage with a visible slope) contig coverage (Suppl. Files 1-3, 6-8).

We inspected the Oscillospiraceae *NSJ-51* positive classifications with substantially larger enrichments in Pre-ABX compared to the CDI condition (Fig. 2C) and found that the WC coverage of the *NSJ-51* contigs in the Pre-ABX condition was very low (∼1-3x) in comparison to the VLP-fractions (∼20-30x). Almost all positively classified *NSJ-51* contigs were characterized by uniform or sloping VLP-fraction coverage, however there were no distinctive patterns that could be used to hypothesize the transduction mechanism at play (Suppl. Files 1-3 and 6-8).

All of the highly abundant and enriched Butyricicoccaceae positive classifications were associated with the *QXXE01* genus. Butyricicoccaceae positive classifications were highly abundant in the CDI transductome however they were enriched in the transductomes of both the Pre-ABX and CDI conditions relative to the WCs (Fig. 2A-C) suggesting that despite the much lower relative abundance of *QXXE01* in Pre-ABX WCs (0.03%) compared to CDI WCs (0.9%), the associated packaging frequency did not change (Fig. 2C). All of the *QXXE01* contigs in the non-redundant contig set were positively classified by TrIdent in at least one of the Pre-ABX or CDI replicates and most contigs had read coverage patterns characterized by uniform or sloping VLP-fraction contig coverage (Supp files 1-8). There were no distinctive patterns that clearly suggested a specific transduction mechanism.

### Mobile genetic elements (MGEs) detected in the transductomes

#### GTAs may be responsible for some of the most abundant and enriched VLP-packaged DNA

Several of the most enriched or abundant positive classifications had coverage patterns similar to those produced by known GTAs which motivated us to analyze all contigs in the respective bacterial host metagenome assembled genome (MAG) for the presence of GTAs. GTAs are non-infective phage-like particles encoded and produced by bacteria as a means of gene transfer amongst GTA-producing bacterial populations. GTA induction leads to cell lysis and therefore GTA producers are highly regulated and only induce GTAs in a small portion of the population^41–43^. While GTAs package the entire bacterial genome, some regions are preferentially packaged thus generating peaks in coverage at those sites^44–46^. GTAs are typically encoded in several gene clusters with distinct functions spread throughout the bacterial genome and can not preferentially replicate or mobilize themselves^47–49^. Additionally, GTAs originate from functional prophage^50–52^ and can therefore be associated with cryptic or remnant phage genes^53^. Since there is no standardized bioinformatics method for GTA detection in genomes, we based our detection strategy on these characteristics. We derived a set of flexible criteria for detecting GTA gene cluster candidates in MAGs that were - 1) classified as a prophage by geNomad^54^ and 2) did not contain gene annotations in the ‘integration and excision’ and ‘DNA, RNA and nucleotide metabolism’ categories provided by Pharokka^55^. While, theoretically, GTAs should not contain genes for either DNA replication or integration/excision, we only disqualified predicted prophage from consideration as GTA if they contained genes associated with both functions to account for the possibility of remnant phage genes^53^.

The VLP-fraction coverage pattern on Anaerotignaceae *Coprocola* contig k141_22490 was more enriched in CDI relative to the Pre-ABX condition and resembled the sloping peaks and valleys of coverage produced by generalized transduction^10,56^ and some GTAs^44,45^ (Fig. 3B). To explore whether a *Coprocola* species in our samples was a GTA producer, we screened all contigs from the *Coprocola* MAG containing contig k141_22490 for prophage. Contig k141_7543, the only *Coprocola* contig with predicted prophages, had two prophages, both of which fit the described GTA criteria (Fig. S13). Contig k141_7543’s VLP-fraction had a large and distinct sloping read coverage pattern that spanned the contig with no major differences in coverage at either of the GTA gene cluster candidate locations (Fig. S14). The first 3.6 kbp GTA gene cluster candidate contained a holin and endolysin while the second 14 kbp candidate contained a large and small terminase in addition to a variety of phage structural genes (Fig. S15). Both the gene content and organization of these two clusters are characteristic of GTAs^57^.

The VLP-fraction coverage pattern on Oscillospiraceae *Pelethomonas* contig k141_34947 was highly enriched and abundant in both the Pre-ABX and CDI transductomes. Contig k141_34947 contained a large coverage peak (∼150 kbp, Fig. 3C), similar to the broad coverage peaks produced by *Bartonella* GTAs (BaGTAs)^46,53^. During BaGTA induction, a remnant prophage’s replication machinery is activated causing run-off replication (ROR) of the bacterial genome surrounding the prophage origin of replication. The amplified DNA is cleaved and packed into BaGTA particles along with the rest of the genome^58^. Based on the similarity in coverage pattern to BaGTA, we speculated that contig k141_34947 may also have remnant prophage DNA replication genes at the peak location and ran it through Pharokka^55^. We found a phage derived DNA polymerase, DNA helicase, DnaC-like helicase loader and replication initiation protein directly at the peak’s apex (Fig. S16). We inspected the coverage patterns of other positively classified contigs in the same MAG as contig k141_34947 and while most contigs had uniform VLP-fraction read coverage, one contig, k141_11357, had a broad coverage peak (>100 kbp) that was ∼2x less abundant than the coverage peak on contig k141_34947. There were two phage-derived DNA topoisomerases located near the peak’s apex, however there was no helicase in close proximity (Fig. S17). We searched for GTA candidates in the *Pelethomonas* contigs found in the same MAG as contigs k141_11357 and k141_34947 using the same search strategy used for the *Coporocola* contigs. While we did not find any strong GTA gene cluster candidates based on our criteria, we did find an unknown MGE whose gene content resembles a capsid-forming phage satellite^59^ (Fig. S18).

The positively classified Butyricicoccaceae *QXXE01* contigs, which were part of three MAGs, were highly abundant and enriched in both Pre-ABX and CDI transductomes and were largely characterized by contigs with uniform or sloping coverage. There were no *QXXE01* contigs with distinct peaks and valleys of read coverage like those observed in the *Coprocola* or *Pelethomonas* contigs (Fig 3B-C) and we therefore could not limit our search for GTAs to a single *QXXE01* MAG. We searched for GTA gene cluster candidates in all Butyricicoccaceae *QXXE01* contigs and found three that met our criteria (Fig. S19). We excluded two of these three candidates because one had a VLP-fraction coverage pattern indicative of prophage induction and the other was located on the edge of its contig and it was therefore unclear if the associated sequence was complete. The final 12.7 kbp cluster was fully contained on contig k141_34042, and encoded a transcriptional activator, small and large terminases, several capsid associated structural proteins, and a holin and endolysin, and did not have any difference in read coverage at its contig location (Fig. S20). The gene content of this candidate is similar to capsid-forming phage satellites^59^.

#### Highly enriched plasmid in the transductome during CDI

Contig k141_442 was classified as a plasmid by geNomad and the associated VLP-fraction coverage pattern showed a large increase in enrichment in the CDI transductomes (Fig. 2C). The read coverage pattern of the contig differed between the Pre-ABX and CDI conditions with the Pre-ABX showing a large decrease in coverage in the core region of the contig with high coverage at the edges compared to uniform contig coverage in CDI (Fig. 4A). The reads mapping to the contig edges in Pre-ABX showed a high number of nucleotide differences when compared to the reference contig which was derived from the CDI condition (Fig. 4A). The contigs edges contained a variety of genes associated with recombination and gene expression such as a helix-turn-helix (DNA binding) and MobA/VirD2-like nuclease (DNA processing) domain-containing proteins and a recombinase family protein, for example (Fig. S21). The core region had far less read mapping variants than the contig edges indicating that while the core region may be lost or recombined, the plasmid is conserved as a whole. The contig encoded all genes needed for conjugation (i.e. a relaxase, coupling protein, and type 4 secretion system^60^) in the core region in addition to two virulence factors- a tetracycline resistance protein and an anti-CRISPR protein identified with DefenseFinder^61–63^. In addition to the density of Pre-ABX read variants at the contig edges, there were also a large number of variants at the tetracycline resistance gene suggesting that while the plasmid’s core remained largely the same between Pre-ABX and CDI conditions, these other regions are more evolutionarily variable.

**Figure 4.**
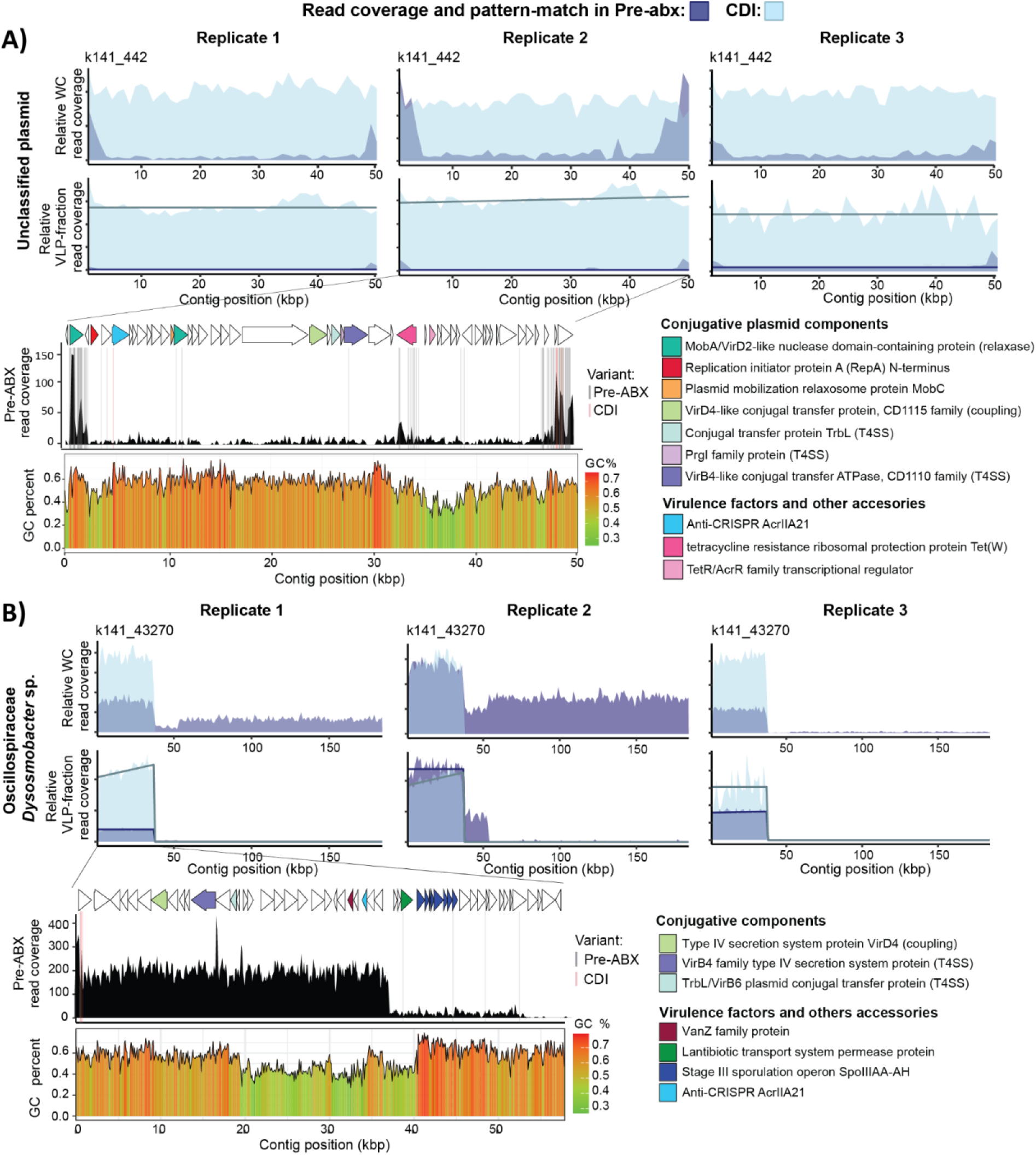
Read coverage patterns associated with packaging of conjugative elements. The read coverage for both the whole-community (WC) and viral-like particle (VLP) fractions were normalized by dividing the coverage values by the number of filtered and decontaminated sequencing reads obtained for the respective sample. The read coverage from the pre-antibiotic (Pre-ABX) condition is displayed in dark blue while the read coverage from the corresponding replicate mouse in the *C. difficile* infection (CDI) condition is overlaid in light blue. Gene annotations are only provided for select open reading frames associated with conjugative components or virulence factors. The sample’s associated TrIdent pattern-match used for classification is overlaid as a dark (Pre-ABX) or light (CDI) blue line. The contig’s GC content, averaged over 100 bp sliding windows, is displayed under the corresponding coverage plot. Read mapping variants (i.e. reads that differ from the reference contig) generated in both Pre-ABX (grey) and CDI (red) conditions are displayed as vertical lines on the coverage plot inset. The coverage plot inset was generated using pileup files with 100 bp windows for better resolution of coverage patterns. A) Contig k141_442 (originates from CDI replicate 3 WC assembly). B) Contig k141_43270 (originates from Pre-ABX replicate 2 WC assembly).

#### A partial Oscillospiraceae *Dysosmobacter* sp. ICE was enriched in the CDI transductome and present in both Pre-ABX and CDI conditions

*Dysosmobacter* contig k141_43270 had VLP-fraction read coverage that was substantially more enriched in the CDI transductome compared to the Pre-ABX transductome (Fig. 2C). The VLP-fraction coverage was associated with a defined ‘block’ of read coverage from 0-36.6 kbp in both the Pre-ABX and CDI conditions; however, there was an additional 16.4 kbp region from 36.6-53 kbp that had a relevant but lower amount of coverage in only the Pre-ABX transductome (Fig. 4B). A short section of the first region (8.9-16.6 kbp) was classified as a partial ICE by ICEscreen^64^ based solely on the start and stop locations of type 4 secretion system (T4SS) proteins VirD4 and VirB4, respectively, however based on the coverage pattern, it is likely that the ICE extends further. There were multiple virulence factors and defense-associated genes encoded on both regions including VanZ family protein (antibiotic resistance^65^) and an anti-CRISPR protein in the higher coverage region and lantibiotic resistance protein in the lower coverage region. The GC content of contig k141_43270 averaged around 59%, however there was considerable GC heterogeneity and a lower GC average (45%) within the region from 20-40 kbp (Fig. 4B). While contig k141_43270 was classified as an Oscillospiraceae *Dysosmobacter* by CAT^66^, many of the gene annotations associated with the elevated coverage region were either classified as Oscillospiraceae *Pelethomonas* or could not be classified past the Oscillospiraceae class level (Fig. S22). The lack of WC contig coverage outside the high coverage region in the CDI samples could be consistent with the partial ICE and associated DNA originating from a different bacterial host in the CDI samples. We searched the sequence associated with the region from 0-53 kbp against the combined non-filtered (redundant) contig set using BLASTn^67^ to determine if and where it was assembled in the CDI samples. We found that the region from 0-33.6 kbp (69% query overlap) was assembled in all of the CDI samples (Fig. S23) and the associated contigs were classified by CAT^66^ to the genus level as Oscillospiraceae *Pelethomonas*. Based on the large query overlaps and presence in two different Oscillospiraceae genera, the partial ICE appears to be present in two genera and thus have a wide host range.

#### An unknown MGE in Anaeroplasmataceae *Pelethenecus* sp. was highly abundant in Pre-ABX transductomes and showed signs of recent mobility within the population

Anaeroplasmataceae *Pelethenecus* contig k141_267558 had a ∼12 kbp ‘block’ of VLP-fraction read coverage from 31.9-44.2 kbp that was highly abundant and enriched in the Pre-ABX transductomes of replicates 2 and 3 (Fig. 5A) suggesting that this genome region harbors an MGE that is mobilized into VLPs. The WC read coverage of this MGE was highly variable between replicates with there being no read coverage in replicate 1, elevated coverage (∼2.5x the average baseline) in replicate 2 and slightly depressed coverage in replicate 3 (∼0.5x the average baseline). The variable coverage is consistent with strain-level variation in the presence or absence of this MGE in the bacterial host genome between replicates. This notion was further supported by WC reads in replicates 2 and 3 with inserts that spanned the entire MGE, consistent with deletion of the MGE, and read pair orientations (Forward-reverse and reverse-forward) which are consistent with translocation or duplication of the MGE in part of the host population (Fig. S24). The MGE encodes for XerD, which is a tyrosine recombinase^68^ capable of site-specific insertion of MGEs at tRNA genes^69^ and, accordingly, this MGE was inserted into the 3’ region of the Arg tRNA gene. Additionally, we found 25 bp direct repeats (DRs) (5’ GTTCGAATCCTCTAGGGTGCGCCAT 3’) flanking the MGE which are a hallmark of site-specific recombination and transposition^70^ (Fig. S25). Finally, the MGE had a notably lower GC content (28%) than the rest of the contig (32%) (Fig. 5A). The MGE had relatively few annotated genes of which most were associated with DNA and RNA processing functionalities. Despite the annotation as a toxin, the zona occludens toxin (Zot), is involved in the morphogenesis of filamentous Inoviruses and while it can have a toxigenic domain^71^, the Zot protein sequence in this MGE did not contain the 8 amino acid motif responsible for Zot receptor binding^72^ and is therefore likely non-toxigenic (Fig. S26).

**Figure 5.**
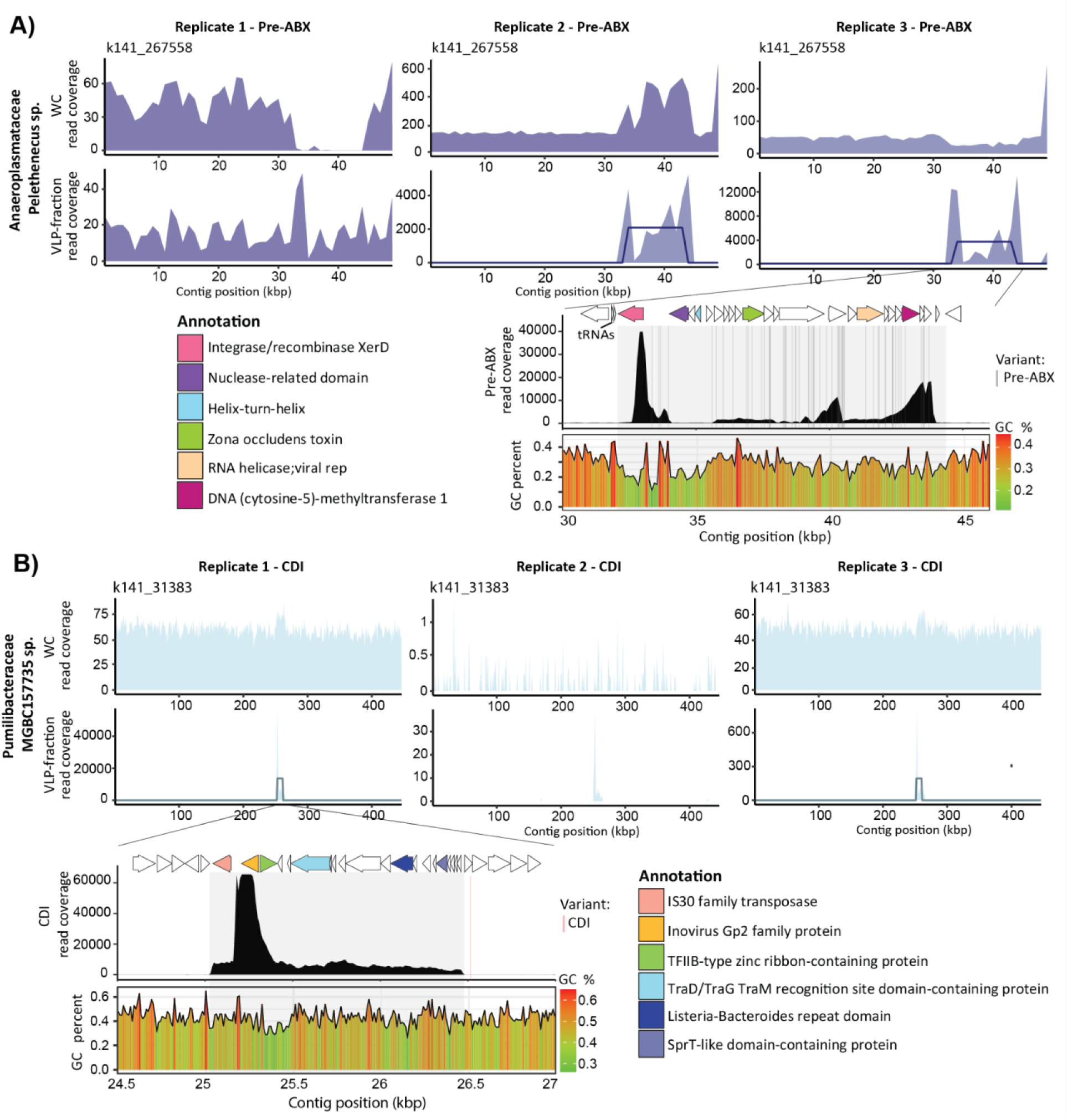
Read coverage patterns associated with packaging of unknown mobile genetic elements. Whole-community (WC) and virus-like particle (VLP)-fraction read coverage plots with overlaid TrIdent pattern-matches (dark blue =Pre-ABX and light blue=CDI) and zoomed-in subsets with associated GC content averaged across 100 bp rolling windows and gene calls with annotations obtained from Mantis and Bakta. Gene annotations are only provided for the ORFs within the grey highlighted area of the plots. ORFs colored white were hypothetical or not assessed. Read mapping variants generated in both Pre-ABX (grey) and CDI (red) conditions are displayed as vertical lines on the coverage plot inset. The coverage plot inset was generated using pileup files with 100 bp windows for better resolution of coverage patterns. A) Contig k141_267558 (originates from Pre-ABX replicate 4 WC assembly). B) Contig k141_31383 (originates from CDI replicate 3 WC assembly).

#### An unknown MGE in Pumilibacteraceae *MGBC157735* sp. was highly abundant in CDI transductomes and was integrated into a bacterial host with variable abundance across replicates

Pumilibacteraceae *MGBC157735* contig k141_31383 had a ∼15 kbp ‘block’ of VLP-fraction read coverage from 250-264.6 kbp that was highly abundant and enriched in CDI replicates 1 and 3 suggesting that this genome region harbors an MGE that is mobilized into VLPs. ISEscan^73^ identified a partial IS30 transposon at the edge of the MGE containing 19 bp imperfect terminal inverted repeats (IRs) (5’ ATAATCGAATATTTCGCGG 3’ (IRL) and 5’ ATAATGAGATAAATCGCGG 3’ (IRR)) with start locations of 250,493 and 251,365 bp, respectively. The ‘partial’ classification was likely due to a divergence in size between the identified IS30 transposon (872 bp) and known IS30 transposons (1-1.7 kbp^74^). While transposases can mobilize neighboring DNA in a composite transposon fashion, to mobilize the entire 15 kbp region, there would need to be an inverted repeat (IR) flanking the MGE’s right border around 264.6 kbp. We scanned contig k141_31383 for a sequence matching either of the transposon’s IRs (IRL (left) and IRR (right)) on both the forward and reverse strands, but did not find any hits close to 264.6 kbp. However, when we relaxed the search parameters and allowed up to 6 mismatches, we found two IRL hits on the forward strand at the read coverage peak’s apex at 252 kbp, and two IRL hits on the reverse strand flanking the peak (Fig. S27). We suspect that they lead to alternative transposase excision sites causing the sharp increase in read coverage at this site. To rule out the possibility that the elevated coverage was the result of ambiguous read mapping, we searched the MGE with BLASTn against the non-filtered (redundant) dataset and found the MGE in only one contig per sample with 100% query overlap and sequence identity indicating that the increased coverage is not caused by the presence of multiple copies of the MGE in the population. Despite not having flanking IS30 IRs, the MGE had flanking 15 bp DRs (5’ GCGTATATGTGGACA 3’) which is a result of site-specific recombination (Fig. S28). Finally, the MGE was found to be inserted into the 3’ region of the YlmC/YmxH family sporulation protein gene rather than a tRNA or other more common housekeeping gene^75^.

## Discussion

While we had designed our experiment with perturbations that we hypothesized would lead to large increases in DNA packaging into VLPs, we found that the transductomes were far more complex than anticipated. In both the baseline (Pre-ABX) and the inflammatory (Post-ABX and CDI) conditions, the transductomes differed vastly from the respective WC metagenomes and were dominated by DNA from specific bacterial taxa (Fig. 1C). The far higher engagement in transduction by specific bacterial taxa opens questions about the associated selective pressures that drive higher mobilization of DNA into VLPs by some bacteria. The transductomes in the antibiotic condition (Post-ABX) consisted entirely of Enterococcaceae contigs which also made up a majority of the highly simplified WC in these samples and contained almost no positive TrIdent classifications. The near absence of transduction is likely due to a lack of bacteria in the WC to serve as sources for transduction. Interestingly, there were a comparable amount of trimmed and decontaminated reads in the Post-ABX VLP-fractions compared to the Pre-ABX and CDI conditions which may indicate high VLP abundance despite the lack of transduction (Fig. S12).

The bacterial taxa we identified as major contributors to the Pre-ABX and CDI transductomes and/or as hosts to active MGEs (i.e. the Oscillospiraceae *Pelethomonas*, Oscillospiraceae *Dysosmobacter,* Turicibactericeae *Turicibacter*, Anaerotignaceae *Coprocola*, Anaeroplasmataceae *Pelethenecus*, Pumilibacteraceae *MGBC157735*, and Butyricicoccaceae *QXXE01*) are largely uncharacterized and the fact that we identify them as potentially major transduction “hubs” makes them important targets for future study. Additionally, the considerable overlap between positively classified contigs identified in Pre-ABX and CDI (Fig. 1D) suggests that certain transduction mechanisms are consistently active in the intestinal microbiome independent of environmental conditions. There were also a number of contigs that were only positively classified in the Pre-ABX or CDI condition; however, it was unclear if the associated DNA packaging mechanisms were not active in a specific condition or if the specific bacterial subpopulations involved were just not present. For example, while there are *Turicibacter* in both the Pre-ABX and CDI WCs, the *Turicibacter* are only enriched in the Pre-ABX transductomes (Fig. S9). It’s possible that the specific *Turicibacter* species or strain involved in transduction was not present in the CDI condition or that the regulatory mechanism associated with *Turicibacter* DNA carriage in VLPs was not active in the CDI condition. Better taxonomic resolution achieved through deeper-sequencing and/or long-read sequencing will help answer this question by determining if the same species or strains capable of transduction are found across conditions.

The transductomes of the mice displayed increased inter-individual heterogeneity compared to the WCs (Fig. S29). This is in line with prior observations that both the viromes and mobilomes of the gut microbiome are individual-specific^76,77^. Since transductomics works on the contig level, we may be capturing blooms of specific bacterial subpopulations, VLPs and MGEs occurring in each individual sample that are not captured at the MAG level. Alternatively, the homogenization produced by coprophagy may occur on a longer time scale than the more rapid responses of the transductome and associated mobilome. Ultimately, our results show that murine gut transductomes are highly diverse and dynamic and may be a source of fast genetic diversification amongst specific bacterial subpopulations. Interestingly, the Pre-ABX transductomes were more diverse than the CDI transductomes despite the unchanging WC Shannon diversities (Fig. S4, Fig. S7). This may indicate that a more stable, less perturbed gut microbiome is associated with a more diverse transductome.

Below, we discuss the systematic extraction and analysis of highly abundant and/or enriched positive TrIdent classifications (Fig. 2). While this analysis revealed the major bacterial contributors to the murine gut transductome, abundances and frequencies of DNA carriage, and several interesting MGEs, a large portion of the transductomes remain unexplored and there are many interesting VLP-fraction read coverage patterns that warrant further study (Suppl. files 1-8).

### Our data shows evidence for uncharacterized mechanisms of bacterial DNA carriage that mobilize large genome regions at high frequencies

Several bacterial families and genres (the Butyricicoccaceae *QXXE01*, Turicibacericeae *Turicibacter*, Oscillospiraceae *NSJ-51* and Muribaculaceae *Duncaniella*) were associated with highly abundant and/or enriched positive classifications characterized by uniform or sloping VLP-fraction contig coverage. Since these large DNA carriage events did not have distinctive coverage patterns like those seen with the Anaerotignaceae *Coprocola* and Oscillospiraceae *Pelethomonas* (Fig. 3A and Fig. 3B), it is difficult to hypothesize about the transduction mechanism that might lead to the mobilization of this DNA in the VLP fraction. The uniform contig coverage associated with these classifications is consistent with generalized and lateral transduction, MV-mediated HGT^36–38^ and some types of GTAs^39,40^ (e.g. see Butyricicoccaceae *QXXE01* contigs in Suppl. Files 1-3 and 6-8). However, there is currently no way to distinguish MV-mediated and phage-mediated “transduction” in complex samples, and it is possible that additional mechanisms for the inclusion of bacterial DNA in VLPs exist that produce similar patterns.

We found evidence that Anaerotignaceae *Coprocola* and Oscillospiraceae *Pelethomonas* are GTA producers. While the *Coporocola* VLP-fraction read coverage indicating potential GTA DNA carriage was present in both the Pre-ABX and CDI conditions, the *Pelethomonas* GTA-mediated DNA carriage was uniquely associated with the Pre-ABX condition. In Anaerotignaceae *Coprocola,* we identified strong evidence for two GTA gene clusters- a ‘core’ cluster which contains a small and large terminase in addition to a variety of structural genes and a cell lysis cluster located on the same contig. Interestingly, based on the sharp discontinuity of the VLP-fraction read coverage and decreased WC read coverage towards the end of contig k141_22490 in addition to several single nucleotide polymorphisms directly at the coverage discontinuity (Fig. 3B, Fig. S30, Fig. S31), there appears to be a non-transducing subpopulation of *Coprocola* with a structural genome variant. It’s possible that structural variation (SV) at this region interrupts a GTA gene cluster or packaging signal that prevents GTA production in a *Coprocola* subpopulation. The evolution of non-GTA producing ‘cheaters’ in a GTA producing population is a common occurrence due to the ability for cheaters to benefit from GTAs without suffering the costs of being a producer^78^.

In Oscillospiraceae *Pelethomonas,* we were unable to find strong evidence for GTA gene clusters using our GTA search criteria, however, we hypothesize that the coverage peak on *Pelethomonas* contig k141_34947 is generated through run-off replication (ROR) and packaging of the amplified DNA into GTA particles, similar to the BaGTA^46,53^. It’s unclear if the second coverage peak on contig k141_11357 is also generated through ROR since there are a lack of phage-derived DNA replication genes in close proximity to the peak, however, the genome region is still being bidirectionally amplified and packaged indicating that there may be some ROR-like mechanism at play^46^. While the identification of the putative GTA gene clusters and associated packaging mechanisms require further investigation, our hypothesis that *Pelethomonas* species are GTA producers is supported by a recent study which found that members of the Oscillospiraceae family were prevalent GTA producers in the gut microbiome^79^.

The criteria we used for GTA gene cluster candidate detection have some overlap with the characteristics of phage satellites, a widespread class of MGEs that hijacks structural components from a helper phage for mobilization. Unlike GTAs, phage satellites can integrate, excise and replicate, but are often misclassified as prophage. We therefore suspect that some of the GTA gene cluster candidates we identified, especially those that are not restricted to one function (i.e. not multilocus), are phage satellites or another form of MGE (Fig. S18, Fig. S19).

### Some MGEs may be spreading through both conjugation and transduction

We inspected the coverage patterns of two highly enriched positive classifications in the CDI condition associated with a partial ICE and plasmid. Both the partial ICE and plasmid encode conjugative elements which leads us to speculate that certain MGEs with the necessary genes for conjugation may be capable of transfer by both transduction and conjugation thereby expanding their host range and overall prevalence. MGEs often share traits or pirate functions of others- for example, ICEs and IMEs inserted in-tandem can form composite MGEs that mobilize together^80,81^, IMEs^82^ and various plasmids^83,84^ use the mating pore made by ICEs, and phage satellites pirate capsids or tails from functional phage^85,86^. While MGEs are generally thought to transfer intercellularly via a defined route of HGT, the crossovers between MGE classes present ample opportunities for MGEs to co-opt each other’s mechanisms of transfer. As is common for MGEs, both the partial ICE and plasmid encoded virulence factors and other accessory genes that likely serve to enhance bacterial fitness. Additionally, both elements showed evidence of SV between conditions which allowed us to identify genes or regions that were important for the elements’ fitness in each condition. For example, the conserved core of the Oscillospirales plasmid on contig k141_442 (Fig. 4A) encodes the conjugative machinery that controls the plasmid’s host range. Since the coverage of this region is decreased in Pre-ABX replicates, we suspect that the bacterial host(s) of the plasmid on contig k141_442 are present at a low abundance in the Pre-ABX condition. Alternatively, the virulence factors encoded on the core region may have given the k141_442 plasmid a greater fitness advantage in CDI compared to Pre-ABX. Interestingly, the presence of the second relaxase in the core region suggests that it may be an independent conjugative plasmid that integrated into an existing mobilizable plasmid^87^. Similarly, we speculate that the ‘accessory’ region associated with the VLP-packaged partial ICE on Oscillospiraceae *Dysosmobacter* contig k141_43270 (Fig. 4B) is a transiently occurring genetic element, like an IME, inserted in tandem with the partial ICE^69^ that provided a fitness advantage during Pre-ABX that it no longer provided in CDI. While a coverage pattern similar to that on contig k141_43270 could be caused by a chimeric contig assembly, the potentially chimeric junction region on contig k141_43270 is strongly supported by paired read coverage mapped with 100% minimum identity which is physical evidence that this sequence existed within the sample (Fig. S32). Overall, our results are in accordance with previous literature describing conjugative elements as mosaic, modular, and highly adaptable to the bacterial hosts and environmental conditions^60,88–90^, and we show that some conjugative elements seem to spread via transduction which would increase their host range, rate of transfer and subsequent prevalence in microbial communities.

#### Transduction of Uncharacterized MGEs

MGEs are difficult to identify and characterize due to their unstable nature in which they can rapidly form and degrade through consecutive insertions, deletions and recombination with other MGEs^69,88,91^. While considerable variability exists, MGEs typically have a different GC content than the core genome, are flanked by DRs as a result of recombination or transposition, are often inserted into the 3’ region of a housekeeping gene, and encode an integrase or transposase for mobilization^70,75,92,93^. The VLP-packaged MGE integrated into Anaeroplasmataceae *Pelethenecus* contig k141_267558 (Fig. 5A) fit all of the described MGE characteristics and based on the presence of Zot, a signature protein required for particle extrusion in filamentous Inoviruses^71^, combined with the lack of phage structural genes, is likely a phage satellite. Phage satellites do not encode their own structural components and instead pirate a ‘helper’ page for packaging. Zot may be a clue that this satellite pirates an Inovirus as its helper. Interestingly, the satellite appears highly unstable due to divergences in WC coverage, read variant density, and evidence for deletion, translocation and duplication.

The VLP-packaged MGE on Pumilibactericeae *MGBC157735* contig k141_31383 (Fig. 5B) is unique for several reasons. First, we were unable to determine the MGE’s mechanism of mobilization due to the lack of an integrase or flanking transposon. Second, the MGE has features associated with several different types of MGEs- transposons (IS30), phage (inovirus gp2 family protein), and conjugative elements (TraG/TraD TraM recognition site domain containing protein) and third, it was inserted into the 3’ region of the YlmC/YmxH family sporulation protein gene rather than a common housekeeping gene, like a tRNA. It has been hypothesized that tRNAs are often used as insertion sites because they are highly conserved and serve a critical role in the cell which means there are guaranteed integration sites in diverse hosts and the genome regions on which they are encoded is stable^75,93^. The YlmC/YmxH family sporulation protein is not only conserved, but is transcribed during sporulation and promoted by the sigma E RNA polymerase^94^. The genes for the sigma E and G RNA polymerases, both of which are involved in sporulation, are located directly upstream of the MGE and therefore may be co-transcribed with the MGE (Fig. S33). We wonder if the MGE is exploiting some process involved in sporulation as there is some evidence that phage can hijack forespores for the encapsidation of their own DNA and may manipulate the sporulation process, however research in these areas is limited^95,96^.

## Conclusions

HGT is an essential process for the generation of genetic diversity, especially in the gut microbiome where it is involved in microbiota resilience to perturbations^97^ and the reshaping of microbiota traits based on influx of genes from transient microbes and changing selective pressures such as diet, disease or medical drugs^98^. HGT in the gut microbiome may thus ultimately have important implications for host health. Our study provides insights into the roles and extent of transduction in HGT in the gut microbiome. The enrichment of specific MGEs and large DNA transfer events in the gut transductomes pre- and post- perturbation suggest that there are regulatory mechanisms driving these changes and thus transduction is highly responsive to perturbations. Additionally, specific community members appear to play a major role in gene exchange through both transduction and the hosting of highly active MGEs. These microbes may be key hubs of HGT in the gut microbiome that warrant further investigation. Finally, we identified several highly abundant and enriched positive TrIdent classifications in both Pre-ABX and CDI conditions associated with uncharacterized MGEs, conjugative plasmids, ICEs, and IMEs. Conjugative MGEs in the transductome suggest that some MGEs may spread through multiple routes of HGT which raises important questions regarding the mechanisms controlling which route is used and under what circumstances.

Our study provides an early example of applying the transductomics toolbox in a real, complex biological setting. Although overall generalizability of our results is limited by the relatively small size of our study, the potential for transductomics as a discovery platform for MGEs was clearly demonstrated. Despite the modest number of samples in our study, we still faced challenges in analyzing and interpreting the amount of potentially interesting patterns obtained with transductomics. While contigs with positive TrIdent classifications only made up 3-20% of the non-redundant dataset in Pre-ABX and CDI conditions, there were still hundreds of coverage patterns associated with VLP-packaged DNA identified. To deal with the large number of detected coverage patterns, we systematically analyzed the data by focusing on the most abundant and enriched patterns, however, this analysis strategy is not the only option.

Furthermore, we used just a few of the tools available for MGE detection and characterization, other tools^74,99–101^ may allow for more discoveries. In future transductomics studies, we hope to take advantage of long-read sequencing to better resolve long coverage patterns associated with generalized, lateral or GTA-mediated transduction and reduce risk of the associated coverage patterns getting split across multiple contigs (Suppl. Text discussion). Additionally, techniques like inverse PCR^102^ and MetaHi-C^103,104^ can validate the integration of MGEs (and eliminate concern about assembly chimeras) and the hosts of extrachromosomal non-integrative MGEs, respectively. Finally, transfer of VLP-fractions into *in vivo* or *in vitro* microbiota might allow us to track actual integration of VLP-packaged DNA into new bacterial hosts by pre- and post-infection sequencing. While our work has provided many important insights into the gut transductome, many questions remain to be answered. We wonder about the rate of integration of VLP-packaged DNA, the timescale of transfer events, the mechanisms by which MGEs are transduced, and how the presence/absence of specific MGEs or bacterial transducers affects the microbial community as a whole. Finally, metagenomics alone often cannot identify the particles encapsulating DNA and therefore techniques such as proteomics or electron microscopy will help in the future to identify the primary transducing VLPs.

## Methods

### Animal housing, treatments and sampling

C57BL/6J mice (male, 4-5 weeks old) purchased from Jackson Laboratories were used for the experimental infections. Mice were housed with autoclaved bedding and water and irradiated food. Cage changes were performed weekly in a laminar flow hood. All mice were subjected to a 12-hour light and 12-hour dark cycle. Animal experiments were conducted in the Laboratory Animal Facilities located on the NCSU CVM campus. The animal facilities are equipped with a full-time animal care staff coordinated by the Laboratory Animal Resources (LAR) division at NCSU. The NCSU CVM is accredited by the Association for the Assessment and Accreditation of Laboratory Animal Care International (AAALAC). Trained animal handlers in the facility fed and assessed the status of animals several times per day. Those assessed as moribund were humanely euthanized by CO_2_ asphyxiation. This protocol is approved by NC State’s Institutional Animal Care and Use Committee (IACUC).

### Experimental design and sampling

The generation of the sequencing data was done as described in Maier *et al.* ^12^. We generated transductomics datasets from murine fecal samples collected during an experiment in which mice (n=4 C57BL/6J males 5-6 weeks old) were treated with antibiotics prior to challenge with *C. difficile*^36^. We collected fecal pellets from each mouse before an antibiotic treatment in which cefoperazone (0.5 g/mL) was provided in the drinking water for 5 days. After 2 days of antibiotic-washout with fresh drinking water, fecal pellets were collected again to represent the post-antibiotic condition. The mice were orally gavaged with ∼1.0 x 10^5^ *C. difficile* spores (strain 630) to induce *C. difficile* infection (CDI). On day 4 of CDI, we collected a final round of fecal pellets from all 4 mice. Each sample collection consisted of two fecal pellets from each mouse replicate. The pellets were manually homogenized in 1.2 mL of SM buffer (100mM NaCl, 8mM MgSO4-7H2O, 50mM TrisHCl) using sterile inoculating loops. After homogenization, the samples were brought up to 2 mL total volume using SM buffer. The fecal homogenates were split into two aliquots, 1.5 mL and 0.5 mL, for WC and VLP-fraction DNA extraction, respectively. Each VLP-fraction was split into two 0.75mL aliquots and one aliquot from each sample was processed with CsCl density gradient ultrancentrifugation.

### DNA extraction and sequencing of transductomics datasets

#### Whole-community DNA extraction

The DNA from the WC aliquots was extracted with the QIAamp PowerFecal Pro DNA kit (Qiagen, catalog no: 51804) according to the manufacturer’s instructions for cells that are difficult to lyse. Unless specified, all centrifugation steps were carried out at 15,000 xG for 1 minute. Briefly, we added the homogenized stool samples to the provided bead beat tubes, added 800µL of CD1 buffer and briefly vortexed to mix. The samples were incubated at 65C for 10 minutes before being vortexed at max speed for 10 minutes. The tubes were centrifuged and the supernatant was transferred to a clean tube. We added 200µL of CD2 buffer and briefly vortexed before centrifuging and transferring supernatant to a clean tube. Next, we added 600µL of CD3 buffer and briefly vortexed before loading the sample on to the provided MB column spin tube and centrifuging. After the entire sample was loaded, the flow through was discarded and the column was placed into a clean collection tube. We added 500µL of EA to the spin tubes and centrifuged before discarding the flow-through. Then, we added 500µL of C5 buffer to the columns, centrifuged, and then placed the column into a clean collection tube where it was centrifuged again at 16,000 xG for 2 minutes. The columns were moved to clean collection tubes where we added 50µL of C6 buffer to the spin columns and centrifuged. The final extracted DNA product was quantified with the Qubit dsDNA HS assay (Thermo Fisher, catalog no: Q32851).

#### VLP-fraction purification

The VLP-fractions were centrifuged at 2,500 xG for 5 minutes and the supernatant was removed and centrifuged for 5,000 xG for an additional 5 minutes to pellet debris. The supernatant was removed, 0.2 µm syringe filtered (VWR, catalog no. 76479-020), treated with 100 U of DNase 1 (Thermo Scientific, catalog no. PI90083) and incubated at 37 C for 2 hours. We set up density-gradients using 1.7, 1.5, and 1.35 g/mL cesium chloride (CsCl) density layers prepared in SM buffer in Beckman ultra-clear tubes (14×89 mm, catalog no. 344059). We marked the interface between the 1.5 and 1.35 g/mL density layers. We loaded the digested VLP-fractions on top of the gradients and ultracentrifuged the tubes at 68,600 xG for 24 hours at 4C. After centrifugation, we used a 27 gauge needle to puncture the tubes ∼2mm below the marked interface and extracted ∼3mL of liquid. We used a 10 Kda Amicon ultra centrifugal filter (Millipore Sigma, catalog no: UFC8010) to wash the samples 3x with SM buffer and concentrate them down to ∼500 µL each.

#### VLP-fraction DNA extraction

First, the VLP samples were lyophilized overnight followed by resuspension in 100µL of PCR-grade water. Next, the resuspended VLPs were combined 1:1 with lysis mix (2% SDS, 90 ug/mL Proteinase K (Thermo Scientific, catalog no. FEREO0491)) and incubated at 55C for 1 hour. The samples were cooled to room temperature before being transferred to pre-spun 5prime light phase-lock tubes (QuantaBio, catalog no: 2302820). We added 500µL of phenol:chloroform:isamyl (25:24:1) and inverted the tubes to mix before centrifuging at 12,000 xG for 5 minutes. The aqueous phase was poured into a new pre-spun phase lock tube. We added 500µL of chloroform:isoamyl (24:1) and inverted to mix before centrifuging at 12,000 xG for 5 minutes. We poured the aqueous phase into a clean microcentrifuge tube and used the MinElute PCR Purification kit (Qiagen, catalog no: 28004) according to the manufacturer’s instructions to clean and concentrate the extracted DNA. Briefly, we diluted the DNA samples 5:1 in buffer PB and applied the diluted sample to a MinElute spin column and centrifuged at 17,900 xG for 1 minute. After the samples were loaded onto the columns, they were washed with 750µL of buffer PE and centrifuged at 17,900 xg for 2 minutes. The columns were placed into clean collection tubes where 20µL of buffer EV was added to the column membranes. The columns were allowed to sit for 1 minute prior to centrifuging at 17,900 xG for 1 minute. The final DNA product was quantified with the Qubit dsDNA HS assay (Thermo Fisher, catalog no: Q32851).

#### Sequencing and read QC for transductomics analysis

DNA Libraries were generated with the Watchmaker DNA library prep kit. Genomic DNA was fragmented using a mastermix of Watchmaker Frag/AT Buffer and Frag/AT enzyme mix. IDTxGen UDI primers and IDT Stubby Adapters were added to each sample followed by 16 cycles of PCR. The final DNA libraries were purified with AMPure magnetic beads and eluted in nuclease-free water. All samples were sequenced with the Illumina NovaSeq X Plus platform at 2×150bp. We received between 50-160M reads for the WC samples and between 35-180M reads for the VLP-fractions. We used TrimGalore to trim the raw reads of Illumina adaptors and polyG sequences identified through FastQC. Next, we used BBSplit with ambig2=split to remove reads that mapped to the host mouse (GCF_000001635.27) or PhiX (NCBI: NC_001422.1) genome.

### Assembly and binning of WC metagenomes

We assembled the WC metagenomes using MEGAHIT^52^ with default parameters. We mapped all of the trimmed and decontaminated whole-community (WC) reads from all replicates and all conditions to each of the WC assemblies using BBMap with default parameters. The resulting .bam files were sorted with SAMtools^105^ then fed into MetaBAT2^106^ for contig binning into metagenome assembled genomes (MAGs) using runMetaBat with default parameters. We performed taxonomic classification of the resulting MAGs using GTDB-Tk’s^107^ classify_wf with the skip-ani-screen parameter.

### Validating CDI and inflammatory response

To validate that *C. difficile* was present on the third sampling day and was not on the first two sampling days, we mapped the whole-community reads to the *C. difficile* 630 genome sequence (NC_009089.1) using BBSplit with ambig2=split and minid=0.98. Next, we used BBSplit again to map the reads that mapped to the *C. difficile* genome back to the whole-community assemblies to determine which sampling days and contigs the *C. difficile* reads were associated with. We calculated *C. difficile* abundance across samples using contigs with greater than 0.1% unambiguous coverage of *C. difficile* reads. As an indirect confirmation of CDI, we quantified the amount of reads in each sample that mapped to the host-mouse genome (described above) to determine inflammatory response.

### Transductomics and TrIdent analysis

Our complete transductomics pipeline is summarized in figure S34.

#### Generation of non-redundant contig set

We generated a non-redundant contig set by concatenating the individual WC assemblies from each replicate in each condition and filtering the combined fasta for contigs greater than 30 kbp. We ran the combined fasta file through the redundancy reduction module of the Redundans^108^ tool with identity=0.95 and overlap=0.70. The filtered contig set was reduced from a total of 8,763 to 3,603 contigs after redundancy reduction. We tested several different minimum overlap cutoff values and found that 70% overlap provided a balance between redundancy removal and preservation of structural variants (SVs) (Suppl. text).

#### Gene annotations and MGE classification in non-redundant contig set

We ran Bakta^109^ in meta mode to obtain gene calls and annotations of the non-redundant contig set. We ran the gene calls output by Pyrodigal (run in Bakta) through Mantis^110^ to generate additional annotations. To obtain final annotations used in figures, we used Bakta^109^ annotations and for any hypothetical proteins, we used the Mantis^110^ annotation if there was one provided. We ran geNomad’s^54^ end-to-end pipeline with score calibration for virus and plasmid identification and ISEscan^73^ for IS identification using the non-redundant contig set as input. We ran ICEscreen^64^ for ICE and IME identification using the Bakta^109^ output .gff file as input. We ran geNomad’s output viral.fna file through Pharokka^55^ in meta mode to obtain phage-associated gene annotations and we ran individual contigs of interest through Pharokka^55^ using g=prodigal (pyrodigal).

#### Taxonomic classification of non-redundant contig set

We performed taxonomic classification of the MAGs using GTDB-Tk’s^107^ classify_wf with the skip-ani-screen parameter. We assigned contig taxonomy using the taxonomy of the MAG they were binned into and for unbinned contigs, we used taxonomy assigned by CAT with the GTDB-Tk^107^ database. Most (3,193 out of 3,603) of the contigs in the non-redundant contig set were contained in MAGs.

#### Relative abundance quantification of whole-community metagenomes and transductomes

We mapped WC and VLP-fraction reads to the non-redundant contig set using BBMap (ambiguous=random, minid=0.97). To obtain the relative abundance of each MAG in the WCs and transductomes, we summed the reads by MAG for each WC and VLP metagenome and divided by the total number of trimmed and decontaminated reads for the respective sample (Fig. S35). MAGs with less than 0.05% relative abundance in a given sample were removed from that sample prior to diversity analyses. We removed viral and prophage-containing contigs (classified with geNomad) prior to determining relative abundances. The enrichment of bacterial families in transductomes relative to the associated WC metagenomes was quantified as a simple ratio of the relative abundances for a bacterial family in the transductome to the relative abundance of that bacterial family in the matched WC metagenome. Unbinned contigs were not included when determining the taxonomic composition of WC metagenomes and transductomes, however they were included when TrIdent was run on the non-redundant contig set. If an unbinned contig was positively classified by TrIdent, its taxonomic classification was obtained with CAT.

#### Diversity calculations

**Alpha diversity:** We calculated Shannon diversity using MAG abundances in the WC and transductome. We used the vegan R package and tested significance with a one-way ANOVA and post-hoc analysis with Tukey’s HSD. We calculated richness by quantifying the number of MAGs represented in each WC and transductome sample. We tested significance with a Kuskal-Wallis test and performed post-hoc analysis with a Dunn test.

**Beta diversity:** We calculated beta diversity with a Bray-Curtis test using MAG abundances in the WC and transductome. We used the vegan R package for the calculation.

#### Generating mapping files for TrIdent analysis

To generate the pileups used as input for TrIdent, we mapped sequencing reads from the WCs and VLP-fractions of replicates 1-3 from each condition (Pre-ABX, Post-ABX and CDI) to the non-redundant contig set using BBMap.sh with the bincov output and ambiguous=all, minid=0.97, and covbinsize=100^12^. While the WC of replicate 4 sequenced well (Table. S1) and was used in generation of the non-redundant contig set, the VLP-fraction sequenced poorly across all three conditions (Table. S2) and was therefore excluded from analysis. We also performed the read mapping against the more redundant contig set (identity=0.97 and overlap=0.95) using secondary=t, ambiguous=random, and minid=0.97 to generate .bam files for visualization of read pair orientation and secondary mappings in IGV^111,112^. Redundant reads can create read mapping artifacts that can lead to false positive pattern-matches. However, by mapping ambiguous reads to all possible mapping locations (ambiguous=all), we smoothed artifacts created by redundancy (Suppl. text). Finally, we mapped reads against the non-redundant contig set using secondary=t, ambiguous=all, and minid=1.0 to generate .bam files for visualization of read coverage at potentially chimeric sites in IGV. We sorted and indexed the .bam files with SAMtools^105,113^ sort and index functions, respectively.

#### TrIdent analysis in R

We ran TrIdent’s TrIdentClassifier() function on all 9 transductomics datasets and merged the output summaryTables. We supplied the number of filtered and decontaminated mapping reads for the respective sample to the TrIdentClassifier() ‘VLPReads’ and ‘WCReads’ parameters. We excluded contigs classified as viral or as containing a prophage by geNomad^54^ for the entire analysis with the exception of quantification of MGEs in positively classified contigs and for the GTA search. We obtained family-level taxonomic classifications for the contigs from their associated metagenome bin. If the contig was unbinned or its bin did not have a family assigned, we used the CAT^66^ annotation.

To determine the top 30 most abundant and enriched positive classifications in Pre-ABX and CDI conditions, we filtered for contigs with positive classifications in at least two replicates in a condition and averaged the abundance and enrichment values across replicates. To calculate the log2 enrichment fold-change between contigs, we filtered for contigs with positive classifications in at least two replicates in both the Pre-ABX and CDI conditions, averaged the associated RTEV values and tools the log2 of (Pre-ABX RTEV avg./CDI RTEV avg.).

We used the ggplot2, tidyr, dplyr, data.table, stringr, ggVennDiagram, gggenes, patchwork, glmmTMB, emmeans, seqinr, multcomp, multcompview, and pheatmap R packages to analyze and visualize the resulting data. We calculated and plotted average GC content across sliding windows of 100 bp.

#### Relative abundance of specific positive TrIdent classifications

We determined positive classification abundance by extracting the maximum pattern-match value for each of TrIdent’s pattern-matches and dividing that value by the number of trimmed and decontaminated VLP-fraction reads for the respective sample 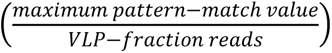. The TrIdent pattern-matches are derived from the VLP-fraction coverage values and therefore the maximum pattern-match value is a reflection of the abundance of the associated coverage pattern.

#### Relative transduction enrichment value

We determined the relative transduction enrichment value (RTEV) by dividing the relative abundance value for a positive TrIdent classification (described above) by the median WC read coverage value for the respective contig divided by the number of trimmed and decontaminated WC reads for the respective sample 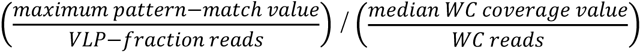. The RTEVs were used to determine the most enriched positive classifications and the positive classifications with the largest change in enrichment.

### MGE-specific analysis

We used the ORF2LCA.txt output from CAT^66^ to identify potentially chimeric contigs by checking to see if a contig’s read coverage pattern was associated with gene annotations assigned to a different taxa than the rest of the contig. We searched for IRs and DRs on select contigs using the pydna_repeatfinder tool^114^ with minlength=10. We created a blastdb of the combined, non-filtered contig set to search for other contigs containing highly abundant and enriched MGEs using BLASTn^67^ (perc_identity=90, qcov_dsp_perc=30). We extracted the protein sequence for the Zot annotation in contig k141_267558 and aligned it with the amino acid motif responsible for Zot receptor binding (GRLCVQDG) ^72^ using a pairwise alignment in the BioEdit software^115^. We used the mpileup and call functions from BCFtools^113^ to call read mapping variants in the .bam files used for the transductomics analysis.

## Abbreviations

VLP: Virus-like particle
WC: Whole-community
MGE: Mobile genetic element
HGT: Horizontal gene transfer
GTA: Gene transfer agent
MV: Membrane vesicle
ABX: Antibiotics
CDI: *C. difficile* infection
IR: Inverted repeat
DR: Direct repeat
IS: Insertion sequence
RTEV: Relative transduction enrichment value
ICE: Integrative conjugative element
IME: Integrative mobile element
GC: Guanine-Cytosine

## Data availability

The raw sequence datasets generated are available in the Sequence Read Archive (SRA) repository under BioProject PRJNA1404639. The assemblies and MAGs used to generate taxonomy are deposited in Dryad (DOI: 10.5061/dryad.76hdr7t9b).

The TrIdent R package can be installed through Bioconductor - https://bioconductor.org/packages/devel/bioc/html/TrIdent.html. Documentation and vignette can be found at github/jlmaier12/TrIdent.

## Supplemental files

**Supplemental Figures, Table and Text: SupplementalMaterial**

**Visual representations of all positive TrIdent pattern-matches for replicates 1-3 in each sampling condition:** Supplemental files 1-8.

## Author contributions

**Jessie Maier:** Conceptualization, Data Curation, Data Analysis, Investigation, Methodology, Visualization, Writing- original draft, Writing- editing and review

**Benjamin Callahan:** Conceptualization, Writing- editing and review

**Breck Duerkop:** Conceptualization, Writing- editing and review

**Manuel Kleiner:** Conceptualization, Data Analysis, Funding Acquisition, Resources, Writing-editing and review

## Acknowledgements

The authors would like to thank the members of the Kleiner Lab for their consistent support and feedback throughout this work. The authors would also like to thank Casey Theriot and Cypress Perkins for their resources and assistance with running the CDI mouse model and collecting fecal pellets for our work.

## References

1. Jyoti & Dey, P. Mechanisms and implications of the gut microbial modulation of intestinal metabolic processes. Npj Metab. Health Dis. 3, 24 (2025).

2. Vos, W. M. de, Tilg, H., Hul, M. V. & Cani, P. D. Gut microbiome and health: mechanistic insights. Gut 71, 1020–1032 (2022).

3. Brito, I. L. Examining horizontal gene transfer in microbial communities. Nat. Rev. Microbiol. 19, 442–453 (2021).

4. Soucy, S. M., Huang, J. & Gogarten, J. P. Horizontal gene transfer: building the web of life. Nat. Rev. Genet. 16, 472–482 (2015).

5. Zhu, S., Hong, J. & Wang, T. Horizontal gene transfer is predicted to overcome the diversity limit of competing microbial species. Nat. Commun. 15, 800 (2024).

6. Peng, H. et al. Longitudinal gut microbiota tracking reveals the dynamics of horizontal gene transfer. Nat. Commun. 16, 11543 (2025).

7. Lerner, A., Matthias, T. & Aminov, R. Potential Effects of Horizontal Gene Exchange in the Human Gut. Front. Immunol. 8, 1630 (2017).

8. Shoeb, E. et al. Horizontal gene transfer of stress resistance genes through plasmid transport. World J. Microbiol. Biotechnol. 28, 1021–1025 (2012).

9. Zhaxybayeva, O. & Nesbø, C. L. Impact of Horizontal Gene Transfer on Adaptations to Extreme Environments. J. Mol. Biol. 438, 169403 (2026).

10. Kleiner, M., Bushnell, B., Sanderson, K. E., Hooper, L. V. & Duerkop, B. A. Transductomics: sequencing-based detection and analysis of transduced DNA in pure cultures and microbial communities. Microbiome 8, 158 (2020).

11. Hyman, P., Trubl, G. & Abedon, S. T. Virus-Like Particle: Evolving Meanings in Different Disciplines. Phage 2, 11–15 (2021).

12. Maier, J. et al. TrIdent - An R package to automate transductomics analysis of virus-like particle mediated DNA mobilization. 2026.03.31.715651 Preprint at 10.64898/2026.03.31.715651 (2026).

13. Eppley, J. M., Biller, S. J., Luo, E., Burger, A. & DeLong, E. F. Marine viral particles reveal an expansive repertoire of phage-parasitizing mobile elements. Proc. Natl. Acad. Sci. 119, e2212722119 (2022).

14. Biller, S. J. et al. Distinct horizontal gene transfer potential of extracellular vesicles versus viral-like particles in marine habitats. Nat. Commun. 16, 2126 (2025).

15. Li, Z. et al. The potential role of viruses in antibiotic resistance gene dissemination in activated sludge viromes. J. Hazard. Mater. 486, 137046 (2025).

16. Mukhopadhyay, A., Choudhury, S. & Kumar, M. Metaviromic analyses of DNA virus community from sediments of the N-Choe stream, North India. Virus Res. 330, 199110 (2023).

17. Diard, M. et al. Inflammation boosts bacteriophage transfer between Salmonella spp. Science 355, 1211–1215 (2017).

18. Fulsundar, S. et al. Gene transfer potential of outer membrane vesicles of Acinetobacter baylyi and effects of stress on vesiculation. Appl. Environ. Microbiol. 80, 3469–3483 (2014).

19. Stanton, T. B., Humphrey, S. B., Sharma, V. K. & Zuerner, R. L. Collateral Effects of Antibiotics: Carbadox and Metronidazole Induce VSH-1 and Facilitate Gene Transfer among Brachyspira hyodysenteriae Strains. Appl. Environ. Microbiol. 74, 2950–2956 (2008).

20. Botka, T. et al. Virion content unpacked by long-read sequencing: stress-induced changes in transmitted staphylococcal mobilome due to phage-satellite interactions. Nucleic Acids Res. 53, gkaf1165 (2025).

21. Sutcliffe, S. G., Shamash, M., Hynes, A. P. & Maurice, C. F. Common Oral Medications Lead to Prophage Induction in Bacterial Isolates from the Human Gut. Viruses 13, 455 (2021).

22. Jiang, X., Hall, A. B., Xavier, R. J. & Alm, E. J. Comprehensive analysis of chromosomal mobile genetic elements in the gut microbiome reveals phylum-level niche-adaptive gene pools. PLOS ONE 14, e0223680 (2019).

23. Tang, Y. et al. Landscape of mobile genetic elements and their functional cargo across the gastrointestinal tract microbiomes in ruminants. Microbiome 13, 162 (2025).

24. Schubert, K., Shosanya, T. & García-Bayona, L. The role of mobile genetic elements in adaptation of the microbiota to the dynamic human gut ecosystem. Curr. Opin. Microbiol. 88, 102675 (2025).

25. Manrique, P. et al. Healthy human gut phageome. Proc. Natl. Acad. Sci. 113, 10400–10405 (2016).

26. Hoyles, L. et al. Characterization of virus-like particles associated with the human faecal and caecal microbiota. Res. Microbiol. 165, 803–812 (2014).

27. Moon, R. C. et al. Epidemiology and Economic Burden of Acute Infectious Gastroenteritis Among Adults Treated in Outpatient Settings in US Health Systems. Off. J. Am. Coll. Gastroenterol. ACG 118, 1069 (2023).

28. Klein, E. Y., et al. Global trends in antibiotic consumption during 2016–2023 and future projections through 2030. Proc. Natl. Acad. Sci. 121, e2411919121 (2024).

29. Theriot, C. M. et al. Cefoperazone-treated mice as an experimental platform to assess differential virulence of Clostridium difficile strains. Gut Microbes 2, 326–334 (2011).

30. Fletcher, J. R. et al. Clostridioides difficile exploits toxin-mediated inflammation to alter the host nutritional landscape and exclude competitors from the gut microbiota. Nat. Commun. 12, 462 (2021).

31. Vincent, C., Mehrotra, S., Loo, V. G., Dewar, K. & Manges, A. R. Excretion of Host DNA in Feces Is Associated with Risk of Clostridium difficile Infection. J. Immunol. Res. 2015, 246203 (2015).

32. Kim, M. et al. Bacterial Interactions with the Host Epithelium. Cell Host Microbe 8, 20–35 (2010).

33. Low, A., Soh, M., Miyake, S. & Seedorf, H. Host Age Prediction from Fecal Microbiota Composition in Male C57BL/6J Mice. Microbiol. Spectr. 10, e00735–22 (2022).

34. Guo, J. et al. Characteristics of gut microbiota in representative mice strains: Implications for biological research. Anim. Models Exp. Med. 5, 337–349 (2022).

35. Borton, M. A. et al. Chemical and pathogen-induced inflammation disrupt the murine intestinal microbiome. Microbiome 5, 47 (2017).

36. Wang, H., et al. Dinoroseobacter shibae Outer Membrane Vesicles Are Enriched for the Chromosome Dimer Resolution Site dif. mSystems 6, 10.1128/msystems.00693-20 (2021).

37. Bitto, N. J. et al. Bacterial membrane vesicles transport their DNA cargo into host cells. Sci. Rep. 7, 7072 (2017).

38. Takano, S. et al. Enrichment of horizontally transferred gene clusters in bacterial extracellular vesicles via non lytic mechanisms. ISME J. 19, wraf193 (2025).

39. Lücking, D., Mercier, C., Alarcón-Schumacher, T. & Erdmann, S. Extracellular vesicles are the main contributor to the non-viral protected extracellular sequence space. ISME Commun. 3, 1–10 (2023).

40. Hynes, A. P., Mercer, R. G., Watton, D. E., Buckley, C. B. & Lang, A. S. DNA packaging bias and differential expression of gene transfer agent genes within a population during production and release of the Rhodobacter capsulatus gene transfer agent, RcGTA. Mol. Microbiol. 85, 314–325 (2012).

41. Fogg, P. C. M. Identification and characterization of a direct activator of a gene transfer agent. Nat. Commun. 10, 595 (2019).

42. Hynes, A. P., Mercer, R. G., Watton, D. E., Buckley, C. B. & Lang, A. S. DNA packaging bias and differential expression of gene transfer agent genes within a population during production and release of the Rhodobacter capsulatus gene transfer agent, RcGTA. Mol. Microbiol. 85, 314–325 (2012).

43. Québatte, M. & Dehio, C. Bartonella gene transfer agent: Evolution, function, and proposed role in host adaptation. Cell. Microbiol. 21, e13068 (2019).

44. Tomasch, J. et al. Packaging of Dinoroseobacter shibae DNA into Gene Transfer Agent Particles Is Not Random. Genome Biol. Evol. 10, 359–369 (2018).

45. Gozzi, K., Tran, N. T., Modell, J. W., Le, T. B. K. & Laub, M. T. Prophage-like gene transfer agents promote Caulobacter crescentus survival and DNA repair during stationary phase. PLOS Biol. 20, e3001790 (2022).

46. Berglund, E. C. et al. Run-Off Replication of Host-Adaptability Genes Is Associated with Gene Transfer Agents in the Genome of Mouse-Infecting Bartonella grahamii. PLOS Genet. 5, e1000546 (2009).

47. Lang, A. S. & Beatty, J. T. Genetic analysis of a bacterial genetic exchange element: The gene transfer agent of Rhodobacter capsulatus. Proc. Natl. Acad. Sci. 97, 859–864 (2000).

48. Hynes, A. P. et al. Functional and Evolutionary Characterization of a Gene Transfer Agent’s Multilocus “Genome”. Mol. Biol. Evol. 33, 2530–2543 (2016).

49. Craske, M. W., Wilson, J. S. & Fogg, P. C. M. Gene transfer agents: structural and functional properties of domesticated viruses. Trends Microbiol. 32, 1200–1211 (2024).

50. Tamarit, D., Neuvonen, M.-M., Engel, P., Guy, L. & Andersson, S. G. E. Origin and Evolution of the Bartonella Gene Transfer Agent. Mol. Biol. Evol. 35, 451–464 (2018).

51. Kogay, R. et al. Formal recognition and classification of gene transfer agents as viriforms. Virus Evol. 8, veac100 (2022).

52. Xu, Y., Liu, B., Jiao, N., Liu, J. & Chen, F. New evidence supports the prophage origin of RcGTA. Appl. Environ. Microbiol. 90, e00434–24 (2024).

53. Québatte, M. et al. Gene Transfer Agent Promotes Evolvability within the Fittest Subpopulation of a Bacterial Pathogen. Cell Syst. 4, 611–621.e6 (2017).

54. Camargo, A. P. et al. Identification of mobile genetic elements with geNomad. Nat. Biotechnol. 42, 1303–1312 (2024).

55. Bouras, G. et al. Pharokka: a fast scalable bacteriophage annotation tool. Bioinformatics 39, btac776 (2023).

56. Fillol-Salom, A. et al. Lateral transduction is inherent to the life cycle of the archetypical Salmonella phage P22. Nat. Commun. 12, 6510 (2021).

57. Banks, E. J. & Le, T. B. K. Co-opting bacterial viruses for DNA exchange: structure and regulation of gene transfer agents. Curr. Opin. Microbiol. 78, 102431 (2024).

58. Tamarit, D., Neuvonen, M.-M., Engel, P., Guy, L. & Andersson, S. G. E. Origin and Evolution of the Bartonella Gene Transfer Agent. Mol. Biol. Evol. 35, 451–464 (2018).

59. Alqurainy, N. et al. A widespread family of phage-inducible chromosomal islands only steals bacteriophage tails to spread in nature. Cell Host Microbe 31, 69–82.e5 (2023).

60. Guglielmini, J., Quintais, L., Garcillán-Barcia, M. P., Cruz, F. de la & Rocha, E. P. C. The Repertoire of ICE in Prokaryotes Underscores the Unity, Diversity, and Ubiquity of Conjugation. PLOS Genet. 7, e1002222 (2011).

61. Tesson, F. et al. Systematic and quantitative view of the antiviral arsenal of prokaryotes. Nat. Commun. 13, 2561 (2022).

62. Néron, B. et al. MacSyFinder v2: Improved modelling and search engine to identify molecular systems in genomes. Peer Community J. 3, (2023).

63. Tesson, F. et al. A Comprehensive Resource for Exploring Antiphage Defense: DefenseFinder Webservice,Wiki and Databases. Peer Community J. 4, (2024).

64. Lao, J., et al. ICEscreen: a tool to detect Firmicute ICEs and IMEs, isolated or enclosed in composite structures. NAR Genomics Bioinforma. 4, lqac079 (2022).

65. Vimberg, V., Zieglerová, L., Buriánková, K., Branny, P. & Balíková Novotná, G. VanZ Reduces the Binding of Lipoglycopeptide Antibiotics to Staphylococcus aureus and Streptococcus pneumoniae Cells. Front. Microbiol. 11, 566 (2020).

66. von Meijenfeldt, F. A. B., Arkhipova, K., Cambuy, D. D., Coutinho, F. H. & Dutilh, B. E. Robust taxonomic classification of uncharted microbial sequences and bins with CAT and BAT. Genome Biol. 20, 217 (2019).

67. Camacho, C. et al. BLAST+: architecture and applications. BMC Bioinformatics 10, 421 (2009).

68. Esposito, D. & Scocca, J. J. The integrase family of tyrosine recombinases: evolution of a conserved active site domain. Nucleic Acids Res. 25, 3605–3614 (1997).

69. Lao, J. et al. Abundance, Diversity and Role of ICEs and IMEs in the Adaptation of Streptococcus salivarius to the Environment. Genes 11, 999 (2020).

70. Parks, A. R. & Peters, J. E. Mobile DNA: Mechanisms, Utility, and Consequences. in Molecular Life Sciences 769–790 (Springer, New York, NY, 2018). doi:10.1007/978-1-4614-1531-2_157.

71. Hay, I. D. & Lithgow, T. Filamentous phages: masters of a microbial sharing economy. EMBO Rep. 20, e47427 (2019).

72. Di Pierro, M. et al. Zonula Occludens Toxin Structure-Function Analysis: IDENTIFICATION OF THE FRAGMENT BIOLOGICALLY ACTIVE ON TIGHT JUNCTIONS AND OF THE ZONULIN RECEPTOR BINDING DOMAIN*. J. Biol. Chem. 276, 19160–19165 (2001).

73. Xie, Z. & Tang, H. ISEScan: automated identification of insertion sequence elements in prokaryotic genomes. Bioinformatics 33, 3340–3347 (2017).

74. Siguier, P., Perochon, J., Lestrade, L., Mahillon, J. & Chandler, M. ISfinder: the reference centre for bacterial insertion sequences. Nucleic Acids Res. 34, D32–D36 (2006).

75. Williams, K. P. Integration sites for genetic elements in prokaryotic tRNA and tmRNA genes: sublocation preference of integrase subfamilies. Nucleic Acids Res. 30, 866–875 (2002).

76. Brito, I. L. et al. Mobile genes in the human microbiome are structured from global to individual scales. Nature 535, 435–439 (2016).

77. Shkoporov, A. N. et al. The Human Gut Virome Is Highly Diverse, Stable, and Individual Specific. Cell Host Microbe 26, 527–541.e5 (2019).

78. Redfield, R. J. & Soucy, S. M. Evolution of Bacterial Gene Transfer Agents. Front. Microbiol. 9, (2018).

79. Borodovich, T. et al. Large-scale capsid-mediated mobilisation of bacterial genomic DNA in the gut microbiome. Nat. Commun. 10.1038/s41467-026-68726-4 (2026) doi:10.1038/s41467-026-68726-4.

80. Pavlovic, G., Burrus, V., Gintz, B., Decaris, B. & Guédon, G. Evolution of genomic islands by deletion and tandem accretion by site-specific recombination: ICESt1-related elements from Streptococcus thermophilus. Microbiology 150, 759–774 (2004).

81. Bellanger, X. et al. Site-specific accretion of an integrative conjugative element together with a related genomic island leads to cis mobilization and gene capture. Mol. Microbiol. 81, 912–925 (2011).

82. Guédon, G., Libante, V., Coluzzi, C., Payot, S. & Leblond-Bourget, N. The Obscure World of Integrative and Mobilizable Elements, Highly Widespread Elements that Pirate Bacterial Conjugative Systems. Genes 8, 337 (2017).

83. Lee, C. A., Thomas, J. & Grossman, A. D. The Bacillus subtilis Conjugative Transposon ICEBs1 Mobilizes Plasmids Lacking Dedicated Mobilization Functions. J. Bacteriol. 194, 3165–3172 (2012).

84. Ares-Arroyo, M., Coluzzi, C. & Rocha, E. P. C. Origins of transfer establish networks of functional dependencies for plasmid transfer by conjugation. Nucleic Acids Res. 51, 3001–3016 (2022).

85. He, L. et al. Chimeric infective particles expand species boundaries in phage-inducible chromosomal island mobilization. Cell 188, 6636–6653.e17 (2025).

86. de Sousa, J. A. M., Fillol-Salom, A., Penadés, J. R. & Rocha, E. P. C. Identification and characterization of thousands of bacteriophage satellites across bacteria. Nucleic Acids Res. 51, 2759–2777 (2023).

87. Coluzzi, C., Garcillán-Barcia, M. P., de la Cruz, F. & Rocha, E. P. C. Evolution of Plasmid Mobility: Origin and Fate of Conjugative and Nonconjugative Plasmids. 10.1093/molbev/msac115.

88. Osborn, A. M. & Böltner, D. When phage, plasmids, and transposons collide: genomic islands, and conjugative- and mobilizable-transposons as a mosaic continuum. Plasmid 48, 202–212 (2002).

89. Wozniak, R. A. F. & Waldor, M. K. Integrative and conjugative elements: mosaic mobile genetic elements enabling dynamic lateral gene flow. Nat. Rev. Microbiol. 8, 552–563 (2010).

90. Norberg, P., Bergström, M., Jethava, V., Dubhashi, D. & Hermansson, M. The IncP-1 plasmid backbone adapts to different host bacterial species and evolves through homologous recombination. Nat. Commun. 2, 268 (2011).

91. Horne, T., Orr, V. T. & Hall, J. P. How do interactions between mobile genetic elements affect horizontal gene transfer? Curr. Opin. Microbiol. 73, 102282 (2023).

92. Almpanis, A., Swain, M., Gatherer, D. & McEwan, N. Correlation between bacterial G+C content, genome size and the G+C content of associated plasmids and bacteriophages. *Microb*. Genomics 4, e000168 (2018).

93. Reiter, W. D., Palm, P. & Yeats, S. Transfer RNA genes frequently serve as integration sites for prokaryotic genetic elements. Nucleic Acids Res. 17, 1907–1914 (1989).

94. Eichenberger, P. et al. The σE Regulon and the Identification of Additional Sporulation Genes in *Bacillus subtilis*. J. Mol. Biol. 327, 945–972 (2003).

95. Butala, M. & Dragoš, A. Unique relationships between phages and endospore-forming hosts. Trends Microbiol. 31, 498–510 (2023).

96. Schwartz, D. A. et al. Human-Gut Phages Harbor Sporulation Genes. mBio 14, e00182–23 (2023).

97. Coyte, K. Z. et al. Horizontal gene transfer and ecological interactions jointly control microbiome stability. PLOS Biol. 20, e3001847 (2022).

98. Lerner, A., Matthias, T. & Aminov, R. Potential Effects of Horizontal Gene Exchange in the Human Gut. Front. Immunol. 8, (2017).

99. Guo, J. et al. VirSorter2: a multi-classifier, expert-guided approach to detect diverse DNA and RNA viruses. Microbiome 9, 37 (2021).

100. Kieft, K., Zhou, Z. & Anantharaman, K. VIBRANT: automated recovery, annotation and curation of microbial viruses, and evaluation of viral community function from genomic sequences. Microbiome 8, 90 (2020).

101. Cho, Y., Kim, E., Kim, M. & Rho, M. DeepMobilome: predicting mobile genetic elements using sequencing reads of microbiomes. Brief. Bioinform. 26, bbaf450 (2025).

102. Tansirichaiya, S., Winje, E., Wigand, J. & Al-Haroni, M. Inverse PCR-based detection reveal novel mobile genetic elements and their associated genes in the human oral metagenome. BMC Oral Health 22, 210 (2022).

103. Marbouty, M., Thierry, A., Millot, G. A. & Koszul, R. MetaHiC phage-bacteria infection network reveals active cycling phages of the healthy human gut. eLife 10, e60608.

104. Wu, R. et al. Hi-C metagenome sequencing reveals soil phage–host interactions. Nat. Commun. 14, 7666 (2023).

105. Li, H. et al. The Sequence Alignment/Map format and SAMtools. Bioinformatics 25, 2078–2079 (2009).

106. Kang, D. D. et al. MetaBAT 2: an adaptive binning algorithm for robust and efficient genome reconstruction from metagenome assemblies. PeerJ 7, e7359 (2019).

107. Chaumeil, P.-A., Mussig, A. J., Hugenholtz, P. & Parks, D. H. GTDB-Tk: a toolkit to classify genomes with the Genome Taxonomy Database. Bioinformatics 36, 1925–1927 (2020).

108. Pryszcz, L. P. & Gabaldón, T. Redundans: an assembly pipeline for highly heterozygous genomes. Nucleic Acids Res. 44, e113 (2016).

109. Schwengers, O. et al. Bakta: rapid and standardized annotation of bacterial genomes via alignment-free sequence identification. *Microb*. Genomics 7, 000685 (2021).

110. Queirós, P., Delogu, F., Hickl, O., May, P. & Wilmes, P. Mantis: flexible and consensus-driven genome annotation. GigaScience 10, giab042 (2021).

111. Robinson, J. T. et al. Integrative Genomics Viewer. Nat. Biotechnol. 29, 24–26 (2011).

112. Thorvaldsdóttir, H., Robinson, J. T. & Mesirov, J. P. Integrative Genomics Viewer (IGV): high-performance genomics data visualization and exploration. Brief. Bioinform. 14, 178–192 (2013).

113. Danecek, P. et al. Twelve years of SAMtools and BCFtools. GigaScience 10, giab008 (2021).

114. Benson, G. Tandem repeats finder: a program to analyze DNA sequences. Nucleic Acids Res. 27, 573–580 (1999).

115. Hall, T. BioEdit: A User-Friendly Biological Sequence Alignment Editor and Analysis Program for Windows 95/98/NT. Nucleic Acids Symp. Ser. 41, 95–98 (1999).

