## Supplemental file 6 for "Perturbations shift the composition of bacterial DNA carried by virus-like particles in the murine gut microbiome"

**File S6) All positive TrIdent pattern-matches for CDI replicate 1.** The sample from which a contig was assembled is indicated by the following prefixes in the contig accession number: WC1-1 = Pre-ABX replicate 1, WC1-2 = Pre-ABX replicate 2, WC1-6 = Pre-ABX replicate 3, WC1-7 = Pre-ABX replicate 4, WC1-1 = CDI replicate 1, WC1-2 = CDI replicate 2, WC1-6 = CDI replicate 3, WC1-7 = CDI replicate 4. Pre-ABX = Pre-antibiotics, Post-ABX= post-antbiotics, CDI= *Clostridioides difficile* infection. Contig taxonomy and geNomad prophage classification in plot subtitle. Plots are listed alphabetically by family name.

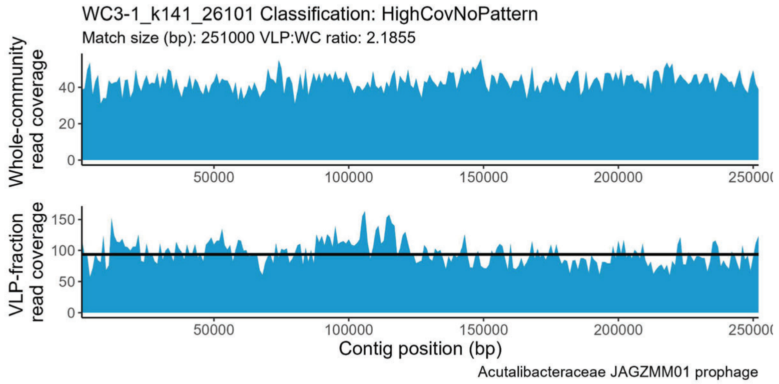


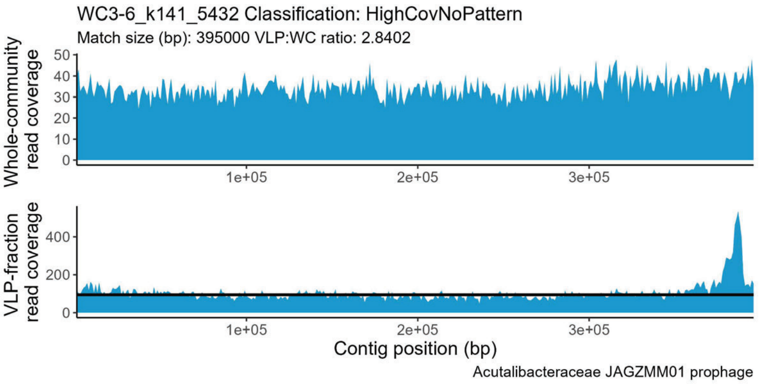


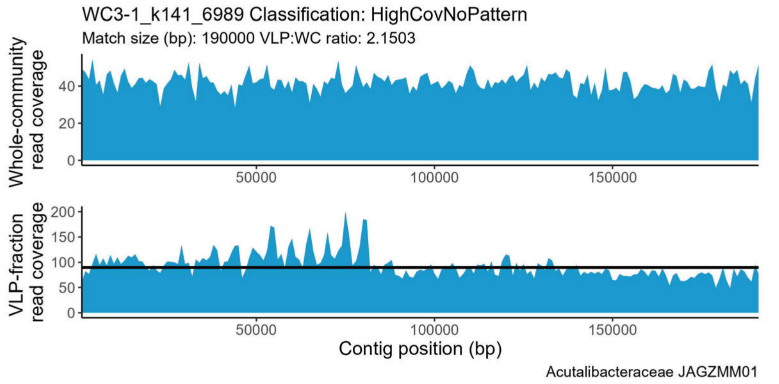


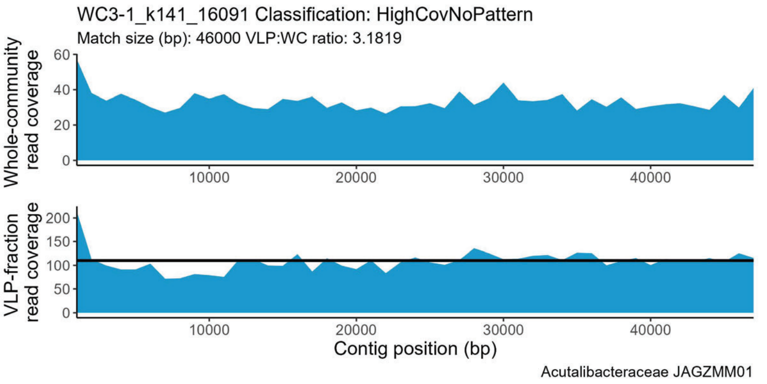


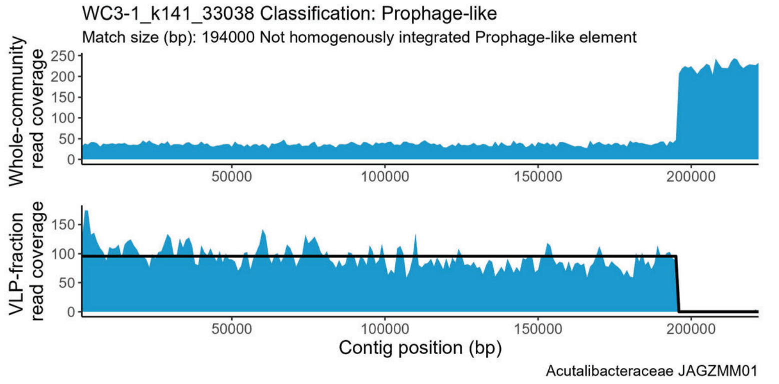


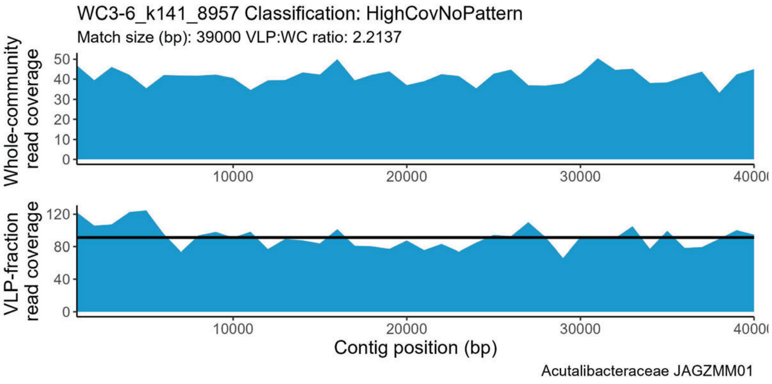


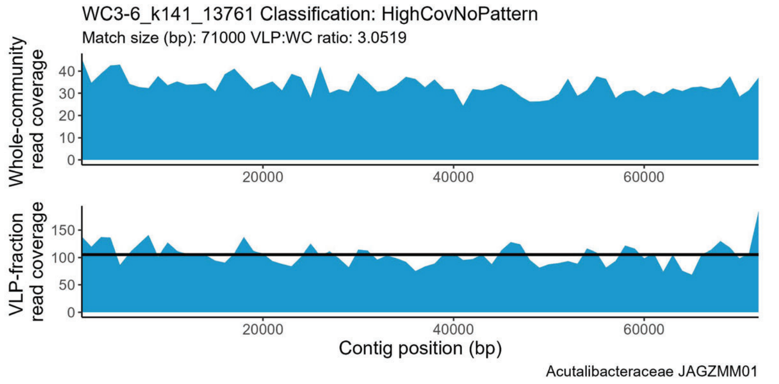


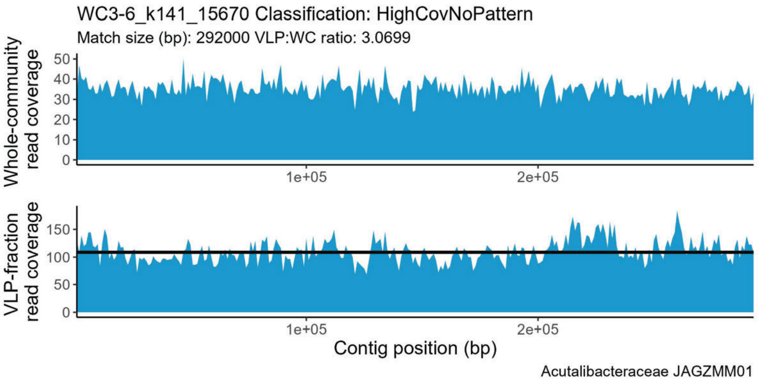

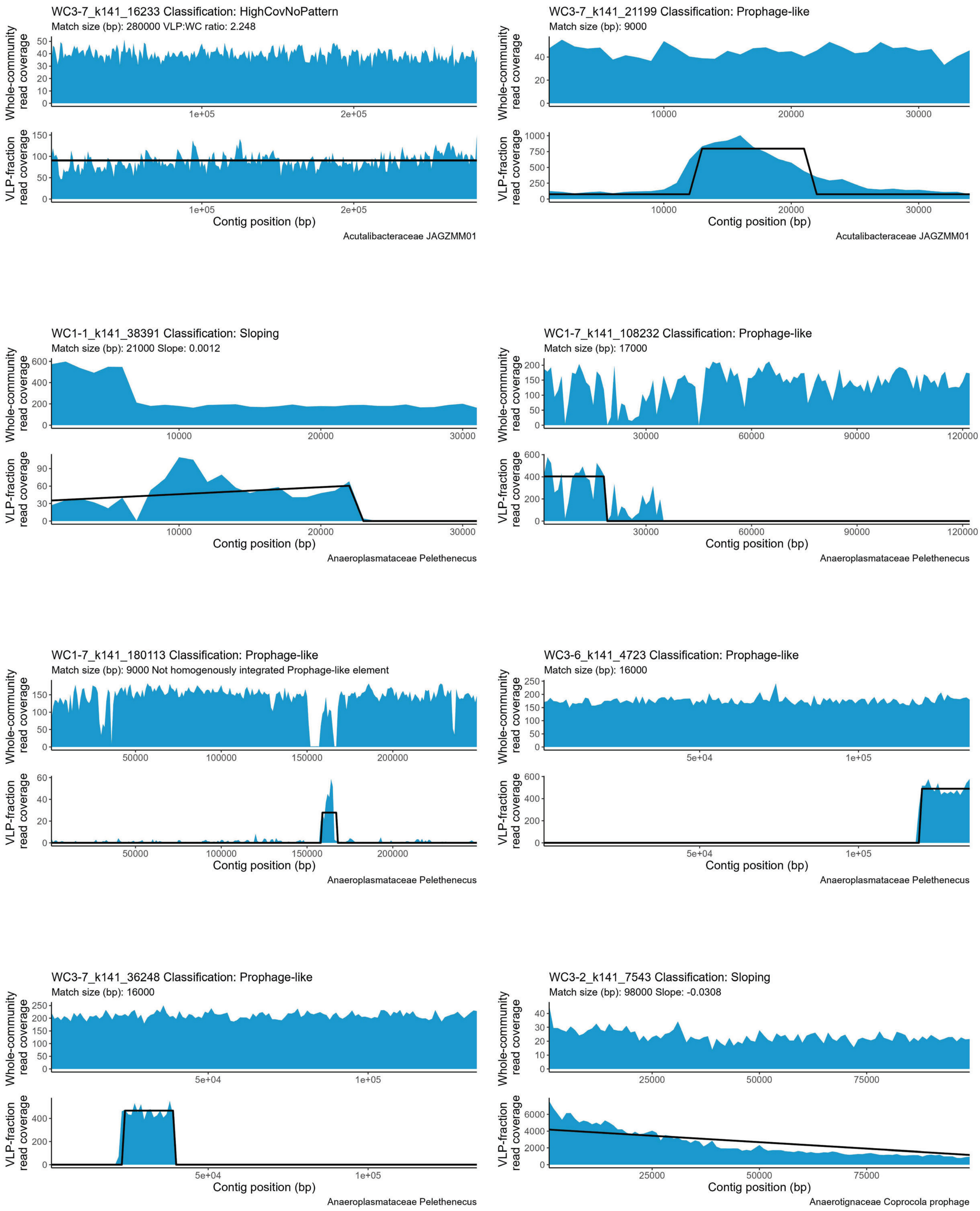
WC3-7_k141_16233 Classification: HighCovNoPattern
Match size (bp): 280000 VLP:WC ratio: 2.248
Acutalibacteraceae JAGZMM01
WC3-7_k141_21199 Classification: Prophage-like
Match size (bp): 9000
Acutalibacteraceae JAGZMM01
WC1-1_k141_38391 Classification: Sloping
Match size (bp): 21000 Slope: 0.0012
Anaeroplasmataceae Pelethenecus
WC1-7_k141_108232 Classification: Prophage-like
Match size (bp): 17000
Anaeroplasmataceae Pelethenecus
WC1-7_k141_180113 Classification: Prophage-like
Match size (bp): 9000 Not homogenously integrated Prophage-like element
Anaeroplasmataceae Pelethenecus
WC3-6_k141_4723 Classification: Prophage-like
Match size (bp): 16000
Anaeroplasmataceae Pelethenecus
WC3-7_k141_36248 Classification: Prophage-like
Match size (bp): 16000
Anaeroplasmataceae Pelethenecus
WC3-2_k141_7543 Classification: Sloping
Match size (bp): 98000 Slope: -0.0308
Anaerotignaceae Coprocola prophage
Whole-community read coverage
VLP-fraction read coverage
Contig position (bp)

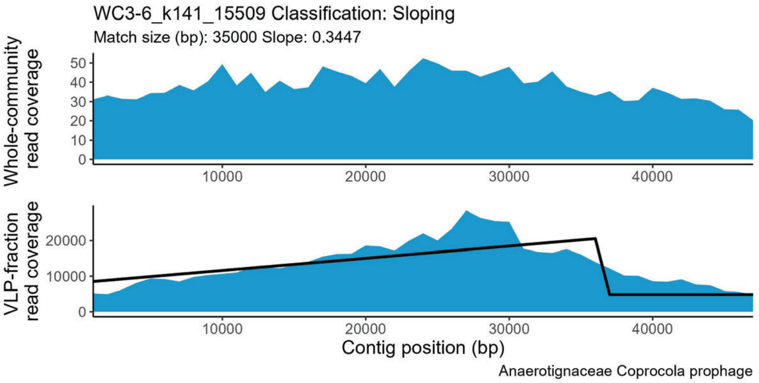
WC3-6_k141_15509 Classification: Sloping
Match size (bp): 35000 Slope: 0.3447
Whole-community read coverage
0 10 20 30 40 50
VLP-fraction read coverage
0 10000 20000
10000 20000 30000 40000
Contig position (bp)
Anaerotignaceae Coprocola prophage

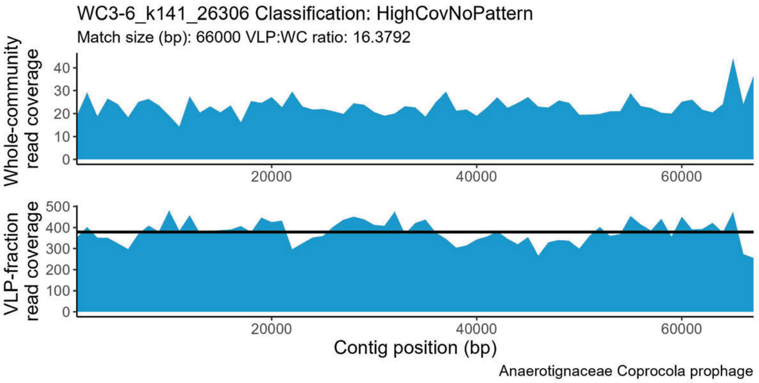
WC3-6_k141_26306 Classification: HighCovNoPattern
Match size (bp): 66000 VLP:WC ratio: 16.3792
Whole-community read coverage
0 10 20 30 40
VLP-fraction read coverage
0 100 200 300 400 500
20000 40000 60000
Contig position (bp)
Anaerotignaceae Coprocola prophage

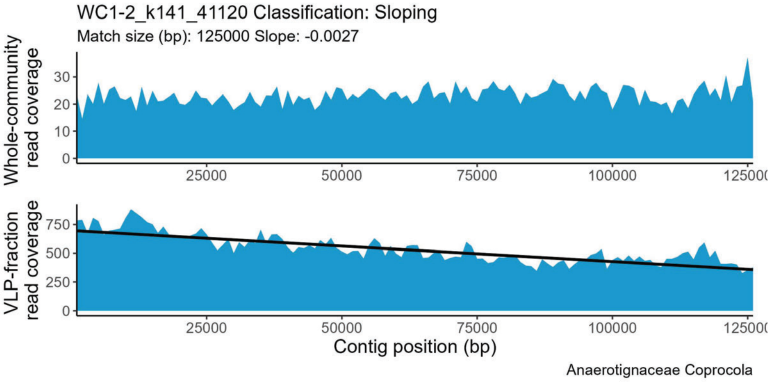
WC1-2_k141_41120 Classification: Sloping
Match size (bp): 125000 Slope: -0.0027
Whole-community read coverage
0 10 20 30
VLP-fraction read coverage
0 250 500 750
25000 50000 75000 100000 125000
Contig position (bp)
Anaerotignaceae Coprocola

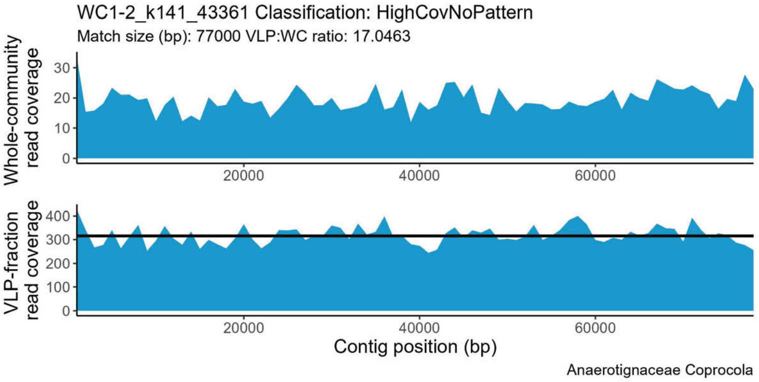
WC1-2_k141_43361 Classification: HighCovNoPattern
Match size (bp): 77000 VLP:WC ratio: 17.0463
Whole-community read coverage
0 10 20 30
VLP-fraction read coverage
0 100 200 300 400
20000 40000 60000
Contig position (bp)
Anaerotignaceae Coprocola

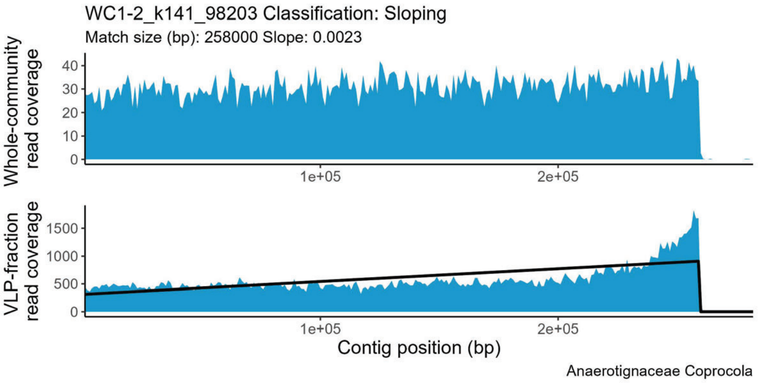
WC1-2_k141_98203 Classification: Sloping
Match size (bp): 258000 Slope: 0.0023
Whole-community read coverage
0 10 20 30 40
VLP-fraction read coverage
0 500 1000 1500
1e+05 2e+05
Contig position (bp)
Anaerotignaceae Coprocola

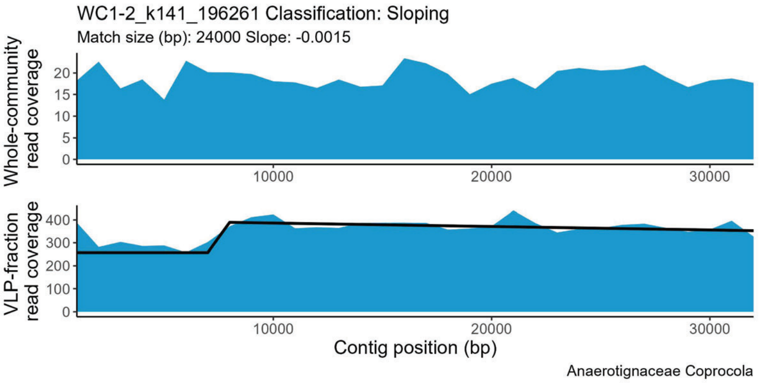
WC1-2_k141_196261 Classification: Sloping
Match size (bp): 24000 Slope: -0.0015
Whole-community read coverage
0 5 10 15 20
VLP-fraction read coverage
0 100 200 300 400
10000 20000 30000
Contig position (bp)
Anaerotignaceae Coprocola

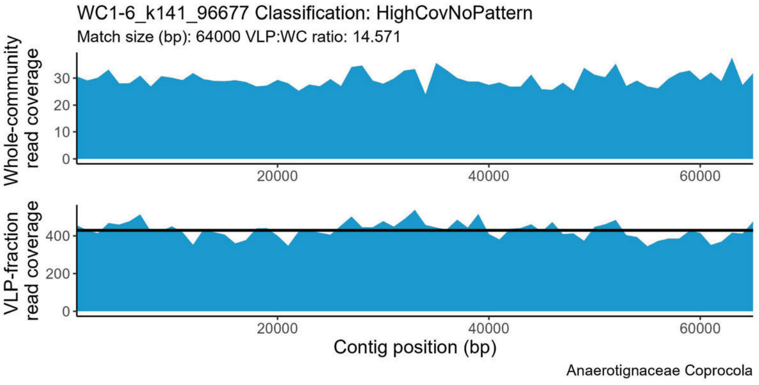
WC1-6_k141_96677 Classification: HighCovNoPattern
Match size (bp): 64000 VLP:WC ratio: 14.571
Whole-community read coverage
0 10 20 30
VLP-fraction read coverage
0 200 400
20000 40000 60000
Contig position (bp)
Anaerotignaceae Coprocola

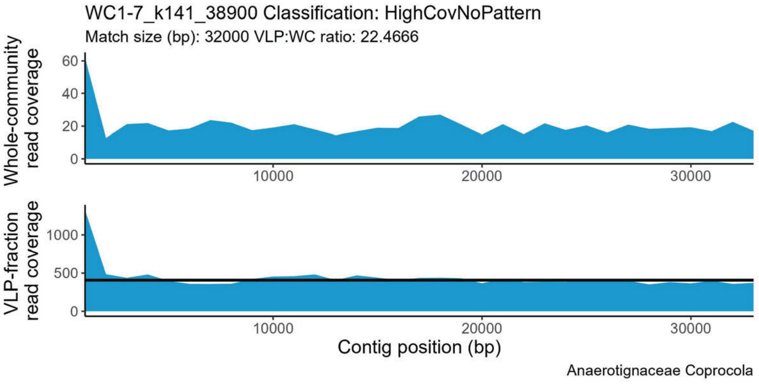
WC1-7_k141_38900 Classification: HighCovNoPattern
Match size (bp): 32000 VLP:WC ratio: 22.4666
Whole-community read coverage
0 20 40 60
VLP-fraction read coverage
0 500 1000
10000 20000 30000
Contig position (bp)
Anaerotignaceae Coprocola

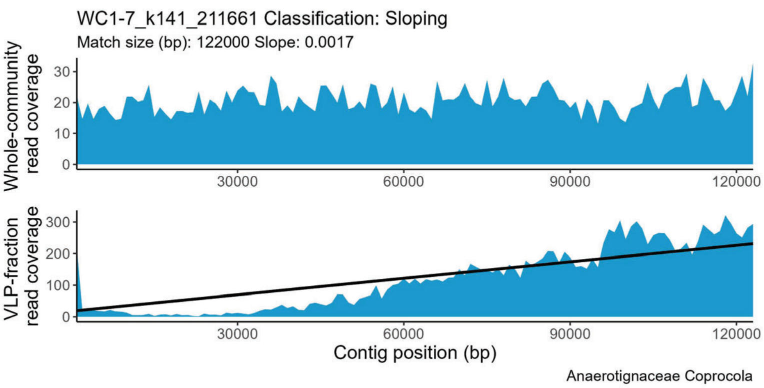
WC1-7_k141_211661 Classification: Sloping
Match size (bp): 122000 Slope: 0.0017
Whole-community read coverage
VLP-fraction read coverage
Contig position (bp)
Anaerotignaceae Coprocola

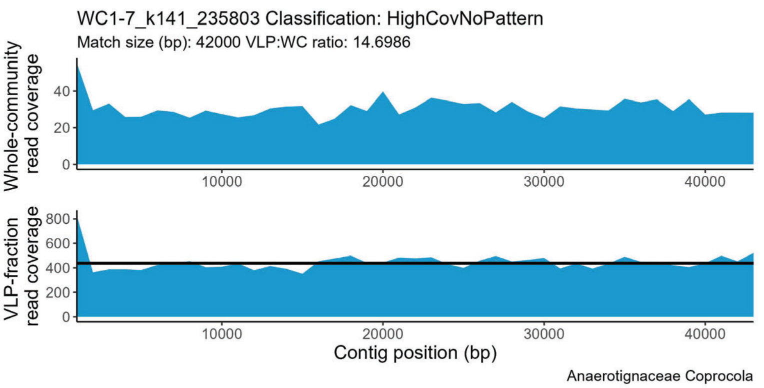
WC1-7_k141_235803 Classification: HighCovNoPattern
Match size (bp): 42000 VLP:WC ratio: 14.6986
Whole-community read coverage
VLP-fraction read coverage
Contig position (bp)
Anaerotignaceae Coprocola

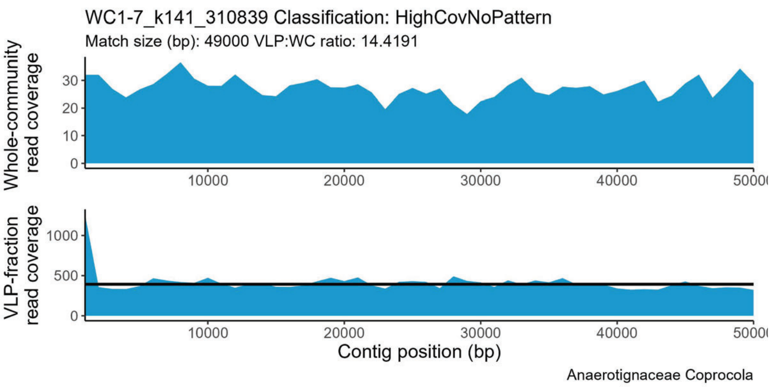
WC1-7_k141_310839 Classification: HighCovNoPattern
Match size (bp): 49000 VLP:WC ratio: 14.4191
Whole-community read coverage
VLP-fraction read coverage
Contig position (bp)
Anaerotignaceae Coprocola

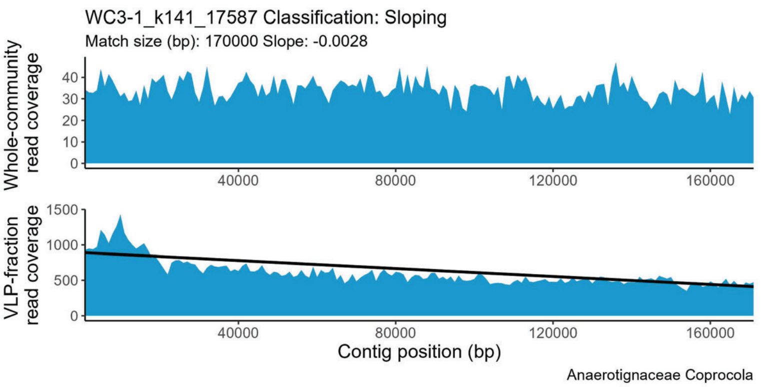
WC3-1_k141_17587 Classification: Sloping
Match size (bp): 170000 Slope: -0.0028
Whole-community read coverage
VLP-fraction read coverage
Contig position (bp)
Anaerotignaceae Coprocola

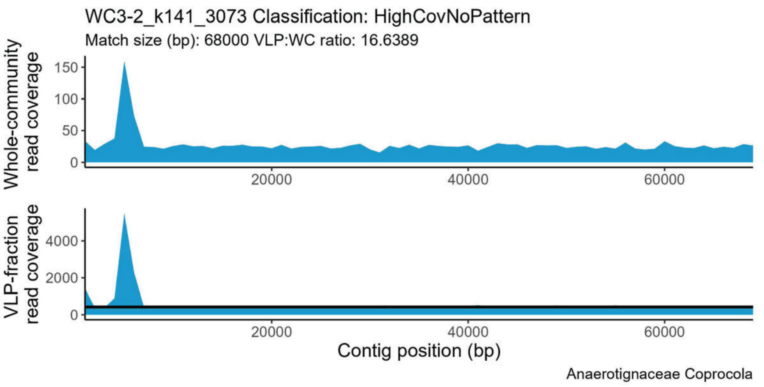
WC3-2_k141_3073 Classification: HighCovNoPattern
Match size (bp): 68000 VLP:WC ratio: 16.6389
Whole-community read coverage
VLP-fraction read coverage
Contig position (bp)
Anaerotignaceae Coprocola

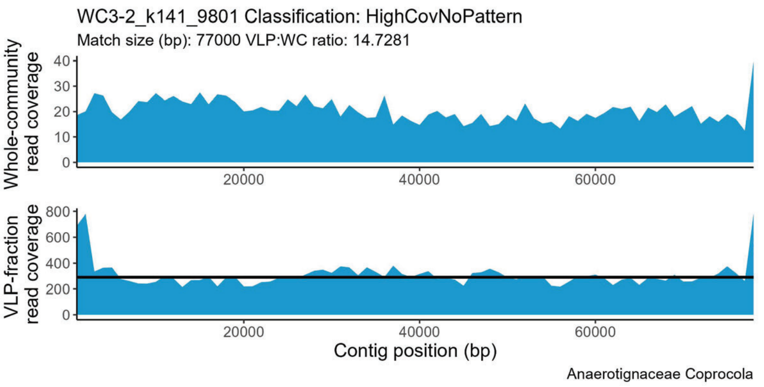
WC3-2_k141_9801 Classification: HighCovNoPattern
Match size (bp): 77000 VLP:WC ratio: 14.7281
Whole-community read coverage
VLP-fraction read coverage
Contig position (bp)
Anaerotignaceae Coprocola

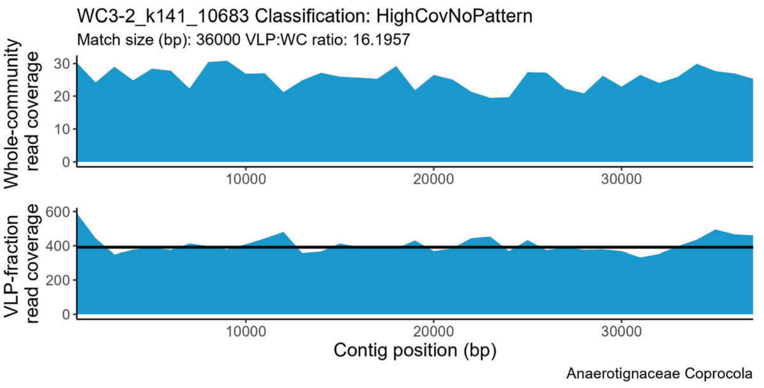
WC3-2_k141_10683 Classification: HighCovNoPattern
Match size (bp): 36000 VLP:WC ratio: 16.1957
Whole-community read coverage
VLP-fraction read coverage
Contig position (bp)
Anaerotignaceae Coprocola

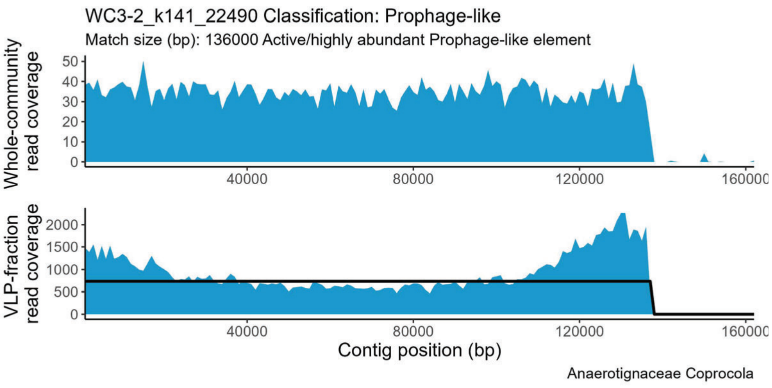
WC3-2_k141_22490 Classification: Prophage-like
Match size (bp): 136000 Active/highly abundant Prophage-like element
Whole-community read coverage
VLP-fraction read coverage
Contig position (bp)
Anaerotignaceae Coprocola

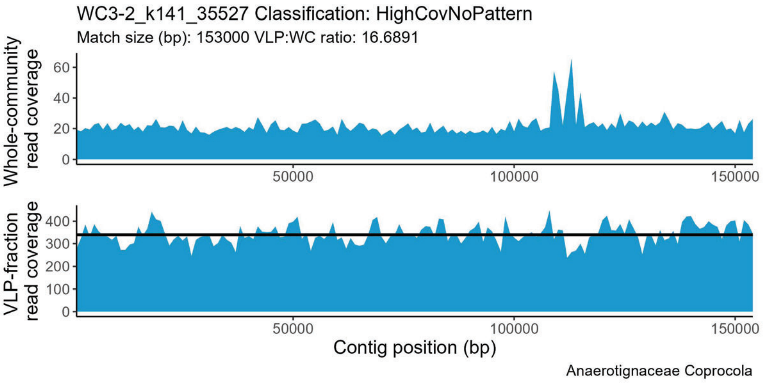

WC3-2_k141_35527 Classification: HighCovNoPattern
Match size (bp): 153000 VLP:WC ratio: 16.6891
Whole-community read coverage
0
20
40
60
50000
100000
150000
VLP-fraction read coverage
0
100
200
300
400
Contig position (bp)
Anaerotignaceae Coprocola


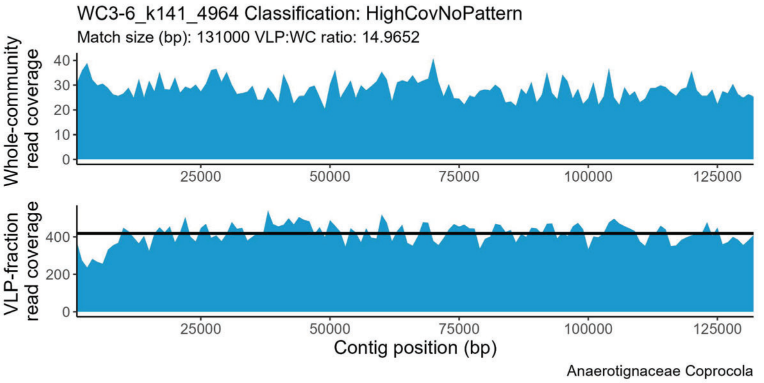

WC3-6_k141_4964 Classification: HighCovNoPattern
Match size (bp): 131000 VLP:WC ratio: 14.9652
Whole-community read coverage
0
10
20
30
40
25000
50000
75000
100000
125000
VLP-fraction read coverage
0
200
400
Contig position (bp)
Anaerotignaceae Coprocola


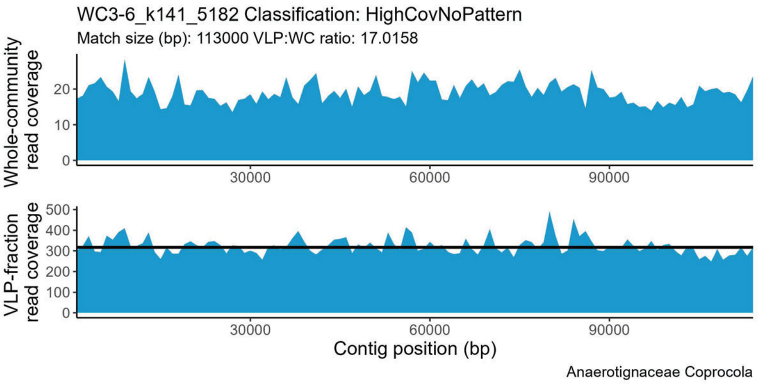

WC3-6_k141_5182 Classification: HighCovNoPattern
Match size (bp): 113000 VLP:WC ratio: 17.0158
Whole-community read coverage
0
10
20
30000
60000
90000
VLP-fraction read coverage
0
100
200
300
400
500
Contig position (bp)
Anaerotignaceae Coprocola


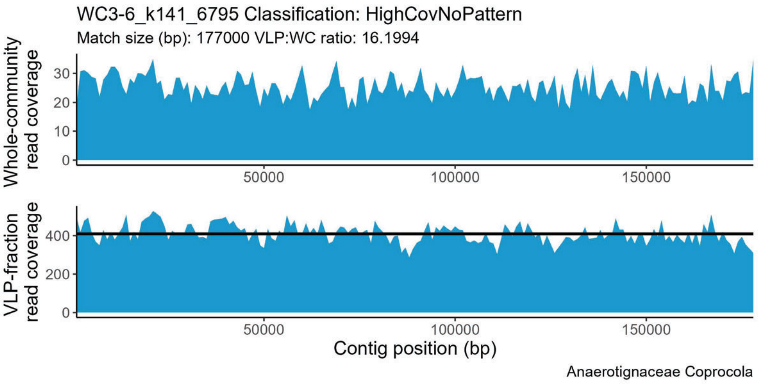

WC3-6_k141_6795 Classification: HighCovNoPattern
Match size (bp): 177000 VLP:WC ratio: 16.1994
Whole-community read coverage
0
10
20
30
50000
100000
150000
VLP-fraction read coverage
0
200
400
Contig position (bp)
Anaerotignaceae Coprocola


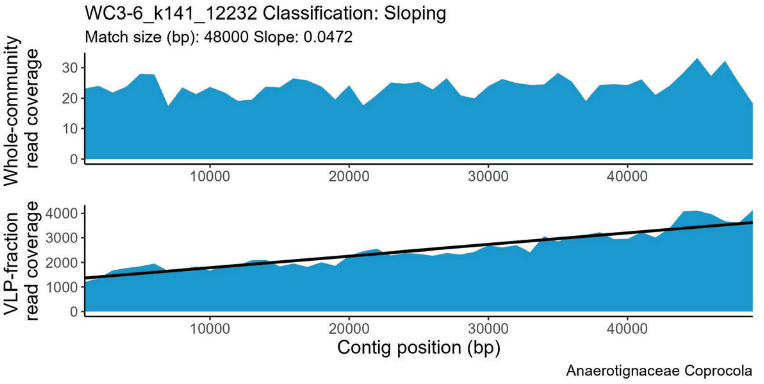

WC3-6_k141_12232 Classification: Sloping
Match size (bp): 48000 Slope: 0.0472
Whole-community read coverage
0
10
20
30
10000
20000
30000
40000
VLP-fraction read coverage
0
1000
2000
3000
4000
Contig position (bp)
Anaerotignaceae Coprocola


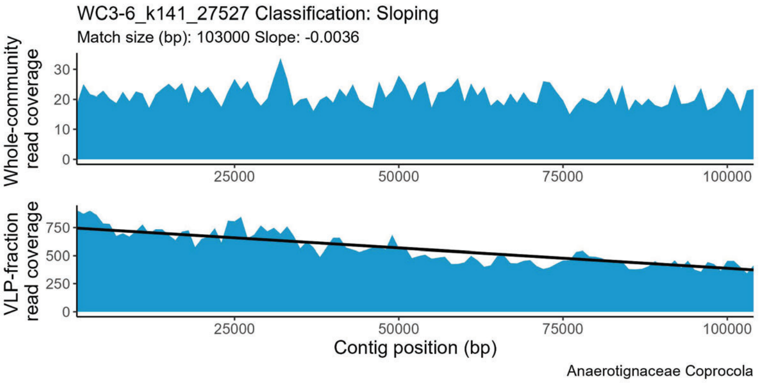

WC3-6_k141_27527 Classification: Sloping
Match size (bp): 103000 Slope: -0.0036
Whole-community read coverage
0
10
20
30
25000
50000
75000
100000
VLP-fraction read coverage
0
250
500
750
Contig position (bp)
Anaerotignaceae Coprocola


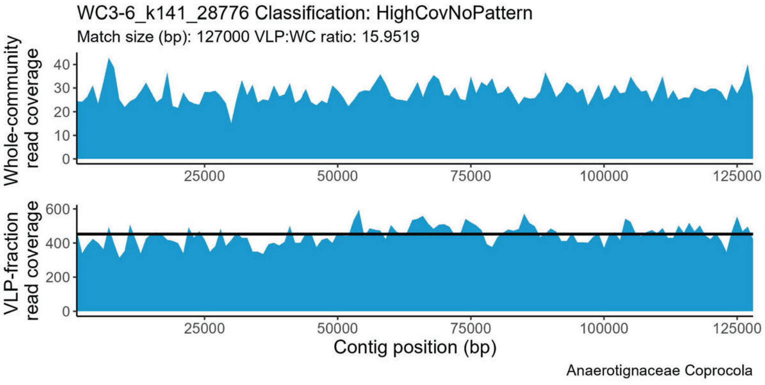

WC3-6_k141_28776 Classification: HighCovNoPattern
Match size (bp): 127000 VLP:WC ratio: 15.9519
Whole-community read coverage
0
10
20
30
40
25000
50000
75000
100000
125000
VLP-fraction read coverage
0
200
400
600
Contig position (bp)
Anaerotignaceae Coprocola


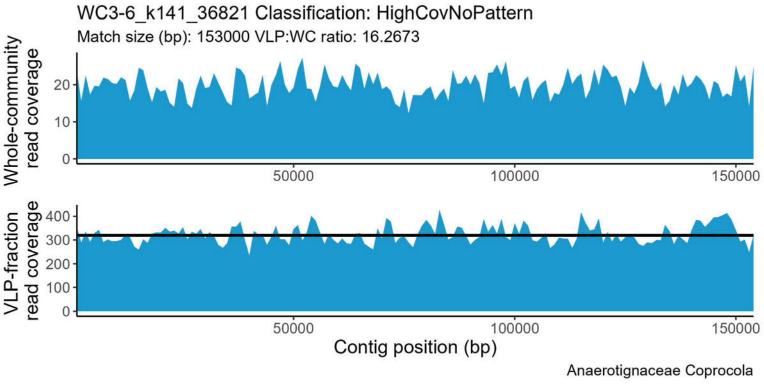

WC3-6_k141_36821 Classification: HighCovNoPattern
Match size (bp): 153000 VLP:WC ratio: 16.2673
Whole-community read coverage
0
10
20
50000
100000
150000
VLP-fraction read coverage
0
100
200
300
400
Contig position (bp)
Anaerotignaceae Coprocola

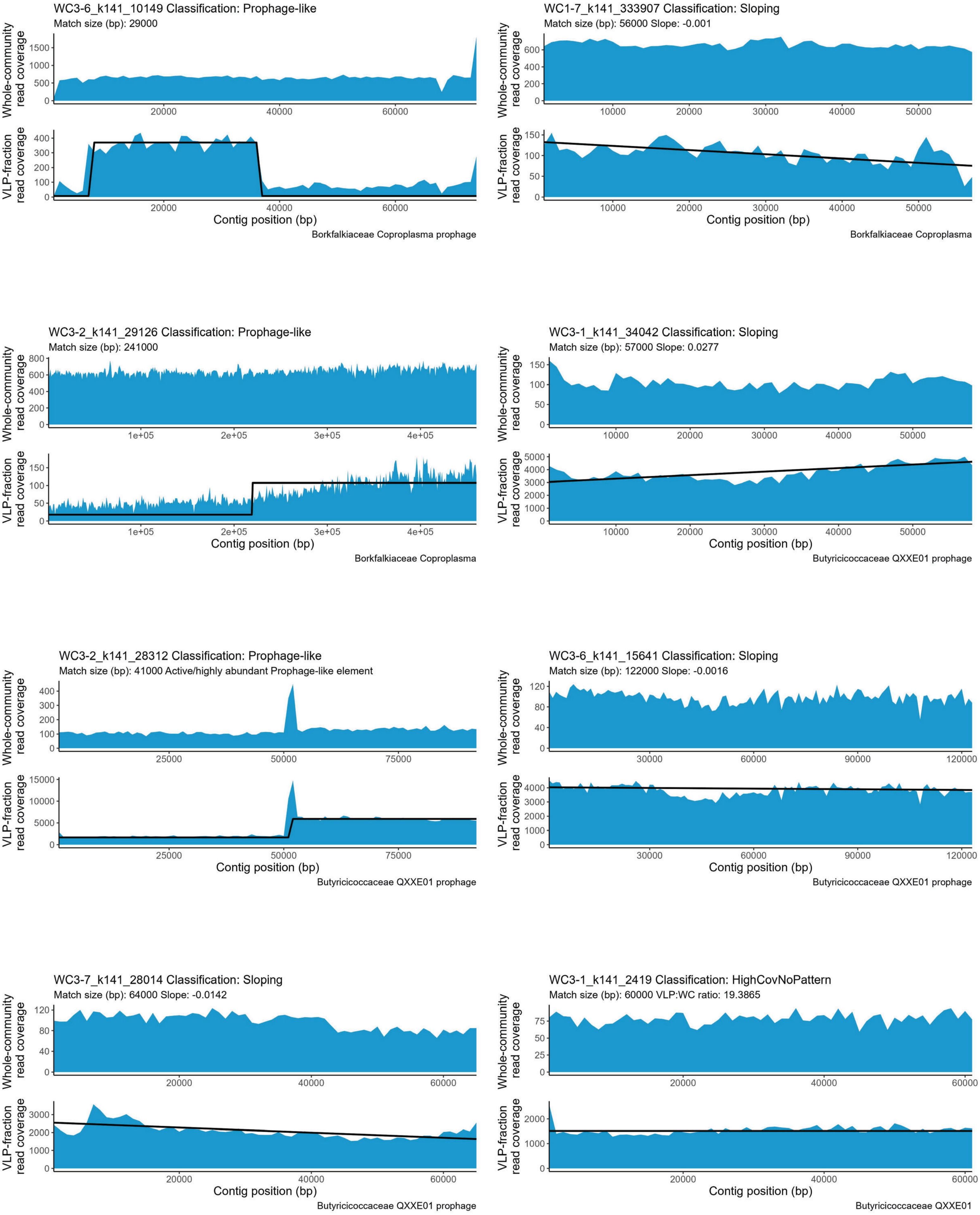
WC3-6_k141_10149 Classification: Prophage-like
Match size (bp): 29000
Whole-community read coverage
VLP-fraction read coverage
Contig position (bp)
Borkfalkiaceae Coproplasma prophage
WC1-7_k141_333907 Classification: Sloping
Match size (bp): 56000 Slope: -0.001
Whole-community read coverage
VLP-fraction read coverage
Contig position (bp)
Borkfalkiaceae Coproplasma
WC3-2_k141_29126 Classification: Prophage-like
Match size (bp): 241000
Whole-community read coverage
VLP-fraction read coverage
Contig position (bp)
Borkfalkiaceae Coproplasma
WC3-1_k141_34042 Classification: Sloping
Match size (bp): 57000 Slope: 0.0277
Whole-community read coverage
VLP-fraction read coverage
Contig position (bp)
Butyricicoccaceae QXXE01 prophage
WC3-2_k141_28312 Classification: Prophage-like
Match size (bp): 41000 Active/highly abundant Prophage-like element
Whole-community read coverage
VLP-fraction read coverage
Contig position (bp)
Butyricicoccaceae QXXE01 prophage
WC3-6_k141_15641 Classification: Sloping
Match size (bp): 122000 Slope: -0.0016
Whole-community read coverage
VLP-fraction read coverage
Contig position (bp)
Butyricicoccaceae QXXE01 prophage
WC3-7_k141_28014 Classification: Sloping
Match size (bp): 64000 Slope: -0.0142
Whole-community read coverage
VLP-fraction read coverage
Contig position (bp)
Butyricicoccaceae QXXE01 prophage
WC3-1_k141_2419 Classification: HighCovNoPattern
Match size (bp): 60000 VLP:WC ratio: 19.3865
Whole-community read coverage
VLP-fraction read coverage
Contig position (bp)
Butyricicoccaceae QXXE01

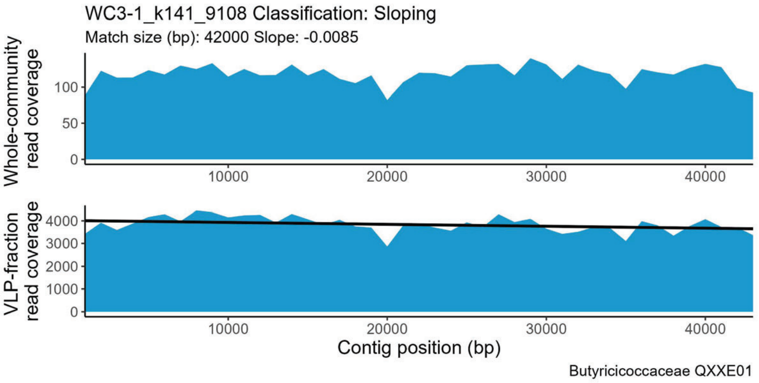
WC3-1_k141_9108 Classification: Sloping
Match size (bp): 42000 Slope: -0.0085
Whole-community read coverage
VLP-fraction read coverage
Contig position (bp)
Butyricicoccaceae QXXE01

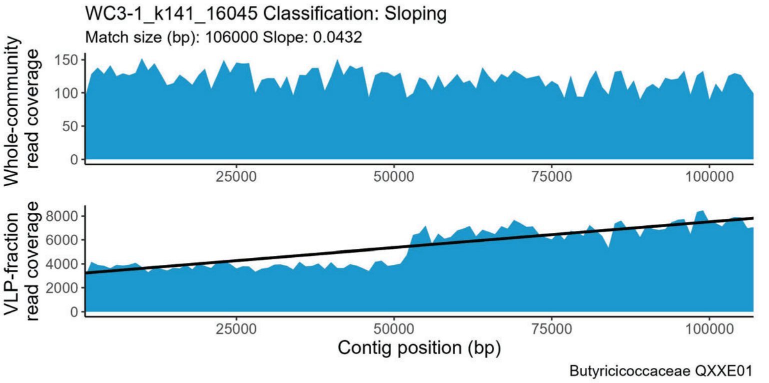
WC3-1_k141_16045 Classification: Sloping
Match size (bp): 106000 Slope: 0.0432
Whole-community read coverage
VLP-fraction read coverage
Contig position (bp)
Butyricicoccaceae QXXE01

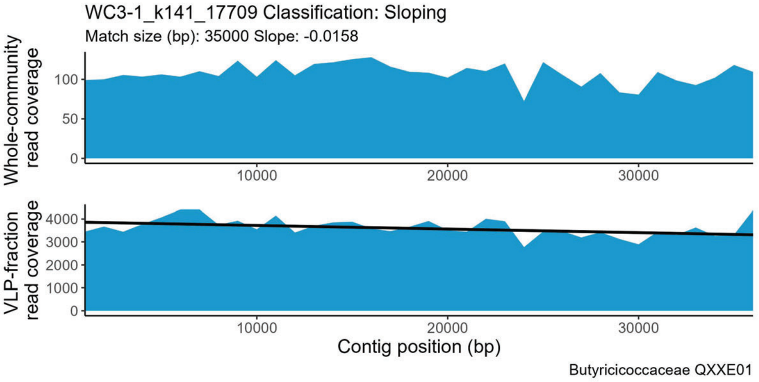
WC3-1_k141_17709 Classification: Sloping
Match size (bp): 35000 Slope: -0.0158
Whole-community read coverage
VLP-fraction read coverage
Contig position (bp)
Butyricicoccaceae QXXE01

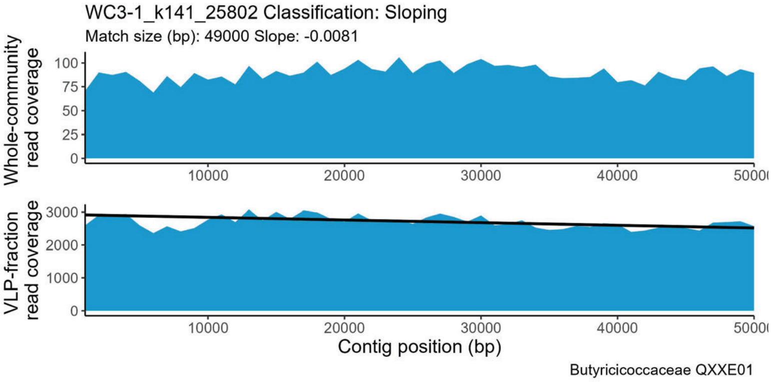
WC3-1_k141_25802 Classification: Sloping
Match size (bp): 49000 Slope: -0.0081
Whole-community read coverage
VLP-fraction read coverage
Contig position (bp)
Butyricicoccaceae QXXE01

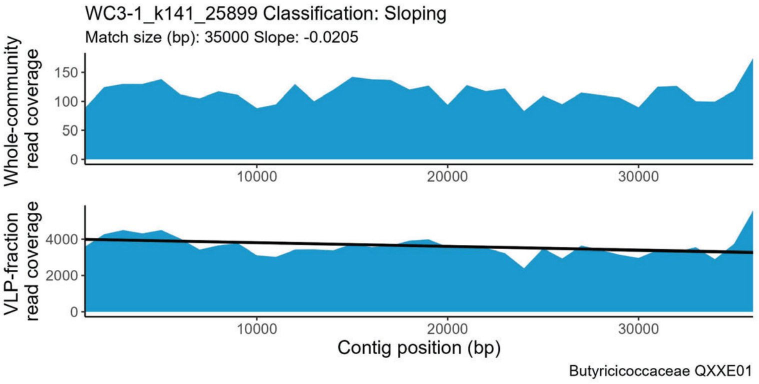
WC3-1_k141_25899 Classification: Sloping
Match size (bp): 35000 Slope: -0.0205
Whole-community read coverage
VLP-fraction read coverage
Contig position (bp)
Butyricicoccaceae QXXE01

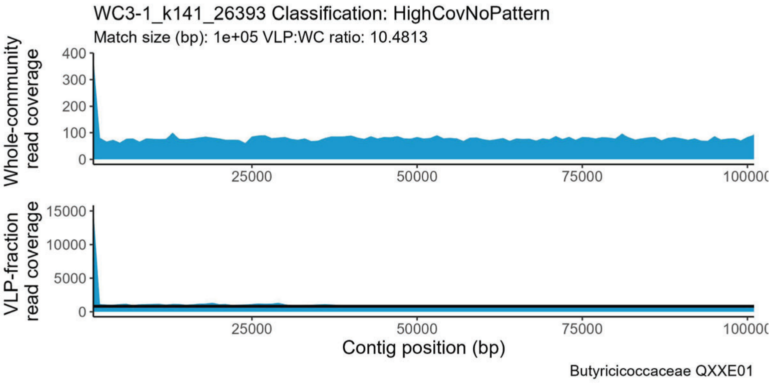
WC3-1_k141_26393 Classification: HighCovNoPattern
Match size (bp): 1e+05 VLP:WC ratio: 10.4813
Whole-community read coverage
VLP-fraction read coverage
Contig position (bp)
Butyricicoccaceae QXXE01

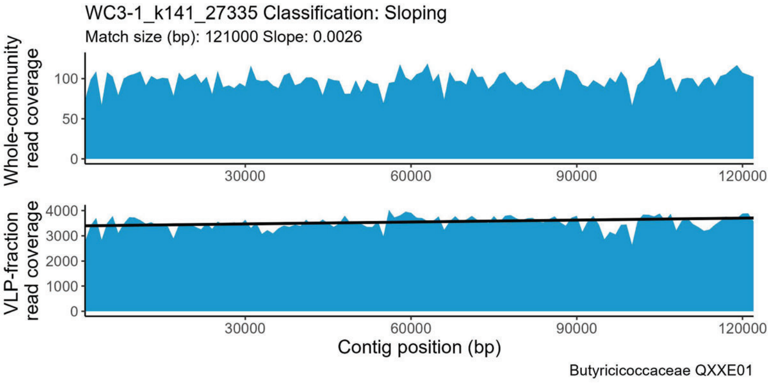
WC3-1_k141_27335 Classification: Sloping
Match size (bp): 121000 Slope: 0.0026
Whole-community read coverage
VLP-fraction read coverage
Contig position (bp)
Butyricicoccaceae QXXE01

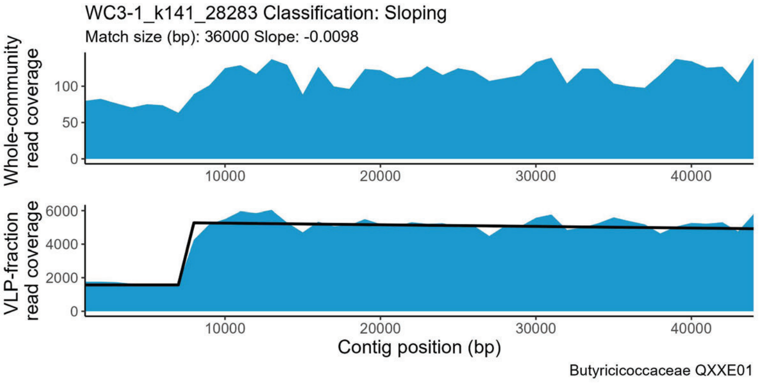
WC3-1_k141_28283 Classification: Sloping
Match size (bp): 36000 Slope: -0.0098
Whole-community read coverage
VLP-fraction read coverage
Contig position (bp)
Butyricicoccaceae QXXE01

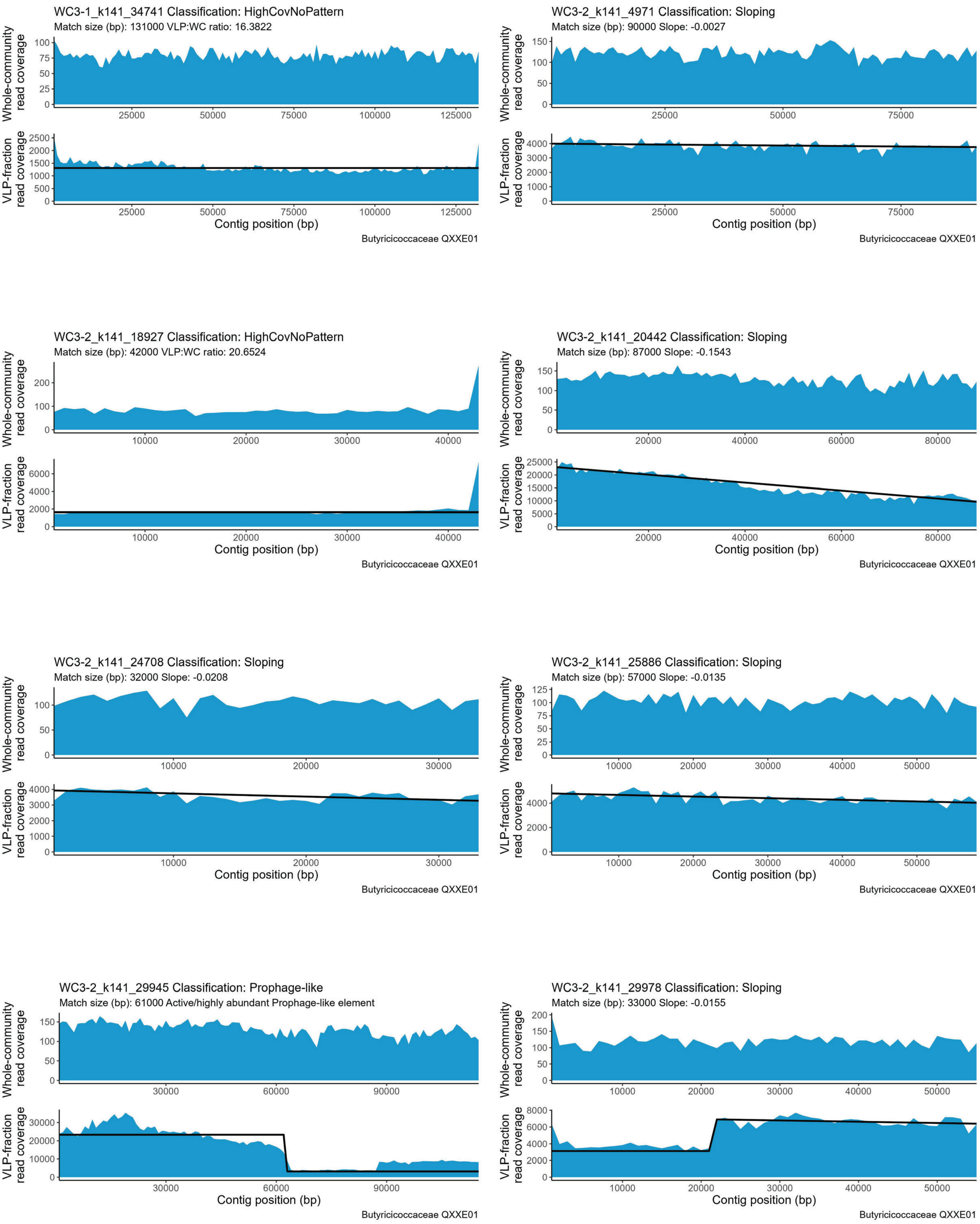
WC3-1_k141_34741 Classification: HighCovNoPattern
Match size (bp): 131000 VLP:WC ratio: 16.3822
Whole-community read coverage
VLP-fraction read coverage
Contig position (bp)
Butyricicoccaceae QXXE01
WC3-2_k141_4971 Classification: Sloping
Match size (bp): 90000 Slope: -0.0027
Whole-community read coverage
VLP-fraction read coverage
Contig position (bp)
Butyricicoccaceae QXXE01
WC3-2_k141_18927 Classification: HighCovNoPattern
Match size (bp): 42000 VLP:WC ratio: 20.6524
Whole-community read coverage
VLP-fraction read coverage
Contig position (bp)
Butyricicoccaceae QXXE01
WC3-2_k141_20442 Classification: Sloping
Match size (bp): 87000 Slope: -0.1543
Whole-community read coverage
VLP-fraction read coverage
Contig position (bp)
Butyricicoccaceae QXXE01
WC3-2_k141_24708 Classification: Sloping
Match size (bp): 32000 Slope: -0.0208
Whole-community read coverage
VLP-fraction read coverage
Contig position (bp)
Butyricicoccaceae QXXE01
WC3-2_k141_25886 Classification: Sloping
Match size (bp): 57000 Slope: -0.0135
Whole-community read coverage
VLP-fraction read coverage
Contig position (bp)
Butyricicoccaceae QXXE01
WC3-2_k141_29945 Classification: Prophage-like
Match size (bp): 61000 Active/highly abundant Prophage-like element
Whole-community read coverage
VLP-fraction read coverage
Contig position (bp)
Butyricicoccaceae QXXE01
WC3-2_k141_29978 Classification: Sloping
Match size (bp): 33000 Slope: -0.0155
Whole-community read coverage
VLP-fraction read coverage
Contig position (bp)
Butyricicoccaceae QXXE01

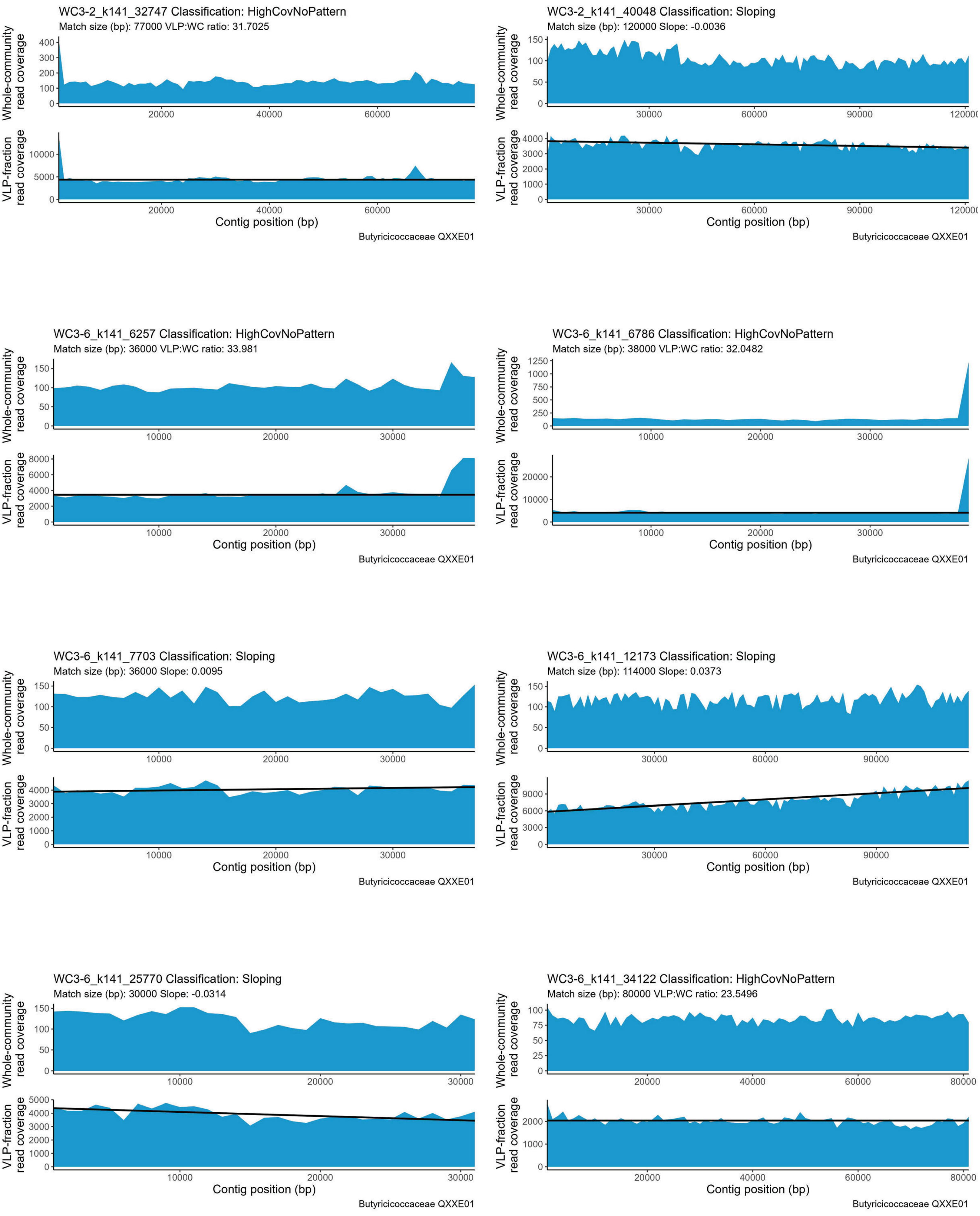
WC3-2_k141_32747 Classification: HighCovNoPattern
Match size (bp): 77000 VLP:WC ratio: 31.7025
WC3-2_k141_40048 Classification: Sloping
Match size (bp): 120000 Slope: -0.0036
WC3-6_k141_6257 Classification: HighCovNoPattern
Match size (bp): 36000 VLP:WC ratio: 33.981
WC3-6_k141_6786 Classification: HighCovNoPattern
Match size (bp): 38000 VLP:WC ratio: 32.0482
WC3-6_k141_7703 Classification: Sloping
Match size (bp): 36000 Slope: 0.0095
WC3-6_k141_12173 Classification: Sloping
Match size (bp): 114000 Slope: 0.0373
WC3-6_k141_25770 Classification: Sloping
Match size (bp): 30000 Slope: -0.0314
WC3-6_k141_34122 Classification: HighCovNoPattern
Match size (bp): 80000 VLP:WC ratio: 23.5496
Whole-community read coverage
VLP-fraction read coverage
Contig position (bp)
Butyricicoccaceae QXXE01

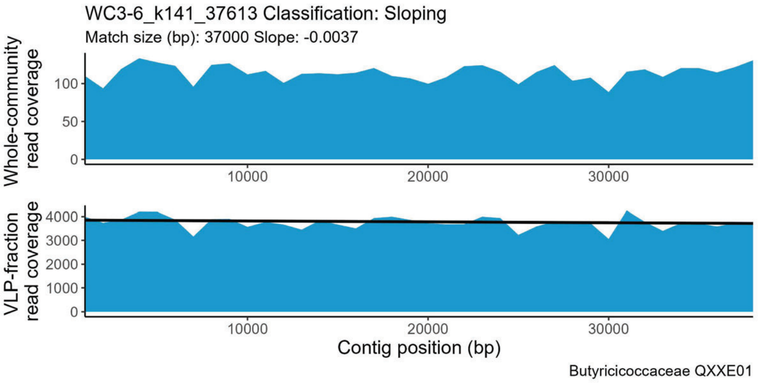
WC3-6_k141_37613 Classification: Sloping
Match size (bp): 37000 Slope: -0.0037
Whole-community read coverage
VLP-fraction read coverage
Contig position (bp)
Butyricicoccaceae QXXE01

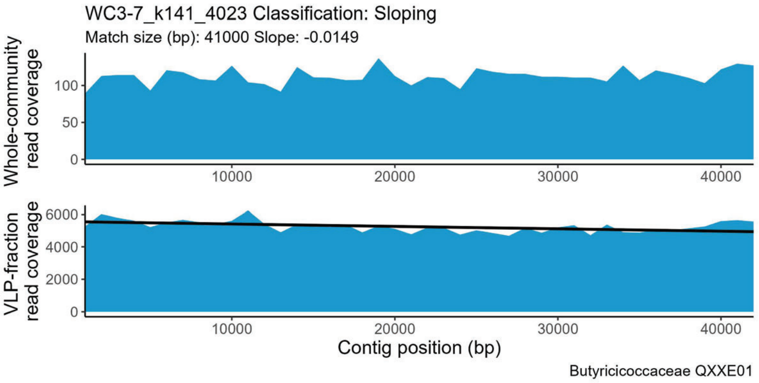
WC3-7_k141_4023 Classification: Sloping
Match size (bp): 41000 Slope: -0.0149
Whole-community read coverage
VLP-fraction read coverage
Contig position (bp)
Butyricicoccaceae QXXE01

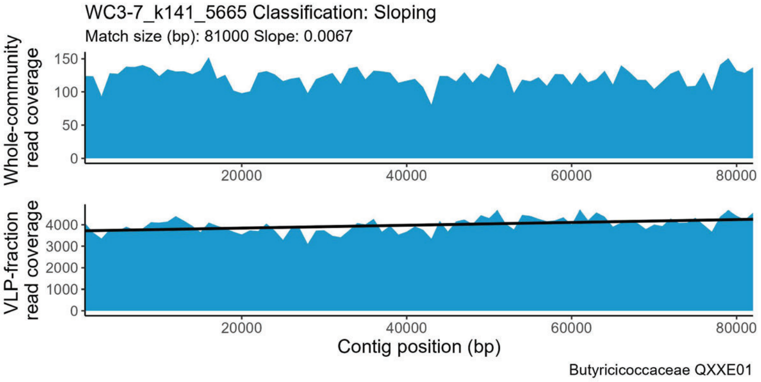
WC3-7_k141_5665 Classification: Sloping
Match size (bp): 81000 Slope: 0.0067
Whole-community read coverage
VLP-fraction read coverage
Contig position (bp)
Butyricicoccaceae QXXE01

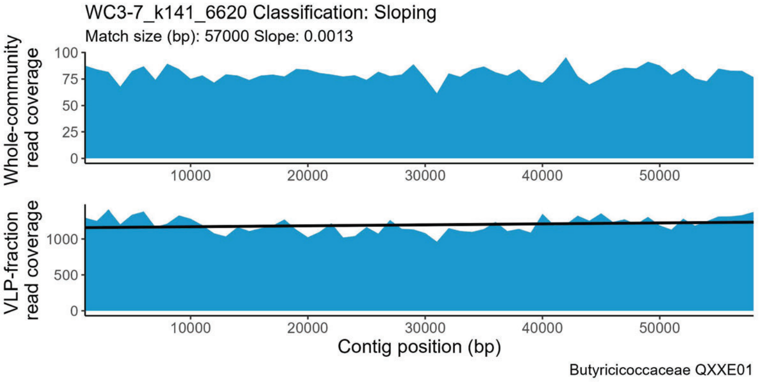
WC3-7_k141_6620 Classification: Sloping
Match size (bp): 57000 Slope: 0.0013
Whole-community read coverage
VLP-fraction read coverage
Contig position (bp)
Butyricicoccaceae QXXE01

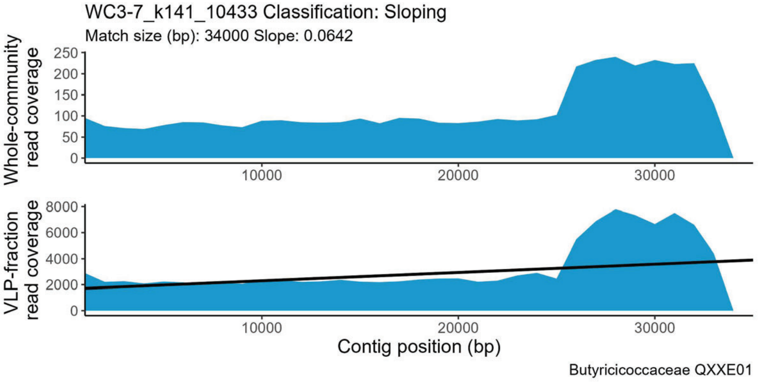
WC3-7_k141_10433 Classification: Sloping
Match size (bp): 34000 Slope: 0.0642
Whole-community read coverage
VLP-fraction read coverage
Contig position (bp)
Butyricicoccaceae QXXE01

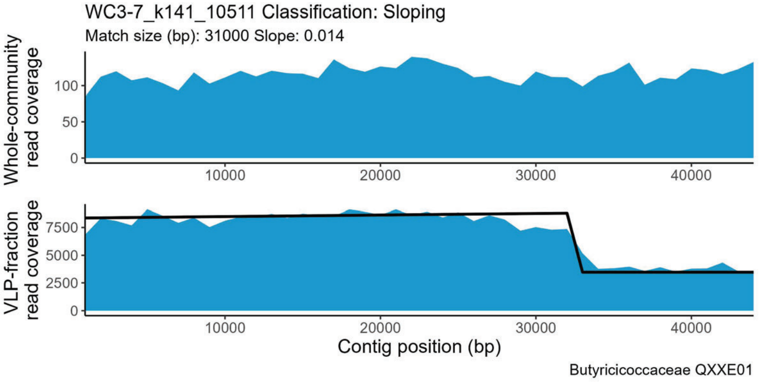
WC3-7_k141_10511 Classification: Sloping
Match size (bp): 31000 Slope: 0.014
Whole-community read coverage
VLP-fraction read coverage
Contig position (bp)
Butyricicoccaceae QXXE01

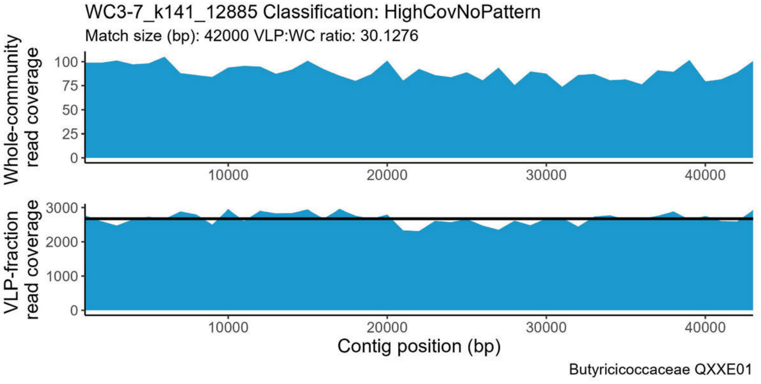
WC3-7_k141_12885 Classification: HighCovNoPattern
Match size (bp): 42000 VLP:WC ratio: 30.1276
Whole-community read coverage
VLP-fraction read coverage
Contig position (bp)
Butyricicoccaceae QXXE01

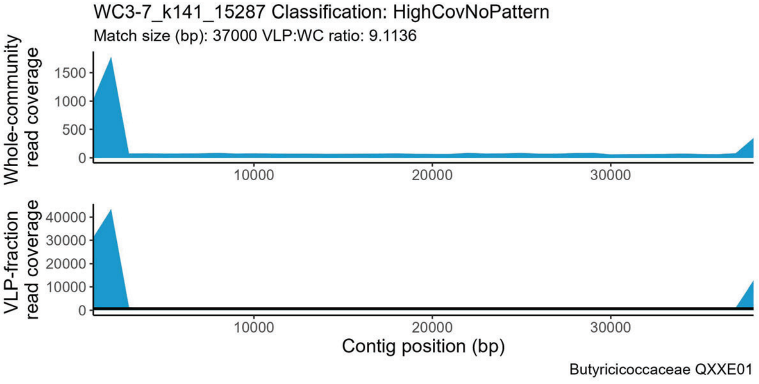
WC3-7_k141_15287 Classification: HighCovNoPattern
Match size (bp): 37000 VLP:WC ratio: 9.1136
Whole-community read coverage
VLP-fraction read coverage
Contig position (bp)
Butyricicoccaceae QXXE01

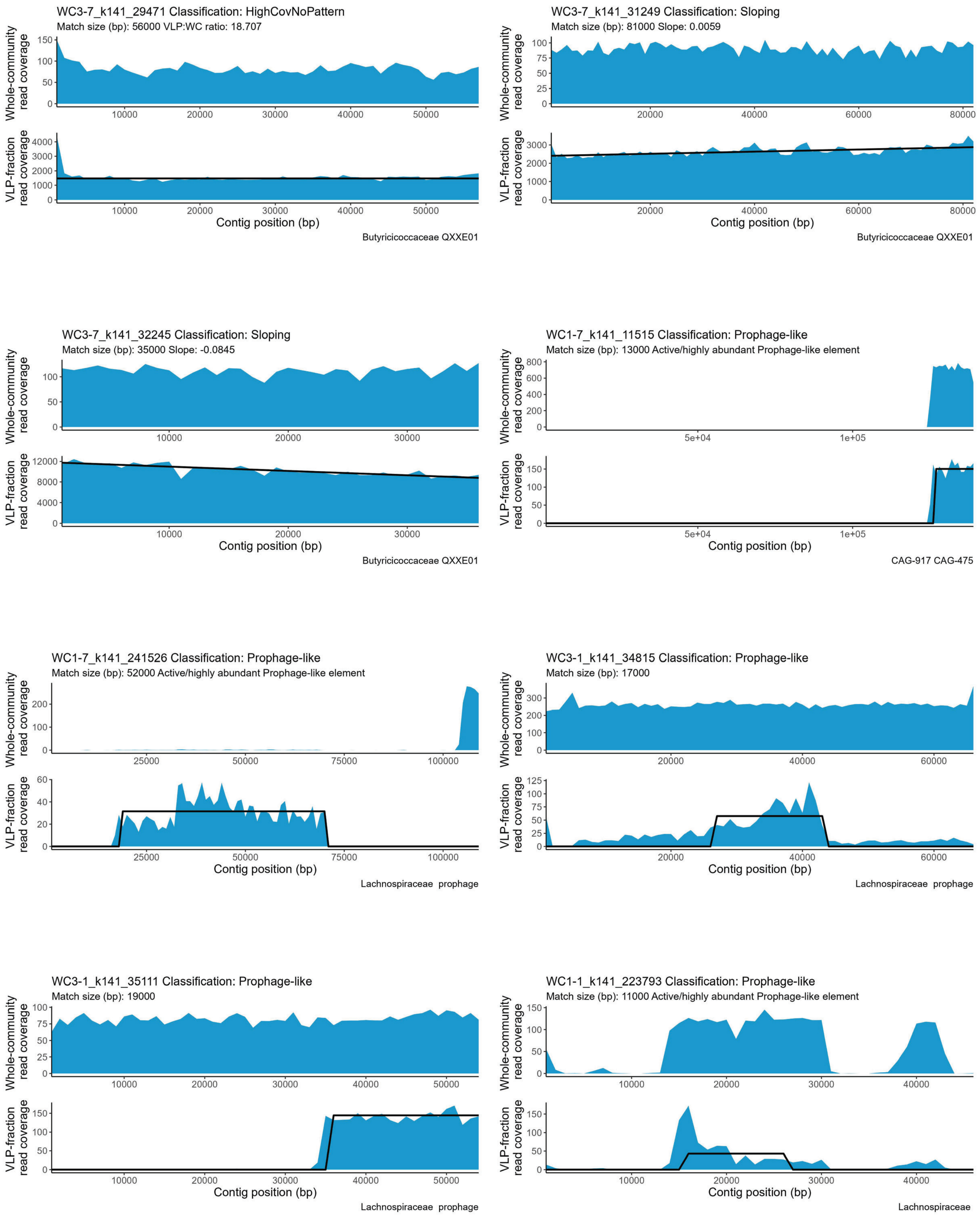
WC3-7_k141_29471 Classification: HighCovNoPattern
Match size (bp): 56000 VLP:WC ratio: 18.707
Whole-community read coverage
VLP-fraction read coverage
Contig position (bp)
Butyricicoccaceae QXXE01
WC3-7_k141_31249 Classification: Sloping
Match size (bp): 81000 Slope: 0.0059
Whole-community read coverage
VLP-fraction read coverage
Contig position (bp)
Butyricicoccaceae QXXE01
WC3-7_k141_32245 Classification: Sloping
Match size (bp): 35000 Slope: -0.0845
Whole-community read coverage
VLP-fraction read coverage
Contig position (bp)
Butyricicoccaceae QXXE01
WC1-7_k141_11515 Classification: Prophage-like
Match size (bp): 13000 Active/highly abundant Prophage-like element
Whole-community read coverage
VLP-fraction read coverage
Contig position (bp)
CAG-917 CAG-475
WC1-7_k141_241526 Classification: Prophage-like
Match size (bp): 52000 Active/highly abundant Prophage-like element
Whole-community read coverage
VLP-fraction read coverage
Contig position (bp)
Lachnospiraceae prophage
WC3-1_k141_34815 Classification: Prophage-like
Match size (bp): 17000
Whole-community read coverage
VLP-fraction read coverage
Contig position (bp)
Lachnospiraceae prophage
WC3-1_k141_35111 Classification: Prophage-like
Match size (bp): 19000
Whole-community read coverage
VLP-fraction read coverage
Contig position (bp)
Lachnospiraceae prophage
WC1-1_k141_223793 Classification: Prophage-like
Match size (bp): 11000 Active/highly abundant Prophage-like element
Whole-community read coverage
VLP-fraction read coverage
Contig position (bp)
Lachnospiraceae

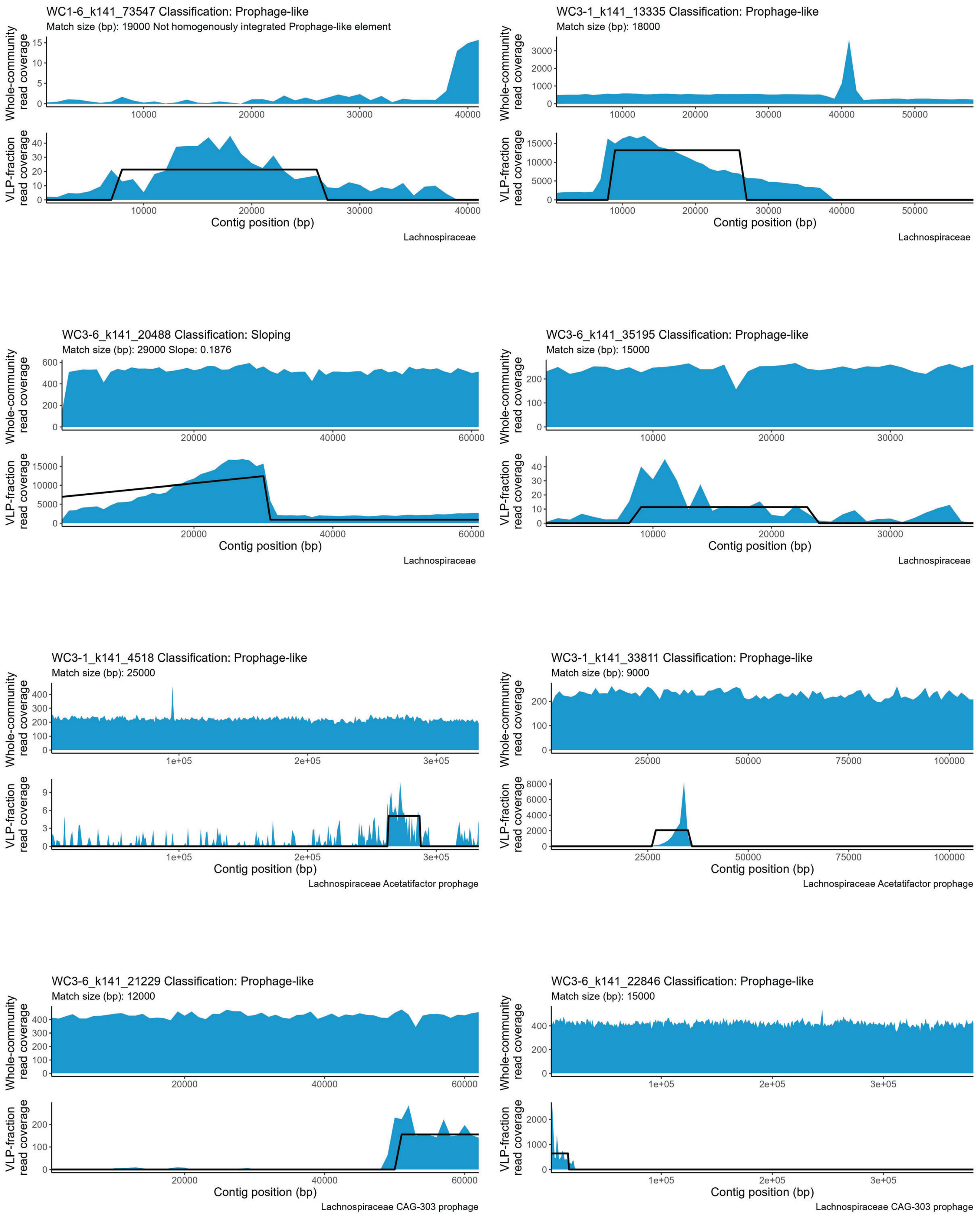
WC1-6_k141_73547 Classification: Prophage-like
Match size (bp): 19000 Not homogenously integrated Prophage-like element
Whole-community read coverage
VLP-fraction read coverage
Contig position (bp)
Lachnospiraceae
WC3-1_k141_13335 Classification: Prophage-like
Match size (bp): 18000
Whole-community read coverage
VLP-fraction read coverage
Contig position (bp)
Lachnospiraceae
WC3-6_k141_20488 Classification: Sloping
Match size (bp): 29000 Slope: 0.1876
Whole-community read coverage
VLP-fraction read coverage
Contig position (bp)
Lachnospiraceae
WC3-6_k141_35195 Classification: Prophage-like
Match size (bp): 15000
Whole-community read coverage
VLP-fraction read coverage
Contig position (bp)
Lachnospiraceae
WC3-1_k141_4518 Classification: Prophage-like
Match size (bp): 25000
Whole-community read coverage
VLP-fraction read coverage
Contig position (bp)
Lachnospiraceae Acetatifactor prophage
WC3-1_k141_33811 Classification: Prophage-like
Match size (bp): 9000
Whole-community read coverage
VLP-fraction read coverage
Contig position (bp)
Lachnospiraceae Acetatifactor prophage
WC3-6_k141_21229 Classification: Prophage-like
Match size (bp): 12000
Whole-community read coverage
VLP-fraction read coverage
Contig position (bp)
Lachnospiraceae CAG-303 prophage
WC3-6_k141_22846 Classification: Prophage-like
Match size (bp): 15000
Whole-community read coverage
VLP-fraction read coverage
Contig position (bp)
Lachnospiraceae CAG-303 prophage

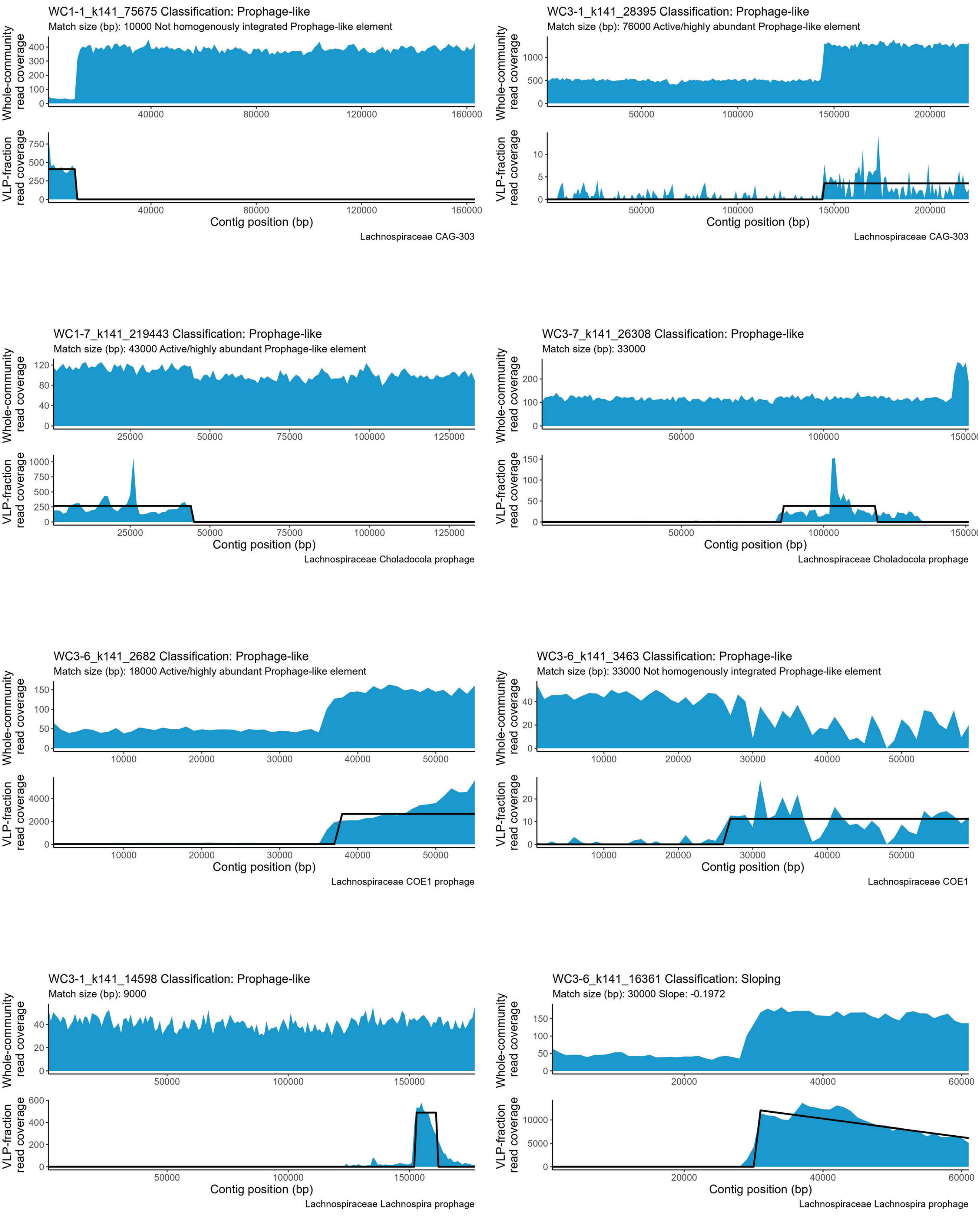
WC1-1_k141_75675 Classification: Prophage-like
Match size (bp): 10000 Not homogenously integrated Prophage-like element
Whole-community read coverage
VLP-fraction read coverage
Contig position (bp)
Lachnospiraceae CAG-303
WC3-1_k141_28395 Classification: Prophage-like
Match size (bp): 76000 Active/highly abundant Prophage-like element
Whole-community read coverage
VLP-fraction read coverage
Contig position (bp)
Lachnospiraceae CAG-303
WC1-7_k141_219443 Classification: Prophage-like
Match size (bp): 43000 Active/highly abundant Prophage-like element
Whole-community read coverage
VLP-fraction read coverage
Contig position (bp)
Lachnospiraceae Choladocola prophage
WC3-7_k141_26308 Classification: Prophage-like
Match size (bp): 33000
Whole-community read coverage
VLP-fraction read coverage
Contig position (bp)
Lachnospiraceae Choladocola prophage
WC3-6_k141_2682 Classification: Prophage-like
Match size (bp): 18000 Active/highly abundant Prophage-like element
Whole-community read coverage
VLP-fraction read coverage
Contig position (bp)
Lachnospiraceae COE1 prophage
WC3-6_k141_3463 Classification: Prophage-like
Match size (bp): 33000 Not homogenously integrated Prophage-like element
Whole-community read coverage
VLP-fraction read coverage
Contig position (bp)
Lachnospiraceae COE1
WC3-1_k141_14598 Classification: Prophage-like
Match size (bp): 9000
Whole-community read coverage
VLP-fraction read coverage
Contig position (bp)
Lachnospiraceae Lachnospira prophage
WC3-6_k141_16361 Classification: Sloping
Match size (bp): 30000 Slope: -0.1972
Whole-community read coverage
VLP-fraction read coverage
Contig position (bp)
Lachnospiraceae Lachnospira prophage

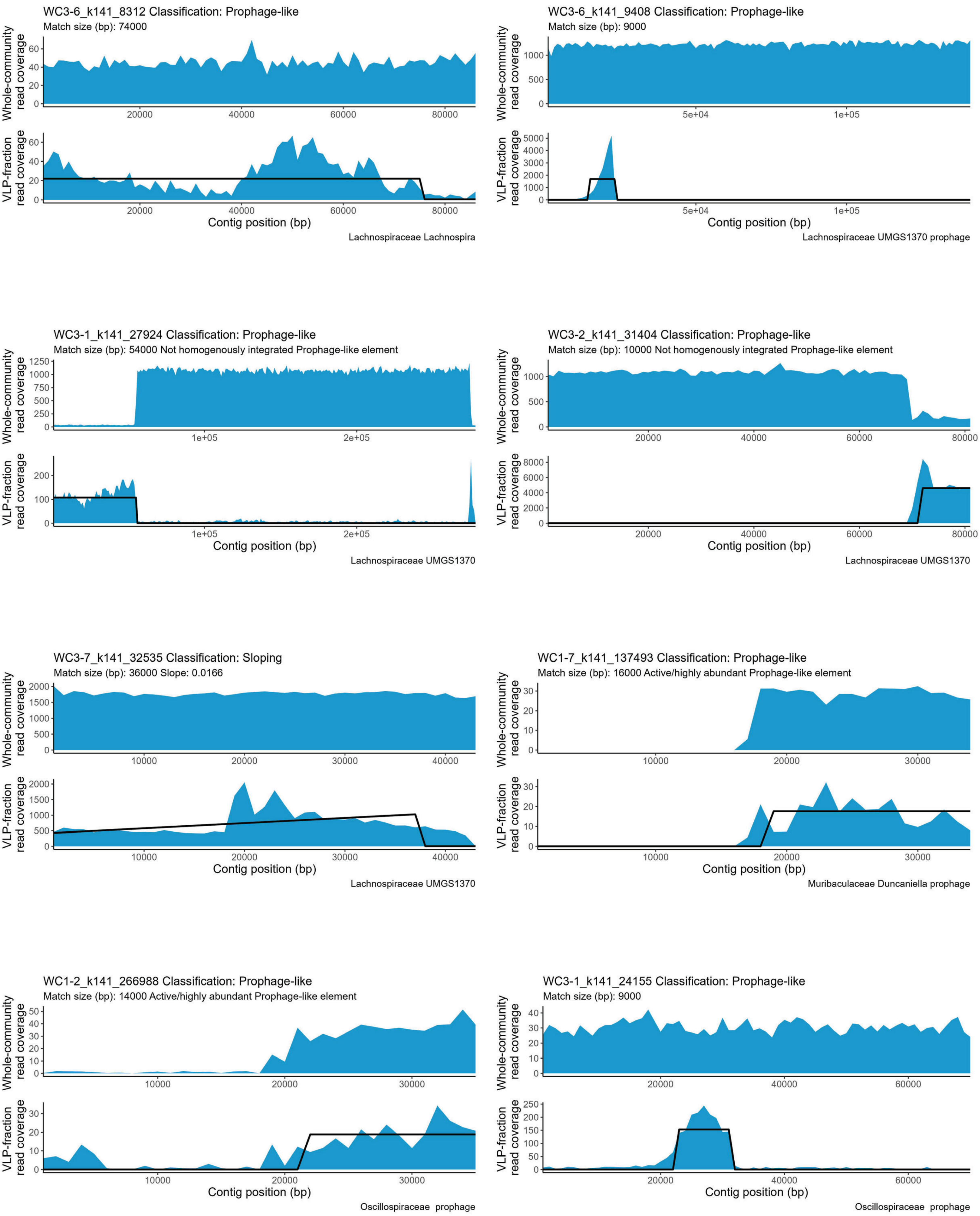
WC3-6_k141_8312 Classification: Prophage-like
Match size (bp): 74000
Lachnospiraceae Lachnospira
WC3-6_k141_9408 Classification: Prophage-like
Match size (bp): 9000
Lachnospiraceae UMGS1370 prophage
WC3-1_k141_27924 Classification: Prophage-like
Match size (bp): 54000 Not homogenously integrated Prophage-like element
Lachnospiraceae UMGS1370
WC3-2_k141_31404 Classification: Prophage-like
Match size (bp): 10000 Not homogenously integrated Prophage-like element
Lachnospiraceae UMGS1370
WC3-7_k141_32535 Classification: Sloping
Match size (bp): 36000 Slope: 0.0166
Lachnospiraceae UMGS1370
WC1-7_k141_137493 Classification: Prophage-like
Match size (bp): 16000 Active/highly abundant Prophage-like element
Muribaculaceae Duncaniella prophage
WC1-2_k141_266988 Classification: Prophage-like
Match size (bp): 14000 Active/highly abundant Prophage-like element
Oscillospiraceae prophage
WC3-1_k141_24155 Classification: Prophage-like
Match size (bp): 9000
Oscillospiraceae prophage
Whole-community read coverage
VLP-fraction read coverage
Contig position (bp)

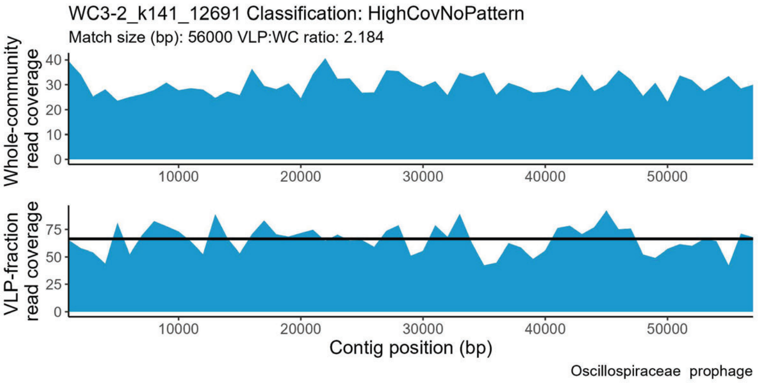
WC3-2_k141_12691 Classification: HighCovNoPattern
Match size (bp): 56000 VLP:WC ratio: 2.184
Whole-community read coverage
VLP-fraction read coverage
Contig position (bp)
Oscillospiraceae prophage

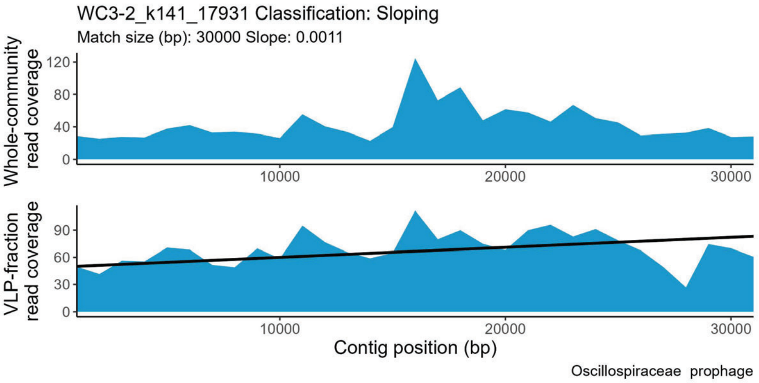
WC3-2_k141_17931 Classification: Sloping
Match size (bp): 30000 Slope: 0.0011
Whole-community read coverage
VLP-fraction read coverage
Contig position (bp)
Oscillospiraceae prophage

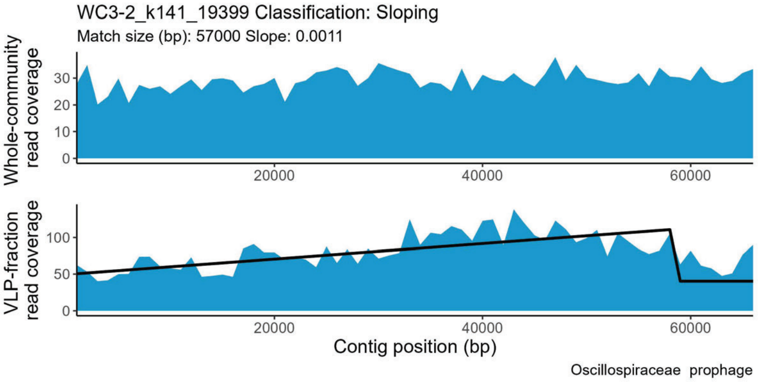
WC3-2_k141_19399 Classification: Sloping
Match size (bp): 57000 Slope: 0.0011
Whole-community read coverage
VLP-fraction read coverage
Contig position (bp)
Oscillospiraceae prophage

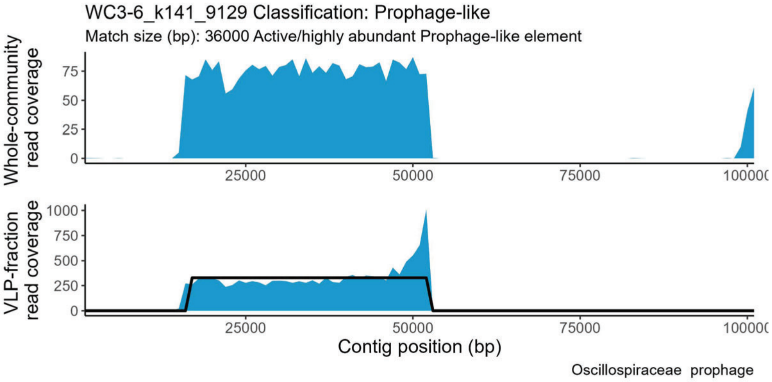
WC3-6_k141_9129 Classification: Prophage-like
Match size (bp): 36000 Active/highly abundant Prophage-like element
Whole-community read coverage
VLP-fraction read coverage
Contig position (bp)
Oscillospiraceae prophage

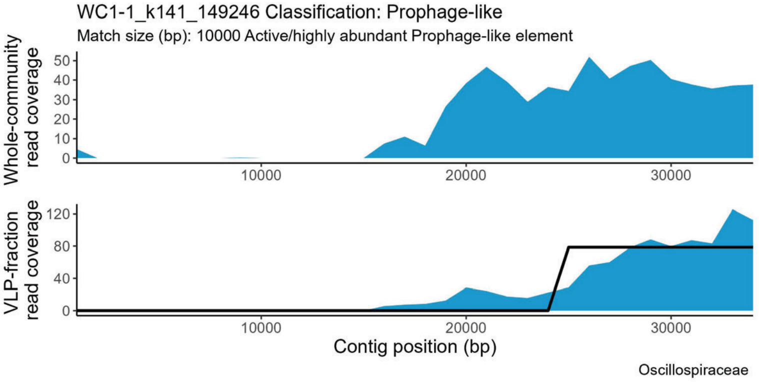
WC1-1_k141_149246 Classification: Prophage-like
Match size (bp): 10000 Active/highly abundant Prophage-like element
Whole-community read coverage
VLP-fraction read coverage
Contig position (bp)
Oscillospiraceae

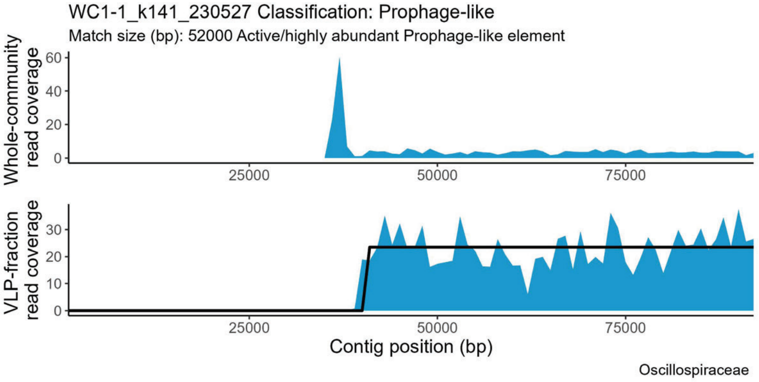
WC1-1_k141_230527 Classification: Prophage-like
Match size (bp): 52000 Active/highly abundant Prophage-like element
Whole-community read coverage
VLP-fraction read coverage
Contig position (bp)
Oscillospiraceae

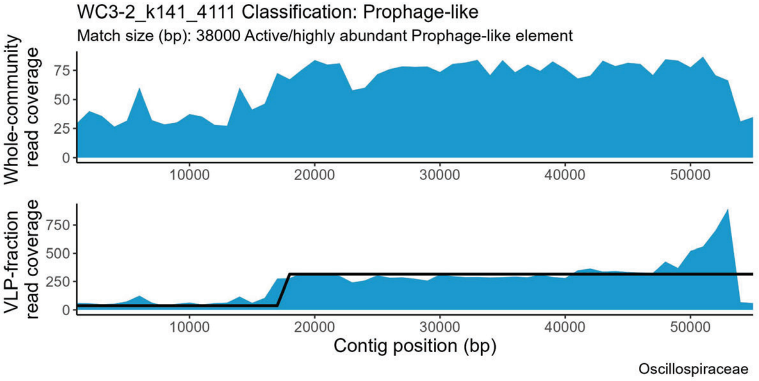
WC3-2_k141_4111 Classification: Prophage-like
Match size (bp): 38000 Active/highly abundant Prophage-like element
Whole-community read coverage
VLP-fraction read coverage
Contig position (bp)
Oscillospiraceae

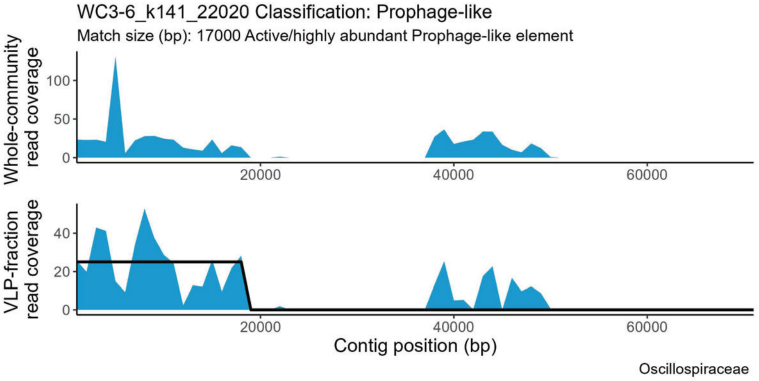
WC3-6_k141_22020 Classification: Prophage-like
Match size (bp): 17000 Active/highly abundant Prophage-like element
Whole-community read coverage
VLP-fraction read coverage
Contig position (bp)
Oscillospiraceae

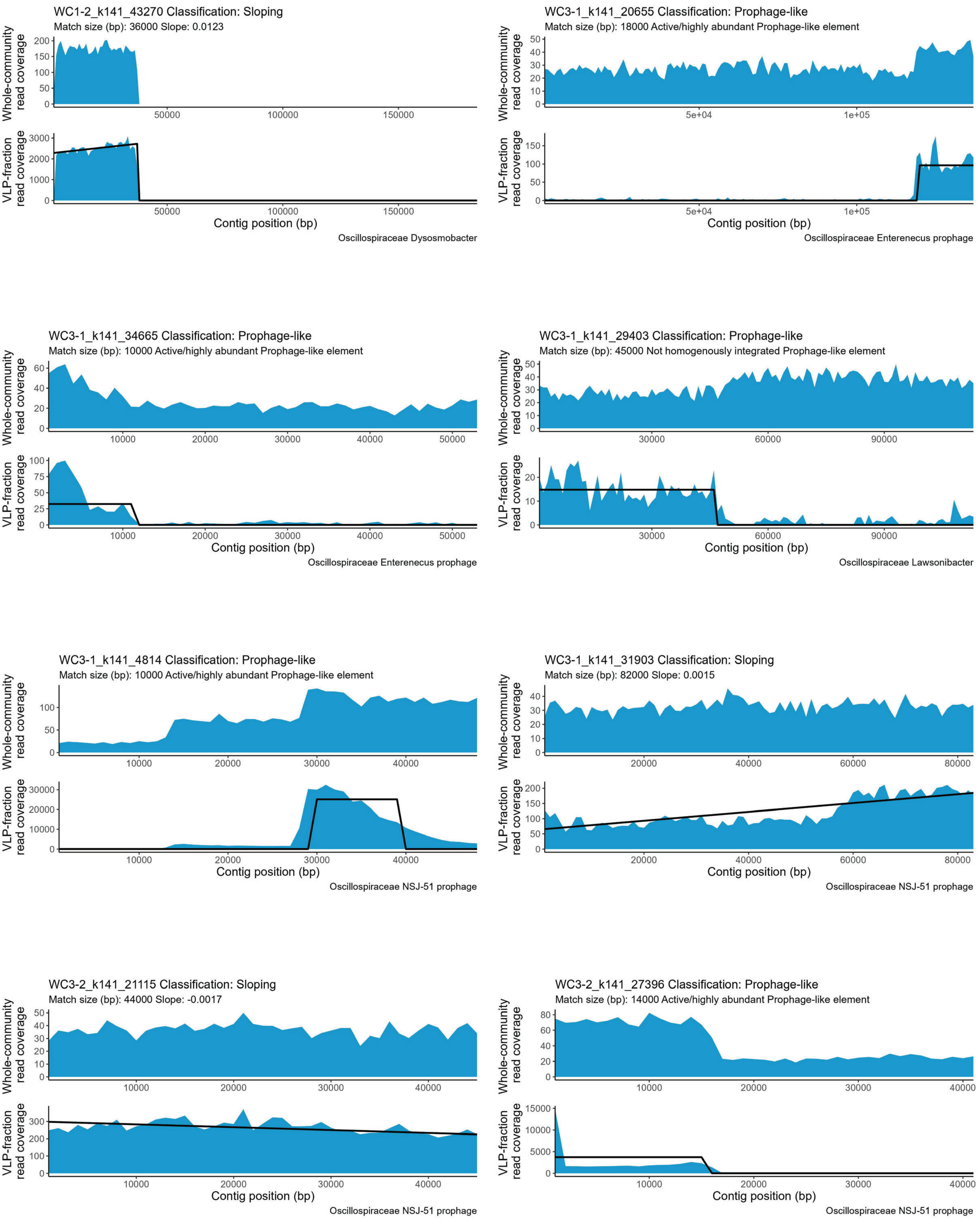
WC1-2_k141_43270 Classification: Sloping
Match size (bp): 36000 Slope: 0.0123
Whole-community read coverage
VLP-fraction read coverage
Contig position (bp)
Oscillospiraceae Dysosmobacter
WC3-1_k141_20655 Classification: Prophage-like
Match size (bp): 18000 Active/highly abundant Prophage-like element
Whole-community read coverage
VLP-fraction read coverage
Contig position (bp)
Oscillospiraceae Enterenecus prophage
WC3-1_k141_34665 Classification: Prophage-like
Match size (bp): 10000 Active/highly abundant Prophage-like element
Whole-community read coverage
VLP-fraction read coverage
Contig position (bp)
Oscillospiraceae Enterenecus prophage
WC3-1_k141_29403 Classification: Prophage-like
Match size (bp): 45000 Not homogenously integrated Prophage-like element
Whole-community read coverage
VLP-fraction read coverage
Contig position (bp)
Oscillospiraceae Lawsonibacter
WC3-1_k141_4814 Classification: Prophage-like
Match size (bp): 10000 Active/highly abundant Prophage-like element
Whole-community read coverage
VLP-fraction read coverage
Contig position (bp)
Oscillospiraceae NSJ-51 prophage
WC3-1_k141_31903 Classification: Sloping
Match size (bp): 82000 Slope: 0.0015
Whole-community read coverage
VLP-fraction read coverage
Contig position (bp)
Oscillospiraceae NSJ-51 prophage
WC3-2_k141_21115 Classification: Sloping
Match size (bp): 44000 Slope: -0.0017
Whole-community read coverage
VLP-fraction read coverage
Contig position (bp)
Oscillospiraceae NSJ-51 prophage
WC3-2_k141_27396 Classification: Prophage-like
Match size (bp): 14000 Active/highly abundant Prophage-like element
Whole-community read coverage
VLP-fraction read coverage
Contig position (bp)
Oscillospiraceae NSJ-51 prophage

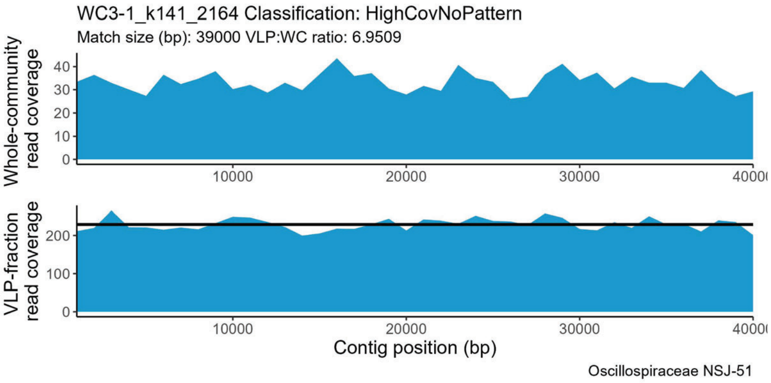
WC3-1_k141_2164 Classification: HighCovNoPattern
Match size (bp): 39000 VLP:WC ratio: 6.9509
Whole-community read coverage
VLP-fraction read coverage
Contig position (bp)
Oscillospiraceae NSJ-51

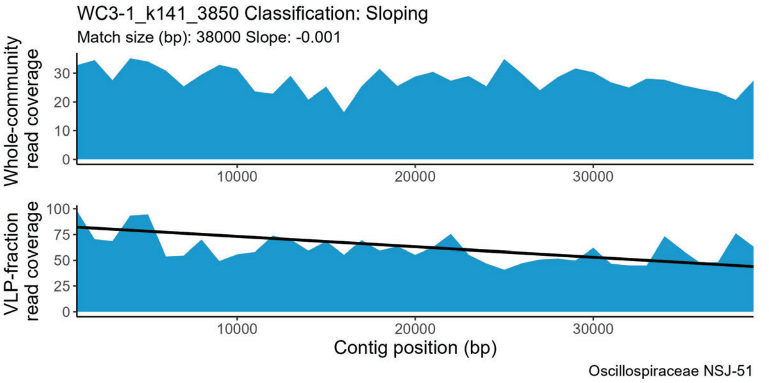
WC3-1_k141_3850 Classification: Sloping
Match size (bp): 38000 Slope: -0.001
Whole-community read coverage
VLP-fraction read coverage
Contig position (bp)
Oscillospiraceae NSJ-51

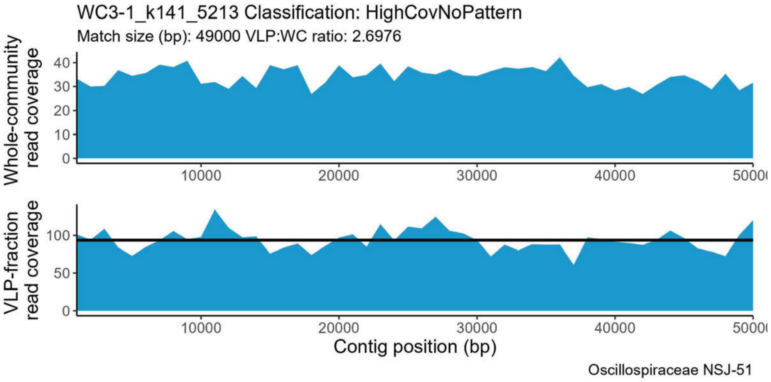
WC3-1_k141_5213 Classification: HighCovNoPattern
Match size (bp): 49000 VLP:WC ratio: 2.6976
Whole-community read coverage
VLP-fraction read coverage
Contig position (bp)
Oscillospiraceae NSJ-51

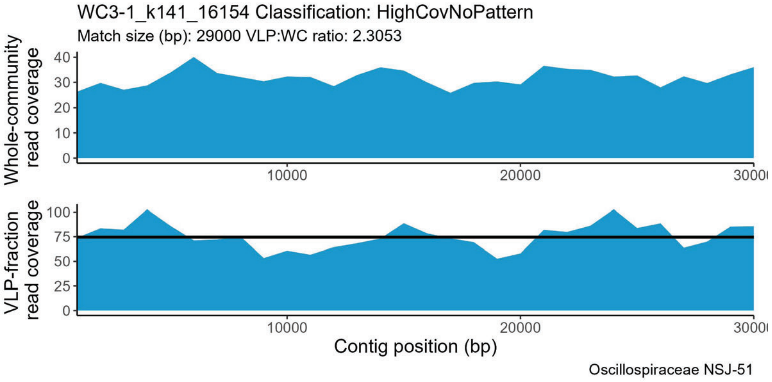
WC3-1_k141_16154 Classification: HighCovNoPattern
Match size (bp): 29000 VLP:WC ratio: 2.3053
Whole-community read coverage
VLP-fraction read coverage
Contig position (bp)
Oscillospiraceae NSJ-51

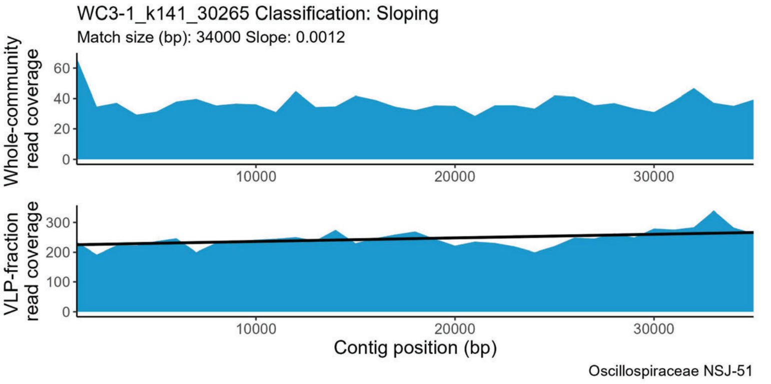
WC3-1_k141_30265 Classification: Sloping
Match size (bp): 34000 Slope: 0.0012
Whole-community read coverage
VLP-fraction read coverage
Contig position (bp)
Oscillospiraceae NSJ-51

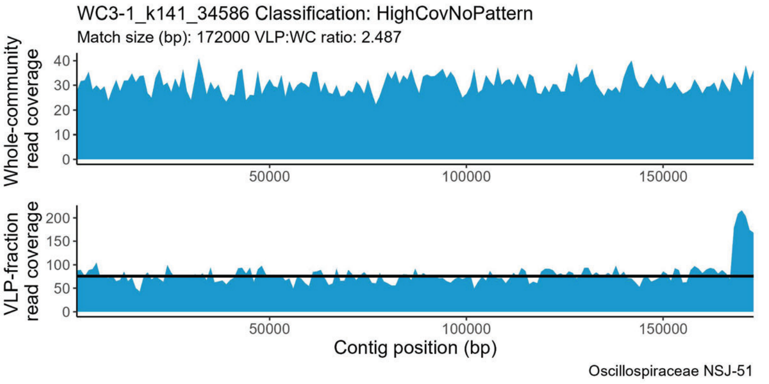
WC3-1_k141_34586 Classification: HighCovNoPattern
Match size (bp): 172000 VLP:WC ratio: 2.487
Whole-community read coverage
VLP-fraction read coverage
Contig position (bp)
Oscillospiraceae NSJ-51

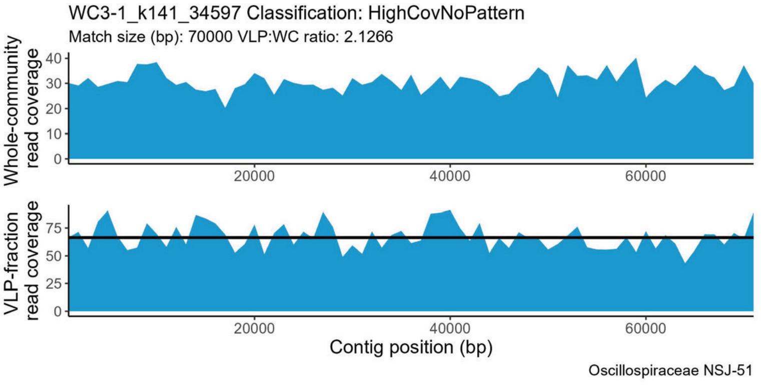
WC3-1_k141_34597 Classification: HighCovNoPattern
Match size (bp): 70000 VLP:WC ratio: 2.1266
Whole-community read coverage
VLP-fraction read coverage
Contig position (bp)
Oscillospiraceae NSJ-51

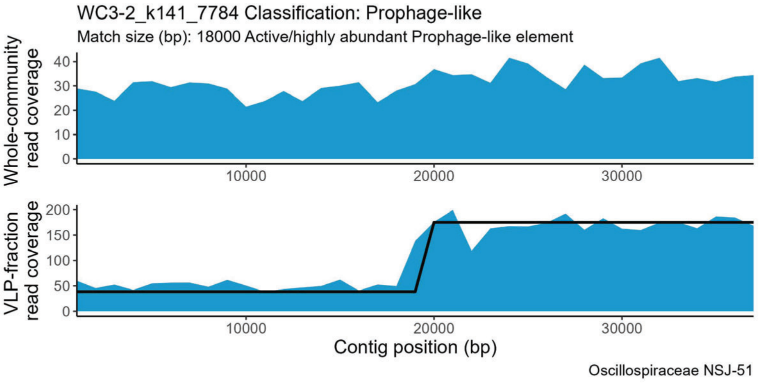
WC3-2_k141_7784 Classification: Prophage-like
Match size (bp): 18000 Active/highly abundant Prophage-like element
Whole-community read coverage
VLP-fraction read coverage
Contig position (bp)
Oscillospiraceae NSJ-51

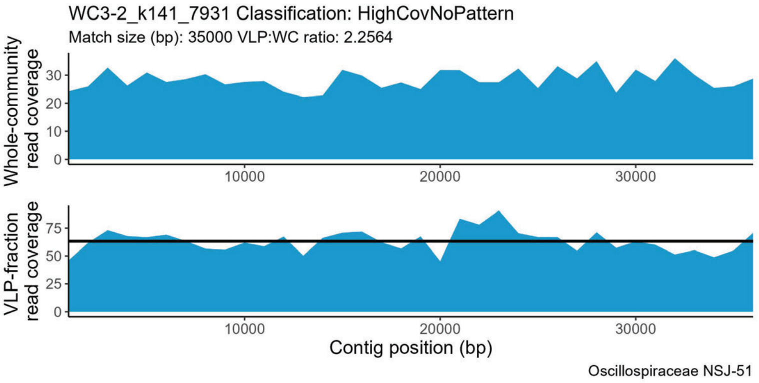
WC3-2_k141_7931 Classification: HighCovNoPattern
Match size (bp): 35000 VLP:WC ratio: 2.2564
Whole-community read coverage
VLP-fraction read coverage
Contig position (bp)
Oscillospiraceae NSJ-51

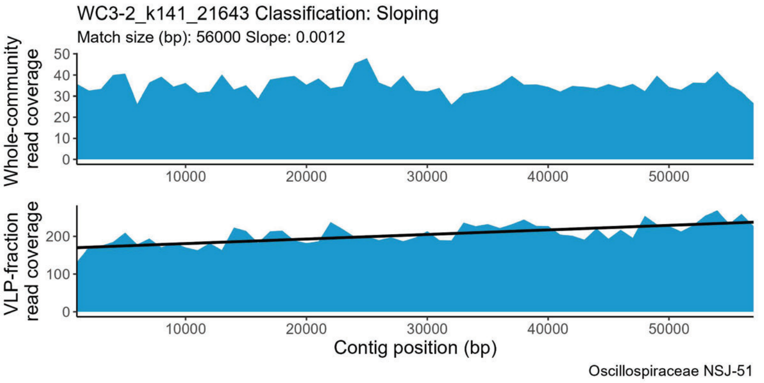
WC3-2_k141_21643 Classification: Sloping
Match size (bp): 56000 Slope: 0.0012
Whole-community read coverage
VLP-fraction read coverage
Contig position (bp)
Oscillospiraceae NSJ-51

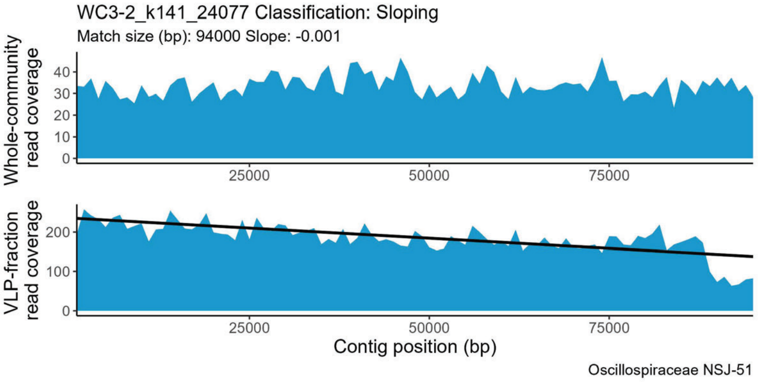
WC3-2_k141_24077 Classification: Sloping
Match size (bp): 94000 Slope: -0.001
Whole-community read coverage
VLP-fraction read coverage
Contig position (bp)
Oscillospiraceae NSJ-51

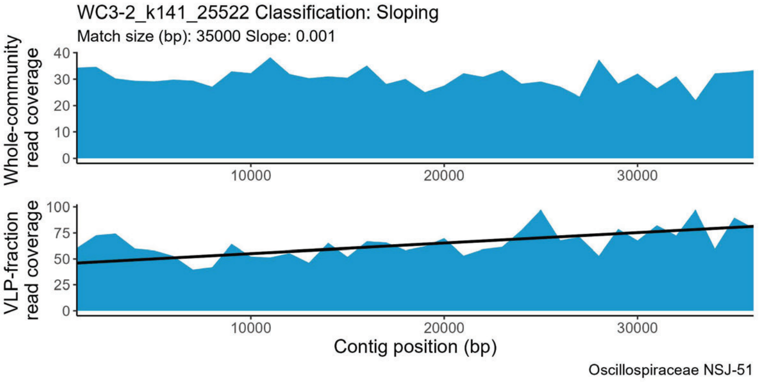
WC3-2_k141_25522 Classification: Sloping
Match size (bp): 35000 Slope: 0.001
Whole-community read coverage
VLP-fraction read coverage
Contig position (bp)
Oscillospiraceae NSJ-51

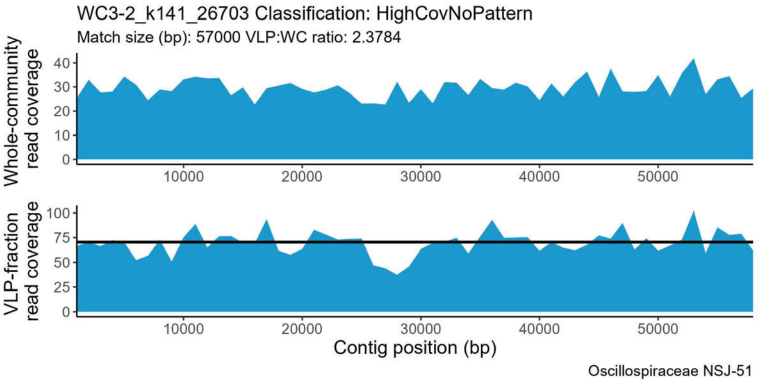
WC3-2_k141_26703 Classification: HighCovNoPattern
Match size (bp): 57000 VLP:WC ratio: 2.3784
Whole-community read coverage
VLP-fraction read coverage
Contig position (bp)
Oscillospiraceae NSJ-51

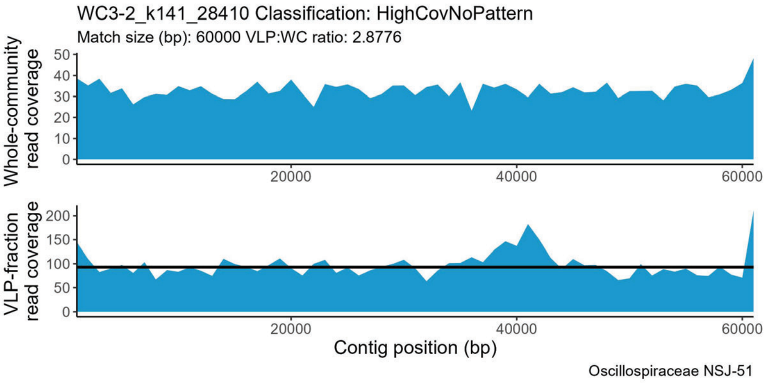
WC3-2_k141_28410 Classification: HighCovNoPattern
Match size (bp): 60000 VLP:WC ratio: 2.8776
Whole-community read coverage
VLP-fraction read coverage
Contig position (bp)
Oscillospiraceae NSJ-51

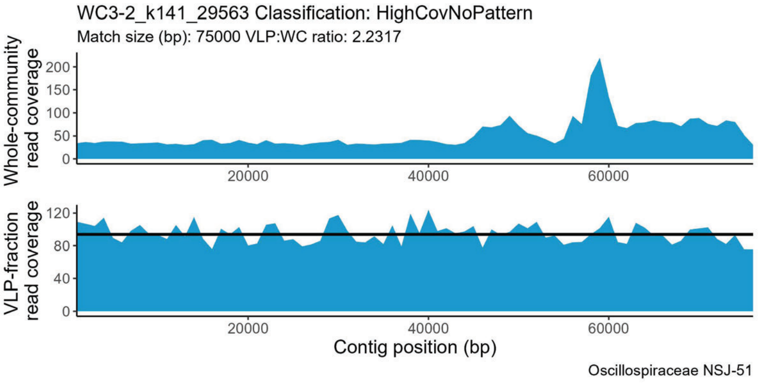
WC3-2_k141_29563 Classification: HighCovNoPattern
Match size (bp): 75000 VLP:WC ratio: 2.2317
Whole-community read coverage
VLP-fraction read coverage
Contig position (bp)
Oscillospiraceae NSJ-51

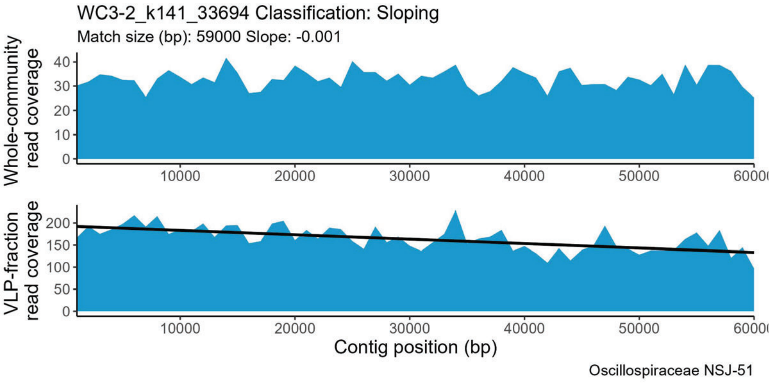
WC3-2_k141_33694 Classification: Sloping
Match size (bp): 59000 Slope: -0.001
Whole-community read coverage
VLP-fraction read coverage
Contig position (bp)
Oscillospiraceae NSJ-51

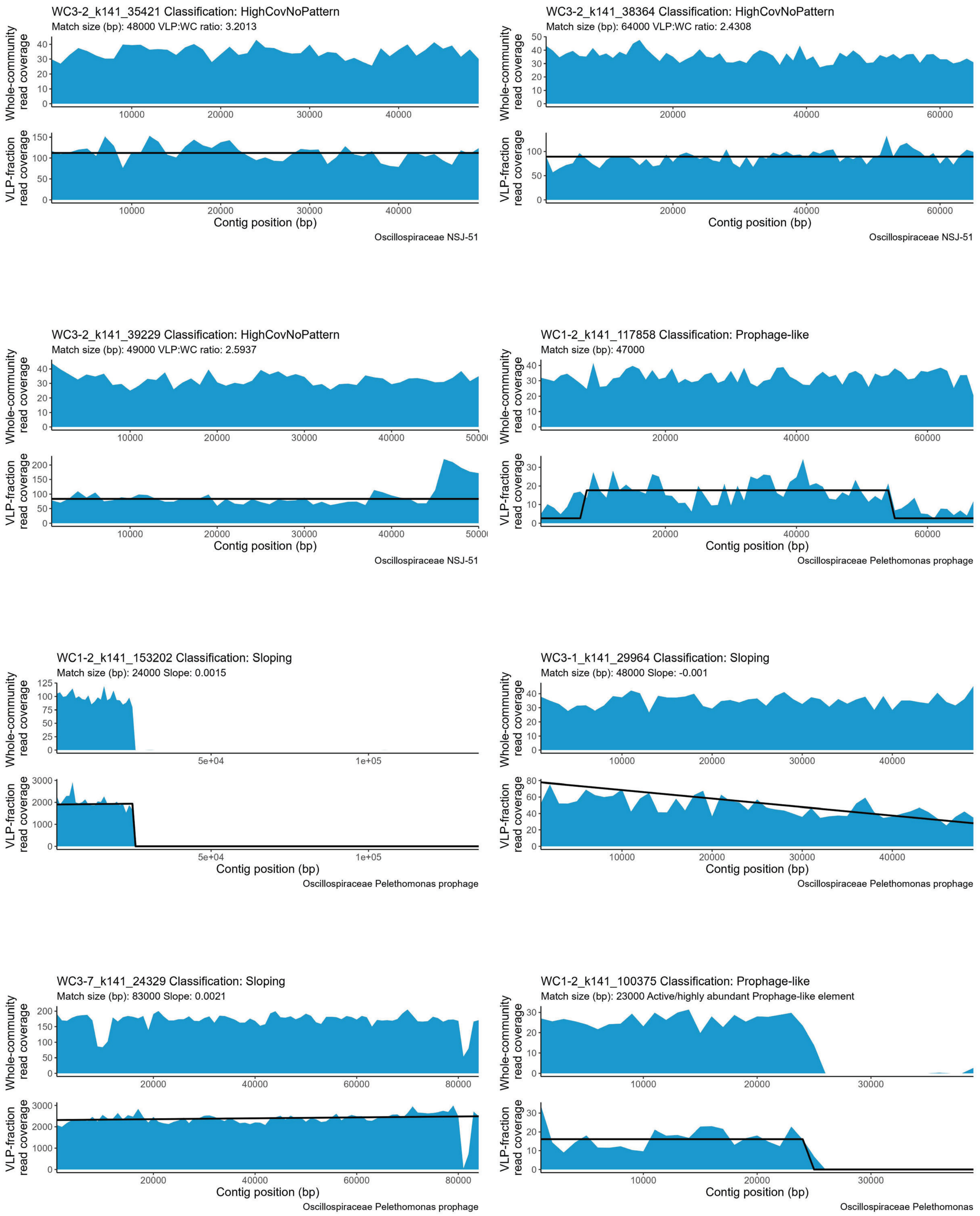

WC3-2_k141_35421 Classification: HighCovNoPattern
Match size (bp): 48000 VLP:WC ratio: 3.2013
Whole-community read coverage
VLP-fraction read coverage
Contig position (bp)
Oscillospiraceae NSJ-51
WC3-2_k141_38364 Classification: HighCovNoPattern
Match size (bp): 64000 VLP:WC ratio: 2.4308
Whole-community read coverage
VLP-fraction read coverage
Contig position (bp)
Oscillospiraceae NSJ-51
WC3-2_k141_39229 Classification: HighCovNoPattern
Match size (bp): 49000 VLP:WC ratio: 2.5937
Whole-community read coverage
VLP-fraction read coverage
Contig position (bp)
Oscillospiraceae NSJ-51
WC1-2_k141_117858 Classification: Prophage-like
Match size (bp): 47000
Whole-community read coverage
VLP-fraction read coverage
Contig position (bp)
Oscillospiraceae Pelethomonas prophage
WC1-2_k141_153202 Classification: Sloping
Match size (bp): 24000 Slope: 0.0015
Whole-community read coverage
VLP-fraction read coverage
Contig position (bp)
Oscillospiraceae Pelethomonas prophage
WC3-1_k141_29964 Classification: Sloping
Match size (bp): 48000 Slope: -0.001
Whole-community read coverage
VLP-fraction read coverage
Contig position (bp)
Oscillospiraceae Pelethomonas prophage
WC3-7_k141_24329 Classification: Sloping
Match size (bp): 83000 Slope: 0.0021
Whole-community read coverage
VLP-fraction read coverage
Contig position (bp)
Oscillospiraceae Pelethomonas prophage
WC1-2_k141_100375 Classification: Prophage-like
Match size (bp): 23000 Active/highly abundant Prophage-like element
Whole-community read coverage
VLP-fraction read coverage
Contig position (bp)
Oscillospiraceae Pelethomonas

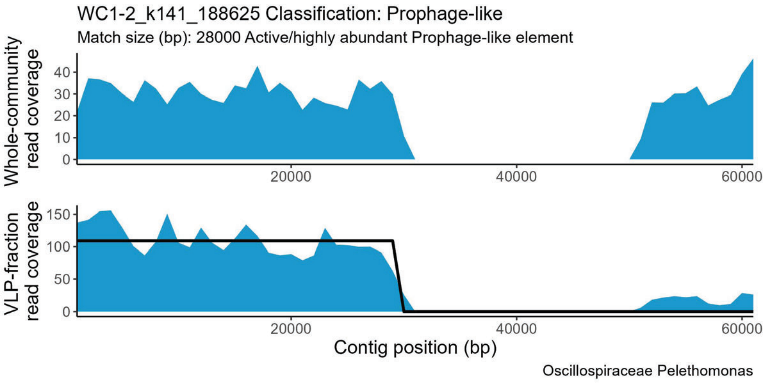
WC1-2_k141_188625 Classification: Prophage-like
Match size (bp): 28000 Active/highly abundant Prophage-like element
Whole-community read coverage
VLP-fraction read coverage
Contig position (bp)
Oscillospiraceae Pelethomonas

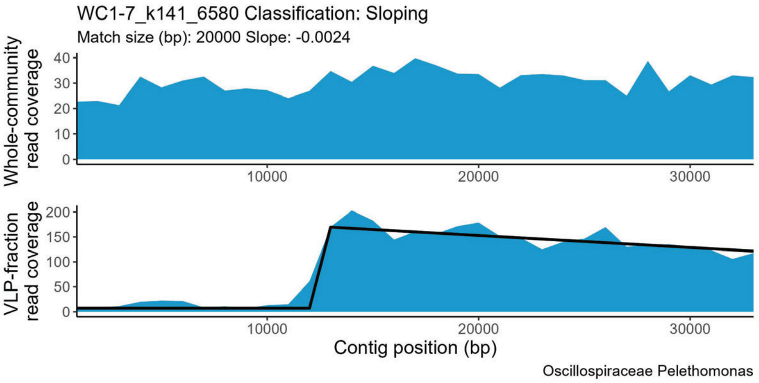
WC1-7_k141_6580 Classification: Sloping
Match size (bp): 20000 Slope: -0.0024
Whole-community read coverage
VLP-fraction read coverage
Contig position (bp)
Oscillospiraceae Pelethomonas

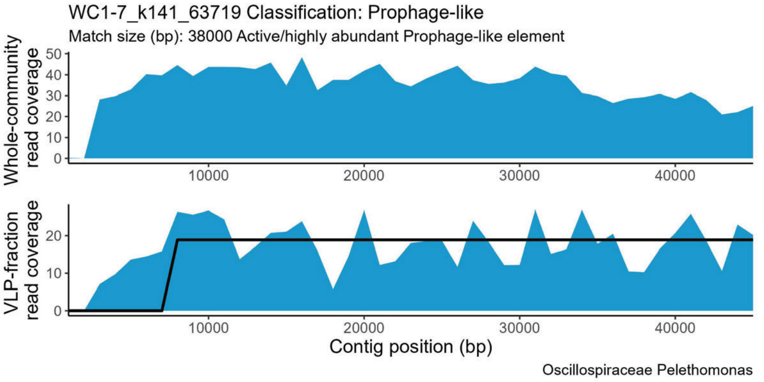
WC1-7_k141_63719 Classification: Prophage-like
Match size (bp): 38000 Active/highly abundant Prophage-like element
Whole-community read coverage
VLP-fraction read coverage
Contig position (bp)
Oscillospiraceae Pelethomonas

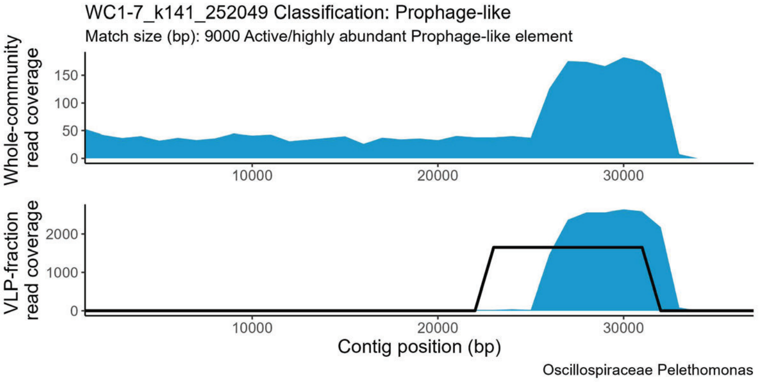
WC1-7_k141_252049 Classification: Prophage-like
Match size (bp): 9000 Active/highly abundant Prophage-like element
Whole-community read coverage
VLP-fraction read coverage
Contig position (bp)
Oscillospiraceae Pelethomonas

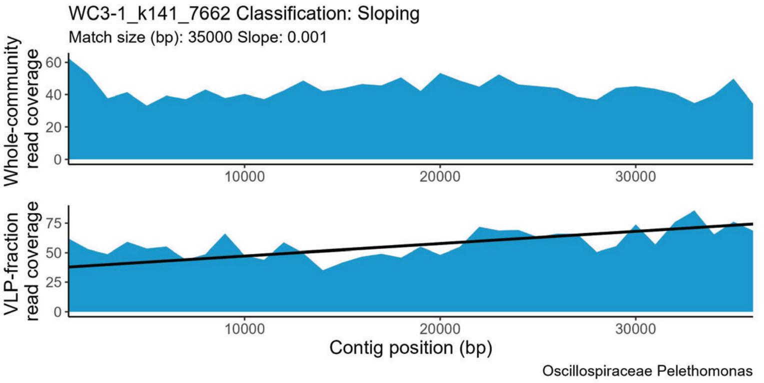
WC3-1_k141_7662 Classification: Sloping
Match size (bp): 35000 Slope: 0.001
Whole-community read coverage
VLP-fraction read coverage
Contig position (bp)
Oscillospiraceae Pelethomonas

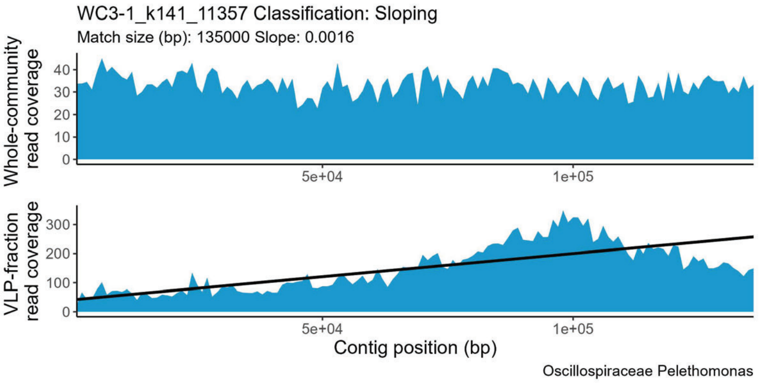
WC3-1_k141_11357 Classification: Sloping
Match size (bp): 135000 Slope: 0.0016
Whole-community read coverage
VLP-fraction read coverage
Contig position (bp)
Oscillospiraceae Pelethomonas

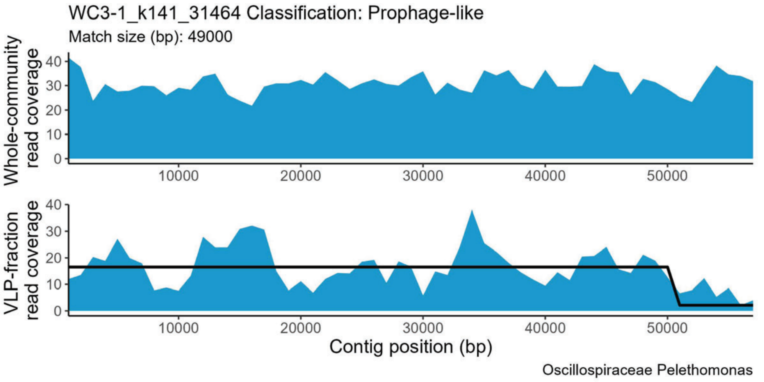
WC3-1_k141_31464 Classification: Prophage-like
Match size (bp): 49000
Whole-community read coverage
VLP-fraction read coverage
Contig position (bp)
Oscillospiraceae Pelethomonas

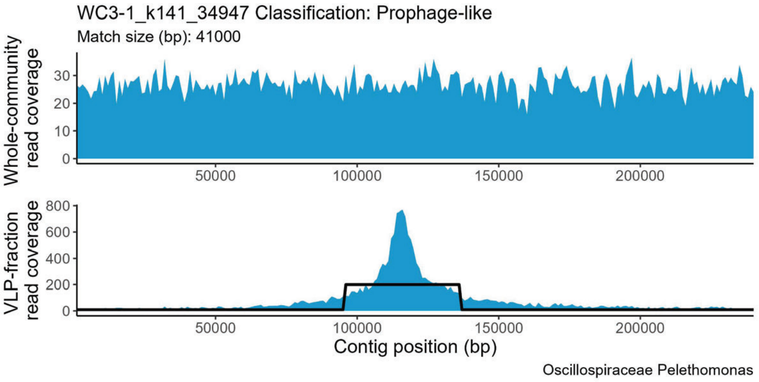
WC3-1_k141_34947 Classification: Prophage-like
Match size (bp): 41000
Whole-community read coverage
VLP-fraction read coverage
Contig position (bp)
Oscillospiraceae Pelethomonas

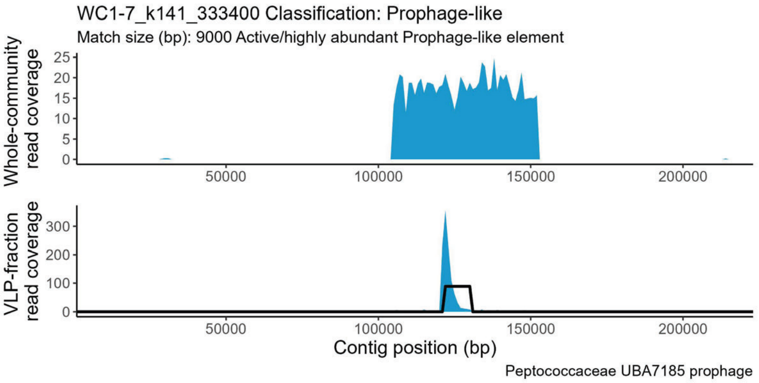
WC1-7_k141_333400 Classification: Prophage-like
Match size (bp): 9000 Active/highly abundant Prophage-like element
Whole-community read coverage
VLP-fraction read coverage
Contig position (bp)
Peptococcaceae UBA7185 prophage

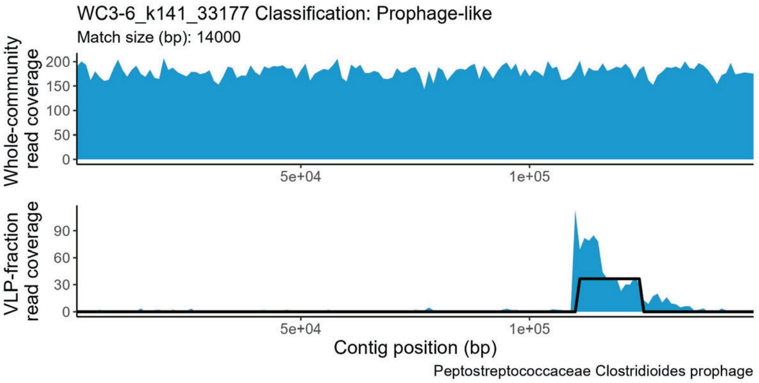
WC3-6_k141_33177 Classification: Prophage-like
Match size (bp): 14000
Whole-community read coverage
VLP-fraction read coverage
Contig position (bp)
Peptostreptococcaceae Clostridioides prophage

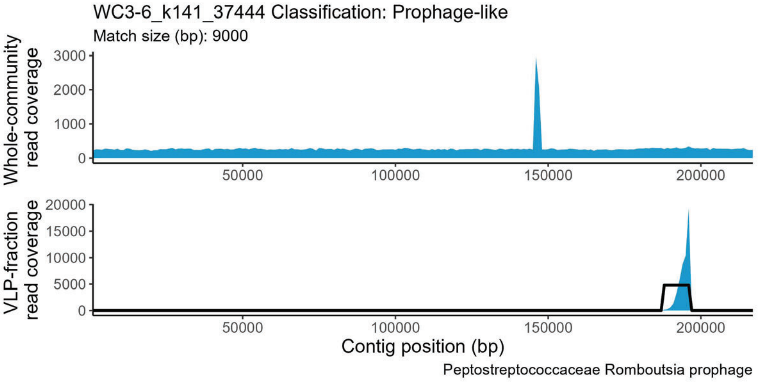
WC3-6_k141_37444 Classification: Prophage-like
Match size (bp): 9000
Whole-community read coverage
VLP-fraction read coverage
Contig position (bp)
Peptostreptococcaceae Romboutsia prophage

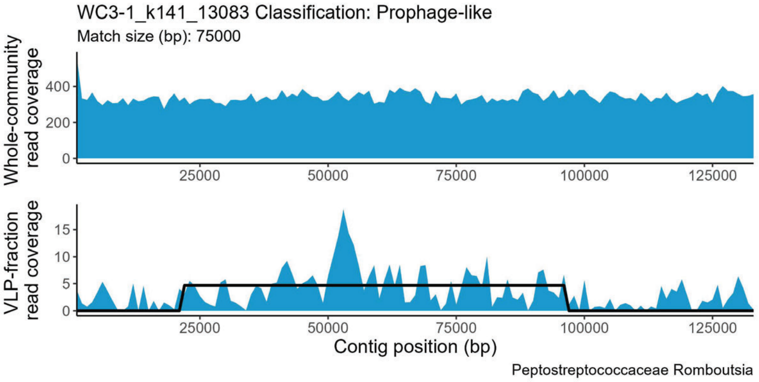
WC3-1_k141_13083 Classification: Prophage-like
Match size (bp): 75000
Whole-community read coverage
VLP-fraction read coverage
Contig position (bp)
Peptostreptococcaceae Romboutsia

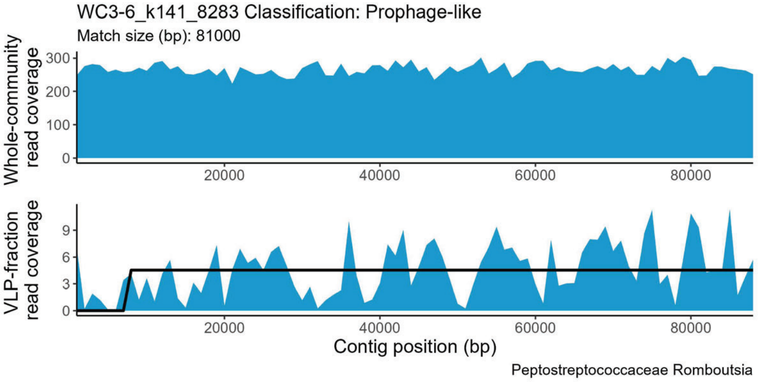
WC3-6_k141_8283 Classification: Prophage-like
Match size (bp): 81000
Whole-community read coverage
VLP-fraction read coverage
Contig position (bp)
Peptostreptococcaceae Romboutsia

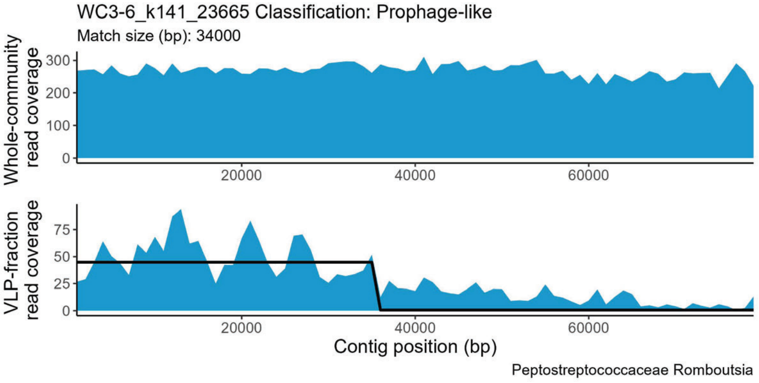
WC3-6_k141_23665 Classification: Prophage-like
Match size (bp): 34000
Whole-community read coverage
VLP-fraction read coverage
Contig position (bp)
Peptostreptococcaceae Romboutsia

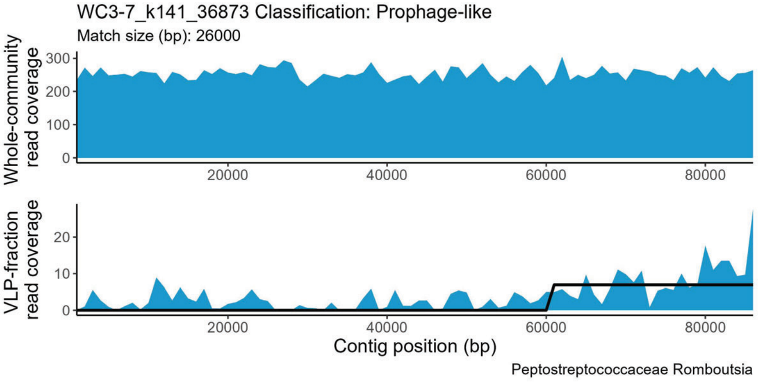
WC3-7_k141_36873 Classification: Prophage-like
Match size (bp): 26000
Whole-community read coverage
VLP-fraction read coverage
Contig position (bp)
Peptostreptococcaceae Romboutsia

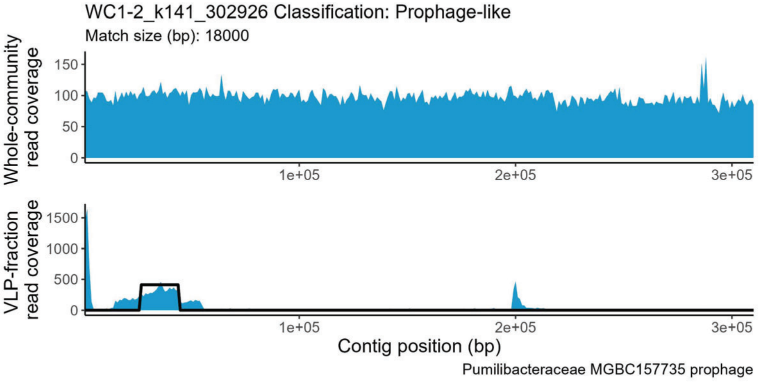
WC1-2_k141_302926 Classification: Prophage-like
Match size (bp): 18000
Whole-community read coverage
VLP-fraction read coverage
Contig position (bp)
Pumilibacteraceae MGBC157735 prophage

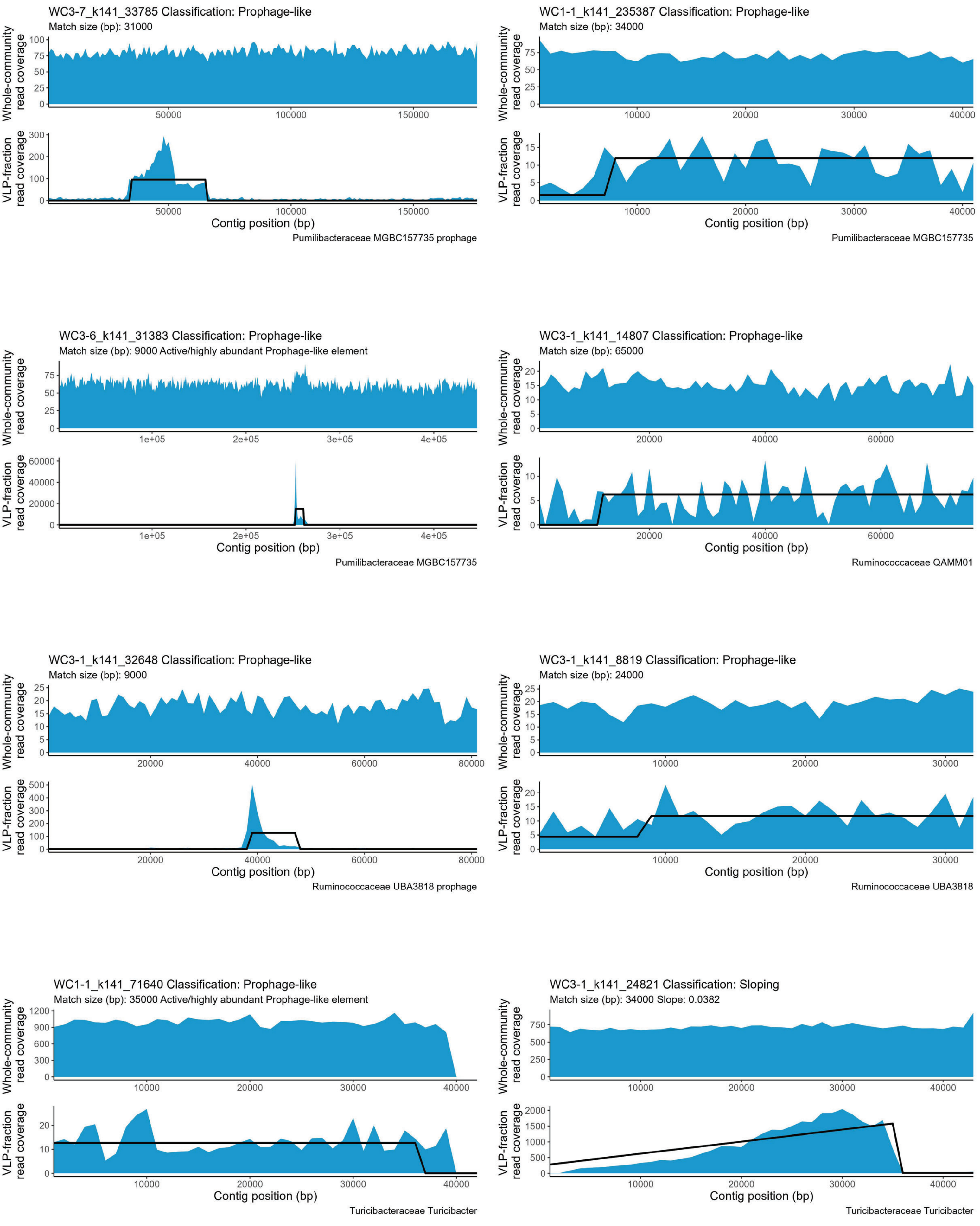
WC3-7_k141_33785 Classification: Prophage-like
Match size (bp): 31000
Whole-community read coverage
VLP-fraction read coverage
Contig position (bp)
Pumilibacteraceae MGBC157735 prophage
WC1-1_k141_235387 Classification: Prophage-like
Match size (bp): 34000
Whole-community read coverage
VLP-fraction read coverage
Contig position (bp)
Pumilibacteraceae MGBC157735
WC3-6_k141_31383 Classification: Prophage-like
Match size (bp): 9000 Active/highly abundant Prophage-like element
Whole-community read coverage
VLP-fraction read coverage
Contig position (bp)
Pumilibacteraceae MGBC157735
WC3-1_k141_14807 Classification: Prophage-like
Match size (bp): 65000
Whole-community read coverage
VLP-fraction read coverage
Contig position (bp)
Ruminococcaceae QAMM01
WC3-1_k141_32648 Classification: Prophage-like
Match size (bp): 9000
Whole-community read coverage
VLP-fraction read coverage
Contig position (bp)
Ruminococcaceae UBA3818 prophage
WC3-1_k141_8819 Classification: Prophage-like
Match size (bp): 24000
Whole-community read coverage
VLP-fraction read coverage
Contig position (bp)
Ruminococcaceae UBA3818
WC1-1_k141_71640 Classification: Prophage-like
Match size (bp): 35000 Active/highly abundant Prophage-like element
Whole-community read coverage
VLP-fraction read coverage
Contig position (bp)
Turicibacteraceae Turicibacter
WC3-1_k141_24821 Classification: Sloping
Match size (bp): 34000 Slope: 0.0382
Whole-community read coverage
VLP-fraction read coverage
Contig position (bp)
Turicibacteraceae Turicibacter

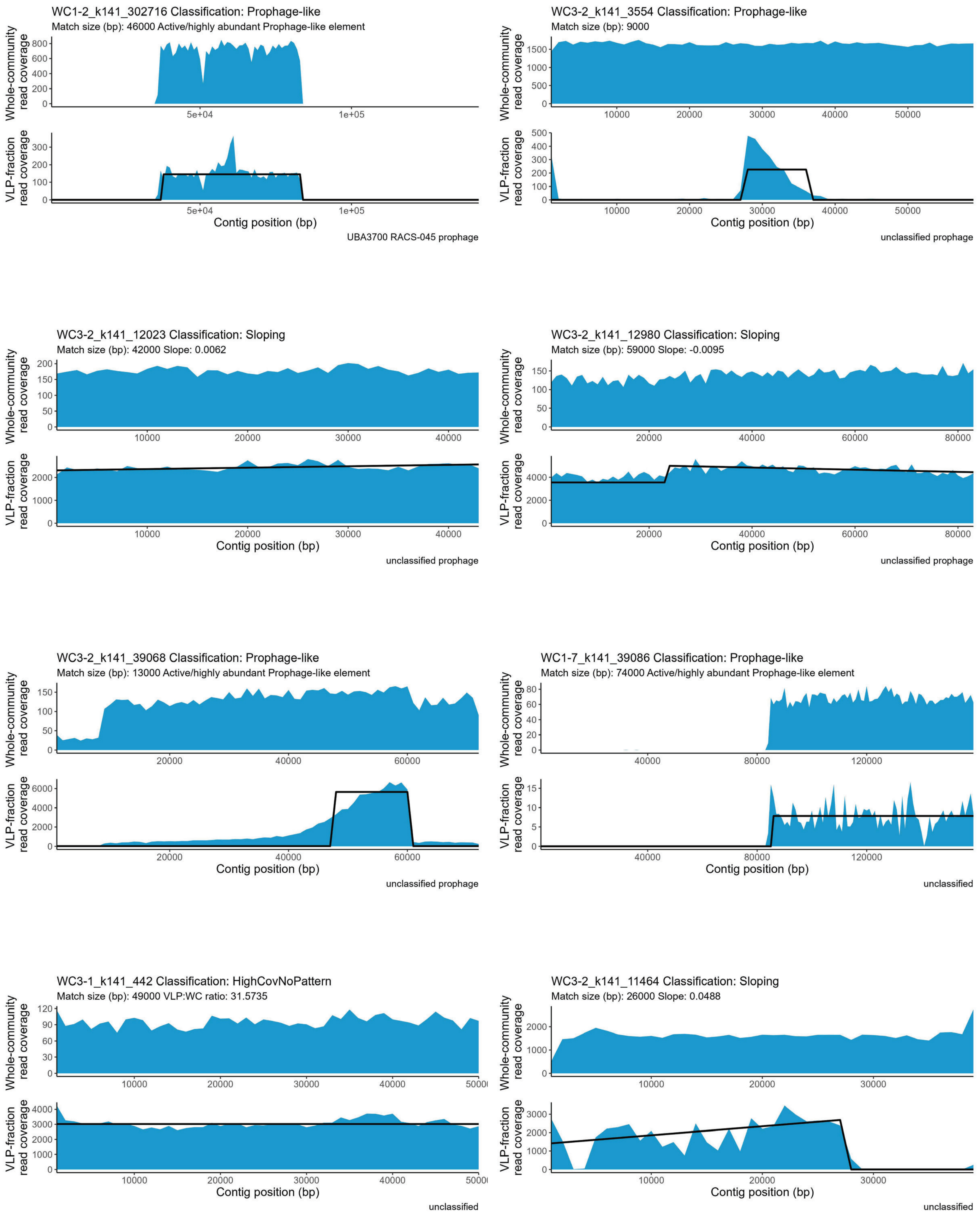
WC1-2_k141_302716 Classification: Prophage-like
Match size (bp): 46000 Active/highly abundant Prophage-like element
Whole-community read coverage
VLP-fraction read coverage
Contig position (bp)
UBA3700 RACS-045 prophage
WC3-2_k141_3554 Classification: Prophage-like
Match size (bp): 9000
Whole-community read coverage
VLP-fraction read coverage
Contig position (bp)
unclassified prophage
WC3-2_k141_12023 Classification: Sloping
Match size (bp): 42000 Slope: 0.0062
Whole-community read coverage
VLP-fraction read coverage
Contig position (bp)
unclassified prophage
WC3-2_k141_12980 Classification: Sloping
Match size (bp): 59000 Slope: -0.0095
Whole-community read coverage
VLP-fraction read coverage
Contig position (bp)
unclassified prophage
WC3-2_k141_39068 Classification: Prophage-like
Match size (bp): 13000 Active/highly abundant Prophage-like element
Whole-community read coverage
VLP-fraction read coverage
Contig position (bp)
unclassified prophage
WC1-7_k141_39086 Classification: Prophage-like
Match size (bp): 74000 Active/highly abundant Prophage-like element
Whole-community read coverage
VLP-fraction read coverage
Contig position (bp)
unclassified
WC3-1_k141_442 Classification: HighCovNoPattern
Match size (bp): 49000 VLP:WC ratio: 31.5735
Whole-community read coverage
VLP-fraction read coverage
Contig position (bp)
unclassified
WC3-2_k141_11464 Classification: Sloping
Match size (bp): 26000 Slope: 0.0488
Whole-community read coverage
VLP-fraction read coverage
Contig position (bp)
unclassified

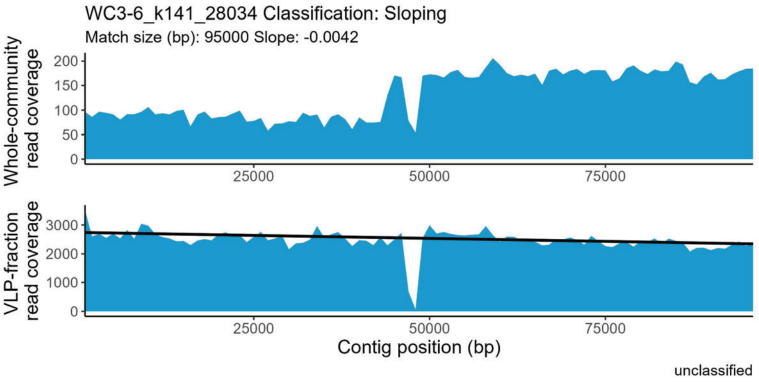
WC3-6_k141_28034 Classification: Sloping
Match size (bp): 95000 Slope: -0.0042
Whole-community read coverage
VLP-fraction read coverage
Contig position (bp)
unclassified

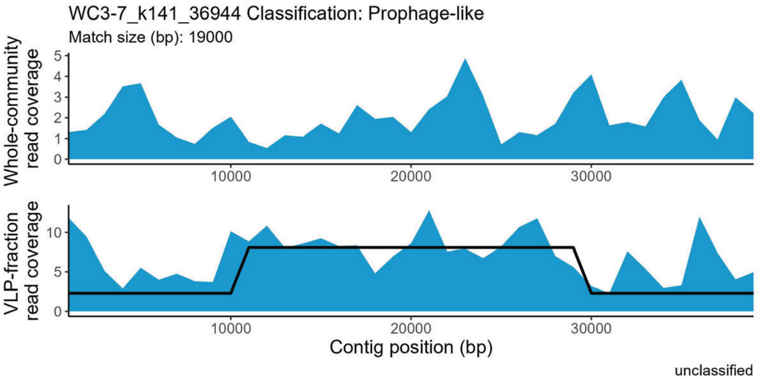
WC3-7_k141_36944 Classification: Prophage-like
Match size (bp): 19000
Whole-community read coverage
VLP-fraction read coverage
Contig position (bp)
unclassified

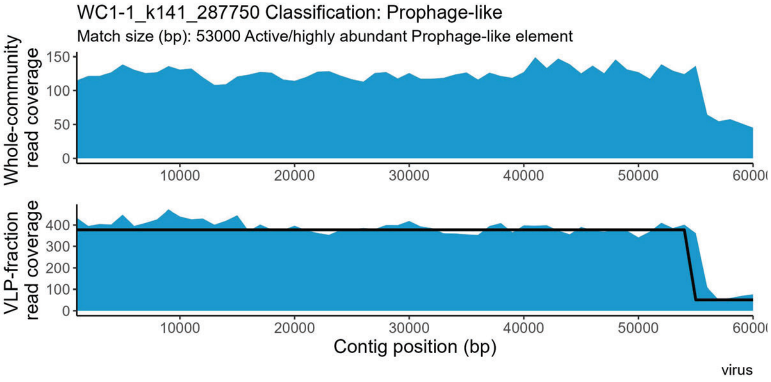
WC1-1_k141_287750 Classification: Prophage-like
Match size (bp): 53000 Active/highly abundant Prophage-like element
Whole-community read coverage
VLP-fraction read coverage
Contig position (bp)
virus

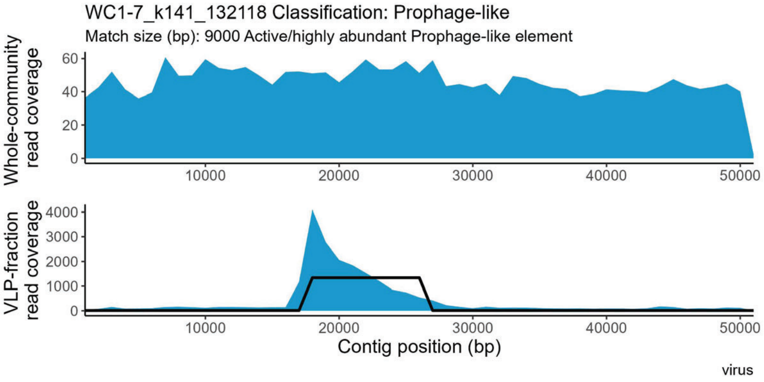
WC1-7_k141_132118 Classification: Prophage-like
Match size (bp): 9000 Active/highly abundant Prophage-like element
Whole-community read coverage
VLP-fraction read coverage
Contig position (bp)
virus

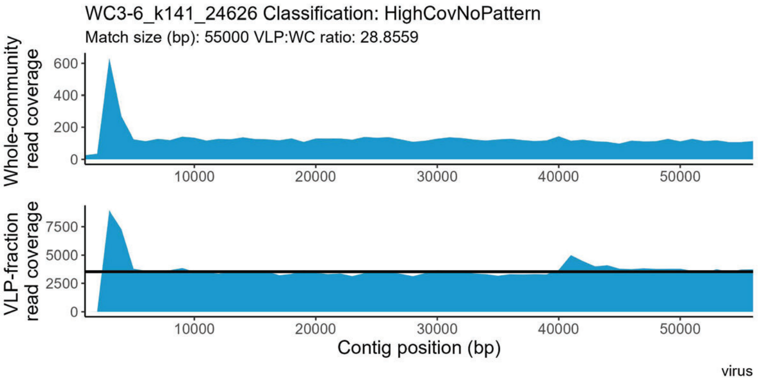
WC3-6_k141_24626 Classification: HighCovNoPattern
Match size (bp): 55000 VLP:WC ratio: 28.8559
Whole-community read coverage
VLP-fraction read coverage
Contig position (bp)
virus
