## Supplemental file 1 for "Perturbations shift the composition of bacterial DNA carried by virus-like particles in the murine gut microbiome"

**File S1) All positive TrIdent pattern-matches for Pre-ABX replicate 1.** The sample from which a contig was assembled is indicated by the following prefixes in the contig accession number: WC1-1 = Pre-ABX replicate 1, WC1-2 = Pre-ABX replicate 2, WC1-6 = Pre-ABX replicate 3, WC1-7 = Pre-ABX replicate 4, WC1-1 = CDI replicate 1, WC1-2 = CDI replicate 2, WC1-6 = CDI replicate 3, WC1-7 = CDI replicate 4. Pre-ABX = Pre-antibiotics, Post-ABX= post-antbiotics, CDI= *Clostridioides difficile* infection. Contig taxonomy and geNomad prophage classification in plot subtitle. Plots are listed alphabetically by family name.

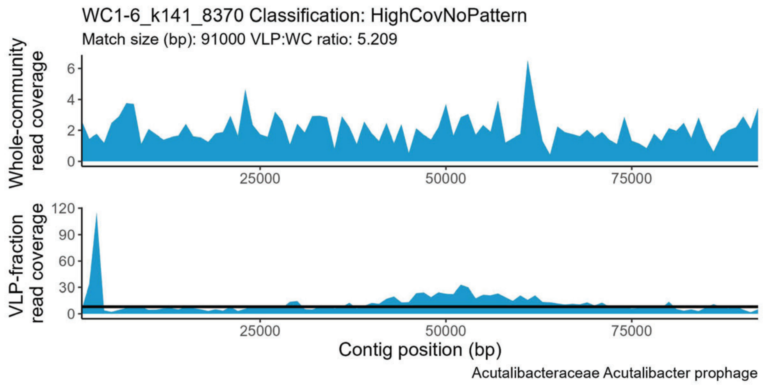


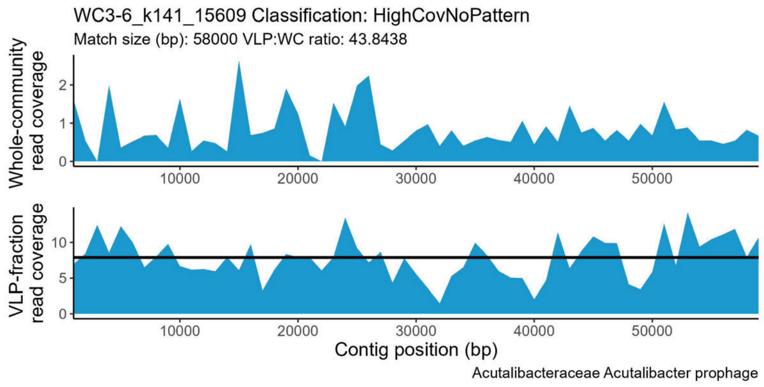


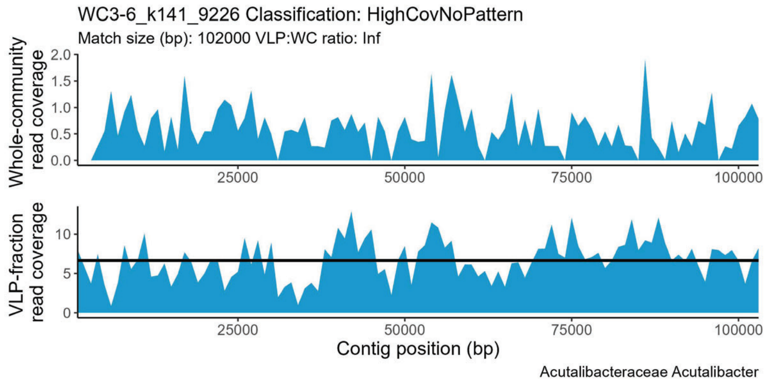


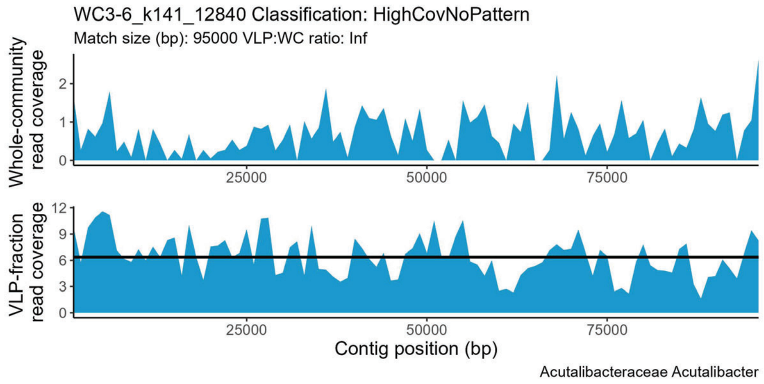


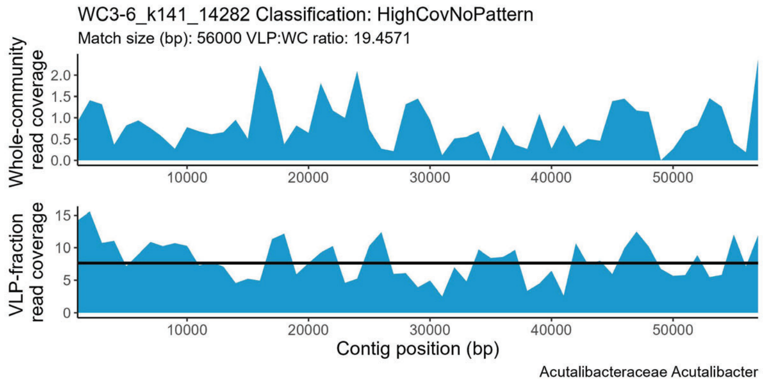


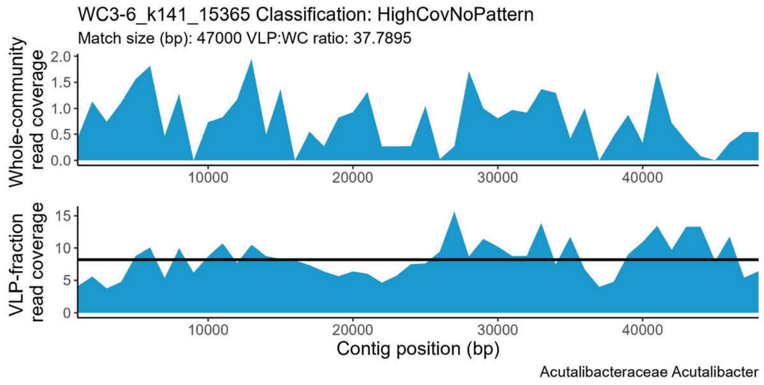


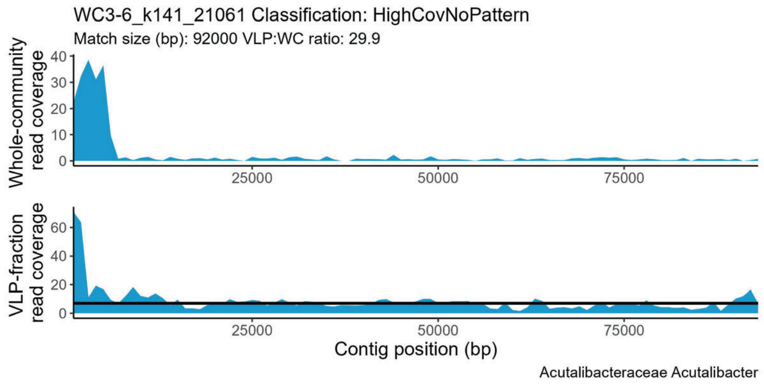


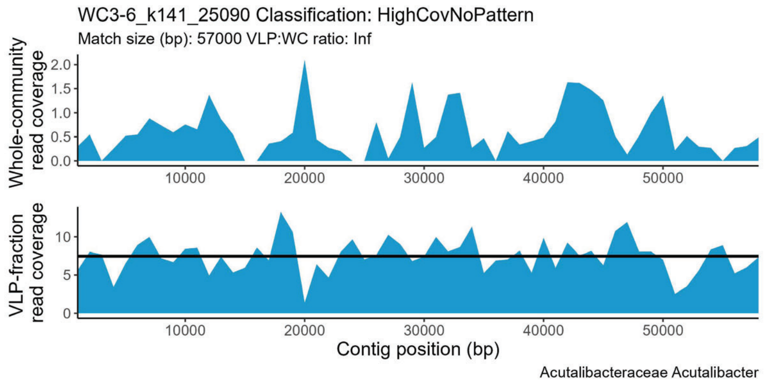

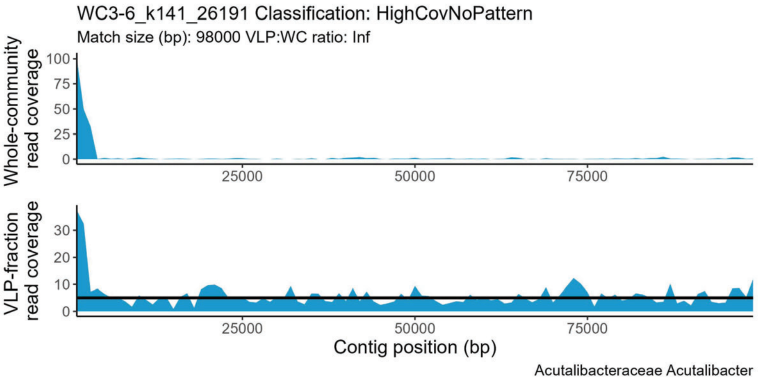
WC3-6_k141_26191 Classification: HighCovNoPattern
Match size (bp): 98000 VLP:WC ratio: Inf
Whole-community read coverage
100
75
50
25
0
25000
50000
75000
VLP-fraction read coverage
30
20
10
0
Contig position (bp)
Acutalibacteraceae Acutalibacter

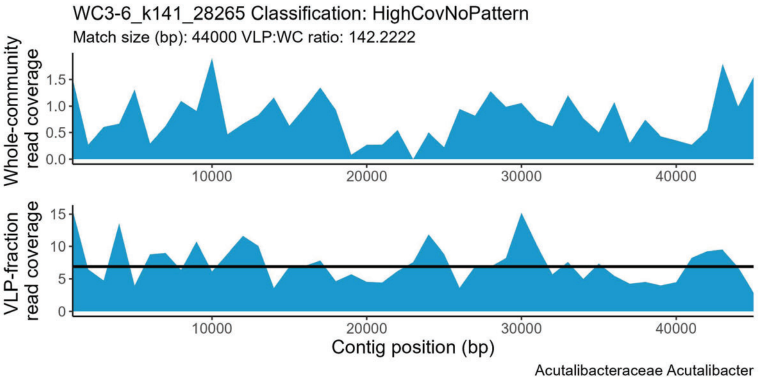
WC3-6_k141_28265 Classification: HighCovNoPattern
Match size (bp): 44000 VLP:WC ratio: 142.2222
Whole-community read coverage
1.5
1.0
0.5
0.0
10000
20000
30000
40000
VLP-fraction read coverage
15
10
5
0
Contig position (bp)
Acutalibacteraceae Acutalibacter

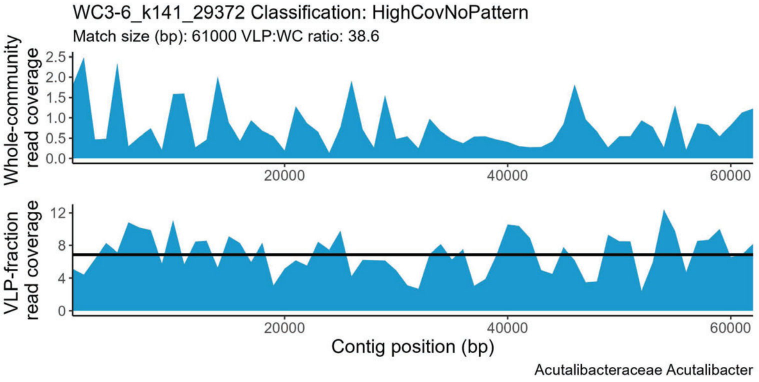
WC3-6_k141_29372 Classification: HighCovNoPattern
Match size (bp): 61000 VLP:WC ratio: 38.6
Whole-community read coverage
2.5
2.0
1.5
1.0
0.5
0.0
20000
40000
60000
VLP-fraction read coverage
12
8
4
0
Contig position (bp)
Acutalibacteraceae Acutalibacter

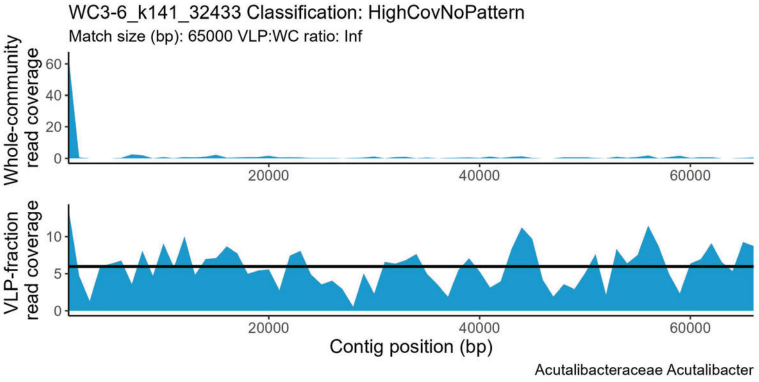
WC3-6_k141_32433 Classification: HighCovNoPattern
Match size (bp): 65000 VLP:WC ratio: Inf
Whole-community read coverage
60
40
20
0
20000
40000
60000
VLP-fraction read coverage
10
5
0
Contig position (bp)
Acutalibacteraceae Acutalibacter

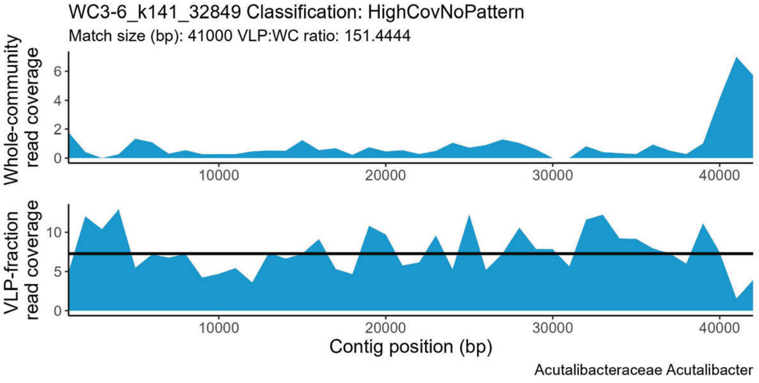
WC3-6_k141_32849 Classification: HighCovNoPattern
Match size (bp): 41000 VLP:WC ratio: 151.4444
Whole-community read coverage
6
4
2
0
10000
20000
30000
40000
VLP-fraction read coverage
10
5
0
Contig position (bp)
Acutalibacteraceae Acutalibacter

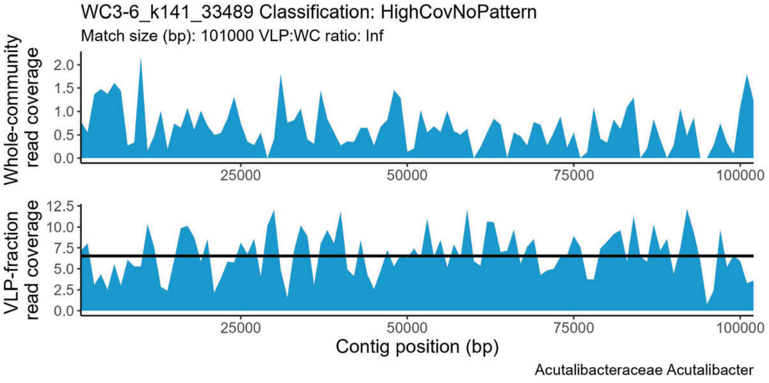
WC3-6_k141_33489 Classification: HighCovNoPattern
Match size (bp): 101000 VLP:WC ratio: Inf
Whole-community read coverage
2.0
1.5
1.0
0.5
0.0
25000
50000
75000
100000
VLP-fraction read coverage
12.5
10.0
7.5
5.0
2.5
0.0
Contig position (bp)
Acutalibacteraceae Acutalibacter

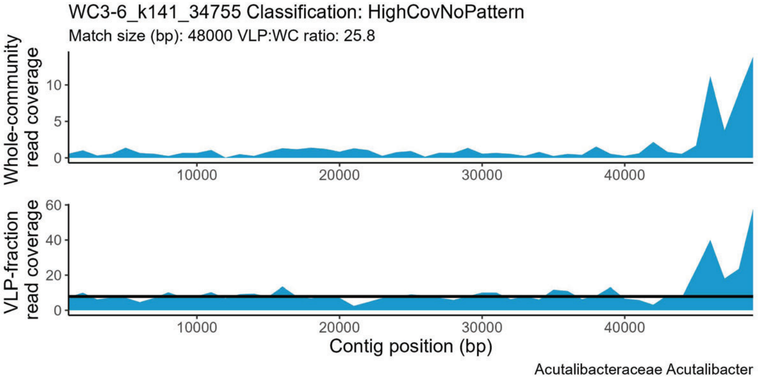
WC3-6_k141_34755 Classification: HighCovNoPattern
Match size (bp): 48000 VLP:WC ratio: 25.8
Whole-community read coverage
10
5
0
10000
20000
30000
40000
VLP-fraction read coverage
60
40
20
0
Contig position (bp)
Acutalibacteraceae Acutalibacter

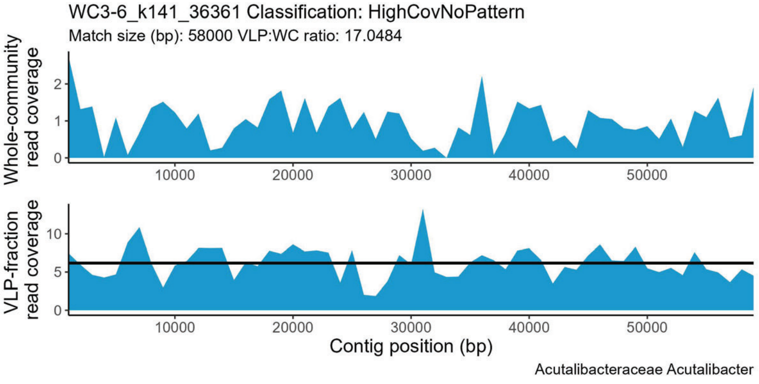
WC3-6_k141_36361 Classification: HighCovNoPattern
Match size (bp): 58000 VLP:WC ratio: 17.0484
Whole-community read coverage
2
1
0
10000
20000
30000
40000
50000
VLP-fraction read coverage
10
5
0
Contig position (bp)
Acutalibacteraceae Acutalibacter

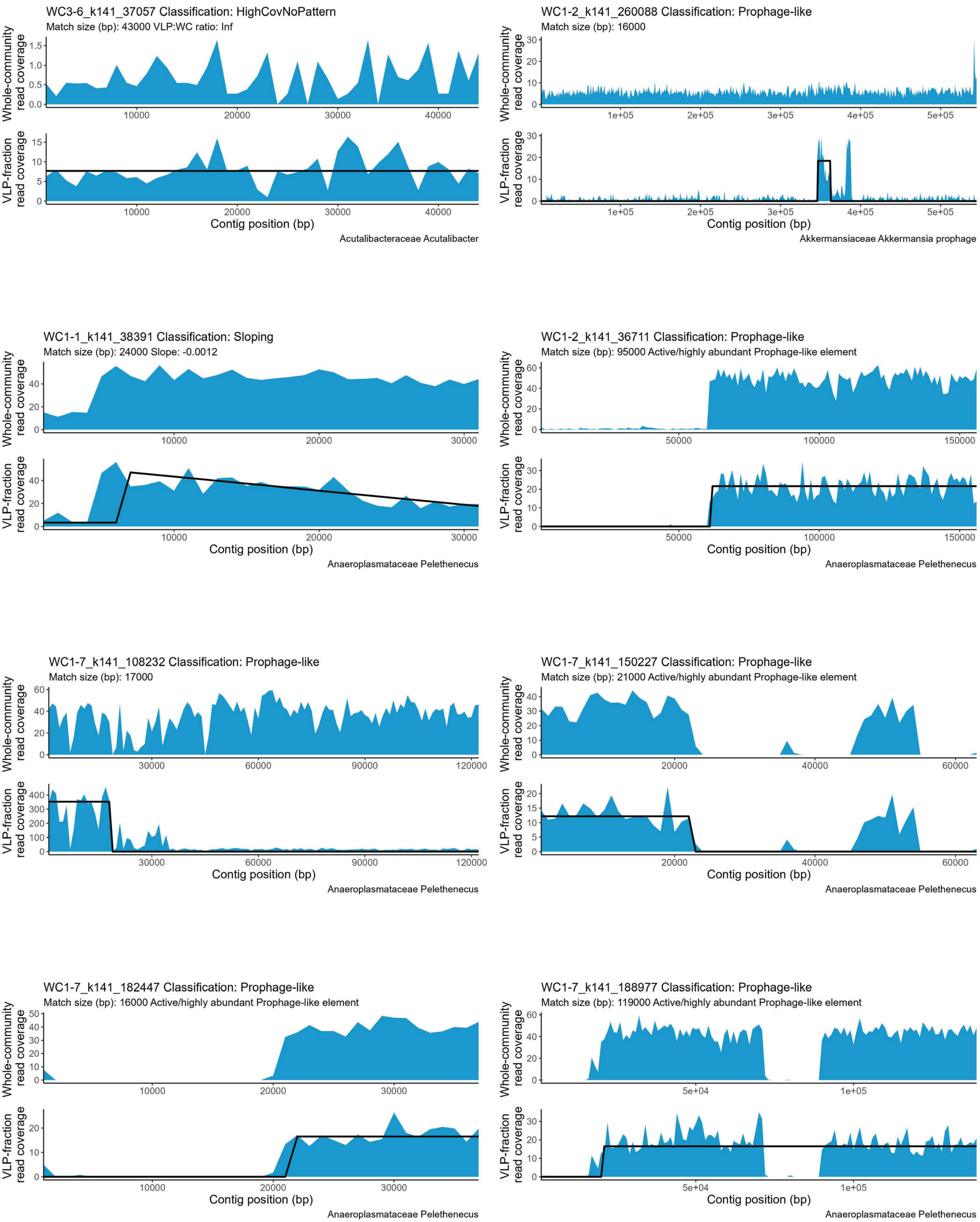
WC3-6_k141_37057 Classification: HighCovNoPattern
Match size (bp): 43000 VLP:WC ratio: Inf
Whole-community read coverage
VLP-fraction read coverage
Contig position (bp)
Acutalibacteraceae Acutalibacter
WC1-2_k141_260088 Classification: Prophage-like
Match size (bp): 16000
Whole-community read coverage
VLP-fraction read coverage
Contig position (bp)
Akkermansiaceae Akkermansia prophage
WC1-1_k141_38391 Classification: Sloping
Match size (bp): 24000 Slope: -0.0012
Whole-community read coverage
VLP-fraction read coverage
Contig position (bp)
Anaeroplasmataceae Pelethenecus
WC1-2_k141_36711 Classification: Prophage-like
Match size (bp): 95000 Active/highly abundant Prophage-like element
Whole-community read coverage
VLP-fraction read coverage
Contig position (bp)
Anaeroplasmataceae Pelethenecus
WC1-7_k141_108232 Classification: Prophage-like
Match size (bp): 17000
Whole-community read coverage
VLP-fraction read coverage
Contig position (bp)
Anaeroplasmataceae Pelethenecus
WC1-7_k141_150227 Classification: Prophage-like
Match size (bp): 21000 Active/highly abundant Prophage-like element
Whole-community read coverage
VLP-fraction read coverage
Contig position (bp)
Anaeroplasmataceae Pelethenecus
WC1-7_k141_182447 Classification: Prophage-like
Match size (bp): 16000 Active/highly abundant Prophage-like element
Whole-community read coverage
VLP-fraction read coverage
Contig position (bp)
Anaeroplasmataceae Pelethenecus
WC1-7_k141_188977 Classification: Prophage-like
Match size (bp): 119000 Active/highly abundant Prophage-like element
Whole-community read coverage
VLP-fraction read coverage
Contig position (bp)
Anaeroplasmataceae Pelethenecus

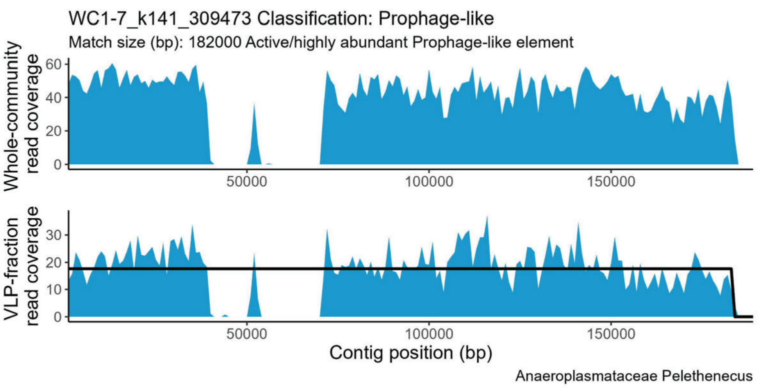
WC1-7_k141_309473 Classification: Prophage-like
Match size (bp): 182000 Active/highly abundant Prophage-like element
Whole-community read coverage
VLP-fraction read coverage
Contig position (bp)
Anaeroplasmataceae Pelethenecus

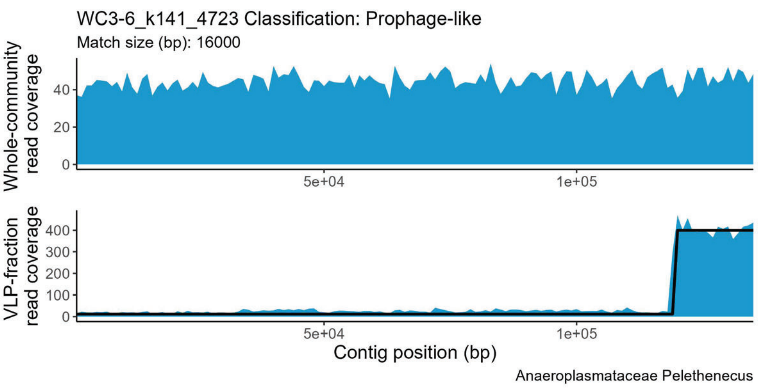
WC3-6_k141_4723 Classification: Prophage-like
Match size (bp): 16000
Whole-community read coverage
VLP-fraction read coverage
Contig position (bp)
Anaeroplasmataceae Pelethenecus

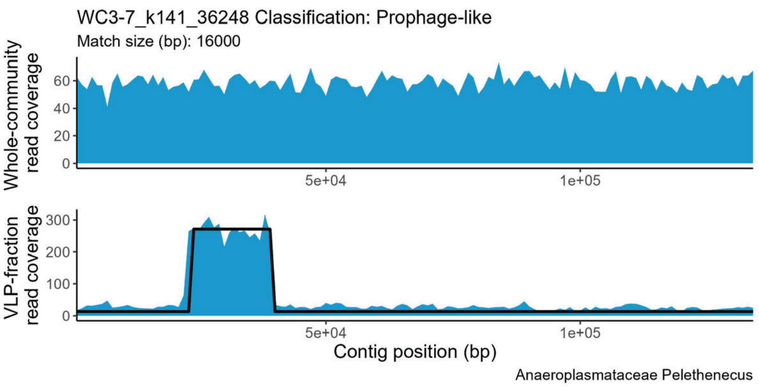
WC3-7_k141_36248 Classification: Prophage-like
Match size (bp): 16000
Whole-community read coverage
VLP-fraction read coverage
Contig position (bp)
Anaeroplasmataceae Pelethenecus

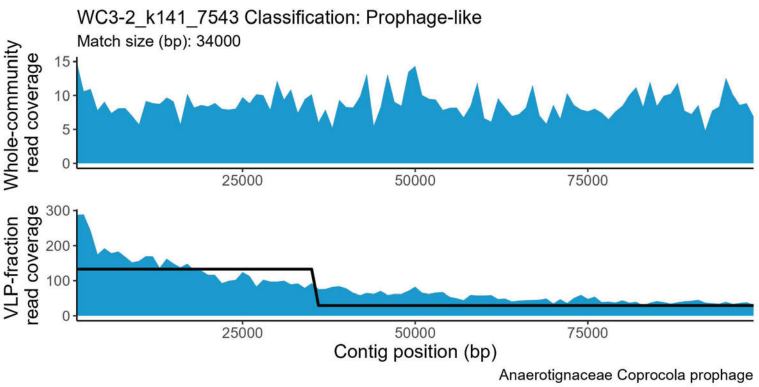
WC3-2_k141_7543 Classification: Prophage-like
Match size (bp): 34000
Whole-community read coverage
VLP-fraction read coverage
Contig position (bp)
Anaerotignaceae Coprocola prophage

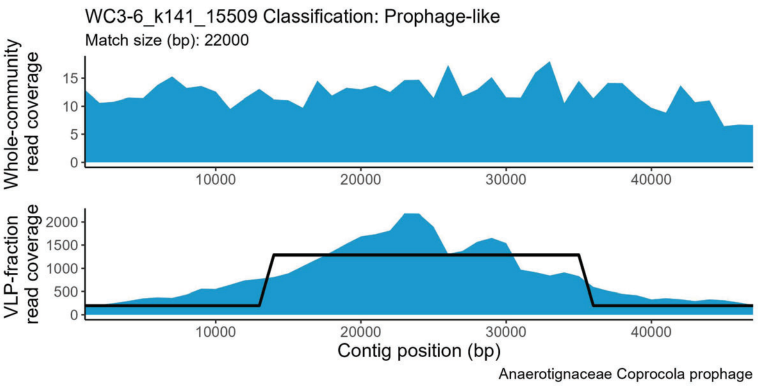
WC3-6_k141_15509 Classification: Prophage-like
Match size (bp): 22000
Whole-community read coverage
VLP-fraction read coverage
Contig position (bp)
Anaerotignaceae Coprocola prophage

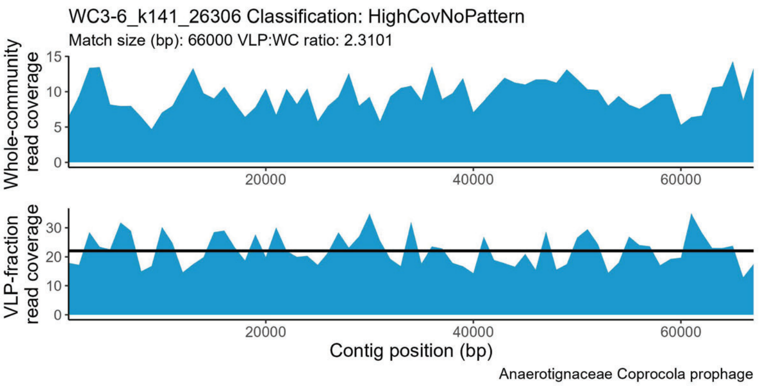
WC3-6_k141_26306 Classification: HighCovNoPattern
Match size (bp): 66000 VLP:WC ratio: 2.3101
Whole-community read coverage
VLP-fraction read coverage
Contig position (bp)
Anaerotignaceae Coprocola prophage

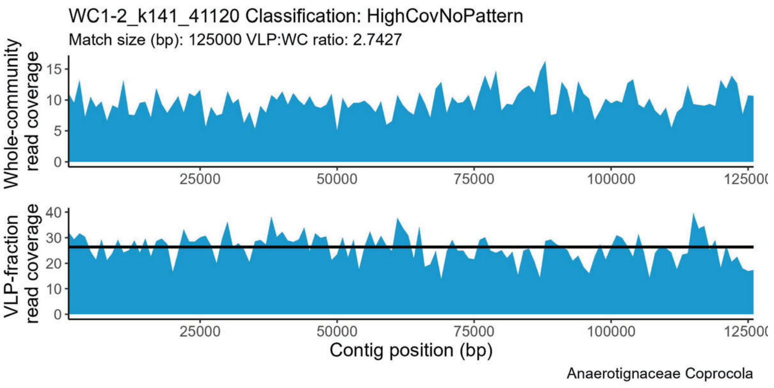
WC1-2_k141_41120 Classification: HighCovNoPattern
Match size (bp): 125000 VLP:WC ratio: 2.7427
Whole-community read coverage
VLP-fraction read coverage
Contig position (bp)
Anaerotignaceae Coprocola

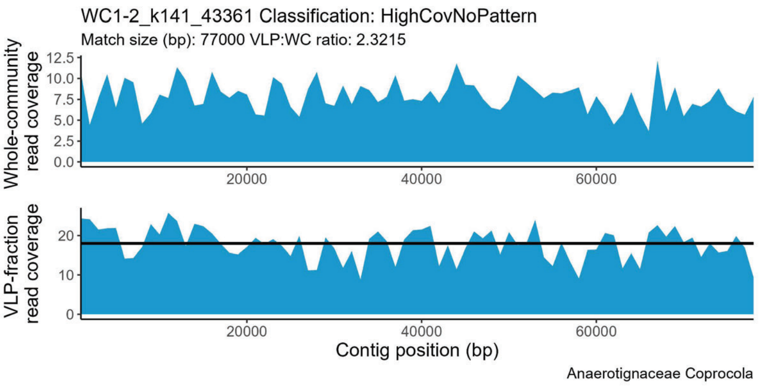
WC1-2_k141_43361 Classification: HighCovNoPattern
Match size (bp): 77000 VLP:WC ratio: 2.3215
Whole-community read coverage
VLP-fraction read coverage
Contig position (bp)
Anaerotignaceae Coprocola

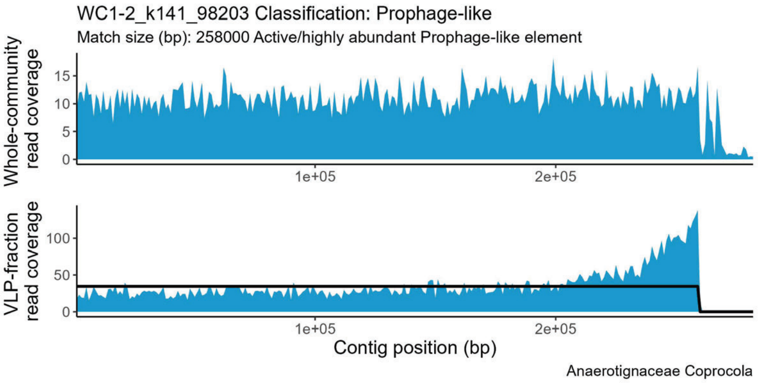
WC1-2_k141_98203 Classification: Prophage-like
Match size (bp): 258000 Active/highly abundant Prophage-like element
Whole-community read coverage
0
5
10
15
1e+05
2e+05
VLP-fraction read coverage
0
50
100
Contig position (bp)
Anaerotignaceae Coprocola

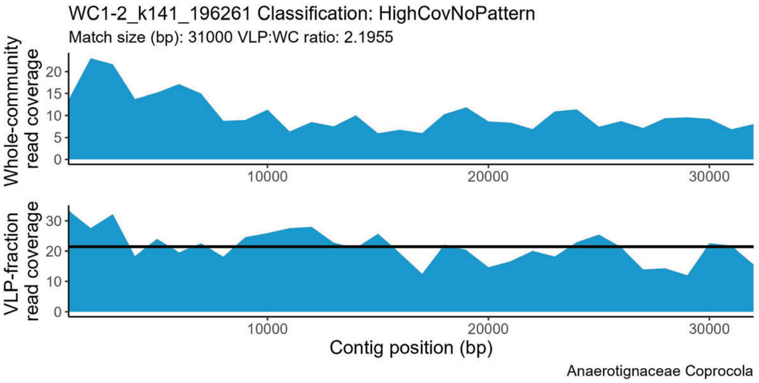
WC1-2_k141_196261 Classification: HighCovNoPattern
Match size (bp): 31000 VLP:WC ratio: 2.1955
Whole-community read coverage
0
5
10
15
20
10000
20000
30000
VLP-fraction read coverage
0
10
20
30
Contig position (bp)
Anaerotignaceae Coprocola

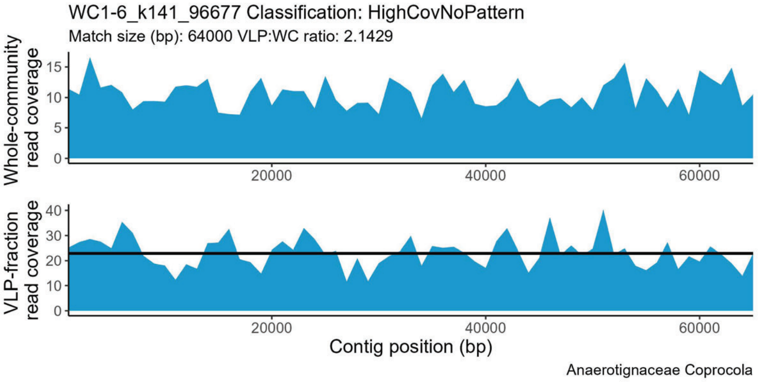
WC1-6_k141_96677 Classification: HighCovNoPattern
Match size (bp): 64000 VLP:WC ratio: 2.1429
Whole-community read coverage
0
5
10
15
20000
40000
60000
VLP-fraction read coverage
0
10
20
30
40
Contig position (bp)
Anaerotignaceae Coprocola

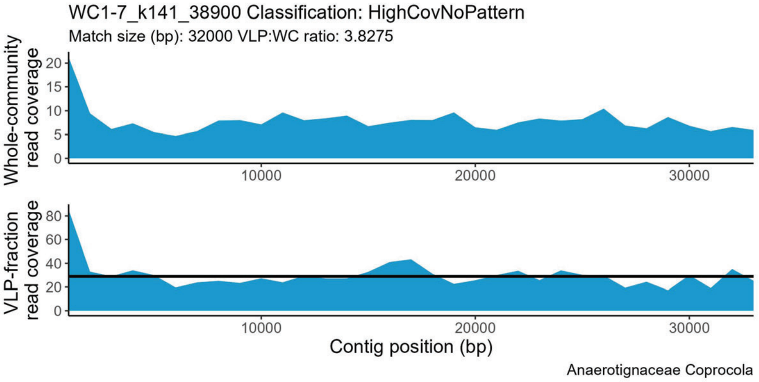
WC1-7_k141_38900 Classification: HighCovNoPattern
Match size (bp): 32000 VLP:WC ratio: 3.8275
Whole-community read coverage
0
5
10
15
20
10000
20000
30000
VLP-fraction read coverage
0
20
40
60
80
Contig position (bp)
Anaerotignaceae Coprocola

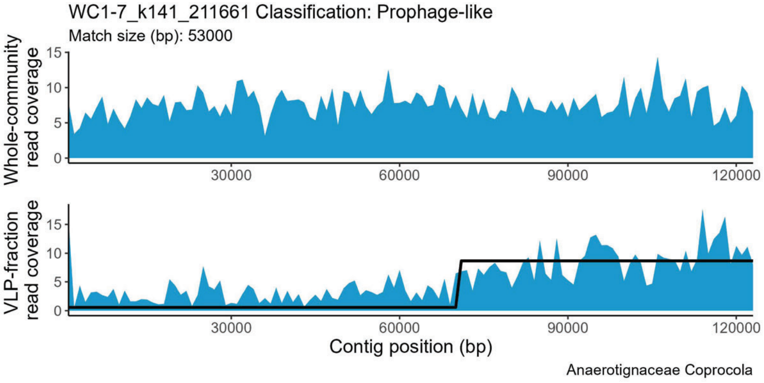
WC1-7_k141_211661 Classification: Prophage-like
Match size (bp): 53000
Whole-community read coverage
0
5
10
15
30000
60000
90000
120000
VLP-fraction read coverage
0
5
10
15
Contig position (bp)
Anaerotignaceae Coprocola

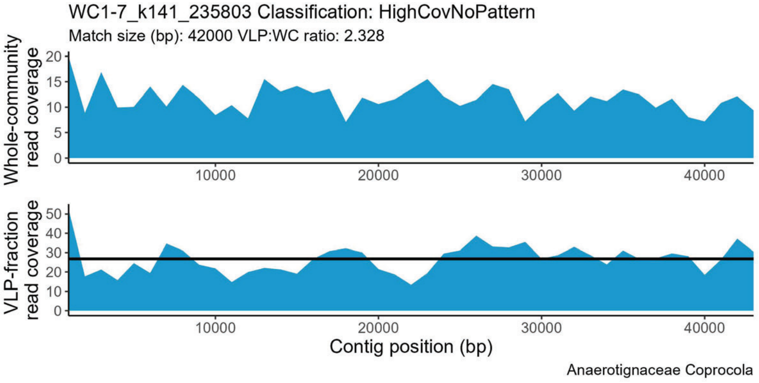
WC1-7_k141_235803 Classification: HighCovNoPattern
Match size (bp): 42000 VLP:WC ratio: 2.328
Whole-community read coverage
0
5
10
15
20
10000
20000
30000
40000
VLP-fraction read coverage
0
10
20
30
40
50
Contig position (bp)
Anaerotignaceae Coprocola

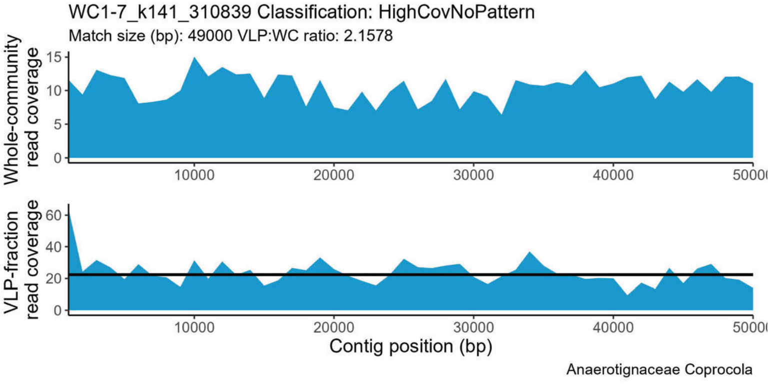
WC1-7_k141_310839 Classification: HighCovNoPattern
Match size (bp): 49000 VLP:WC ratio: 2.1578
Whole-community read coverage
0
5
10
15
10000
20000
30000
40000
VLP-fraction read coverage
0
20
40
60
Contig position (bp)
Anaerotignaceae Coprocola

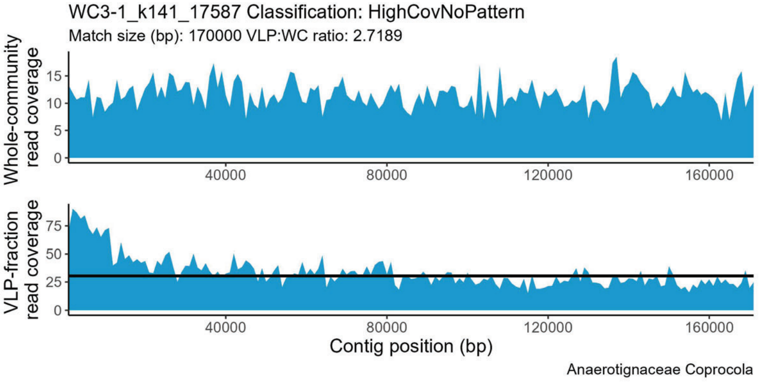
WC3-1_k141_17587 Classification: HighCovNoPattern
Match size (bp): 170000 VLP:WC ratio: 2.7189
Whole-community read coverage
0
5
10
15
40000
80000
120000
160000
VLP-fraction read coverage
0
25
50
75
Contig position (bp)
Anaerotignaceae Coprocola

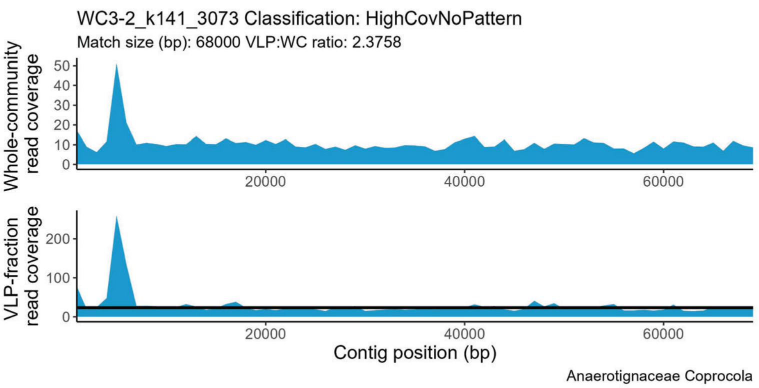
WC3-2_k141_3073 Classification: HighCovNoPattern
Match size (bp): 68000 VLP:WC ratio: 2.3758
Whole-community read coverage
VLP-fraction read coverage
Contig position (bp)
Anaerotignaceae Coprocola

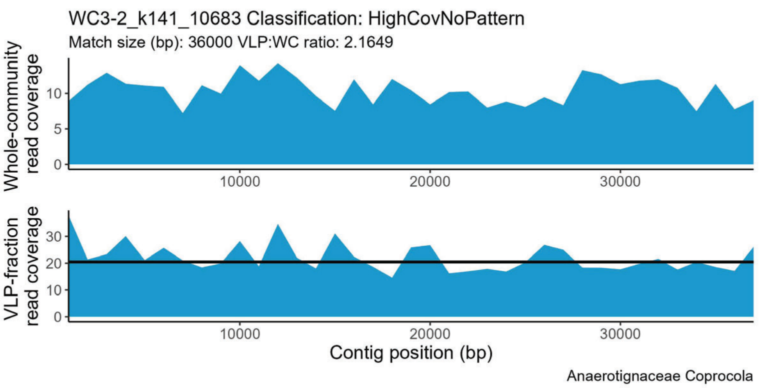
WC3-2_k141_10683 Classification: HighCovNoPattern
Match size (bp): 36000 VLP:WC ratio: 2.1649
Whole-community read coverage
VLP-fraction read coverage
Contig position (bp)
Anaerotignaceae Coprocola

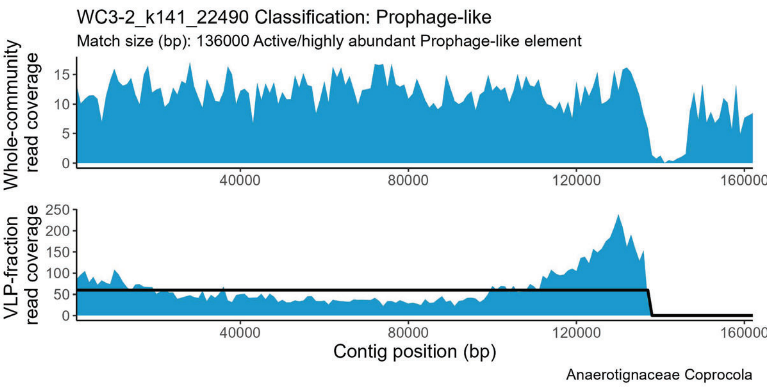
WC3-2_k141_22490 Classification: Prophage-like
Match size (bp): 136000 Active/highly abundant Prophage-like element
Whole-community read coverage
VLP-fraction read coverage
Contig position (bp)
Anaerotignaceae Coprocola

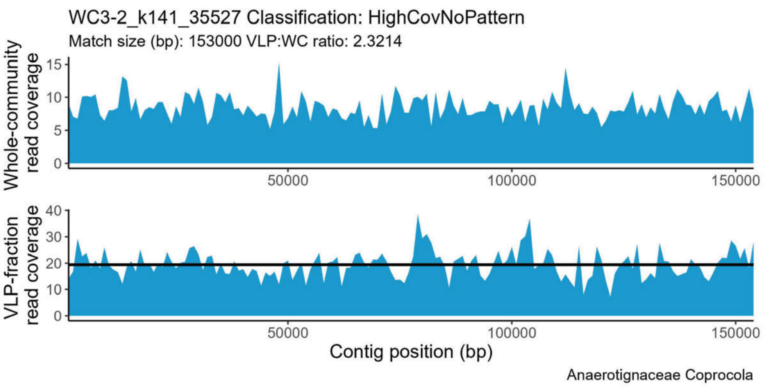
WC3-2_k141_35527 Classification: HighCovNoPattern
Match size (bp): 153000 VLP:WC ratio: 2.3214
Whole-community read coverage
VLP-fraction read coverage
Contig position (bp)
Anaerotignaceae Coprocola

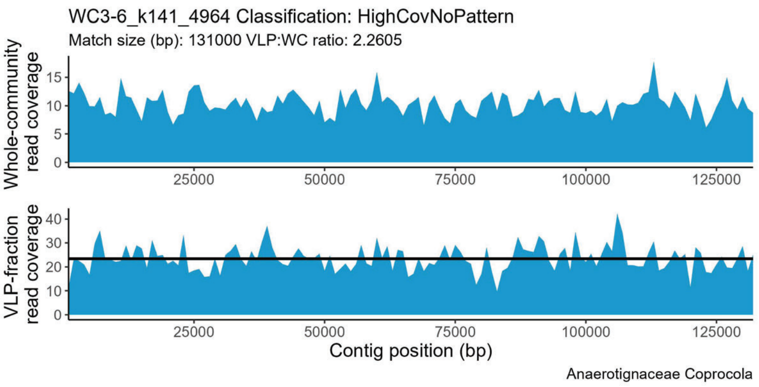
WC3-6_k141_4964 Classification: HighCovNoPattern
Match size (bp): 131000 VLP:WC ratio: 2.2605
Whole-community read coverage
VLP-fraction read coverage
Contig position (bp)
Anaerotignaceae Coprocola

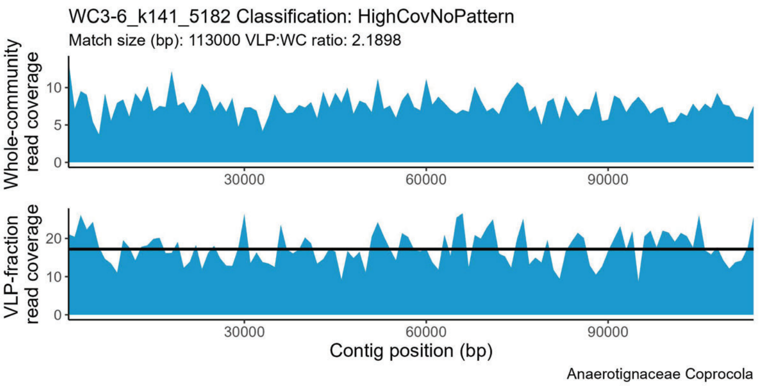
WC3-6_k141_5182 Classification: HighCovNoPattern
Match size (bp): 113000 VLP:WC ratio: 2.1898
Whole-community read coverage
VLP-fraction read coverage
Contig position (bp)
Anaerotignaceae Coprocola

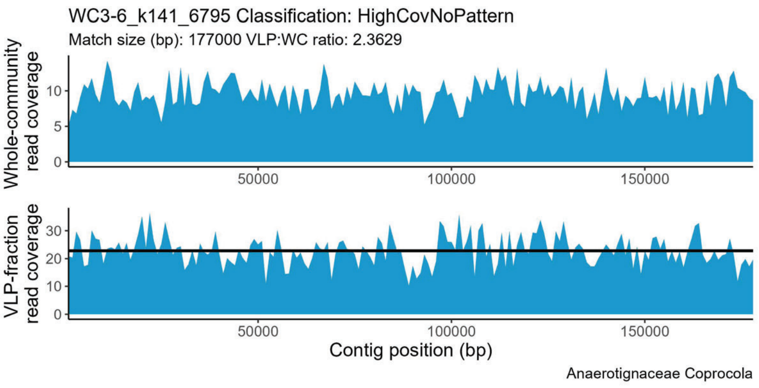
WC3-6_k141_6795 Classification: HighCovNoPattern
Match size (bp): 177000 VLP:WC ratio: 2.3629
Whole-community read coverage
VLP-fraction read coverage
Contig position (bp)
Anaerotignaceae Coprocola

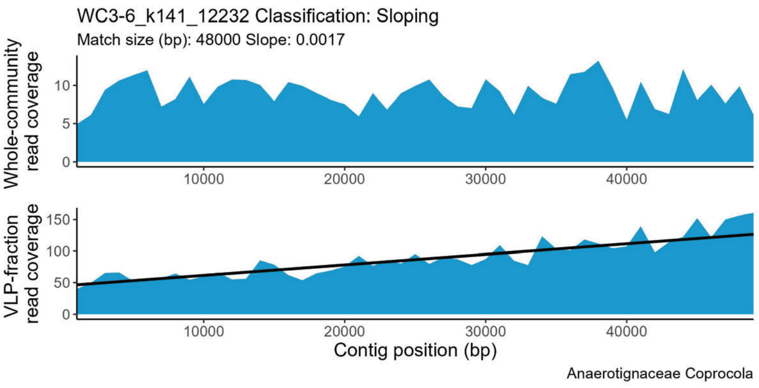
WC3-6_k141_12232 Classification: Sloping
Match size (bp): 48000 Slope: 0.0017
Whole-community read coverage
VLP-fraction read coverage
Contig position (bp)
Anaerotignaceae Coprocola

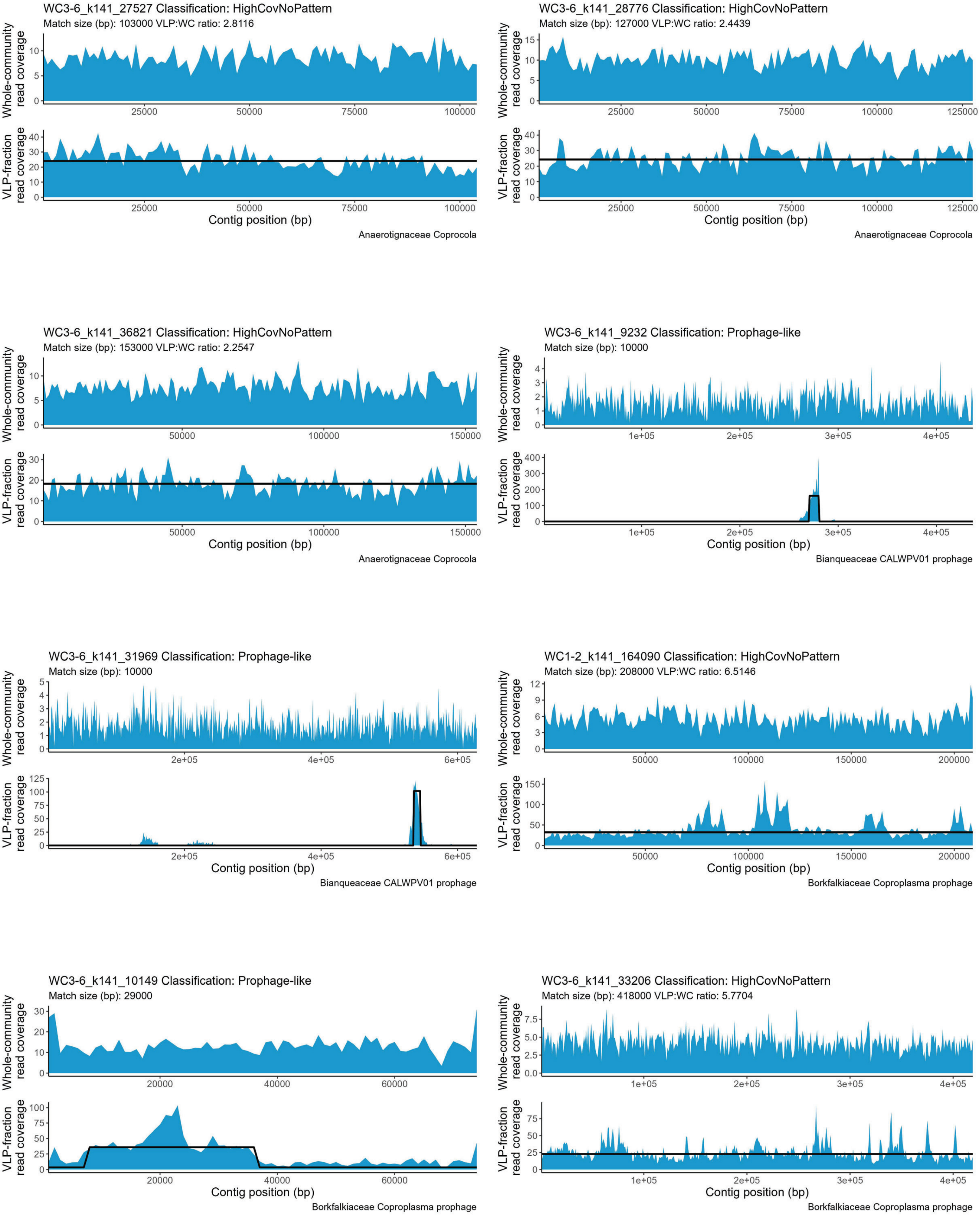
WC3-6_k141_27527 Classification: HighCovNoPattern
Match size (bp): 103000 VLP:WC ratio: 2.8116
Whole-community read coverage
VLP-fraction read coverage
Contig position (bp)
Anaerotignaceae Coprocola
WC3-6_k141_28776 Classification: HighCovNoPattern
Match size (bp): 127000 VLP:WC ratio: 2.4439
Whole-community read coverage
VLP-fraction read coverage
Contig position (bp)
Anaerotignaceae Coprocola
WC3-6_k141_36821 Classification: HighCovNoPattern
Match size (bp): 153000 VLP:WC ratio: 2.2547
Whole-community read coverage
VLP-fraction read coverage
Contig position (bp)
Anaerotignaceae Coprocola
WC3-6_k141_9232 Classification: Prophage-like
Match size (bp): 10000
Whole-community read coverage
VLP-fraction read coverage
Contig position (bp)
Bianqueaceae CALWPV01 prophage
WC3-6_k141_31969 Classification: Prophage-like
Match size (bp): 10000
Whole-community read coverage
VLP-fraction read coverage
Contig position (bp)
Bianqueaceae CALWPV01 prophage
WC1-2_k141_164090 Classification: HighCovNoPattern
Match size (bp): 208000 VLP:WC ratio: 6.5146
Whole-community read coverage
VLP-fraction read coverage
Contig position (bp)
Borkfalkiaceae Coproplasma prophage
WC3-6_k141_10149 Classification: Prophage-like
Match size (bp): 29000
Whole-community read coverage
VLP-fraction read coverage
Contig position (bp)
Borkfalkiaceae Coproplasma prophage
WC3-6_k141_33206 Classification: HighCovNoPattern
Match size (bp): 418000 VLP:WC ratio: 5.7704
Whole-community read coverage
VLP-fraction read coverage
Contig position (bp)
Borkfalkiaceae Coproplasma prophage

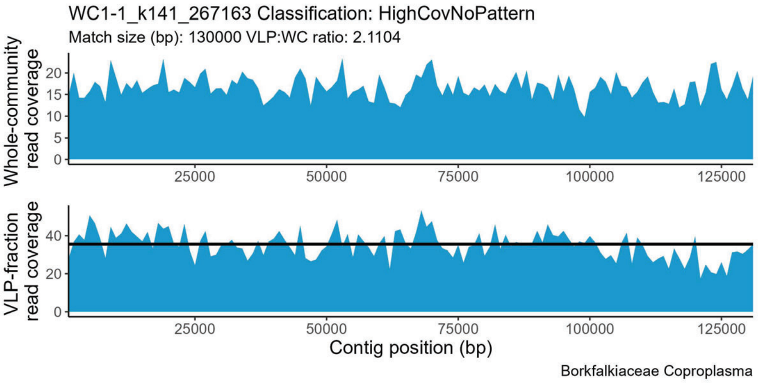
WC1-1_k141_267163 Classification: HighCovNoPattern
Match size (bp): 130000 VLP:WC ratio: 2.1104
Whole-community read coverage
VLP-fraction read coverage
Contig position (bp)
Borkfalkiaceae Coproplasma

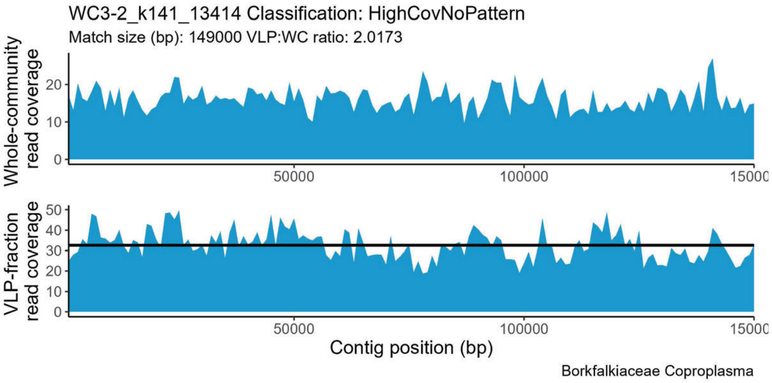
WC3-2_k141_13414 Classification: HighCovNoPattern
Match size (bp): 149000 VLP:WC ratio: 2.0173
Whole-community read coverage
VLP-fraction read coverage
Contig position (bp)
Borkfalkiaceae Coproplasma

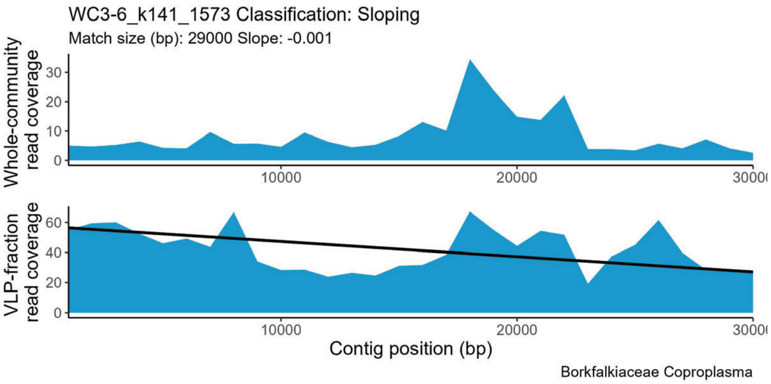
WC3-6_k141_1573 Classification: Sloping
Match size (bp): 29000 Slope: -0.001
Whole-community read coverage
VLP-fraction read coverage
Contig position (bp)
Borkfalkiaceae Coproplasma

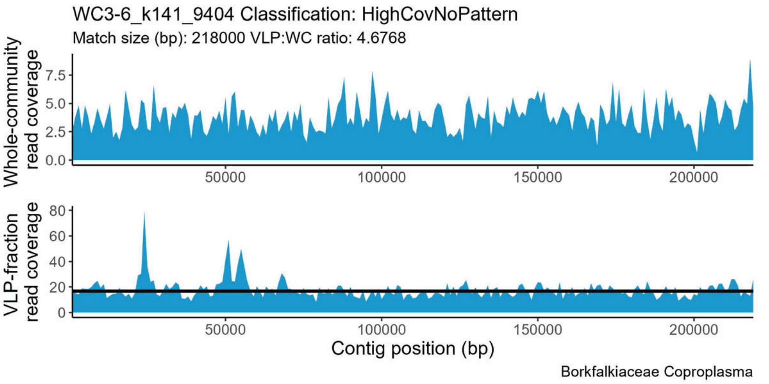
WC3-6_k141_9404 Classification: HighCovNoPattern
Match size (bp): 218000 VLP:WC ratio: 4.6768
Whole-community read coverage
VLP-fraction read coverage
Contig position (bp)
Borkfalkiaceae Coproplasma

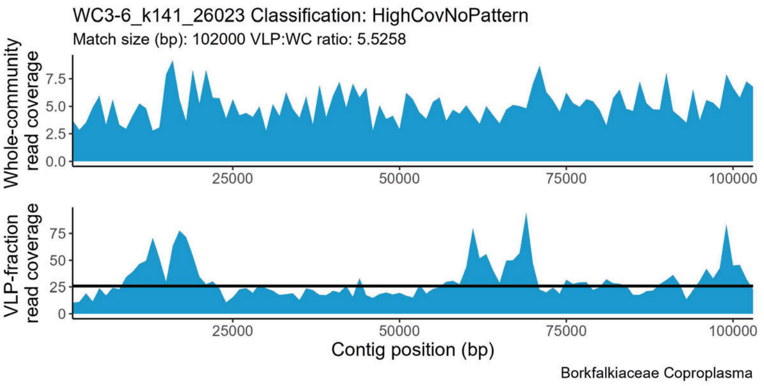
WC3-6_k141_26023 Classification: HighCovNoPattern
Match size (bp): 102000 VLP:WC ratio: 5.5258
Whole-community read coverage
VLP-fraction read coverage
Contig position (bp)
Borkfalkiaceae Coproplasma

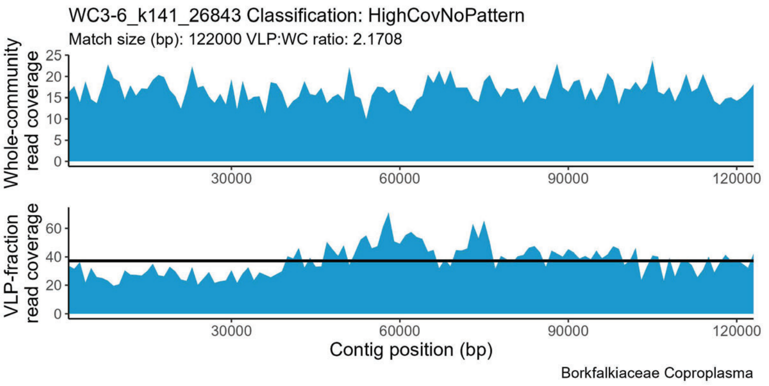
WC3-6_k141_26843 Classification: HighCovNoPattern
Match size (bp): 122000 VLP:WC ratio: 2.1708
Whole-community read coverage
VLP-fraction read coverage
Contig position (bp)
Borkfalkiaceae Coproplasma

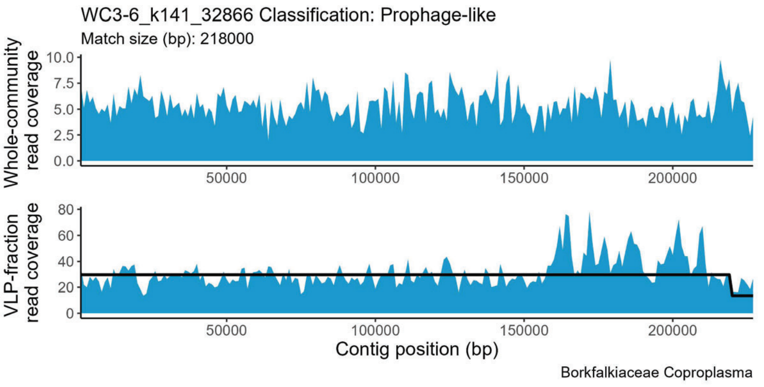
WC3-6_k141_32866 Classification: Prophage-like
Match size (bp): 218000
Whole-community read coverage
VLP-fraction read coverage
Contig position (bp)
Borkfalkiaceae Coproplasma

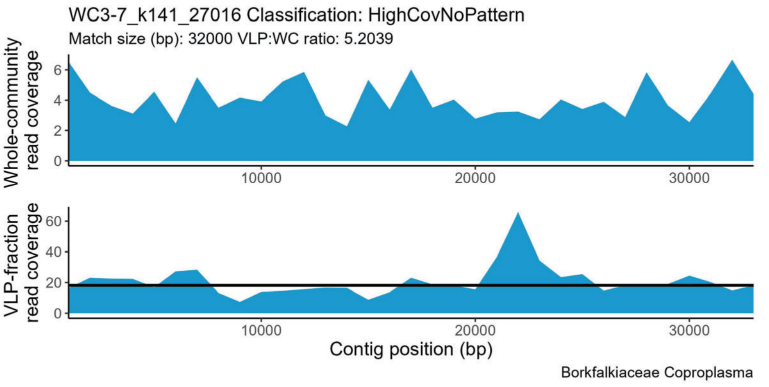
WC3-7_k141_27016 Classification: HighCovNoPattern
Match size (bp): 32000 VLP:WC ratio: 5.2039
Whole-community read coverage
VLP-fraction read coverage
Contig position (bp)
Borkfalkiaceae Coproplasma

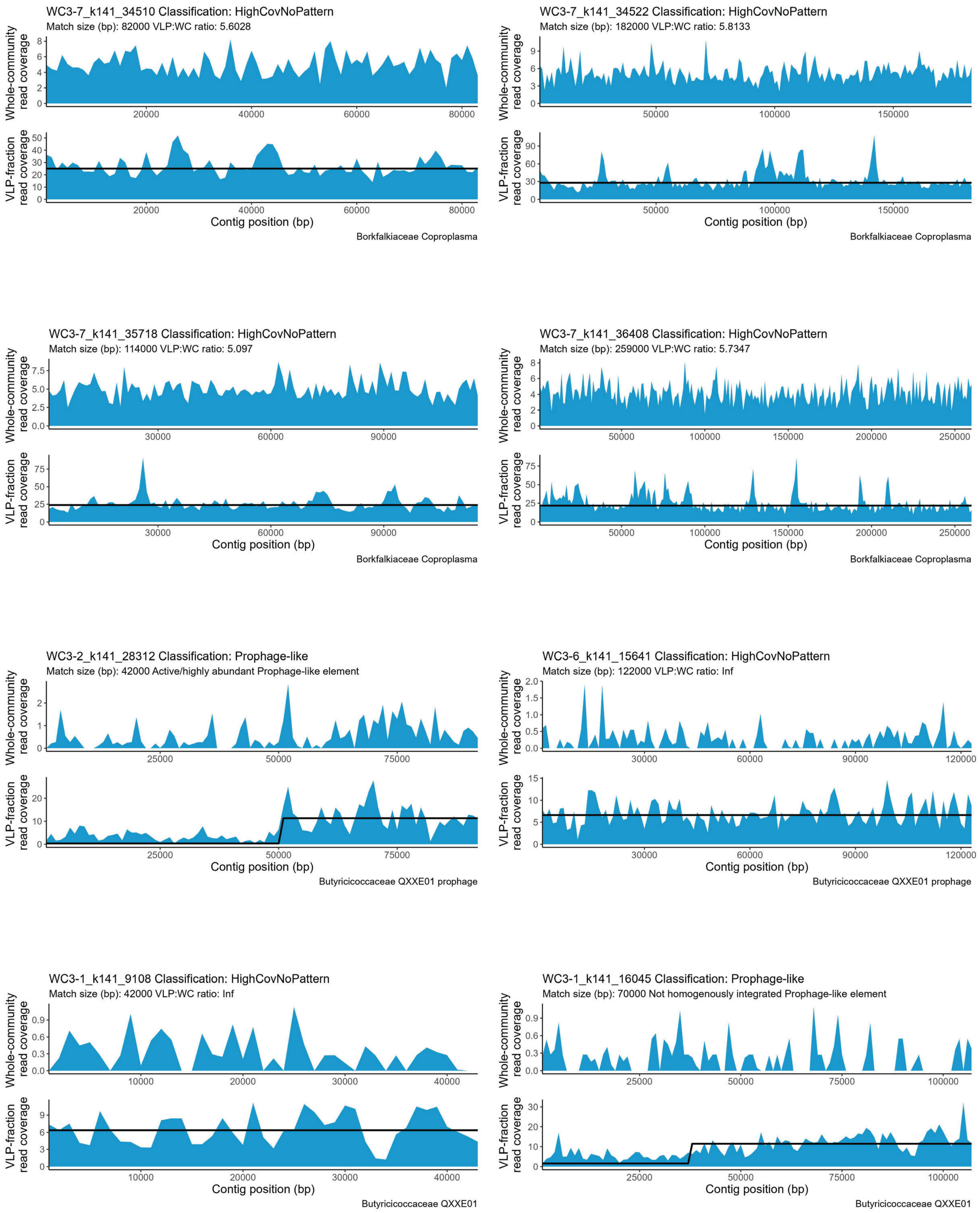
WC3-7_k141_34510 Classification: HighCovNoPattern
Match size (bp): 82000 VLP:WC ratio: 5.6028
Whole-community read coverage
VLP-fraction read coverage
Contig position (bp)
Borkfalkiaceae Coproplasma
WC3-7_k141_34522 Classification: HighCovNoPattern
Match size (bp): 182000 VLP:WC ratio: 5.8133
Whole-community read coverage
VLP-fraction read coverage
Contig position (bp)
Borkfalkiaceae Coproplasma
WC3-7_k141_35718 Classification: HighCovNoPattern
Match size (bp): 114000 VLP:WC ratio: 5.097
Whole-community read coverage
VLP-fraction read coverage
Contig position (bp)
Borkfalkiaceae Coproplasma
WC3-7_k141_36408 Classification: HighCovNoPattern
Match size (bp): 259000 VLP:WC ratio: 5.7347
Whole-community read coverage
VLP-fraction read coverage
Contig position (bp)
Borkfalkiaceae Coproplasma
WC3-2_k141_28312 Classification: Prophage-like
Match size (bp): 42000 Active/highly abundant Prophage-like element
Whole-community read coverage
VLP-fraction read coverage
Contig position (bp)
Butyricicoccaceae QXXE01 prophage
WC3-6_k141_15641 Classification: HighCovNoPattern
Match size (bp): 122000 VLP:WC ratio: Inf
Whole-community read coverage
VLP-fraction read coverage
Contig position (bp)
Butyricicoccaceae QXXE01 prophage
WC3-1_k141_9108 Classification: HighCovNoPattern
Match size (bp): 42000 VLP:WC ratio: Inf
Whole-community read coverage
VLP-fraction read coverage
Contig position (bp)
Butyricicoccaceae QXXE01
WC3-1_k141_16045 Classification: Prophage-like
Match size (bp): 70000 Not homogenously integrated Prophage-like element
Whole-community read coverage
VLP-fraction read coverage
Contig position (bp)
Butyricicoccaceae QXXE01

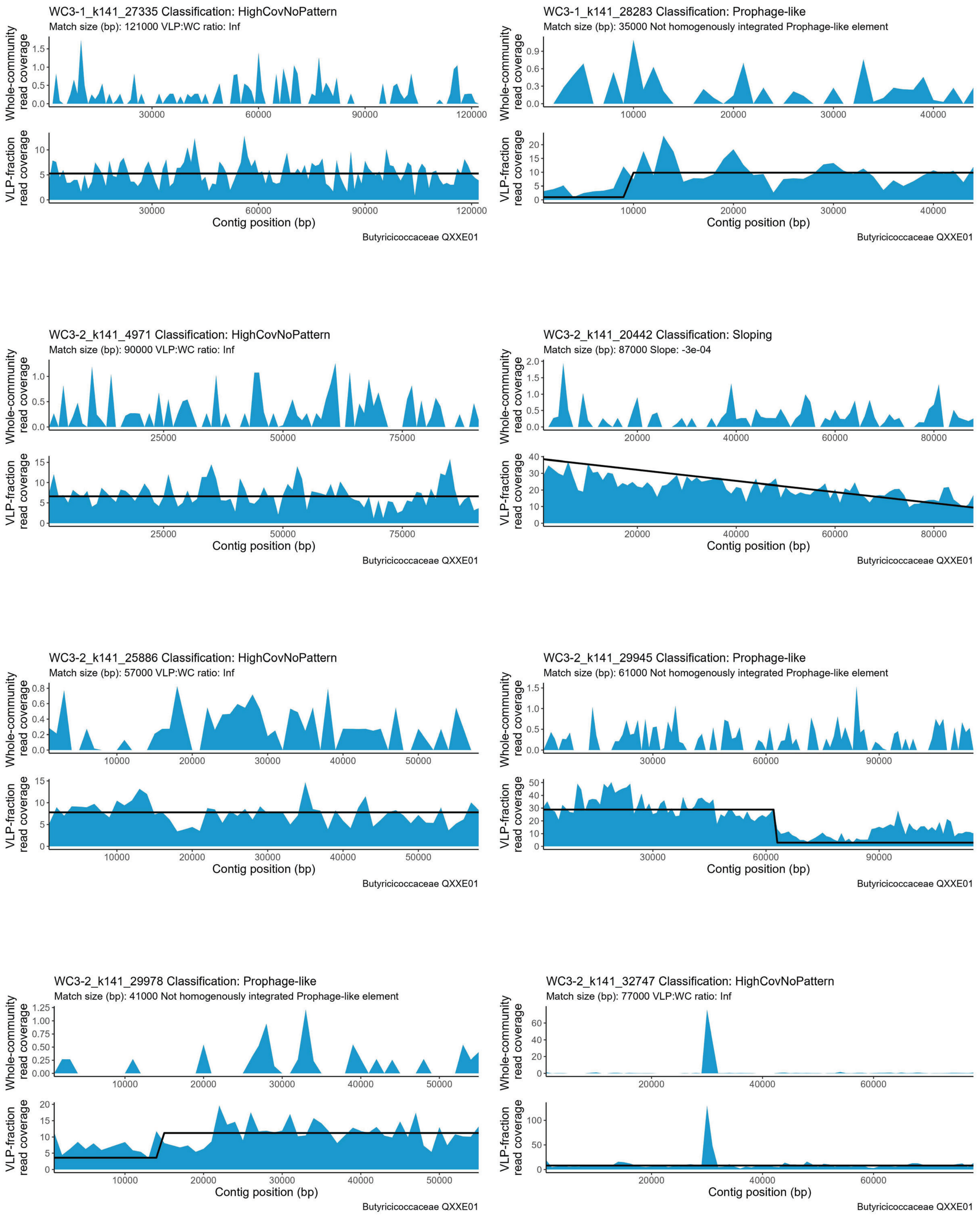
WC3-1_k141_27335 Classification: HighCovNoPattern
Match size (bp): 121000 VLP:WC ratio: Inf
WC3-1_k141_28283 Classification: Prophage-like
Match size (bp): 35000 Not homogenously integrated Prophage-like element
WC3-2_k141_4971 Classification: HighCovNoPattern
Match size (bp): 90000 VLP:WC ratio: Inf
WC3-2_k141_20442 Classification: Sloping
Match size (bp): 87000 Slope: -3e-04
WC3-2_k141_25886 Classification: HighCovNoPattern
Match size (bp): 57000 VLP:WC ratio: Inf
WC3-2_k141_29945 Classification: Prophage-like
Match size (bp): 61000 Not homogenously integrated Prophage-like element
WC3-2_k141_29978 Classification: Prophage-like
Match size (bp): 41000 Not homogenously integrated Prophage-like element
WC3-2_k141_32747 Classification: HighCovNoPattern
Match size (bp): 77000 VLP:WC ratio: Inf
Whole-community read coverage
VLP-fraction read coverage
Contig position (bp)
Butyricicoccaceae QXXE01

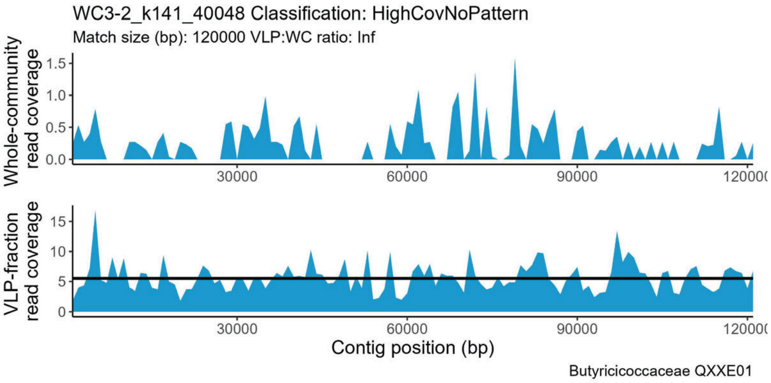
WC3-2_k141_40048 Classification: HighCovNoPattern
Match size (bp): 120000 VLP:WC ratio: Inf
Whole-community read coverage
VLP-fraction read coverage
Contig position (bp)
Butyricicoccaceae QXXE01

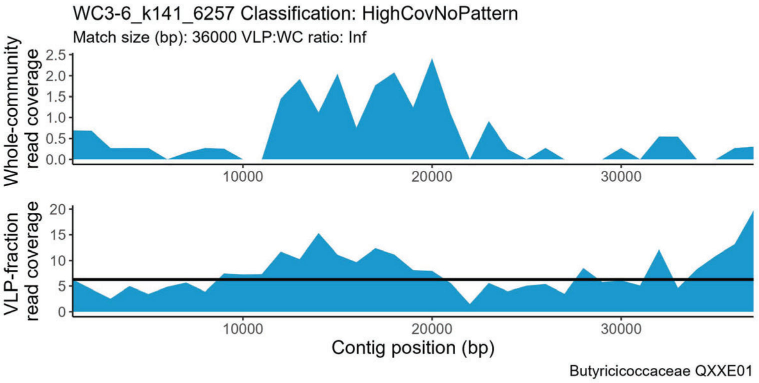
WC3-6_k141_6257 Classification: HighCovNoPattern
Match size (bp): 36000 VLP:WC ratio: Inf
Whole-community read coverage
VLP-fraction read coverage
Contig position (bp)
Butyricicoccaceae QXXE01

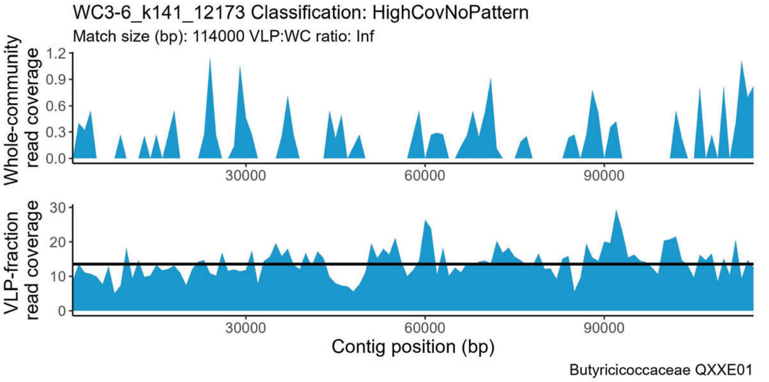
WC3-6_k141_12173 Classification: HighCovNoPattern
Match size (bp): 114000 VLP:WC ratio: Inf
Whole-community read coverage
VLP-fraction read coverage
Contig position (bp)
Butyricicoccaceae QXXE01

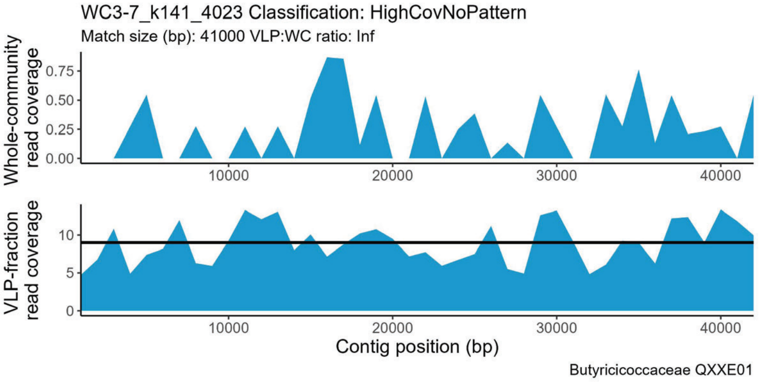
WC3-7_k141_4023 Classification: HighCovNoPattern
Match size (bp): 41000 VLP:WC ratio: Inf
Whole-community read coverage
VLP-fraction read coverage
Contig position (bp)
Butyricicoccaceae QXXE01

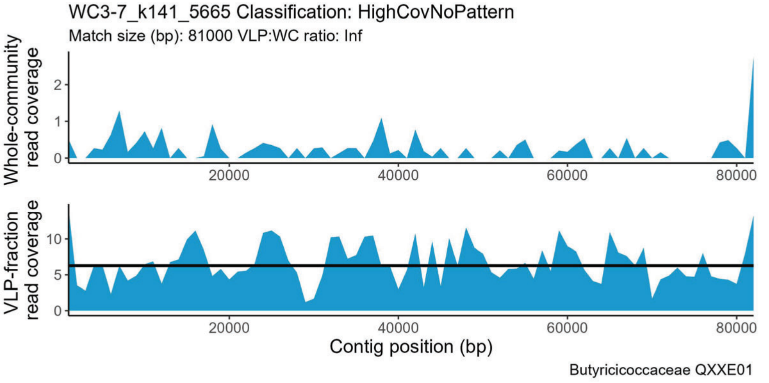
WC3-7_k141_5665 Classification: HighCovNoPattern
Match size (bp): 81000 VLP:WC ratio: Inf
Whole-community read coverage
VLP-fraction read coverage
Contig position (bp)
Butyricicoccaceae QXXE01

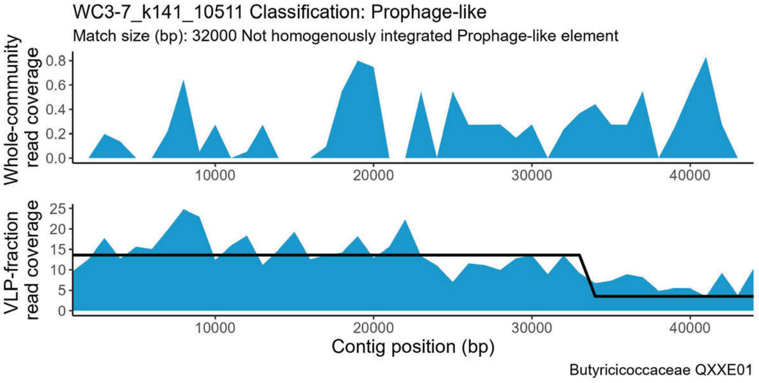
WC3-7_k141_10511 Classification: Prophage-like
Match size (bp): 32000 Not homogenously integrated Prophage-like element
Whole-community read coverage
VLP-fraction read coverage
Contig position (bp)
Butyricicoccaceae QXXE01

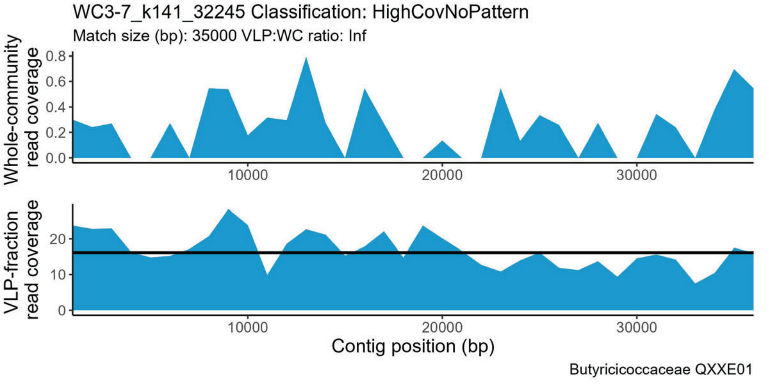
WC3-7_k141_32245 Classification: HighCovNoPattern
Match size (bp): 35000 VLP:WC ratio: Inf
Whole-community read coverage
VLP-fraction read coverage
Contig position (bp)
Butyricicoccaceae QXXE01

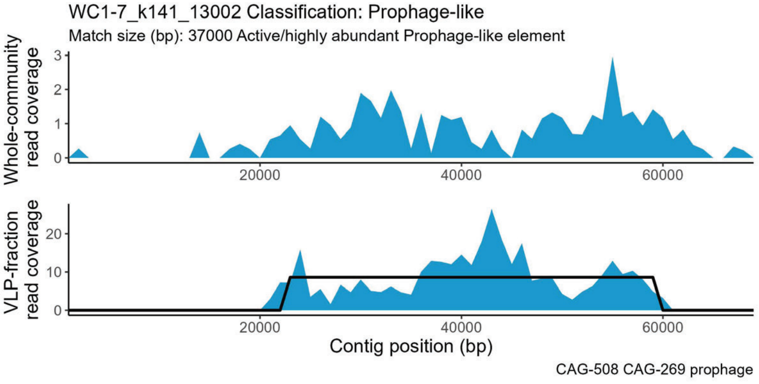
WC1-7_k141_13002 Classification: Prophage-like
Match size (bp): 37000 Active/highly abundant Prophage-like element
Whole-community read coverage
VLP-fraction read coverage
Contig position (bp)
CAG-508 CAG-269 prophage

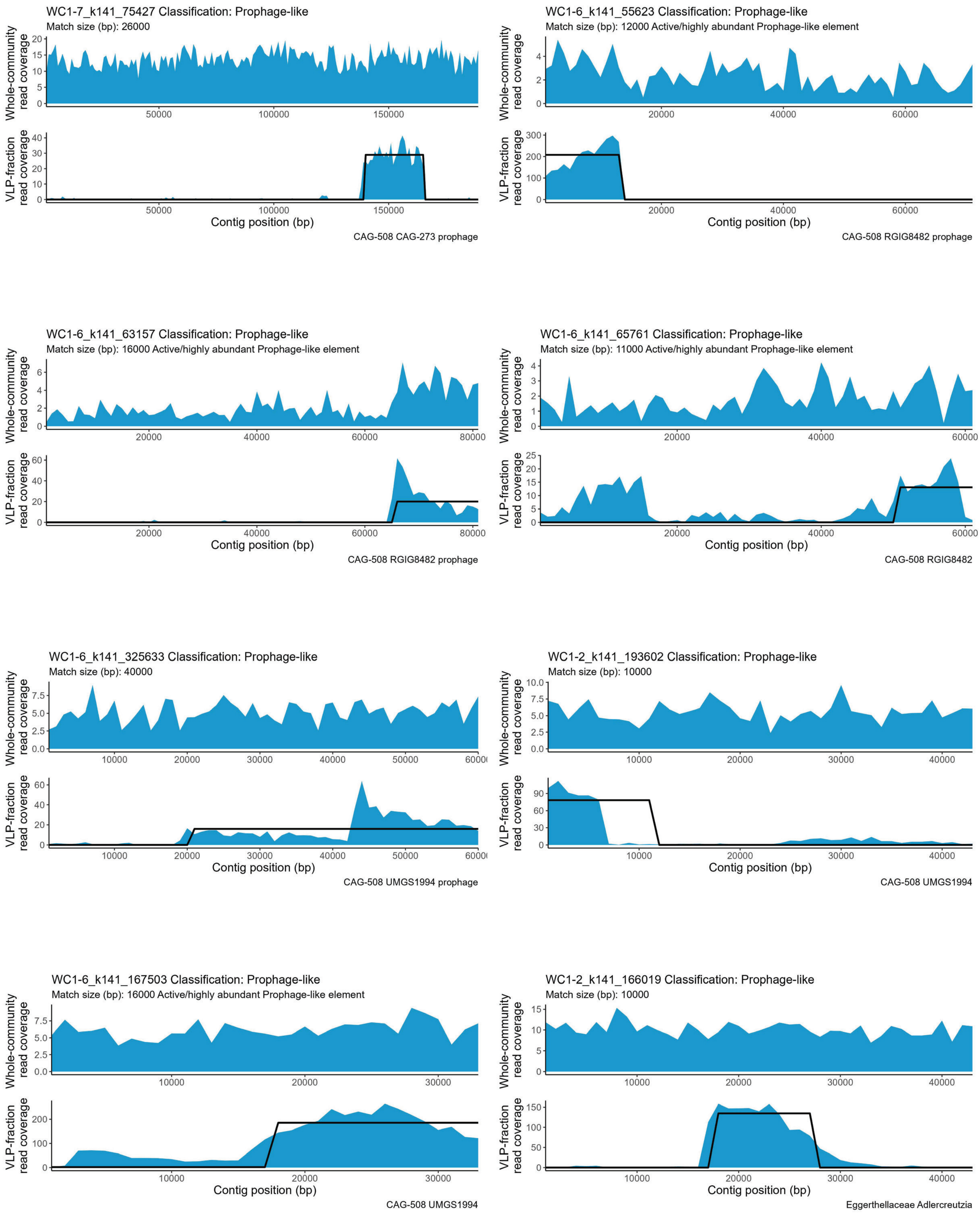
WC1-7_k141_75427 Classification: Prophage-like
Match size (bp): 26000
Whole-community read coverage
0 5 10 15 20
50000 100000 150000
VLP-fraction read coverage
0 10 20 30 40
50000 100000 150000
Contig position (bp)
CAG-508 CAG-273 prophage
WC1-6_k141_55623 Classification: Prophage-like
Match size (bp): 12000 Active/highly abundant Prophage-like element
Whole-community read coverage
0 2 4
20000 40000 60000
VLP-fraction read coverage
0 100 200 300
20000 40000 60000
Contig position (bp)
CAG-508 RGIG8482 prophage
WC1-6_k141_63157 Classification: Prophage-like
Match size (bp): 16000 Active/highly abundant Prophage-like element
Whole-community read coverage
0 2 4 6
20000 40000 60000 80000
VLP-fraction read coverage
0 20 40 60
20000 40000 60000 80000
Contig position (bp)
CAG-508 RGIG8482 prophage
WC1-6_k141_65761 Classification: Prophage-like
Match size (bp): 11000 Active/highly abundant Prophage-like element
Whole-community read coverage
0 1 2 3 4
20000 40000 60000
VLP-fraction read coverage
0 5 10 15 20 25
20000 40000 60000
Contig position (bp)
CAG-508 RGIG8482
WC1-6_k141_325633 Classification: Prophage-like
Match size (bp): 40000
Whole-community read coverage
0.0 2.5 5.0 7.5
10000 20000 30000 40000 50000 60000
VLP-fraction read coverage
0 20 40 60
10000 20000 30000 40000 50000 60000
Contig position (bp)
CAG-508 UMGS1994 prophage
WC1-2_k141_193602 Classification: Prophage-like
Match size (bp): 10000
Whole-community read coverage
0.0 2.5 5.0 7.5 10.0
10000 20000 30000 40000
VLP-fraction read coverage
0 30 60 90
10000 20000 30000 40000
Contig position (bp)
CAG-508 UMGS1994
WC1-6_k141_167503 Classification: Prophage-like
Match size (bp): 16000 Active/highly abundant Prophage-like element
Whole-community read coverage
0.0 2.5 5.0 7.5
10000 20000 30000
VLP-fraction read coverage
0 100 200
10000 20000 30000
Contig position (bp)
CAG-508 UMGS1994
WC1-2_k141_166019 Classification: Prophage-like
Match size (bp): 10000
Whole-community read coverage
0 5 10 15
10000 20000 30000 40000
VLP-fraction read coverage
0 50 100 150
10000 20000 30000 40000
Contig position (bp)
Eggerthellaceae Adlercreutzia

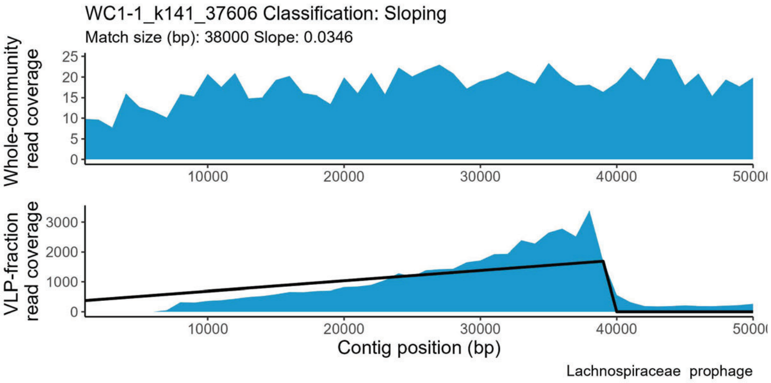
WC1-1_k141_37606 Classification: Sloping
Match size (bp): 38000 Slope: 0.0346
Whole-community read coverage
VLP-fraction read coverage
Contig position (bp)
Lachnospiraceae prophage

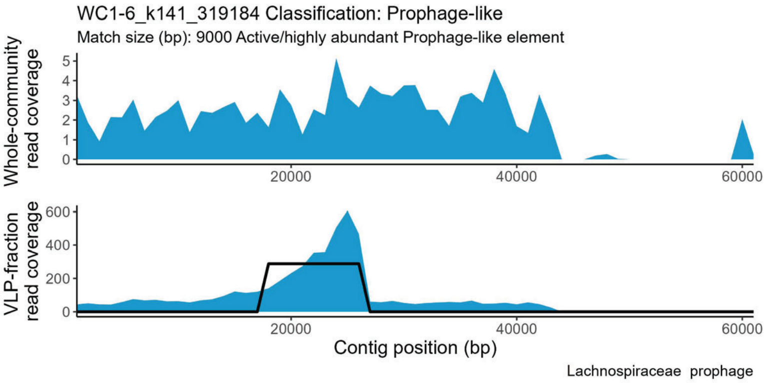
WC1-6_k141_319184 Classification: Prophage-like
Match size (bp): 9000 Active/highly abundant Prophage-like element
Whole-community read coverage
VLP-fraction read coverage
Contig position (bp)
Lachnospiraceae prophage

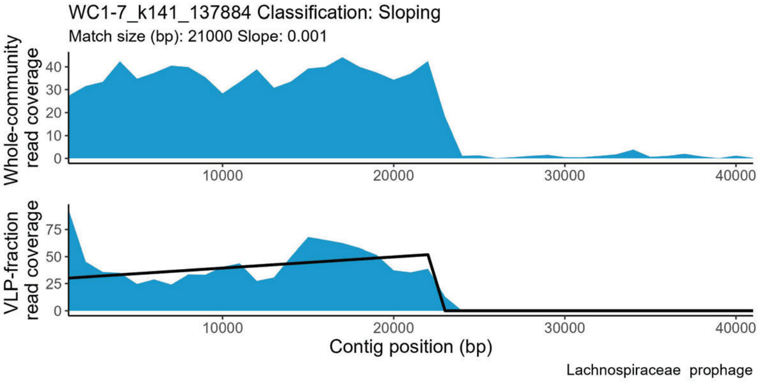
WC1-7_k141_137884 Classification: Sloping
Match size (bp): 21000 Slope: 0.001
Whole-community read coverage
VLP-fraction read coverage
Contig position (bp)
Lachnospiraceae prophage

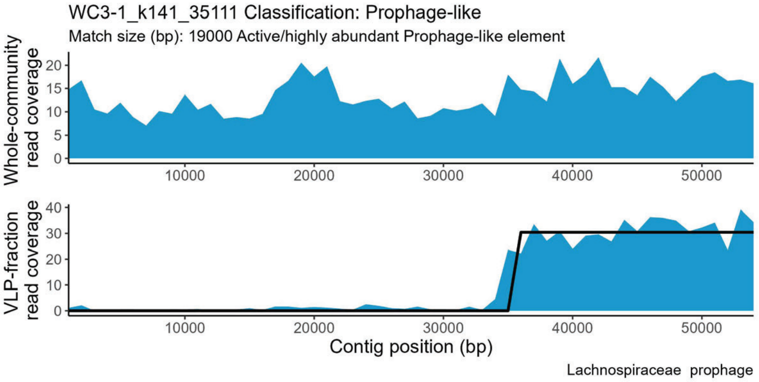
WC3-1_k141_35111 Classification: Prophage-like
Match size (bp): 19000 Active/highly abundant Prophage-like element
Whole-community read coverage
VLP-fraction read coverage
Contig position (bp)
Lachnospiraceae prophage

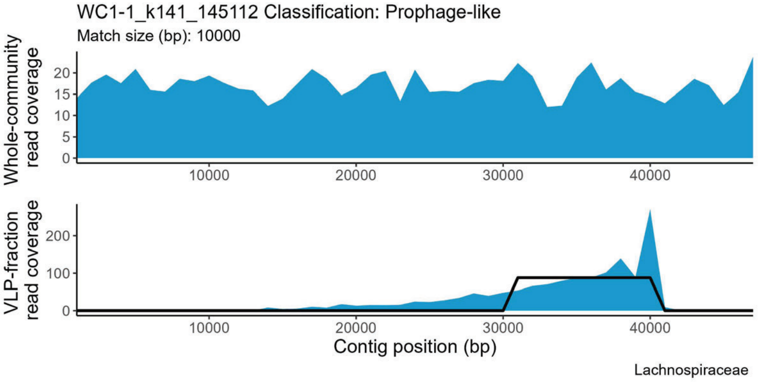
WC1-1_k141_145112 Classification: Prophage-like
Match size (bp): 10000
Whole-community read coverage
VLP-fraction read coverage
Contig position (bp)
Lachnospiraceae

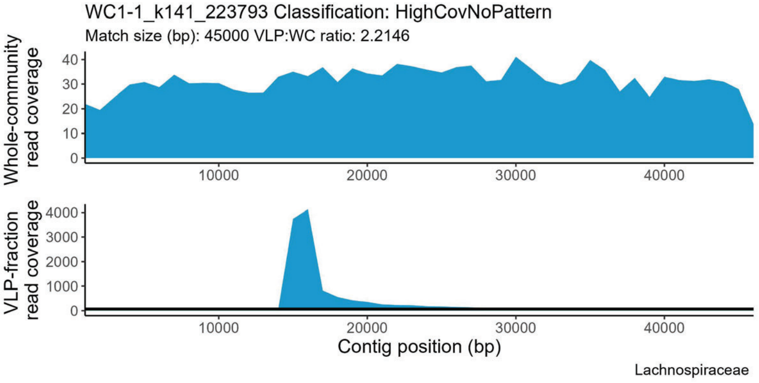
WC1-1_k141_223793 Classification: HighCovNoPattern
Match size (bp): 45000 VLP:WC ratio: 2.2146
Whole-community read coverage
VLP-fraction read coverage
Contig position (bp)
Lachnospiraceae

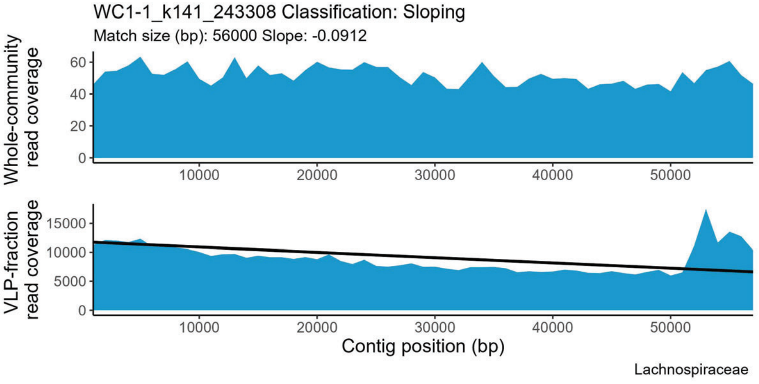
WC1-1_k141_243308 Classification: Sloping
Match size (bp): 56000 Slope: -0.0912
Whole-community read coverage
VLP-fraction read coverage
Contig position (bp)
Lachnospiraceae

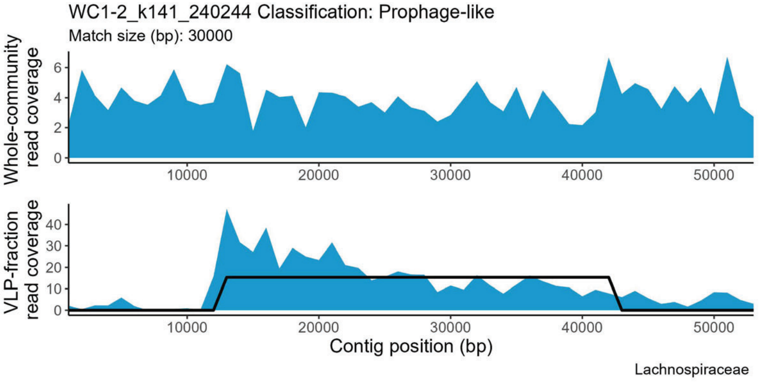
WC1-2_k141_240244 Classification: Prophage-like
Match size (bp): 30000
Whole-community read coverage
VLP-fraction read coverage
Contig position (bp)
Lachnospiraceae

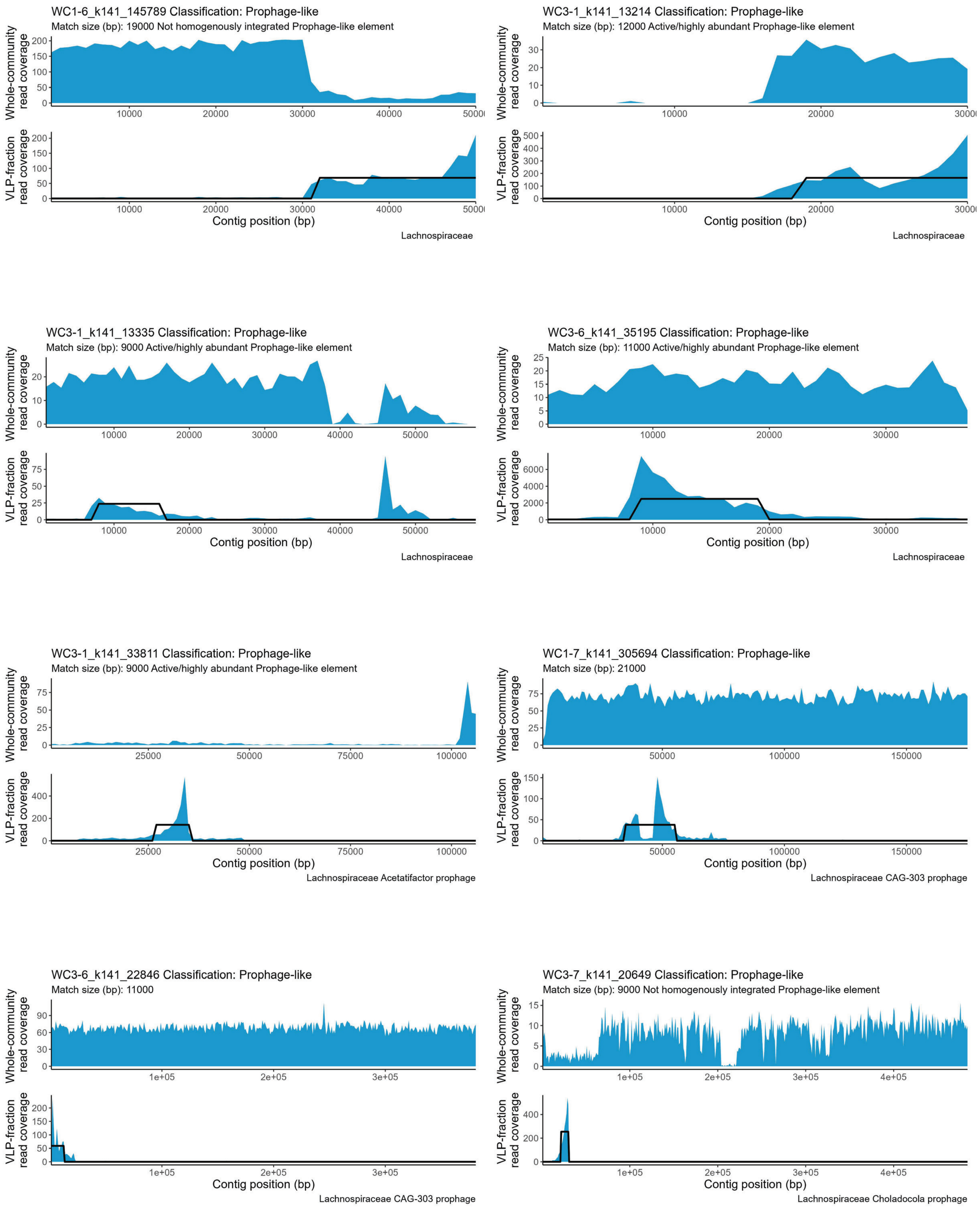

WC1-6_k141_145789 Classification: Prophage-like
Match size (bp): 19000 Not homogenously integrated Prophage-like element
Whole-community read coverage
VLP-fraction read coverage
Contig position (bp)
Lachnospiraceae
WC3-1_k141_13214 Classification: Prophage-like
Match size (bp): 12000 Active/highly abundant Prophage-like element
Whole-community read coverage
VLP-fraction read coverage
Contig position (bp)
Lachnospiraceae
WC3-1_k141_13335 Classification: Prophage-like
Match size (bp): 9000 Active/highly abundant Prophage-like element
Whole-community read coverage
VLP-fraction read coverage
Contig position (bp)
Lachnospiraceae
WC3-6_k141_35195 Classification: Prophage-like
Match size (bp): 11000 Active/highly abundant Prophage-like element
Whole-community read coverage
VLP-fraction read coverage
Contig position (bp)
Lachnospiraceae
WC3-1_k141_33811 Classification: Prophage-like
Match size (bp): 9000 Active/highly abundant Prophage-like element
Whole-community read coverage
VLP-fraction read coverage
Contig position (bp)
Lachnospiraceae Acetatifactor prophage
WC1-7_k141_305694 Classification: Prophage-like
Match size (bp): 21000
Whole-community read coverage
VLP-fraction read coverage
Contig position (bp)
Lachnospiraceae CAG-303 prophage
WC3-6_k141_22846 Classification: Prophage-like
Match size (bp): 11000
Whole-community read coverage
VLP-fraction read coverage
Contig position (bp)
Lachnospiraceae CAG-303 prophage
WC3-7_k141_20649 Classification: Prophage-like
Match size (bp): 9000 Not homogenously integrated Prophage-like element
Whole-community read coverage
VLP-fraction read coverage
Contig position (bp)
Lachnospiraceae Choladocola prophage

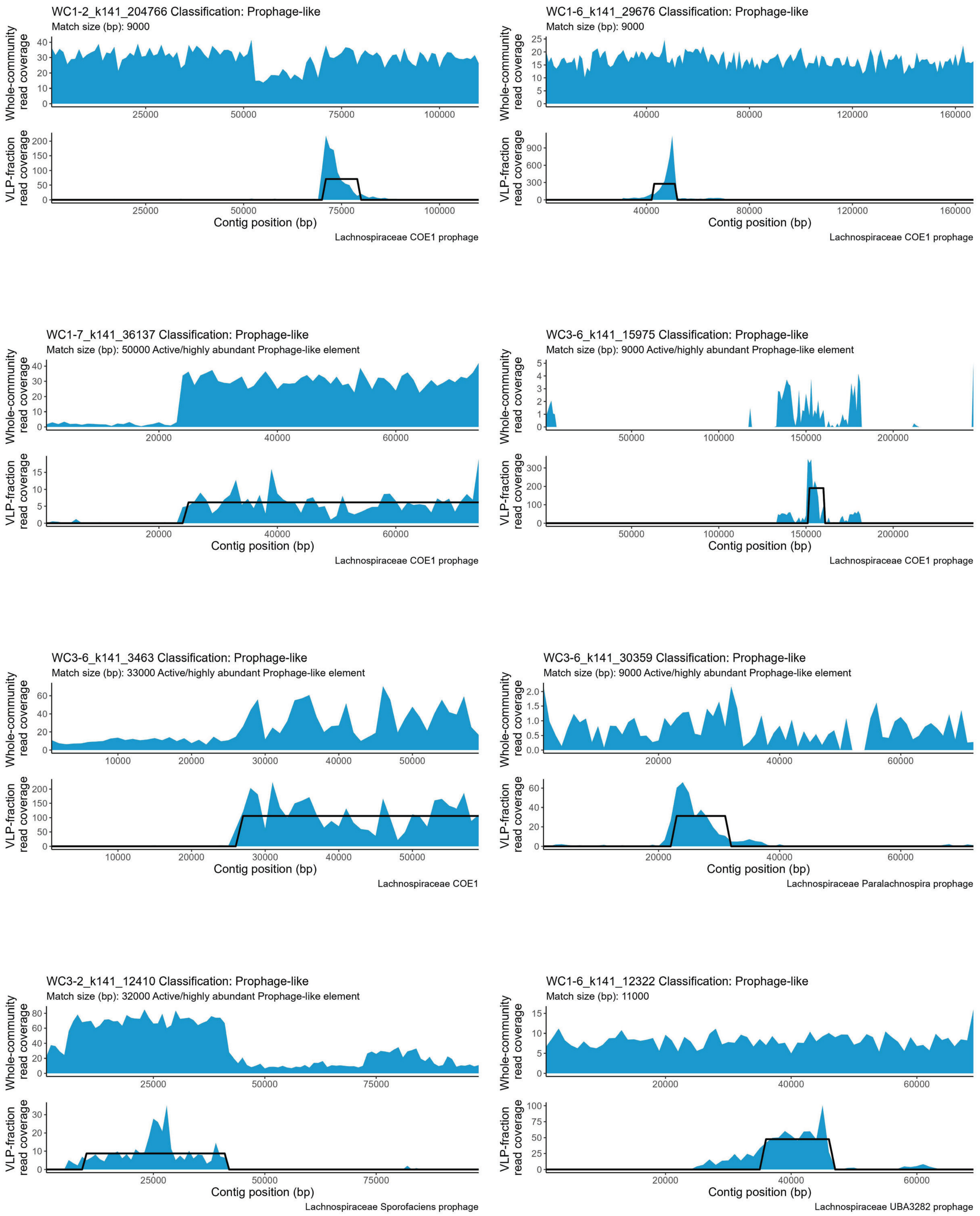

WC1-2_k141_204766 Classification: Prophage-like
Match size (bp): 9000
Whole-community read coverage
VLP-fraction read coverage
Contig position (bp)
Lachnospiraceae COE1 prophage
WC1-6_k141_29676 Classification: Prophage-like
Match size (bp): 9000
Whole-community read coverage
VLP-fraction read coverage
Contig position (bp)
Lachnospiraceae COE1 prophage
WC1-7_k141_36137 Classification: Prophage-like
Match size (bp): 50000 Active/highly abundant Prophage-like element
Whole-community read coverage
VLP-fraction read coverage
Contig position (bp)
Lachnospiraceae COE1 prophage
WC3-6_k141_15975 Classification: Prophage-like
Match size (bp): 9000 Active/highly abundant Prophage-like element
Whole-community read coverage
VLP-fraction read coverage
Contig position (bp)
Lachnospiraceae COE1 prophage
WC3-6_k141_3463 Classification: Prophage-like
Match size (bp): 33000 Active/highly abundant Prophage-like element
Whole-community read coverage
VLP-fraction read coverage
Contig position (bp)
Lachnospiraceae COE1
WC3-6_k141_30359 Classification: Prophage-like
Match size (bp): 9000 Active/highly abundant Prophage-like element
Whole-community read coverage
VLP-fraction read coverage
Contig position (bp)
Lachnospiraceae Paralachnospira prophage
WC3-2_k141_12410 Classification: Prophage-like
Match size (bp): 32000 Active/highly abundant Prophage-like element
Whole-community read coverage
VLP-fraction read coverage
Contig position (bp)
Lachnospiraceae Sporofaciens prophage
WC1-6_k141_12322 Classification: Prophage-like
Match size (bp): 11000
Whole-community read coverage
VLP-fraction read coverage
Contig position (bp)
Lachnospiraceae UBA3282 prophage

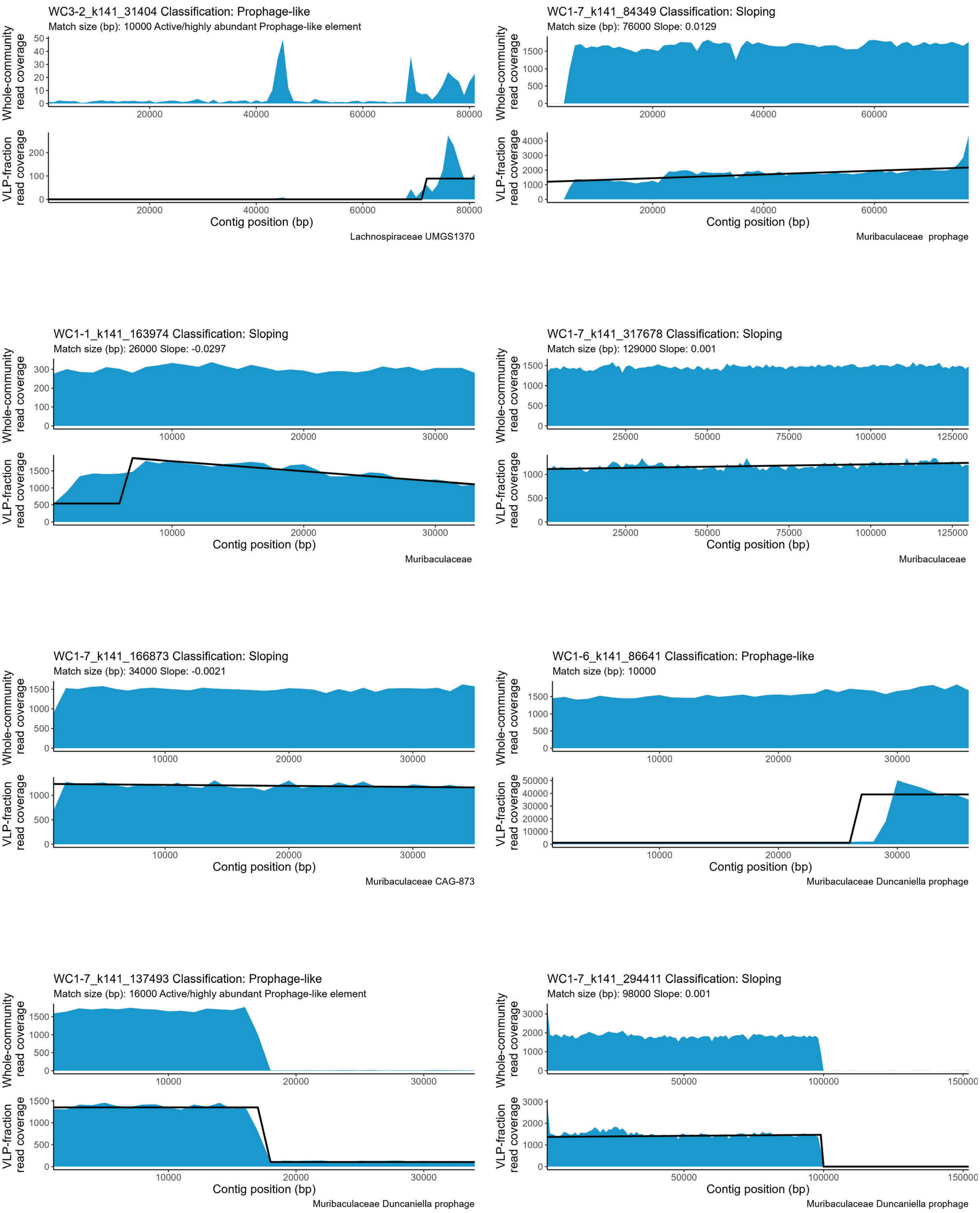
WC3-2_k141_31404 Classification: Prophage-like
Match size (bp): 10000 Active/highly abundant Prophage-like element
Whole-community read coverage
VLP-fraction read coverage
Contig position (bp)
Lachnospiraceae UMGS1370
WC1-7_k141_84349 Classification: Sloping
Match size (bp): 76000 Slope: 0.0129
Whole-community read coverage
VLP-fraction read coverage
Contig position (bp)
Muribaculaceae prophage
WC1-1_k141_163974 Classification: Sloping
Match size (bp): 26000 Slope: -0.0297
Whole-community read coverage
VLP-fraction read coverage
Contig position (bp)
Muribaculaceae
WC1-7_k141_317678 Classification: Sloping
Match size (bp): 129000 Slope: 0.001
Whole-community read coverage
VLP-fraction read coverage
Contig position (bp)
Muribaculaceae
WC1-7_k141_166873 Classification: Sloping
Match size (bp): 34000 Slope: -0.0021
Whole-community read coverage
VLP-fraction read coverage
Contig position (bp)
Muribaculaceae CAG-873
WC1-6_k141_86641 Classification: Prophage-like
Match size (bp): 10000
Whole-community read coverage
VLP-fraction read coverage
Contig position (bp)
Muribaculaceae Duncaniella prophage
WC1-7_k141_137493 Classification: Prophage-like
Match size (bp): 16000 Active/highly abundant Prophage-like element
Whole-community read coverage
VLP-fraction read coverage
Contig position (bp)
Muribaculaceae Duncaniella prophage
WC1-7_k141_294411 Classification: Sloping
Match size (bp): 98000 Slope: 0.001
Whole-community read coverage
VLP-fraction read coverage
Contig position (bp)
Muribaculaceae Duncaniella prophage

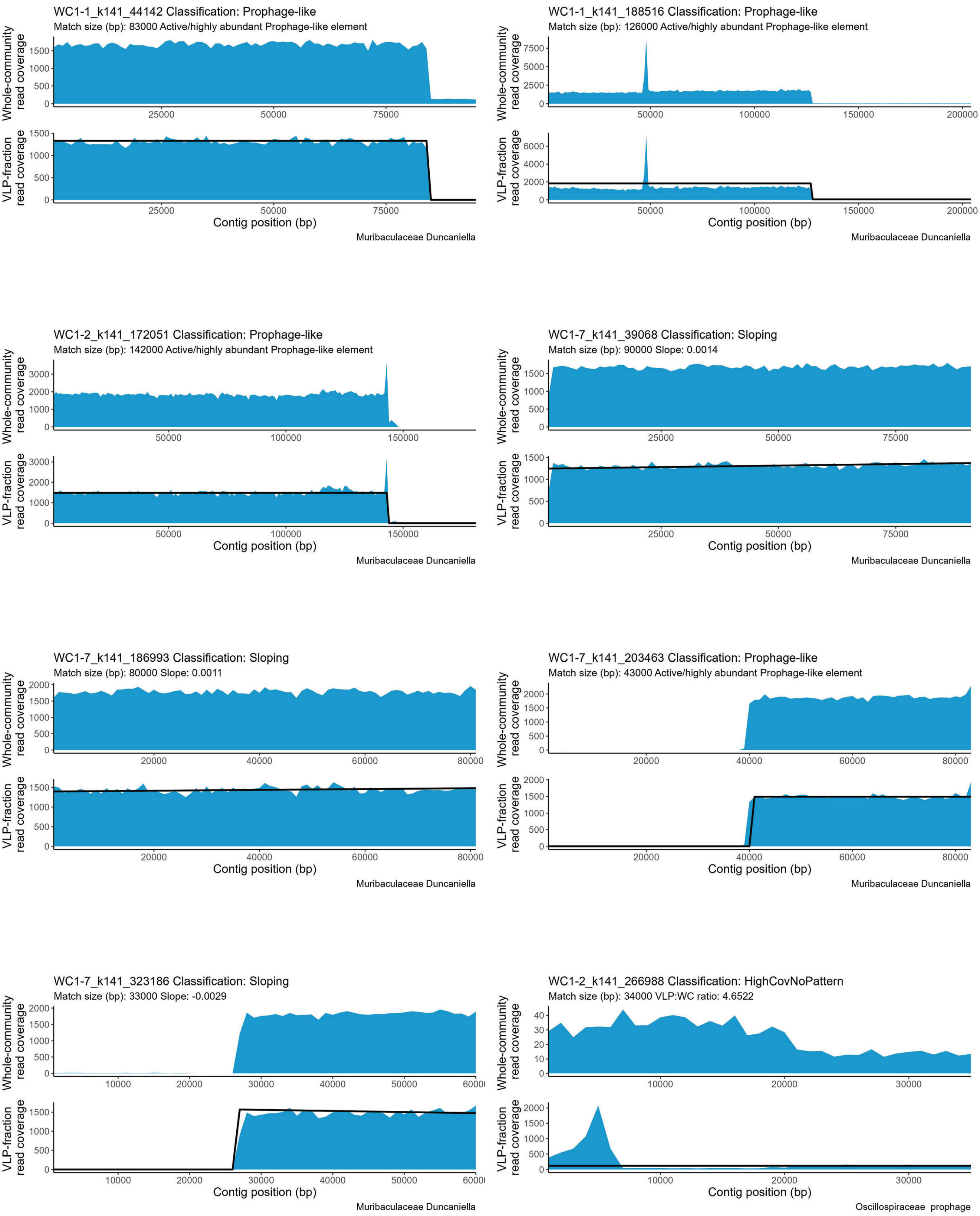

WC1-1_k141_44142 Classification: Prophage-like
Match size (bp): 83000 Active/highly abundant Prophage-like element
Whole-community read coverage
VLP-fraction read coverage
Contig position (bp)
Muribaculaceae Duncaniella
WC1-1_k141_188516 Classification: Prophage-like
Match size (bp): 126000 Active/highly abundant Prophage-like element
Whole-community read coverage
VLP-fraction read coverage
Contig position (bp)
Muribaculaceae Duncaniella
WC1-2_k141_172051 Classification: Prophage-like
Match size (bp): 142000 Active/highly abundant Prophage-like element
Whole-community read coverage
VLP-fraction read coverage
Contig position (bp)
Muribaculaceae Duncaniella
WC1-7_k141_39068 Classification: Sloping
Match size (bp): 90000 Slope: 0.0014
Whole-community read coverage
VLP-fraction read coverage
Contig position (bp)
Muribaculaceae Duncaniella
WC1-7_k141_186993 Classification: Sloping
Match size (bp): 80000 Slope: 0.0011
Whole-community read coverage
VLP-fraction read coverage
Contig position (bp)
Muribaculaceae Duncaniella
WC1-7_k141_203463 Classification: Prophage-like
Match size (bp): 43000 Active/highly abundant Prophage-like element
Whole-community read coverage
VLP-fraction read coverage
Contig position (bp)
Muribaculaceae Duncaniella
WC1-7_k141_323186 Classification: Sloping
Match size (bp): 33000 Slope: -0.0029
Whole-community read coverage
VLP-fraction read coverage
Contig position (bp)
Muribaculaceae Duncaniella
WC1-2_k141_266988 Classification: HighCovNoPattern
Match size (bp): 34000 VLP:WC ratio: 4.6522
Whole-community read coverage
VLP-fraction read coverage
Contig position (bp)
Oscillospiraceae prophage

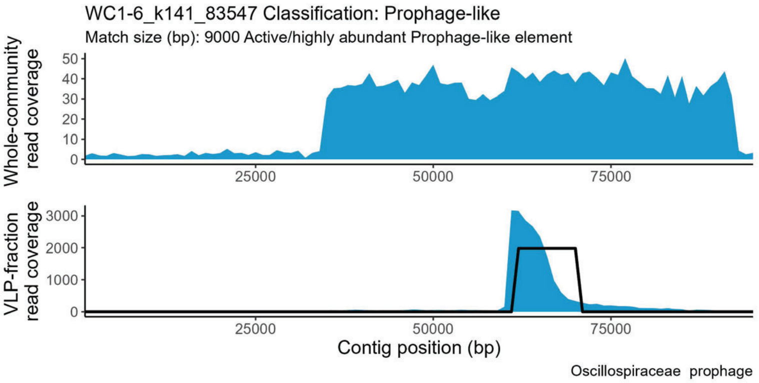
WC1-6_k141_83547 Classification: Prophage-like
Match size (bp): 9000 Active/highly abundant Prophage-like element
Whole-community read coverage
VLP-fraction read coverage
25000
50000
75000
Contig position (bp)
Oscillospiraceae prophage

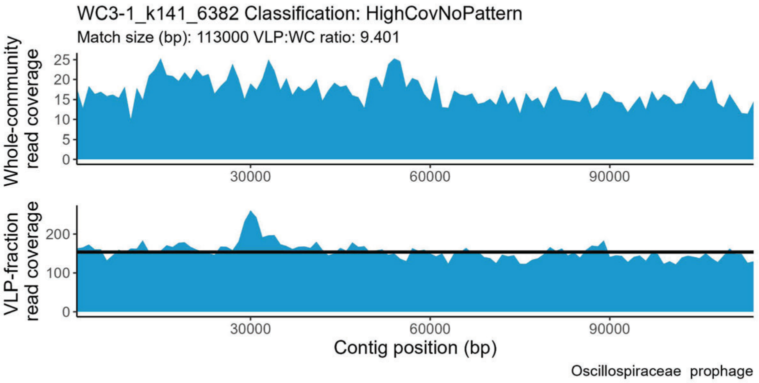
WC3-1_k141_6382 Classification: HighCovNoPattern
Match size (bp): 113000 VLP:WC ratio: 9.401
Whole-community read coverage
VLP-fraction read coverage
30000
60000
90000
Contig position (bp)
Oscillospiraceae prophage

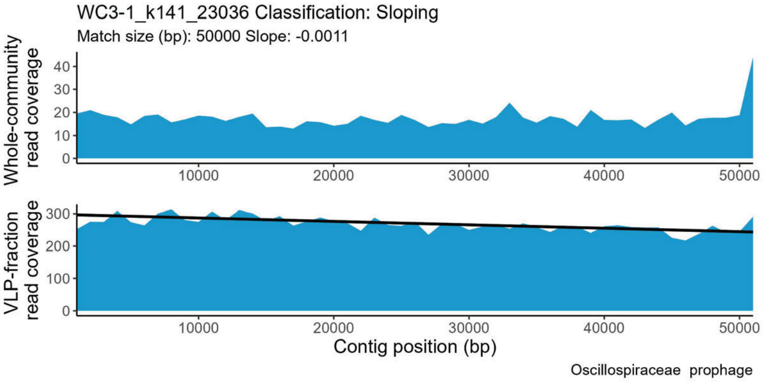
WC3-1_k141_23036 Classification: Sloping
Match size (bp): 50000 Slope: -0.0011
Whole-community read coverage
VLP-fraction read coverage
10000
20000
30000
40000
50000
Contig position (bp)
Oscillospiraceae prophage

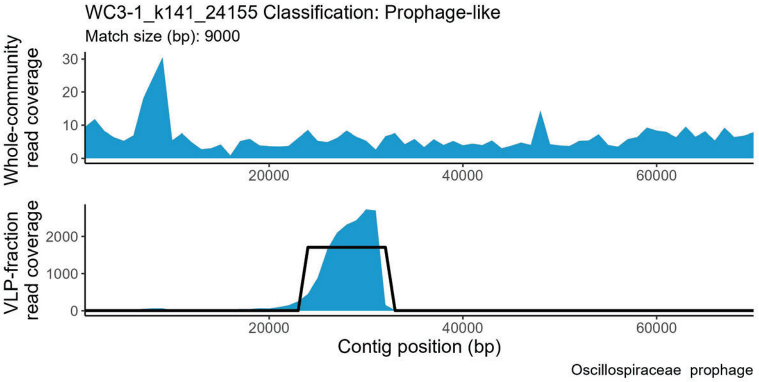
WC3-1_k141_24155 Classification: Prophage-like
Match size (bp): 9000
Whole-community read coverage
VLP-fraction read coverage
20000
40000
60000
Contig position (bp)
Oscillospiraceae prophage

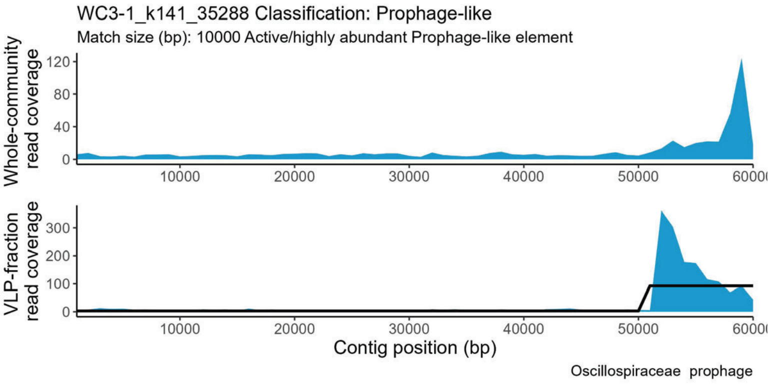
WC3-1_k141_35288 Classification: Prophage-like
Match size (bp): 10000 Active/highly abundant Prophage-like element
Whole-community read coverage
VLP-fraction read coverage
10000
20000
30000
40000
50000
Contig position (bp)
Oscillospiraceae prophage

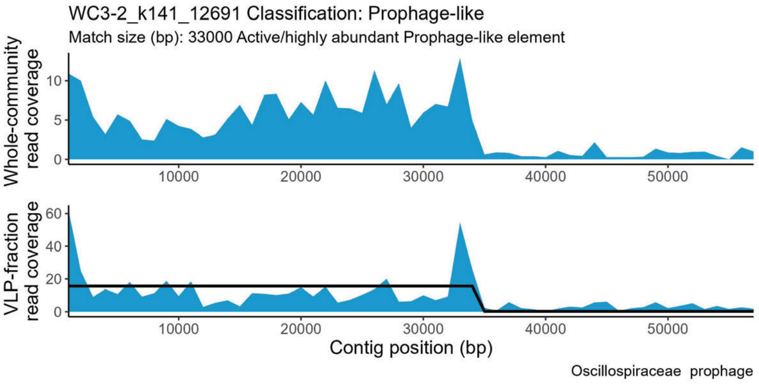
WC3-2_k141_12691 Classification: Prophage-like
Match size (bp): 33000 Active/highly abundant Prophage-like element
Whole-community read coverage
VLP-fraction read coverage
10000
20000
30000
40000
50000
Contig position (bp)
Oscillospiraceae prophage

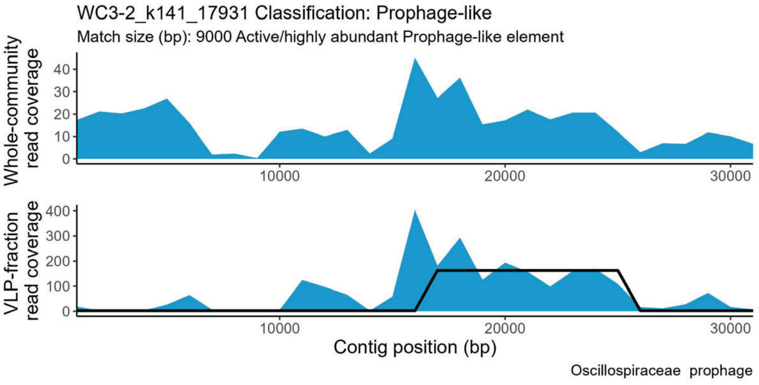
WC3-2_k141_17931 Classification: Prophage-like
Match size (bp): 9000 Active/highly abundant Prophage-like element
Whole-community read coverage
VLP-fraction read coverage
10000
20000
30000
Contig position (bp)
Oscillospiraceae prophage

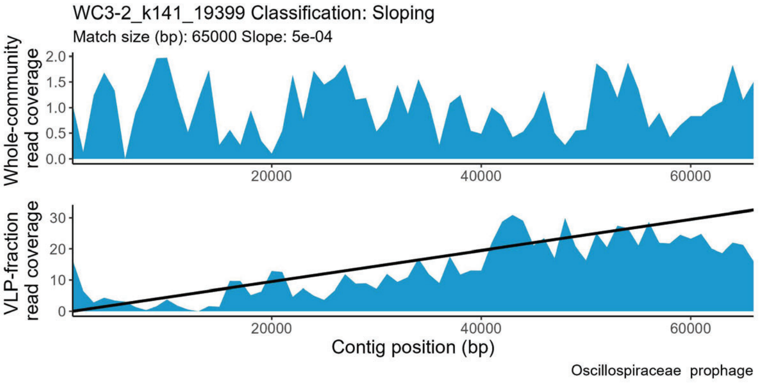
WC3-2_k141_19399 Classification: Sloping
Match size (bp): 65000 Slope: 5e-04
Whole-community read coverage
VLP-fraction read coverage
20000
40000
60000
Contig position (bp)
Oscillospiraceae prophage

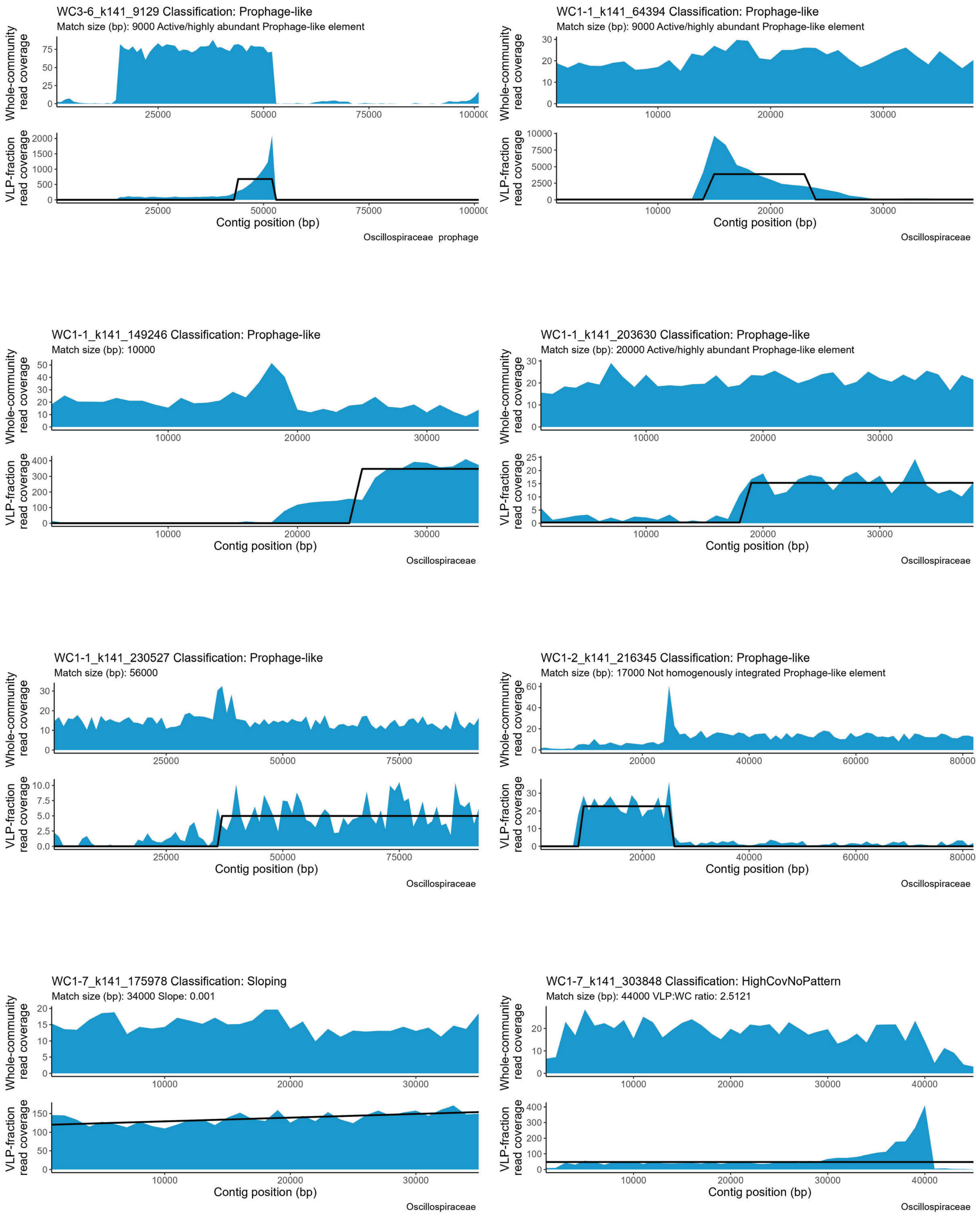

WC3-6_k141_9129 Classification: Prophage-like
Match size (bp): 9000 Active/highly abundant Prophage-like element
Whole-community read coverage
VLP-fraction read coverage
Contig position (bp)
Oscillospiraceae prophage
WC1-1_k141_64394 Classification: Prophage-like
Match size (bp): 9000 Active/highly abundant Prophage-like element
Whole-community read coverage
VLP-fraction read coverage
Contig position (bp)
Oscillospiraceae
WC1-1_k141_149246 Classification: Prophage-like
Match size (bp): 10000
Whole-community read coverage
VLP-fraction read coverage
Contig position (bp)
Oscillospiraceae
WC1-1_k141_203630 Classification: Prophage-like
Match size (bp): 20000 Active/highly abundant Prophage-like element
Whole-community read coverage
VLP-fraction read coverage
Contig position (bp)
Oscillospiraceae
WC1-1_k141_230527 Classification: Prophage-like
Match size (bp): 56000
Whole-community read coverage
VLP-fraction read coverage
Contig position (bp)
Oscillospiraceae
WC1-2_k141_216345 Classification: Prophage-like
Match size (bp): 17000 Not homogenously integrated Prophage-like element
Whole-community read coverage
VLP-fraction read coverage
Contig position (bp)
Oscillospiraceae
WC1-7_k141_175978 Classification: Sloping
Match size (bp): 34000 Slope: 0.001
Whole-community read coverage
VLP-fraction read coverage
Contig position (bp)
Oscillospiraceae
WC1-7_k141_303848 Classification: HighCovNoPattern
Match size (bp): 44000 VLP:WC ratio: 2.5121
Whole-community read coverage
VLP-fraction read coverage
Contig position (bp)
Oscillospiraceae

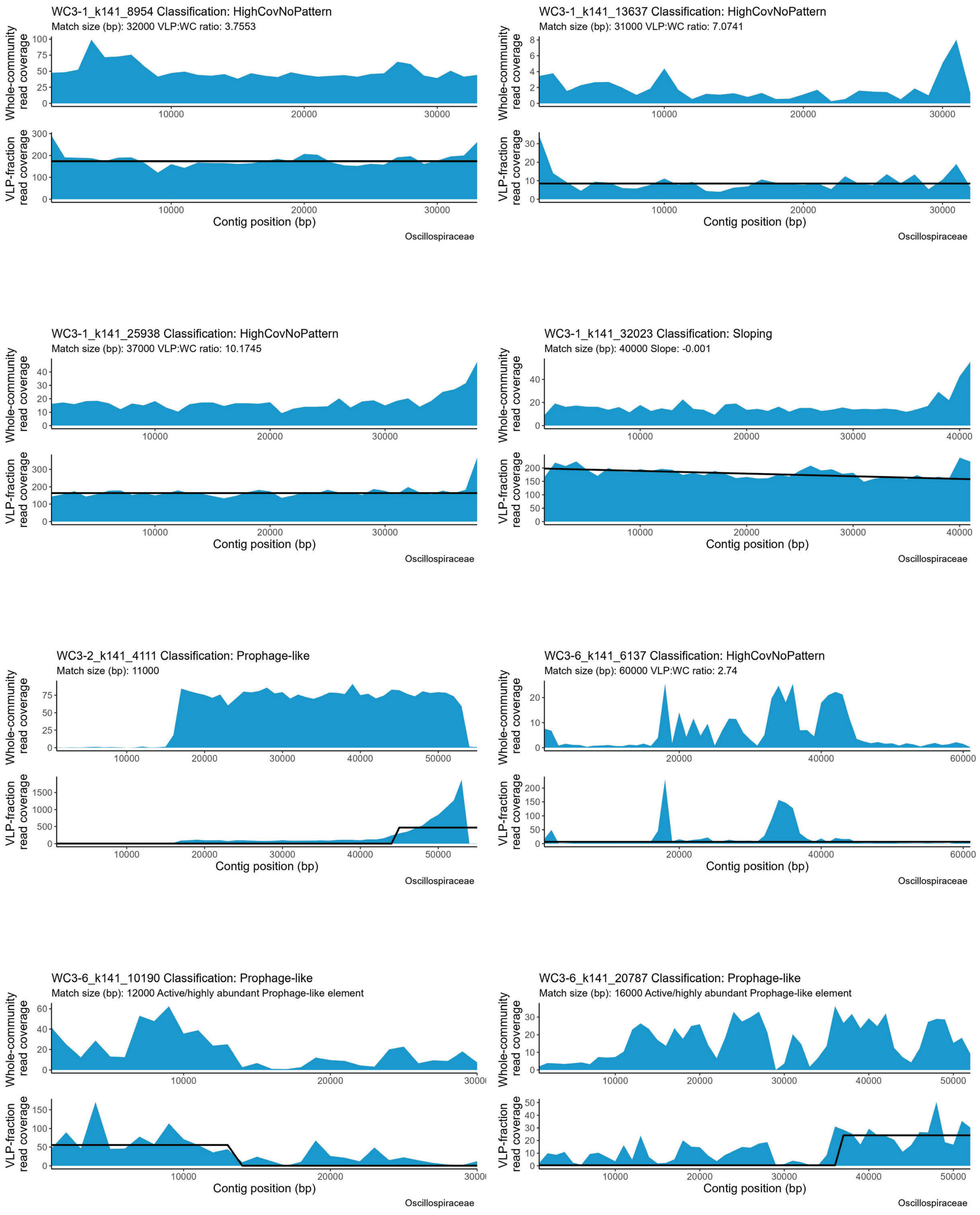
WC3-1_k141_8954 Classification: HighCovNoPattern
Match size (bp): 32000 VLP:WC ratio: 3.7553
Whole-community read coverage
VLP-fraction read coverage
Contig position (bp)
Oscillospiraceae
WC3-1_k141_13637 Classification: HighCovNoPattern
Match size (bp): 31000 VLP:WC ratio: 7.0741
Whole-community read coverage
VLP-fraction read coverage
Contig position (bp)
Oscillospiraceae
WC3-1_k141_25938 Classification: HighCovNoPattern
Match size (bp): 37000 VLP:WC ratio: 10.1745
Whole-community read coverage
VLP-fraction read coverage
Contig position (bp)
Oscillospiraceae
WC3-1_k141_32023 Classification: Sloping
Match size (bp): 40000 Slope: -0.001
Whole-community read coverage
VLP-fraction read coverage
Contig position (bp)
Oscillospiraceae
WC3-2_k141_4111 Classification: Prophage-like
Match size (bp): 11000
Whole-community read coverage
VLP-fraction read coverage
Contig position (bp)
Oscillospiraceae
WC3-6_k141_6137 Classification: HighCovNoPattern
Match size (bp): 60000 VLP:WC ratio: 2.74
Whole-community read coverage
VLP-fraction read coverage
Contig position (bp)
Oscillospiraceae
WC3-6_k141_10190 Classification: Prophage-like
Match size (bp): 12000 Active/highly abundant Prophage-like element
Whole-community read coverage
VLP-fraction read coverage
Contig position (bp)
Oscillospiraceae
WC3-6_k141_20787 Classification: Prophage-like
Match size (bp): 16000 Active/highly abundant Prophage-like element
Whole-community read coverage
VLP-fraction read coverage
Contig position (bp)
Oscillospiraceae

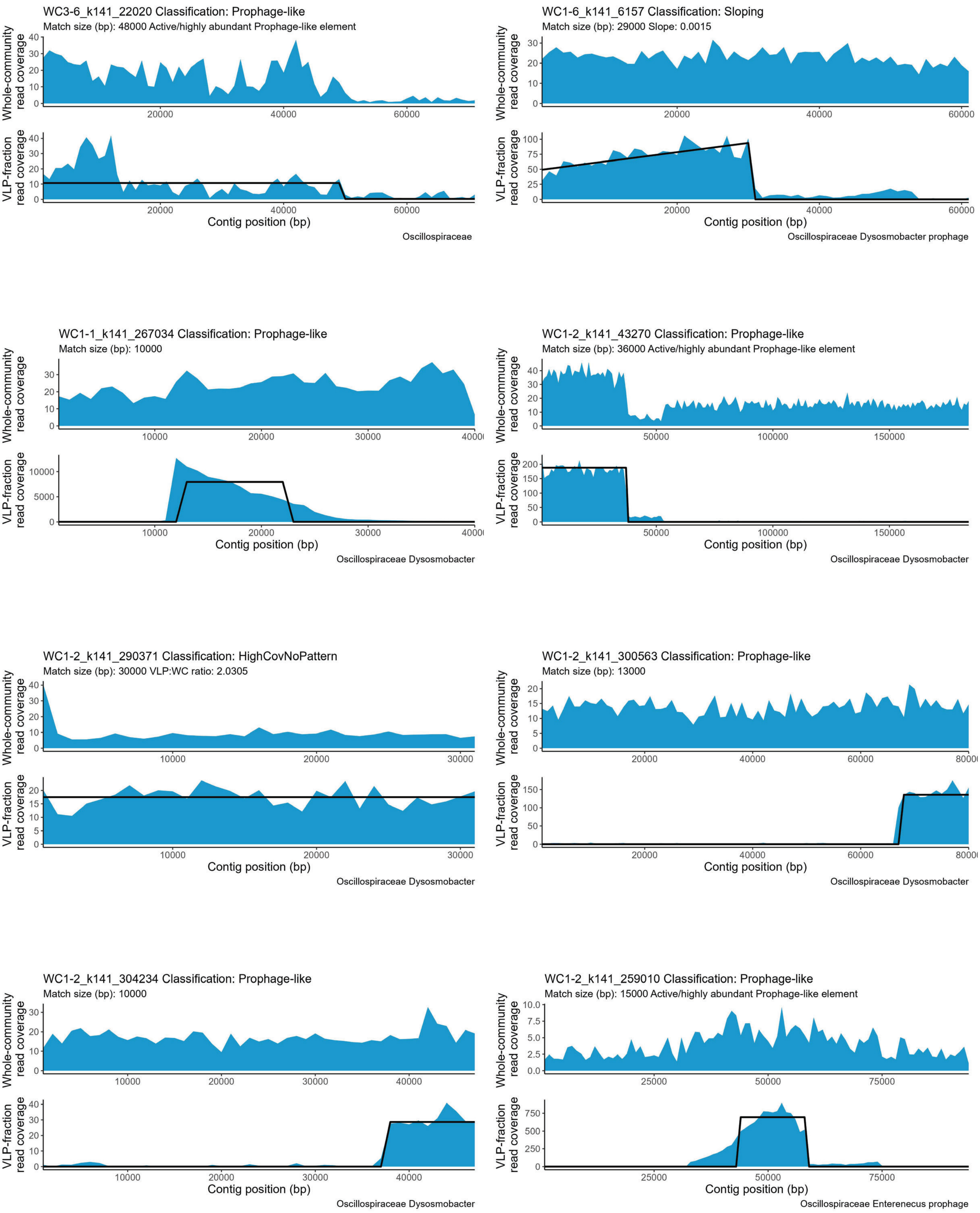
WC3-6_k141_22020 Classification: Prophage-like
Match size (bp): 48000 Active/highly abundant Prophage-like element
Whole-community read coverage
VLP-fraction read coverage
Contig position (bp)
Oscillospiraceae
WC1-6_k141_6157 Classification: Sloping
Match size (bp): 29000 Slope: 0.0015
Whole-community read coverage
VLP-fraction read coverage
Contig position (bp)
Oscillospiraceae Dysosmobacter prophage
WC1-1_k141_267034 Classification: Prophage-like
Match size (bp): 10000
Whole-community read coverage
VLP-fraction read coverage
Contig position (bp)
Oscillospiraceae Dysosmobacter
WC1-2_k141_43270 Classification: Prophage-like
Match size (bp): 36000 Active/highly abundant Prophage-like element
Whole-community read coverage
VLP-fraction read coverage
Contig position (bp)
Oscillospiraceae Dysosmobacter
WC1-2_k141_290371 Classification: HighCovNoPattern
Match size (bp): 30000 VLP:WC ratio: 2.0305
Whole-community read coverage
VLP-fraction read coverage
Contig position (bp)
Oscillospiraceae Dysosmobacter
WC1-2_k141_300563 Classification: Prophage-like
Match size (bp): 13000
Whole-community read coverage
VLP-fraction read coverage
Contig position (bp)
Oscillospiraceae Dysosmobacter
WC1-2_k141_304234 Classification: Prophage-like
Match size (bp): 10000
Whole-community read coverage
VLP-fraction read coverage
Contig position (bp)
Oscillospiraceae Dysosmobacter
WC1-2_k141_259010 Classification: Prophage-like
Match size (bp): 15000 Active/highly abundant Prophage-like element
Whole-community read coverage
VLP-fraction read coverage
Contig position (bp)
Oscillospiraceae Enterenecus prophage

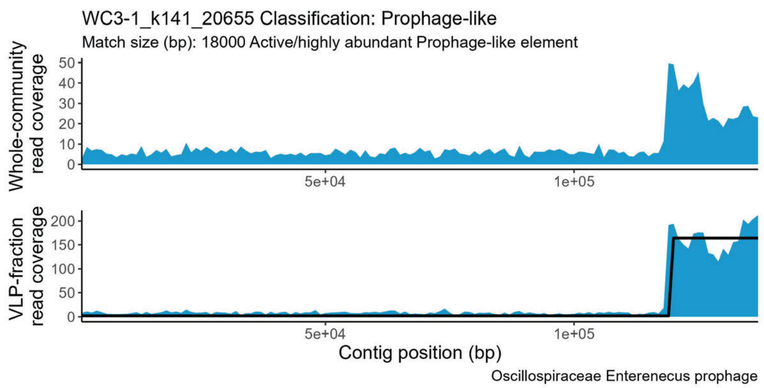
WC3-1_k141_20655 Classification: Prophage-like
Match size (bp): 18000 Active/highly abundant Prophage-like element
Whole-community read coverage
VLP-fraction read coverage
Contig position (bp)
Oscillospiraceae Enterenecus prophage

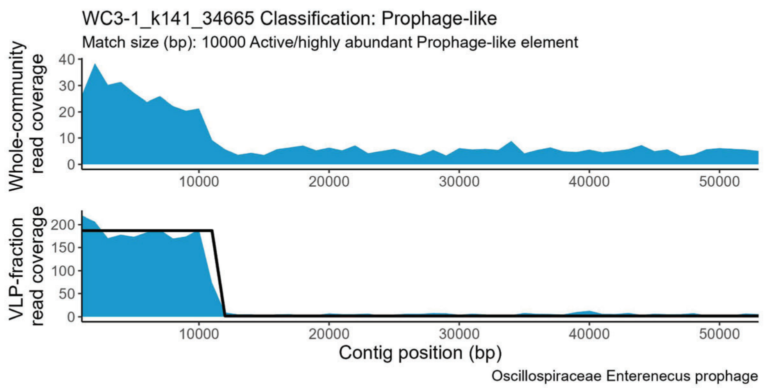
WC3-1_k141_34665 Classification: Prophage-like
Match size (bp): 10000 Active/highly abundant Prophage-like element
Whole-community read coverage
VLP-fraction read coverage
Contig position (bp)
Oscillospiraceae Enterenecus prophage

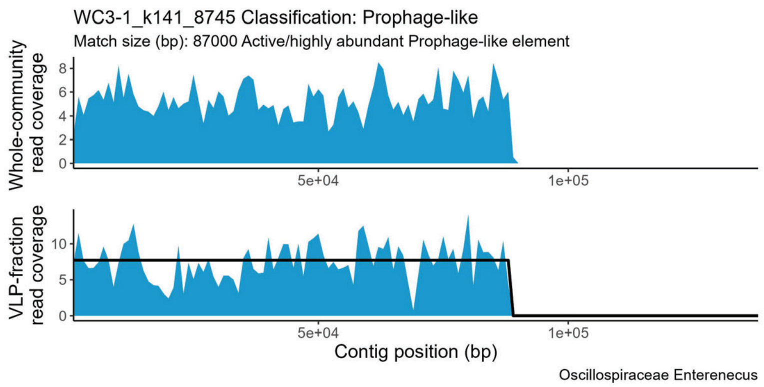
WC3-1_k141_8745 Classification: Prophage-like
Match size (bp): 87000 Active/highly abundant Prophage-like element
Whole-community read coverage
VLP-fraction read coverage
Contig position (bp)
Oscillospiraceae Enterenecus

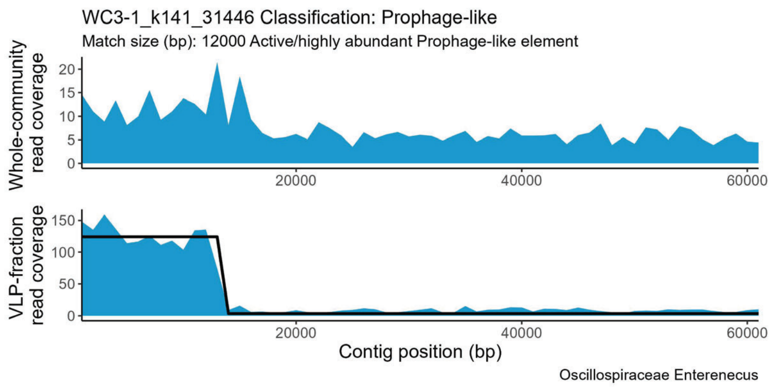
WC3-1_k141_31446 Classification: Prophage-like
Match size (bp): 12000 Active/highly abundant Prophage-like element
Whole-community read coverage
VLP-fraction read coverage
Contig position (bp)
Oscillospiraceae Enterenecus

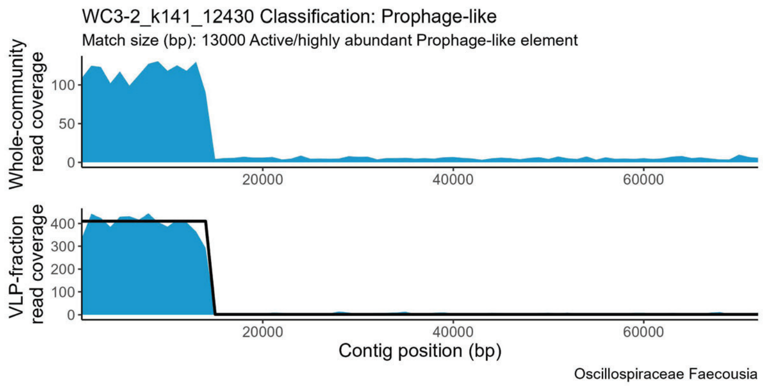
WC3-2_k141_12430 Classification: Prophage-like
Match size (bp): 13000 Active/highly abundant Prophage-like element
Whole-community read coverage
VLP-fraction read coverage
Contig position (bp)
Oscillospiraceae Faecousia

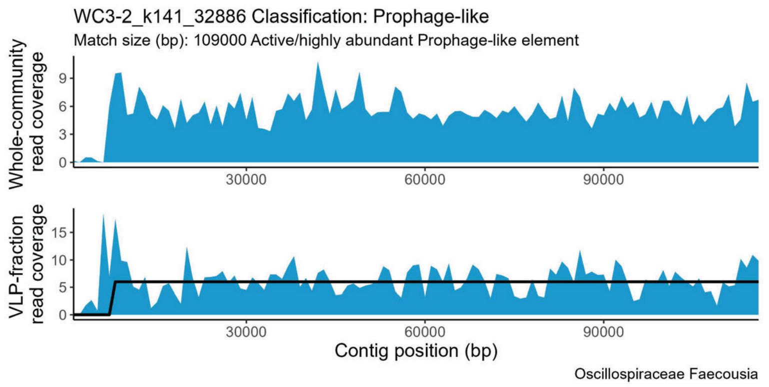
WC3-2_k141_32886 Classification: Prophage-like
Match size (bp): 109000 Active/highly abundant Prophage-like element
Whole-community read coverage
VLP-fraction read coverage
Contig position (bp)
Oscillospiraceae Faecousia

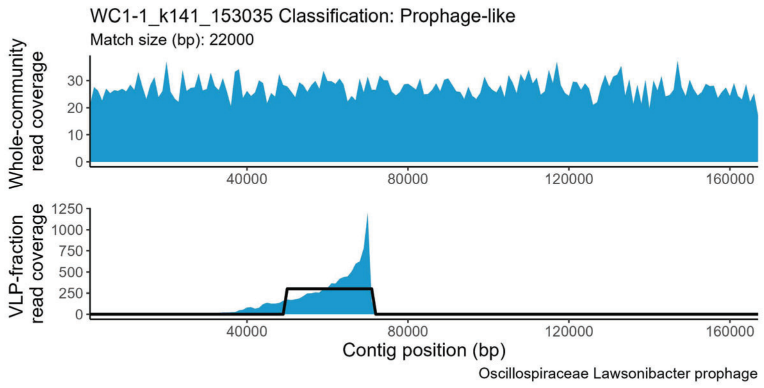
WC1-1_k141_153035 Classification: Prophage-like
Match size (bp): 22000
Whole-community read coverage
VLP-fraction read coverage
Contig position (bp)
Oscillospiraceae Lawsonibacter prophage

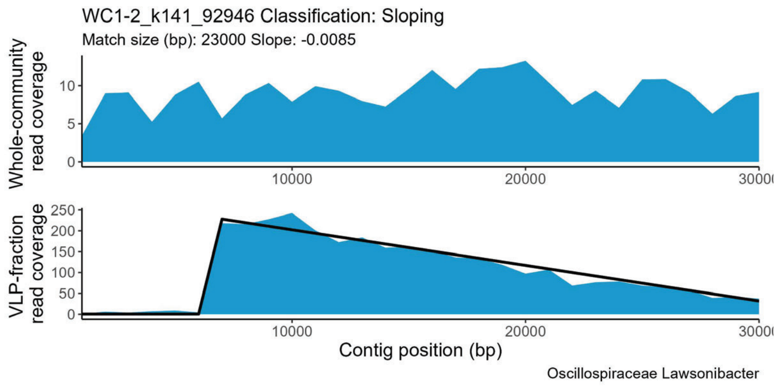
WC1-2_k141_92946 Classification: Sloping
Match size (bp): 23000 Slope: -0.0085
Whole-community read coverage
VLP-fraction read coverage
Contig position (bp)
Oscillospiraceae Lawsonibacter

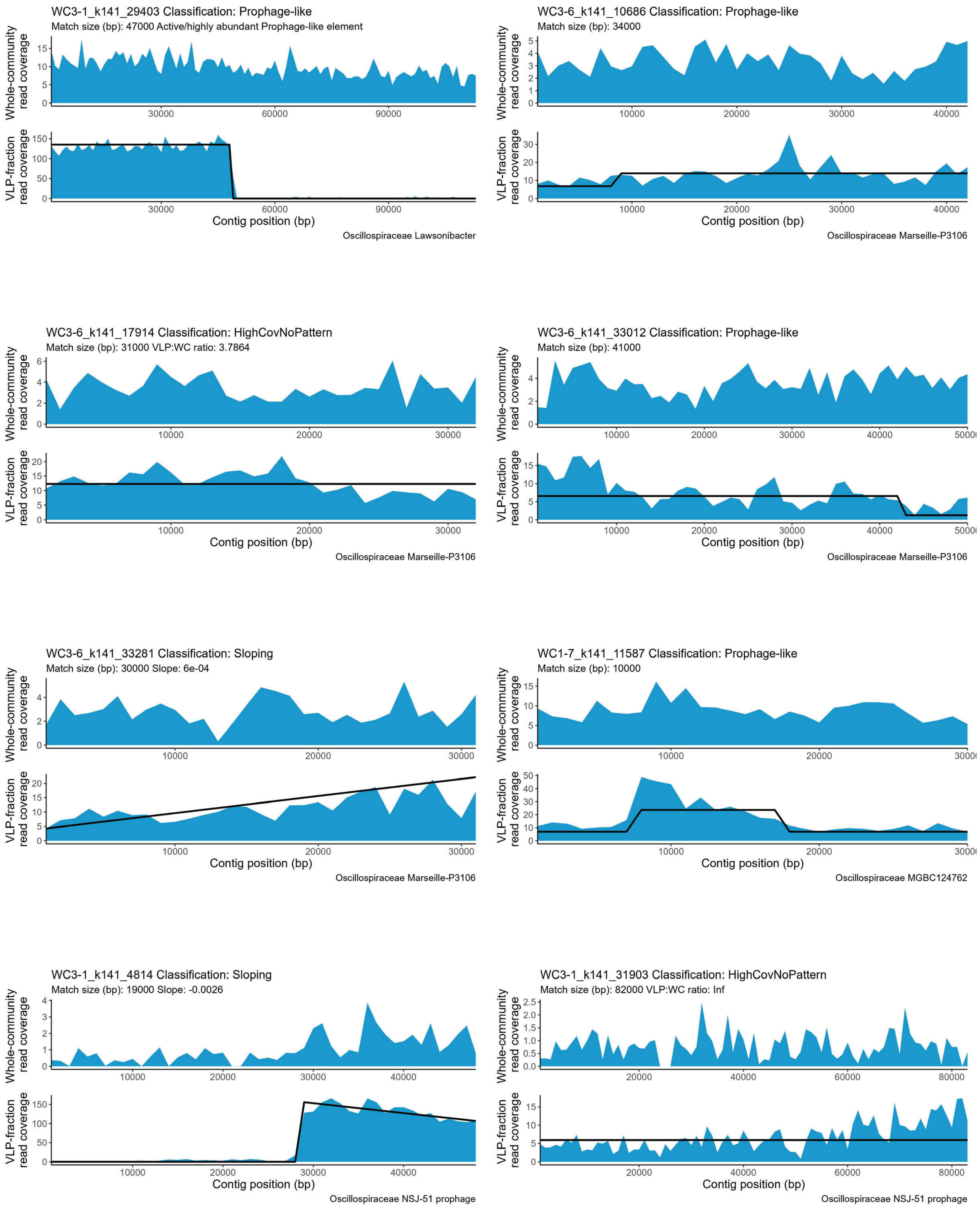

WC3-1_k141_29403 Classification: Prophage-like
Match size (bp): 47000 Active/highly abundant Prophage-like element
Oscillospiraceae Lawsonibacter
WC3-6_k141_10686 Classification: Prophage-like
Match size (bp): 34000
Oscillospiraceae Marseille-P3106
WC3-6_k141_17914 Classification: HighCovNoPattern
Match size (bp): 31000 VLP:WC ratio: 3.7864
Oscillospiraceae Marseille-P3106
WC3-6_k141_33012 Classification: Prophage-like
Match size (bp): 41000
Oscillospiraceae Marseille-P3106
WC3-6_k141_33281 Classification: Sloping
Match size (bp): 30000 Slope: 6e-04
Oscillospiraceae Marseille-P3106
WC1-7_k141_11587 Classification: Prophage-like
Match size (bp): 10000
Oscillospiraceae MGBC124762
WC3-1_k141_4814 Classification: Sloping
Match size (bp): 19000 Slope: -0.0026
Oscillospiraceae NSJ-51 prophage
WC3-1_k141_31903 Classification: HighCovNoPattern
Match size (bp): 82000 VLP:WC ratio: Inf
Oscillospiraceae NSJ-51 prophage
Whole-community read coverage
VLP-fraction read coverage
Contig position (bp)

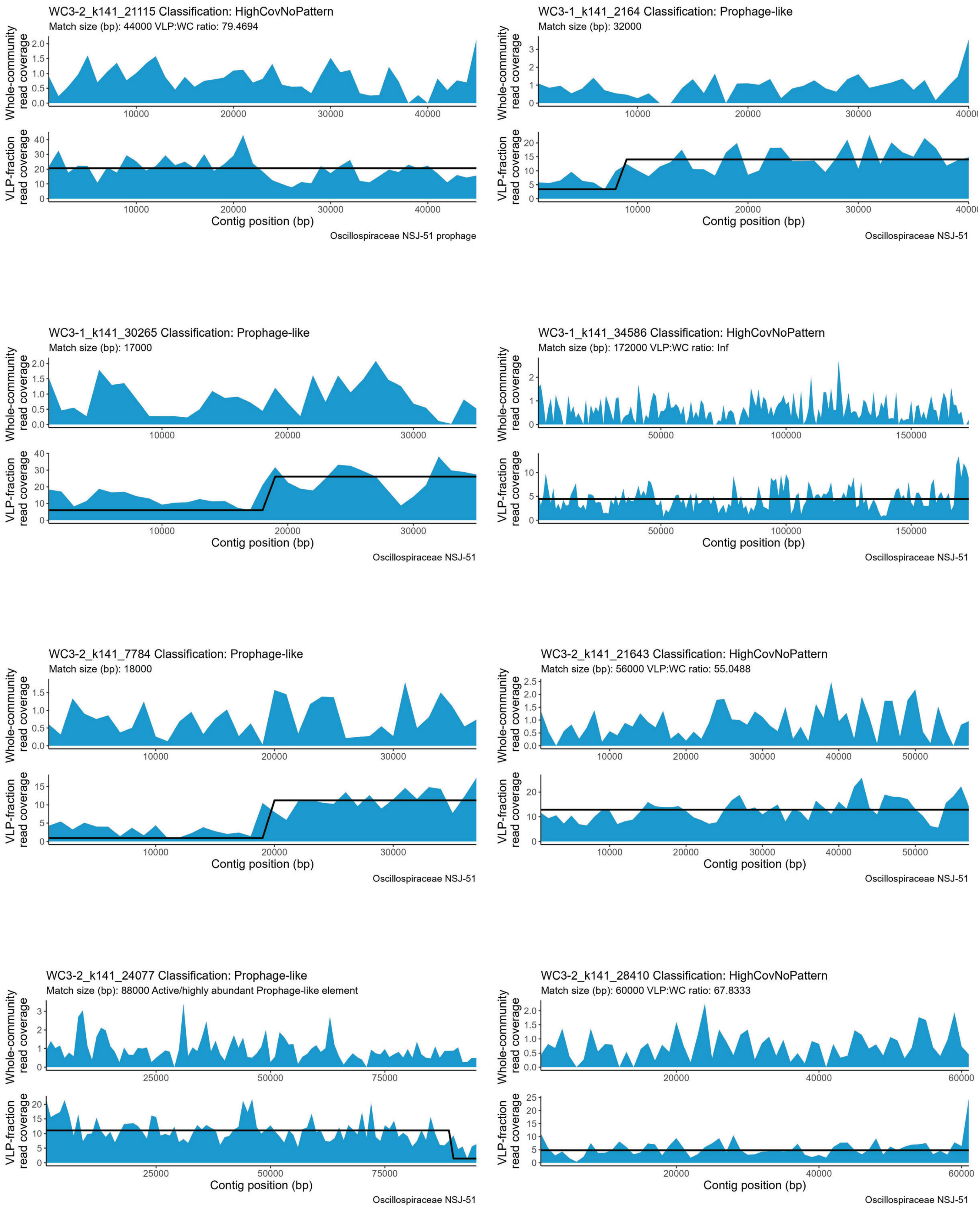
WC3-2_k141_21115 Classification: HighCovNoPattern
Match size (bp): 44000 VLP:WC ratio: 79.4694
Whole-community read coverage
VLP-fraction read coverage
Contig position (bp)
Oscillospiraceae NSJ-51 prophage
WC3-1_k141_2164 Classification: Prophage-like
Match size (bp): 32000
Whole-community read coverage
VLP-fraction read coverage
Contig position (bp)
Oscillospiraceae NSJ-51
WC3-1_k141_30265 Classification: Prophage-like
Match size (bp): 17000
Whole-community read coverage
VLP-fraction read coverage
Contig position (bp)
Oscillospiraceae NSJ-51
WC3-1_k141_34586 Classification: HighCovNoPattern
Match size (bp): 172000 VLP:WC ratio: Inf
Whole-community read coverage
VLP-fraction read coverage
Contig position (bp)
Oscillospiraceae NSJ-51
WC3-2_k141_7784 Classification: Prophage-like
Match size (bp): 18000
Whole-community read coverage
VLP-fraction read coverage
Contig position (bp)
Oscillospiraceae NSJ-51
WC3-2_k141_21643 Classification: HighCovNoPattern
Match size (bp): 56000 VLP:WC ratio: 55.0488
Whole-community read coverage
VLP-fraction read coverage
Contig position (bp)
Oscillospiraceae NSJ-51
WC3-2_k141_24077 Classification: Prophage-like
Match size (bp): 88000 Active/highly abundant Prophage-like element
Whole-community read coverage
VLP-fraction read coverage
Contig position (bp)
Oscillospiraceae NSJ-51
WC3-2_k141_28410 Classification: HighCovNoPattern
Match size (bp): 60000 VLP:WC ratio: 67.8333
Whole-community read coverage
VLP-fraction read coverage
Contig position (bp)
Oscillospiraceae NSJ-51

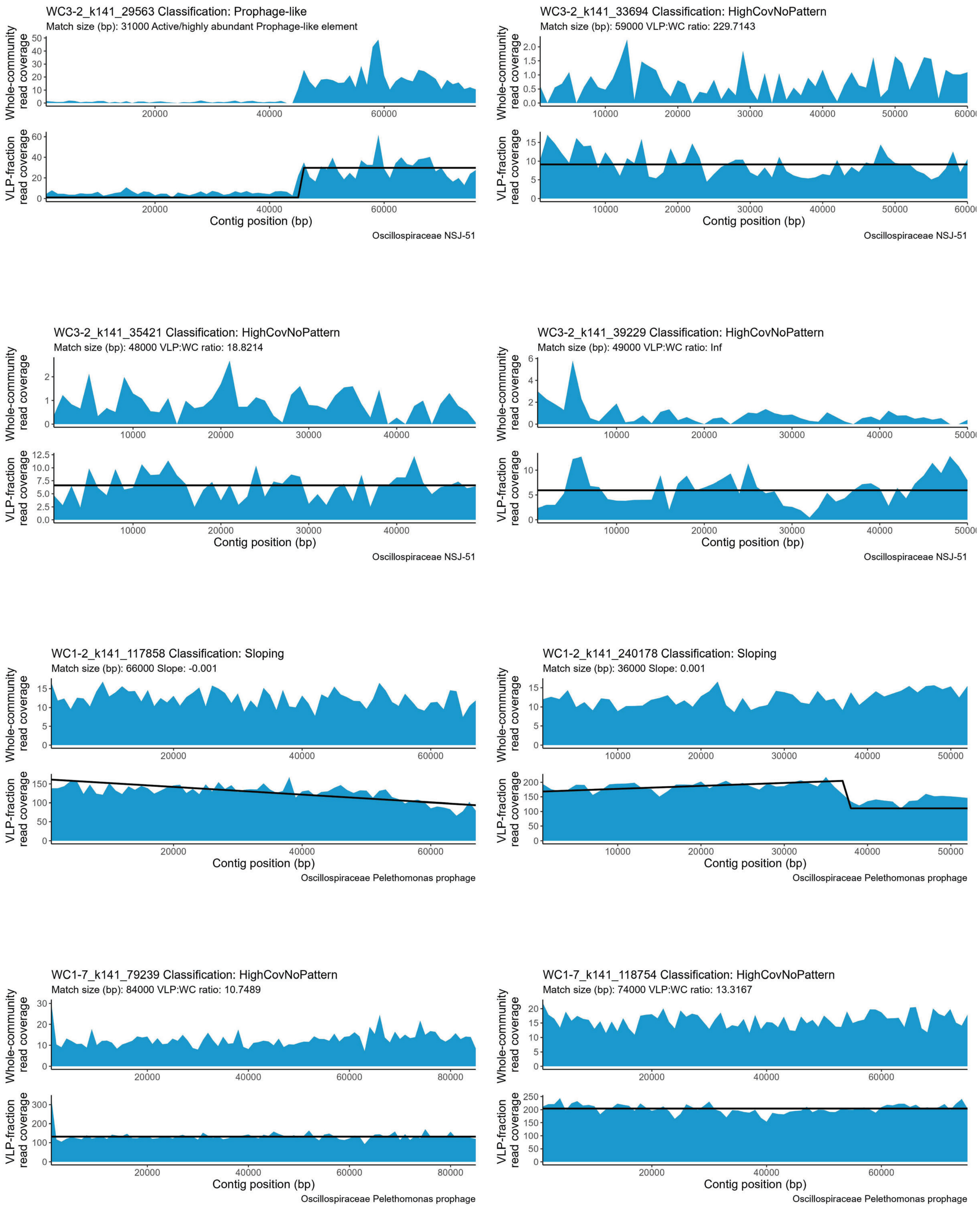

WC3-2_k141_29563 Classification: Prophage-like
Match size (bp): 31000 Active/highly abundant Prophage-like element
Whole-community read coverage
VLP-fraction read coverage
Contig position (bp)
Oscillospiraceae NSJ-51
WC3-2_k141_33694 Classification: HighCovNoPattern
Match size (bp): 59000 VLP:WC ratio: 229.7143
Whole-community read coverage
VLP-fraction read coverage
Contig position (bp)
Oscillospiraceae NSJ-51
WC3-2_k141_35421 Classification: HighCovNoPattern
Match size (bp): 48000 VLP:WC ratio: 18.8214
Whole-community read coverage
VLP-fraction read coverage
Contig position (bp)
Oscillospiraceae NSJ-51
WC3-2_k141_39229 Classification: HighCovNoPattern
Match size (bp): 49000 VLP:WC ratio: Inf
Whole-community read coverage
VLP-fraction read coverage
Contig position (bp)
Oscillospiraceae NSJ-51
WC1-2_k141_117858 Classification: Sloping
Match size (bp): 66000 Slope: -0.001
Whole-community read coverage
VLP-fraction read coverage
Contig position (bp)
Oscillospiraceae Pelethomonas prophage
WC1-2_k141_240178 Classification: Sloping
Match size (bp): 36000 Slope: 0.001
Whole-community read coverage
VLP-fraction read coverage
Contig position (bp)
Oscillospiraceae Pelethomonas prophage
WC1-7_k141_79239 Classification: HighCovNoPattern
Match size (bp): 84000 VLP:WC ratio: 10.7489
Whole-community read coverage
VLP-fraction read coverage
Contig position (bp)
Oscillospiraceae Pelethomonas prophage
WC1-7_k141_118754 Classification: HighCovNoPattern
Match size (bp): 74000 VLP:WC ratio: 13.3167
Whole-community read coverage
VLP-fraction read coverage
Contig position (bp)
Oscillospiraceae Pelethomonas prophage

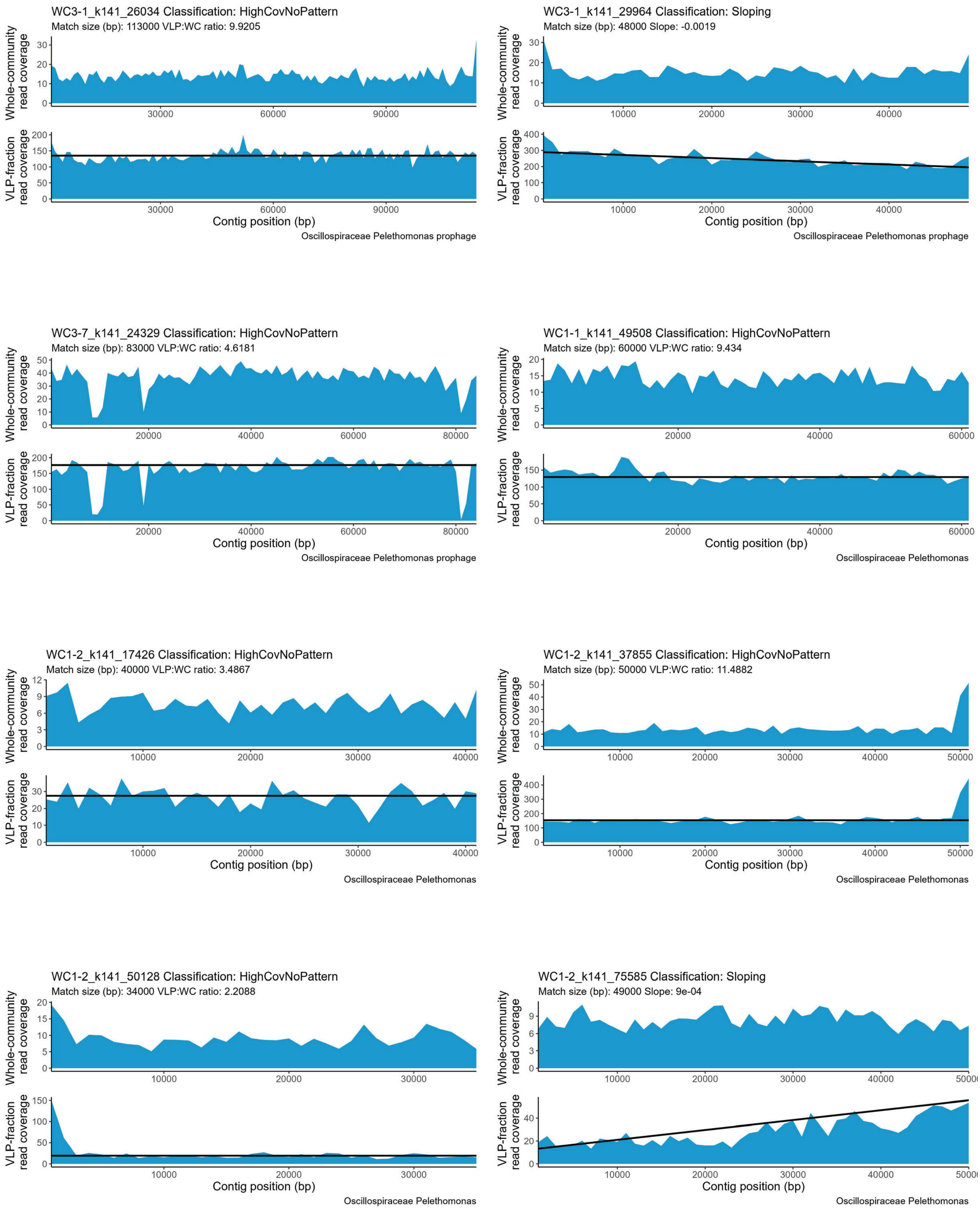
WC3-1_k141_26034 Classification: HighCovNoPattern
Match size (bp): 113000 VLP:WC ratio: 9.9205
Oscillospiraceae Pelethomonas prophage
WC3-1_k141_29964 Classification: Sloping
Match size (bp): 48000 Slope: -0.0019
Oscillospiraceae Pelethomonas prophage
WC3-7_k141_24329 Classification: HighCovNoPattern
Match size (bp): 83000 VLP:WC ratio: 4.6181
Oscillospiraceae Pelethomonas prophage
WC1-1_k141_49508 Classification: HighCovNoPattern
Match size (bp): 60000 VLP:WC ratio: 9.434
Oscillospiraceae Pelethomonas
WC1-2_k141_17426 Classification: HighCovNoPattern
Match size (bp): 40000 VLP:WC ratio: 3.4867
Oscillospiraceae Pelethomonas
WC1-2_k141_37855 Classification: HighCovNoPattern
Match size (bp): 50000 VLP:WC ratio: 11.4882
Oscillospiraceae Pelethomonas
WC1-2_k141_50128 Classification: HighCovNoPattern
Match size (bp): 34000 VLP:WC ratio: 2.2088
Oscillospiraceae Pelethomonas
WC1-2_k141_75585 Classification: Sloping
Match size (bp): 49000 Slope: 9e-04
Oscillospiraceae Pelethomonas
Whole-community read coverage
VLP-fraction read coverage
Contig position (bp)

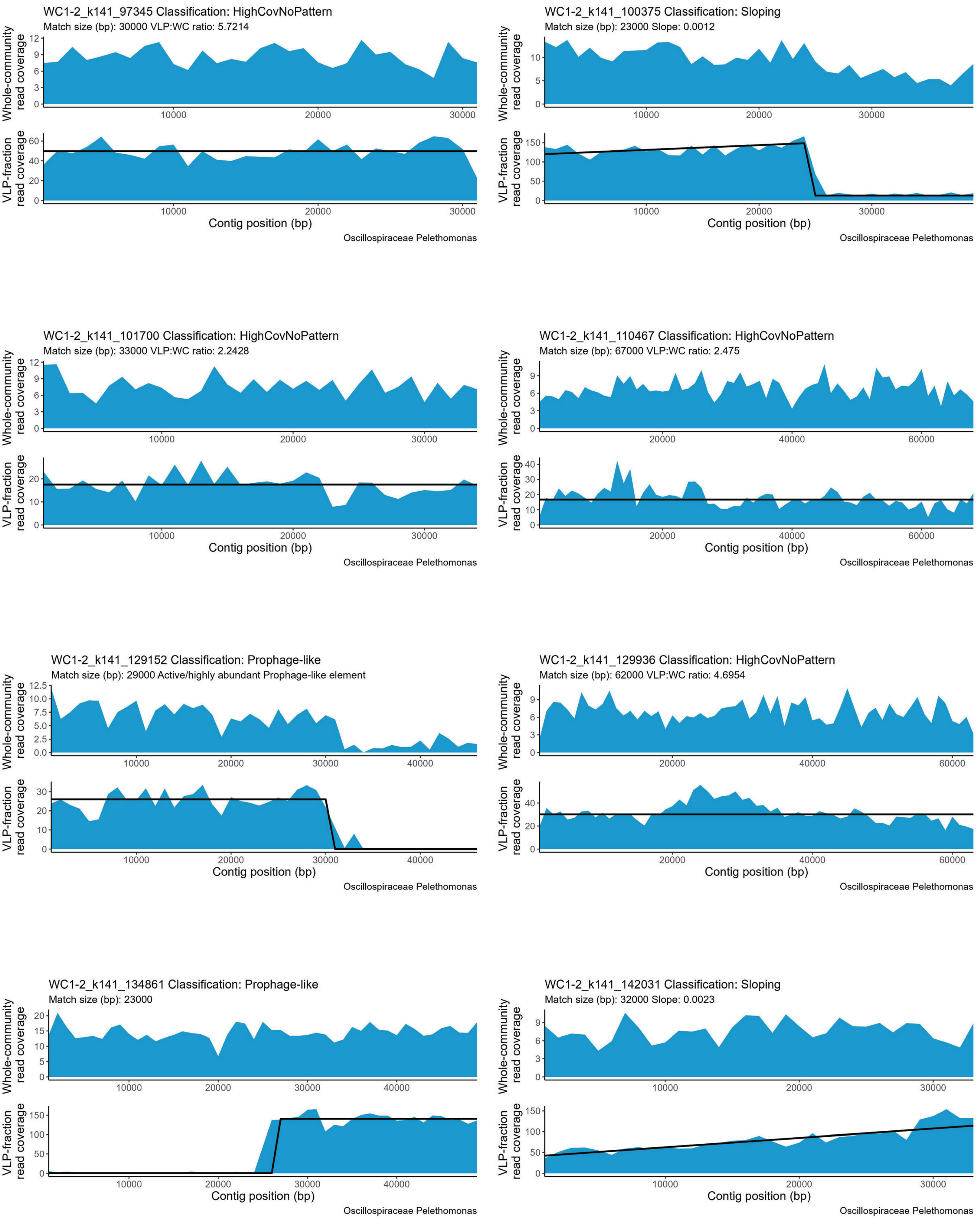
WC1-2_k141_97345 Classification: HighCovNoPattern
Match size (bp): 30000 VLP:WC ratio: 5.7214
Whole-community read coverage
VLP-fraction read coverage
Contig position (bp)
Oscillospiraceae Pelethomonas
WC1-2_k141_100375 Classification: Sloping
Match size (bp): 23000 Slope: 0.0012
Whole-community read coverage
VLP-fraction read coverage
Contig position (bp)
Oscillospiraceae Pelethomonas
WC1-2_k141_101700 Classification: HighCovNoPattern
Match size (bp): 33000 VLP:WC ratio: 2.2428
Whole-community read coverage
VLP-fraction read coverage
Contig position (bp)
Oscillospiraceae Pelethomonas
WC1-2_k141_110467 Classification: HighCovNoPattern
Match size (bp): 67000 VLP:WC ratio: 2.475
Whole-community read coverage
VLP-fraction read coverage
Contig position (bp)
Oscillospiraceae Pelethomonas
WC1-2_k141_129152 Classification: Prophage-like
Match size (bp): 29000 Active/highly abundant Prophage-like element
Whole-community read coverage
VLP-fraction read coverage
Contig position (bp)
Oscillospiraceae Pelethomonas
WC1-2_k141_129936 Classification: HighCovNoPattern
Match size (bp): 62000 VLP:WC ratio: 4.6954
Whole-community read coverage
VLP-fraction read coverage
Contig position (bp)
Oscillospiraceae Pelethomonas
WC1-2_k141_134861 Classification: Prophage-like
Match size (bp): 23000
Whole-community read coverage
VLP-fraction read coverage
Contig position (bp)
Oscillospiraceae Pelethomonas
WC1-2_k141_142031 Classification: Sloping
Match size (bp): 32000 Slope: 0.0023
Whole-community read coverage
VLP-fraction read coverage
Contig position (bp)
Oscillospiraceae Pelethomonas

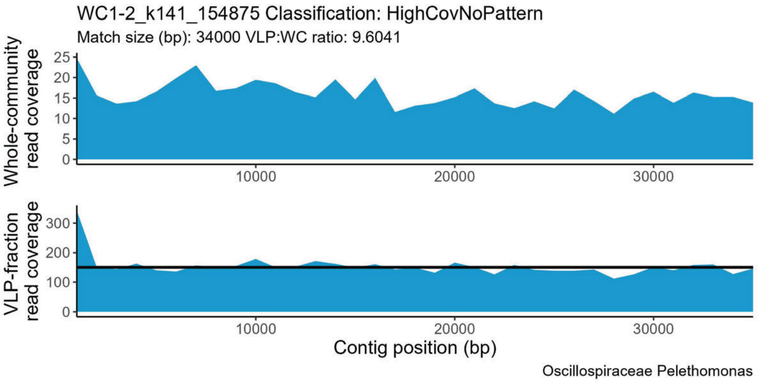
WC1-2_k141_154875 Classification: HighCovNoPattern
Match size (bp): 34000 VLP:WC ratio: 9.6041
Whole-community read coverage
VLP-fraction read coverage
Contig position (bp)
Oscillospiraceae Pelethomonas

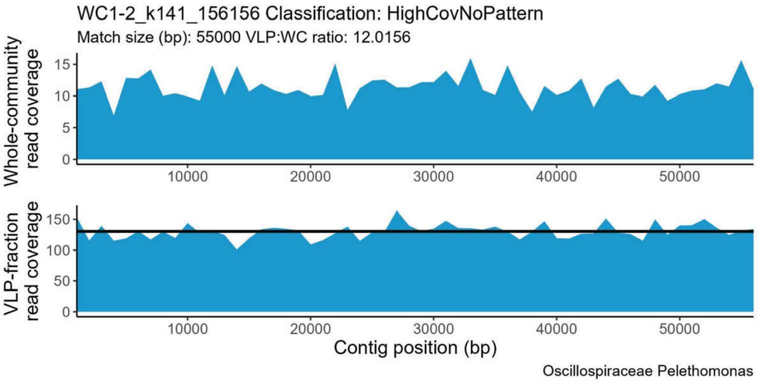
WC1-2_k141_156156 Classification: HighCovNoPattern
Match size (bp): 55000 VLP:WC ratio: 12.0156
Whole-community read coverage
VLP-fraction read coverage
Contig position (bp)
Oscillospiraceae Pelethomonas

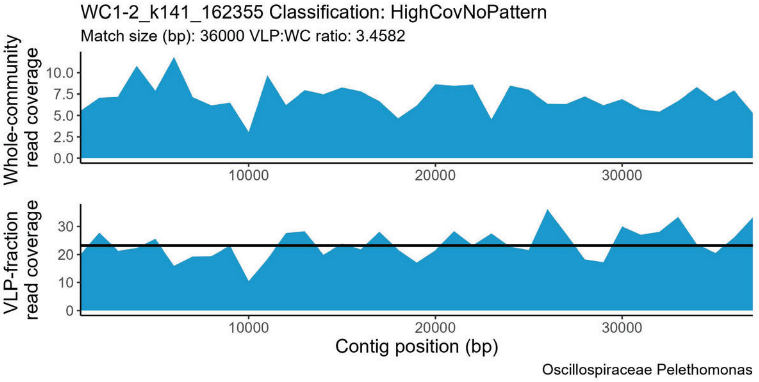
WC1-2_k141_162355 Classification: HighCovNoPattern
Match size (bp): 36000 VLP:WC ratio: 3.4582
Whole-community read coverage
VLP-fraction read coverage
Contig position (bp)
Oscillospiraceae Pelethomonas

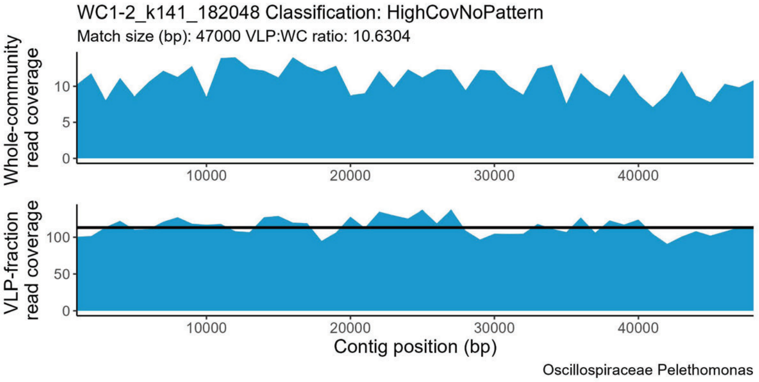
WC1-2_k141_182048 Classification: HighCovNoPattern
Match size (bp): 47000 VLP:WC ratio: 10.6304
Whole-community read coverage
VLP-fraction read coverage
Contig position (bp)
Oscillospiraceae Pelethomonas

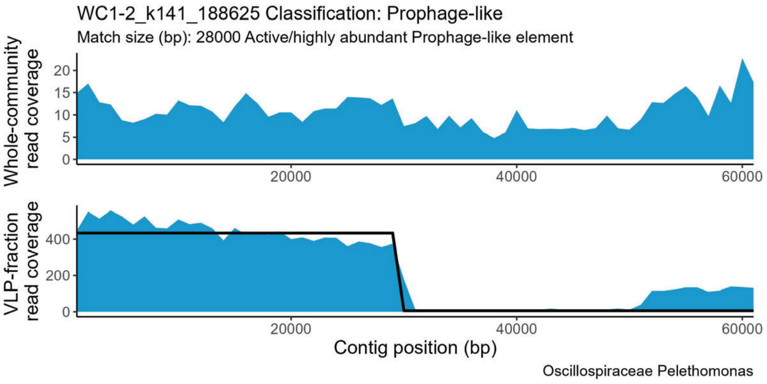
WC1-2_k141_188625 Classification: Prophage-like
Match size (bp): 28000 Active/highly abundant Prophage-like element
Whole-community read coverage
VLP-fraction read coverage
Contig position (bp)
Oscillospiraceae Pelethomonas

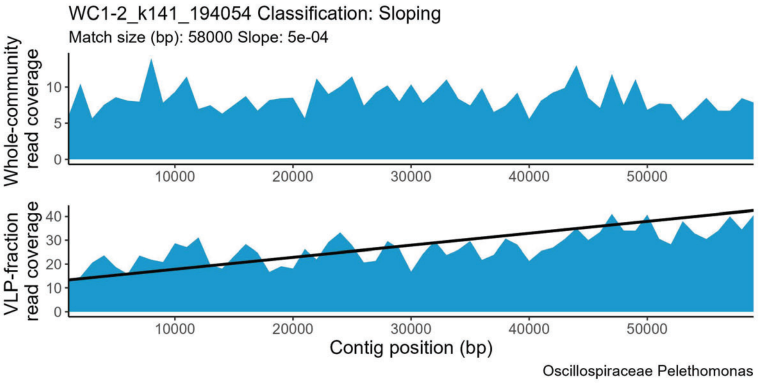
WC1-2_k141_194054 Classification: Sloping
Match size (bp): 58000 Slope: 5e-04
Whole-community read coverage
VLP-fraction read coverage
Contig position (bp)
Oscillospiraceae Pelethomonas

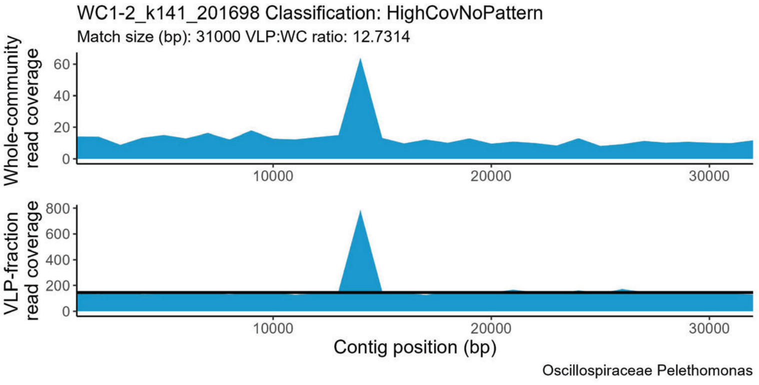
WC1-2_k141_201698 Classification: HighCovNoPattern
Match size (bp): 31000 VLP:WC ratio: 12.7314
Whole-community read coverage
VLP-fraction read coverage
Contig position (bp)
Oscillospiraceae Pelethomonas

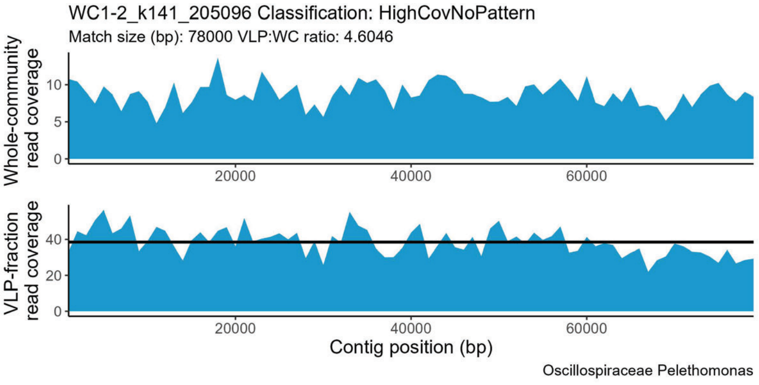
WC1-2_k141_205096 Classification: HighCovNoPattern
Match size (bp): 78000 VLP:WC ratio: 4.6046
Whole-community read coverage
VLP-fraction read coverage
Contig position (bp)
Oscillospiraceae Pelethomonas

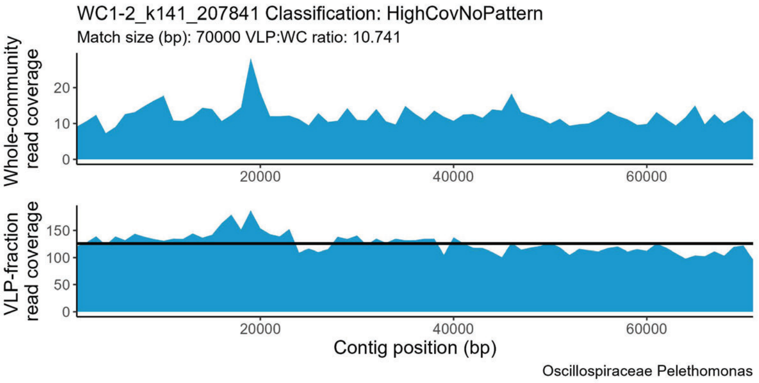
WC1-2_k141_207841 Classification: HighCovNoPattern
Match size (bp): 70000 VLP:WC ratio: 10.741
Whole-community read coverage
VLP-fraction read coverage
Contig position (bp)
Oscillospiraceae Pelethomonas

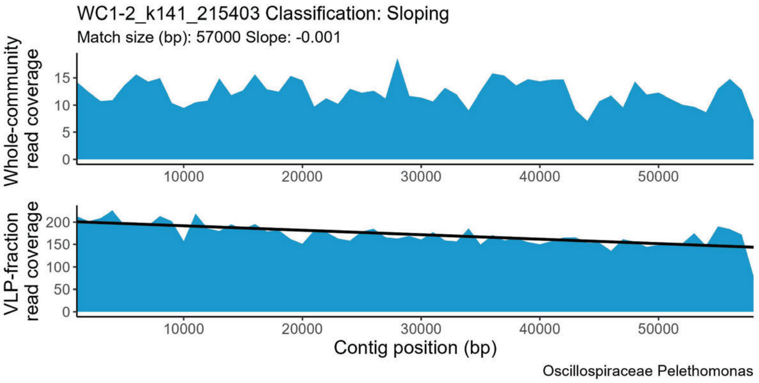
WC1-2_k141_215403 Classification: Sloping
Match size (bp): 57000 Slope: -0.001
Whole-community read coverage
VLP-fraction read coverage
Contig position (bp)
Oscillospiraceae Pelethomonas

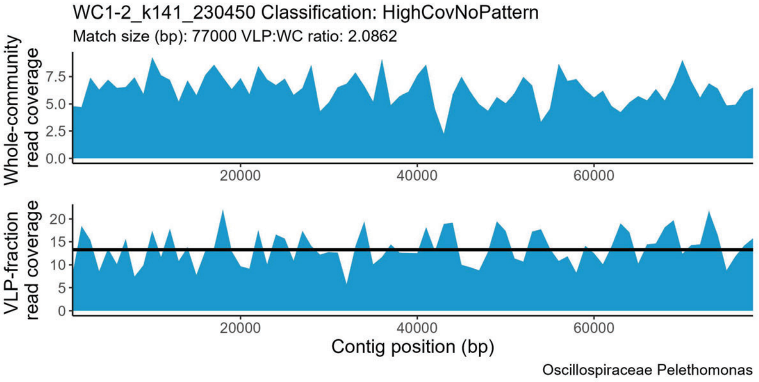
WC1-2_k141_230450 Classification: HighCovNoPattern
Match size (bp): 77000 VLP:WC ratio: 2.0862
Whole-community read coverage
VLP-fraction read coverage
Contig position (bp)
Oscillospiraceae Pelethomonas

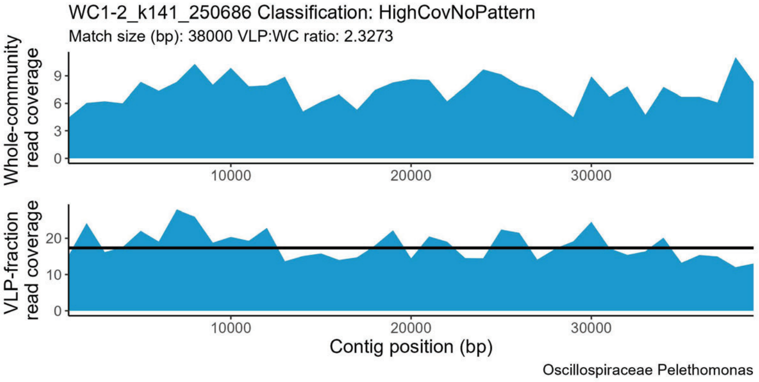
WC1-2_k141_250686 Classification: HighCovNoPattern
Match size (bp): 38000 VLP:WC ratio: 2.3273
Whole-community read coverage
VLP-fraction read coverage
Contig position (bp)
Oscillospiraceae Pelethomonas

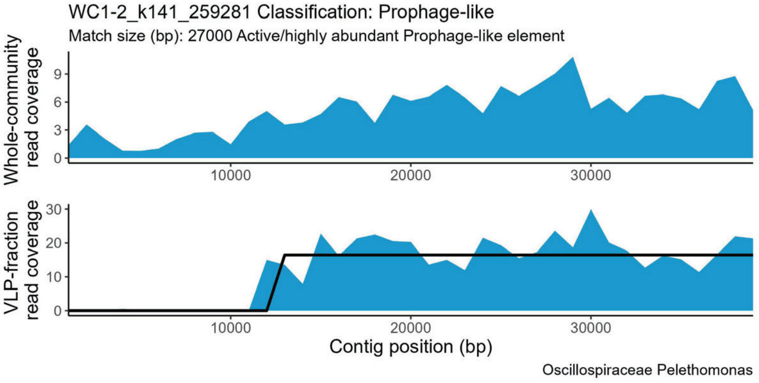
WC1-2_k141_259281 Classification: Prophage-like
Match size (bp): 27000 Active/highly abundant Prophage-like element
Whole-community read coverage
VLP-fraction read coverage
Contig position (bp)
Oscillospiraceae Pelethomonas

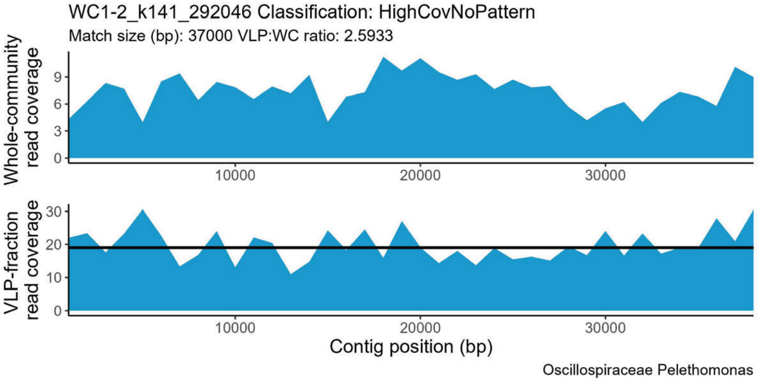
WC1-2_k141_292046 Classification: HighCovNoPattern
Match size (bp): 37000 VLP:WC ratio: 2.5933
Whole-community read coverage
VLP-fraction read coverage
Contig position (bp)
Oscillospiraceae Pelethomonas

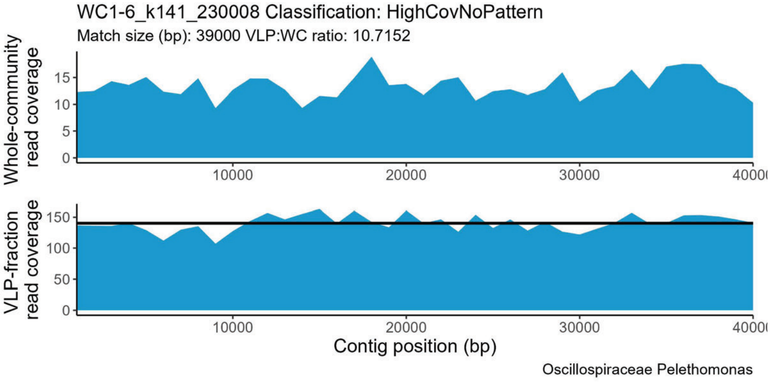
WC1-6_k141_230008 Classification: HighCovNoPattern
Match size (bp): 39000 VLP:WC ratio: 10.7152
Whole-community read coverage
VLP-fraction read coverage
Contig position (bp)
Oscillospiraceae Pelethomonas

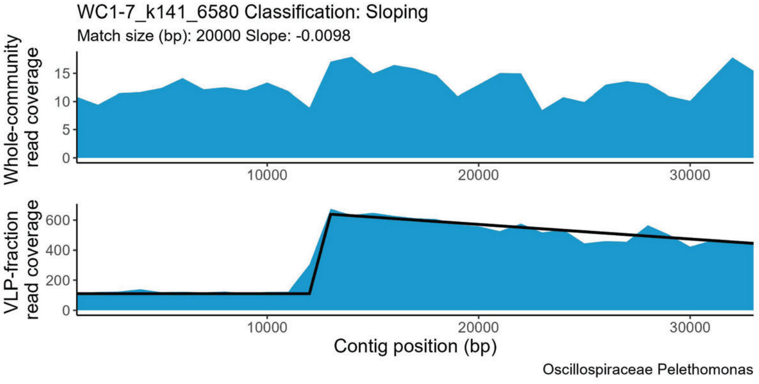
WC1-7_k141_6580 Classification: Sloping
Match size (bp): 20000 Slope: -0.0098
Whole-community read coverage
VLP-fraction read coverage
Contig position (bp)
Oscillospiraceae Pelethomonas

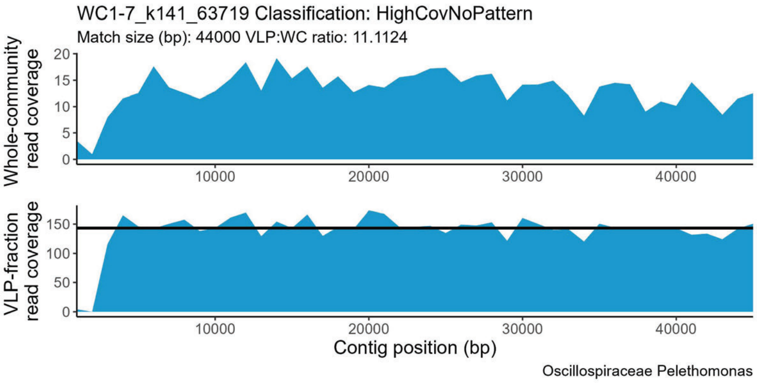
WC1-7_k141_63719 Classification: HighCovNoPattern
Match size (bp): 44000 VLP:WC ratio: 11.1124
Whole-community read coverage
VLP-fraction read coverage
Contig position (bp)
Oscillospiraceae Pelethomonas

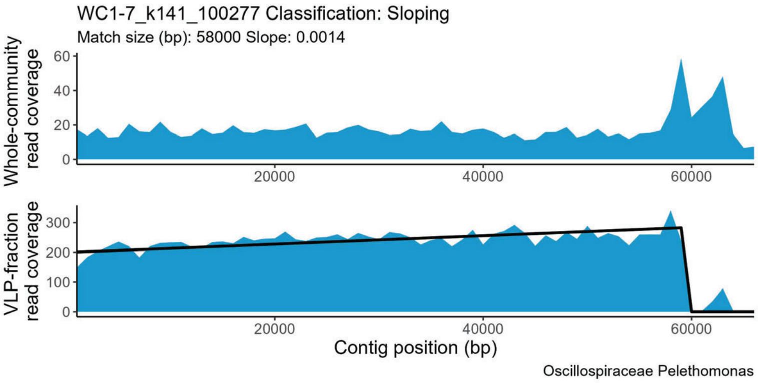
WC1-7_k141_100277 Classification: Sloping
Match size (bp): 58000 Slope: 0.0014
Whole-community read coverage
VLP-fraction read coverage
Contig position (bp)
Oscillospiraceae Pelethomonas

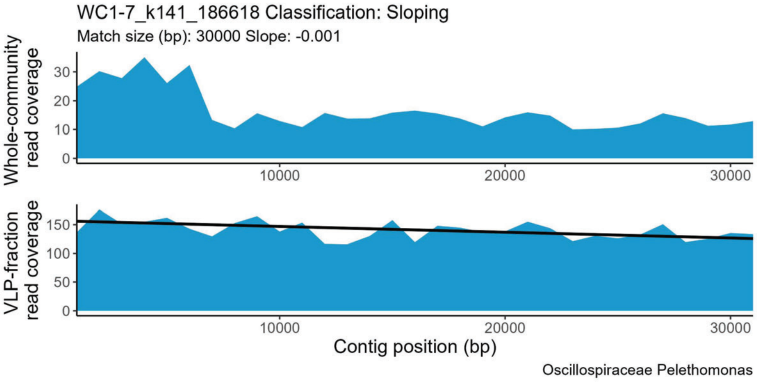
WC1-7_k141_186618 Classification: Sloping
Match size (bp): 30000 Slope: -0.001
Whole-community read coverage
VLP-fraction read coverage
Contig position (bp)
Oscillospiraceae Pelethomonas

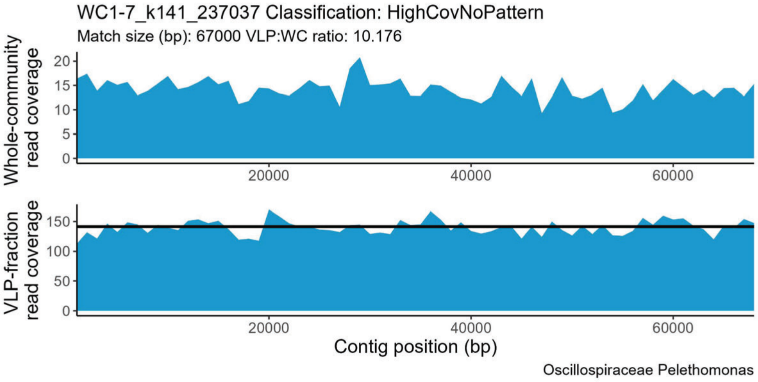
WC1-7_k141_237037 Classification: HighCovNoPattern
Match size (bp): 67000 VLP:WC ratio: 10.176
Whole-community read coverage
VLP-fraction read coverage
Contig position (bp)
Oscillospiraceae Pelethomonas

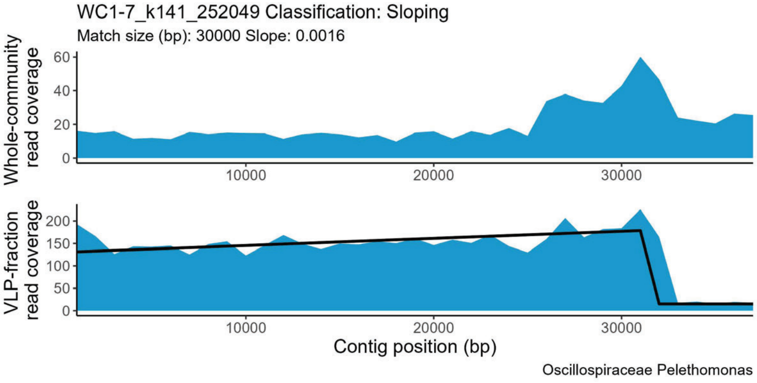
WC1-7_k141_252049 Classification: Sloping
Match size (bp): 30000 Slope: 0.0016
Whole-community read coverage
VLP-fraction read coverage
Contig position (bp)
Oscillospiraceae Pelethomonas

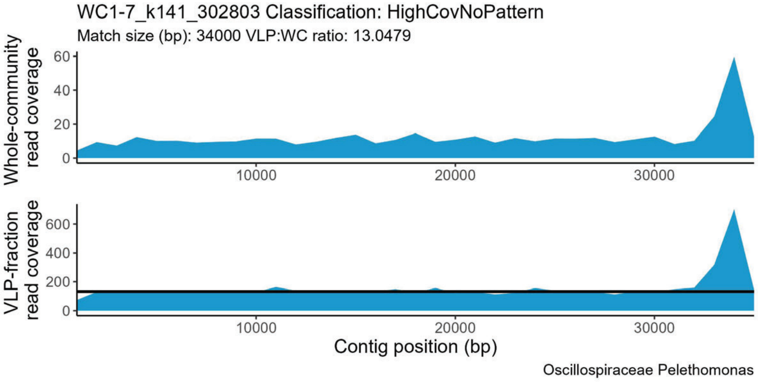
WC1-7_k141_302803 Classification: HighCovNoPattern
Match size (bp): 34000 VLP:WC ratio: 13.0479
Whole-community read coverage
VLP-fraction read coverage
Contig position (bp)
Oscillospiraceae Pelethomonas

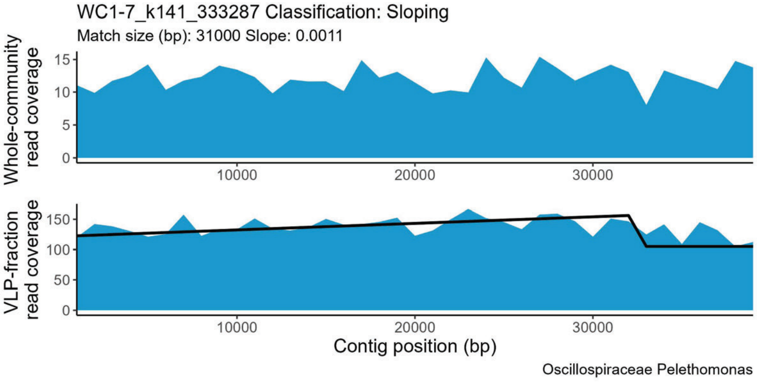
WC1-7_k141_333287 Classification: Sloping
Match size (bp): 31000 Slope: 0.0011
Whole-community read coverage
VLP-fraction read coverage
Contig position (bp)
Oscillospiraceae Pelethomonas

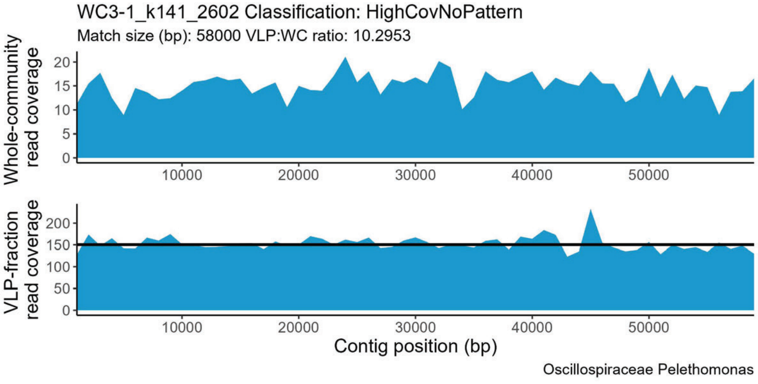
WC3-1_k141_2602 Classification: HighCovNoPattern
Match size (bp): 58000 VLP:WC ratio: 10.2953
Whole-community read coverage
VLP-fraction read coverage
Contig position (bp)
Oscillospiraceae Pelethomonas

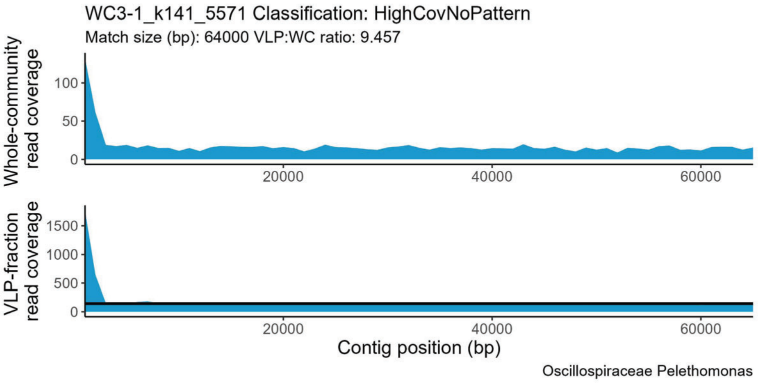
WC3-1_k141_5571 Classification: HighCovNoPattern
Match size (bp): 64000 VLP:WC ratio: 9.457
Whole-community read coverage
VLP-fraction read coverage
Contig position (bp)
Oscillospiraceae Pelethomonas

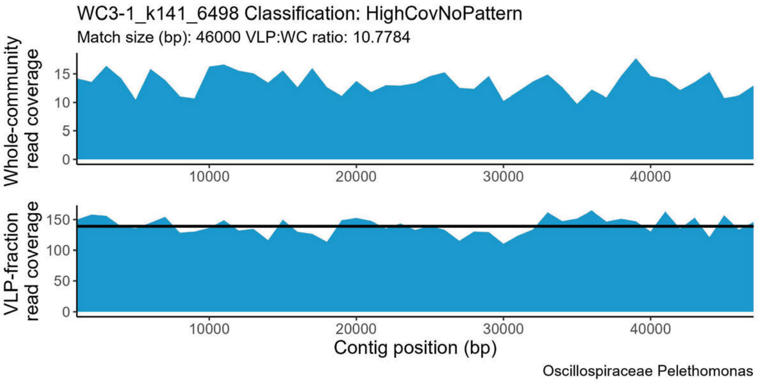
WC3-1_k141_6498 Classification: HighCovNoPattern
Match size (bp): 46000 VLP:WC ratio: 10.7784
Whole-community read coverage
VLP-fraction read coverage
Contig position (bp)
Oscillospiraceae Pelethomonas

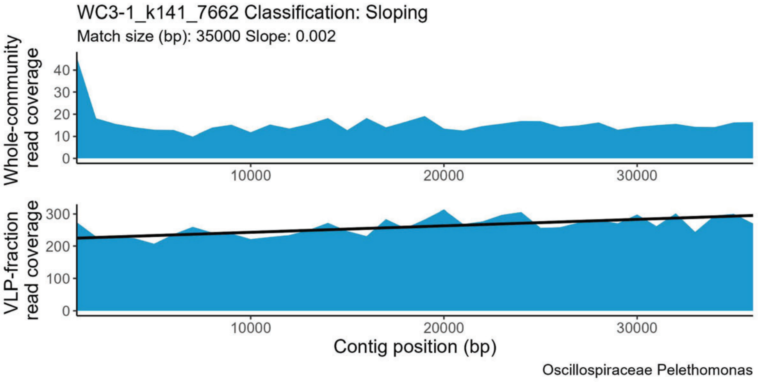
WC3-1_k141_7662 Classification: Sloping
Match size (bp): 35000 Slope: 0.002
Whole-community read coverage
VLP-fraction read coverage
Contig position (bp)
Oscillospiraceae Pelethomonas

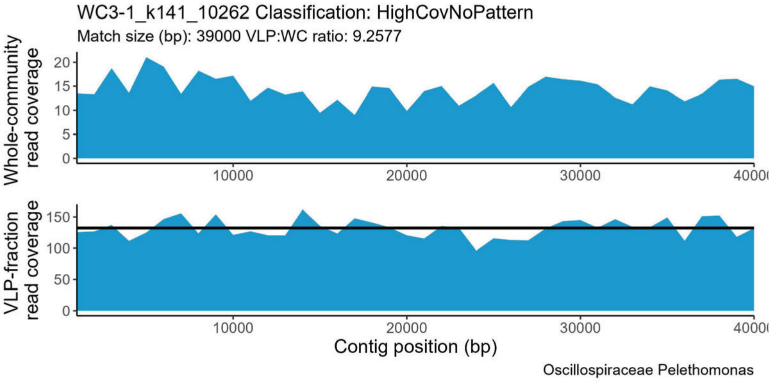
WC3-1_k141_10262 Classification: HighCovNoPattern
Match size (bp): 39000 VLP:WC ratio: 9.2577
Whole-community read coverage
VLP-fraction read coverage
Contig position (bp)
Oscillospiraceae Pelethomonas

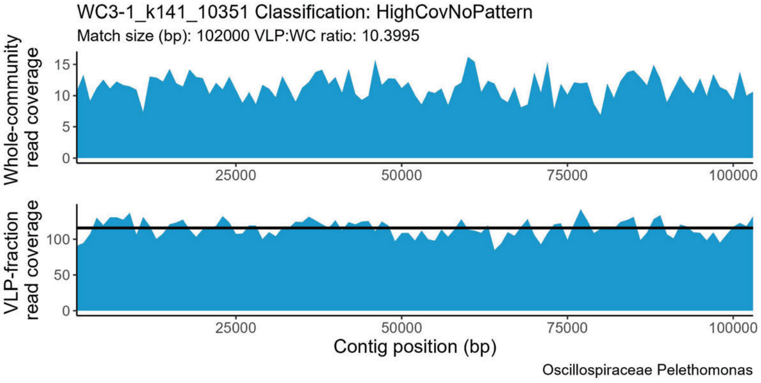
WC3-1_k141_10351 Classification: HighCovNoPattern
Match size (bp): 102000 VLP:WC ratio: 10.3995
Whole-community read coverage
VLP-fraction read coverage
Contig position (bp)
Oscillospiraceae Pelethomonas

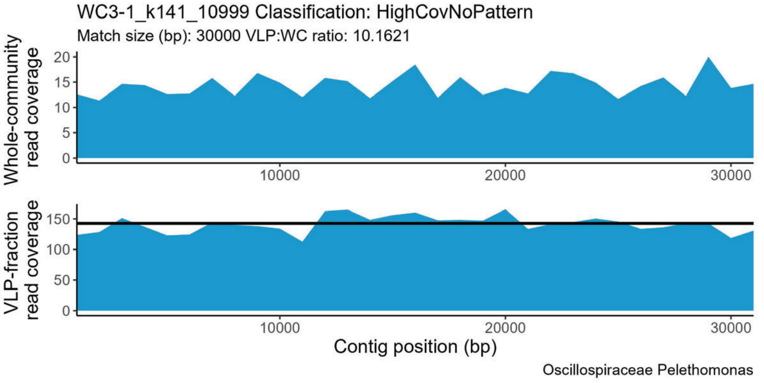
WC3-1_k141_10999 Classification: HighCovNoPattern
Match size (bp): 30000 VLP:WC ratio: 10.1621
Whole-community read coverage
VLP-fraction read coverage
Contig position (bp)
Oscillospiraceae Pelethomonas

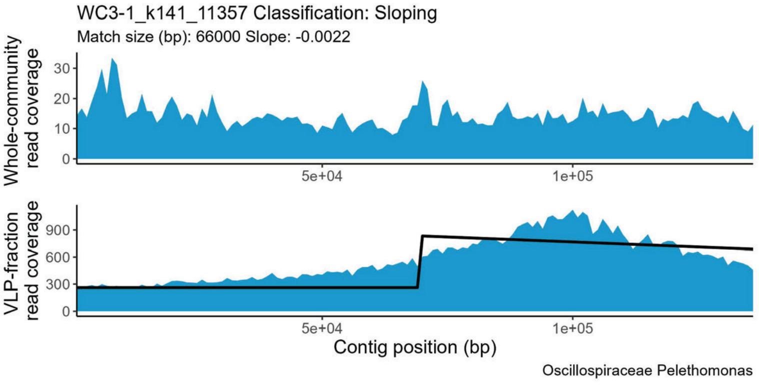
WC3-1_k141_11357 Classification: Sloping
Match size (bp): 66000 Slope: -0.0022
Whole-community read coverage
VLP-fraction read coverage
Contig position (bp)
Oscillospiraceae Pelethomonas

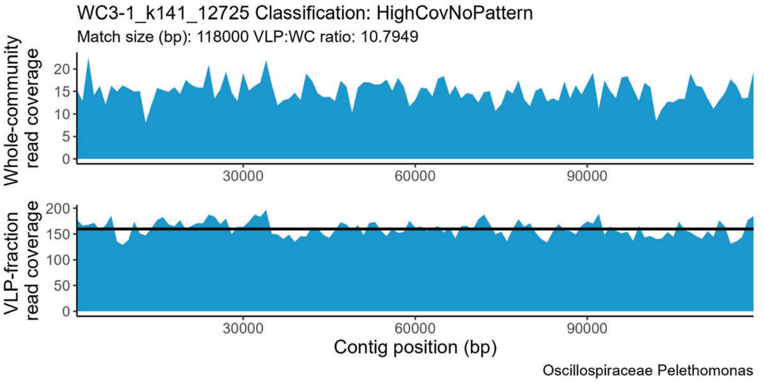
WC3-1_k141_12725 Classification: HighCovNoPattern
Match size (bp): 118000 VLP:WC ratio: 10.7949
Whole-community read coverage
VLP-fraction read coverage
Contig position (bp)
Oscillospiraceae Pelethomonas

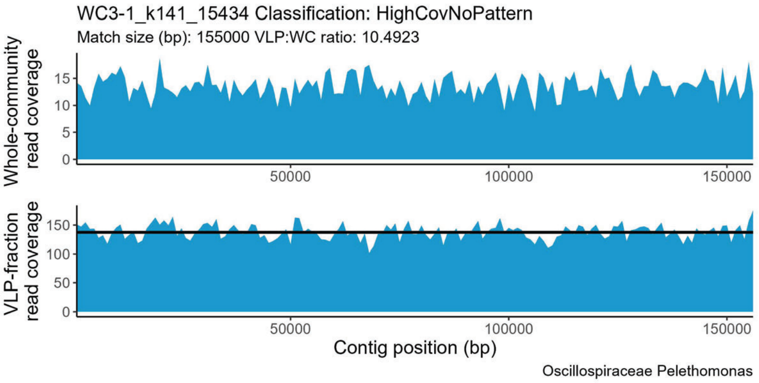
WC3-1_k141_15434 Classification: HighCovNoPattern
Match size (bp): 155000 VLP:WC ratio: 10.4923
Whole-community read coverage
VLP-fraction read coverage
Contig position (bp)
Oscillospiraceae Pelethomonas

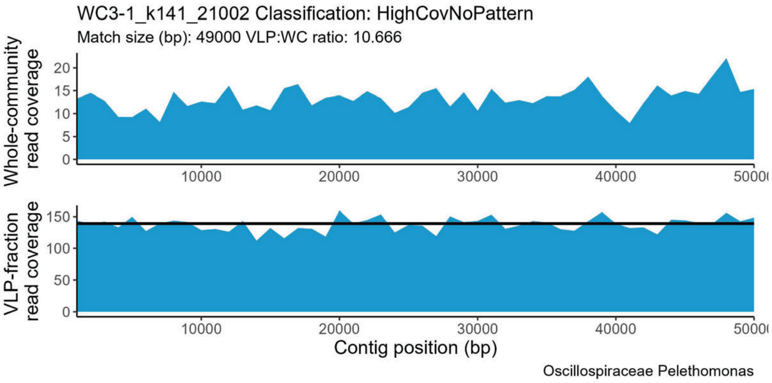
WC3-1_k141_21002 Classification: HighCovNoPattern
Match size (bp): 49000 VLP:WC ratio: 10.666
Whole-community read coverage
VLP-fraction read coverage
Contig position (bp)
Oscillospiraceae Pelethomonas

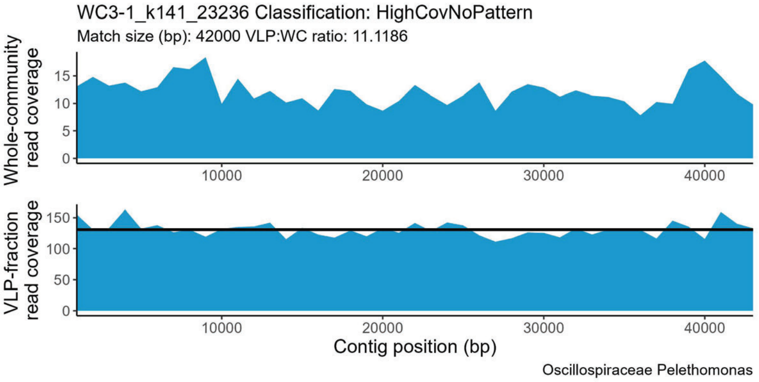
WC3-1_k141_23236 Classification: HighCovNoPattern
Match size (bp): 42000 VLP:WC ratio: 11.1186
Whole-community read coverage
VLP-fraction read coverage
Contig position (bp)
Oscillospiraceae Pelethomonas

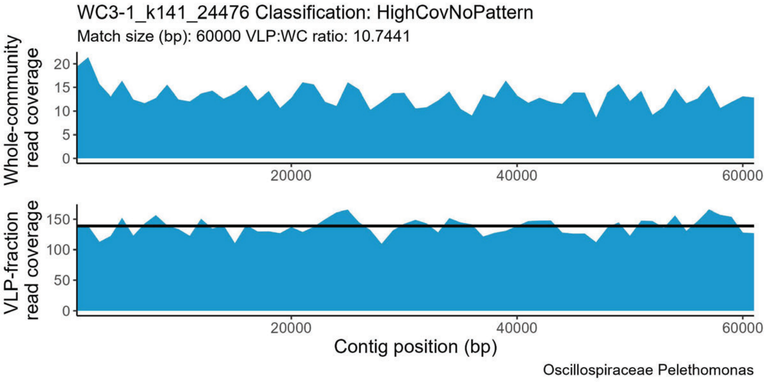
WC3-1_k141_24476 Classification: HighCovNoPattern
Match size (bp): 60000 VLP:WC ratio: 10.7441
Whole-community read coverage
VLP-fraction read coverage
Contig position (bp)
Oscillospiraceae Pelethomonas

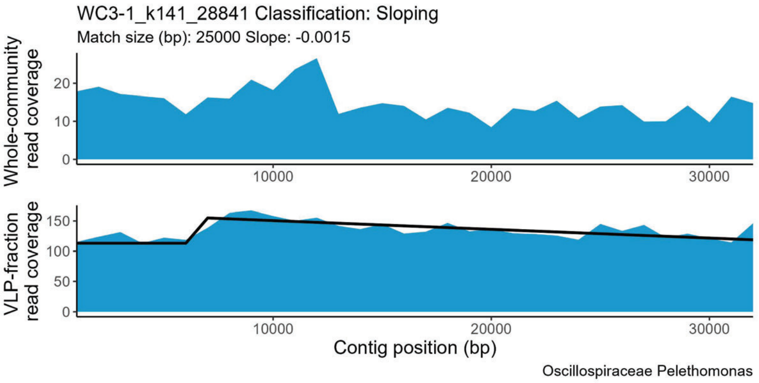
WC3-1_k141_28841 Classification: Sloping
Match size (bp): 25000 Slope: -0.0015
Whole-community read coverage
VLP-fraction read coverage
Contig position (bp)
Oscillospiraceae Pelethomonas

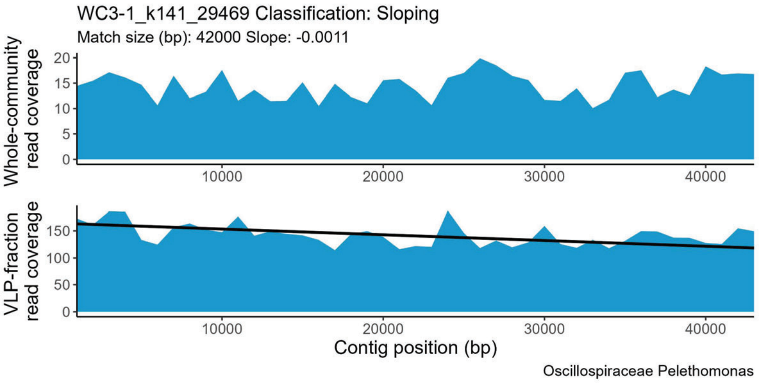
WC3-1_k141_29469 Classification: Sloping
Match size (bp): 42000 Slope: -0.0011
Whole-community read coverage
VLP-fraction read coverage
Contig position (bp)
Oscillospiraceae Pelethomonas

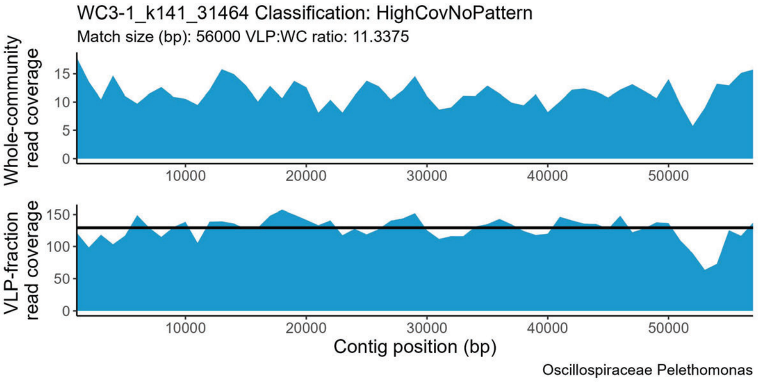
WC3-1_k141_31464 Classification: HighCovNoPattern
Match size (bp): 56000 VLP:WC ratio: 11.3375
Whole-community read coverage
VLP-fraction read coverage
Contig position (bp)
Oscillospiraceae Pelethomonas

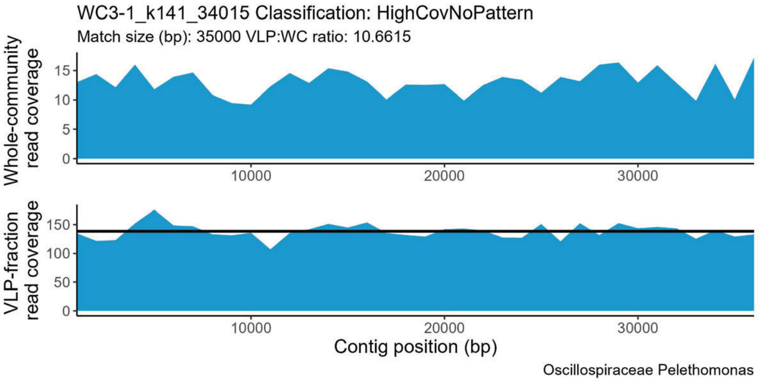
WC3-1_k141_34015 Classification: HighCovNoPattern
Match size (bp): 35000 VLP:WC ratio: 10.6615
Whole-community read coverage
VLP-fraction read coverage
Contig position (bp)
Oscillospiraceae Pelethomonas

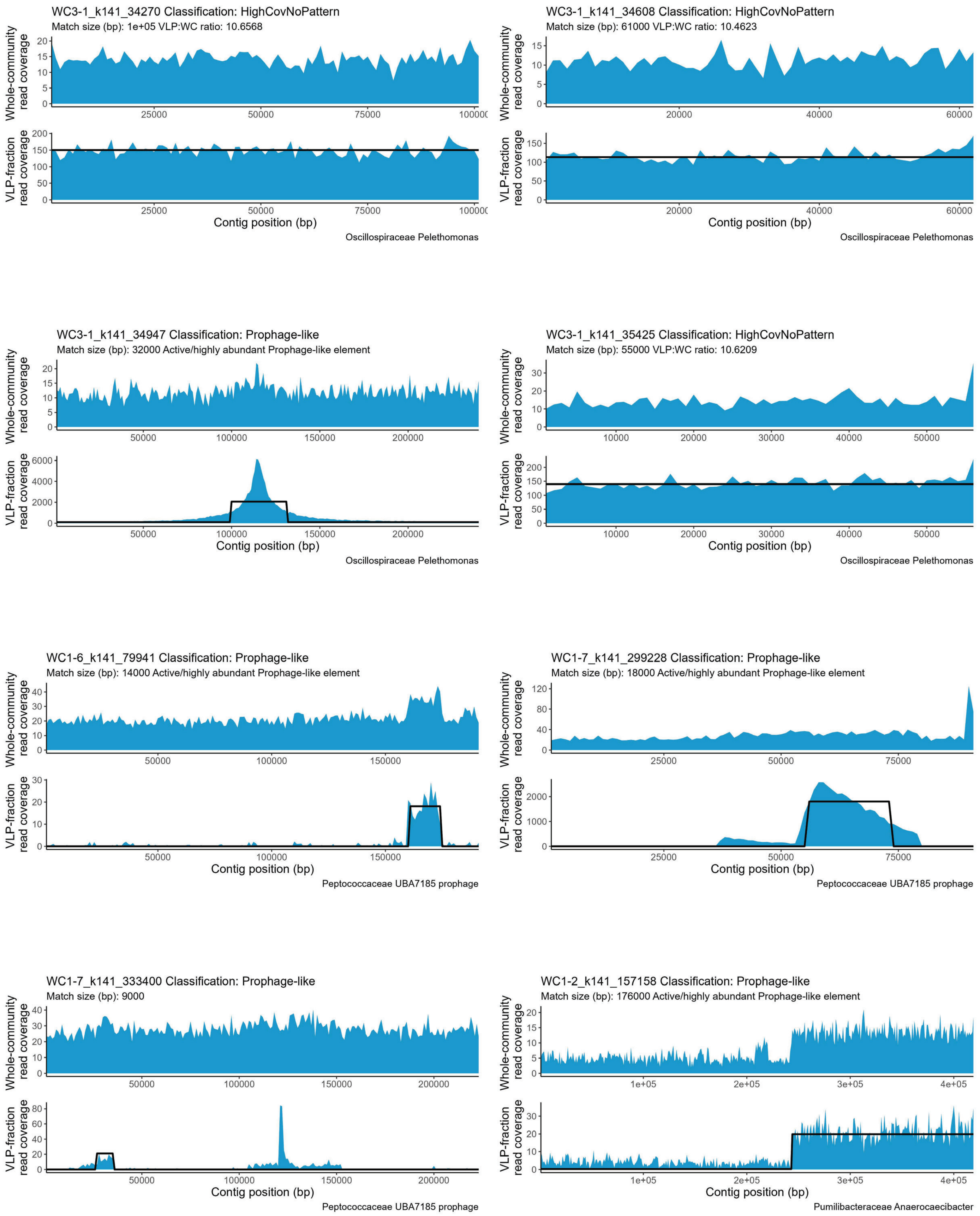

WC3-1_k141_34270 Classification: HighCovNoPattern
Match size (bp): 1e+05 VLP:WC ratio: 10.6568
Whole-community read coverage
VLP-fraction read coverage
Contig position (bp)
Oscillospiraceae Pelethomonas
WC3-1_k141_34608 Classification: HighCovNoPattern
Match size (bp): 61000 VLP:WC ratio: 10.4623
Whole-community read coverage
VLP-fraction read coverage
Contig position (bp)
Oscillospiraceae Pelethomonas
WC3-1_k141_34947 Classification: Prophage-like
Match size (bp): 32000 Active/highly abundant Prophage-like element
Whole-community read coverage
VLP-fraction read coverage
Contig position (bp)
Oscillospiraceae Pelethomonas
WC3-1_k141_35425 Classification: HighCovNoPattern
Match size (bp): 55000 VLP:WC ratio: 10.6209
Whole-community read coverage
VLP-fraction read coverage
Contig position (bp)
Oscillospiraceae Pelethomonas
WC1-6_k141_79941 Classification: Prophage-like
Match size (bp): 14000 Active/highly abundant Prophage-like element
Whole-community read coverage
VLP-fraction read coverage
Contig position (bp)
Peptococcaceae UBA7185 prophage
WC1-7_k141_299228 Classification: Prophage-like
Match size (bp): 18000 Active/highly abundant Prophage-like element
Whole-community read coverage
VLP-fraction read coverage
Contig position (bp)
Peptococcaceae UBA7185 prophage
WC1-7_k141_333400 Classification: Prophage-like
Match size (bp): 9000
Whole-community read coverage
VLP-fraction read coverage
Contig position (bp)
Peptococcaceae UBA7185 prophage
WC1-2_k141_157158 Classification: Prophage-like
Match size (bp): 176000 Active/highly abundant Prophage-like element
Whole-community read coverage
VLP-fraction read coverage
Contig position (bp)
Pumilibacteraceae Anaerocaecibacter

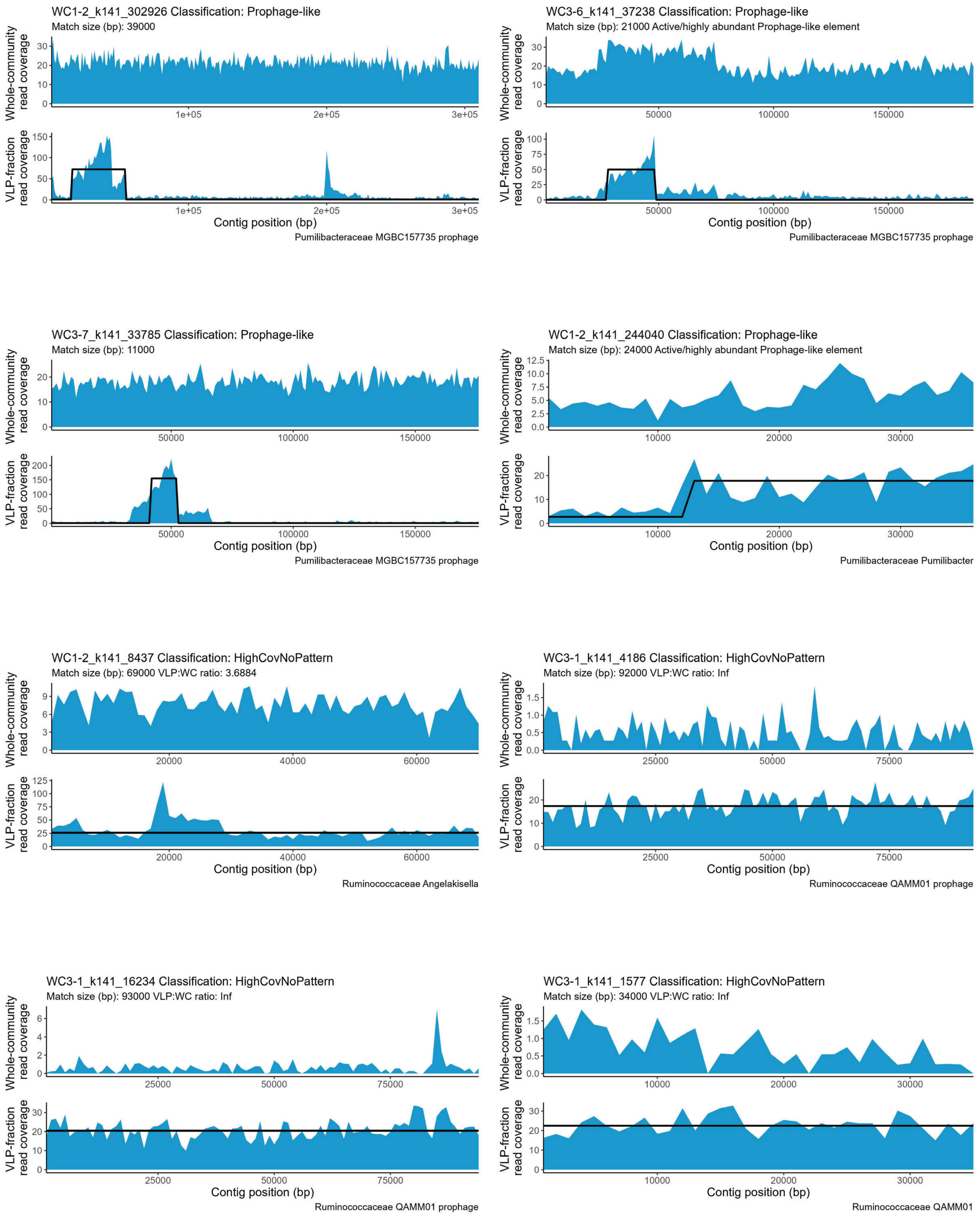

WC1-2_k141_302926 Classification: Prophage-like
Match size (bp): 39000
Whole-community read coverage
VLP-fraction read coverage
Contig position (bp)
Pumilibacteraceae MGBC157735 prophage
WC3-6_k141_37238 Classification: Prophage-like
Match size (bp): 21000 Active/highly abundant Prophage-like element
Whole-community read coverage
VLP-fraction read coverage
Contig position (bp)
Pumilibacteraceae MGBC157735 prophage
WC3-7_k141_33785 Classification: Prophage-like
Match size (bp): 11000
Whole-community read coverage
VLP-fraction read coverage
Contig position (bp)
Pumilibacteraceae MGBC157735 prophage
WC1-2_k141_244040 Classification: Prophage-like
Match size (bp): 24000 Active/highly abundant Prophage-like element
Whole-community read coverage
VLP-fraction read coverage
Contig position (bp)
Pumilibacteraceae Pumilibacter
WC1-2_k141_8437 Classification: HighCovNoPattern
Match size (bp): 69000 VLP:WC ratio: 3.6884
Whole-community read coverage
VLP-fraction read coverage
Contig position (bp)
Ruminococcaceae Angelakisella
WC3-1_k141_4186 Classification: HighCovNoPattern
Match size (bp): 92000 VLP:WC ratio: Inf
Whole-community read coverage
VLP-fraction read coverage
Contig position (bp)
Ruminococcaceae QAMM01 prophage
WC3-1_k141_16234 Classification: HighCovNoPattern
Match size (bp): 93000 VLP:WC ratio: Inf
Whole-community read coverage
VLP-fraction read coverage
Contig position (bp)
Ruminococcaceae QAMM01 prophage
WC3-1_k141_1577 Classification: HighCovNoPattern
Match size (bp): 34000 VLP:WC ratio: Inf
Whole-community read coverage
VLP-fraction read coverage
Contig position (bp)
Ruminococcaceae QAMM01

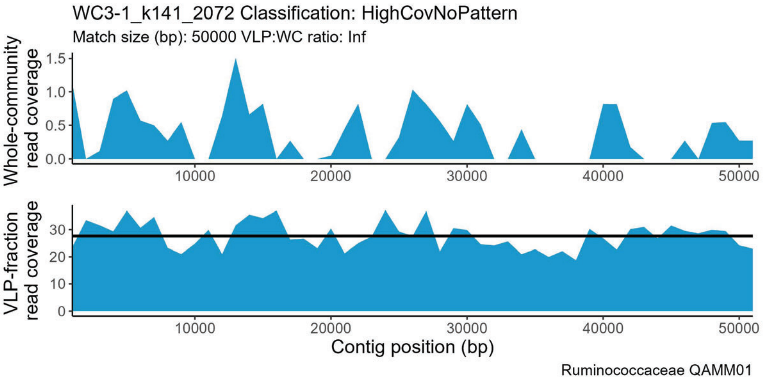
WC3-1_k141_2072 Classification: HighCovNoPattern
Match size (bp): 50000 VLP:WC ratio: Inf
Whole-community read coverage
VLP-fraction read coverage
Contig position (bp)
Ruminococcaceae QAMM01

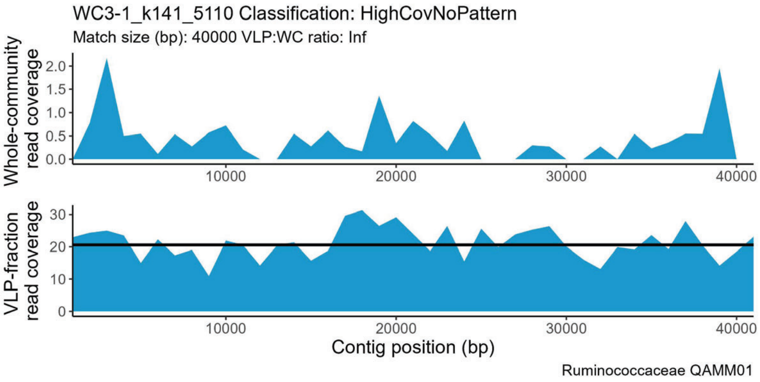
WC3-1_k141_5110 Classification: HighCovNoPattern
Match size (bp): 40000 VLP:WC ratio: Inf
Whole-community read coverage
VLP-fraction read coverage
Contig position (bp)
Ruminococcaceae QAMM01

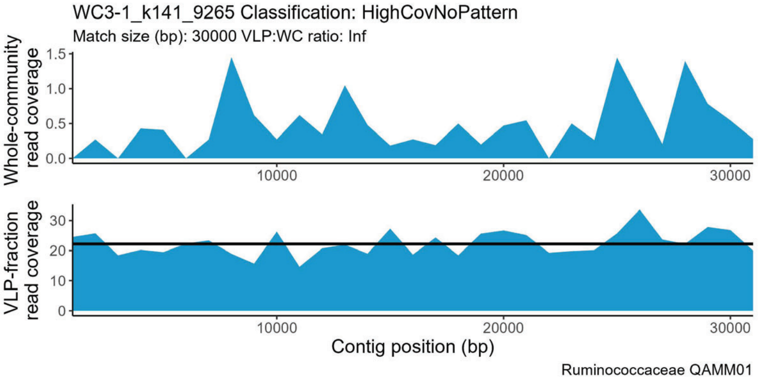
WC3-1_k141_9265 Classification: HighCovNoPattern
Match size (bp): 30000 VLP:WC ratio: Inf
Whole-community read coverage
VLP-fraction read coverage
Contig position (bp)
Ruminococcaceae QAMM01

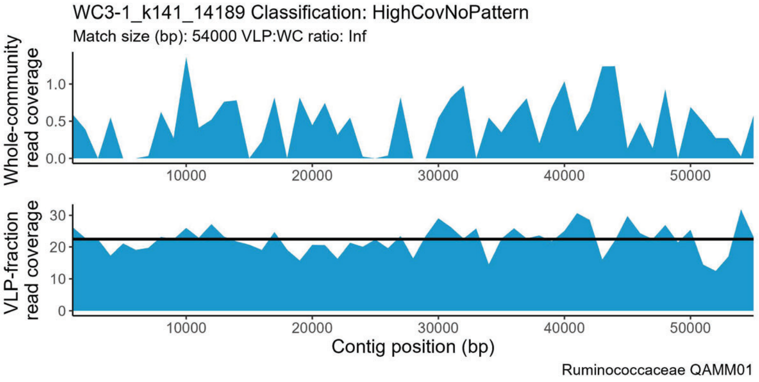
WC3-1_k141_14189 Classification: HighCovNoPattern
Match size (bp): 54000 VLP:WC ratio: Inf
Whole-community read coverage
VLP-fraction read coverage
Contig position (bp)
Ruminococcaceae QAMM01

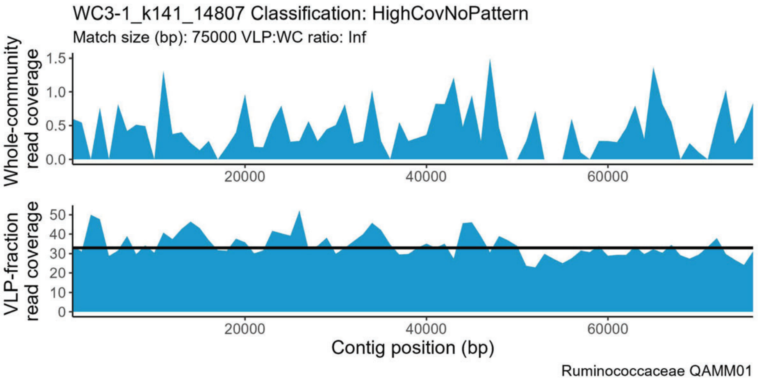
WC3-1_k141_14807 Classification: HighCovNoPattern
Match size (bp): 75000 VLP:WC ratio: Inf
Whole-community read coverage
VLP-fraction read coverage
Contig position (bp)
Ruminococcaceae QAMM01

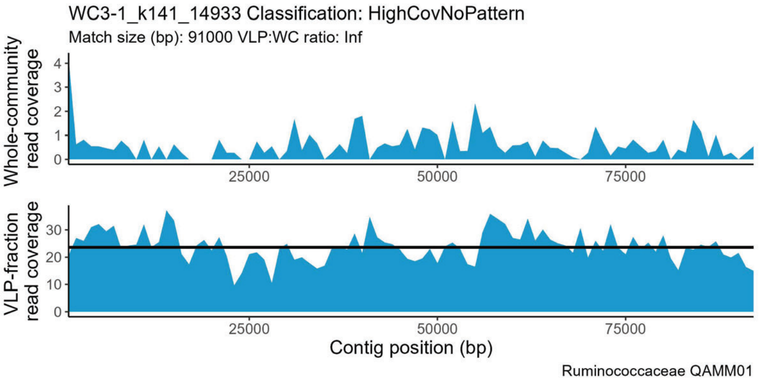
WC3-1_k141_14933 Classification: HighCovNoPattern
Match size (bp): 91000 VLP:WC ratio: Inf
Whole-community read coverage
VLP-fraction read coverage
Contig position (bp)
Ruminococcaceae QAMM01

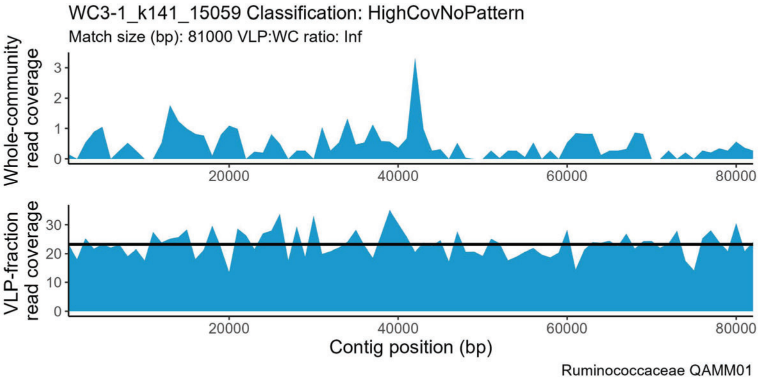
WC3-1_k141_15059 Classification: HighCovNoPattern
Match size (bp): 81000 VLP:WC ratio: Inf
Whole-community read coverage
VLP-fraction read coverage
Contig position (bp)
Ruminococcaceae QAMM01

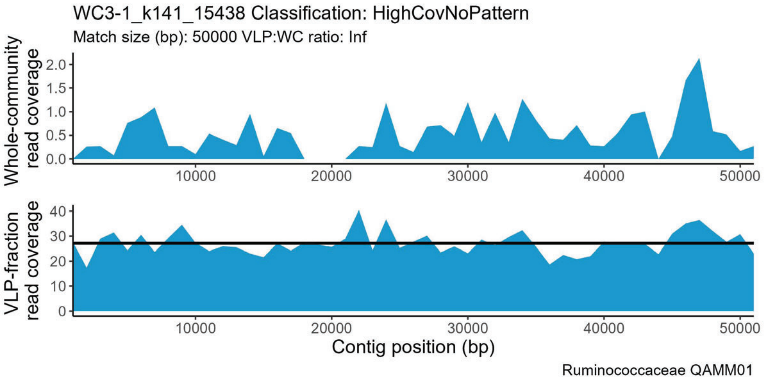
WC3-1_k141_15438 Classification: HighCovNoPattern
Match size (bp): 50000 VLP:WC ratio: Inf
Whole-community read coverage
VLP-fraction read coverage
Contig position (bp)
Ruminococcaceae QAMM01

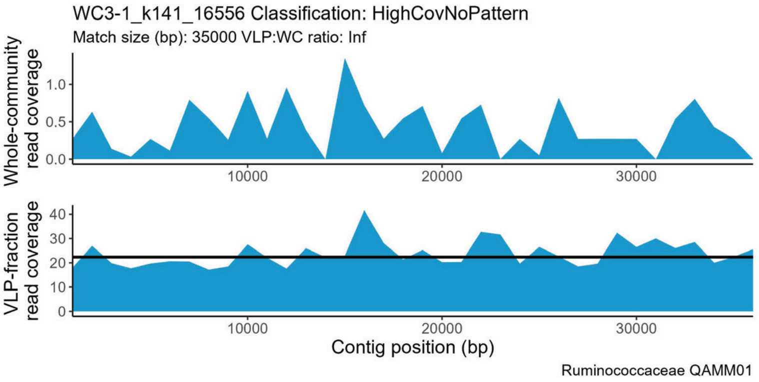
WC3-1_k141_16556 Classification: HighCovNoPattern
Match size (bp): 35000 VLP:WC ratio: Inf
Whole-community read coverage
VLP-fraction read coverage
Contig position (bp)
Ruminococcaceae QAMM01

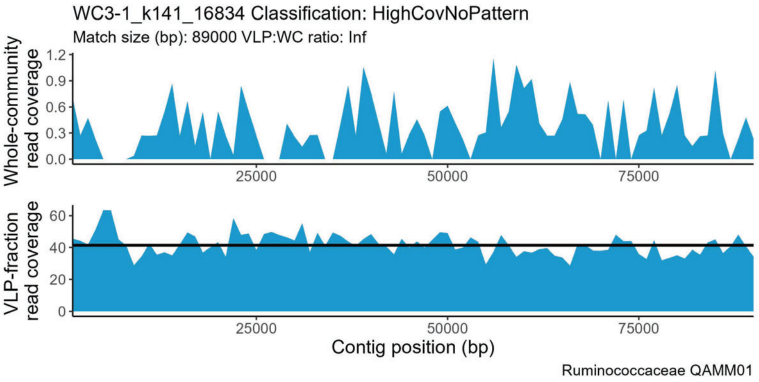
WC3-1_k141_16834 Classification: HighCovNoPattern
Match size (bp): 89000 VLP:WC ratio: Inf
Whole-community read coverage
VLP-fraction read coverage
Contig position (bp)
Ruminococcaceae QAMM01

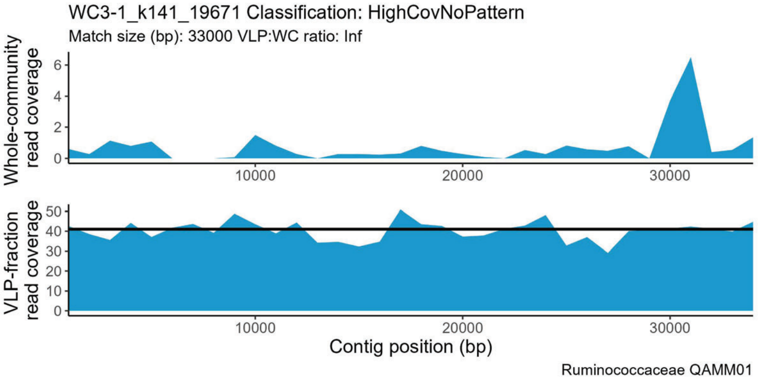
WC3-1_k141_19671 Classification: HighCovNoPattern
Match size (bp): 33000 VLP:WC ratio: Inf
Whole-community read coverage
VLP-fraction read coverage
Contig position (bp)
Ruminococcaceae QAMM01

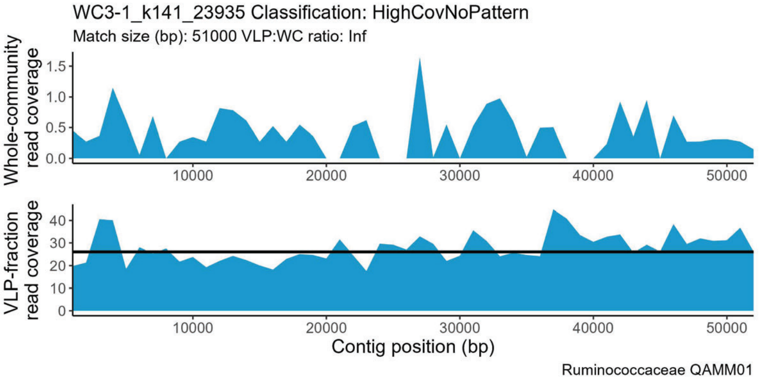
WC3-1_k141_23935 Classification: HighCovNoPattern
Match size (bp): 51000 VLP:WC ratio: Inf
Whole-community read coverage
VLP-fraction read coverage
Contig position (bp)
Ruminococcaceae QAMM01

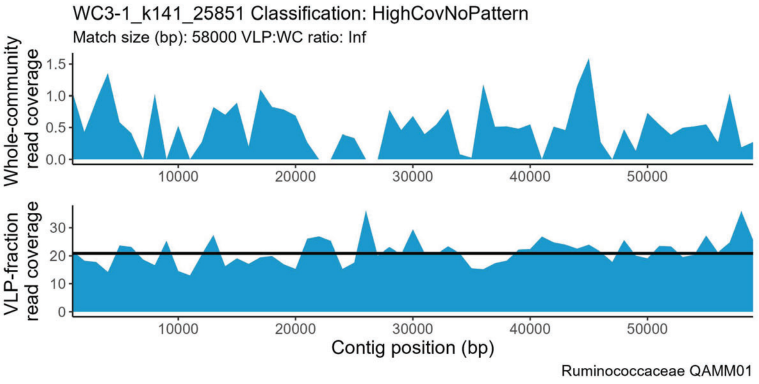
WC3-1_k141_25851 Classification: HighCovNoPattern
Match size (bp): 58000 VLP:WC ratio: Inf
Whole-community read coverage
VLP-fraction read coverage
Contig position (bp)
Ruminococcaceae QAMM01

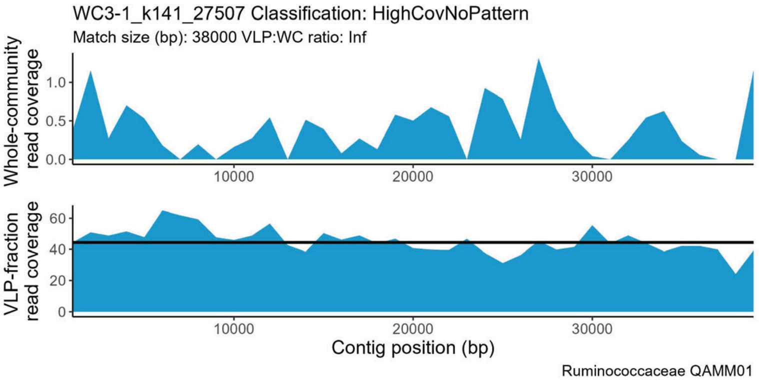
WC3-1_k141_27507 Classification: HighCovNoPattern
Match size (bp): 38000 VLP:WC ratio: Inf
Whole-community read coverage
VLP-fraction read coverage
Contig position (bp)
Ruminococcaceae QAMM01

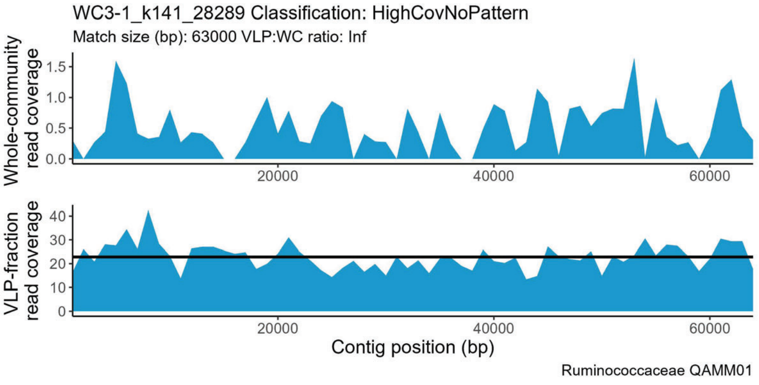
WC3-1_k141_28289 Classification: HighCovNoPattern
Match size (bp): 63000 VLP:WC ratio: Inf
Whole-community read coverage
VLP-fraction read coverage
Contig position (bp)
Ruminococcaceae QAMM01

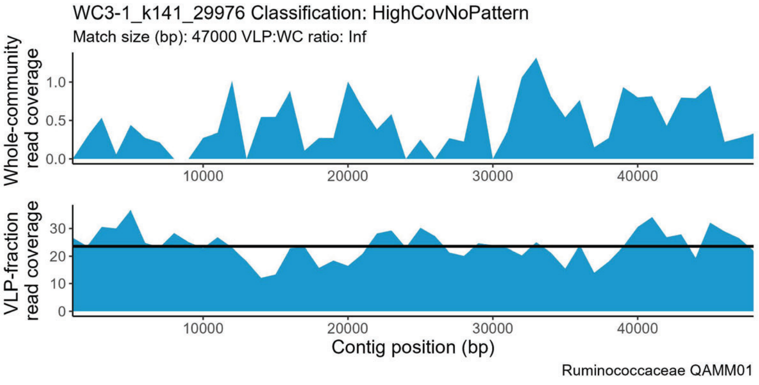
WC3-1_k141_29976 Classification: HighCovNoPattern
Match size (bp): 47000 VLP:WC ratio: Inf
Whole-community read coverage
VLP-fraction read coverage
Contig position (bp)
Ruminococcaceae QAMM01

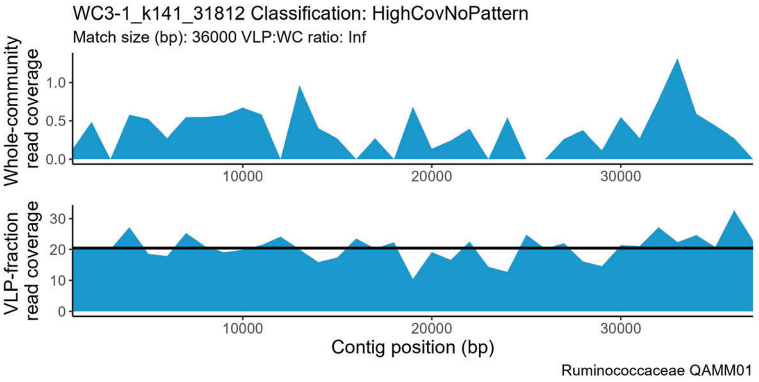
WC3-1_k141_31812 Classification: HighCovNoPattern
Match size (bp): 36000 VLP:WC ratio: Inf
Whole-community read coverage
VLP-fraction read coverage
Contig position (bp)
Ruminococcaceae QAMM01

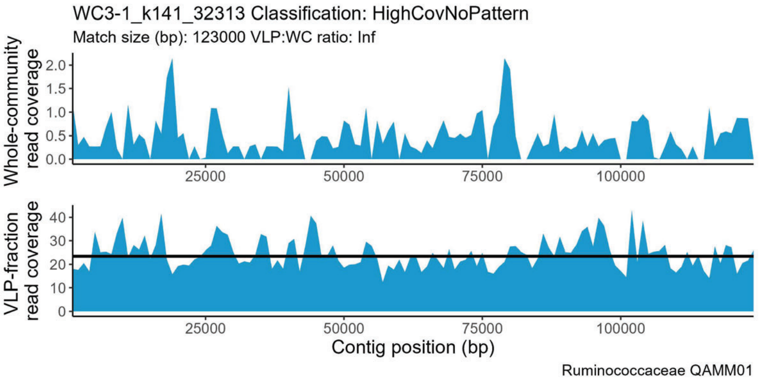
WC3-1_k141_32313 Classification: HighCovNoPattern
Match size (bp): 123000 VLP:WC ratio: Inf
Whole-community read coverage
VLP-fraction read coverage
Contig position (bp)
Ruminococcaceae QAMM01

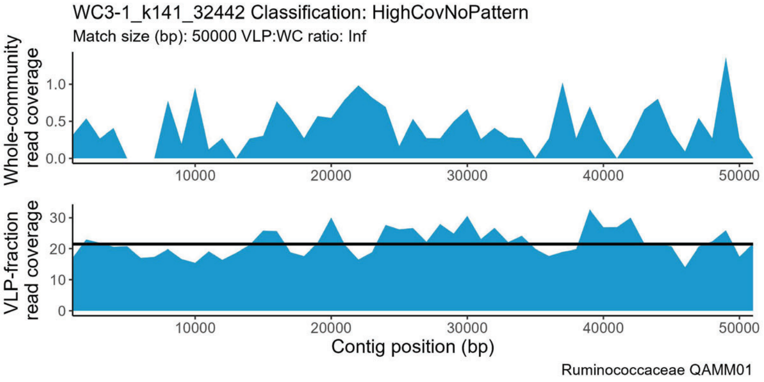
WC3-1_k141_32442 Classification: HighCovNoPattern
Match size (bp): 50000 VLP:WC ratio: Inf
Whole-community read coverage
VLP-fraction read coverage
Contig position (bp)
Ruminococcaceae QAMM01

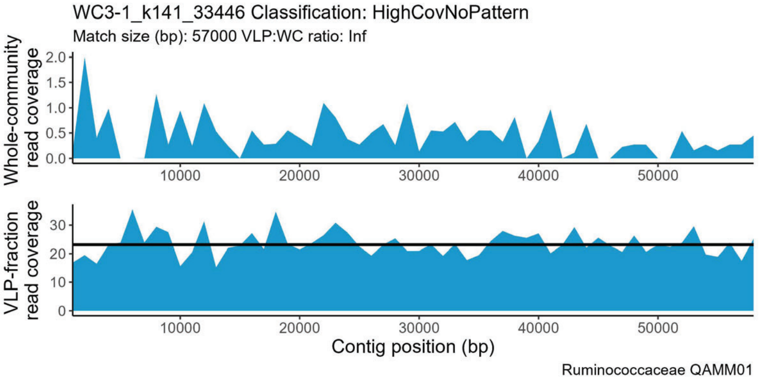
WC3-1_k141_33446 Classification: HighCovNoPattern
Match size (bp): 57000 VLP:WC ratio: Inf
Whole-community read coverage
VLP-fraction read coverage
Contig position (bp)
Ruminococcaceae QAMM01

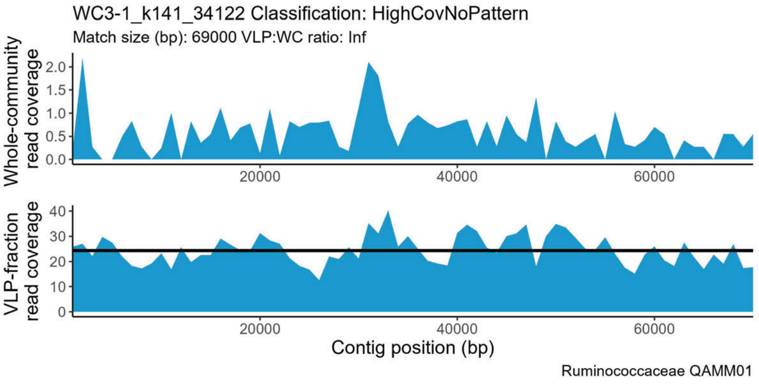
WC3-1_k141_34122 Classification: HighCovNoPattern
Match size (bp): 69000 VLP:WC ratio: Inf
Whole-community read coverage
VLP-fraction read coverage
Contig position (bp)
Ruminococcaceae QAMM01

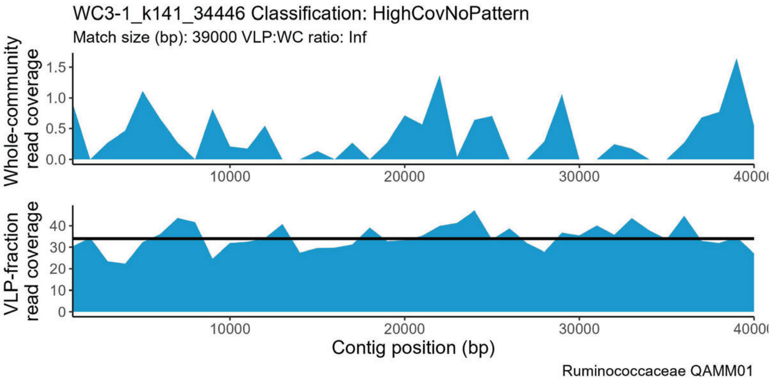
WC3-1_k141_34446 Classification: HighCovNoPattern
Match size (bp): 39000 VLP:WC ratio: Inf
Whole-community read coverage
VLP-fraction read coverage
Contig position (bp)
Ruminococcaceae QAMM01

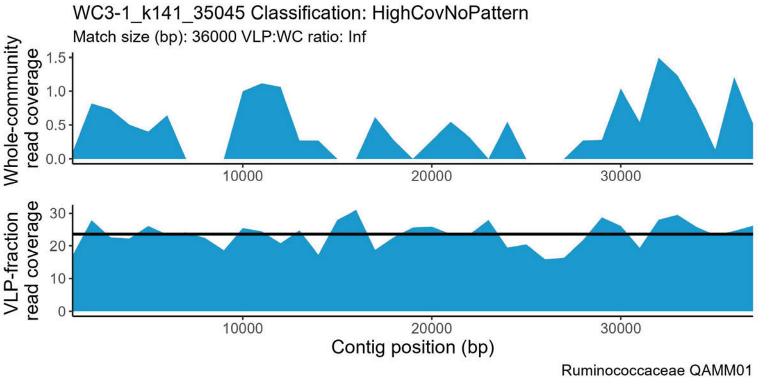
WC3-1_k141_35045 Classification: HighCovNoPattern
Match size (bp): 36000 VLP:WC ratio: Inf
Whole-community read coverage
VLP-fraction read coverage
Contig position (bp)
Ruminococcaceae QAMM01

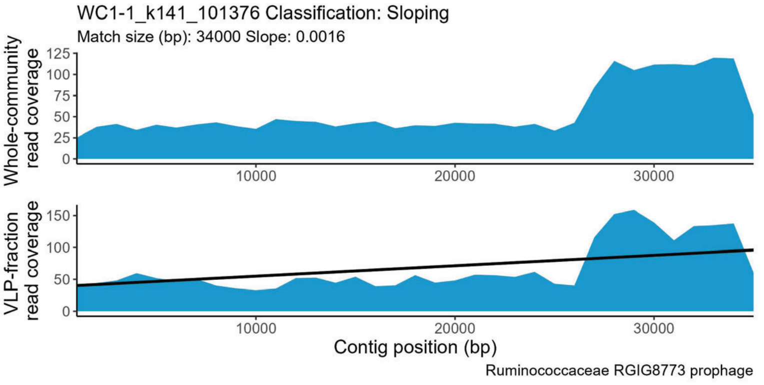
WC1-1_k141_101376 Classification: Sloping
Match size (bp): 34000 Slope: 0.0016
Whole-community read coverage
VLP-fraction read coverage
Contig position (bp)
Ruminococcaceae RGIG8773 prophage

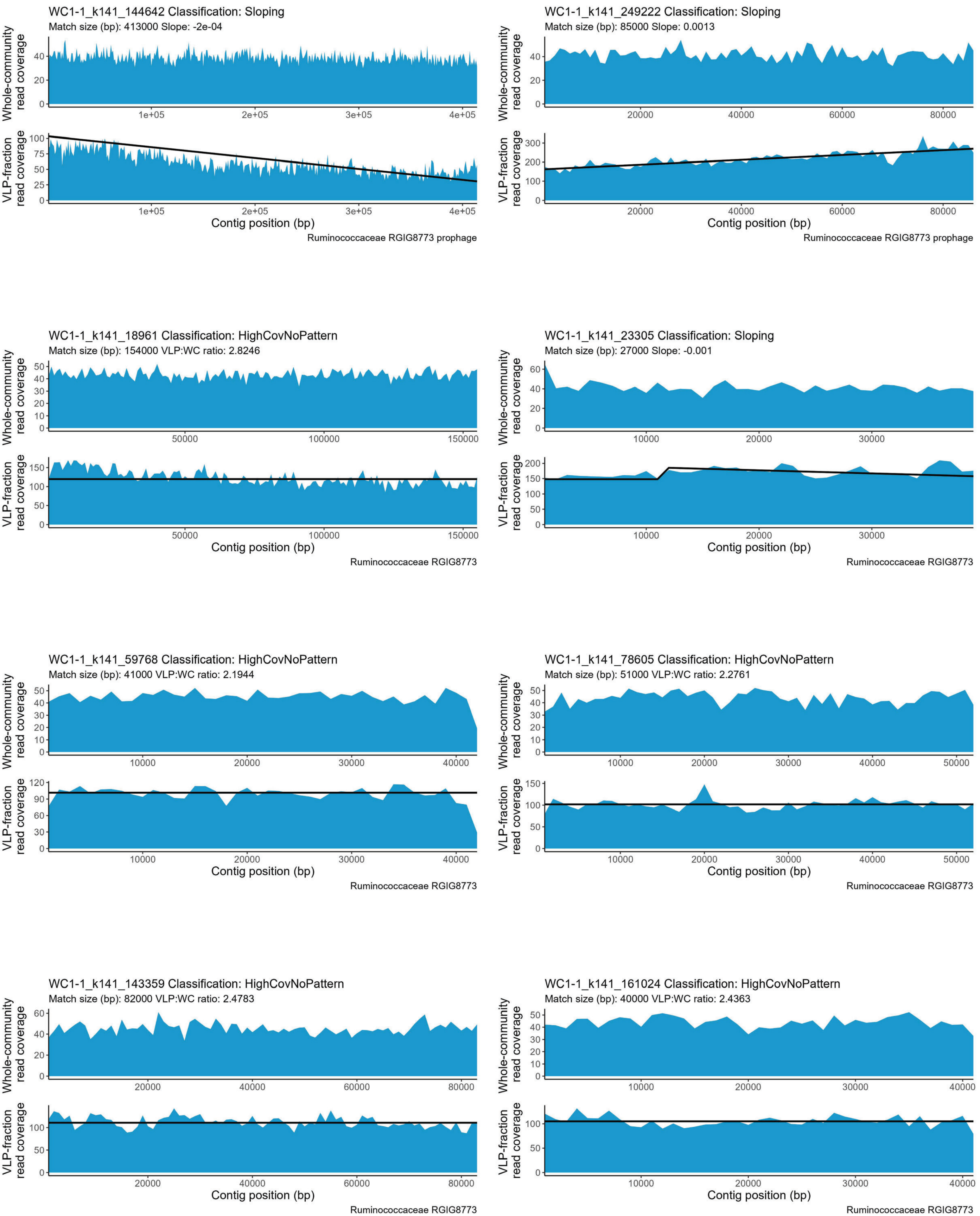
WC1-1_k141_144642 Classification: Sloping
Match size (bp): 413000 Slope: -2e-04
Ruminococcaceae RGIG8773 prophage
WC1-1_k141_249222 Classification: Sloping
Match size (bp): 85000 Slope: 0.0013
Ruminococcaceae RGIG8773 prophage
WC1-1_k141_18961 Classification: HighCovNoPattern
Match size (bp): 154000 VLP:WC ratio: 2.8246
Ruminococcaceae RGIG8773
WC1-1_k141_23305 Classification: Sloping
Match size (bp): 27000 Slope: -0.001
Ruminococcaceae RGIG8773
WC1-1_k141_59768 Classification: HighCovNoPattern
Match size (bp): 41000 VLP:WC ratio: 2.1944
Ruminococcaceae RGIG8773
WC1-1_k141_78605 Classification: HighCovNoPattern
Match size (bp): 51000 VLP:WC ratio: 2.2761
Ruminococcaceae RGIG8773
WC1-1_k141_143359 Classification: HighCovNoPattern
Match size (bp): 82000 VLP:WC ratio: 2.4783
Ruminococcaceae RGIG8773
WC1-1_k141_161024 Classification: HighCovNoPattern
Match size (bp): 40000 VLP:WC ratio: 2.4363
Ruminococcaceae RGIG8773
Whole-community read coverage
VLP-fraction read coverage
Contig position (bp)

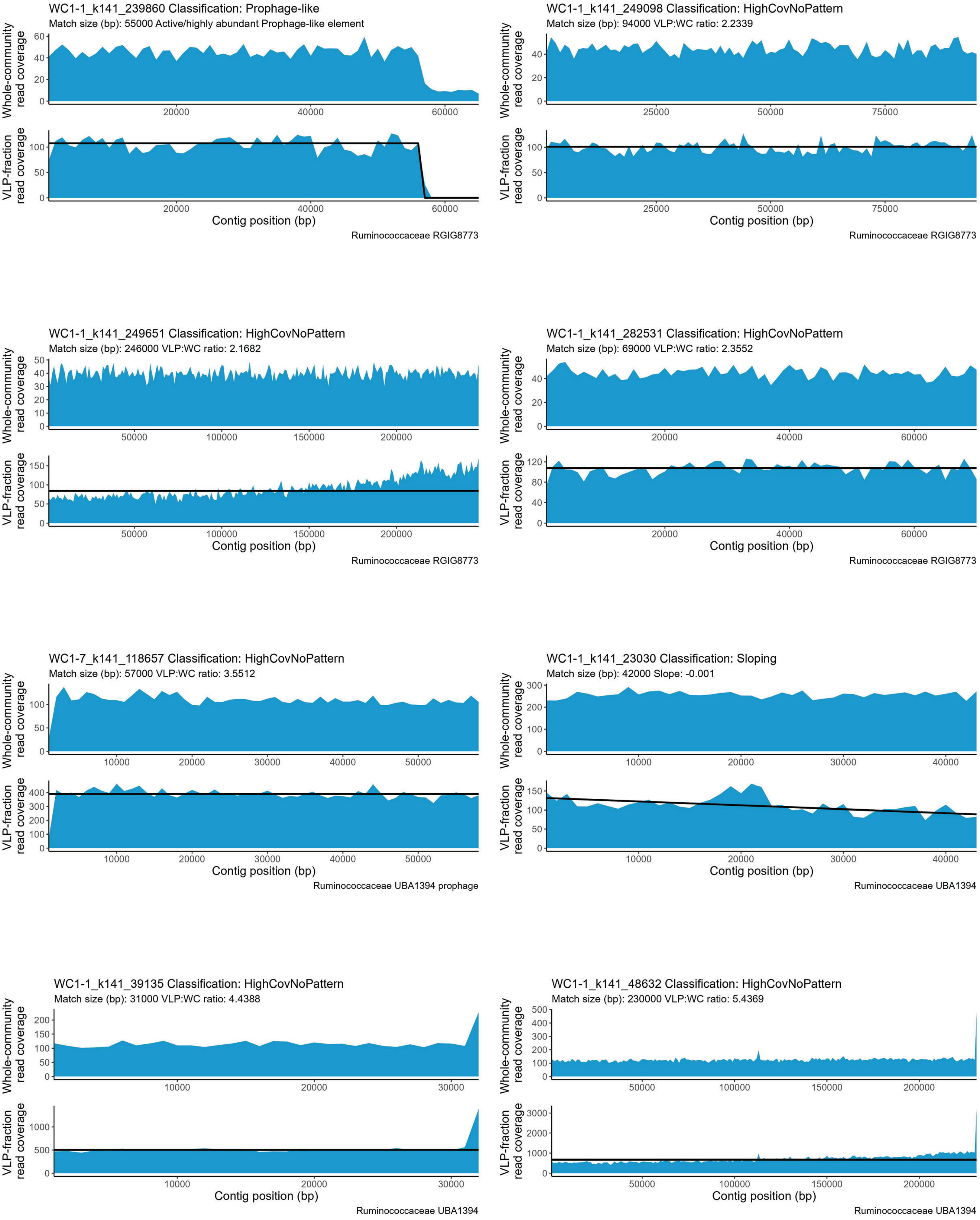
WC1-1_k141_239860 Classification: Prophage-like
Match size (bp): 55000 Active/highly abundant Prophage-like element
Whole-community read coverage
VLP-fraction read coverage
Contig position (bp)
Ruminococcaceae RGIG8773
WC1-1_k141_249098 Classification: HighCovNoPattern
Match size (bp): 94000 VLP:WC ratio: 2.2339
Whole-community read coverage
VLP-fraction read coverage
Contig position (bp)
Ruminococcaceae RGIG8773
WC1-1_k141_249651 Classification: HighCovNoPattern
Match size (bp): 246000 VLP:WC ratio: 2.1682
Whole-community read coverage
VLP-fraction read coverage
Contig position (bp)
Ruminococcaceae RGIG8773
WC1-1_k141_282531 Classification: HighCovNoPattern
Match size (bp): 69000 VLP:WC ratio: 2.3552
Whole-community read coverage
VLP-fraction read coverage
Contig position (bp)
Ruminococcaceae RGIG8773
WC1-7_k141_118657 Classification: HighCovNoPattern
Match size (bp): 57000 VLP:WC ratio: 3.5512
Whole-community read coverage
VLP-fraction read coverage
Contig position (bp)
Ruminococcaceae UBA1394 prophage
WC1-1_k141_23030 Classification: Sloping
Match size (bp): 42000 Slope: -0.001
Whole-community read coverage
VLP-fraction read coverage
Contig position (bp)
Ruminococcaceae UBA1394
WC1-1_k141_39135 Classification: HighCovNoPattern
Match size (bp): 31000 VLP:WC ratio: 4.4388
Whole-community read coverage
VLP-fraction read coverage
Contig position (bp)
Ruminococcaceae UBA1394
WC1-1_k141_48632 Classification: HighCovNoPattern
Match size (bp): 230000 VLP:WC ratio: 5.4369
Whole-community read coverage
VLP-fraction read coverage
Contig position (bp)
Ruminococcaceae UBA1394

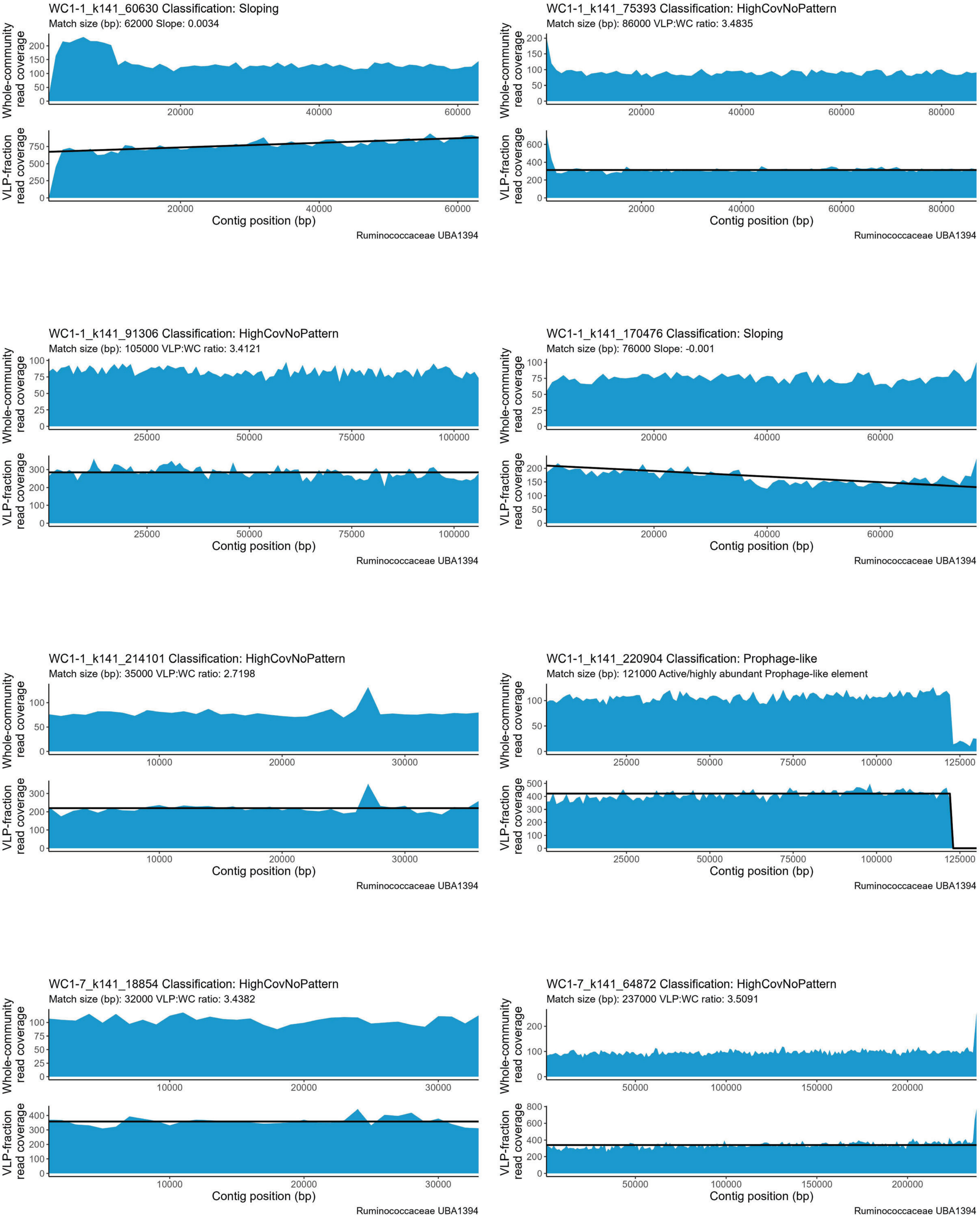
WC1-1_k141_60630 Classification: Sloping
Match size (bp): 62000 Slope: 0.0034
Ruminococcaceae UBA1394
WC1-1_k141_75393 Classification: HighCovNoPattern
Match size (bp): 86000 VLP:WC ratio: 3.4835
Ruminococcaceae UBA1394
WC1-1_k141_91306 Classification: HighCovNoPattern
Match size (bp): 105000 VLP:WC ratio: 3.4121
Ruminococcaceae UBA1394
WC1-1_k141_170476 Classification: Sloping
Match size (bp): 76000 Slope: -0.001
Ruminococcaceae UBA1394
WC1-1_k141_214101 Classification: HighCovNoPattern
Match size (bp): 35000 VLP:WC ratio: 2.7198
Ruminococcaceae UBA1394
WC1-1_k141_220904 Classification: Prophage-like
Match size (bp): 121000 Active/highly abundant Prophage-like element
Ruminococcaceae UBA1394
WC1-7_k141_18854 Classification: HighCovNoPattern
Match size (bp): 32000 VLP:WC ratio: 3.4382
Ruminococcaceae UBA1394
WC1-7_k141_64872 Classification: HighCovNoPattern
Match size (bp): 237000 VLP:WC ratio: 3.5091
Ruminococcaceae UBA1394
Whole-community read coverage
VLP-fraction read coverage
Contig position (bp)

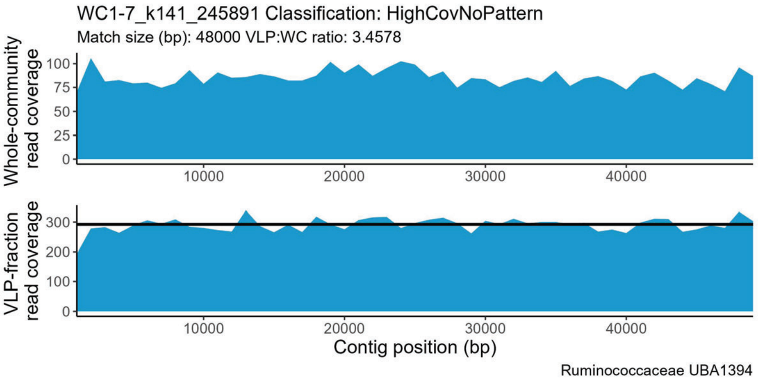
WC1-7_k141_245891 Classification: HighCovNoPattern
Match size (bp): 48000 VLP:WC ratio: 3.4578
Whole-community read coverage
VLP-fraction read coverage
Contig position (bp)
Ruminococcaceae UBA1394

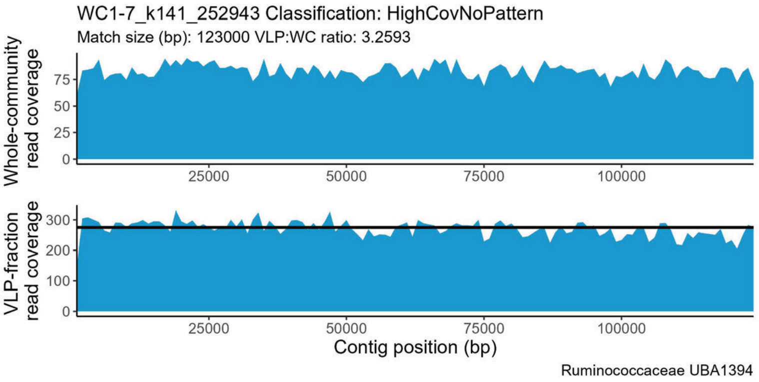
WC1-7_k141_252943 Classification: HighCovNoPattern
Match size (bp): 123000 VLP:WC ratio: 3.2593
Whole-community read coverage
VLP-fraction read coverage
Contig position (bp)
Ruminococcaceae UBA1394

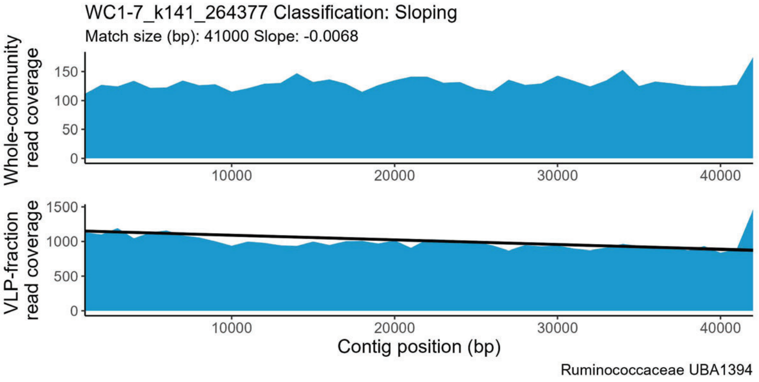
WC1-7_k141_264377 Classification: Sloping
Match size (bp): 41000 Slope: -0.0068
Whole-community read coverage
VLP-fraction read coverage
Contig position (bp)
Ruminococcaceae UBA1394

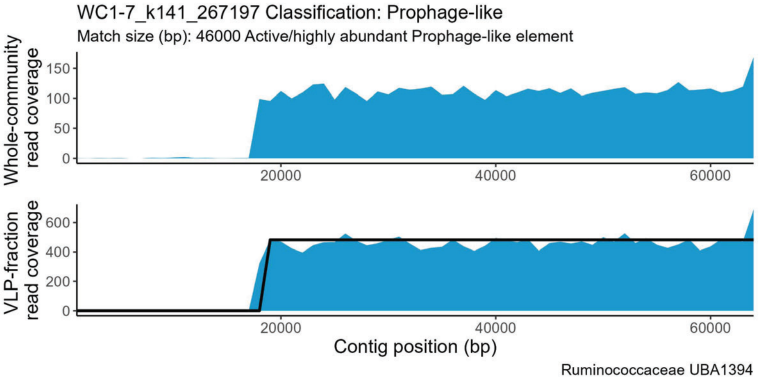
WC1-7_k141_267197 Classification: Prophage-like
Match size (bp): 46000 Active/highly abundant Prophage-like element
Whole-community read coverage
VLP-fraction read coverage
Contig position (bp)
Ruminococcaceae UBA1394

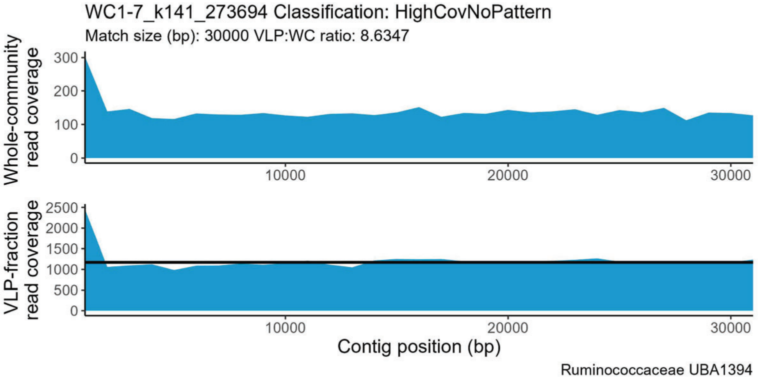
WC1-7_k141_273694 Classification: HighCovNoPattern
Match size (bp): 30000 VLP:WC ratio: 8.6347
Whole-community read coverage
VLP-fraction read coverage
Contig position (bp)
Ruminococcaceae UBA1394

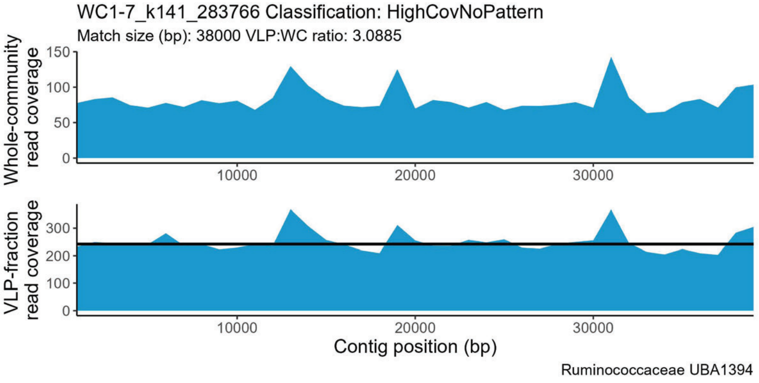
WC1-7_k141_283766 Classification: HighCovNoPattern
Match size (bp): 38000 VLP:WC ratio: 3.0885
Whole-community read coverage
VLP-fraction read coverage
Contig position (bp)
Ruminococcaceae UBA1394

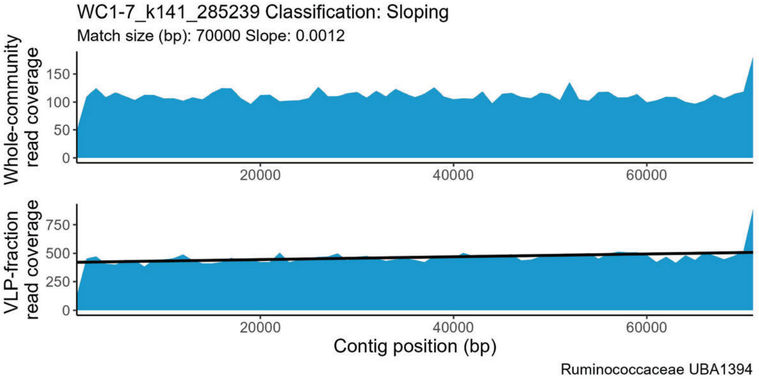
WC1-7_k141_285239 Classification: Sloping
Match size (bp): 70000 Slope: 0.0012
Whole-community read coverage
VLP-fraction read coverage
Contig position (bp)
Ruminococcaceae UBA1394

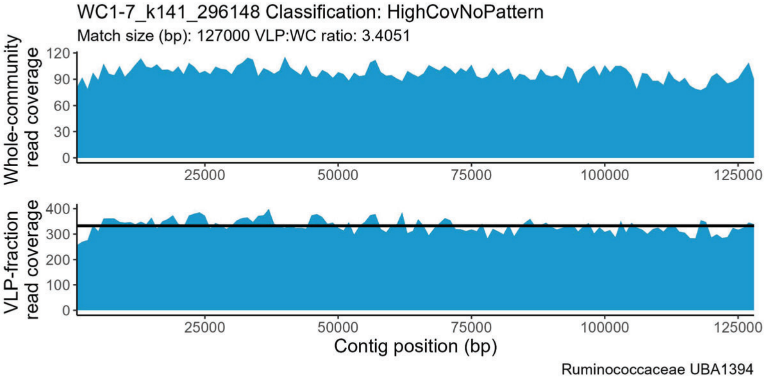
WC1-7_k141_296148 Classification: HighCovNoPattern
Match size (bp): 127000 VLP:WC ratio: 3.4051
Whole-community read coverage
VLP-fraction read coverage
Contig position (bp)
Ruminococcaceae UBA1394

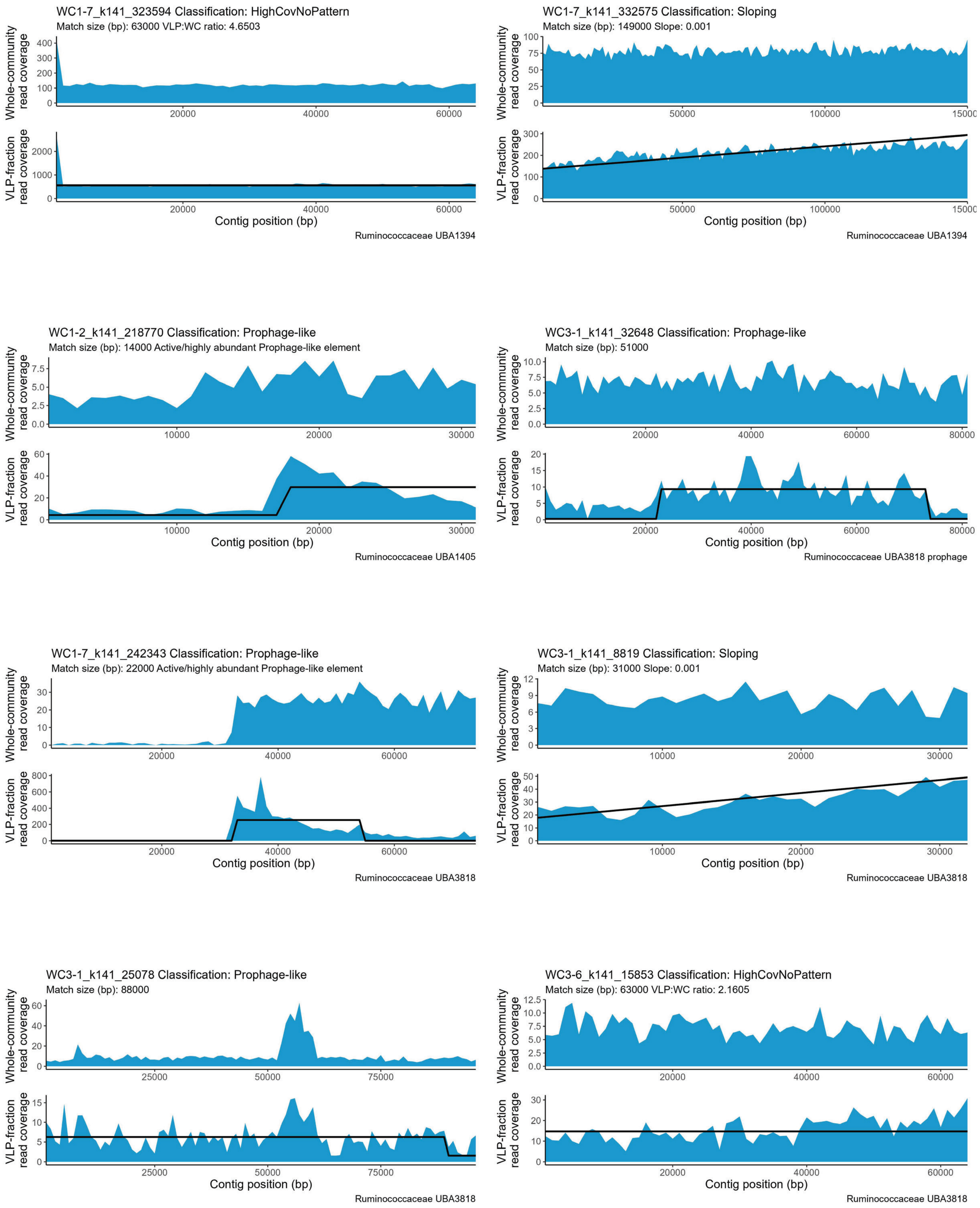

WC1-7_k141_323594 Classification: HighCovNoPattern
Match size (bp): 63000 VLP:WC ratio: 4.6503
Whole-community read coverage
VLP-fraction read coverage
Contig position (bp)
Ruminococcaceae UBA1394
WC1-7_k141_332575 Classification: Sloping
Match size (bp): 149000 Slope: 0.001
Whole-community read coverage
VLP-fraction read coverage
Contig position (bp)
Ruminococcaceae UBA1394
WC1-2_k141_218770 Classification: Prophage-like
Match size (bp): 14000 Active/highly abundant Prophage-like element
Whole-community read coverage
VLP-fraction read coverage
Contig position (bp)
Ruminococcaceae UBA1405
WC3-1_k141_32648 Classification: Prophage-like
Match size (bp): 51000
Whole-community read coverage
VLP-fraction read coverage
Contig position (bp)
Ruminococcaceae UBA3818 prophage
WC1-7_k141_242343 Classification: Prophage-like
Match size (bp): 22000 Active/highly abundant Prophage-like element
Whole-community read coverage
VLP-fraction read coverage
Contig position (bp)
Ruminococcaceae UBA3818
WC3-1_k141_8819 Classification: Sloping
Match size (bp): 31000 Slope: 0.001
Whole-community read coverage
VLP-fraction read coverage
Contig position (bp)
Ruminococcaceae UBA3818
WC3-1_k141_25078 Classification: Prophage-like
Match size (bp): 88000
Whole-community read coverage
VLP-fraction read coverage
Contig position (bp)
Ruminococcaceae UBA3818
WC3-6_k141_15853 Classification: HighCovNoPattern
Match size (bp): 63000 VLP:WC ratio: 2.1605
Whole-community read coverage
VLP-fraction read coverage
Contig position (bp)
Ruminococcaceae UBA3818

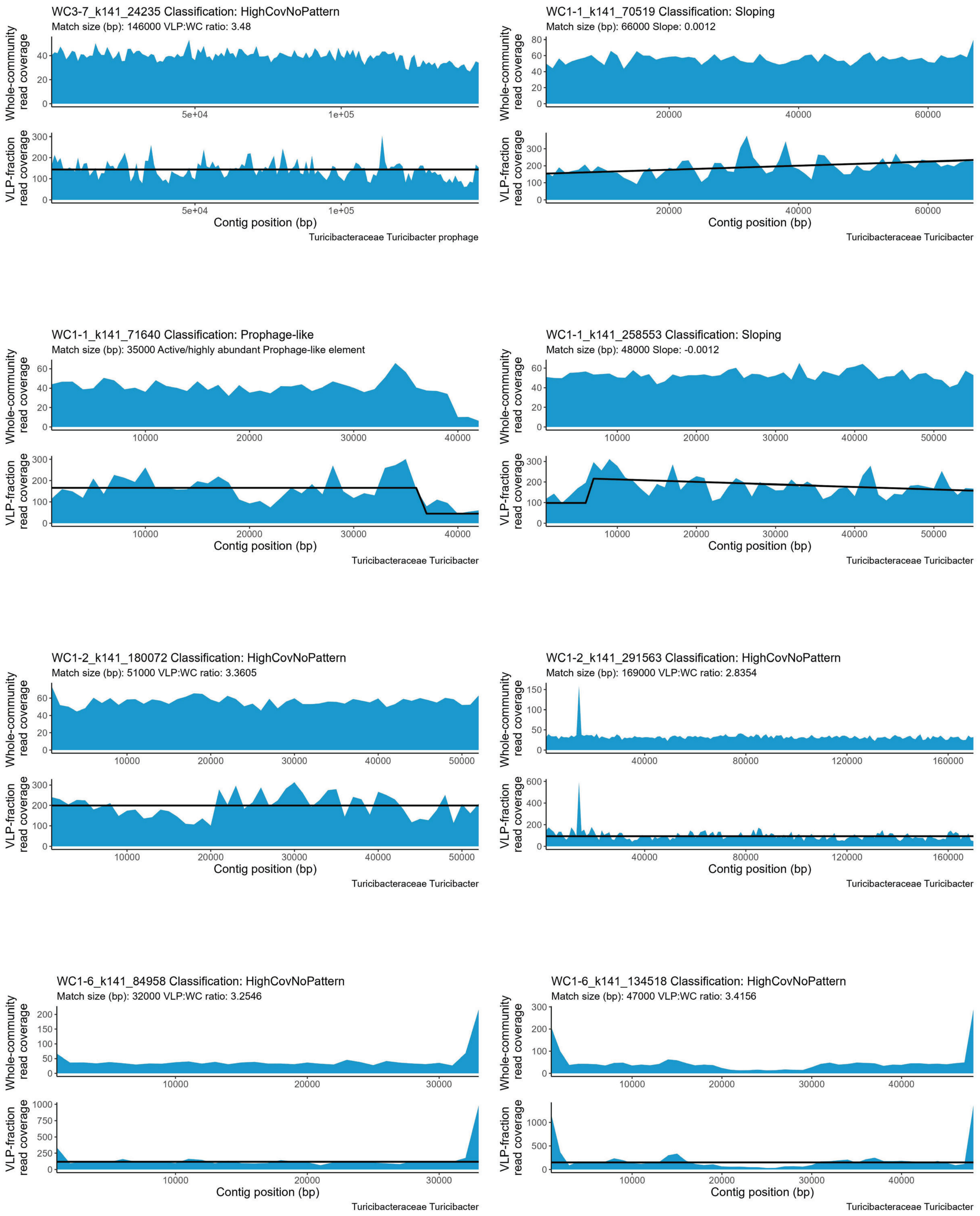
WC3-7_k141_24235 Classification: HighCovNoPattern
Match size (bp): 146000 VLP:WC ratio: 3.48
Whole-community read coverage
VLP-fraction read coverage
Contig position (bp)
Turicibacteraceae Turicibacter prophage
WC1-1_k141_70519 Classification: Sloping
Match size (bp): 66000 Slope: 0.0012
Whole-community read coverage
VLP-fraction read coverage
Contig position (bp)
Turicibacteraceae Turicibacter
WC1-1_k141_71640 Classification: Prophage-like
Match size (bp): 35000 Active/highly abundant Prophage-like element
Whole-community read coverage
VLP-fraction read coverage
Contig position (bp)
Turicibacteraceae Turicibacter
WC1-1_k141_258553 Classification: Sloping
Match size (bp): 48000 Slope: -0.0012
Whole-community read coverage
VLP-fraction read coverage
Contig position (bp)
Turicibacteraceae Turicibacter
WC1-2_k141_180072 Classification: HighCovNoPattern
Match size (bp): 51000 VLP:WC ratio: 3.3605
Whole-community read coverage
VLP-fraction read coverage
Contig position (bp)
Turicibacteraceae Turicibacter
WC1-2_k141_291563 Classification: HighCovNoPattern
Match size (bp): 169000 VLP:WC ratio: 2.8354
Whole-community read coverage
VLP-fraction read coverage
Contig position (bp)
Turicibacteraceae Turicibacter
WC1-6_k141_84958 Classification: HighCovNoPattern
Match size (bp): 32000 VLP:WC ratio: 3.2546
Whole-community read coverage
VLP-fraction read coverage
Contig position (bp)
Turicibacteraceae Turicibacter
WC1-6_k141_134518 Classification: HighCovNoPattern
Match size (bp): 47000 VLP:WC ratio: 3.4156
Whole-community read coverage
VLP-fraction read coverage
Contig position (bp)
Turicibacteraceae Turicibacter

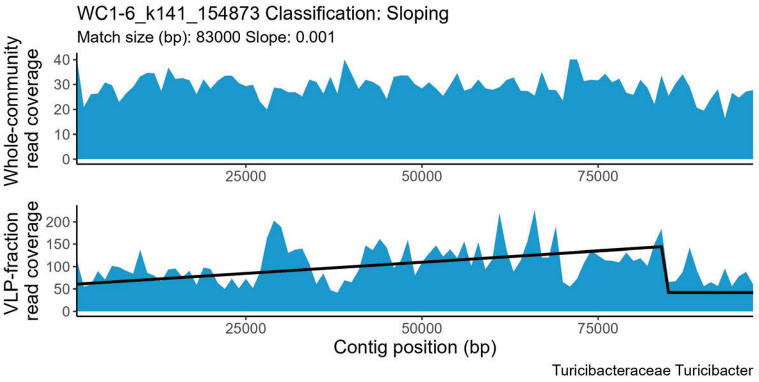
WC1-6_k141_154873 Classification: Sloping
Match size (bp): 83000 Slope: 0.001
Whole-community read coverage
VLP-fraction read coverage
Contig position (bp)
Turicibacteraceae Turicibacter

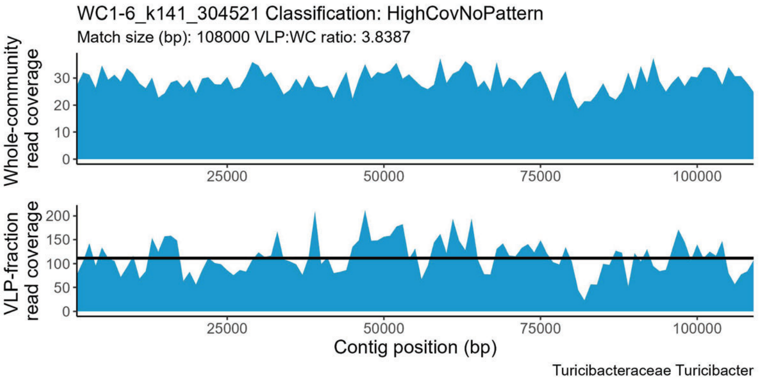
WC1-6_k141_304521 Classification: HighCovNoPattern
Match size (bp): 108000 VLP:WC ratio: 3.8387
Whole-community read coverage
VLP-fraction read coverage
Contig position (bp)
Turicibacteraceae Turicibacter

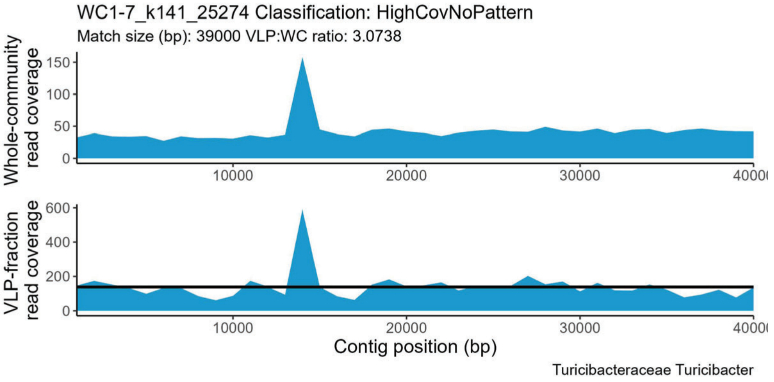
WC1-7_k141_25274 Classification: HighCovNoPattern
Match size (bp): 39000 VLP:WC ratio: 3.0738
Whole-community read coverage
VLP-fraction read coverage
Contig position (bp)
Turicibacteraceae Turicibacter

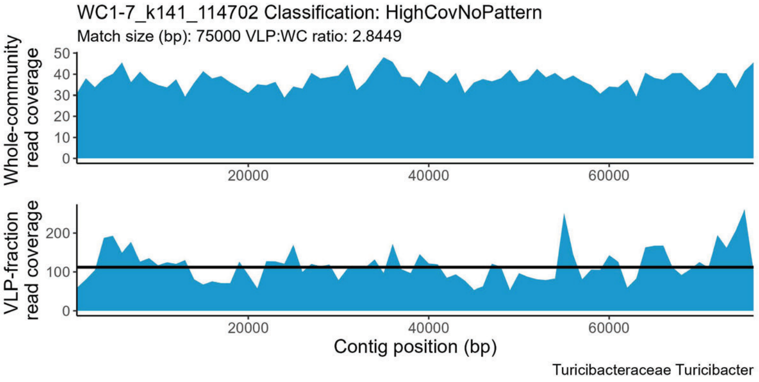
WC1-7_k141_114702 Classification: HighCovNoPattern
Match size (bp): 75000 VLP:WC ratio: 2.8449
Whole-community read coverage
VLP-fraction read coverage
Contig position (bp)
Turicibacteraceae Turicibacter

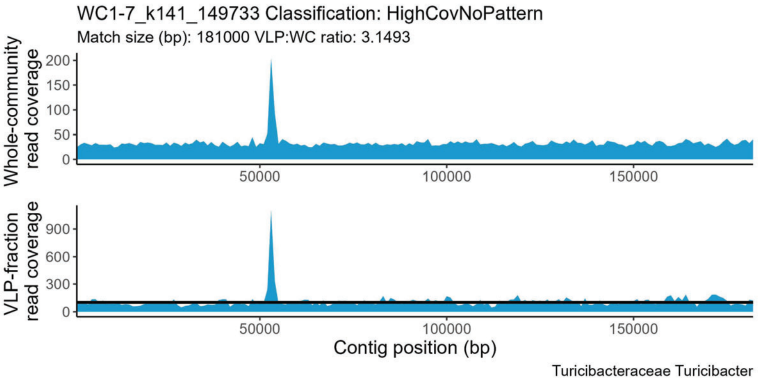
WC1-7_k141_149733 Classification: HighCovNoPattern
Match size (bp): 181000 VLP:WC ratio: 3.1493
Whole-community read coverage
VLP-fraction read coverage
Contig position (bp)
Turicibacteraceae Turicibacter

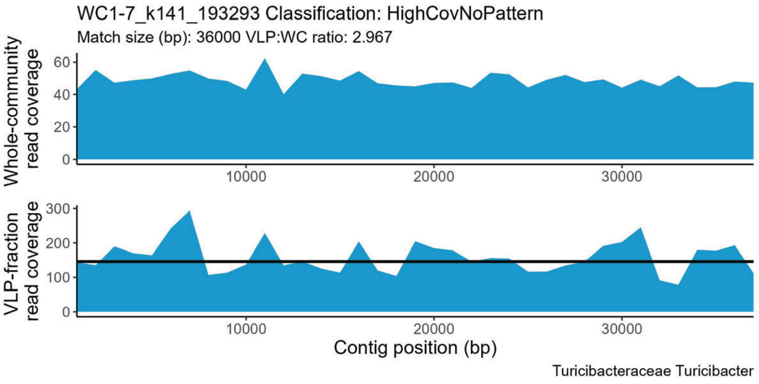
WC1-7_k141_193293 Classification: HighCovNoPattern
Match size (bp): 36000 VLP:WC ratio: 2.967
Whole-community read coverage
VLP-fraction read coverage
Contig position (bp)
Turicibacteraceae Turicibacter

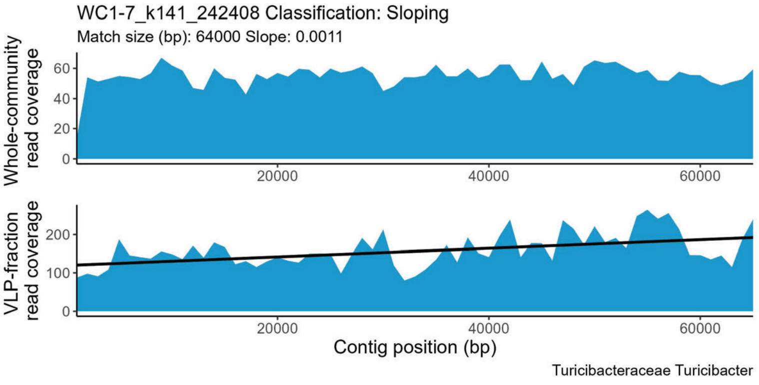
WC1-7_k141_242408 Classification: Sloping
Match size (bp): 64000 Slope: 0.0011
Whole-community read coverage
VLP-fraction read coverage
Contig position (bp)
Turicibacteraceae Turicibacter

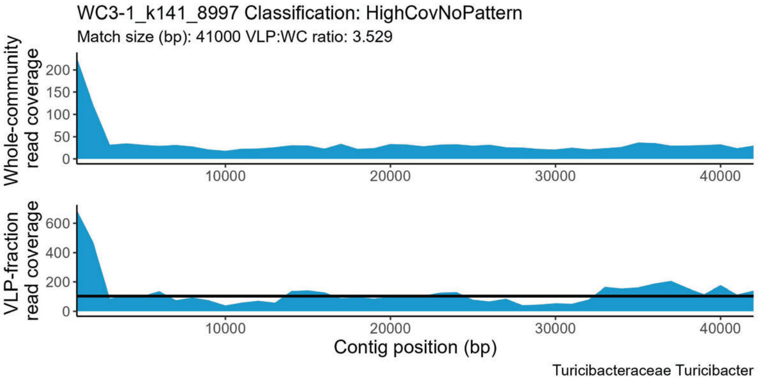
WC3-1_k141_8997 Classification: HighCovNoPattern
Match size (bp): 41000 VLP:WC ratio: 3.529
Whole-community read coverage
VLP-fraction read coverage
Contig position (bp)
Turicibacteraceae Turicibacter

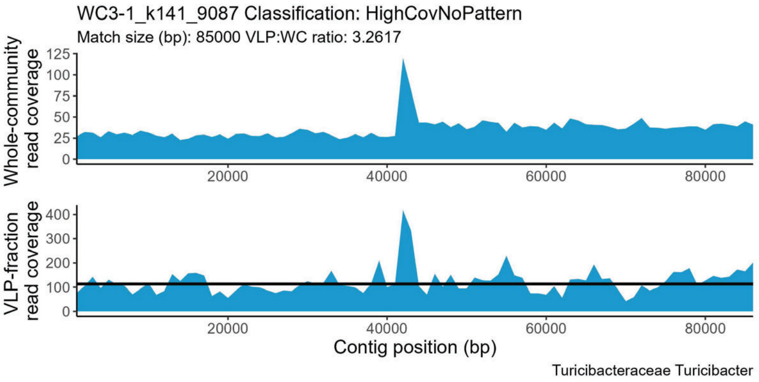
WC3-1_k141_9087 Classification: HighCovNoPattern
Match size (bp): 85000 VLP:WC ratio: 3.2617
Whole-community read coverage
VLP-fraction read coverage
Contig position (bp)
Turicibacteraceae Turicibacter

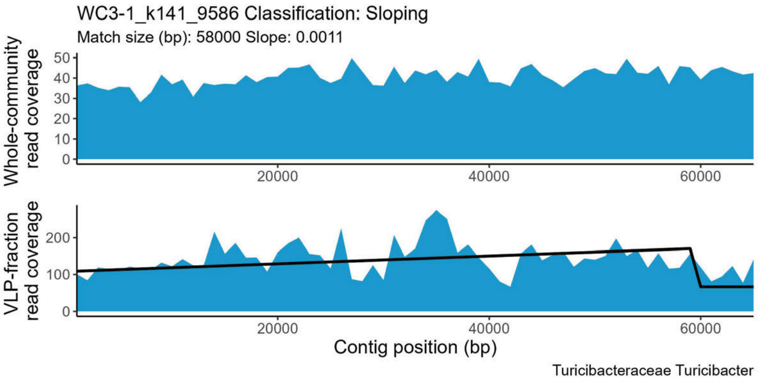
WC3-1_k141_9586 Classification: Sloping
Match size (bp): 58000 Slope: 0.0011
Whole-community read coverage
VLP-fraction read coverage
Contig position (bp)
Turicibacteraceae Turicibacter

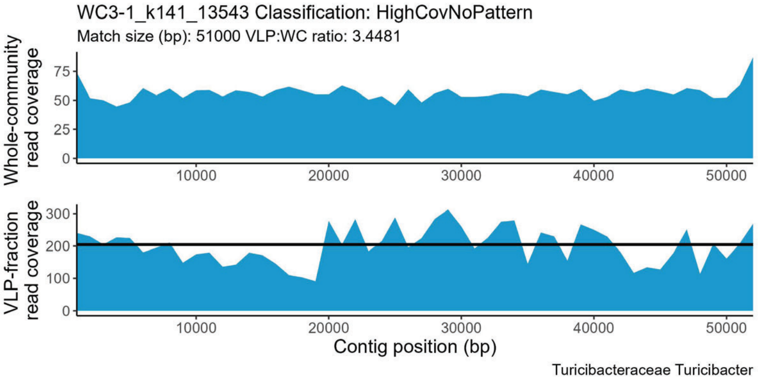
WC3-1_k141_13543 Classification: HighCovNoPattern
Match size (bp): 51000 VLP:WC ratio: 3.4481
Whole-community read coverage
VLP-fraction read coverage
Contig position (bp)
Turicibacteraceae Turicibacter

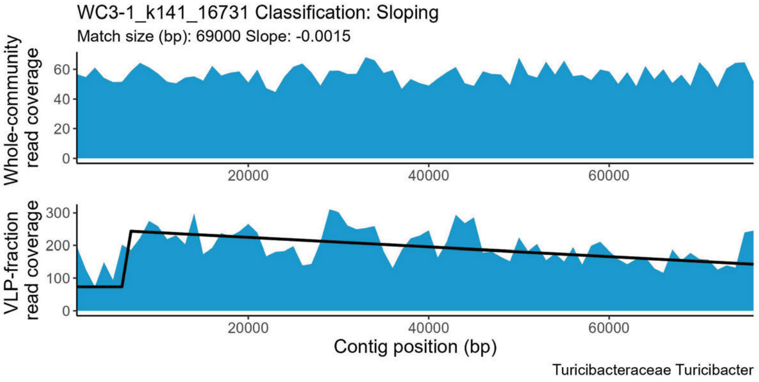
WC3-1_k141_16731 Classification: Sloping
Match size (bp): 69000 Slope: -0.0015
Whole-community read coverage
VLP-fraction read coverage
Contig position (bp)
Turicibacteraceae Turicibacter

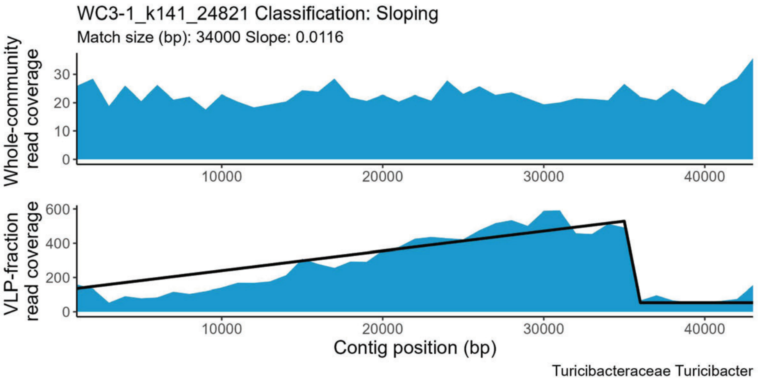
WC3-1_k141_24821 Classification: Sloping
Match size (bp): 34000 Slope: 0.0116
Whole-community read coverage
VLP-fraction read coverage
Contig position (bp)
Turicibacteraceae Turicibacter

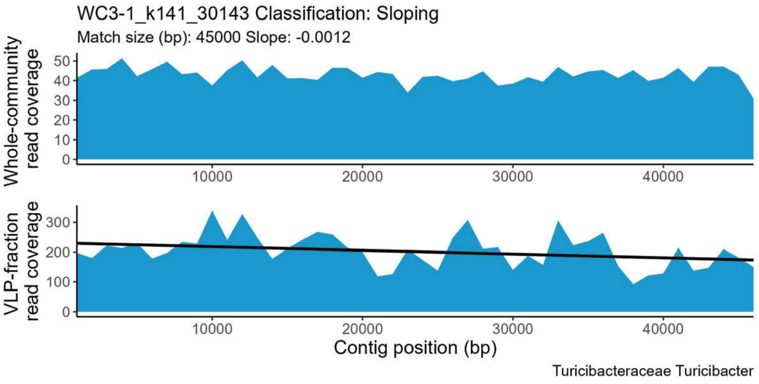
WC3-1_k141_30143 Classification: Sloping
Match size (bp): 45000 Slope: -0.0012
Whole-community read coverage
VLP-fraction read coverage
Contig position (bp)
Turicibacteraceae Turicibacter

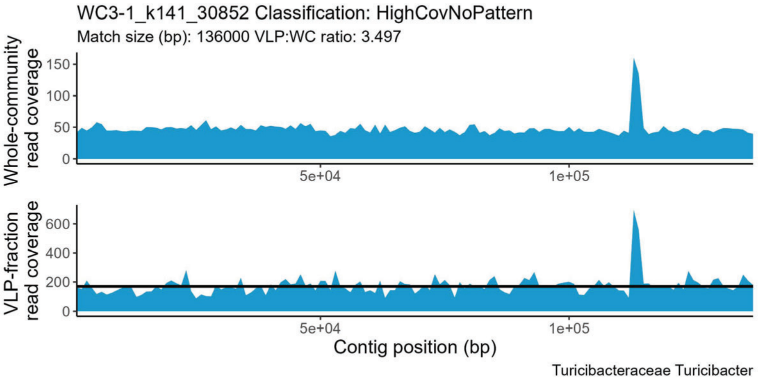
WC3-1_k141_30852 Classification: HighCovNoPattern
Match size (bp): 136000 VLP:WC ratio: 3.497
Whole-community read coverage
VLP-fraction read coverage
Contig position (bp)
Turicibacteraceae Turicibacter

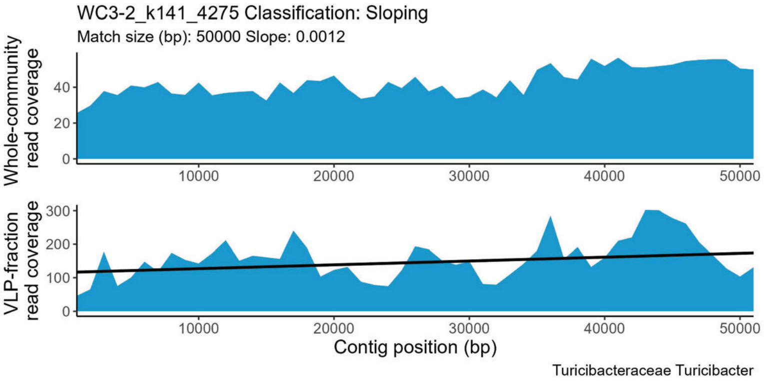
WC3-2_k141_4275 Classification: Sloping
Match size (bp): 50000 Slope: 0.0012
Whole-community read coverage
VLP-fraction read coverage
Contig position (bp)
Turicibacteraceae Turicibacter

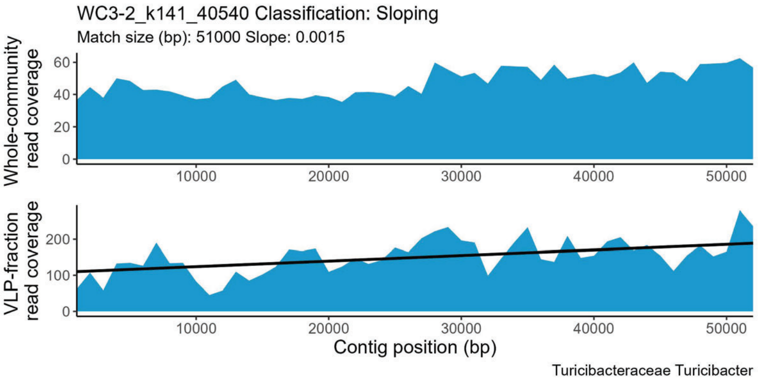
WC3-2_k141_40540 Classification: Sloping
Match size (bp): 51000 Slope: 0.0015
Whole-community read coverage
VLP-fraction read coverage
Contig position (bp)
Turicibacteraceae Turicibacter

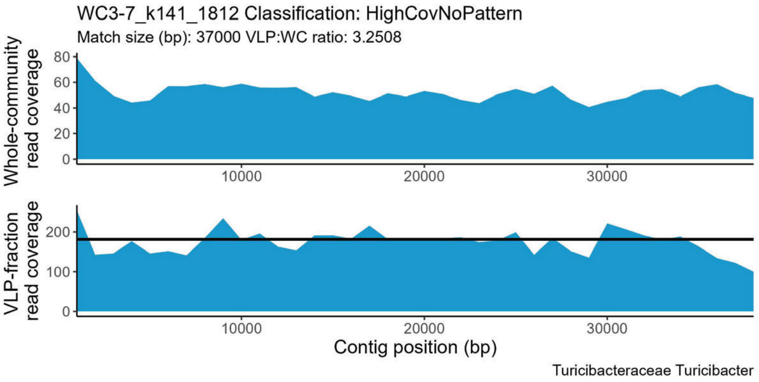
WC3-7_k141_1812 Classification: HighCovNoPattern
Match size (bp): 37000 VLP:WC ratio: 3.2508
Whole-community read coverage
VLP-fraction read coverage
Contig position (bp)
Turicibacteraceae Turicibacter

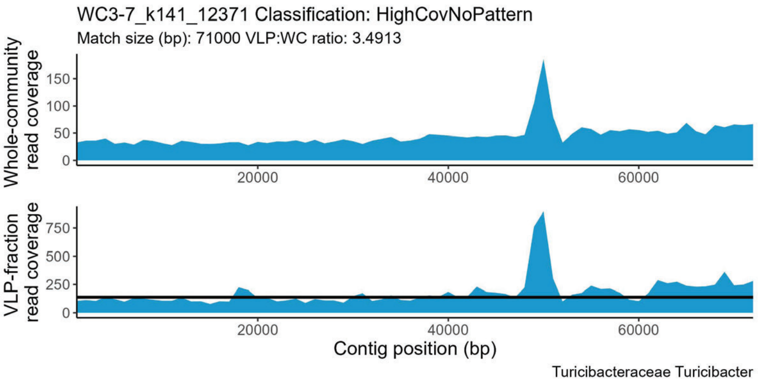
WC3-7_k141_12371 Classification: HighCovNoPattern
Match size (bp): 71000 VLP:WC ratio: 3.4913
Whole-community read coverage
VLP-fraction read coverage
Contig position (bp)
Turicibacteraceae Turicibacter

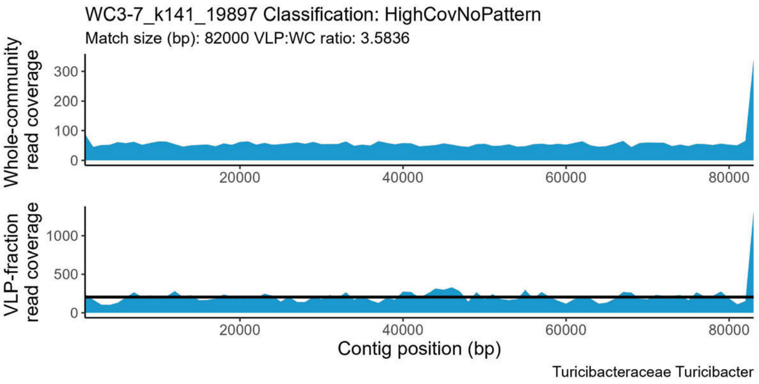
WC3-7_k141_19897 Classification: HighCovNoPattern
Match size (bp): 82000 VLP:WC ratio: 3.5836
Whole-community read coverage
VLP-fraction read coverage
Contig position (bp)
Turicibacteraceae Turicibacter

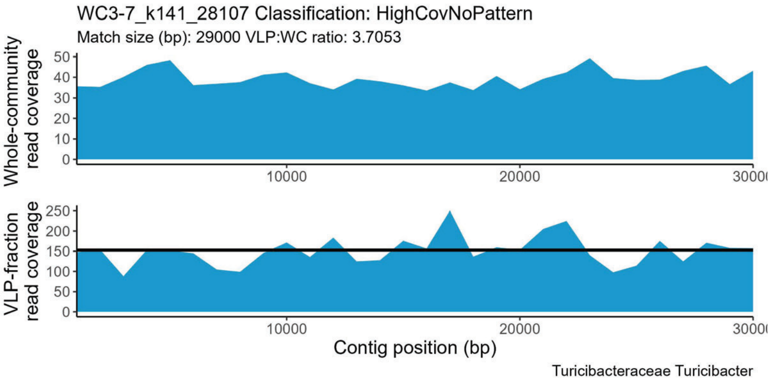
WC3-7_k141_28107 Classification: HighCovNoPattern
Match size (bp): 29000 VLP:WC ratio: 3.7053
Whole-community read coverage
VLP-fraction read coverage
Contig position (bp)
Turicibacteraceae Turicibacter

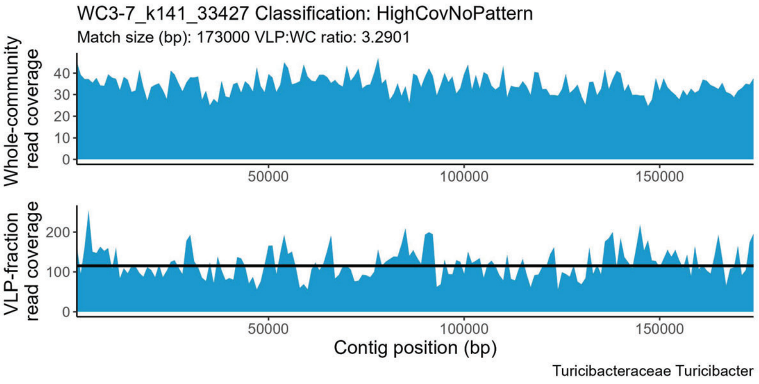
WC3-7_k141_33427 Classification: HighCovNoPattern
Match size (bp): 173000 VLP:WC ratio: 3.2901
Whole-community read coverage
VLP-fraction read coverage
Contig position (bp)
Turicibacteraceae Turicibacter

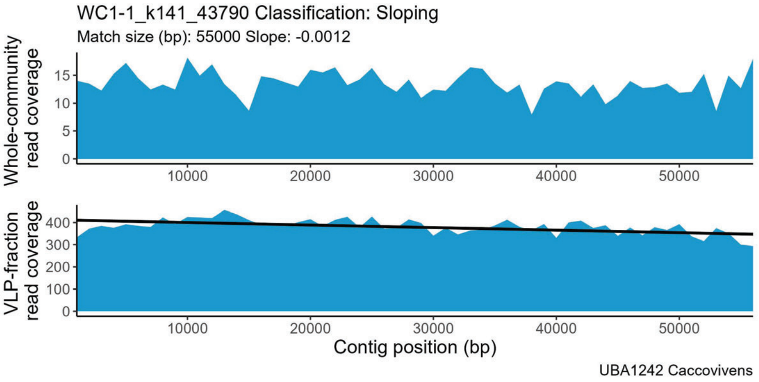
WC1-1_k141_43790 Classification: Sloping
Match size (bp): 55000 Slope: -0.0012
Whole-community read coverage
VLP-fraction read coverage
Contig position (bp)
UBA1242 Caccovivens

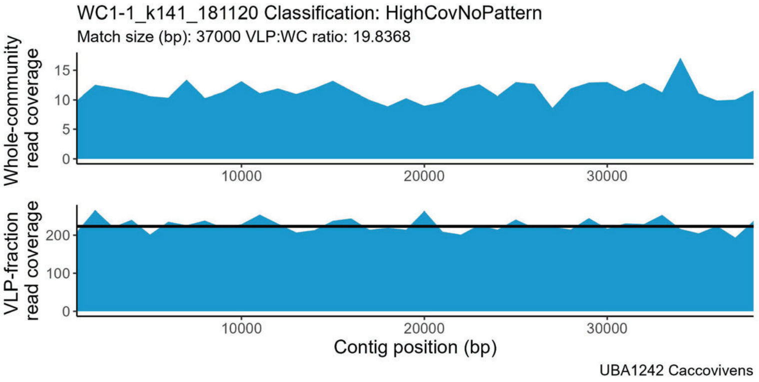
WC1-1_k141_181120 Classification: HighCovNoPattern
Match size (bp): 37000 VLP:WC ratio: 19.8368
Whole-community read coverage
VLP-fraction read coverage
Contig position (bp)
UBA1242 Caccovivens

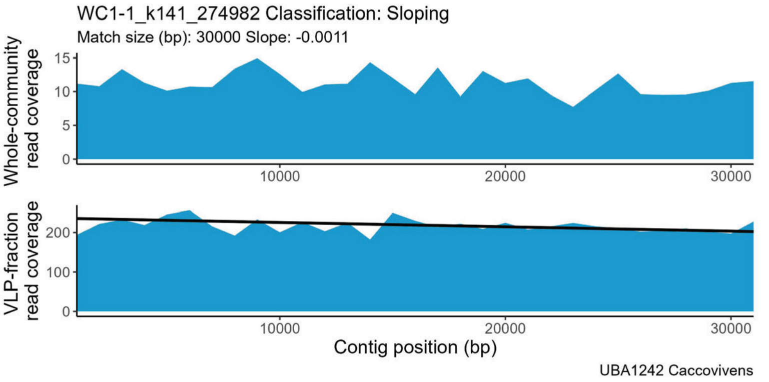
WC1-1_k141_274982 Classification: Sloping
Match size (bp): 30000 Slope: -0.0011
Whole-community read coverage
VLP-fraction read coverage
Contig position (bp)
UBA1242 Caccovivens

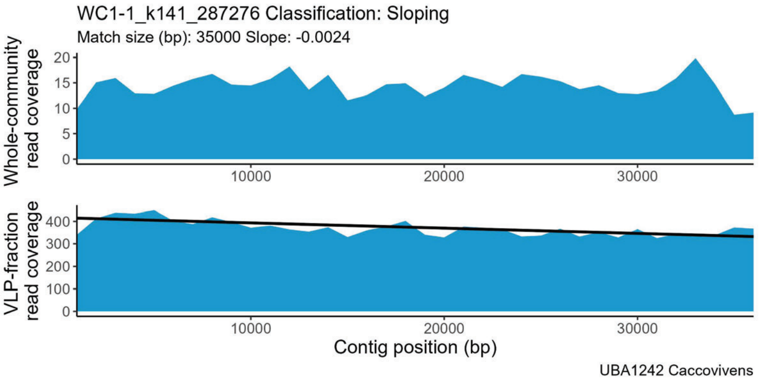
WC1-1_k141_287276 Classification: Sloping
Match size (bp): 35000 Slope: -0.0024
Whole-community read coverage
VLP-fraction read coverage
Contig position (bp)
UBA1242 Caccovivens

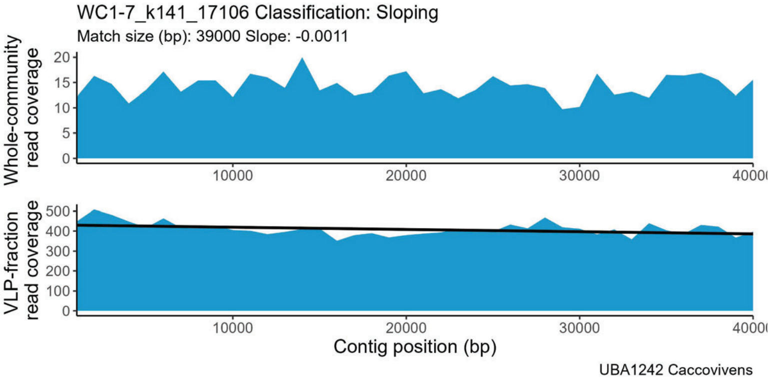
WC1-7_k141_17106 Classification: Sloping
Match size (bp): 39000 Slope: -0.0011
Whole-community read coverage
VLP-fraction read coverage
Contig position (bp)
UBA1242 Caccovivens

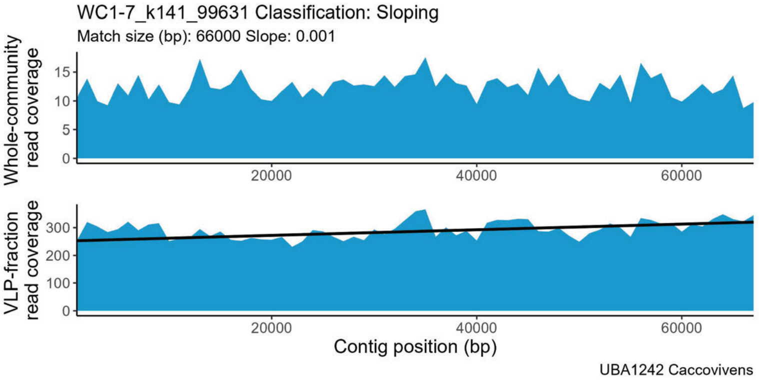
WC1-7_k141_99631 Classification: Sloping
Match size (bp): 66000 Slope: 0.001
Whole-community read coverage
VLP-fraction read coverage
Contig position (bp)
UBA1242 Caccovivens

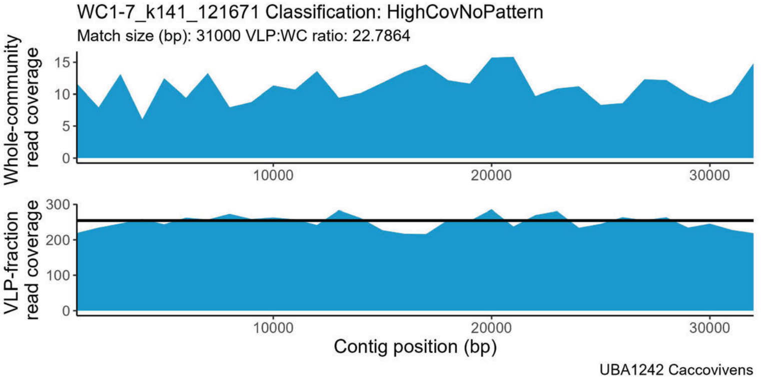
WC1-7_k141_121671 Classification: HighCovNoPattern
Match size (bp): 31000 VLP:WC ratio: 22.7864
Whole-community read coverage
VLP-fraction read coverage
Contig position (bp)
UBA1242 Caccovivens

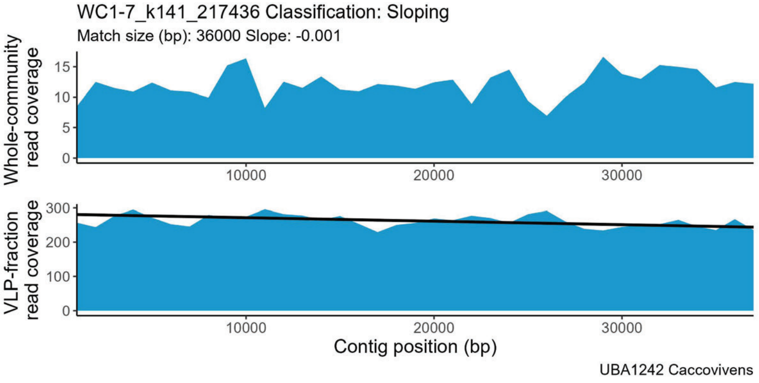
WC1-7_k141_217436 Classification: Sloping
Match size (bp): 36000 Slope: -0.001
Whole-community read coverage
VLP-fraction read coverage
Contig position (bp)
UBA1242 Caccovivens

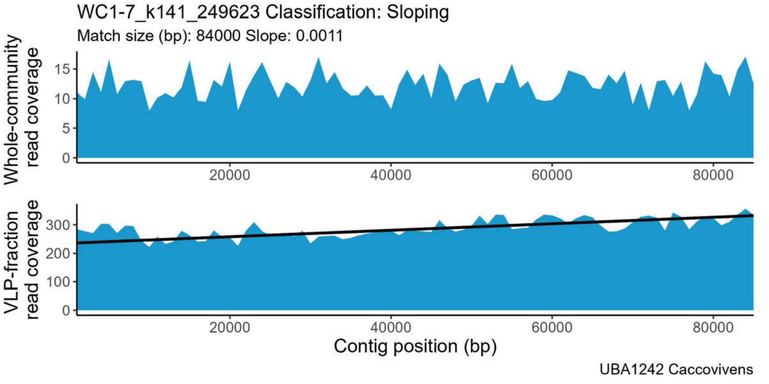
WC1-7_k141_249623 Classification: Sloping
Match size (bp): 84000 Slope: 0.0011
Whole-community read coverage
VLP-fraction read coverage
Contig position (bp)
UBA1242 Caccovivens

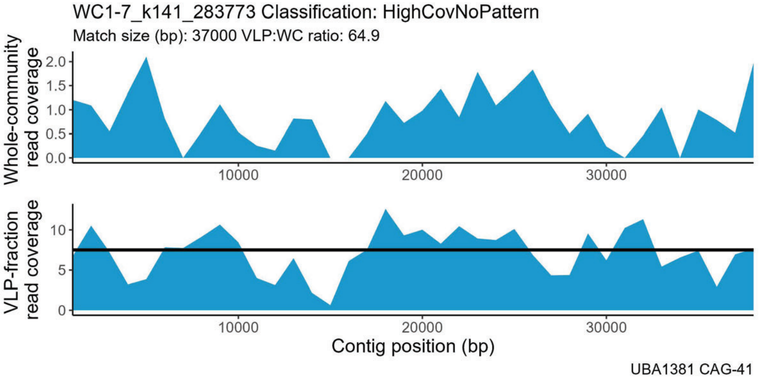
WC1-7_k141_283773 Classification: HighCovNoPattern
Match size (bp): 37000 VLP:WC ratio: 64.9
Whole-community read coverage
VLP-fraction read coverage
Contig position (bp)
UBA1381 CAG-41

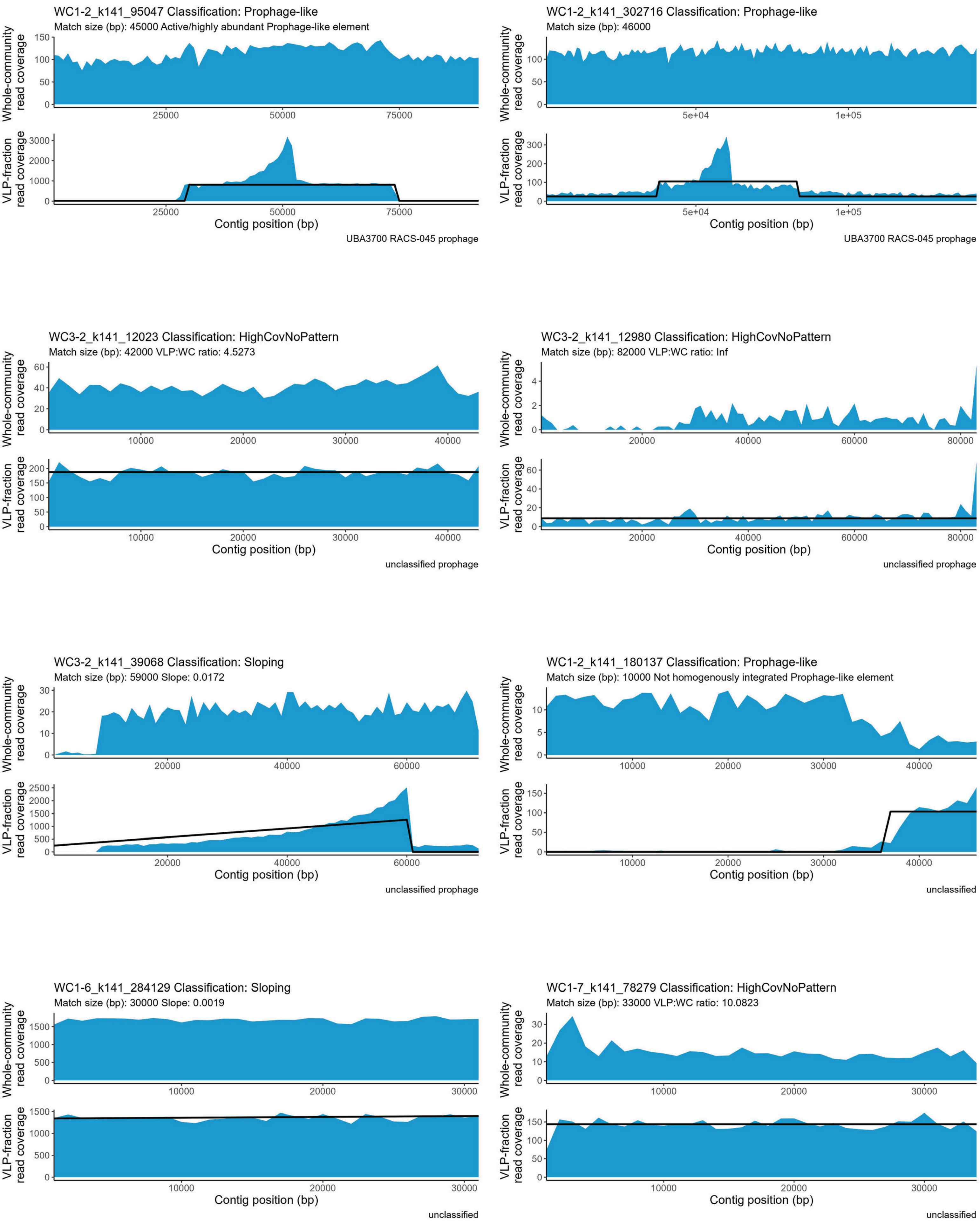
WC1-2_k141_95047 Classification: Prophage-like
Match size (bp): 45000 Active/highly abundant Prophage-like element
Whole-community read coverage
VLP-fraction read coverage
Contig position (bp)
UBA3700 RACS-045 prophage
WC1-2_k141_302716 Classification: Prophage-like
Match size (bp): 46000
Whole-community read coverage
VLP-fraction read coverage
Contig position (bp)
UBA3700 RACS-045 prophage
WC3-2_k141_12023 Classification: HighCovNoPattern
Match size (bp): 42000 VLP:WC ratio: 4.5273
Whole-community read coverage
VLP-fraction read coverage
Contig position (bp)
unclassified prophage
WC3-2_k141_12980 Classification: HighCovNoPattern
Match size (bp): 82000 VLP:WC ratio: Inf
Whole-community read coverage
VLP-fraction read coverage
Contig position (bp)
unclassified prophage
WC3-2_k141_39068 Classification: Sloping
Match size (bp): 59000 Slope: 0.0172
Whole-community read coverage
VLP-fraction read coverage
Contig position (bp)
unclassified prophage
WC1-2_k141_180137 Classification: Prophage-like
Match size (bp): 10000 Not homogenously integrated Prophage-like element
Whole-community read coverage
VLP-fraction read coverage
Contig position (bp)
unclassified
WC1-6_k141_284129 Classification: Sloping
Match size (bp): 30000 Slope: 0.0019
Whole-community read coverage
VLP-fraction read coverage
Contig position (bp)
unclassified
WC1-7_k141_78279 Classification: HighCovNoPattern
Match size (bp): 33000 VLP:WC ratio: 10.0823
Whole-community read coverage
VLP-fraction read coverage
Contig position (bp)
unclassified

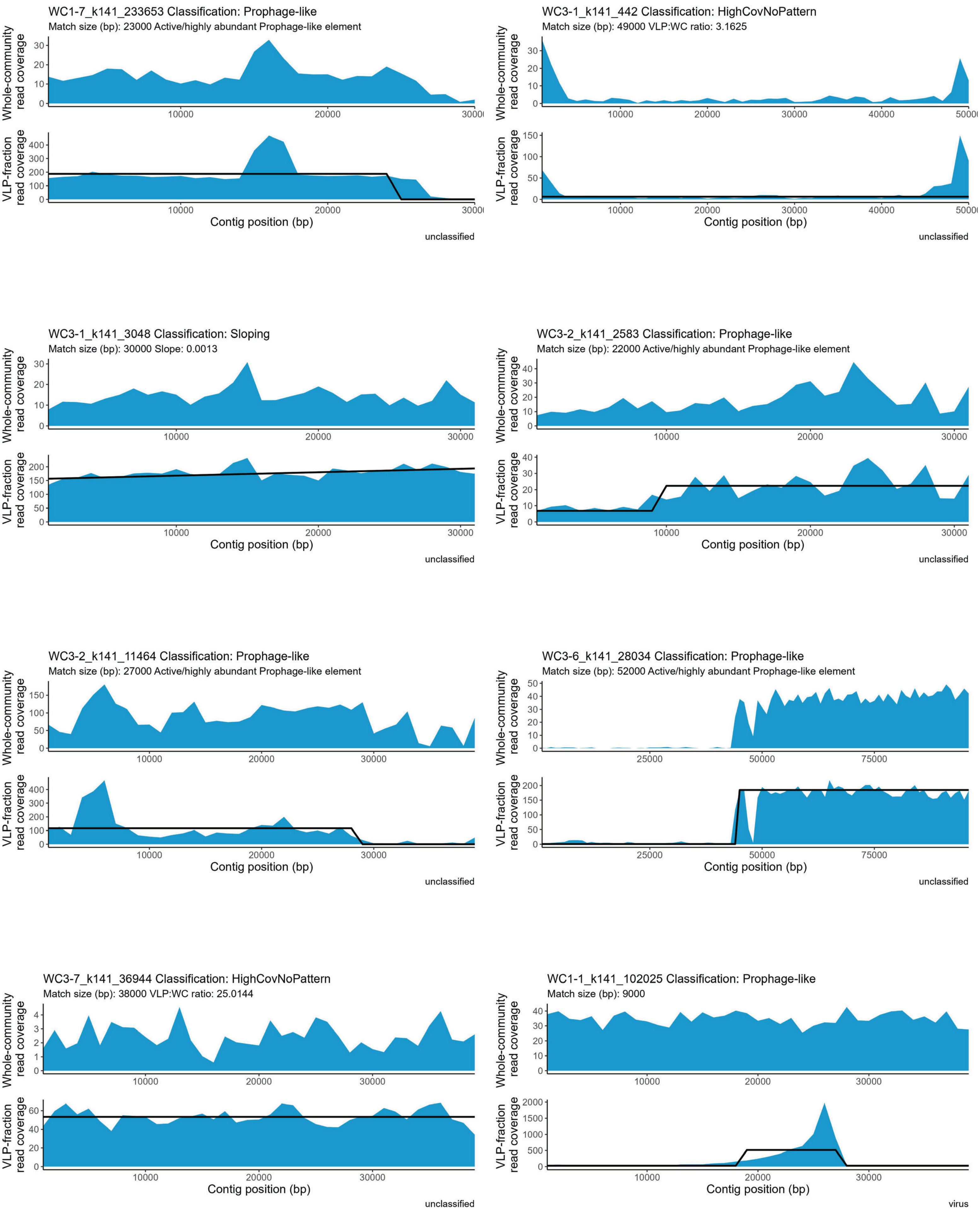
WC1-7_k141_233653 Classification: Prophage-like
Match size (bp): 23000 Active/highly abundant Prophage-like element
Whole-community read coverage
VLP-fraction read coverage
Contig position (bp)
unclassified
WC3-1_k141_442 Classification: HighCovNoPattern
Match size (bp): 49000 VLP:WC ratio: 3.1625
Whole-community read coverage
VLP-fraction read coverage
Contig position (bp)
unclassified
WC3-1_k141_3048 Classification: Sloping
Match size (bp): 30000 Slope: 0.0013
Whole-community read coverage
VLP-fraction read coverage
Contig position (bp)
unclassified
WC3-2_k141_2583 Classification: Prophage-like
Match size (bp): 22000 Active/highly abundant Prophage-like element
Whole-community read coverage
VLP-fraction read coverage
Contig position (bp)
unclassified
WC3-2_k141_11464 Classification: Prophage-like
Match size (bp): 27000 Active/highly abundant Prophage-like element
Whole-community read coverage
VLP-fraction read coverage
Contig position (bp)
unclassified
WC3-6_k141_28034 Classification: Prophage-like
Match size (bp): 52000 Active/highly abundant Prophage-like element
Whole-community read coverage
VLP-fraction read coverage
Contig position (bp)
unclassified
WC3-7_k141_36944 Classification: HighCovNoPattern
Match size (bp): 38000 VLP:WC ratio: 25.0144
Whole-community read coverage
VLP-fraction read coverage
Contig position (bp)
unclassified
WC1-1_k141_102025 Classification: Prophage-like
Match size (bp): 9000
Whole-community read coverage
VLP-fraction read coverage
Contig position (bp)
virus

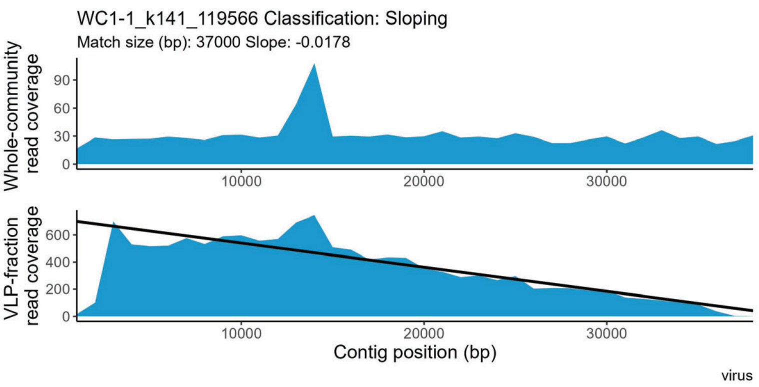
WC1-1_k141_119566 Classification: Sloping
Match size (bp): 37000 Slope: -0.0178
Whole-community read coverage
VLP-fraction read coverage
Contig position (bp)
virus

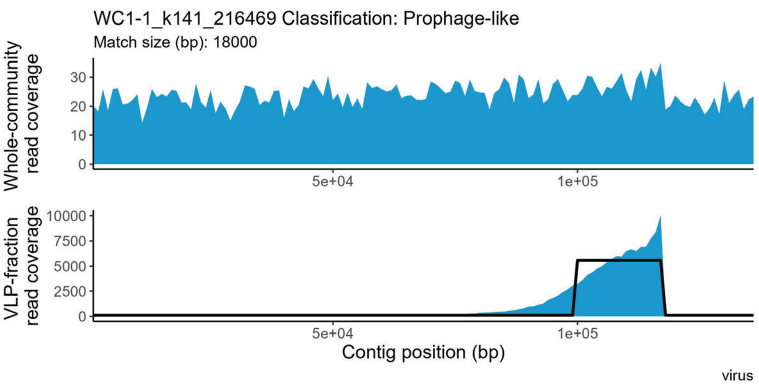
WC1-1_k141_216469 Classification: Prophage-like
Match size (bp): 18000
Whole-community read coverage
VLP-fraction read coverage
Contig position (bp)
virus

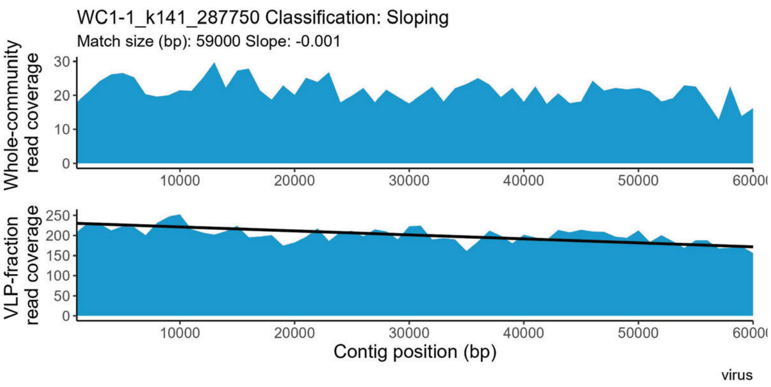
WC1-1_k141_287750 Classification: Sloping
Match size (bp): 59000 Slope: -0.001
Whole-community read coverage
VLP-fraction read coverage
Contig position (bp)
virus

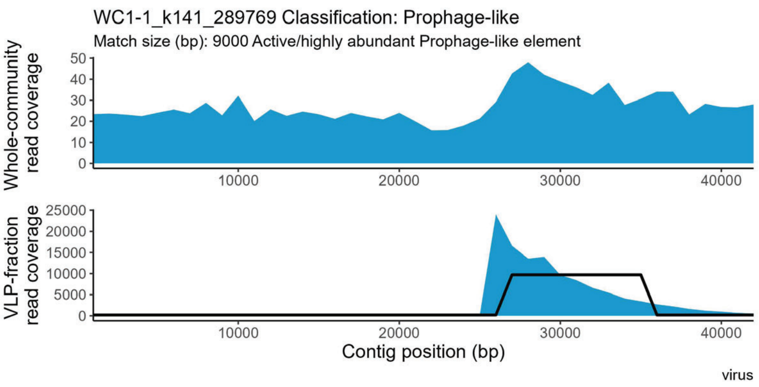
WC1-1_k141_289769 Classification: Prophage-like
Match size (bp): 9000 Active/highly abundant Prophage-like element
Whole-community read coverage
VLP-fraction read coverage
Contig position (bp)
virus

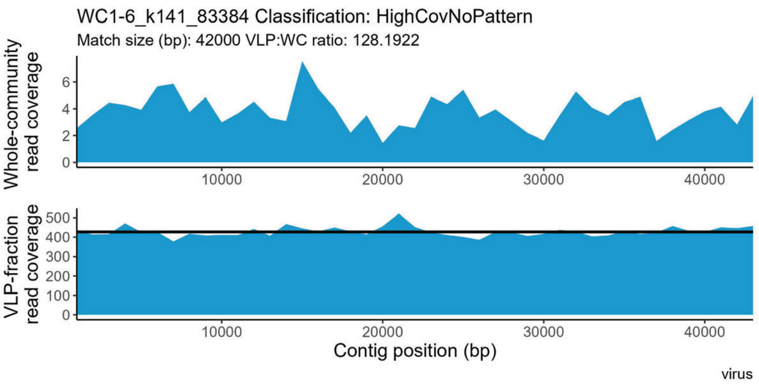
WC1-6_k141_83384 Classification: HighCovNoPattern
Match size (bp): 42000 VLP:WC ratio: 128.1922
Whole-community read coverage
VLP-fraction read coverage
Contig position (bp)
virus

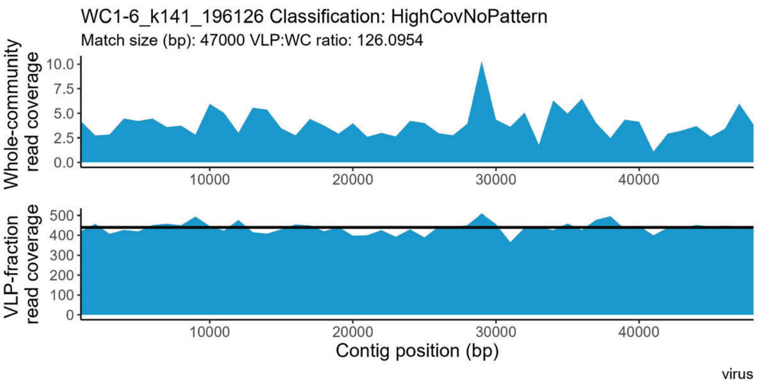
WC1-6_k141_196126 Classification: HighCovNoPattern
Match size (bp): 47000 VLP:WC ratio: 126.0954
Whole-community read coverage
VLP-fraction read coverage
Contig position (bp)
virus

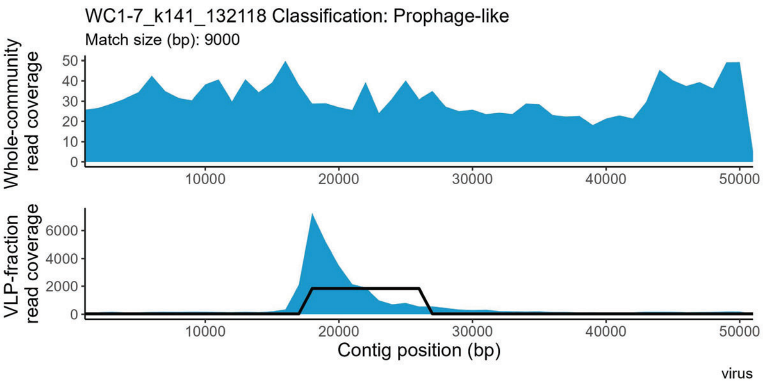
WC1-7_k141_132118 Classification: Prophage-like
Match size (bp): 9000
Whole-community read coverage
VLP-fraction read coverage
Contig position (bp)
virus

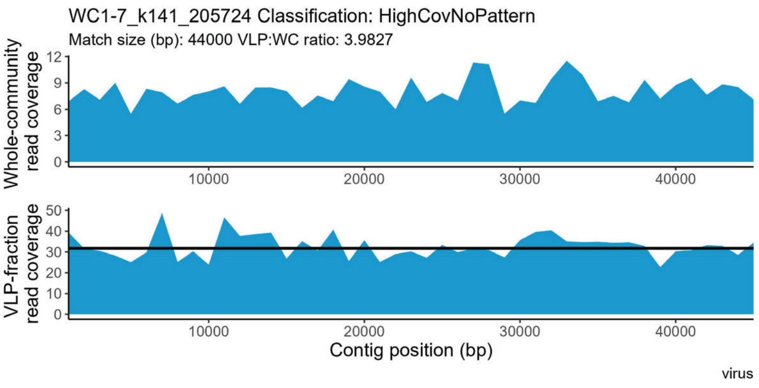
WC1-7_k141_205724 Classification: HighCovNoPattern
Match size (bp): 44000 VLP:WC ratio: 3.9827
Whole-community read coverage
VLP-fraction read coverage
Contig position (bp)
virus

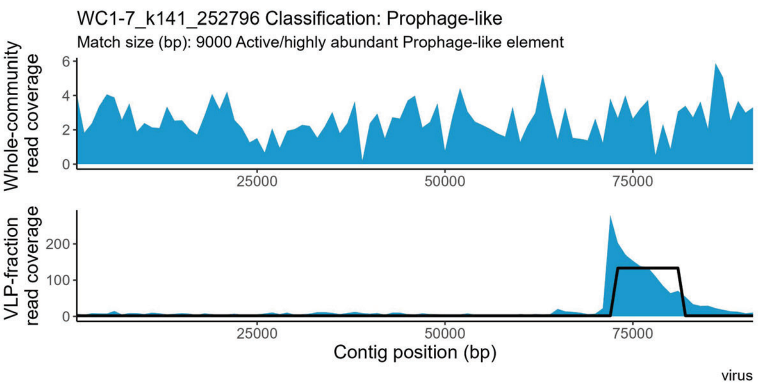
WC1-7_k141_252796 Classification: Prophage-like
Match size (bp): 9000 Active/highly abundant Prophage-like element
Whole-community read coverage
0
2
4
6
25000
50000
75000
VLP-fraction read coverage
0
100
200
25000
50000
75000
Contig position (bp)
virus

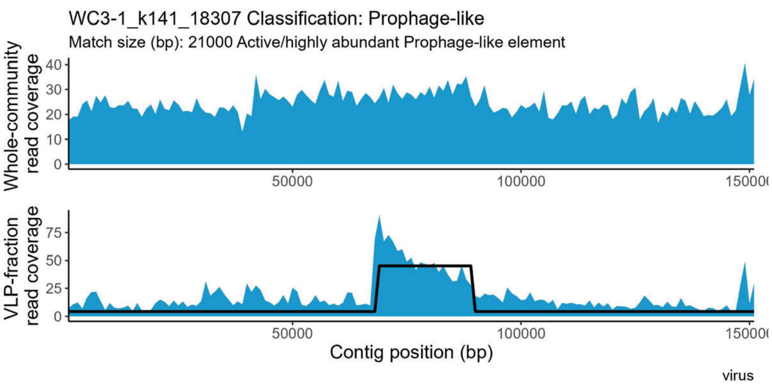
WC3-1_k141_18307 Classification: Prophage-like
Match size (bp): 21000 Active/highly abundant Prophage-like element
Whole-community read coverage
0
10
20
30
40
50000
100000
VLP-fraction read coverage
0
25
50
75
50000
100000
Contig position (bp)
virus
