## Supplemental file 7 for "Perturbations shift the composition of bacterial DNA carried by virus-like particles in the murine gut microbiome"

**File S7) All positive TrIdent pattern-matches for CDI replicate 2.** The sample from which a contig was assembled is indicated by the following prefixes in the contig accession number: WC1-1 = Pre-ABX replicate 1, WC1-2 = Pre-ABX replicate 2, WC1-6 = Pre-ABX replicate 3, WC1-7 = Pre-ABX replicate 4, WC1-1 = CDI replicate 1, WC1-2 = CDI replicate 2, WC1-6 = CDI replicate 3, WC1-7 = CDI replicate 4. Pre-ABX = Pre-antibiotics, Post-ABX= post-antbiotics, CDI= *Clostridioides difficile* infection. Contig taxonomy and geNomad prophage classification in plot subtitle. Plots are listed alphabetically by family name.

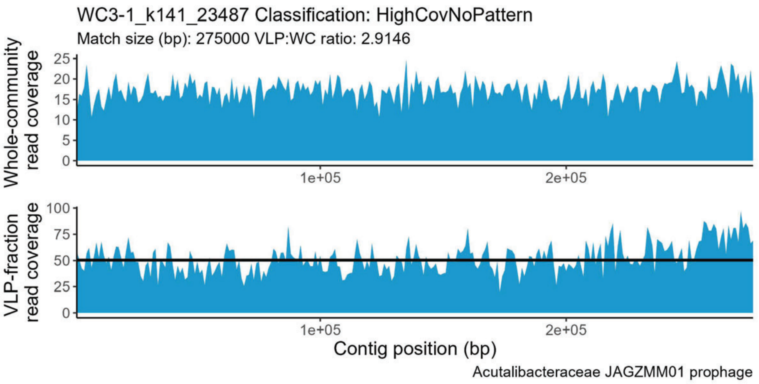


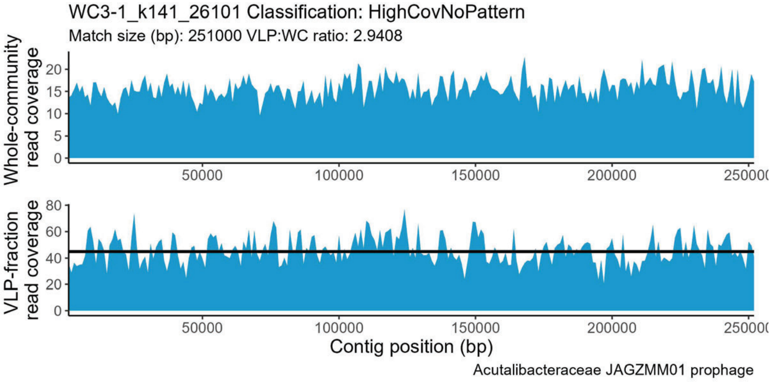


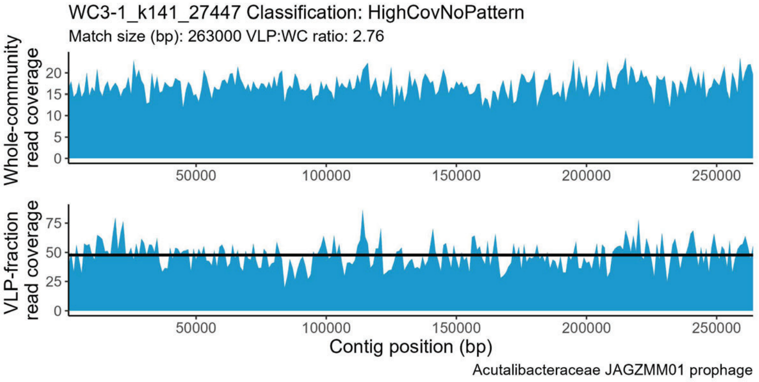


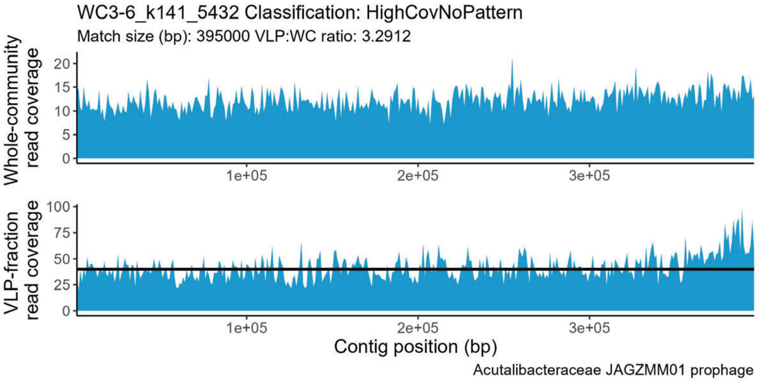


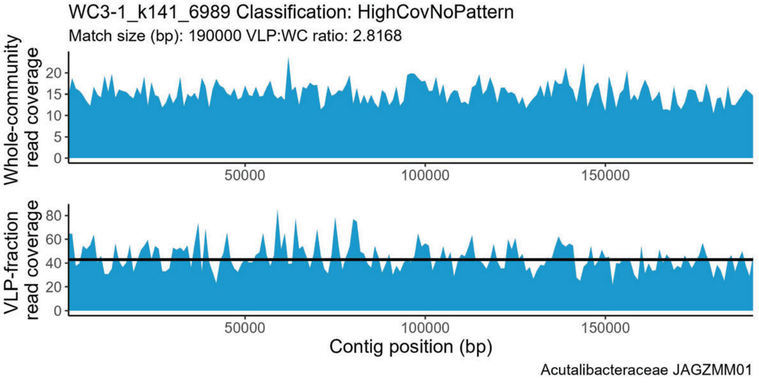


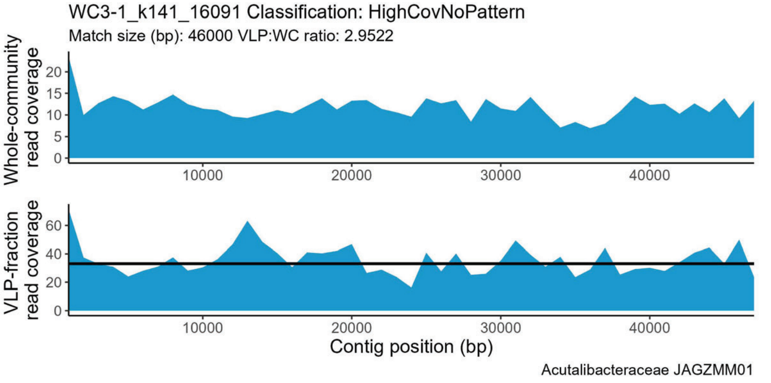


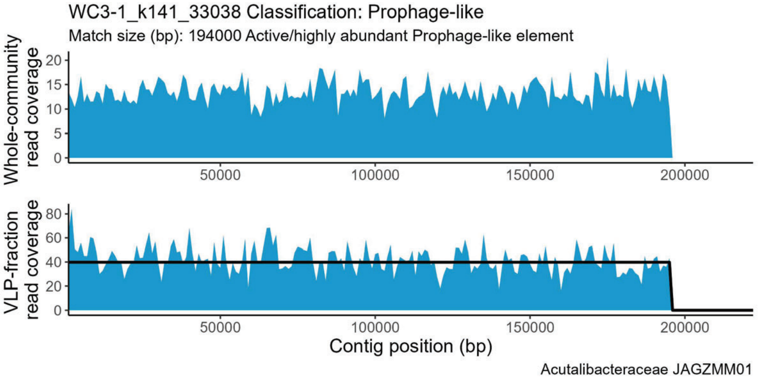


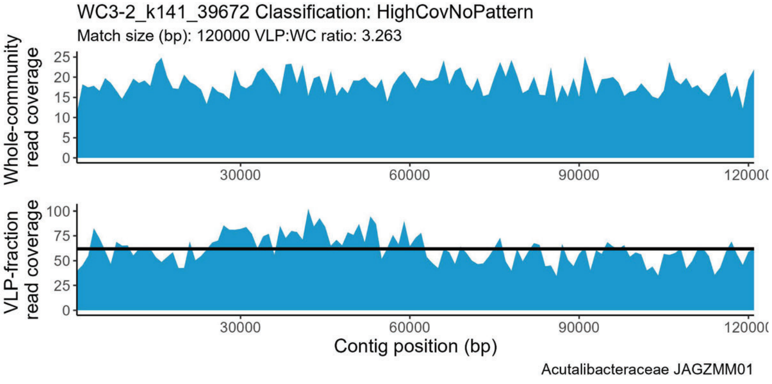

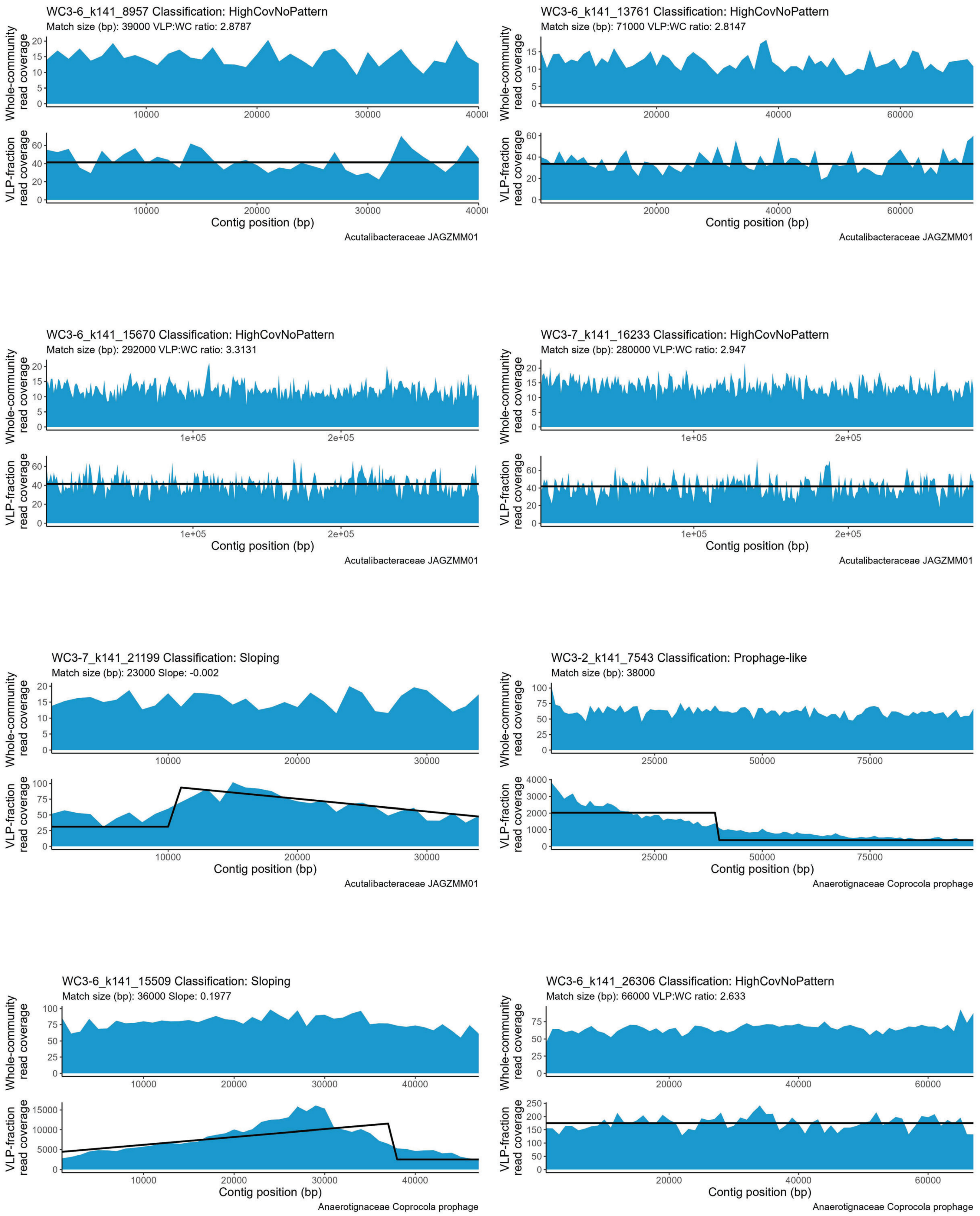

WC3-6_k141_8957 Classification: HighCovNoPattern
Match size (bp): 39000 VLP:WC ratio: 2.8787
Acutalibacteraceae JAGZMM01
WC3-6_k141_13761 Classification: HighCovNoPattern
Match size (bp): 71000 VLP:WC ratio: 2.8147
Acutalibacteraceae JAGZMM01
WC3-6_k141_15670 Classification: HighCovNoPattern
Match size (bp): 292000 VLP:WC ratio: 3.3131
Acutalibacteraceae JAGZMM01
WC3-7_k141_16233 Classification: HighCovNoPattern
Match size (bp): 280000 VLP:WC ratio: 2.947
Acutalibacteraceae JAGZMM01
WC3-7_k141_21199 Classification: Sloping
Match size (bp): 23000 Slope: -0.002
Acutalibacteraceae JAGZMM01
WC3-2_k141_7543 Classification: Prophage-like
Match size (bp): 38000
Anaerotignaceae Coprocola prophage
WC3-6_k141_15509 Classification: Sloping
Match size (bp): 36000 Slope: 0.1977
Anaerotignaceae Coprocola prophage
WC3-6_k141_26306 Classification: HighCovNoPattern
Match size (bp): 66000 VLP:WC ratio: 2.633
Anaerotignaceae Coprocola prophage
Whole-community read coverage
VLP-fraction read coverage
Contig position (bp)

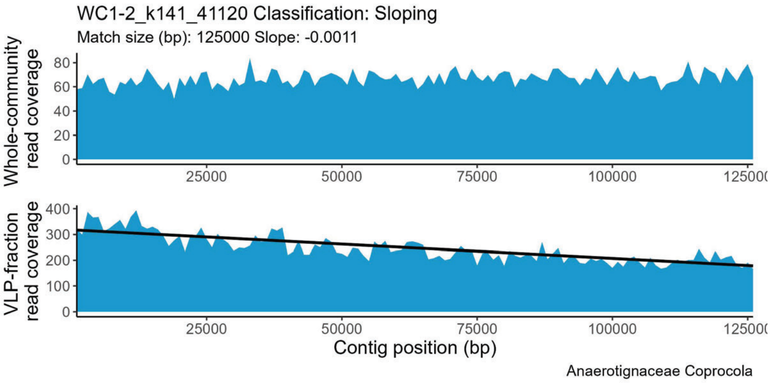
WC1-2_k141_41120 Classification: Sloping
Match size (bp): 125000 Slope: -0.0011
Whole-community read coverage
VLP-fraction read coverage
Contig position (bp)
Anaerotignaceae Coprocola

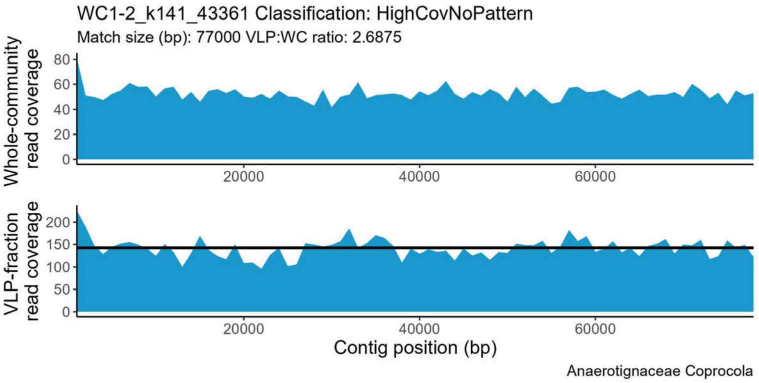
WC1-2_k141_43361 Classification: HighCovNoPattern
Match size (bp): 77000 VLP:WC ratio: 2.6875
Whole-community read coverage
VLP-fraction read coverage
Contig position (bp)
Anaerotignaceae Coprocola

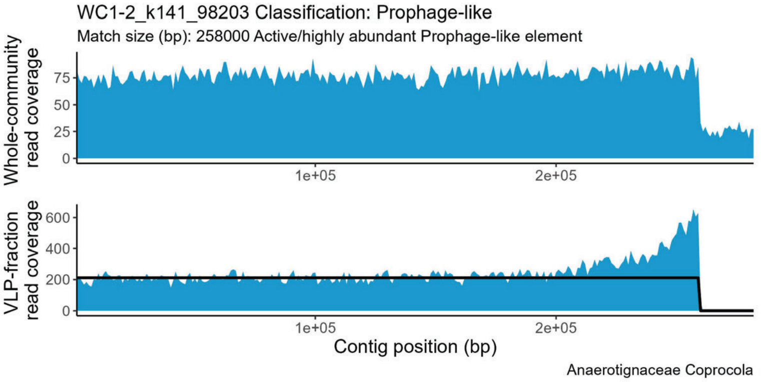
WC1-2_k141_98203 Classification: Prophage-like
Match size (bp): 258000 Active/highly abundant Prophage-like element
Whole-community read coverage
VLP-fraction read coverage
Contig position (bp)
Anaerotignaceae Coprocola

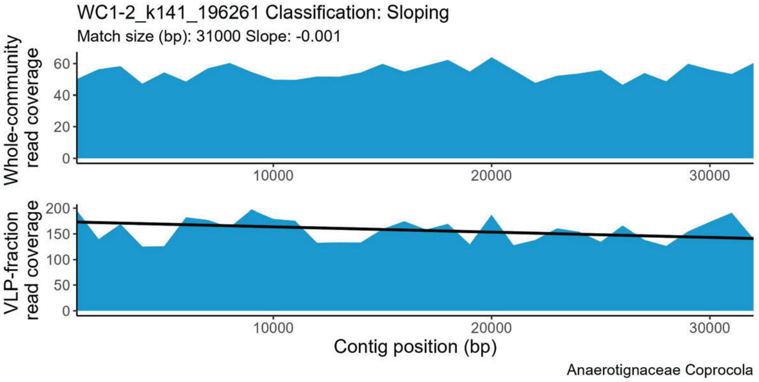
WC1-2_k141_196261 Classification: Sloping
Match size (bp): 31000 Slope: -0.001
Whole-community read coverage
VLP-fraction read coverage
Contig position (bp)
Anaerotignaceae Coprocola

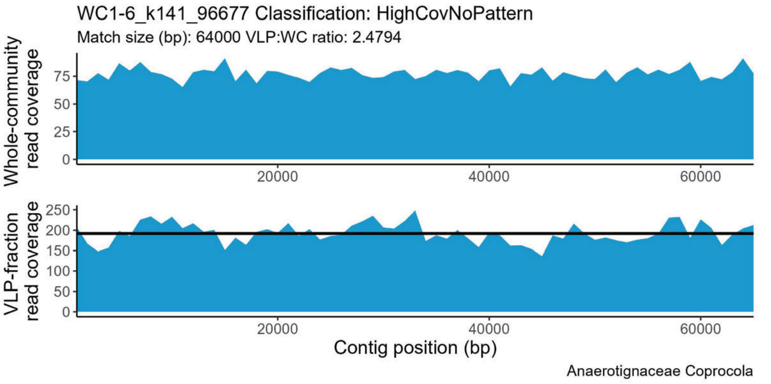
WC1-6_k141_96677 Classification: HighCovNoPattern
Match size (bp): 64000 VLP:WC ratio: 2.4794
Whole-community read coverage
VLP-fraction read coverage
Contig position (bp)
Anaerotignaceae Coprocola

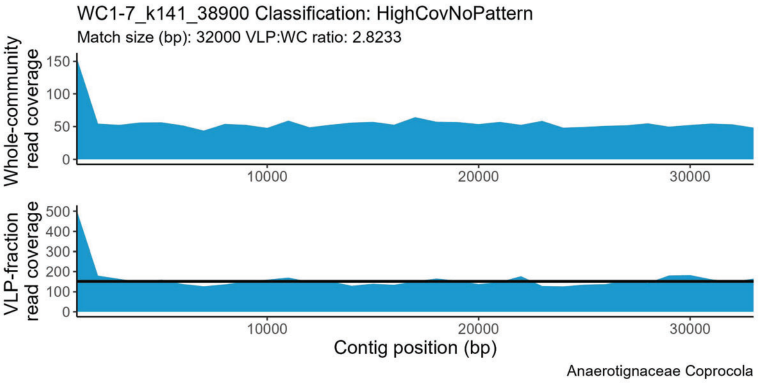
WC1-7_k141_38900 Classification: HighCovNoPattern
Match size (bp): 32000 VLP:WC ratio: 2.8233
Whole-community read coverage
VLP-fraction read coverage
Contig position (bp)
Anaerotignaceae Coprocola

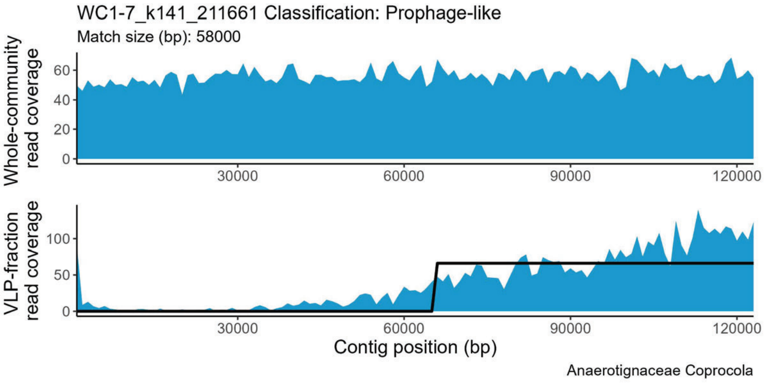
WC1-7_k141_211661 Classification: Prophage-like
Match size (bp): 58000
Whole-community read coverage
VLP-fraction read coverage
Contig position (bp)
Anaerotignaceae Coprocola

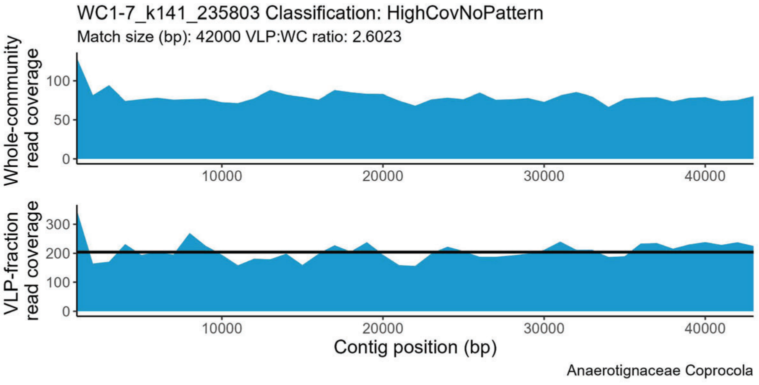
WC1-7_k141_235803 Classification: HighCovNoPattern
Match size (bp): 42000 VLP:WC ratio: 2.6023
Whole-community read coverage
VLP-fraction read coverage
Contig position (bp)
Anaerotignaceae Coprocola

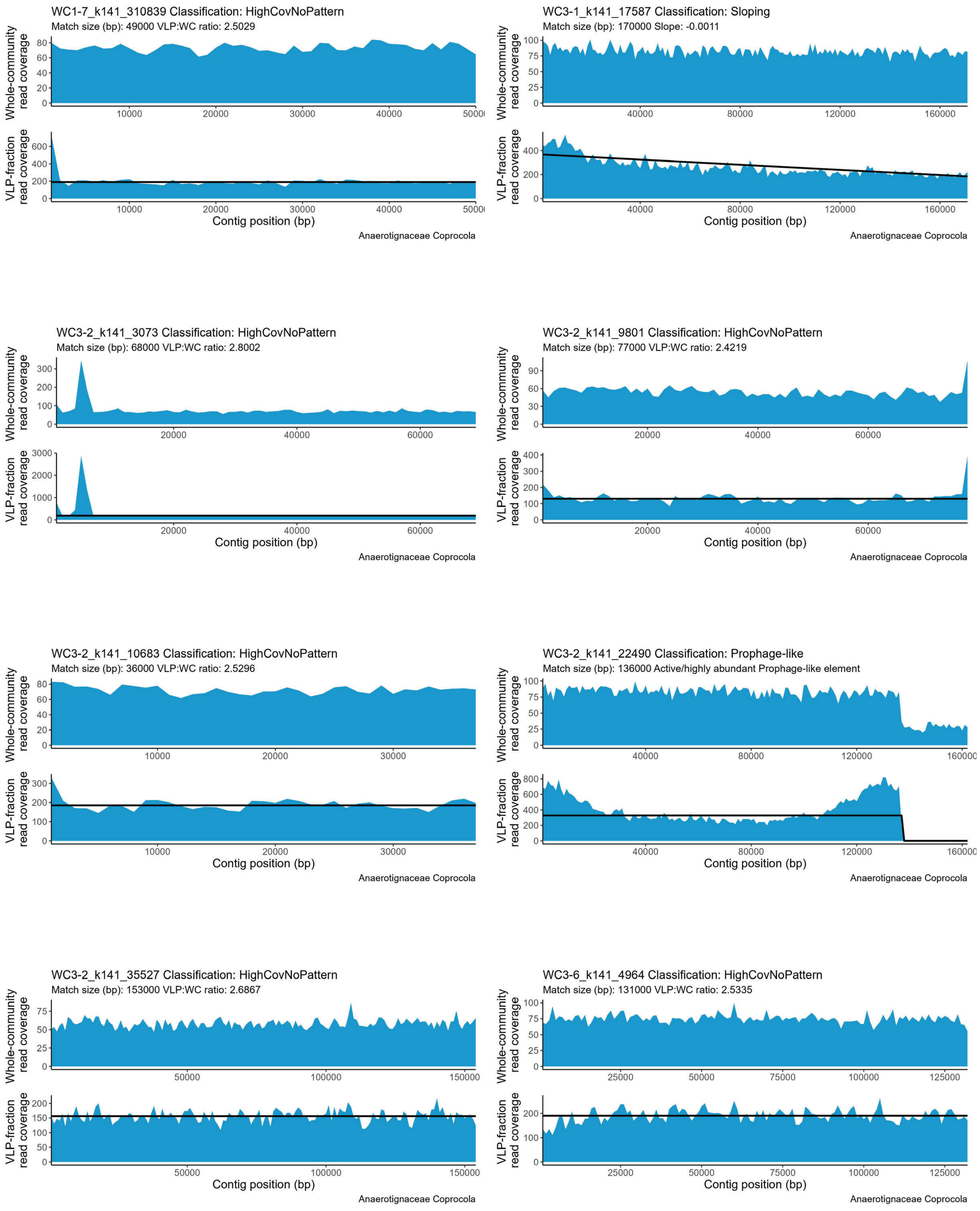
WC1-7_k141_310839 Classification: HighCovNoPattern
Match size (bp): 49000 VLP:WC ratio: 2.5029
Whole-community read coverage
VLP-fraction read coverage
Contig position (bp)
Anaerotignaceae Coprocola
WC3-1_k141_17587 Classification: Sloping
Match size (bp): 170000 Slope: -0.0011
Whole-community read coverage
VLP-fraction read coverage
Contig position (bp)
Anaerotignaceae Coprocola
WC3-2_k141_3073 Classification: HighCovNoPattern
Match size (bp): 68000 VLP:WC ratio: 2.8002
Whole-community read coverage
VLP-fraction read coverage
Contig position (bp)
Anaerotignaceae Coprocola
WC3-2_k141_9801 Classification: HighCovNoPattern
Match size (bp): 77000 VLP:WC ratio: 2.4219
Whole-community read coverage
VLP-fraction read coverage
Contig position (bp)
Anaerotignaceae Coprocola
WC3-2_k141_10683 Classification: HighCovNoPattern
Match size (bp): 36000 VLP:WC ratio: 2.5296
Whole-community read coverage
VLP-fraction read coverage
Contig position (bp)
Anaerotignaceae Coprocola
WC3-2_k141_22490 Classification: Prophage-like
Match size (bp): 136000 Active/highly abundant Prophage-like element
Whole-community read coverage
VLP-fraction read coverage
Contig position (bp)
Anaerotignaceae Coprocola
WC3-2_k141_35527 Classification: HighCovNoPattern
Match size (bp): 153000 VLP:WC ratio: 2.6867
Whole-community read coverage
VLP-fraction read coverage
Contig position (bp)
Anaerotignaceae Coprocola
WC3-6_k141_4964 Classification: HighCovNoPattern
Match size (bp): 131000 VLP:WC ratio: 2.5335
Whole-community read coverage
VLP-fraction read coverage
Contig position (bp)
Anaerotignaceae Coprocola

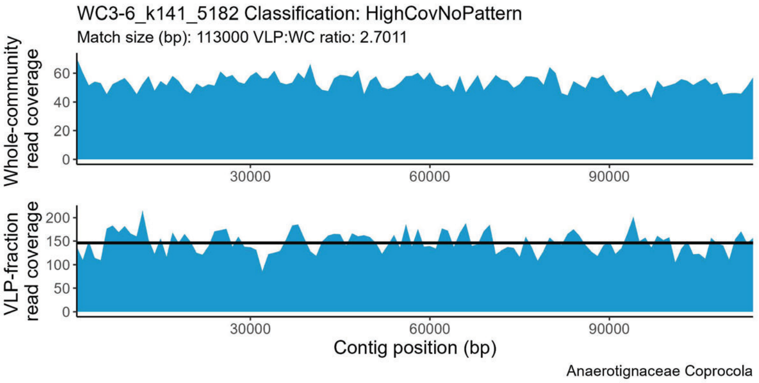
WC3-6_k141_5182 Classification: HighCovNoPattern
Match size (bp): 113000 VLP:WC ratio: 2.7011
Whole-community read coverage
0
20
40
60
30000
60000
90000
VLP-fraction read coverage
0
50
100
150
200
Contig position (bp)
Anaerotignaceae Coprocola

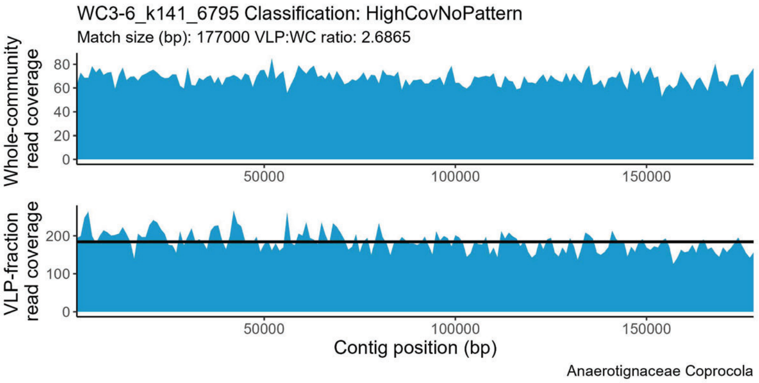
WC3-6_k141_6795 Classification: HighCovNoPattern
Match size (bp): 177000 VLP:WC ratio: 2.6865
Whole-community read coverage
0
20
40
60
80
50000
100000
150000
VLP-fraction read coverage
0
100
200
Contig position (bp)
Anaerotignaceae Coprocola

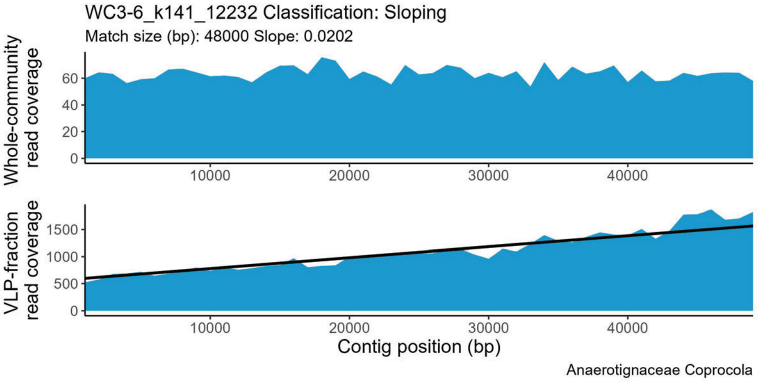
WC3-6_k141_12232 Classification: Sloping
Match size (bp): 48000 Slope: 0.0202
Whole-community read coverage
0
20
40
60
10000
20000
30000
40000
VLP-fraction read coverage
0
500
1000
1500
Contig position (bp)
Anaerotignaceae Coprocola

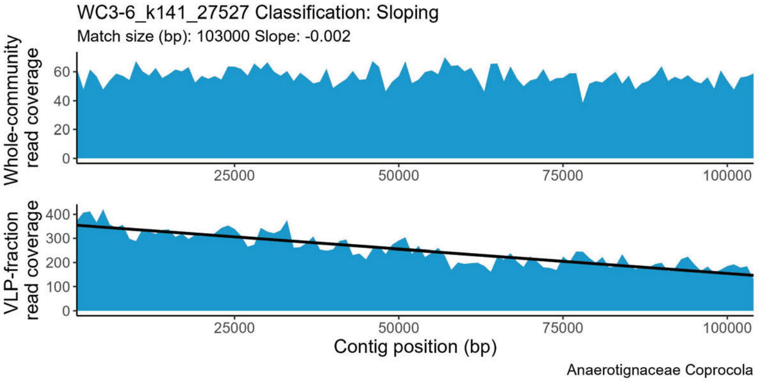
WC3-6_k141_27527 Classification: Sloping
Match size (bp): 103000 Slope: -0.002
Whole-community read coverage
0
20
40
60
25000
50000
75000
100000
VLP-fraction read coverage
0
100
200
300
400
Contig position (bp)
Anaerotignaceae Coprocola

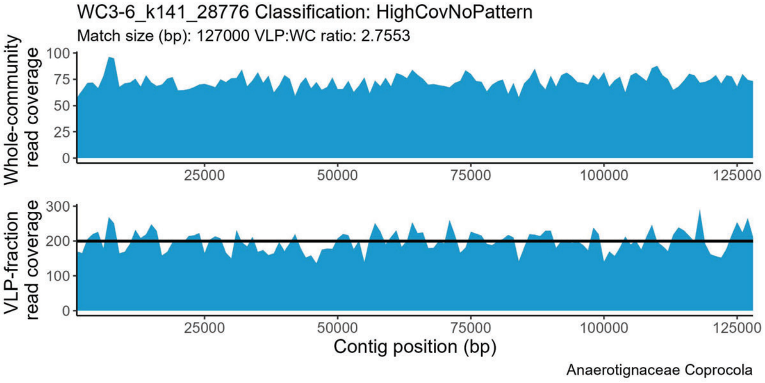
WC3-6_k141_28776 Classification: HighCovNoPattern
Match size (bp): 127000 VLP:WC ratio: 2.7553
Whole-community read coverage
0
25
50
75
100
25000
50000
75000
100000
125000
VLP-fraction read coverage
0
100
200
300
Contig position (bp)
Anaerotignaceae Coprocola

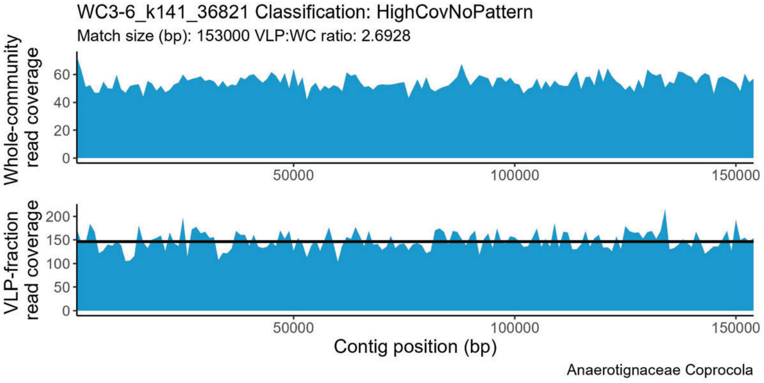
WC3-6_k141_36821 Classification: HighCovNoPattern
Match size (bp): 153000 VLP:WC ratio: 2.6928
Whole-community read coverage
0
20
40
60
50000
100000
150000
VLP-fraction read coverage
0
50
100
150
200
Contig position (bp)
Anaerotignaceae Coprocola

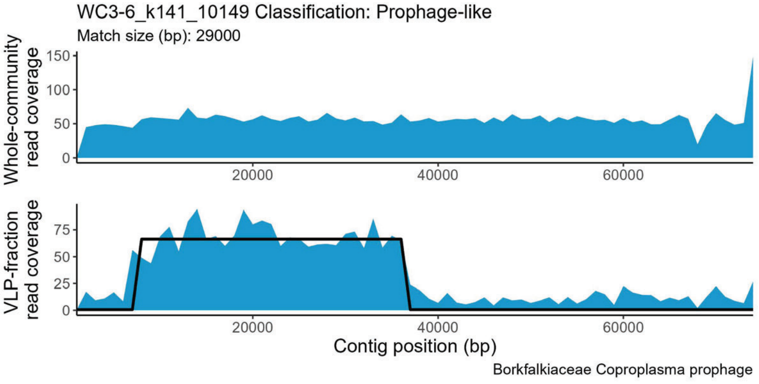
WC3-6_k141_10149 Classification: Prophage-like
Match size (bp): 29000
Whole-community read coverage
0
50
100
150
20000
40000
60000
VLP-fraction read coverage
0
25
50
75
Contig position (bp)
Borkfalkiaceae Coproplasma prophage

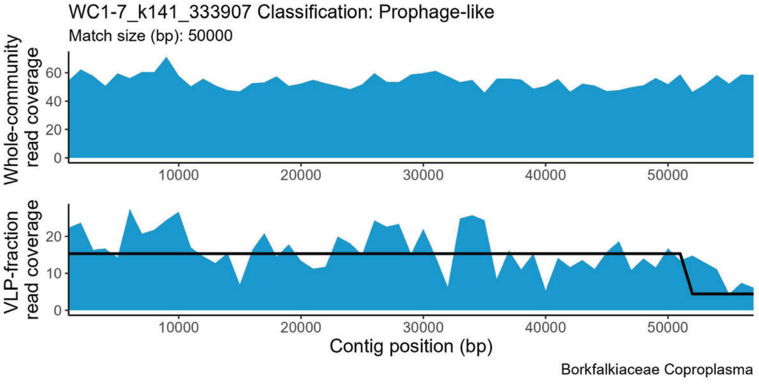
WC1-7_k141_333907 Classification: Prophage-like
Match size (bp): 50000
Whole-community read coverage
0
20
40
60
10000
20000
30000
40000
50000
VLP-fraction read coverage
0
10
20
Contig position (bp)
Borkfalkiaceae Coproplasma

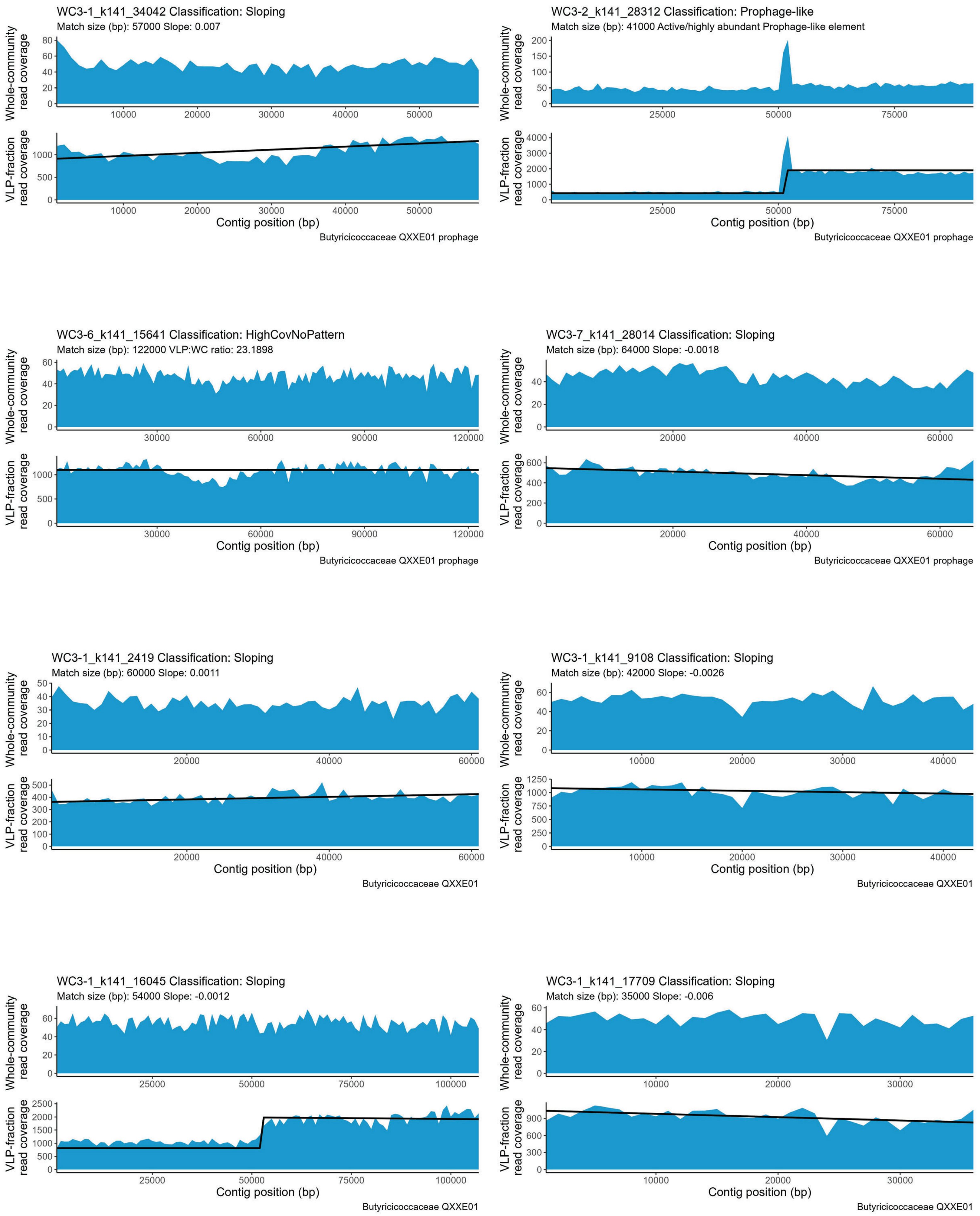

WC3-1_k141_34042 Classification: Sloping
Match size (bp): 57000 Slope: 0.007
Whole-community read coverage
VLP-fraction read coverage
Contig position (bp)
Butyricicoccaceae QXXE01 prophage
WC3-2_k141_28312 Classification: Prophage-like
Match size (bp): 41000 Active/highly abundant Prophage-like element
Whole-community read coverage
VLP-fraction read coverage
Contig position (bp)
Butyricicoccaceae QXXE01 prophage
WC3-6_k141_15641 Classification: HighCovNoPattern
Match size (bp): 122000 VLP:WC ratio: 23.1898
Whole-community read coverage
VLP-fraction read coverage
Contig position (bp)
Butyricicoccaceae QXXE01 prophage
WC3-7_k141_28014 Classification: Sloping
Match size (bp): 64000 Slope: -0.0018
Whole-community read coverage
VLP-fraction read coverage
Contig position (bp)
Butyricicoccaceae QXXE01 prophage
WC3-1_k141_2419 Classification: Sloping
Match size (bp): 60000 Slope: 0.0011
Whole-community read coverage
VLP-fraction read coverage
Contig position (bp)
Butyricicoccaceae QXXE01
WC3-1_k141_9108 Classification: Sloping
Match size (bp): 42000 Slope: -0.0026
Whole-community read coverage
VLP-fraction read coverage
Contig position (bp)
Butyricicoccaceae QXXE01
WC3-1_k141_16045 Classification: Sloping
Match size (bp): 54000 Slope: -0.0012
Whole-community read coverage
VLP-fraction read coverage
Contig position (bp)
Butyricicoccaceae QXXE01
WC3-1_k141_17709 Classification: Sloping
Match size (bp): 35000 Slope: -0.006
Whole-community read coverage
VLP-fraction read coverage
Contig position (bp)
Butyricicoccaceae QXXE01

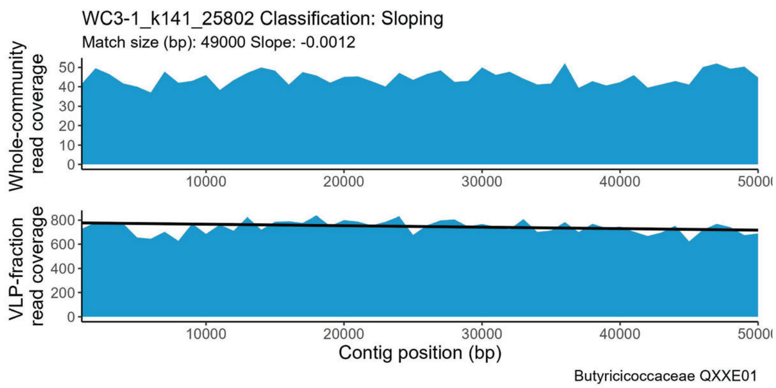
WC3-1_k141_25802 Classification: Sloping
Match size (bp): 49000 Slope: -0.0012
Whole-community read coverage
VLP-fraction read coverage
Contig position (bp)
Butyricicoccaceae QXXE01

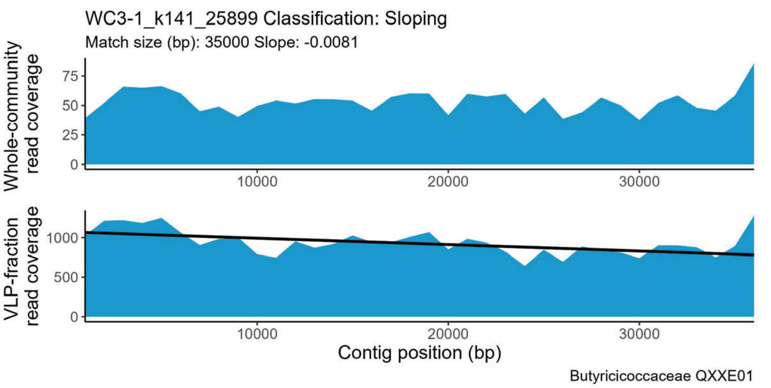
WC3-1_k141_25899 Classification: Sloping
Match size (bp): 35000 Slope: -0.0081
Whole-community read coverage
VLP-fraction read coverage
Contig position (bp)
Butyricicoccaceae QXXE01

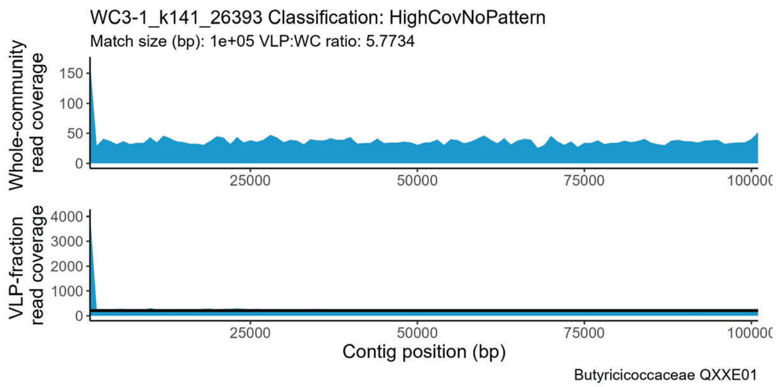
WC3-1_k141_26393 Classification: HighCovNoPattern
Match size (bp): 1e+05 VLP:WC ratio: 5.7734
Whole-community read coverage
VLP-fraction read coverage
Contig position (bp)
Butyricicoccaceae QXXE01

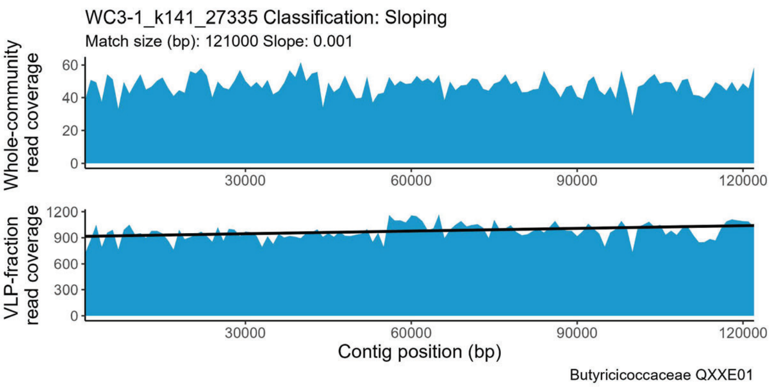
WC3-1_k141_27335 Classification: Sloping
Match size (bp): 121000 Slope: 0.001
Whole-community read coverage
VLP-fraction read coverage
Contig position (bp)
Butyricicoccaceae QXXE01

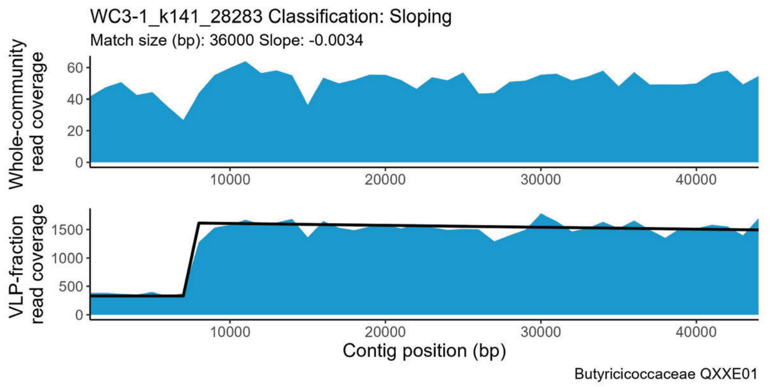
WC3-1_k141_28283 Classification: Sloping
Match size (bp): 36000 Slope: -0.0034
Whole-community read coverage
VLP-fraction read coverage
Contig position (bp)
Butyricicoccaceae QXXE01

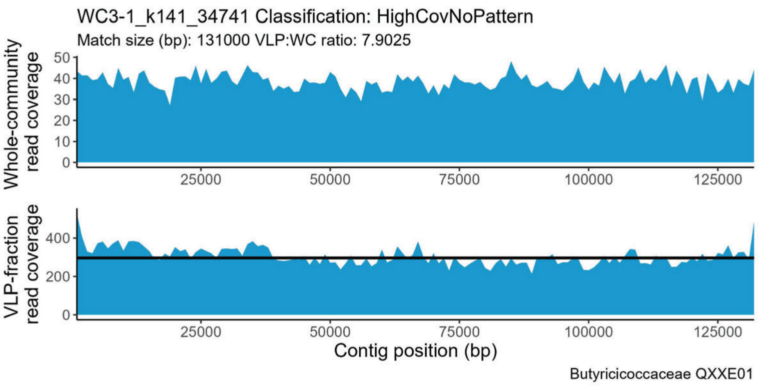
WC3-1_k141_34741 Classification: HighCovNoPattern
Match size (bp): 131000 VLP:WC ratio: 7.9025
Whole-community read coverage
VLP-fraction read coverage
Contig position (bp)
Butyricicoccaceae QXXE01

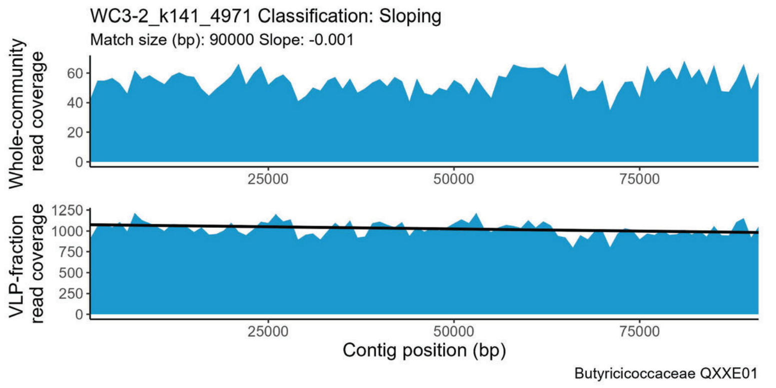
WC3-2_k141_4971 Classification: Sloping
Match size (bp): 90000 Slope: -0.001
Whole-community read coverage
VLP-fraction read coverage
Contig position (bp)
Butyricicoccaceae QXXE01

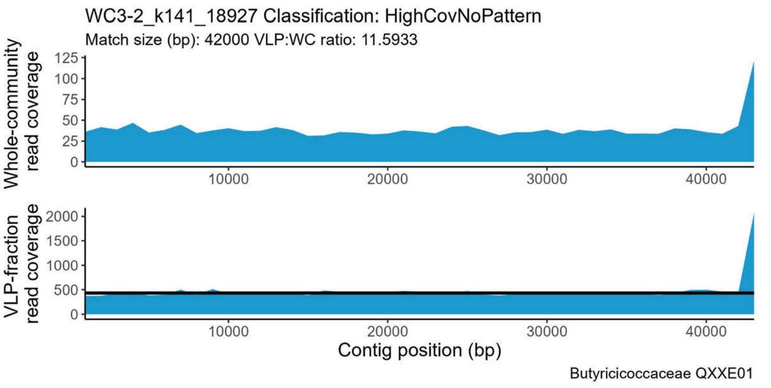
WC3-2_k141_18927 Classification: HighCovNoPattern
Match size (bp): 42000 VLP:WC ratio: 11.5933
Whole-community read coverage
VLP-fraction read coverage
Contig position (bp)
Butyricicoccaceae QXXE01

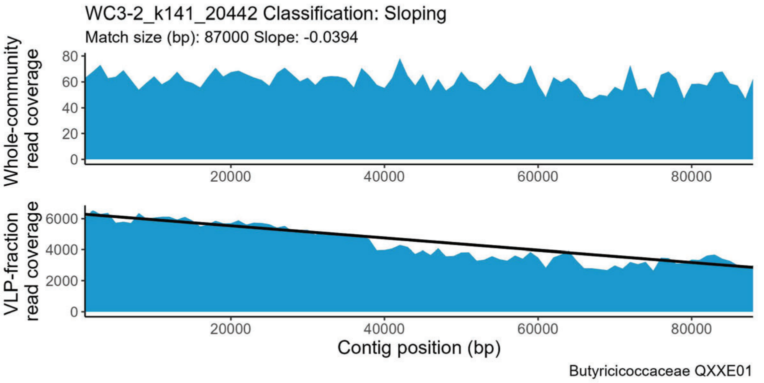
WC3-2_k141_20442 Classification: Sloping
Match size (bp): 87000 Slope: -0.0394
Whole-community read coverage
VLP-fraction read coverage
Contig position (bp)
Butyricicoccaceae QXXE01

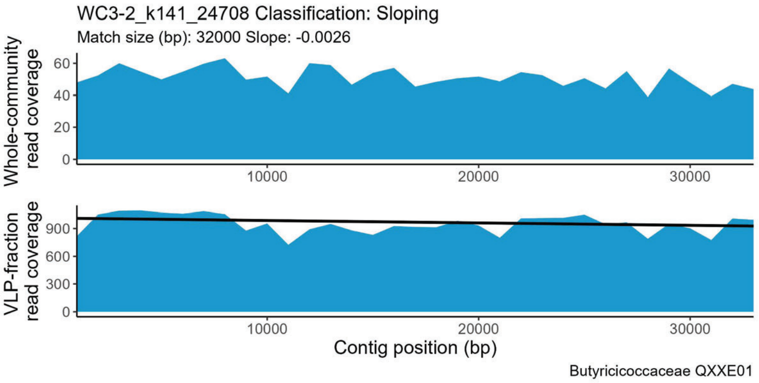
WC3-2_k141_24708 Classification: Sloping
Match size (bp): 32000 Slope: -0.0026
Whole-community read coverage
VLP-fraction read coverage
Contig position (bp)
Butyricicoccaceae QXXE01

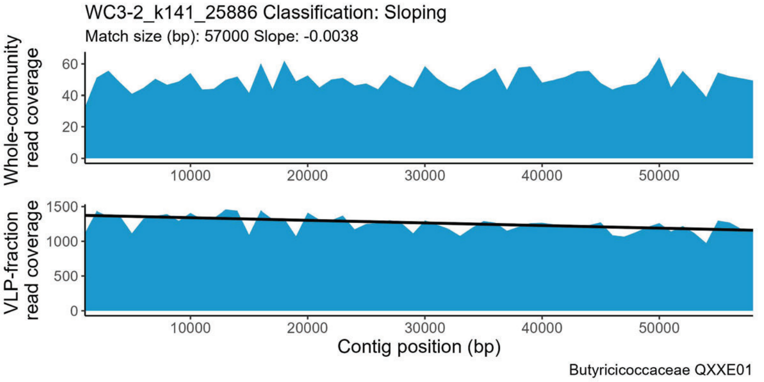
WC3-2_k141_25886 Classification: Sloping
Match size (bp): 57000 Slope: -0.0038
Whole-community read coverage
VLP-fraction read coverage
Contig position (bp)
Butyricicoccaceae QXXE01

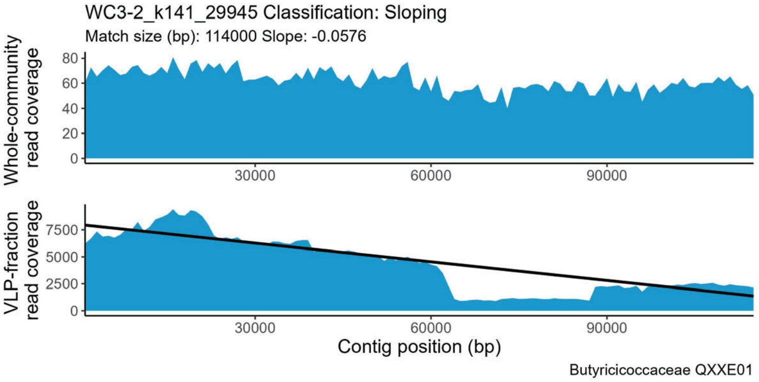
WC3-2_k141_29945 Classification: Sloping
Match size (bp): 114000 Slope: -0.0576
Whole-community read coverage
VLP-fraction read coverage
Contig position (bp)
Butyricicoccaceae QXXE01

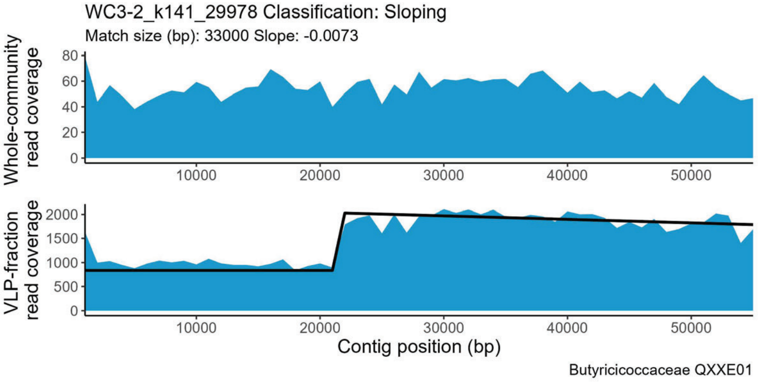
WC3-2_k141_29978 Classification: Sloping
Match size (bp): 33000 Slope: -0.0073
Whole-community read coverage
VLP-fraction read coverage
Contig position (bp)
Butyricicoccaceae QXXE01

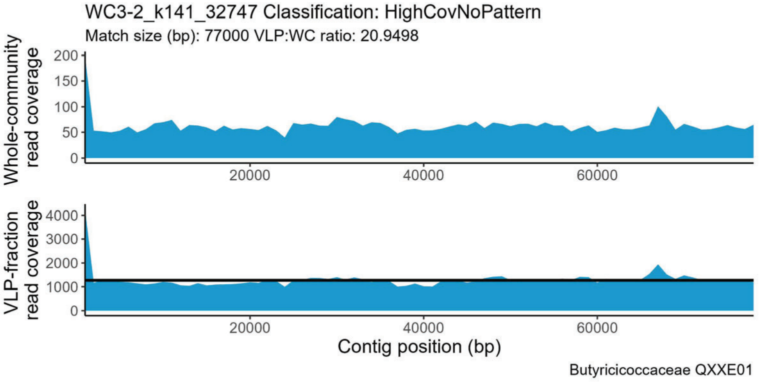
WC3-2_k141_32747 Classification: HighCovNoPattern
Match size (bp): 77000 VLP:WC ratio: 20.9498
Whole-community read coverage
VLP-fraction read coverage
Contig position (bp)
Butyricicoccaceae QXXE01

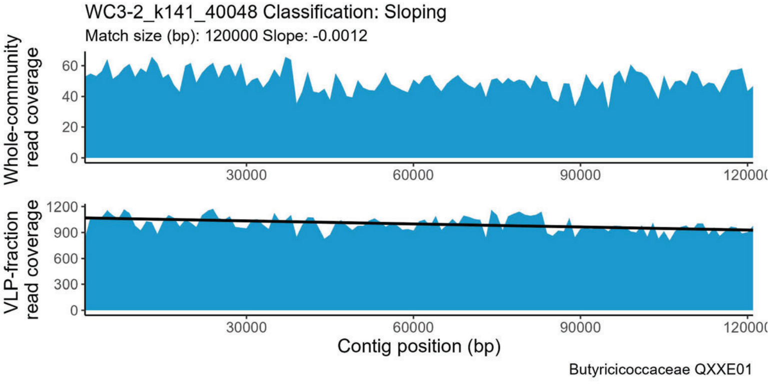
WC3-2_k141_40048 Classification: Sloping
Match size (bp): 120000 Slope: -0.0012
Whole-community read coverage
VLP-fraction read coverage
Contig position (bp)
Butyricicoccaceae QXXE01

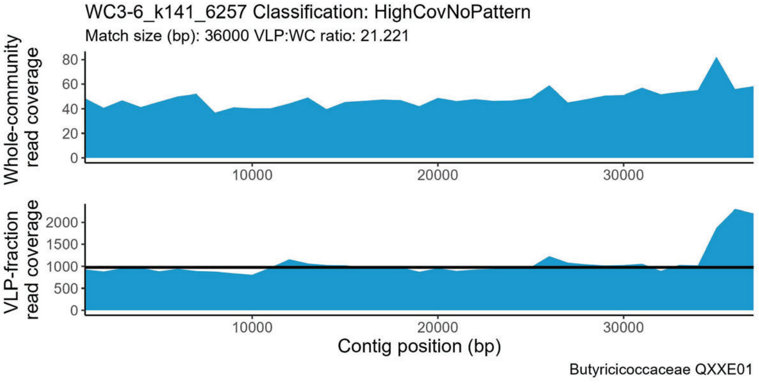
WC3-6_k141_6257 Classification: HighCovNoPattern
Match size (bp): 36000 VLP:WC ratio: 21.221
Whole-community read coverage
VLP-fraction read coverage
Contig position (bp)
Butyricicoccaceae QXXE01

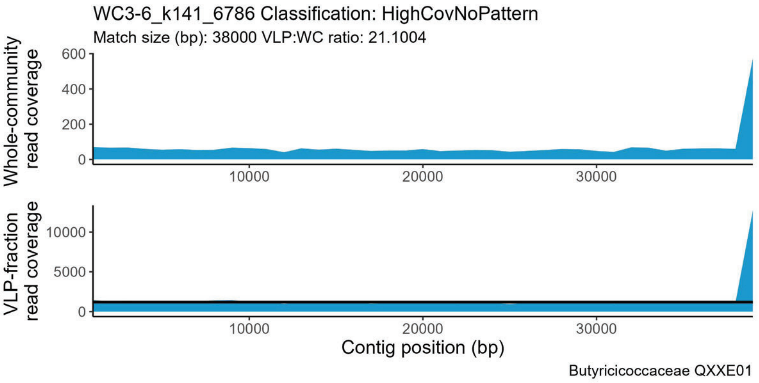
WC3-6_k141_6786 Classification: HighCovNoPattern
Match size (bp): 38000 VLP:WC ratio: 21.1004
Whole-community read coverage
VLP-fraction read coverage
Contig position (bp)
Butyricicoccaceae QXXE01

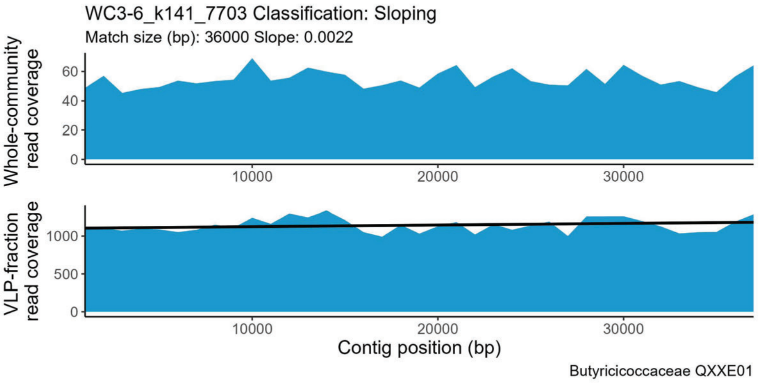
WC3-6_k141_7703 Classification: Sloping
Match size (bp): 36000 Slope: 0.0022
Whole-community read coverage
VLP-fraction read coverage
Contig position (bp)
Butyricicoccaceae QXXE01

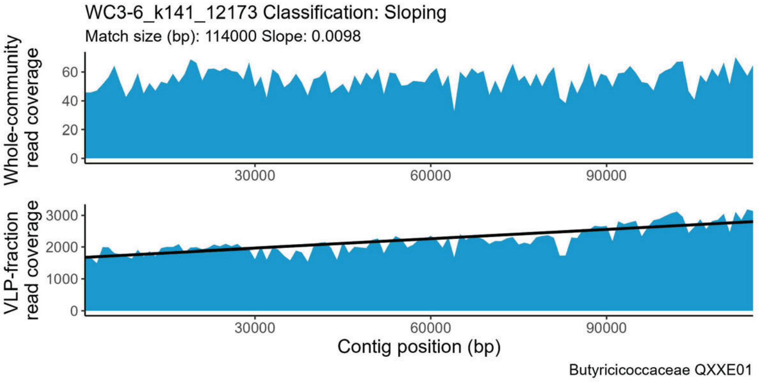
WC3-6_k141_12173 Classification: Sloping
Match size (bp): 114000 Slope: 0.0098
Whole-community read coverage
VLP-fraction read coverage
Contig position (bp)
Butyricicoccaceae QXXE01

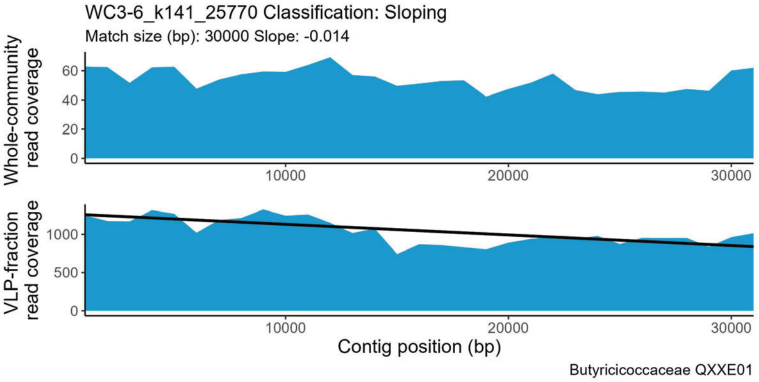
WC3-6_k141_25770 Classification: Sloping
Match size (bp): 30000 Slope: -0.014
Whole-community read coverage
VLP-fraction read coverage
Contig position (bp)
Butyricicoccaceae QXXE01

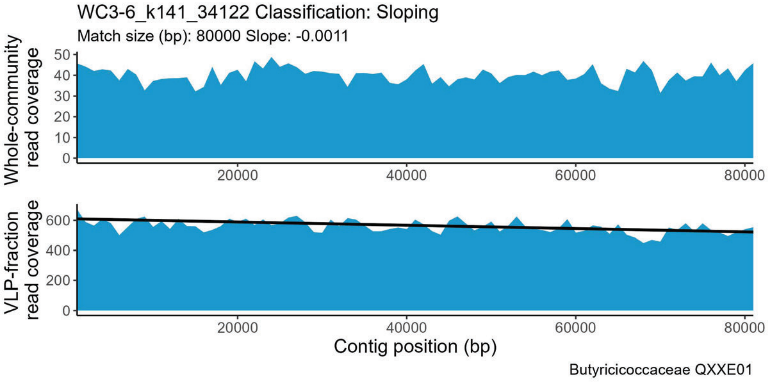
WC3-6_k141_34122 Classification: Sloping
Match size (bp): 80000 Slope: -0.0011
Whole-community read coverage
VLP-fraction read coverage
Contig position (bp)
Butyricicoccaceae QXXE01

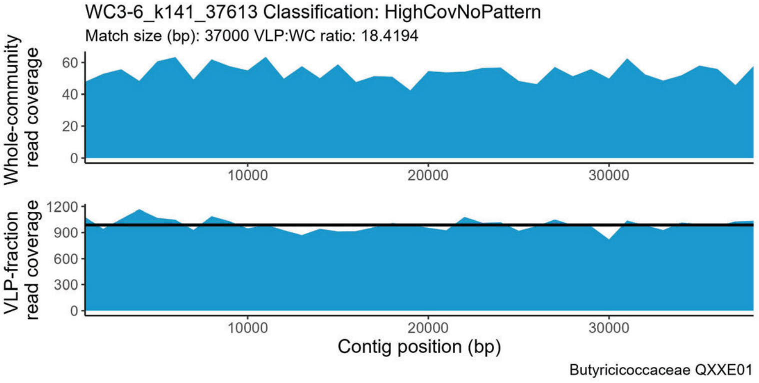
WC3-6_k141_37613 Classification: HighCovNoPattern
Match size (bp): 37000 VLP:WC ratio: 18.4194
Whole-community read coverage
VLP-fraction read coverage
Contig position (bp)
Butyricicoccaceae QXXE01

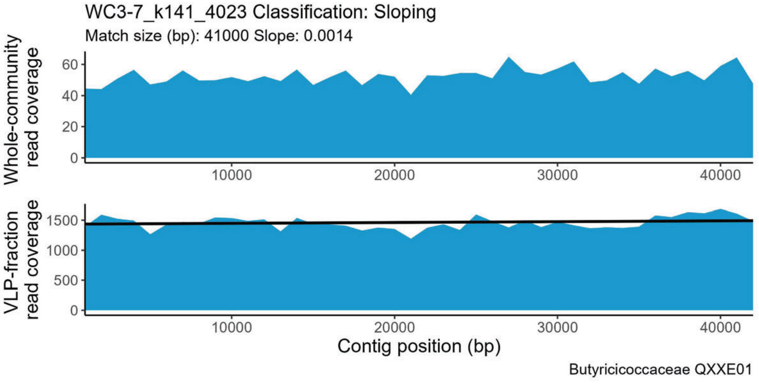
WC3-7_k141_4023 Classification: Sloping
Match size (bp): 41000 Slope: 0.0014
Whole-community read coverage
VLP-fraction read coverage
Contig position (bp)
Butyricicoccaceae QXXE01

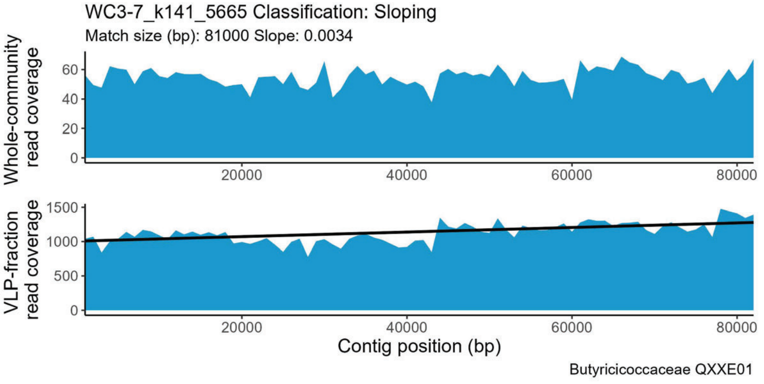
WC3-7_k141_5665 Classification: Sloping
Match size (bp): 81000 Slope: 0.0034
Whole-community read coverage
VLP-fraction read coverage
Contig position (bp)
Butyricicoccaceae QXXE01

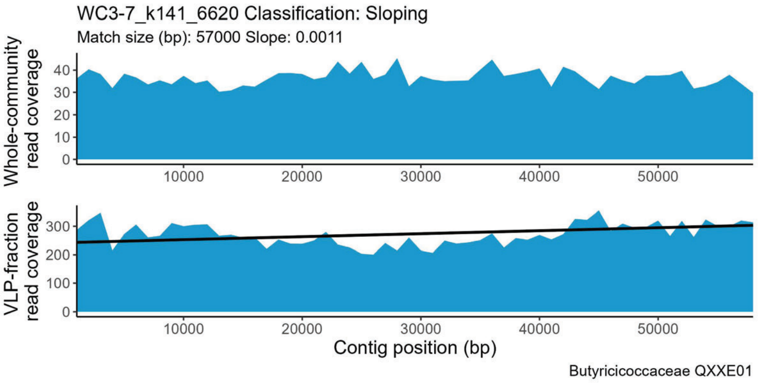
WC3-7_k141_6620 Classification: Sloping
Match size (bp): 57000 Slope: 0.0011
Whole-community read coverage
VLP-fraction read coverage
Contig position (bp)
Butyricicoccaceae QXXE01

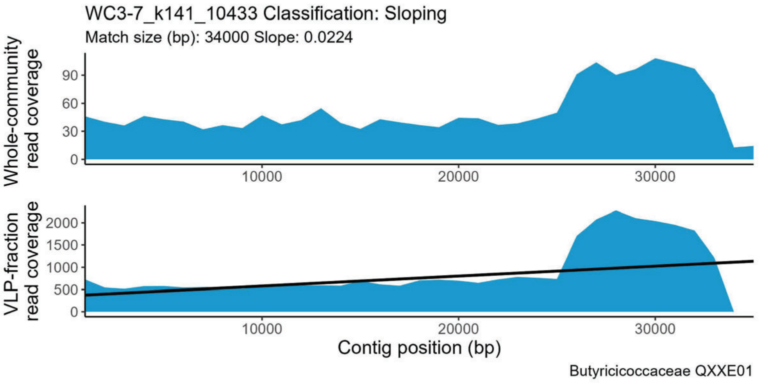
WC3-7_k141_10433 Classification: Sloping
Match size (bp): 34000 Slope: 0.0224
Whole-community read coverage
VLP-fraction read coverage
Contig position (bp)
Butyricicoccaceae QXXE01

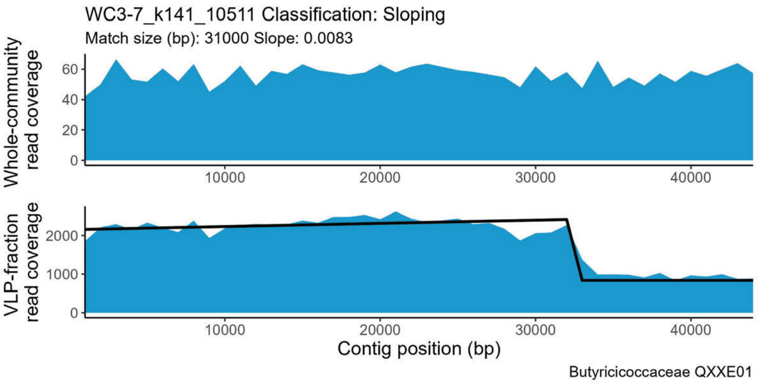
WC3-7_k141_10511 Classification: Sloping
Match size (bp): 31000 Slope: 0.0083
Whole-community read coverage
VLP-fraction read coverage
Contig position (bp)
Butyricicoccaceae QXXE01

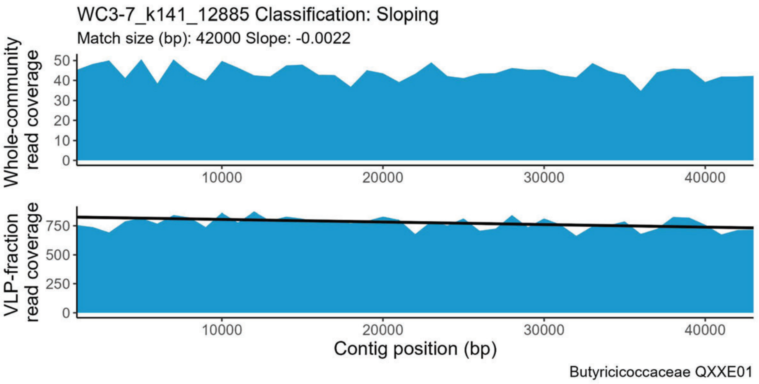
WC3-7_k141_12885 Classification: Sloping
Match size (bp): 42000 Slope: -0.0022
Whole-community read coverage
VLP-fraction read coverage
Contig position (bp)
Butyricicoccaceae QXXE01

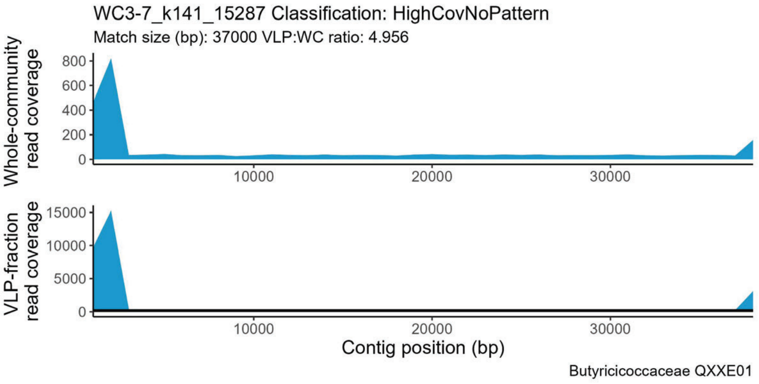
WC3-7_k141_15287 Classification: HighCovNoPattern
Match size (bp): 37000 VLP:WC ratio: 4.956
Whole-community read coverage
VLP-fraction read coverage
Contig position (bp)
Butyricicoccaceae QXXE01

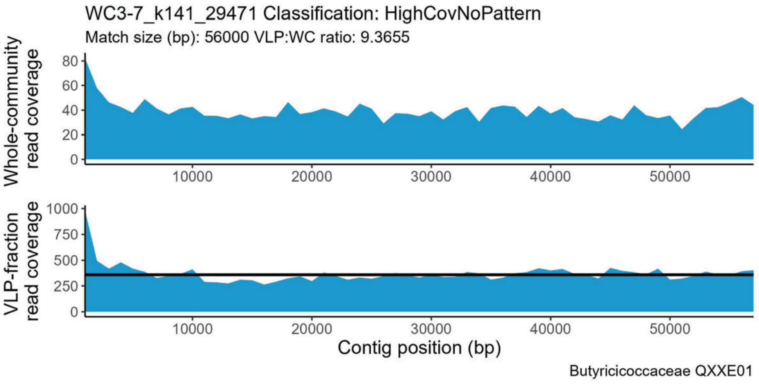
WC3-7_k141_29471 Classification: HighCovNoPattern
Match size (bp): 56000 VLP:WC ratio: 9.3655
Whole-community read coverage
VLP-fraction read coverage
Contig position (bp)
Butyricicoccaceae QXXE01

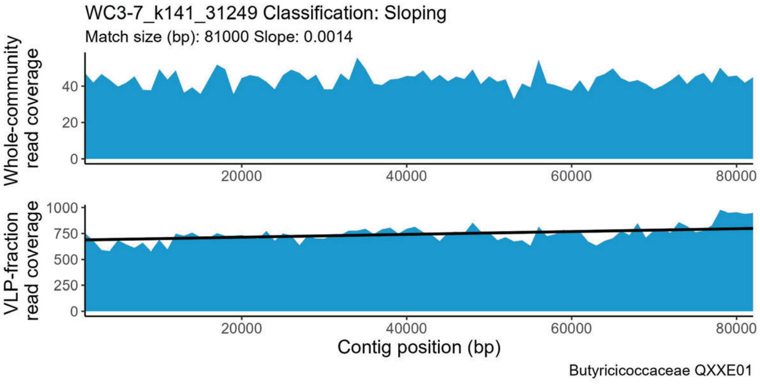
WC3-7_k141_31249 Classification: Sloping
Match size (bp): 81000 Slope: 0.0014
Whole-community read coverage
VLP-fraction read coverage
Contig position (bp)
Butyricicoccaceae QXXE01

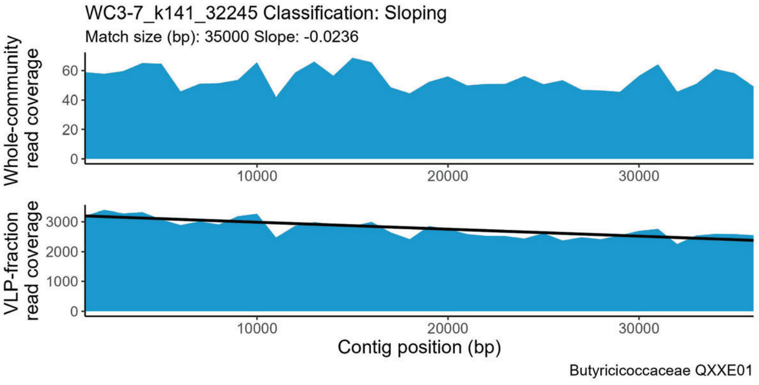
WC3-7_k141_32245 Classification: Sloping
Match size (bp): 35000 Slope: -0.0236
Whole-community read coverage
VLP-fraction read coverage
Contig position (bp)
Butyricicoccaceae QXXE01

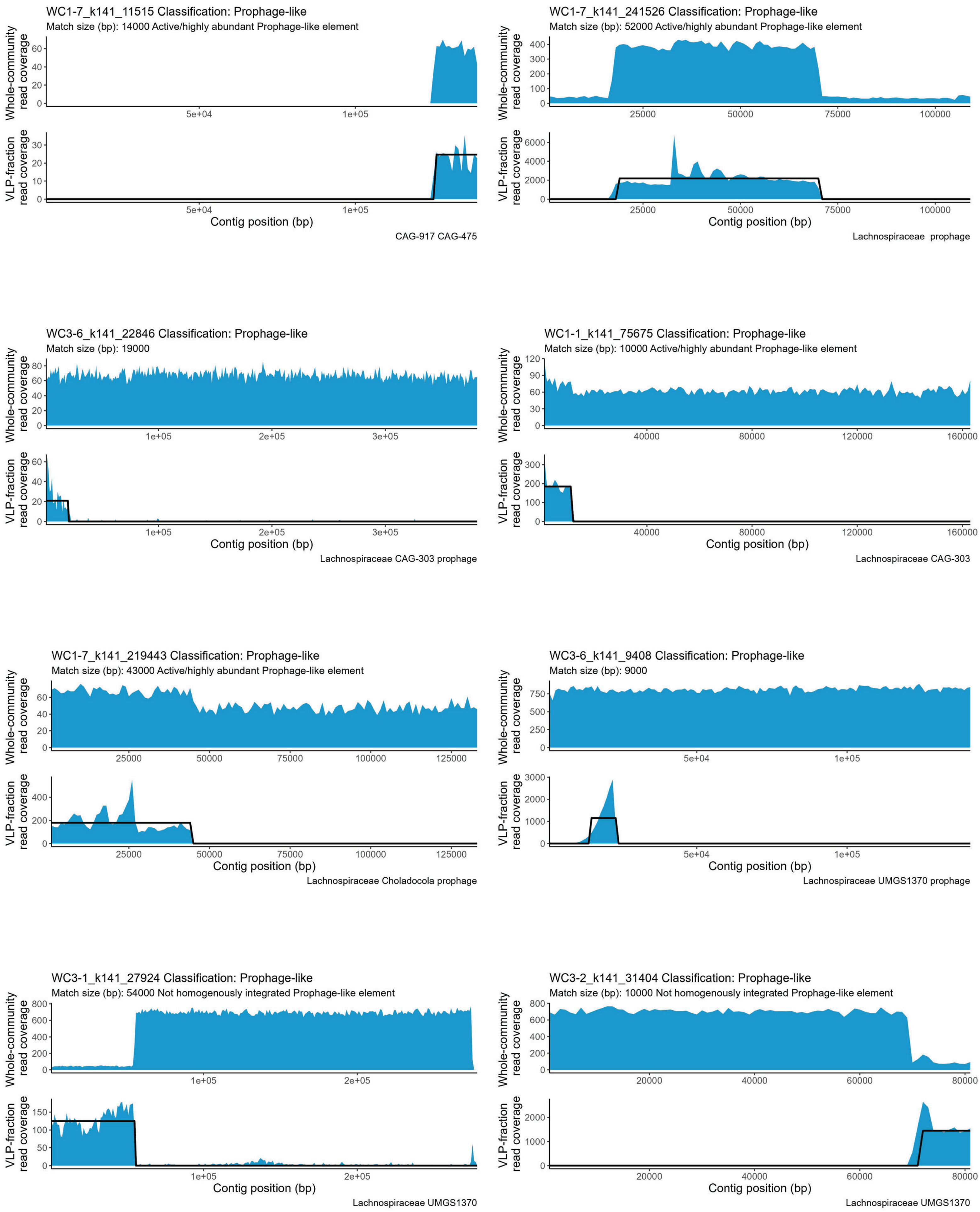

WC1-7_k141_11515 Classification: Prophage-like
Match size (bp): 14000 Active/highly abundant Prophage-like element
Whole-community read coverage
VLP-fraction read coverage
Contig position (bp)
CAG-917 CAG-475
WC1-7_k141_241526 Classification: Prophage-like
Match size (bp): 52000 Active/highly abundant Prophage-like element
Lachnospiraceae prophage
WC3-6_k141_22846 Classification: Prophage-like
Match size (bp): 19000
Lachnospiraceae CAG-303 prophage
WC1-1_k141_75675 Classification: Prophage-like
Match size (bp): 10000 Active/highly abundant Prophage-like element
Lachnospiraceae CAG-303
WC1-7_k141_219443 Classification: Prophage-like
Match size (bp): 43000 Active/highly abundant Prophage-like element
Lachnospiraceae Choladocola prophage
WC3-6_k141_9408 Classification: Prophage-like
Match size (bp): 9000
Lachnospiraceae UMGS1370 prophage
WC3-1_k141_27924 Classification: Prophage-like
Match size (bp): 54000 Not homogenously integrated Prophage-like element
Lachnospiraceae UMGS1370
WC3-2_k141_31404 Classification: Prophage-like
Match size (bp): 10000 Not homogenously integrated Prophage-like element
Lachnospiraceae UMGS1370

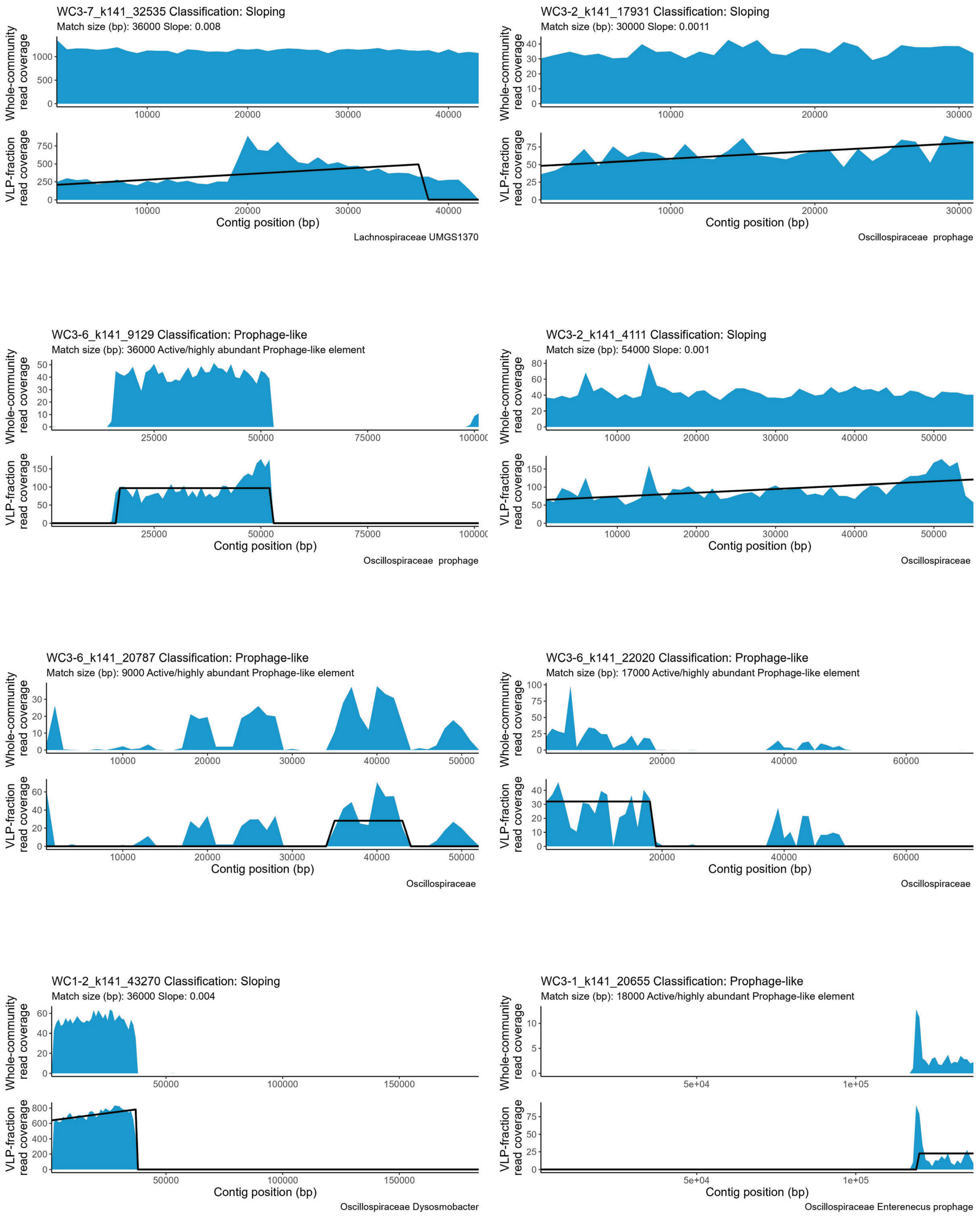

WC3-7_k141_32535 Classification: Sloping
Match size (bp): 36000 Slope: 0.008
Whole-community read coverage
VLP-fraction read coverage
Contig position (bp)
Lachnospiraceae UMGS1370
WC3-2_k141_17931 Classification: Sloping
Match size (bp): 30000 Slope: 0.0011
Whole-community read coverage
VLP-fraction read coverage
Contig position (bp)
Oscillospiraceae prophage
WC3-6_k141_9129 Classification: Prophage-like
Match size (bp): 36000 Active/highly abundant Prophage-like element
Whole-community read coverage
VLP-fraction read coverage
Contig position (bp)
Oscillospiraceae prophage
WC3-2_k141_4111 Classification: Sloping
Match size (bp): 54000 Slope: 0.001
Whole-community read coverage
VLP-fraction read coverage
Contig position (bp)
Oscillospiraceae
WC3-6_k141_20787 Classification: Prophage-like
Match size (bp): 9000 Active/highly abundant Prophage-like element
Whole-community read coverage
VLP-fraction read coverage
Contig position (bp)
Oscillospiraceae
WC3-6_k141_22020 Classification: Prophage-like
Match size (bp): 17000 Active/highly abundant Prophage-like element
Whole-community read coverage
VLP-fraction read coverage
Contig position (bp)
Oscillospiraceae
WC1-2_k141_43270 Classification: Sloping
Match size (bp): 36000 Slope: 0.004
Whole-community read coverage
VLP-fraction read coverage
Contig position (bp)
Oscillospiraceae Dysosmobacter
WC3-1_k141_20655 Classification: Prophage-like
Match size (bp): 18000 Active/highly abundant Prophage-like element
Whole-community read coverage
VLP-fraction read coverage
Contig position (bp)
Oscillospiraceae Enterenecus prophage

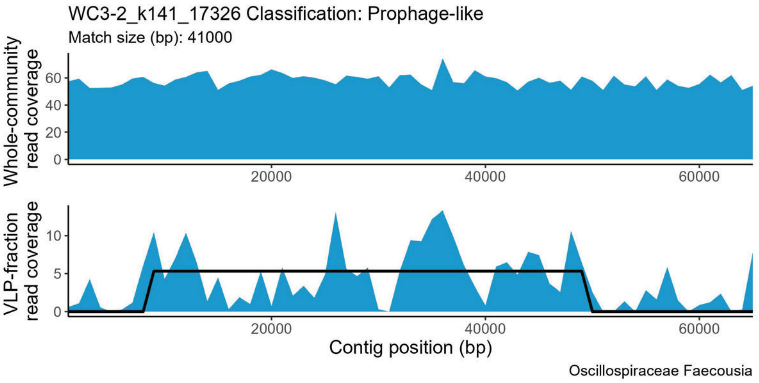
WC3-2_k141_17326 Classification: Prophage-like
Match size (bp): 41000
Whole-community read coverage
VLP-fraction read coverage
Contig position (bp)
Oscillospiraceae Faecousia

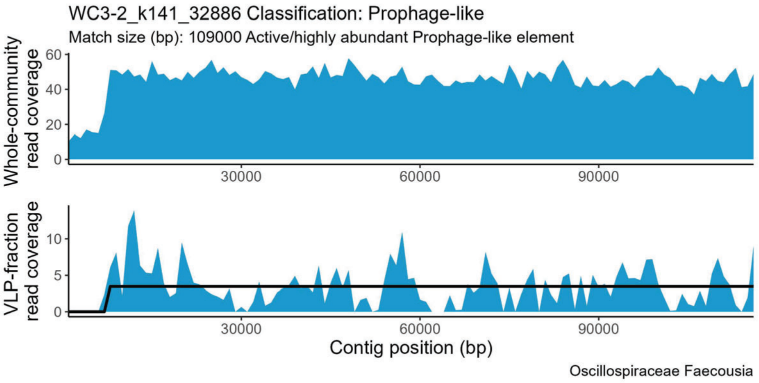
WC3-2_k141_32886 Classification: Prophage-like
Match size (bp): 109000 Active/highly abundant Prophage-like element
Whole-community read coverage
VLP-fraction read coverage
Contig position (bp)
Oscillospiraceae Faecousia

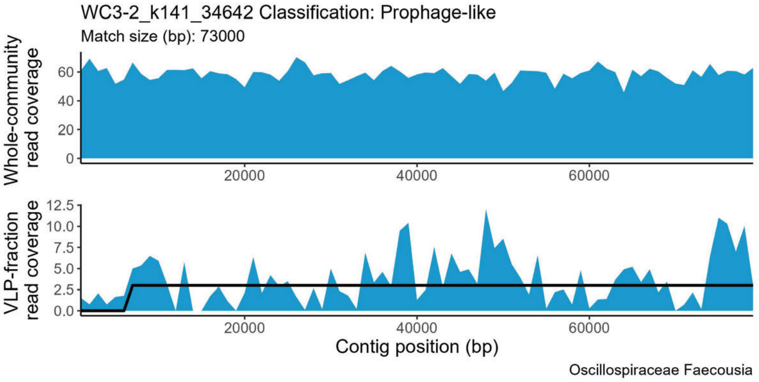
WC3-2_k141_34642 Classification: Prophage-like
Match size (bp): 73000
Whole-community read coverage
VLP-fraction read coverage
Contig position (bp)
Oscillospiraceae Faecousia

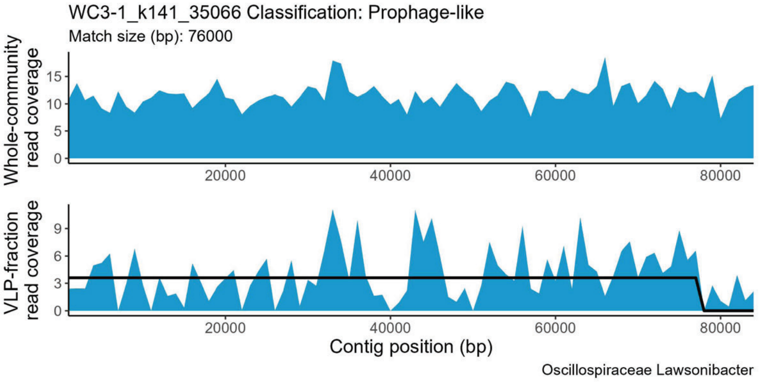
WC3-1_k141_35066 Classification: Prophage-like
Match size (bp): 76000
Whole-community read coverage
VLP-fraction read coverage
Contig position (bp)
Oscillospiraceae Lawsonibacter

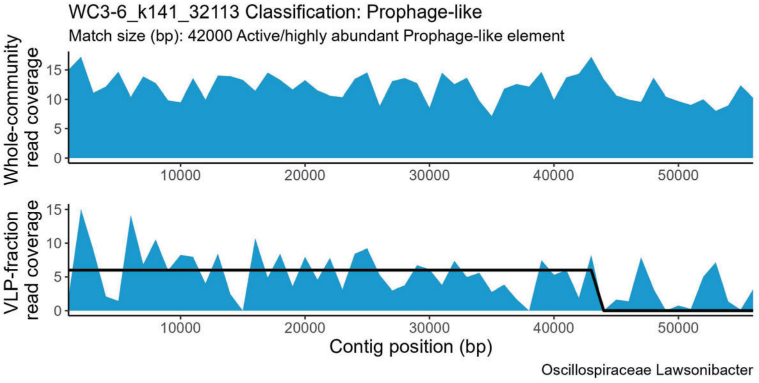
WC3-6_k141_32113 Classification: Prophage-like
Match size (bp): 42000 Active/highly abundant Prophage-like element
Whole-community read coverage
VLP-fraction read coverage
Contig position (bp)
Oscillospiraceae Lawsonibacter

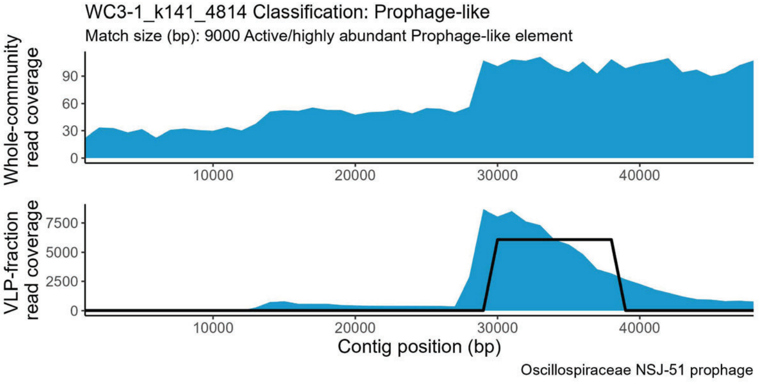
WC3-1_k141_4814 Classification: Prophage-like
Match size (bp): 9000 Active/highly abundant Prophage-like element
Whole-community read coverage
VLP-fraction read coverage
Contig position (bp)
Oscillospiraceae NSJ-51 prophage

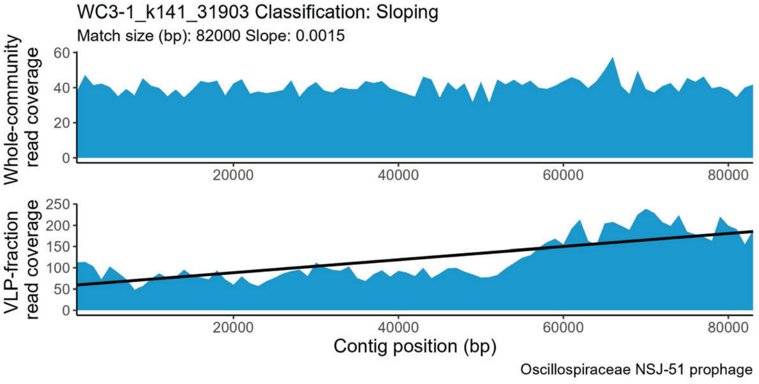
WC3-1_k141_31903 Classification: Sloping
Match size (bp): 82000 Slope: 0.0015
Whole-community read coverage
VLP-fraction read coverage
Contig position (bp)
Oscillospiraceae NSJ-51 prophage

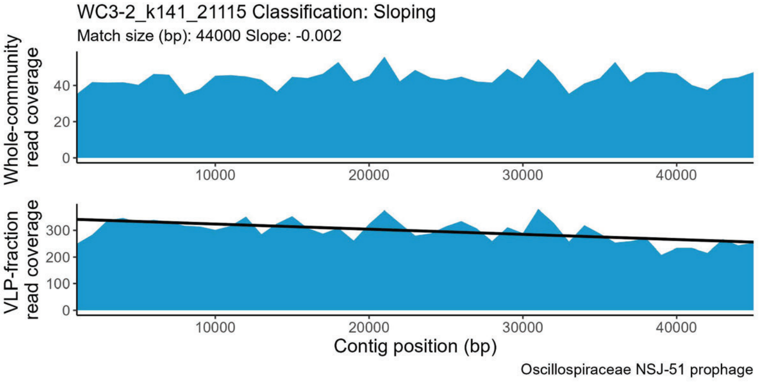
WC3-2_k141_21115 Classification: Sloping
Match size (bp): 44000 Slope: -0.002
Whole-community read coverage
VLP-fraction read coverage
Contig position (bp)
Oscillospiraceae NSJ-51 prophage

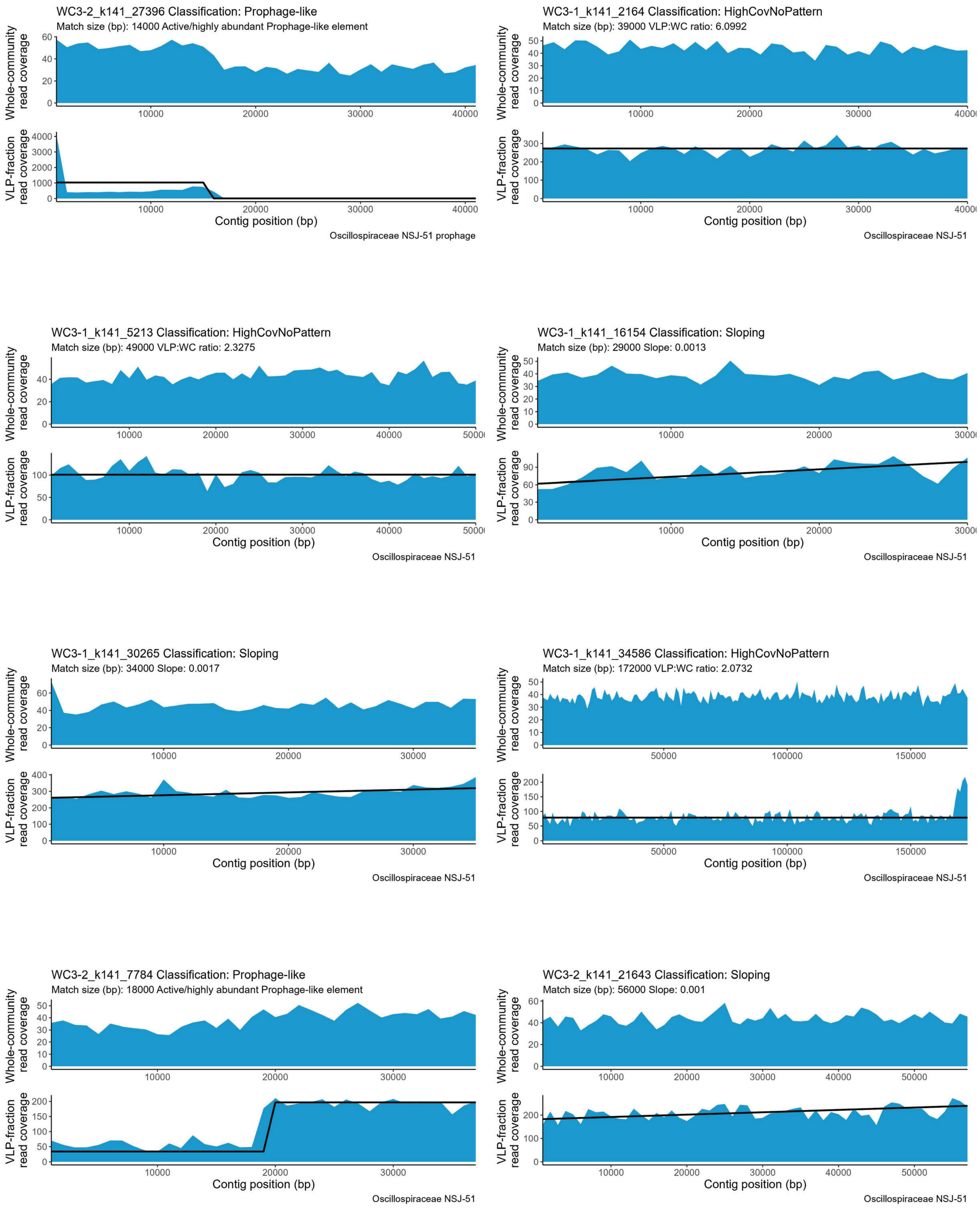

WC3-2_k141_27396 Classification: Prophage-like
Match size (bp): 14000 Active/highly abundant Prophage-like element
Whole-community read coverage
VLP-fraction read coverage
Contig position (bp)
Oscillospiraceae NSJ-51 prophage
WC3-1_k141_2164 Classification: HighCovNoPattern
Match size (bp): 39000 VLP:WC ratio: 6.0992
Whole-community read coverage
VLP-fraction read coverage
Contig position (bp)
Oscillospiraceae NSJ-51
WC3-1_k141_5213 Classification: HighCovNoPattern
Match size (bp): 49000 VLP:WC ratio: 2.3275
Whole-community read coverage
VLP-fraction read coverage
Contig position (bp)
Oscillospiraceae NSJ-51
WC3-1_k141_16154 Classification: Sloping
Match size (bp): 29000 Slope: 0.0013
Whole-community read coverage
VLP-fraction read coverage
Contig position (bp)
Oscillospiraceae NSJ-51
WC3-1_k141_30265 Classification: Sloping
Match size (bp): 34000 Slope: 0.0017
Whole-community read coverage
VLP-fraction read coverage
Contig position (bp)
Oscillospiraceae NSJ-51
WC3-1_k141_34586 Classification: HighCovNoPattern
Match size (bp): 172000 VLP:WC ratio: 2.0732
Whole-community read coverage
VLP-fraction read coverage
Contig position (bp)
Oscillospiraceae NSJ-51
WC3-2_k141_7784 Classification: Prophage-like
Match size (bp): 18000 Active/highly abundant Prophage-like element
Whole-community read coverage
VLP-fraction read coverage
Contig position (bp)
Oscillospiraceae NSJ-51
WC3-2_k141_21643 Classification: Sloping
Match size (bp): 56000 Slope: 0.001
Whole-community read coverage
VLP-fraction read coverage
Contig position (bp)
Oscillospiraceae NSJ-51

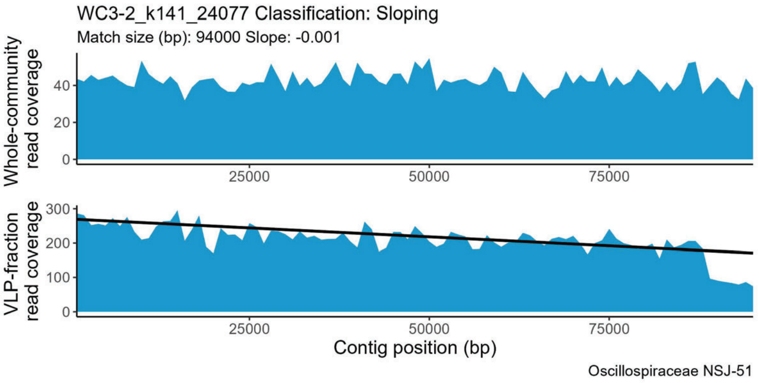
WC3-2_k141_24077 Classification: Sloping
Match size (bp): 94000 Slope: -0.001
Whole-community read coverage
VLP-fraction read coverage
Contig position (bp)
Oscillospiraceae NSJ-51

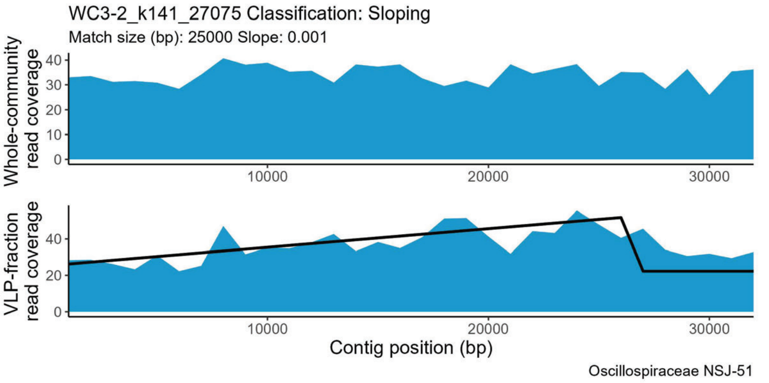
WC3-2_k141_27075 Classification: Sloping
Match size (bp): 25000 Slope: 0.001
Whole-community read coverage
VLP-fraction read coverage
Contig position (bp)
Oscillospiraceae NSJ-51

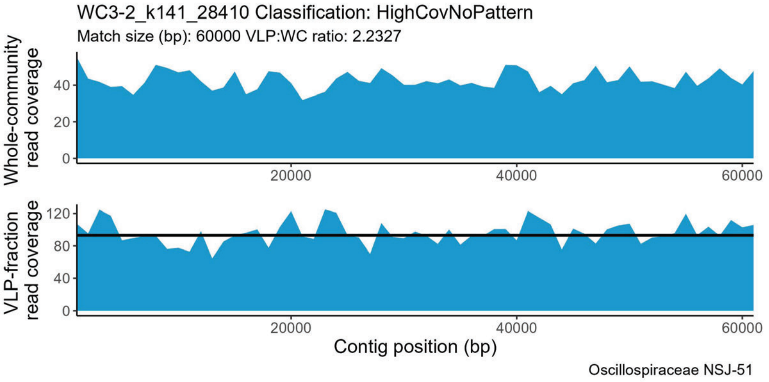
WC3-2_k141_28410 Classification: HighCovNoPattern
Match size (bp): 60000 VLP:WC ratio: 2.2327
Whole-community read coverage
VLP-fraction read coverage
Contig position (bp)
Oscillospiraceae NSJ-51

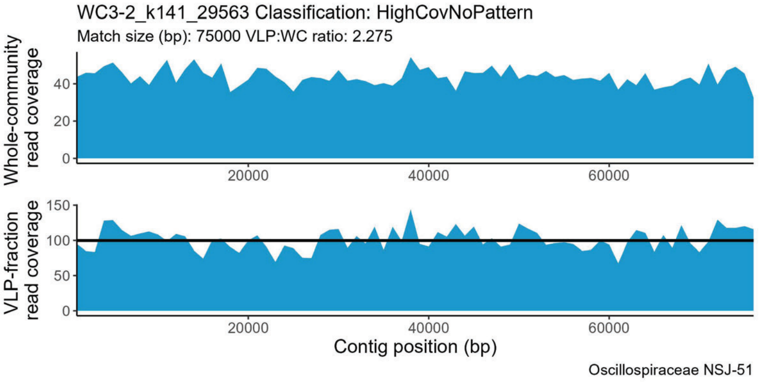
WC3-2_k141_29563 Classification: HighCovNoPattern
Match size (bp): 75000 VLP:WC ratio: 2.275
Whole-community read coverage
VLP-fraction read coverage
Contig position (bp)
Oscillospiraceae NSJ-51

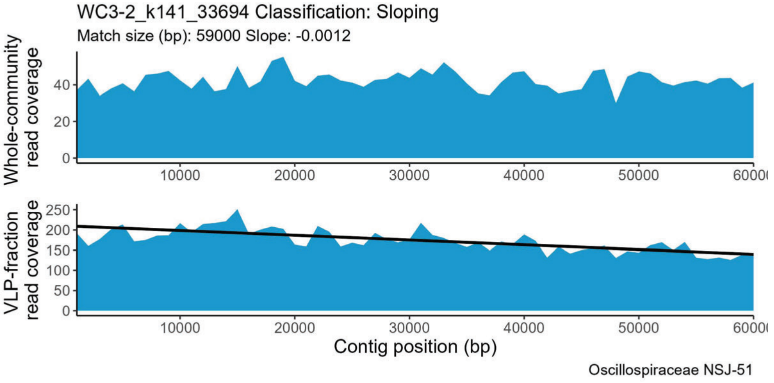
WC3-2_k141_33694 Classification: Sloping
Match size (bp): 59000 Slope: -0.0012
Whole-community read coverage
VLP-fraction read coverage
Contig position (bp)
Oscillospiraceae NSJ-51

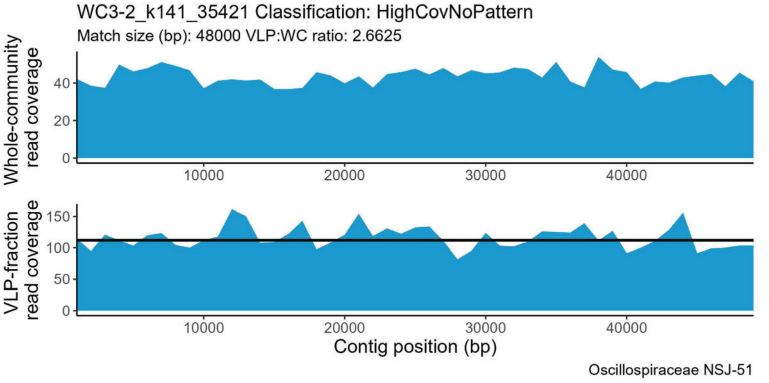
WC3-2_k141_35421 Classification: HighCovNoPattern
Match size (bp): 48000 VLP:WC ratio: 2.6625
Whole-community read coverage
VLP-fraction read coverage
Contig position (bp)
Oscillospiraceae NSJ-51

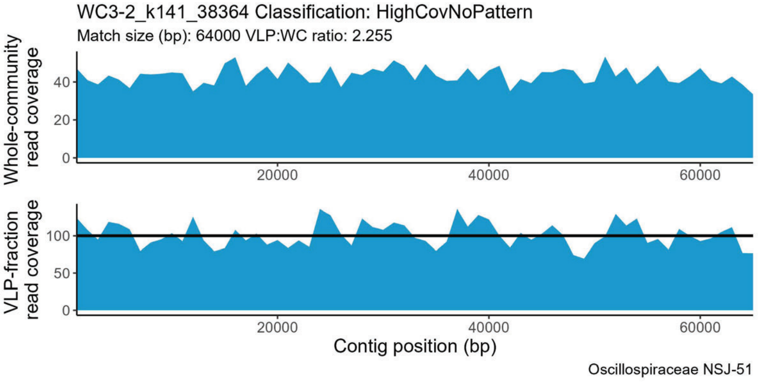
WC3-2_k141_38364 Classification: HighCovNoPattern
Match size (bp): 64000 VLP:WC ratio: 2.255
Whole-community read coverage
VLP-fraction read coverage
Contig position (bp)
Oscillospiraceae NSJ-51

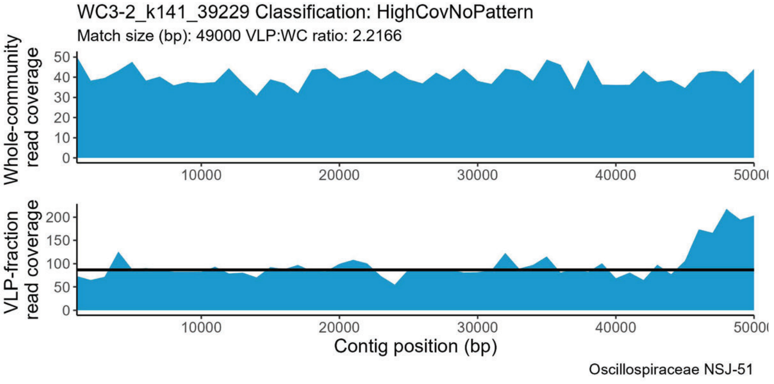
WC3-2_k141_39229 Classification: HighCovNoPattern
Match size (bp): 49000 VLP:WC ratio: 2.2166
Whole-community read coverage
VLP-fraction read coverage
Contig position (bp)
Oscillospiraceae NSJ-51

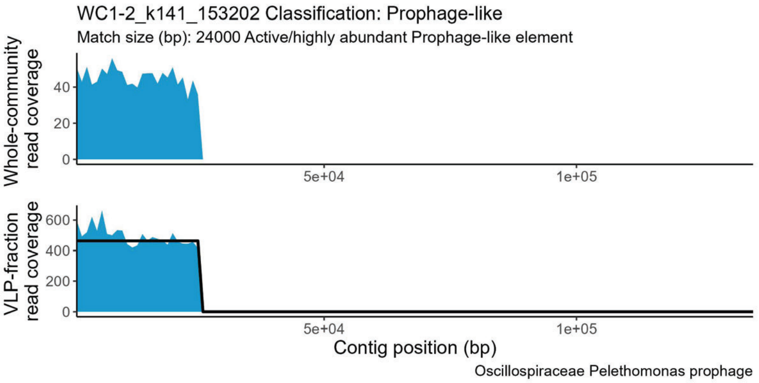
WC1-2_k141_153202 Classification: Prophage-like
Match size (bp): 24000 Active/highly abundant Prophage-like element
Whole-community read coverage
VLP-fraction read coverage
Contig position (bp)
Oscillospiraceae Pelethomonas prophage

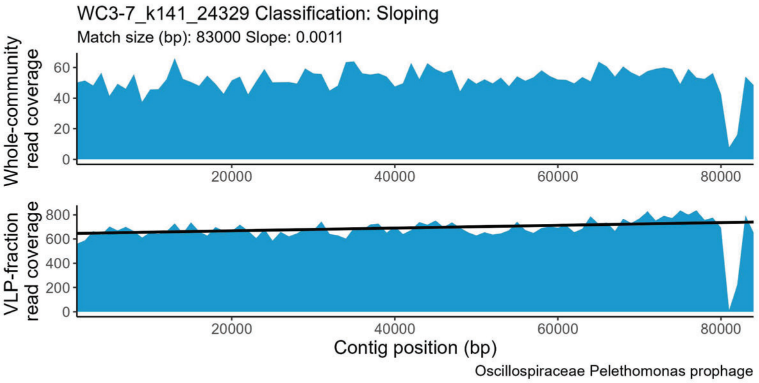
WC3-7_k141_24329 Classification: Sloping
Match size (bp): 83000 Slope: 0.0011
Whole-community read coverage
VLP-fraction read coverage
Contig position (bp)
Oscillospiraceae Pelethomonas prophage

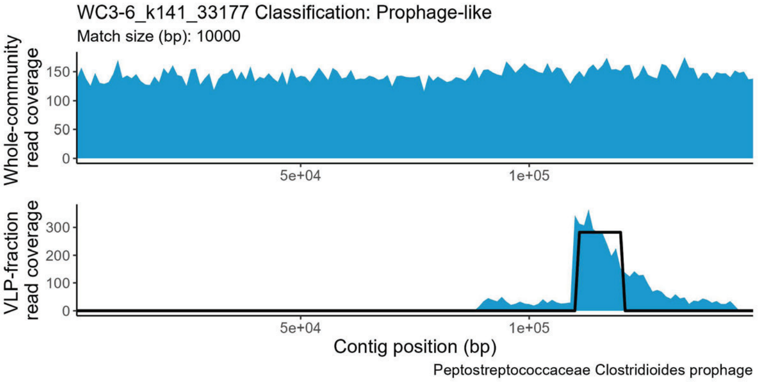
WC3-6_k141_33177 Classification: Prophage-like
Match size (bp): 10000
Whole-community read coverage
VLP-fraction read coverage
Contig position (bp)
Peptostreptococcaceae Clostridioides prophage

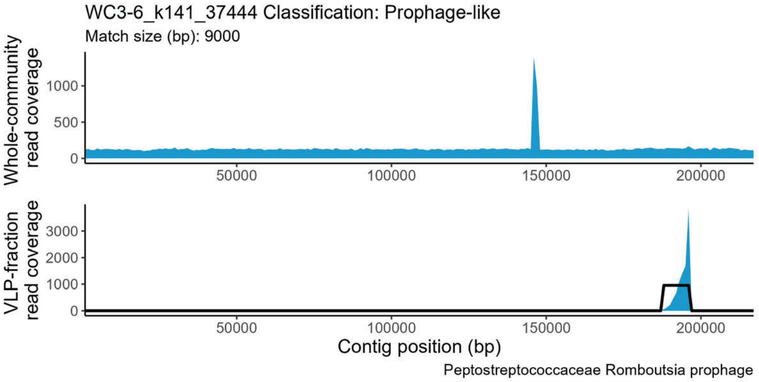
WC3-6_k141_37444 Classification: Prophage-like
Match size (bp): 9000
Whole-community read coverage
VLP-fraction read coverage
Contig position (bp)
Peptostreptococcaceae Romboutsia prophage

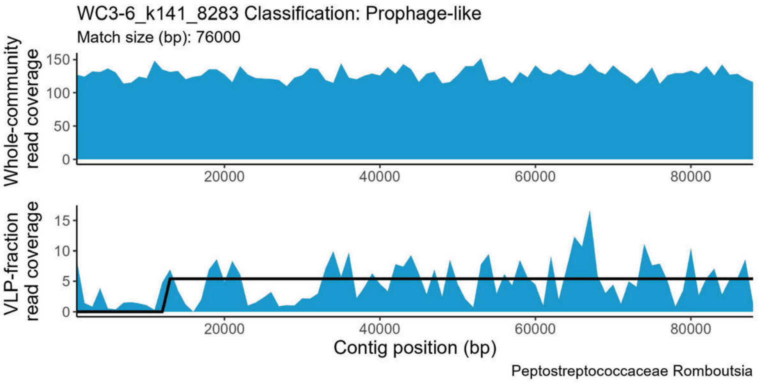
WC3-6_k141_8283 Classification: Prophage-like
Match size (bp): 76000
Whole-community read coverage
VLP-fraction read coverage
Contig position (bp)
Peptostreptococcaceae Romboutsia

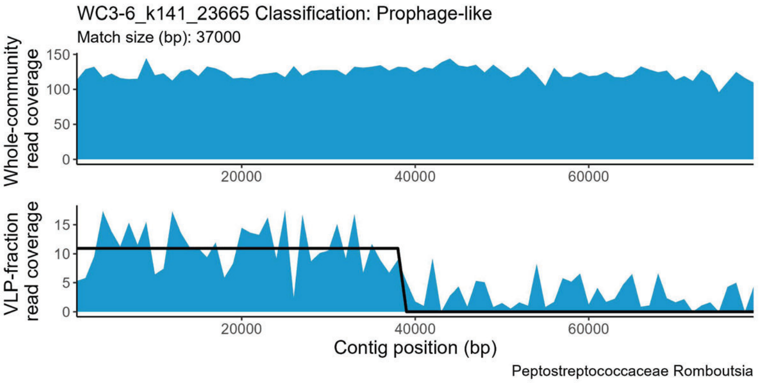
WC3-6_k141_23665 Classification: Prophage-like
Match size (bp): 37000
Whole-community read coverage
VLP-fraction read coverage
Contig position (bp)
Peptostreptococcaceae Romboutsia

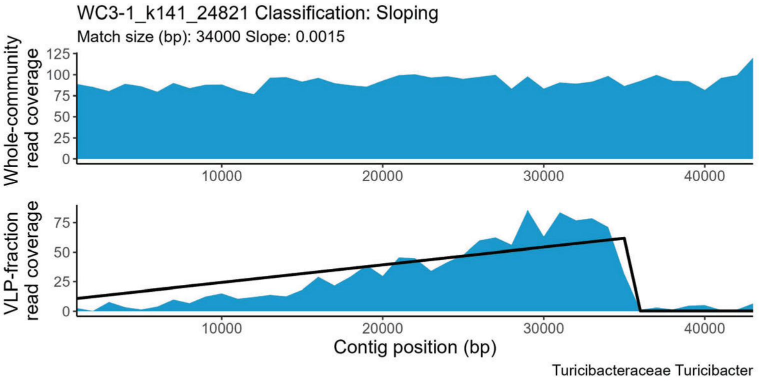
WC3-1_k141_24821 Classification: Sloping
Match size (bp): 34000 Slope: 0.0015
Whole-community read coverage
VLP-fraction read coverage
Contig position (bp)
Turicibacteraceae Turicibacter

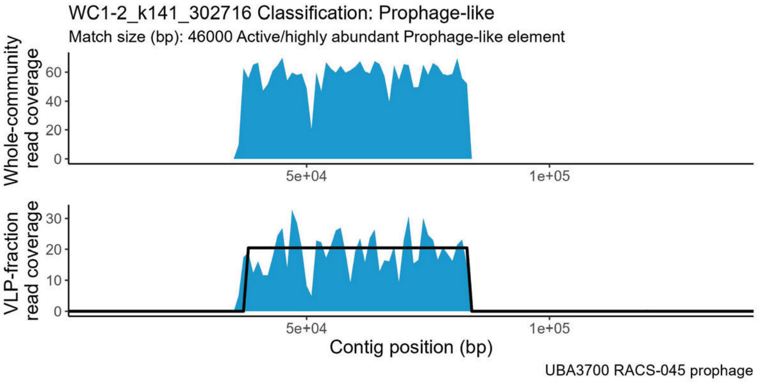
WC1-2_k141_302716 Classification: Prophage-like
Match size (bp): 46000 Active/highly abundant Prophage-like element
Whole-community read coverage
VLP-fraction read coverage
Contig position (bp)
UBA3700 RACS-045 prophage

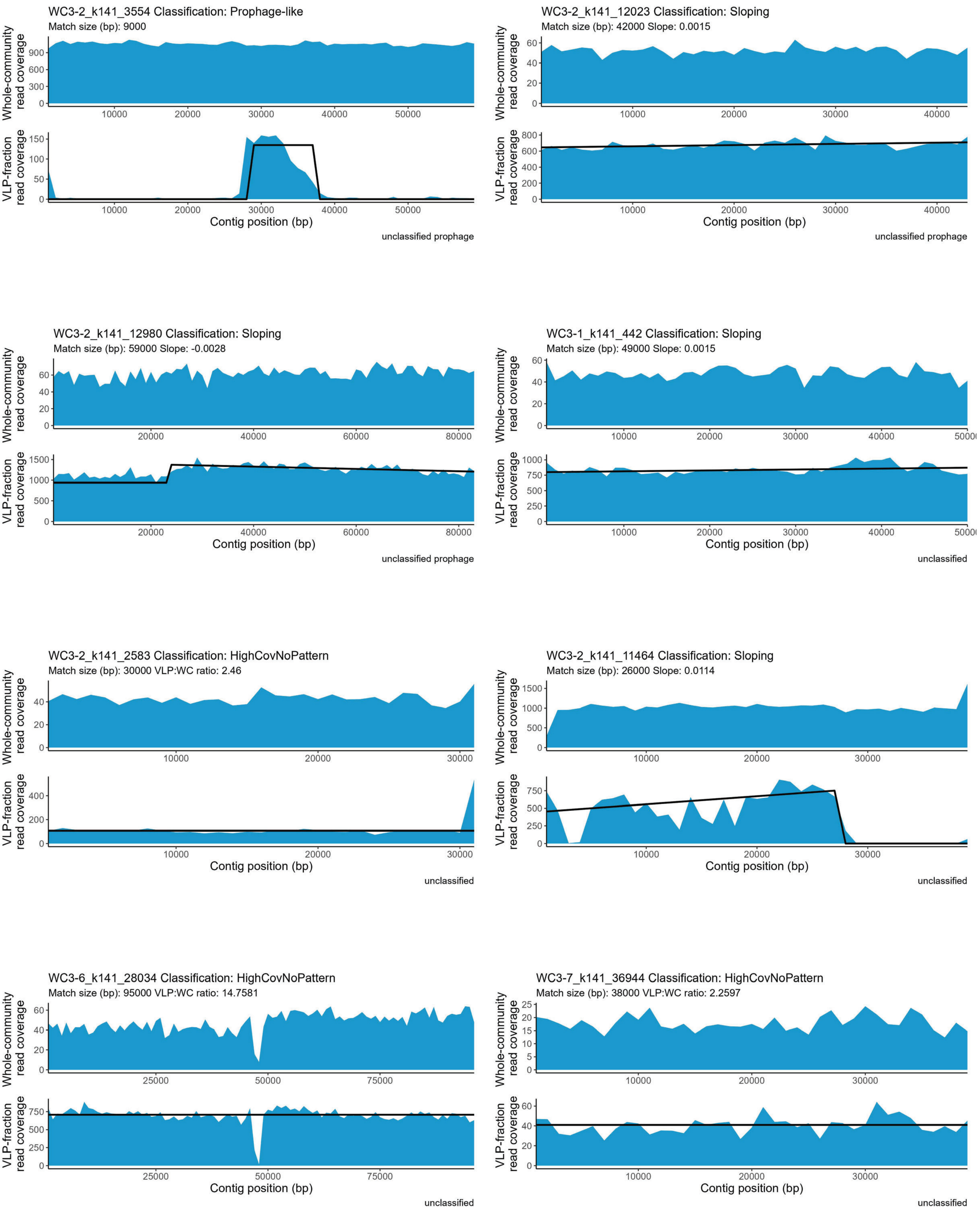

WC3-2_k141_3554 Classification: Prophage-like
Match size (bp): 9000
WC3-2_k141_12023 Classification: Sloping
Match size (bp): 42000 Slope: 0.0015
WC3-2_k141_12980 Classification: Sloping
Match size (bp): 59000 Slope: -0.0028
WC3-1_k141_442 Classification: Sloping
Match size (bp): 49000 Slope: 0.0015
WC3-2_k141_2583 Classification: HighCovNoPattern
Match size (bp): 30000 VLP:WC ratio: 2.46
WC3-2_k141_11464 Classification: Sloping
Match size (bp): 26000 Slope: 0.0114
WC3-6_k141_28034 Classification: HighCovNoPattern
Match size (bp): 95000 VLP:WC ratio: 14.7581
WC3-7_k141_36944 Classification: HighCovNoPattern
Match size (bp): 38000 VLP:WC ratio: 2.2597
Whole-community read coverage
VLP-fraction read coverage
Contig position (bp)
unclassified prophage
unclassified

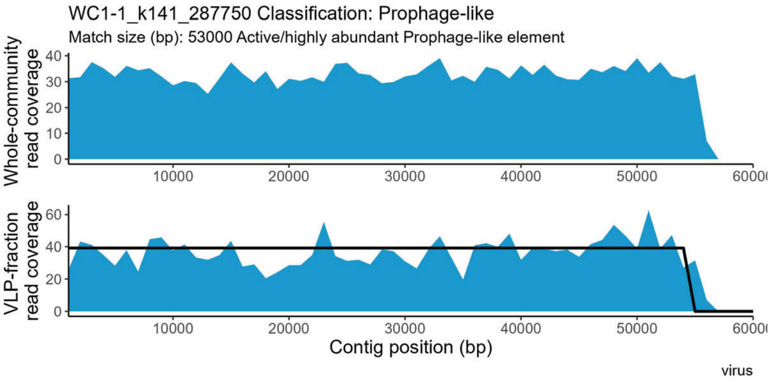

WC1-1_k141_287750 Classification: Prophage-like
Match size (bp): 53000 Active/highly abundant Prophage-like element
Whole-community read coverage
0
10
20
30
40
10000
20000
30000
40000
50000
60000
VLP-fraction read coverage
0
20
40
60
Contig position (bp)
virus


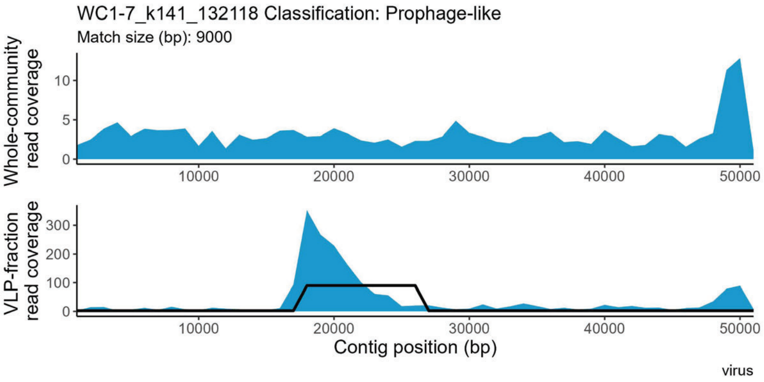

WC1-7_k141_132118 Classification: Prophage-like
Match size (bp): 9000
Whole-community read coverage
0
5
10
10000
20000
30000
40000
50000
VLP-fraction read coverage
0
100
200
300
Contig position (bp)
virus


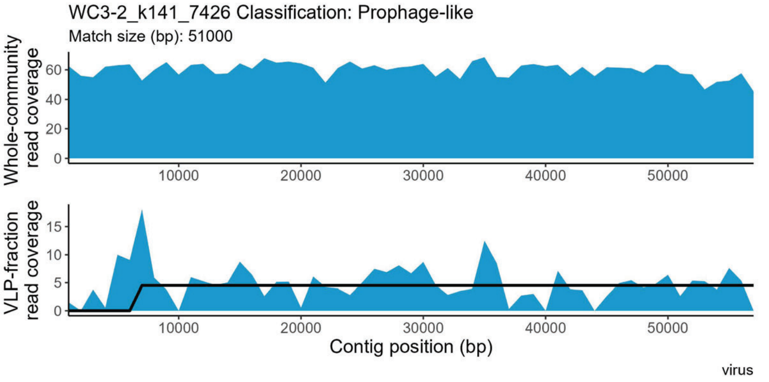

WC3-2_k141_7426 Classification: Prophage-like
Match size (bp): 51000
Whole-community read coverage
0
20
40
60
10000
20000
30000
40000
50000
VLP-fraction read coverage
0
5
10
15
Contig position (bp)
virus


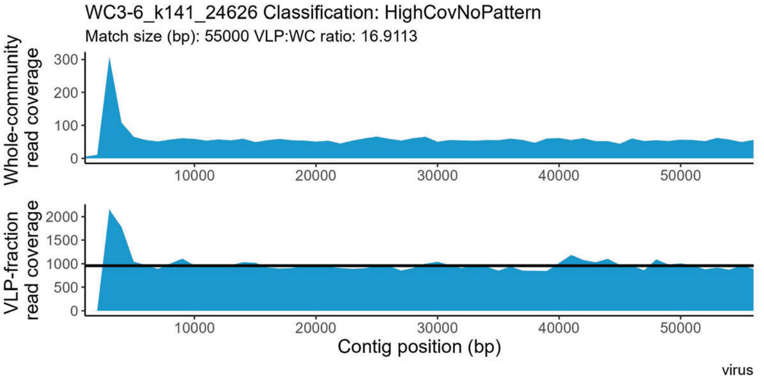

WC3-6_k141_24626 Classification: HighCovNoPattern
Match size (bp): 55000 VLP:WC ratio: 16.9113
Whole-community read coverage
0
100
200
300
10000
20000
30000
40000
50000
VLP-fraction read coverage
0
500
1000
1500
2000
Contig position (bp)
virus
