## Supplemental file 2 for "Perturbations shift the composition of bacterial DNA carried by virus-like particles in the murine gut microbiome"

**File S2) All positive TrIdent pattern-matches for Pre-ABX replicate 2.** The sample from which a contig was assembled is indicated by the following prefixes in the contig accession number: WC1-1 = Pre-ABX replicate 1, WC1-2 = Pre-ABX replicate 2 WC1-6 = Pre-ABX replicate 3, WC1-7 = Pre-ABX replicate 4, WC1-1 = CDI replicate 1, WC1-2 = CDI replicate 2, WC1-6 = CDI replicate 3, WC1-7 = CDI replicate 4. Pre-ABX = Pre-antibiotics, Post-ABX= post-antbiotics, CDI= *Clostridioides difficile* infection. Contig taxonomy and geNomad prophage classification in plot subtitle. Plots are listed alphabetically by family name.

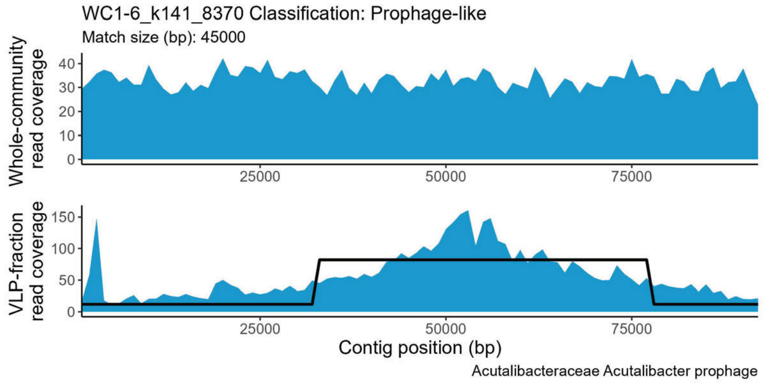


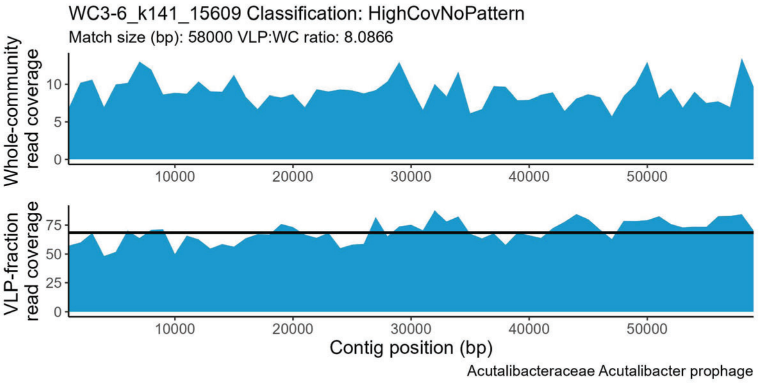


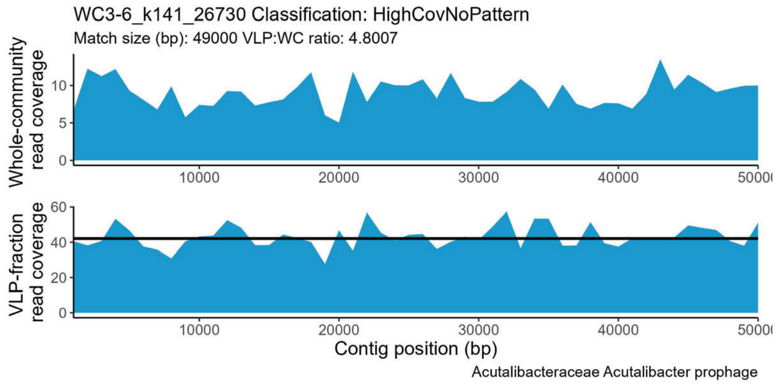


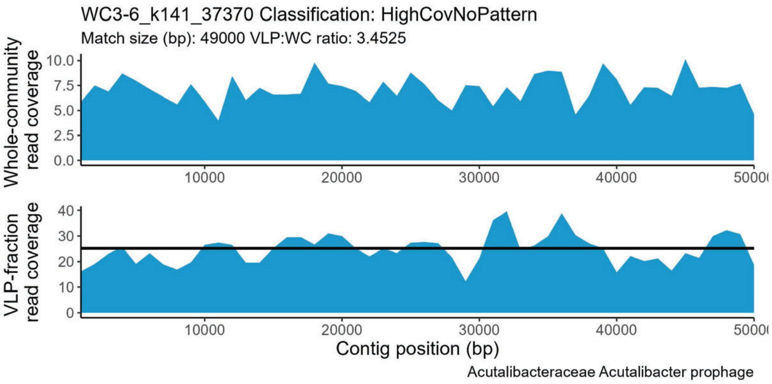


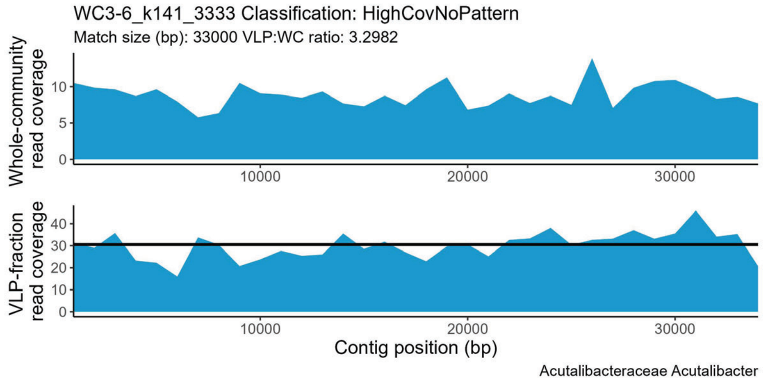


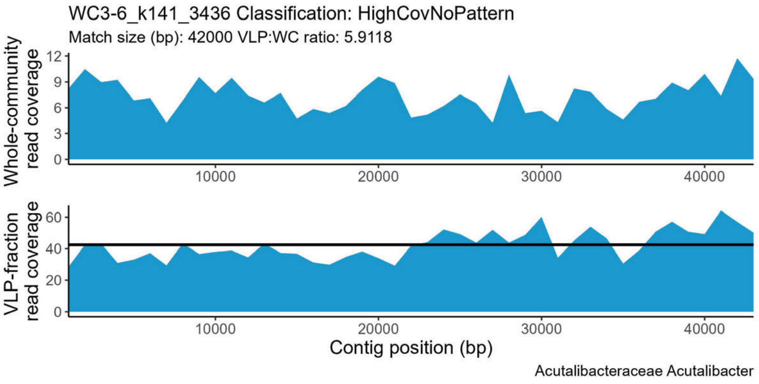


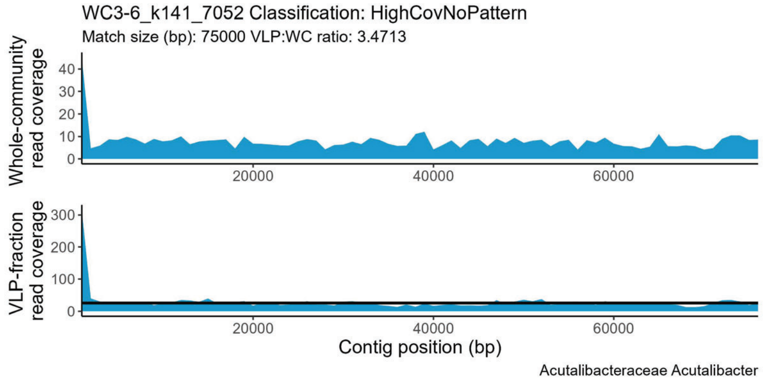


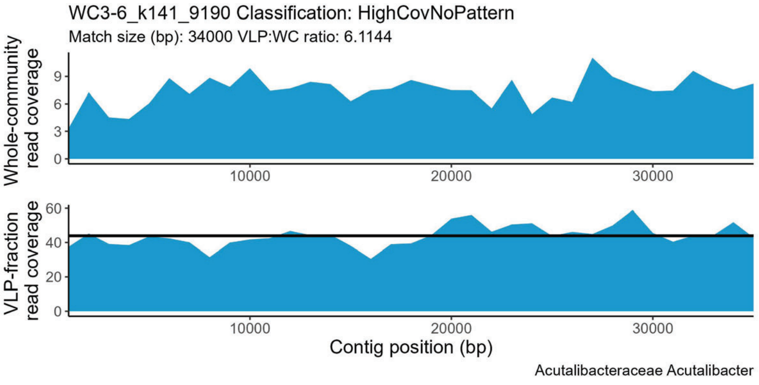

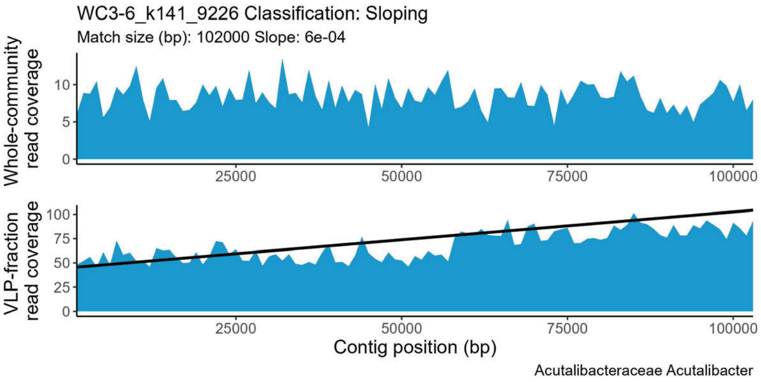
WC3-6_k141_9226 Classification: Sloping
Match size (bp): 102000 Slope: 6e-04
Whole-community read coverage
0 5 10
25000 50000 75000 100000
VLP-fraction read coverage
0 25 50 75 100
25000 50000 75000 100000
Contig position (bp)
Acutalibacteraceae Acutalibacter

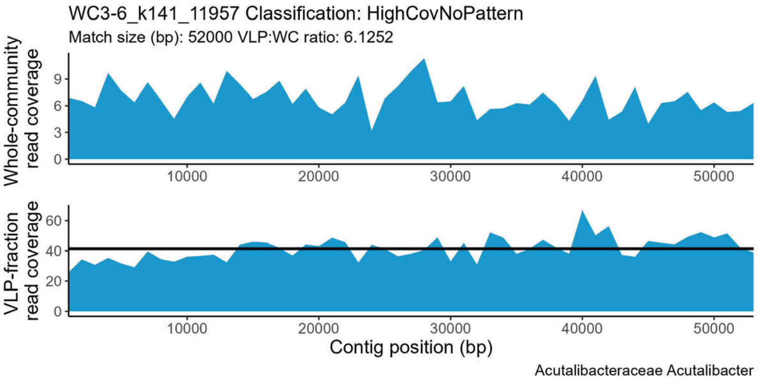
WC3-6_k141_11957 Classification: HighCovNoPattern
Match size (bp): 52000 VLP:WC ratio: 6.1252
Whole-community read coverage
0 3 6 9
10000 20000 30000 40000 50000
VLP-fraction read coverage
0 20 40 60
10000 20000 30000 40000 50000
Contig position (bp)
Acutalibacteraceae Acutalibacter

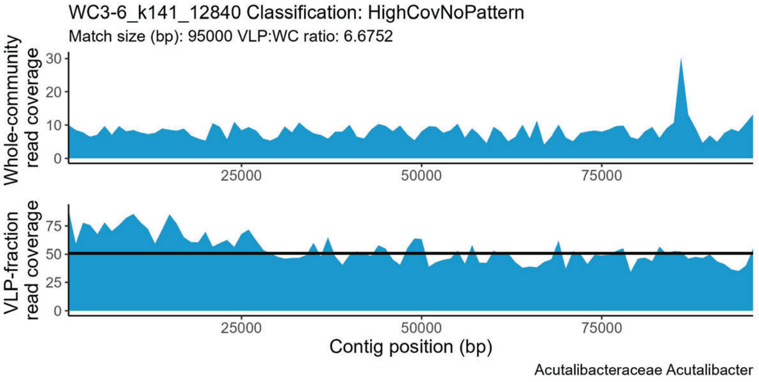
WC3-6_k141_12840 Classification: HighCovNoPattern
Match size (bp): 95000 VLP:WC ratio: 6.6752
Whole-community read coverage
0 10 20 30
25000 50000 75000
VLP-fraction read coverage
0 25 50 75
25000 50000 75000
Contig position (bp)
Acutalibacteraceae Acutalibacter

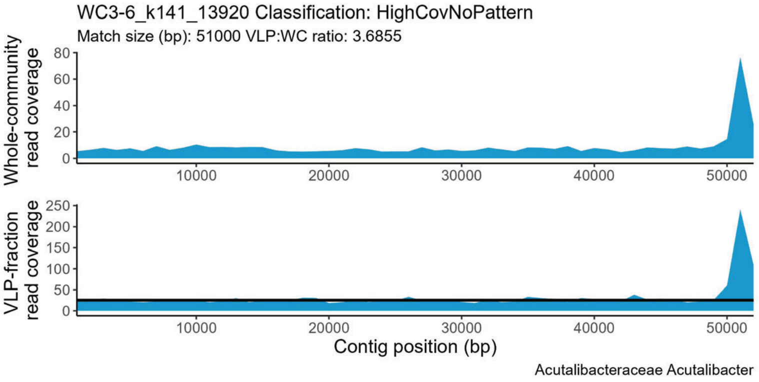
WC3-6_k141_13920 Classification: HighCovNoPattern
Match size (bp): 51000 VLP:WC ratio: 3.6855
Whole-community read coverage
0 20 40 60 80
10000 20000 30000 40000 50000
VLP-fraction read coverage
0 50 100 150 200 250
10000 20000 30000 40000 50000
Contig position (bp)
Acutalibacteraceae Acutalibacter

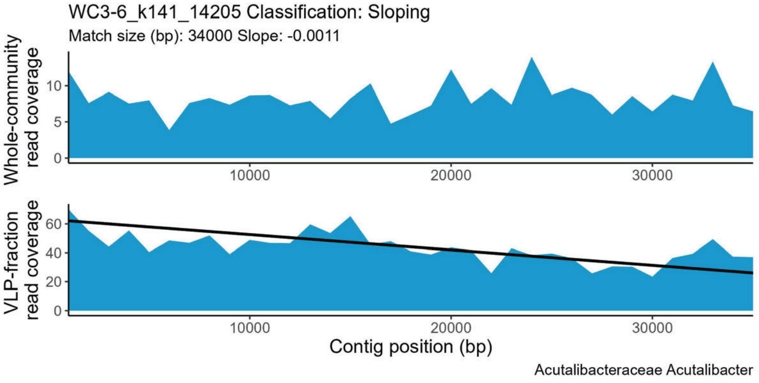
WC3-6_k141_14205 Classification: Sloping
Match size (bp): 34000 Slope: -0.0011
Whole-community read coverage
0 5 10
10000 20000 30000
VLP-fraction read coverage
0 20 40 60
10000 20000 30000
Contig position (bp)
Acutalibacteraceae Acutalibacter

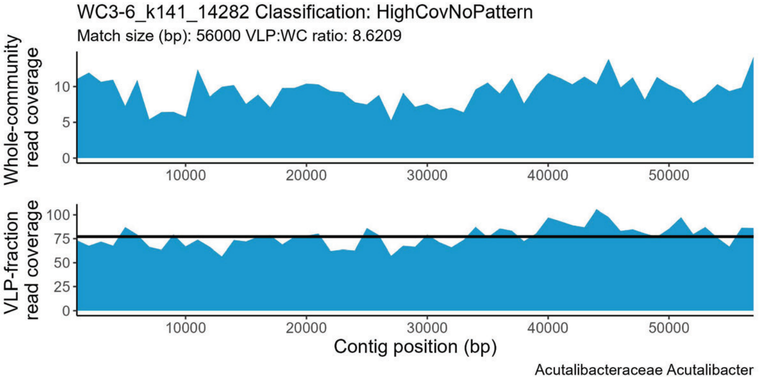
WC3-6_k141_14282 Classification: HighCovNoPattern
Match size (bp): 56000 VLP:WC ratio: 8.6209
Whole-community read coverage
0 5 10
10000 20000 30000 40000 50000
VLP-fraction read coverage
0 25 50 75 100
10000 20000 30000 40000 50000
Contig position (bp)
Acutalibacteraceae Acutalibacter

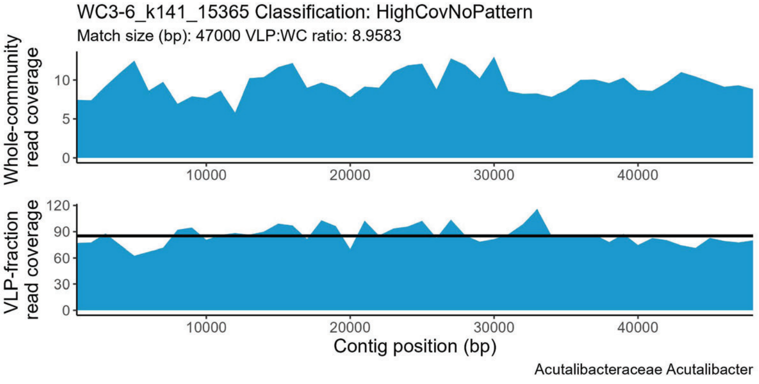
WC3-6_k141_15365 Classification: HighCovNoPattern
Match size (bp): 47000 VLP:WC ratio: 8.9583
Whole-community read coverage
0 5 10
10000 20000 30000 40000
VLP-fraction read coverage
0 30 60 90 120
10000 20000 30000 40000
Contig position (bp)
Acutalibacteraceae Acutalibacter

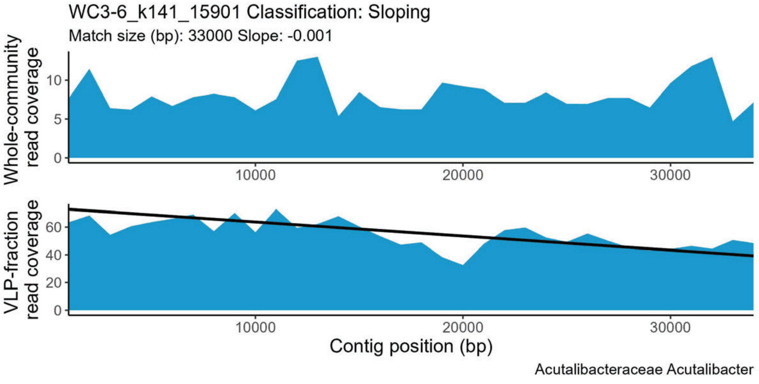
WC3-6_k141_15901 Classification: Sloping
Match size (bp): 33000 Slope: -0.001
Whole-community read coverage
0 5 10
10000 20000 30000
VLP-fraction read coverage
0 20 40 60
10000 20000 30000
Contig position (bp)
Acutalibacteraceae Acutalibacter

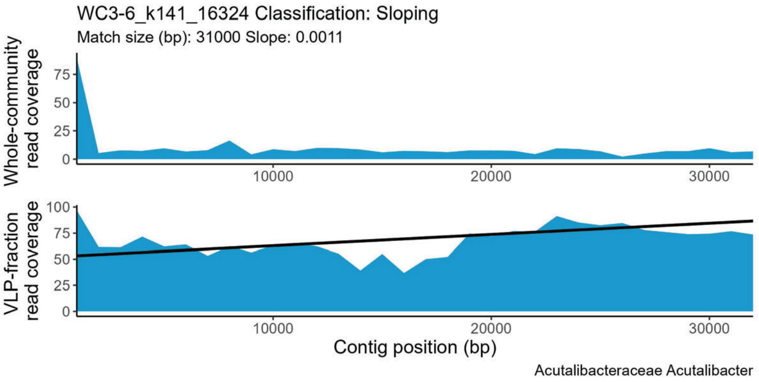
WC3-6_k141_16324 Classification: Sloping
Match size (bp): 31000 Slope: 0.0011
Whole-community read coverage
0 25 50 75
VLP-fraction read coverage
0 25 50 75 100
10000 20000 30000
Contig position (bp)
Acutalibacteraceae Acutalibacter

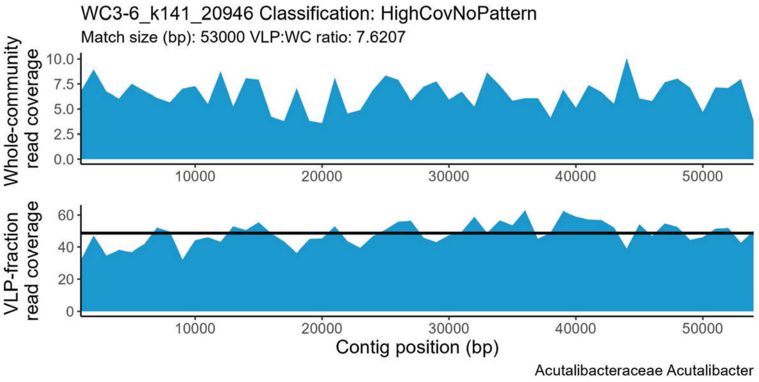
WC3-6_k141_20946 Classification: HighCovNoPattern
Match size (bp): 53000 VLP:WC ratio: 7.6207
Whole-community read coverage
0.0 2.5 5.0 7.5 10.0
VLP-fraction read coverage
0 20 40 60
10000 20000 30000 40000 50000
Contig position (bp)
Acutalibacteraceae Acutalibacter

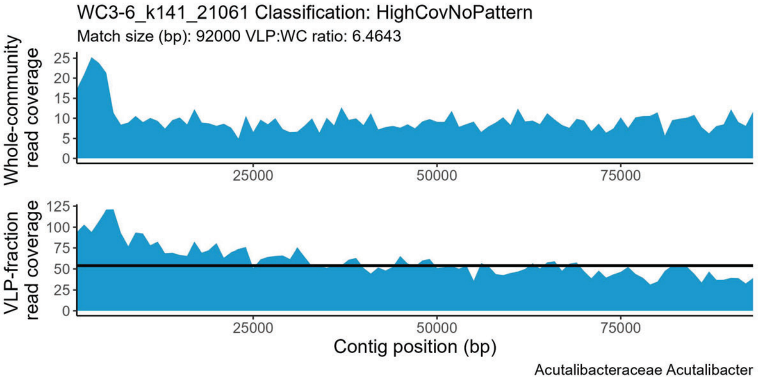
WC3-6_k141_21061 Classification: HighCovNoPattern
Match size (bp): 92000 VLP:WC ratio: 6.4643
Whole-community read coverage
0 5 10 15 20 25
VLP-fraction read coverage
0 25 50 75 100 125
25000 50000 75000
Contig position (bp)
Acutalibacteraceae Acutalibacter

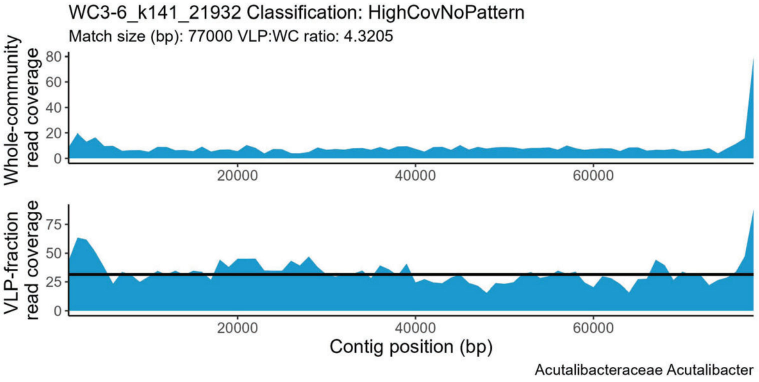
WC3-6_k141_21932 Classification: HighCovNoPattern
Match size (bp): 77000 VLP:WC ratio: 4.3205
Whole-community read coverage
0 20 40 60 80
VLP-fraction read coverage
0 25 50 75
20000 40000 60000
Contig position (bp)
Acutalibacteraceae Acutalibacter

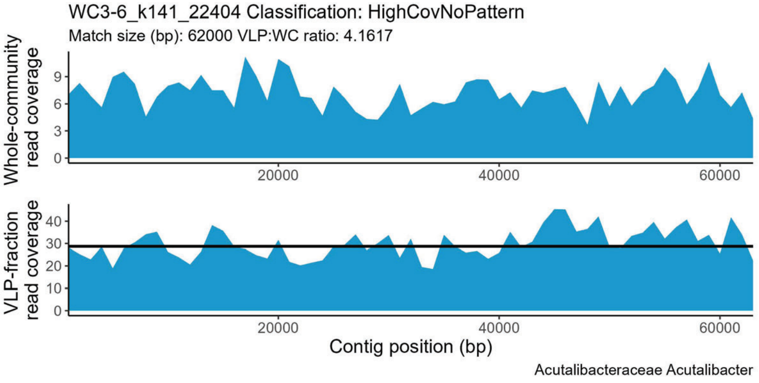
WC3-6_k141_22404 Classification: HighCovNoPattern
Match size (bp): 62000 VLP:WC ratio: 4.1617
Whole-community read coverage
0 3 6 9
VLP-fraction read coverage
0 10 20 30 40
20000 40000 60000
Contig position (bp)
Acutalibacteraceae Acutalibacter

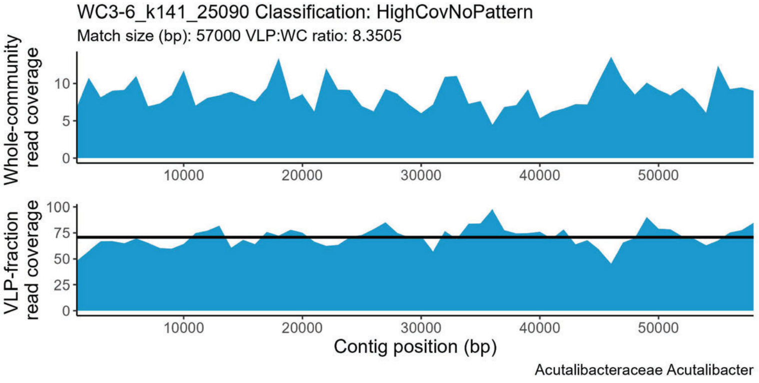
WC3-6_k141_25090 Classification: HighCovNoPattern
Match size (bp): 57000 VLP:WC ratio: 8.3505
Whole-community read coverage
0 5 10
VLP-fraction read coverage
0 25 50 75 100
10000 20000 30000 40000 50000
Contig position (bp)
Acutalibacteraceae Acutalibacter

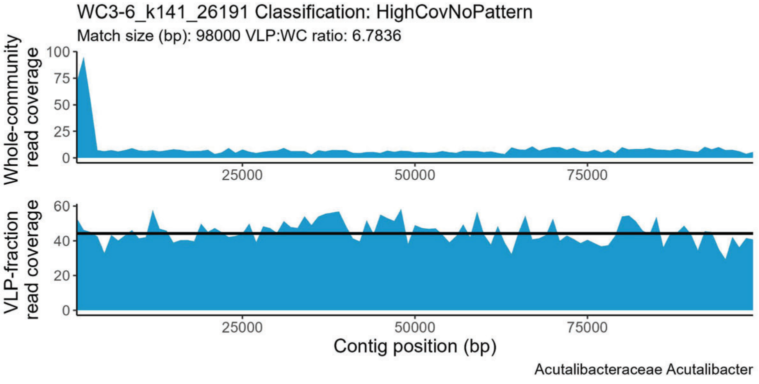
WC3-6_k141_26191 Classification: HighCovNoPattern
Match size (bp): 98000 VLP:WC ratio: 6.7836
Whole-community read coverage
0 25 50 75 100
VLP-fraction read coverage
0 20 40 60
25000 50000 75000
Contig position (bp)
Acutalibacteraceae Acutalibacter

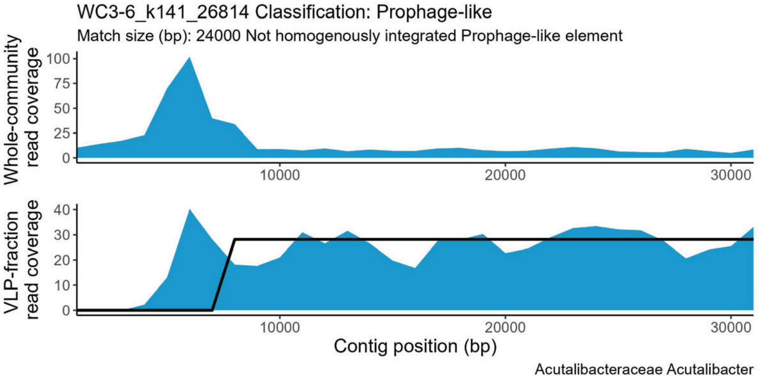
WC3-6_k141_26814 Classification: Prophage-like
Match size (bp): 24000 Not homogenously integrated Prophage-like element
Whole-community read coverage
0 25 50 75 100
VLP-fraction read coverage
0 10 20 30 40
10000 20000 30000
Contig position (bp)
Acutalibacteraceae Acutalibacter

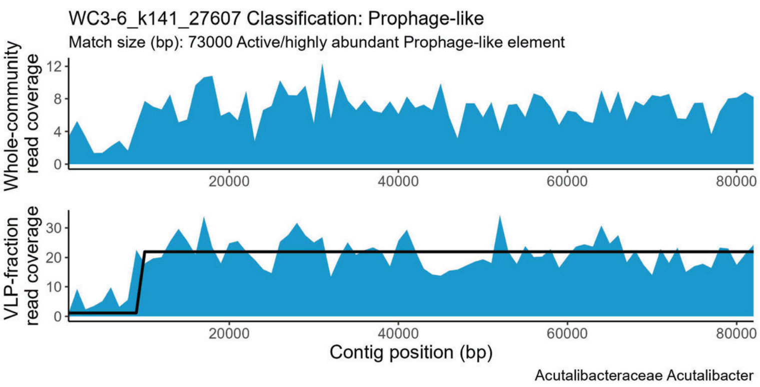
WC3-6_k141_27607 Classification: Prophage-like
Match size (bp): 73000 Active/highly abundant Prophage-like element
Whole-community read coverage
VLP-fraction read coverage
Contig position (bp)
Acutalibacteraceae Acutalibacter

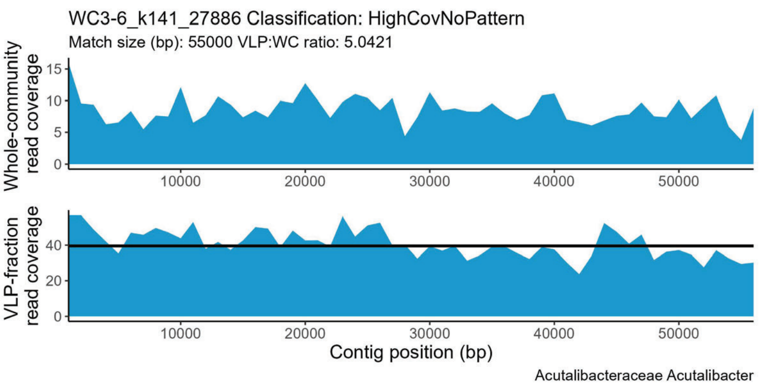
WC3-6_k141_27886 Classification: HighCovNoPattern
Match size (bp): 55000 VLP:WC ratio: 5.0421
Whole-community read coverage
VLP-fraction read coverage
Contig position (bp)
Acutalibacteraceae Acutalibacter

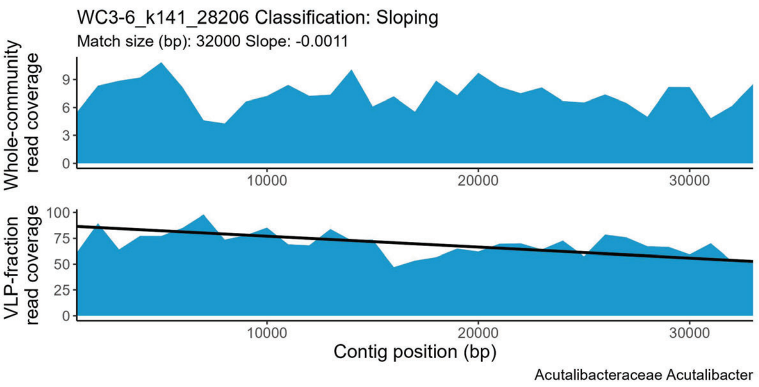
WC3-6_k141_28206 Classification: Sloping
Match size (bp): 32000 Slope: -0.0011
Whole-community read coverage
VLP-fraction read coverage
Contig position (bp)
Acutalibacteraceae Acutalibacter

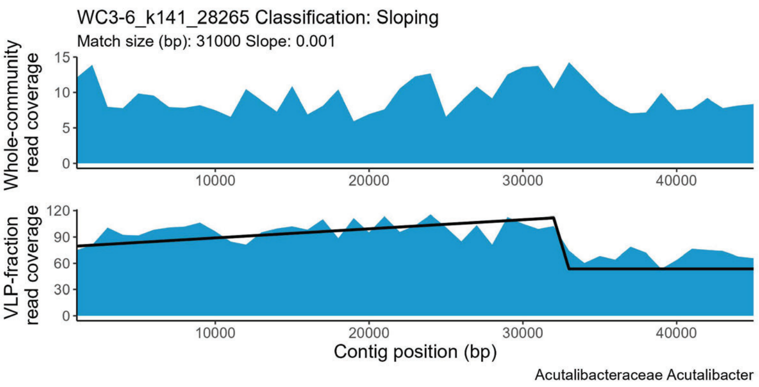
WC3-6_k141_28265 Classification: Sloping
Match size (bp): 31000 Slope: 0.001
Whole-community read coverage
VLP-fraction read coverage
Contig position (bp)
Acutalibacteraceae Acutalibacter

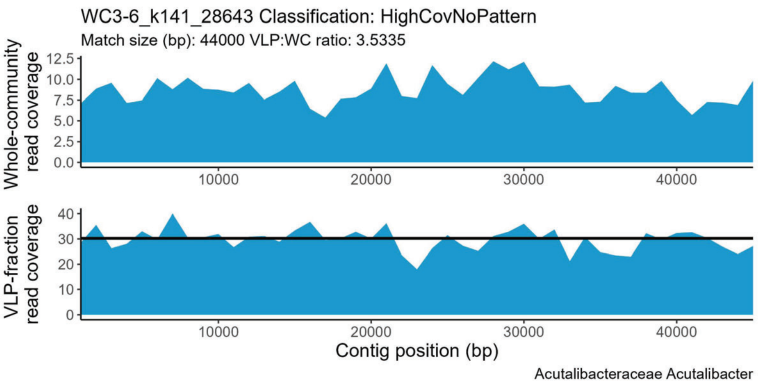
WC3-6_k141_28643 Classification: HighCovNoPattern
Match size (bp): 44000 VLP:WC ratio: 3.5335
Whole-community read coverage
VLP-fraction read coverage
Contig position (bp)
Acutalibacteraceae Acutalibacter

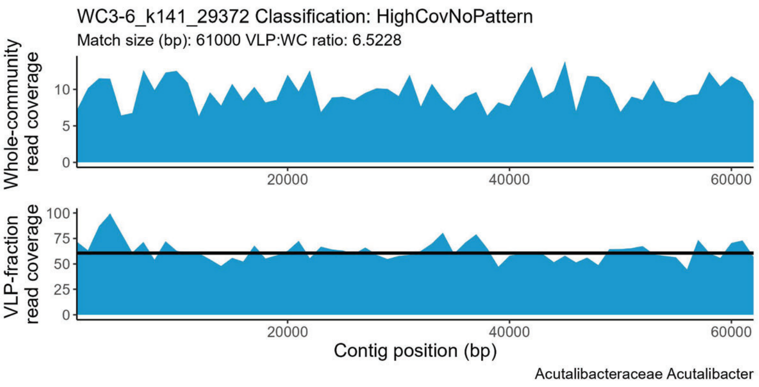
WC3-6_k141_29372 Classification: HighCovNoPattern
Match size (bp): 61000 VLP:WC ratio: 6.5228
Whole-community read coverage
VLP-fraction read coverage
Contig position (bp)
Acutalibacteraceae Acutalibacter

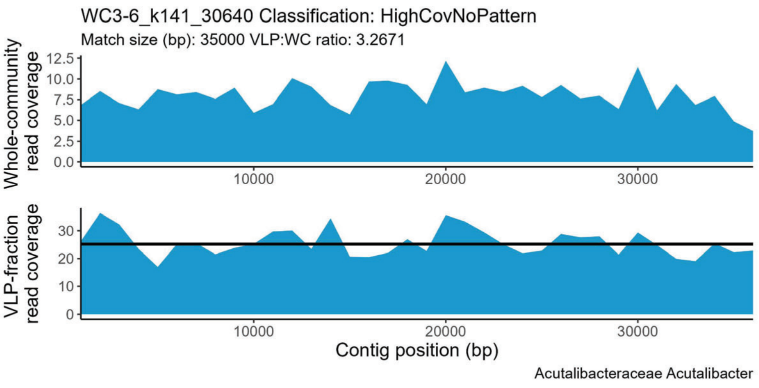
WC3-6_k141_30640 Classification: HighCovNoPattern
Match size (bp): 35000 VLP:WC ratio: 3.2671
Whole-community read coverage
VLP-fraction read coverage
Contig position (bp)
Acutalibacteraceae Acutalibacter

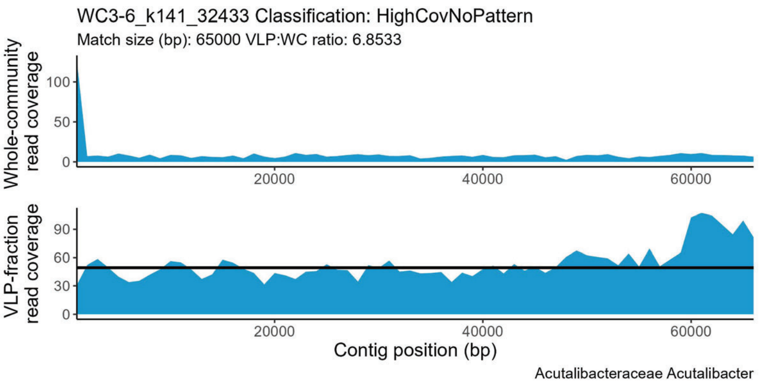
WC3-6_k141_32433 Classification: HighCovNoPattern
Match size (bp): 65000 VLP:WC ratio: 6.8533
Whole-community read coverage
VLP-fraction read coverage
Contig position (bp)
Acutalibacteraceae Acutalibacter

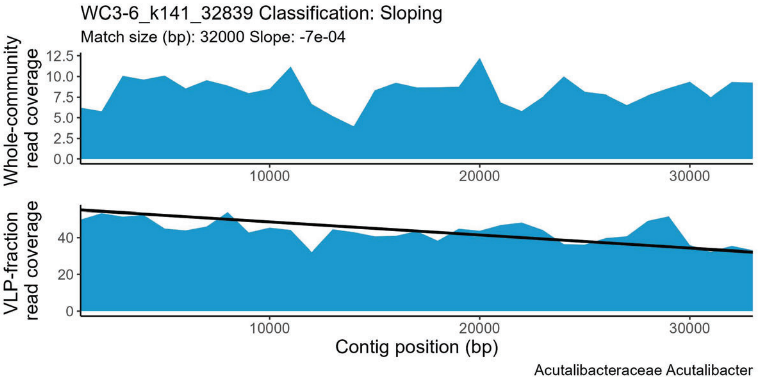
WC3-6_k141_32839 Classification: Sloping
Match size (bp): 32000 Slope: -7e-04
Whole-community read coverage
VLP-fraction read coverage
Contig position (bp)
Acutalibacteraceae Acutalibacter

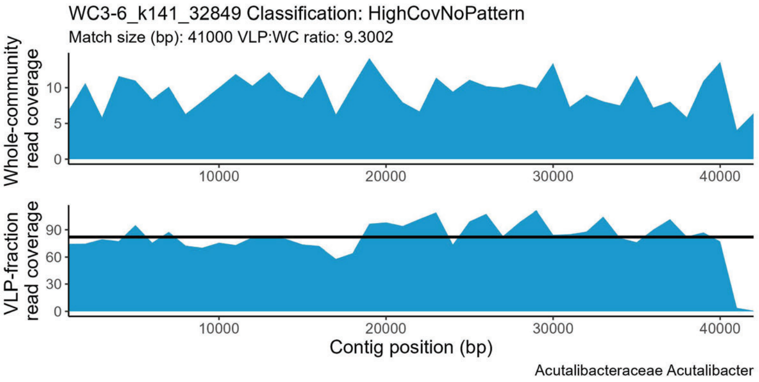
WC3-6_k141_32849 Classification: HighCovNoPattern
Match size (bp): 41000 VLP:WC ratio: 9.3002
Whole-community read coverage
VLP-fraction read coverage
Contig position (bp)
Acutalibacteraceae Acutalibacter

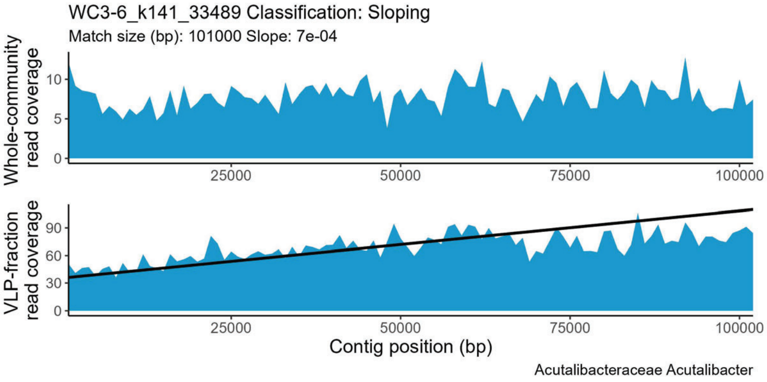
WC3-6_k141_33489 Classification: Sloping
Match size (bp): 101000 Slope: 7e-04
Whole-community read coverage
VLP-fraction read coverage
Contig position (bp)
Acutalibacteraceae Acutalibacter

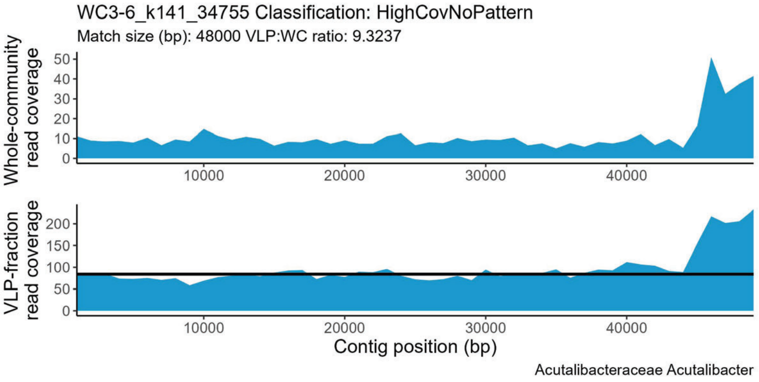
WC3-6_k141_34755 Classification: HighCovNoPattern
Match size (bp): 48000 VLP:WC ratio: 9.3237
Whole-community read coverage
VLP-fraction read coverage
Contig position (bp)
Acutalibacteraceae Acutalibacter

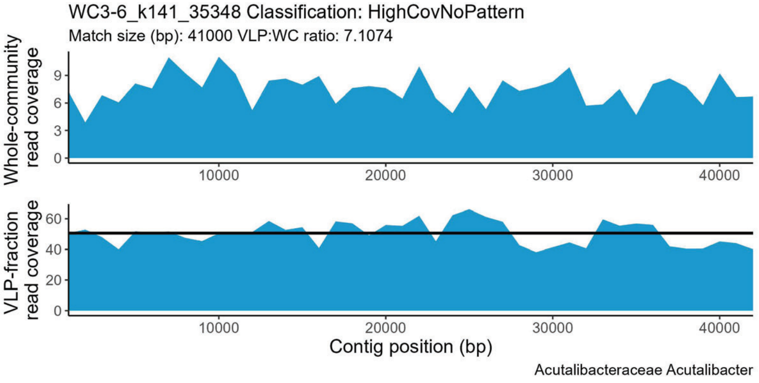
WC3-6_k141_35348 Classification: HighCovNoPattern
Match size (bp): 41000 VLP:WC ratio: 7.1074
Whole-community read coverage
VLP-fraction read coverage
Contig position (bp)
Acutalibacteraceae Acutalibacter

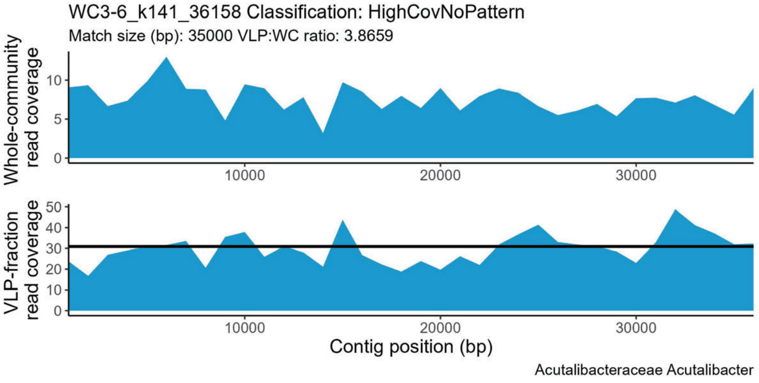
WC3-6_k141_36158 Classification: HighCovNoPattern
Match size (bp): 35000 VLP:WC ratio: 3.8659
Whole-community read coverage
VLP-fraction read coverage
Contig position (bp)
Acutalibacteraceae Acutalibacter

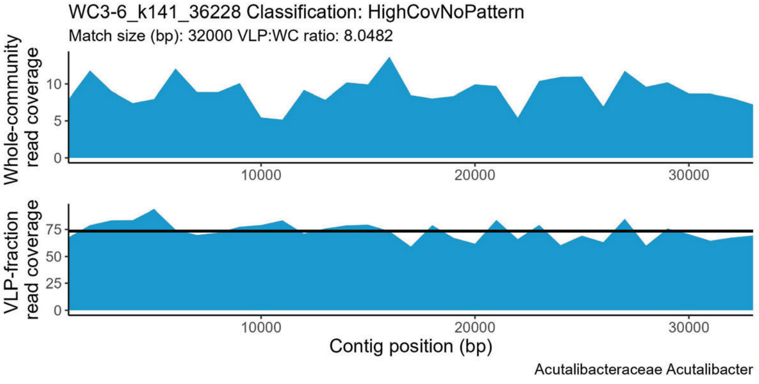
WC3-6_k141_36228 Classification: HighCovNoPattern
Match size (bp): 32000 VLP:WC ratio: 8.0482
Whole-community read coverage
VLP-fraction read coverage
Contig position (bp)
Acutalibacteraceae Acutalibacter

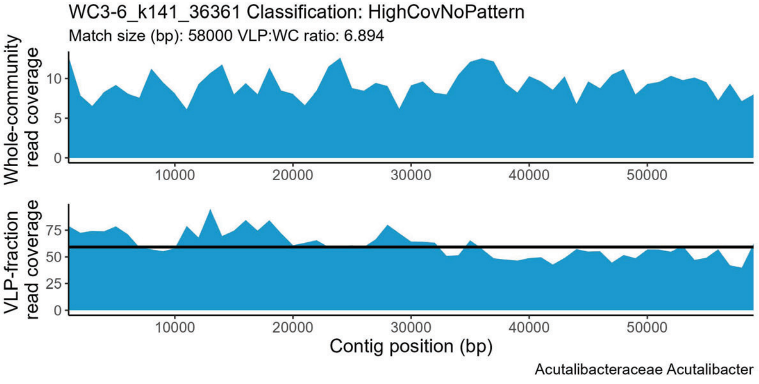
WC3-6_k141_36361 Classification: HighCovNoPattern
Match size (bp): 58000 VLP:WC ratio: 6.894
Whole-community read coverage
VLP-fraction read coverage
Contig position (bp)
Acutalibacteraceae Acutalibacter

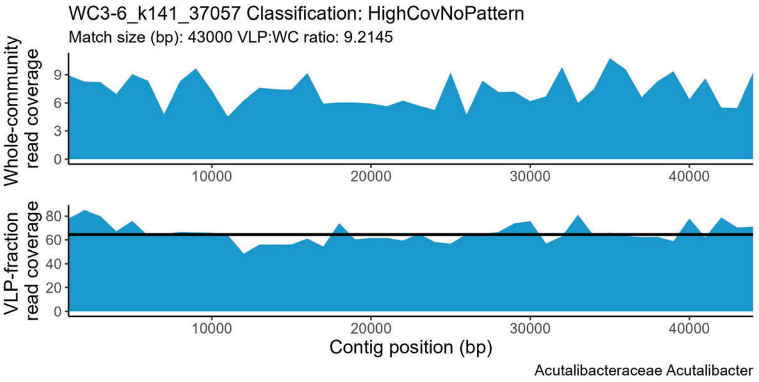
WC3-6_k141_37057 Classification: HighCovNoPattern
Match size (bp): 43000 VLP:WC ratio: 9.2145
Whole-community read coverage
VLP-fraction read coverage
Contig position (bp)
Acutalibacteraceae Acutalibacter

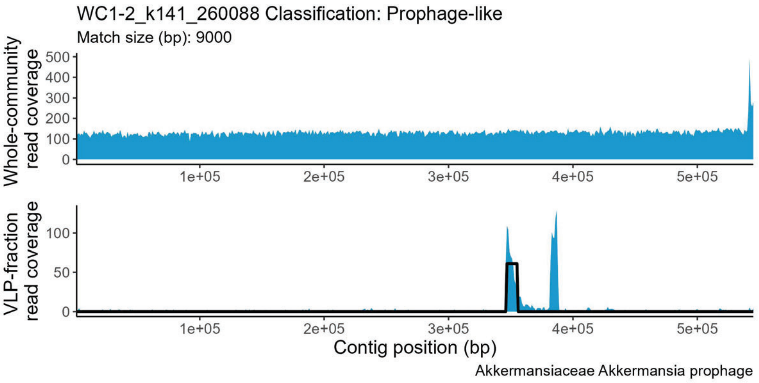
WC1-2_k141_260088 Classification: Prophage-like
Match size (bp): 9000
Whole-community read coverage
VLP-fraction read coverage
Contig position (bp)
Akkermansiaceae Akkermansia prophage

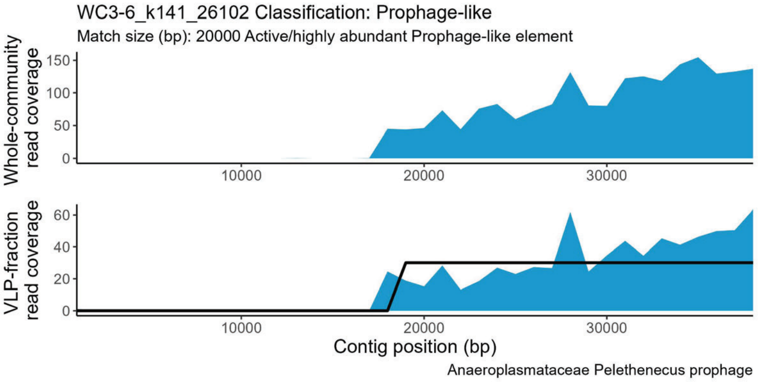
WC3-6_k141_26102 Classification: Prophage-like
Match size (bp): 20000 Active/highly abundant Prophage-like element
Whole-community read coverage
VLP-fraction read coverage
Contig position (bp)
Anaeroplasmataceae Pelethenecus prophage

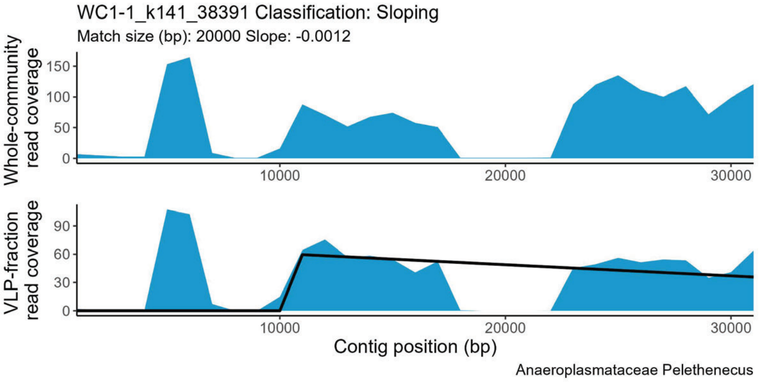
WC1-1_k141_38391 Classification: Sloping
Match size (bp): 20000 Slope: -0.0012
Whole-community read coverage
VLP-fraction read coverage
Contig position (bp)
Anaeroplasmataceae Pelethenecus

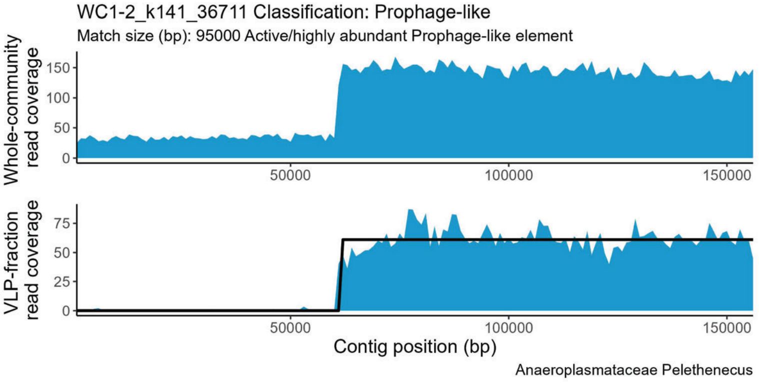
WC1-2_k141_36711 Classification: Prophage-like
Match size (bp): 95000 Active/highly abundant Prophage-like element
Whole-community read coverage
VLP-fraction read coverage
Contig position (bp)
Anaeroplasmataceae Pelethenecus

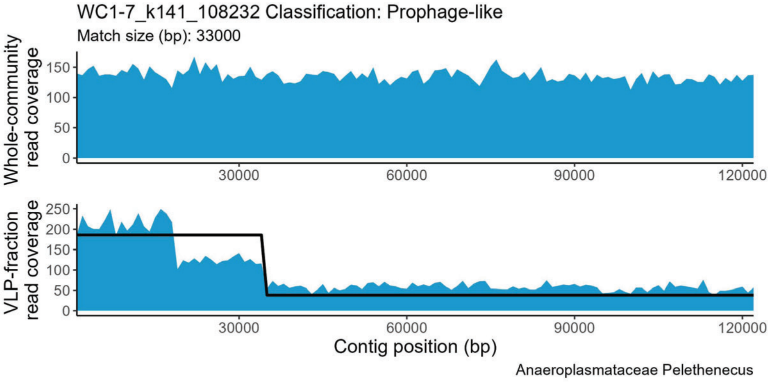
WC1-7_k141_108232 Classification: Prophage-like
Match size (bp): 33000
Whole-community read coverage
VLP-fraction read coverage
Contig position (bp)
Anaeroplasmataceae Pelethenecus

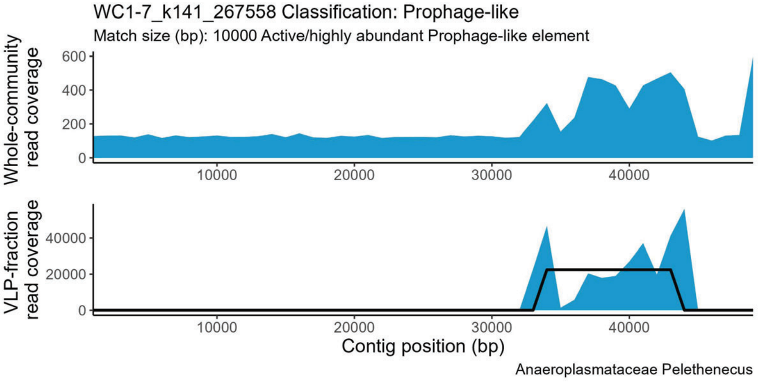
WC1-7_k141_267558 Classification: Prophage-like
Match size (bp): 10000 Active/highly abundant Prophage-like element
Whole-community read coverage
VLP-fraction read coverage
Contig position (bp)
Anaeroplasmataceae Pelethenecus

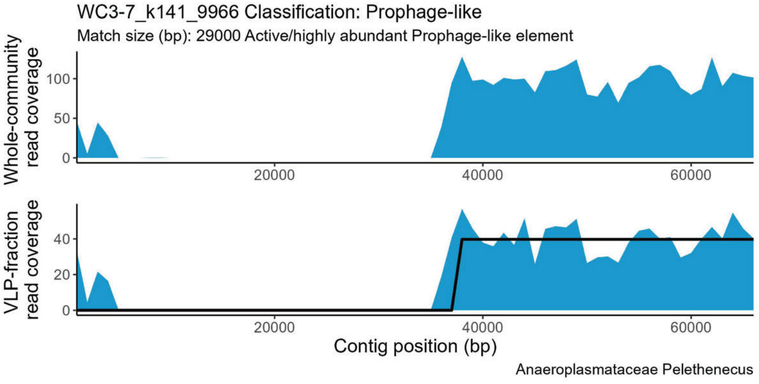
WC3-7_k141_9966 Classification: Prophage-like
Match size (bp): 29000 Active/highly abundant Prophage-like element
Whole-community read coverage
VLP-fraction read coverage
Contig position (bp)
Anaeroplasmataceae Pelethenecus

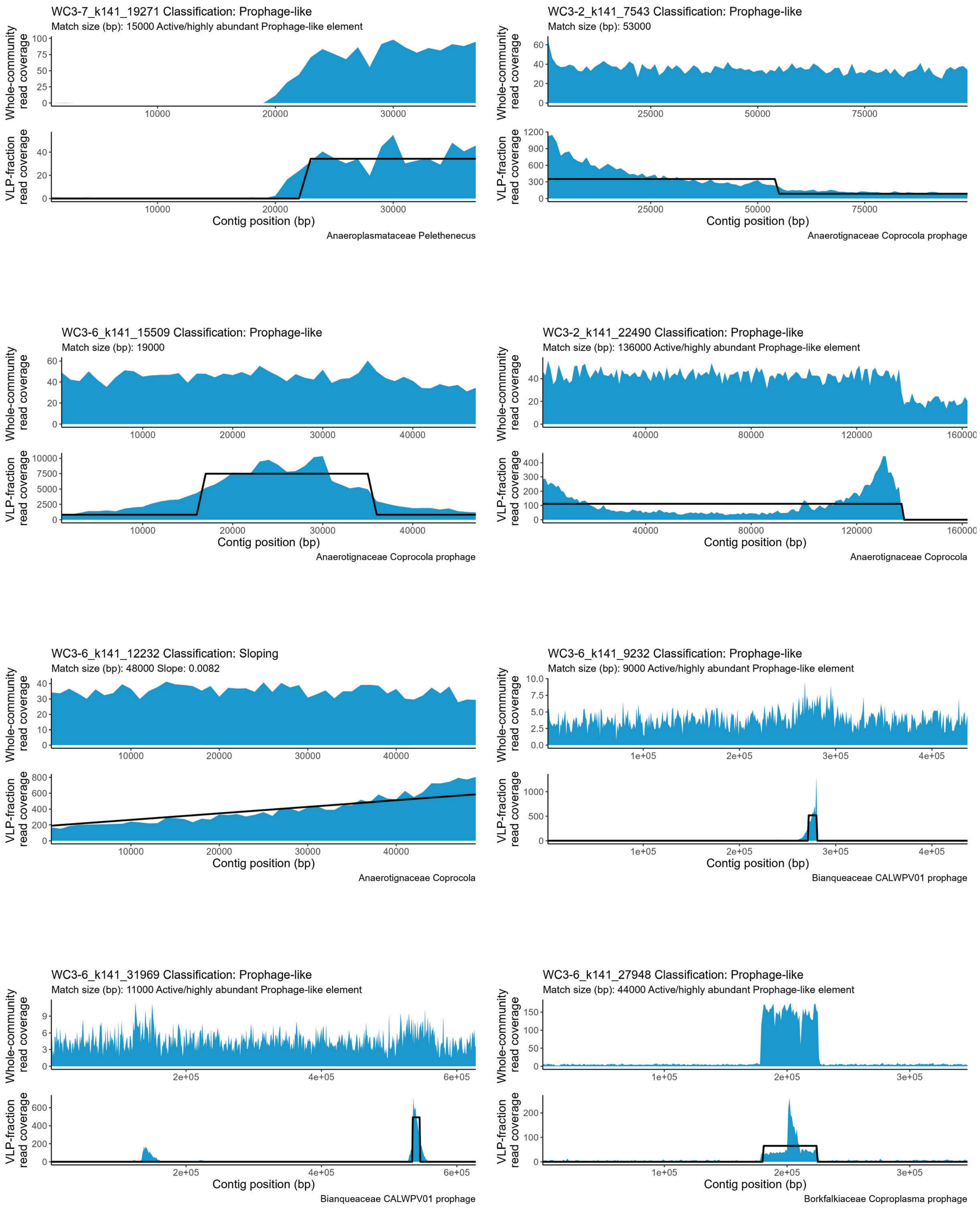

WC3-7_k141_19271 Classification: Prophage-like
Match size (bp): 15000 Active/highly abundant Prophage-like element
Whole-community read coverage
VLP-fraction read coverage
Contig position (bp)
Anaeroplasmataceae Pelethenecus
WC3-2_k141_7543 Classification: Prophage-like
Match size (bp): 53000
Whole-community read coverage
VLP-fraction read coverage
Contig position (bp)
Anaerotignaceae Coprocola prophage
WC3-6_k141_15509 Classification: Prophage-like
Match size (bp): 19000
Whole-community read coverage
VLP-fraction read coverage
Contig position (bp)
Anaerotignaceae Coprocola prophage
WC3-2_k141_22490 Classification: Prophage-like
Match size (bp): 136000 Active/highly abundant Prophage-like element
Whole-community read coverage
VLP-fraction read coverage
Contig position (bp)
Anaerotignaceae Coprocola
WC3-6_k141_12232 Classification: Sloping
Match size (bp): 48000 Slope: 0.0082
Whole-community read coverage
VLP-fraction read coverage
Contig position (bp)
Anaerotignaceae Coprocola
WC3-6_k141_9232 Classification: Prophage-like
Match size (bp): 9000 Active/highly abundant Prophage-like element
Whole-community read coverage
VLP-fraction read coverage
Contig position (bp)
Bianqueaceae CALWPV01 prophage
WC3-6_k141_31969 Classification: Prophage-like
Match size (bp): 11000 Active/highly abundant Prophage-like element
Whole-community read coverage
VLP-fraction read coverage
Contig position (bp)
Bianqueaceae CALWPV01 prophage
WC3-6_k141_27948 Classification: Prophage-like
Match size (bp): 44000 Active/highly abundant Prophage-like element
Whole-community read coverage
VLP-fraction read coverage
Contig position (bp)
Borkfalkiaceae Coproplasma prophage

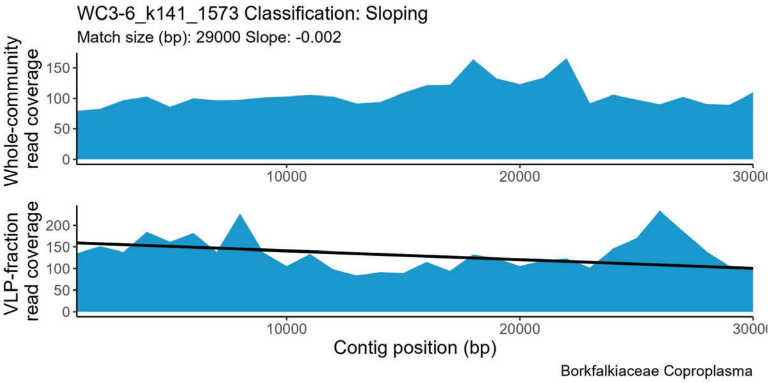
WC3-6_k141_1573 Classification: Sloping
Match size (bp): 29000 Slope: -0.002
Whole-community read coverage
VLP-fraction read coverage
Contig position (bp)
Borkfalkiaceae Coproplasma

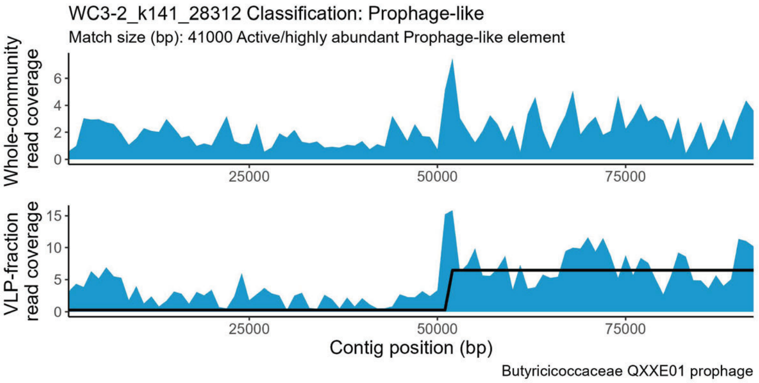
WC3-2_k141_28312 Classification: Prophage-like
Match size (bp): 41000 Active/highly abundant Prophage-like element
Whole-community read coverage
VLP-fraction read coverage
Contig position (bp)
Butyricicoccaceae QXXE01 prophage

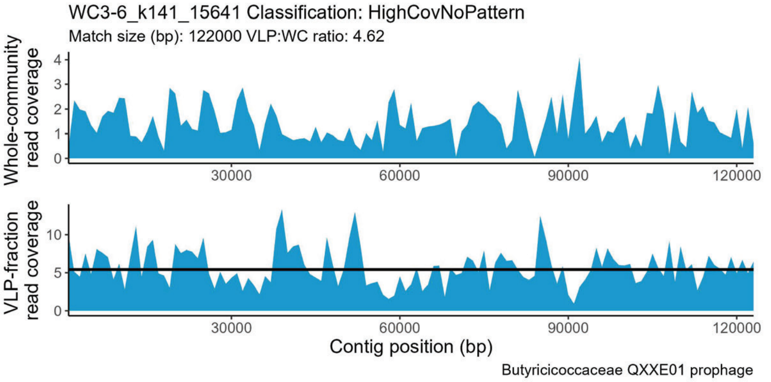
WC3-6_k141_15641 Classification: HighCovNoPattern
Match size (bp): 122000 VLP:WC ratio: 4.62
Whole-community read coverage
VLP-fraction read coverage
Contig position (bp)
Butyricicoccaceae QXXE01 prophage

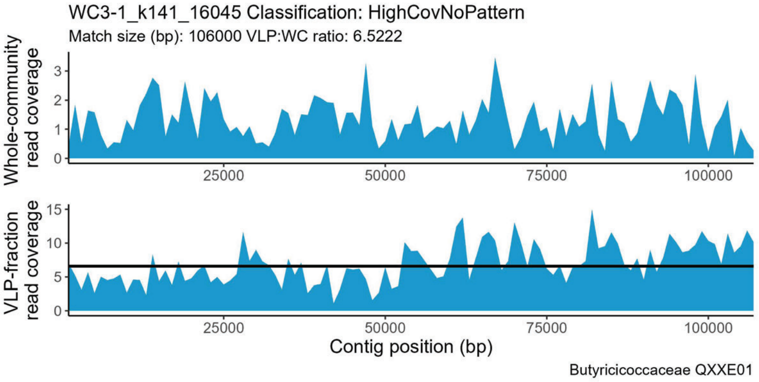
WC3-1_k141_16045 Classification: HighCovNoPattern
Match size (bp): 106000 VLP:WC ratio: 6.5222
Whole-community read coverage
VLP-fraction read coverage
Contig position (bp)
Butyricicoccaceae QXXE01

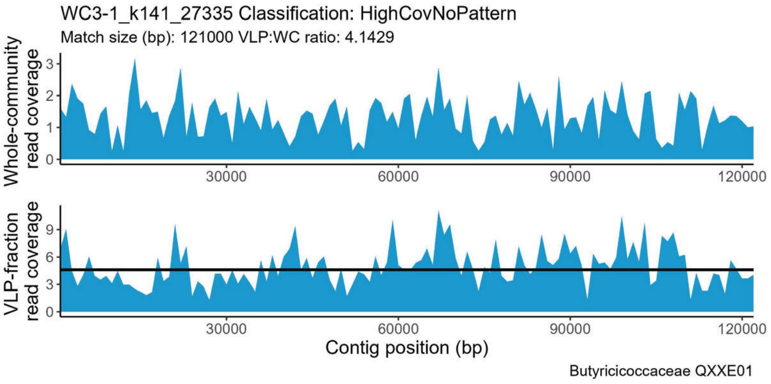
WC3-1_k141_27335 Classification: HighCovNoPattern
Match size (bp): 121000 VLP:WC ratio: 4.1429
Whole-community read coverage
VLP-fraction read coverage
Contig position (bp)
Butyricicoccaceae QXXE01

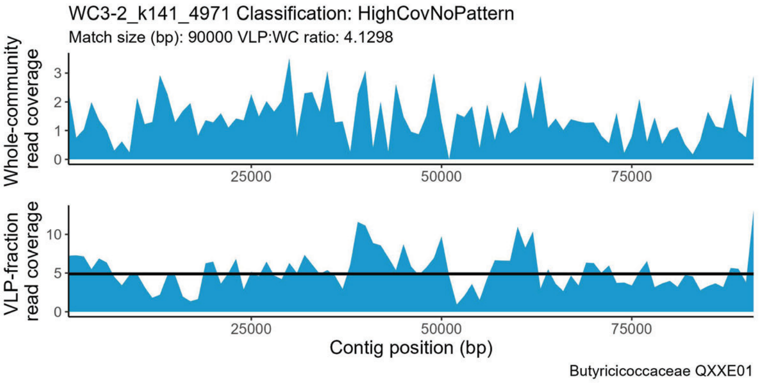
WC3-2_k141_4971 Classification: HighCovNoPattern
Match size (bp): 90000 VLP:WC ratio: 4.1298
Whole-community read coverage
VLP-fraction read coverage
Contig position (bp)
Butyricicoccaceae QXXE01

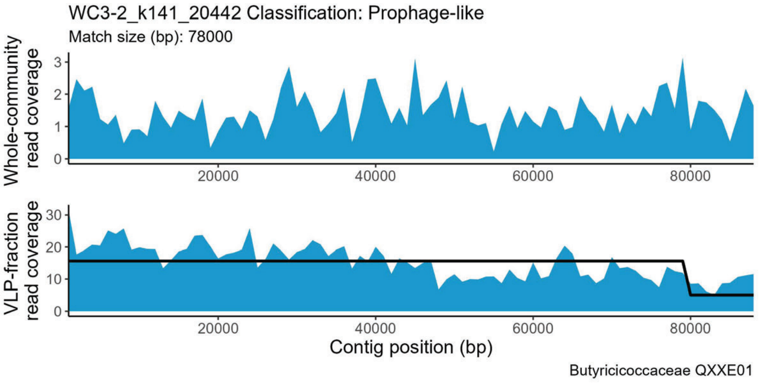
WC3-2_k141_20442 Classification: Prophage-like
Match size (bp): 78000
Whole-community read coverage
VLP-fraction read coverage
Contig position (bp)
Butyricicoccaceae QXXE01

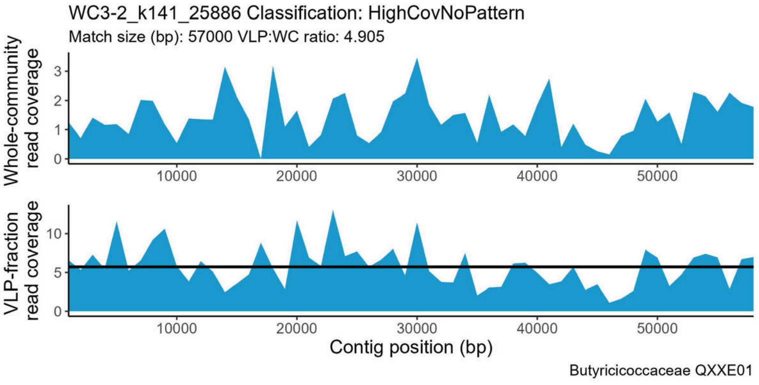
WC3-2_k141_25886 Classification: HighCovNoPattern
Match size (bp): 57000 VLP:WC ratio: 4.905
Whole-community read coverage
VLP-fraction read coverage
Contig position (bp)
Butyricicoccaceae QXXE01

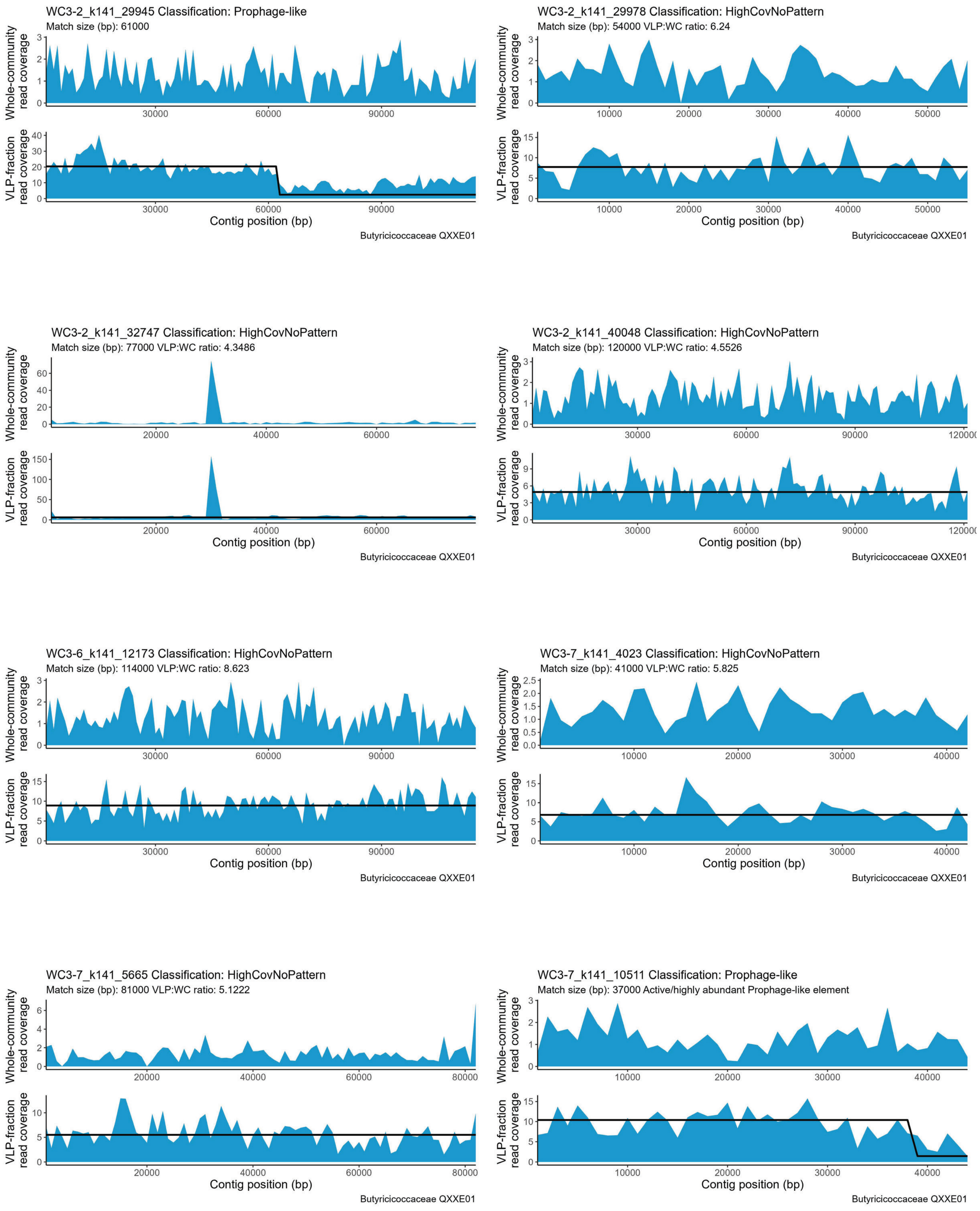
WC3-2_k141_29945 Classification: Prophage-like
Match size (bp): 61000
Whole-community read coverage
VLP-fraction read coverage
Contig position (bp)
Butyricicoccaceae QXXE01
WC3-2_k141_29978 Classification: HighCovNoPattern
Match size (bp): 54000 VLP:WC ratio: 6.24
Whole-community read coverage
VLP-fraction read coverage
Contig position (bp)
Butyricicoccaceae QXXE01
WC3-2_k141_32747 Classification: HighCovNoPattern
Match size (bp): 77000 VLP:WC ratio: 4.3486
Whole-community read coverage
VLP-fraction read coverage
Contig position (bp)
Butyricicoccaceae QXXE01
WC3-2_k141_40048 Classification: HighCovNoPattern
Match size (bp): 120000 VLP:WC ratio: 4.5526
Whole-community read coverage
VLP-fraction read coverage
Contig position (bp)
Butyricicoccaceae QXXE01
WC3-6_k141_12173 Classification: HighCovNoPattern
Match size (bp): 114000 VLP:WC ratio: 8.623
Whole-community read coverage
VLP-fraction read coverage
Contig position (bp)
Butyricicoccaceae QXXE01
WC3-7_k141_4023 Classification: HighCovNoPattern
Match size (bp): 41000 VLP:WC ratio: 5.825
Whole-community read coverage
VLP-fraction read coverage
Contig position (bp)
Butyricicoccaceae QXXE01
WC3-7_k141_5665 Classification: HighCovNoPattern
Match size (bp): 81000 VLP:WC ratio: 5.1222
Whole-community read coverage
VLP-fraction read coverage
Contig position (bp)
Butyricicoccaceae QXXE01
WC3-7_k141_10511 Classification: Prophage-like
Match size (bp): 37000 Active/highly abundant Prophage-like element
Whole-community read coverage
VLP-fraction read coverage
Contig position (bp)
Butyricicoccaceae QXXE01

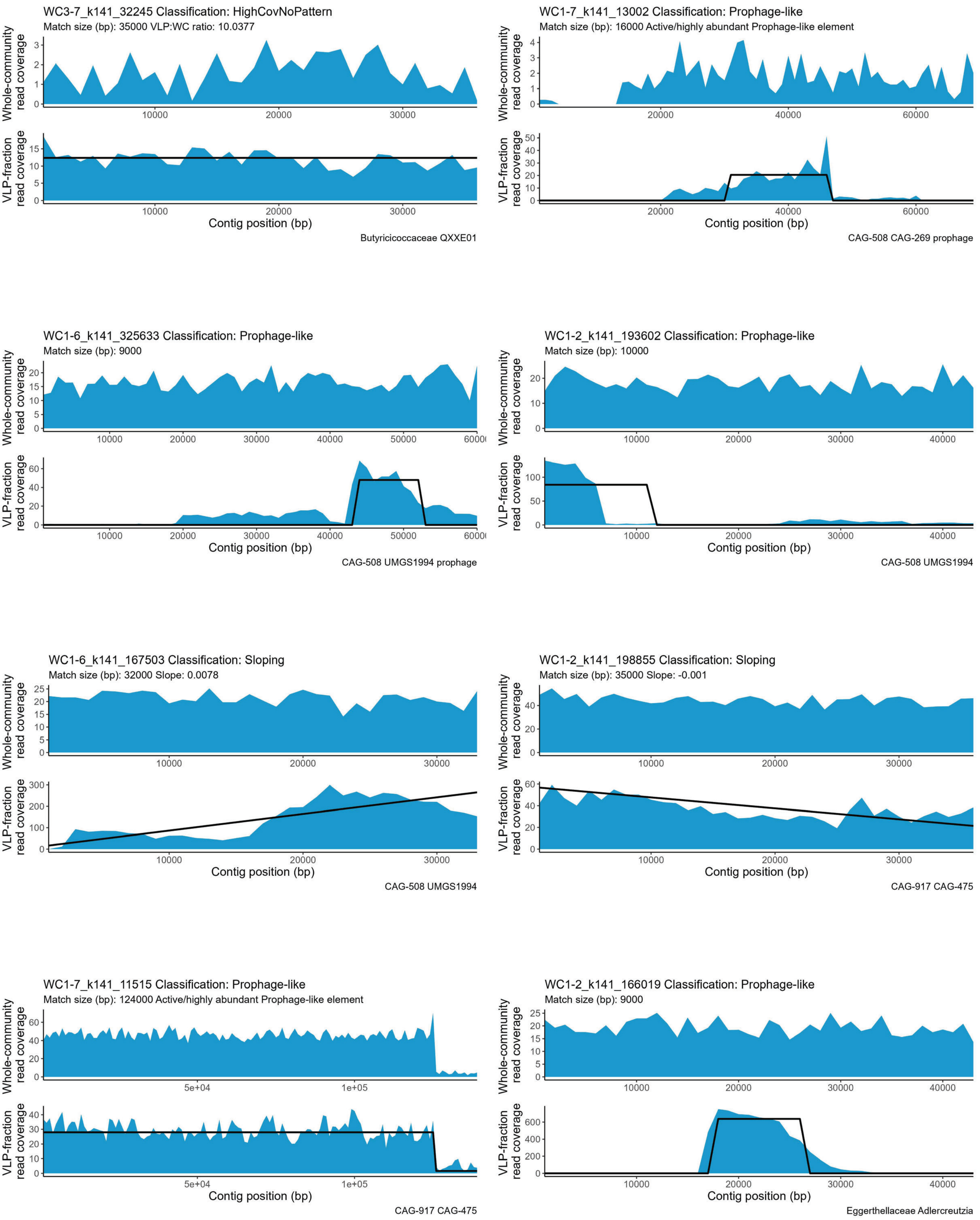

WC3-7_k141_32245 Classification: HighCovNoPattern
Match size (bp): 35000 VLP:WC ratio: 10.0377
Whole-community read coverage
VLP-fraction read coverage
Contig position (bp)
Butyricicoccaceae QXXE01
WC1-7_k141_13002 Classification: Prophage-like
Match size (bp): 16000 Active/highly abundant Prophage-like element
Whole-community read coverage
VLP-fraction read coverage
Contig position (bp)
CAG-508 CAG-269 prophage
WC1-6_k141_325633 Classification: Prophage-like
Match size (bp): 9000
Whole-community read coverage
VLP-fraction read coverage
Contig position (bp)
CAG-508 UMGS1994 prophage
WC1-2_k141_193602 Classification: Prophage-like
Match size (bp): 10000
Whole-community read coverage
VLP-fraction read coverage
Contig position (bp)
CAG-508 UMGS1994
WC1-6_k141_167503 Classification: Sloping
Match size (bp): 32000 Slope: 0.0078
Whole-community read coverage
VLP-fraction read coverage
Contig position (bp)
CAG-508 UMGS1994
WC1-2_k141_198855 Classification: Sloping
Match size (bp): 35000 Slope: -0.001
Whole-community read coverage
VLP-fraction read coverage
Contig position (bp)
CAG-917 CAG-475
WC1-7_k141_11515 Classification: Prophage-like
Match size (bp): 124000 Active/highly abundant Prophage-like element
Whole-community read coverage
VLP-fraction read coverage
Contig position (bp)
CAG-917 CAG-475
WC1-2_k141_166019 Classification: Prophage-like
Match size (bp): 9000
Whole-community read coverage
VLP-fraction read coverage
Contig position (bp)
Eggerthellaceae Adlercreutzia

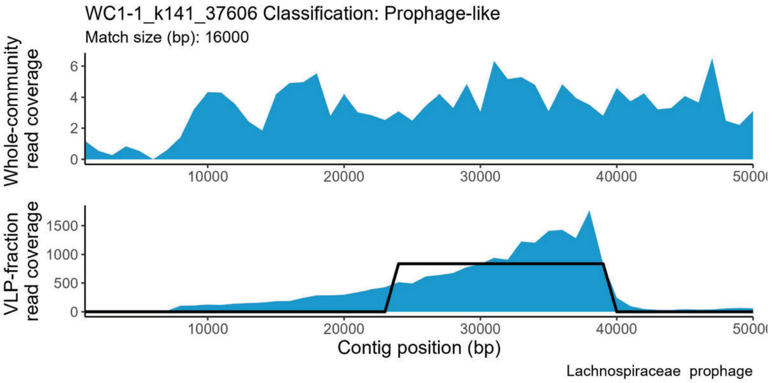
WC1-1_k141_37606 Classification: Prophage-like
Match size (bp): 16000
Whole-community read coverage
VLP-fraction read coverage
Contig position (bp)
Lachnospiraceae prophage

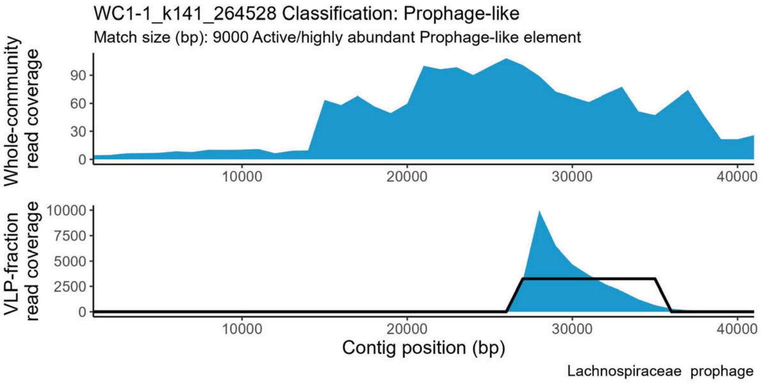
WC1-1_k141_264528 Classification: Prophage-like
Match size (bp): 9000 Active/highly abundant Prophage-like element
Whole-community read coverage
VLP-fraction read coverage
Contig position (bp)
Lachnospiraceae prophage

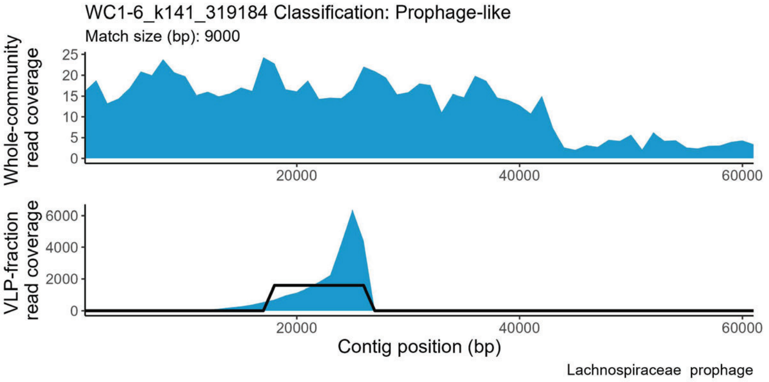
WC1-6_k141_319184 Classification: Prophage-like
Match size (bp): 9000
Whole-community read coverage
VLP-fraction read coverage
Contig position (bp)
Lachnospiraceae prophage

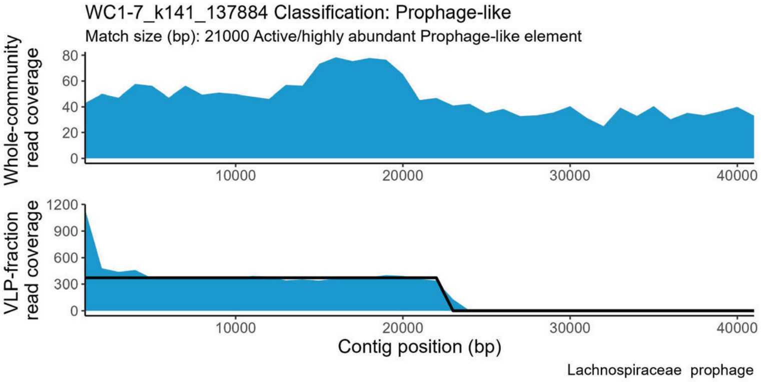
WC1-7_k141_137884 Classification: Prophage-like
Match size (bp): 21000 Active/highly abundant Prophage-like element
Whole-community read coverage
VLP-fraction read coverage
Contig position (bp)
Lachnospiraceae prophage

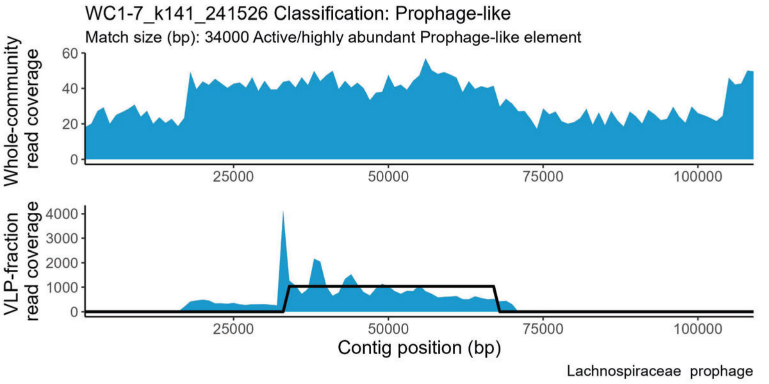
WC1-7_k141_241526 Classification: Prophage-like
Match size (bp): 34000 Active/highly abundant Prophage-like element
Whole-community read coverage
VLP-fraction read coverage
Contig position (bp)
Lachnospiraceae prophage

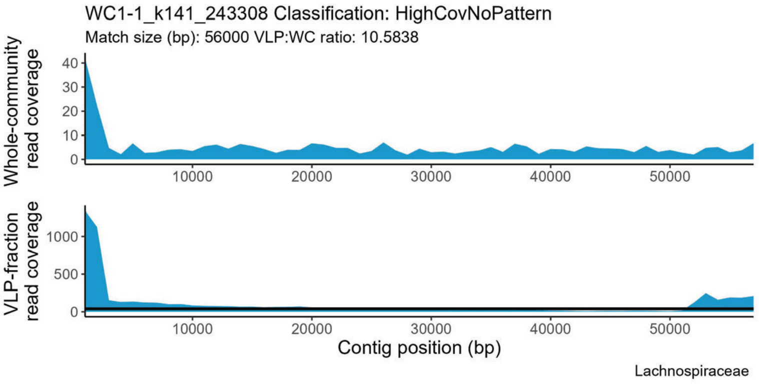
WC1-1_k141_243308 Classification: HighCovNoPattern
Match size (bp): 56000 VLP:WC ratio: 10.5838
Whole-community read coverage
VLP-fraction read coverage
Contig position (bp)
Lachnospiraceae

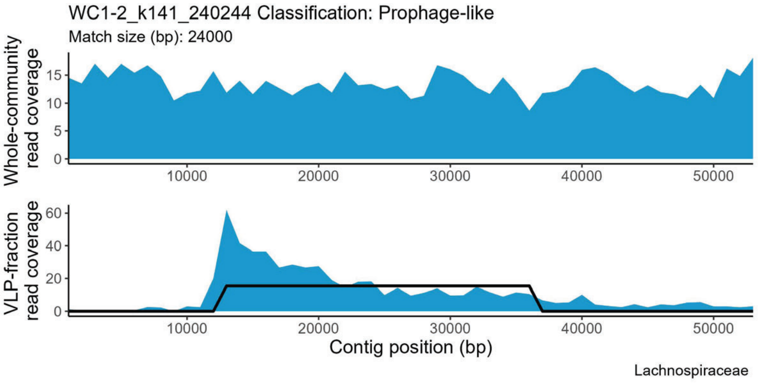
WC1-2_k141_240244 Classification: Prophage-like
Match size (bp): 24000
Whole-community read coverage
VLP-fraction read coverage
Contig position (bp)
Lachnospiraceae

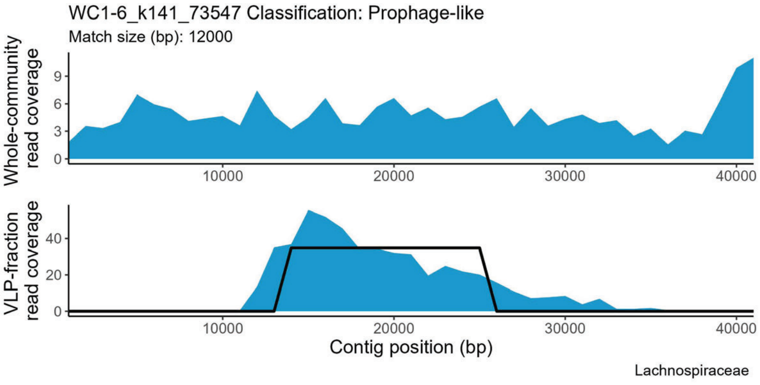
WC1-6_k141_73547 Classification: Prophage-like
Match size (bp): 12000
Whole-community read coverage
VLP-fraction read coverage
Contig position (bp)
Lachnospiraceae

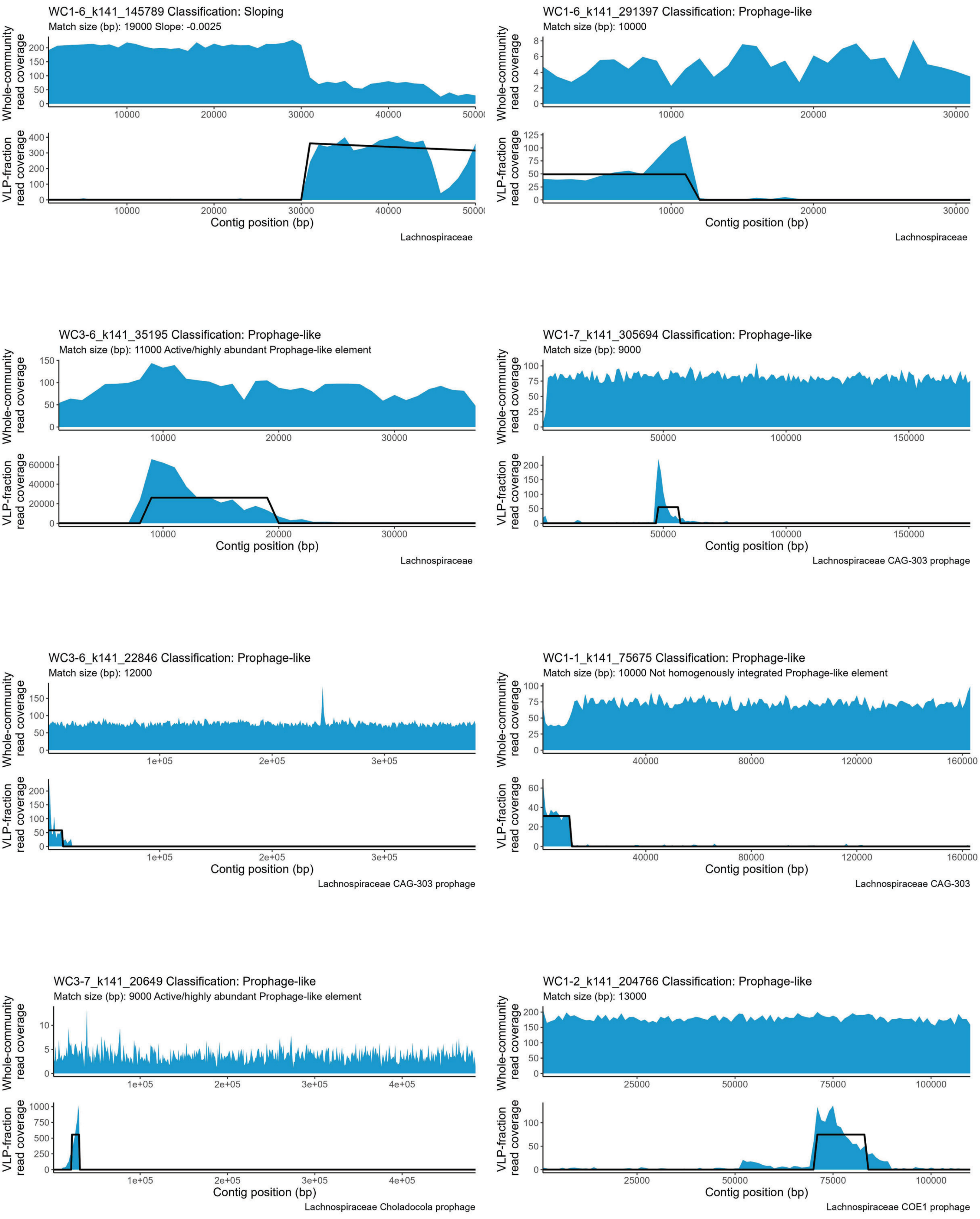

WC1-6_k141_145789 Classification: Sloping
Match size (bp): 19000 Slope: -0.0025
Whole-community read coverage
VLP-fraction read coverage
Contig position (bp)
Lachnospiraceae
WC1-6_k141_291397 Classification: Prophage-like
Match size (bp): 10000
Whole-community read coverage
VLP-fraction read coverage
Contig position (bp)
Lachnospiraceae
WC3-6_k141_35195 Classification: Prophage-like
Match size (bp): 11000 Active/highly abundant Prophage-like element
Whole-community read coverage
VLP-fraction read coverage
Contig position (bp)
Lachnospiraceae
WC1-7_k141_305694 Classification: Prophage-like
Match size (bp): 9000
Whole-community read coverage
VLP-fraction read coverage
Contig position (bp)
Lachnospiraceae CAG-303 prophage
WC3-6_k141_22846 Classification: Prophage-like
Match size (bp): 12000
Whole-community read coverage
VLP-fraction read coverage
Contig position (bp)
Lachnospiraceae CAG-303 prophage
WC1-1_k141_75675 Classification: Prophage-like
Match size (bp): 10000 Not homogenously integrated Prophage-like element
Whole-community read coverage
VLP-fraction read coverage
Contig position (bp)
Lachnospiraceae CAG-303
WC3-7_k141_20649 Classification: Prophage-like
Match size (bp): 9000 Active/highly abundant Prophage-like element
Whole-community read coverage
VLP-fraction read coverage
Contig position (bp)
Lachnospiraceae Choladocola prophage
WC1-2_k141_204766 Classification: Prophage-like
Match size (bp): 13000
Whole-community read coverage
VLP-fraction read coverage
Contig position (bp)
Lachnospiraceae COE1 prophage

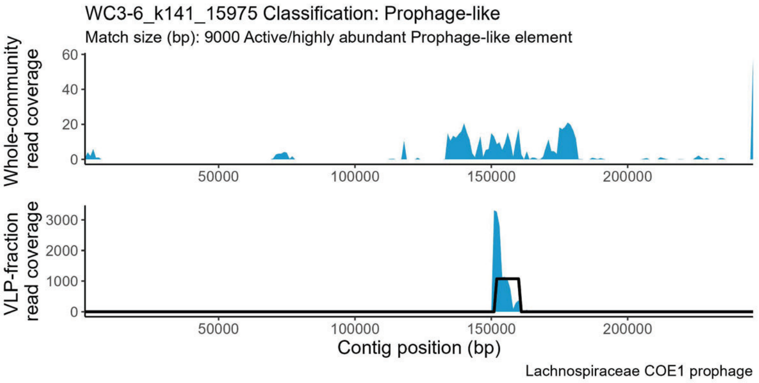
WC3-6_k141_15975 Classification: Prophage-like
Match size (bp): 9000 Active/highly abundant Prophage-like element
Whole-community read coverage
VLP-fraction read coverage
Contig position (bp)
Lachnospiraceae COE1 prophage

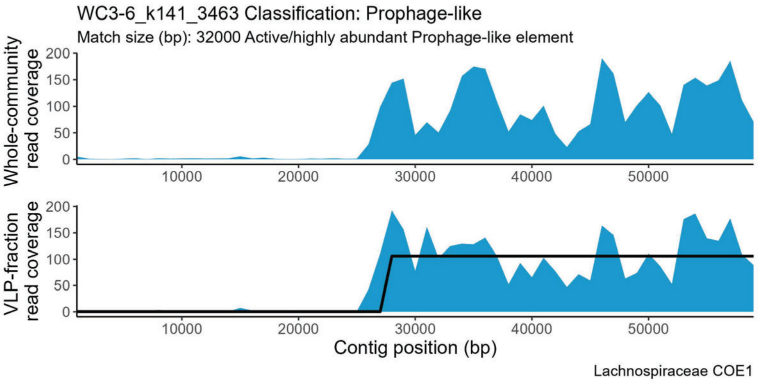
WC3-6_k141_3463 Classification: Prophage-like
Match size (bp): 32000 Active/highly abundant Prophage-like element
Whole-community read coverage
VLP-fraction read coverage
Contig position (bp)
Lachnospiraceae COE1

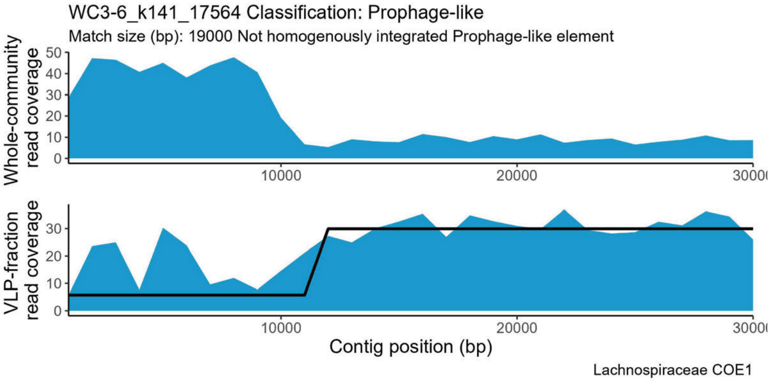
WC3-6_k141_17564 Classification: Prophage-like
Match size (bp): 19000 Not homogenously integrated Prophage-like element
Whole-community read coverage
VLP-fraction read coverage
Contig position (bp)
Lachnospiraceae COE1

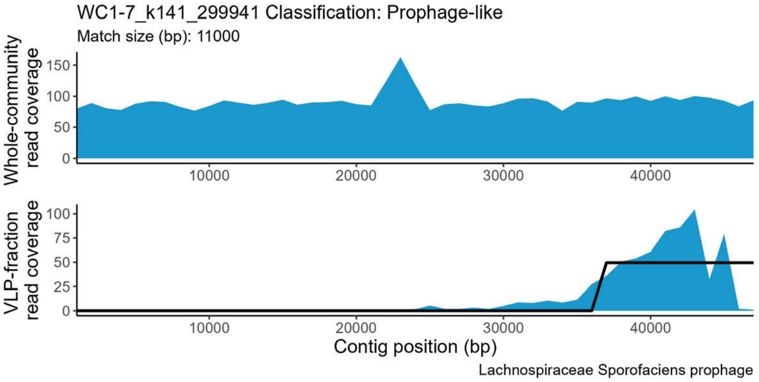
WC1-7_k141_299941 Classification: Prophage-like
Match size (bp): 11000
Whole-community read coverage
VLP-fraction read coverage
Contig position (bp)
Lachnospiraceae Sporofaciens prophage

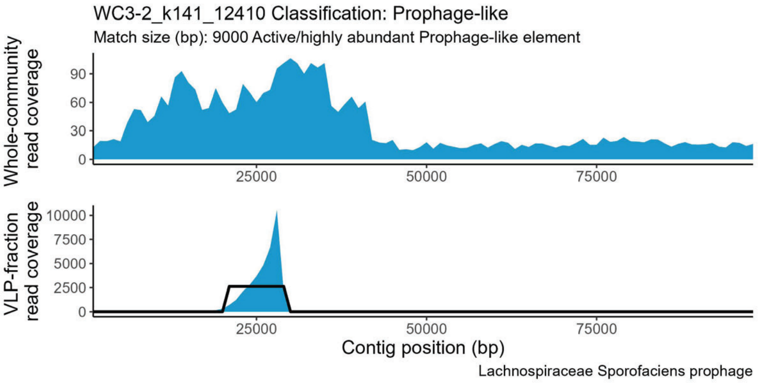
WC3-2_k141_12410 Classification: Prophage-like
Match size (bp): 9000 Active/highly abundant Prophage-like element
Whole-community read coverage
VLP-fraction read coverage
Contig position (bp)
Lachnospiraceae Sporofaciens prophage

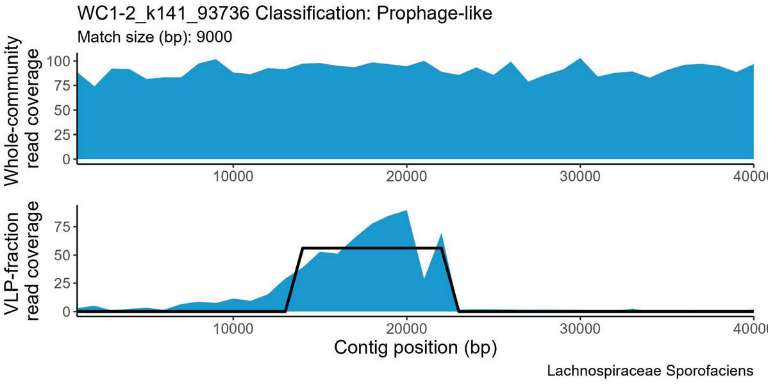
WC1-2_k141_93736 Classification: Prophage-like
Match size (bp): 9000
Whole-community read coverage
VLP-fraction read coverage
Contig position (bp)
Lachnospiraceae Sporofaciens

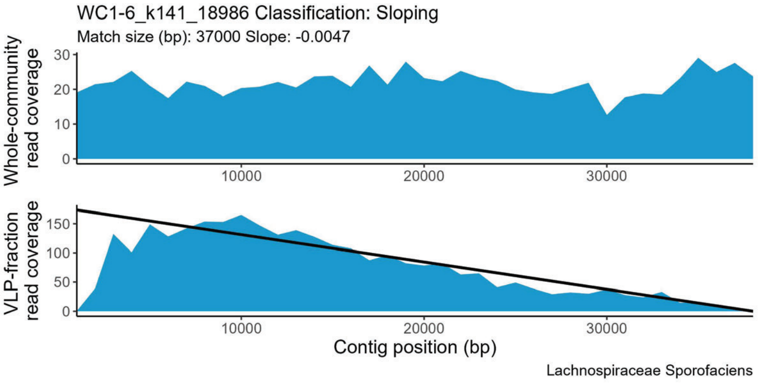
WC1-6_k141_18986 Classification: Sloping
Match size (bp): 37000 Slope: -0.0047
Whole-community read coverage
VLP-fraction read coverage
Contig position (bp)
Lachnospiraceae Sporofaciens

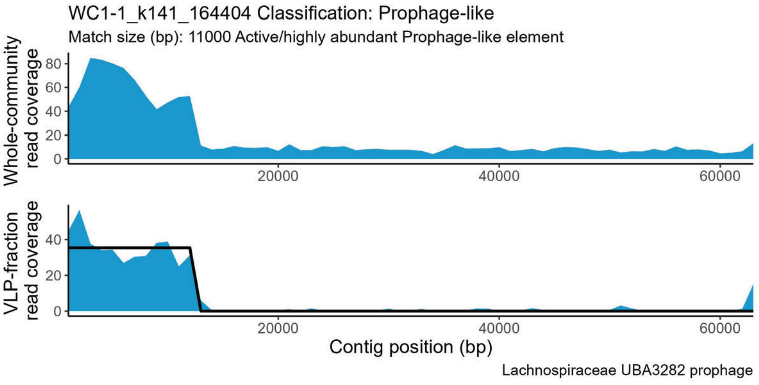
WC1-1_k141_164404 Classification: Prophage-like
Match size (bp): 11000 Active/highly abundant Prophage-like element
Whole-community read coverage
VLP-fraction read coverage
Contig position (bp)
Lachnospiraceae UBA3282 prophage

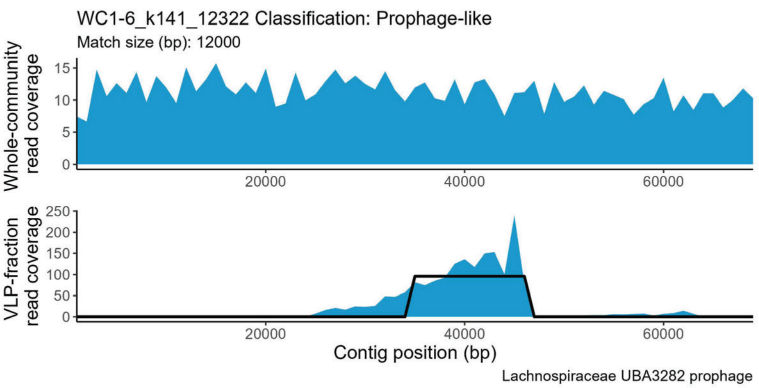
WC1-6_k141_12322 Classification: Prophage-like
Match size (bp): 12000
Whole-community read coverage
0
5
10
15
20000
40000
60000
VLP-fraction read coverage
0
50
100
150
200
250
Contig position (bp)
Lachnospiraceae UBA3282 prophage

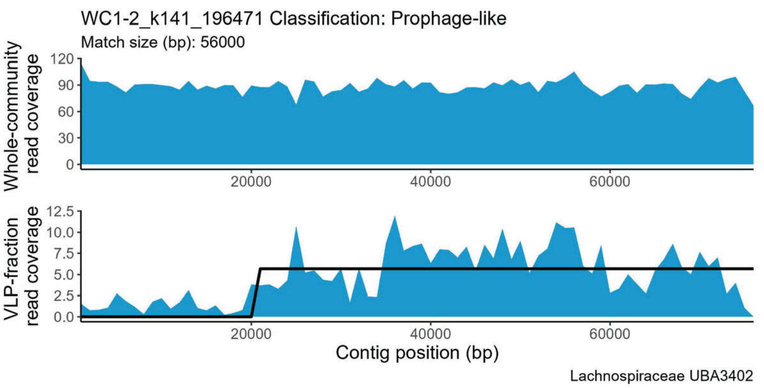
WC1-2_k141_196471 Classification: Prophage-like
Match size (bp): 56000
Whole-community read coverage
0
30
60
90
120
20000
40000
60000
VLP-fraction read coverage
0.0
2.5
5.0
7.5
10.0
12.5
Contig position (bp)
Lachnospiraceae UBA3402

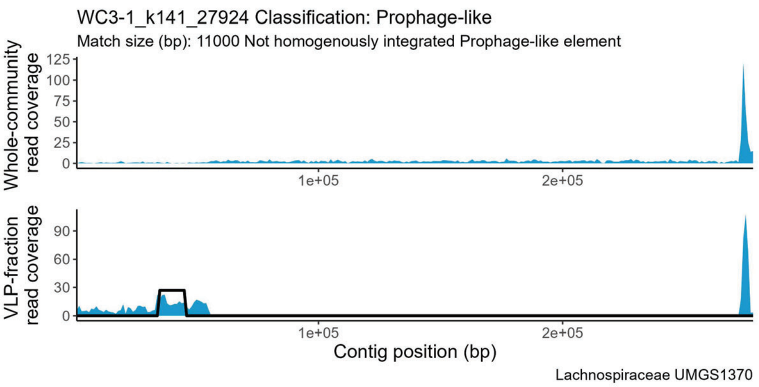
WC3-1_k141_27924 Classification: Prophage-like
Match size (bp): 11000 Not homogenously integrated Prophage-like element
Whole-community read coverage
0
25
50
75
100
125
1e+05
2e+05
VLP-fraction read coverage
0
30
60
90
Contig position (bp)
Lachnospiraceae UMGS1370

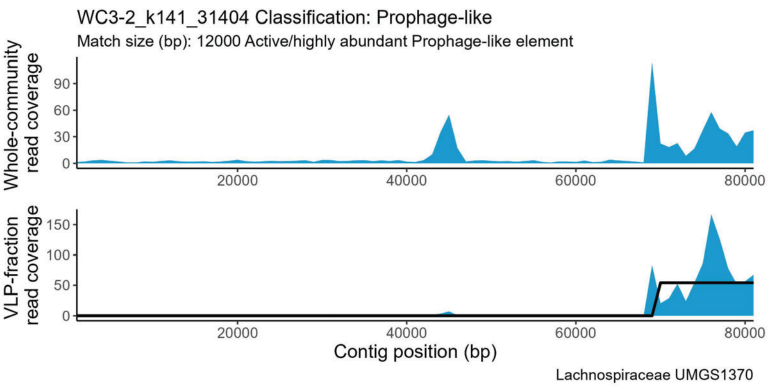
WC3-2_k141_31404 Classification: Prophage-like
Match size (bp): 12000 Active/highly abundant Prophage-like element
Whole-community read coverage
0
30
60
90
20000
40000
60000
80000
VLP-fraction read coverage
0
50
100
150
Contig position (bp)
Lachnospiraceae UMGS1370

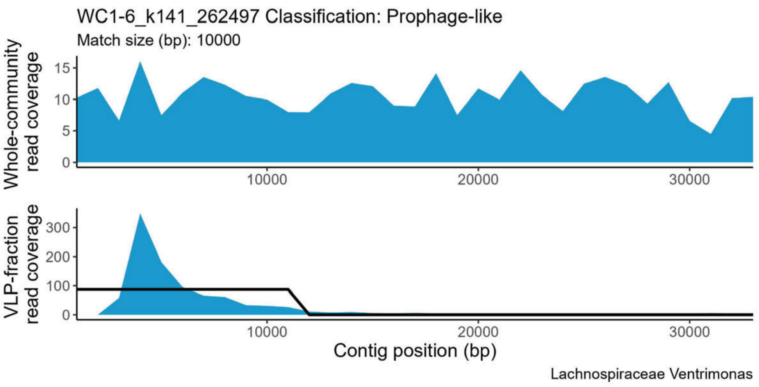
WC1-6_k141_262497 Classification: Prophage-like
Match size (bp): 10000
Whole-community read coverage
0
5
10
15
10000
20000
30000
VLP-fraction read coverage
0
100
200
300
Contig position (bp)
Lachnospiraceae Ventrimonas

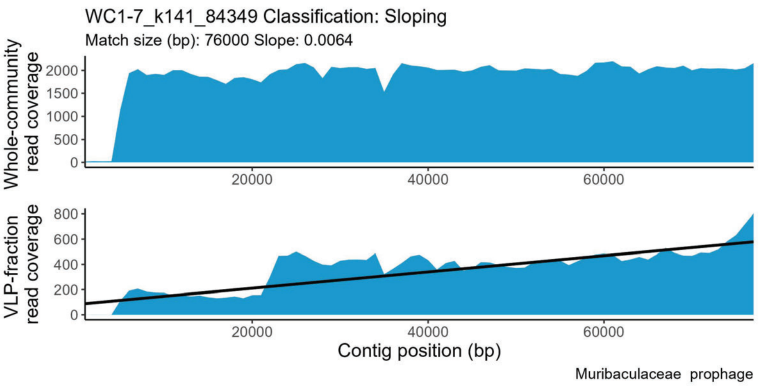
WC1-7_k141_84349 Classification: Sloping
Match size (bp): 76000 Slope: 0.0064
Whole-community read coverage
0
500
1000
1500
2000
20000
40000
60000
VLP-fraction read coverage
0
200
400
600
800
Contig position (bp)
Muribaculaceae prophage

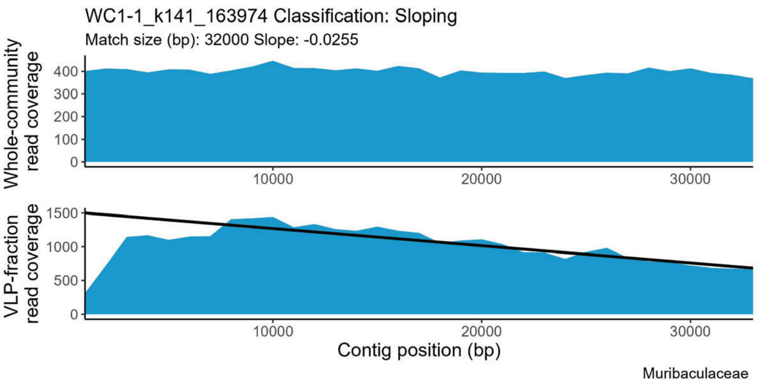
WC1-1_k141_163974 Classification: Sloping
Match size (bp): 32000 Slope: -0.0255
Whole-community read coverage
0
100
200
300
400
10000
20000
30000
VLP-fraction read coverage
0
500
1000
1500
Contig position (bp)
Muribaculaceae

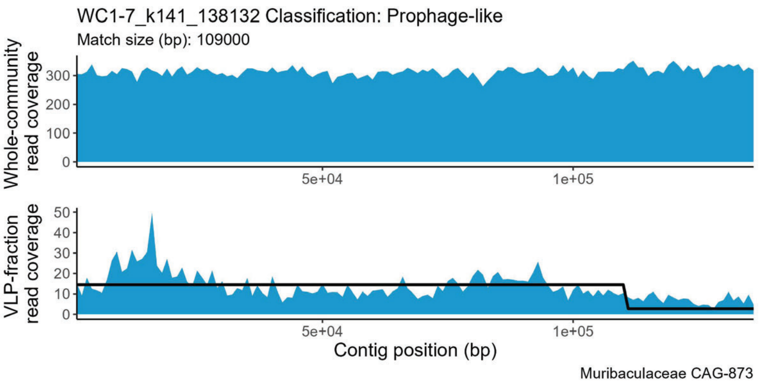
WC1-7_k141_138132 Classification: Prophage-like
Match size (bp): 109000
Whole-community read coverage
0
100
200
300
5e+04
1e+05
VLP-fraction read coverage
0
10
20
30
40
50
Contig position (bp)
Muribaculaceae CAG-873

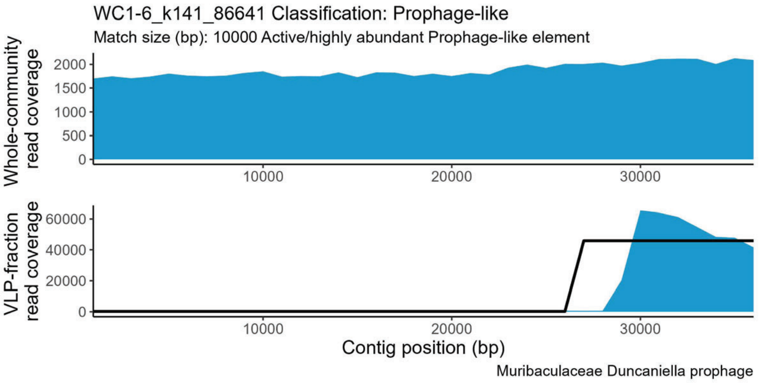
WC1-6_k141_86641 Classification: Prophage-like
Match size (bp): 10000 Active/highly abundant Prophage-like element
Whole-community read coverage
VLP-fraction read coverage
Contig position (bp)
Muribaculaceae Duncaniella prophage

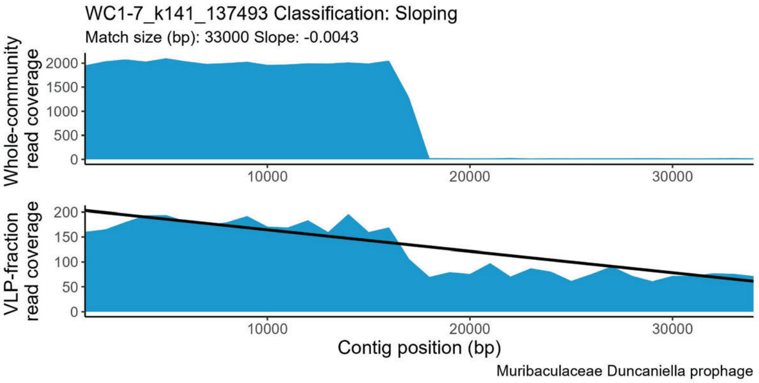
WC1-7_k141_137493 Classification: Sloping
Match size (bp): 33000 Slope: -0.0043
Whole-community read coverage
VLP-fraction read coverage
Contig position (bp)
Muribaculaceae Duncaniella prophage

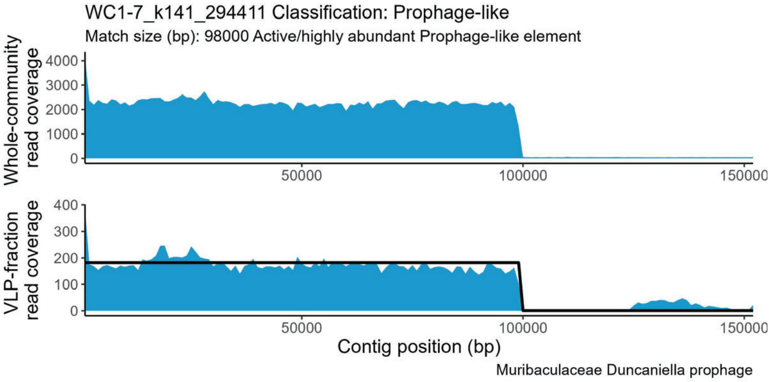
WC1-7_k141_294411 Classification: Prophage-like
Match size (bp): 98000 Active/highly abundant Prophage-like element
Whole-community read coverage
VLP-fraction read coverage
Contig position (bp)
Muribaculaceae Duncaniella prophage

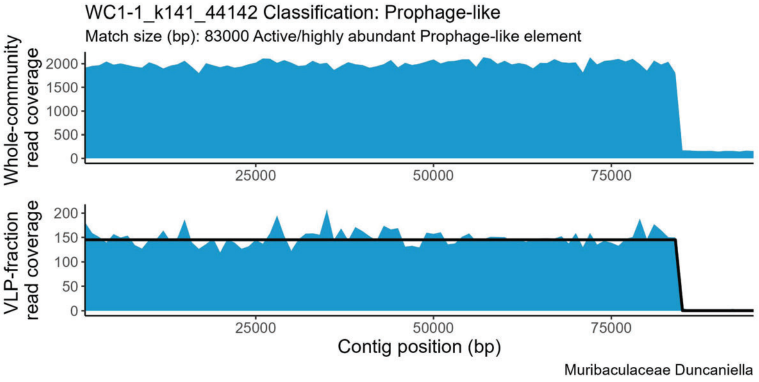
WC1-1_k141_44142 Classification: Prophage-like
Match size (bp): 83000 Active/highly abundant Prophage-like element
Whole-community read coverage
VLP-fraction read coverage
Contig position (bp)
Muribaculaceae Duncaniella

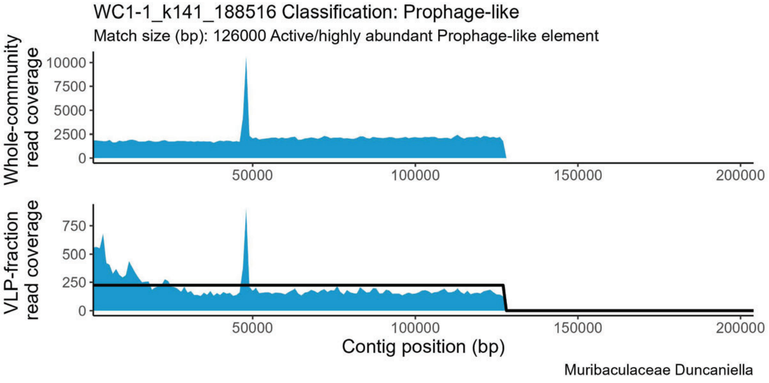
WC1-1_k141_188516 Classification: Prophage-like
Match size (bp): 126000 Active/highly abundant Prophage-like element
Whole-community read coverage
VLP-fraction read coverage
Contig position (bp)
Muribaculaceae Duncaniella

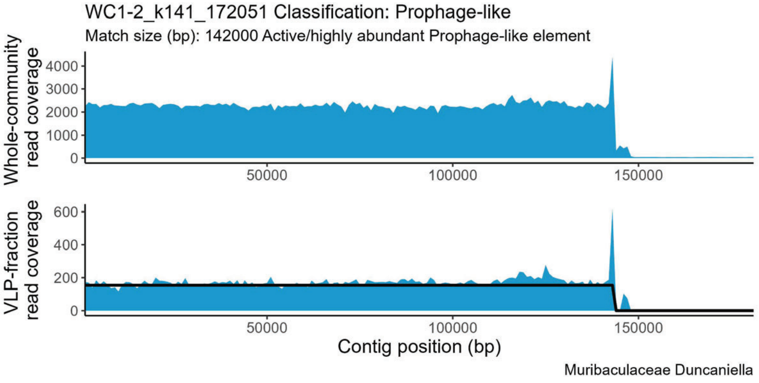
WC1-2_k141_172051 Classification: Prophage-like
Match size (bp): 142000 Active/highly abundant Prophage-like element
Whole-community read coverage
VLP-fraction read coverage
Contig position (bp)
Muribaculaceae Duncaniella

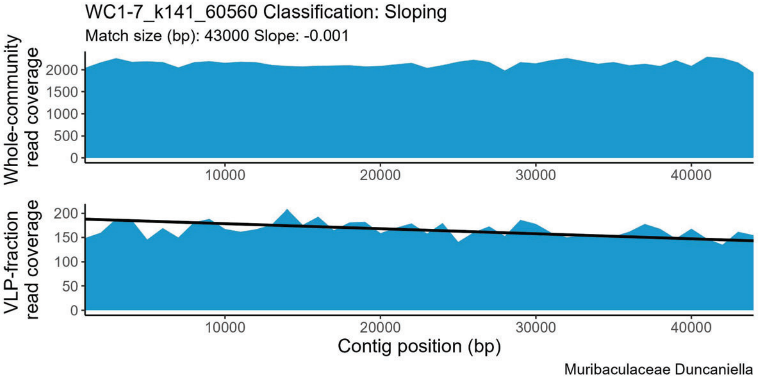
WC1-7_k141_60560 Classification: Sloping
Match size (bp): 43000 Slope: -0.001
Whole-community read coverage
VLP-fraction read coverage
Contig position (bp)
Muribaculaceae Duncaniella

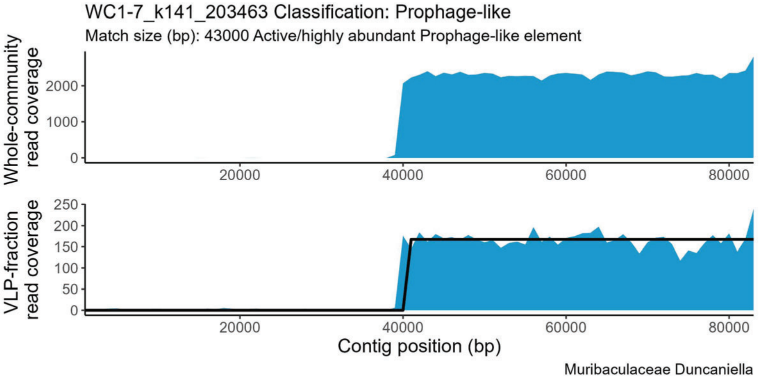
WC1-7_k141_203463 Classification: Prophage-like
Match size (bp): 43000 Active/highly abundant Prophage-like element
Whole-community read coverage
VLP-fraction read coverage
Contig position (bp)
Muribaculaceae Duncaniella

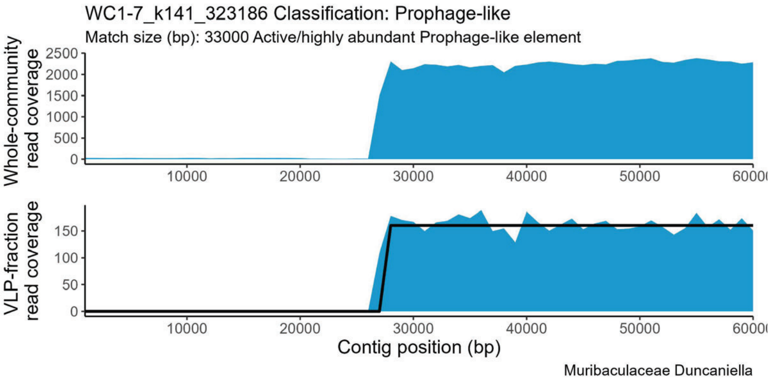
WC1-7_k141_323186 Classification: Prophage-like
Match size (bp): 33000 Active/highly abundant Prophage-like element
Whole-community read coverage
2500 2000 1500 1000 500 0
VLP-fraction read coverage
150 100 50 0
10000 20000 30000 40000 50000
Contig position (bp)
Muribaculaceae Duncaniella

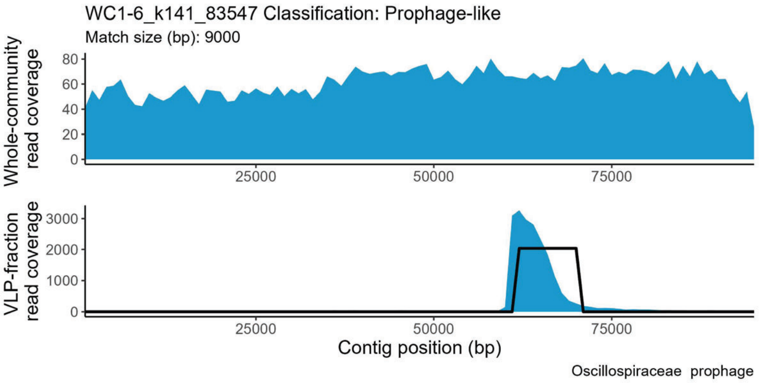
WC1-6_k141_83547 Classification: Prophage-like
Match size (bp): 9000
Whole-community read coverage
80 60 40 20 0
VLP-fraction read coverage
3000 2000 1000 0
25000 50000 75000
Contig position (bp)
Oscillospiraceae prophage

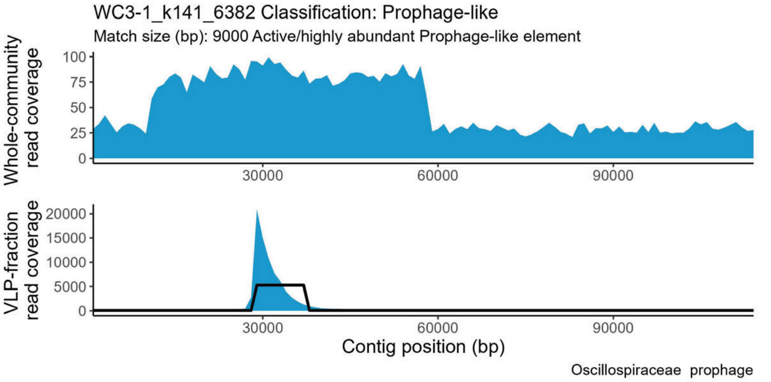
WC3-1_k141_6382 Classification: Prophage-like
Match size (bp): 9000 Active/highly abundant Prophage-like element
Whole-community read coverage
100 75 50 25 0
VLP-fraction read coverage
20000 15000 10000 5000 0
30000 60000 90000
Contig position (bp)
Oscillospiraceae prophage

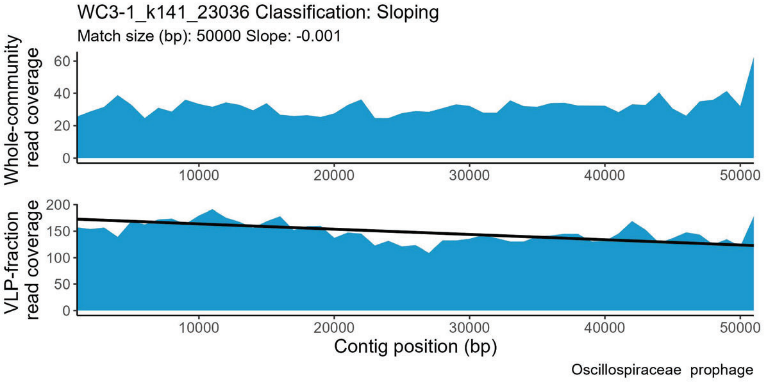
WC3-1_k141_23036 Classification: Sloping
Match size (bp): 50000 Slope: -0.001
Whole-community read coverage
60 40 20 0
VLP-fraction read coverage
200 150 100 50 0
10000 20000 30000 40000 50000
Contig position (bp)
Oscillospiraceae prophage

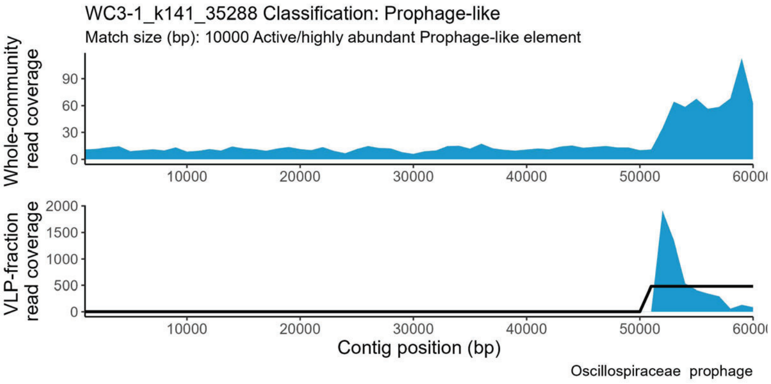
WC3-1_k141_35288 Classification: Prophage-like
Match size (bp): 10000 Active/highly abundant Prophage-like element
Whole-community read coverage
90 60 30 0
VLP-fraction read coverage
2000 1500 1000 500 0
10000 20000 30000 40000 50000
Contig position (bp)
Oscillospiraceae prophage

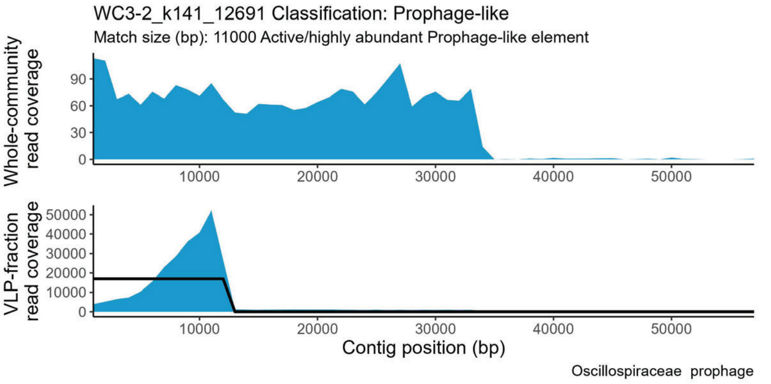
WC3-2_k141_12691 Classification: Prophage-like
Match size (bp): 11000 Active/highly abundant Prophage-like element
Whole-community read coverage
90 60 30 0
VLP-fraction read coverage
50000 40000 30000 20000 10000 0
10000 20000 30000 40000 50000
Contig position (bp)
Oscillospiraceae prophage

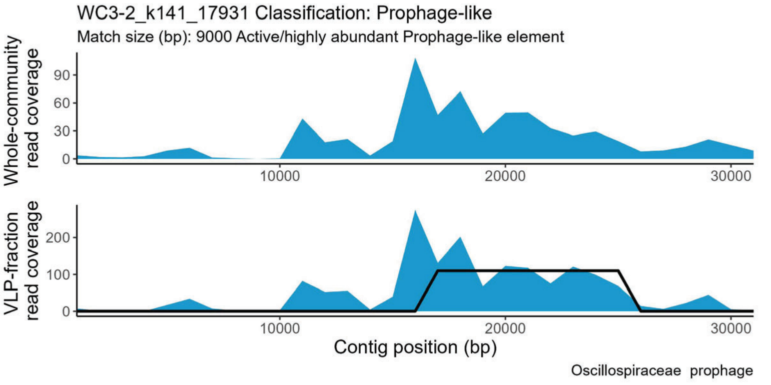
WC3-2_k141_17931 Classification: Prophage-like
Match size (bp): 9000 Active/highly abundant Prophage-like element
Whole-community read coverage
90 60 30 0
VLP-fraction read coverage
200 100 0
10000 20000 30000
Contig position (bp)
Oscillospiraceae prophage

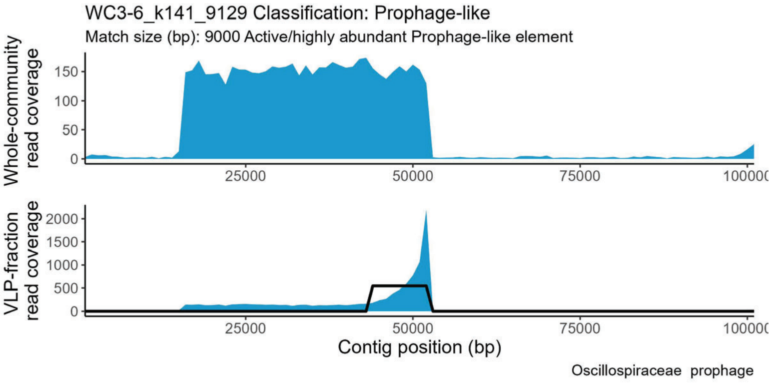
WC3-6_k141_9129 Classification: Prophage-like
Match size (bp): 9000 Active/highly abundant Prophage-like element
Whole-community read coverage
150 100 50 0
VLP-fraction read coverage
2000 1500 1000 500 0
25000 50000 75000
Contig position (bp)
Oscillospiraceae prophage

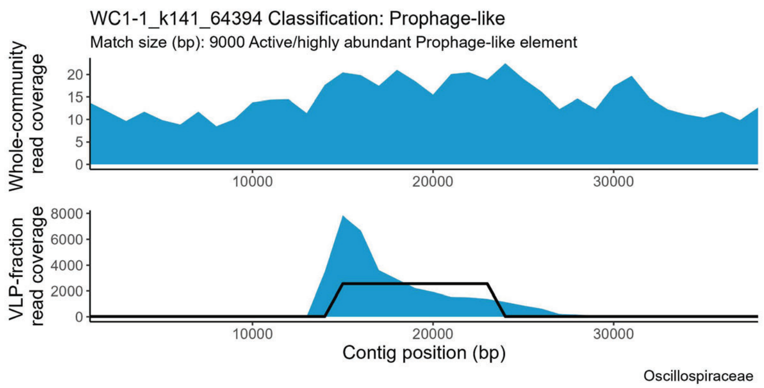
WC1-1_k141_64394 Classification: Prophage-like
Match size (bp): 9000 Active/highly abundant Prophage-like element
Whole-community read coverage
VLP-fraction read coverage
Contig position (bp)
Oscillospiraceae

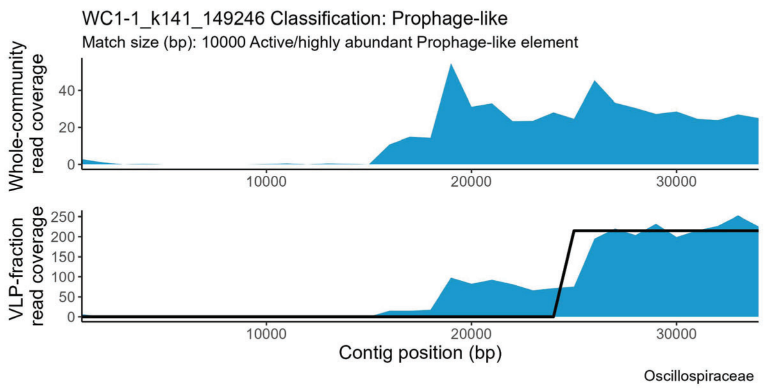
WC1-1_k141_149246 Classification: Prophage-like
Match size (bp): 10000 Active/highly abundant Prophage-like element
Whole-community read coverage
VLP-fraction read coverage
Contig position (bp)
Oscillospiraceae

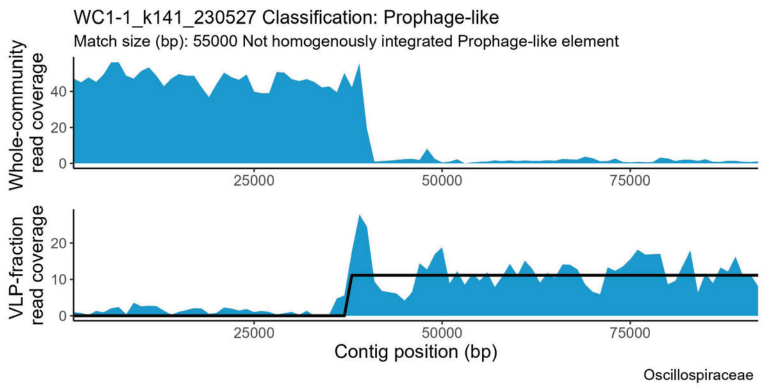
WC1-1_k141_230527 Classification: Prophage-like
Match size (bp): 55000 Not homogenously integrated Prophage-like element
Whole-community read coverage
VLP-fraction read coverage
Contig position (bp)
Oscillospiraceae

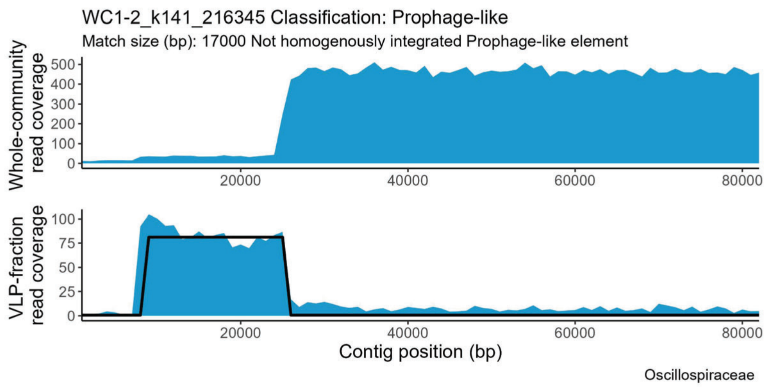
WC1-2_k141_216345 Classification: Prophage-like
Match size (bp): 17000 Not homogenously integrated Prophage-like element
Whole-community read coverage
VLP-fraction read coverage
Contig position (bp)
Oscillospiraceae

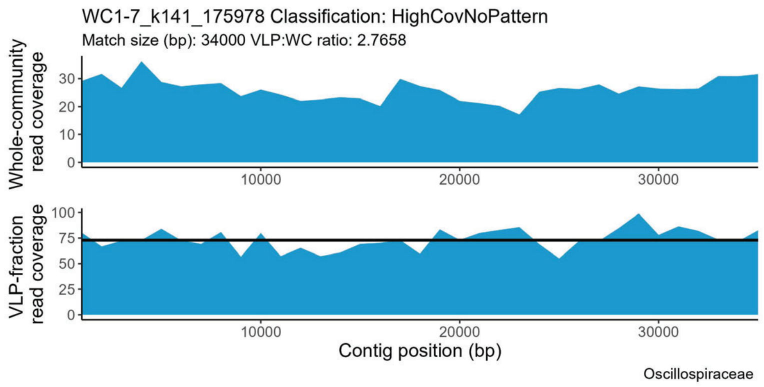
WC1-7_k141_175978 Classification: HighCovNoPattern
Match size (bp): 34000 VLP:WC ratio: 2.7658
Whole-community read coverage
VLP-fraction read coverage
Contig position (bp)
Oscillospiraceae

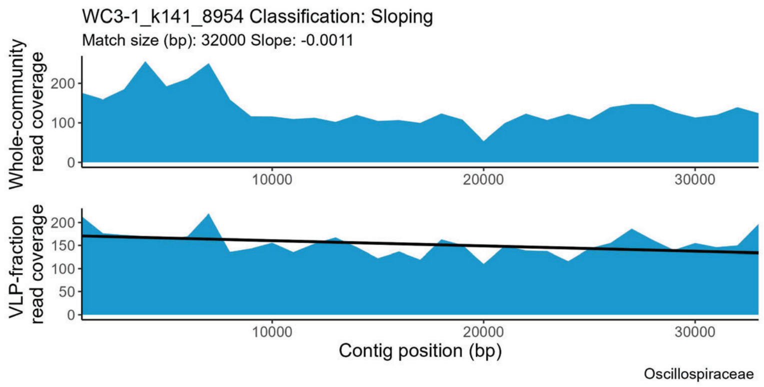
WC3-1_k141_8954 Classification: Sloping
Match size (bp): 32000 Slope: -0.0011
Whole-community read coverage
VLP-fraction read coverage
Contig position (bp)
Oscillospiraceae

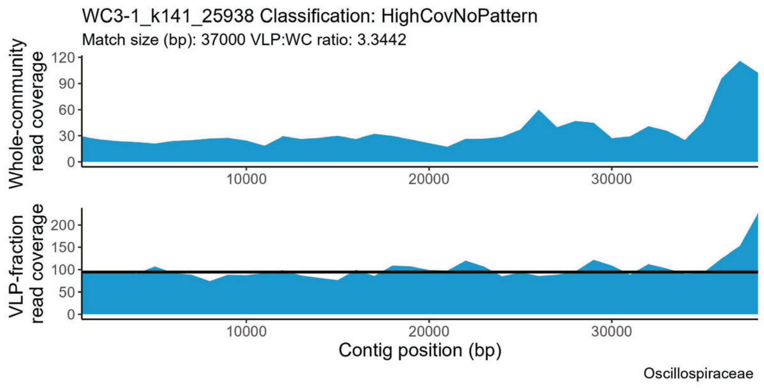
WC3-1_k141_25938 Classification: HighCovNoPattern
Match size (bp): 37000 VLP:WC ratio: 3.3442
Whole-community read coverage
VLP-fraction read coverage
Contig position (bp)
Oscillospiraceae

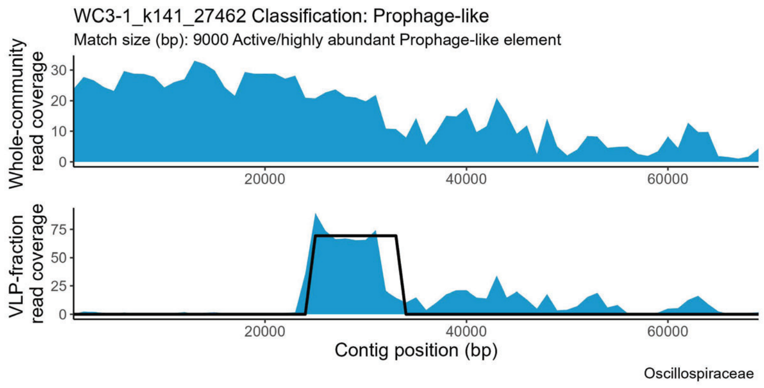
WC3-1_k141_27462 Classification: Prophage-like
Match size (bp): 9000 Active/highly abundant Prophage-like element
Whole-community read coverage
VLP-fraction read coverage
Contig position (bp)
Oscillospiraceae

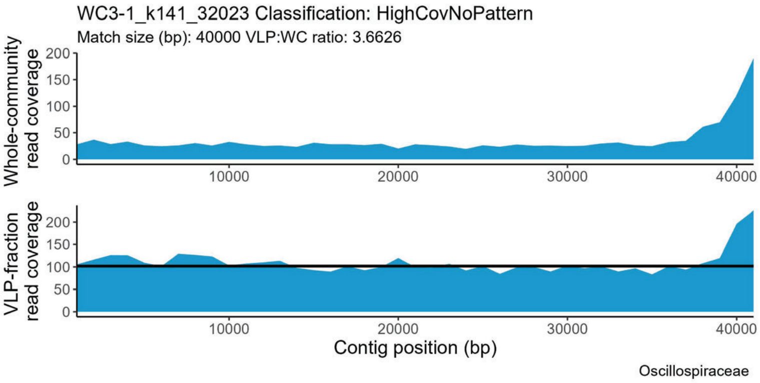
WC3-1_k141_32023 Classification: HighCovNoPattern
Match size (bp): 40000 VLP:WC ratio: 3.6626
Whole-community read coverage
VLP-fraction read coverage
Contig position (bp)
Oscillospiraceae

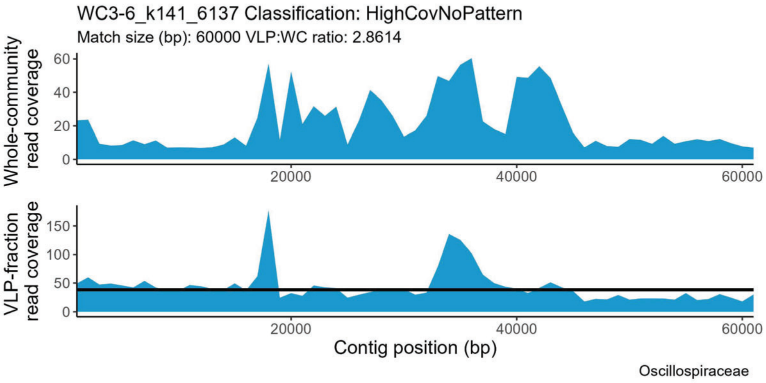
WC3-6_k141_6137 Classification: HighCovNoPattern
Match size (bp): 60000 VLP:WC ratio: 2.8614
Whole-community read coverage
VLP-fraction read coverage
Contig position (bp)
Oscillospiraceae

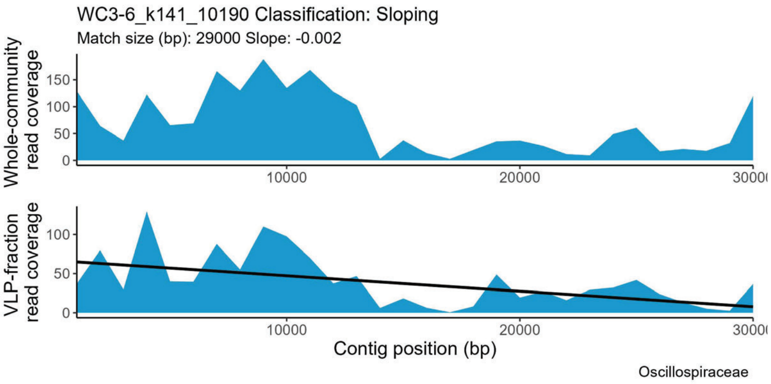
WC3-6_k141_10190 Classification: Sloping
Match size (bp): 29000 Slope: -0.002
Whole-community read coverage
VLP-fraction read coverage
Contig position (bp)
Oscillospiraceae

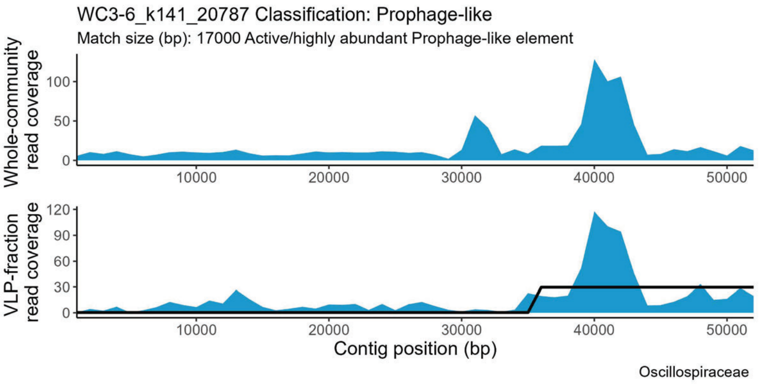
WC3-6_k141_20787 Classification: Prophage-like
Match size (bp): 17000 Active/highly abundant Prophage-like element
Whole-community read coverage
VLP-fraction read coverage
Contig position (bp)
Oscillospiraceae

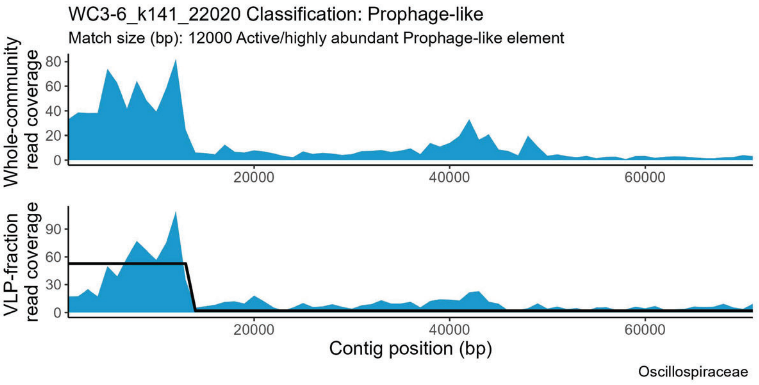
WC3-6_k141_22020 Classification: Prophage-like
Match size (bp): 12000 Active/highly abundant Prophage-like element
Whole-community read coverage
VLP-fraction read coverage
Contig position (bp)
Oscillospiraceae

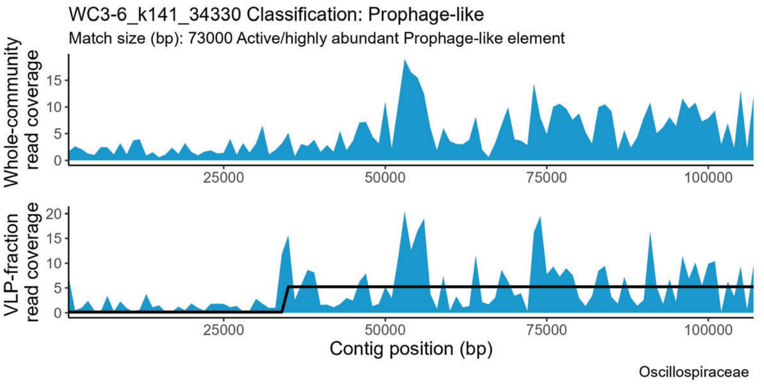
WC3-6_k141_34330 Classification: Prophage-like
Match size (bp): 73000 Active/highly abundant Prophage-like element
Whole-community read coverage
VLP-fraction read coverage
Contig position (bp)
Oscillospiraceae

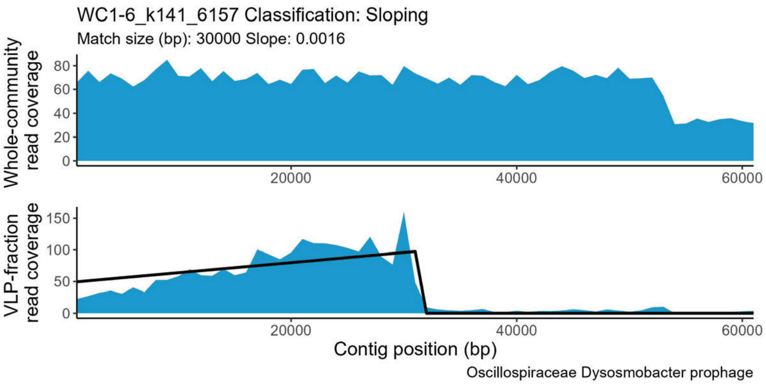
WC1-6_k141_6157 Classification: Sloping
Match size (bp): 30000 Slope: 0.0016
Whole-community read coverage
VLP-fraction read coverage
Contig position (bp)
Oscillospiraceae Dysosmobacter prophage

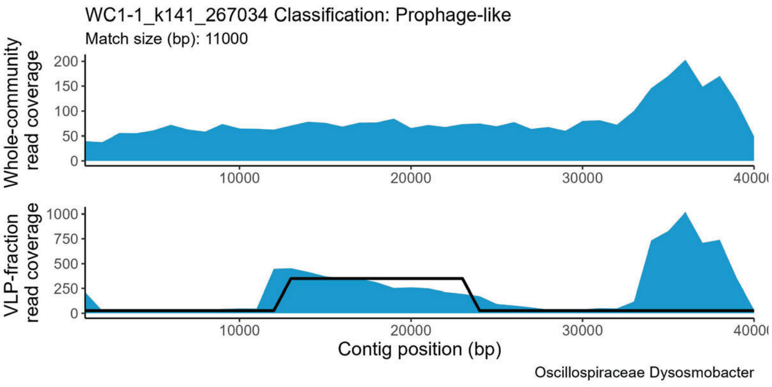
WC1-1_k141_267034 Classification: Prophage-like
Match size (bp): 11000
Whole-community read coverage
VLP-fraction read coverage
Contig position (bp)
Oscillospiraceae Dysosmobacter

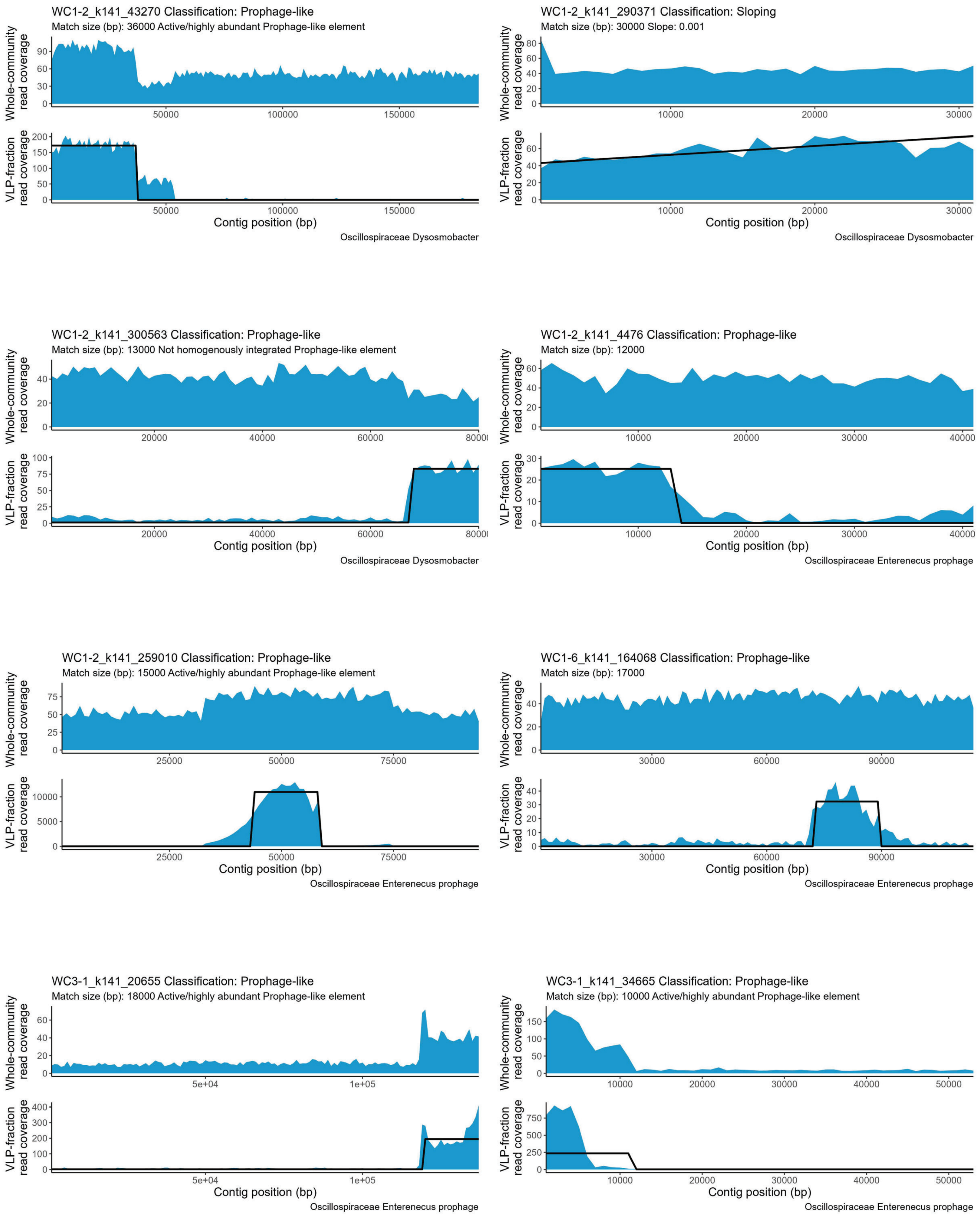

WC1-2_k141_43270 Classification: Prophage-like
Match size (bp): 36000 Active/highly abundant Prophage-like element
Whole-community read coverage
VLP-fraction read coverage
Contig position (bp)
Oscillospiraceae Dysosmobacter
WC1-2_k141_290371 Classification: Sloping
Match size (bp): 30000 Slope: 0.001
Whole-community read coverage
VLP-fraction read coverage
Contig position (bp)
Oscillospiraceae Dysosmobacter
WC1-2_k141_300563 Classification: Prophage-like
Match size (bp): 13000 Not homogenously integrated Prophage-like element
Whole-community read coverage
VLP-fraction read coverage
Contig position (bp)
Oscillospiraceae Dysosmobacter
WC1-2_k141_4476 Classification: Prophage-like
Match size (bp): 12000
Whole-community read coverage
VLP-fraction read coverage
Contig position (bp)
Oscillospiraceae Enterenecus prophage
WC1-2_k141_259010 Classification: Prophage-like
Match size (bp): 15000 Active/highly abundant Prophage-like element
Whole-community read coverage
VLP-fraction read coverage
Contig position (bp)
Oscillospiraceae Enterenecus prophage
WC1-6_k141_164068 Classification: Prophage-like
Match size (bp): 17000
Whole-community read coverage
VLP-fraction read coverage
Contig position (bp)
Oscillospiraceae Enterenecus prophage
WC3-1_k141_20655 Classification: Prophage-like
Match size (bp): 18000 Active/highly abundant Prophage-like element
Whole-community read coverage
VLP-fraction read coverage
Contig position (bp)
Oscillospiraceae Enterenecus prophage
WC3-1_k141_34665 Classification: Prophage-like
Match size (bp): 10000 Active/highly abundant Prophage-like element
Whole-community read coverage
VLP-fraction read coverage
Contig position (bp)
Oscillospiraceae Enterenecus prophage

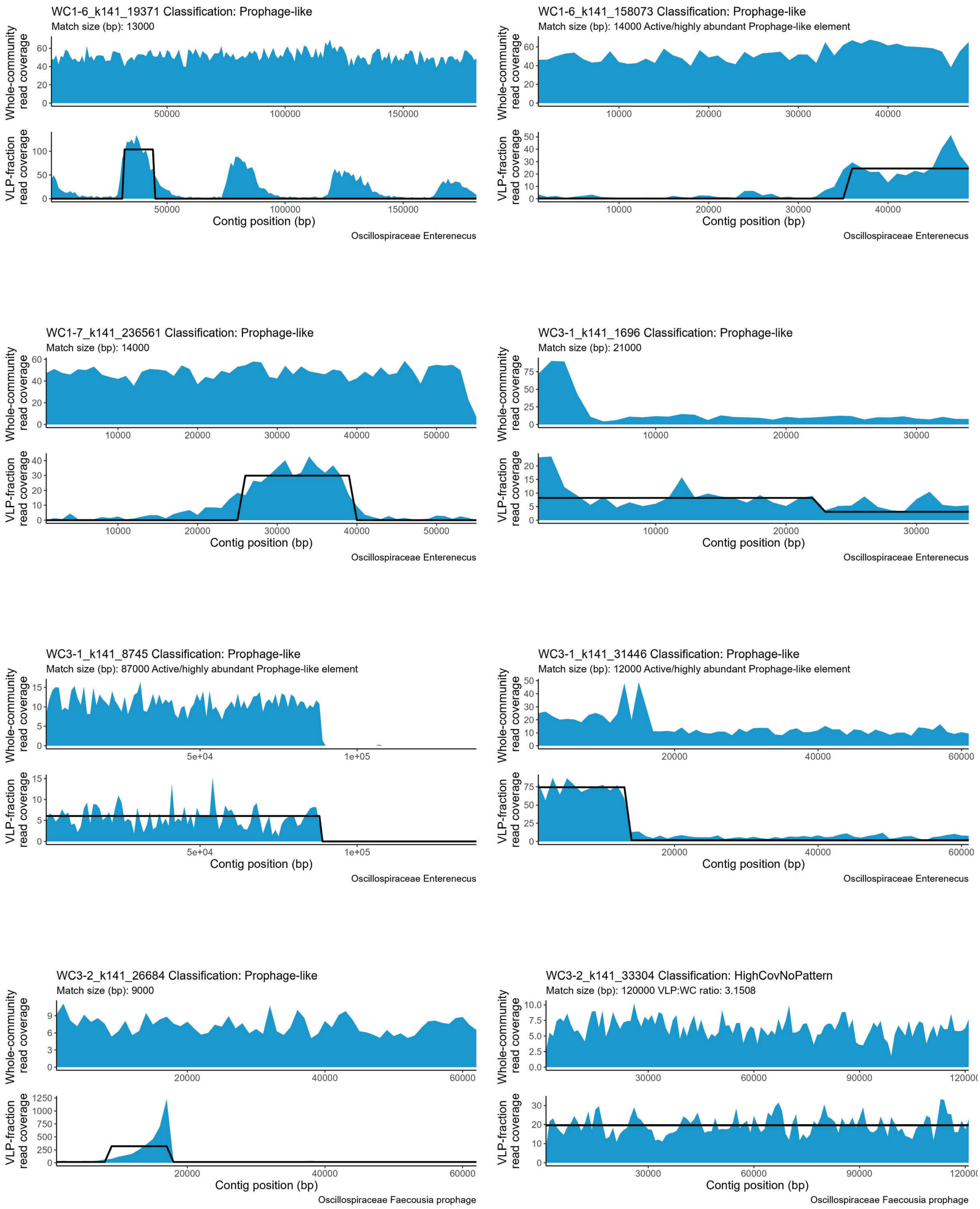
WC1-6_k141_19371 Classification: Prophage-like
Match size (bp): 13000
Whole-community read coverage
VLP-fraction read coverage
Contig position (bp)
Oscillospiraceae Enterenecus
WC1-6_k141_158073 Classification: Prophage-like
Match size (bp): 14000 Active/highly abundant Prophage-like element
Whole-community read coverage
VLP-fraction read coverage
Contig position (bp)
Oscillospiraceae Enterenecus
WC1-7_k141_236561 Classification: Prophage-like
Match size (bp): 14000
Whole-community read coverage
VLP-fraction read coverage
Contig position (bp)
Oscillospiraceae Enterenecus
WC3-1_k141_1696 Classification: Prophage-like
Match size (bp): 21000
Whole-community read coverage
VLP-fraction read coverage
Contig position (bp)
Oscillospiraceae Enterenecus
WC3-1_k141_8745 Classification: Prophage-like
Match size (bp): 87000 Active/highly abundant Prophage-like element
Whole-community read coverage
VLP-fraction read coverage
Contig position (bp)
Oscillospiraceae Enterenecus
WC3-1_k141_31446 Classification: Prophage-like
Match size (bp): 12000 Active/highly abundant Prophage-like element
Whole-community read coverage
VLP-fraction read coverage
Contig position (bp)
Oscillospiraceae Enterenecus
WC3-2_k141_26684 Classification: Prophage-like
Match size (bp): 9000
Whole-community read coverage
VLP-fraction read coverage
Contig position (bp)
Oscillospiraceae Faecousia prophage
WC3-2_k141_33304 Classification: HighCovNoPattern
Match size (bp): 120000 VLP:WC ratio: 3.1508
Whole-community read coverage
VLP-fraction read coverage
Contig position (bp)
Oscillospiraceae Faecousia prophage

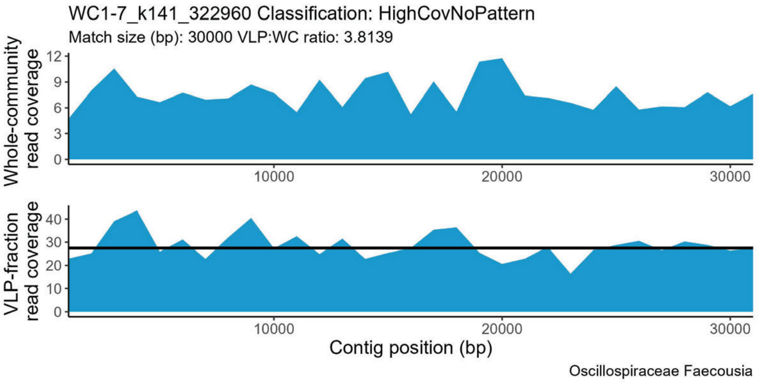

WC1-7_k141_322960 Classification: HighCovNoPattern
Match size (bp): 30000 VLP:WC ratio: 3.8139
Whole-community read coverage
VLP-fraction read coverage
Contig position (bp)
Oscillospiraceae Faecousia


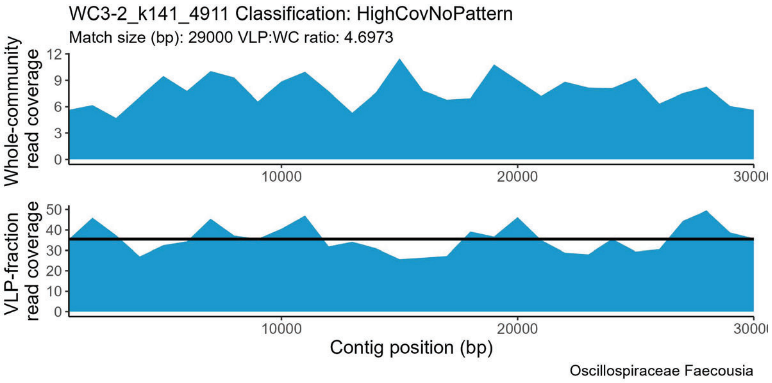

WC3-2_k141_4911 Classification: HighCovNoPattern
Match size (bp): 29000 VLP:WC ratio: 4.6973
Whole-community read coverage
VLP-fraction read coverage
Contig position (bp)
Oscillospiraceae Faecousia


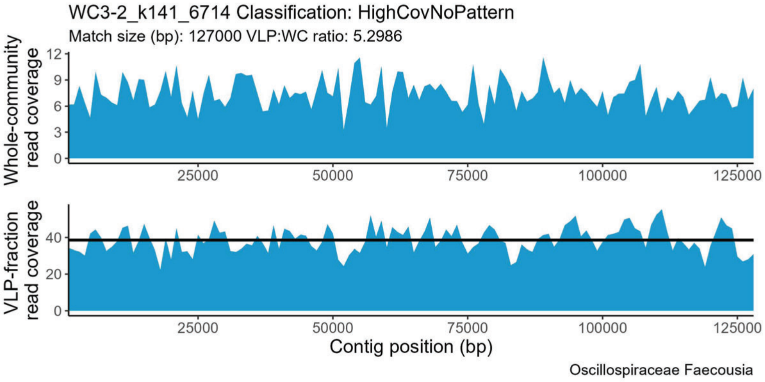

WC3-2_k141_6714 Classification: HighCovNoPattern
Match size (bp): 127000 VLP:WC ratio: 5.2986
Whole-community read coverage
VLP-fraction read coverage
Contig position (bp)
Oscillospiraceae Faecousia


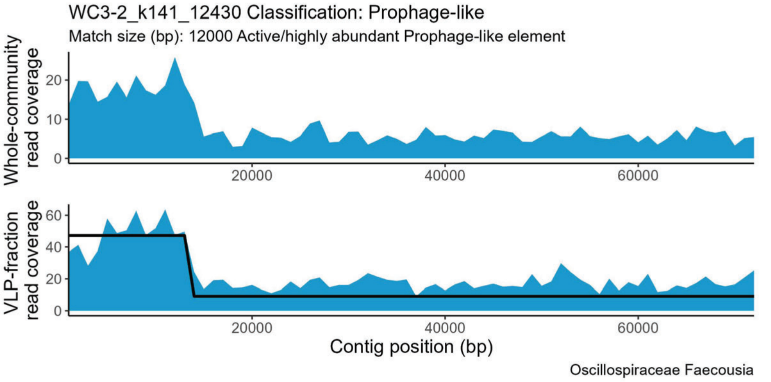

WC3-2_k141_12430 Classification: Prophage-like
Match size (bp): 12000 Active/highly abundant Prophage-like element
Whole-community read coverage
VLP-fraction read coverage
Contig position (bp)
Oscillospiraceae Faecousia


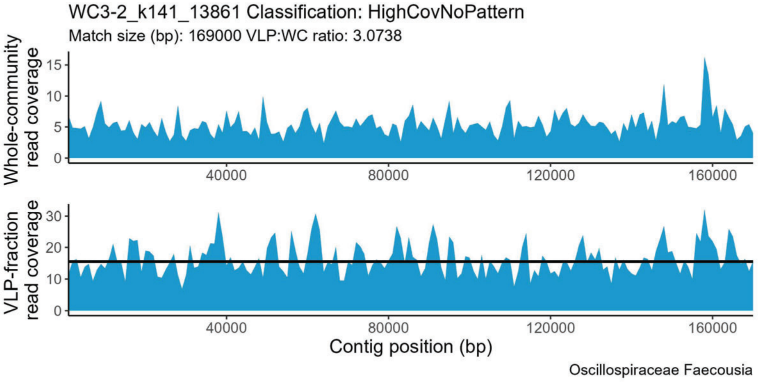

WC3-2_k141_13861 Classification: HighCovNoPattern
Match size (bp): 169000 VLP:WC ratio: 3.0738
Whole-community read coverage
VLP-fraction read coverage
Contig position (bp)
Oscillospiraceae Faecousia


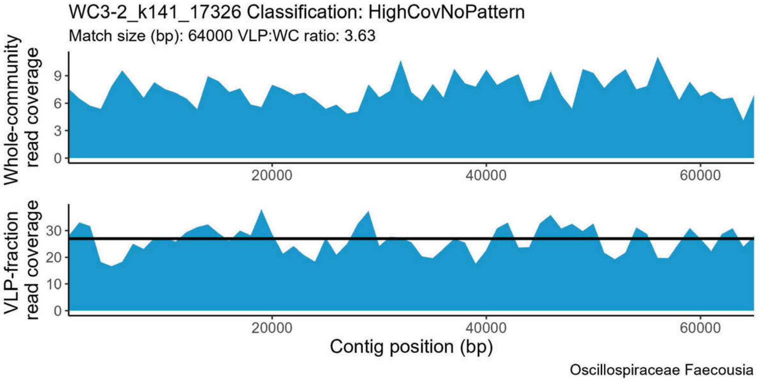

WC3-2_k141_17326 Classification: HighCovNoPattern
Match size (bp): 64000 VLP:WC ratio: 3.63
Whole-community read coverage
VLP-fraction read coverage
Contig position (bp)
Oscillospiraceae Faecousia


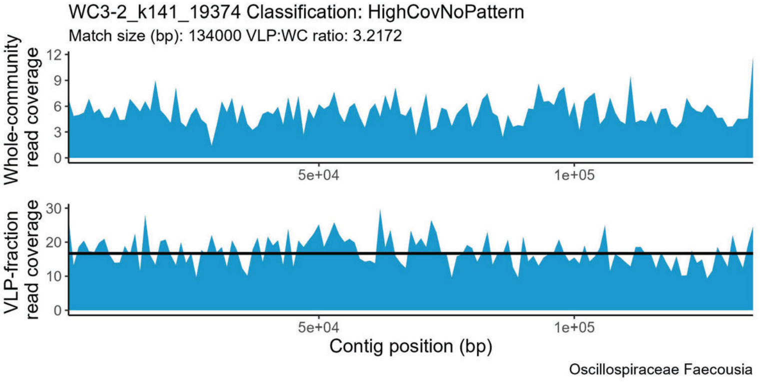

WC3-2_k141_19374 Classification: HighCovNoPattern
Match size (bp): 134000 VLP:WC ratio: 3.2172
Whole-community read coverage
VLP-fraction read coverage
Contig position (bp)
Oscillospiraceae Faecousia


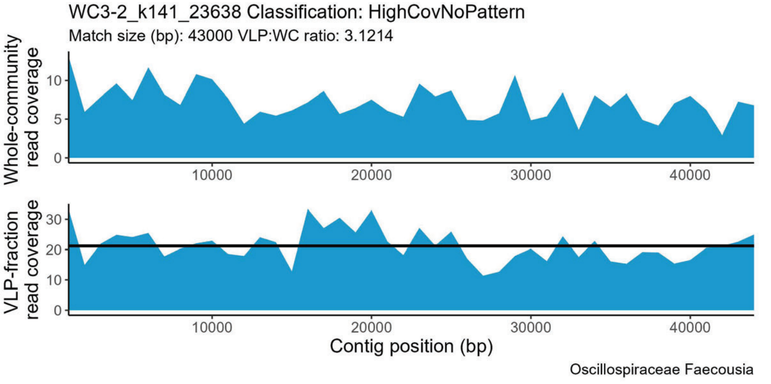

WC3-2_k141_23638 Classification: HighCovNoPattern
Match size (bp): 43000 VLP:WC ratio: 3.1214
Whole-community read coverage
VLP-fraction read coverage
Contig position (bp)
Oscillospiraceae Faecousia

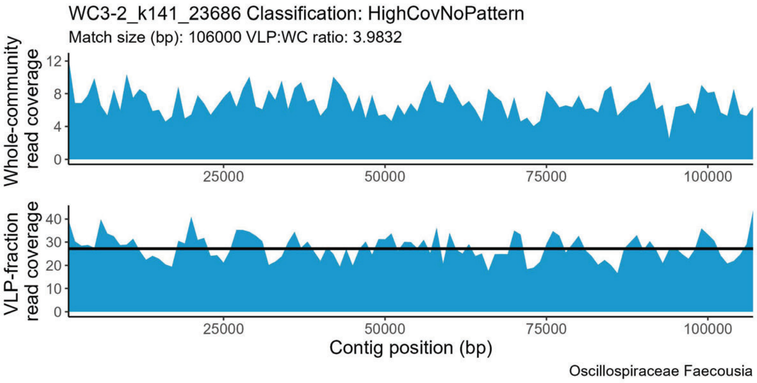
WC3-2_k141_23686 Classification: HighCovNoPattern
Match size (bp): 106000 VLP:WC ratio: 3.9832
Whole-community read coverage
VLP-fraction read coverage
Contig position (bp)
Oscillospiraceae Faecousia

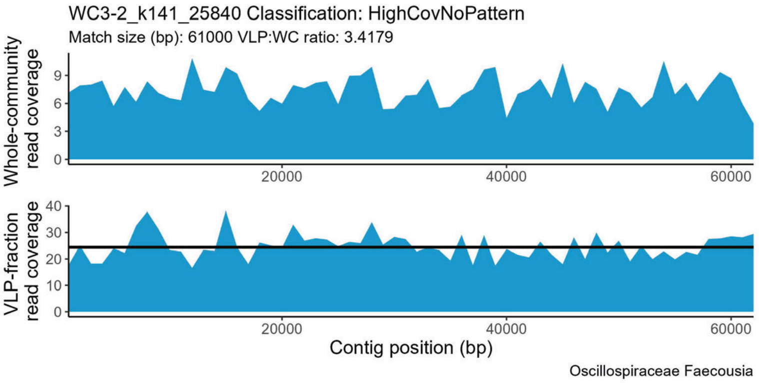
WC3-2_k141_25840 Classification: HighCovNoPattern
Match size (bp): 61000 VLP:WC ratio: 3.4179
Whole-community read coverage
VLP-fraction read coverage
Contig position (bp)
Oscillospiraceae Faecousia

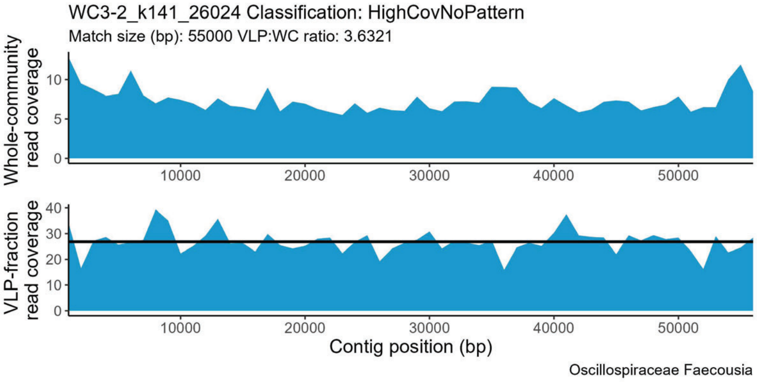
WC3-2_k141_26024 Classification: HighCovNoPattern
Match size (bp): 55000 VLP:WC ratio: 3.6321
Whole-community read coverage
VLP-fraction read coverage
Contig position (bp)
Oscillospiraceae Faecousia

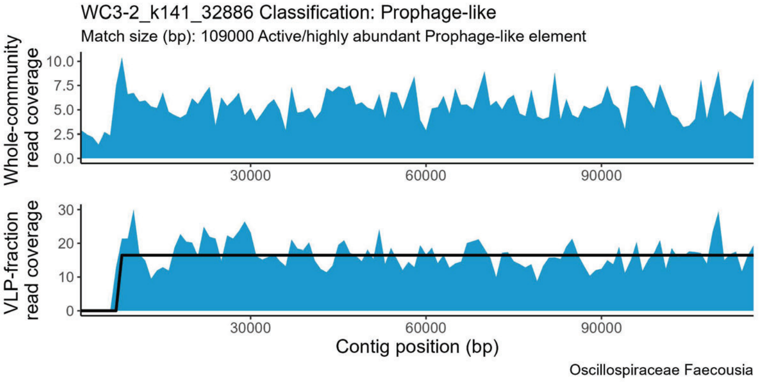
WC3-2_k141_32886 Classification: Prophage-like
Match size (bp): 109000 Active/highly abundant Prophage-like element
Whole-community read coverage
VLP-fraction read coverage
Contig position (bp)
Oscillospiraceae Faecousia

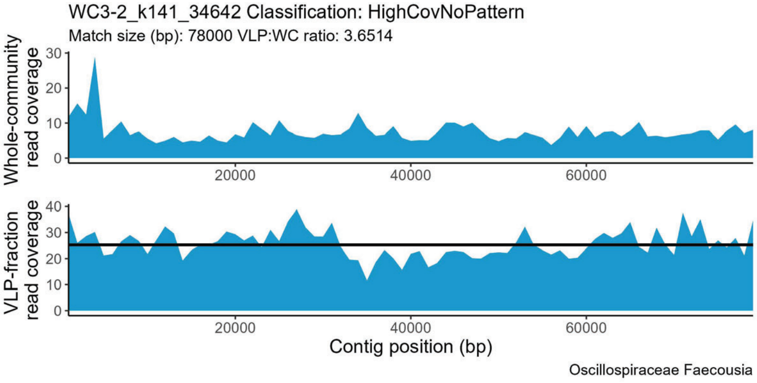
WC3-2_k141_34642 Classification: HighCovNoPattern
Match size (bp): 78000 VLP:WC ratio: 3.6514
Whole-community read coverage
VLP-fraction read coverage
Contig position (bp)
Oscillospiraceae Faecousia

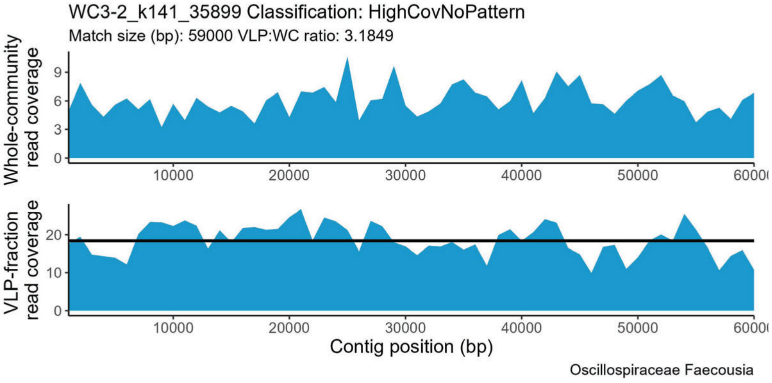
WC3-2_k141_35899 Classification: HighCovNoPattern
Match size (bp): 59000 VLP:WC ratio: 3.1849
Whole-community read coverage
VLP-fraction read coverage
Contig position (bp)
Oscillospiraceae Faecousia

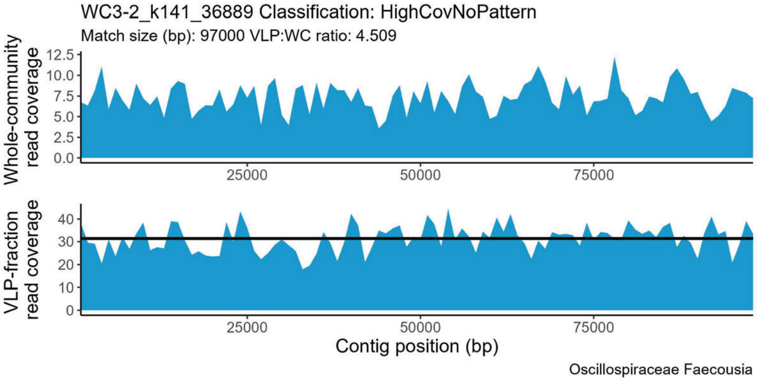
WC3-2_k141_36889 Classification: HighCovNoPattern
Match size (bp): 97000 VLP:WC ratio: 4.509
Whole-community read coverage
VLP-fraction read coverage
Contig position (bp)
Oscillospiraceae Faecousia

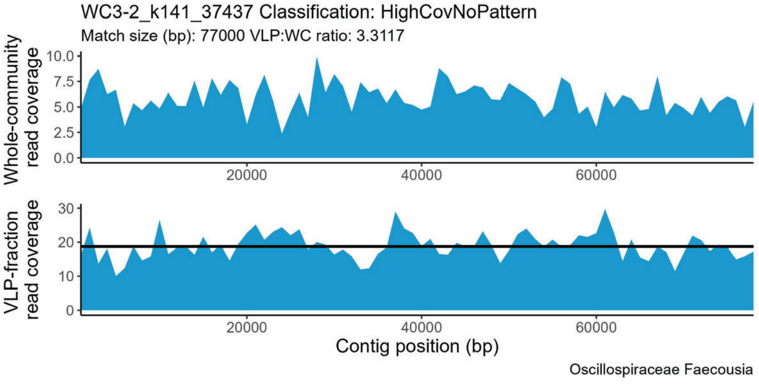
WC3-2_k141_37437 Classification: HighCovNoPattern
Match size (bp): 77000 VLP:WC ratio: 3.3117
Whole-community read coverage
VLP-fraction read coverage
Contig position (bp)
Oscillospiraceae Faecousia

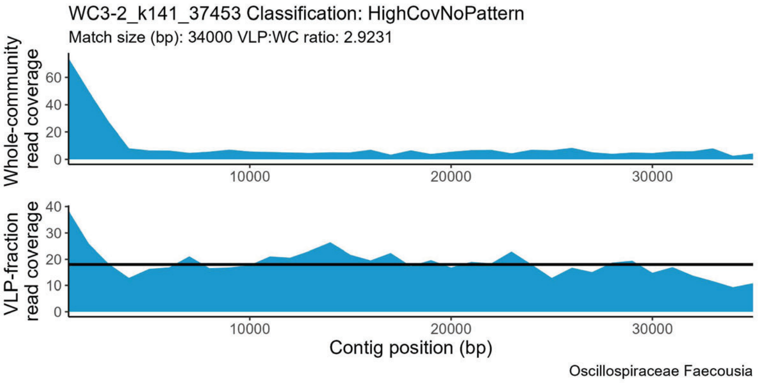
WC3-2_k141_37453 Classification: HighCovNoPattern
Match size (bp): 34000 VLP:WC ratio: 2.9231
Whole-community read coverage
VLP-fraction read coverage
Contig position (bp)
Oscillospiraceae Faecousia

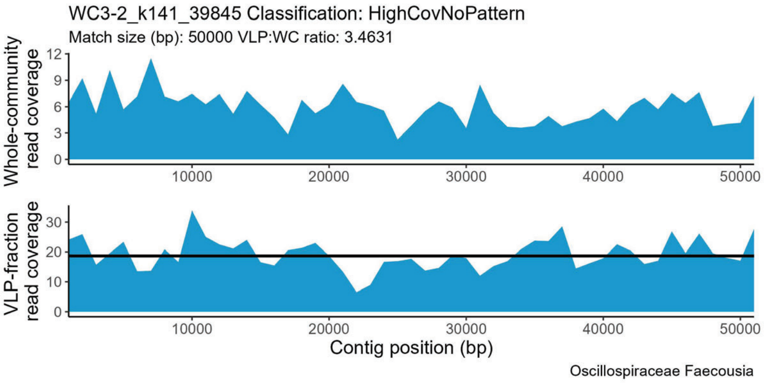
WC3-2_k141_39845 Classification: HighCovNoPattern
Match size (bp): 50000 VLP:WC ratio: 3.4631
Whole-community read coverage
VLP-fraction read coverage
Contig position (bp)
Oscillospiraceae Faecousia

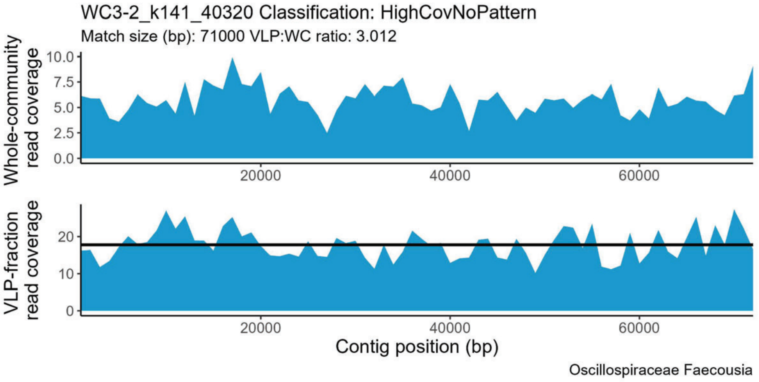
WC3-2_k141_40320 Classification: HighCovNoPattern
Match size (bp): 71000 VLP:WC ratio: 3.012
Whole-community read coverage
VLP-fraction read coverage
Contig position (bp)
Oscillospiraceae Faecousia

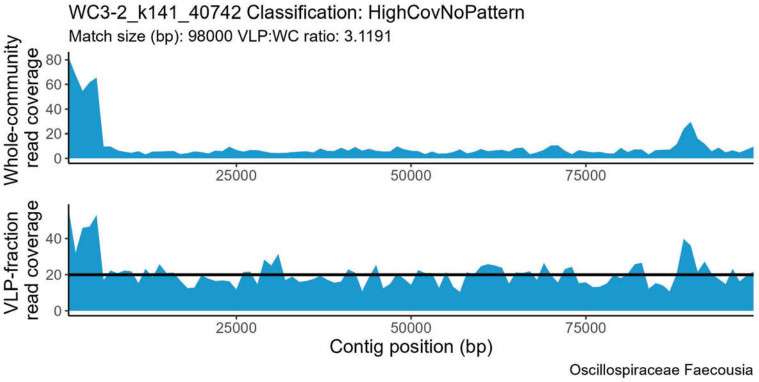
WC3-2_k141_40742 Classification: HighCovNoPattern
Match size (bp): 98000 VLP:WC ratio: 3.1191
Whole-community read coverage
VLP-fraction read coverage
Contig position (bp)
Oscillospiraceae Faecousia

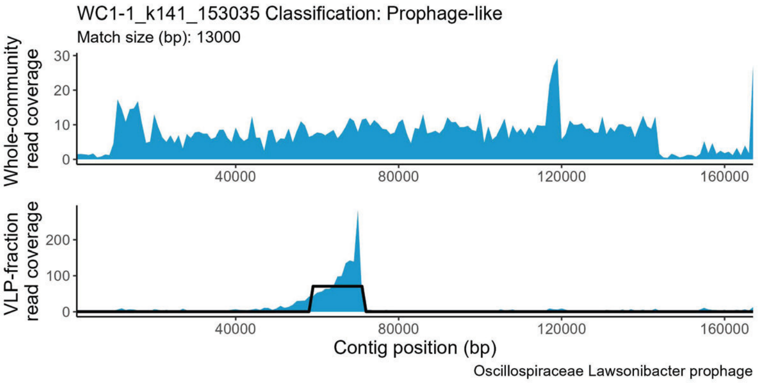
WC1-1_k141_153035 Classification: Prophage-like
Match size (bp): 13000
Whole-community read coverage
VLP-fraction read coverage
Contig position (bp)
Oscillospiraceae Lawsonibacter prophage

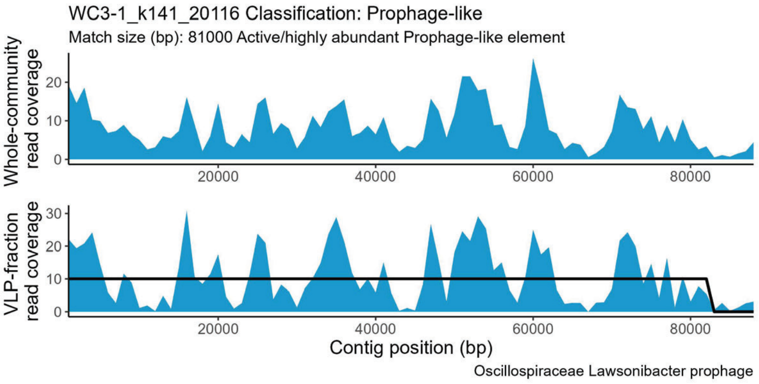
WC3-1_k141_20116 Classification: Prophage-like
Match size (bp): 81000 Active/highly abundant Prophage-like element
Whole-community read coverage
VLP-fraction read coverage
Contig position (bp)
Oscillospiraceae Lawsonibacter prophage

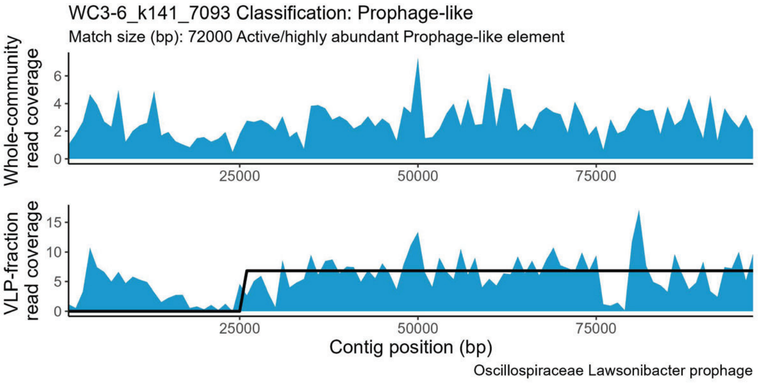
WC3-6_k141_7093 Classification: Prophage-like
Match size (bp): 72000 Active/highly abundant Prophage-like element
Whole-community read coverage
VLP-fraction read coverage
Contig position (bp)
Oscillospiraceae Lawsonibacter prophage

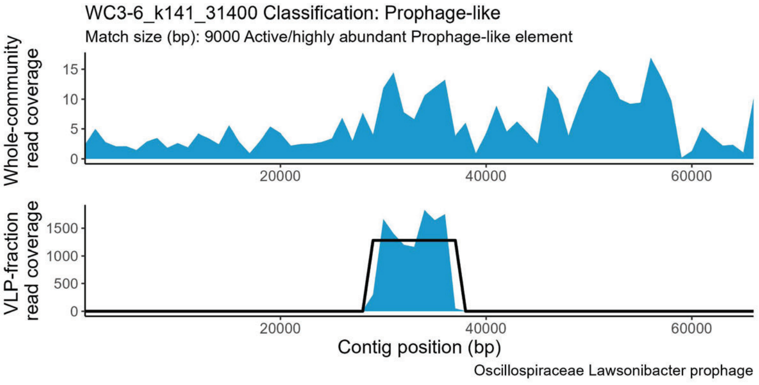
WC3-6_k141_31400 Classification: Prophage-like
Match size (bp): 9000 Active/highly abundant Prophage-like element
Whole-community read coverage
VLP-fraction read coverage
Contig position (bp)
Oscillospiraceae Lawsonibacter prophage

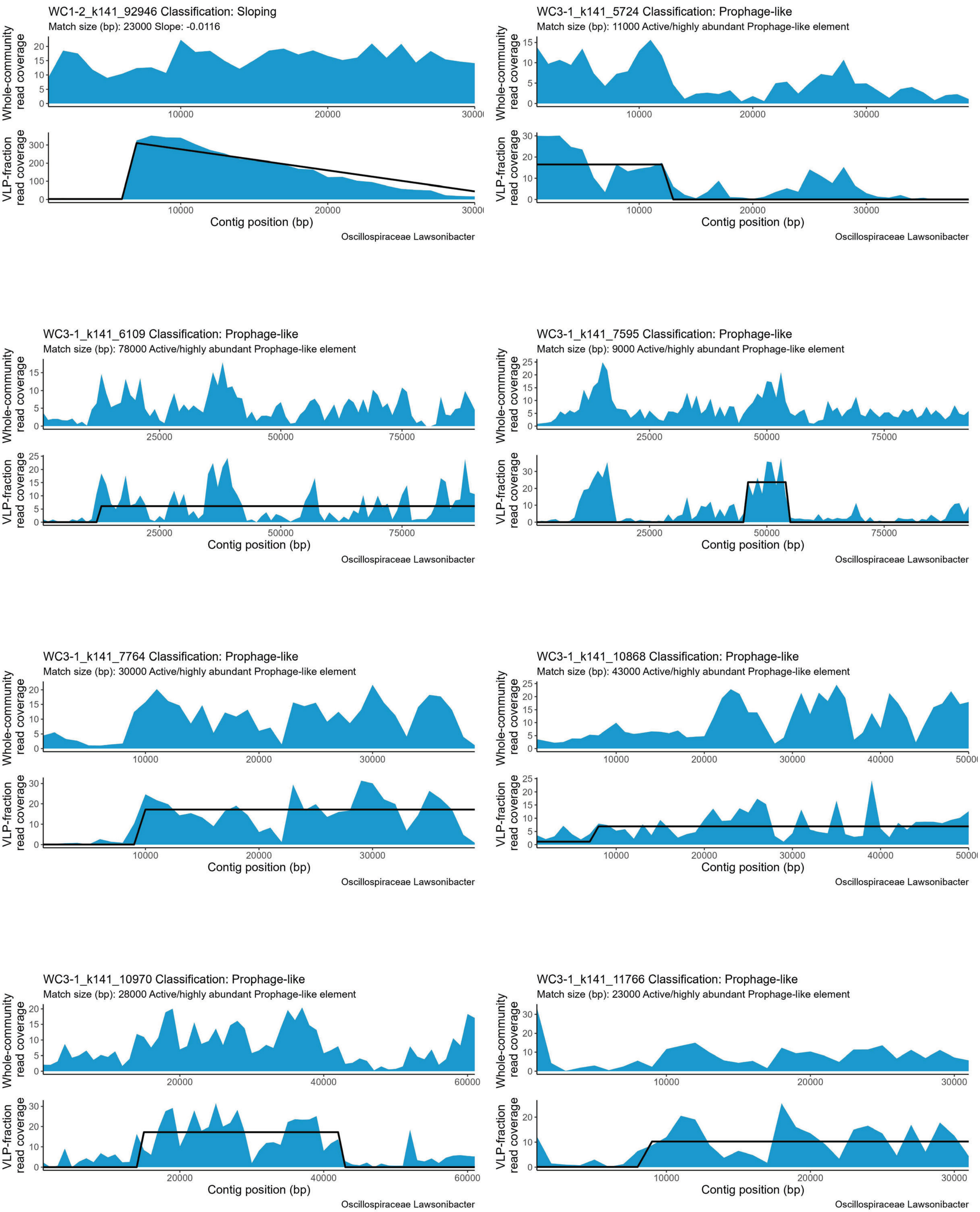
WC1-2_k141_92946 Classification: Sloping
Match size (bp): 23000 Slope: -0.0116
Whole-community read coverage
VLP-fraction read coverage
Contig position (bp)
Oscillospiraceae Lawsonibacter
WC3-1_k141_5724 Classification: Prophage-like
Match size (bp): 11000 Active/highly abundant Prophage-like element
Whole-community read coverage
VLP-fraction read coverage
Contig position (bp)
Oscillospiraceae Lawsonibacter
WC3-1_k141_6109 Classification: Prophage-like
Match size (bp): 78000 Active/highly abundant Prophage-like element
Whole-community read coverage
VLP-fraction read coverage
Contig position (bp)
Oscillospiraceae Lawsonibacter
WC3-1_k141_7595 Classification: Prophage-like
Match size (bp): 9000 Active/highly abundant Prophage-like element
Whole-community read coverage
VLP-fraction read coverage
Contig position (bp)
Oscillospiraceae Lawsonibacter
WC3-1_k141_7764 Classification: Prophage-like
Match size (bp): 30000 Active/highly abundant Prophage-like element
Whole-community read coverage
VLP-fraction read coverage
Contig position (bp)
Oscillospiraceae Lawsonibacter
WC3-1_k141_10868 Classification: Prophage-like
Match size (bp): 43000 Active/highly abundant Prophage-like element
Whole-community read coverage
VLP-fraction read coverage
Contig position (bp)
Oscillospiraceae Lawsonibacter
WC3-1_k141_10970 Classification: Prophage-like
Match size (bp): 28000 Active/highly abundant Prophage-like element
Whole-community read coverage
VLP-fraction read coverage
Contig position (bp)
Oscillospiraceae Lawsonibacter
WC3-1_k141_11766 Classification: Prophage-like
Match size (bp): 23000 Active/highly abundant Prophage-like element
Whole-community read coverage
VLP-fraction read coverage
Contig position (bp)
Oscillospiraceae Lawsonibacter

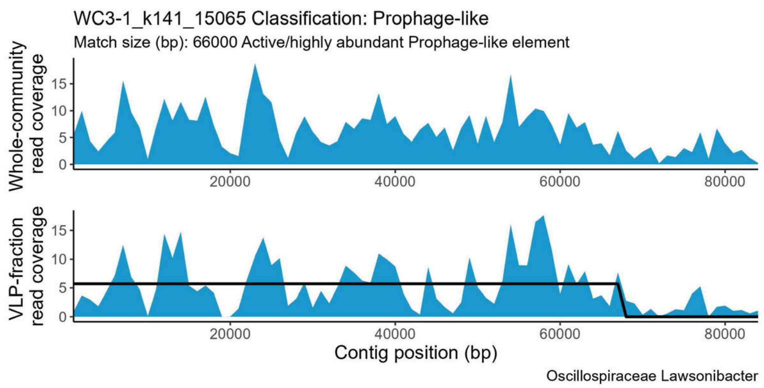
WC3-1_k141_15065 Classification: Prophage-like
Match size (bp): 66000 Active/highly abundant Prophage-like element
Whole-community read coverage
VLP-fraction read coverage
Contig position (bp)
Oscillospiraceae Lawsonibacter

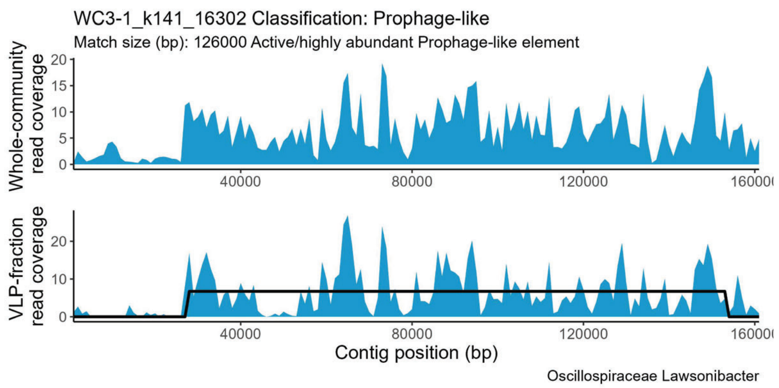
WC3-1_k141_16302 Classification: Prophage-like
Match size (bp): 126000 Active/highly abundant Prophage-like element
Whole-community read coverage
VLP-fraction read coverage
Contig position (bp)
Oscillospiraceae Lawsonibacter

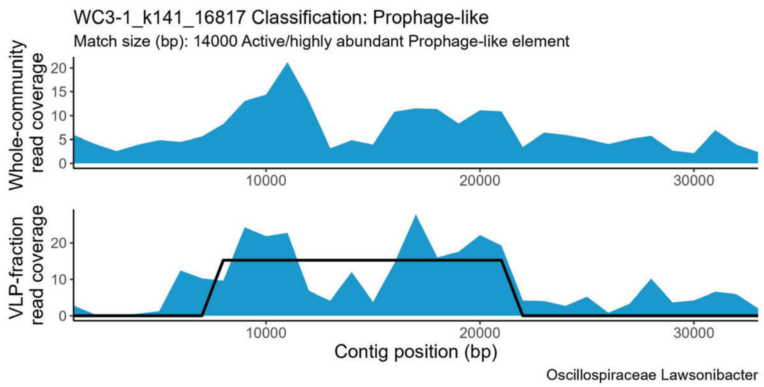
WC3-1_k141_16817 Classification: Prophage-like
Match size (bp): 14000 Active/highly abundant Prophage-like element
Whole-community read coverage
VLP-fraction read coverage
Contig position (bp)
Oscillospiraceae Lawsonibacter

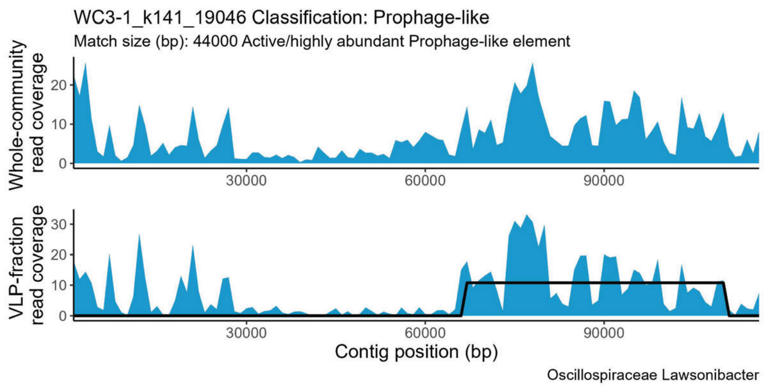
WC3-1_k141_19046 Classification: Prophage-like
Match size (bp): 44000 Active/highly abundant Prophage-like element
Whole-community read coverage
VLP-fraction read coverage
Contig position (bp)
Oscillospiraceae Lawsonibacter

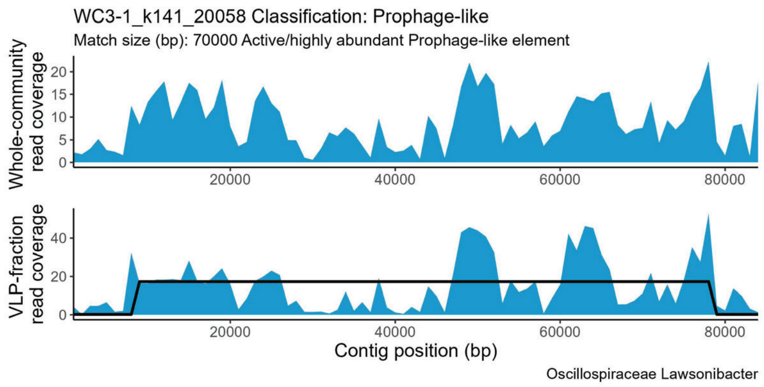
WC3-1_k141_20058 Classification: Prophage-like
Match size (bp): 70000 Active/highly abundant Prophage-like element
Whole-community read coverage
VLP-fraction read coverage
Contig position (bp)
Oscillospiraceae Lawsonibacter

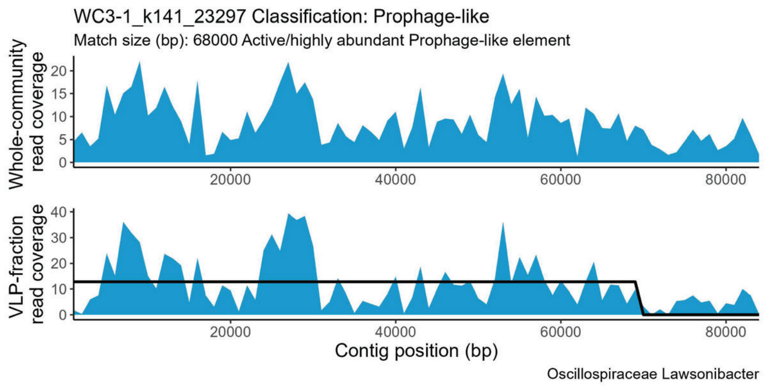
WC3-1_k141_23297 Classification: Prophage-like
Match size (bp): 68000 Active/highly abundant Prophage-like element
Whole-community read coverage
VLP-fraction read coverage
Contig position (bp)
Oscillospiraceae Lawsonibacter

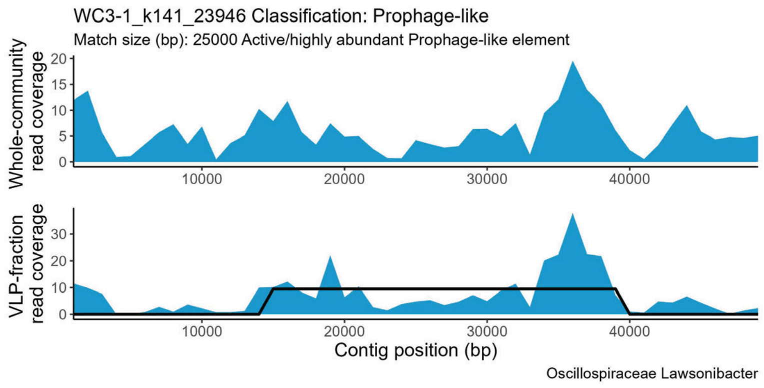
WC3-1_k141_23946 Classification: Prophage-like
Match size (bp): 25000 Active/highly abundant Prophage-like element
Whole-community read coverage
VLP-fraction read coverage
Contig position (bp)
Oscillospiraceae Lawsonibacter

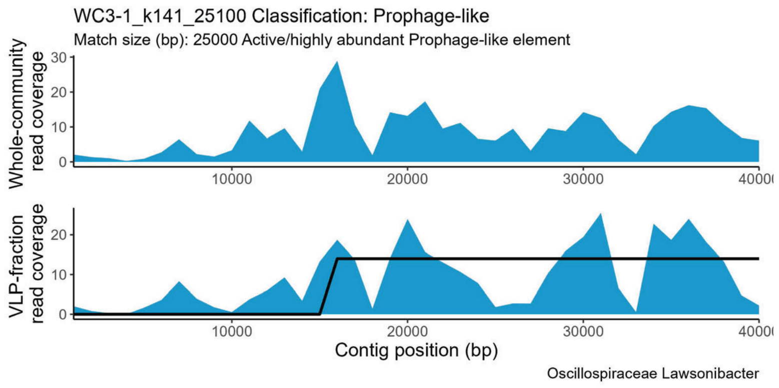
WC3-1_k141_25100 Classification: Prophage-like
Match size (bp): 25000 Active/highly abundant Prophage-like element
Whole-community read coverage
VLP-fraction read coverage
Contig position (bp)
Oscillospiraceae Lawsonibacter

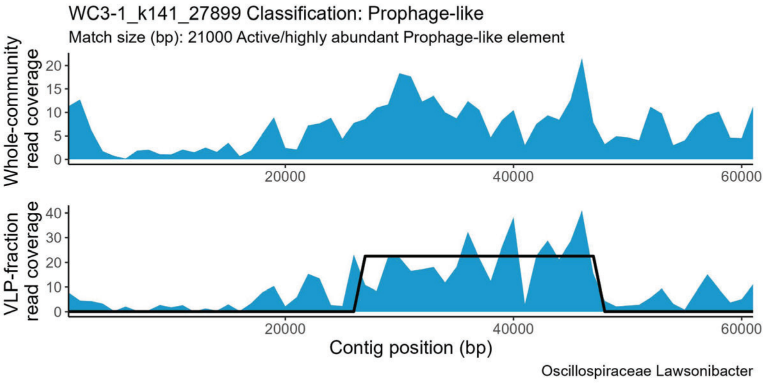
WC3-1_k141_27899 Classification: Prophage-like
Match size (bp): 21000 Active/highly abundant Prophage-like element
Whole-community read coverage
VLP-fraction read coverage
Contig position (bp)
Oscillospiraceae Lawsonibacter

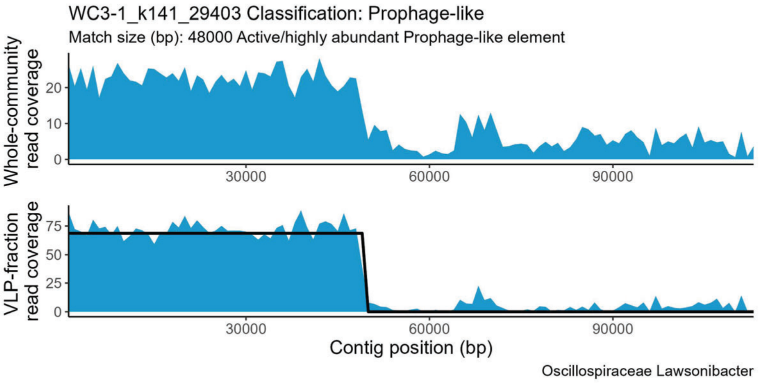
WC3-1_k141_29403 Classification: Prophage-like
Match size (bp): 48000 Active/highly abundant Prophage-like element
Whole-community read coverage
VLP-fraction read coverage
Contig position (bp)
Oscillospiraceae Lawsonibacter

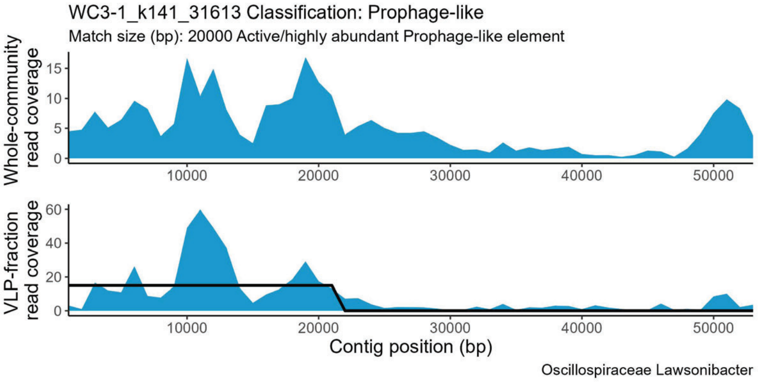
WC3-1_k141_31613 Classification: Prophage-like
Match size (bp): 20000 Active/highly abundant Prophage-like element
Whole-community read coverage
VLP-fraction read coverage
Contig position (bp)
Oscillospiraceae Lawsonibacter

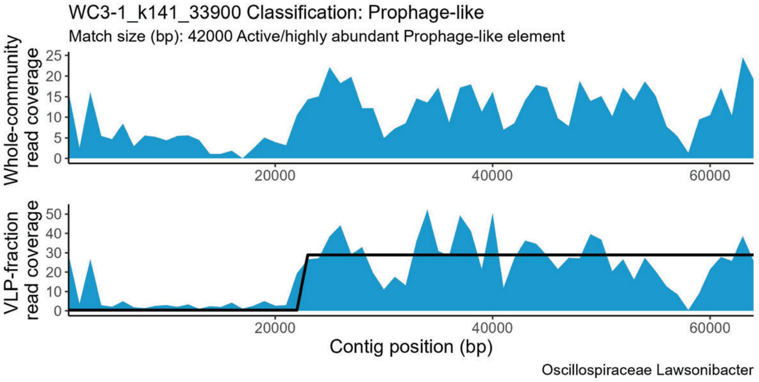
WC3-1_k141_33900 Classification: Prophage-like
Match size (bp): 42000 Active/highly abundant Prophage-like element
Whole-community read coverage
VLP-fraction read coverage
Contig position (bp)
Oscillospiraceae Lawsonibacter

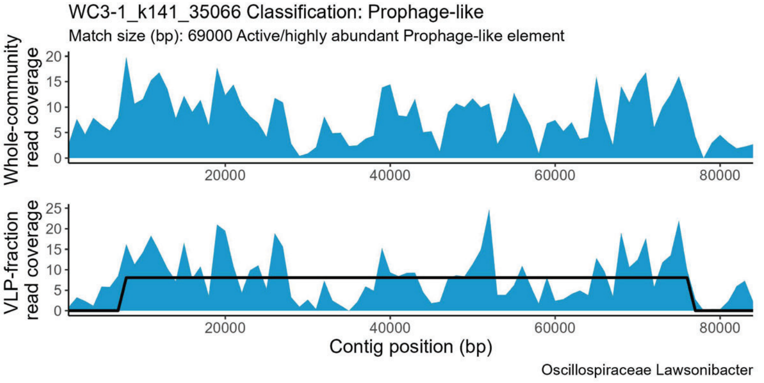
WC3-1_k141_35066 Classification: Prophage-like
Match size (bp): 69000 Active/highly abundant Prophage-like element
Whole-community read coverage
VLP-fraction read coverage
Contig position (bp)
Oscillospiraceae Lawsonibacter

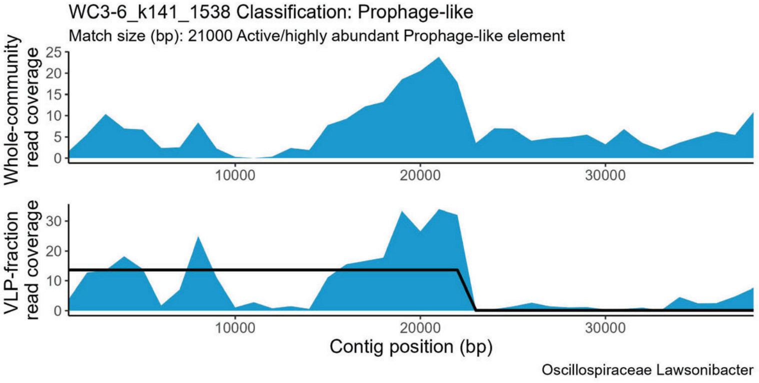
WC3-6_k141_1538 Classification: Prophage-like
Match size (bp): 21000 Active/highly abundant Prophage-like element
Whole-community read coverage
VLP-fraction read coverage
Contig position (bp)
Oscillospiraceae Lawsonibacter

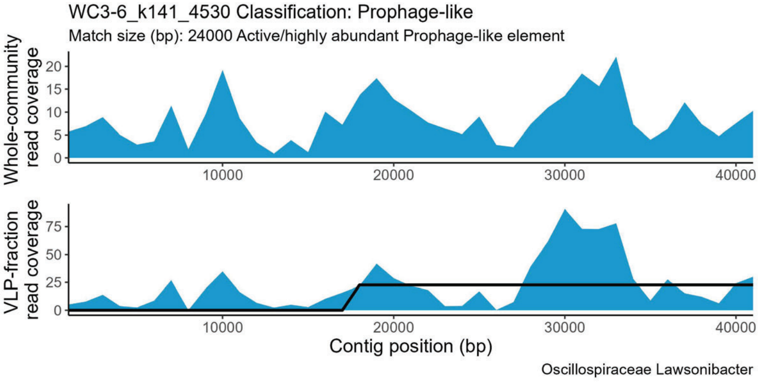
WC3-6_k141_4530 Classification: Prophage-like
Match size (bp): 24000 Active/highly abundant Prophage-like element
Whole-community read coverage
VLP-fraction read coverage
Contig position (bp)
Oscillospiraceae Lawsonibacter

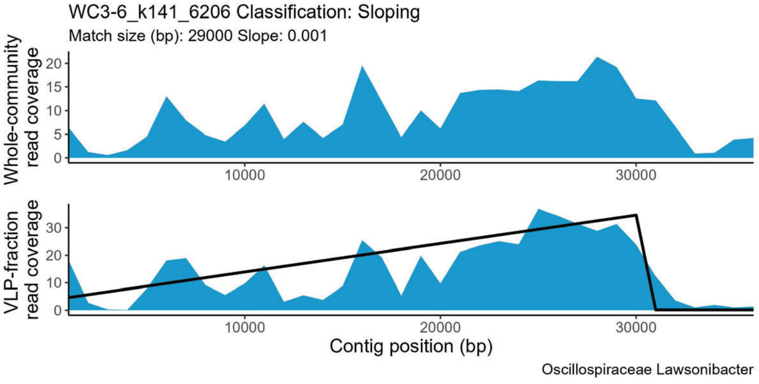
WC3-6_k141_6206 Classification: Sloping
Match size (bp): 29000 Slope: 0.001
Whole-community read coverage
VLP-fraction read coverage
Contig position (bp)
Oscillospiraceae Lawsonibacter

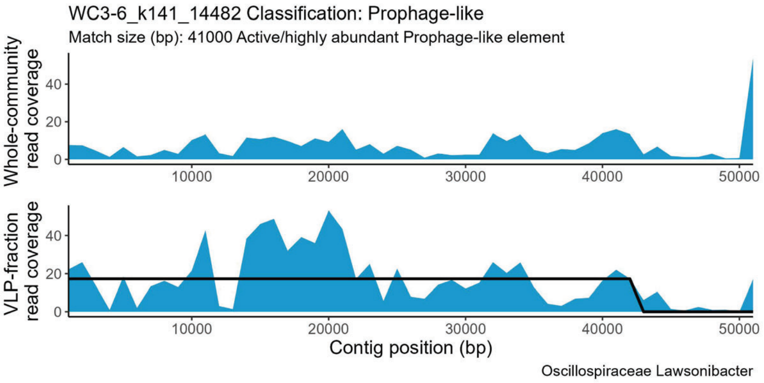
WC3-6_k141_14482 Classification: Prophage-like
Match size (bp): 41000 Active/highly abundant Prophage-like element
Whole-community read coverage
VLP-fraction read coverage
Contig position (bp)
Oscillospiraceae Lawsonibacter

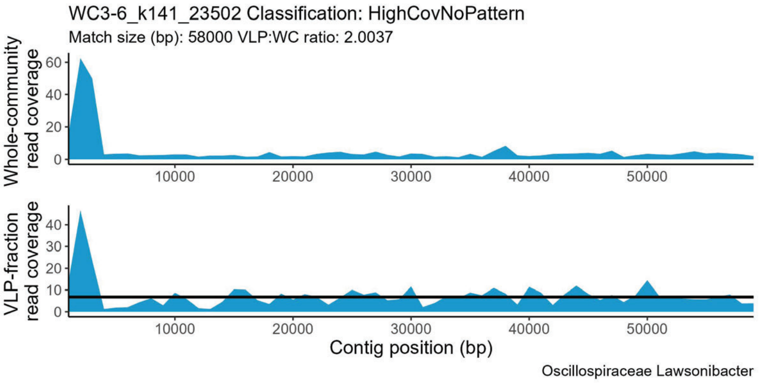
WC3-6_k141_23502 Classification: HighCovNoPattern
Match size (bp): 58000 VLP:WC ratio: 2.0037
Whole-community read coverage
VLP-fraction read coverage
Contig position (bp)
Oscillospiraceae Lawsonibacter

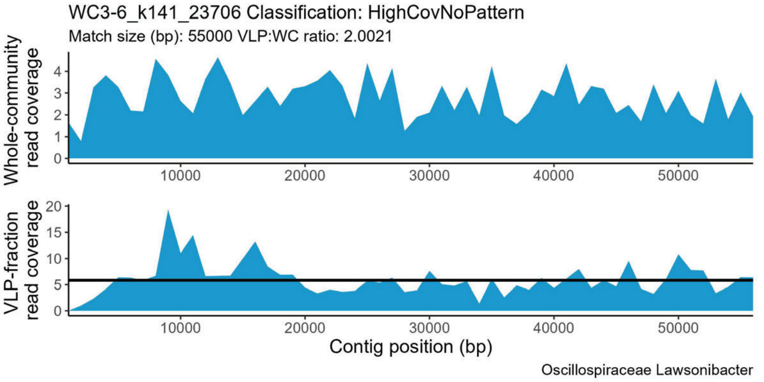
WC3-6_k141_23706 Classification: HighCovNoPattern
Match size (bp): 55000 VLP:WC ratio: 2.0021
Whole-community read coverage
VLP-fraction read coverage
Contig position (bp)
Oscillospiraceae Lawsonibacter

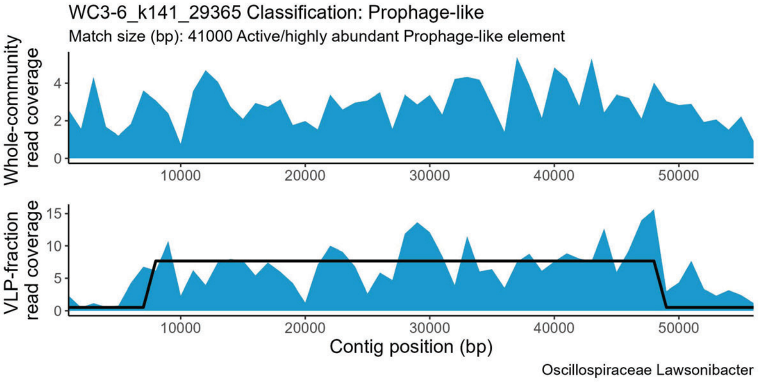
WC3-6_k141_29365 Classification: Prophage-like
Match size (bp): 41000 Active/highly abundant Prophage-like element
Whole-community read coverage
VLP-fraction read coverage
Contig position (bp)
Oscillospiraceae Lawsonibacter

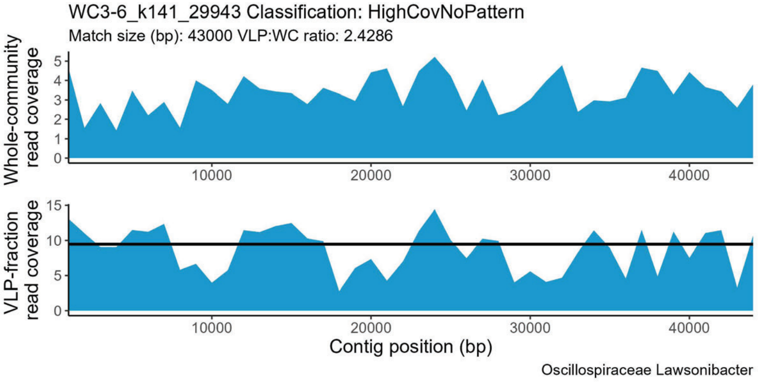
WC3-6_k141_29943 Classification: HighCovNoPattern
Match size (bp): 43000 VLP:WC ratio: 2.4286
Whole-community read coverage
VLP-fraction read coverage
Contig position (bp)
Oscillospiraceae Lawsonibacter

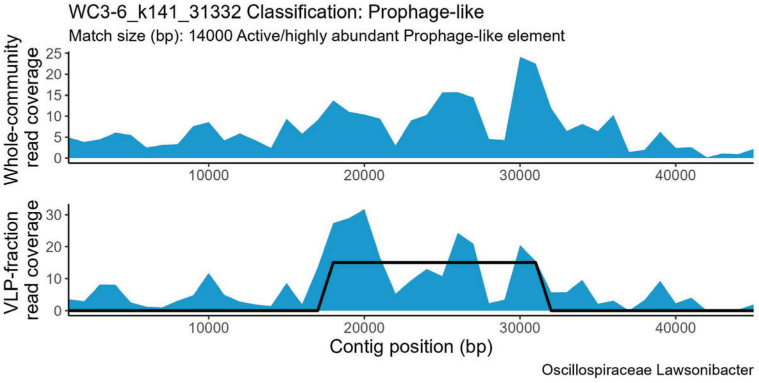
WC3-6_k141_31332 Classification: Prophage-like
Match size (bp): 14000 Active/highly abundant Prophage-like element
Whole-community read coverage
VLP-fraction read coverage
Contig position (bp)
Oscillospiraceae Lawsonibacter

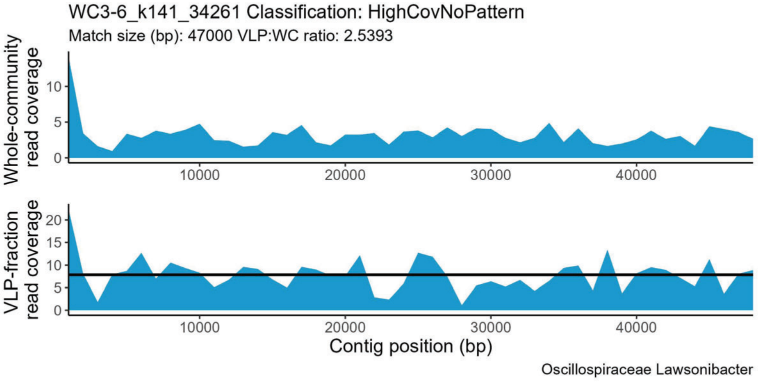
WC3-6_k141_34261 Classification: HighCovNoPattern
Match size (bp): 47000 VLP:WC ratio: 2.5393
Whole-community read coverage
VLP-fraction read coverage
Contig position (bp)
Oscillospiraceae Lawsonibacter

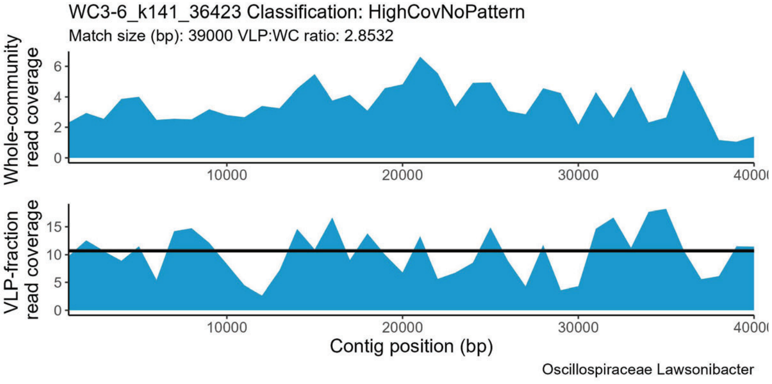
WC3-6_k141_36423 Classification: HighCovNoPattern
Match size (bp): 39000 VLP:WC ratio: 2.8532
Whole-community read coverage
VLP-fraction read coverage
Contig position (bp)
Oscillospiraceae Lawsonibacter

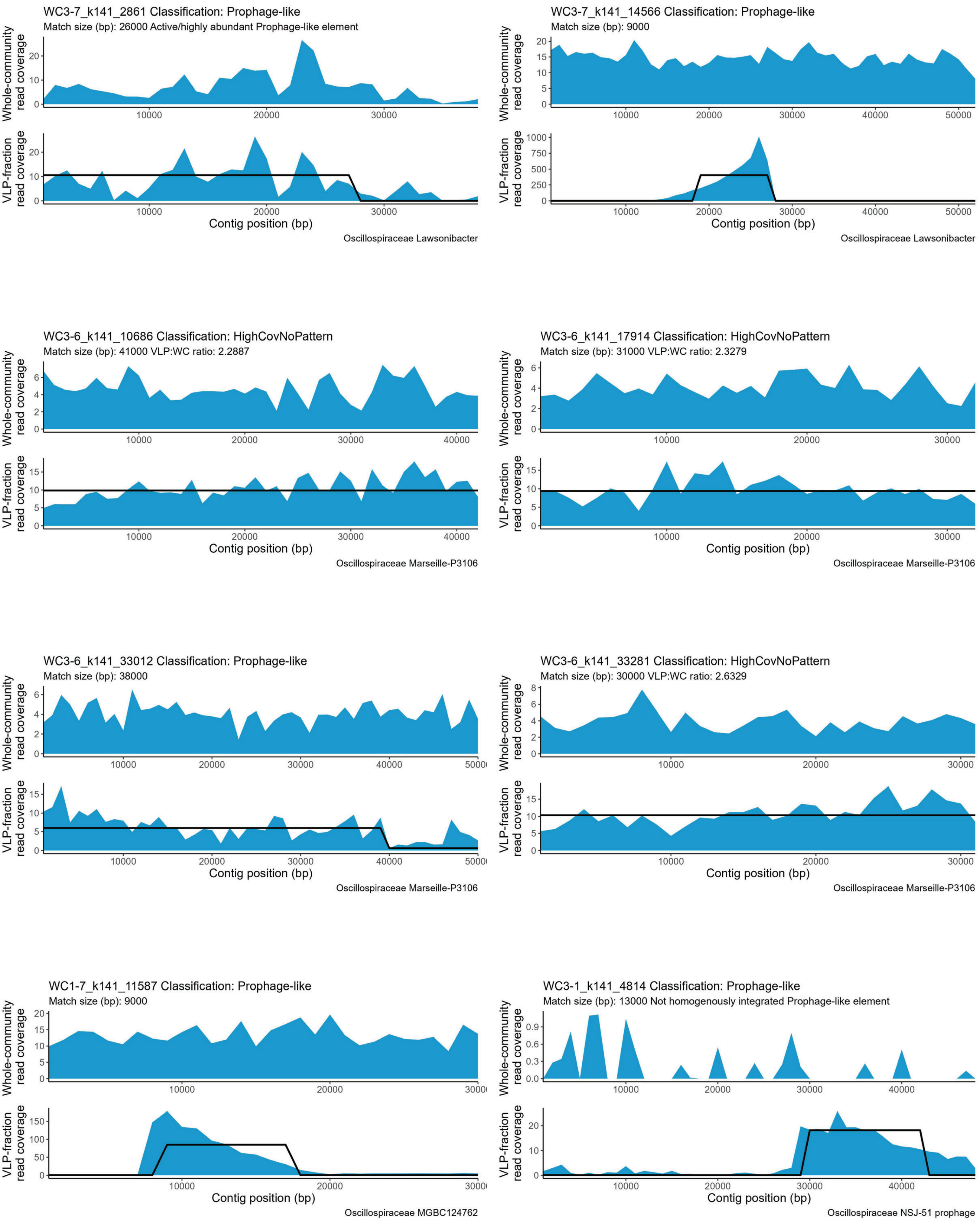
WC3-7_k141_2861 Classification: Prophage-like
Match size (bp): 26000 Active/highly abundant Prophage-like element
Oscillospiraceae Lawsonibacter
WC3-7_k141_14566 Classification: Prophage-like
Match size (bp): 9000
Oscillospiraceae Lawsonibacter
WC3-6_k141_10686 Classification: HighCovNoPattern
Match size (bp): 41000 VLP:WC ratio: 2.2887
Oscillospiraceae Marseille-P3106
WC3-6_k141_17914 Classification: HighCovNoPattern
Match size (bp): 31000 VLP:WC ratio: 2.3279
Oscillospiraceae Marseille-P3106
WC3-6_k141_33012 Classification: Prophage-like
Match size (bp): 38000
Oscillospiraceae Marseille-P3106
WC3-6_k141_33281 Classification: HighCovNoPattern
Match size (bp): 30000 VLP:WC ratio: 2.6329
Oscillospiraceae Marseille-P3106
WC1-7_k141_11587 Classification: Prophage-like
Match size (bp): 9000
Oscillospiraceae MGBC124762
WC3-1_k141_4814 Classification: Prophage-like
Match size (bp): 13000 Not homogenously integrated Prophage-like element
Oscillospiraceae NSJ-51 prophage
Whole-community read coverage
VLP-fraction read coverage
Contig position (bp)

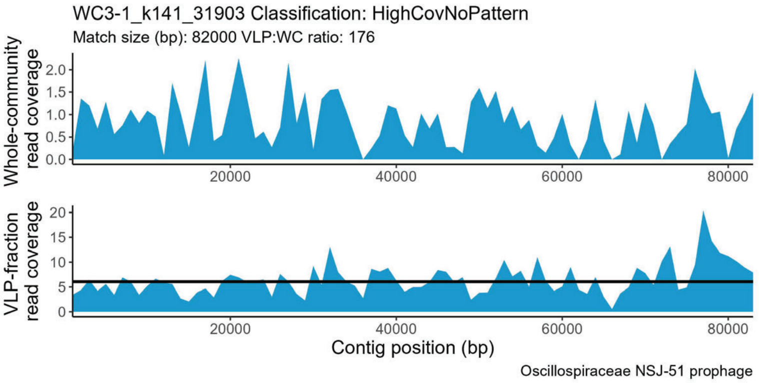
WC3-1_k141_31903 Classification: HighCovNoPattern
Match size (bp): 82000 VLP:WC ratio: 176
Whole-community read coverage
VLP-fraction read coverage
Contig position (bp)
Oscillospiraceae NSJ-51 prophage

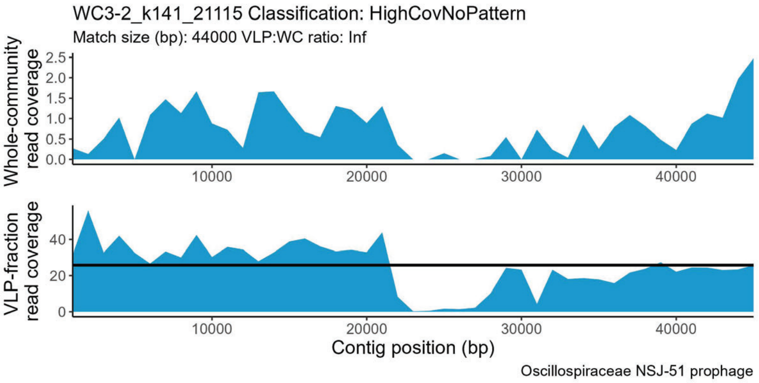
WC3-2_k141_21115 Classification: HighCovNoPattern
Match size (bp): 44000 VLP:WC ratio: Inf
Whole-community read coverage
VLP-fraction read coverage
Contig position (bp)
Oscillospiraceae NSJ-51 prophage

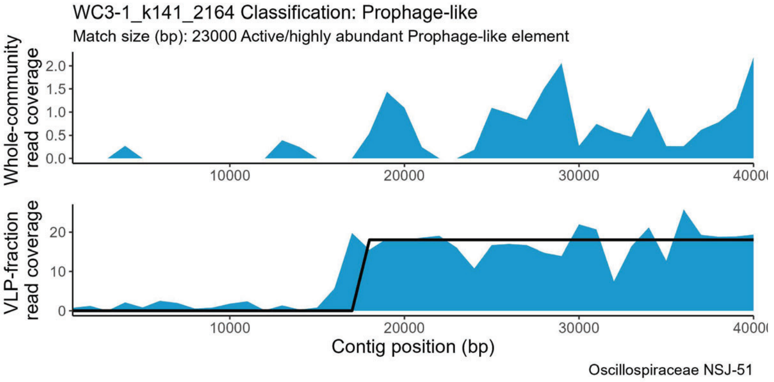
WC3-1_k141_2164 Classification: Prophage-like
Match size (bp): 23000 Active/highly abundant Prophage-like element
Whole-community read coverage
VLP-fraction read coverage
Contig position (bp)
Oscillospiraceae NSJ-51

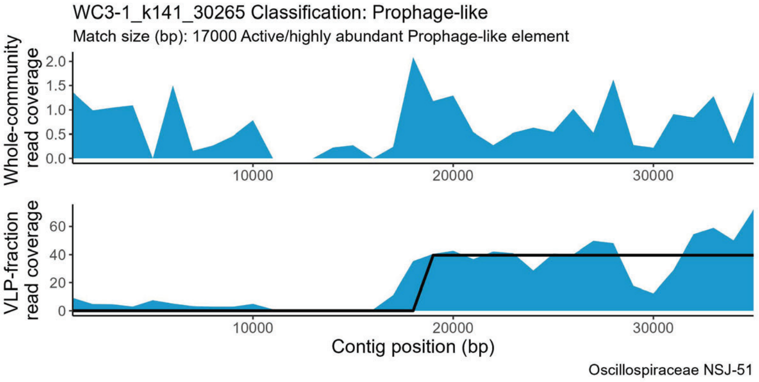
WC3-1_k141_30265 Classification: Prophage-like
Match size (bp): 17000 Active/highly abundant Prophage-like element
Whole-community read coverage
VLP-fraction read coverage
Contig position (bp)
Oscillospiraceae NSJ-51

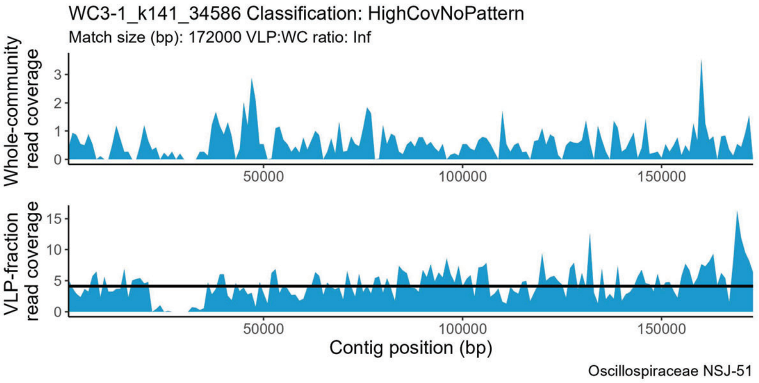
WC3-1_k141_34586 Classification: HighCovNoPattern
Match size (bp): 172000 VLP:WC ratio: Inf
Whole-community read coverage
VLP-fraction read coverage
Contig position (bp)
Oscillospiraceae NSJ-51

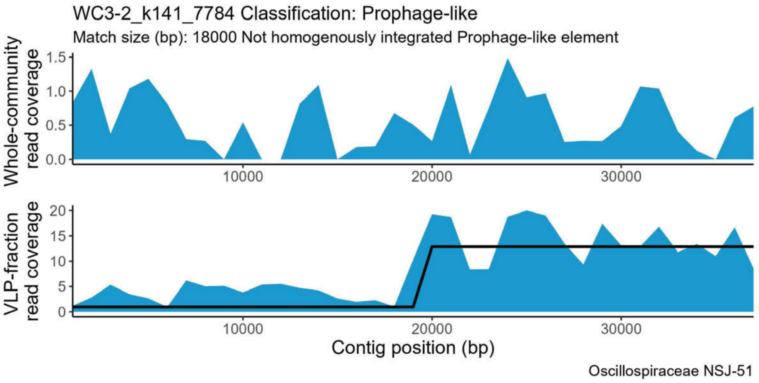
WC3-2_k141_7784 Classification: Prophage-like
Match size (bp): 18000 Not homogenously integrated Prophage-like element
Whole-community read coverage
VLP-fraction read coverage
Contig position (bp)
Oscillospiraceae NSJ-51

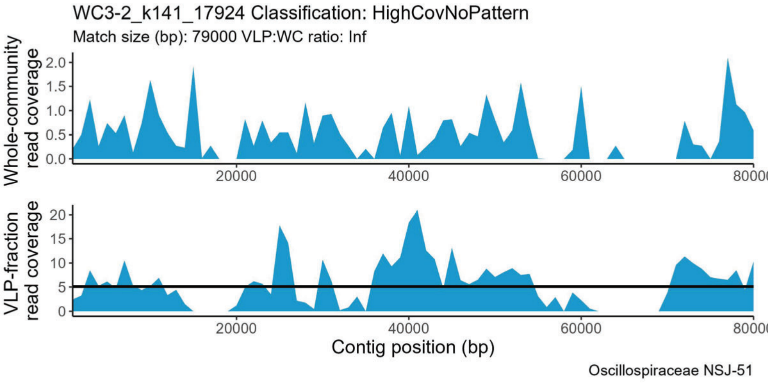
WC3-2_k141_17924 Classification: HighCovNoPattern
Match size (bp): 79000 VLP:WC ratio: Inf
Whole-community read coverage
VLP-fraction read coverage
Contig position (bp)
Oscillospiraceae NSJ-51

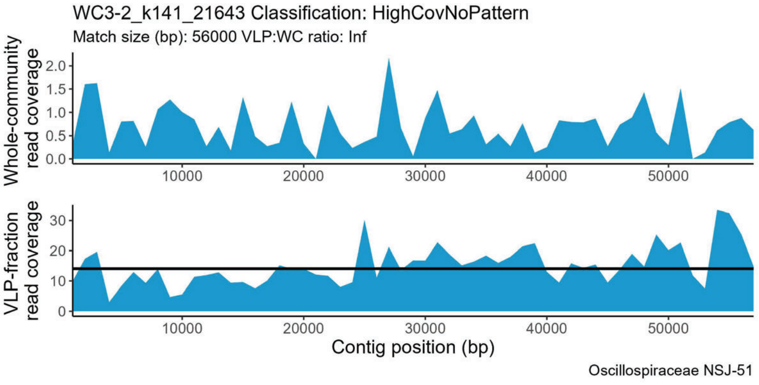
WC3-2_k141_21643 Classification: HighCovNoPattern
Match size (bp): 56000 VLP:WC ratio: Inf
Whole-community read coverage
VLP-fraction read coverage
Contig position (bp)
Oscillospiraceae NSJ-51

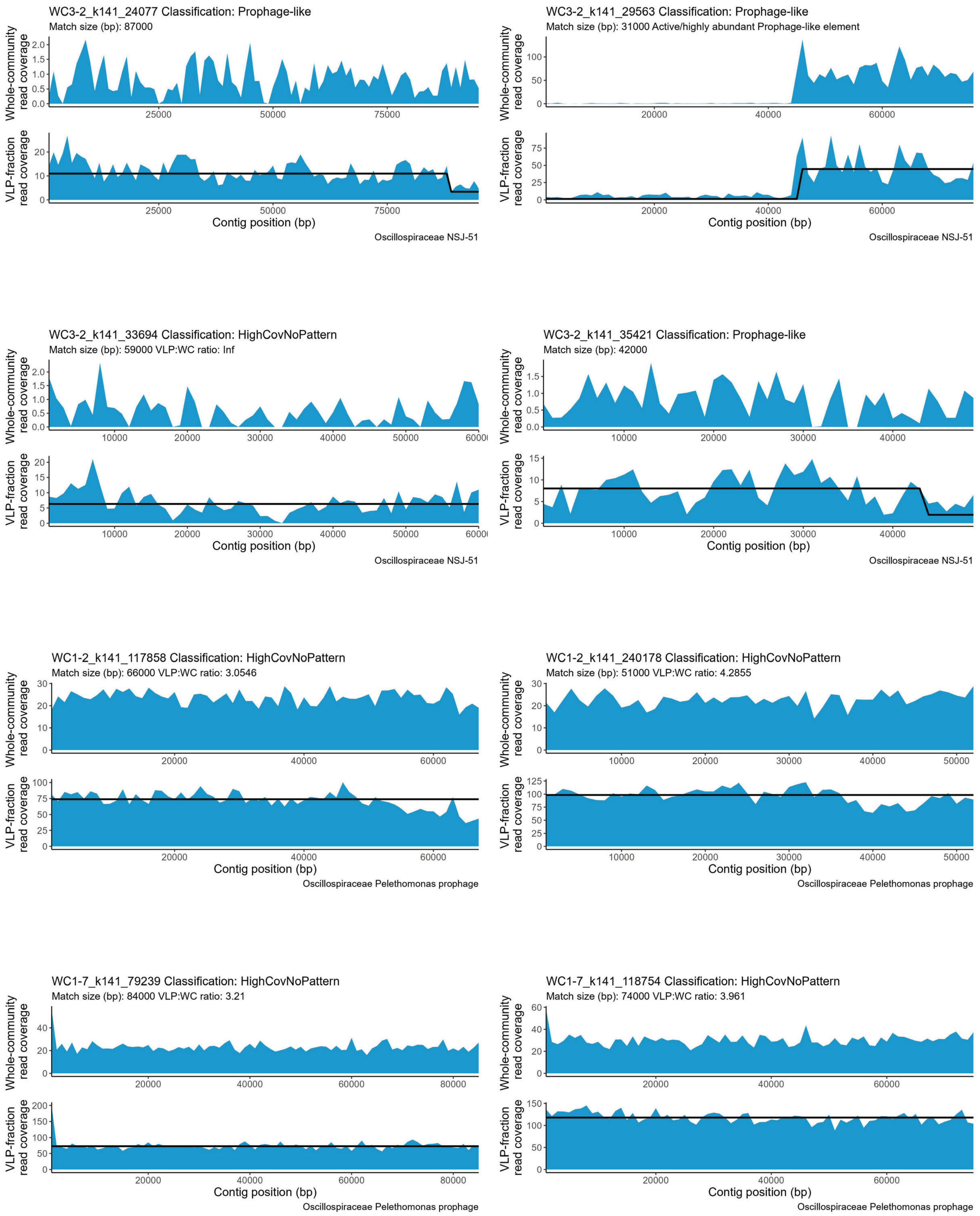
WC3-2_k141_24077 Classification: Prophage-like
Match size (bp): 87000
Whole-community read coverage
VLP-fraction read coverage
Contig position (bp)
Oscillospiraceae NSJ-51
WC3-2_k141_29563 Classification: Prophage-like
Match size (bp): 31000 Active/highly abundant Prophage-like element
Whole-community read coverage
VLP-fraction read coverage
Contig position (bp)
Oscillospiraceae NSJ-51
WC3-2_k141_33694 Classification: HighCovNoPattern
Match size (bp): 59000 VLP:WC ratio: Inf
Whole-community read coverage
VLP-fraction read coverage
Contig position (bp)
Oscillospiraceae NSJ-51
WC3-2_k141_35421 Classification: Prophage-like
Match size (bp): 42000
Whole-community read coverage
VLP-fraction read coverage
Contig position (bp)
Oscillospiraceae NSJ-51
WC1-2_k141_117858 Classification: HighCovNoPattern
Match size (bp): 66000 VLP:WC ratio: 3.0546
Whole-community read coverage
VLP-fraction read coverage
Contig position (bp)
Oscillospiraceae Pelethomonas prophage
WC1-2_k141_240178 Classification: HighCovNoPattern
Match size (bp): 51000 VLP:WC ratio: 4.2855
Whole-community read coverage
VLP-fraction read coverage
Contig position (bp)
Oscillospiraceae Pelethomonas prophage
WC1-7_k141_79239 Classification: HighCovNoPattern
Match size (bp): 84000 VLP:WC ratio: 3.21
Whole-community read coverage
VLP-fraction read coverage
Contig position (bp)
Oscillospiraceae Pelethomonas prophage
WC1-7_k141_118754 Classification: HighCovNoPattern
Match size (bp): 74000 VLP:WC ratio: 3.961
Whole-community read coverage
VLP-fraction read coverage
Contig position (bp)
Oscillospiraceae Pelethomonas prophage

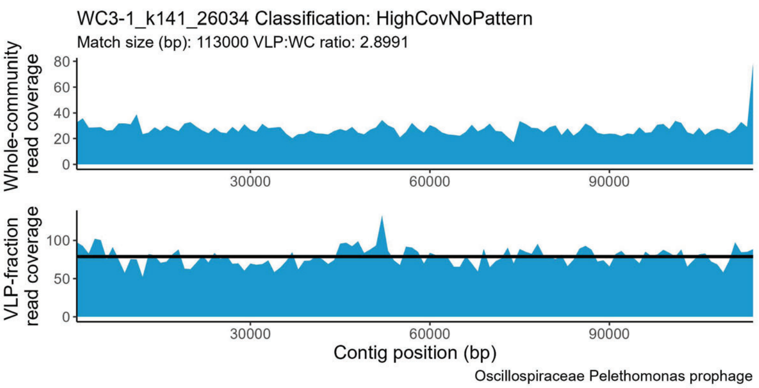
WC3-1_k141_26034 Classification: HighCovNoPattern
Match size (bp): 113000 VLP:WC ratio: 2.8991
Whole-community read coverage
VLP-fraction read coverage
Contig position (bp)
Oscillospiraceae Pelethomonas prophage

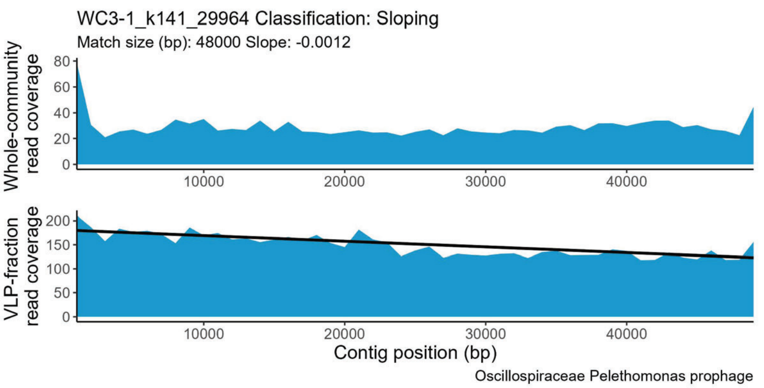
WC3-1_k141_29964 Classification: Sloping
Match size (bp): 48000 Slope: -0.0012
Whole-community read coverage
VLP-fraction read coverage
Contig position (bp)
Oscillospiraceae Pelethomonas prophage

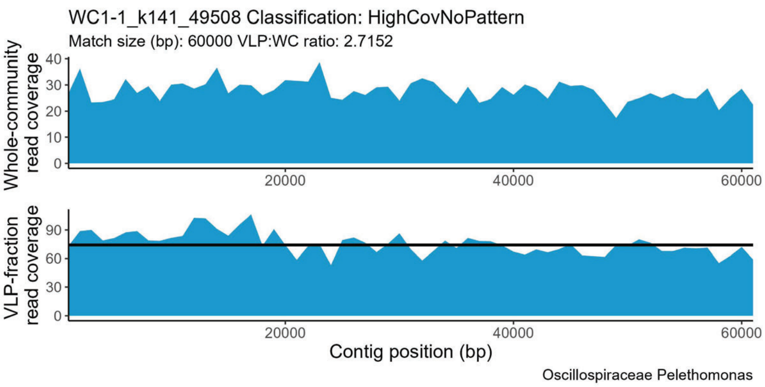
WC1-1_k141_49508 Classification: HighCovNoPattern
Match size (bp): 60000 VLP:WC ratio: 2.7152
Whole-community read coverage
VLP-fraction read coverage
Contig position (bp)
Oscillospiraceae Pelethomonas

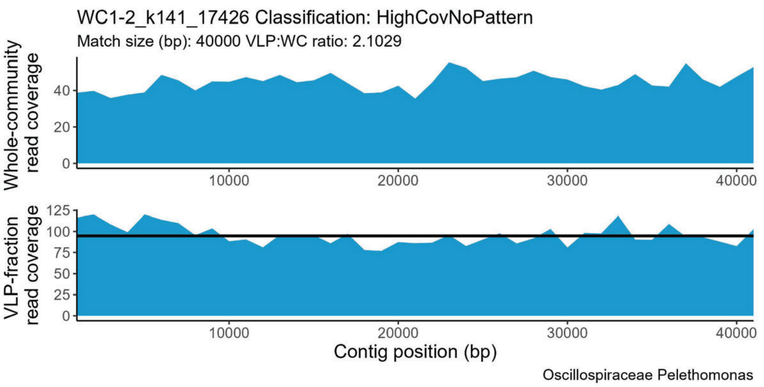
WC1-2_k141_17426 Classification: HighCovNoPattern
Match size (bp): 40000 VLP:WC ratio: 2.1029
Whole-community read coverage
VLP-fraction read coverage
Contig position (bp)
Oscillospiraceae Pelethomonas

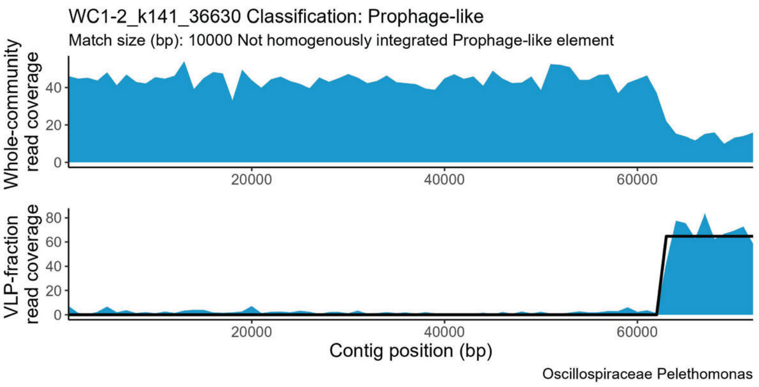
WC1-2_k141_36630 Classification: Prophage-like
Match size (bp): 10000 Not homogenously integrated Prophage-like element
Whole-community read coverage
VLP-fraction read coverage
Contig position (bp)
Oscillospiraceae Pelethomonas

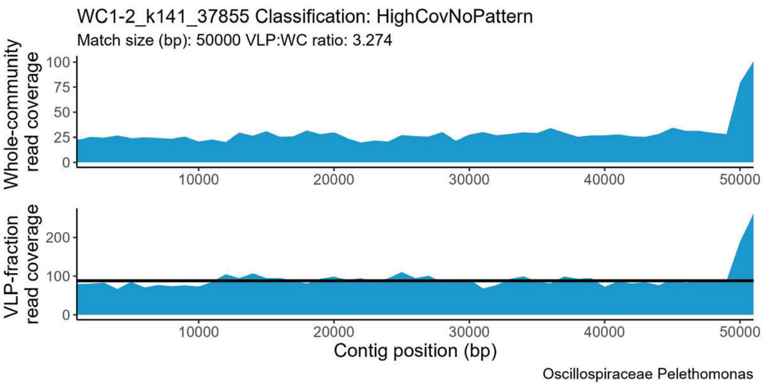
WC1-2_k141_37855 Classification: HighCovNoPattern
Match size (bp): 50000 VLP:WC ratio: 3.274
Whole-community read coverage
VLP-fraction read coverage
Contig position (bp)
Oscillospiraceae Pelethomonas

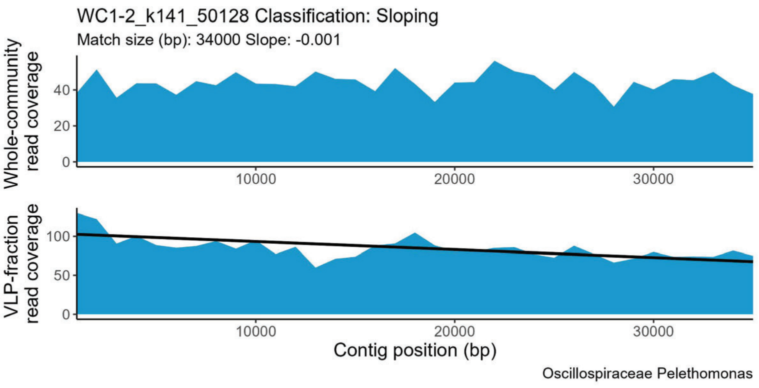
WC1-2_k141_50128 Classification: Sloping
Match size (bp): 34000 Slope: -0.001
Whole-community read coverage
VLP-fraction read coverage
Contig position (bp)
Oscillospiraceae Pelethomonas

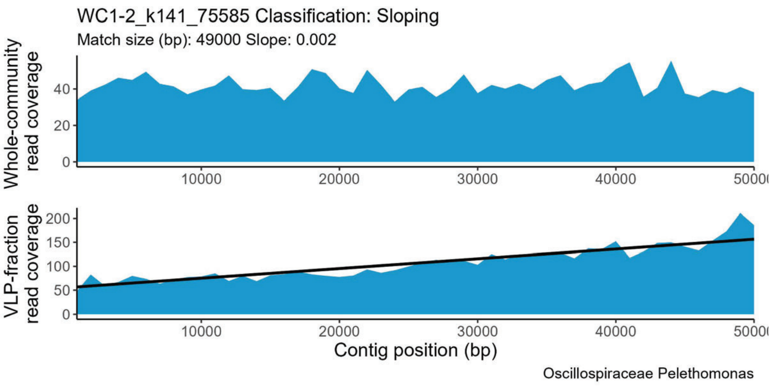
WC1-2_k141_75585 Classification: Sloping
Match size (bp): 49000 Slope: 0.002
Whole-community read coverage
VLP-fraction read coverage
Contig position (bp)
Oscillospiraceae Pelethomonas

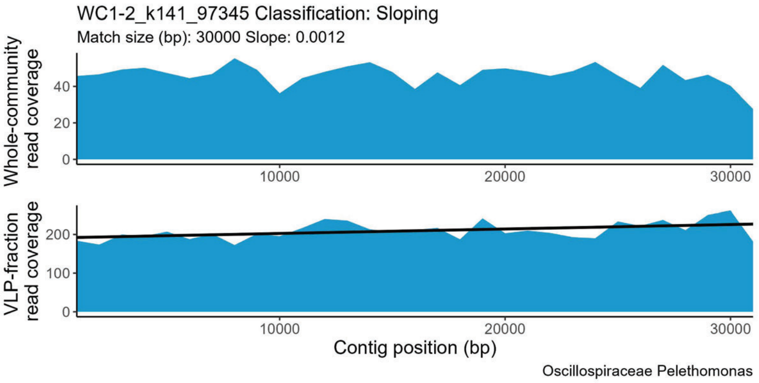
WC1-2_k141_97345 Classification: Sloping
Match size (bp): 30000 Slope: 0.0012
Whole-community read coverage
VLP-fraction read coverage
Contig position (bp)
Oscillospiraceae Pelethomonas

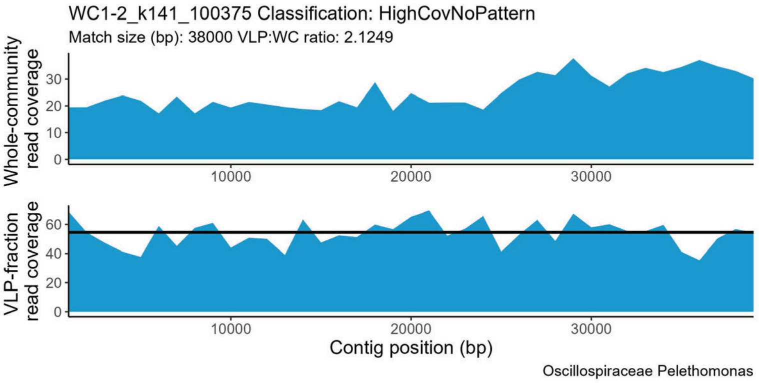
WC1-2_k141_100375 Classification: HighCovNoPattern
Match size (bp): 38000 VLP:WC ratio: 2.1249
Whole-community read coverage
VLP-fraction read coverage
Contig position (bp)
Oscillospiraceae Pelethomonas

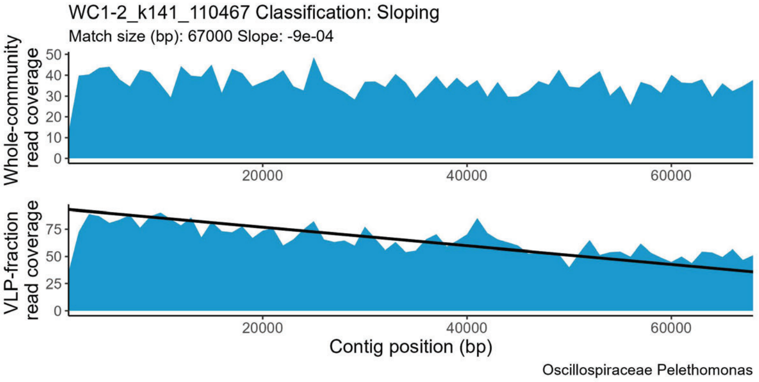
WC1-2_k141_110467 Classification: Sloping
Match size (bp): 67000 Slope: -9e-04
Whole-community read coverage
VLP-fraction read coverage
Contig position (bp)
Oscillospiraceae Pelethomonas

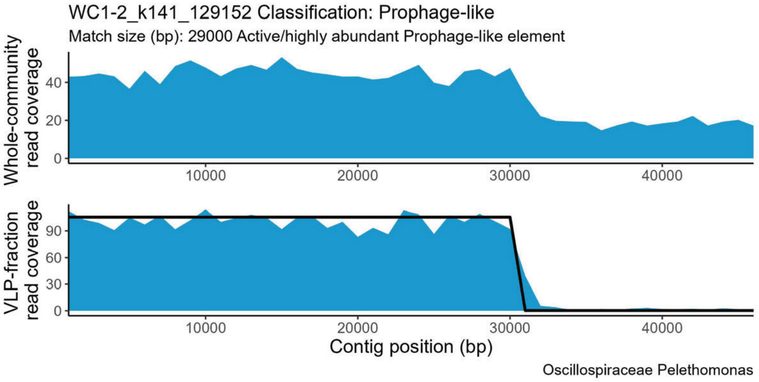
WC1-2_k141_129152 Classification: Prophage-like
Match size (bp): 29000 Active/highly abundant Prophage-like element
Whole-community read coverage
VLP-fraction read coverage
Contig position (bp)
Oscillospiraceae Pelethomonas

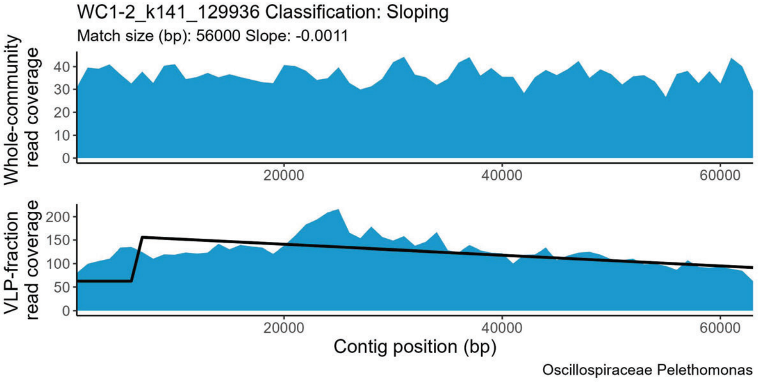
WC1-2_k141_129936 Classification: Sloping
Match size (bp): 56000 Slope: -0.0011
Whole-community read coverage
VLP-fraction read coverage
Contig position (bp)
Oscillospiraceae Pelethomonas

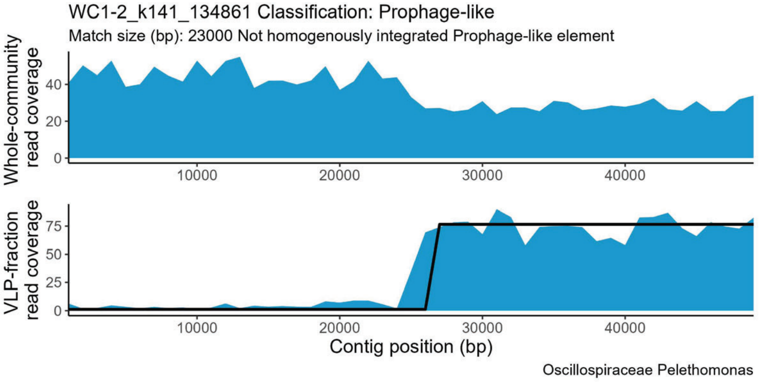
WC1-2_k141_134861 Classification: Prophage-like
Match size (bp): 23000 Not homogenously integrated Prophage-like element
Whole-community read coverage
VLP-fraction read coverage
Contig position (bp)
Oscillospiraceae Pelethomonas

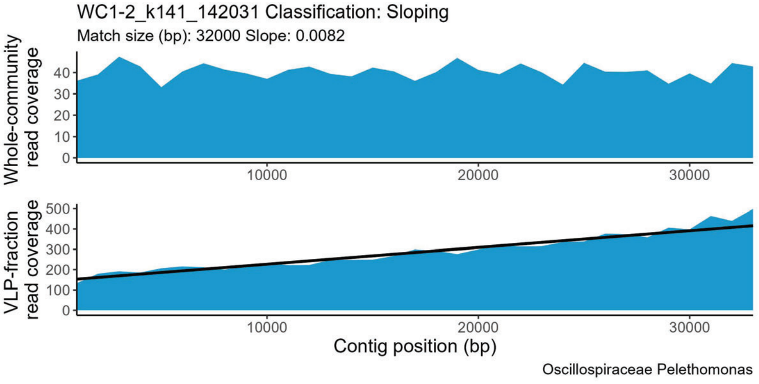
WC1-2_k141_142031 Classification: Sloping
Match size (bp): 32000 Slope: 0.0082
Whole-community read coverage
VLP-fraction read coverage
Contig position (bp)
Oscillospiraceae Pelethomonas

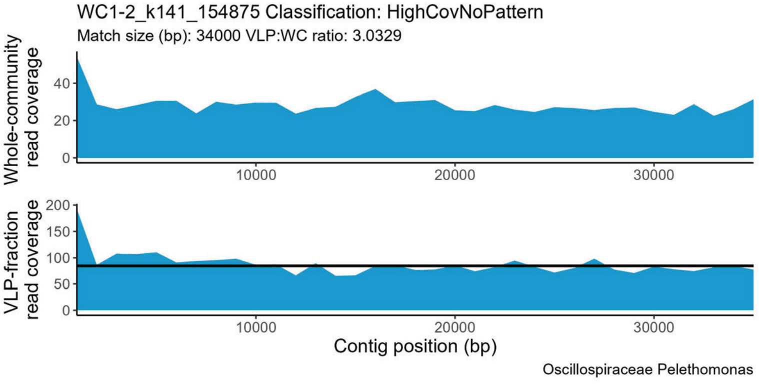
WC1-2_k141_154875 Classification: HighCovNoPattern
Match size (bp): 34000 VLP:WC ratio: 3.0329
Whole-community read coverage
VLP-fraction read coverage
Contig position (bp)
Oscillospiraceae Pelethomonas

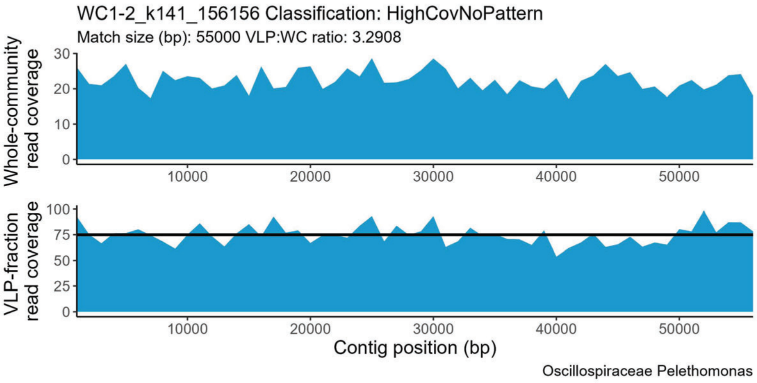
WC1-2_k141_156156 Classification: HighCovNoPattern
Match size (bp): 55000 VLP:WC ratio: 3.2908
Whole-community read coverage
VLP-fraction read coverage
Contig position (bp)
Oscillospiraceae Pelethomonas

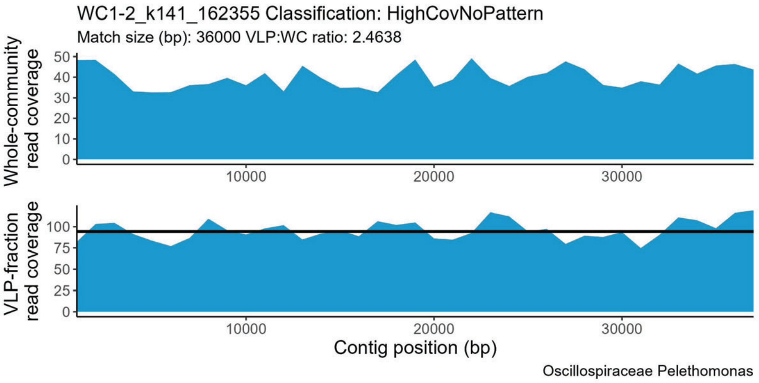
WC1-2_k141_162355 Classification: HighCovNoPattern
Match size (bp): 36000 VLP:WC ratio: 2.4638
Whole-community read coverage
VLP-fraction read coverage
Contig position (bp)
Oscillospiraceae Pelethomonas

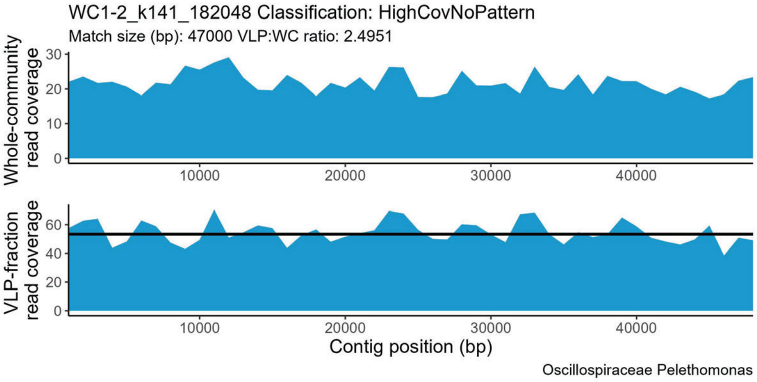
WC1-2_k141_182048 Classification: HighCovNoPattern
Match size (bp): 47000 VLP:WC ratio: 2.4951
Whole-community read coverage
VLP-fraction read coverage
Contig position (bp)
Oscillospiraceae Pelethomonas

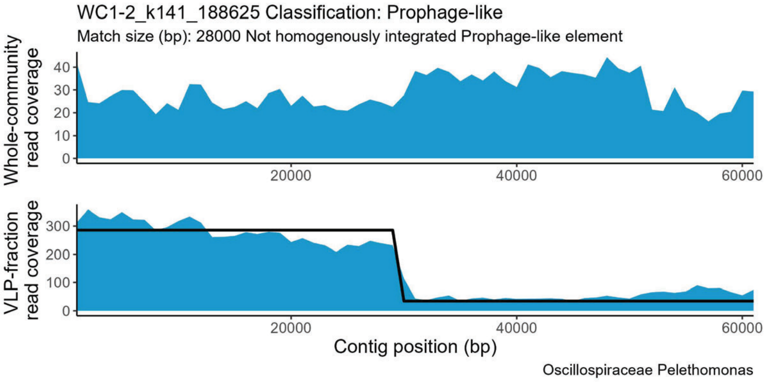
WC1-2_k141_188625 Classification: Prophage-like
Match size (bp): 28000 Not homogenously integrated Prophage-like element
Whole-community read coverage
VLP-fraction read coverage
Contig position (bp)
Oscillospiraceae Pelethomonas

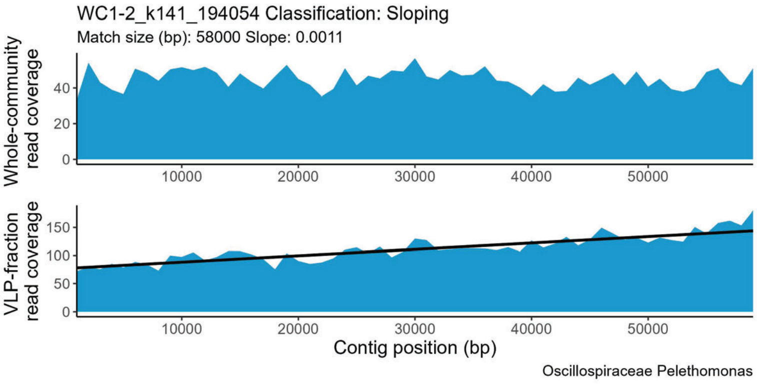
WC1-2_k141_194054 Classification: Sloping
Match size (bp): 58000 Slope: 0.0011
Whole-community read coverage
VLP-fraction read coverage
Contig position (bp)
Oscillospiraceae Pelethomonas

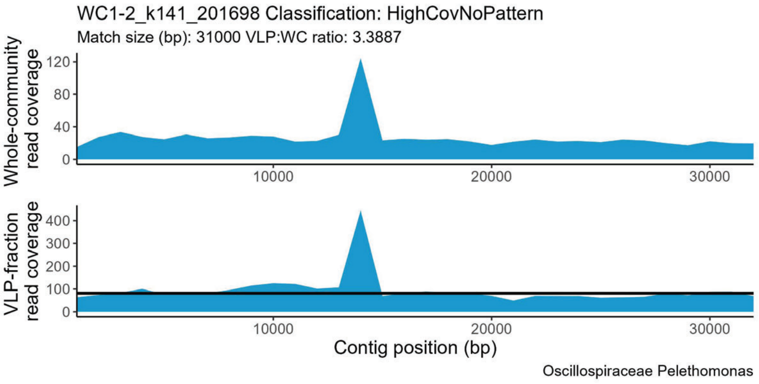
WC1-2_k141_201698 Classification: HighCovNoPattern
Match size (bp): 31000 VLP:WC ratio: 3.3887
Whole-community read coverage
VLP-fraction read coverage
Contig position (bp)
Oscillospiraceae Pelethomonas

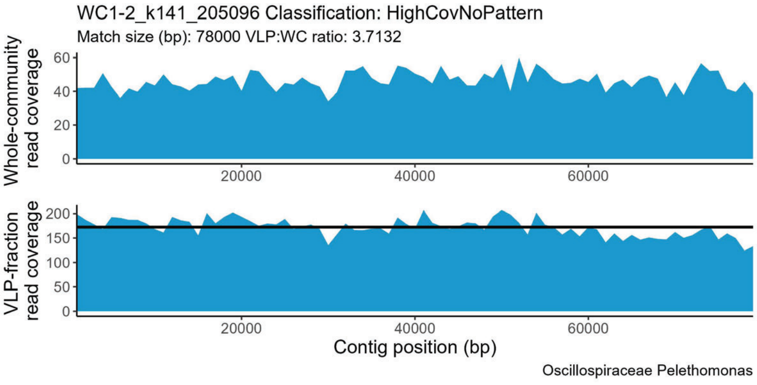
WC1-2_k141_205096 Classification: HighCovNoPattern
Match size (bp): 78000 VLP:WC ratio: 3.7132
Whole-community read coverage
VLP-fraction read coverage
Contig position (bp)
Oscillospiraceae Pelethomonas

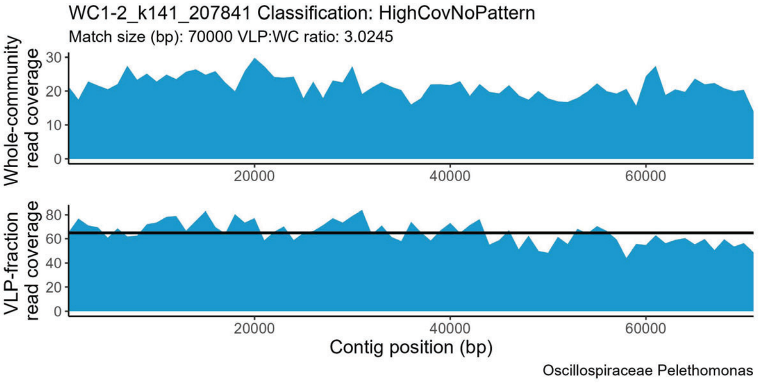
WC1-2_k141_207841 Classification: HighCovNoPattern
Match size (bp): 70000 VLP:WC ratio: 3.0245
Whole-community read coverage
VLP-fraction read coverage
Contig position (bp)
Oscillospiraceae Pelethomonas

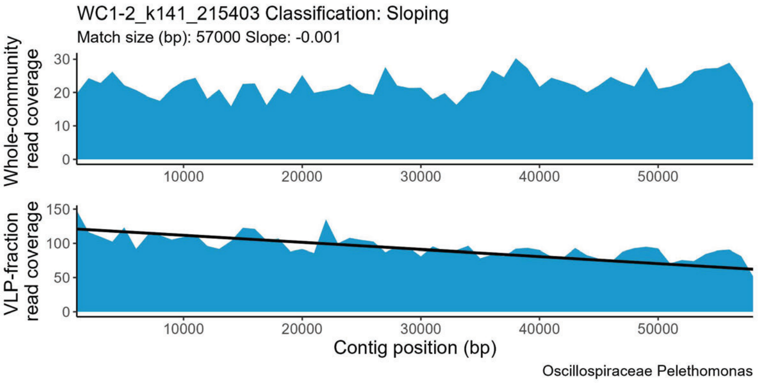
WC1-2_k141_215403 Classification: Sloping
Match size (bp): 57000 Slope: -0.001
Whole-community read coverage
VLP-fraction read coverage
Contig position (bp)
Oscillospiraceae Pelethomonas

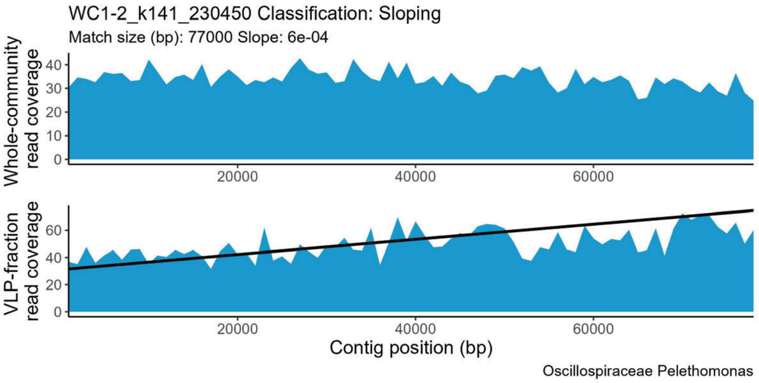
WC1-2_k141_230450 Classification: Sloping
Match size (bp): 77000 Slope: 6e-04
Whole-community read coverage
VLP-fraction read coverage
Contig position (bp)
Oscillospiraceae Pelethomonas

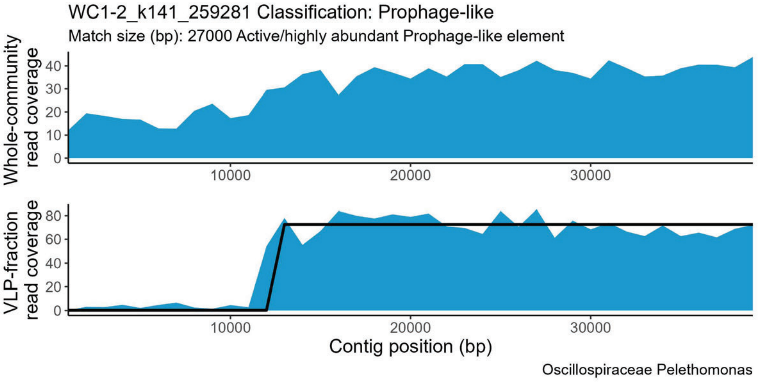
WC1-2_k141_259281 Classification: Prophage-like
Match size (bp): 27000 Active/highly abundant Prophage-like element
Whole-community read coverage
VLP-fraction read coverage
Contig position (bp)
Oscillospiraceae Pelethomonas

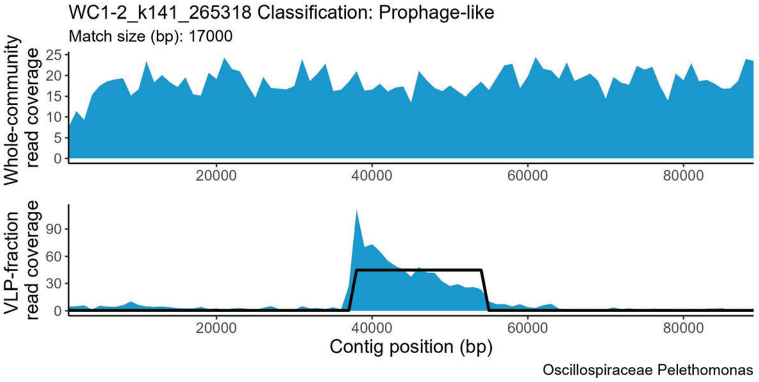
WC1-2_k141_265318 Classification: Prophage-like
Match size (bp): 17000
Whole-community read coverage
VLP-fraction read coverage
Contig position (bp)
Oscillospiraceae Pelethomonas

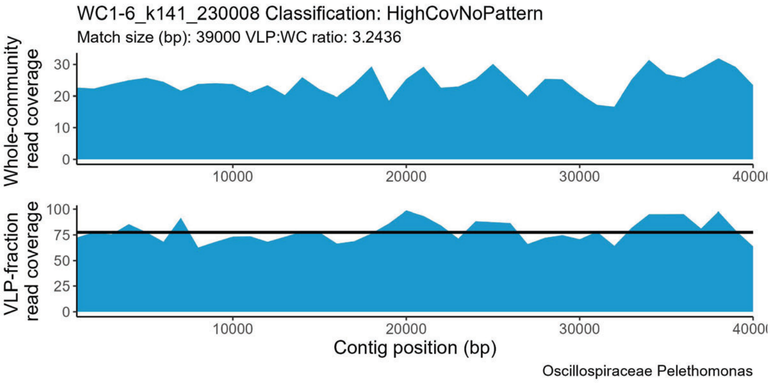
WC1-6_k141_230008 Classification: HighCovNoPattern
Match size (bp): 39000 VLP:WC ratio: 3.2436
Whole-community read coverage
VLP-fraction read coverage
Contig position (bp)
Oscillospiraceae Pelethomonas

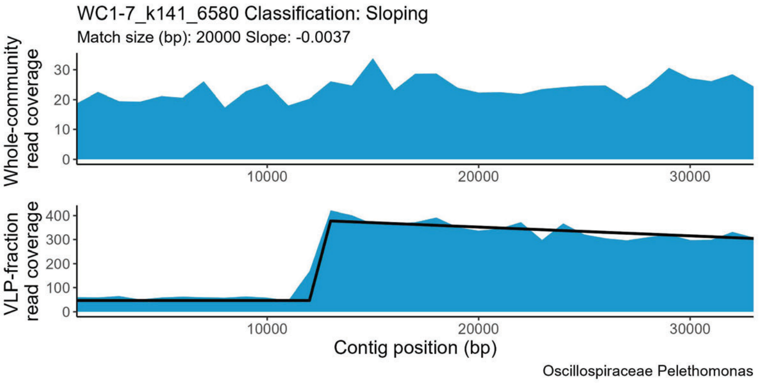
WC1-7_k141_6580 Classification: Sloping
Match size (bp): 20000 Slope: -0.0037
Whole-community read coverage
VLP-fraction read coverage
Contig position (bp)
Oscillospiraceae Pelethomonas

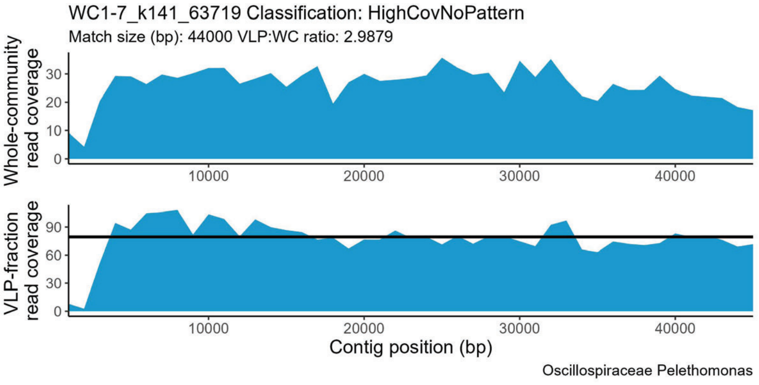
WC1-7_k141_63719 Classification: HighCovNoPattern
Match size (bp): 44000 VLP:WC ratio: 2.9879
Whole-community read coverage
VLP-fraction read coverage
Contig position (bp)
Oscillospiraceae Pelethomonas

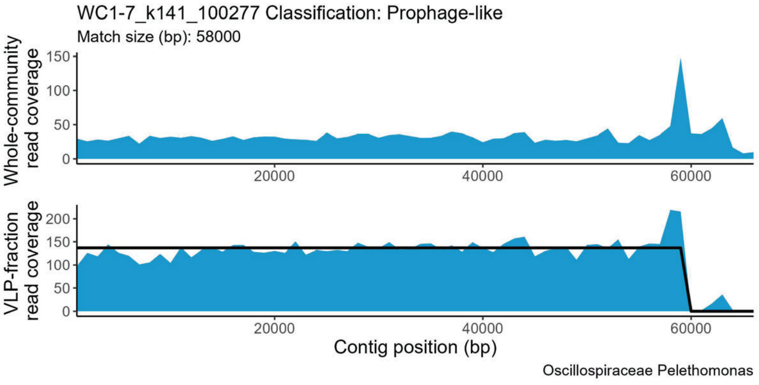
WC1-7_k141_100277 Classification: Prophage-like
Match size (bp): 58000
Whole-community read coverage
VLP-fraction read coverage
Contig position (bp)
Oscillospiraceae Pelethomonas

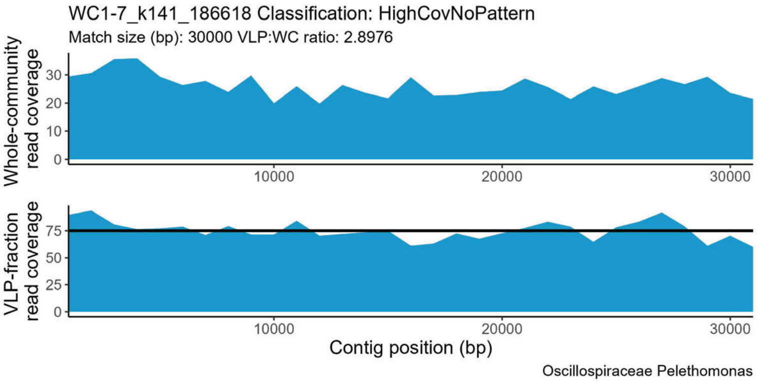
WC1-7_k141_186618 Classification: HighCovNoPattern
Match size (bp): 30000 VLP:WC ratio: 2.8976
Whole-community read coverage
0
10
20
30
10000
20000
30000
VLP-fraction read coverage
0
25
50
75
10000
20000
30000
Contig position (bp)
Oscillospiraceae Pelethomonas

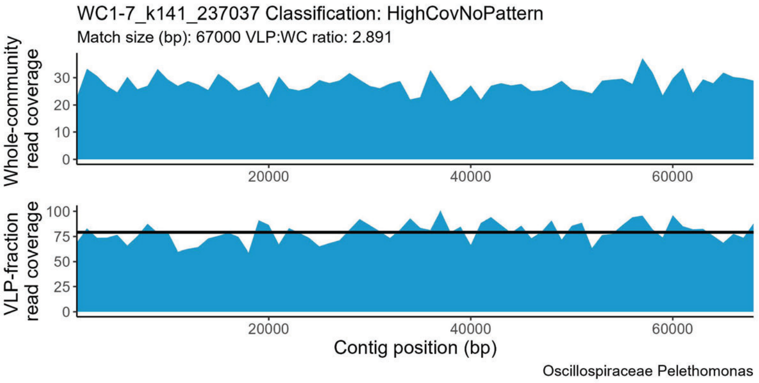
WC1-7_k141_237037 Classification: HighCovNoPattern
Match size (bp): 67000 VLP:WC ratio: 2.891
Whole-community read coverage
0
10
20
30
20000
40000
60000
VLP-fraction read coverage
0
25
50
75
100
20000
40000
60000
Contig position (bp)
Oscillospiraceae Pelethomonas

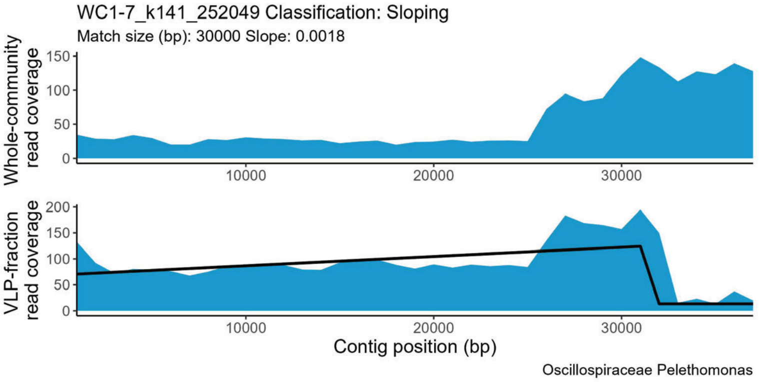
WC1-7_k141_252049 Classification: Sloping
Match size (bp): 30000 Slope: 0.0018
Whole-community read coverage
0
50
100
150
10000
20000
30000
VLP-fraction read coverage
0
50
100
150
200
10000
20000
30000
Contig position (bp)
Oscillospiraceae Pelethomonas

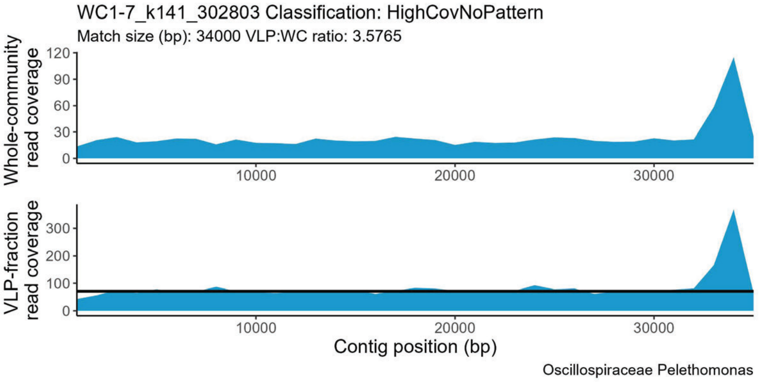
WC1-7_k141_302803 Classification: HighCovNoPattern
Match size (bp): 34000 VLP:WC ratio: 3.5765
Whole-community read coverage
0
30
60
90
120
10000
20000
30000
VLP-fraction read coverage
0
100
200
300
10000
20000
30000
Contig position (bp)
Oscillospiraceae Pelethomonas

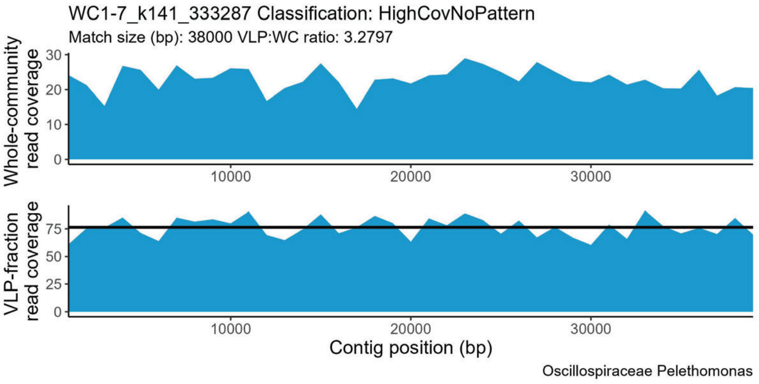
WC1-7_k141_333287 Classification: HighCovNoPattern
Match size (bp): 38000 VLP:WC ratio: 3.2797
Whole-community read coverage
0
10
20
30
10000
20000
30000
VLP-fraction read coverage
0
25
50
75
10000
20000
30000
Contig position (bp)
Oscillospiraceae Pelethomonas

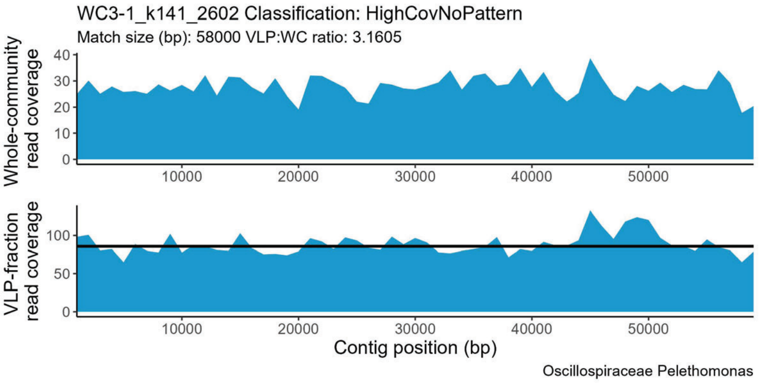
WC3-1_k141_2602 Classification: HighCovNoPattern
Match size (bp): 58000 VLP:WC ratio: 3.1605
Whole-community read coverage
0
10
20
30
40
10000
20000
30000
40000
50000
VLP-fraction read coverage
0
50
100
10000
20000
30000
40000
50000
Contig position (bp)
Oscillospiraceae Pelethomonas

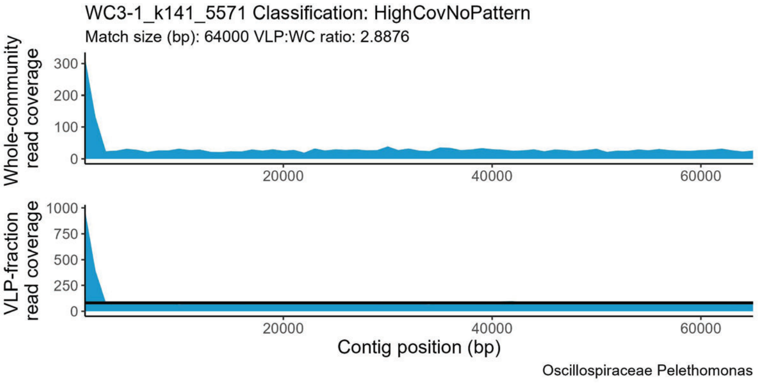
WC3-1_k141_5571 Classification: HighCovNoPattern
Match size (bp): 64000 VLP:WC ratio: 2.8876
Whole-community read coverage
0
100
200
300
20000
40000
60000
VLP-fraction read coverage
0
250
500
750
1000
20000
40000
60000
Contig position (bp)
Oscillospiraceae Pelethomonas

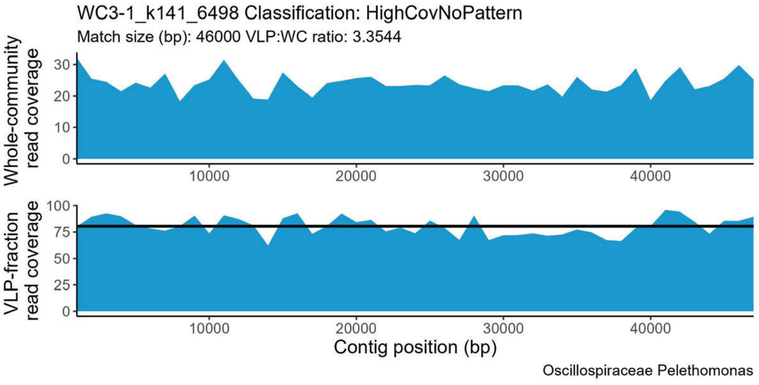
WC3-1_k141_6498 Classification: HighCovNoPattern
Match size (bp): 46000 VLP:WC ratio: 3.3544
Whole-community read coverage
0
10
20
30
10000
20000
30000
40000
VLP-fraction read coverage
0
25
50
75
100
10000
20000
30000
40000
Contig position (bp)
Oscillospiraceae Pelethomonas

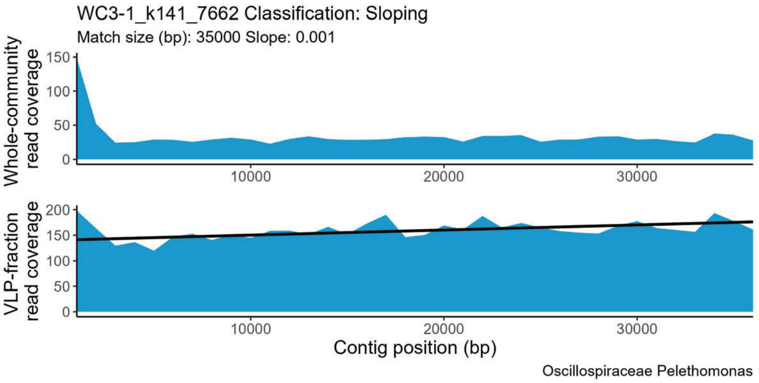
WC3-1_k141_7662 Classification: Sloping
Match size (bp): 35000 Slope: 0.001
Whole-community read coverage
VLP-fraction read coverage
Contig position (bp)
Oscillospiraceae Pelethomonas

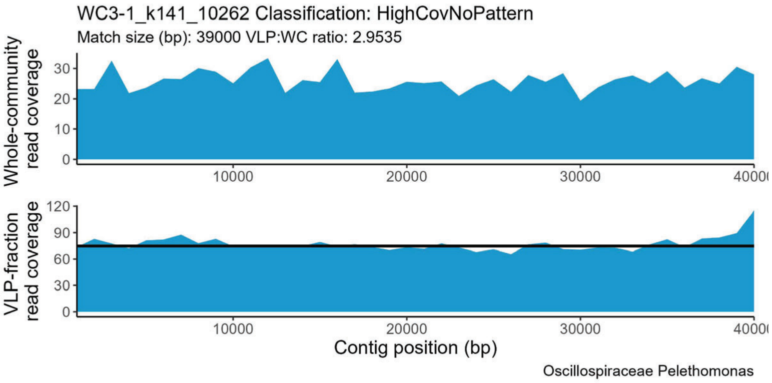
WC3-1_k141_10262 Classification: HighCovNoPattern
Match size (bp): 39000 VLP:WC ratio: 2.9535
Whole-community read coverage
VLP-fraction read coverage
Contig position (bp)
Oscillospiraceae Pelethomonas

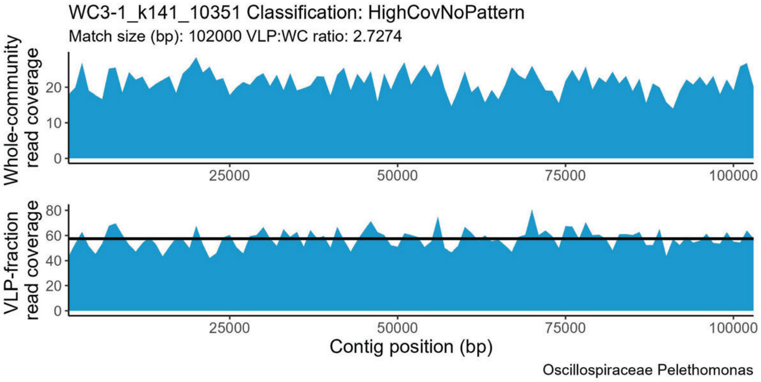
WC3-1_k141_10351 Classification: HighCovNoPattern
Match size (bp): 102000 VLP:WC ratio: 2.7274
Whole-community read coverage
VLP-fraction read coverage
Contig position (bp)
Oscillospiraceae Pelethomonas

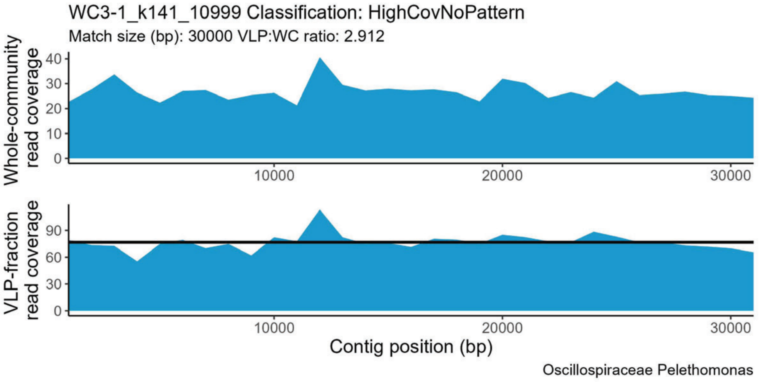
WC3-1_k141_10999 Classification: HighCovNoPattern
Match size (bp): 30000 VLP:WC ratio: 2.912
Whole-community read coverage
VLP-fraction read coverage
Contig position (bp)
Oscillospiraceae Pelethomonas

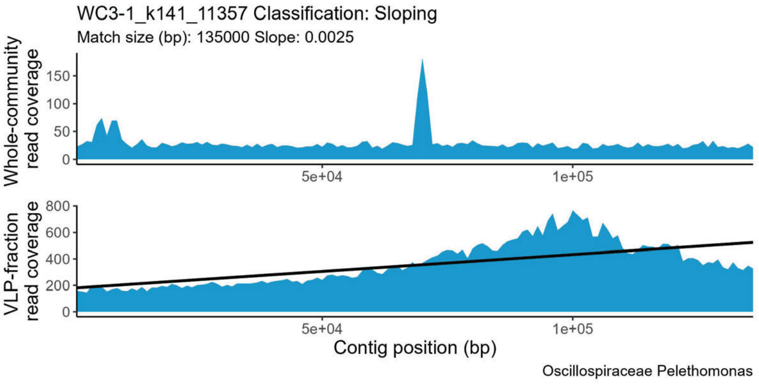
WC3-1_k141_11357 Classification: Sloping
Match size (bp): 135000 Slope: 0.0025
Whole-community read coverage
VLP-fraction read coverage
Contig position (bp)
Oscillospiraceae Pelethomonas

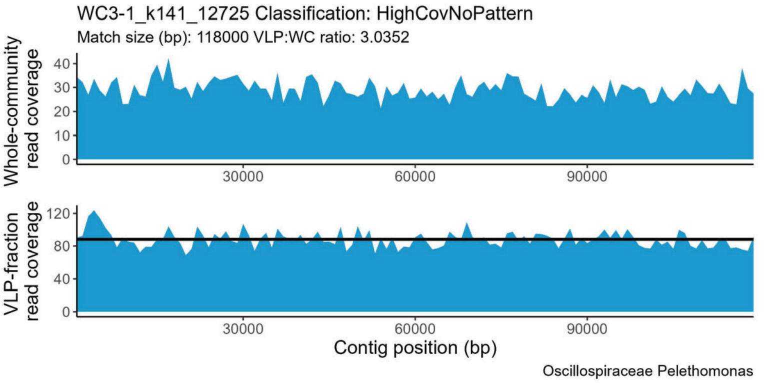
WC3-1_k141_12725 Classification: HighCovNoPattern
Match size (bp): 118000 VLP:WC ratio: 3.0352
Whole-community read coverage
VLP-fraction read coverage
Contig position (bp)
Oscillospiraceae Pelethomonas

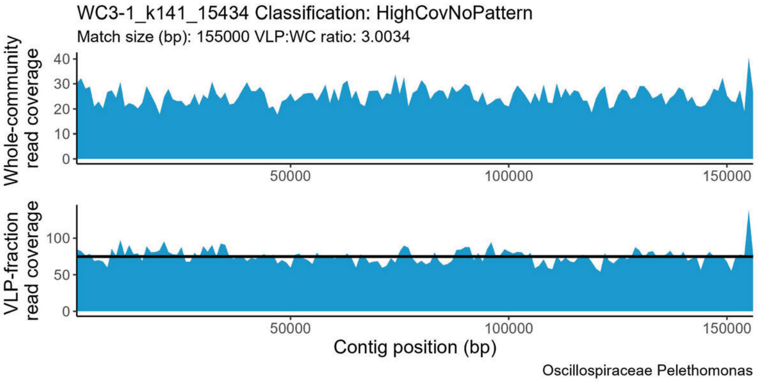
WC3-1_k141_15434 Classification: HighCovNoPattern
Match size (bp): 155000 VLP:WC ratio: 3.0034
Whole-community read coverage
VLP-fraction read coverage
Contig position (bp)
Oscillospiraceae Pelethomonas

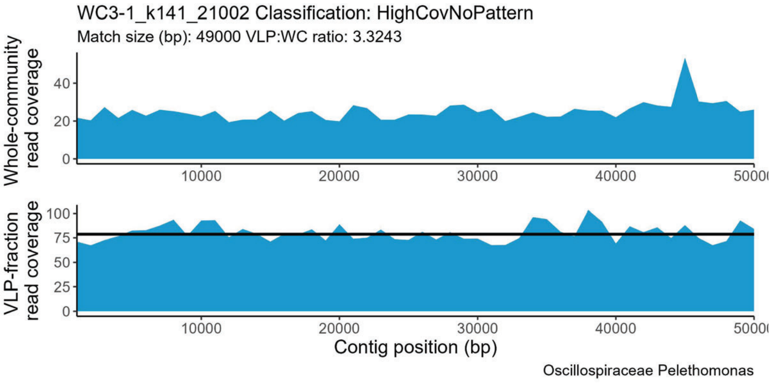
WC3-1_k141_21002 Classification: HighCovNoPattern
Match size (bp): 49000 VLP:WC ratio: 3.3243
Whole-community read coverage
VLP-fraction read coverage
Contig position (bp)
Oscillospiraceae Pelethomonas

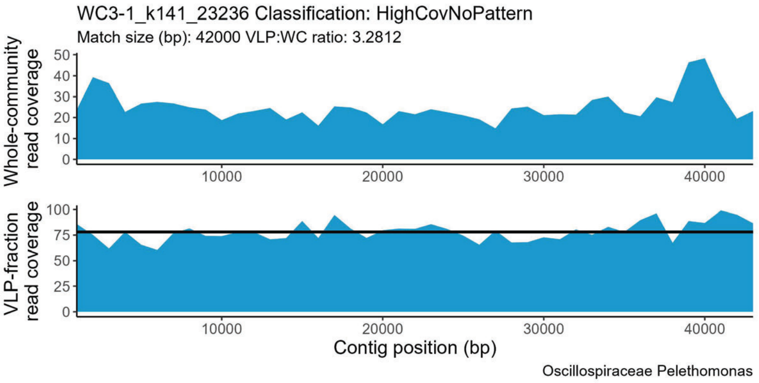
WC3-1_k141_23236 Classification: HighCovNoPattern
Match size (bp): 42000 VLP:WC ratio: 3.2812
Whole-community read coverage
VLP-fraction read coverage
Contig position (bp)
Oscillospiraceae Pelethomonas

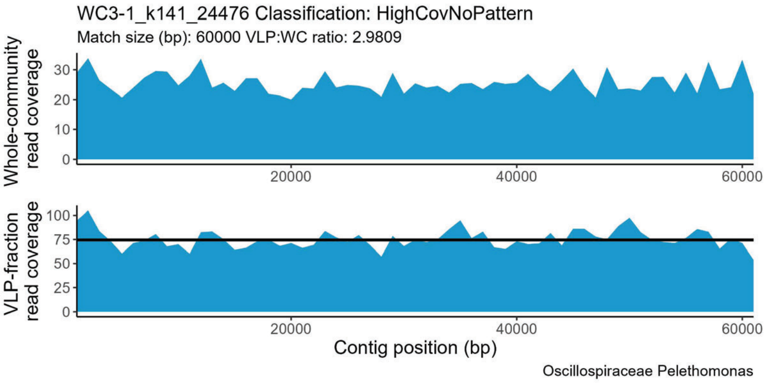
WC3-1_k141_24476 Classification: HighCovNoPattern
Match size (bp): 60000 VLP:WC ratio: 2.9809
Whole-community read coverage
VLP-fraction read coverage
Contig position (bp)
Oscillospiraceae Pelethomonas

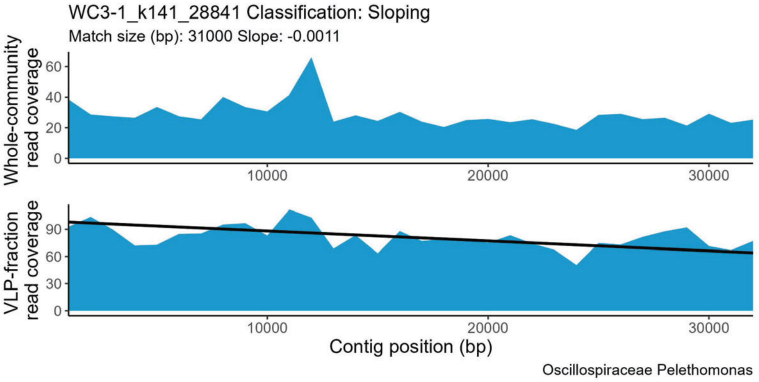
WC3-1_k141_28841 Classification: Sloping
Match size (bp): 31000 Slope: -0.0011
Whole-community read coverage
VLP-fraction read coverage
Contig position (bp)
Oscillospiraceae Pelethomonas

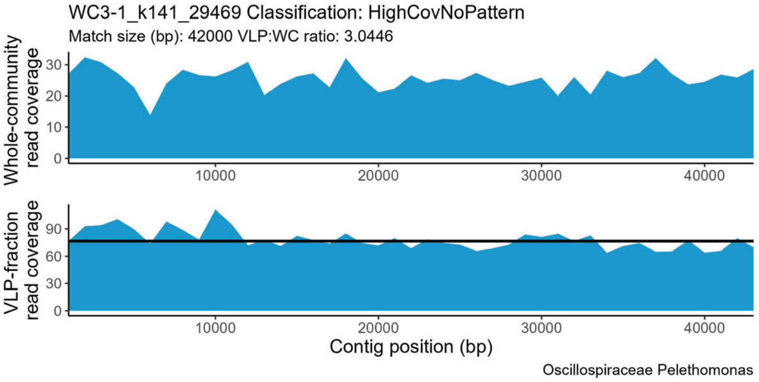
WC3-1_k141_29469 Classification: HighCovNoPattern
Match size (bp): 42000 VLP:WC ratio: 3.0446
Whole-community read coverage
VLP-fraction read coverage
Contig position (bp)
Oscillospiraceae Pelethomonas

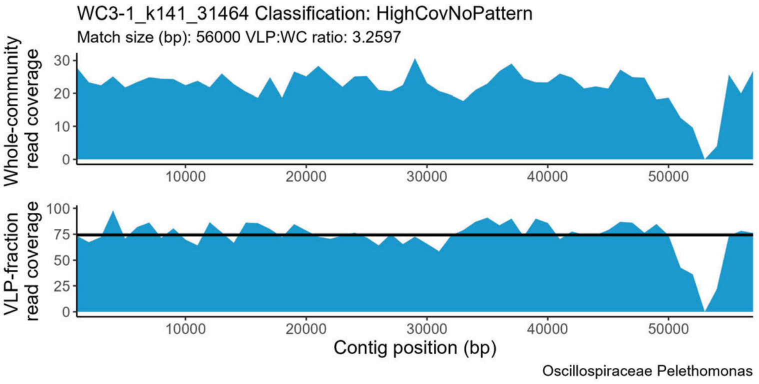
WC3-1_k141_31464 Classification: HighCovNoPattern
Match size (bp): 56000 VLP:WC ratio: 3.2597
Whole-community read coverage
VLP-fraction read coverage
Contig position (bp)
Oscillospiraceae Pelethomonas

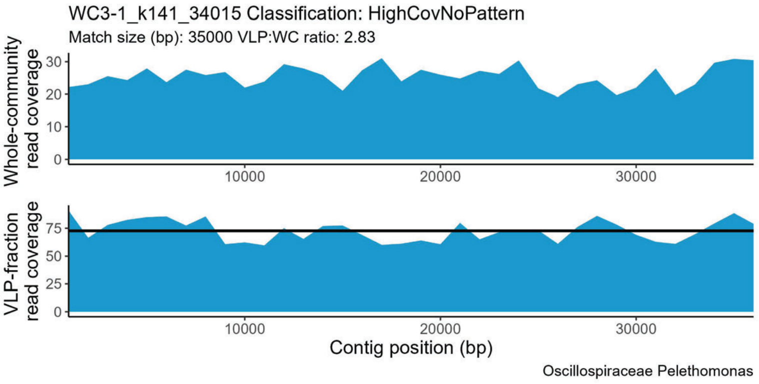
WC3-1_k141_34015 Classification: HighCovNoPattern
Match size (bp): 35000 VLP:WC ratio: 2.83
Whole-community read coverage
VLP-fraction read coverage
Contig position (bp)
Oscillospiraceae Pelethomonas

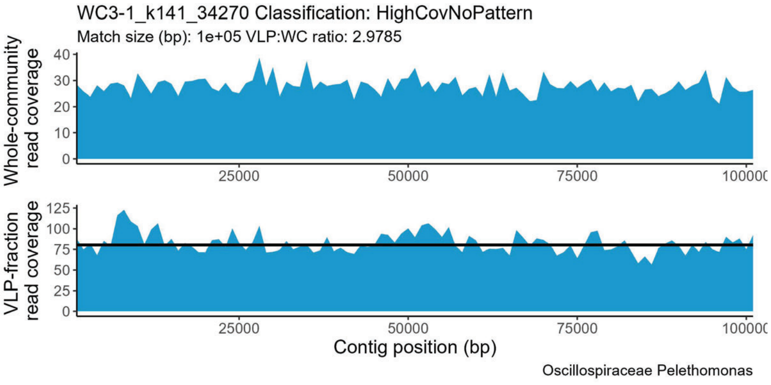
WC3-1_k141_34270 Classification: HighCovNoPattern
Match size (bp): 1e+05 VLP:WC ratio: 2.9785
Whole-community read coverage
VLP-fraction read coverage
Contig position (bp)
Oscillospiraceae Pelethomonas

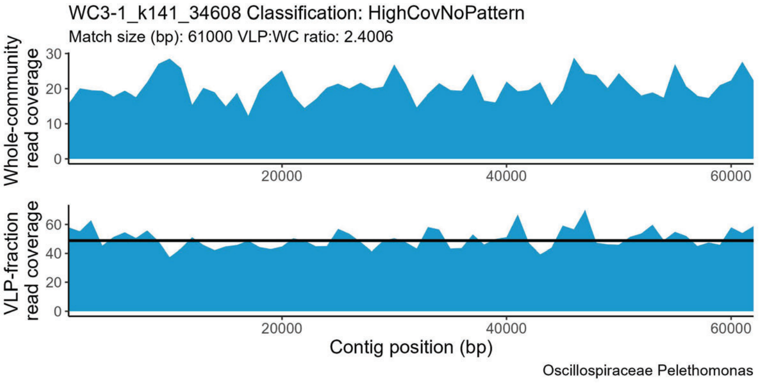
WC3-1_k141_34608 Classification: HighCovNoPattern
Match size (bp): 61000 VLP:WC ratio: 2.4006
Whole-community read coverage
VLP-fraction read coverage
Contig position (bp)
Oscillospiraceae Pelethomonas

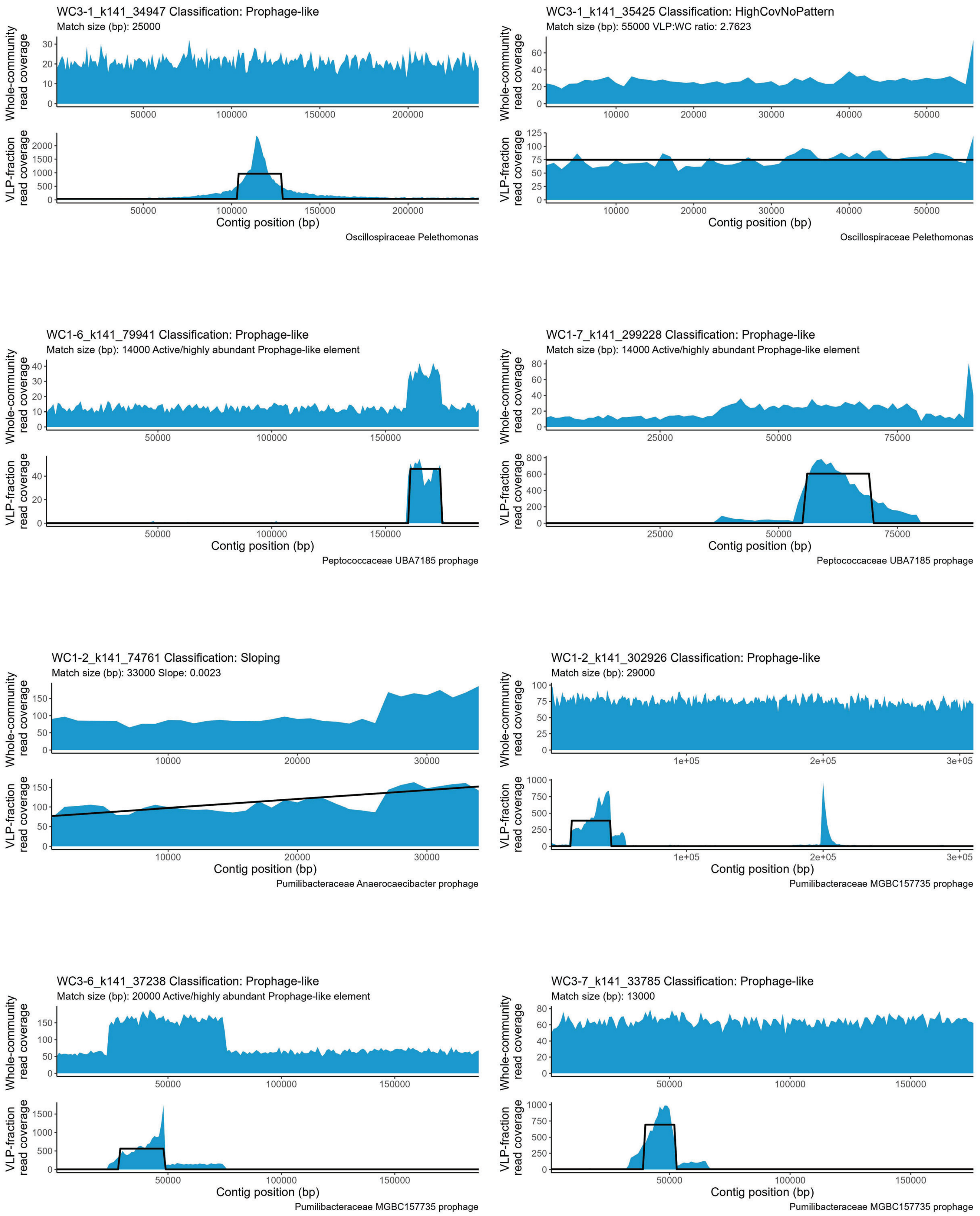
WC3-1_k141_34947 Classification: Prophage-like
Match size (bp): 25000
Whole-community read coverage
VLP-fraction read coverage
Contig position (bp)
Oscillospiraceae Pelethomonas
WC3-1_k141_35425 Classification: HighCovNoPattern
Match size (bp): 55000 VLP:WC ratio: 2.7623
Whole-community read coverage
VLP-fraction read coverage
Contig position (bp)
Oscillospiraceae Pelethomonas
WC1-6_k141_79941 Classification: Prophage-like
Match size (bp): 14000 Active/highly abundant Prophage-like element
Whole-community read coverage
VLP-fraction read coverage
Contig position (bp)
Peptococcaceae UBA7185 prophage
WC1-7_k141_299228 Classification: Prophage-like
Match size (bp): 14000 Active/highly abundant Prophage-like element
Whole-community read coverage
VLP-fraction read coverage
Contig position (bp)
Peptococcaceae UBA7185 prophage
WC1-2_k141_74761 Classification: Sloping
Match size (bp): 33000 Slope: 0.0023
Whole-community read coverage
VLP-fraction read coverage
Contig position (bp)
Pumilibacteraceae Anaerocaecibacter prophage
WC1-2_k141_302926 Classification: Prophage-like
Match size (bp): 29000
Whole-community read coverage
VLP-fraction read coverage
Contig position (bp)
Pumilibacteraceae MGBC157735 prophage
WC3-6_k141_37238 Classification: Prophage-like
Match size (bp): 20000 Active/highly abundant Prophage-like element
Whole-community read coverage
VLP-fraction read coverage
Contig position (bp)
Pumilibacteraceae MGBC157735 prophage
WC3-7_k141_33785 Classification: Prophage-like
Match size (bp): 13000
Whole-community read coverage
VLP-fraction read coverage
Contig position (bp)
Pumilibacteraceae MGBC157735 prophage

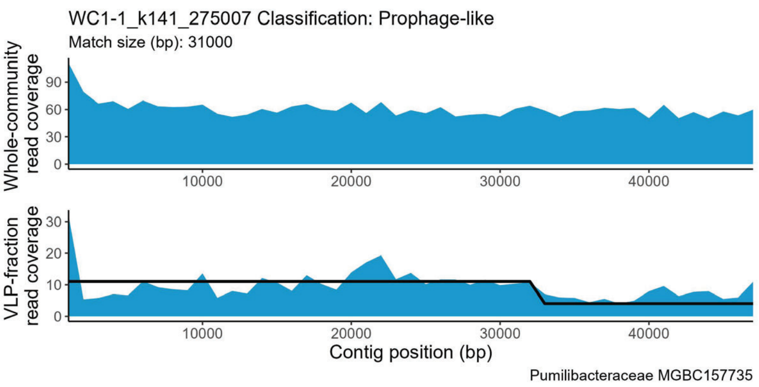
WC1-1_k141_275007 Classification: Prophage-like
Match size (bp): 31000
Whole-community read coverage
VLP-fraction read coverage
Contig position (bp)
Pumilibacteraceae MGBC157735

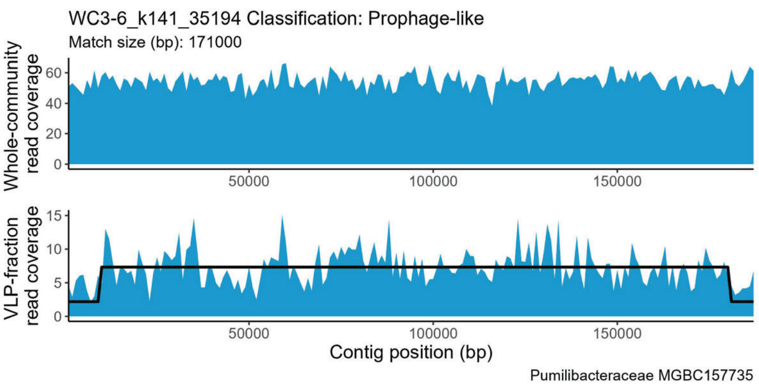
WC3-6_k141_35194 Classification: Prophage-like
Match size (bp): 171000
Whole-community read coverage
VLP-fraction read coverage
Contig position (bp)
Pumilibacteraceae MGBC157735

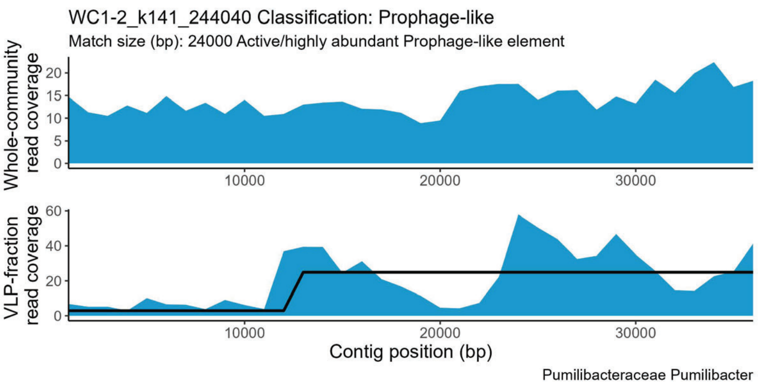
WC1-2_k141_244040 Classification: Prophage-like
Match size (bp): 24000 Active/highly abundant Prophage-like element
Whole-community read coverage
VLP-fraction read coverage
Contig position (bp)
Pumilibacteraceae Pumilibacter

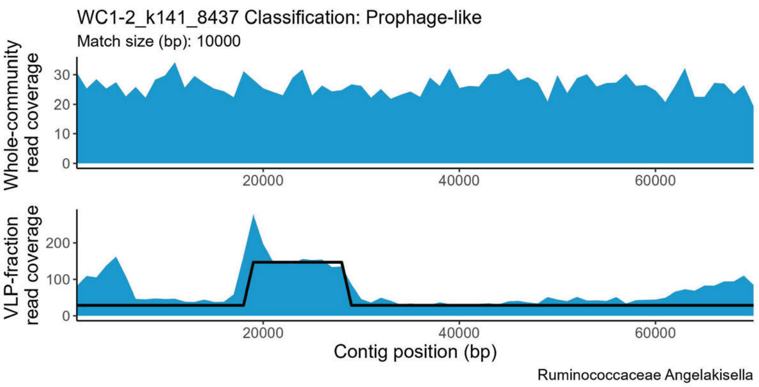
WC1-2_k141_8437 Classification: Prophage-like
Match size (bp): 10000
Whole-community read coverage
VLP-fraction read coverage
Contig position (bp)
Ruminococcaceae Angelakisella

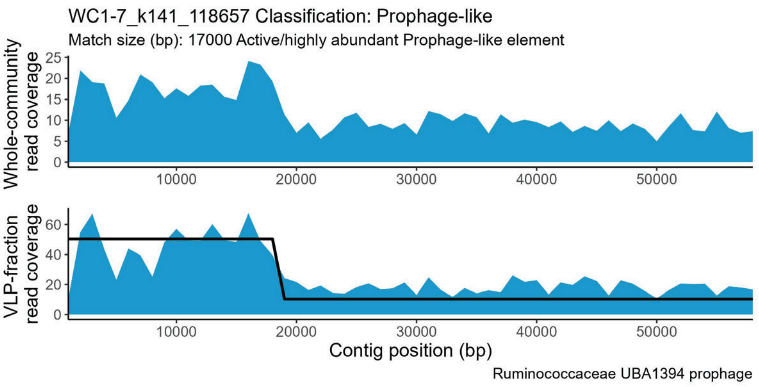
WC1-7_k141_118657 Classification: Prophage-like
Match size (bp): 17000 Active/highly abundant Prophage-like element
Whole-community read coverage
VLP-fraction read coverage
Contig position (bp)
Ruminococcaceae UBA1394 prophage

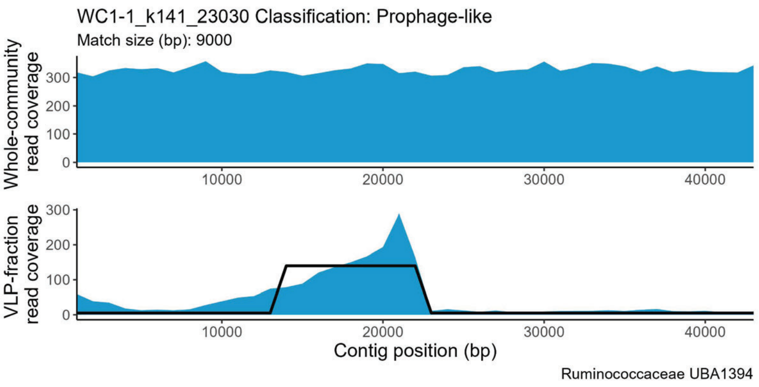
WC1-1_k141_23030 Classification: Prophage-like
Match size (bp): 9000
Whole-community read coverage
VLP-fraction read coverage
Contig position (bp)
Ruminococcaceae UBA1394

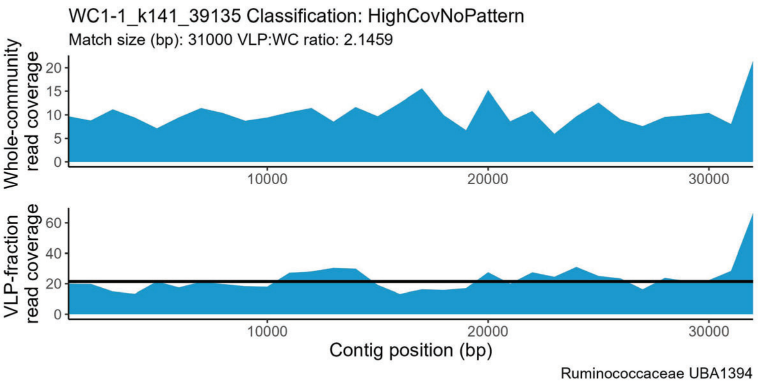
WC1-1_k141_39135 Classification: HighCovNoPattern
Match size (bp): 31000 VLP:WC ratio: 2.1459
Whole-community read coverage
VLP-fraction read coverage
Contig position (bp)
Ruminococcaceae UBA1394

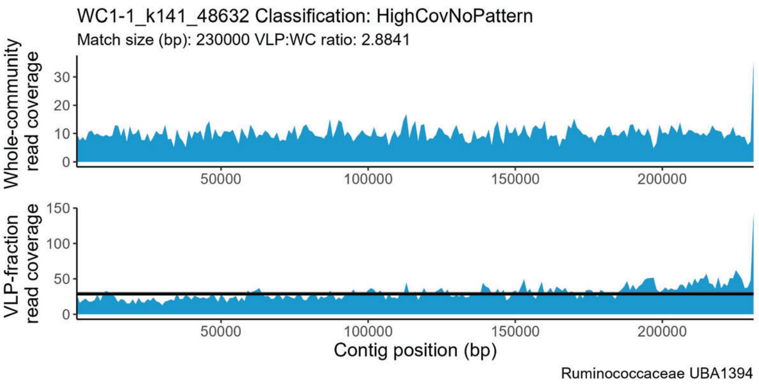
WC1-1_k141_48632 Classification: HighCovNoPattern
Match size (bp): 230000 VLP:WC ratio: 2.8841
Whole-community read coverage
VLP-fraction read coverage
Contig position (bp)
Ruminococcaceae UBA1394

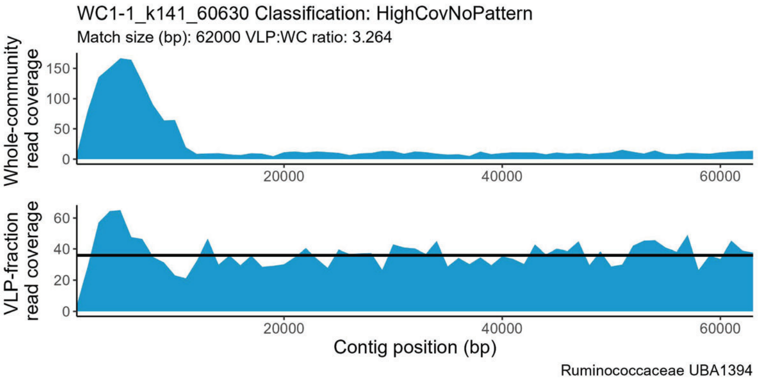
WC1-1_k141_60630 Classification: HighCovNoPattern
Match size (bp): 62000 VLP:WC ratio: 3.264
Whole-community read coverage
VLP-fraction read coverage
Contig position (bp)
Ruminococcaceae UBA1394

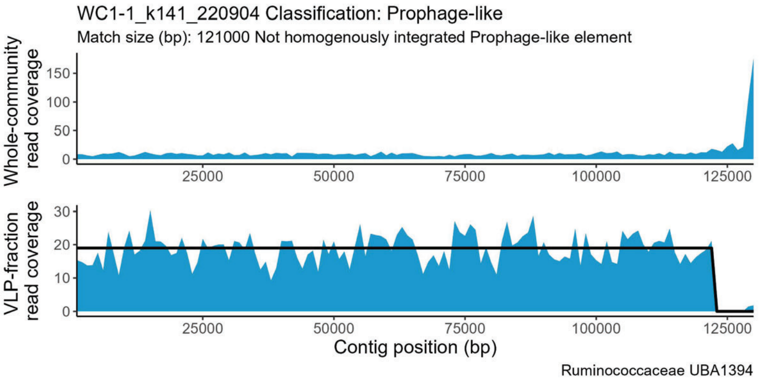
WC1-1_k141_220904 Classification: Prophage-like
Match size (bp): 121000 Not homogenously integrated Prophage-like element
Whole-community read coverage
VLP-fraction read coverage
Contig position (bp)
Ruminococcaceae UBA1394

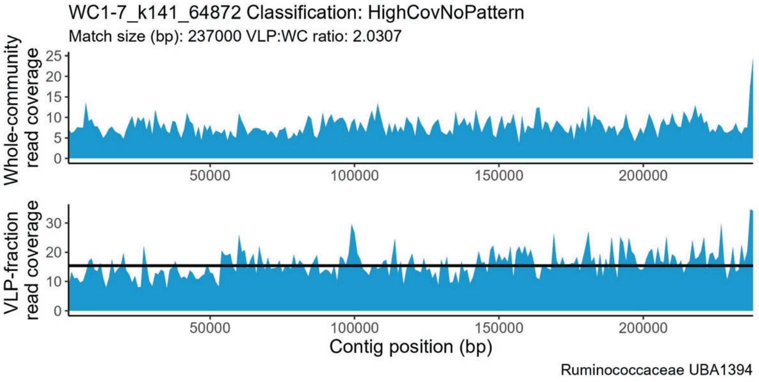
WC1-7_k141_64872 Classification: HighCovNoPattern
Match size (bp): 237000 VLP:WC ratio: 2.0307
Whole-community read coverage
VLP-fraction read coverage
Contig position (bp)
Ruminococcaceae UBA1394

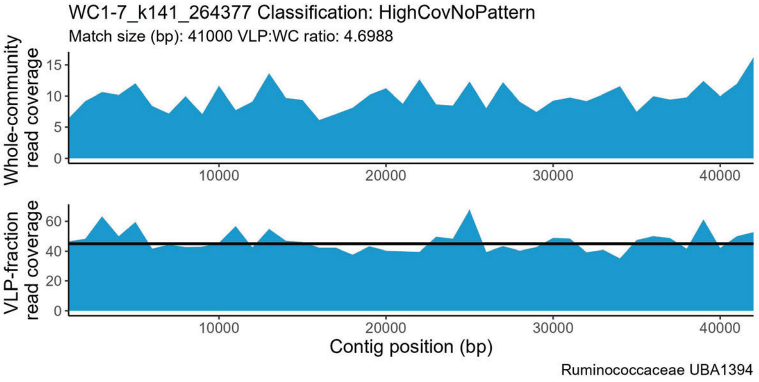
WC1-7_k141_264377 Classification: HighCovNoPattern
Match size (bp): 41000 VLP:WC ratio: 4.6988
Whole-community read coverage
VLP-fraction read coverage
Contig position (bp)
Ruminococcaceae UBA1394

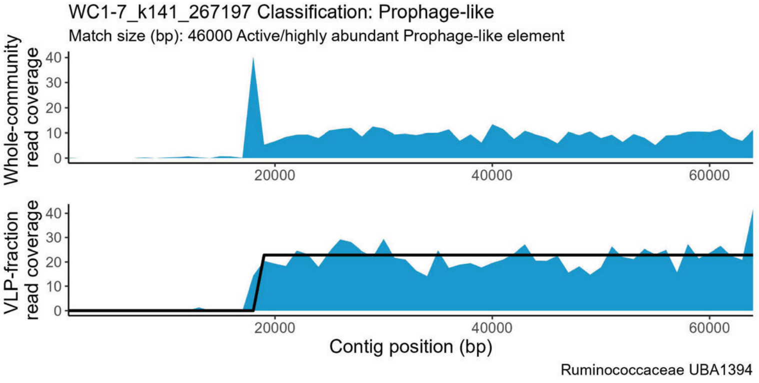
WC1-7_k141_267197 Classification: Prophage-like
Match size (bp): 46000 Active/highly abundant Prophage-like element
Whole-community read coverage
VLP-fraction read coverage
Contig position (bp)
Ruminococcaceae UBA1394

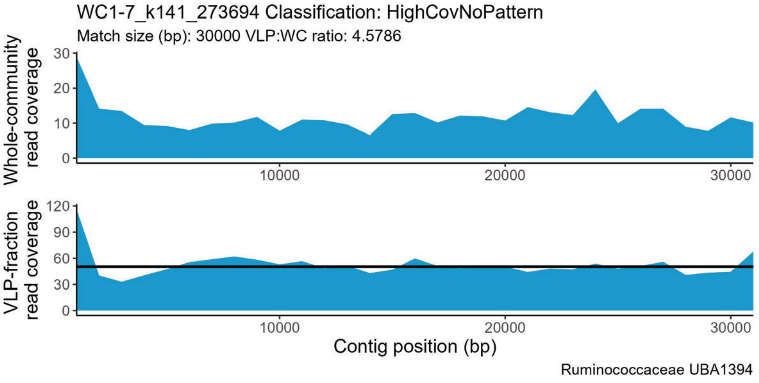
WC1-7_k141_273694 Classification: HighCovNoPattern
Match size (bp): 30000 VLP:WC ratio: 4.5786
Whole-community read coverage
VLP-fraction read coverage
Contig position (bp)
Ruminococcaceae UBA1394

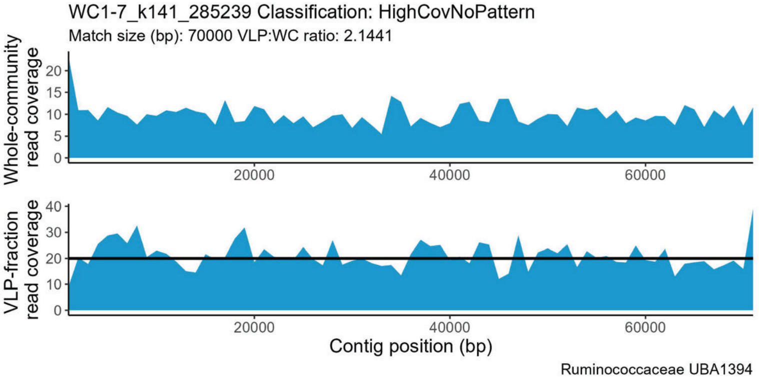
WC1-7_k141_285239 Classification: HighCovNoPattern
Match size (bp): 70000 VLP:WC ratio: 2.1441
Whole-community read coverage
VLP-fraction read coverage
Contig position (bp)
Ruminococcaceae UBA1394

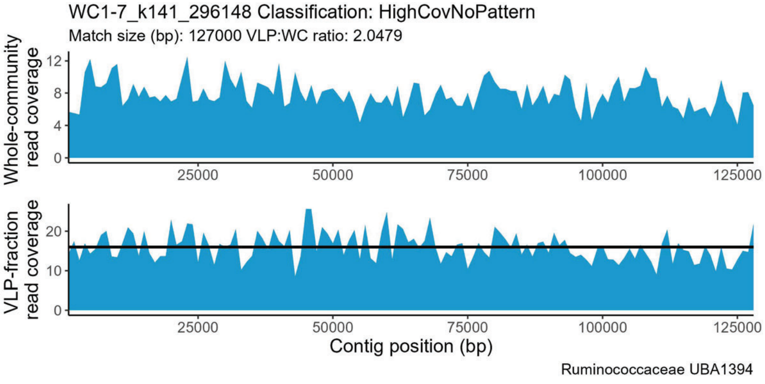
WC1-7_k141_296148 Classification: HighCovNoPattern
Match size (bp): 127000 VLP:WC ratio: 2.0479
Whole-community read coverage
VLP-fraction read coverage
Contig position (bp)
Ruminococcaceae UBA1394

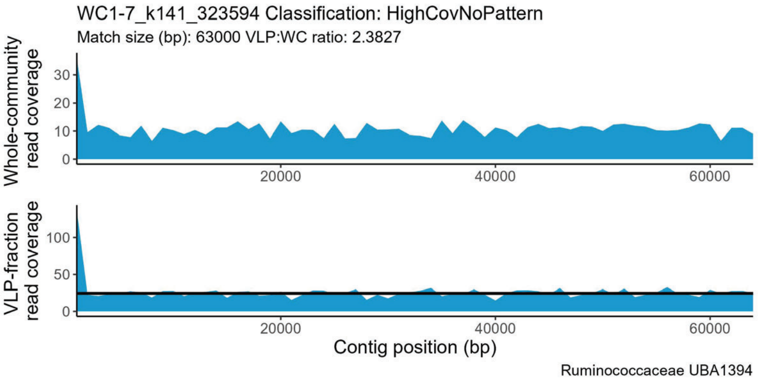
WC1-7_k141_323594 Classification: HighCovNoPattern
Match size (bp): 63000 VLP:WC ratio: 2.3827
Whole-community read coverage
VLP-fraction read coverage
Contig position (bp)
Ruminococcaceae UBA1394

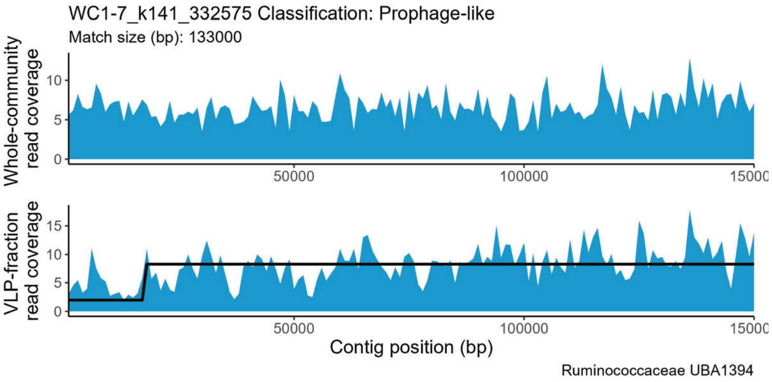
WC1-7_k141_332575 Classification: Prophage-like
Match size (bp): 133000
Whole-community read coverage
VLP-fraction read coverage
Contig position (bp)
Ruminococcaceae UBA1394

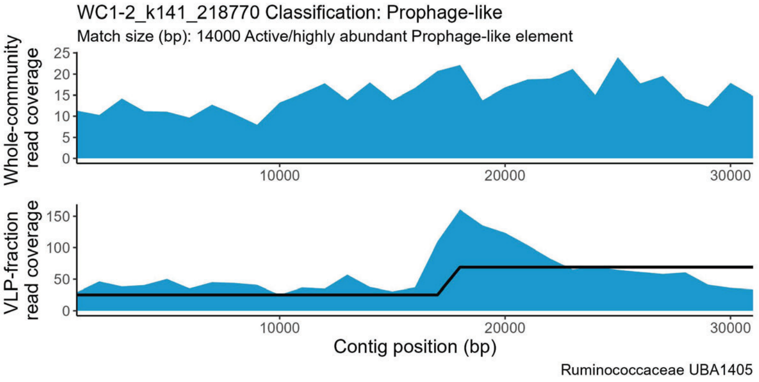
WC1-2_k141_218770 Classification: Prophage-like
Match size (bp): 14000 Active/highly abundant Prophage-like element
Whole-community read coverage
VLP-fraction read coverage
Contig position (bp)
Ruminococcaceae UBA1405

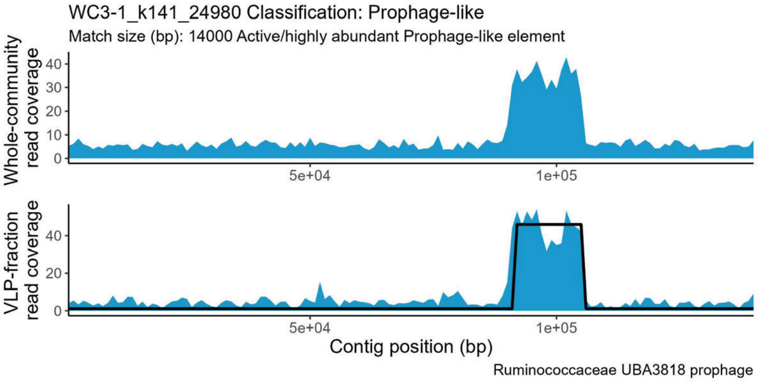
WC3-1_k141_24980 Classification: Prophage-like
Match size (bp): 14000 Active/highly abundant Prophage-like element
Whole-community read coverage
VLP-fraction read coverage
Contig position (bp)
Ruminococcaceae UBA3818 prophage

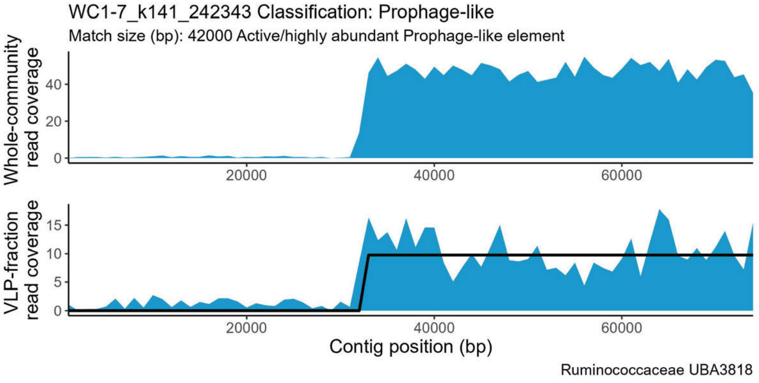
WC1-7_k141_242343 Classification: Prophage-like
Match size (bp): 42000 Active/highly abundant Prophage-like element
Whole-community read coverage
VLP-fraction read coverage
Contig position (bp)
Ruminococcaceae UBA3818

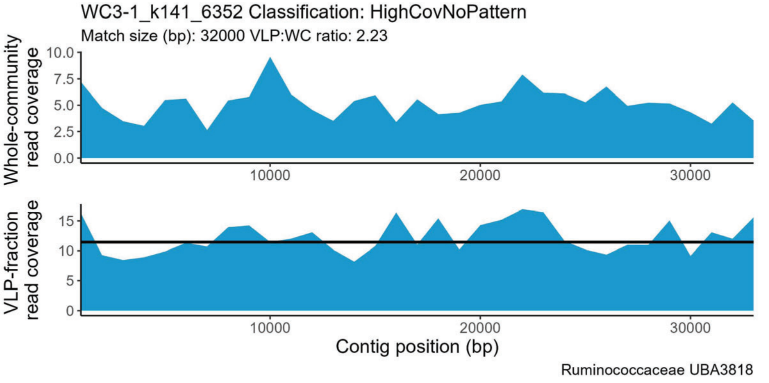
WC3-1_k141_6352 Classification: HighCovNoPattern
Match size (bp): 32000 VLP:WC ratio: 2.23
Whole-community read coverage
VLP-fraction read coverage
Contig position (bp)
Ruminococcaceae UBA3818

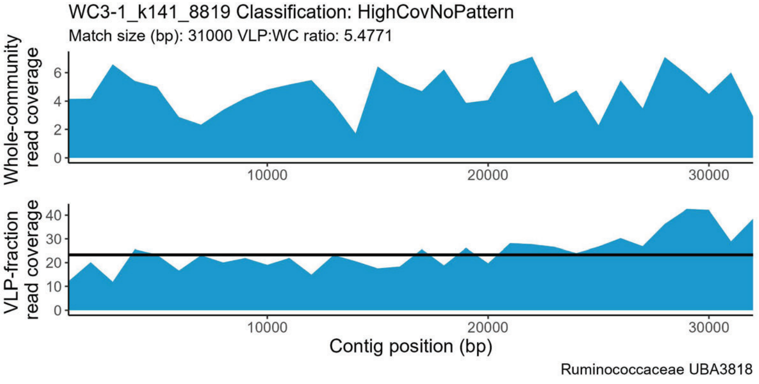
WC3-1_k141_8819 Classification: HighCovNoPattern
Match size (bp): 31000 VLP:WC ratio: 5.4771
Whole-community read coverage
VLP-fraction read coverage
Contig position (bp)
Ruminococcaceae UBA3818

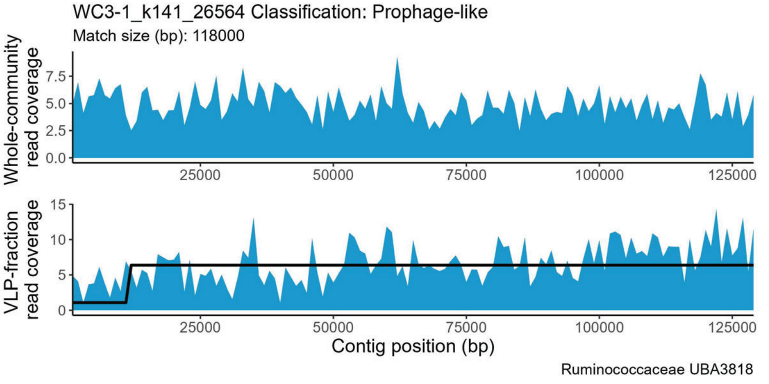
WC3-1_k141_26564 Classification: Prophage-like
Match size (bp): 118000
Whole-community read coverage
VLP-fraction read coverage
Contig position (bp)
Ruminococcaceae UBA3818

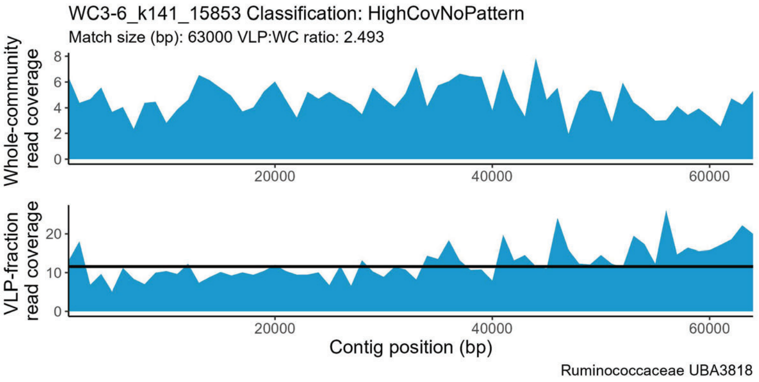
WC3-6_k141_15853 Classification: HighCovNoPattern
Match size (bp): 63000 VLP:WC ratio: 2.493
Whole-community read coverage
0 2 4 6 8
20000 40000 60000
VLP-fraction read coverage
0 10 20
20000 40000 60000
Contig position (bp)
Ruminococcaceae UBA3818

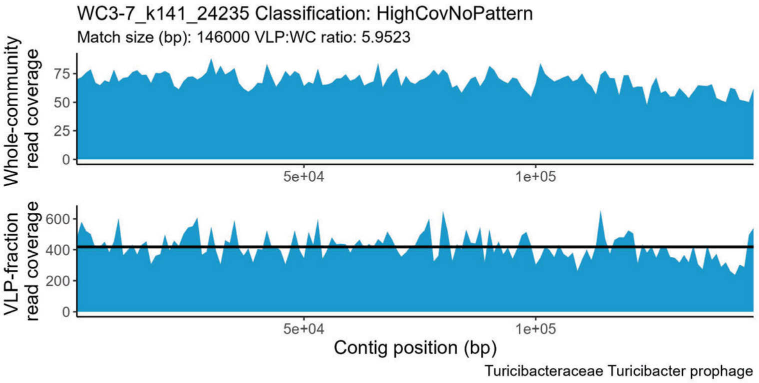
WC3-7_k141_24235 Classification: HighCovNoPattern
Match size (bp): 146000 VLP:WC ratio: 5.9523
Whole-community read coverage
0 25 50 75
5e+04 1e+05
VLP-fraction read coverage
0 200 400 600
5e+04 1e+05
Contig position (bp)
Turicibacteraceae Turicibacter prophage

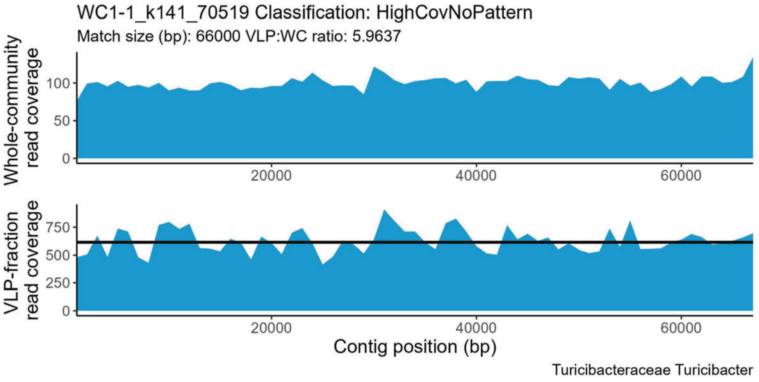
WC1-1_k141_70519 Classification: HighCovNoPattern
Match size (bp): 66000 VLP:WC ratio: 5.9637
Whole-community read coverage
0 50 100
20000 40000 60000
VLP-fraction read coverage
0 250 500 750
20000 40000 60000
Contig position (bp)
Turicibacteraceae Turicibacter

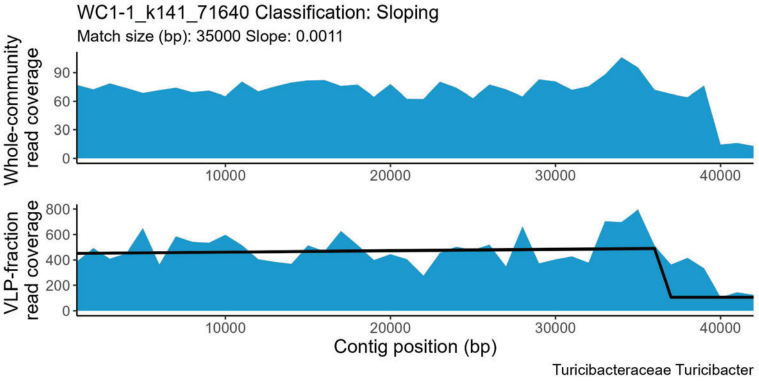
WC1-1_k141_71640 Classification: Sloping
Match size (bp): 35000 Slope: 0.0011
Whole-community read coverage
0 30 60 90
10000 20000 30000 40000
VLP-fraction read coverage
0 200 400 600 800
10000 20000 30000 40000
Contig position (bp)
Turicibacteraceae Turicibacter

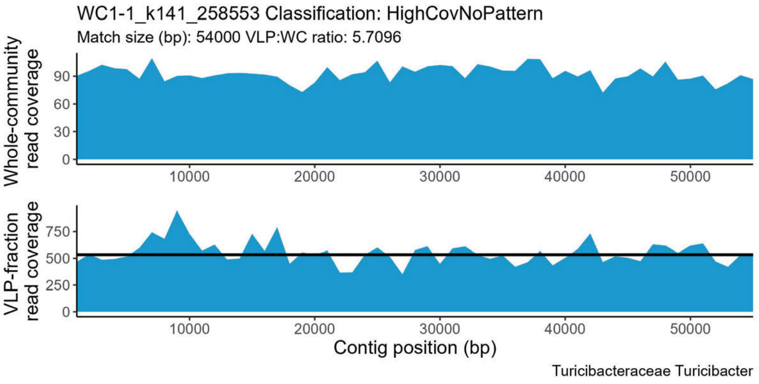
WC1-1_k141_258553 Classification: HighCovNoPattern
Match size (bp): 54000 VLP:WC ratio: 5.7096
Whole-community read coverage
0 30 60 90
10000 20000 30000 40000 50000
VLP-fraction read coverage
0 250 500 750
10000 20000 30000 40000 50000
Contig position (bp)
Turicibacteraceae Turicibacter

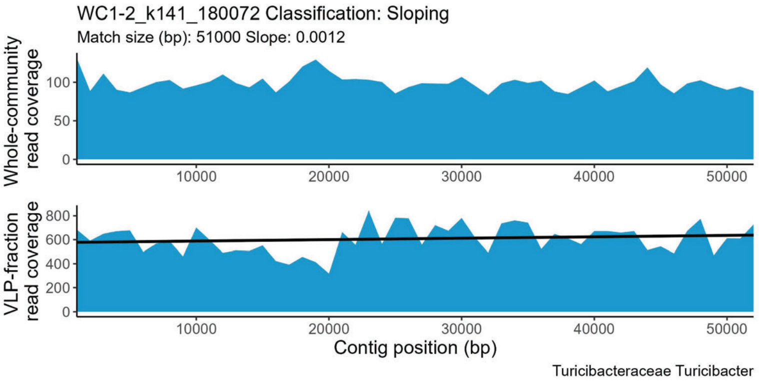
WC1-2_k141_180072 Classification: Sloping
Match size (bp): 51000 Slope: 0.0012
Whole-community read coverage
0 50 100
10000 20000 30000 40000 50000
VLP-fraction read coverage
0 200 400 600 800
10000 20000 30000 40000 50000
Contig position (bp)
Turicibacteraceae Turicibacter

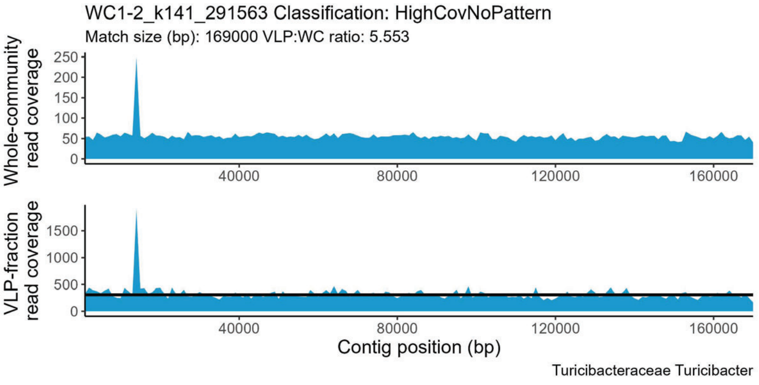
WC1-2_k141_291563 Classification: HighCovNoPattern
Match size (bp): 169000 VLP:WC ratio: 5.553
Whole-community read coverage
0 50 100 150 200 250
40000 80000 120000 160000
VLP-fraction read coverage
0 500 1000 1500
40000 80000 120000 160000
Contig position (bp)
Turicibacteraceae Turicibacter

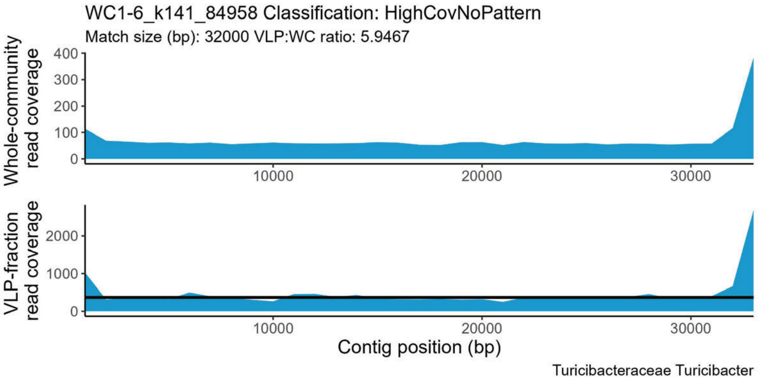
WC1-6_k141_84958 Classification: HighCovNoPattern
Match size (bp): 32000 VLP:WC ratio: 5.9467
Whole-community read coverage
0 100 200 300 400
10000 20000 30000
VLP-fraction read coverage
0 1000 2000
10000 20000 30000
Contig position (bp)
Turicibacteraceae Turicibacter

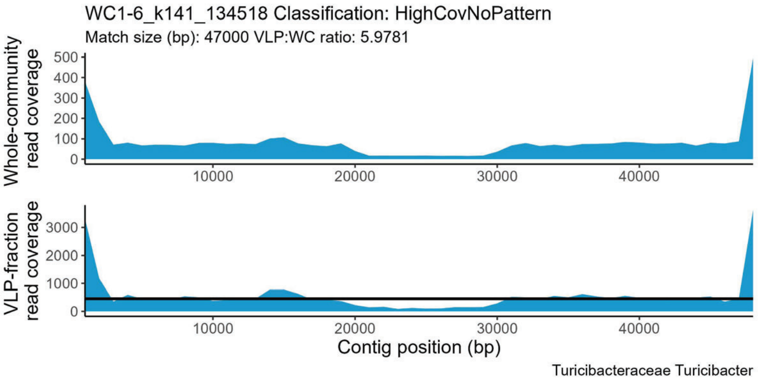
WC1-6_k141_134518 Classification: HighCovNoPattern
Match size (bp): 47000 VLP:WC ratio: 5.9781
Whole-community read coverage
VLP-fraction read coverage
Contig position (bp)
Turicibacteraceae Turicibacter

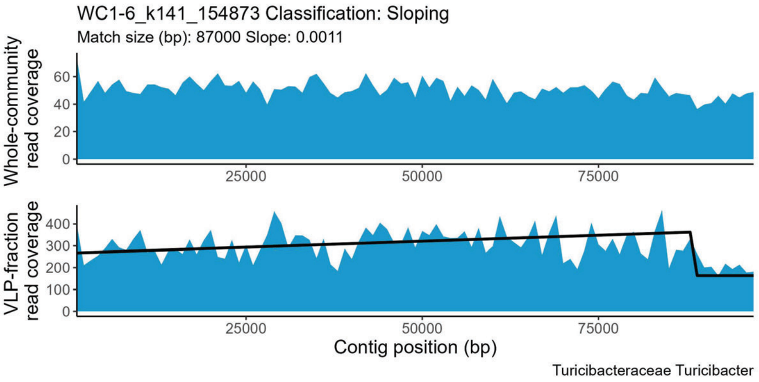
WC1-6_k141_154873 Classification: Sloping
Match size (bp): 87000 Slope: 0.0011
Whole-community read coverage
VLP-fraction read coverage
Contig position (bp)
Turicibacteraceae Turicibacter

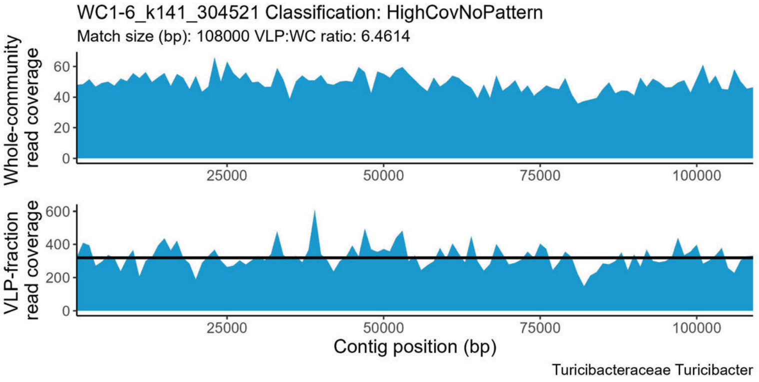
WC1-6_k141_304521 Classification: HighCovNoPattern
Match size (bp): 108000 VLP:WC ratio: 6.4614
Whole-community read coverage
VLP-fraction read coverage
Contig position (bp)
Turicibacteraceae Turicibacter

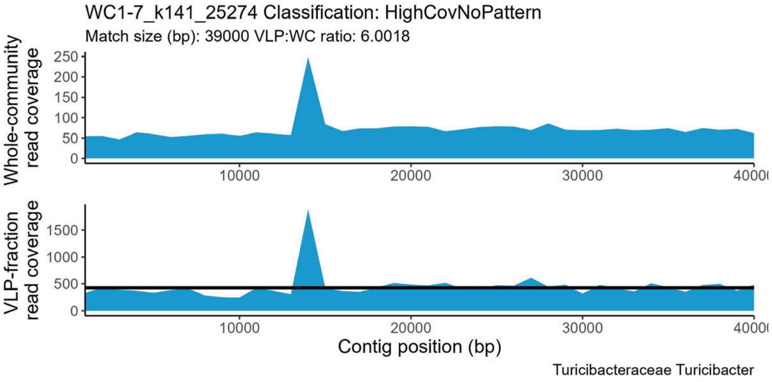
WC1-7_k141_25274 Classification: HighCovNoPattern
Match size (bp): 39000 VLP:WC ratio: 6.0018
Whole-community read coverage
VLP-fraction read coverage
Contig position (bp)
Turicibacteraceae Turicibacter

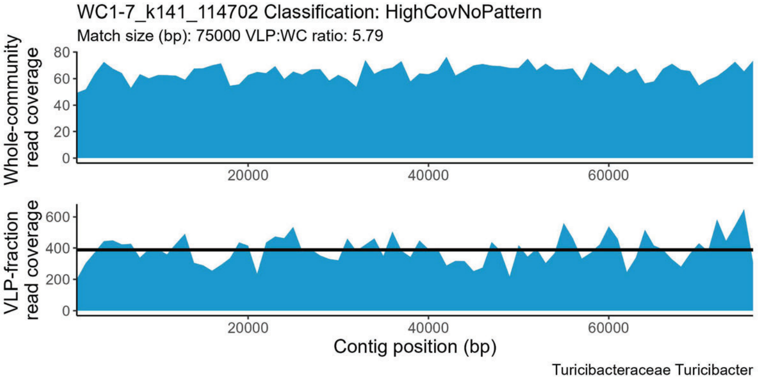
WC1-7_k141_114702 Classification: HighCovNoPattern
Match size (bp): 75000 VLP:WC ratio: 5.79
Whole-community read coverage
VLP-fraction read coverage
Contig position (bp)
Turicibacteraceae Turicibacter

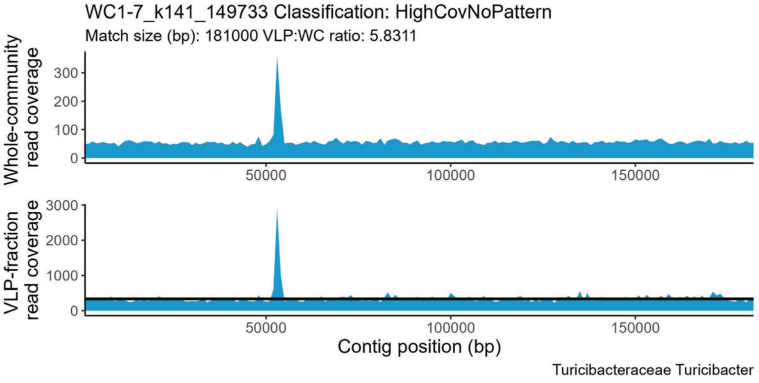
WC1-7_k141_149733 Classification: HighCovNoPattern
Match size (bp): 181000 VLP:WC ratio: 5.8311
Whole-community read coverage
VLP-fraction read coverage
Contig position (bp)
Turicibacteraceae Turicibacter

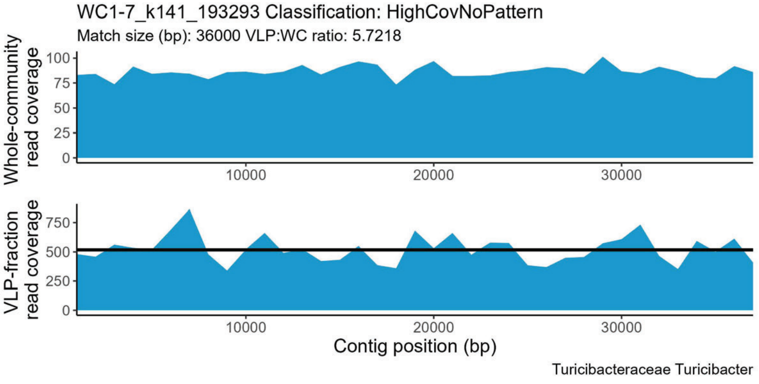
WC1-7_k141_193293 Classification: HighCovNoPattern
Match size (bp): 36000 VLP:WC ratio: 5.7218
Whole-community read coverage
VLP-fraction read coverage
Contig position (bp)
Turicibacteraceae Turicibacter

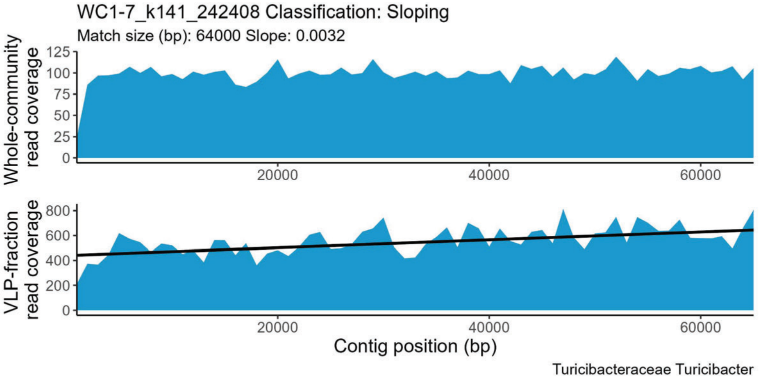
WC1-7_k141_242408 Classification: Sloping
Match size (bp): 64000 Slope: 0.0032
Whole-community read coverage
VLP-fraction read coverage
Contig position (bp)
Turicibacteraceae Turicibacter

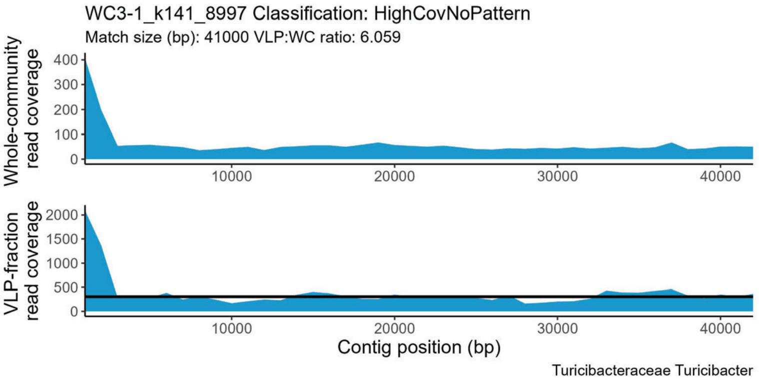
WC3-1_k141_8997 Classification: HighCovNoPattern
Match size (bp): 41000 VLP:WC ratio: 6.059
Whole-community read coverage
VLP-fraction read coverage
Contig position (bp)
Turicibacteraceae Turicibacter

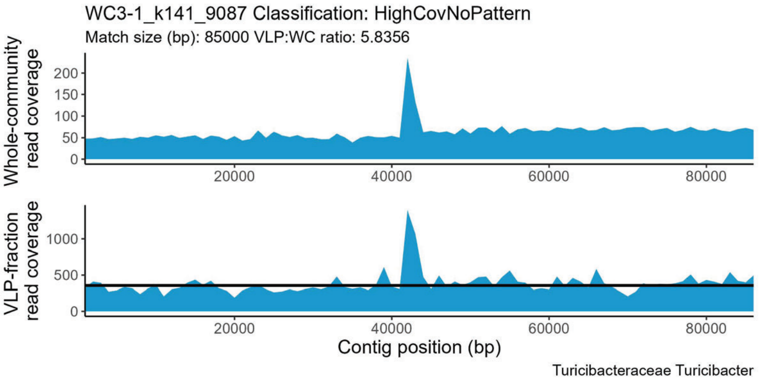
WC3-1_k141_9087 Classification: HighCovNoPattern
Match size (bp): 85000 VLP:WC ratio: 5.8356
Whole-community read coverage
VLP-fraction read coverage
Contig position (bp)
Turicibacteraceae Turicibacter

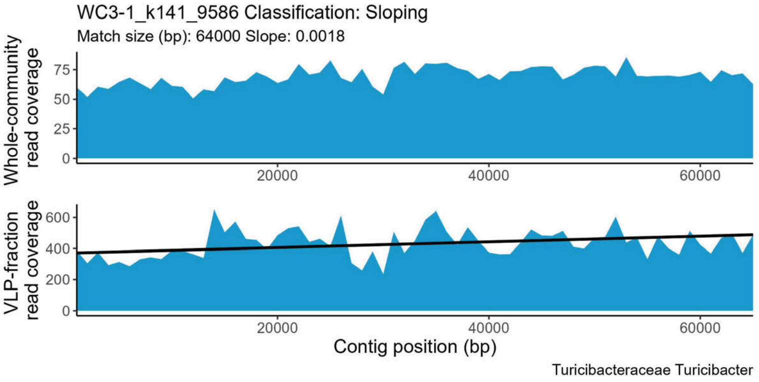
WC3-1_k141_9586 Classification: Sloping
Match size (bp): 64000 Slope: 0.0018
Whole-community read coverage
VLP-fraction read coverage
Contig position (bp)
Turicibacteraceae Turicibacter

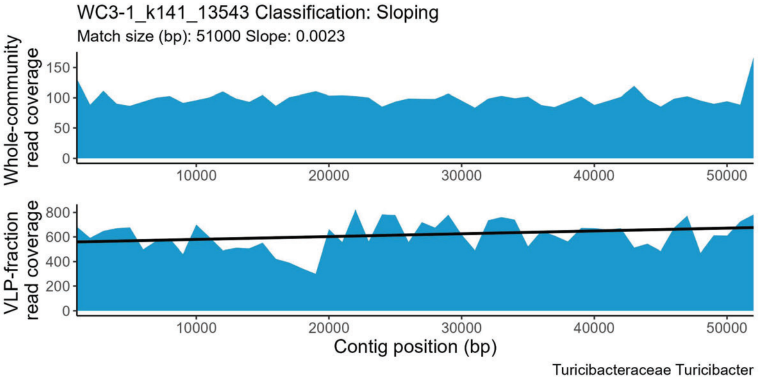
WC3-1_k141_13543 Classification: Sloping
Match size (bp): 51000 Slope: 0.0023
Whole-community read coverage
VLP-fraction read coverage
Contig position (bp)
Turicibacteraceae Turicibacter

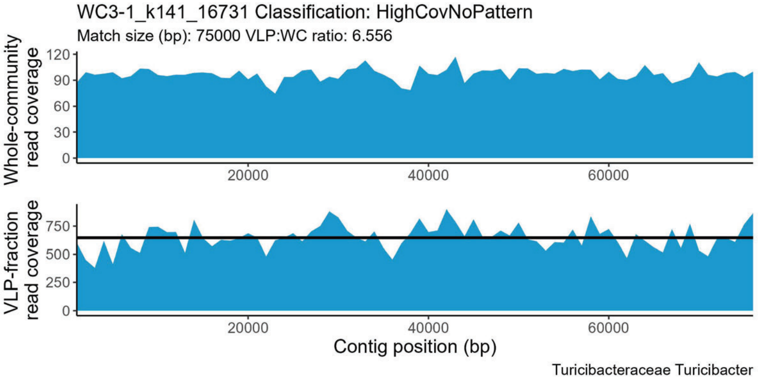
WC3-1_k141_16731 Classification: HighCovNoPattern
Match size (bp): 75000 VLP:WC ratio: 6.556
Whole-community read coverage
VLP-fraction read coverage
Contig position (bp)
Turicibacteraceae Turicibacter

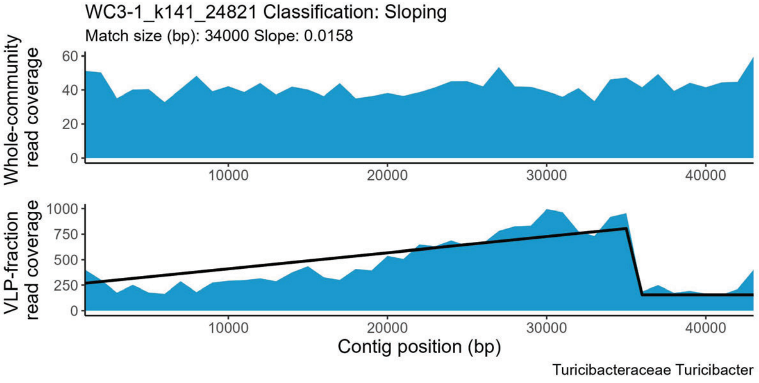
WC3-1_k141_24821 Classification: Sloping
Match size (bp): 34000 Slope: 0.0158
Whole-community read coverage
VLP-fraction read coverage
Contig position (bp)
Turicibacteraceae Turicibacter

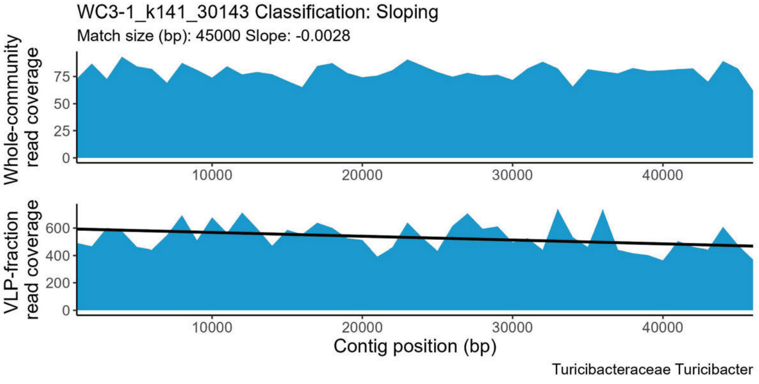
WC3-1_k141_30143 Classification: Sloping
Match size (bp): 45000 Slope: -0.0028
Whole-community read coverage
VLP-fraction read coverage
Contig position (bp)
Turicibacteraceae Turicibacter

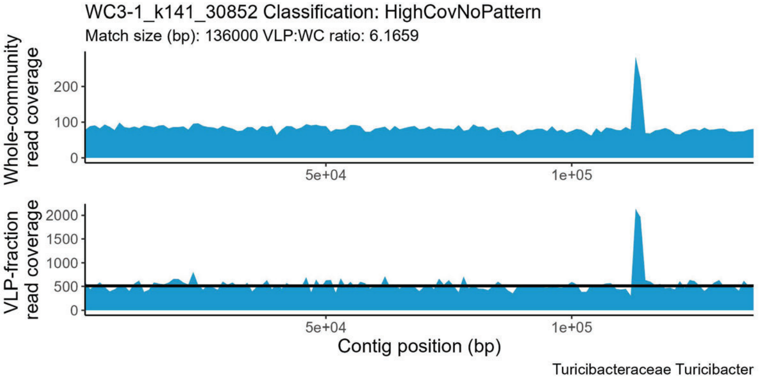
WC3-1_k141_30852 Classification: HighCovNoPattern
Match size (bp): 136000 VLP:WC ratio: 6.1659
Whole-community read coverage
VLP-fraction read coverage
Contig position (bp)
Turicibacteraceae Turicibacter

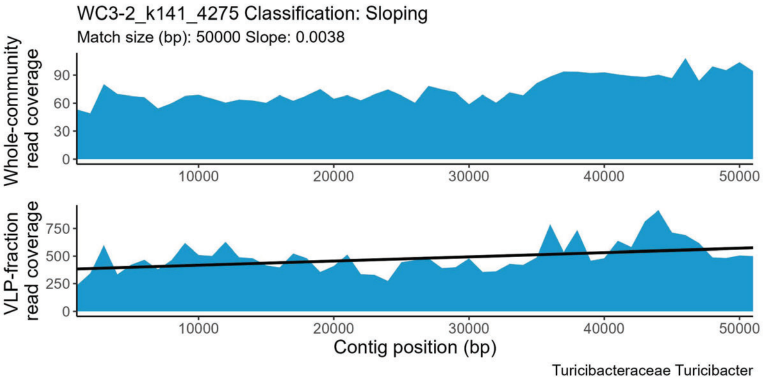
WC3-2_k141_4275 Classification: Sloping
Match size (bp): 50000 Slope: 0.0038
Whole-community read coverage
VLP-fraction read coverage
Contig position (bp)
Turicibacteraceae Turicibacter

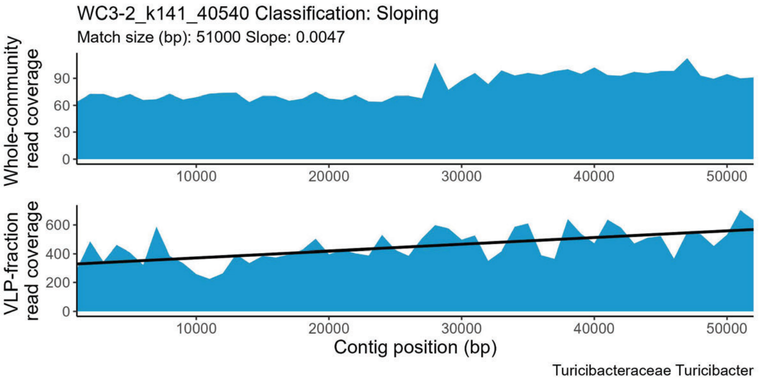
WC3-2_k141_40540 Classification: Sloping
Match size (bp): 51000 Slope: 0.0047
Whole-community read coverage
VLP-fraction read coverage
Contig position (bp)
Turicibacteraceae Turicibacter

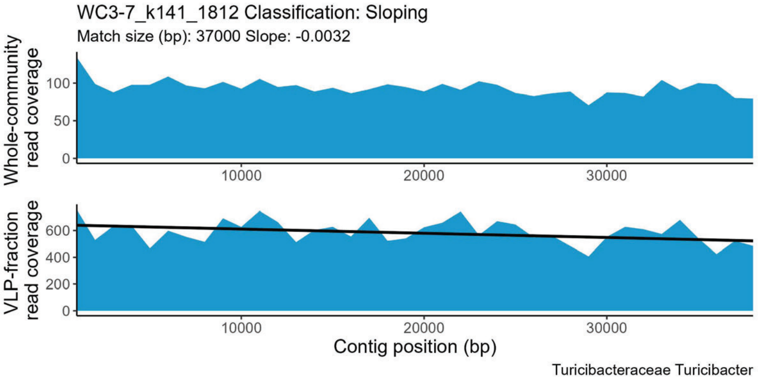
WC3-7_k141_1812 Classification: Sloping
Match size (bp): 37000 Slope: -0.0032
Whole-community read coverage
VLP-fraction read coverage
Contig position (bp)
Turicibacteraceae Turicibacter

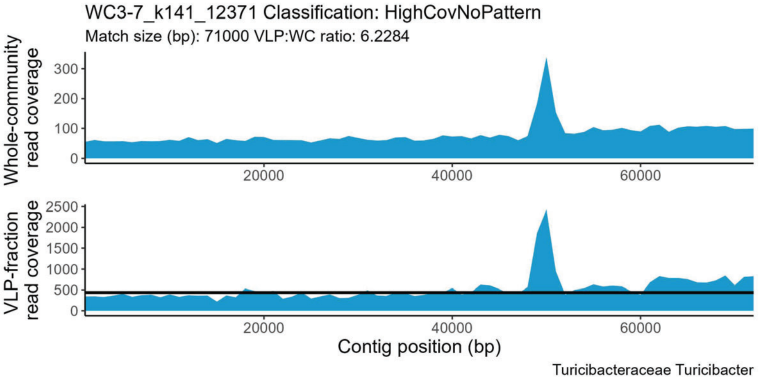
WC3-7_k141_12371 Classification: HighCovNoPattern
Match size (bp): 71000 VLP:WC ratio: 6.2284
Whole-community read coverage
VLP-fraction read coverage
Contig position (bp)
Turicibacteraceae Turicibacter

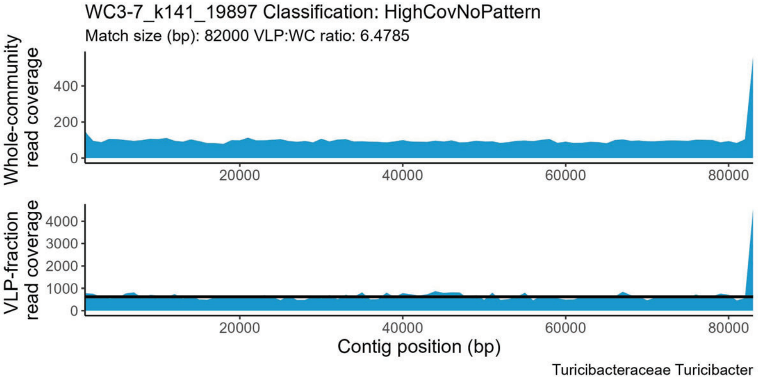
WC3-7_k141_19897 Classification: HighCovNoPattern
Match size (bp): 82000 VLP:WC ratio: 6.4785
Whole-community read coverage
VLP-fraction read coverage
Contig position (bp)
Turicibacteraceae Turicibacter

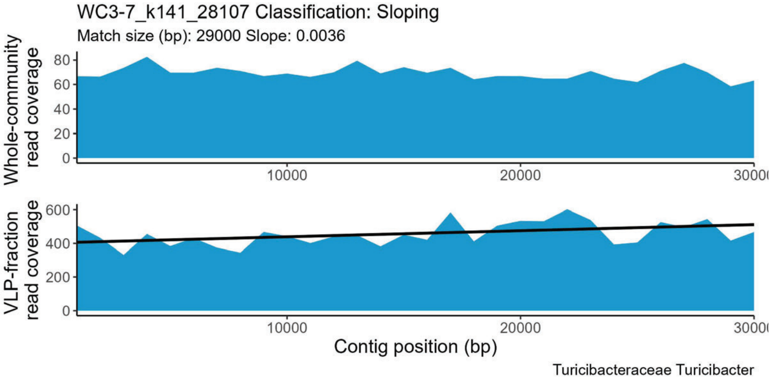
WC3-7_k141_28107 Classification: Sloping
Match size (bp): 29000 Slope: 0.0036
Whole-community read coverage
VLP-fraction read coverage
Contig position (bp)
Turicibacteraceae Turicibacter

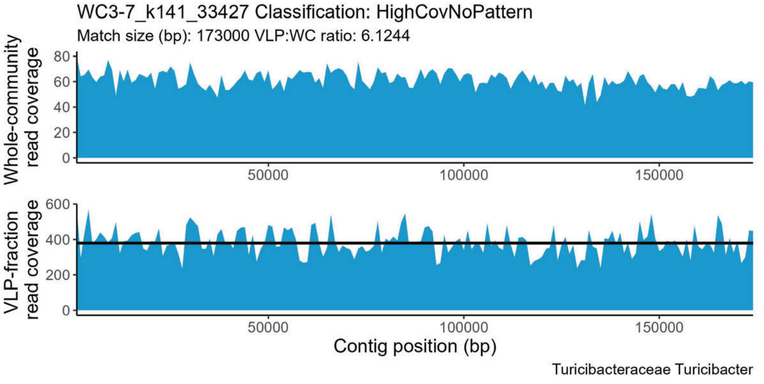
WC3-7_k141_33427 Classification: HighCovNoPattern
Match size (bp): 173000 VLP:WC ratio: 6.1244
Whole-community read coverage
VLP-fraction read coverage
Contig position (bp)
Turicibacteraceae Turicibacter

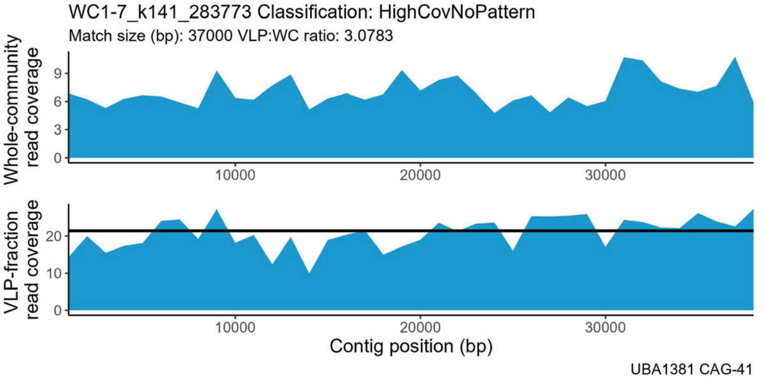
WC1-7_k141_283773 Classification: HighCovNoPattern
Match size (bp): 37000 VLP:WC ratio: 3.0783
Whole-community read coverage
VLP-fraction read coverage
Contig position (bp)
UBA1381 CAG-41

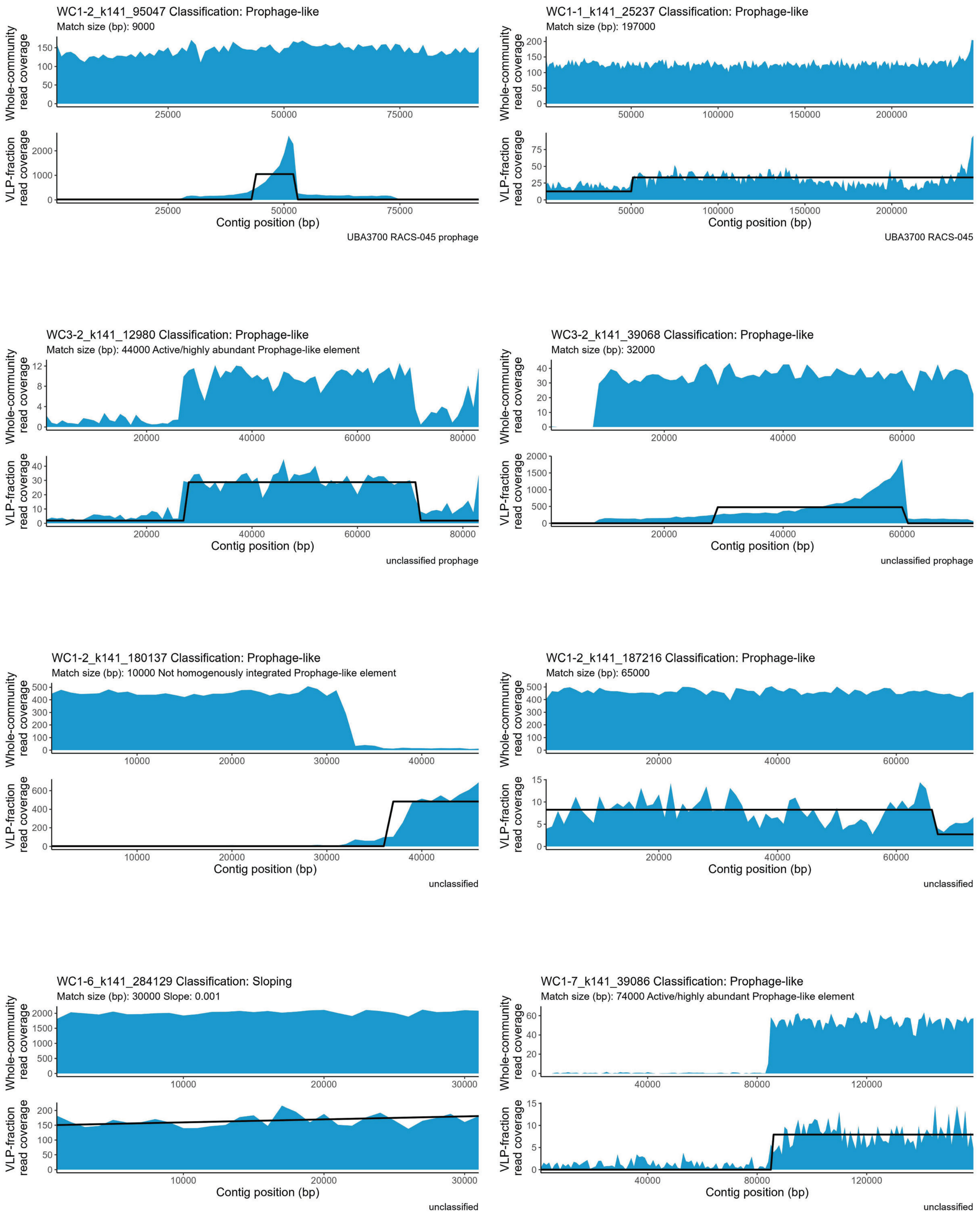

WC1-2_k141_95047 Classification: Prophage-like
Match size (bp): 9000
Whole-community read coverage
VLP-fraction read coverage
Contig position (bp)
UBA3700 RACS-045 prophage
WC1-1_k141_25237 Classification: Prophage-like
Match size (bp): 197000
Whole-community read coverage
VLP-fraction read coverage
Contig position (bp)
UBA3700 RACS-045
WC3-2_k141_12980 Classification: Prophage-like
Match size (bp): 44000 Active/highly abundant Prophage-like element
Whole-community read coverage
VLP-fraction read coverage
Contig position (bp)
unclassified prophage
WC3-2_k141_39068 Classification: Prophage-like
Match size (bp): 32000
Whole-community read coverage
VLP-fraction read coverage
Contig position (bp)
unclassified prophage
WC1-2_k141_180137 Classification: Prophage-like
Match size (bp): 10000 Not homogenously integrated Prophage-like element
Whole-community read coverage
VLP-fraction read coverage
Contig position (bp)
unclassified
WC1-2_k141_187216 Classification: Prophage-like
Match size (bp): 65000
Whole-community read coverage
VLP-fraction read coverage
Contig position (bp)
unclassified
WC1-6_k141_284129 Classification: Sloping
Match size (bp): 30000 Slope: 0.001
Whole-community read coverage
VLP-fraction read coverage
Contig position (bp)
unclassified
WC1-7_k141_39086 Classification: Prophage-like
Match size (bp): 74000 Active/highly abundant Prophage-like element
Whole-community read coverage
VLP-fraction read coverage
Contig position (bp)
unclassified

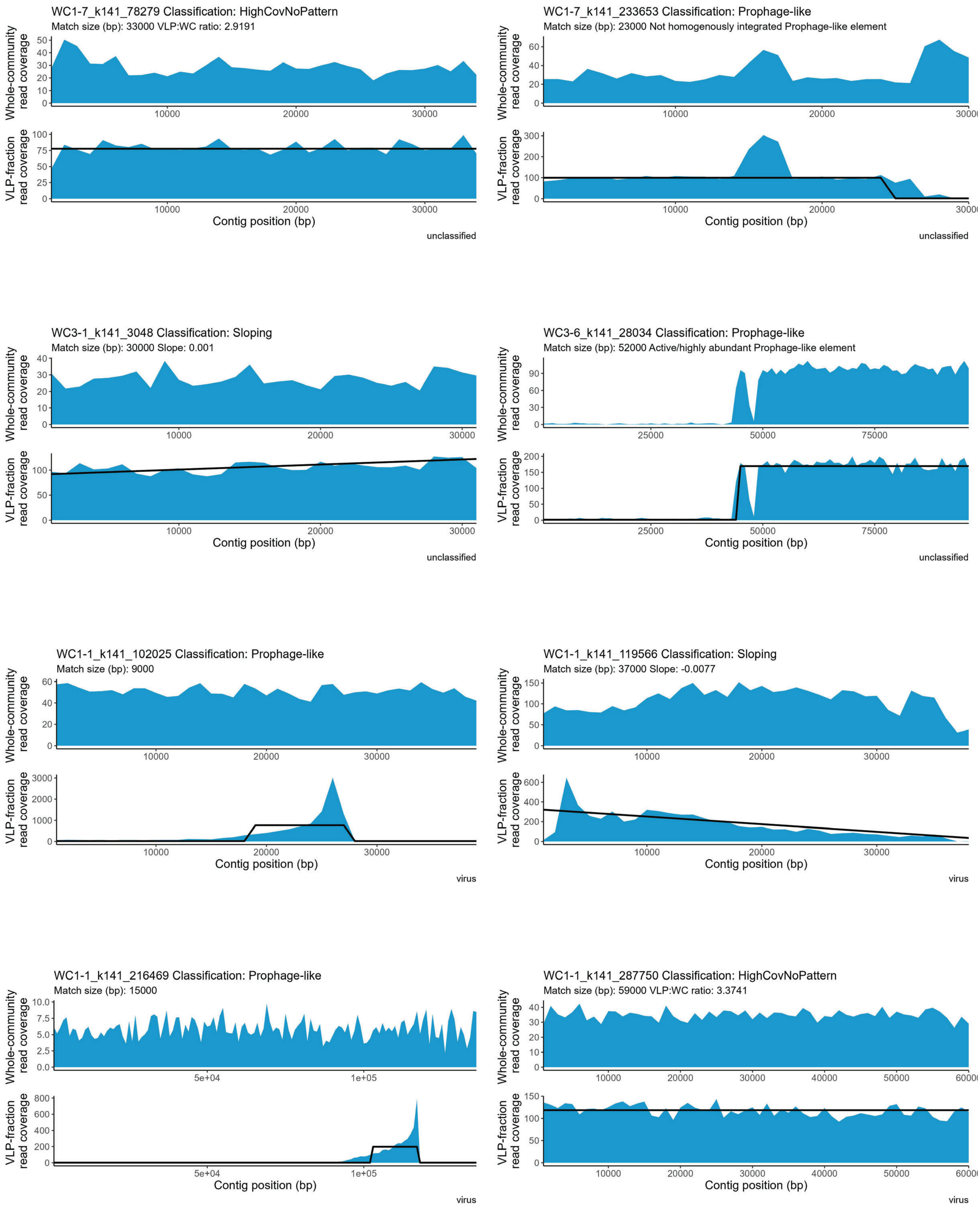
WC1-7_k141_78279 Classification: HighCovNoPattern
Match size (bp): 33000 VLP:WC ratio: 2.9191
unclassified
WC1-7_k141_233653 Classification: Prophage-like
Match size (bp): 23000 Not homogenously integrated Prophage-like element
unclassified
WC3-1_k141_3048 Classification: Sloping
Match size (bp): 30000 Slope: 0.001
unclassified
WC3-6_k141_28034 Classification: Prophage-like
Match size (bp): 52000 Active/highly abundant Prophage-like element
unclassified
WC1-1_k141_102025 Classification: Prophage-like
Match size (bp): 9000
virus
WC1-1_k141_119566 Classification: Sloping
Match size (bp): 37000 Slope: -0.0077
virus
WC1-1_k141_216469 Classification: Prophage-like
Match size (bp): 15000
virus
WC1-1_k141_287750 Classification: HighCovNoPattern
Match size (bp): 59000 VLP:WC ratio: 3.3741
virus
Whole-community read coverage
VLP-fraction read coverage
Contig position (bp)

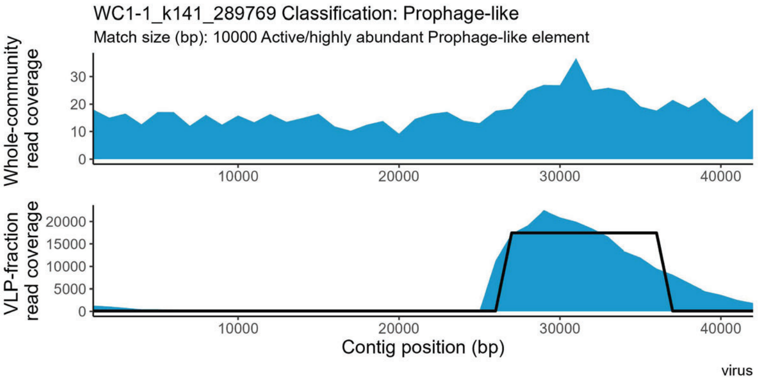
WC1-1_k141_289769 Classification: Prophage-like
Match size (bp): 10000 Active/highly abundant Prophage-like element
Whole-community read coverage
VLP-fraction read coverage
Contig position (bp)
virus

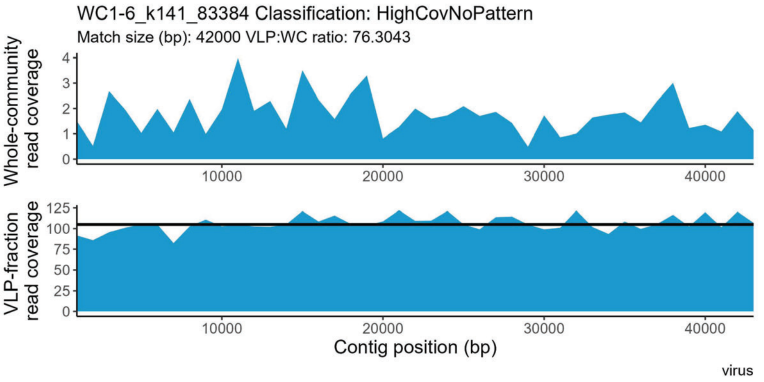
WC1-6_k141_83384 Classification: HighCovNoPattern
Match size (bp): 42000 VLP:WC ratio: 76.3043
Whole-community read coverage
VLP-fraction read coverage
Contig position (bp)
virus

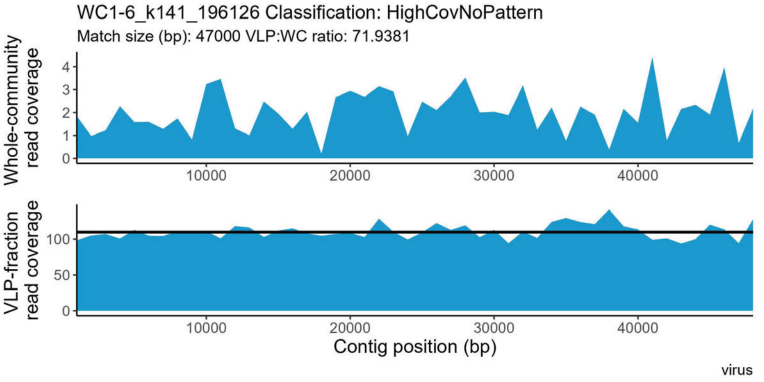
WC1-6_k141_196126 Classification: HighCovNoPattern
Match size (bp): 47000 VLP:WC ratio: 71.9381
Whole-community read coverage
VLP-fraction read coverage
Contig position (bp)
virus

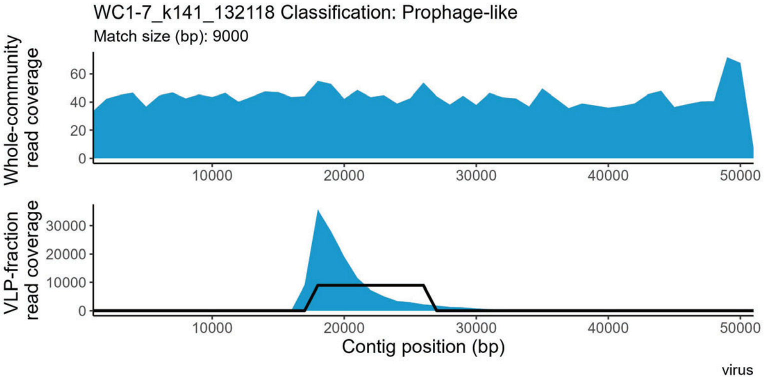
WC1-7_k141_132118 Classification: Prophage-like
Match size (bp): 9000
Whole-community read coverage
VLP-fraction read coverage
Contig position (bp)
virus

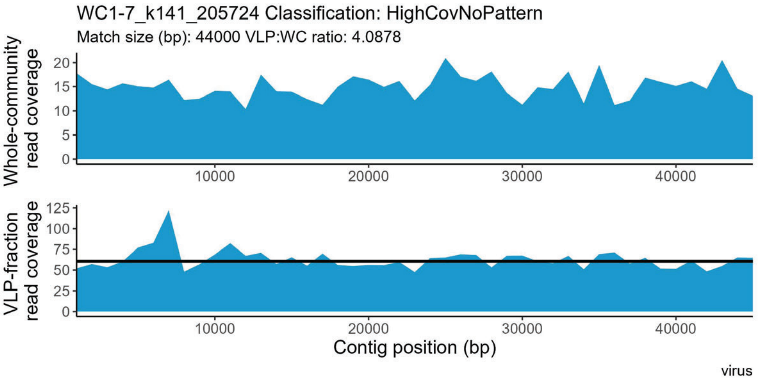
WC1-7_k141_205724 Classification: HighCovNoPattern
Match size (bp): 44000 VLP:WC ratio: 4.0878
Whole-community read coverage
VLP-fraction read coverage
Contig position (bp)
virus

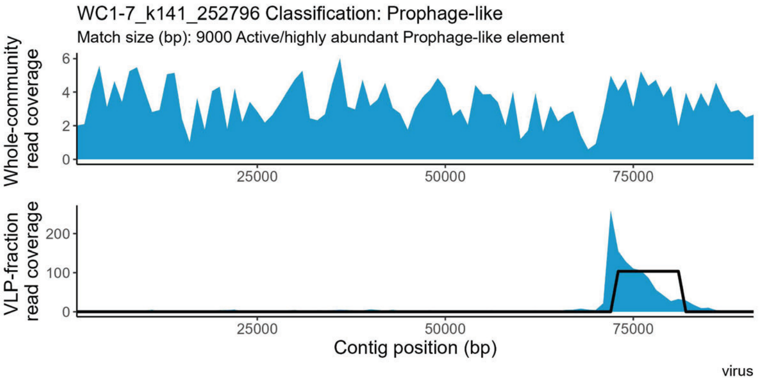
WC1-7_k141_252796 Classification: Prophage-like
Match size (bp): 9000 Active/highly abundant Prophage-like element
Whole-community read coverage
VLP-fraction read coverage
Contig position (bp)
virus

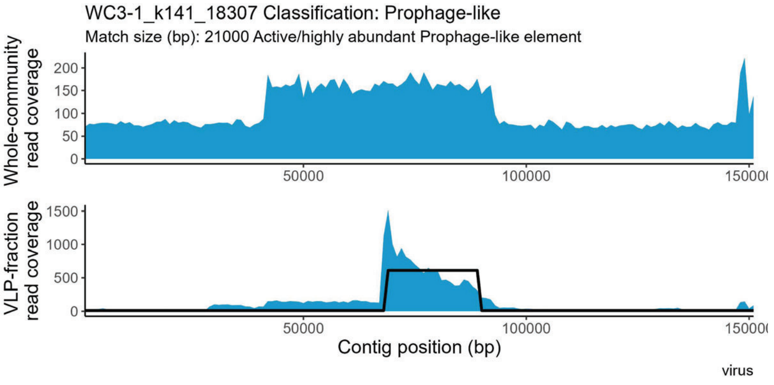
WC3-1_k141_18307 Classification: Prophage-like
Match size (bp): 21000 Active/highly abundant Prophage-like element
Whole-community read coverage
VLP-fraction read coverage
Contig position (bp)
virus

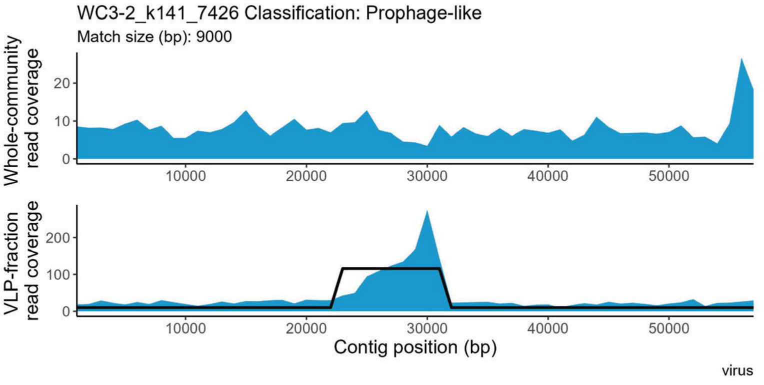
WC3-2_k141_7426 Classification: Prophage-like
Match size (bp): 9000
Whole-community read coverage
VLP-fraction read coverage
Contig position (bp)
virus
