## Supplemental file 8 for "Perturbations shift the composition of bacterial DNA carried by virus-like particles in the murine gut microbiome"

**File S8) All positive TrIdent pattern-matches for CDI replicate 3.** The sample from which a contig was assembled is indicated by the following prefixes in the contig accession number: WC1-1 = Pre-ABX replicate 1, WC1-2 = Pre-ABX replicate 2, WC1-6 = Pre-ABX replicate 3, WC1-7 = Pre-ABX replicate 4, WC1-1 = CDI replicate 1, WC1-2 = CDI replicate 2, WC1-6 = CDI replicate 3, WC1-7 = CDI replicate 4. Pre-ABX = Pre-antibiotics, Post-ABX= post-antbiotics, CDI= *Clostridioides difficile* infection. Contig taxonomy and geNomad prophage classification in plot subtitle. Plots are listed alphabetically by family name.

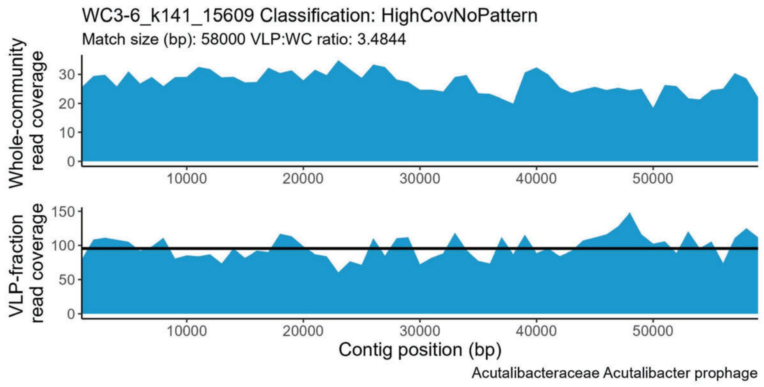


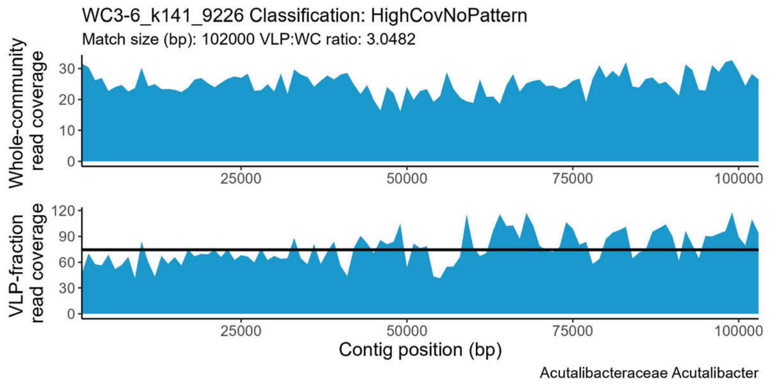


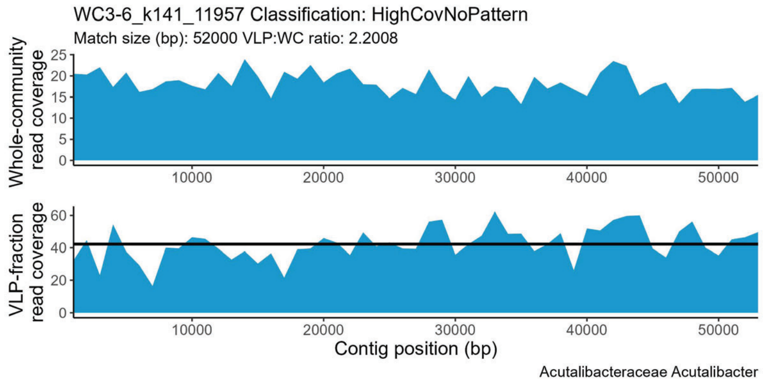


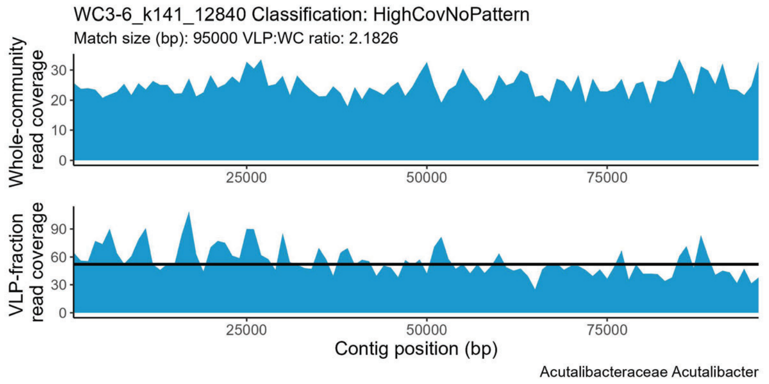


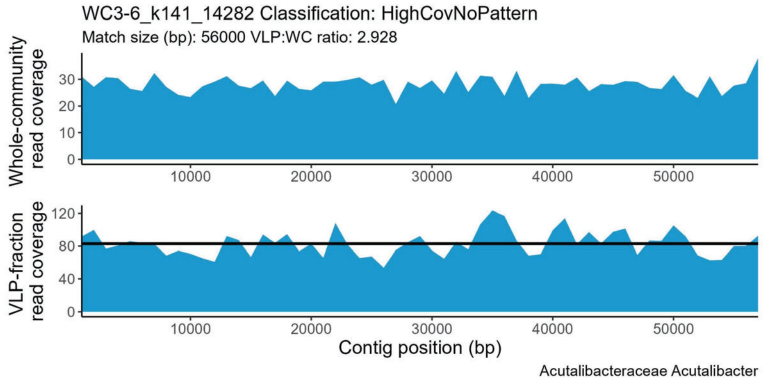


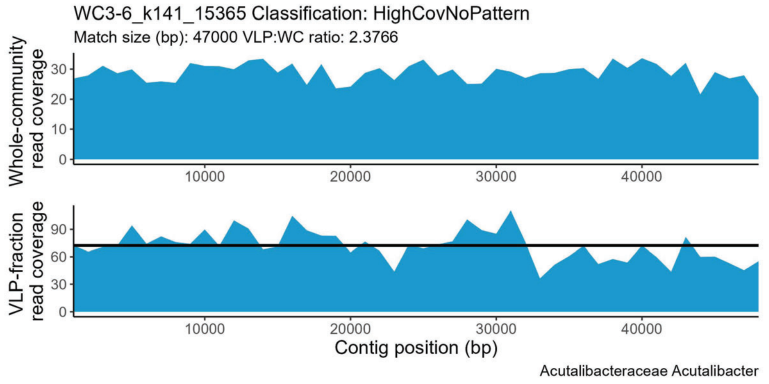


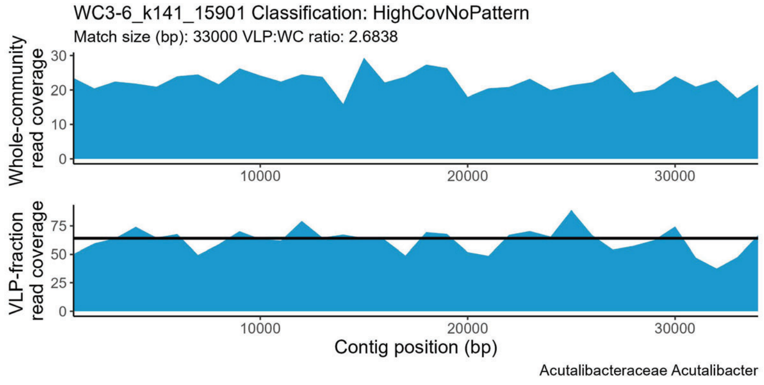


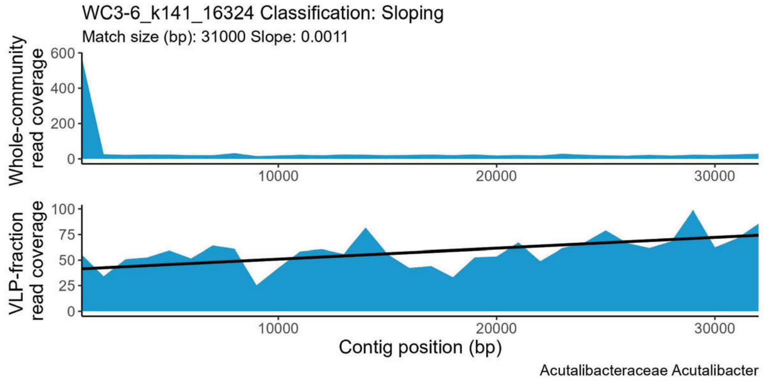

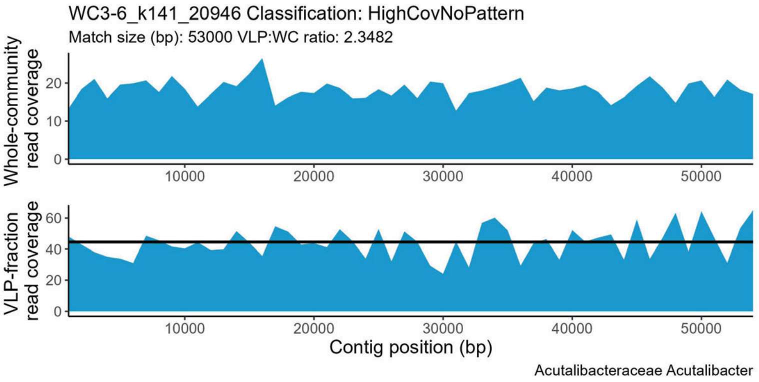
WC3-6_k141_20946 Classification: HighCovNoPattern
Match size (bp): 53000 VLP:WC ratio: 2.3482
Whole-community read coverage
VLP-fraction read coverage
Contig position (bp)
Acutalibacteraceae Acutalibacter

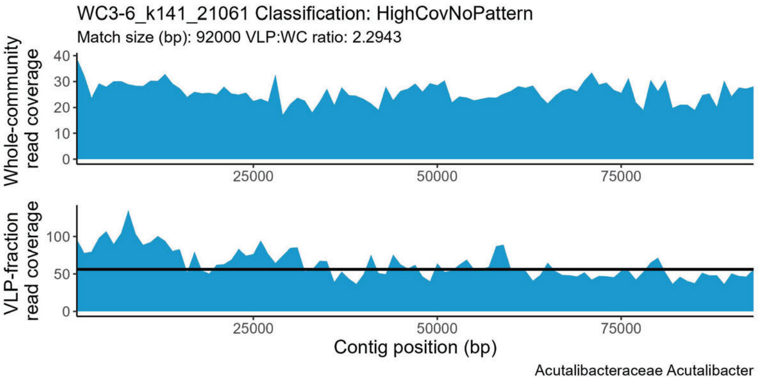
WC3-6_k141_21061 Classification: HighCovNoPattern
Match size (bp): 92000 VLP:WC ratio: 2.2943
Whole-community read coverage
VLP-fraction read coverage
Contig position (bp)
Acutalibacteraceae Acutalibacter

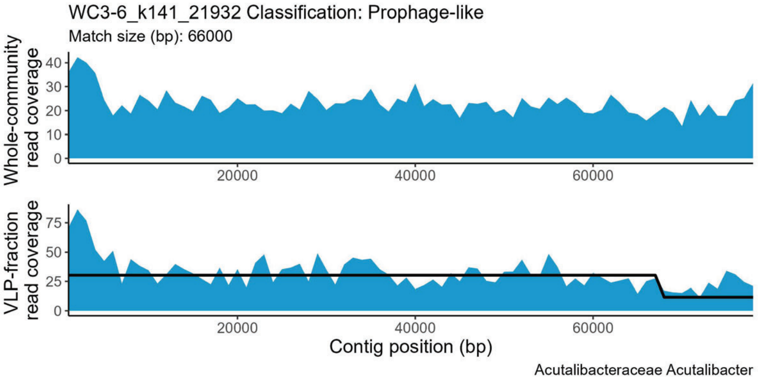
WC3-6_k141_21932 Classification: Prophage-like
Match size (bp): 66000
Whole-community read coverage
VLP-fraction read coverage
Contig position (bp)
Acutalibacteraceae Acutalibacter

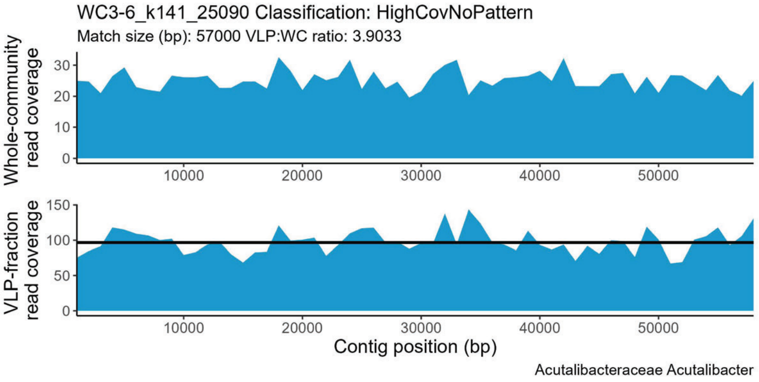
WC3-6_k141_25090 Classification: HighCovNoPattern
Match size (bp): 57000 VLP:WC ratio: 3.9033
Whole-community read coverage
VLP-fraction read coverage
Contig position (bp)
Acutalibacteraceae Acutalibacter

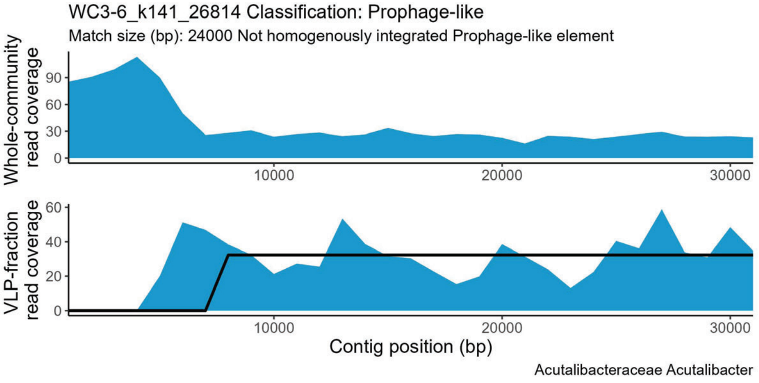
WC3-6_k141_26814 Classification: Prophage-like
Match size (bp): 24000 Not homogenously integrated Prophage-like element
Whole-community read coverage
VLP-fraction read coverage
Contig position (bp)
Acutalibacteraceae Acutalibacter

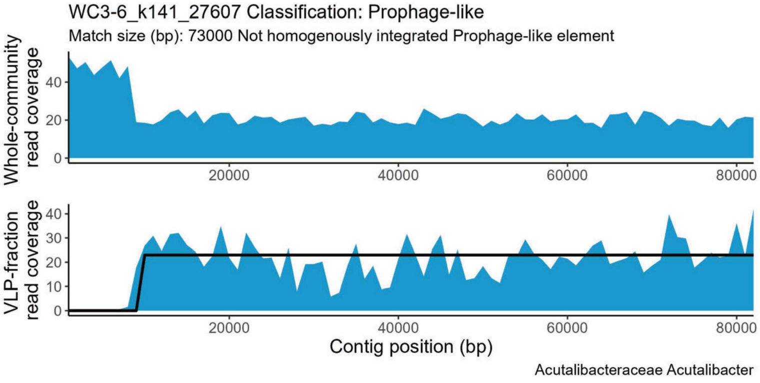
WC3-6_k141_27607 Classification: Prophage-like
Match size (bp): 73000 Not homogenously integrated Prophage-like element
Whole-community read coverage
VLP-fraction read coverage
Contig position (bp)
Acutalibacteraceae Acutalibacter

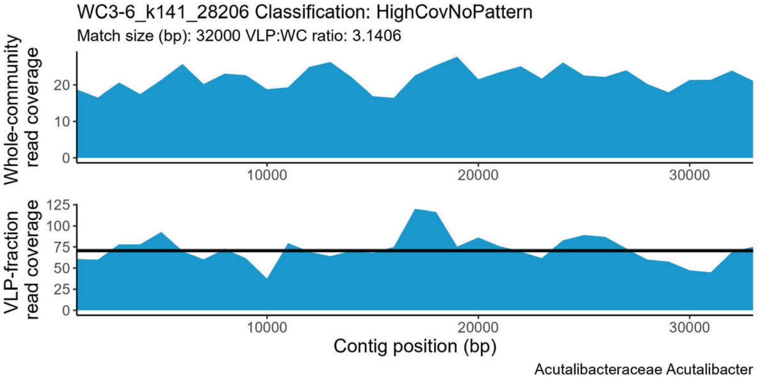
WC3-6_k141_28206 Classification: HighCovNoPattern
Match size (bp): 32000 VLP:WC ratio: 3.1406
Whole-community read coverage
VLP-fraction read coverage
Contig position (bp)
Acutalibacteraceae Acutalibacter

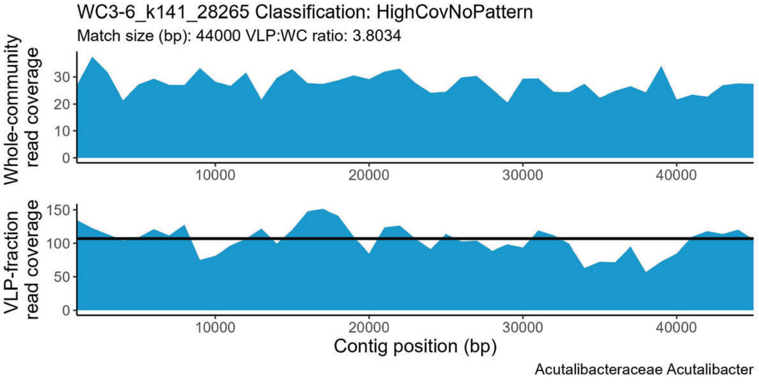
WC3-6_k141_28265 Classification: HighCovNoPattern
Match size (bp): 44000 VLP:WC ratio: 3.8034
Whole-community read coverage
VLP-fraction read coverage
Contig position (bp)
Acutalibacteraceae Acutalibacter

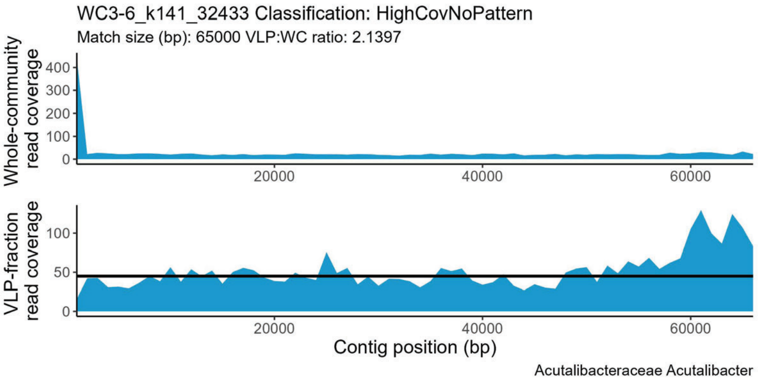
WC3-6_k141_32433 Classification: HighCovNoPattern
Match size (bp): 65000 VLP:WC ratio: 2.1397
Whole-community read coverage
VLP-fraction read coverage
Contig position (bp)
Acutalibacteraceae Acutalibacter

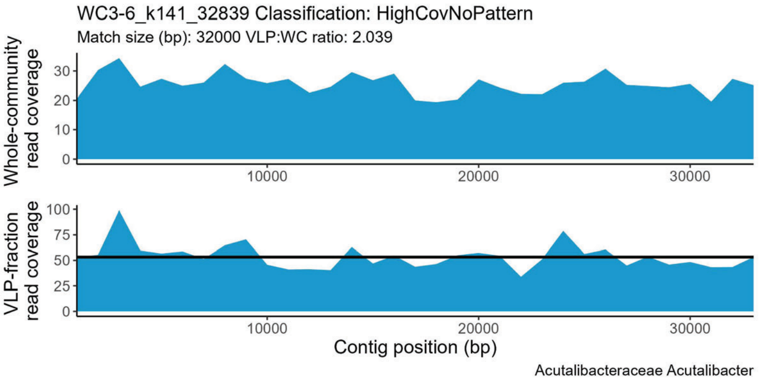
WC3-6_k141_32839 Classification: HighCovNoPattern
Match size (bp): 32000 VLP:WC ratio: 2.039
Whole-community read coverage
VLP-fraction read coverage
Contig position (bp)
Acutalibacteraceae Acutalibacter

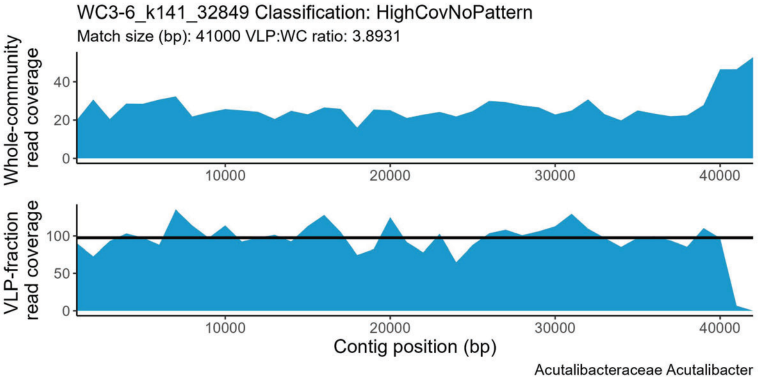
WC3-6_k141_32849 Classification: HighCovNoPattern
Match size (bp): 41000 VLP:WC ratio: 3.8931
Whole-community read coverage
VLP-fraction read coverage
Contig position (bp)
Acutalibacteraceae Acutalibacter

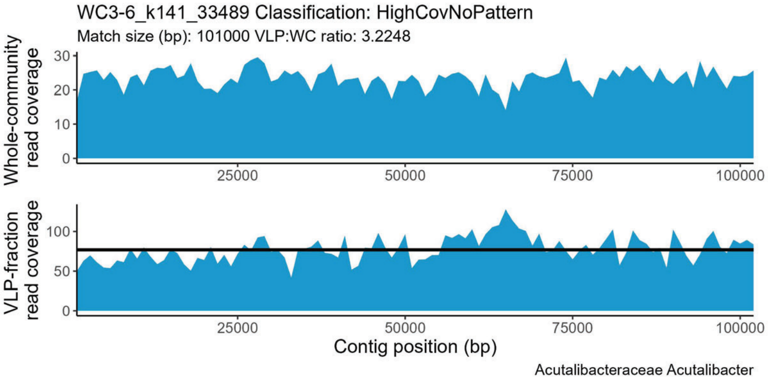
WC3-6_k141_33489 Classification: HighCovNoPattern
Match size (bp): 101000 VLP:WC ratio: 3.2248
Whole-community read coverage
VLP-fraction read coverage
Contig position (bp)
Acutalibacteraceae Acutalibacter

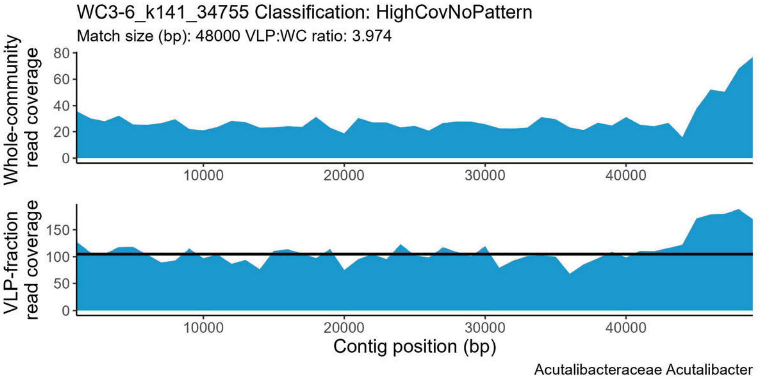
WC3-6_k141_34755 Classification: HighCovNoPattern
Match size (bp): 48000 VLP:WC ratio: 3.974
Whole-community read coverage
VLP-fraction read coverage
Contig position (bp)
Acutalibacteraceae Acutalibacter

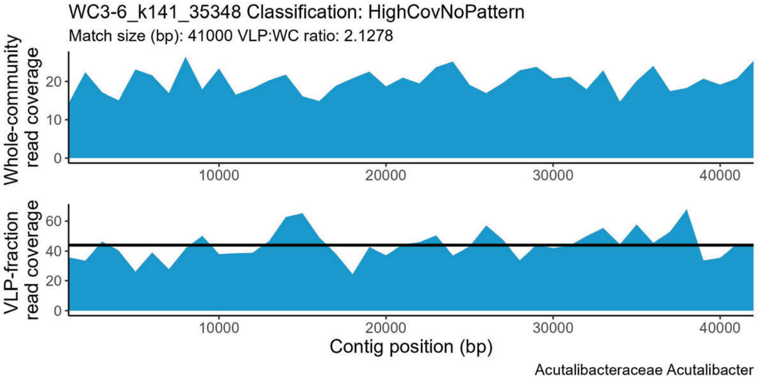
WC3-6_k141_35348 Classification: HighCovNoPattern
Match size (bp): 41000 VLP:WC ratio: 2.1278
Whole-community read coverage
VLP-fraction read coverage
Contig position (bp)
Acutalibacteraceae Acutalibacter

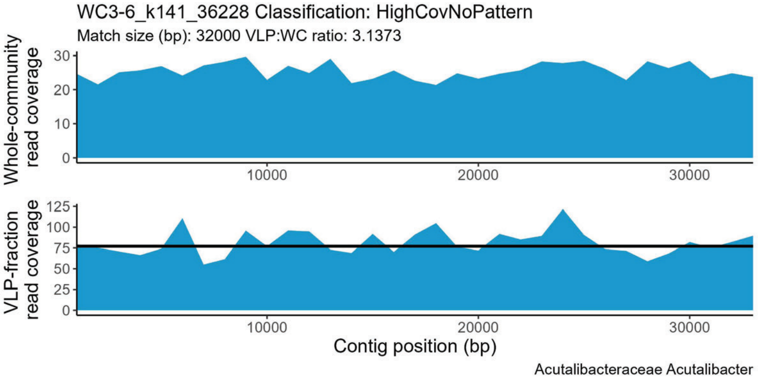
WC3-6_k141_36228 Classification: HighCovNoPattern
Match size (bp): 32000 VLP:WC ratio: 3.1373
Whole-community read coverage
VLP-fraction read coverage
Contig position (bp)
Acutalibacteraceae Acutalibacter

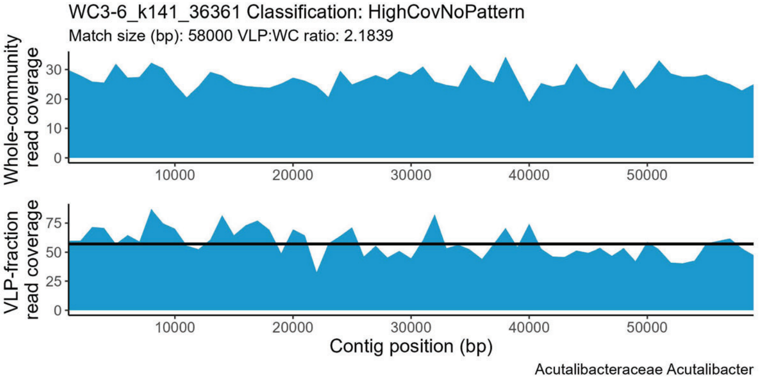
WC3-6_k141_36361 Classification: HighCovNoPattern
Match size (bp): 58000 VLP:WC ratio: 2.1839
Whole-community read coverage
VLP-fraction read coverage
Contig position (bp)
Acutalibacteraceae Acutalibacter

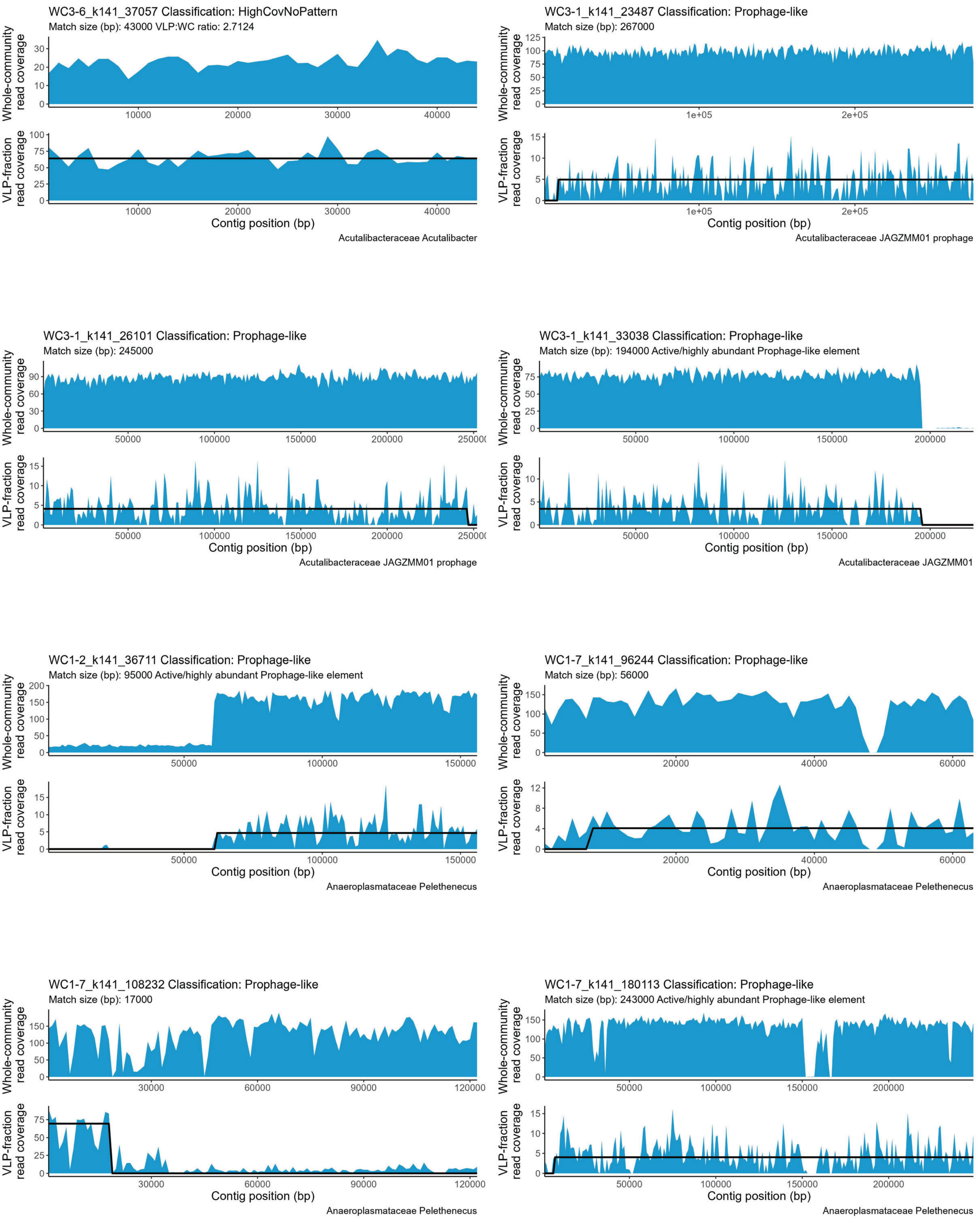
WC3-6_k141_37057 Classification: HighCovNoPattern
Match size (bp): 43000 VLP:WC ratio: 2.7124
Whole-community read coverage
VLP-fraction read coverage
Contig position (bp)
Acutalibacteraceae Acutalibacter
WC3-1_k141_23487 Classification: Prophage-like
Match size (bp): 267000
Whole-community read coverage
VLP-fraction read coverage
Contig position (bp)
Acutalibacteraceae JAGZMM01 prophage
WC3-1_k141_26101 Classification: Prophage-like
Match size (bp): 245000
Whole-community read coverage
VLP-fraction read coverage
Contig position (bp)
Acutalibacteraceae JAGZMM01 prophage
WC3-1_k141_33038 Classification: Prophage-like
Match size (bp): 194000 Active/highly abundant Prophage-like element
Whole-community read coverage
VLP-fraction read coverage
Contig position (bp)
Acutalibacteraceae JAGZMM01
WC1-2_k141_36711 Classification: Prophage-like
Match size (bp): 95000 Active/highly abundant Prophage-like element
Whole-community read coverage
VLP-fraction read coverage
Contig position (bp)
Anaeroplasmataceae Pelethenecus
WC1-7_k141_96244 Classification: Prophage-like
Match size (bp): 56000
Whole-community read coverage
VLP-fraction read coverage
Contig position (bp)
Anaeroplasmataceae Pelethenecus
WC1-7_k141_108232 Classification: Prophage-like
Match size (bp): 17000
Whole-community read coverage
VLP-fraction read coverage
Contig position (bp)
Anaeroplasmataceae Pelethenecus
WC1-7_k141_180113 Classification: Prophage-like
Match size (bp): 243000 Active/highly abundant Prophage-like element
Whole-community read coverage
VLP-fraction read coverage
Contig position (bp)
Anaeroplasmataceae Pelethenecus

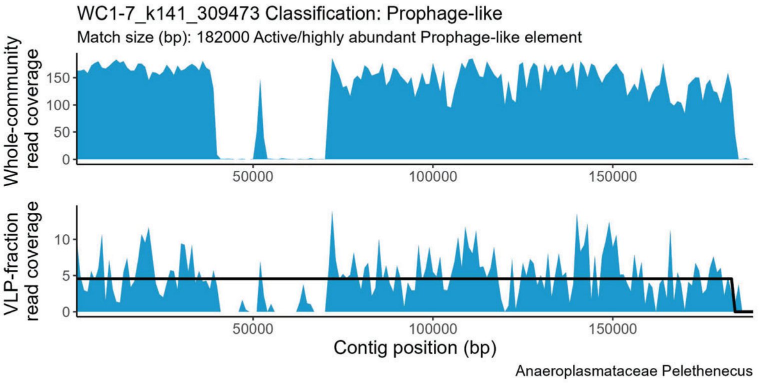
WC1-7_k141_309473 Classification: Prophage-like
Match size (bp): 182000 Active/highly abundant Prophage-like element
Whole-community read coverage
VLP-fraction read coverage
Contig position (bp)
50000
100000
150000
Anaeroplasmataceae Pelethenecus

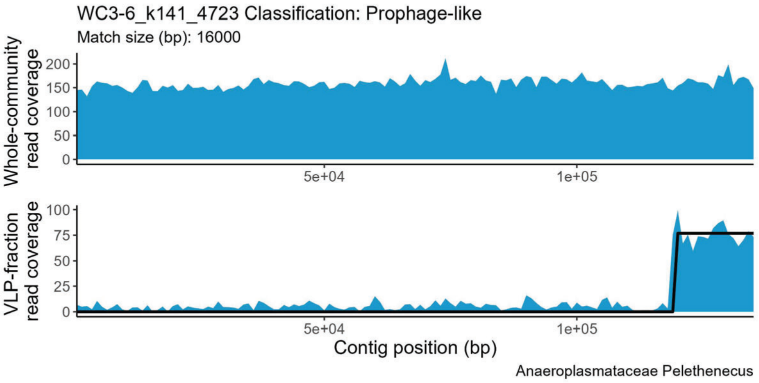
WC3-6_k141_4723 Classification: Prophage-like
Match size (bp): 16000
Whole-community read coverage
VLP-fraction read coverage
Contig position (bp)
5e+04
1e+05
Anaeroplasmataceae Pelethenecus

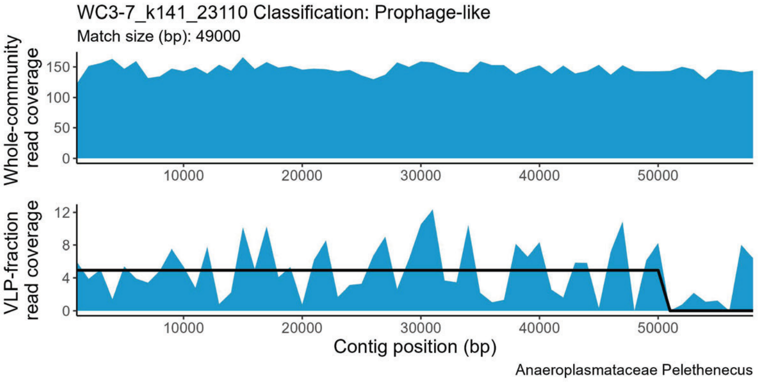
WC3-7_k141_23110 Classification: Prophage-like
Match size (bp): 49000
Whole-community read coverage
VLP-fraction read coverage
Contig position (bp)
10000
20000
30000
40000
50000
Anaeroplasmataceae Pelethenecus

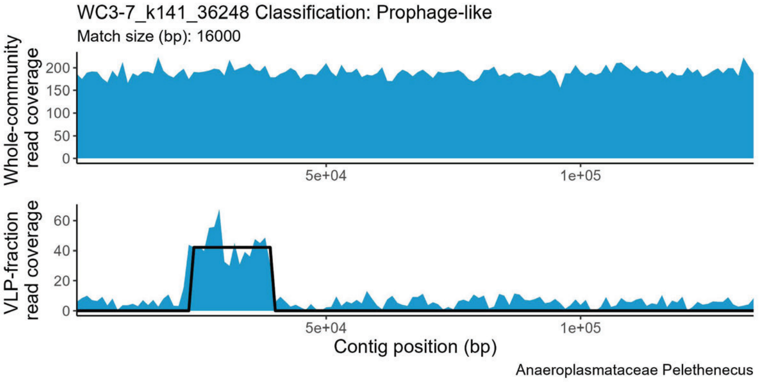
WC3-7_k141_36248 Classification: Prophage-like
Match size (bp): 16000
Whole-community read coverage
VLP-fraction read coverage
Contig position (bp)
5e+04
1e+05
Anaeroplasmataceae Pelethenecus

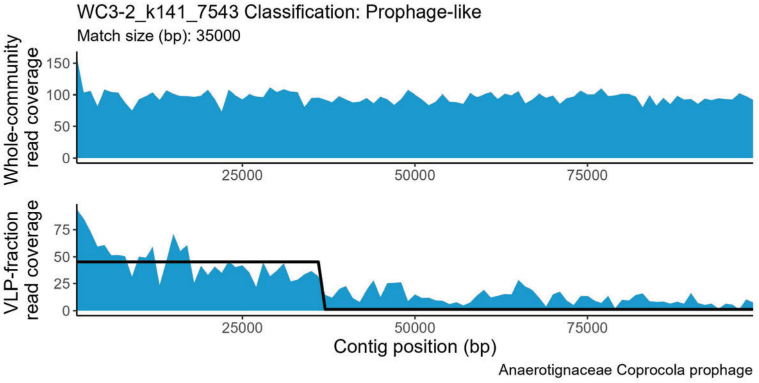
WC3-2_k141_7543 Classification: Prophage-like
Match size (bp): 35000
Whole-community read coverage
VLP-fraction read coverage
Contig position (bp)
25000
50000
75000
Anaerotignaceae Coprocola prophage

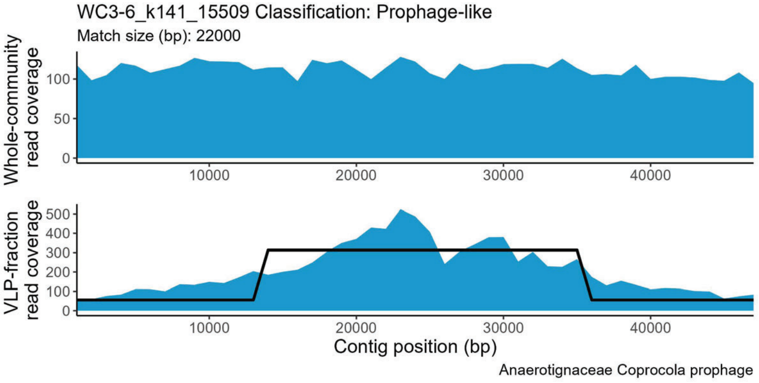
WC3-6_k141_15509 Classification: Prophage-like
Match size (bp): 22000
Whole-community read coverage
VLP-fraction read coverage
Contig position (bp)
10000
20000
30000
40000
Anaerotignaceae Coprocola prophage

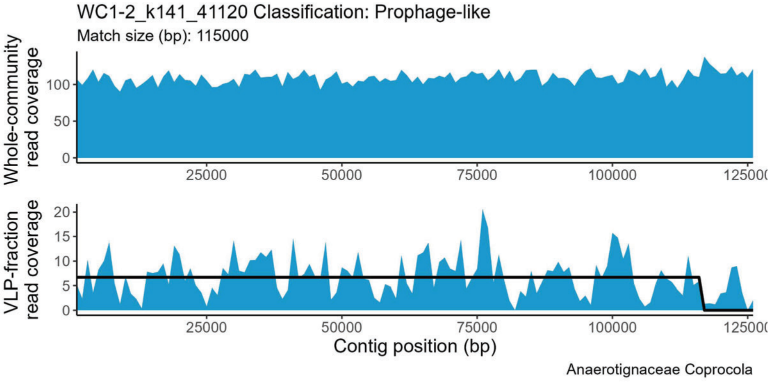
WC1-2_k141_41120 Classification: Prophage-like
Match size (bp): 115000
Whole-community read coverage
VLP-fraction read coverage
Contig position (bp)
25000
50000
75000
100000
Anaerotignaceae Coprocola

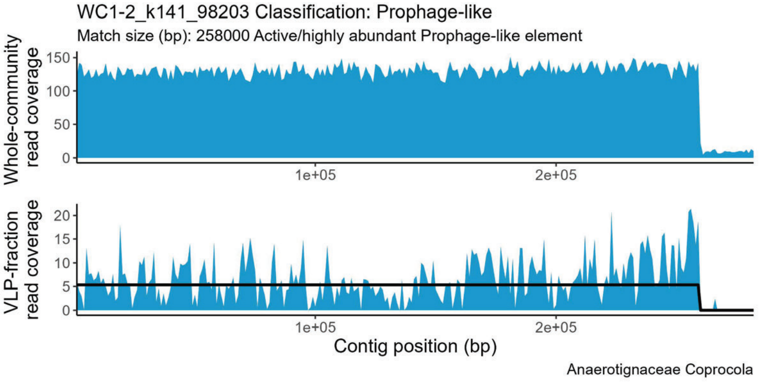
WC1-2_k141_98203 Classification: Prophage-like
Match size (bp): 258000 Active/highly abundant Prophage-like element
Whole-community read coverage
VLP-fraction read coverage
Contig position (bp)
1e+05
2e+05
Anaerotignaceae Coprocola

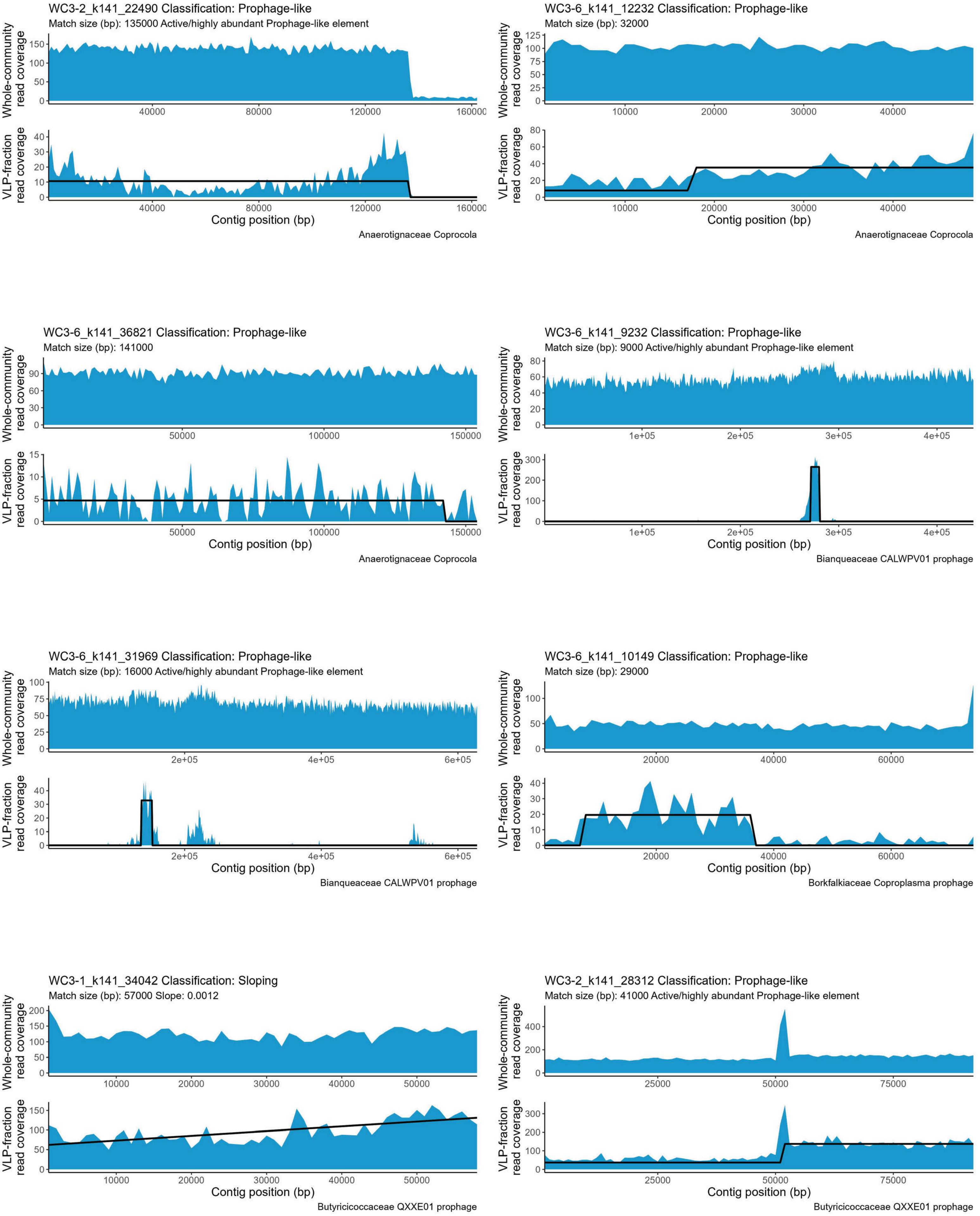
WC3-2_k141_22490 Classification: Prophage-like
Match size (bp): 135000 Active/highly abundant Prophage-like element
Whole-community read coverage
VLP-fraction read coverage
Contig position (bp)
Anaerotignaceae Coprocola
WC3-6_k141_12232 Classification: Prophage-like
Match size (bp): 32000
Whole-community read coverage
VLP-fraction read coverage
Contig position (bp)
Anaerotignaceae Coprocola
WC3-6_k141_36821 Classification: Prophage-like
Match size (bp): 141000
Whole-community read coverage
VLP-fraction read coverage
Contig position (bp)
Anaerotignaceae Coprocola
WC3-6_k141_9232 Classification: Prophage-like
Match size (bp): 9000 Active/highly abundant Prophage-like element
Whole-community read coverage
VLP-fraction read coverage
Contig position (bp)
Bianqueaceae CALWPV01 prophage
WC3-6_k141_31969 Classification: Prophage-like
Match size (bp): 16000 Active/highly abundant Prophage-like element
Whole-community read coverage
VLP-fraction read coverage
Contig position (bp)
Bianqueaceae CALWPV01 prophage
WC3-6_k141_10149 Classification: Prophage-like
Match size (bp): 29000
Whole-community read coverage
VLP-fraction read coverage
Contig position (bp)
Borkfalkiaceae Coproplasma prophage
WC3-1_k141_34042 Classification: Sloping
Match size (bp): 57000 Slope: 0.0012
Whole-community read coverage
VLP-fraction read coverage
Contig position (bp)
Butyricicoccaceae QXXE01 prophage
WC3-2_k141_28312 Classification: Prophage-like
Match size (bp): 41000 Active/highly abundant Prophage-like element
Whole-community read coverage
VLP-fraction read coverage
Contig position (bp)
Butyricicoccaceae QXXE01 prophage

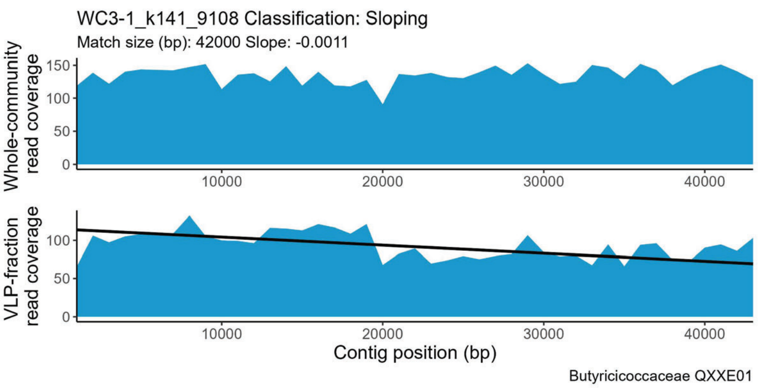
WC3-1_k141_9108 Classification: Sloping
Match size (bp): 42000 Slope: -0.0011
Whole-community read coverage
0
50
100
150
VLP-fraction read coverage
0
50
100
10000
20000
30000
40000
Contig position (bp)
Butyricicoccaceae QXXE01

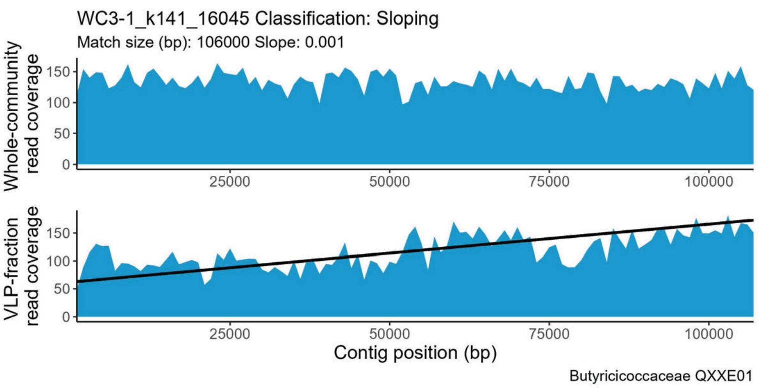
WC3-1_k141_16045 Classification: Sloping
Match size (bp): 106000 Slope: 0.001
Whole-community read coverage
0
50
100
150
VLP-fraction read coverage
0
50
100
150
25000
50000
75000
100000
Contig position (bp)
Butyricicoccaceae QXXE01

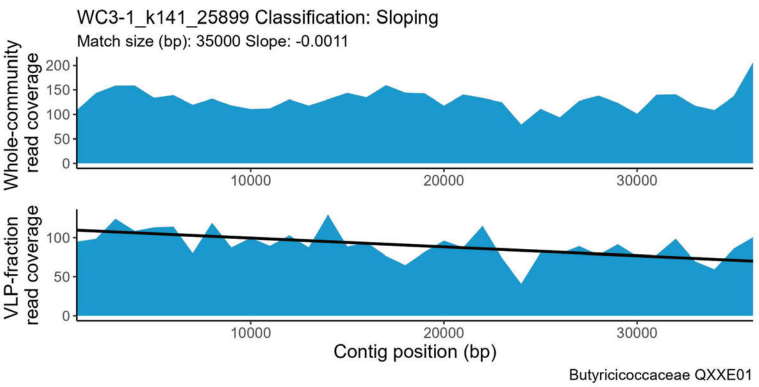
WC3-1_k141_25899 Classification: Sloping
Match size (bp): 35000 Slope: -0.0011
Whole-community read coverage
0
50
100
150
200
VLP-fraction read coverage
0
50
100
10000
20000
30000
Contig position (bp)
Butyricicoccaceae QXXE01

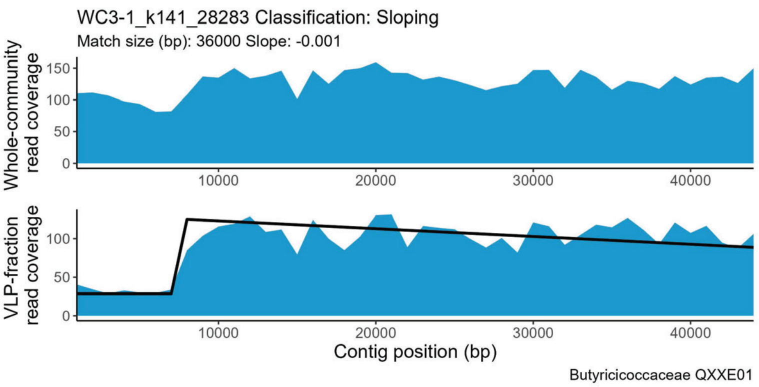
WC3-1_k141_28283 Classification: Sloping
Match size (bp): 36000 Slope: -0.001
Whole-community read coverage
0
50
100
150
VLP-fraction read coverage
0
50
100
10000
20000
30000
40000
Contig position (bp)
Butyricicoccaceae QXXE01

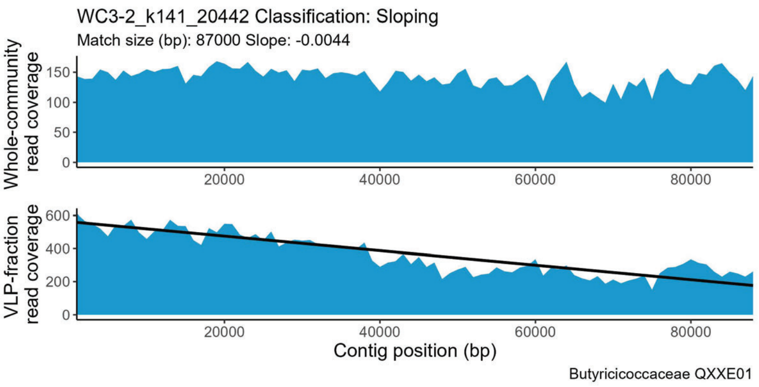
WC3-2_k141_20442 Classification: Sloping
Match size (bp): 87000 Slope: -0.0044
Whole-community read coverage
0
50
100
150
VLP-fraction read coverage
0
200
400
600
20000
40000
60000
80000
Contig position (bp)
Butyricicoccaceae QXXE01

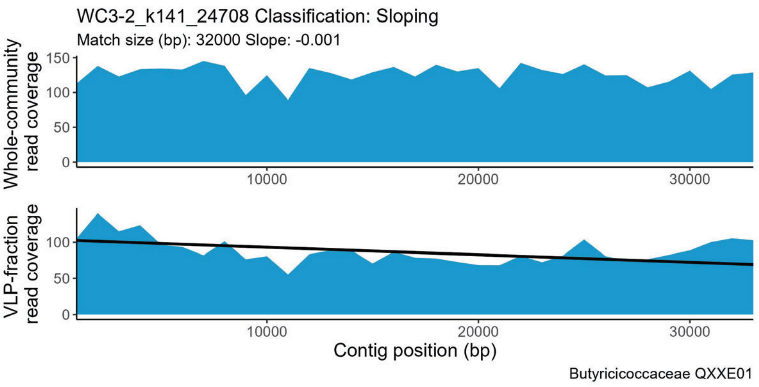
WC3-2_k141_24708 Classification: Sloping
Match size (bp): 32000 Slope: -0.001
Whole-community read coverage
0
50
100
150
VLP-fraction read coverage
0
50
100
10000
20000
30000
Contig position (bp)
Butyricicoccaceae QXXE01

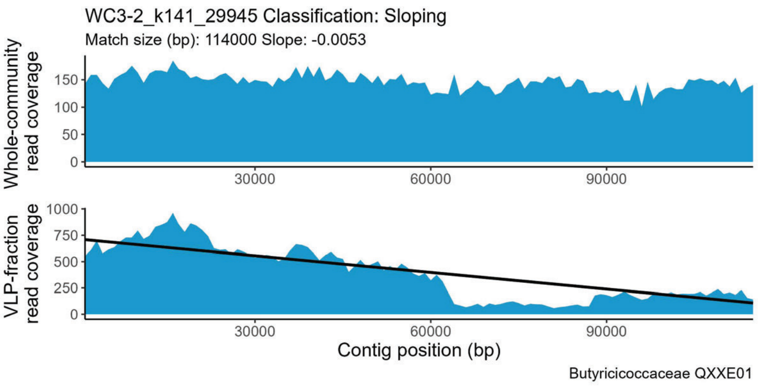
WC3-2_k141_29945 Classification: Sloping
Match size (bp): 114000 Slope: -0.0053
Whole-community read coverage
0
50
100
150
VLP-fraction read coverage
0
250
500
750
1000
30000
60000
90000
Contig position (bp)
Butyricicoccaceae QXXE01

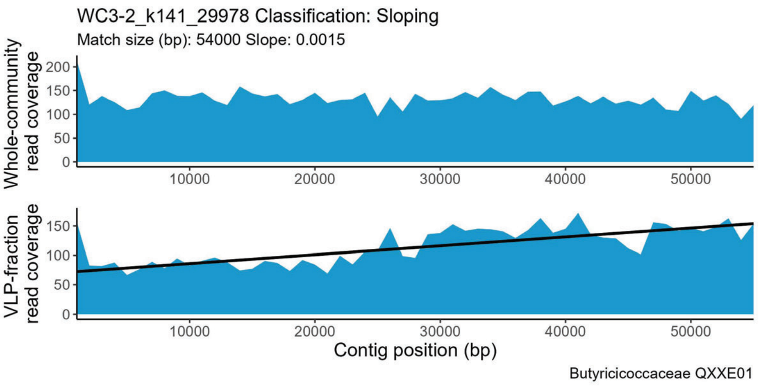
WC3-2_k141_29978 Classification: Sloping
Match size (bp): 54000 Slope: 0.0015
Whole-community read coverage
0
50
100
150
200
VLP-fraction read coverage
0
50
100
150
10000
20000
30000
40000
50000
Contig position (bp)
Butyricicoccaceae QXXE01

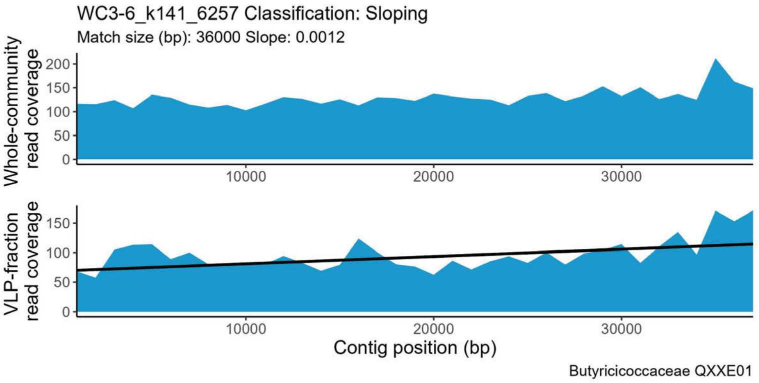
WC3-6_k141_6257 Classification: Sloping
Match size (bp): 36000 Slope: 0.0012
Whole-community read coverage
VLP-fraction read coverage
Contig position (bp)
Butyricicoccaceae QXXE01

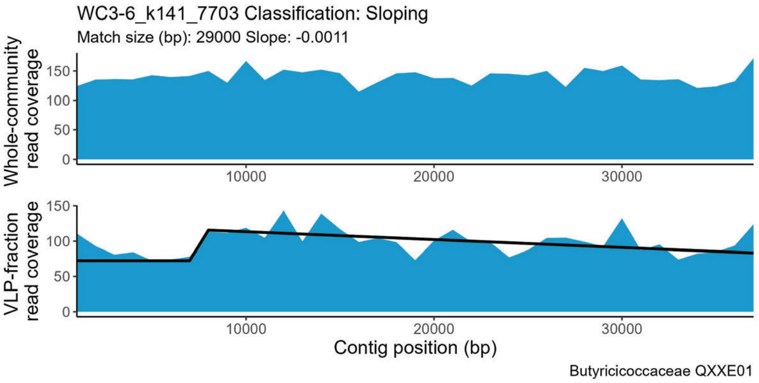
WC3-6_k141_7703 Classification: Sloping
Match size (bp): 29000 Slope: -0.0011
Whole-community read coverage
VLP-fraction read coverage
Contig position (bp)
Butyricicoccaceae QXXE01

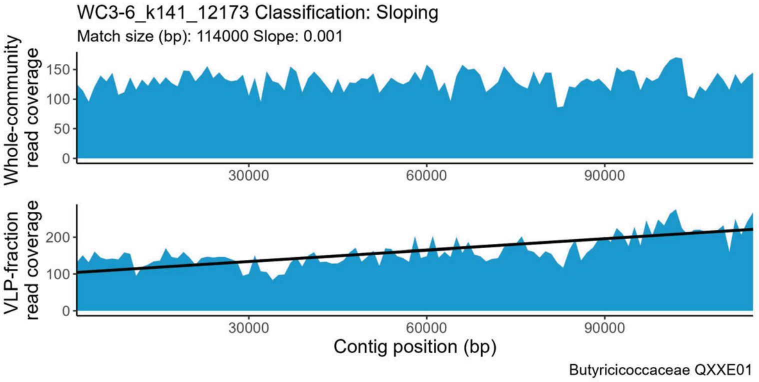
WC3-6_k141_12173 Classification: Sloping
Match size (bp): 114000 Slope: 0.001
Whole-community read coverage
VLP-fraction read coverage
Contig position (bp)
Butyricicoccaceae QXXE01

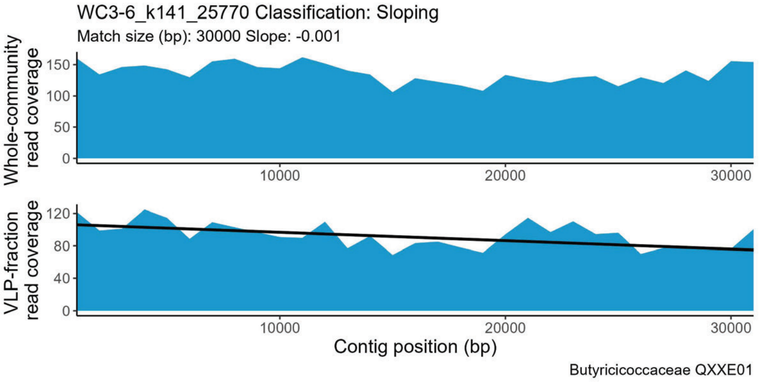
WC3-6_k141_25770 Classification: Sloping
Match size (bp): 30000 Slope: -0.001
Whole-community read coverage
VLP-fraction read coverage
Contig position (bp)
Butyricicoccaceae QXXE01

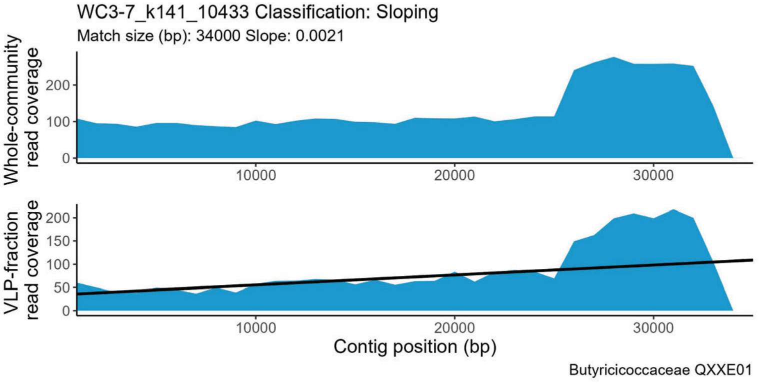
WC3-7_k141_10433 Classification: Sloping
Match size (bp): 34000 Slope: 0.0021
Whole-community read coverage
VLP-fraction read coverage
Contig position (bp)
Butyricicoccaceae QXXE01

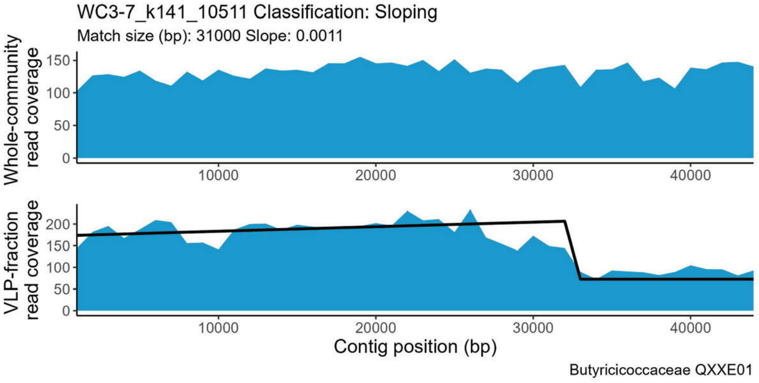
WC3-7_k141_10511 Classification: Sloping
Match size (bp): 31000 Slope: 0.0011
Whole-community read coverage
VLP-fraction read coverage
Contig position (bp)
Butyricicoccaceae QXXE01

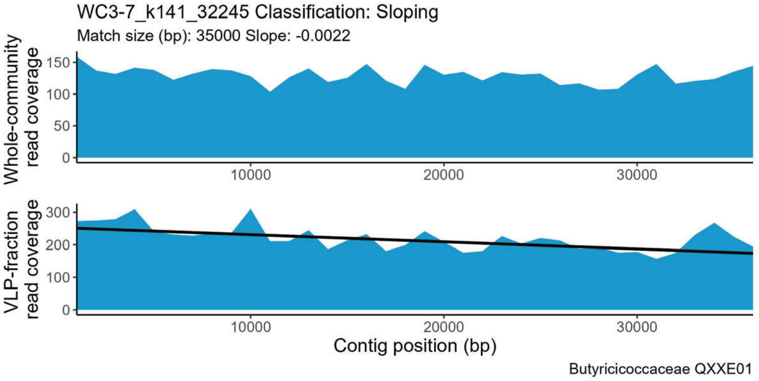
WC3-7_k141_32245 Classification: Sloping
Match size (bp): 35000 Slope: -0.0022
Whole-community read coverage
VLP-fraction read coverage
Contig position (bp)
Butyricicoccaceae QXXE01

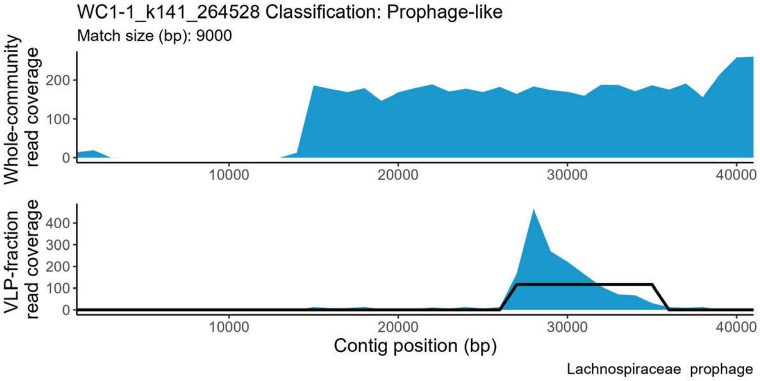
WC1-1_k141_264528 Classification: Prophage-like
Match size (bp): 9000
Whole-community read coverage
VLP-fraction read coverage
Contig position (bp)
Lachnospiraceae prophage

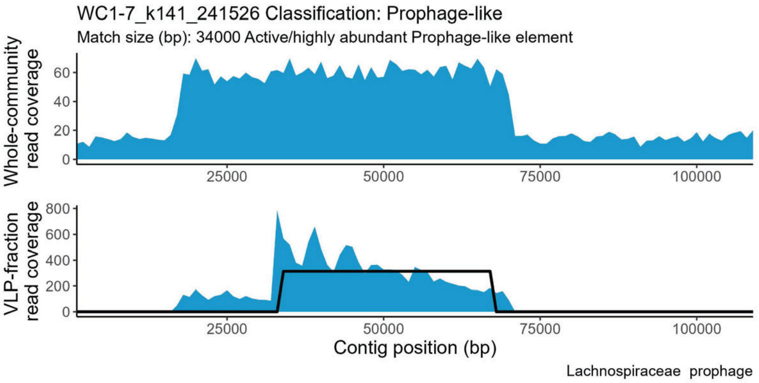
WC1-7_k141_241526 Classification: Prophage-like
Match size (bp): 34000 Active/highly abundant Prophage-like element
Whole-community read coverage
VLP-fraction read coverage
Contig position (bp)
Lachnospiraceae  prophage

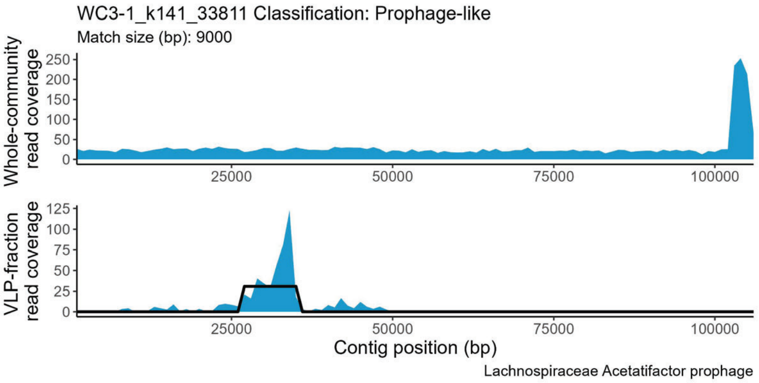
WC3-1_k141_33811 Classification: Prophage-like
Match size (bp): 9000
Whole-community read coverage
VLP-fraction read coverage
Contig position (bp)
Lachnospiraceae Acetatifactor prophage

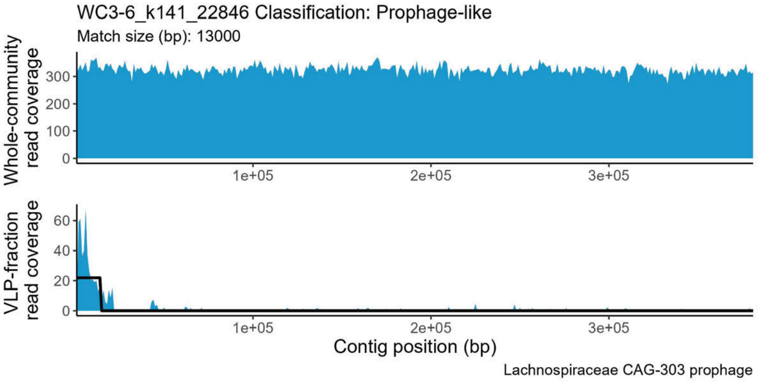
WC3-6_k141_22846 Classification: Prophage-like
Match size (bp): 13000
Whole-community read coverage
VLP-fraction read coverage
Contig position (bp)
Lachnospiraceae CAG-303 prophage

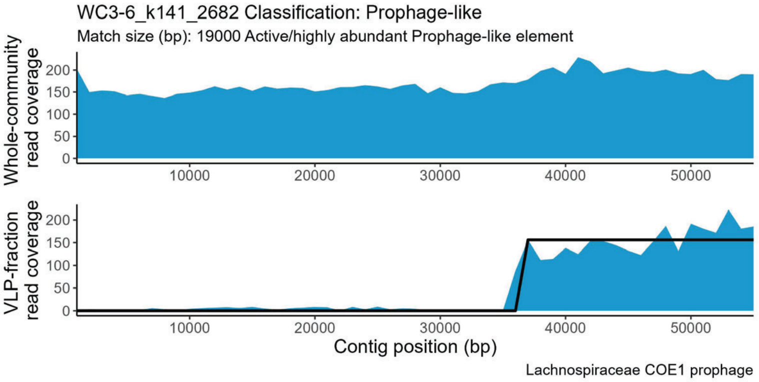
WC3-6_k141_2682 Classification: Prophage-like
Match size (bp): 19000 Active/highly abundant Prophage-like element
Whole-community read coverage
VLP-fraction read coverage
Contig position (bp)
Lachnospiraceae COE1 prophage

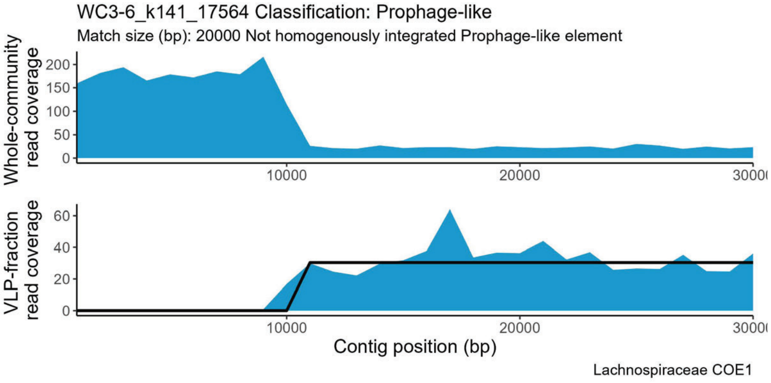
WC3-6_k141_17564 Classification: Prophage-like
Match size (bp): 20000 Not homogenously integrated Prophage-like element
Whole-community read coverage
VLP-fraction read coverage
Contig position (bp)
Lachnospiraceae COE1

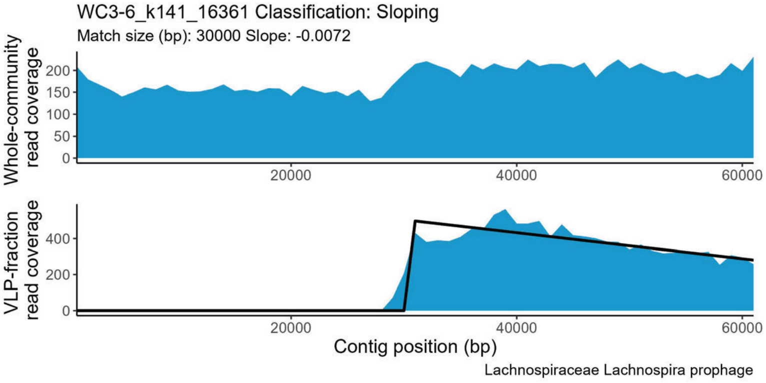
WC3-6_k141_16361 Classification: Sloping
Match size (bp): 30000 Slope: -0.0072
Whole-community read coverage
VLP-fraction read coverage
Contig position (bp)
Lachnospiraceae Lachnospira prophage

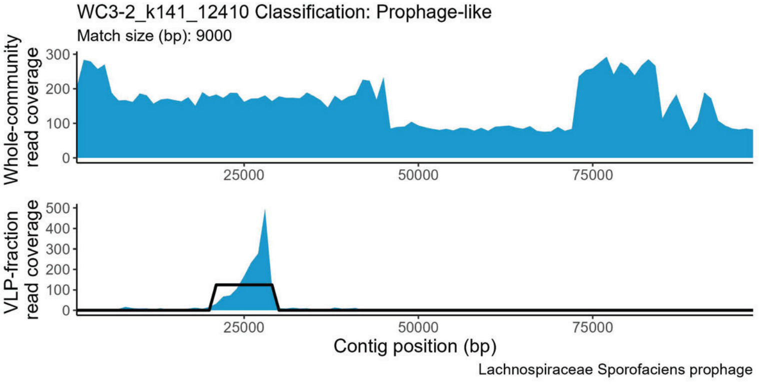
WC3-2_k141_12410 Classification: Prophage-like
Match size (bp): 9000
Whole-community read coverage
VLP-fraction read coverage
Contig position (bp)
Lachnospiraceae Sporofaciens prophage

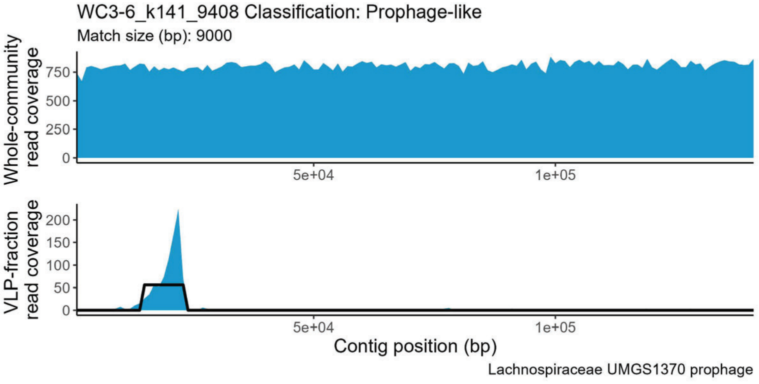
WC3-6_k141_9408 Classification: Prophage-like
Match size (bp): 9000
Whole-community read coverage
VLP-fraction read coverage
Contig position (bp)
Lachnospiraceae UMGS1370 prophage

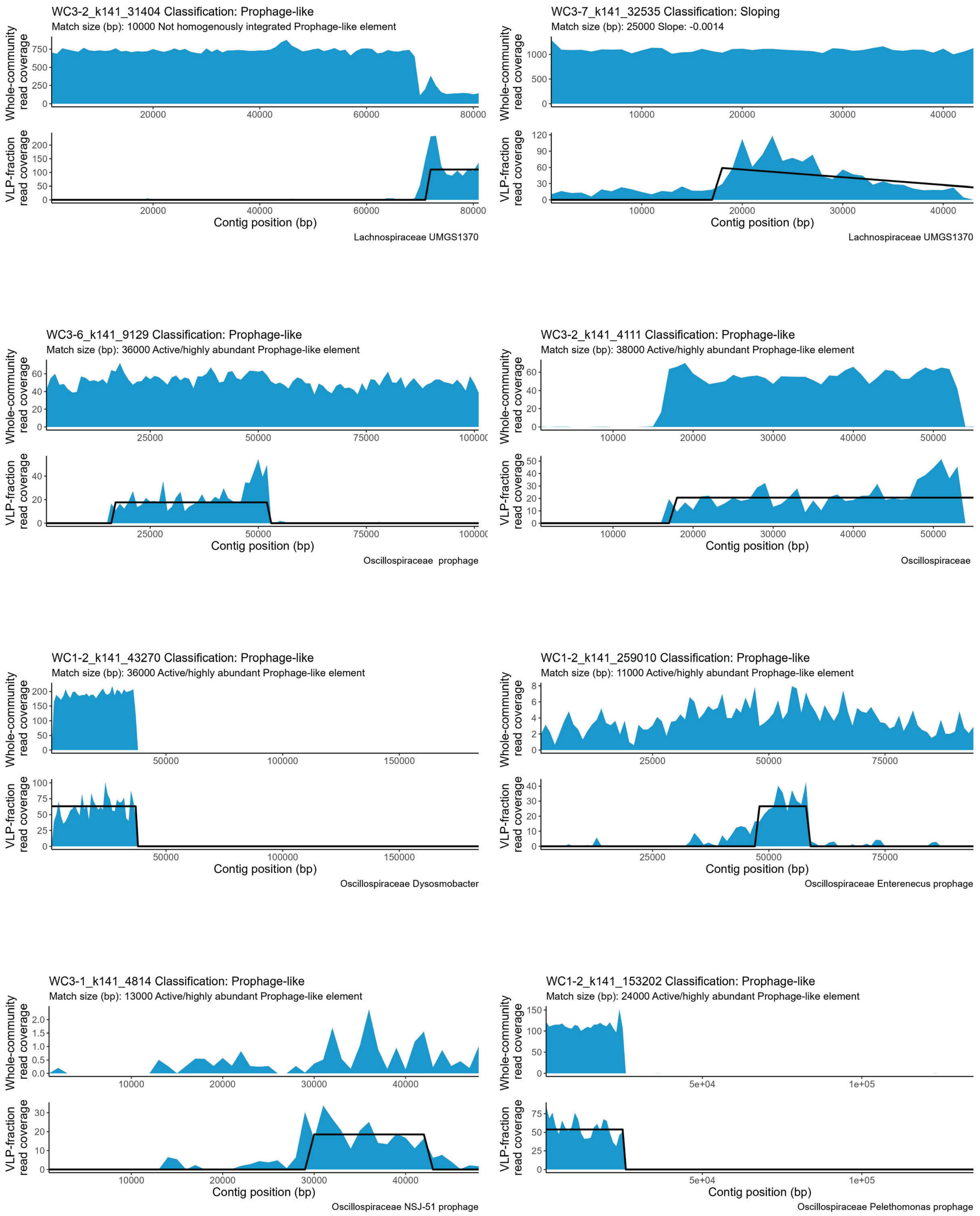
WC3-2_k141_31404 Classification: Prophage-like
Match size (bp): 10000 Not homogenously integrated Prophage-like element
Whole-community read coverage
VLP-fraction read coverage
Contig position (bp)
Lachnospiraceae UMGS1370
WC3-7_k141_32535 Classification: Sloping
Match size (bp): 25000 Slope: -0.0014
Whole-community read coverage
VLP-fraction read coverage
Contig position (bp)
Lachnospiraceae UMGS1370
WC3-6_k141_9129 Classification: Prophage-like
Match size (bp): 36000 Active/highly abundant Prophage-like element
Whole-community read coverage
VLP-fraction read coverage
Contig position (bp)
Oscillospiraceae prophage
WC3-2_k141_4111 Classification: Prophage-like
Match size (bp): 38000 Active/highly abundant Prophage-like element
Whole-community read coverage
VLP-fraction read coverage
Contig position (bp)
Oscillospiraceae
WC1-2_k141_43270 Classification: Prophage-like
Match size (bp): 36000 Active/highly abundant Prophage-like element
Whole-community read coverage
VLP-fraction read coverage
Contig position (bp)
Oscillospiraceae Dysosmobacter
WC1-2_k141_259010 Classification: Prophage-like
Match size (bp): 11000 Active/highly abundant Prophage-like element
Whole-community read coverage
VLP-fraction read coverage
Contig position (bp)
Oscillospiraceae Enterenecus prophage
WC3-1_k141_4814 Classification: Prophage-like
Match size (bp): 13000 Active/highly abundant Prophage-like element
Whole-community read coverage
VLP-fraction read coverage
Contig position (bp)
Oscillospiraceae NSJ-51 prophage
WC1-2_k141_153202 Classification: Prophage-like
Match size (bp): 24000 Active/highly abundant Prophage-like element
Whole-community read coverage
VLP-fraction read coverage
Contig position (bp)
Oscillospiraceae Pelethomonas prophage

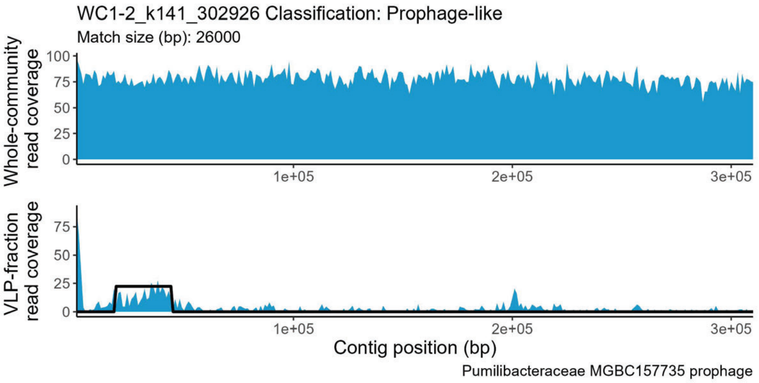
WC1-2_k141_302926 Classification: Prophage-like
Match size (bp): 26000
Whole-community read coverage
VLP-fraction read coverage
Contig position (bp)
Pumilibacteraceae MGBC157735 prophage

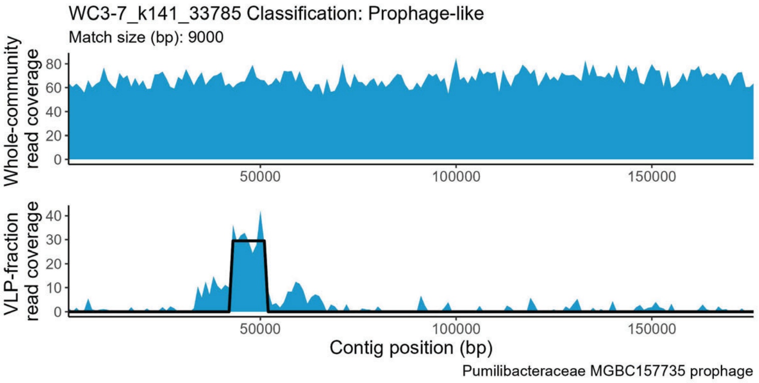
WC3-7_k141_33785 Classification: Prophage-like
Match size (bp): 9000
Whole-community read coverage
VLP-fraction read coverage
Contig position (bp)
Pumilibacteraceae MGBC157735 prophage

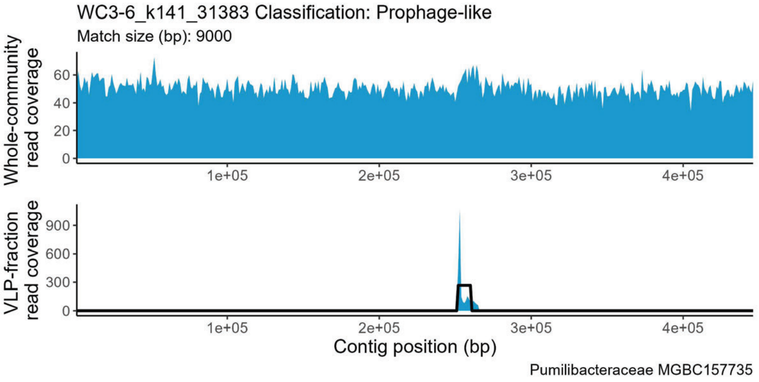
WC3-6_k141_31383 Classification: Prophage-like
Match size (bp): 9000
Whole-community read coverage
VLP-fraction read coverage
Contig position (bp)
Pumilibacteraceae MGBC157735

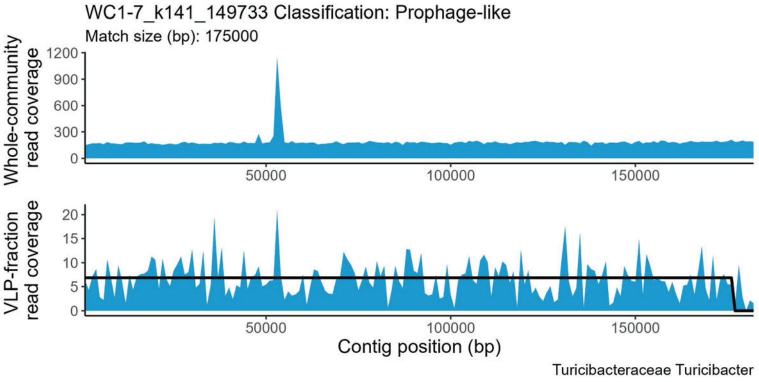
WC1-7_k141_149733 Classification: Prophage-like
Match size (bp): 175000
Whole-community read coverage
VLP-fraction read coverage
Contig position (bp)
Turicibacteraceae Turicibacter

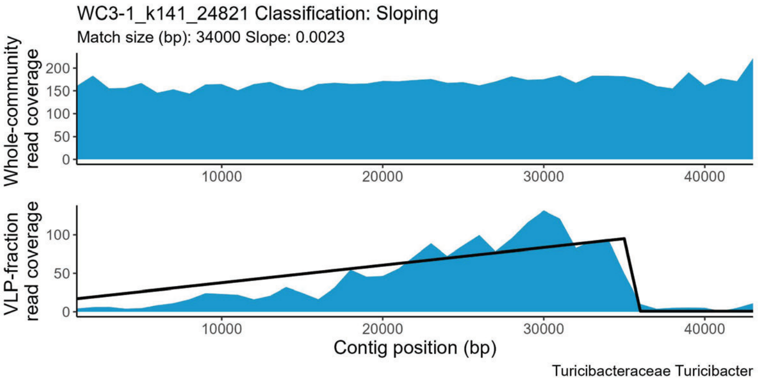
WC3-1_k141_24821 Classification: Sloping
Match size (bp): 34000 Slope: 0.0023
Whole-community read coverage
VLP-fraction read coverage
Contig position (bp)
Turicibacteraceae Turicibacter

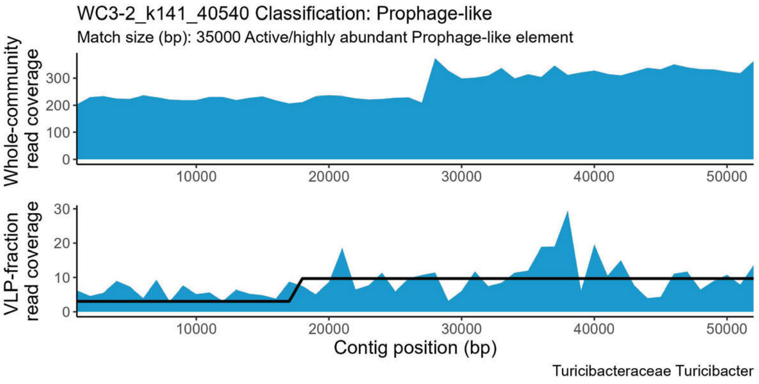
WC3-2_k141_40540 Classification: Prophage-like
Match size (bp): 35000 Active/highly abundant Prophage-like element
Whole-community read coverage
VLP-fraction read coverage
Contig position (bp)
Turicibacteraceae Turicibacter

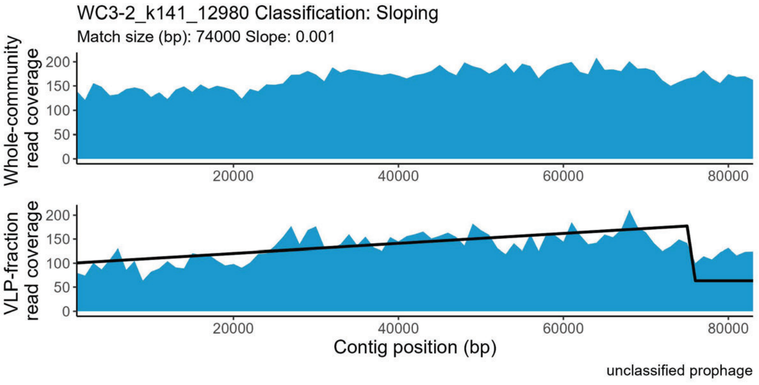
WC3-2_k141_12980 Classification: Sloping
Match size (bp): 74000 Slope: 0.001
Whole-community read coverage
VLP-fraction read coverage
Contig position (bp)
unclassified prophage

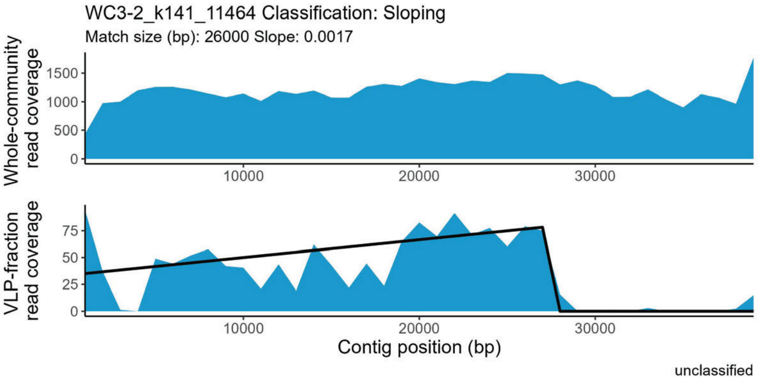
WC3-2_k141_11464 Classification: Sloping
Match size (bp): 26000 Slope: 0.0017
Whole-community read coverage
VLP-fraction read coverage
Contig position (bp)
unclassified
