## Supplemental file 5 for "Perturbations shift the composition of bacterial DNA carried by virus-like particles in the murine gut microbiome"

**File S5) All positive TrIdent pattern-matches for Post-ABX replicate 3.** The sample from which a contig was assembled is indicated by the following prefixes in the contig accession number: WC1-1 = Pre-ABX replicate 1, WC1-2 = Pre-ABX replicate 2, WC1-6 = Pre-ABX replicate 3, WC1-7 = Pre-ABX replicate 4, WC1-1 = CDI replicate 1, WC1-2 = CDI replicate 2, WC1-6 = CDI replicate 3, WC1-7 = CDI replicate 4. Pre-ABX = Pre-antibiotics, Post-ABX= post-antbiotics, CDI= *Clostridioides difficile* infection. Contig taxonomy and geNomad prophage classification in plot subtitle. Plots are listed alphabetically by family name.

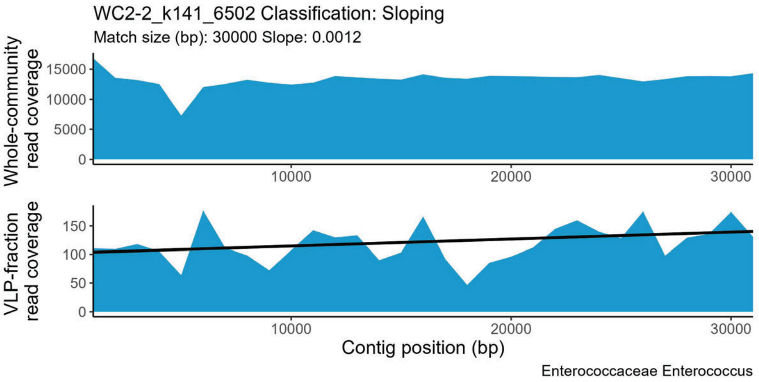


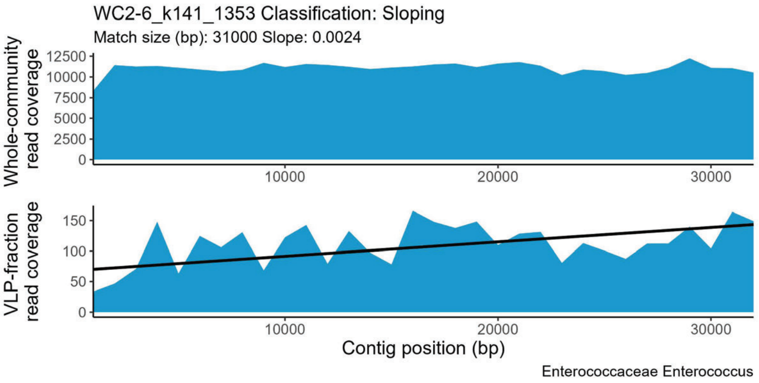


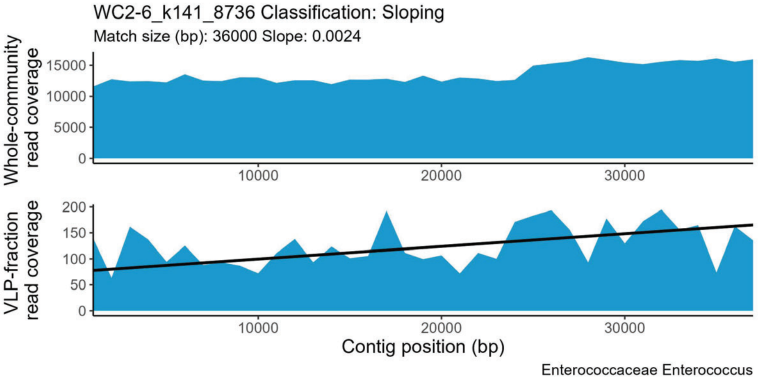


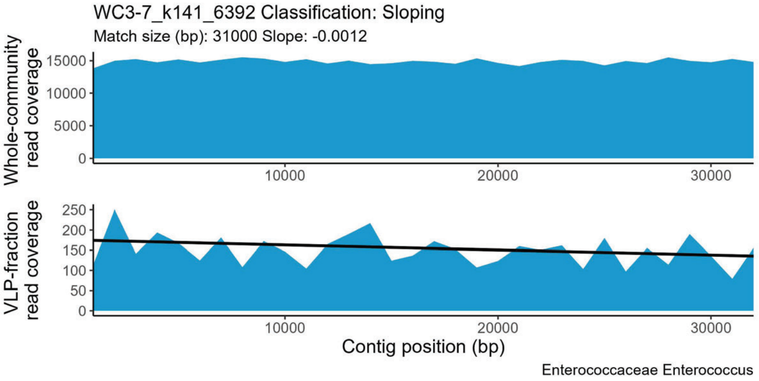


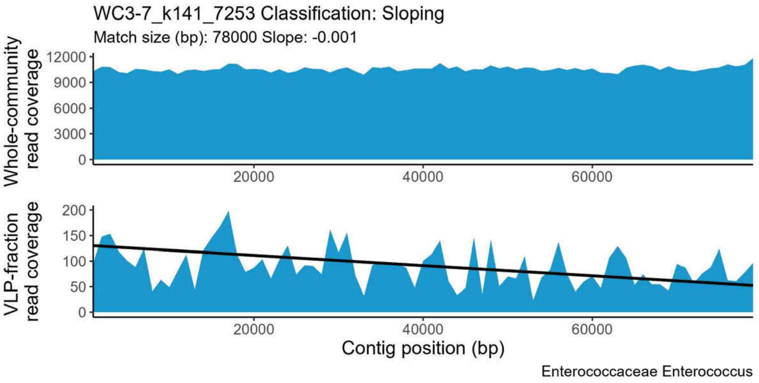


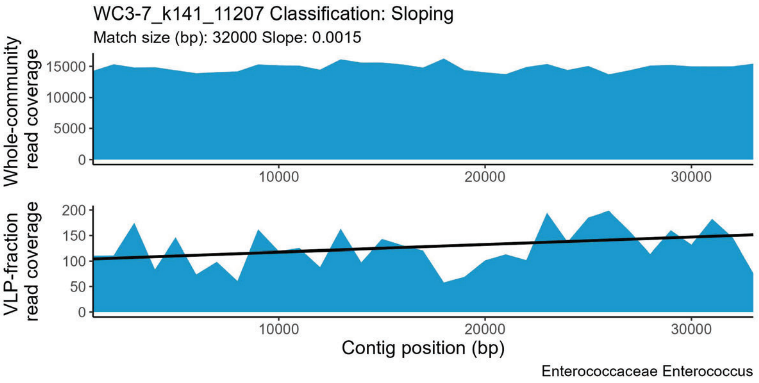


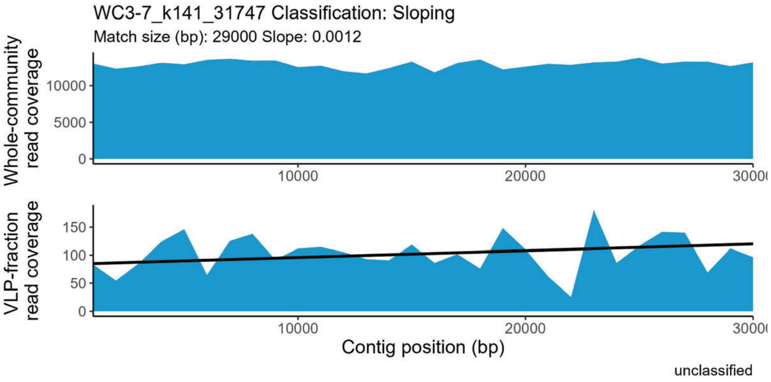
