## Supplemental file 3 for "Perturbations shift the composition of bacterial DNA carried by virus-like particles in the murine gut microbiome"

**File S3) All positive TrIdent pattern-matches for Pre-ABX replicate 3.** The sample from which a contig was assembled is indicated by the following prefixes in the contig accession number: WC1-1 = Pre-ABX replicate 1, WC1-2 = Pre-ABX replicate 2, WC1-6 = Pre-ABX replicate 3, WC1-7 = Pre-ABX replicate 4, WC1-1 = CDI replicate 1, WC1-2 = CDI replicate 2, WC1-6 = CDI replicate 3, WC1-7 = CDI replicate 4. Pre-ABX = Pre-antibiotics, Post-ABX= post-antbiotics, CDI= C*lostridioides difficile* infection. Contig taxonomy and geNomad prophage classification in plot subtitle. Plots are listed alphabetically by family name.

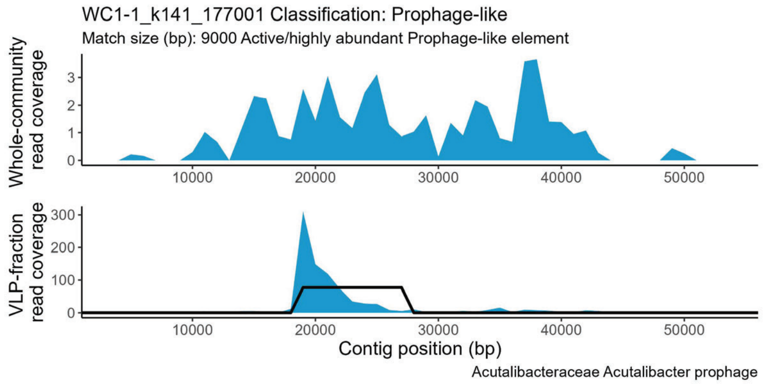


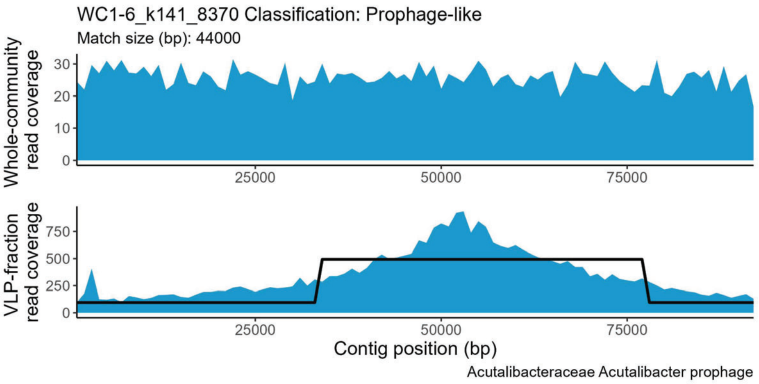


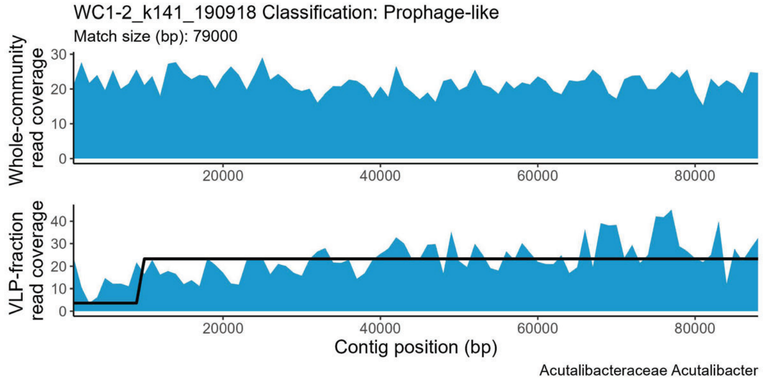


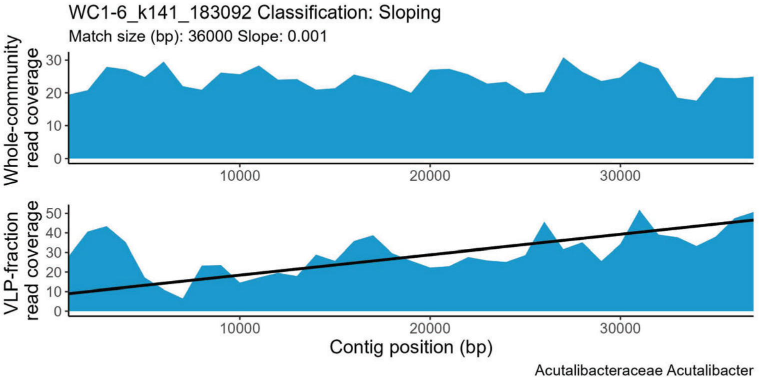


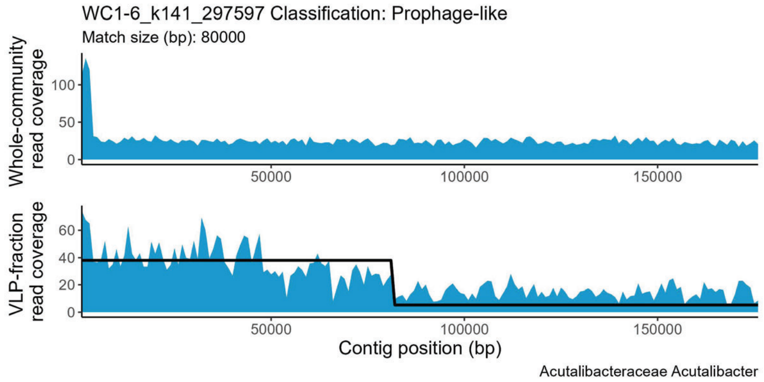


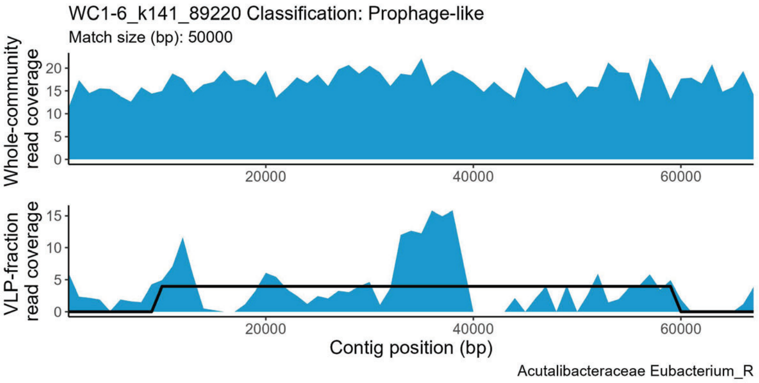


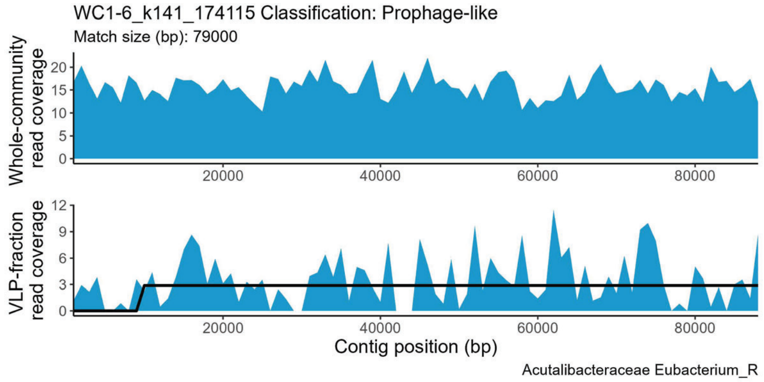


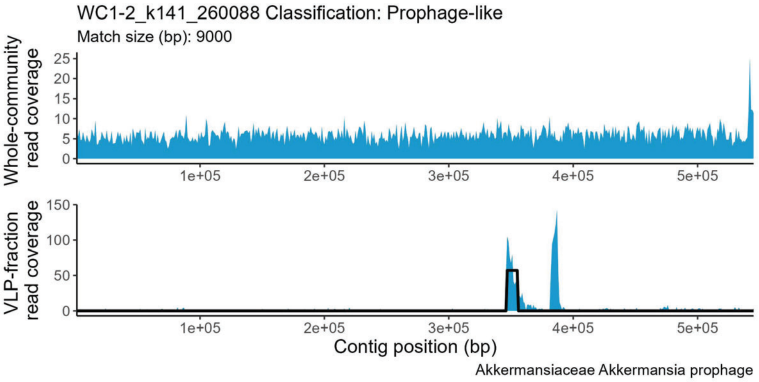

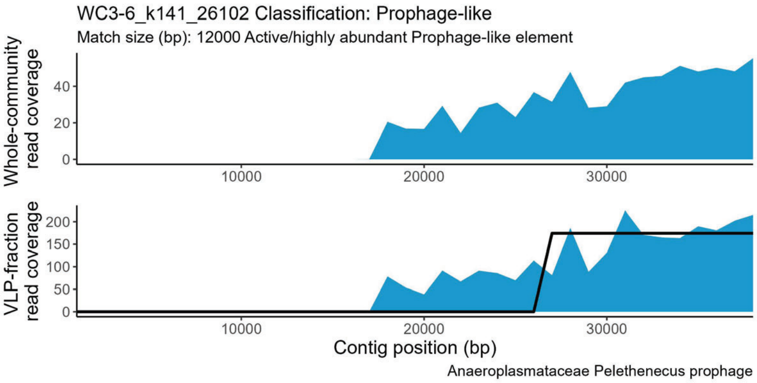
WC3-6_k141_26102 Classification: Prophage-like
Match size (bp): 12000 Active/highly abundant Prophage-like element
Whole-community read coverage
VLP-fraction read coverage
Contig position (bp)
Anaeroplasmataceae Pelethenecus prophage

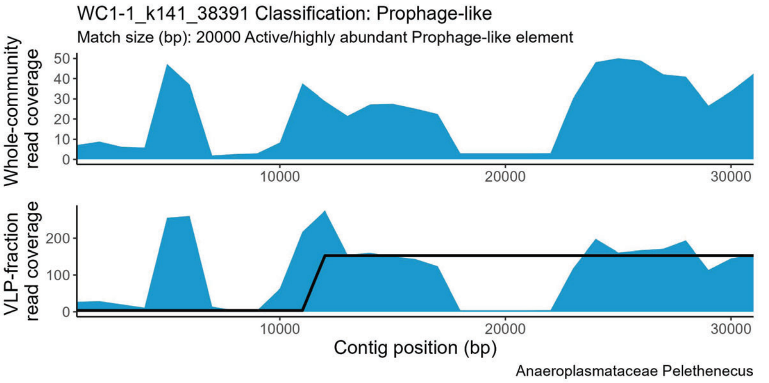
WC1-1_k141_38391 Classification: Prophage-like
Match size (bp): 20000 Active/highly abundant Prophage-like element
Whole-community read coverage
VLP-fraction read coverage
Contig position (bp)
Anaeroplasmataceae Pelethenecus

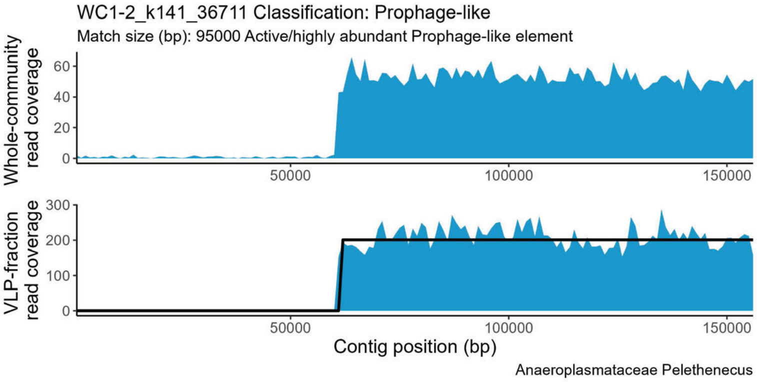
WC1-2_k141_36711 Classification: Prophage-like
Match size (bp): 95000 Active/highly abundant Prophage-like element
Whole-community read coverage
VLP-fraction read coverage
Contig position (bp)
Anaeroplasmataceae Pelethenecus

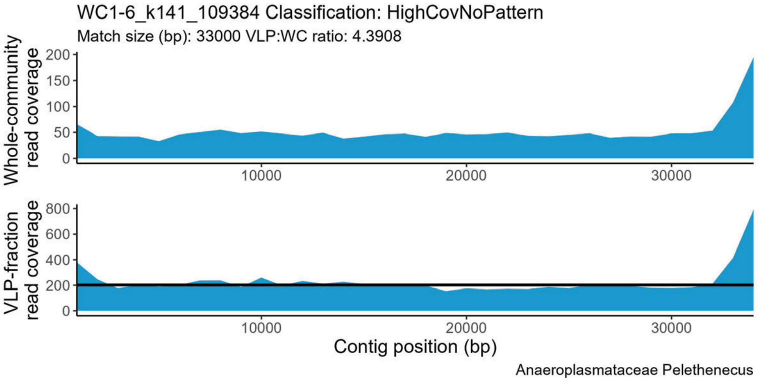
WC1-6_k141_109384 Classification: HighCovNoPattern
Match size (bp): 33000 VLP:WC ratio: 4.3908
Whole-community read coverage
VLP-fraction read coverage
Contig position (bp)
Anaeroplasmataceae Pelethenecus

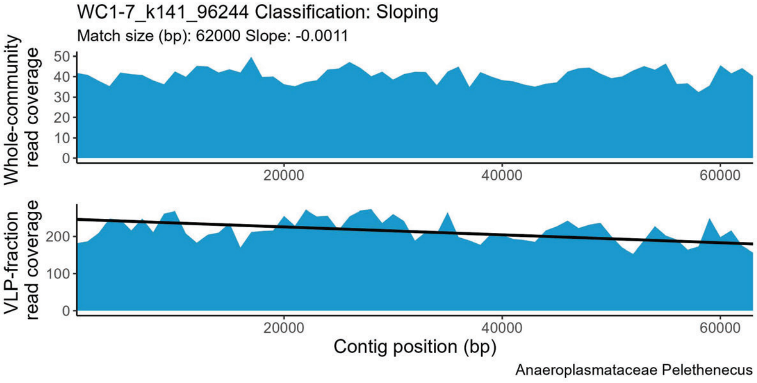
WC1-7_k141_96244 Classification: Sloping
Match size (bp): 62000 Slope: -0.0011
Whole-community read coverage
VLP-fraction read coverage
Contig position (bp)
Anaeroplasmataceae Pelethenecus

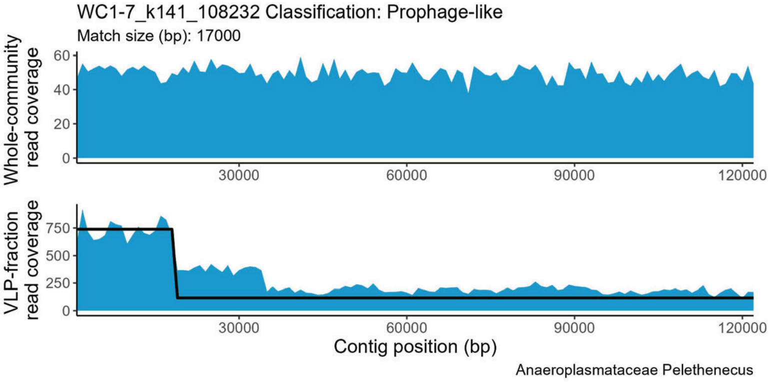
WC1-7_k141_108232 Classification: Prophage-like
Match size (bp): 17000
Whole-community read coverage
VLP-fraction read coverage
Contig position (bp)
Anaeroplasmataceae Pelethenecus

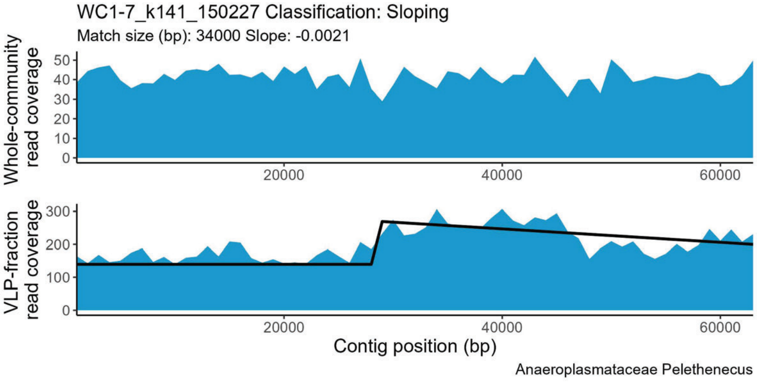
WC1-7_k141_150227 Classification: Sloping
Match size (bp): 34000 Slope: -0.0021
Whole-community read coverage
VLP-fraction read coverage
Contig position (bp)
Anaeroplasmataceae Pelethenecus

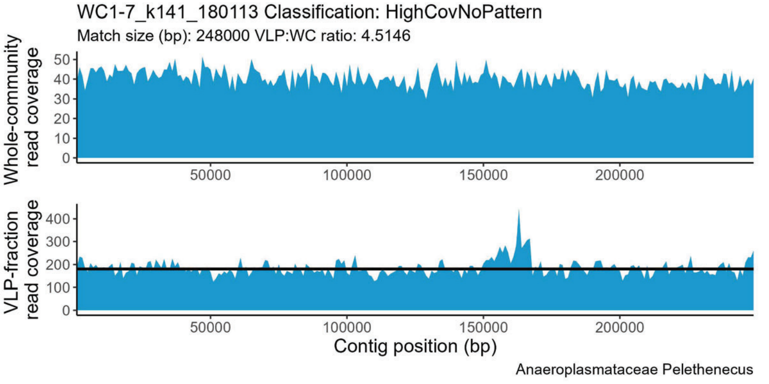
WC1-7_k141_180113 Classification: HighCovNoPattern
Match size (bp): 248000 VLP:WC ratio: 4.5146
Whole-community read coverage
VLP-fraction read coverage
Contig position (bp)
Anaeroplasmataceae Pelethenecus

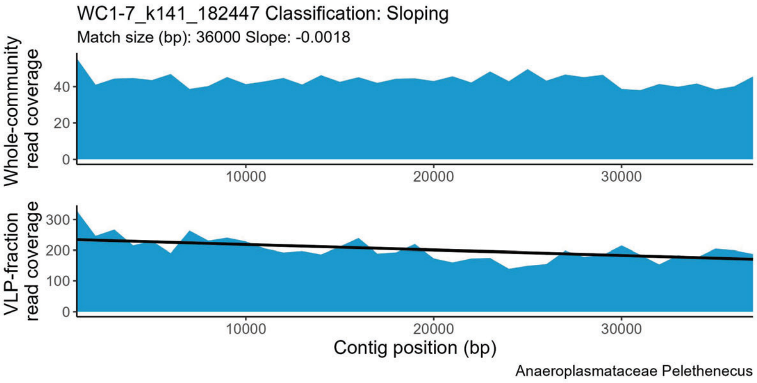
WC1-7_k141_182447 Classification: Sloping
Match size (bp): 36000 Slope: -0.0018
Whole-community read coverage
0
20
40
10000
20000
30000
VLP-fraction read coverage
0
100
200
300
10000
20000
30000
Contig position (bp)
Anaeroplasmataceae Pelethenecus

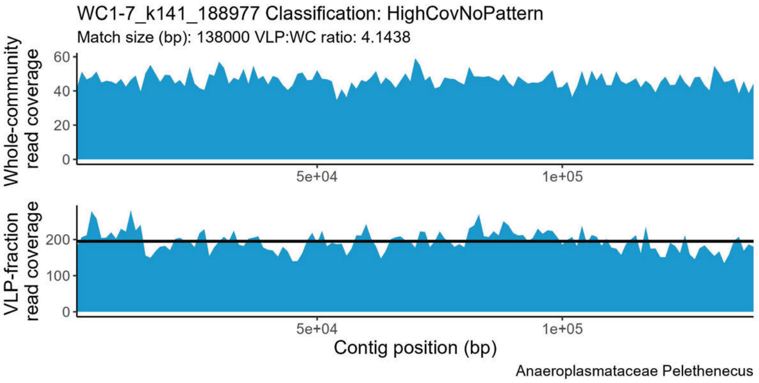
WC1-7_k141_188977 Classification: HighCovNoPattern
Match size (bp): 138000 VLP:WC ratio: 4.1438
Whole-community read coverage
0
20
40
60
5e+04
1e+05
VLP-fraction read coverage
0
100
200
5e+04
1e+05
Contig position (bp)
Anaeroplasmataceae Pelethenecus

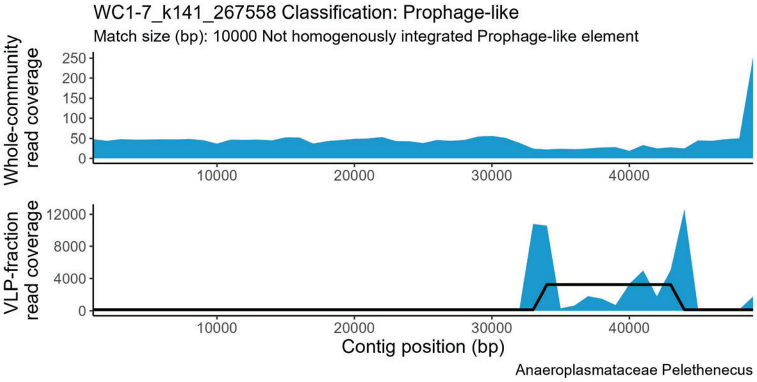
WC1-7_k141_267558 Classification: Prophage-like
Match size (bp): 10000 Not homogenously integrated Prophage-like element
Whole-community read coverage
0
50
100
150
200
250
10000
20000
30000
40000
VLP-fraction read coverage
0
4000
8000
12000
10000
20000
30000
40000
Contig position (bp)
Anaeroplasmataceae Pelethenecus

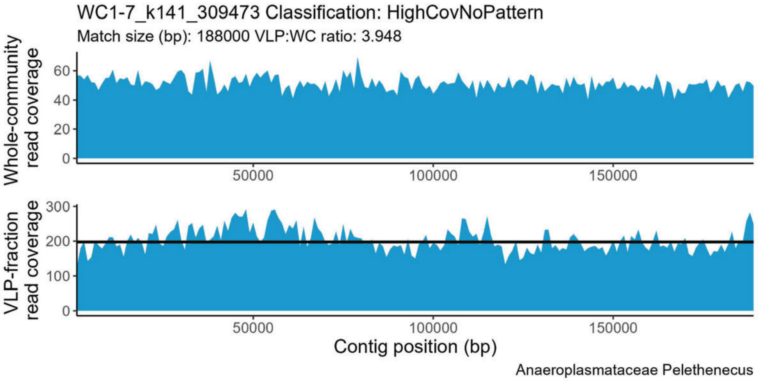
WC1-7_k141_309473 Classification: HighCovNoPattern
Match size (bp): 188000 VLP:WC ratio: 3.948
Whole-community read coverage
0
20
40
60
50000
100000
150000
VLP-fraction read coverage
0
100
200
300
50000
100000
150000
Contig position (bp)
Anaeroplasmataceae Pelethenecus

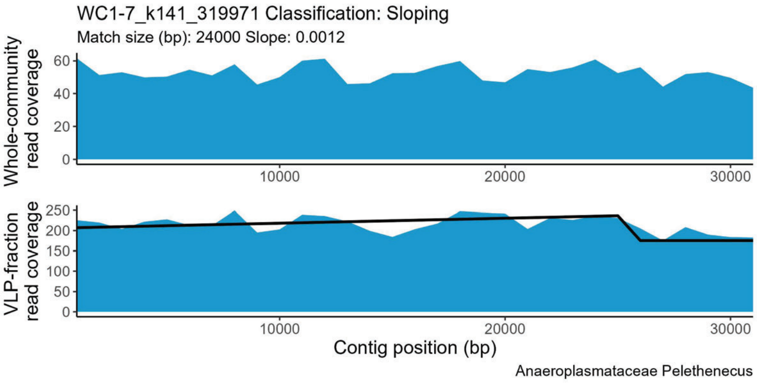
WC1-7_k141_319971 Classification: Sloping
Match size (bp): 24000 Slope: 0.0012
Whole-community read coverage
0
20
40
60
10000
20000
30000
VLP-fraction read coverage
0
50
100
150
200
250
10000
20000
30000
Contig position (bp)
Anaeroplasmataceae Pelethenecus

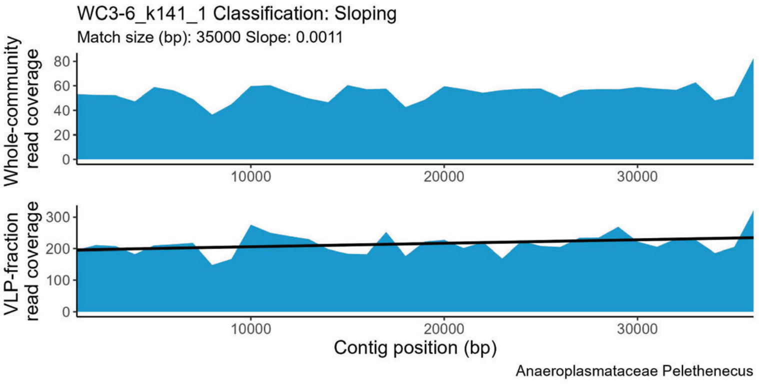
WC3-6_k141_1 Classification: Sloping
Match size (bp): 35000 Slope: 0.0011
Whole-community read coverage
0
20
40
60
80
10000
20000
30000
VLP-fraction read coverage
0
100
200
300
10000
20000
30000
Contig position (bp)
Anaeroplasmataceae Pelethenecus

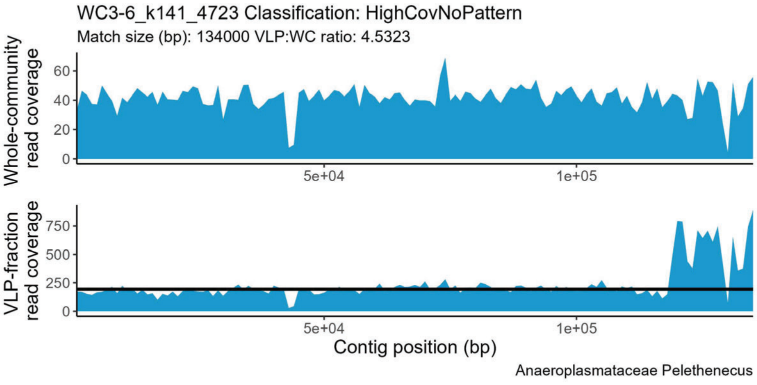
WC3-6_k141_4723 Classification: HighCovNoPattern
Match size (bp): 134000 VLP:WC ratio: 4.5323
Whole-community read coverage
0
20
40
60
5e+04
1e+05
VLP-fraction read coverage
0
250
500
750
5e+04
1e+05
Contig position (bp)
Anaeroplasmataceae Pelethenecus

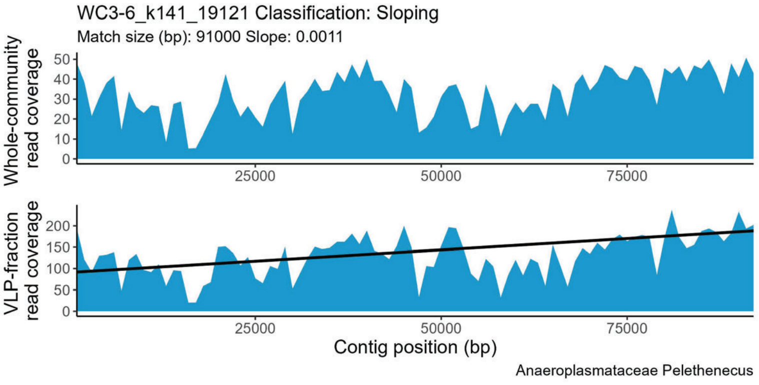
WC3-6_k141_19121 Classification: Sloping
Match size (bp): 91000 Slope: 0.0011
Whole-community read coverage
0
10
20
30
40
50
25000
50000
75000
VLP-fraction read coverage
0
50
100
150
200
25000
50000
75000
Contig position (bp)
Anaeroplasmataceae Pelethenecus

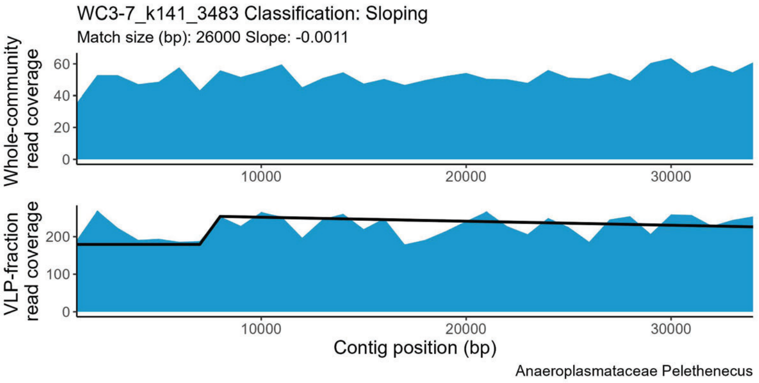
WC3-7_k141_3483 Classification: Sloping
Match size (bp): 26000 Slope: -0.0011
Whole-community read coverage
VLP-fraction read coverage
Contig position (bp)
Anaeroplasmataceae Pelethenecus

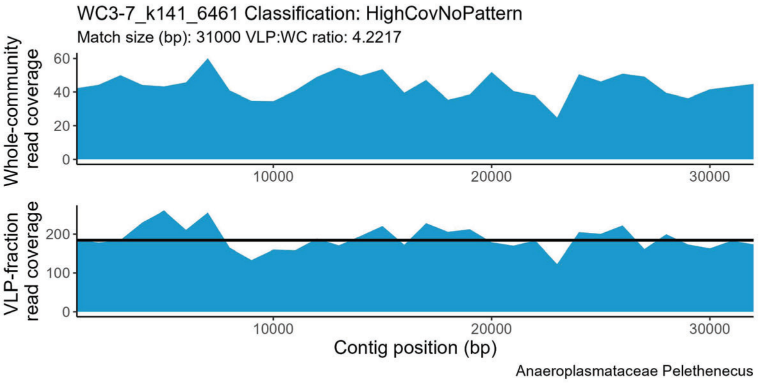
WC3-7_k141_6461 Classification: HighCovNoPattern
Match size (bp): 31000 VLP:WC ratio: 4.2217
Whole-community read coverage
VLP-fraction read coverage
Contig position (bp)
Anaeroplasmataceae Pelethenecus

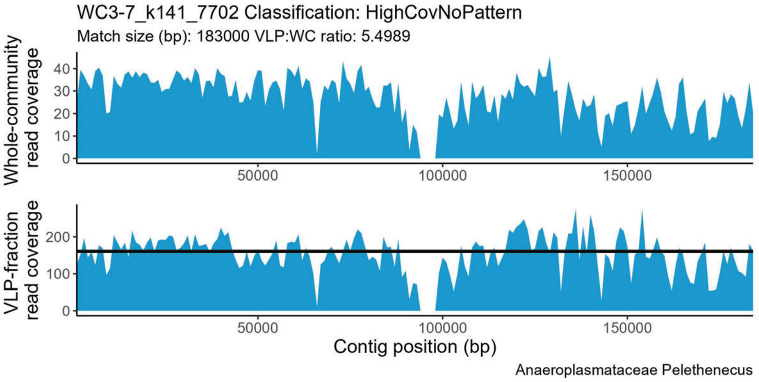
WC3-7_k141_7702 Classification: HighCovNoPattern
Match size (bp): 183000 VLP:WC ratio: 5.4989
Whole-community read coverage
VLP-fraction read coverage
Contig position (bp)
Anaeroplasmataceae Pelethenecus

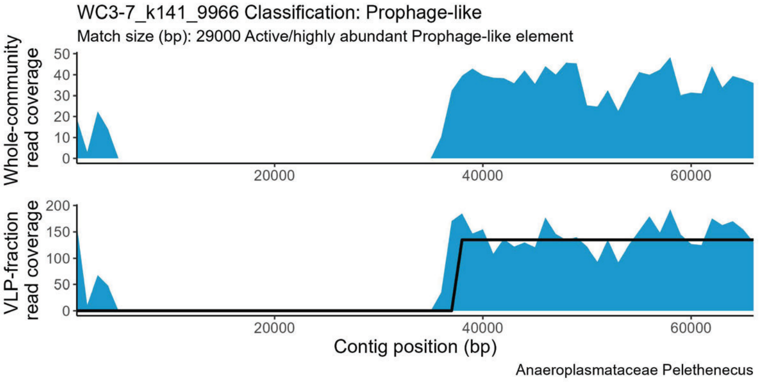
WC3-7_k141_9966 Classification: Prophage-like
Match size (bp): 29000 Active/highly abundant Prophage-like element
Whole-community read coverage
VLP-fraction read coverage
Contig position (bp)
Anaeroplasmataceae Pelethenecus

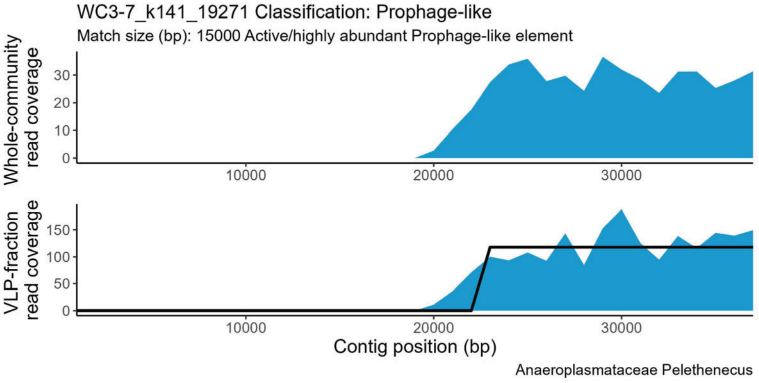
WC3-7_k141_19271 Classification: Prophage-like
Match size (bp): 15000 Active/highly abundant Prophage-like element
Whole-community read coverage
VLP-fraction read coverage
Contig position (bp)
Anaeroplasmataceae Pelethenecus

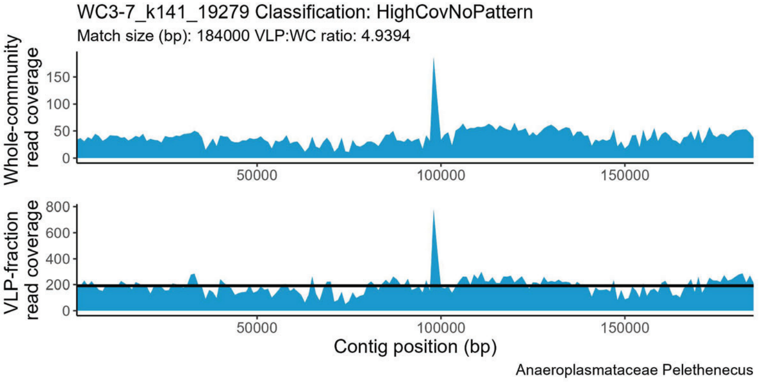
WC3-7_k141_19279 Classification: HighCovNoPattern
Match size (bp): 184000 VLP:WC ratio: 4.9394
Whole-community read coverage
VLP-fraction read coverage
Contig position (bp)
Anaeroplasmataceae Pelethenecus

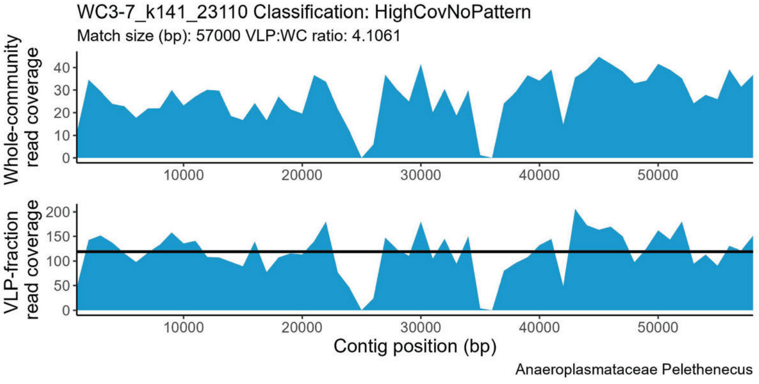
WC3-7_k141_23110 Classification: HighCovNoPattern
Match size (bp): 57000 VLP:WC ratio: 4.1061
Whole-community read coverage
VLP-fraction read coverage
Contig position (bp)
Anaeroplasmataceae Pelethenecus

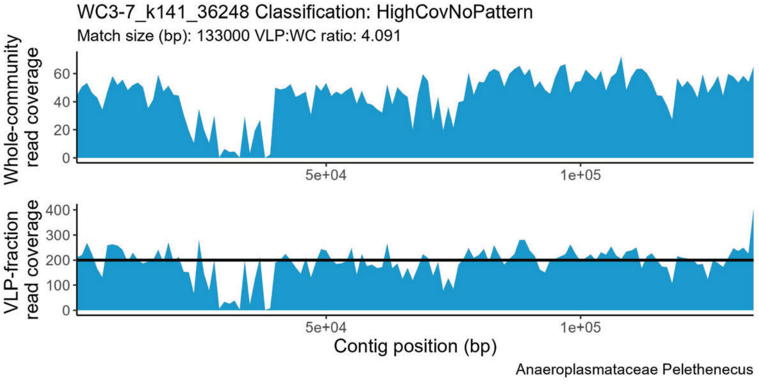
WC3-7_k141_36248 Classification: HighCovNoPattern
Match size (bp): 133000 VLP:WC ratio: 4.091
Whole-community read coverage
VLP-fraction read coverage
Contig position (bp)
Anaeroplasmataceae Pelethenecus

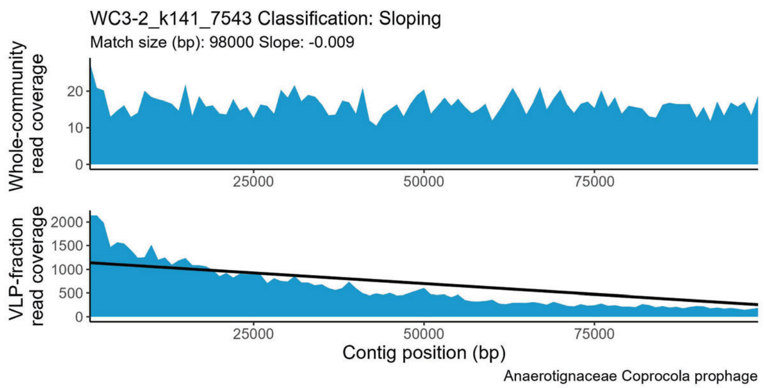
WC3-2_k141_7543 Classification: Sloping
Match size (bp): 98000 Slope: -0.009
Whole-community read coverage
VLP-fraction read coverage
Contig position (bp)
Anaerotignaceae Coprocola prophage

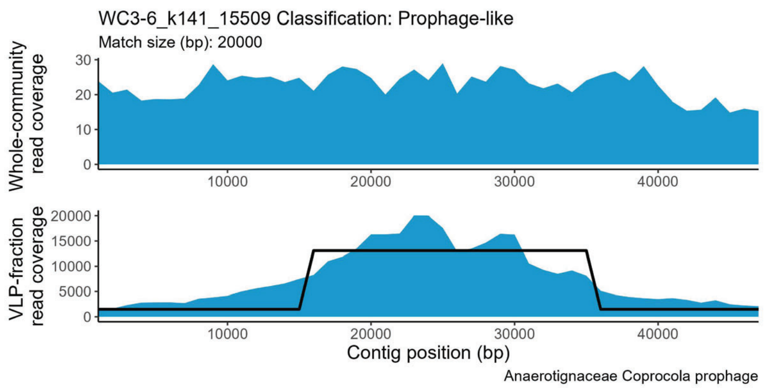
WC3-6_k141_15509 Classification: Prophage-like
Match size (bp): 20000
Whole-community read coverage
VLP-fraction read coverage
Contig position (bp)
Anaerotignaceae Coprocola prophage

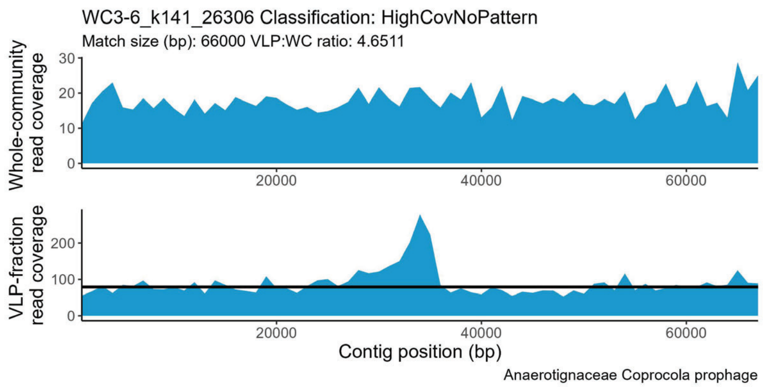
WC3-6_k141_26306 Classification: HighCovNoPattern
Match size (bp): 66000 VLP:WC ratio: 4.6511
Whole-community read coverage
VLP-fraction read coverage
Contig position (bp)
Anaerotignaceae Coprocola prophage

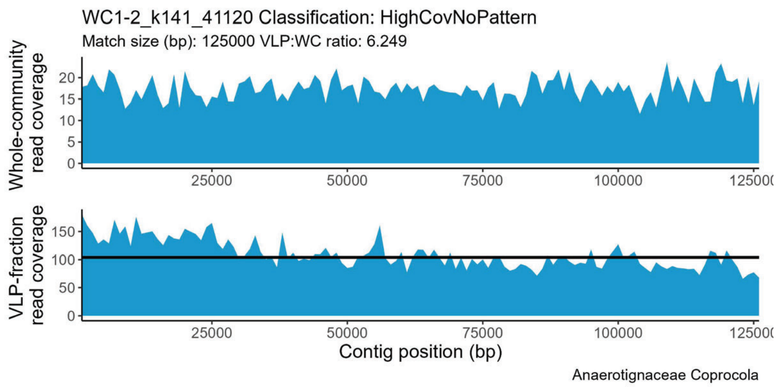
WC1-2_k141_41120 Classification: HighCovNoPattern
Match size (bp): 125000 VLP:WC ratio: 6.249
Whole-community read coverage
VLP-fraction read coverage
Contig position (bp)
Anaerotignaceae Coprocola

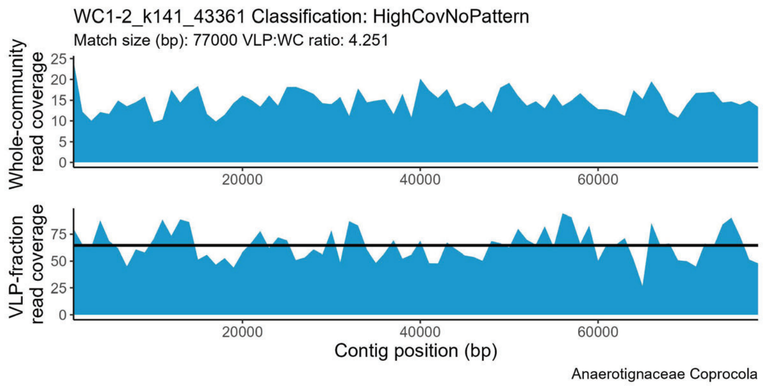
WC1-2_k141_43361 Classification: HighCovNoPattern
Match size (bp): 77000 VLP:WC ratio: 4.251
Whole-community read coverage
VLP-fraction read coverage
Contig position (bp)
Anaerotignaceae Coprocola

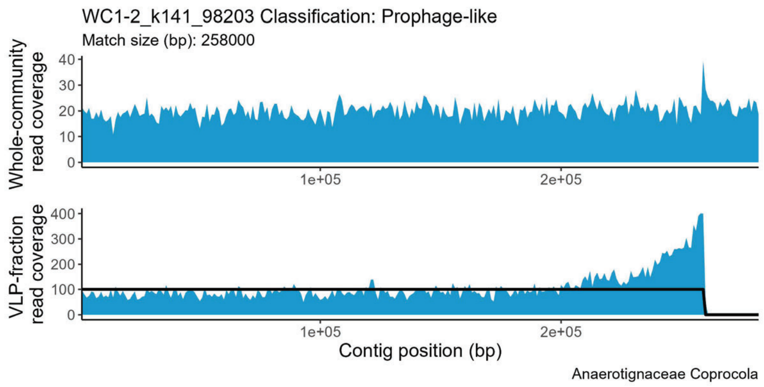
WC1-2_k141_98203 Classification: Prophage-like
Match size (bp): 258000
Whole-community read coverage
VLP-fraction read coverage
Contig position (bp)
Anaerotignaceae Coprocola

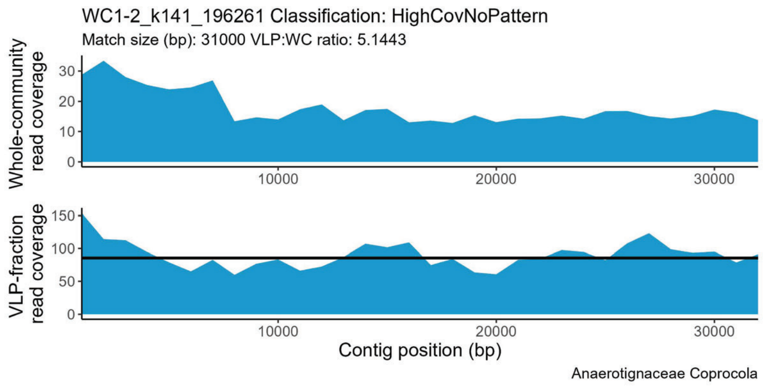
WC1-2_k141_196261 Classification: HighCovNoPattern
Match size (bp): 31000 VLP:WC ratio: 5.1443
Whole-community read coverage
VLP-fraction read coverage
Contig position (bp)
Anaerotignaceae Coprocola

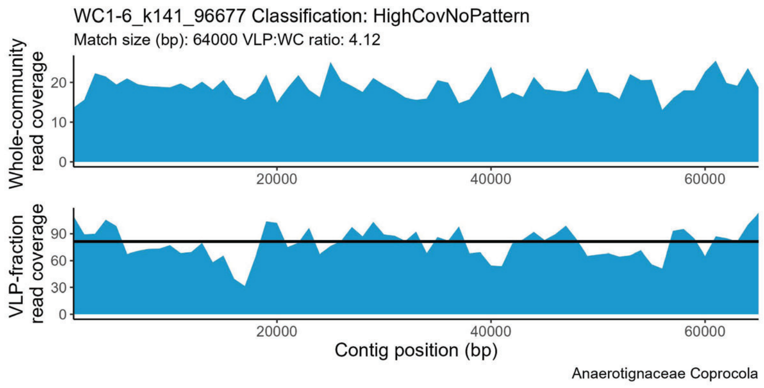
WC1-6_k141_96677 Classification: HighCovNoPattern
Match size (bp): 64000 VLP:WC ratio: 4.12
Whole-community read coverage
VLP-fraction read coverage
Contig position (bp)
Anaerotignaceae Coprocola

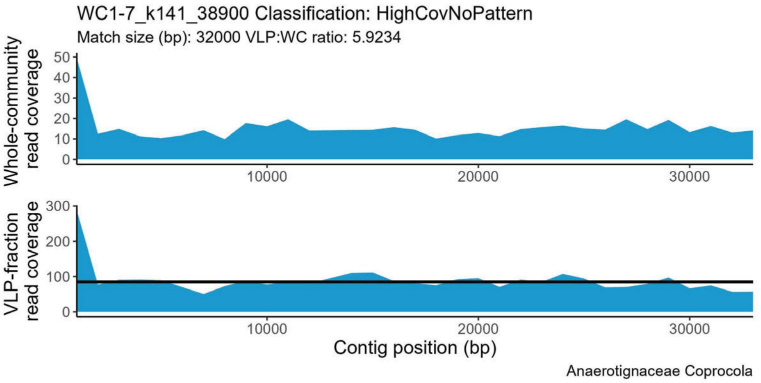
WC1-7_k141_38900 Classification: HighCovNoPattern
Match size (bp): 32000 VLP:WC ratio: 5.9234
Whole-community read coverage
VLP-fraction read coverage
Contig position (bp)
Anaerotignaceae Coprocola

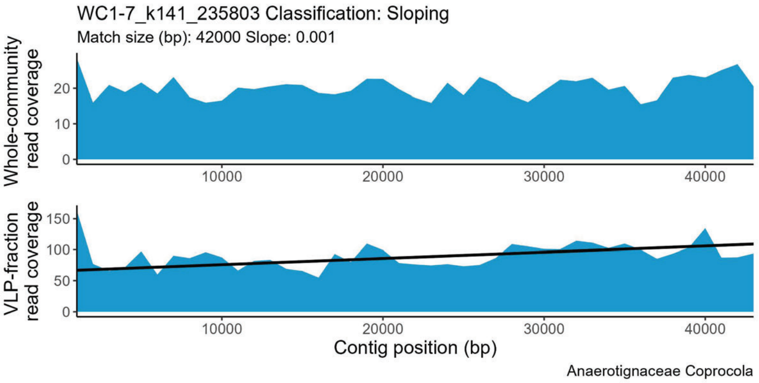
WC1-7_k141_235803 Classification: Sloping
Match size (bp): 42000 Slope: 0.001
Whole-community read coverage
VLP-fraction read coverage
Contig position (bp)
Anaerotignaceae Coprocola

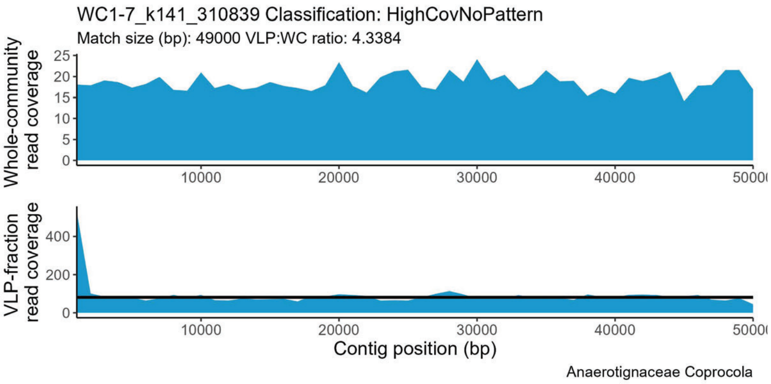
WC1-7_k141_310839 Classification: HighCovNoPattern
Match size (bp): 49000 VLP:WC ratio: 4.3384
Whole-community read coverage
VLP-fraction read coverage
Contig position (bp)
Anaerotignaceae Coprocola

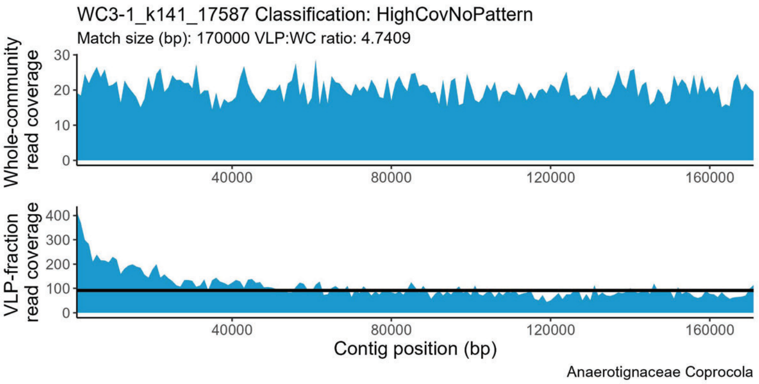
WC3-1_k141_17587 Classification: HighCovNoPattern
Match size (bp): 170000 VLP:WC ratio: 4.7409
Whole-community read coverage
VLP-fraction read coverage
Contig position (bp)
Anaerotignaceae Coprocola

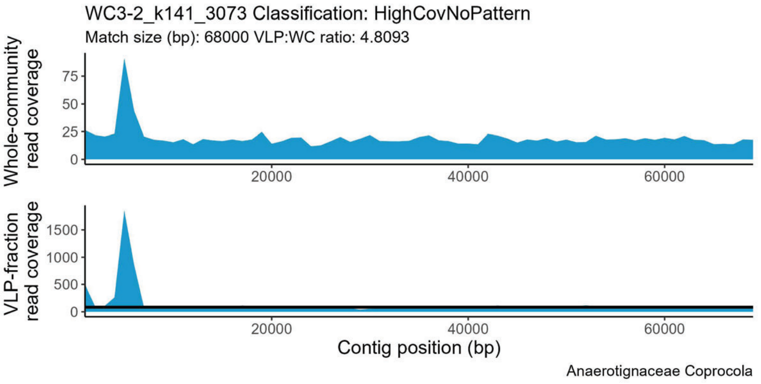
WC3-2_k141_3073 Classification: HighCovNoPattern
Match size (bp): 68000 VLP:WC ratio: 4.8093
Whole-community read coverage
VLP-fraction read coverage
Contig position (bp)
Anaerotignaceae Coprocola

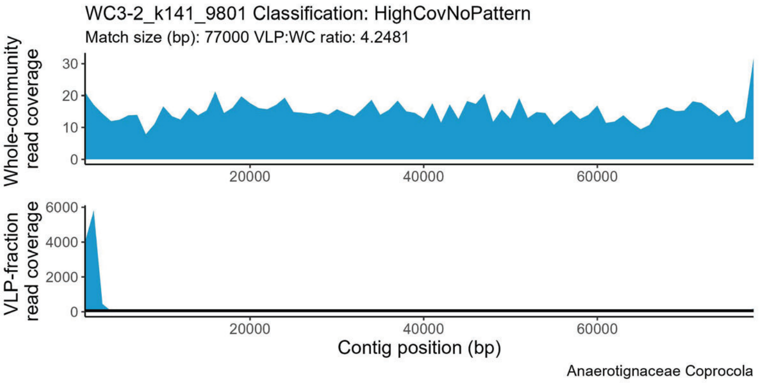
WC3-2_k141_9801 Classification: HighCovNoPattern
Match size (bp): 77000 VLP:WC ratio: 4.2481
Whole-community read coverage
VLP-fraction read coverage
Contig position (bp)
Anaerotignaceae Coprocola

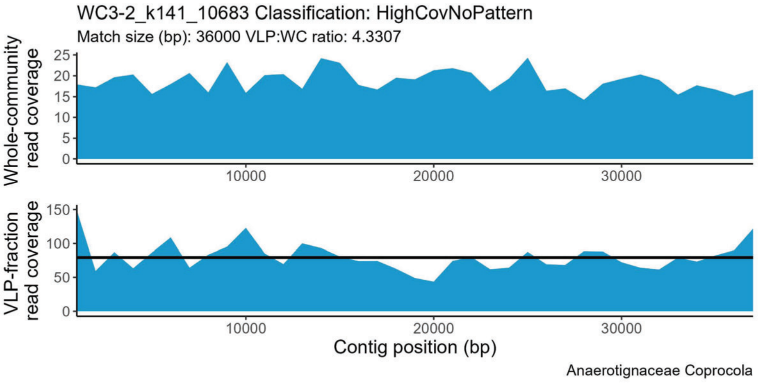
WC3-2_k141_10683 Classification: HighCovNoPattern
Match size (bp): 36000 VLP:WC ratio: 4.3307
Whole-community read coverage
VLP-fraction read coverage
Contig position (bp)
Anaerotignaceae Coprocola

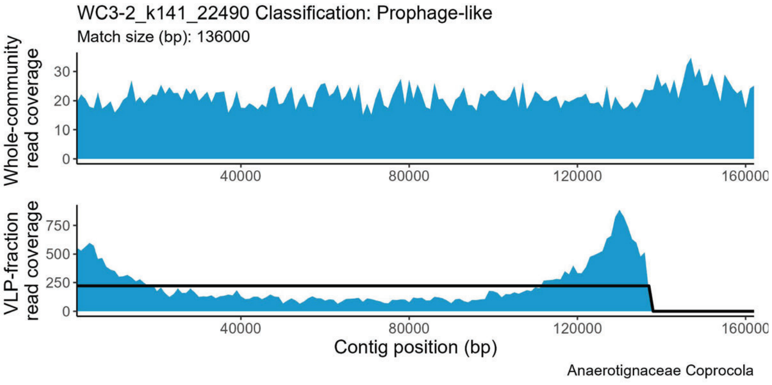
WC3-2_k141_22490 Classification: Prophage-like
Match size (bp): 136000
Whole-community read coverage
VLP-fraction read coverage
Contig position (bp)
Anaerotignaceae Coprocola

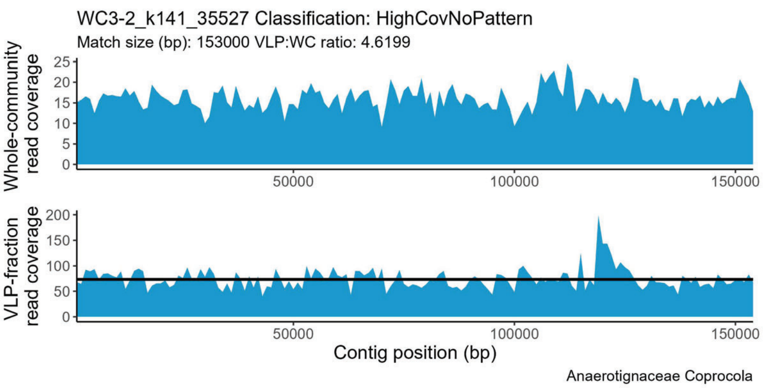
WC3-2_k141_35527 Classification: HighCovNoPattern
Match size (bp): 153000 VLP:WC ratio: 4.6199
Whole-community read coverage
0 5 10 15 20 25
50000 100000 150000
VLP-fraction read coverage
0 50 100 150 200
50000 100000 150000
Contig position (bp)
Anaerotignaceae Coprocola

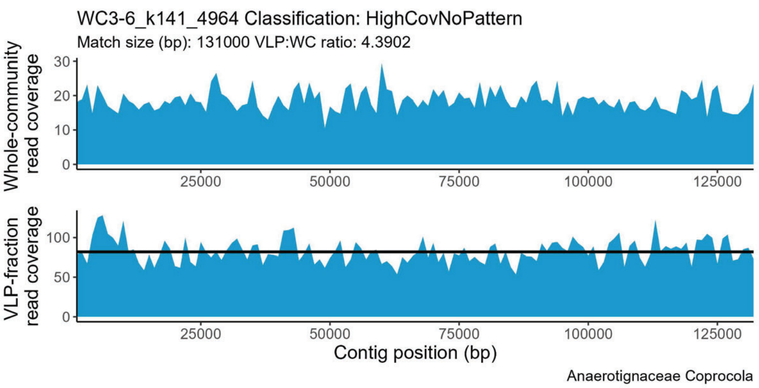
WC3-6_k141_4964 Classification: HighCovNoPattern
Match size (bp): 131000 VLP:WC ratio: 4.3902
Whole-community read coverage
0 10 20 30
25000 50000 75000 100000 125000
VLP-fraction read coverage
0 50 100
25000 50000 75000 100000 125000
Contig position (bp)
Anaerotignaceae Coprocola

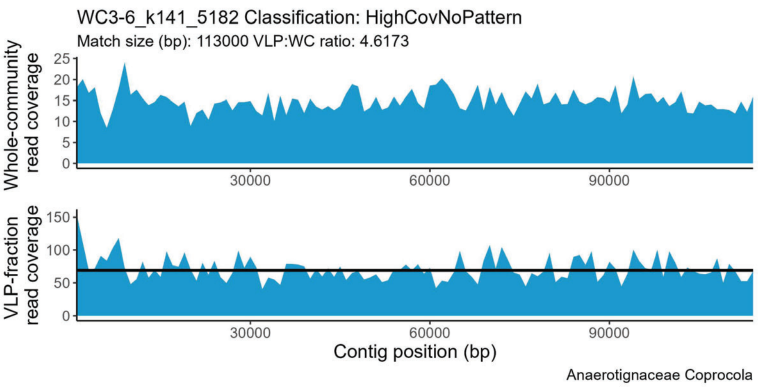
WC3-6_k141_5182 Classification: HighCovNoPattern
Match size (bp): 113000 VLP:WC ratio: 4.6173
Whole-community read coverage
0 5 10 15 20 25
30000 60000 90000
VLP-fraction read coverage
0 50 100 150
30000 60000 90000
Contig position (bp)
Anaerotignaceae Coprocola

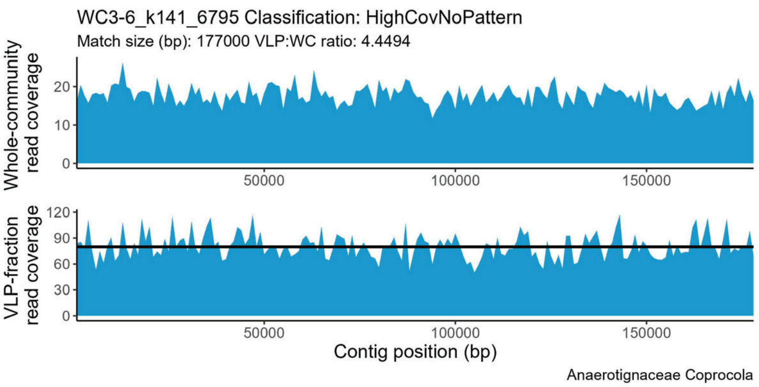
WC3-6_k141_6795 Classification: HighCovNoPattern
Match size (bp): 177000 VLP:WC ratio: 4.4494
Whole-community read coverage
0 10 20
50000 100000 150000
VLP-fraction read coverage
0 30 60 90 120
50000 100000 150000
Contig position (bp)
Anaerotignaceae Coprocola

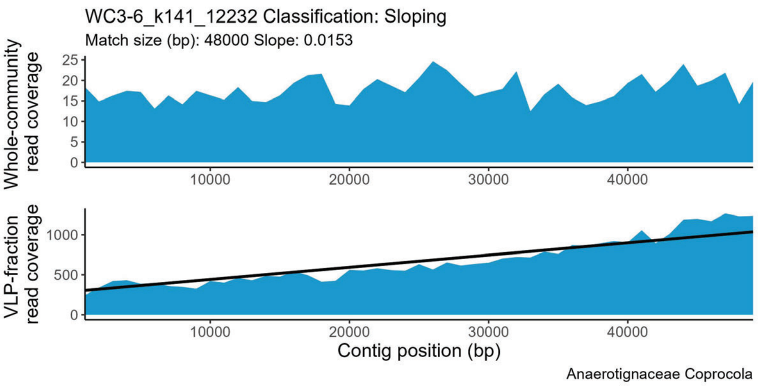
WC3-6_k141_12232 Classification: Sloping
Match size (bp): 48000 Slope: 0.0153
Whole-community read coverage
0 5 10 15 20 25
10000 20000 30000 40000
VLP-fraction read coverage
0 500 1000
10000 20000 30000 40000
Contig position (bp)
Anaerotignaceae Coprocola

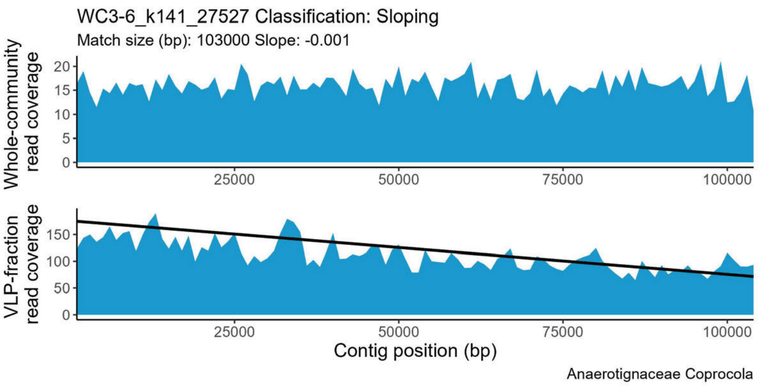
WC3-6_k141_27527 Classification: Sloping
Match size (bp): 103000 Slope: -0.001
Whole-community read coverage
0 5 10 15 20
25000 50000 75000 100000
VLP-fraction read coverage
0 50 100 150
25000 50000 75000 100000
Contig position (bp)
Anaerotignaceae Coprocola

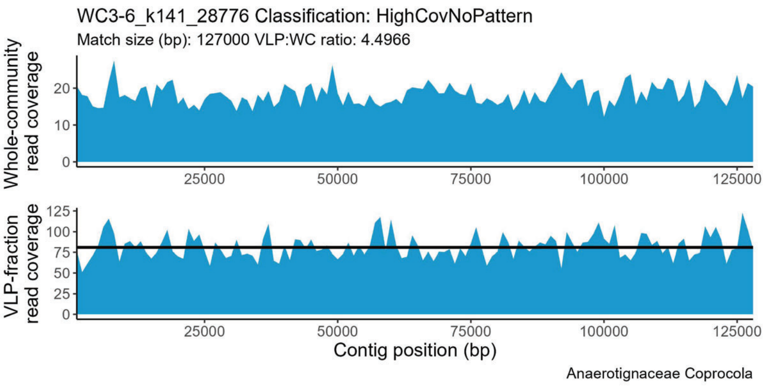
WC3-6_k141_28776 Classification: HighCovNoPattern
Match size (bp): 127000 VLP:WC ratio: 4.4966
Whole-community read coverage
0 10 20
25000 50000 75000 100000 125000
VLP-fraction read coverage
0 25 50 75 100 125
25000 50000 75000 100000 125000
Contig position (bp)
Anaerotignaceae Coprocola

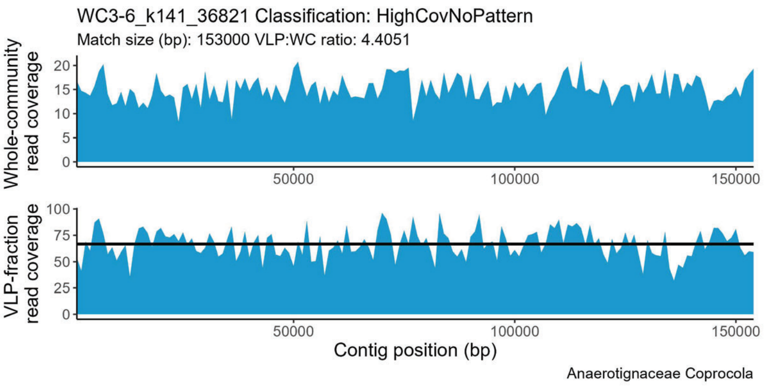
WC3-6_k141_36821 Classification: HighCovNoPattern
Match size (bp): 153000 VLP:WC ratio: 4.4051
Whole-community read coverage
0 5 10 15 20
50000 100000 150000
VLP-fraction read coverage
0 25 50 75 100
50000 100000 150000
Contig position (bp)
Anaerotignaceae Coprocola

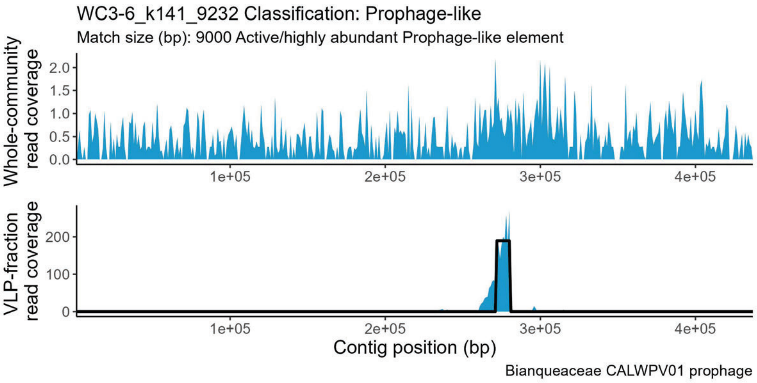
WC3-6_k141_9232 Classification: Prophage-like
Match size (bp): 9000 Active/highly abundant Prophage-like element
Whole-community read coverage
0.0
0.5
1.0
1.5
2.0
1e+05
2e+05
3e+05
4e+05
VLP-fraction read coverage
0
100
200
Contig position (bp)
Bianqueaceae CALWPV01 prophage

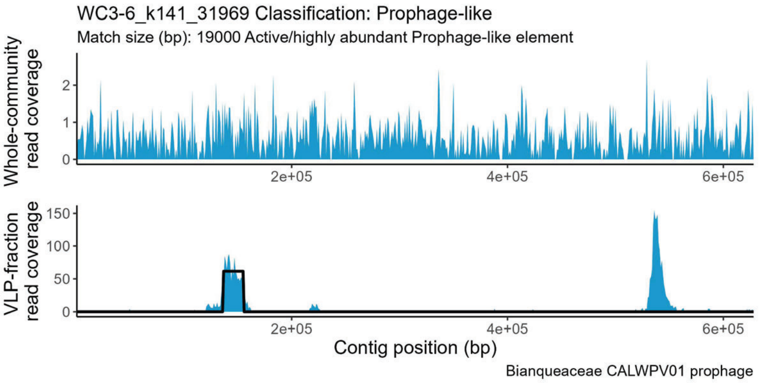
WC3-6_k141_31969 Classification: Prophage-like
Match size (bp): 19000 Active/highly abundant Prophage-like element
Whole-community read coverage
0
1
2
2e+05
4e+05
6e+05
VLP-fraction read coverage
0
50
100
150
Contig position (bp)
Bianqueaceae CALWPV01 prophage

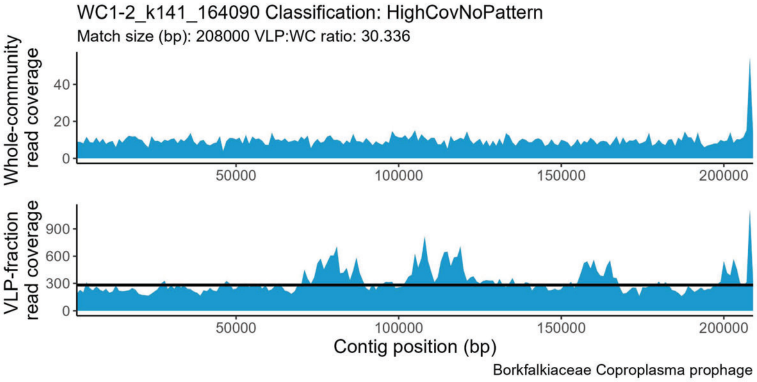
WC1-2_k141_164090 Classification: HighCovNoPattern
Match size (bp): 208000 VLP:WC ratio: 30.336
Whole-community read coverage
0
20
40
50000
100000
150000
200000
VLP-fraction read coverage
0
300
600
900
Contig position (bp)
Borkfalkiaceae Coproplasma prophage

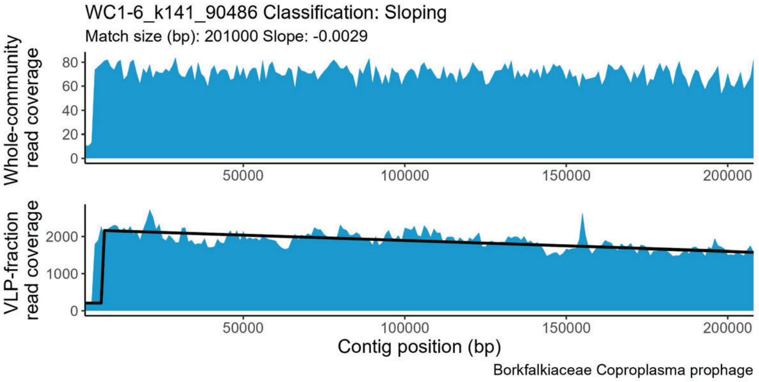
WC1-6_k141_90486 Classification: Sloping
Match size (bp): 201000 Slope: -0.0029
Whole-community read coverage
0
20
40
60
80
50000
100000
150000
200000
VLP-fraction read coverage
0
1000
2000
Contig position (bp)
Borkfalkiaceae Coproplasma prophage

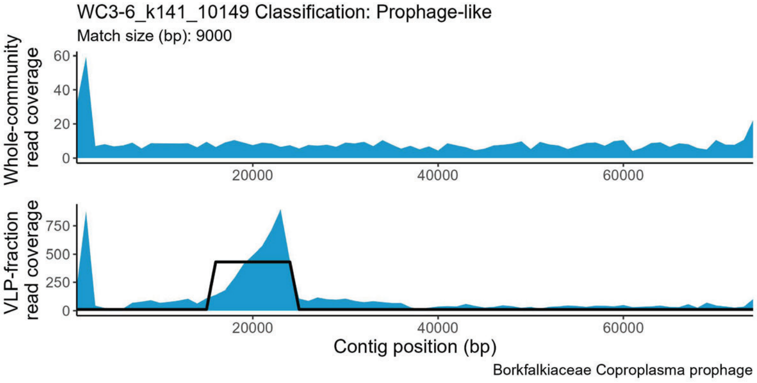
WC3-6_k141_10149 Classification: Prophage-like
Match size (bp): 9000
Whole-community read coverage
0
20
40
60
20000
40000
60000
VLP-fraction read coverage
0
250
500
750
Contig position (bp)
Borkfalkiaceae Coproplasma prophage

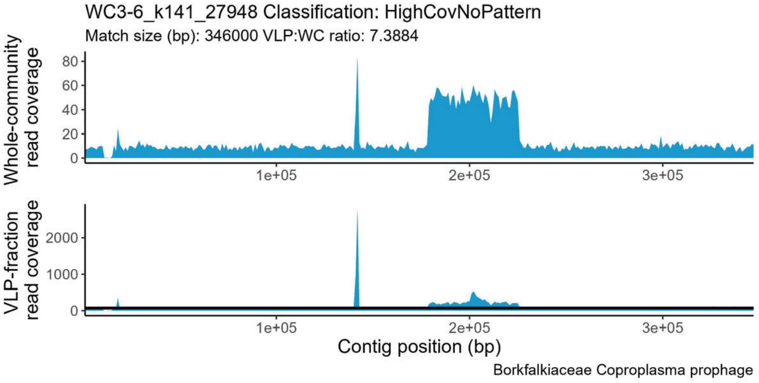
WC3-6_k141_27948 Classification: HighCovNoPattern
Match size (bp): 346000 VLP:WC ratio: 7.3884
Whole-community read coverage
0
20
40
60
80
1e+05
2e+05
3e+05
VLP-fraction read coverage
0
1000
2000
Contig position (bp)
Borkfalkiaceae Coproplasma prophage

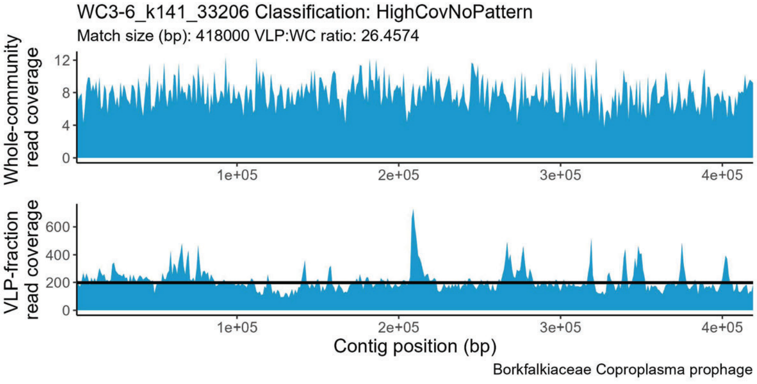
WC3-6_k141_33206 Classification: HighCovNoPattern
Match size (bp): 418000 VLP:WC ratio: 26.4574
Whole-community read coverage
0
4
8
12
1e+05
2e+05
3e+05
4e+05
VLP-fraction read coverage
0
200
400
600
Contig position (bp)
Borkfalkiaceae Coproplasma prophage

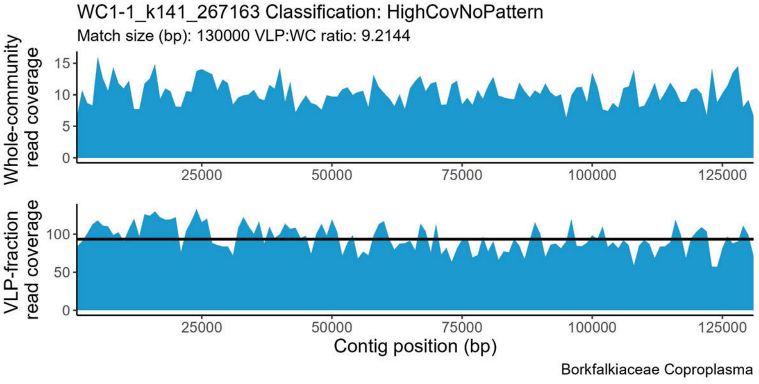
WC1-1_k141_267163 Classification: HighCovNoPattern
Match size (bp): 130000 VLP:WC ratio: 9.2144
Whole-community read coverage
0
5
10
15
25000
50000
75000
100000
125000
VLP-fraction read coverage
0
50
100
Contig position (bp)
Borkfalkiaceae Coproplasma

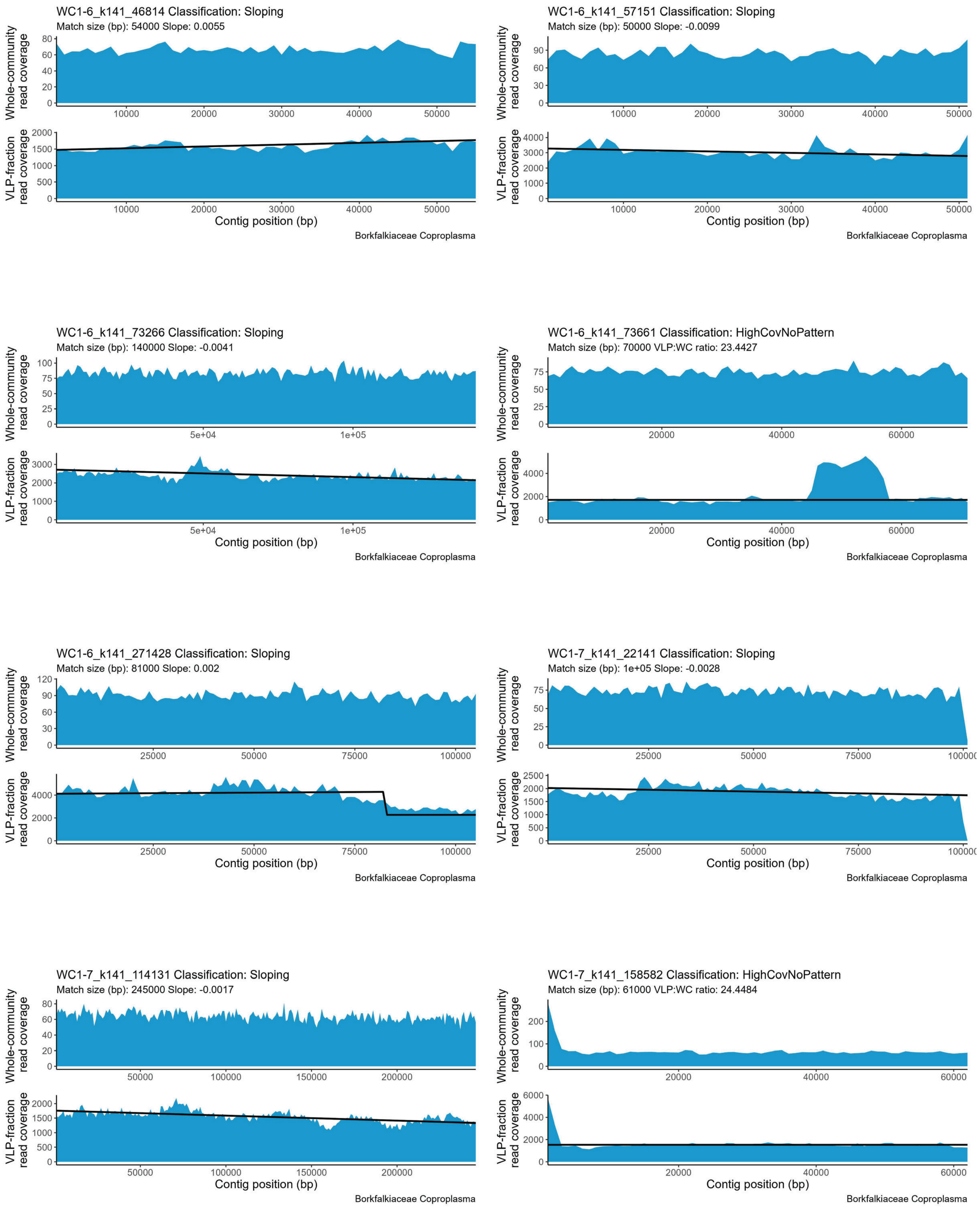
WC1-6_k141_46814 Classification: Sloping
Match size (bp): 54000 Slope: 0.0055
Whole-community read coverage
VLP-fraction read coverage
Contig position (bp)
Borkfalkiaceae Coproplasma
WC1-6_k141_57151 Classification: Sloping
Match size (bp): 50000 Slope: -0.0099
Whole-community read coverage
VLP-fraction read coverage
Contig position (bp)
Borkfalkiaceae Coproplasma
WC1-6_k141_73266 Classification: Sloping
Match size (bp): 140000 Slope: -0.0041
Whole-community read coverage
VLP-fraction read coverage
Contig position (bp)
Borkfalkiaceae Coproplasma
WC1-6_k141_73661 Classification: HighCovNoPattern
Match size (bp): 70000 VLP:WC ratio: 23.4427
Whole-community read coverage
VLP-fraction read coverage
Contig position (bp)
Borkfalkiaceae Coproplasma
WC1-6_k141_271428 Classification: Sloping
Match size (bp): 81000 Slope: 0.002
Whole-community read coverage
VLP-fraction read coverage
Contig position (bp)
Borkfalkiaceae Coproplasma
WC1-7_k141_22141 Classification: Sloping
Match size (bp): 1e+05 Slope: -0.0028
Whole-community read coverage
VLP-fraction read coverage
Contig position (bp)
Borkfalkiaceae Coproplasma
WC1-7_k141_114131 Classification: Sloping
Match size (bp): 245000 Slope: -0.0017
Whole-community read coverage
VLP-fraction read coverage
Contig position (bp)
Borkfalkiaceae Coproplasma
WC1-7_k141_158582 Classification: HighCovNoPattern
Match size (bp): 61000 VLP:WC ratio: 24.4484
Whole-community read coverage
VLP-fraction read coverage
Contig position (bp)
Borkfalkiaceae Coproplasma

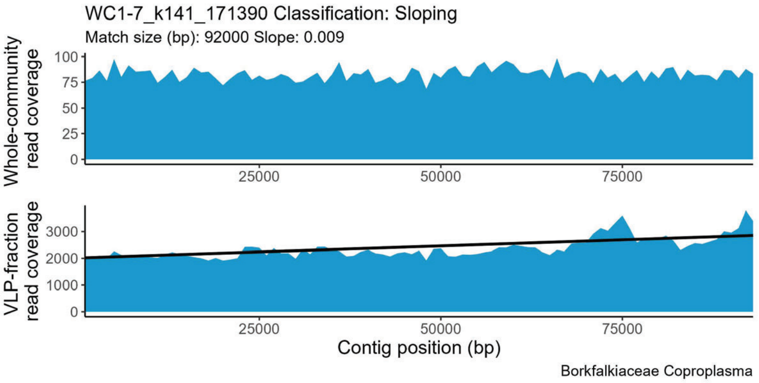
WC1-7_k141_171390 Classification: Sloping
Match size (bp): 92000 Slope: 0.009
Whole-community read coverage
VLP-fraction read coverage
Contig position (bp)
Borkfalkiaceae Coproplasma

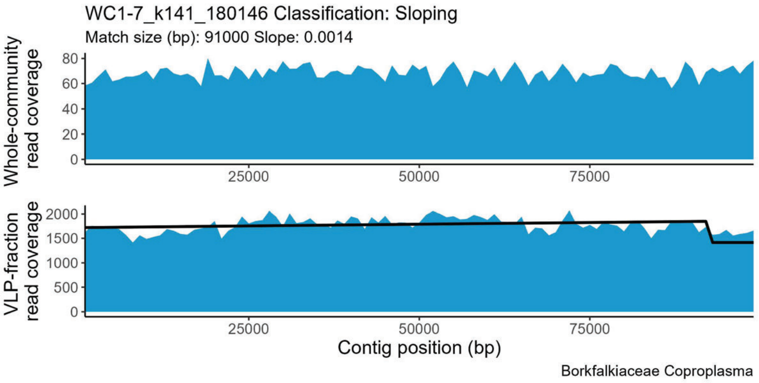
WC1-7_k141_180146 Classification: Sloping
Match size (bp): 91000 Slope: 0.0014
Whole-community read coverage
VLP-fraction read coverage
Contig position (bp)
Borkfalkiaceae Coproplasma

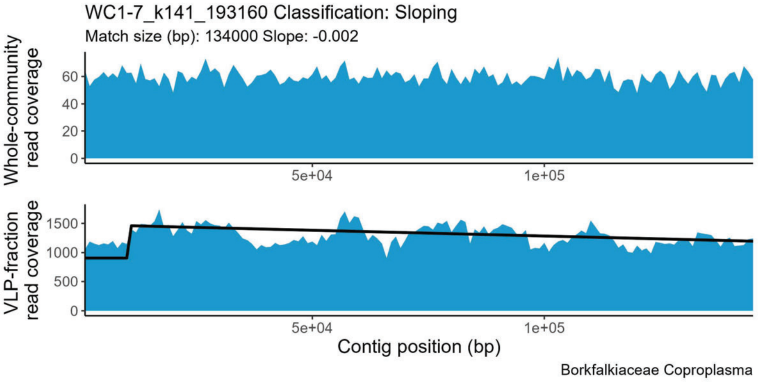
WC1-7_k141_193160 Classification: Sloping
Match size (bp): 134000 Slope: -0.002
Whole-community read coverage
VLP-fraction read coverage
Contig position (bp)
Borkfalkiaceae Coproplasma

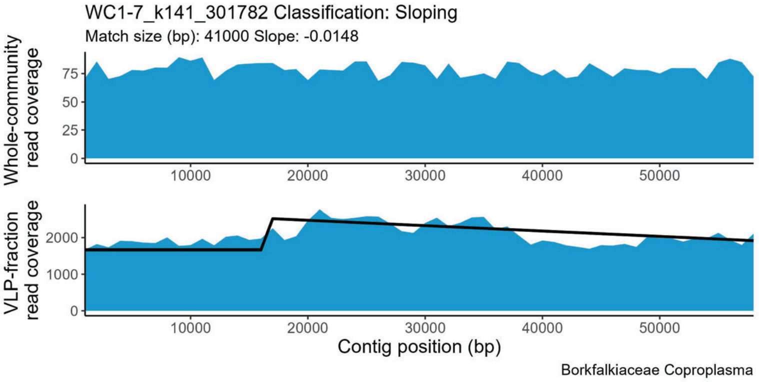
WC1-7_k141_301782 Classification: Sloping
Match size (bp): 41000 Slope: -0.0148
Whole-community read coverage
VLP-fraction read coverage
Contig position (bp)
Borkfalkiaceae Coproplasma

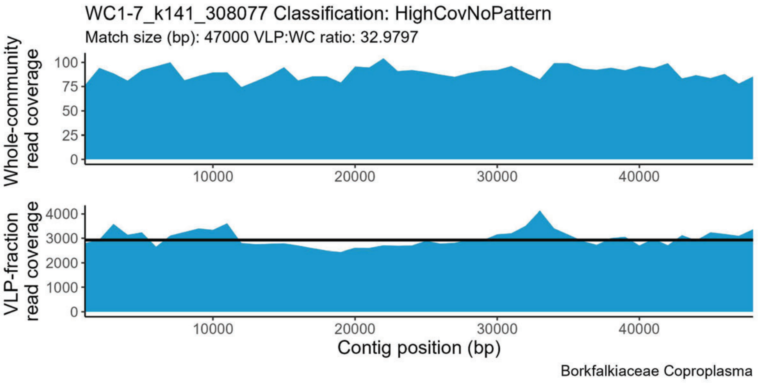
WC1-7_k141_308077 Classification: HighCovNoPattern
Match size (bp): 47000 VLP:WC ratio: 32.9797
Whole-community read coverage
VLP-fraction read coverage
Contig position (bp)
Borkfalkiaceae Coproplasma

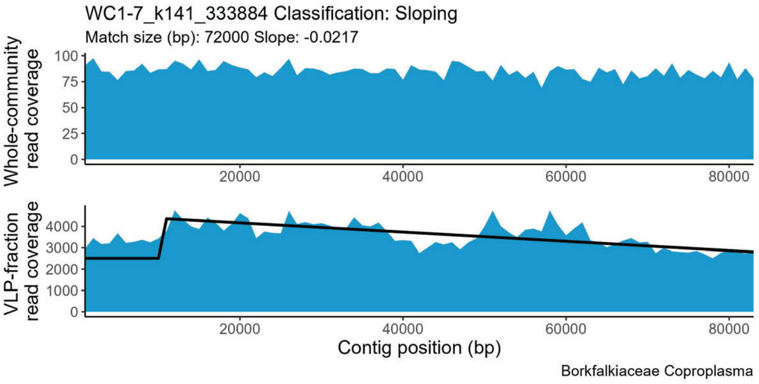
WC1-7_k141_333884 Classification: Sloping
Match size (bp): 72000 Slope: -0.0217
Whole-community read coverage
VLP-fraction read coverage
Contig position (bp)
Borkfalkiaceae Coproplasma

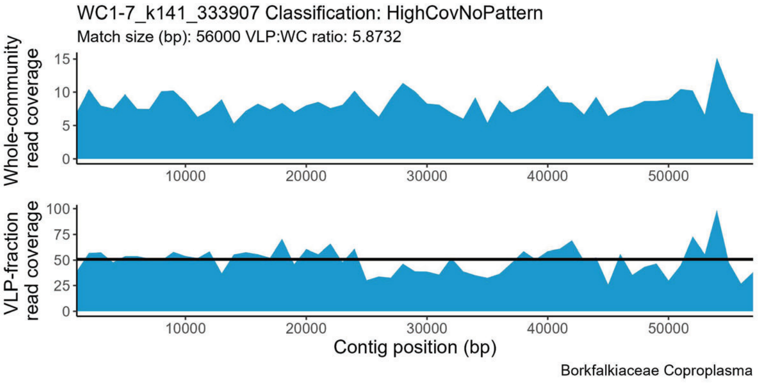
WC1-7_k141_333907 Classification: HighCovNoPattern
Match size (bp): 56000 VLP:WC ratio: 5.8732
Whole-community read coverage
VLP-fraction read coverage
Contig position (bp)
Borkfalkiaceae Coproplasma

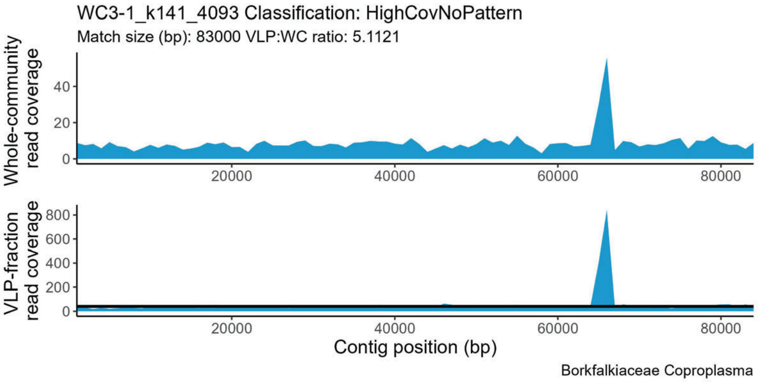
WC3-1_k141_4093 Classification: HighCovNoPattern
Match size (bp): 83000 VLP:WC ratio: 5.1121
Whole-community read coverage
VLP-fraction read coverage
Contig position (bp)
Borkfalkiaceae Coproplasma

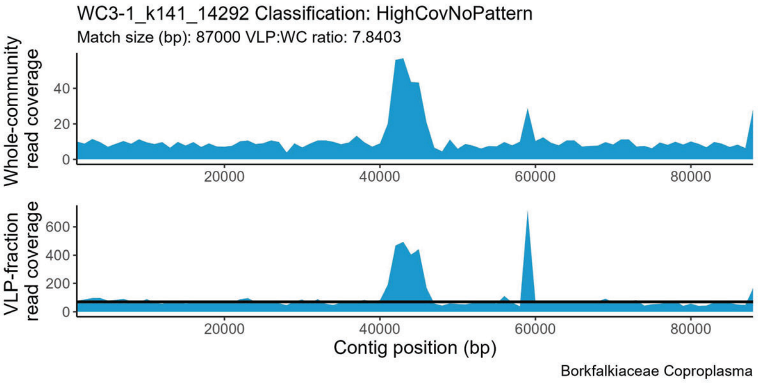
WC3-1_k141_14292 Classification: HighCovNoPattern
Match size (bp): 87000 VLP:WC ratio: 7.8403
Whole-community read coverage
VLP-fraction read coverage
Contig position (bp)
Borkfalkiaceae Coproplasma

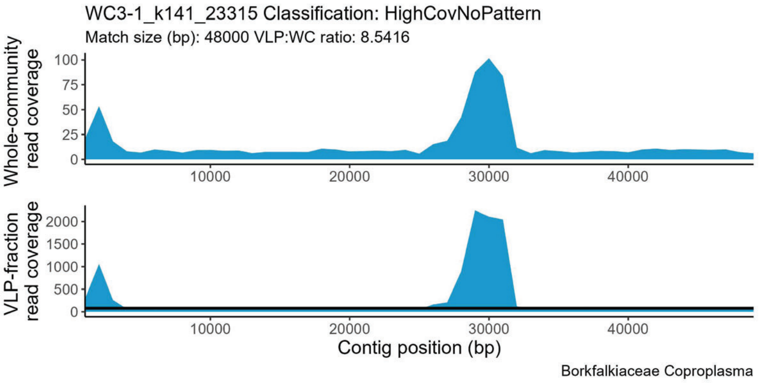
WC3-1_k141_23315 Classification: HighCovNoPattern
Match size (bp): 48000 VLP:WC ratio: 8.5416
Whole-community read coverage
VLP-fraction read coverage
Contig position (bp)
Borkfalkiaceae Coproplasma

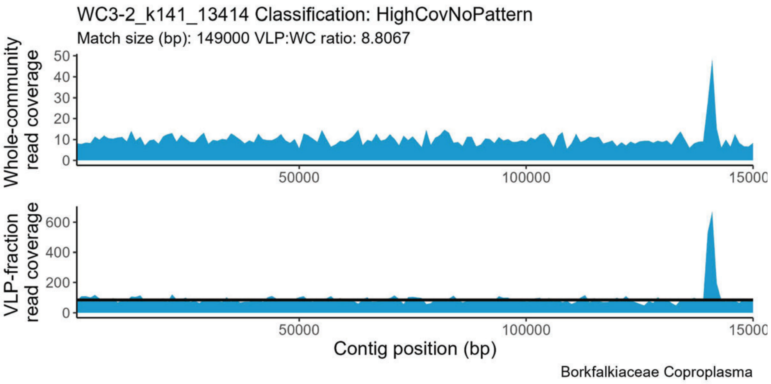
WC3-2_k141_13414 Classification: HighCovNoPattern
Match size (bp): 149000 VLP:WC ratio: 8.8067
Whole-community read coverage
VLP-fraction read coverage
Contig position (bp)
Borkfalkiaceae Coproplasma

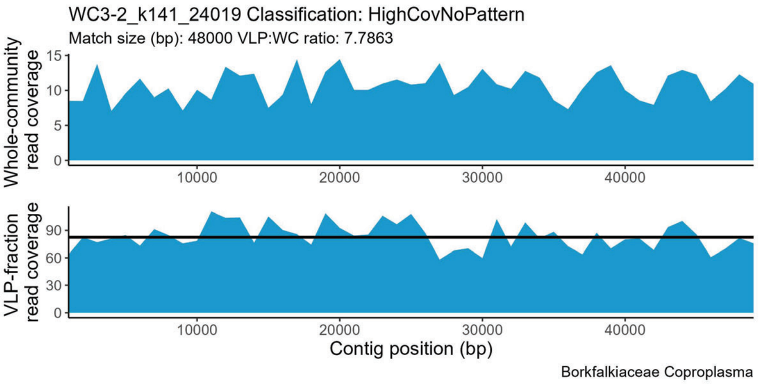
WC3-2_k141_24019 Classification: HighCovNoPattern
Match size (bp): 48000 VLP:WC ratio: 7.7863
Whole-community read coverage
VLP-fraction read coverage
Contig position (bp)
Borkfalkiaceae Coproplasma

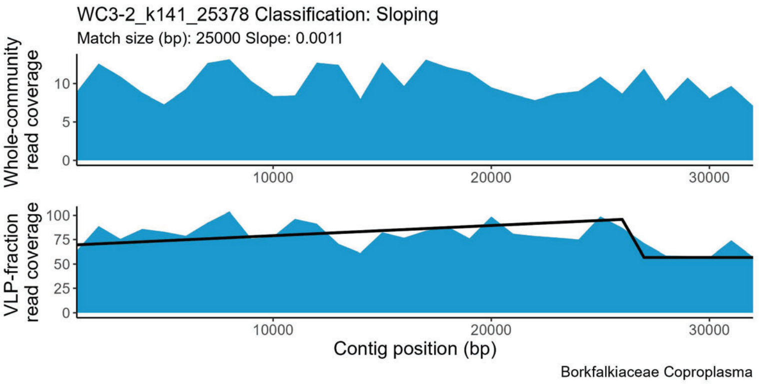
WC3-2_k141_25378 Classification: Sloping
Match size (bp): 25000 Slope: 0.0011
Whole-community read coverage
VLP-fraction read coverage
Contig position (bp)
Borkfalkiaceae Coproplasma

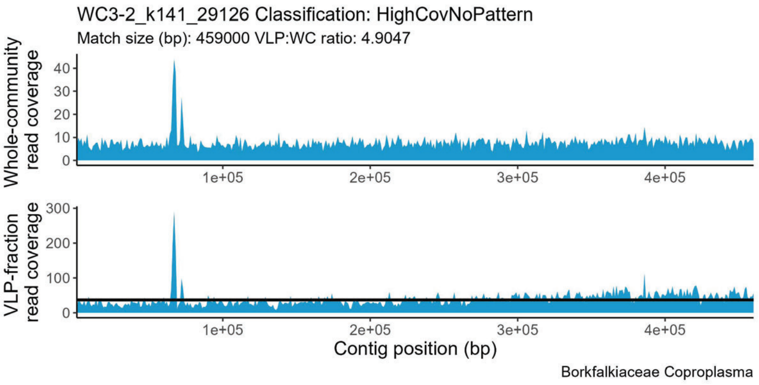
WC3-2_k141_29126 Classification: HighCovNoPattern
Match size (bp): 459000 VLP:WC ratio: 4.9047
Whole-community read coverage
VLP-fraction read coverage
Contig position (bp)
Borkfalkiaceae Coproplasma

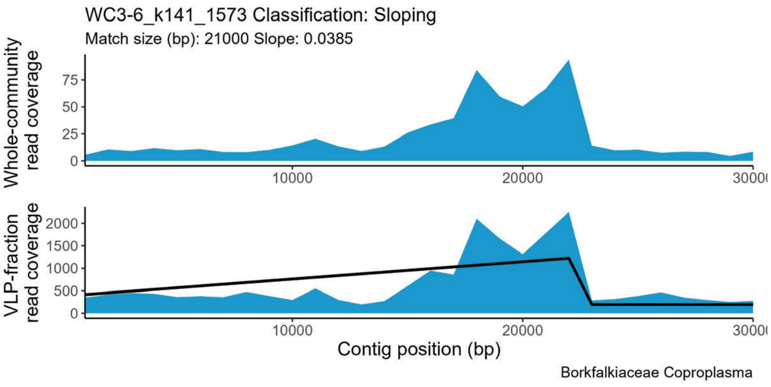
WC3-6_k141_1573 Classification: Sloping
Match size (bp): 21000 Slope: 0.0385
Whole-community read coverage
VLP-fraction read coverage
Contig position (bp)
Borkfalkiaceae Coproplasma

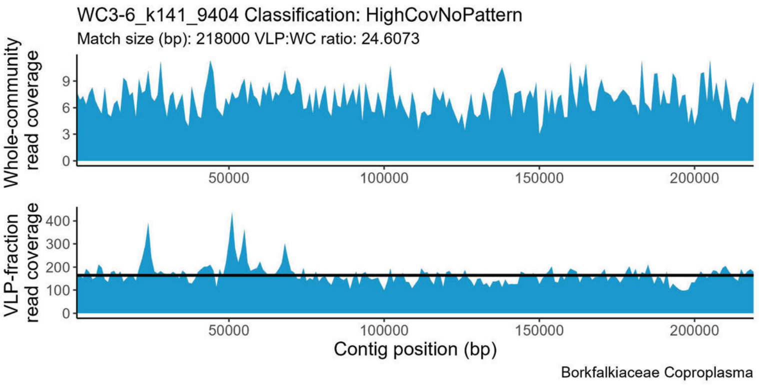
WC3-6_k141_9404 Classification: HighCovNoPattern
Match size (bp): 218000 VLP:WC ratio: 24.6073
Whole-community read coverage
VLP-fraction read coverage
Contig position (bp)
Borkfalkiaceae Coproplasma

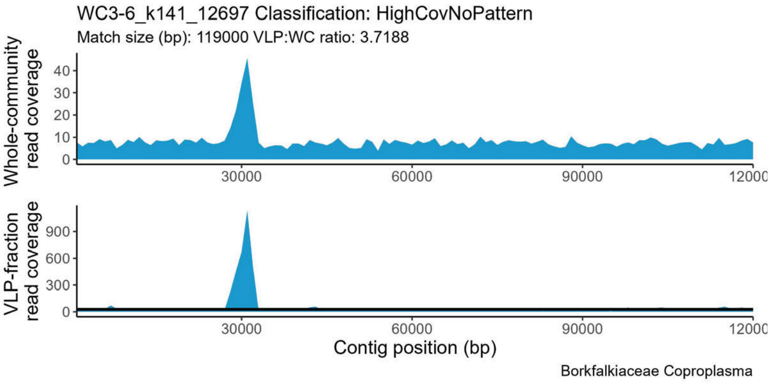
WC3-6_k141_12697 Classification: HighCovNoPattern
Match size (bp): 119000 VLP:WC ratio: 3.7188
Whole-community read coverage
VLP-fraction read coverage
Contig position (bp)
Borkfalkiaceae Coproplasma

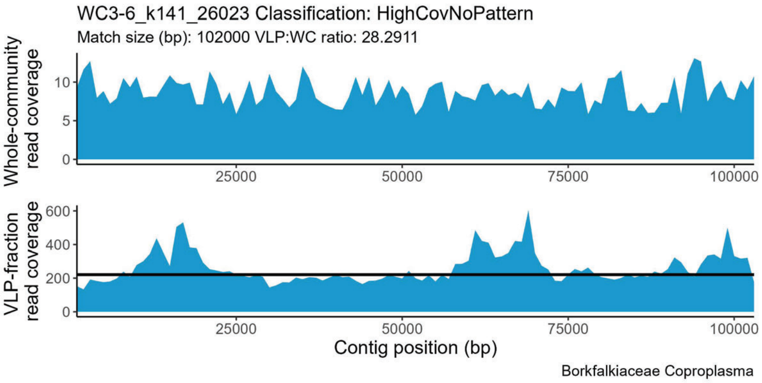
WC3-6_k141_26023 Classification: HighCovNoPattern
Match size (bp): 102000 VLP:WC ratio: 28.2911
Whole-community read coverage
VLP-fraction read coverage
Contig position (bp)
Borkfalkiaceae Coproplasma

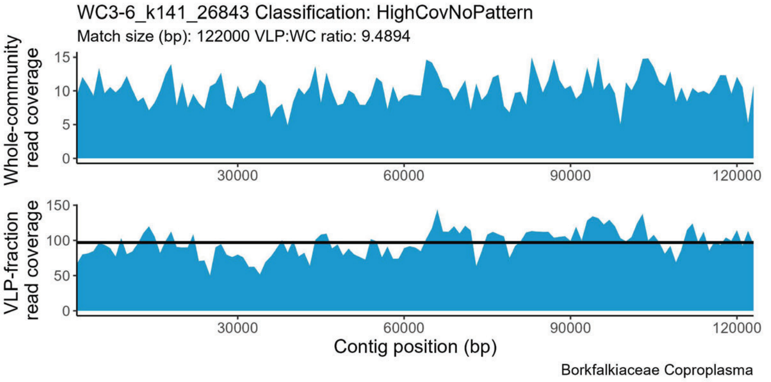
WC3-6_k141_26843 Classification: HighCovNoPattern
Match size (bp): 122000 VLP:WC ratio: 9.4894
Whole-community read coverage
VLP-fraction read coverage
Contig position (bp)
Borkfalkiaceae Coproplasma

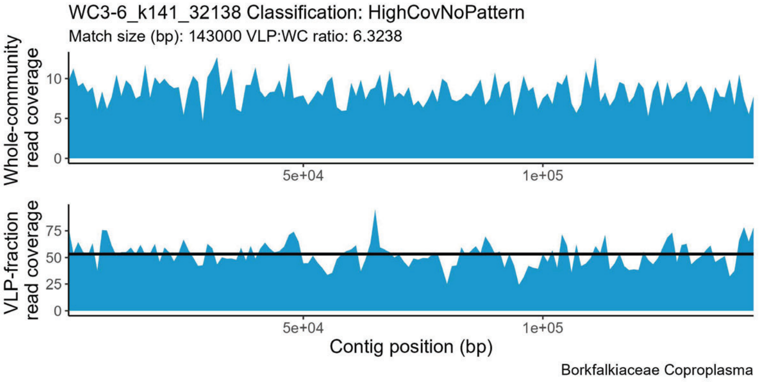
WC3-6_k141_32138 Classification: HighCovNoPattern
Match size (bp): 143000 VLP:WC ratio: 6.3238
Whole-community read coverage
VLP-fraction read coverage
Contig position (bp)
Borkfalkiaceae Coproplasma

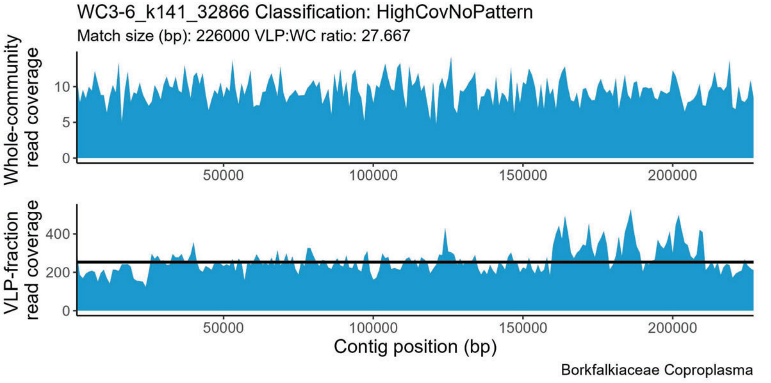
WC3-6_k141_32866 Classification: HighCovNoPattern
Match size (bp): 226000 VLP:WC ratio: 27.667
Whole-community read coverage
VLP-fraction read coverage
Contig position (bp)
Borkfalkiaceae Coproplasma

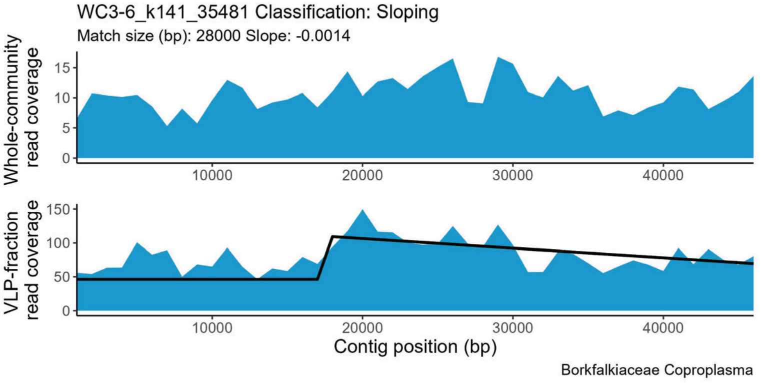
WC3-6_k141_35481 Classification: Sloping
Match size (bp): 28000 Slope: -0.0014
Whole-community read coverage
VLP-fraction read coverage
Contig position (bp)
Borkfalkiaceae Coproplasma

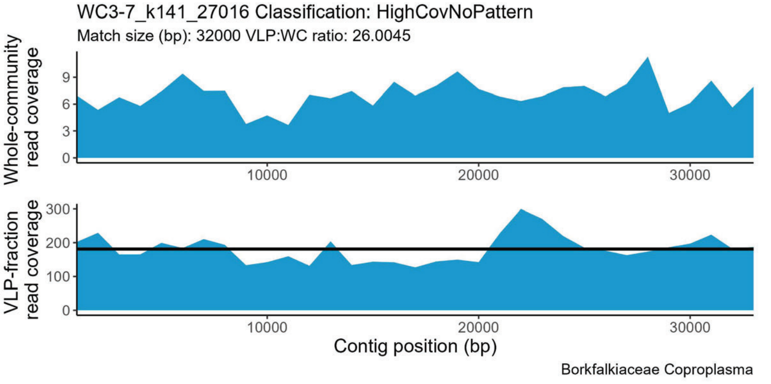
WC3-7_k141_27016 Classification: HighCovNoPattern
Match size (bp): 32000 VLP:WC ratio: 26.0045
Whole-community read coverage
VLP-fraction read coverage
Contig position (bp)
Borkfalkiaceae Coproplasma

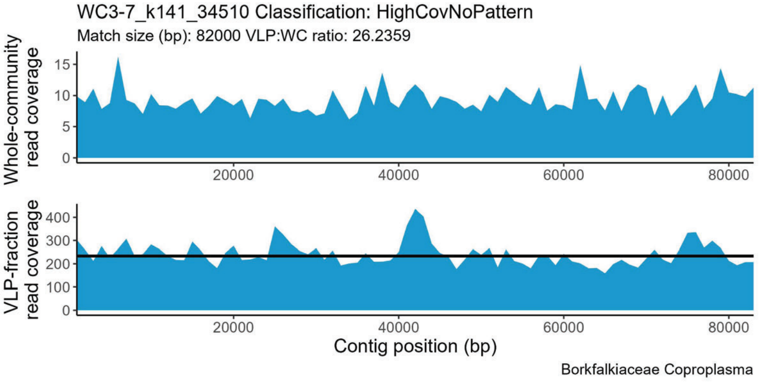
WC3-7_k141_34510 Classification: HighCovNoPattern
Match size (bp): 82000 VLP:WC ratio: 26.2359
Whole-community read coverage
VLP-fraction read coverage
Contig position (bp)
Borkfalkiaceae Coproplasma

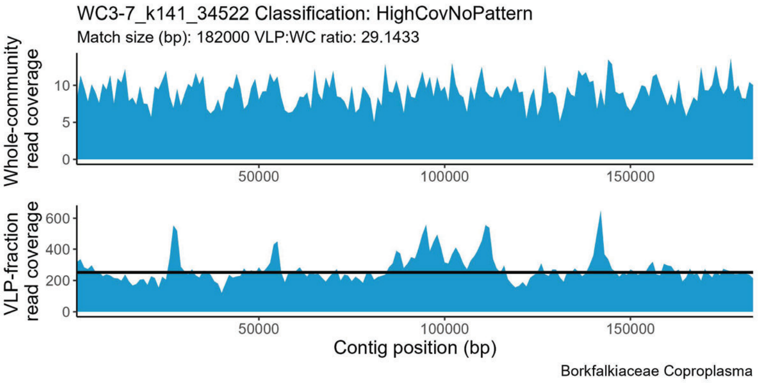
WC3-7_k141_34522 Classification: HighCovNoPattern
Match size (bp): 182000 VLP:WC ratio: 29.1433
Whole-community read coverage
0
5
10
50000
100000
150000
VLP-fraction read coverage
0
200
400
600
Contig position (bp)
Borkfalkiaceae Coproplasma

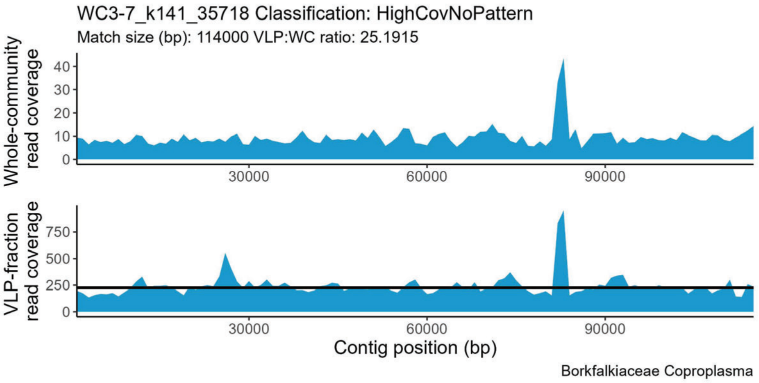
WC3-7_k141_35718 Classification: HighCovNoPattern
Match size (bp): 114000 VLP:WC ratio: 25.1915
Whole-community read coverage
0
10
20
30
40
30000
60000
90000
VLP-fraction read coverage
0
250
500
750
Contig position (bp)
Borkfalkiaceae Coproplasma

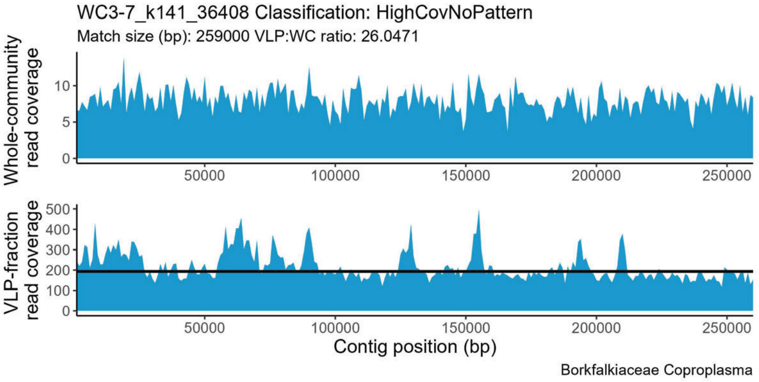
WC3-7_k141_36408 Classification: HighCovNoPattern
Match size (bp): 259000 VLP:WC ratio: 26.0471
Whole-community read coverage
0
5
10
50000
100000
150000
200000
250000
VLP-fraction read coverage
0
100
200
300
400
500
Contig position (bp)
Borkfalkiaceae Coproplasma

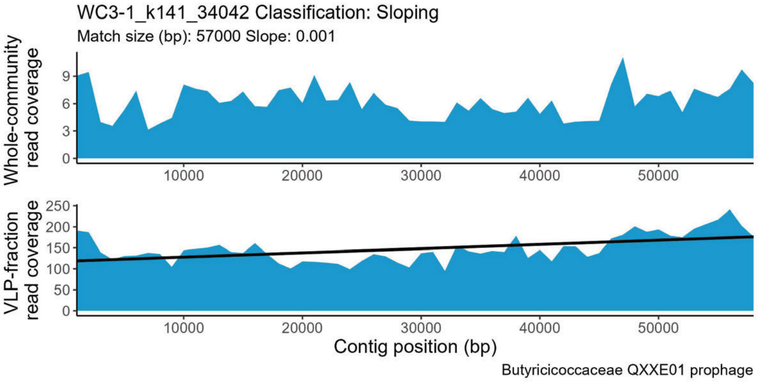
WC3-1_k141_34042 Classification: Sloping
Match size (bp): 57000 Slope: 0.001
Whole-community read coverage
0
3
6
9
10000
20000
30000
40000
50000
VLP-fraction read coverage
0
50
100
150
200
250
Contig position (bp)
Butyricicoccaceae QXXE01 prophage

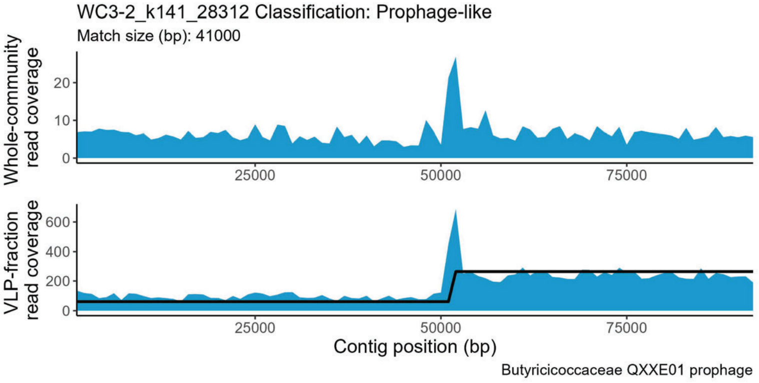
WC3-2_k141_28312 Classification: Prophage-like
Match size (bp): 41000
Whole-community read coverage
0
10
20
25000
50000
75000
VLP-fraction read coverage
0
200
400
600
Contig position (bp)
Butyricicoccaceae QXXE01 prophage

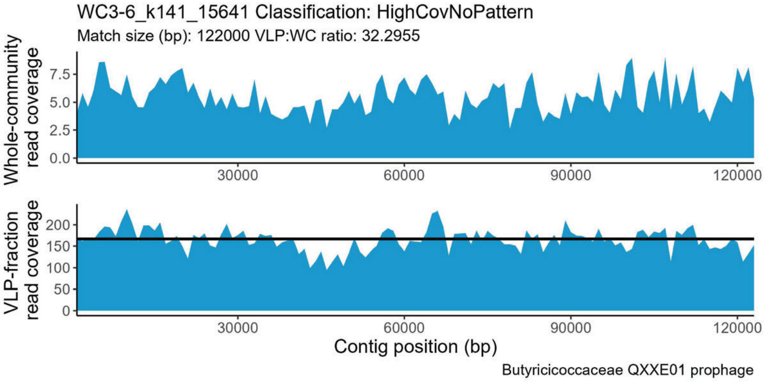
WC3-6_k141_15641 Classification: HighCovNoPattern
Match size (bp): 122000 VLP:WC ratio: 32.2955
Whole-community read coverage
0.0
2.5
5.0
7.5
30000
60000
90000
120000
VLP-fraction read coverage
0
50
100
150
200
Contig position (bp)
Butyricicoccaceae QXXE01 prophage

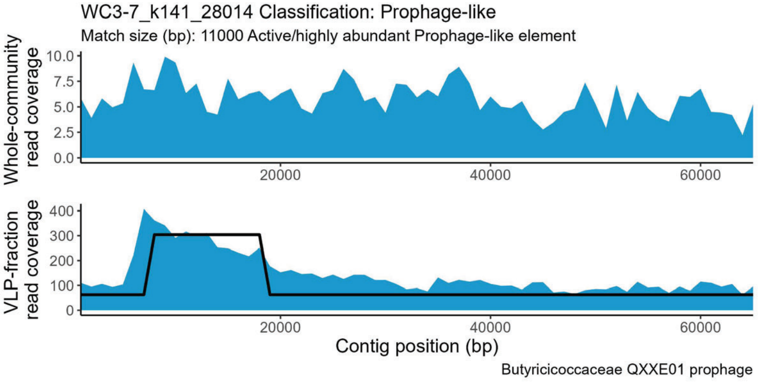
WC3-7_k141_28014 Classification: Prophage-like
Match size (bp): 11000 Active/highly abundant Prophage-like element
Whole-community read coverage
0.0
2.5
5.0
7.5
10.0
20000
40000
60000
VLP-fraction read coverage
0
100
200
300
400
Contig position (bp)
Butyricicoccaceae QXXE01 prophage

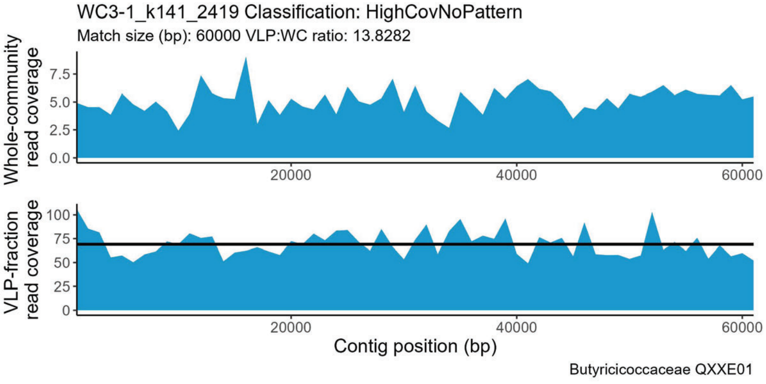
WC3-1_k141_2419 Classification: HighCovNoPattern
Match size (bp): 60000 VLP:WC ratio: 13.8282
Whole-community read coverage
0.0
2.5
5.0
7.5
20000
40000
60000
VLP-fraction read coverage
0
25
50
75
100
Contig position (bp)
Butyricicoccaceae QXXE01

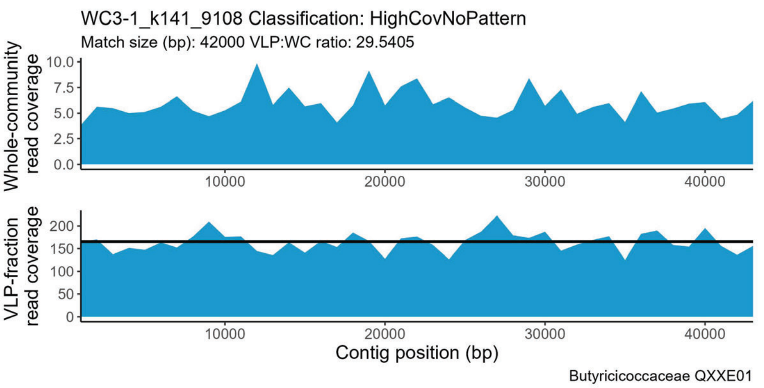
WC3-1_k141_9108 Classification: HighCovNoPattern
Match size (bp): 42000 VLP:WC ratio: 29.5405
Whole-community read coverage
VLP-fraction read coverage
Contig position (bp)
Butyricicoccaceae QXXE01

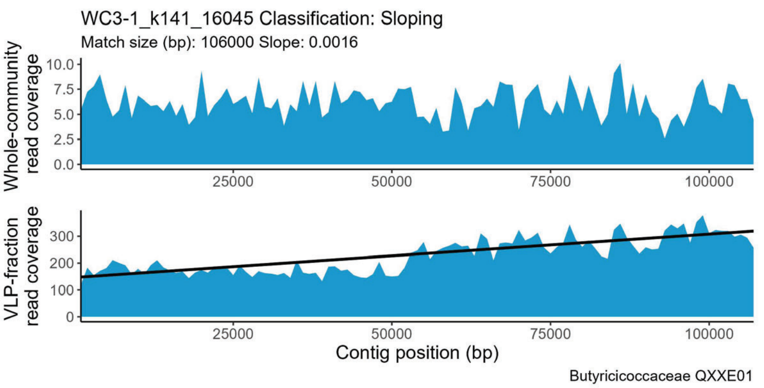
WC3-1_k141_16045 Classification: Sloping
Match size (bp): 106000 Slope: 0.0016
Whole-community read coverage
VLP-fraction read coverage
Contig position (bp)
Butyricicoccaceae QXXE01

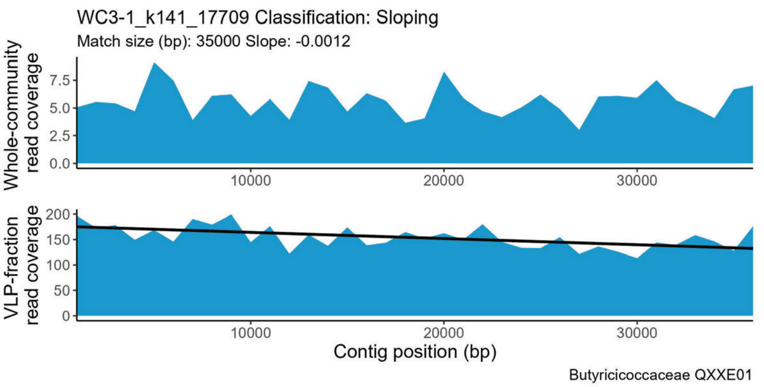
WC3-1_k141_17709 Classification: Sloping
Match size (bp): 35000 Slope: -0.0012
Whole-community read coverage
VLP-fraction read coverage
Contig position (bp)
Butyricicoccaceae QXXE01

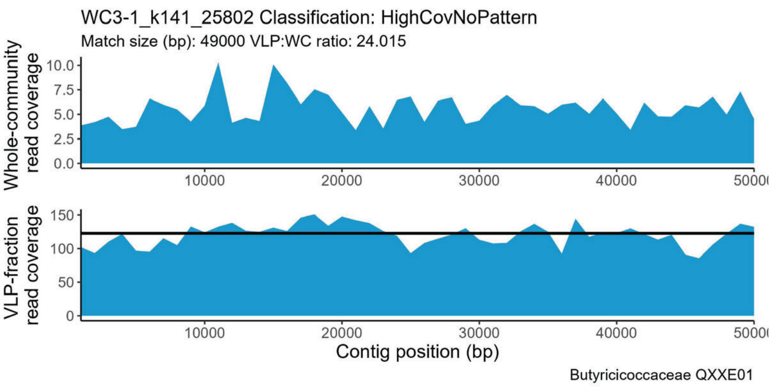
WC3-1_k141_25802 Classification: HighCovNoPattern
Match size (bp): 49000 VLP:WC ratio: 24.015
Whole-community read coverage
VLP-fraction read coverage
Contig position (bp)
Butyricicoccaceae QXXE01

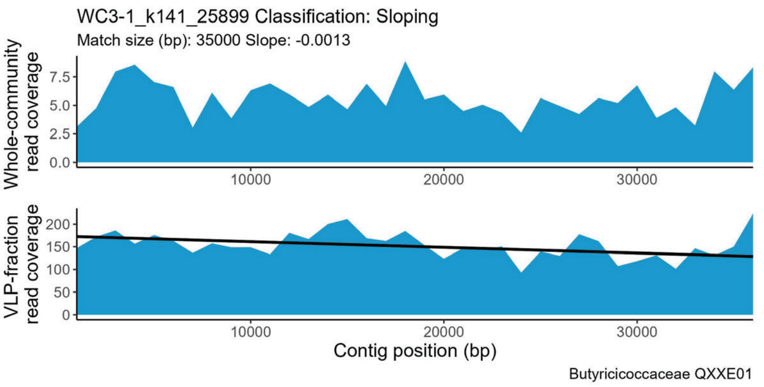
WC3-1_k141_25899 Classification: Sloping
Match size (bp): 35000 Slope: -0.0013
Whole-community read coverage
VLP-fraction read coverage
Contig position (bp)
Butyricicoccaceae QXXE01

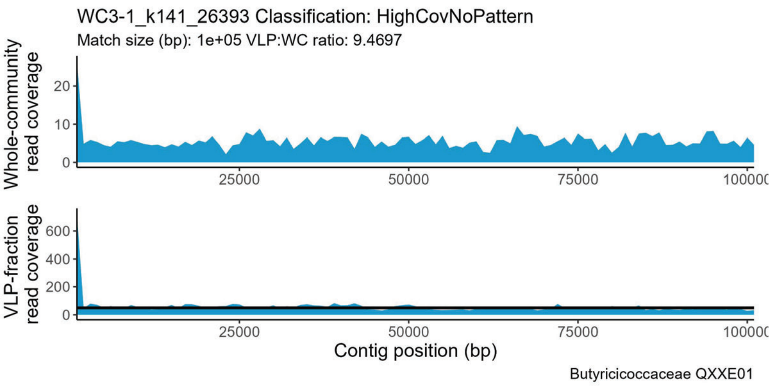
WC3-1_k141_26393 Classification: HighCovNoPattern
Match size (bp): 1e+05 VLP:WC ratio: 9.4697
Whole-community read coverage
VLP-fraction read coverage
Contig position (bp)
Butyricicoccaceae QXXE01

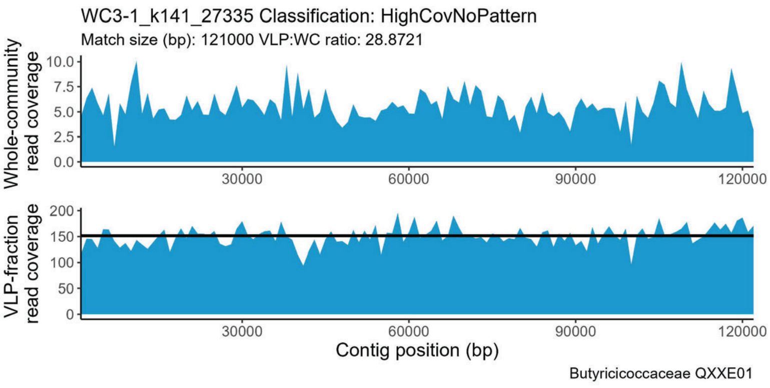
WC3-1_k141_27335 Classification: HighCovNoPattern
Match size (bp): 121000 VLP:WC ratio: 28.8721
Whole-community read coverage
VLP-fraction read coverage
Contig position (bp)
Butyricicoccaceae QXXE01

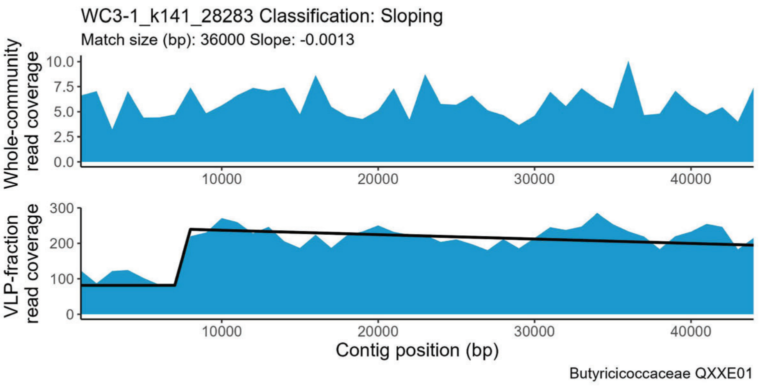
WC3-1_k141_28283 Classification: Sloping
Match size (bp): 36000 Slope: -0.0013
Whole-community read coverage
VLP-fraction read coverage
Contig position (bp)
Butyricicoccaceae QXXE01

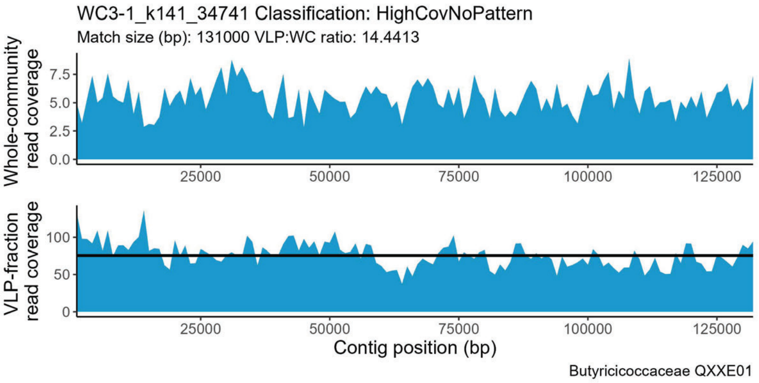
WC3-1_k141_34741 Classification: HighCovNoPattern
Match size (bp): 131000 VLP:WC ratio: 14.4413
Whole-community read coverage
VLP-fraction read coverage
Contig position (bp)
Butyricicoccaceae QXXE01

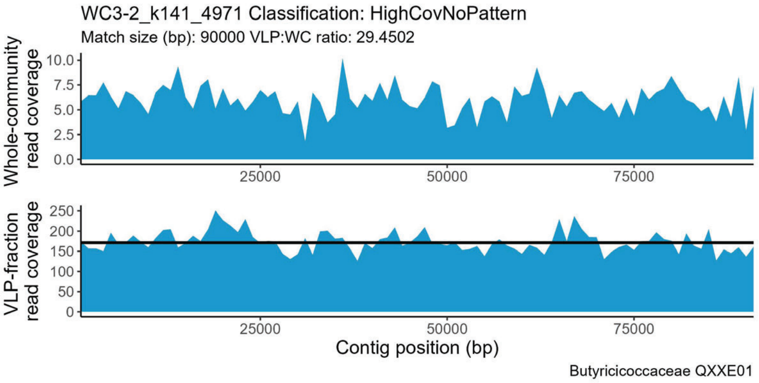
WC3-2_k141_4971 Classification: HighCovNoPattern
Match size (bp): 90000 VLP:WC ratio: 29.4502
Whole-community read coverage
VLP-fraction read coverage
Contig position (bp)
Butyricicoccaceae QXXE01

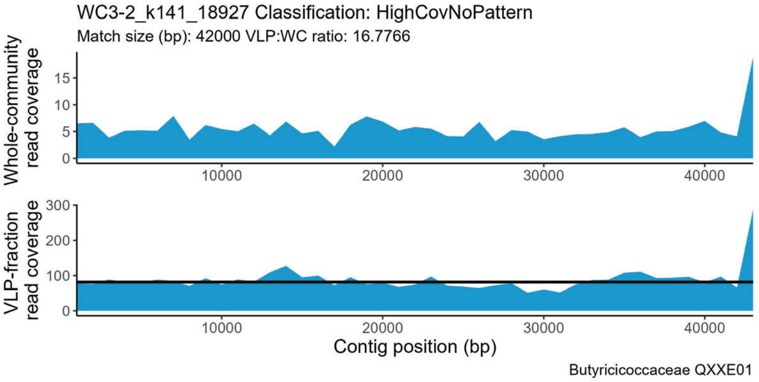
WC3-2_k141_18927 Classification: HighCovNoPattern
Match size (bp): 42000 VLP:WC ratio: 16.7766
Whole-community read coverage
VLP-fraction read coverage
Contig position (bp)
Butyricicoccaceae QXXE01

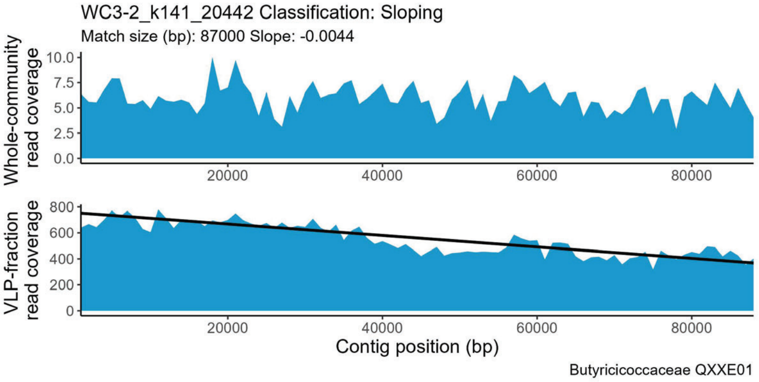
WC3-2_k141_20442 Classification: Sloping
Match size (bp): 87000 Slope: -0.0044
Whole-community read coverage
VLP-fraction read coverage
Contig position (bp)
Butyricicoccaceae QXXE01

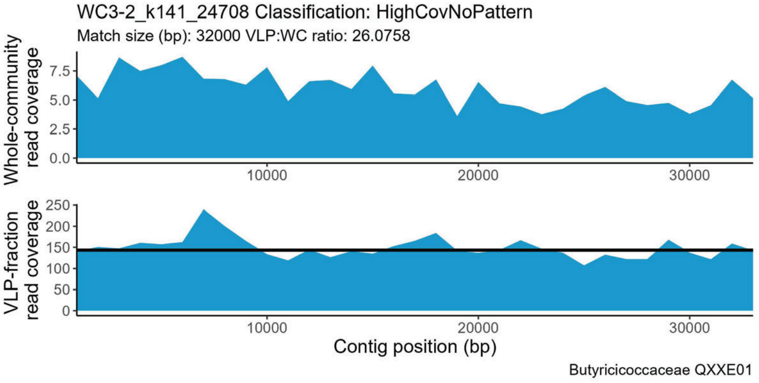
WC3-2_k141_24708 Classification: HighCovNoPattern
Match size (bp): 32000 VLP:WC ratio: 26.0758
Whole-community read coverage
VLP-fraction read coverage
Contig position (bp)
Butyricicoccaceae QXXE01

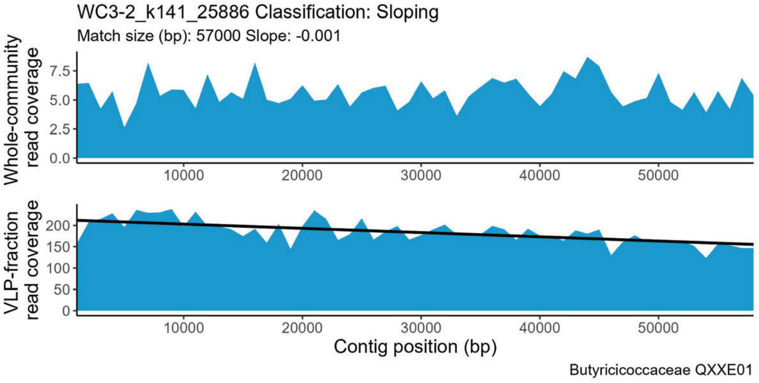
WC3-2_k141_25886 Classification: Sloping
Match size (bp): 57000 Slope: -0.001
Whole-community read coverage
VLP-fraction read coverage
Contig position (bp)
Butyricicoccaceae QXXE01

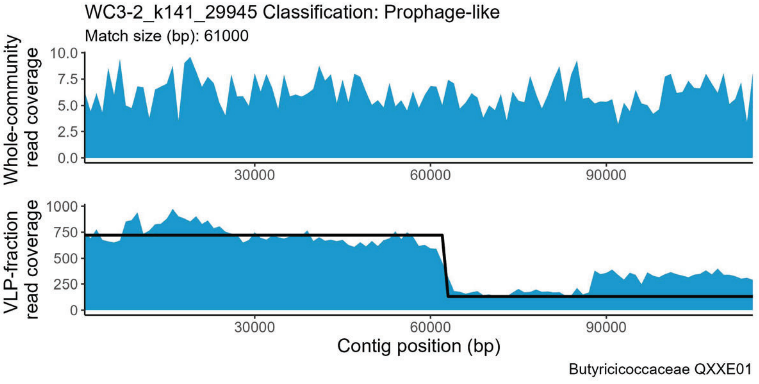
WC3-2_k141_29945 Classification: Prophage-like
Match size (bp): 61000
Whole-community read coverage
VLP-fraction read coverage
Contig position (bp)
Butyricicoccaceae QXXE01

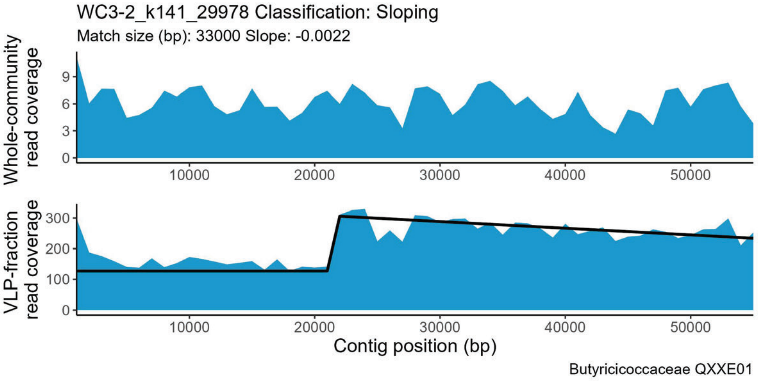
WC3-2_k141_29978 Classification: Sloping
Match size (bp): 33000 Slope: -0.0022
Whole-community read coverage
VLP-fraction read coverage
Contig position (bp)
Butyricicoccaceae QXXE01

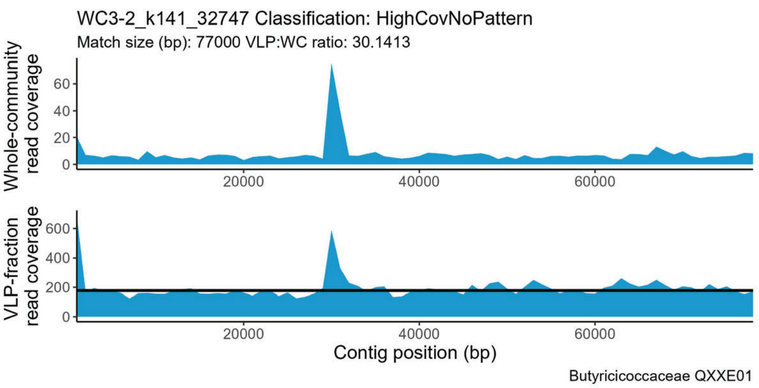
WC3-2_k141_32747 Classification: HighCovNoPattern
Match size (bp): 77000 VLP:WC ratio: 30.1413
Whole-community read coverage
VLP-fraction read coverage
Contig position (bp)
Butyricicoccaceae QXXE01

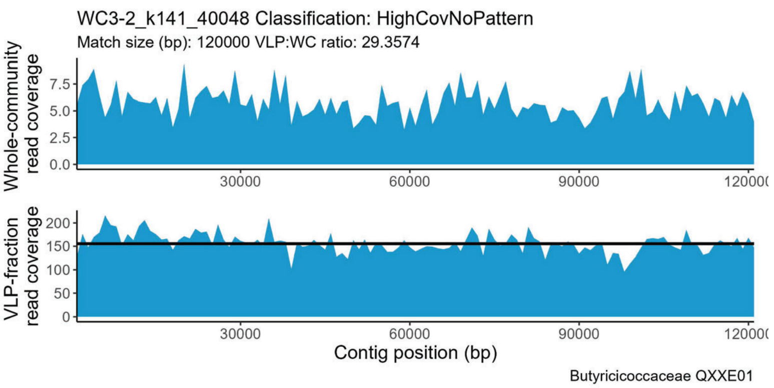
WC3-2_k141_40048 Classification: HighCovNoPattern
Match size (bp): 120000 VLP:WC ratio: 29.3574
Whole-community read coverage
VLP-fraction read coverage
Contig position (bp)
Butyricicoccaceae QXXE01

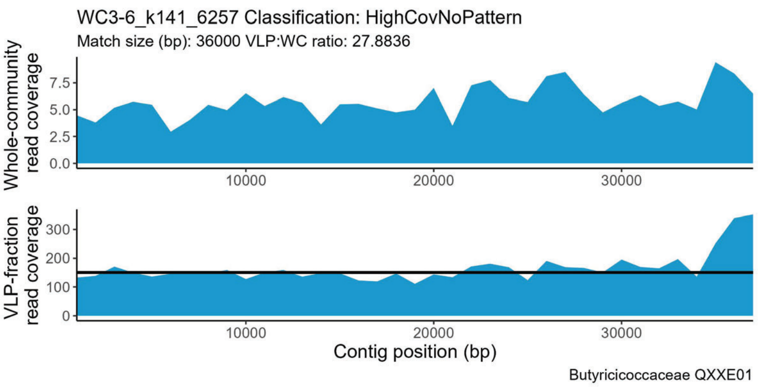
WC3-6_k141_6257 Classification: HighCovNoPattern
Match size (bp): 36000 VLP:WC ratio: 27.8836
Whole-community read coverage
VLP-fraction read coverage
Contig position (bp)
Butyricicoccaceae QXXE01

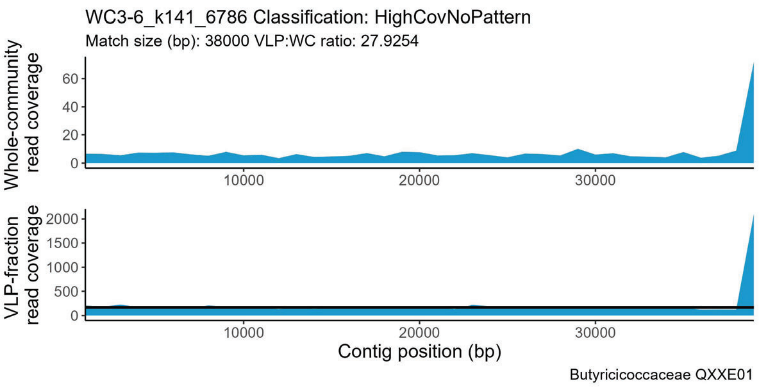
WC3-6_k141_6786 Classification: HighCovNoPattern
Match size (bp): 38000 VLP:WC ratio: 27.9254
Whole-community read coverage
VLP-fraction read coverage
Contig position (bp)
Butyricicoccaceae QXXE01

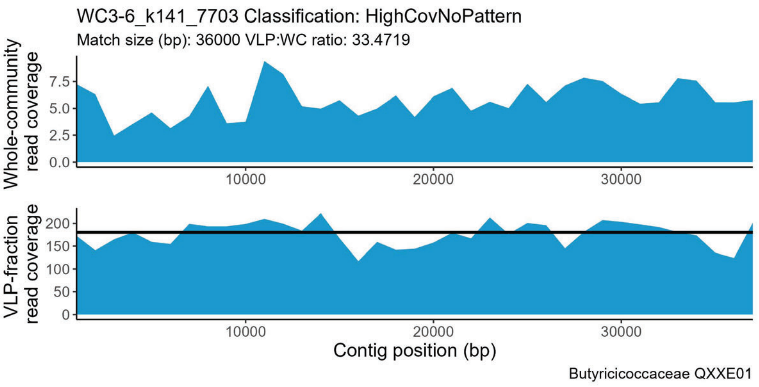
WC3-6_k141_7703 Classification: HighCovNoPattern
Match size (bp): 36000 VLP:WC ratio: 33.4719
Whole-community read coverage
VLP-fraction read coverage
Contig position (bp)
Butyricicoccaceae QXXE01

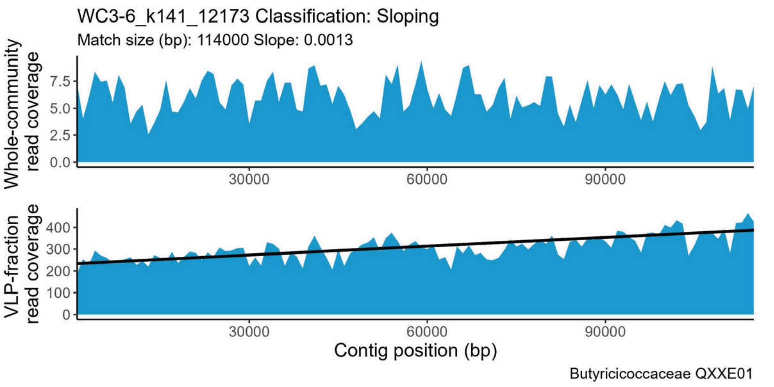
WC3-6_k141_12173 Classification: Sloping
Match size (bp): 114000 Slope: 0.0013
Whole-community read coverage
VLP-fraction read coverage
Contig position (bp)
Butyricicoccaceae QXXE01

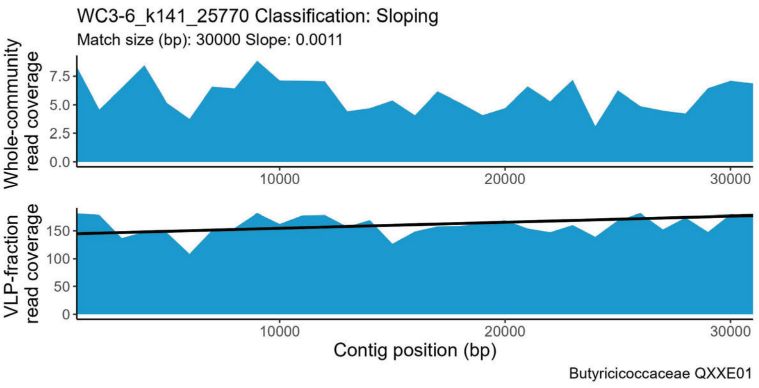
WC3-6_k141_25770 Classification: Sloping
Match size (bp): 30000 Slope: 0.0011
Whole-community read coverage
VLP-fraction read coverage
Contig position (bp)
Butyricicoccaceae QXXE01

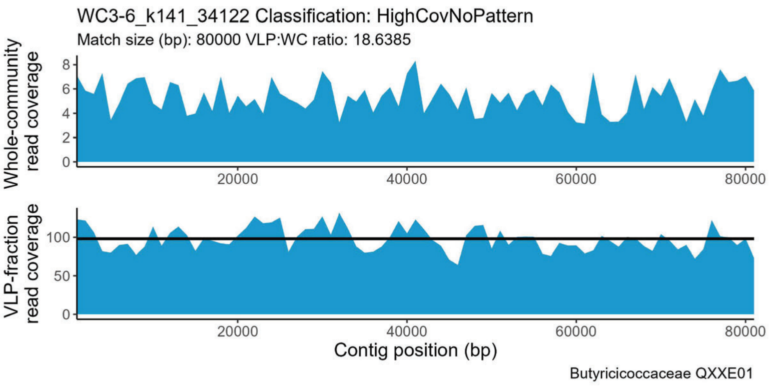
WC3-6_k141_34122 Classification: HighCovNoPattern
Match size (bp): 80000 VLP:WC ratio: 18.6385
Whole-community read coverage
VLP-fraction read coverage
Contig position (bp)
Butyricicoccaceae QXXE01

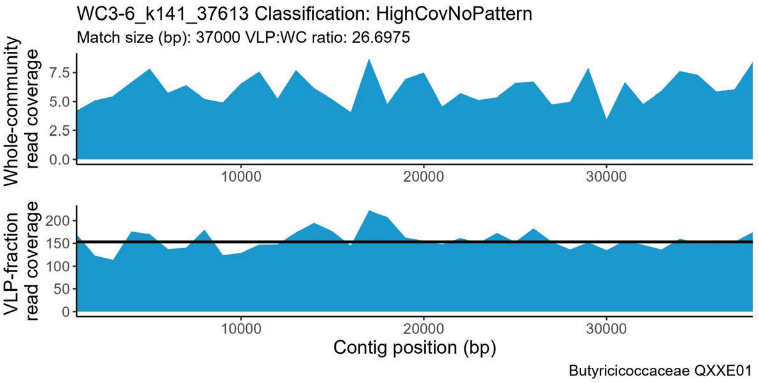
WC3-6_k141_37613 Classification: HighCovNoPattern
Match size (bp): 37000 VLP:WC ratio: 26.6975
Whole-community read coverage
VLP-fraction read coverage
Contig position (bp)
Butyricicoccaceae QXXE01

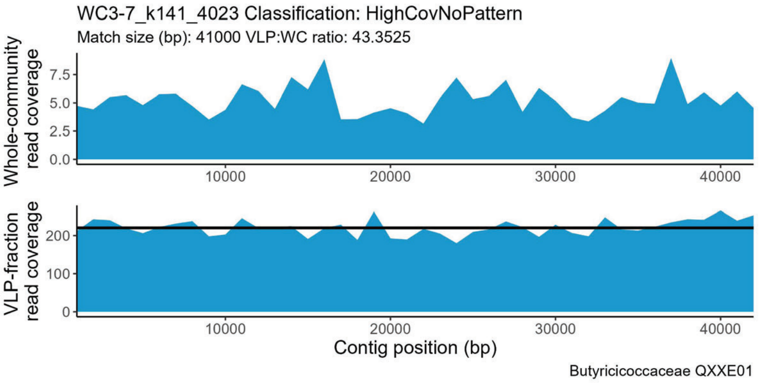
WC3-7_k141_4023 Classification: HighCovNoPattern
Match size (bp): 41000 VLP:WC ratio: 43.3525
Whole-community read coverage
VLP-fraction read coverage
Contig position (bp)
Butyricicoccaceae QXXE01

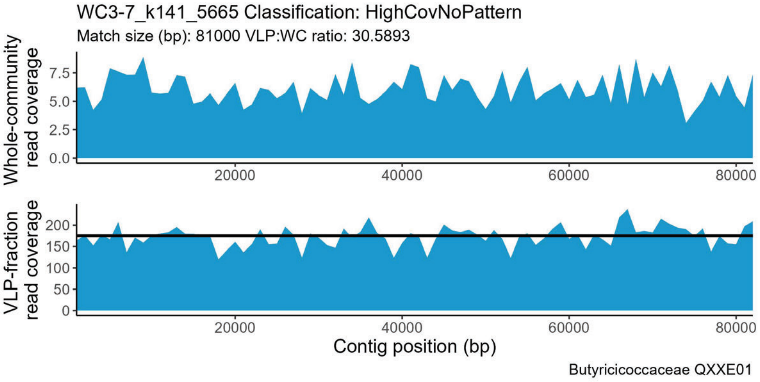
WC3-7_k141_5665 Classification: HighCovNoPattern
Match size (bp): 81000 VLP:WC ratio: 30.5893
Whole-community read coverage
VLP-fraction read coverage
Contig position (bp)
Butyricicoccaceae QXXE01

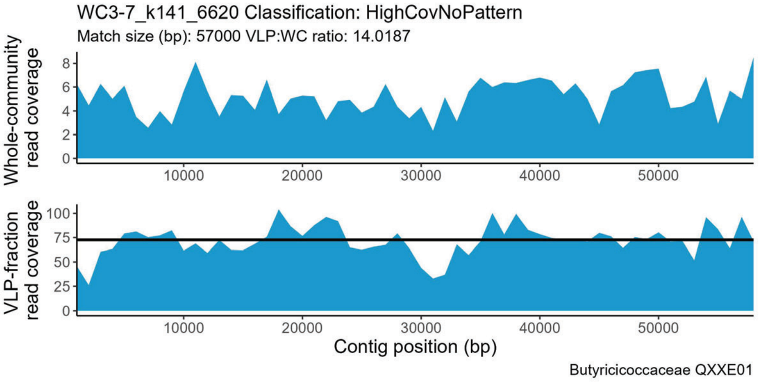
WC3-7_k141_6620 Classification: HighCovNoPattern
Match size (bp): 57000 VLP:WC ratio: 14.0187
Whole-community read coverage
VLP-fraction read coverage
Contig position (bp)
Butyricicoccaceae QXXE01

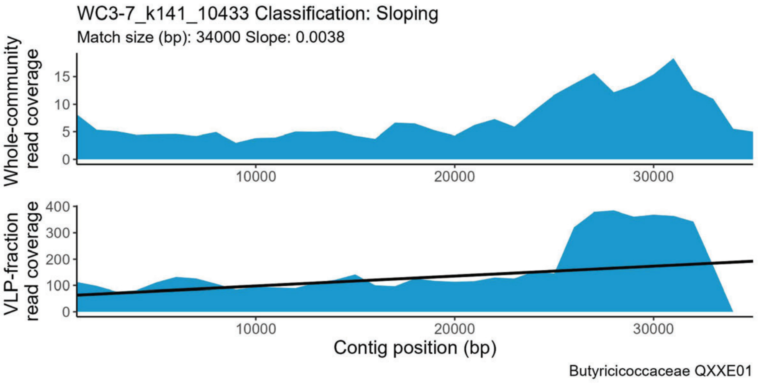
WC3-7_k141_10433 Classification: Sloping
Match size (bp): 34000 Slope: 0.0038
Whole-community read coverage
VLP-fraction read coverage
Contig position (bp)
Butyricicoccaceae QXXE01

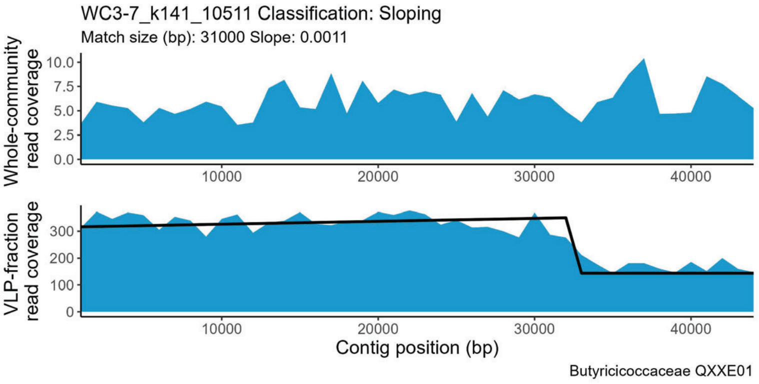
WC3-7_k141_10511 Classification: Sloping
Match size (bp): 31000 Slope: 0.0011
Whole-community read coverage
VLP-fraction read coverage
Contig position (bp)
Butyricicoccaceae QXXE01

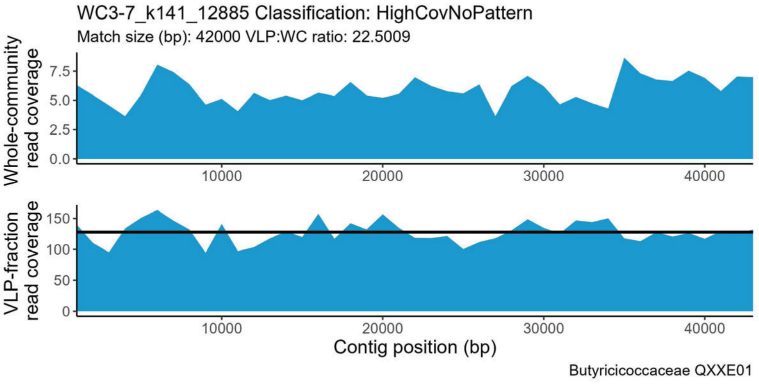
WC3-7_k141_12885 Classification: HighCovNoPattern
Match size (bp): 42000 VLP:WC ratio: 22.5009
Whole-community read coverage
VLP-fraction read coverage
Contig position (bp)
Butyricicoccaceae QXXE01

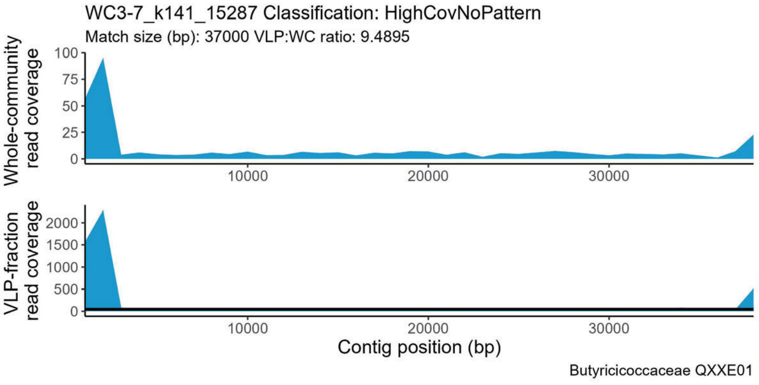
WC3-7_k141_15287 Classification: HighCovNoPattern
Match size (bp): 37000 VLP:WC ratio: 9.4895
Whole-community read coverage
VLP-fraction read coverage
Contig position (bp)
Butyricicoccaceae QXXE01

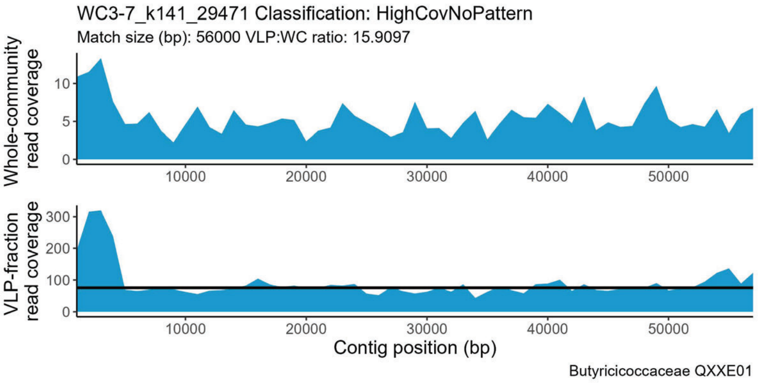
WC3-7_k141_29471 Classification: HighCovNoPattern
Match size (bp): 56000 VLP:WC ratio: 15.9097
Whole-community read coverage
VLP-fraction read coverage
Contig position (bp)
Butyricicoccaceae QXXE01

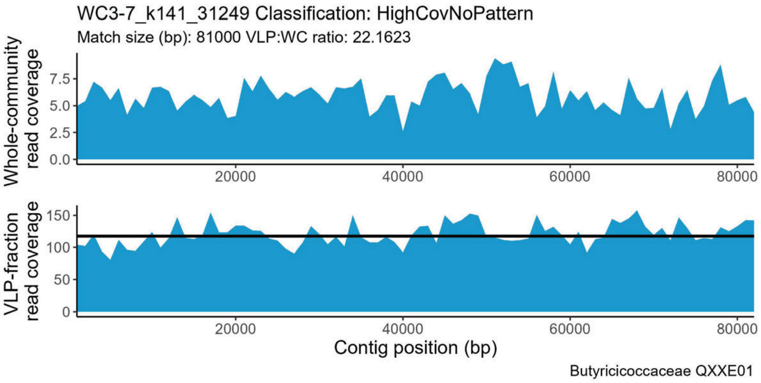
WC3-7_k141_31249 Classification: HighCovNoPattern
Match size (bp): 81000 VLP:WC ratio: 22.1623
Whole-community read coverage
VLP-fraction read coverage
Contig position (bp)
Butyricicoccaceae QXXE01

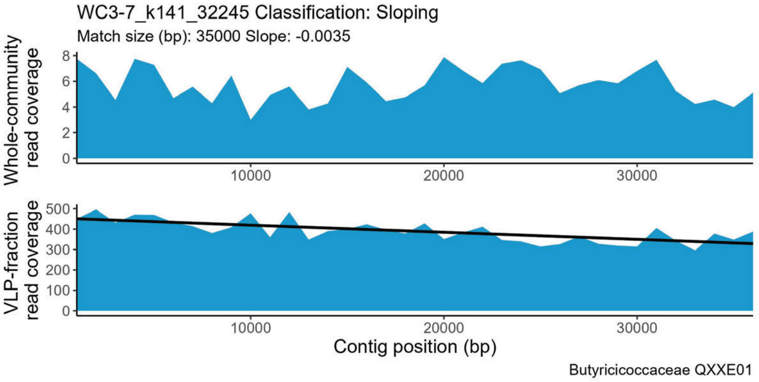
WC3-7_k141_32245 Classification: Sloping
Match size (bp): 35000 Slope: -0.0035
Whole-community read coverage
VLP-fraction read coverage
Contig position (bp)
Butyricicoccaceae QXXE01

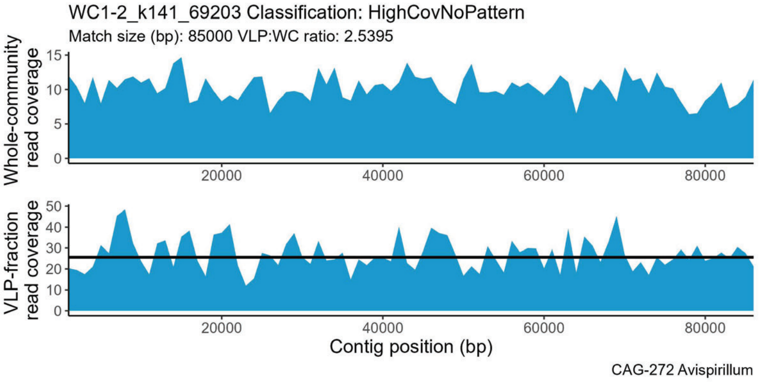
WC1-2_k141_69203 Classification: HighCovNoPattern
Match size (bp): 85000 VLP:WC ratio: 2.5395
Whole-community read coverage
VLP-fraction read coverage
Contig position (bp)
CAG-272 Avispirillum

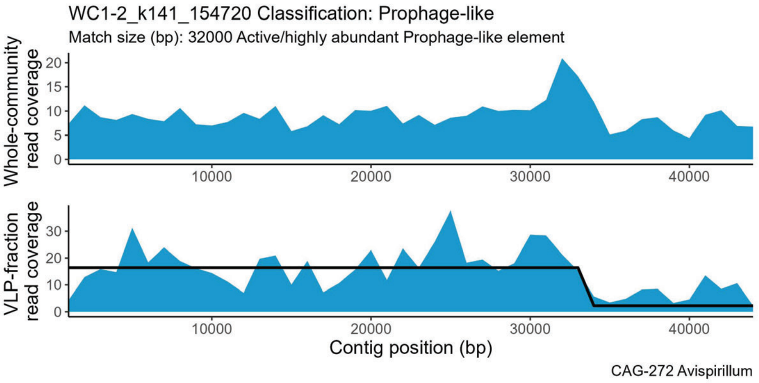
WC1-2_k141_154720 Classification: Prophage-like
Match size (bp): 32000 Active/highly abundant Prophage-like element
Whole-community read coverage
VLP-fraction read coverage
Contig position (bp)
CAG-272 Avispirillum

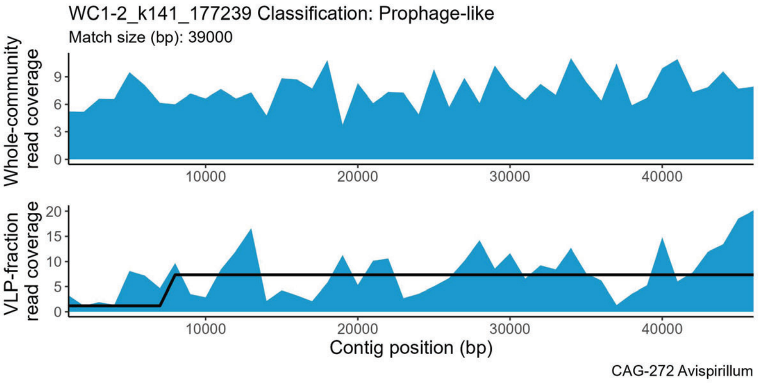
WC1-2_k141_177239 Classification: Prophage-like
Match size (bp): 39000
Whole-community read coverage
VLP-fraction read coverage
Contig position (bp)
CAG-272 Avispirillum

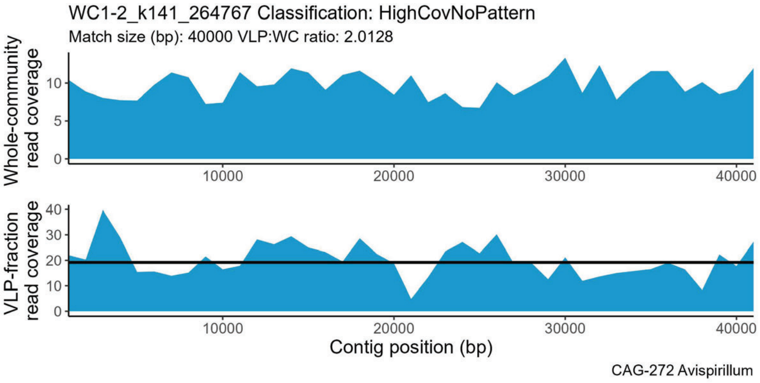
WC1-2_k141_264767 Classification: HighCovNoPattern
Match size (bp): 40000 VLP:WC ratio: 2.0128
Whole-community read coverage
VLP-fraction read coverage
Contig position (bp)
CAG-272 Avispirillum

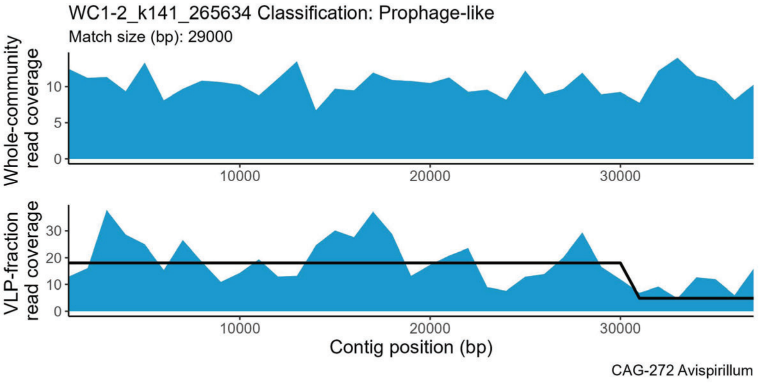
WC1-2_k141_265634 Classification: Prophage-like
Match size (bp): 29000
Whole-community read coverage
VLP-fraction read coverage
Contig position (bp)
CAG-272 Avispirillum

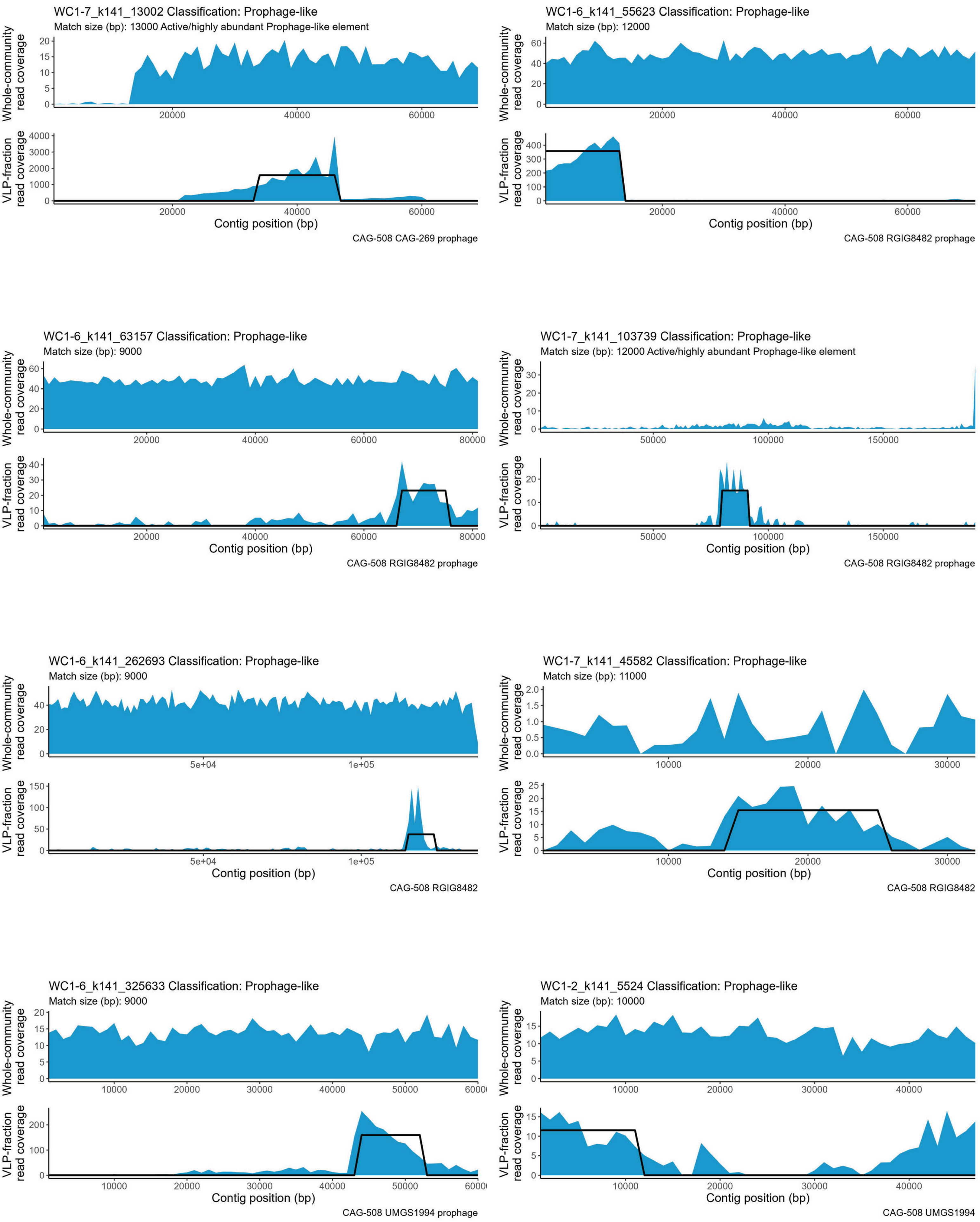

WC1-7_k141_13002 Classification: Prophage-like
Match size (bp): 13000 Active/highly abundant Prophage-like element
Whole-community read coverage
VLP-fraction read coverage
Contig position (bp)
CAG-508 CAG-269 prophage
WC1-6_k141_55623 Classification: Prophage-like
Match size (bp): 12000
Whole-community read coverage
VLP-fraction read coverage
Contig position (bp)
CAG-508 RGIG8482 prophage
WC1-6_k141_63157 Classification: Prophage-like
Match size (bp): 9000
Whole-community read coverage
VLP-fraction read coverage
Contig position (bp)
CAG-508 RGIG8482 prophage
WC1-7_k141_103739 Classification: Prophage-like
Match size (bp): 12000 Active/highly abundant Prophage-like element
Whole-community read coverage
VLP-fraction read coverage
Contig position (bp)
CAG-508 RGIG8482 prophage
WC1-6_k141_262693 Classification: Prophage-like
Match size (bp): 9000
Whole-community read coverage
VLP-fraction read coverage
Contig position (bp)
CAG-508 RGIG8482
WC1-7_k141_45582 Classification: Prophage-like
Match size (bp): 11000
Whole-community read coverage
VLP-fraction read coverage
Contig position (bp)
CAG-508 RGIG8482
WC1-6_k141_325633 Classification: Prophage-like
Match size (bp): 9000
Whole-community read coverage
VLP-fraction read coverage
Contig position (bp)
CAG-508 UMGS1994 prophage
WC1-2_k141_5524 Classification: Prophage-like
Match size (bp): 10000
Whole-community read coverage
VLP-fraction read coverage
Contig position (bp)
CAG-508 UMGS1994

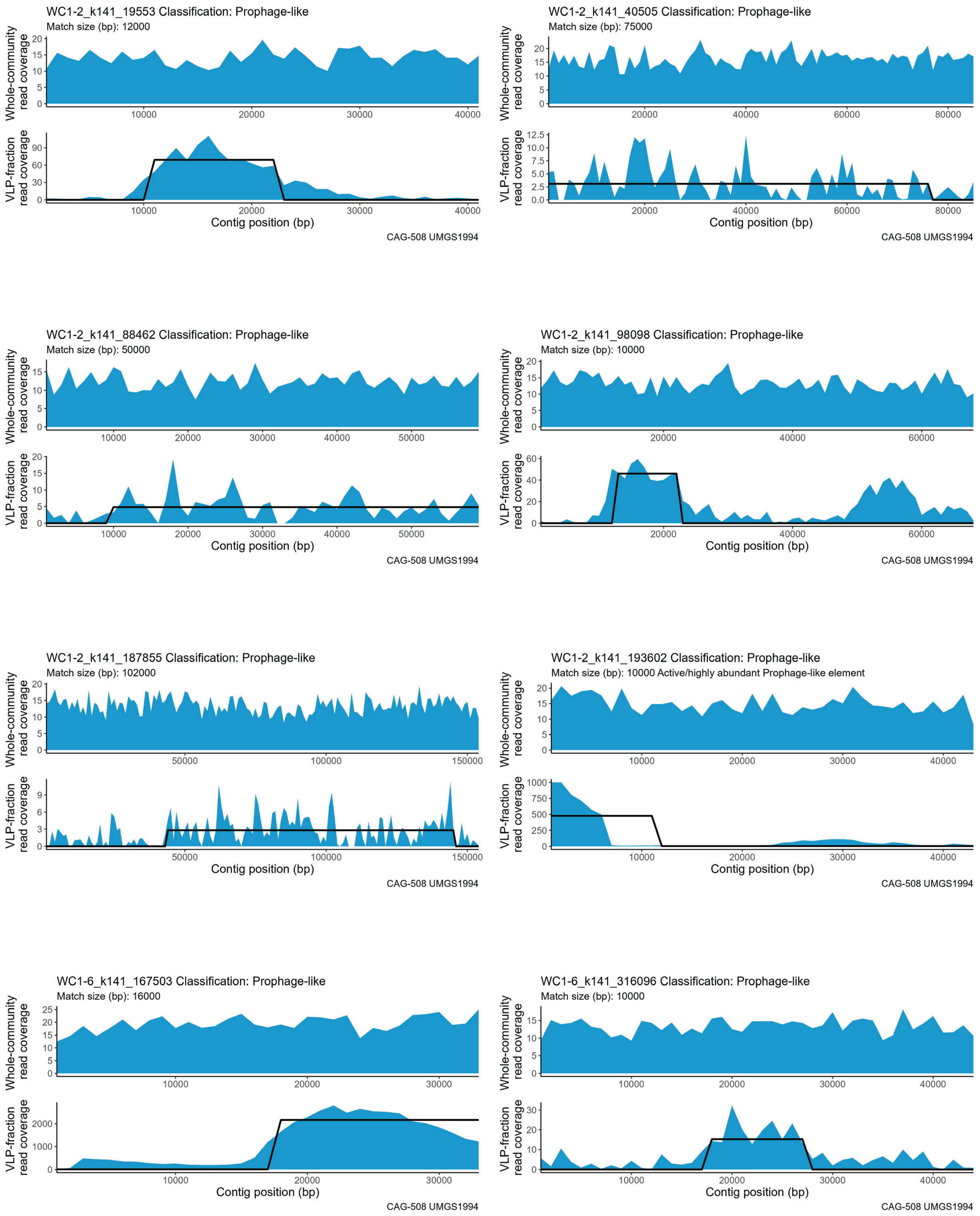
WC1-2_k141_19553 Classification: Prophage-like
Match size (bp): 12000
WC1-2_k141_40505 Classification: Prophage-like
Match size (bp): 75000
WC1-2_k141_88462 Classification: Prophage-like
Match size (bp): 50000
WC1-2_k141_98098 Classification: Prophage-like
Match size (bp): 10000
WC1-2_k141_187855 Classification: Prophage-like
Match size (bp): 102000
WC1-2_k141_193602 Classification: Prophage-like
Match size (bp): 10000 Active/highly abundant Prophage-like element
WC1-6_k141_167503 Classification: Prophage-like
Match size (bp): 16000
WC1-6_k141_316096 Classification: Prophage-like
Match size (bp): 10000
Whole-community read coverage
VLP-fraction read coverage
Contig position (bp)
CAG-508 UMGS1994

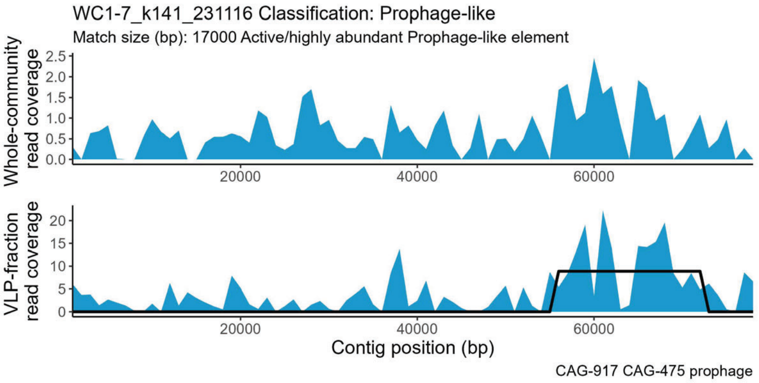
WC1-7_k141_231116 Classification: Prophage-like
Match size (bp): 17000 Active/highly abundant Prophage-like element
Whole-community read coverage
VLP-fraction read coverage
Contig position (bp)
CAG-917 CAG-475 prophage

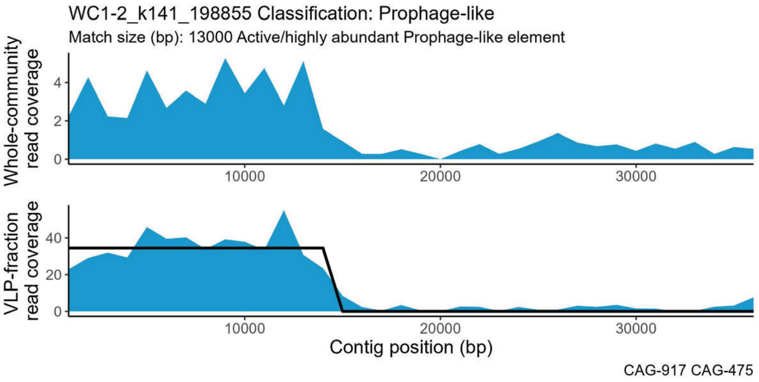
WC1-2_k141_198855 Classification: Prophage-like
Match size (bp): 13000 Active/highly abundant Prophage-like element
Whole-community read coverage
VLP-fraction read coverage
Contig position (bp)
CAG-917 CAG-475

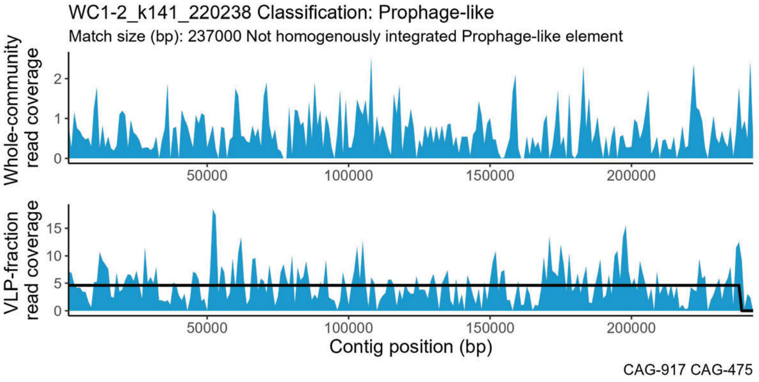
WC1-2_k141_220238 Classification: Prophage-like
Match size (bp): 237000 Not homogenously integrated Prophage-like element
Whole-community read coverage
VLP-fraction read coverage
Contig position (bp)
CAG-917 CAG-475

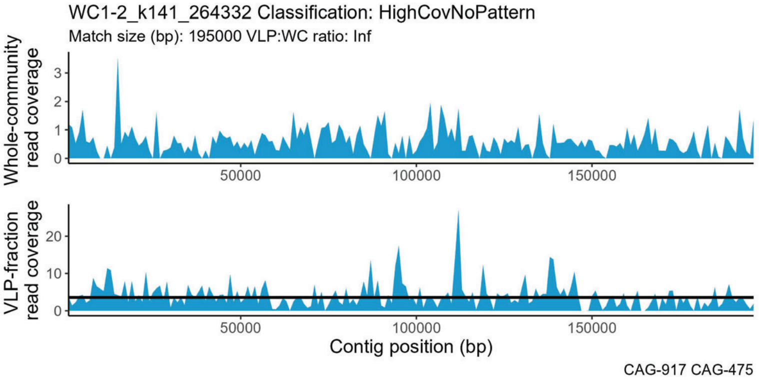
WC1-2_k141_264332 Classification: HighCovNoPattern
Match size (bp): 195000 VLP:WC ratio: Inf
Whole-community read coverage
VLP-fraction read coverage
Contig position (bp)
CAG-917 CAG-475

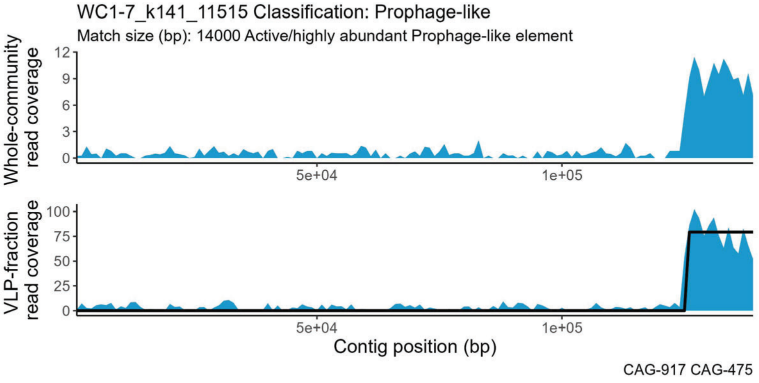
WC1-7_k141_11515 Classification: Prophage-like
Match size (bp): 14000 Active/highly abundant Prophage-like element
Whole-community read coverage
VLP-fraction read coverage
Contig position (bp)
CAG-917 CAG-475

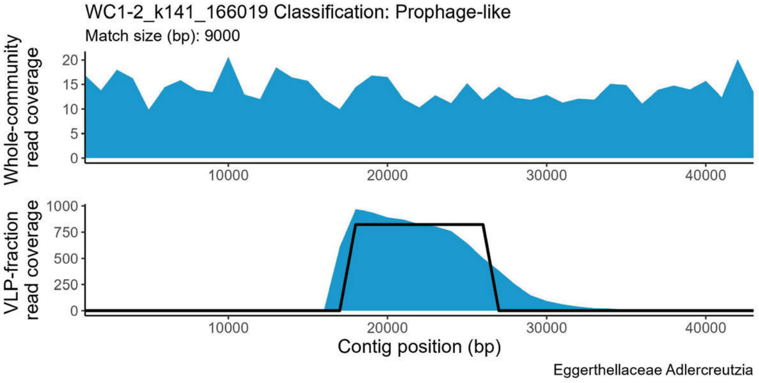
WC1-2_k141_166019 Classification: Prophage-like
Match size (bp): 9000
Whole-community read coverage
VLP-fraction read coverage
Contig position (bp)
Eggerthellaceae Adlercreutzia

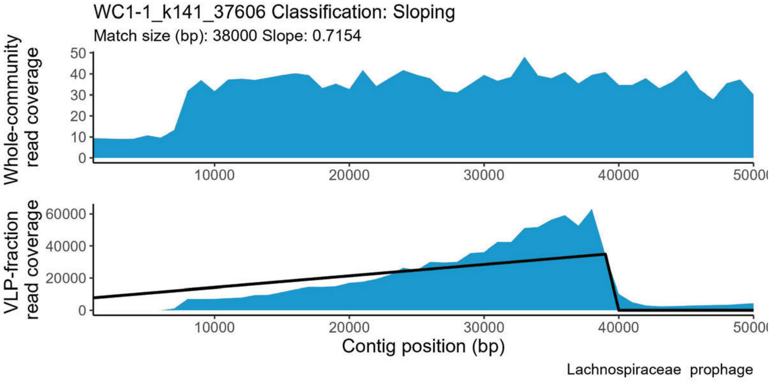
WC1-1_k141_37606 Classification: Sloping
Match size (bp): 38000 Slope: 0.7154
Whole-community read coverage
VLP-fraction read coverage
Contig position (bp)
Lachnospiraceae prophage

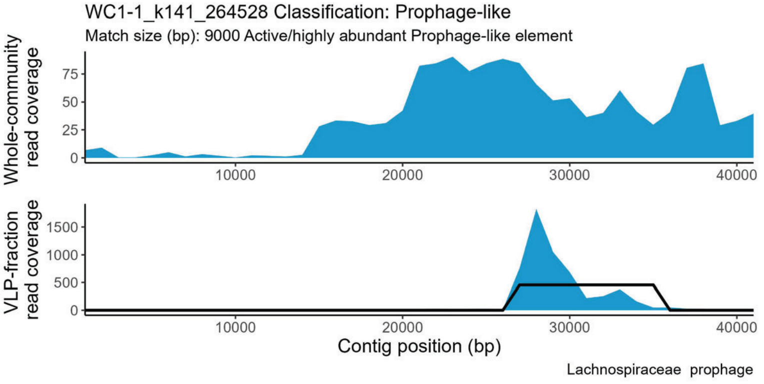
WC1-1_k141_264528 Classification: Prophage-like
Match size (bp): 9000 Active/highly abundant Prophage-like element
Whole-community read coverage
VLP-fraction read coverage
Contig position (bp)
Lachnospiraceae prophage

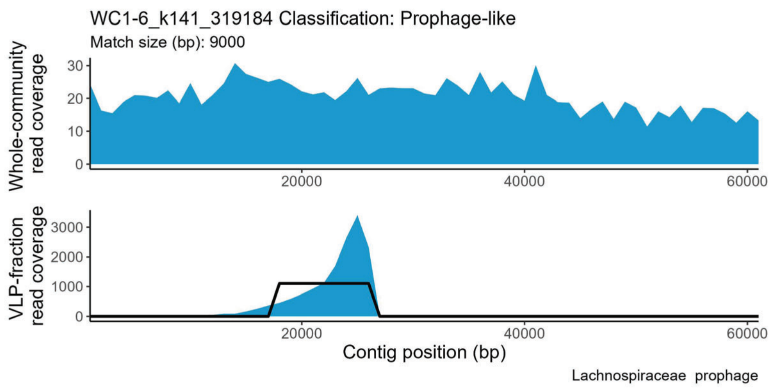
WC1-6_k141_319184 Classification: Prophage-like
Match size (bp): 9000
Whole-community read coverage
VLP-fraction read coverage
Contig position (bp)
Lachnospiraceae prophage

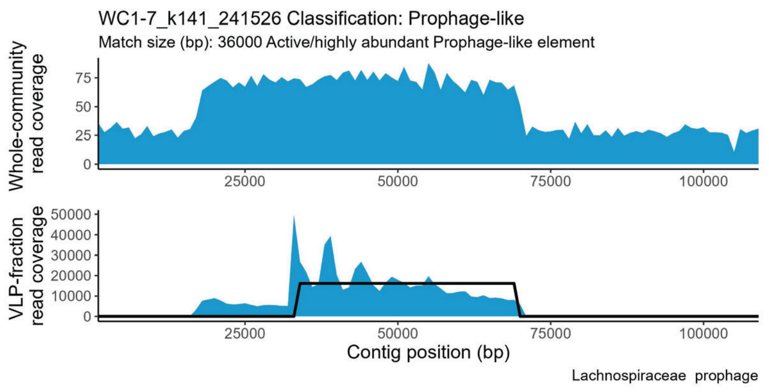
WC1-7_k141_241526 Classification: Prophage-like
Match size (bp): 36000 Active/highly abundant Prophage-like element
Whole-community read coverage
VLP-fraction read coverage
Contig position (bp)
Lachnospiraceae prophage

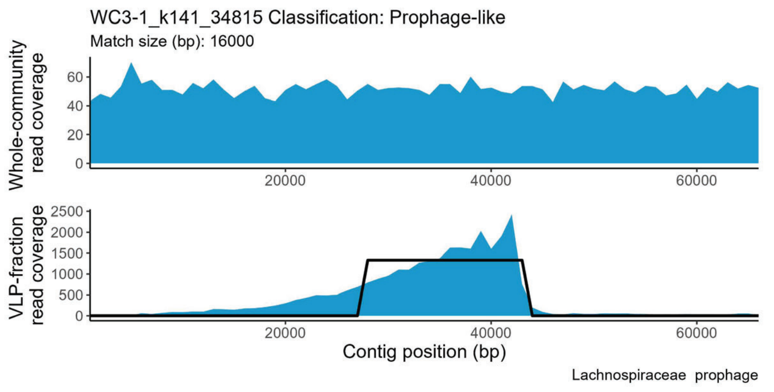
WC3-1_k141_34815 Classification: Prophage-like
Match size (bp): 16000
Whole-community read coverage
VLP-fraction read coverage
Contig position (bp)
Lachnospiraceae prophage

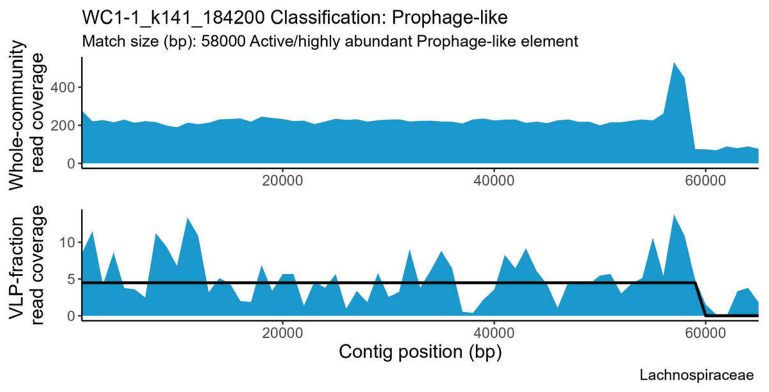
WC1-1_k141_184200 Classification: Prophage-like
Match size (bp): 58000 Active/highly abundant Prophage-like element
Whole-community read coverage
VLP-fraction read coverage
Contig position (bp)
Lachnospiraceae

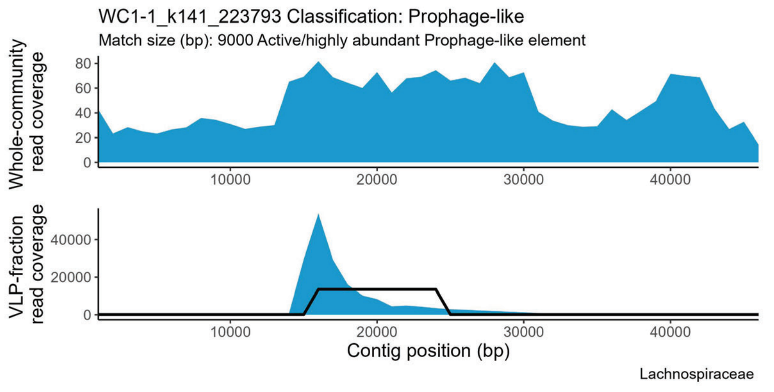
WC1-1_k141_223793 Classification: Prophage-like
Match size (bp): 9000 Active/highly abundant Prophage-like element
Whole-community read coverage
VLP-fraction read coverage
Contig position (bp)
Lachnospiraceae

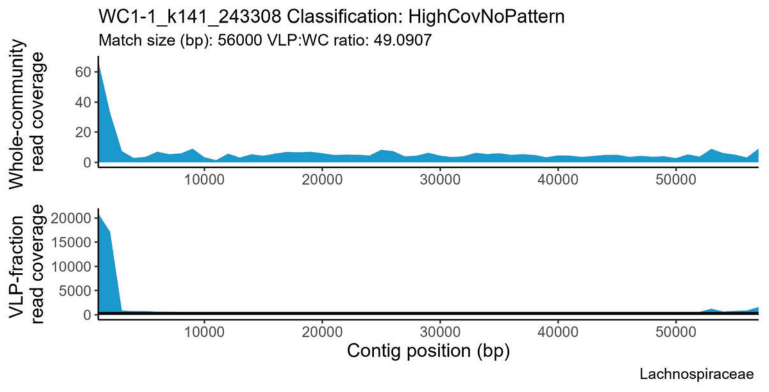
WC1-1_k141_243308 Classification: HighCovNoPattern
Match size (bp): 56000 VLP:WC ratio: 49.0907
Whole-community read coverage
VLP-fraction read coverage
Contig position (bp)
Lachnospiraceae

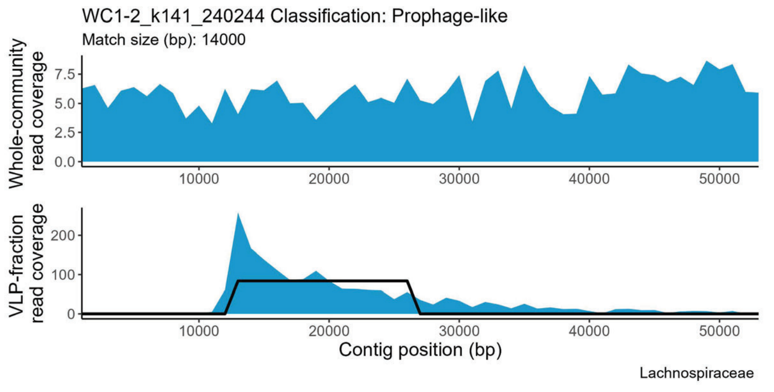
WC1-2_k141_240244 Classification: Prophage-like
Match size (bp): 14000
Whole-community read coverage
VLP-fraction read coverage
Contig position (bp)
Lachnospiraceae

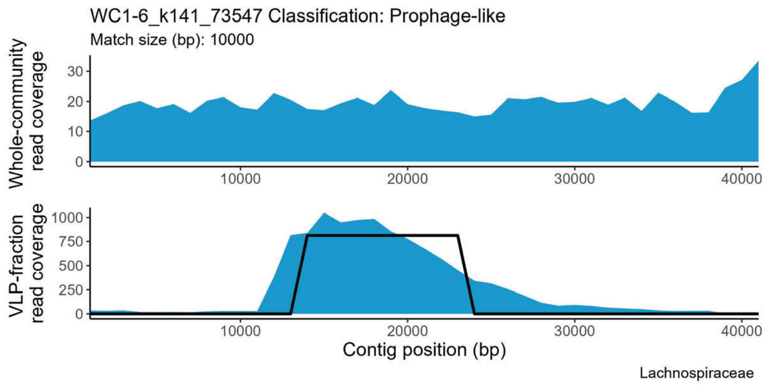
WC1-6_k141_73547 Classification: Prophage-like
Match size (bp): 10000
Whole-community read coverage
VLP-fraction read coverage
Contig position (bp)
Lachnospiraceae

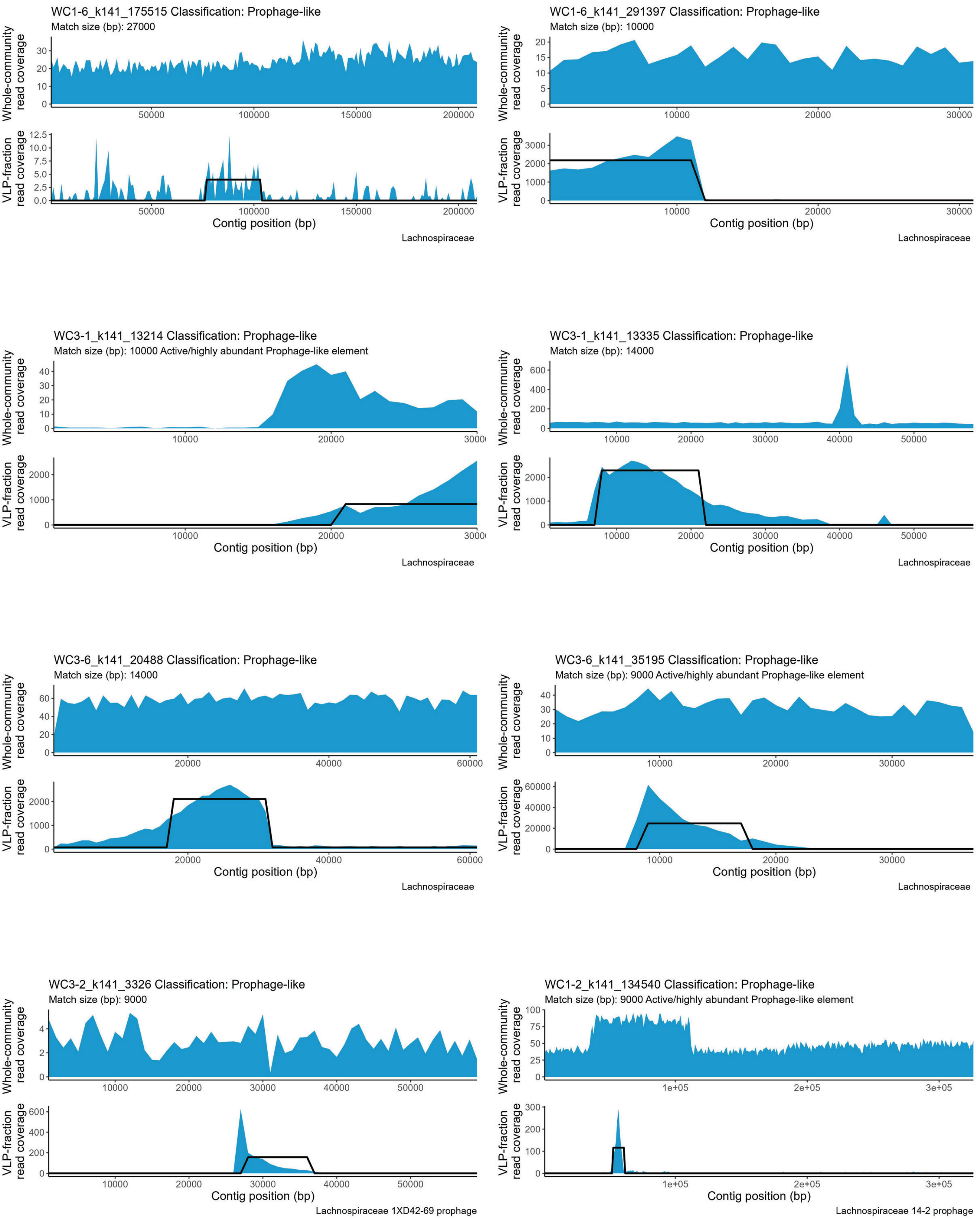
WC1-6_k141_175515 Classification: Prophage-like
Match size (bp): 27000
Whole-community read coverage
VLP-fraction read coverage
Contig position (bp)
Lachnospiraceae
WC1-6_k141_291397 Classification: Prophage-like
Match size (bp): 10000
Whole-community read coverage
VLP-fraction read coverage
Contig position (bp)
Lachnospiraceae
WC3-1_k141_13214 Classification: Prophage-like
Match size (bp): 10000 Active/highly abundant Prophage-like element
Whole-community read coverage
VLP-fraction read coverage
Contig position (bp)
Lachnospiraceae
WC3-1_k141_13335 Classification: Prophage-like
Match size (bp): 14000
Whole-community read coverage
VLP-fraction read coverage
Contig position (bp)
Lachnospiraceae
WC3-6_k141_20488 Classification: Prophage-like
Match size (bp): 14000
Whole-community read coverage
VLP-fraction read coverage
Contig position (bp)
Lachnospiraceae
WC3-6_k141_35195 Classification: Prophage-like
Match size (bp): 9000 Active/highly abundant Prophage-like element
Whole-community read coverage
VLP-fraction read coverage
Contig position (bp)
Lachnospiraceae
WC3-2_k141_3326 Classification: Prophage-like
Match size (bp): 9000
Whole-community read coverage
VLP-fraction read coverage
Contig position (bp)
Lachnospiraceae 1XD42-69 prophage
WC1-2_k141_134540 Classification: Prophage-like
Match size (bp): 9000 Active/highly abundant Prophage-like element
Whole-community read coverage
VLP-fraction read coverage
Contig position (bp)
Lachnospiraceae 14-2 prophage

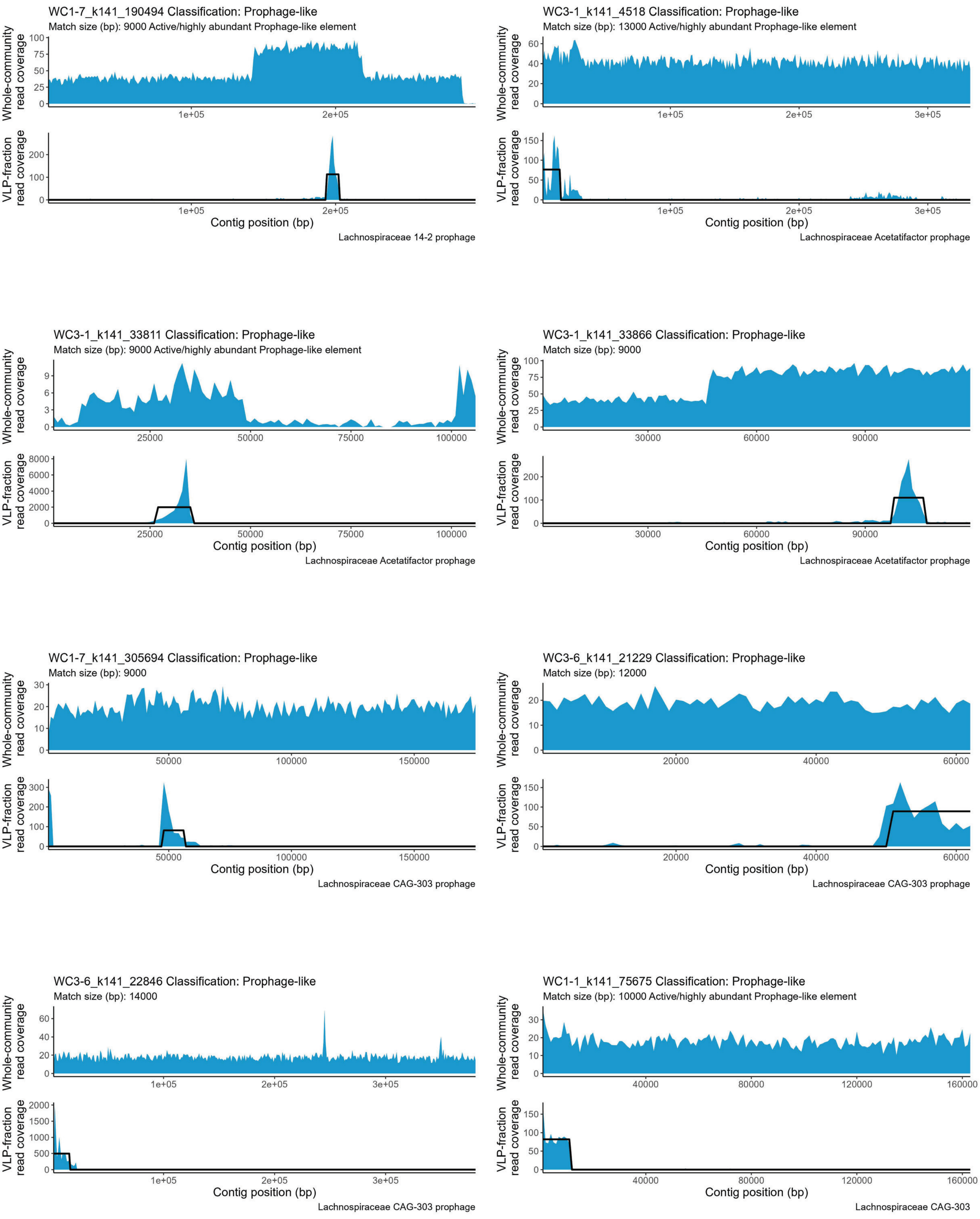

WC1-7_k141_190494 Classification: Prophage-like
Match size (bp): 9000 Active/highly abundant Prophage-like element
Whole-community read coverage
VLP-fraction read coverage
Contig position (bp)
Lachnospiraceae 14-2 prophage
WC3-1_k141_4518 Classification: Prophage-like
Match size (bp): 13000 Active/highly abundant Prophage-like element
Whole-community read coverage
VLP-fraction read coverage
Contig position (bp)
Lachnospiraceae Acetatifactor prophage
WC3-1_k141_33811 Classification: Prophage-like
Match size (bp): 9000 Active/highly abundant Prophage-like element
Whole-community read coverage
VLP-fraction read coverage
Contig position (bp)
Lachnospiraceae Acetatifactor prophage
WC3-1_k141_33866 Classification: Prophage-like
Match size (bp): 9000
Whole-community read coverage
VLP-fraction read coverage
Contig position (bp)
Lachnospiraceae Acetatifactor prophage
WC1-7_k141_305694 Classification: Prophage-like
Match size (bp): 9000
Whole-community read coverage
VLP-fraction read coverage
Contig position (bp)
Lachnospiraceae CAG-303 prophage
WC3-6_k141_21229 Classification: Prophage-like
Match size (bp): 12000
Whole-community read coverage
VLP-fraction read coverage
Contig position (bp)
Lachnospiraceae CAG-303 prophage
WC3-6_k141_22846 Classification: Prophage-like
Match size (bp): 14000
Whole-community read coverage
VLP-fraction read coverage
Contig position (bp)
Lachnospiraceae CAG-303 prophage
WC1-1_k141_75675 Classification: Prophage-like
Match size (bp): 10000 Active/highly abundant Prophage-like element
Whole-community read coverage
VLP-fraction read coverage
Contig position (bp)
Lachnospiraceae CAG-303

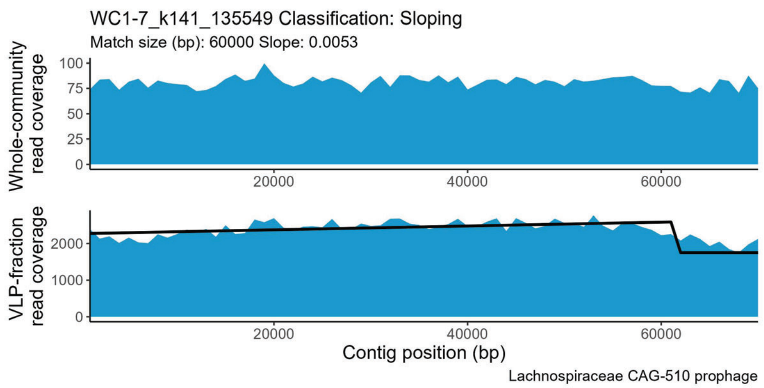
WC1-7_k141_135549 Classification: Sloping
Match size (bp): 60000 Slope: 0.0053
Whole-community read coverage
VLP-fraction read coverage
Contig position (bp)
Lachnospiraceae CAG-510 prophage

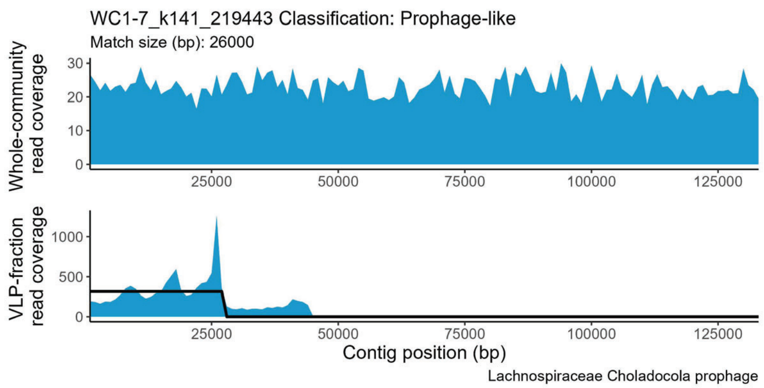
WC1-7_k141_219443 Classification: Prophage-like
Match size (bp): 26000
Whole-community read coverage
VLP-fraction read coverage
Contig position (bp)
Lachnospiraceae Choladocola prophage

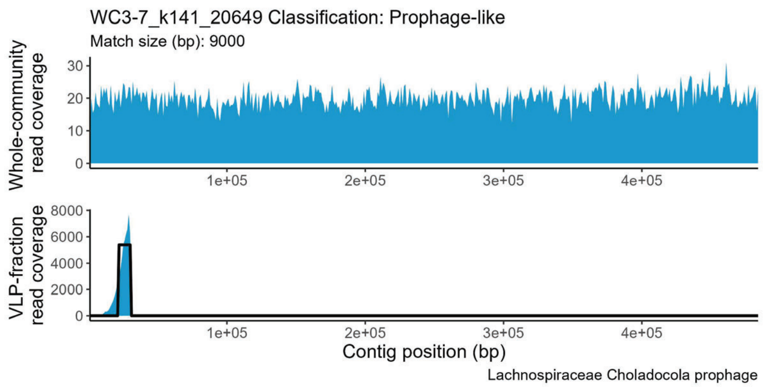
WC3-7_k141_20649 Classification: Prophage-like
Match size (bp): 9000
Whole-community read coverage
VLP-fraction read coverage
Contig position (bp)
Lachnospiraceae Choladocola prophage

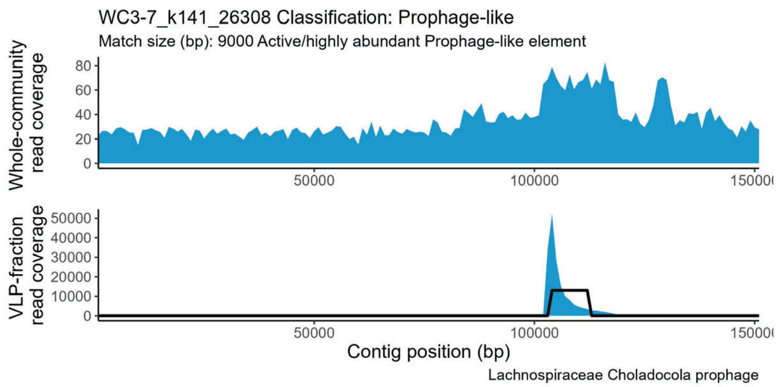
WC3-7_k141_26308 Classification: Prophage-like
Match size (bp): 9000 Active/highly abundant Prophage-like element
Whole-community read coverage
VLP-fraction read coverage
Contig position (bp)
Lachnospiraceae Choladocola prophage

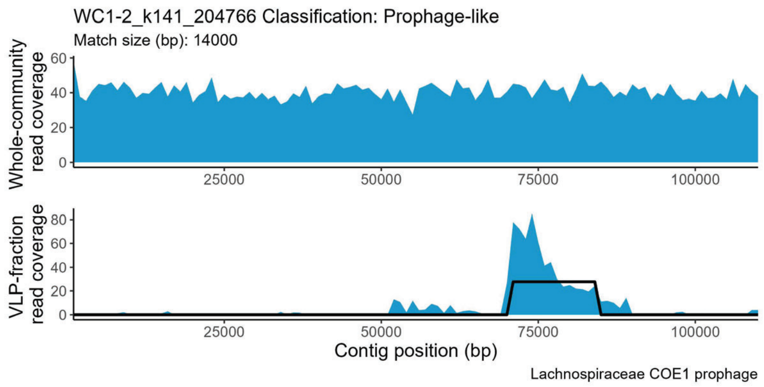
WC1-2_k141_204766 Classification: Prophage-like
Match size (bp): 14000
Whole-community read coverage
VLP-fraction read coverage
Contig position (bp)
Lachnospiraceae COE1 prophage

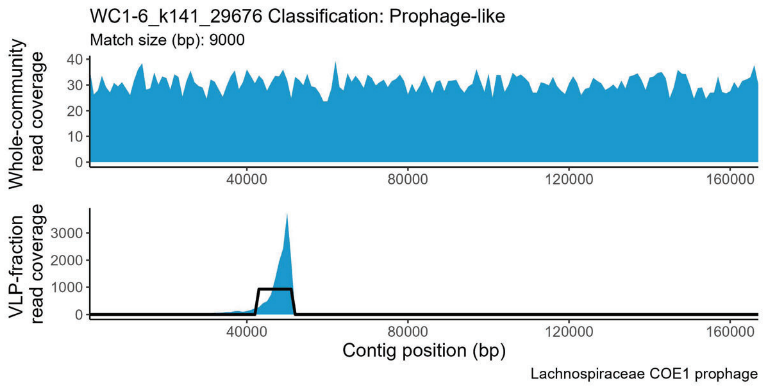
WC1-6_k141_29676 Classification: Prophage-like
Match size (bp): 9000
Whole-community read coverage
VLP-fraction read coverage
Contig position (bp)
Lachnospiraceae COE1 prophage

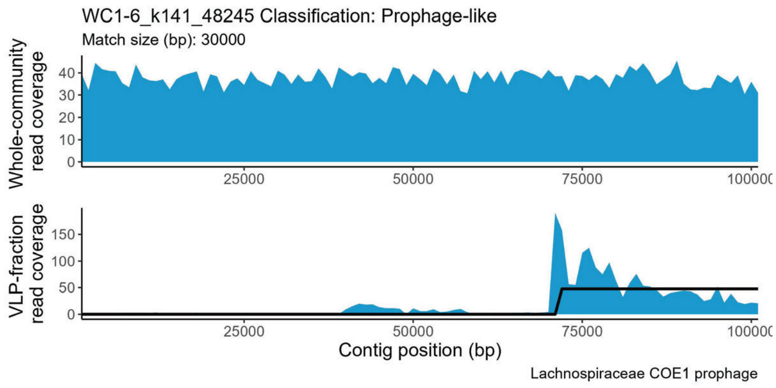
WC1-6_k141_48245 Classification: Prophage-like
Match size (bp): 30000
Whole-community read coverage
VLP-fraction read coverage
Contig position (bp)
Lachnospiraceae COE1 prophage

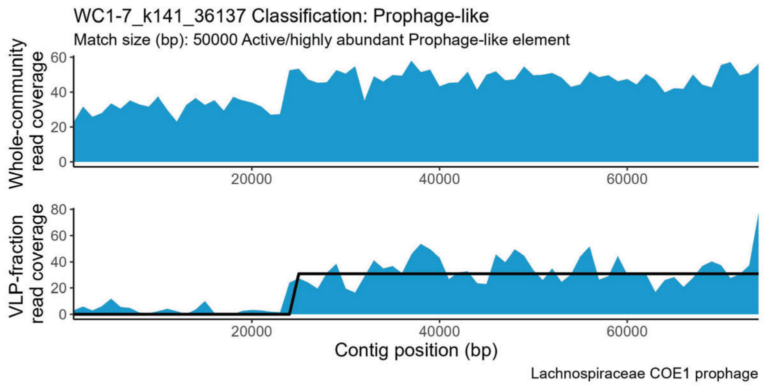
WC1-7_k141_36137 Classification: Prophage-like
Match size (bp): 50000 Active/highly abundant Prophage-like element
Whole-community read coverage
VLP-fraction read coverage
Contig position (bp)
Lachnospiraceae COE1 prophage

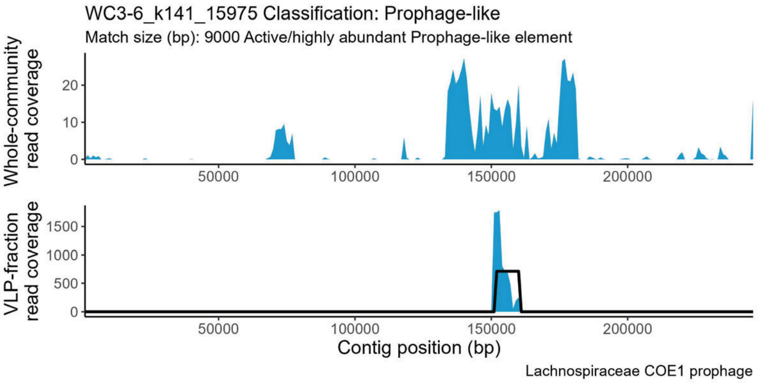
WC3-6_k141_15975 Classification: Prophage-like
Match size (bp): 9000 Active/highly abundant Prophage-like element
Whole-community read coverage
0
10
20
50000
100000
150000
200000
VLP-fraction read coverage
0
500
1000
1500
Contig position (bp)
Lachnospiraceae COE1 prophage

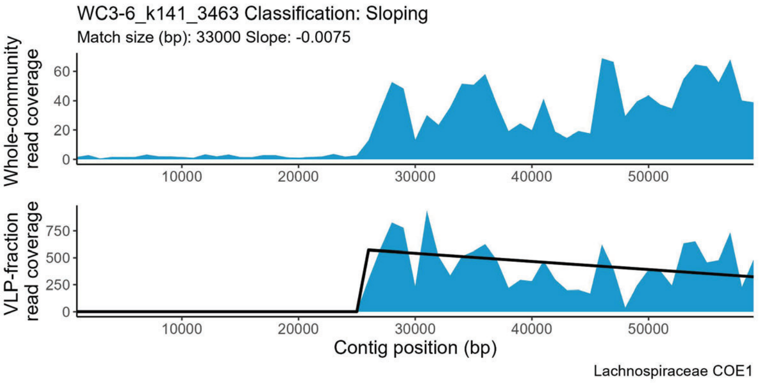
WC3-6_k141_3463 Classification: Sloping
Match size (bp): 33000 Slope: -0.0075
Whole-community read coverage
0
20
40
60
10000
20000
30000
40000
50000
VLP-fraction read coverage
0
250
500
750
Contig position (bp)
Lachnospiraceae COE1

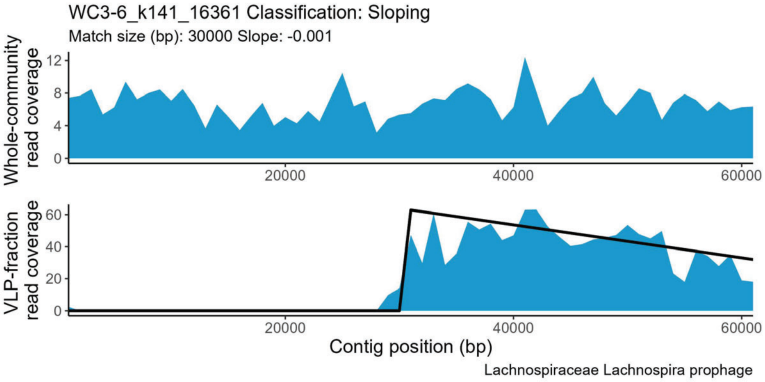
WC3-6_k141_16361 Classification: Sloping
Match size (bp): 30000 Slope: -0.001
Whole-community read coverage
0
4
8
12
20000
40000
60000
VLP-fraction read coverage
0
20
40
60
Contig position (bp)
Lachnospiraceae Lachnospira prophage

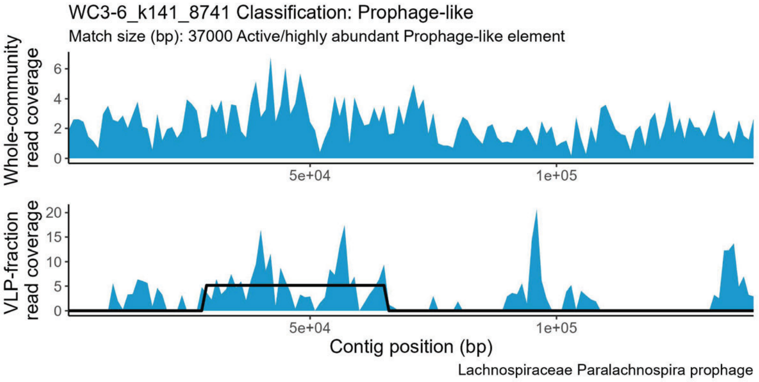
WC3-6_k141_8741 Classification: Prophage-like
Match size (bp): 37000 Active/highly abundant Prophage-like element
Whole-community read coverage
0
2
4
6
5e+04
1e+05
VLP-fraction read coverage
0
5
10
15
20
Contig position (bp)
Lachnospiraceae Paralachnospira prophage

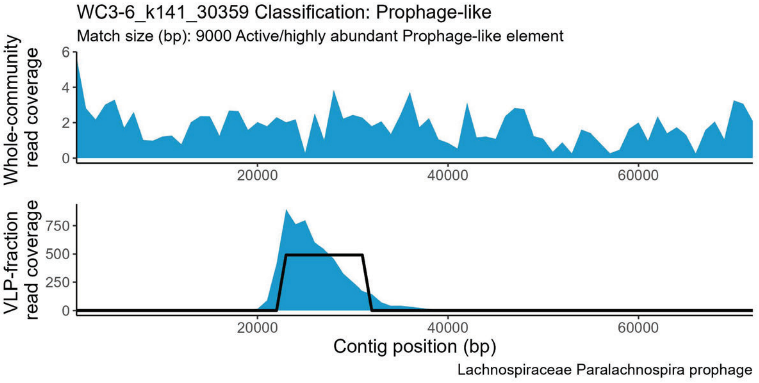
WC3-6_k141_30359 Classification: Prophage-like
Match size (bp): 9000 Active/highly abundant Prophage-like element
Whole-community read coverage
0
2
4
6
20000
40000
60000
VLP-fraction read coverage
0
250
500
750
Contig position (bp)
Lachnospiraceae Paralachnospira prophage

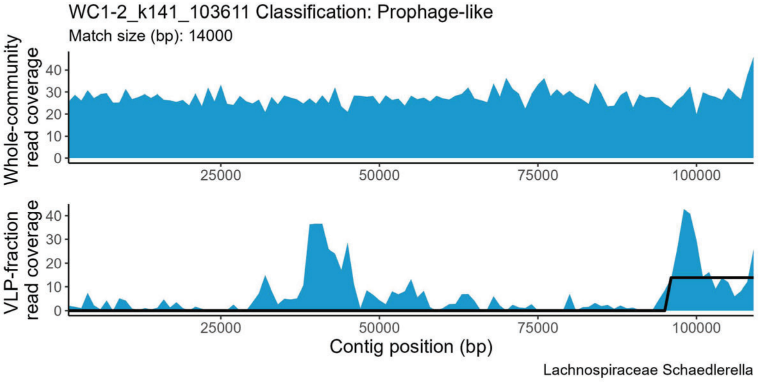
WC1-2_k141_103611 Classification: Prophage-like
Match size (bp): 14000
Whole-community read coverage
0
10
20
30
40
25000
50000
75000
100000
VLP-fraction read coverage
0
10
20
30
40
Contig position (bp)
Lachnospiraceae Schaedlerella

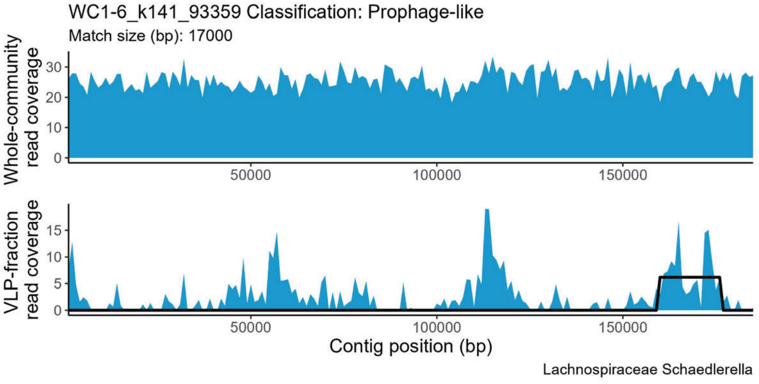
WC1-6_k141_93359 Classification: Prophage-like
Match size (bp): 17000
Whole-community read coverage
0
10
20
30
50000
100000
150000
VLP-fraction read coverage
0
5
10
15
Contig position (bp)
Lachnospiraceae Schaedlerella

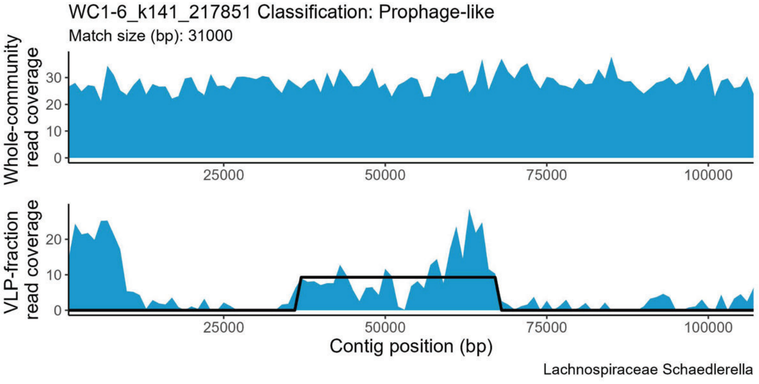
WC1-6_k141_217851 Classification: Prophage-like
Match size (bp): 31000
Whole-community read coverage
0
10
20
30
25000
50000
75000
100000
VLP-fraction read coverage
0
10
20
Contig position (bp)
Lachnospiraceae Schaedlerella

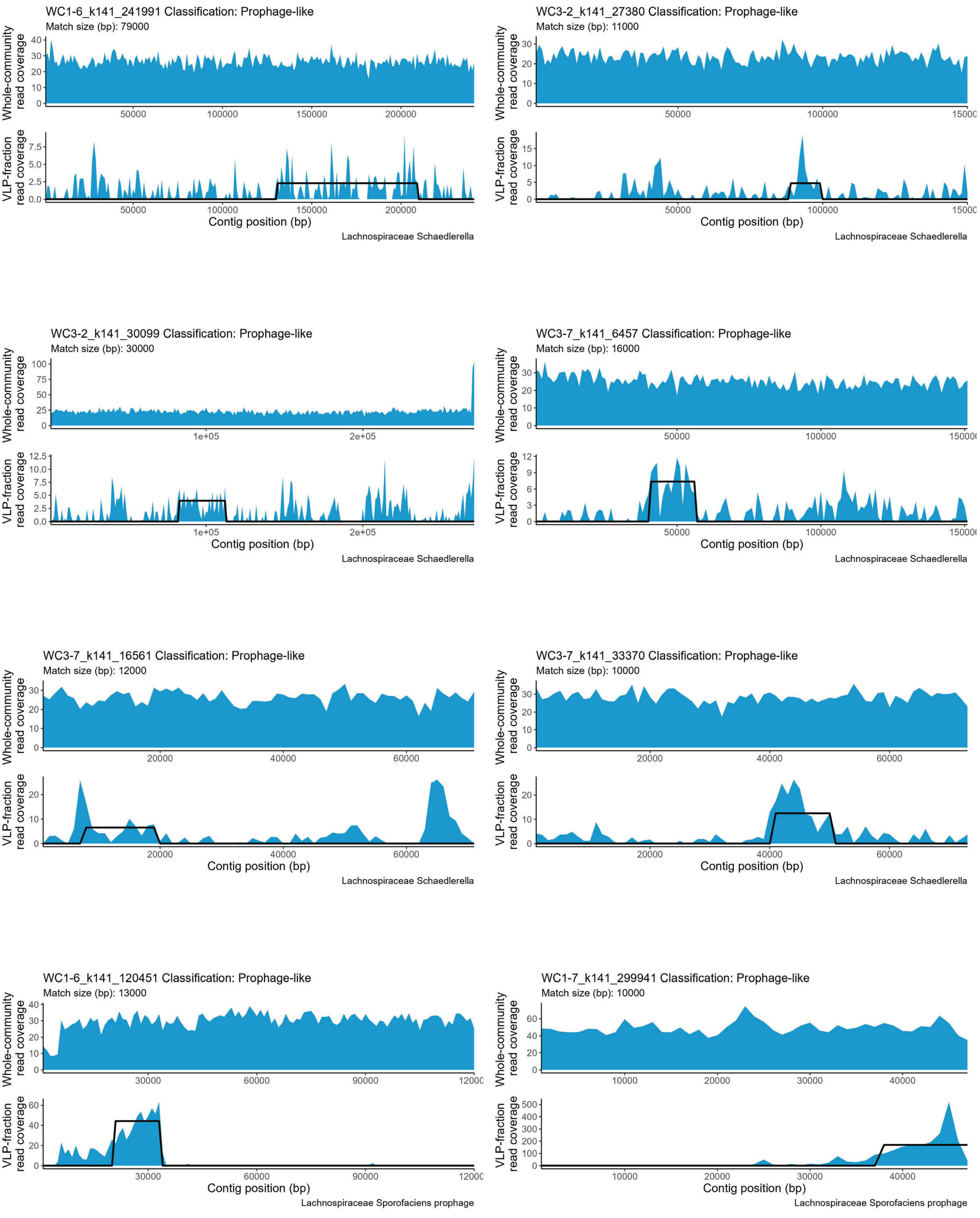

WC1-6_k141_241991 Classification: Prophage-like
Match size (bp): 79000
Whole-community read coverage
VLP-fraction read coverage
Contig position (bp)
Lachnospiraceae Schaedlerella
WC3-2_k141_27380 Classification: Prophage-like
Match size (bp): 11000
Whole-community read coverage
VLP-fraction read coverage
Contig position (bp)
Lachnospiraceae Schaedlerella
WC3-2_k141_30099 Classification: Prophage-like
Match size (bp): 30000
Whole-community read coverage
VLP-fraction read coverage
Contig position (bp)
Lachnospiraceae Schaedlerella
WC3-7_k141_6457 Classification: Prophage-like
Match size (bp): 16000
Whole-community read coverage
VLP-fraction read coverage
Contig position (bp)
Lachnospiraceae Schaedlerella
WC3-7_k141_16561 Classification: Prophage-like
Match size (bp): 12000
Whole-community read coverage
VLP-fraction read coverage
Contig position (bp)
Lachnospiraceae Schaedlerella
WC3-7_k141_33370 Classification: Prophage-like
Match size (bp): 10000
Whole-community read coverage
VLP-fraction read coverage
Contig position (bp)
Lachnospiraceae Schaedlerella
WC1-6_k141_120451 Classification: Prophage-like
Match size (bp): 13000
Whole-community read coverage
VLP-fraction read coverage
Contig position (bp)
Lachnospiraceae Sporofaciens prophage
WC1-7_k141_299941 Classification: Prophage-like
Match size (bp): 10000
Whole-community read coverage
VLP-fraction read coverage
Contig position (bp)
Lachnospiraceae Sporofaciens prophage

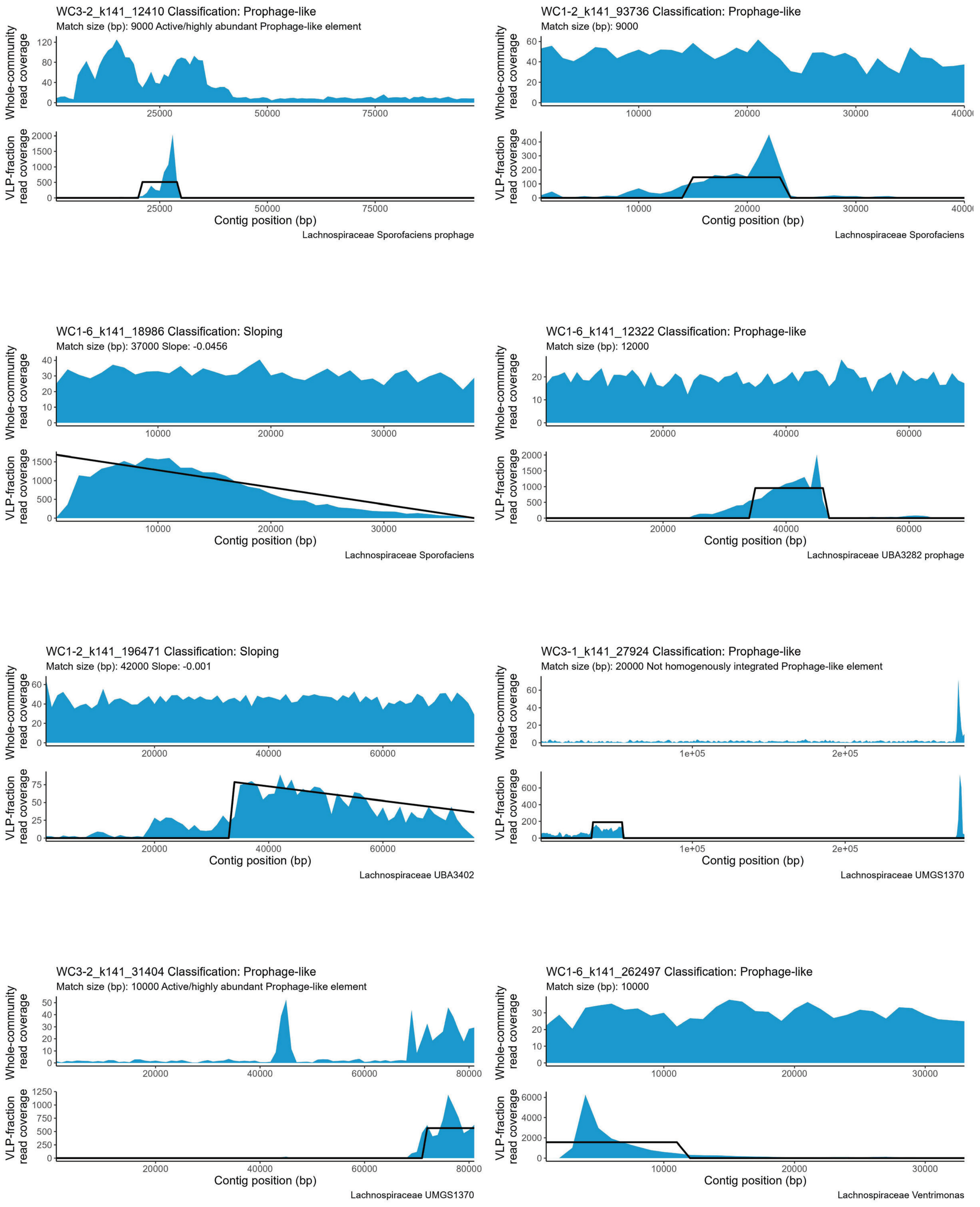
WC3-2_k141_12410 Classification: Prophage-like
Match size (bp): 9000 Active/highly abundant Prophage-like element
Whole-community read coverage
VLP-fraction read coverage
Contig position (bp)
Lachnospiraceae Sporofaciens prophage
WC1-2_k141_93736 Classification: Prophage-like
Match size (bp): 9000
Whole-community read coverage
VLP-fraction read coverage
Contig position (bp)
Lachnospiraceae Sporofaciens
WC1-6_k141_18986 Classification: Sloping
Match size (bp): 37000 Slope: -0.0456
Whole-community read coverage
VLP-fraction read coverage
Contig position (bp)
Lachnospiraceae Sporofaciens
WC1-6_k141_12322 Classification: Prophage-like
Match size (bp): 12000
Whole-community read coverage
VLP-fraction read coverage
Contig position (bp)
Lachnospiraceae UBA3282 prophage
WC1-2_k141_196471 Classification: Sloping
Match size (bp): 42000 Slope: -0.001
Whole-community read coverage
VLP-fraction read coverage
Contig position (bp)
Lachnospiraceae UBA3402
WC3-1_k141_27924 Classification: Prophage-like
Match size (bp): 20000 Not homogenously integrated Prophage-like element
Whole-community read coverage
VLP-fraction read coverage
Contig position (bp)
Lachnospiraceae UMGS1370
WC3-2_k141_31404 Classification: Prophage-like
Match size (bp): 10000 Active/highly abundant Prophage-like element
Whole-community read coverage
VLP-fraction read coverage
Contig position (bp)
Lachnospiraceae UMGS1370
WC1-6_k141_262497 Classification: Prophage-like
Match size (bp): 10000
Whole-community read coverage
VLP-fraction read coverage
Contig position (bp)
Lachnospiraceae Ventrimonas

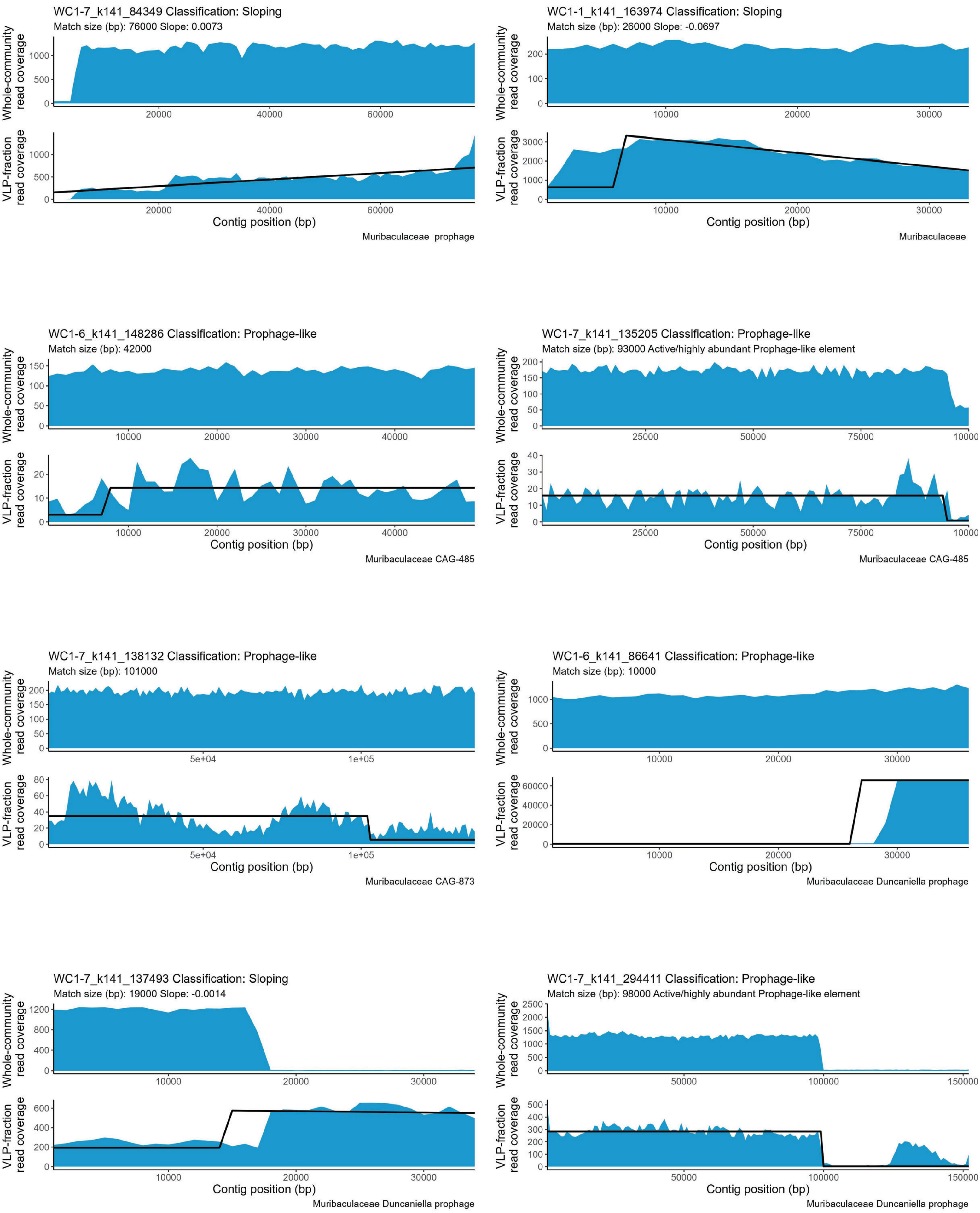

WC1-7_k141_84349 Classification: Sloping
Match size (bp): 76000 Slope: 0.0073
Whole-community read coverage
VLP-fraction read coverage
Contig position (bp)
Muribaculaceae prophage
WC1-1_k141_163974 Classification: Sloping
Match size (bp): 26000 Slope: -0.0697
Whole-community read coverage
VLP-fraction read coverage
Contig position (bp)
Muribaculaceae
WC1-6_k141_148286 Classification: Prophage-like
Match size (bp): 42000
Whole-community read coverage
VLP-fraction read coverage
Contig position (bp)
Muribaculaceae CAG-485
WC1-7_k141_135205 Classification: Prophage-like
Match size (bp): 93000 Active/highly abundant Prophage-like element
Whole-community read coverage
VLP-fraction read coverage
Contig position (bp)
Muribaculaceae CAG-485
WC1-7_k141_138132 Classification: Prophage-like
Match size (bp): 101000
Whole-community read coverage
VLP-fraction read coverage
Contig position (bp)
Muribaculaceae CAG-873
WC1-6_k141_86641 Classification: Prophage-like
Match size (bp): 10000
Whole-community read coverage
VLP-fraction read coverage
Contig position (bp)
Muribaculaceae Duncaniella prophage
WC1-7_k141_137493 Classification: Sloping
Match size (bp): 19000 Slope: -0.0014
Whole-community read coverage
VLP-fraction read coverage
Contig position (bp)
Muribaculaceae Duncaniella prophage
WC1-7_k141_294411 Classification: Prophage-like
Match size (bp): 98000 Active/highly abundant Prophage-like element
Whole-community read coverage
VLP-fraction read coverage
Contig position (bp)
Muribaculaceae Duncaniella prophage

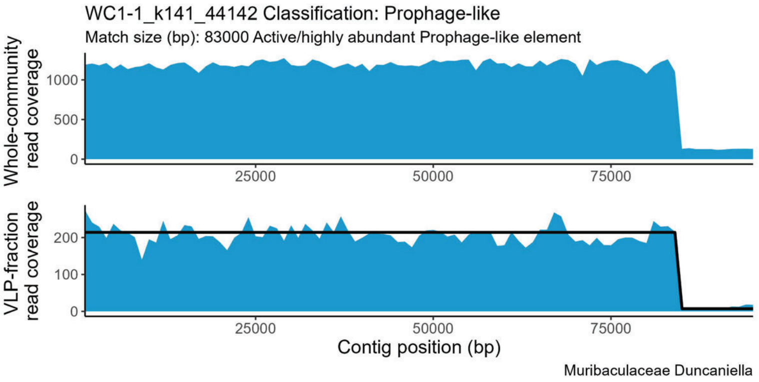
WC1-1_k141_44142 Classification: Prophage-like
Match size (bp): 83000 Active/highly abundant Prophage-like element
Whole-community read coverage
VLP-fraction read coverage
Contig position (bp)
Muribaculaceae Duncaniella

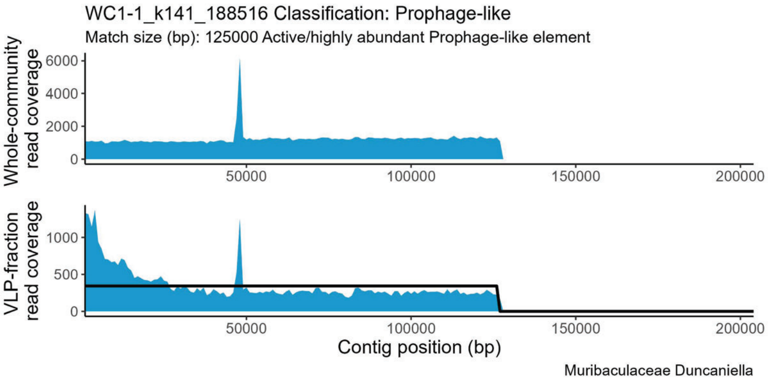
WC1-1_k141_188516 Classification: Prophage-like
Match size (bp): 125000 Active/highly abundant Prophage-like element
Whole-community read coverage
VLP-fraction read coverage
Contig position (bp)
Muribaculaceae Duncaniella

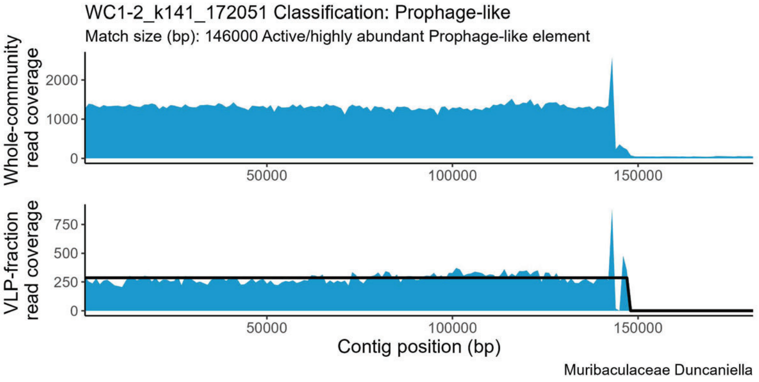
WC1-2_k141_172051 Classification: Prophage-like
Match size (bp): 146000 Active/highly abundant Prophage-like element
Whole-community read coverage
VLP-fraction read coverage
Contig position (bp)
Muribaculaceae Duncaniella

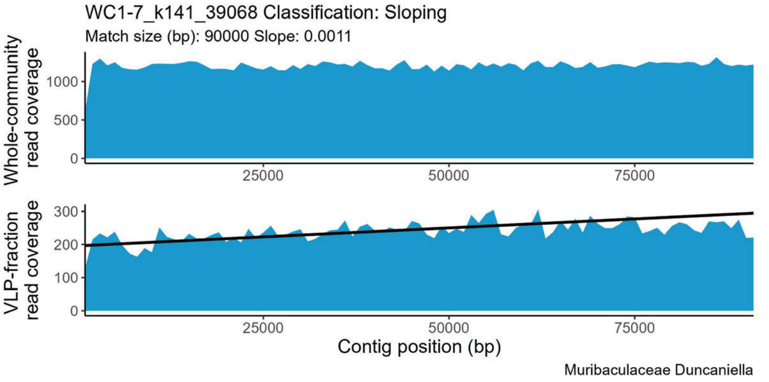
WC1-7_k141_39068 Classification: Sloping
Match size (bp): 90000 Slope: 0.0011
Whole-community read coverage
VLP-fraction read coverage
Contig position (bp)
Muribaculaceae Duncaniella

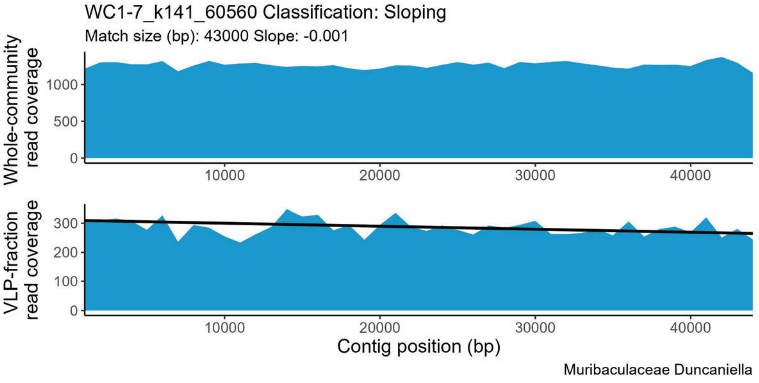
WC1-7_k141_60560 Classification: Sloping
Match size (bp): 43000 Slope: -0.001
Whole-community read coverage
VLP-fraction read coverage
Contig position (bp)
Muribaculaceae Duncaniella

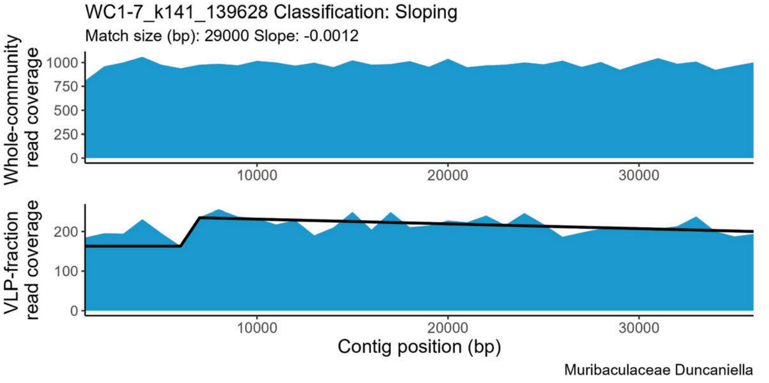
WC1-7_k141_139628 Classification: Sloping
Match size (bp): 29000 Slope: -0.0012
Whole-community read coverage
VLP-fraction read coverage
Contig position (bp)
Muribaculaceae Duncaniella

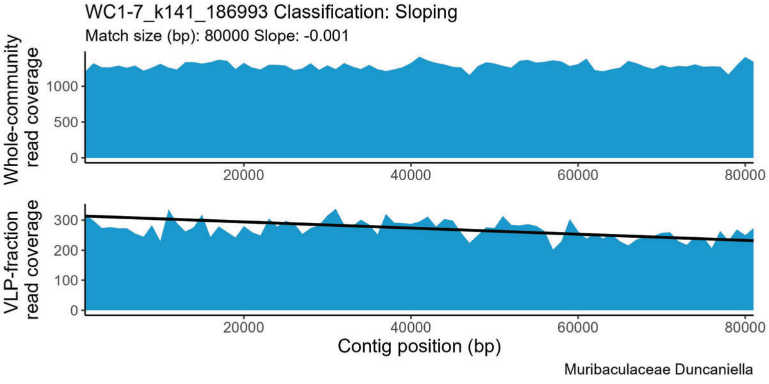
WC1-7_k141_186993 Classification: Sloping
Match size (bp): 80000 Slope: -0.001
Whole-community read coverage
VLP-fraction read coverage
Contig position (bp)
Muribaculaceae Duncaniella

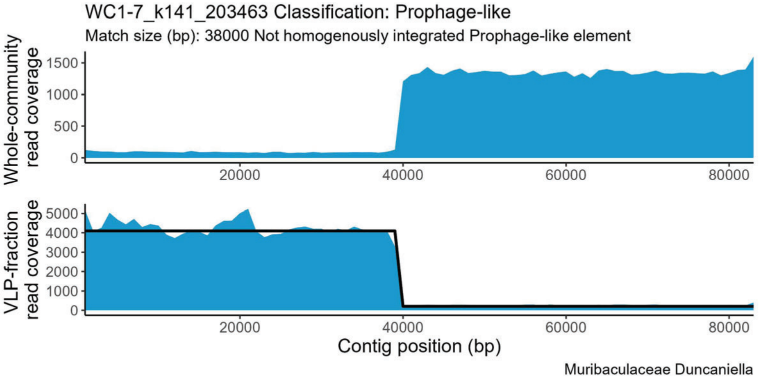
WC1-7_k141_203463 Classification: Prophage-like
Match size (bp): 38000 Not homogenously integrated Prophage-like element
Whole-community read coverage
VLP-fraction read coverage
Contig position (bp)
Muribaculaceae Duncaniella

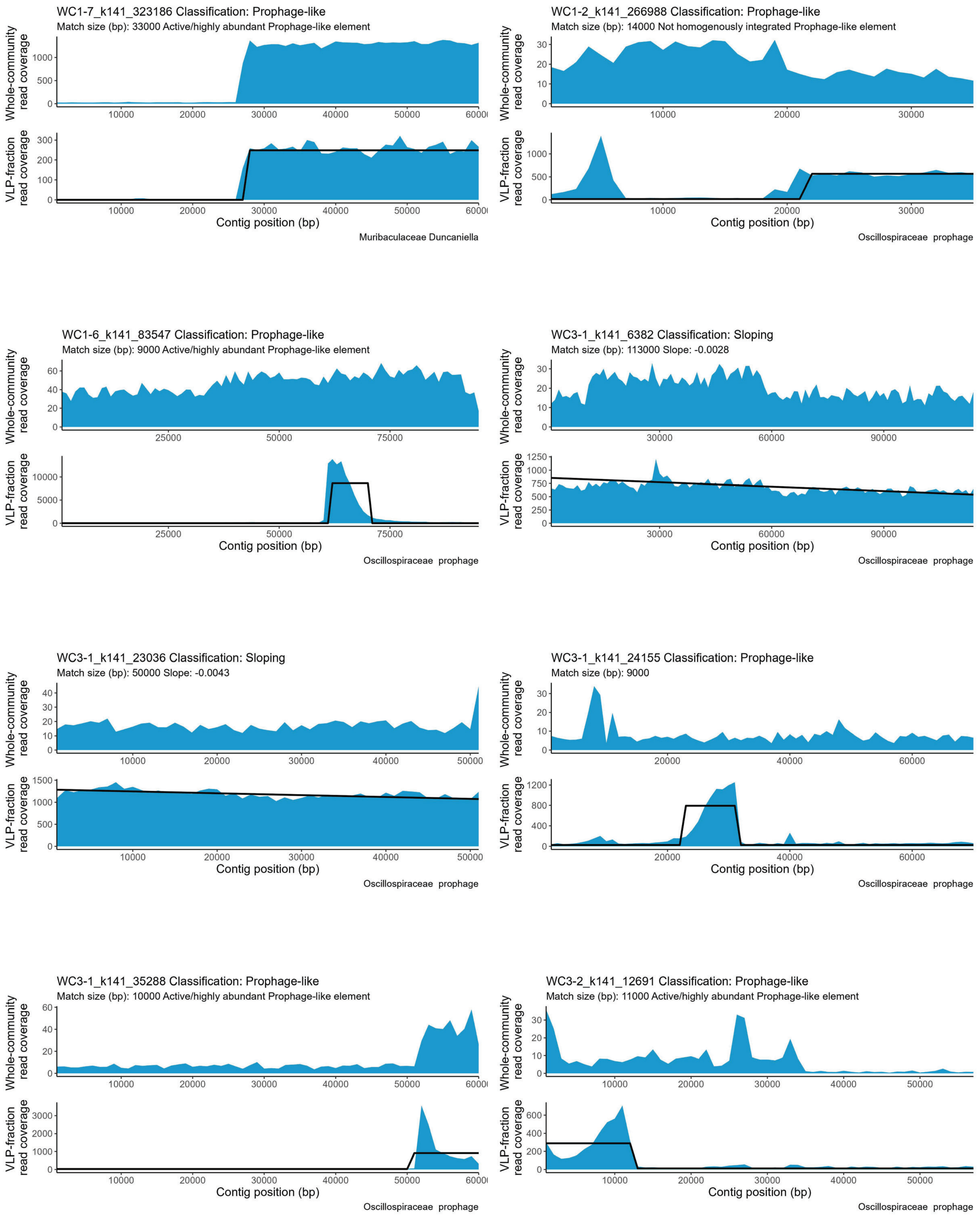
WC1-7_k141_323186 Classification: Prophage-like
Match size (bp): 33000 Active/highly abundant Prophage-like element
Whole-community read coverage
VLP-fraction read coverage
Contig position (bp)
Muribaculaceae Duncaniella
WC1-2_k141_266988 Classification: Prophage-like
Match size (bp): 14000 Not homogenously integrated Prophage-like element
Whole-community read coverage
VLP-fraction read coverage
Contig position (bp)
Oscillospiraceae prophage
WC1-6_k141_83547 Classification: Prophage-like
Match size (bp): 9000 Active/highly abundant Prophage-like element
Whole-community read coverage
VLP-fraction read coverage
Contig position (bp)
Oscillospiraceae prophage
WC3-1_k141_6382 Classification: Sloping
Match size (bp): 113000 Slope: -0.0028
Whole-community read coverage
VLP-fraction read coverage
Contig position (bp)
Oscillospiraceae prophage
WC3-1_k141_23036 Classification: Sloping
Match size (bp): 50000 Slope: -0.0043
Whole-community read coverage
VLP-fraction read coverage
Contig position (bp)
Oscillospiraceae prophage
WC3-1_k141_24155 Classification: Prophage-like
Match size (bp): 9000
Whole-community read coverage
VLP-fraction read coverage
Contig position (bp)
Oscillospiraceae prophage
WC3-1_k141_35288 Classification: Prophage-like
Match size (bp): 10000 Active/highly abundant Prophage-like element
Whole-community read coverage
VLP-fraction read coverage
Contig position (bp)
Oscillospiraceae prophage
WC3-2_k141_12691 Classification: Prophage-like
Match size (bp): 11000 Active/highly abundant Prophage-like element
Whole-community read coverage
VLP-fraction read coverage
Contig position (bp)
Oscillospiraceae prophage

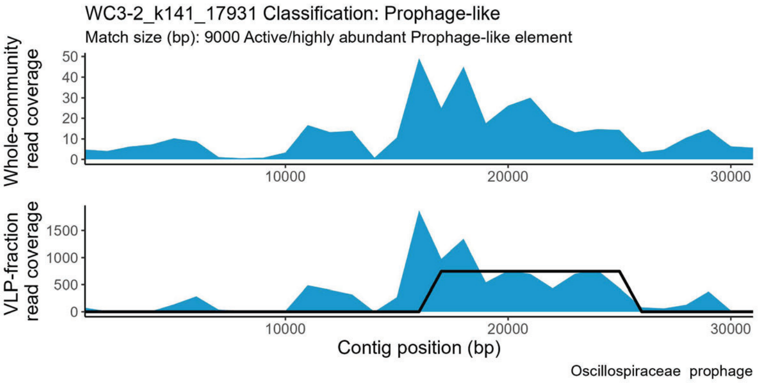
WC3-2_k141_17931 Classification: Prophage-like
Match size (bp): 9000 Active/highly abundant Prophage-like element
Whole-community read coverage
VLP-fraction read coverage
Contig position (bp)
Oscillospiraceae prophage

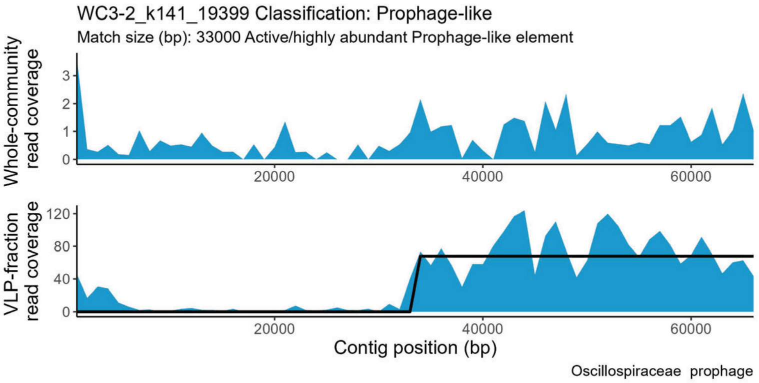
WC3-2_k141_19399 Classification: Prophage-like
Match size (bp): 33000 Active/highly abundant Prophage-like element
Whole-community read coverage
VLP-fraction read coverage
Contig position (bp)
Oscillospiraceae prophage

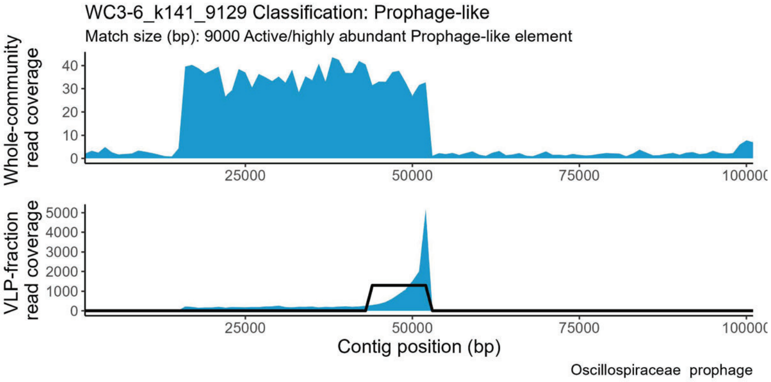
WC3-6_k141_9129 Classification: Prophage-like
Match size (bp): 9000 Active/highly abundant Prophage-like element
Whole-community read coverage
VLP-fraction read coverage
Contig position (bp)
Oscillospiraceae prophage

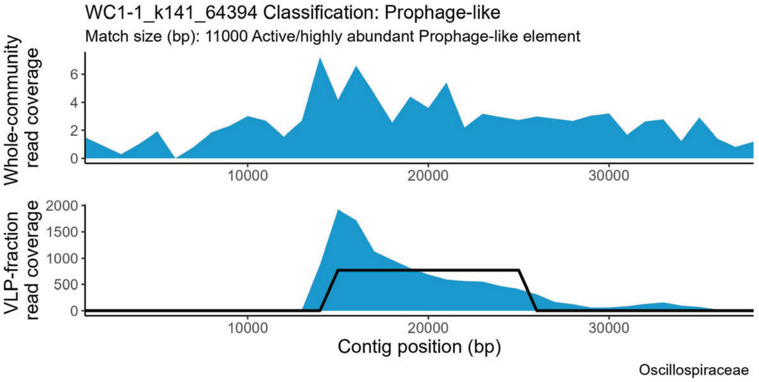
WC1-1_k141_64394 Classification: Prophage-like
Match size (bp): 11000 Active/highly abundant Prophage-like element
Whole-community read coverage
VLP-fraction read coverage
Contig position (bp)
Oscillospiraceae

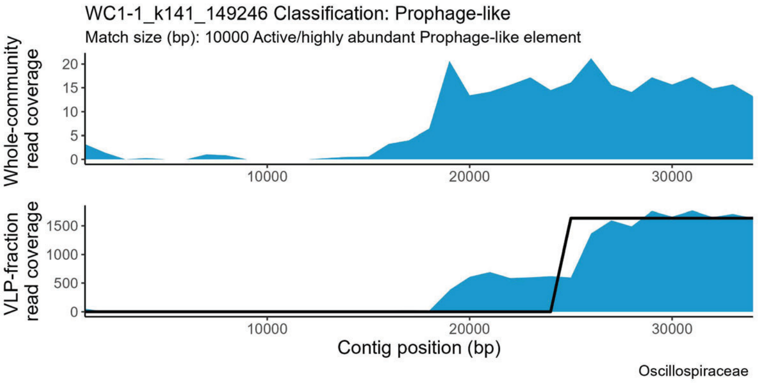
WC1-1_k141_149246 Classification: Prophage-like
Match size (bp): 10000 Active/highly abundant Prophage-like element
Whole-community read coverage
VLP-fraction read coverage
Contig position (bp)
Oscillospiraceae

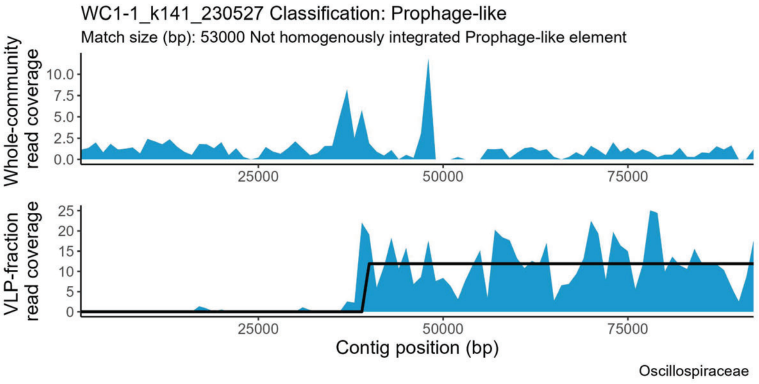
WC1-1_k141_230527 Classification: Prophage-like
Match size (bp): 53000 Not homogenously integrated Prophage-like element
Whole-community read coverage
VLP-fraction read coverage
Contig position (bp)
Oscillospiraceae

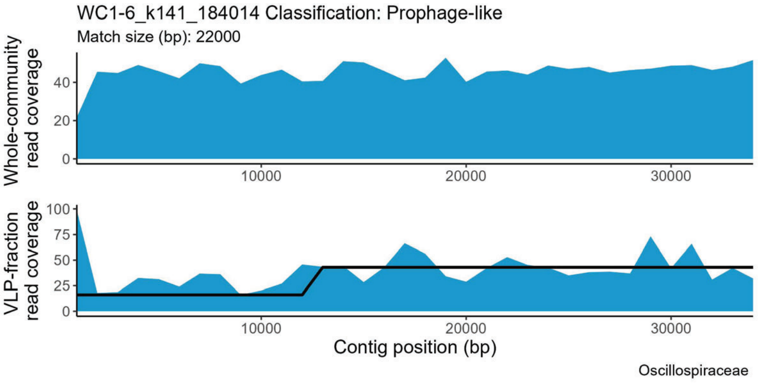
WC1-6_k141_184014 Classification: Prophage-like
Match size (bp): 22000
Whole-community read coverage
VLP-fraction read coverage
Contig position (bp)
Oscillospiraceae

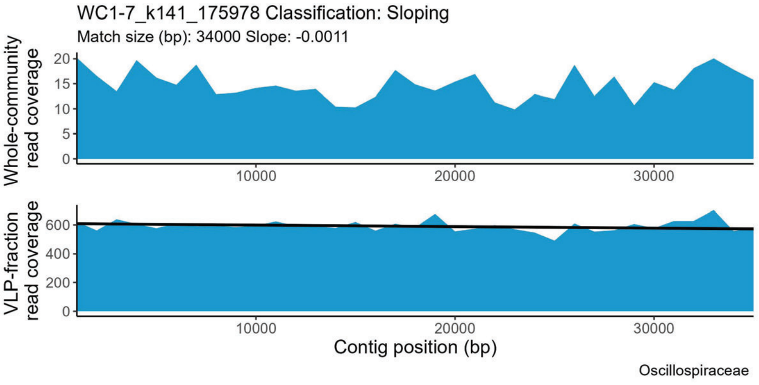
WC1-7_k141_175978 Classification: Sloping
Match size (bp): 34000 Slope: -0.0011
Whole-community read coverage
VLP-fraction read coverage
Contig position (bp)
Oscillospiraceae

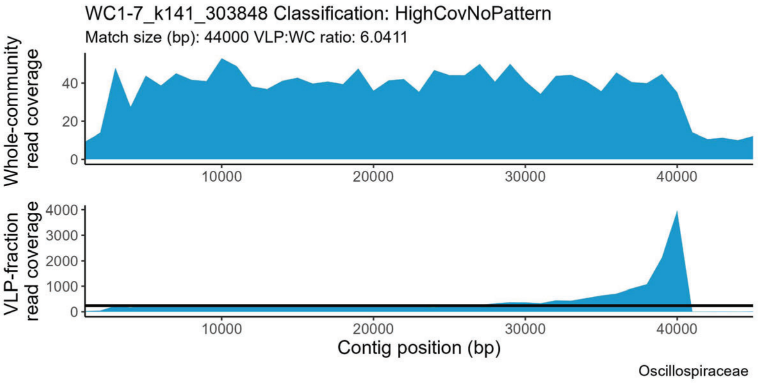
WC1-7_k141_303848 Classification: HighCovNoPattern
Match size (bp): 44000 VLP:WC ratio: 6.0411
Whole-community read coverage
VLP-fraction read coverage
Contig position (bp)
Oscillospiraceae

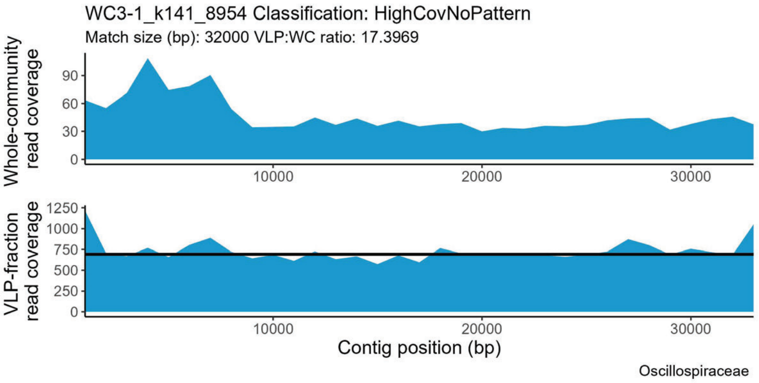
WC3-1_k141_8954 Classification: HighCovNoPattern
Match size (bp): 32000 VLP:WC ratio: 17.3969
Whole-community read coverage
VLP-fraction read coverage
Contig position (bp)
Oscillospiraceae

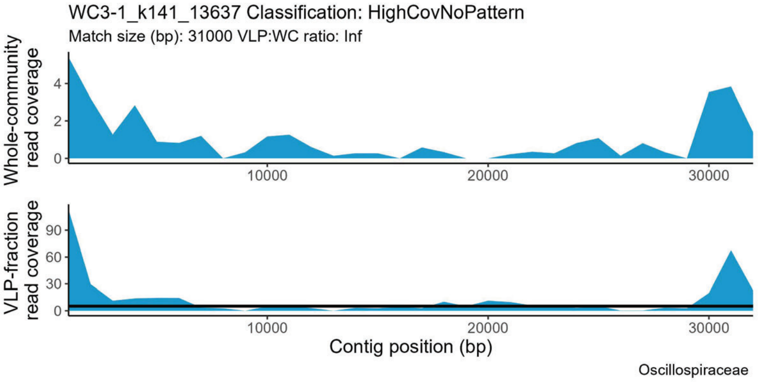
WC3-1_k141_13637 Classification: HighCovNoPattern
Match size (bp): 31000 VLP:WC ratio: Inf
Whole-community read coverage
VLP-fraction read coverage
Contig position (bp)
Oscillospiraceae

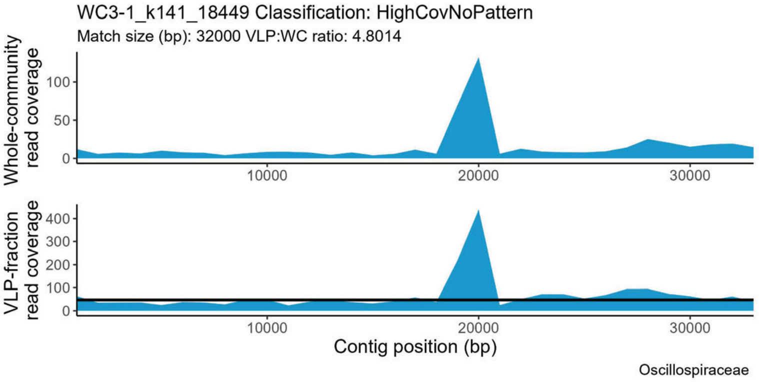
WC3-1_k141_18449 Classification: HighCovNoPattern
Match size (bp): 32000 VLP:WC ratio: 4.8014
Whole-community read coverage
VLP-fraction read coverage
Contig position (bp)
Oscillospiraceae

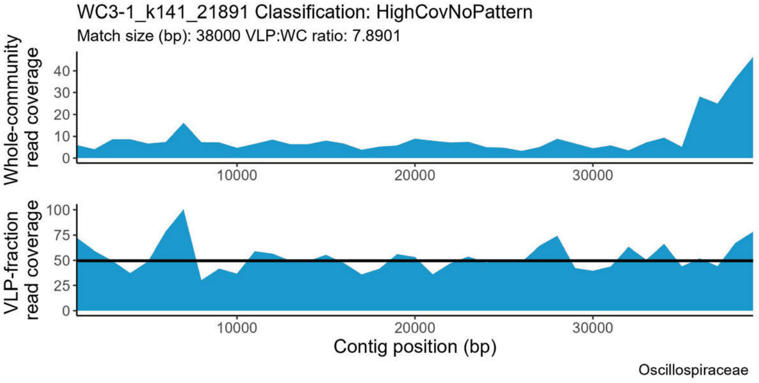
WC3-1_k141_21891 Classification: HighCovNoPattern
Match size (bp): 38000 VLP:WC ratio: 7.8901
Whole-community read coverage
VLP-fraction read coverage
Contig position (bp)
Oscillospiraceae

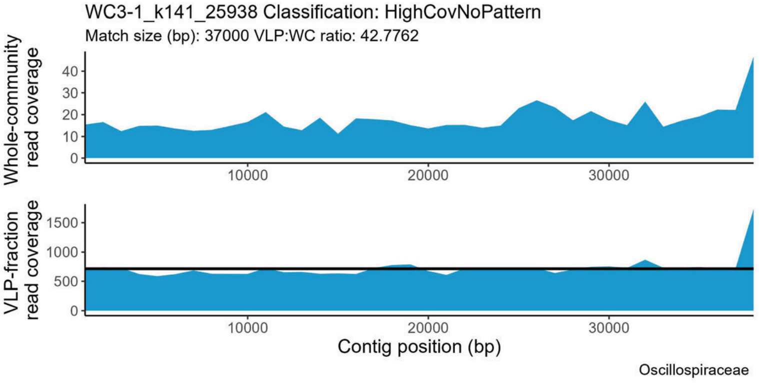
WC3-1_k141_25938 Classification: HighCovNoPattern
Match size (bp): 37000 VLP:WC ratio: 42.7762
Whole-community read coverage
VLP-fraction read coverage
Contig position (bp)
Oscillospiraceae

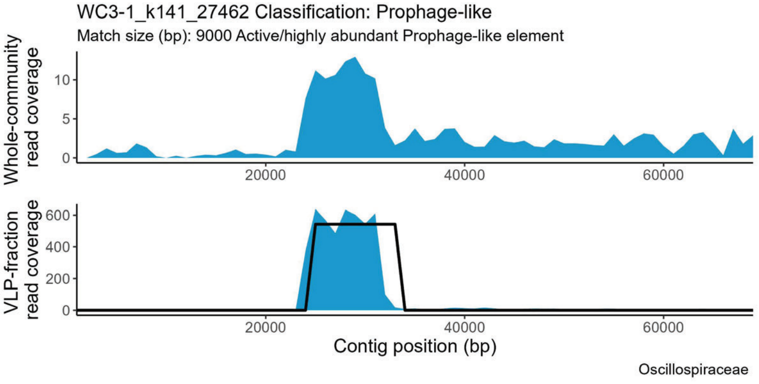
WC3-1_k141_27462 Classification: Prophage-like
Match size (bp): 9000 Active/highly abundant Prophage-like element
Whole-community read coverage
VLP-fraction read coverage
Contig position (bp)
Oscillospiraceae

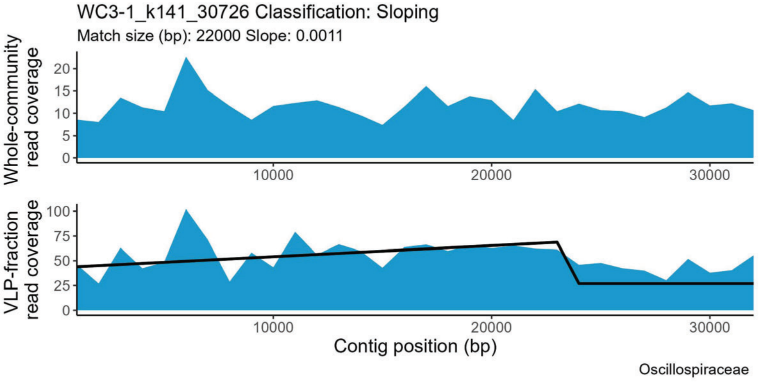
WC3-1_k141_30726 Classification: Sloping
Match size (bp): 22000 Slope: 0.0011
Whole-community read coverage
VLP-fraction read coverage
Contig position (bp)
Oscillospiraceae

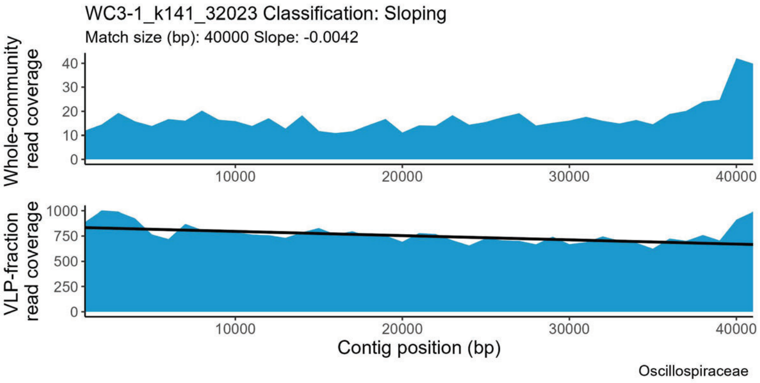
WC3-1_k141_32023 Classification: Sloping
Match size (bp): 40000 Slope: -0.0042
Whole-community read coverage
VLP-fraction read coverage
Contig position (bp)
Oscillospiraceae

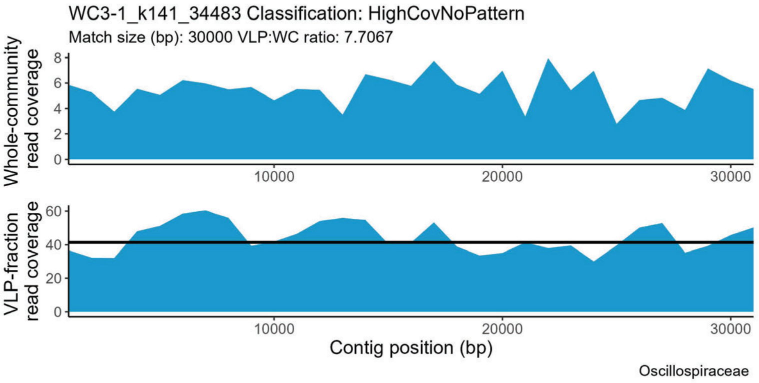
WC3-1_k141_34483 Classification: HighCovNoPattern
Match size (bp): 30000 VLP:WC ratio: 7.7067
Whole-community read coverage
VLP-fraction read coverage
Contig position (bp)
Oscillospiraceae

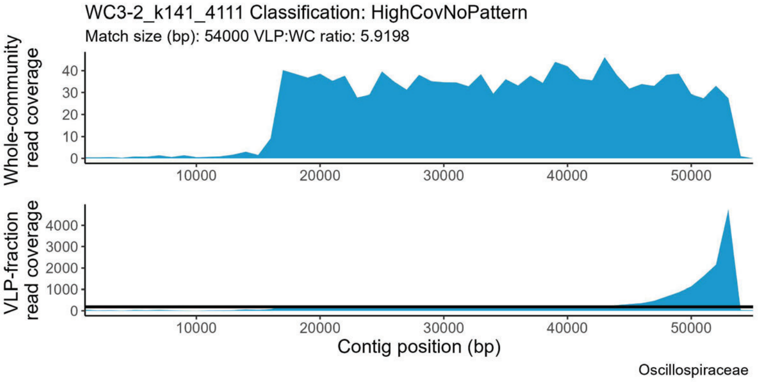
WC3-2_k141_4111 Classification: HighCovNoPattern
Match size (bp): 54000 VLP:WC ratio: 5.9198
Whole-community read coverage
VLP-fraction read coverage
Contig position (bp)
Oscillospiraceae

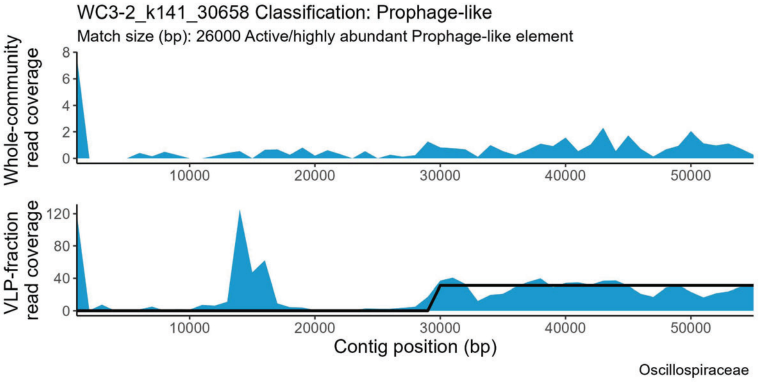
WC3-2_k141_30658 Classification: Prophage-like
Match size (bp): 26000 Active/highly abundant Prophage-like element
Whole-community read coverage
VLP-fraction read coverage
Contig position (bp)
Oscillospiraceae

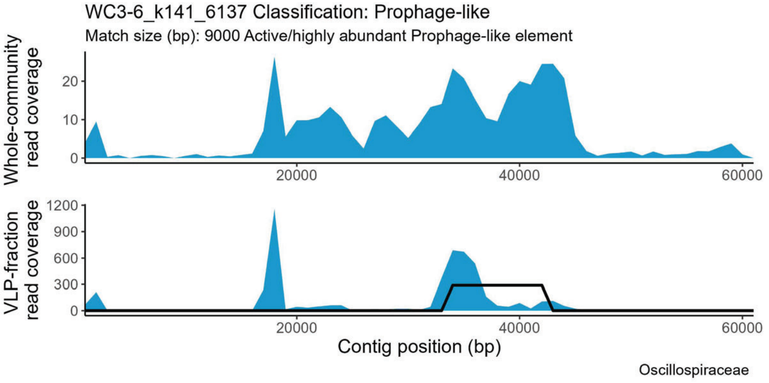
WC3-6_k141_6137 Classification: Prophage-like
Match size (bp): 9000 Active/highly abundant Prophage-like element
Whole-community read coverage
VLP-fraction read coverage
Contig position (bp)
Oscillospiraceae

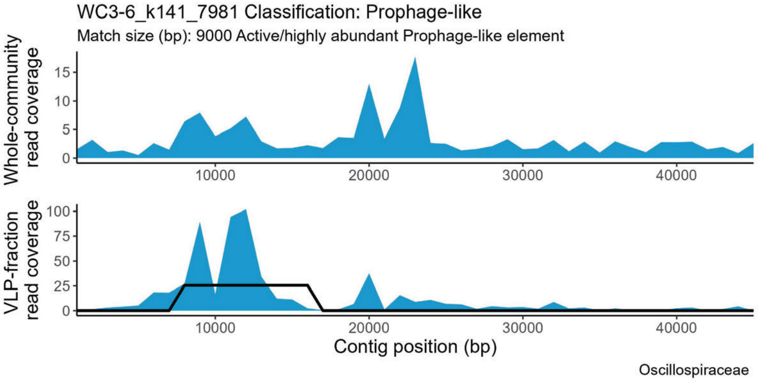
WC3-6_k141_7981 Classification: Prophage-like
Match size (bp): 9000 Active/highly abundant Prophage-like element
Whole-community read coverage
VLP-fraction read coverage
Contig position (bp)
Oscillospiraceae

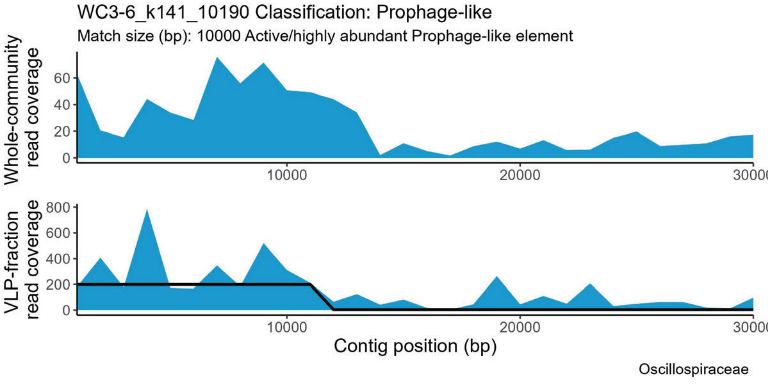
WC3-6_k141_10190 Classification: Prophage-like
Match size (bp): 10000 Active/highly abundant Prophage-like element
Whole-community read coverage
VLP-fraction read coverage
Contig position (bp)
Oscillospiraceae

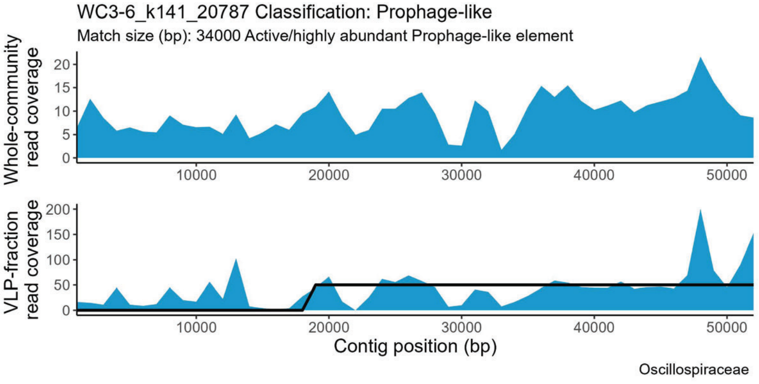
WC3-6_k141_20787 Classification: Prophage-like
Match size (bp): 34000 Active/highly abundant Prophage-like element
Whole-community read coverage
VLP-fraction read coverage
Contig position (bp)
Oscillospiraceae

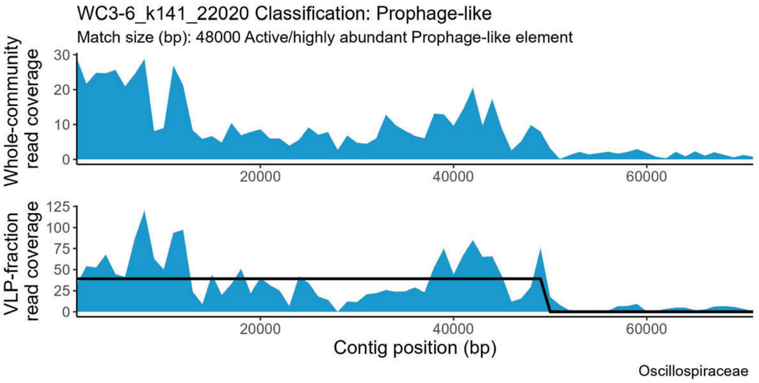
WC3-6_k141_22020 Classification: Prophage-like
Match size (bp): 48000 Active/highly abundant Prophage-like element
Whole-community read coverage
VLP-fraction read coverage
Contig position (bp)
Oscillospiraceae

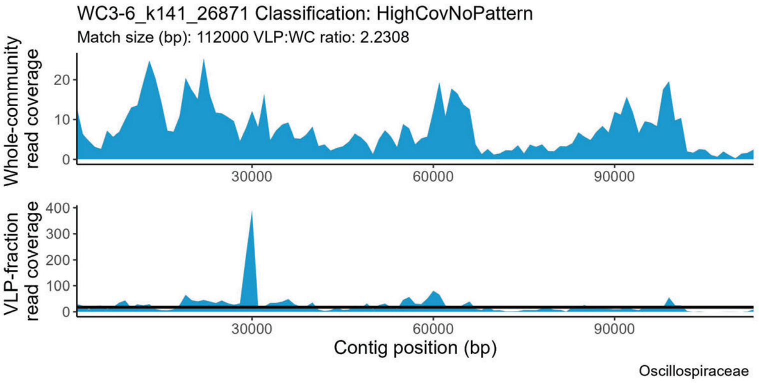
WC3-6_k141_26871 Classification: HighCovNoPattern
Match size (bp): 112000 VLP:WC ratio: 2.2308
Whole-community read coverage
VLP-fraction read coverage
Contig position (bp)
Oscillospiraceae

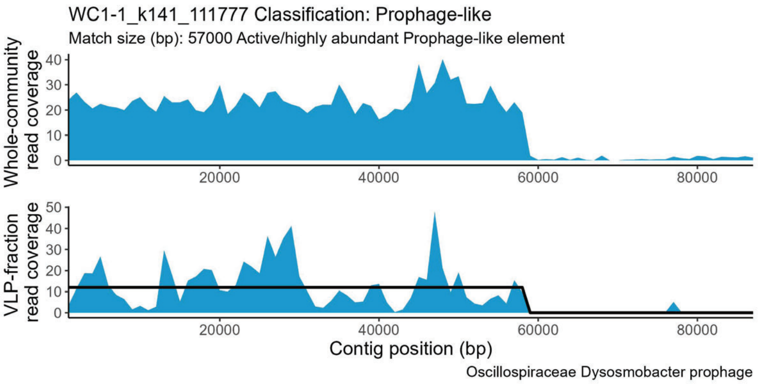
WC1-1_k141_111777 Classification: Prophage-like
Match size (bp): 57000 Active/highly abundant Prophage-like element
Whole-community read coverage
VLP-fraction read coverage
Contig position (bp)
Oscillospiraceae Dysosmobacter prophage

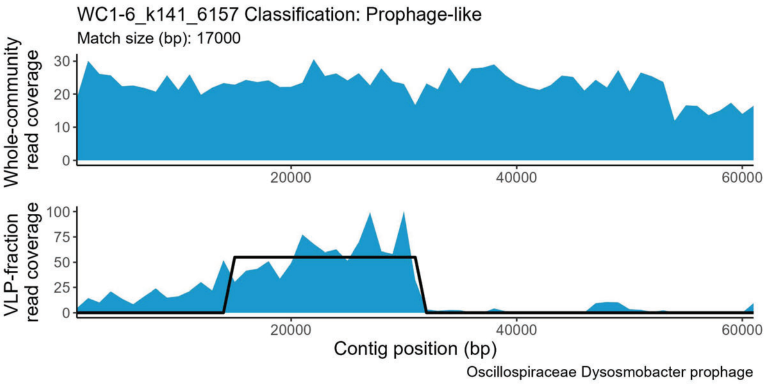
WC1-6_k141_6157 Classification: Prophage-like
Match size (bp): 17000
Whole-community read coverage
VLP-fraction read coverage
Contig position (bp)
Oscillospiraceae Dysosmobacter prophage

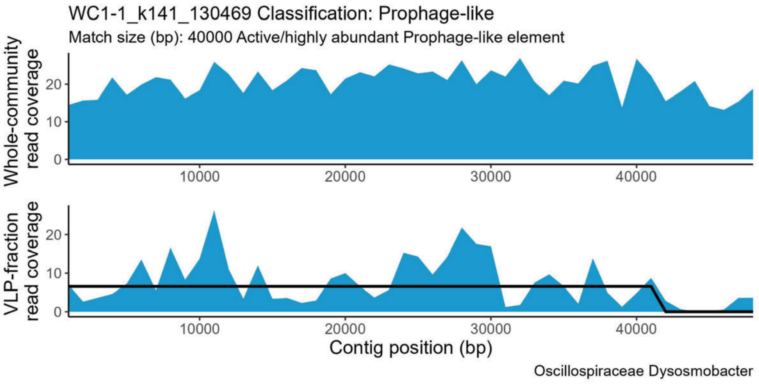
WC1-1_k141_130469 Classification: Prophage-like
Match size (bp): 40000 Active/highly abundant Prophage-like element
Whole-community read coverage
VLP-fraction read coverage
Contig position (bp)
Oscillospiraceae Dysosmobacter

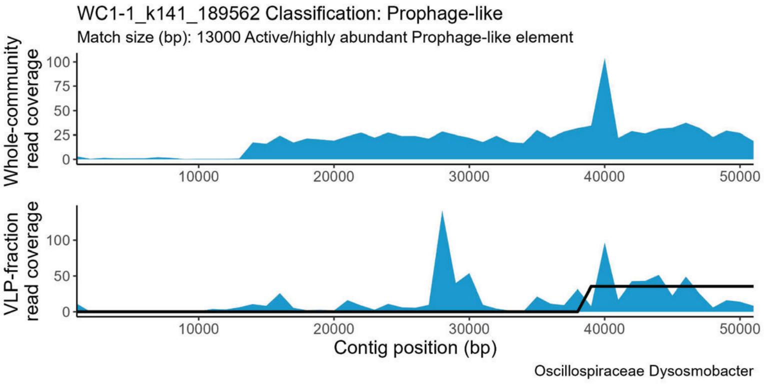
WC1-1_k141_189562 Classification: Prophage-like
Match size (bp): 13000 Active/highly abundant Prophage-like element
Whole-community read coverage
VLP-fraction read coverage
Contig position (bp)
Oscillospiraceae Dysosmobacter

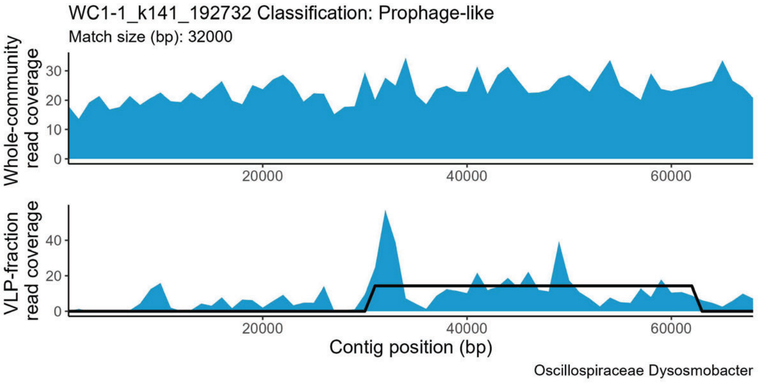
WC1-1_k141_192732 Classification: Prophage-like
Match size (bp): 32000
Whole-community read coverage
VLP-fraction read coverage
Contig position (bp)
Oscillospiraceae Dysosmobacter

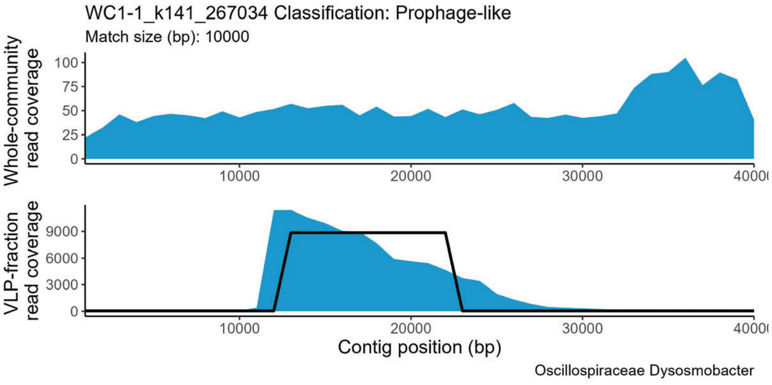
WC1-1_k141_267034 Classification: Prophage-like
Match size (bp): 10000
Whole-community read coverage
VLP-fraction read coverage
Contig position (bp)
Oscillospiraceae Dysosmobacter

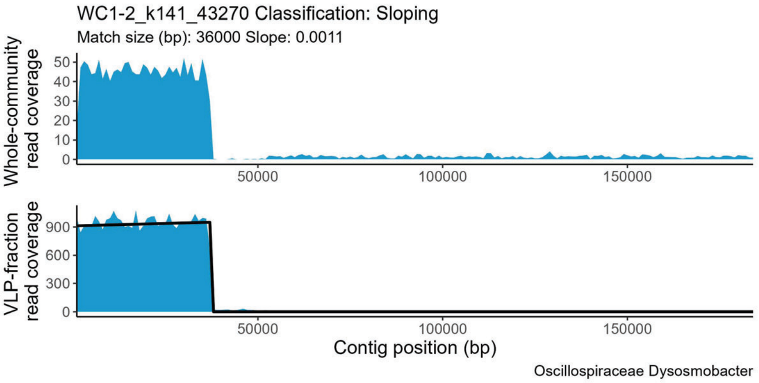
WC1-2_k141_43270 Classification: Sloping
Match size (bp): 36000 Slope: 0.0011
Whole-community read coverage
0
10
20
30
40
50
50000
100000
150000
VLP-fraction read coverage
0
300
600
900
50000
100000
150000
Contig position (bp)
Oscillospiraceae Dysosmobacter

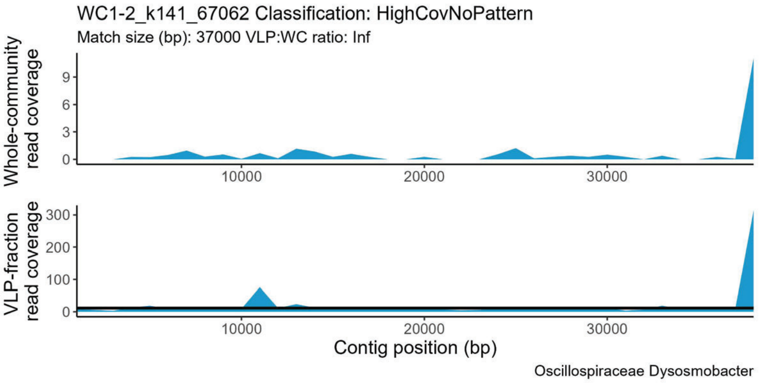
WC1-2_k141_67062 Classification: HighCovNoPattern
Match size (bp): 37000 VLP:WC ratio: Inf
Whole-community read coverage
0
3
6
9
10000
20000
30000
VLP-fraction read coverage
0
100
200
300
10000
20000
30000
Contig position (bp)
Oscillospiraceae Dysosmobacter

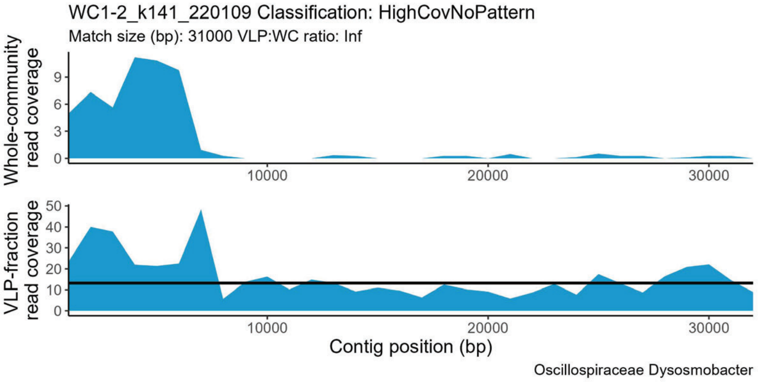
WC1-2_k141_220109 Classification: HighCovNoPattern
Match size (bp): 31000 VLP:WC ratio: Inf
Whole-community read coverage
0
3
6
9
10000
20000
30000
VLP-fraction read coverage
0
10
20
30
40
50
10000
20000
30000
Contig position (bp)
Oscillospiraceae Dysosmobacter

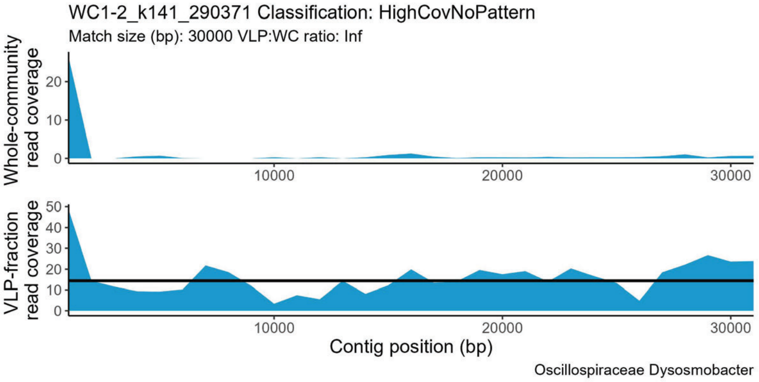
WC1-2_k141_290371 Classification: HighCovNoPattern
Match size (bp): 30000 VLP:WC ratio: Inf
Whole-community read coverage
0
10
20
10000
20000
30000
VLP-fraction read coverage
0
10
20
30
40
50
10000
20000
30000
Contig position (bp)
Oscillospiraceae Dysosmobacter

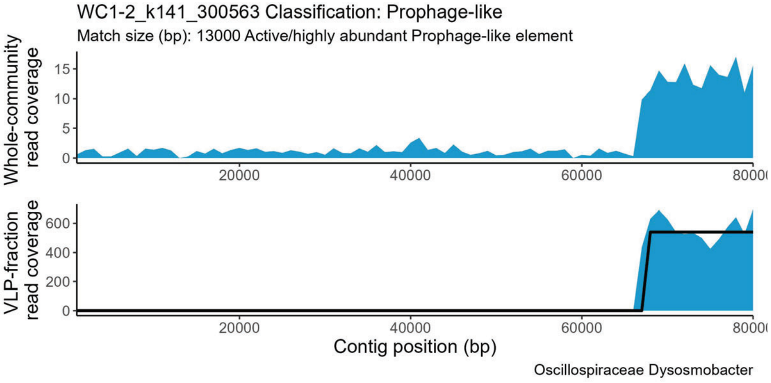
WC1-2_k141_300563 Classification: Prophage-like
Match size (bp): 13000 Active/highly abundant Prophage-like element
Whole-community read coverage
0
5
10
15
20000
40000
60000
VLP-fraction read coverage
0
200
400
600
20000
40000
60000
Contig position (bp)
Oscillospiraceae Dysosmobacter

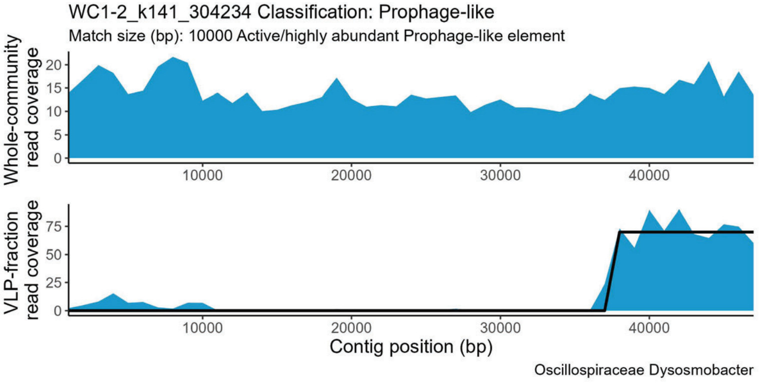
WC1-2_k141_304234 Classification: Prophage-like
Match size (bp): 10000 Active/highly abundant Prophage-like element
Whole-community read coverage
0
5
10
15
20
10000
20000
30000
40000
VLP-fraction read coverage
0
25
50
75
10000
20000
30000
40000
Contig position (bp)
Oscillospiraceae Dysosmobacter

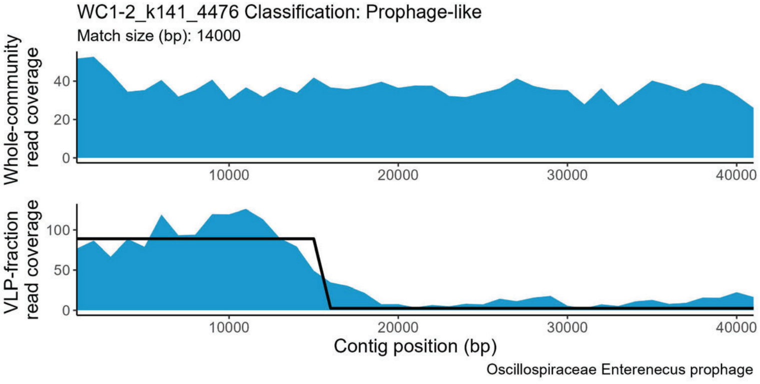
WC1-2_k141_4476 Classification: Prophage-like
Match size (bp): 14000
Whole-community read coverage
0
20
40
10000
20000
30000
40000
VLP-fraction read coverage
0
50
100
10000
20000
30000
40000
Contig position (bp)
Oscillospiraceae Enterenecus prophage

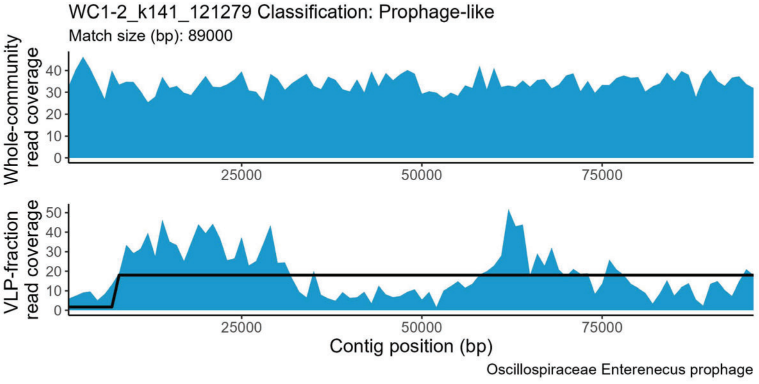
WC1-2_k141_121279 Classification: Prophage-like
Match size (bp): 89000
Whole-community read coverage
0
10
20
30
40
25000
50000
75000
VLP-fraction read coverage
0
10
20
30
40
50
25000
50000
75000
Contig position (bp)
Oscillospiraceae Enterenecus prophage

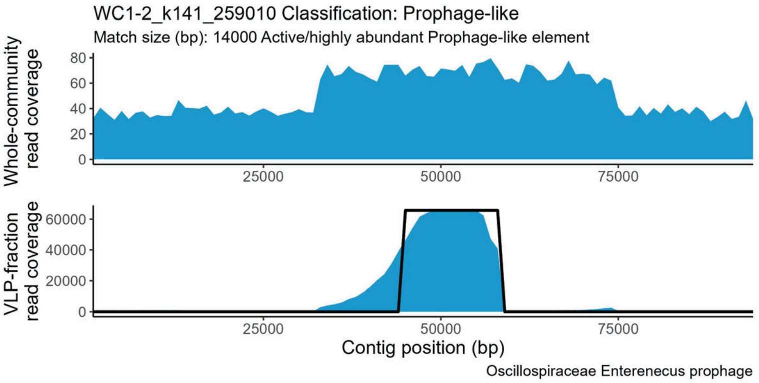
WC1-2_k141_259010 Classification: Prophage-like
Match size (bp): 14000 Active/highly abundant Prophage-like element
Whole-community read coverage
0 20 40 60 80
25000 50000 75000
VLP-fraction read coverage
0 20000 40000 60000
25000 50000 75000
Contig position (bp)
Oscillospiraceae Enterenecus prophage

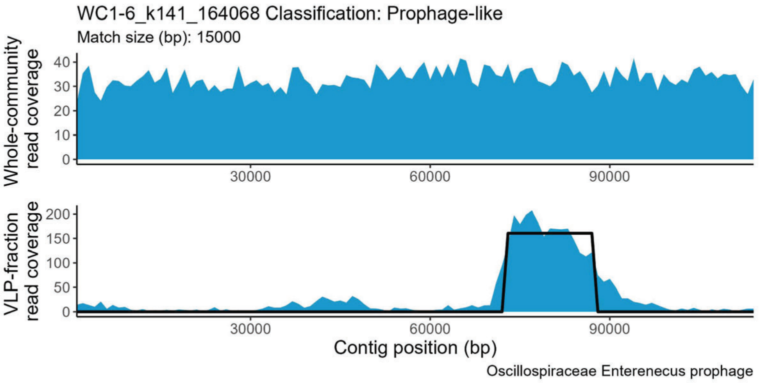
WC1-6_k141_164068 Classification: Prophage-like
Match size (bp): 15000
Whole-community read coverage
0 10 20 30 40
30000 60000 90000
VLP-fraction read coverage
0 50 100 150 200
30000 60000 90000
Contig position (bp)
Oscillospiraceae Enterenecus prophage

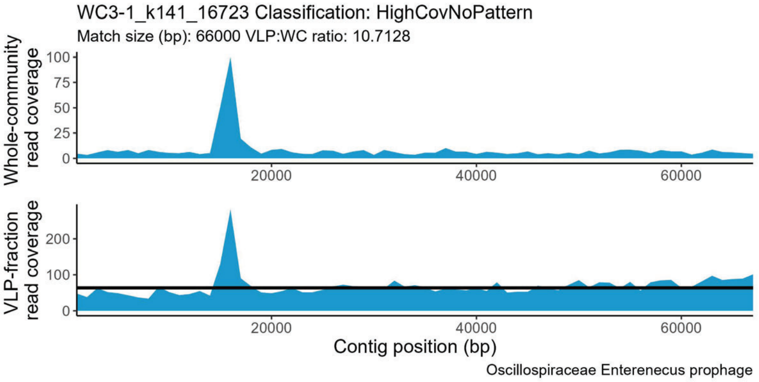
WC3-1_k141_16723 Classification: HighCovNoPattern
Match size (bp): 66000 VLP:WC ratio: 10.7128
Whole-community read coverage
0 25 50 75 100
20000 40000 60000
VLP-fraction read coverage
0 100 200
20000 40000 60000
Contig position (bp)
Oscillospiraceae Enterenecus prophage

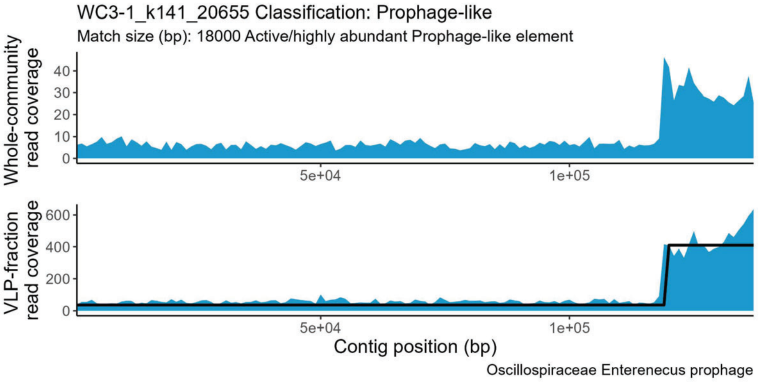
WC3-1_k141_20655 Classification: Prophage-like
Match size (bp): 18000 Active/highly abundant Prophage-like element
Whole-community read coverage
0 10 20 30 40
5e+04 1e+05
VLP-fraction read coverage
0 200 400 600
5e+04 1e+05
Contig position (bp)
Oscillospiraceae Enterenecus prophage

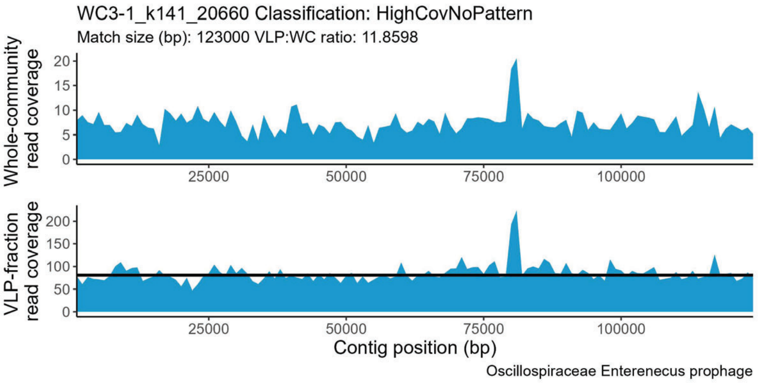
WC3-1_k141_20660 Classification: HighCovNoPattern
Match size (bp): 123000 VLP:WC ratio: 11.8598
Whole-community read coverage
0 5 10 15 20
25000 50000 75000 100000
VLP-fraction read coverage
0 50 100 150 200
25000 50000 75000 100000
Contig position (bp)
Oscillospiraceae Enterenecus prophage

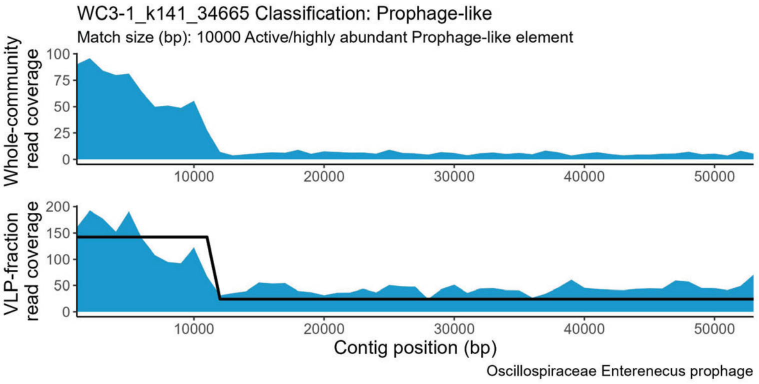
WC3-1_k141_34665 Classification: Prophage-like
Match size (bp): 10000 Active/highly abundant Prophage-like element
Whole-community read coverage
0 25 50 75 100
10000 20000 30000 40000 50000
VLP-fraction read coverage
0 50 100 150 200
10000 20000 30000 40000 50000
Contig position (bp)
Oscillospiraceae Enterenecus prophage

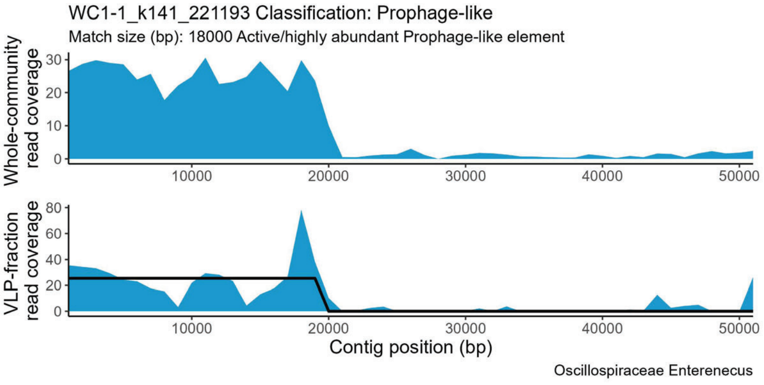
WC1-1_k141_221193 Classification: Prophage-like
Match size (bp): 18000 Active/highly abundant Prophage-like element
Whole-community read coverage
0 10 20 30
10000 20000 30000 40000 50000
VLP-fraction read coverage
0 20 40 60 80
10000 20000 30000 40000 50000
Contig position (bp)
Oscillospiraceae Enterenecus

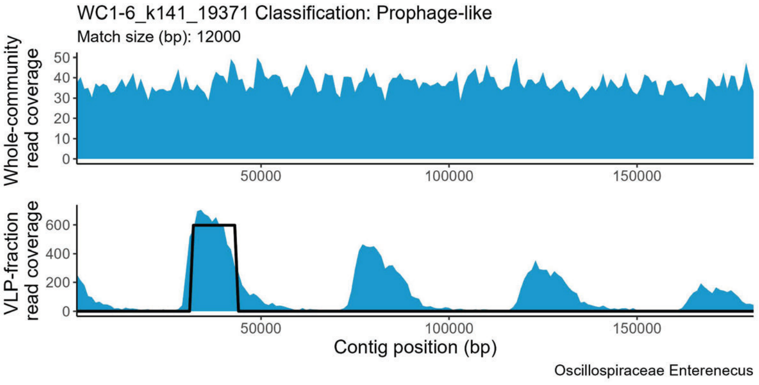
WC1-6_k141_19371 Classification: Prophage-like
Match size (bp): 12000
Whole-community read coverage
0 10 20 30 40 50
50000 100000 150000
VLP-fraction read coverage
0 200 400 600
50000 100000 150000
Contig position (bp)
Oscillospiraceae Enterenecus

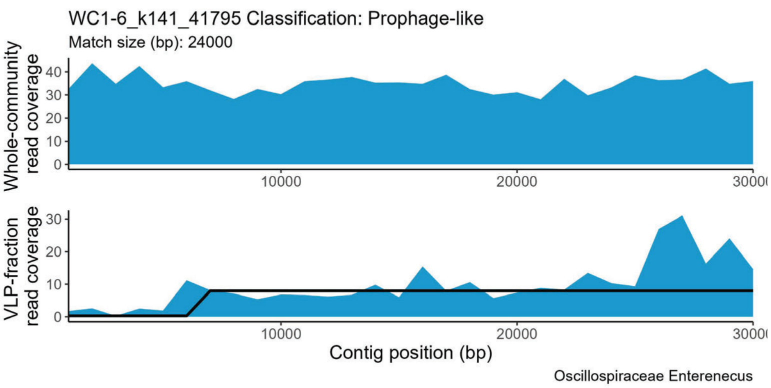
WC1-6_k141_41795 Classification: Prophage-like
Match size (bp): 24000
Whole-community read coverage
VLP-fraction read coverage
Contig position (bp)
Oscillospiraceae Enterenecus

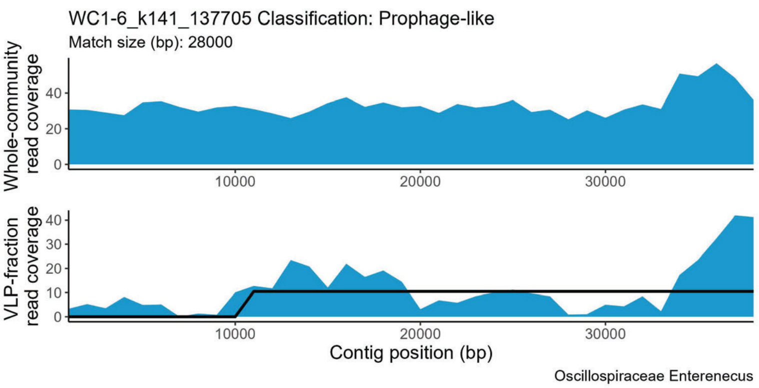
WC1-6_k141_137705 Classification: Prophage-like
Match size (bp): 28000
Whole-community read coverage
VLP-fraction read coverage
Contig position (bp)
Oscillospiraceae Enterenecus

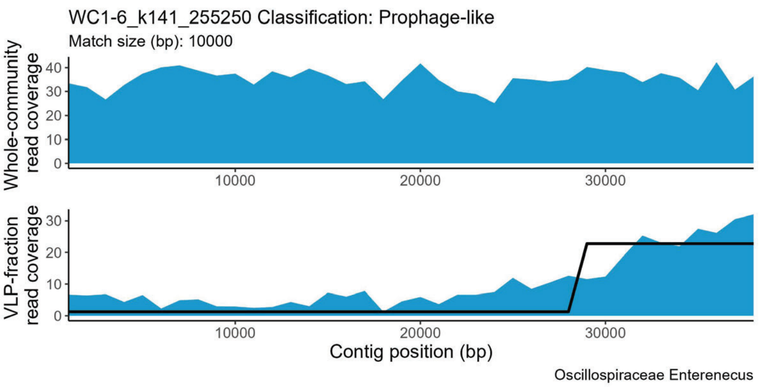
WC1-6_k141_255250 Classification: Prophage-like
Match size (bp): 10000
Whole-community read coverage
VLP-fraction read coverage
Contig position (bp)
Oscillospiraceae Enterenecus

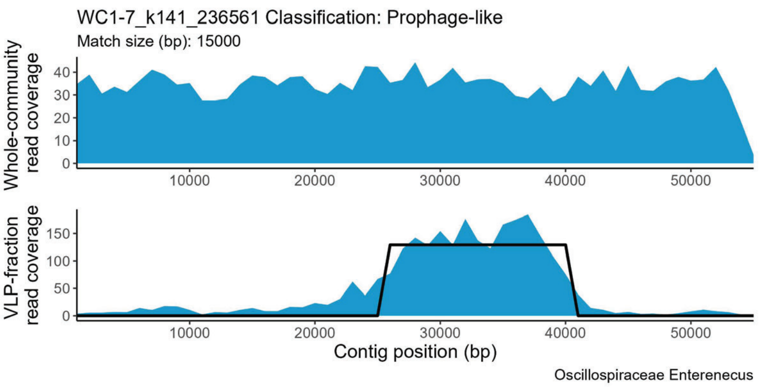
WC1-7_k141_236561 Classification: Prophage-like
Match size (bp): 15000
Whole-community read coverage
VLP-fraction read coverage
Contig position (bp)
Oscillospiraceae Enterenecus

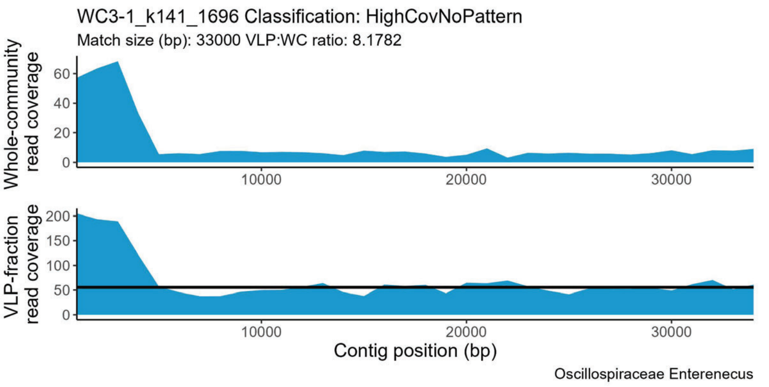
WC3-1_k141_1696 Classification: HighCovNoPattern
Match size (bp): 33000 VLP:WC ratio: 8.1782
Whole-community read coverage
VLP-fraction read coverage
Contig position (bp)
Oscillospiraceae Enterenecus

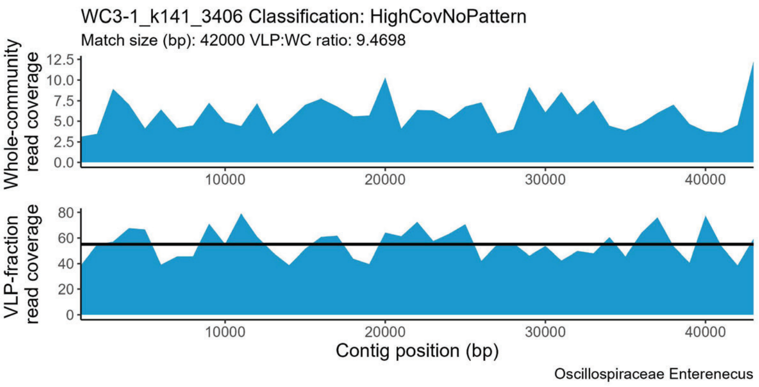
WC3-1_k141_3406 Classification: HighCovNoPattern
Match size (bp): 42000 VLP:WC ratio: 9.4698
Whole-community read coverage
VLP-fraction read coverage
Contig position (bp)
Oscillospiraceae Enterenecus

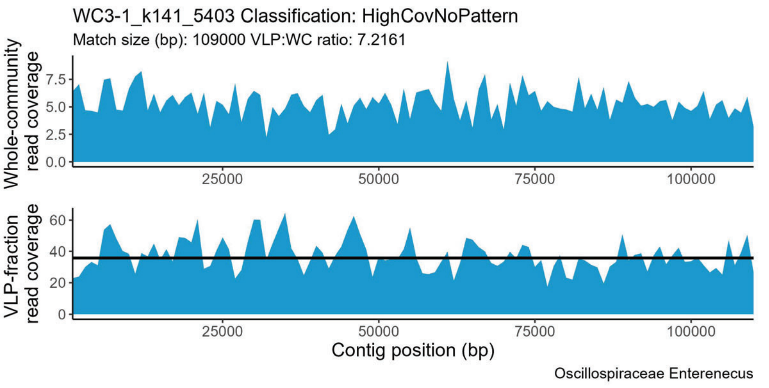
WC3-1_k141_5403 Classification: HighCovNoPattern
Match size (bp): 109000 VLP:WC ratio: 7.2161
Whole-community read coverage
VLP-fraction read coverage
Contig position (bp)
Oscillospiraceae Enterenecus

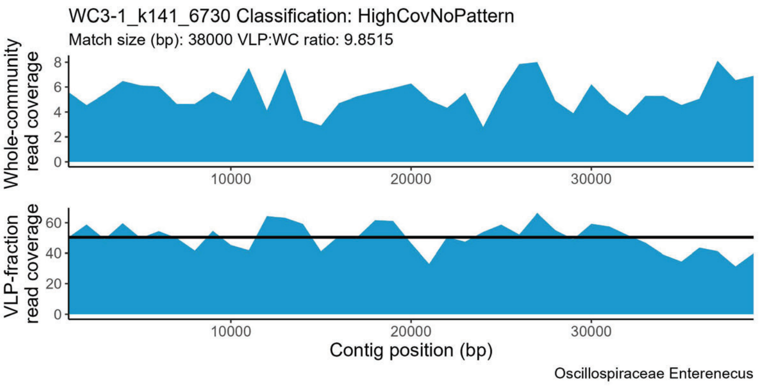
WC3-1_k141_6730 Classification: HighCovNoPattern
Match size (bp): 38000 VLP:WC ratio: 9.8515
Whole-community read coverage
VLP-fraction read coverage
Contig position (bp)
Oscillospiraceae Enterenecus

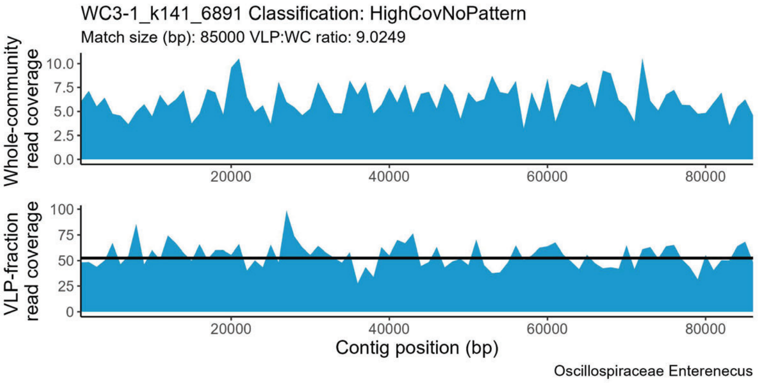
WC3-1_k141_6891 Classification: HighCovNoPattern
Match size (bp): 85000 VLP:WC ratio: 9.0249
Whole-community read coverage
VLP-fraction read coverage
Contig position (bp)
Oscillospiraceae Enterenecus

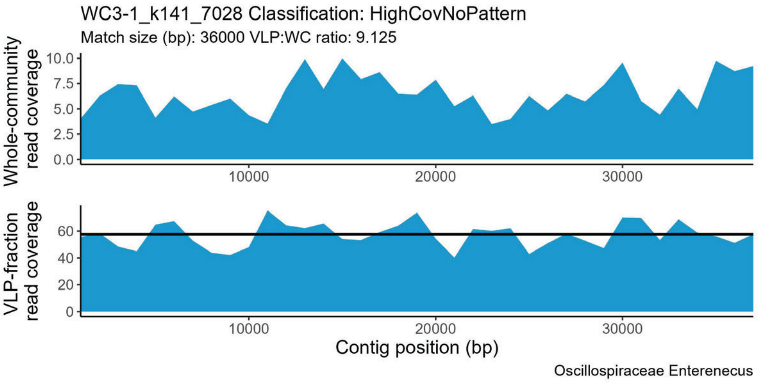
WC3-1_k141_7028 Classification: HighCovNoPattern
Match size (bp): 36000 VLP:WC ratio: 9.125
Whole-community read coverage
VLP-fraction read coverage
Contig position (bp)
Oscillospiraceae Enterenecus

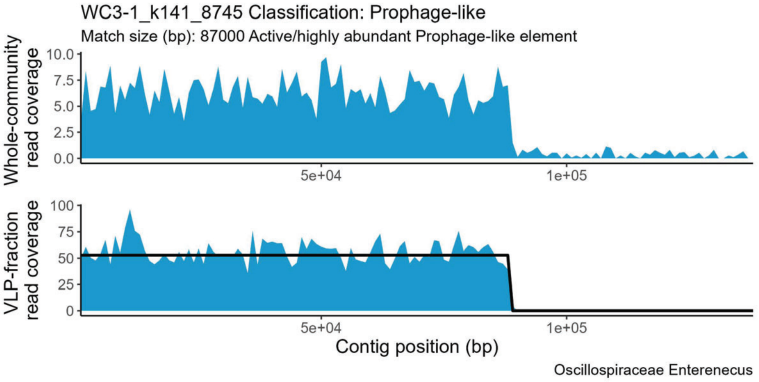
WC3-1_k141_8745 Classification: Prophage-like
Match size (bp): 87000 Active/highly abundant Prophage-like element
Whole-community read coverage
VLP-fraction read coverage
Contig position (bp)
Oscillospiraceae Enterenecus

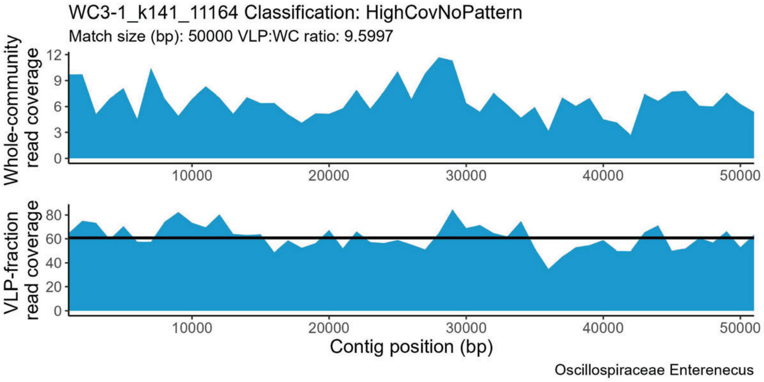
WC3-1_k141_11164 Classification: HighCovNoPattern
Match size (bp): 50000 VLP:WC ratio: 9.5997
Whole-community read coverage
VLP-fraction read coverage
Contig position (bp)
Oscillospiraceae Enterenecus

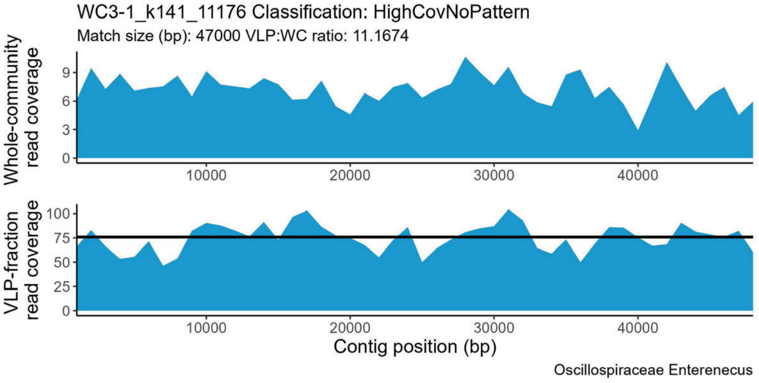
WC3-1_k141_11176 Classification: HighCovNoPattern
Match size (bp): 47000 VLP:WC ratio: 11.1674
Whole-community read coverage
VLP-fraction read coverage
Contig position (bp)
Oscillospiraceae Enterenecus

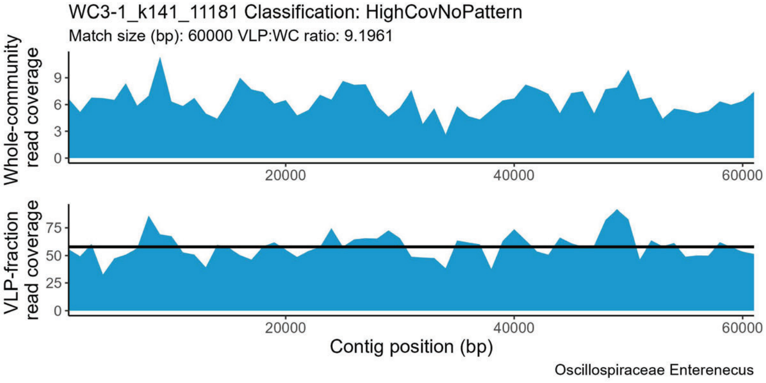
WC3-1_k141_11181 Classification: HighCovNoPattern
Match size (bp): 60000 VLP:WC ratio: 9.1961
Whole-community read coverage
VLP-fraction read coverage
Contig position (bp)
Oscillospiraceae Enterenecus

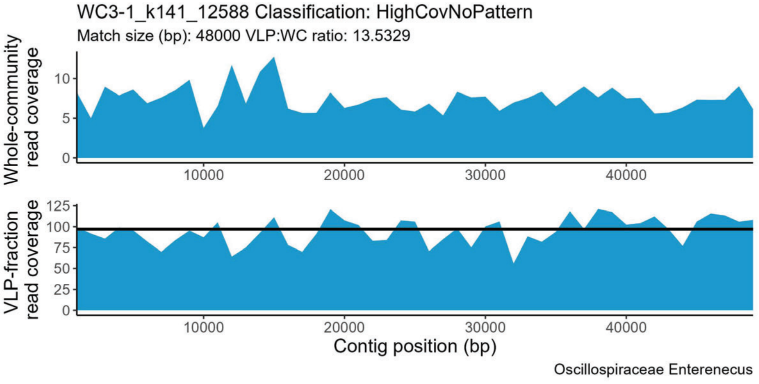
WC3-1_k141_12588 Classification: HighCovNoPattern
Match size (bp): 48000 VLP:WC ratio: 13.5329
Whole-community read coverage
VLP-fraction read coverage
Contig position (bp)
Oscillospiraceae Enterenecus

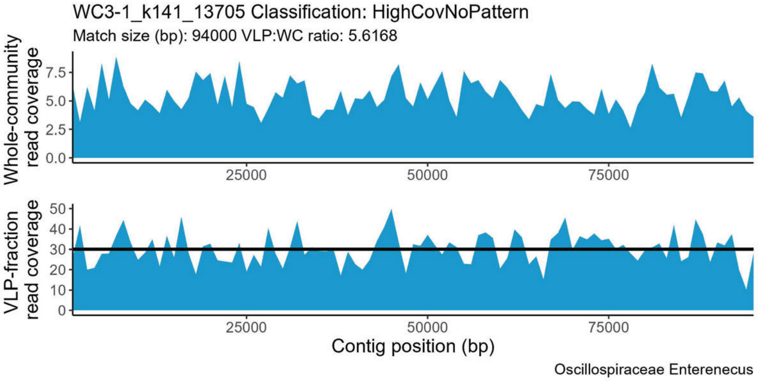
WC3-1_k141_13705 Classification: HighCovNoPattern
Match size (bp): 94000 VLP:WC ratio: 5.6168
Whole-community read coverage
VLP-fraction read coverage
Contig position (bp)
Oscillospiraceae Enterenecus

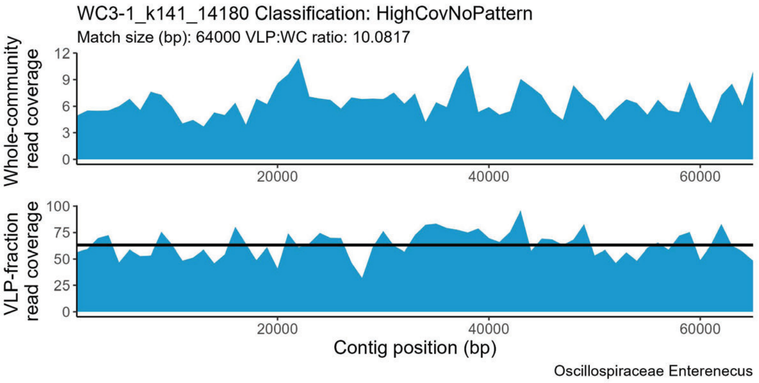
WC3-1_k141_14180 Classification: HighCovNoPattern
Match size (bp): 64000 VLP:WC ratio: 10.0817
Whole-community read coverage
VLP-fraction read coverage
Contig position (bp)
Oscillospiraceae Enterenecus

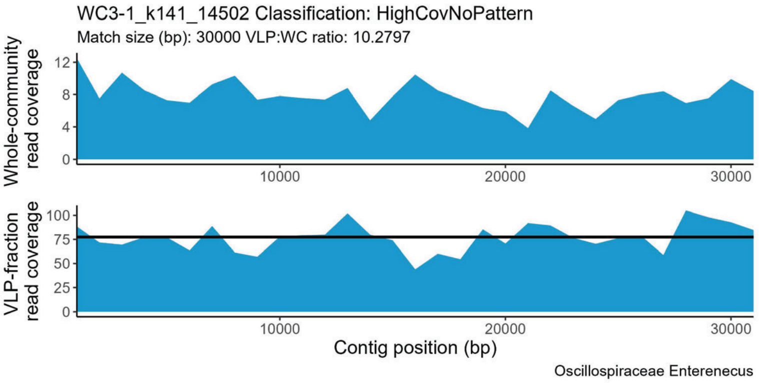
WC3-1_k141_14502 Classification: HighCovNoPattern
Match size (bp): 30000 VLP:WC ratio: 10.2797
Whole-community read coverage
VLP-fraction read coverage
Contig position (bp)
Oscillospiraceae Enterenecus

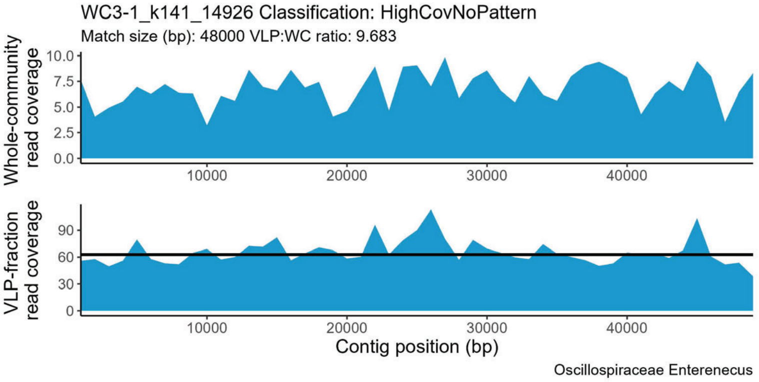
WC3-1_k141_14926 Classification: HighCovNoPattern
Match size (bp): 48000 VLP:WC ratio: 9.683
Whole-community read coverage
VLP-fraction read coverage
Contig position (bp)
Oscillospiraceae Enterenecus

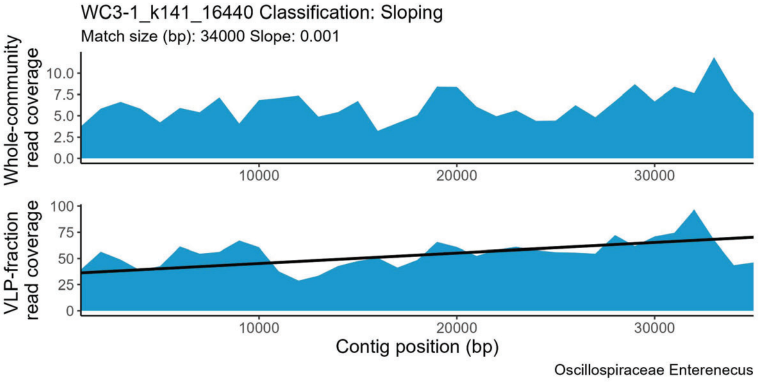
WC3-1_k141_16440 Classification: Sloping
Match size (bp): 34000 Slope: 0.001
Whole-community read coverage
VLP-fraction read coverage
Contig position (bp)
Oscillospiraceae Enterenecus

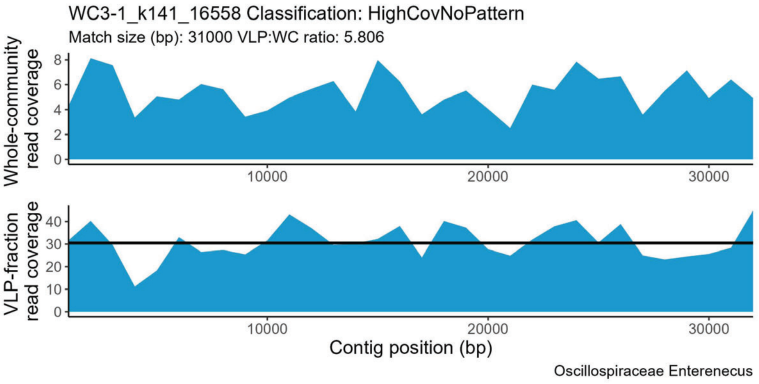
WC3-1_k141_16558 Classification: HighCovNoPattern
Match size (bp): 31000 VLP:WC ratio: 5.806
Whole-community read coverage
VLP-fraction read coverage
Contig position (bp)
Oscillospiraceae Enterenecus

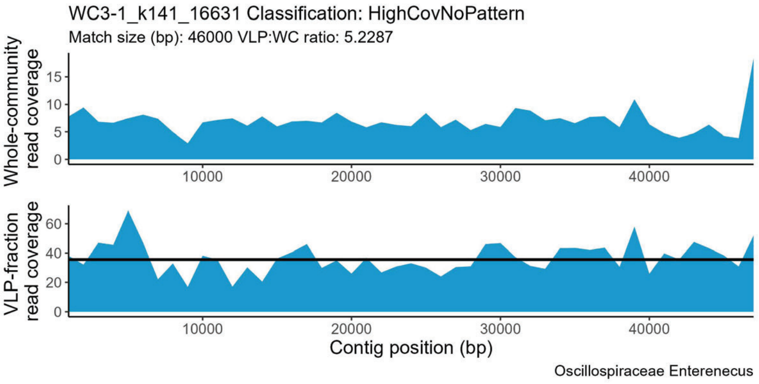
WC3-1_k141_16631 Classification: HighCovNoPattern
Match size (bp): 46000 VLP:WC ratio: 5.2287
Whole-community read coverage
VLP-fraction read coverage
Contig position (bp)
Oscillospiraceae Enterenecus

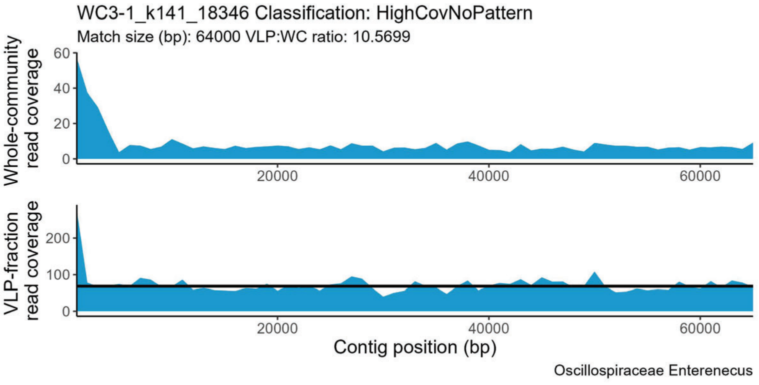
WC3-1_k141_18346 Classification: HighCovNoPattern
Match size (bp): 64000 VLP:WC ratio: 10.5699
Whole-community read coverage
VLP-fraction read coverage
Contig position (bp)
Oscillospiraceae Enterenecus

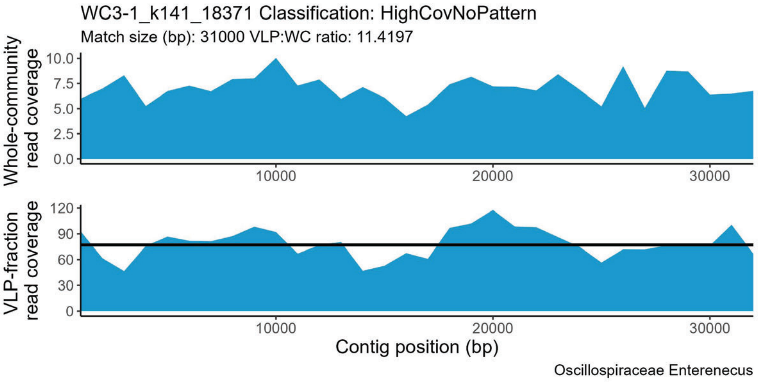
WC3-1_k141_18371 Classification: HighCovNoPattern
Match size (bp): 31000 VLP:WC ratio: 11.4197
Whole-community read coverage
VLP-fraction read coverage
Contig position (bp)
Oscillospiraceae Enterenecus

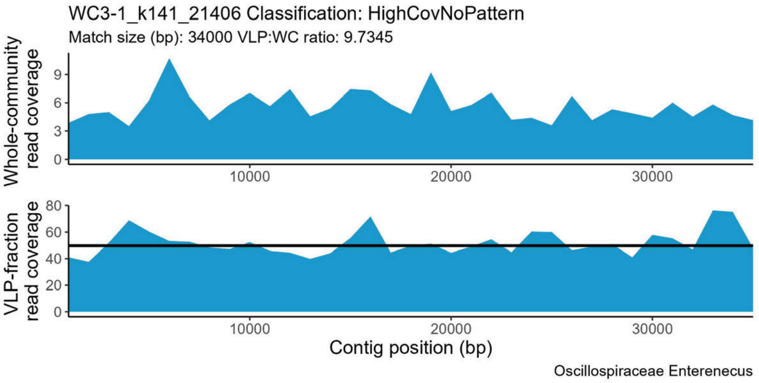
WC3-1_k141_21406 Classification: HighCovNoPattern
Match size (bp): 34000 VLP:WC ratio: 9.7345
Whole-community read coverage
VLP-fraction read coverage
Contig position (bp)
Oscillospiraceae Enterenecus

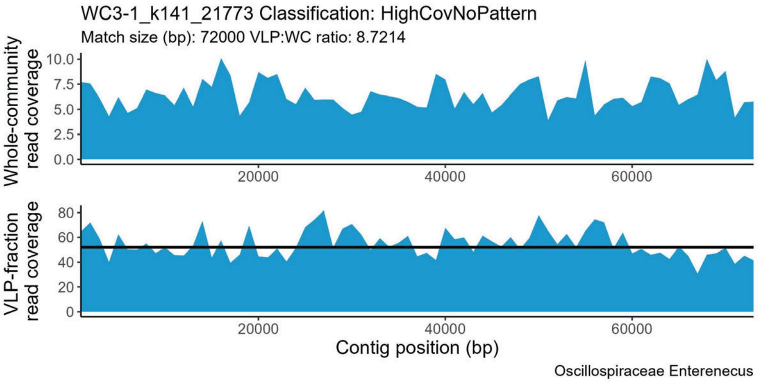
WC3-1_k141_21773 Classification: HighCovNoPattern
Match size (bp): 72000 VLP:WC ratio: 8.7214
Whole-community read coverage
VLP-fraction read coverage
Contig position (bp)
Oscillospiraceae Enterenecus

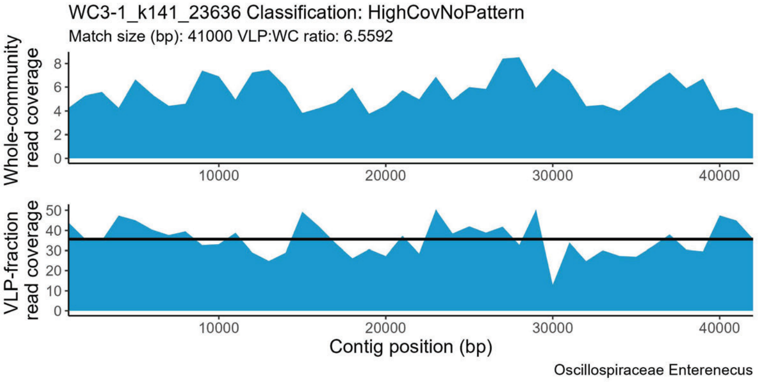
WC3-1_k141_23636 Classification: HighCovNoPattern
Match size (bp): 41000 VLP:WC ratio: 6.5592
Whole-community read coverage
VLP-fraction read coverage
Contig position (bp)
Oscillospiraceae Enterenecus

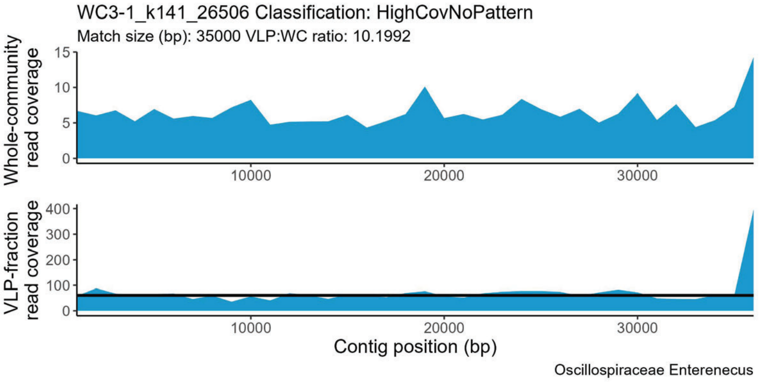
WC3-1_k141_26506 Classification: HighCovNoPattern
Match size (bp): 35000 VLP:WC ratio: 10.1992
Whole-community read coverage
VLP-fraction read coverage
Contig position (bp)
Oscillospiraceae Enterenecus

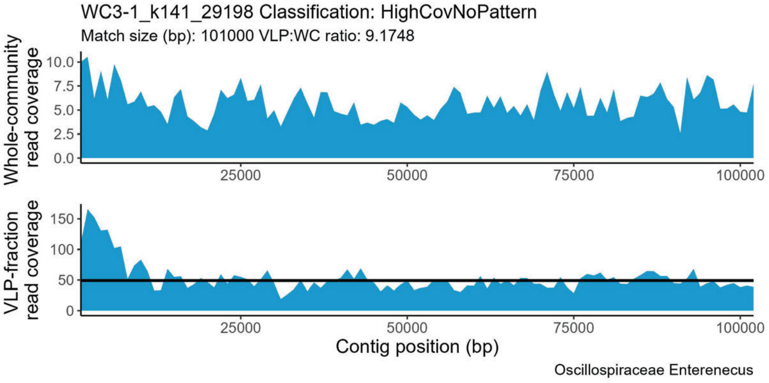
WC3-1_k141_29198 Classification: HighCovNoPattern
Match size (bp): 101000 VLP:WC ratio: 9.1748
Whole-community read coverage
VLP-fraction read coverage
Contig position (bp)
Oscillospiraceae Enterenecus

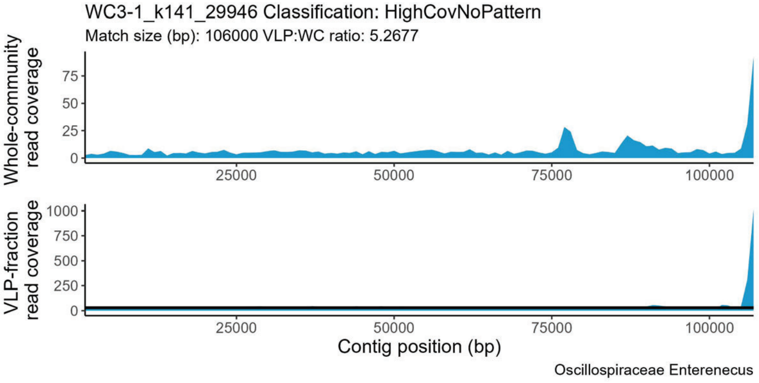
WC3-1_k141_29946 Classification: HighCovNoPattern
Match size (bp): 106000 VLP:WC ratio: 5.2677
Whole-community read coverage
VLP-fraction read coverage
Contig position (bp)
Oscillospiraceae Enterenecus

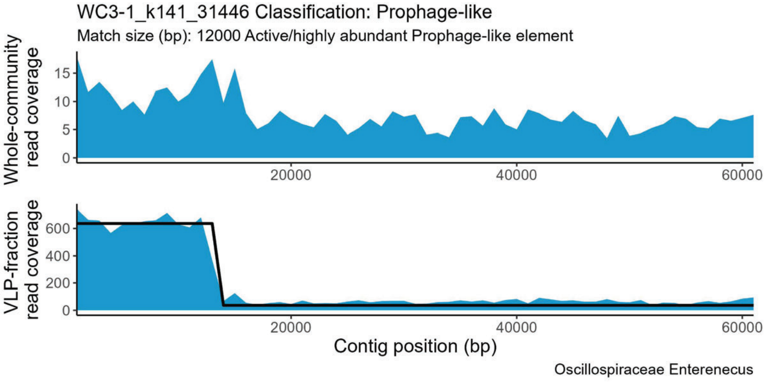
WC3-1_k141_31446 Classification: Prophage-like
Match size (bp): 12000 Active/highly abundant Prophage-like element
Whole-community read coverage
VLP-fraction read coverage
Contig position (bp)
Oscillospiraceae Enterenecus

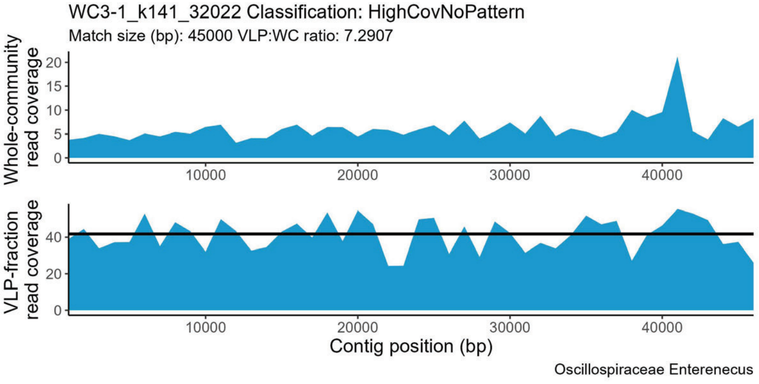
WC3-1_k141_32022 Classification: HighCovNoPattern
Match size (bp): 45000 VLP:WC ratio: 7.2907
Whole-community read coverage
VLP-fraction read coverage
Contig position (bp)
Oscillospiraceae Enterenecus

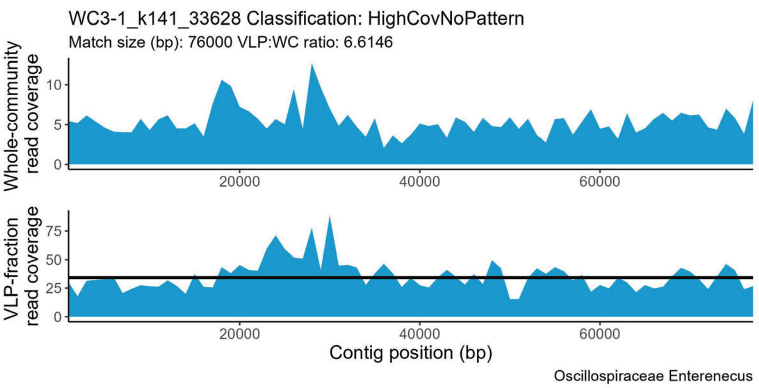
WC3-1_k141_33628 Classification: HighCovNoPattern
Match size (bp): 76000 VLP:WC ratio: 6.6146
Whole-community read coverage
VLP-fraction read coverage
Contig position (bp)
Oscillospiraceae Enterenecus

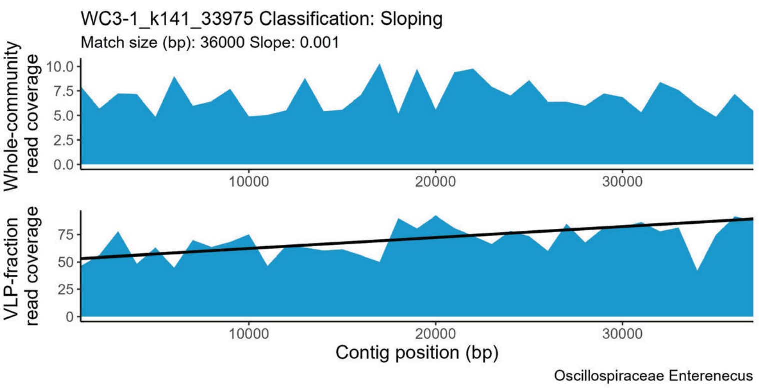
WC3-1_k141_33975 Classification: Sloping
Match size (bp): 36000 Slope: 0.001
Whole-community read coverage
VLP-fraction read coverage
Contig position (bp)
Oscillospiraceae Enterenecus

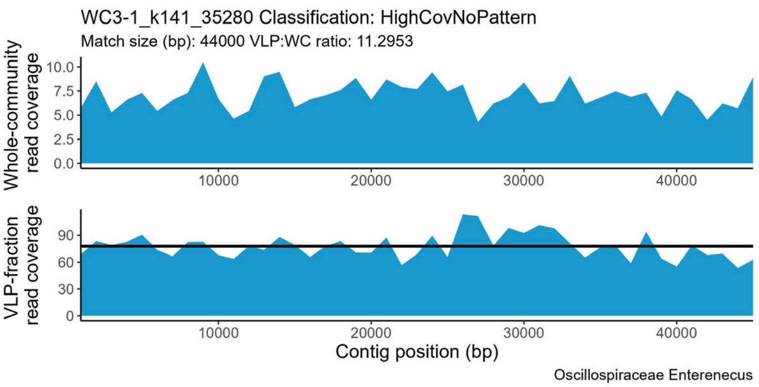
WC3-1_k141_35280 Classification: HighCovNoPattern
Match size (bp): 44000 VLP:WC ratio: 11.2953
Whole-community read coverage
VLP-fraction read coverage
Contig position (bp)
Oscillospiraceae Enterenecus

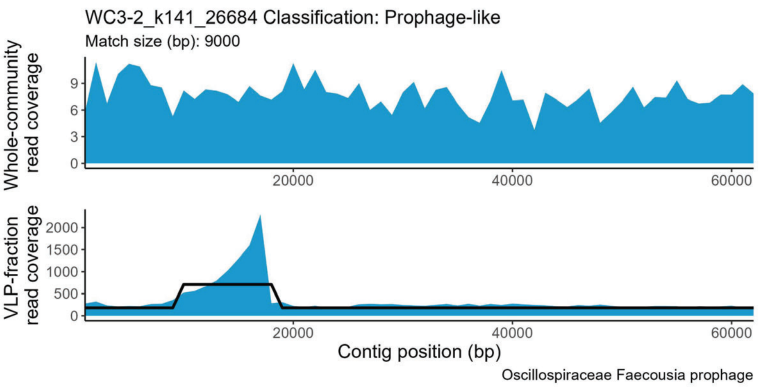
WC3-2_k141_26684 Classification: Prophage-like
Match size (bp): 9000
Whole-community read coverage
VLP-fraction read coverage
Contig position (bp)
Oscillospiraceae Faecousia prophage

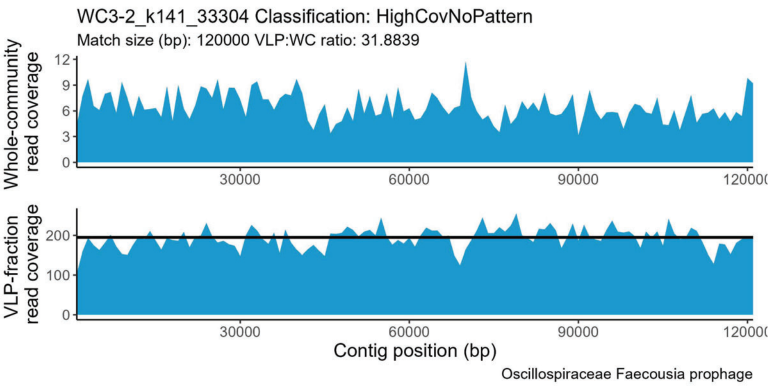
WC3-2_k141_33304 Classification: HighCovNoPattern
Match size (bp): 120000 VLP:WC ratio: 31.8839
Whole-community read coverage
VLP-fraction read coverage
Contig position (bp)
Oscillospiraceae Faecousia prophage

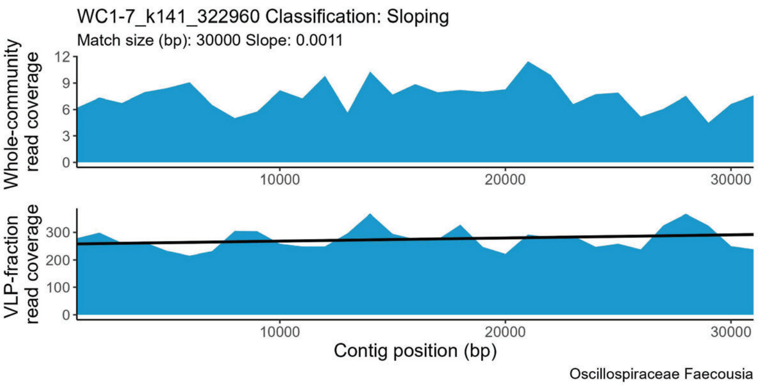
WC1-7_k141_322960 Classification: Sloping
Match size (bp): 30000 Slope: 0.0011
Whole-community read coverage
VLP-fraction read coverage
Contig position (bp)
Oscillospiraceae Faecousia

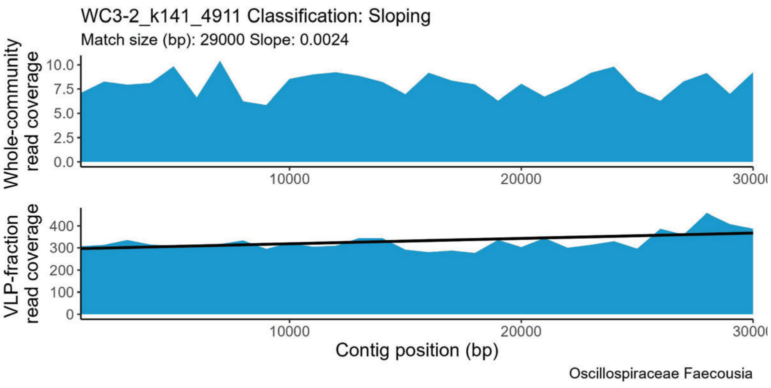
WC3-2_k141_4911 Classification: Sloping
Match size (bp): 29000 Slope: 0.0024
Whole-community read coverage
VLP-fraction read coverage
Contig position (bp)
Oscillospiraceae Faecousia

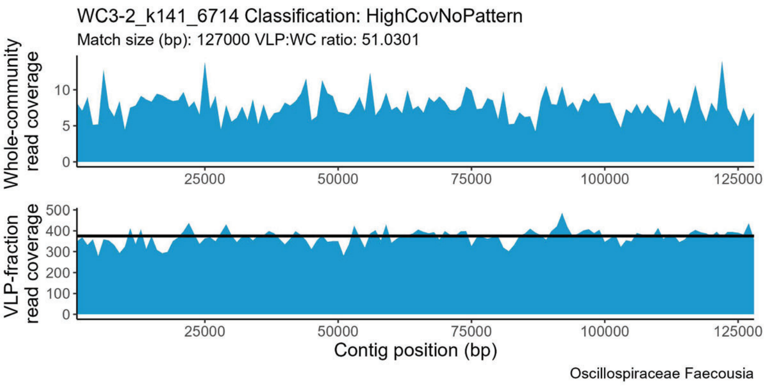
WC3-2_k141_6714 Classification: HighCovNoPattern
Match size (bp): 127000 VLP:WC ratio: 51.0301
Whole-community read coverage
VLP-fraction read coverage
Contig position (bp)
Oscillospiraceae Faecousia

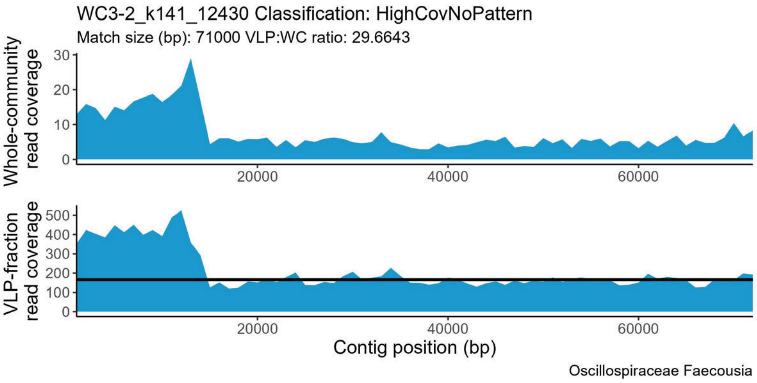
WC3-2_k141_12430 Classification: HighCovNoPattern
Match size (bp): 71000 VLP:WC ratio: 29.6643
Whole-community read coverage
VLP-fraction read coverage
Contig position (bp)
Oscillospiraceae Faecousia

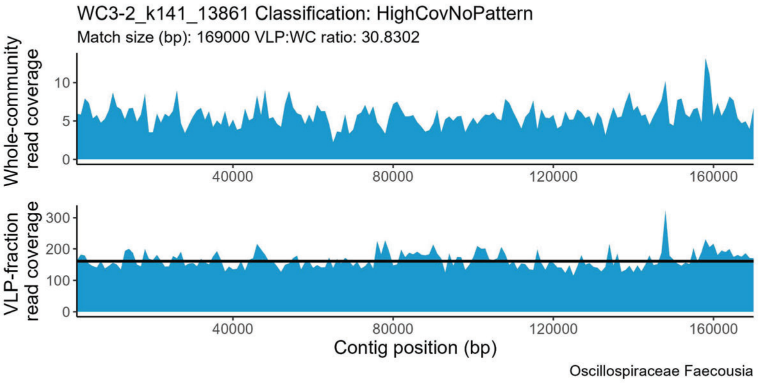
WC3-2_k141_13861 Classification: HighCovNoPattern
Match size (bp): 169000 VLP:WC ratio: 30.8302
Whole-community read coverage
VLP-fraction read coverage
Contig position (bp)
Oscillospiraceae Faecousia

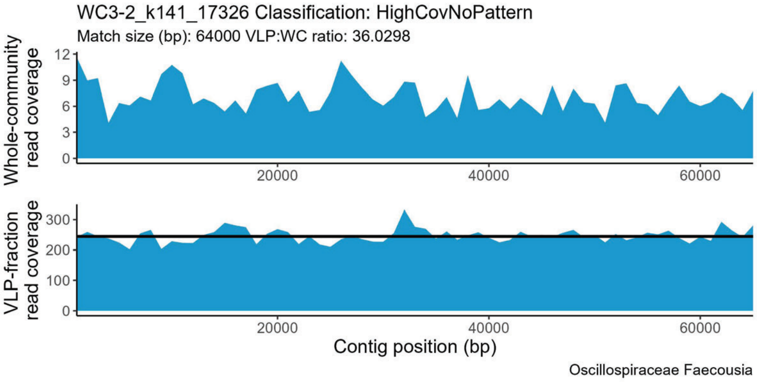
WC3-2_k141_17326 Classification: HighCovNoPattern
Match size (bp): 64000 VLP:WC ratio: 36.0298
Whole-community read coverage
VLP-fraction read coverage
Contig position (bp)
Oscillospiraceae Faecousia

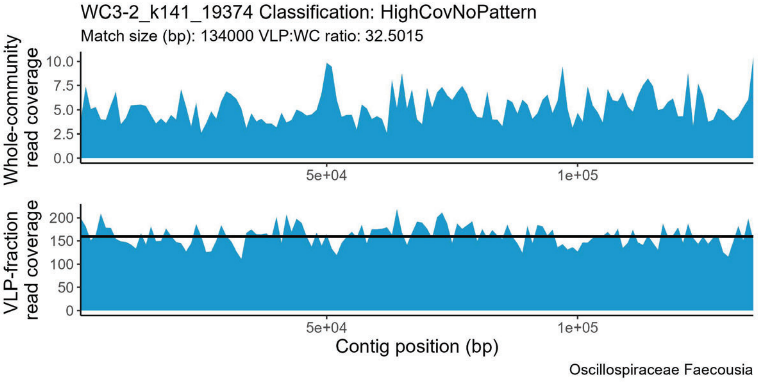
WC3-2_k141_19374 Classification: HighCovNoPattern
Match size (bp): 134000 VLP:WC ratio: 32.5015
Whole-community read coverage
VLP-fraction read coverage
Contig position (bp)
Oscillospiraceae Faecousia

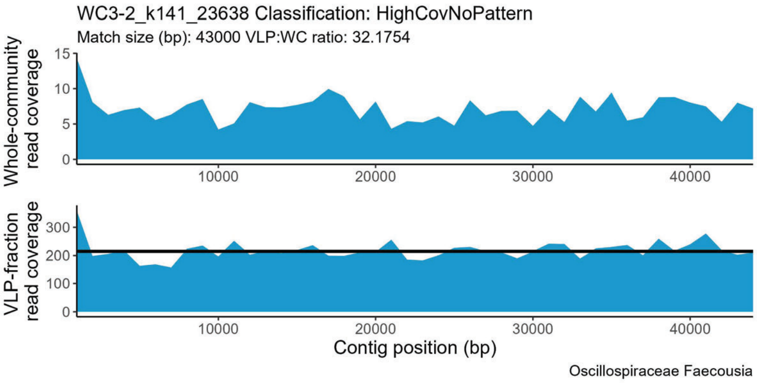
WC3-2_k141_23638 Classification: HighCovNoPattern
Match size (bp): 43000 VLP:WC ratio: 32.1754
Whole-community read coverage
VLP-fraction read coverage
Contig position (bp)
Oscillospiraceae Faecousia

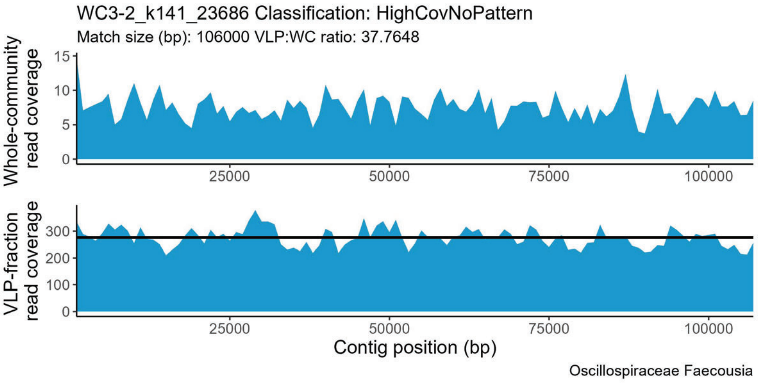
WC3-2_k141_23686 Classification: HighCovNoPattern
Match size (bp): 106000 VLP:WC ratio: 37.7648
Whole-community read coverage
VLP-fraction read coverage
Contig position (bp)
Oscillospiraceae Faecousia

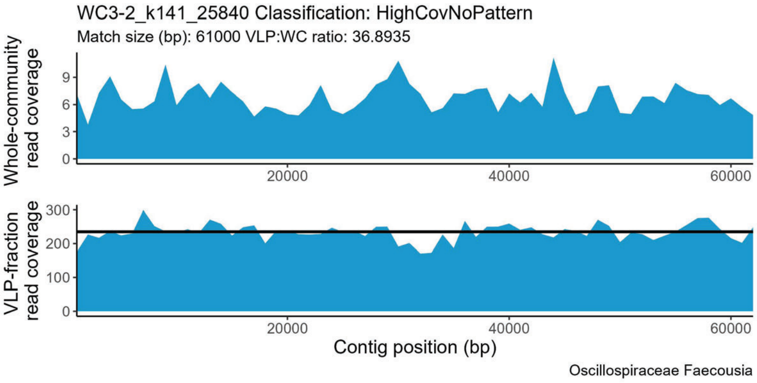
WC3-2_k141_25840 Classification: HighCovNoPattern
Match size (bp): 61000 VLP:WC ratio: 36.8935
Whole-community read coverage
VLP-fraction read coverage
Contig position (bp)
Oscillospiraceae Faecousia

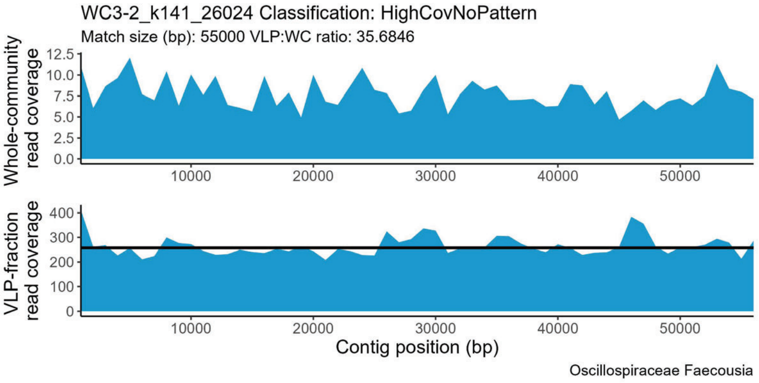
WC3-2_k141_26024 Classification: HighCovNoPattern
Match size (bp): 55000 VLP:WC ratio: 35.6846
Whole-community read coverage
VLP-fraction read coverage
Contig position (bp)
Oscillospiraceae Faecousia

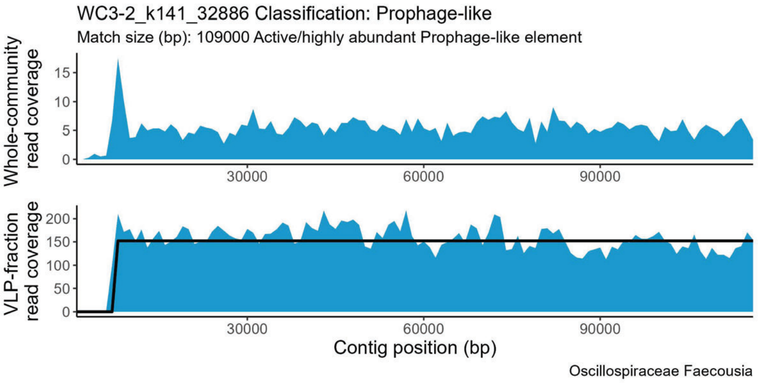
WC3-2_k141_32886 Classification: Prophage-like
Match size (bp): 109000 Active/highly abundant Prophage-like element
Whole-community read coverage
VLP-fraction read coverage
Contig position (bp)
Oscillospiraceae Faecousia

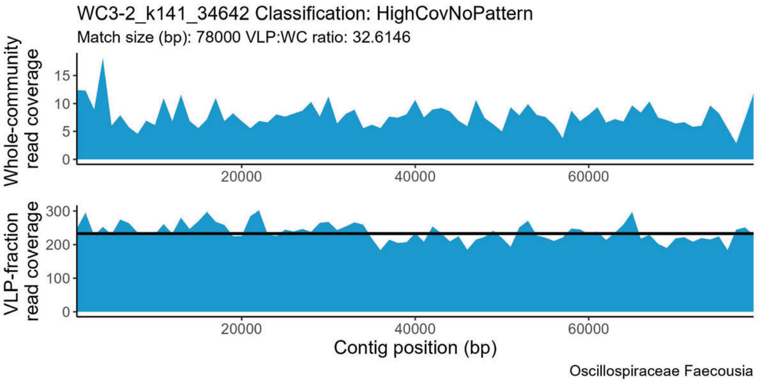
WC3-2_k141_34642 Classification: HighCovNoPattern
Match size (bp): 78000 VLP:WC ratio: 32.6146
Whole-community read coverage
VLP-fraction read coverage
Contig position (bp)
Oscillospiraceae Faecousia

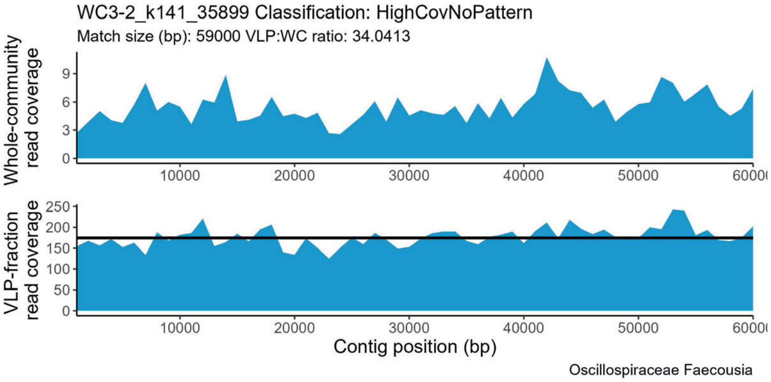
WC3-2_k141_35899 Classification: HighCovNoPattern
Match size (bp): 59000 VLP:WC ratio: 34.0413
Whole-community read coverage
VLP-fraction read coverage
Contig position (bp)
Oscillospiraceae Faecousia

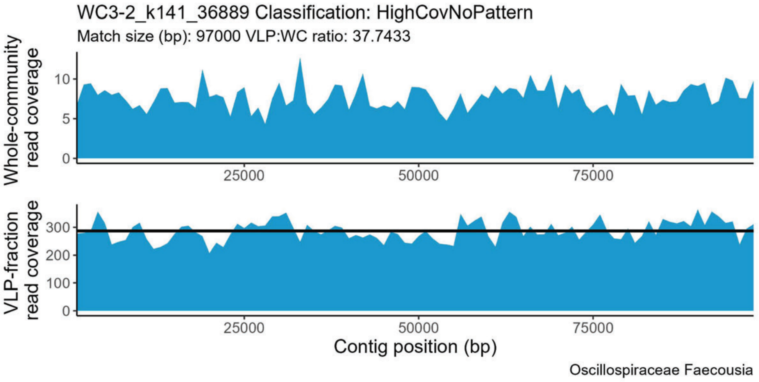
WC3-2_k141_36889 Classification: HighCovNoPattern
Match size (bp): 97000 VLP:WC ratio: 37.7433
Whole-community read coverage
VLP-fraction read coverage
Contig position (bp)
Oscillospiraceae Faecousia

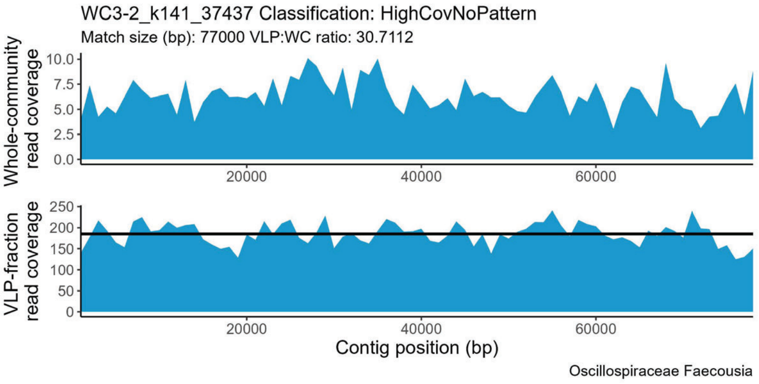
WC3-2_k141_37437 Classification: HighCovNoPattern
Match size (bp): 77000 VLP:WC ratio: 30.7112
Whole-community read coverage
VLP-fraction read coverage
Contig position (bp)
Oscillospiraceae Faecousia

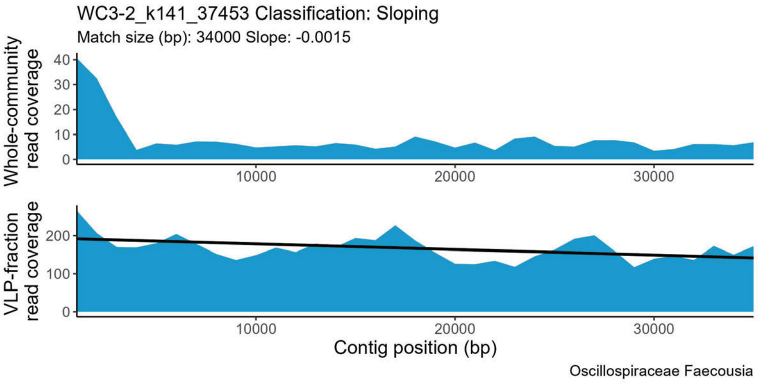
WC3-2_k141_37453 Classification: Sloping
Match size (bp): 34000 Slope: -0.0015
Whole-community read coverage
VLP-fraction read coverage
Contig position (bp)
Oscillospiraceae Faecousia

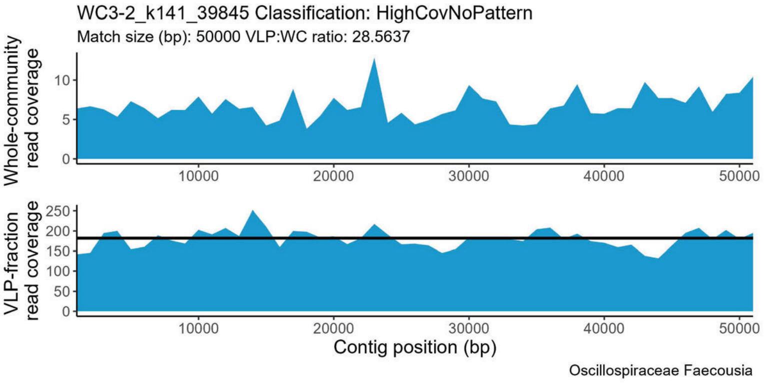
WC3-2_k141_39845 Classification: HighCovNoPattern
Match size (bp): 50000 VLP:WC ratio: 28.5637
Whole-community read coverage
VLP-fraction read coverage
Contig position (bp)
Oscillospiraceae Faecousia

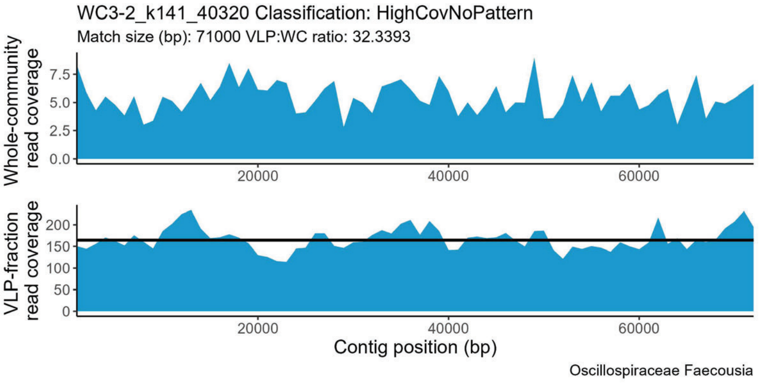
WC3-2_k141_40320 Classification: HighCovNoPattern
Match size (bp): 71000 VLP:WC ratio: 32.3393
Whole-community read coverage
VLP-fraction read coverage
Contig position (bp)
Oscillospiraceae Faecousia

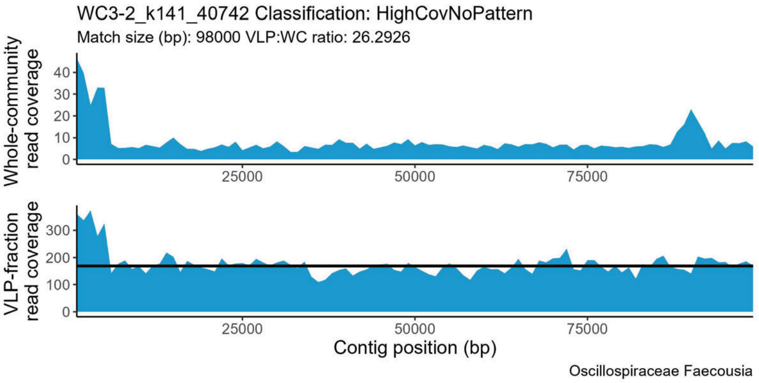
WC3-2_k141_40742 Classification: HighCovNoPattern
Match size (bp): 98000 VLP:WC ratio: 26.2926
Whole-community read coverage
VLP-fraction read coverage
Contig position (bp)
Oscillospiraceae Faecousia

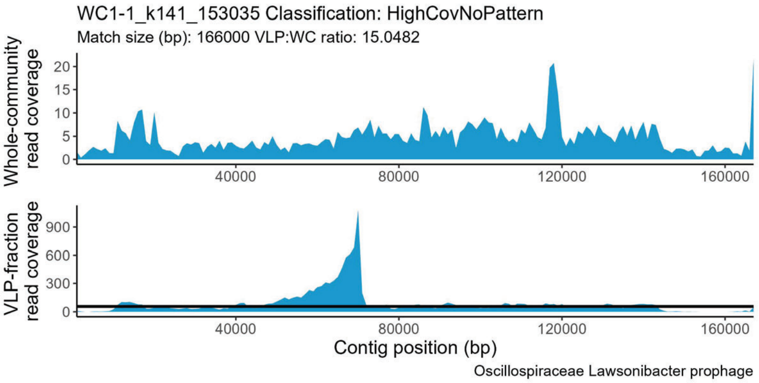
WC1-1_k141_153035 Classification: HighCovNoPattern
Match size (bp): 166000 VLP:WC ratio: 15.0482
Whole-community read coverage
VLP-fraction read coverage
Contig position (bp)
Oscillospiraceae Lawsonibacter prophage

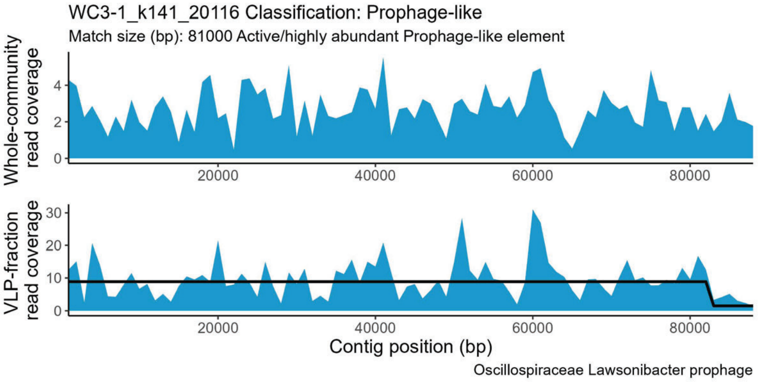
WC3-1_k141_20116 Classification: Prophage-like
Match size (bp): 81000 Active/highly abundant Prophage-like element
Whole-community read coverage
VLP-fraction read coverage
Contig position (bp)
Oscillospiraceae Lawsonibacter prophage

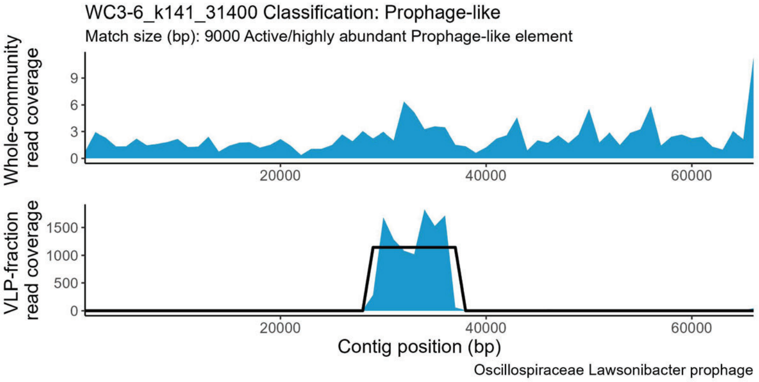
WC3-6_k141_31400 Classification: Prophage-like
Match size (bp): 9000 Active/highly abundant Prophage-like element
Whole-community read coverage
VLP-fraction read coverage
Contig position (bp)
Oscillospiraceae Lawsonibacter prophage

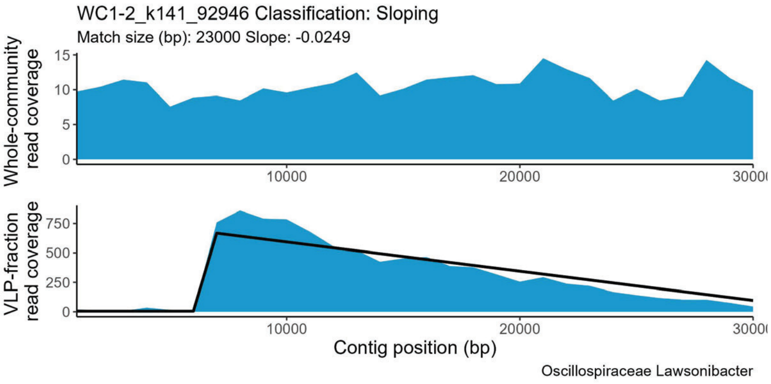
WC1-2_k141_92946 Classification: Sloping
Match size (bp): 23000 Slope: -0.0249
Whole-community read coverage
VLP-fraction read coverage
Contig position (bp)
Oscillospiraceae Lawsonibacter

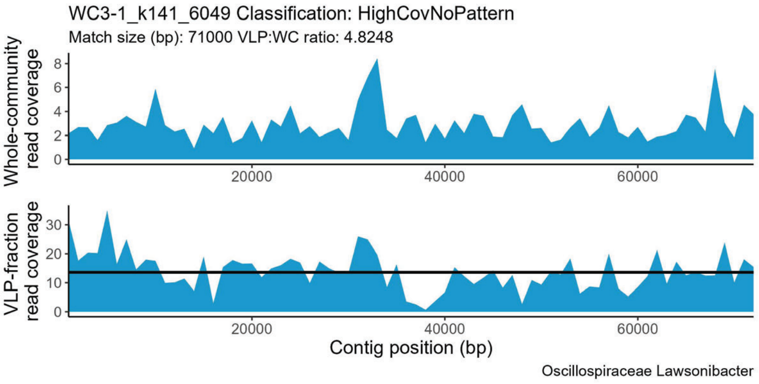
WC3-1_k141_6049 Classification: HighCovNoPattern
Match size (bp): 71000 VLP:WC ratio: 4.8248
Whole-community read coverage
VLP-fraction read coverage
Contig position (bp)
Oscillospiraceae Lawsonibacter

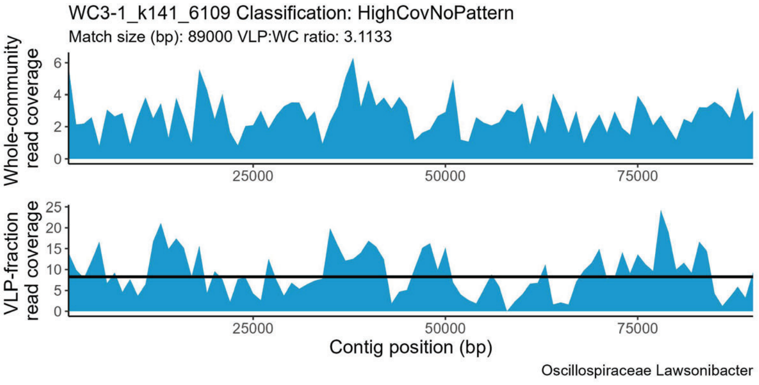
WC3-1_k141_6109 Classification: HighCovNoPattern
Match size (bp): 89000 VLP:WC ratio: 3.1133
Whole-community read coverage
VLP-fraction read coverage
Contig position (bp)
Oscillospiraceae Lawsonibacter

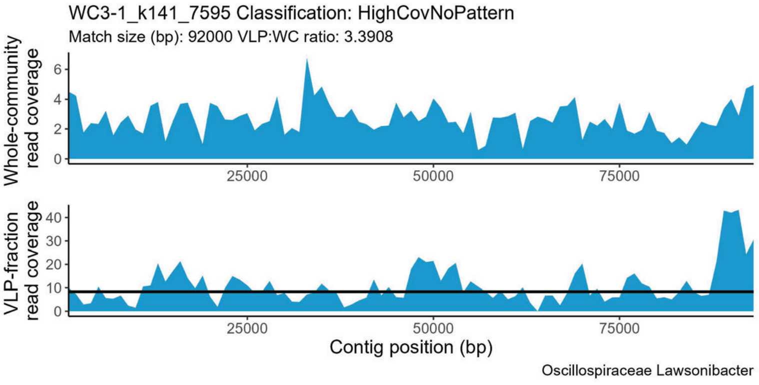
WC3-1_k141_7595 Classification: HighCovNoPattern
Match size (bp): 92000 VLP:WC ratio: 3.3908
Whole-community read coverage
VLP-fraction read coverage
Contig position (bp)
Oscillospiraceae Lawsonibacter

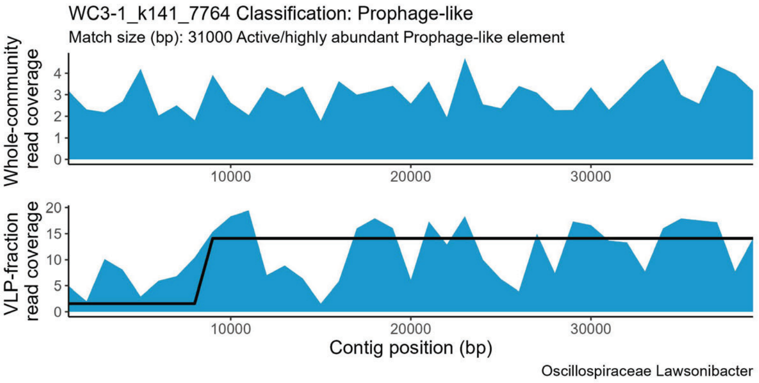
WC3-1_k141_7764 Classification: Prophage-like
Match size (bp): 31000 Active/highly abundant Prophage-like element
Whole-community read coverage
0 1 2 3 4
10000 20000 30000
VLP-fraction read coverage
0 5 10 15 20
10000 20000 30000
Contig position (bp)
Oscillospiraceae Lawsonibacter

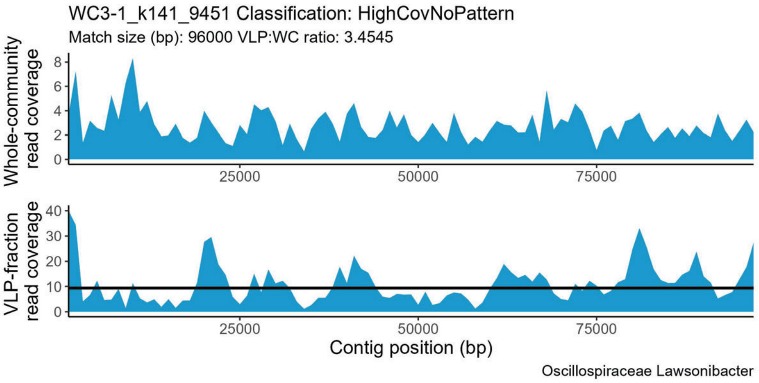
WC3-1_k141_9451 Classification: HighCovNoPattern
Match size (bp): 96000 VLP:WC ratio: 3.4545
Whole-community read coverage
0 2 4 6 8
25000 50000 75000
VLP-fraction read coverage
0 10 20 30 40
25000 50000 75000
Contig position (bp)
Oscillospiraceae Lawsonibacter

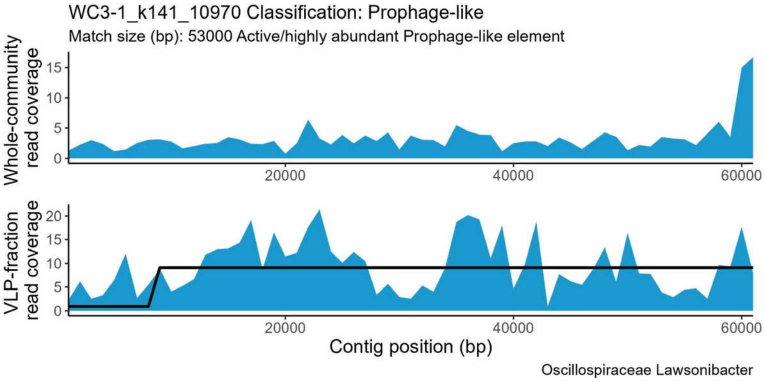
WC3-1_k141_10970 Classification: Prophage-like
Match size (bp): 53000 Active/highly abundant Prophage-like element
Whole-community read coverage
0 5 10 15
20000 40000 60000
VLP-fraction read coverage
0 5 10 15 20
20000 40000 60000
Contig position (bp)
Oscillospiraceae Lawsonibacter

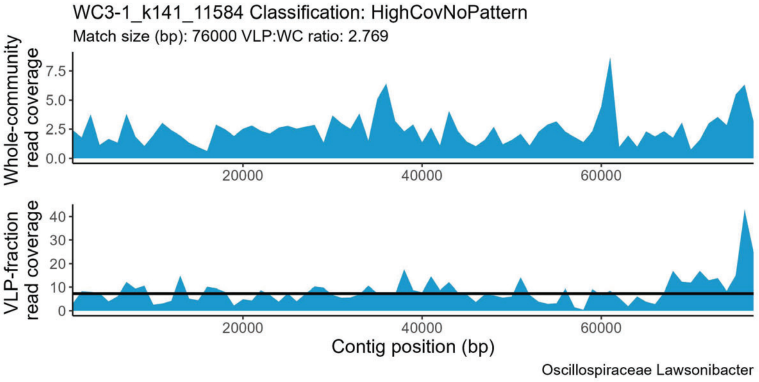
WC3-1_k141_11584 Classification: HighCovNoPattern
Match size (bp): 76000 VLP:WC ratio: 2.769
Whole-community read coverage
0.0 2.5 5.0 7.5
20000 40000 60000
VLP-fraction read coverage
0 10 20 30 40
20000 40000 60000
Contig position (bp)
Oscillospiraceae Lawsonibacter

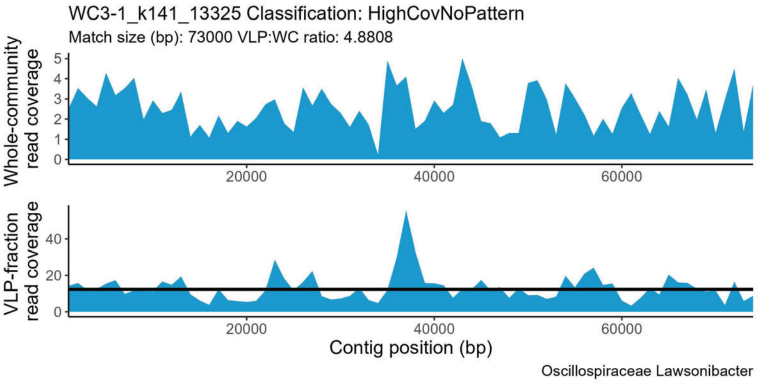
WC3-1_k141_13325 Classification: HighCovNoPattern
Match size (bp): 73000 VLP:WC ratio: 4.8808
Whole-community read coverage
0 1 2 3 4 5
20000 40000 60000
VLP-fraction read coverage
0 20 40
20000 40000 60000
Contig position (bp)
Oscillospiraceae Lawsonibacter

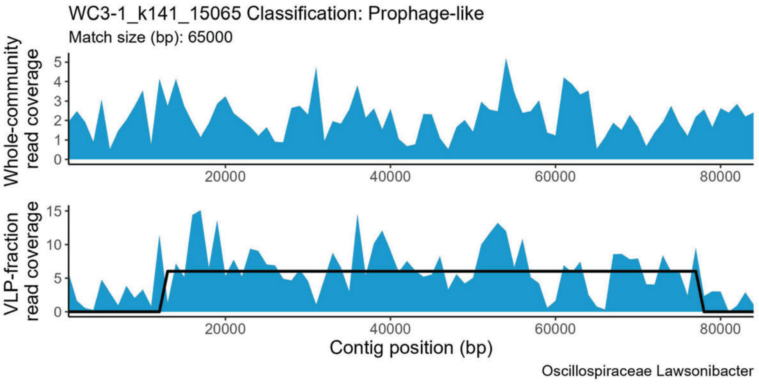
WC3-1_k141_15065 Classification: Prophage-like
Match size (bp): 65000
Whole-community read coverage
0 1 2 3 4 5
20000 40000 60000 80000
VLP-fraction read coverage
0 5 10 15
20000 40000 60000 80000
Contig position (bp)
Oscillospiraceae Lawsonibacter

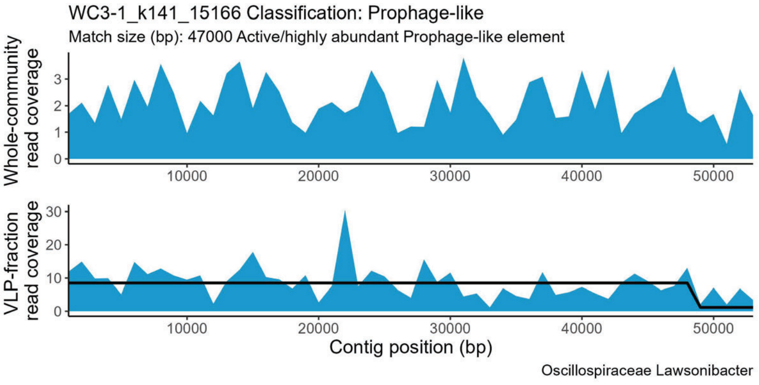
WC3-1_k141_15166 Classification: Prophage-like
Match size (bp): 47000 Active/highly abundant Prophage-like element
Whole-community read coverage
0 1 2 3
10000 20000 30000 40000 50000
VLP-fraction read coverage
0 10 20 30
10000 20000 30000 40000 50000
Contig position (bp)
Oscillospiraceae Lawsonibacter

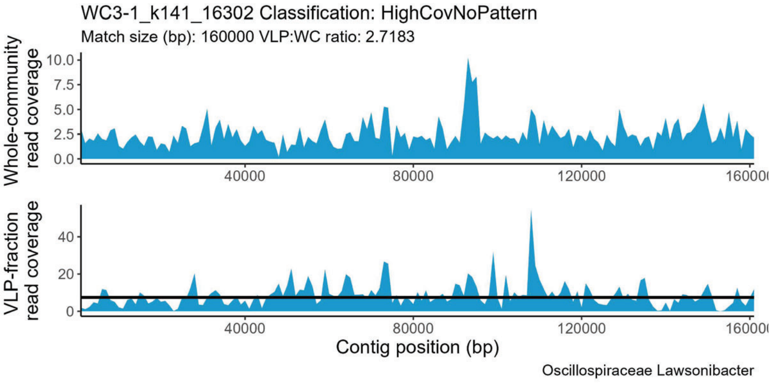
WC3-1_k141_16302 Classification: HighCovNoPattern
Match size (bp): 160000 VLP:WC ratio: 2.7183
Whole-community read coverage
0.0 2.5 5.0 7.5 10.0
40000 80000 120000
VLP-fraction read coverage
0 20 40
40000 80000 120000
Contig position (bp)
Oscillospiraceae Lawsonibacter

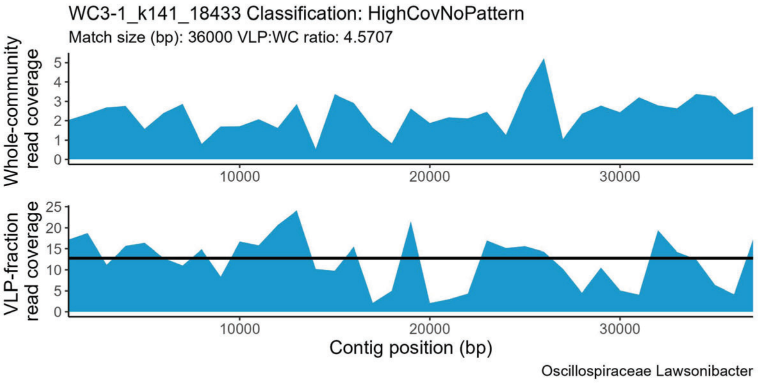
WC3-1_k141_18433 Classification: HighCovNoPattern
Match size (bp): 36000 VLP:WC ratio: 4.5707
Whole-community read coverage
VLP-fraction read coverage
Contig position (bp)
Oscillospiraceae Lawsonibacter

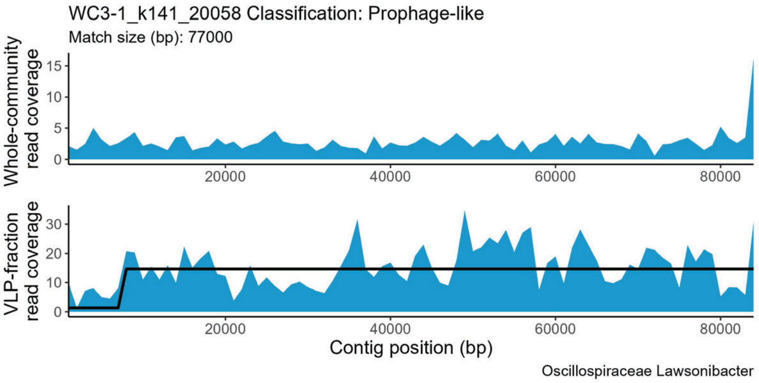
WC3-1_k141_20058 Classification: Prophage-like
Match size (bp): 77000
Whole-community read coverage
VLP-fraction read coverage
Contig position (bp)
Oscillospiraceae Lawsonibacter

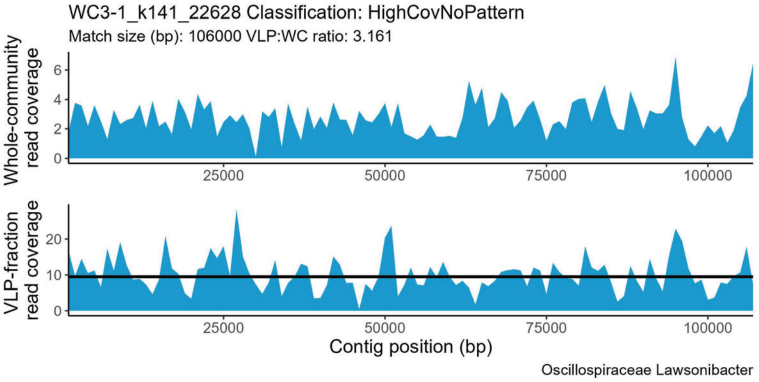
WC3-1_k141_22628 Classification: HighCovNoPattern
Match size (bp): 106000 VLP:WC ratio: 3.161
Whole-community read coverage
VLP-fraction read coverage
Contig position (bp)
Oscillospiraceae Lawsonibacter

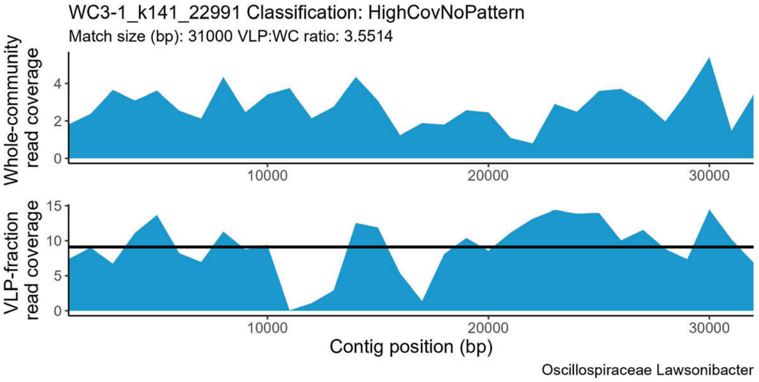
WC3-1_k141_22991 Classification: HighCovNoPattern
Match size (bp): 31000 VLP:WC ratio: 3.5514
Whole-community read coverage
VLP-fraction read coverage
Contig position (bp)
Oscillospiraceae Lawsonibacter

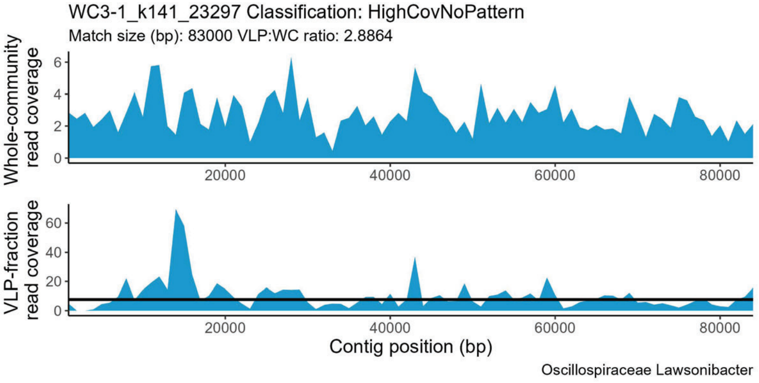
WC3-1_k141_23297 Classification: HighCovNoPattern
Match size (bp): 83000 VLP:WC ratio: 2.8864
Whole-community read coverage
VLP-fraction read coverage
Contig position (bp)
Oscillospiraceae Lawsonibacter

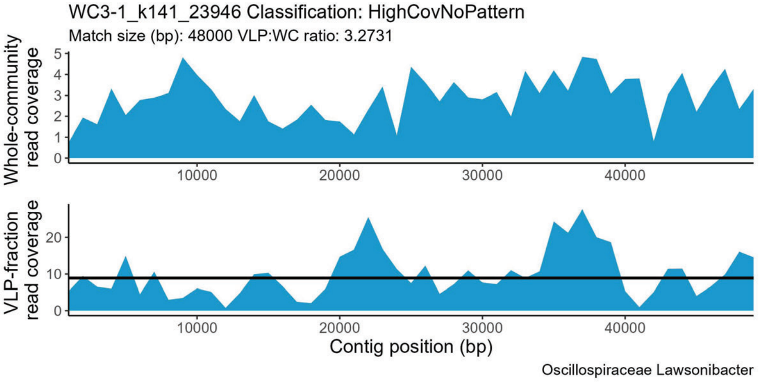
WC3-1_k141_23946 Classification: HighCovNoPattern
Match size (bp): 48000 VLP:WC ratio: 3.2731
Whole-community read coverage
VLP-fraction read coverage
Contig position (bp)
Oscillospiraceae Lawsonibacter

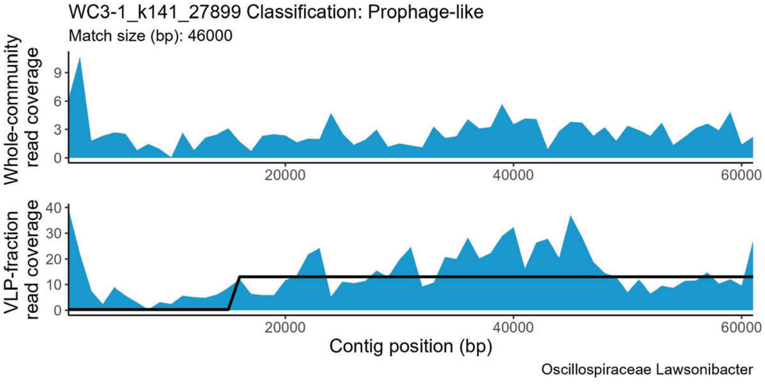
WC3-1_k141_27899 Classification: Prophage-like
Match size (bp): 46000
Whole-community read coverage
VLP-fraction read coverage
Contig position (bp)
Oscillospiraceae Lawsonibacter

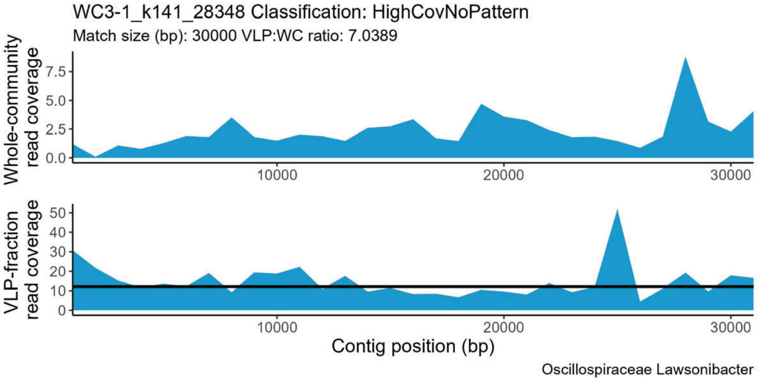
WC3-1_k141_28348 Classification: HighCovNoPattern
Match size (bp): 30000 VLP:WC ratio: 7.0389
Whole-community read coverage
VLP-fraction read coverage
Contig position (bp)
Oscillospiraceae Lawsonibacter

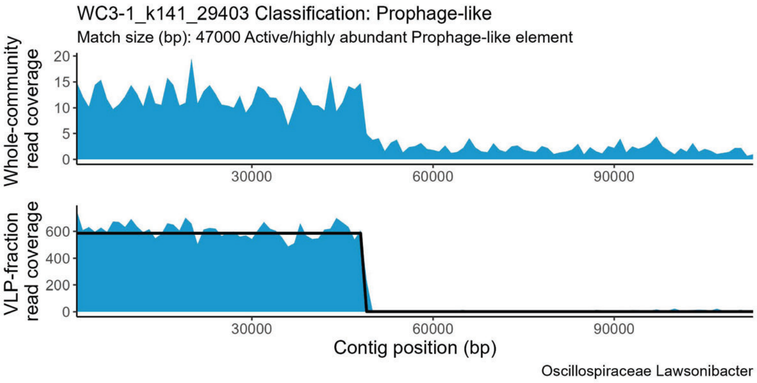
WC3-1_k141_29403 Classification: Prophage-like
Match size (bp): 47000 Active/highly abundant Prophage-like element
Whole-community read coverage
VLP-fraction read coverage
Contig position (bp)
Oscillospiraceae Lawsonibacter

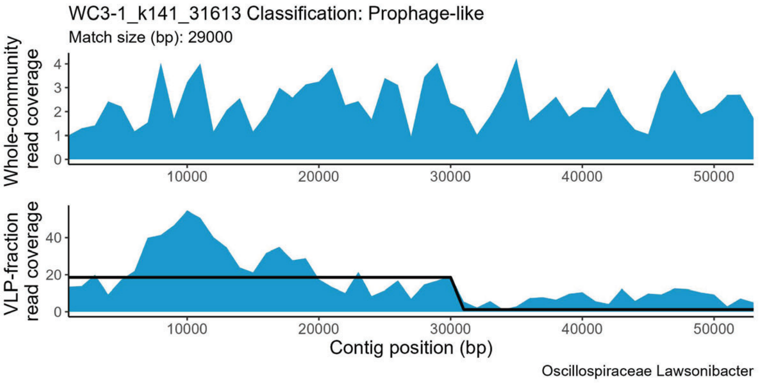
WC3-1_k141_31613 Classification: Prophage-like
Match size (bp): 29000
Whole-community read coverage
VLP-fraction read coverage
Contig position (bp)
Oscillospiraceae Lawsonibacter

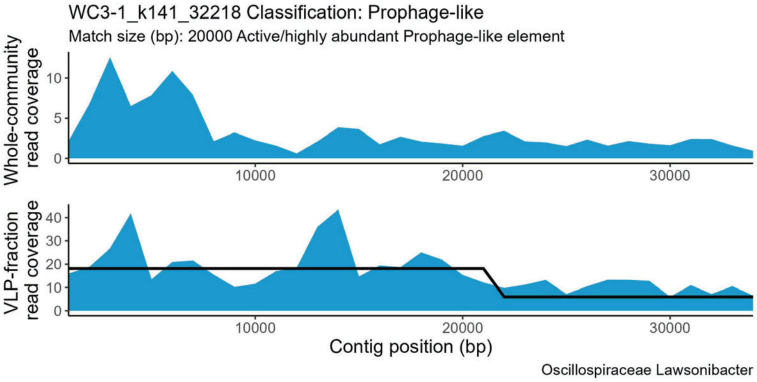
WC3-1_k141_32218 Classification: Prophage-like
Match size (bp): 20000 Active/highly abundant Prophage-like element
Whole-community read coverage
VLP-fraction read coverage
Contig position (bp)
Oscillospiraceae Lawsonibacter

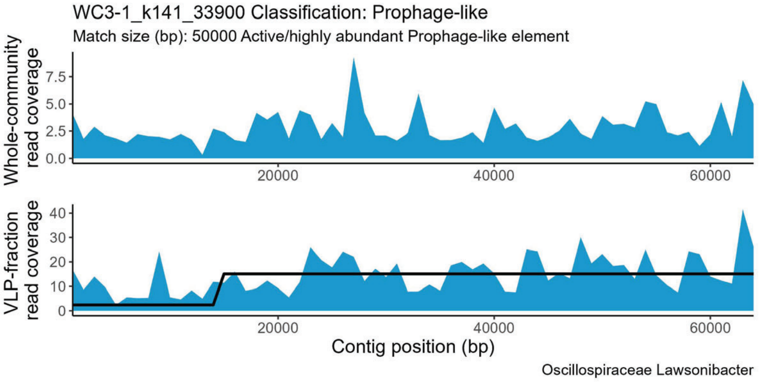
WC3-1_k141_33900 Classification: Prophage-like
Match size (bp): 50000 Active/highly abundant Prophage-like element
Whole-community read coverage
VLP-fraction read coverage
Contig position (bp)
Oscillospiraceae Lawsonibacter

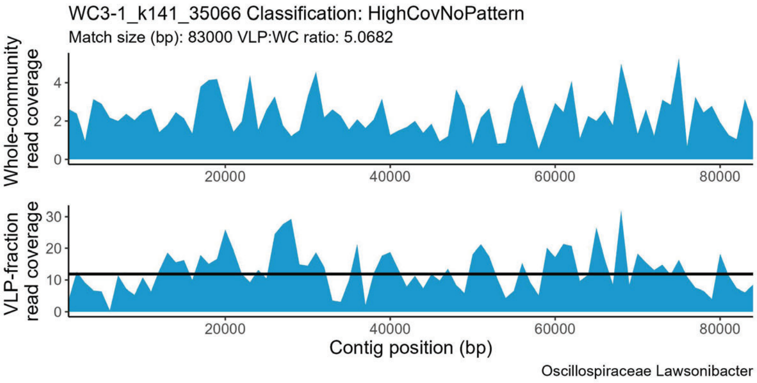
WC3-1_k141_35066 Classification: HighCovNoPattern
Match size (bp): 83000 VLP:WC ratio: 5.0682
Whole-community read coverage
VLP-fraction read coverage
Contig position (bp)
Oscillospiraceae Lawsonibacter

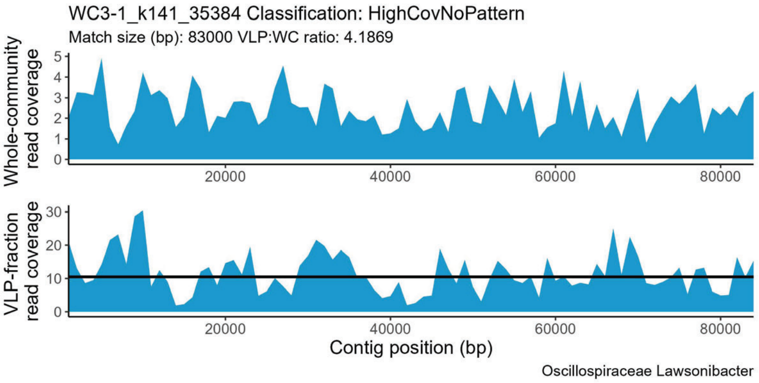
WC3-1_k141_35384 Classification: HighCovNoPattern
Match size (bp): 83000 VLP:WC ratio: 4.1869
Whole-community read coverage
VLP-fraction read coverage
Contig position (bp)
Oscillospiraceae Lawsonibacter

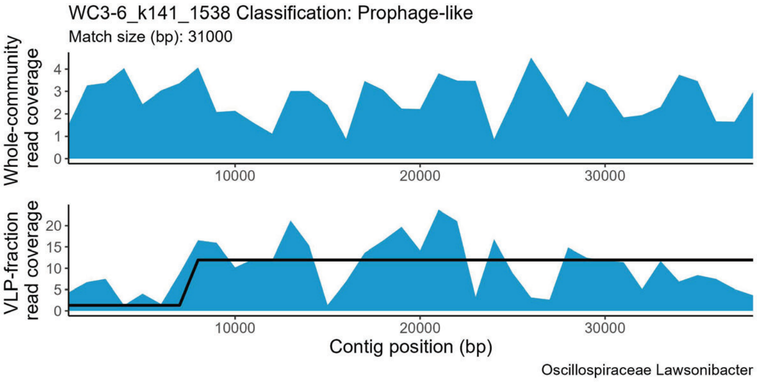
WC3-6_k141_1538 Classification: Prophage-like
Match size (bp): 31000
Whole-community read coverage
VLP-fraction read coverage
Contig position (bp)
Oscillospiraceae Lawsonibacter

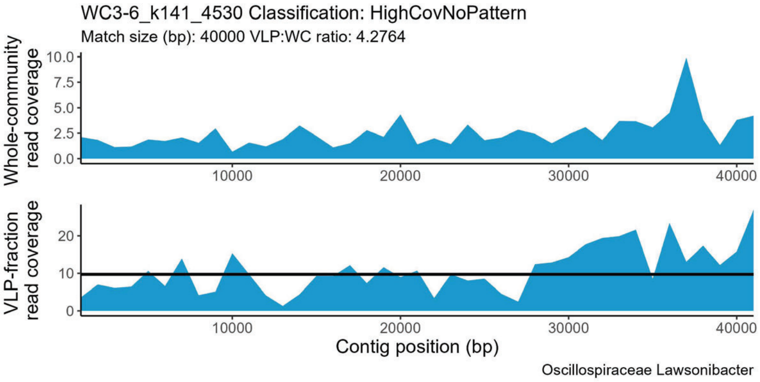
WC3-6_k141_4530 Classification: HighCovNoPattern
Match size (bp): 40000 VLP:WC ratio: 4.2764
Whole-community read coverage
VLP-fraction read coverage
Contig position (bp)
Oscillospiraceae Lawsonibacter

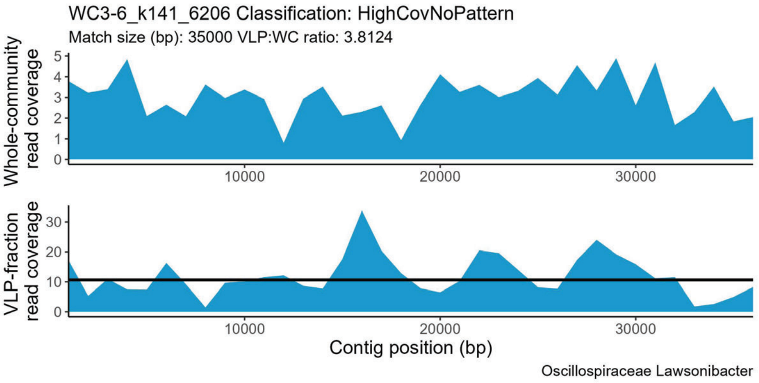
WC3-6_k141_6206 Classification: HighCovNoPattern
Match size (bp): 35000 VLP:WC ratio: 3.8124
Whole-community read coverage
VLP-fraction read coverage
Contig position (bp)
Oscillospiraceae Lawsonibacter

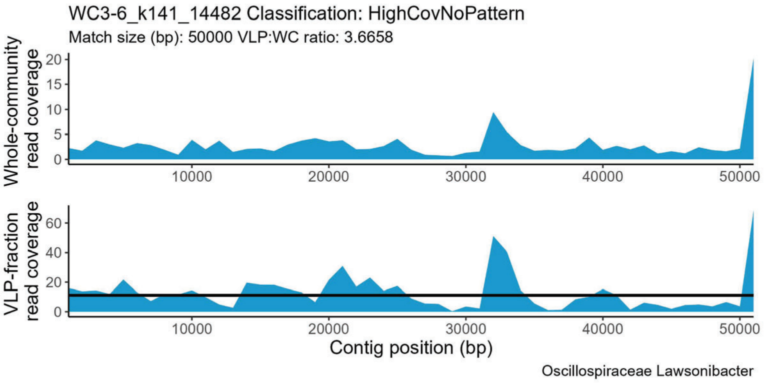
WC3-6_k141_14482 Classification: HighCovNoPattern
Match size (bp): 50000 VLP:WC ratio: 3.6658
Whole-community read coverage
VLP-fraction read coverage
Contig position (bp)
Oscillospiraceae Lawsonibacter

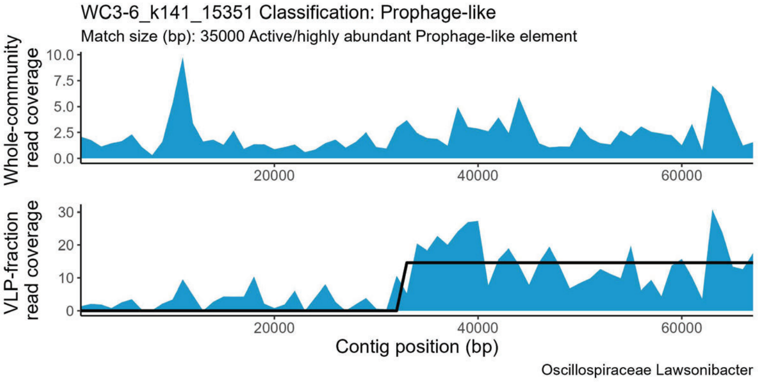
WC3-6_k141_15351 Classification: Prophage-like
Match size (bp): 35000 Active/highly abundant Prophage-like element
Whole-community read coverage
VLP-fraction read coverage
Contig position (bp)
Oscillospiraceae Lawsonibacter

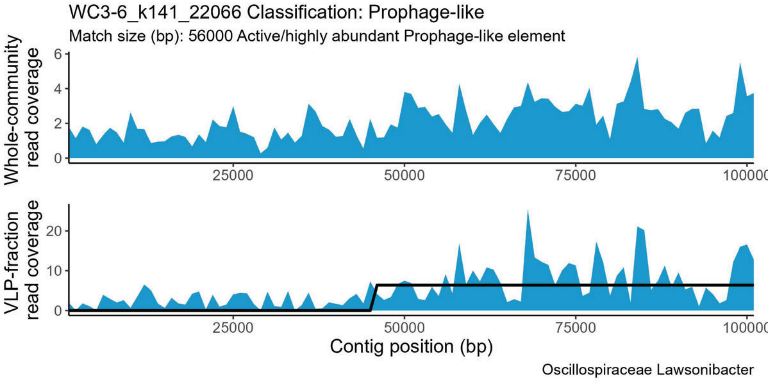
WC3-6_k141_22066 Classification: Prophage-like
Match size (bp): 56000 Active/highly abundant Prophage-like element
Whole-community read coverage
VLP-fraction read coverage
Contig position (bp)
Oscillospiraceae Lawsonibacter

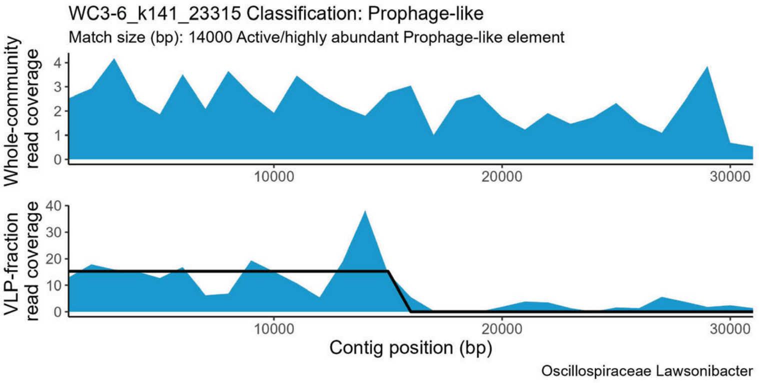
WC3-6_k141_23315 Classification: Prophage-like
Match size (bp): 14000 Active/highly abundant Prophage-like element
Whole-community read coverage
VLP-fraction read coverage
Contig position (bp)
Oscillospiraceae Lawsonibacter

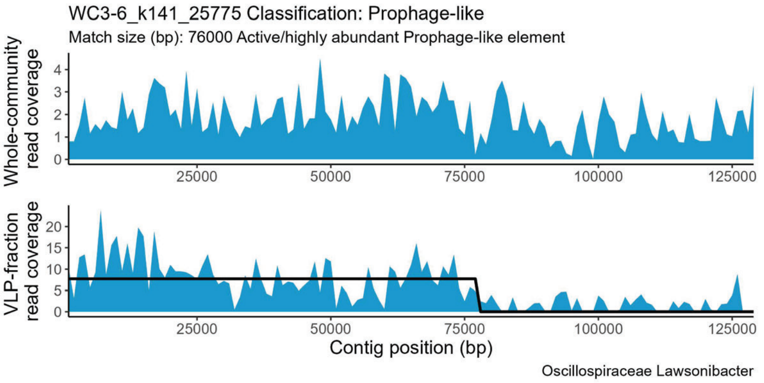
WC3-6_k141_25775 Classification: Prophage-like
Match size (bp): 76000 Active/highly abundant Prophage-like element
Whole-community read coverage
VLP-fraction read coverage
Contig position (bp)
Oscillospiraceae Lawsonibacter

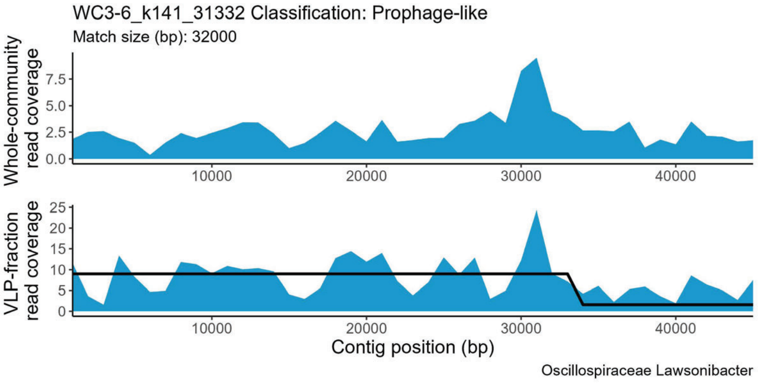
WC3-6_k141_31332 Classification: Prophage-like
Match size (bp): 32000
Whole-community read coverage
VLP-fraction read coverage
Contig position (bp)
Oscillospiraceae Lawsonibacter

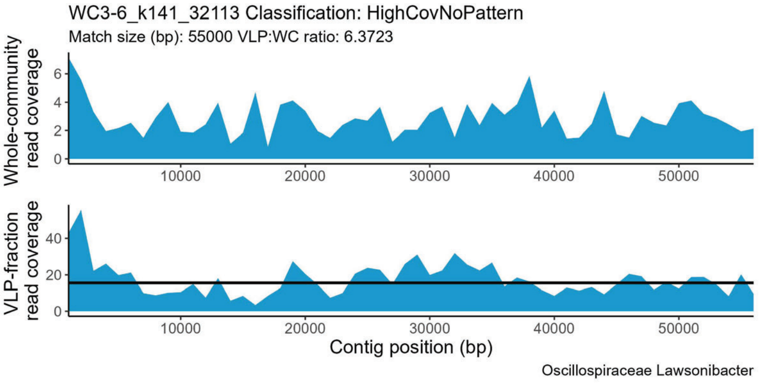
WC3-6_k141_32113 Classification: HighCovNoPattern
Match size (bp): 55000 VLP:WC ratio: 6.3723
Whole-community read coverage
VLP-fraction read coverage
Contig position (bp)
Oscillospiraceae Lawsonibacter

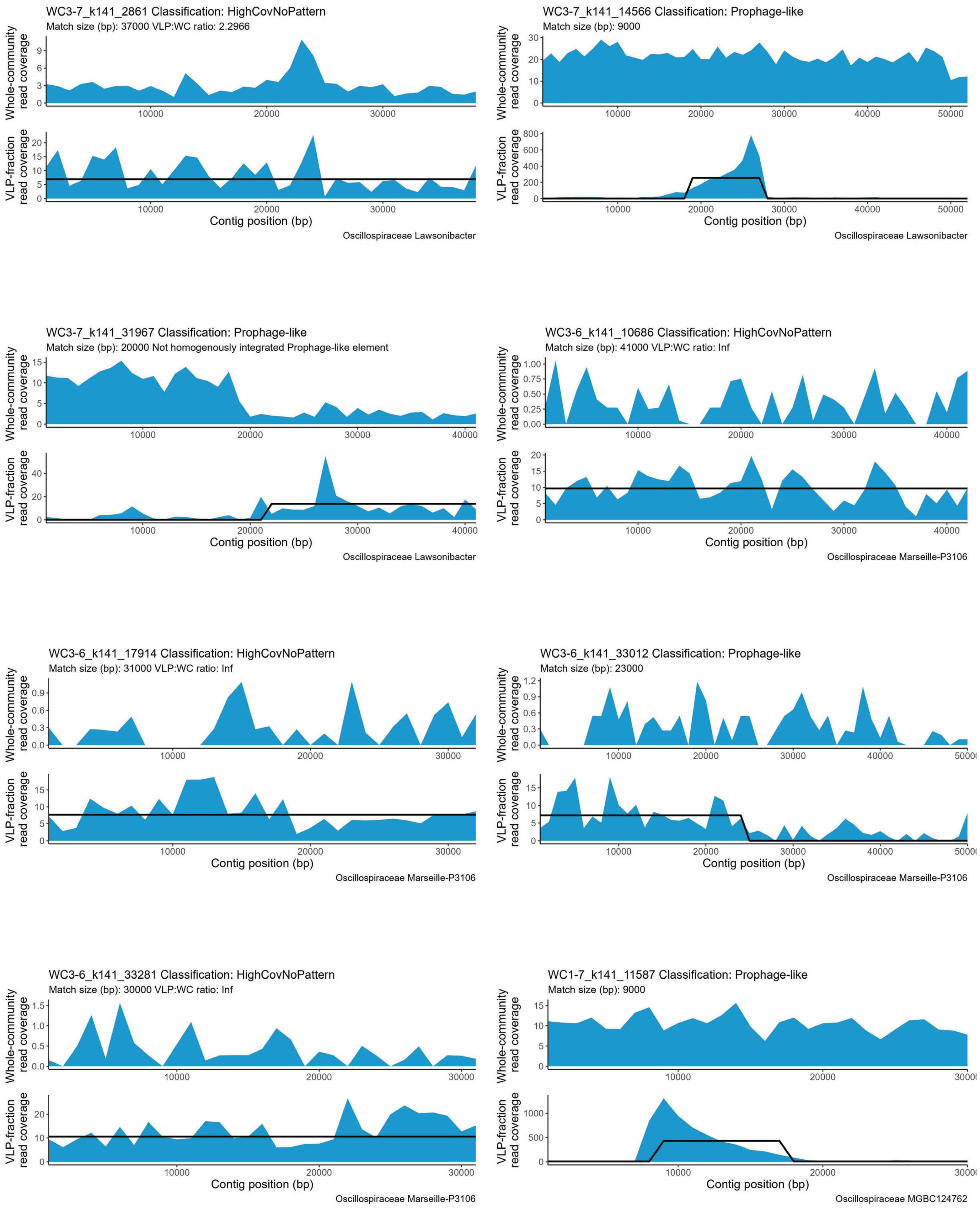
WC3-7_k141_2861 Classification: HighCovNoPattern
Match size (bp): 37000 VLP:WC ratio: 2.2966
Whole-community read coverage
VLP-fraction read coverage
Contig position (bp)
Oscillospiraceae Lawsonibacter
WC3-7_k141_14566 Classification: Prophage-like
Match size (bp): 9000
Whole-community read coverage
VLP-fraction read coverage
Contig position (bp)
Oscillospiraceae Lawsonibacter
WC3-7_k141_31967 Classification: Prophage-like
Match size (bp): 20000 Not homogenously integrated Prophage-like element
Whole-community read coverage
VLP-fraction read coverage
Contig position (bp)
Oscillospiraceae Lawsonibacter
WC3-6_k141_10686 Classification: HighCovNoPattern
Match size (bp): 41000 VLP:WC ratio: Inf
Whole-community read coverage
VLP-fraction read coverage
Contig position (bp)
Oscillospiraceae Marseille-P3106
WC3-6_k141_17914 Classification: HighCovNoPattern
Match size (bp): 31000 VLP:WC ratio: Inf
Whole-community read coverage
VLP-fraction read coverage
Contig position (bp)
Oscillospiraceae Marseille-P3106
WC3-6_k141_33012 Classification: Prophage-like
Match size (bp): 23000
Whole-community read coverage
VLP-fraction read coverage
Contig position (bp)
Oscillospiraceae Marseille-P3106
WC3-6_k141_33281 Classification: HighCovNoPattern
Match size (bp): 30000 VLP:WC ratio: Inf
Whole-community read coverage
VLP-fraction read coverage
Contig position (bp)
Oscillospiraceae Marseille-P3106
WC1-7_k141_11587 Classification: Prophage-like
Match size (bp): 9000
Whole-community read coverage
VLP-fraction read coverage
Contig position (bp)
Oscillospiraceae MGBC124762

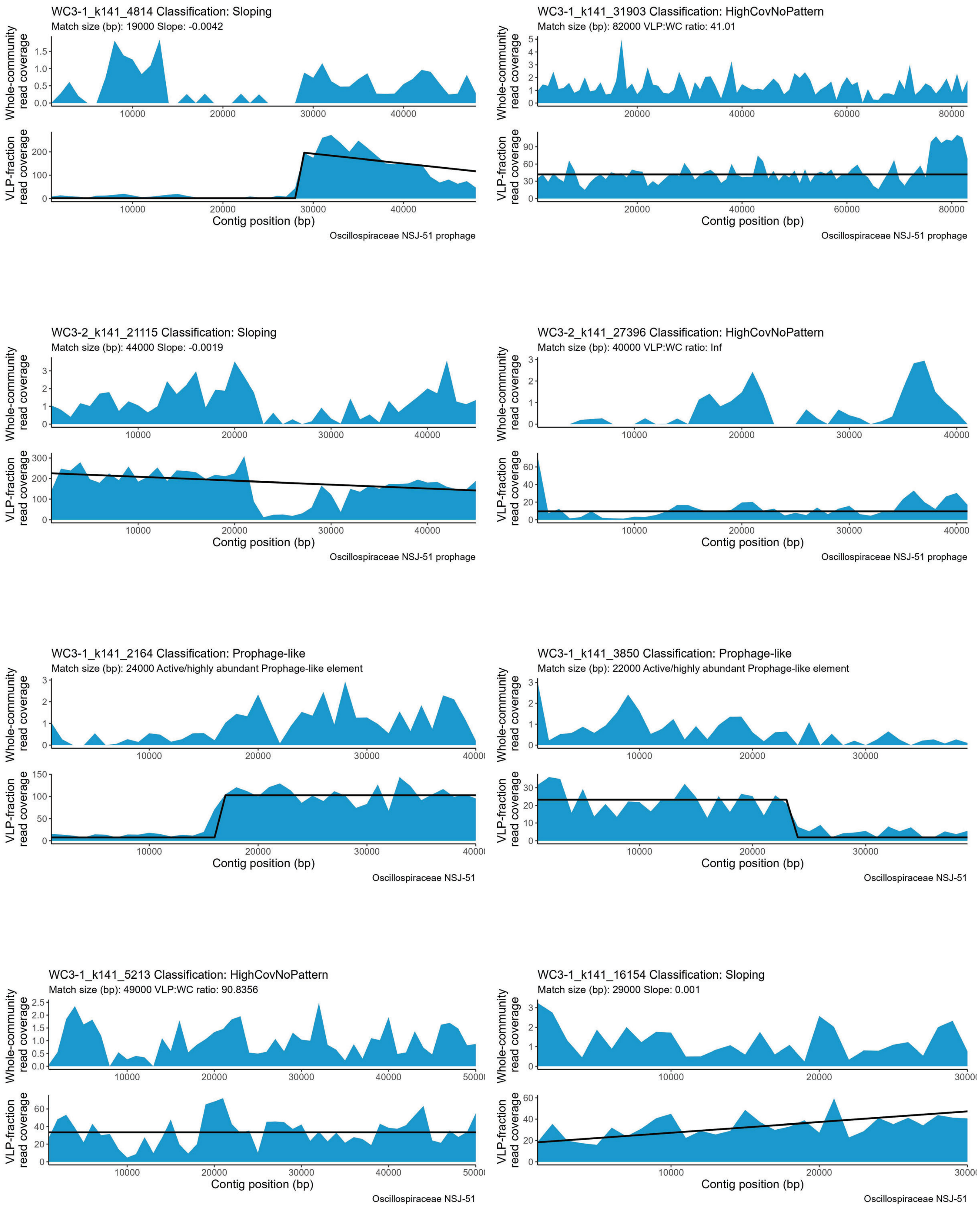

WC3-1_k141_4814 Classification: Sloping
Match size (bp): 19000 Slope: -0.0042
Whole-community read coverage
VLP-fraction read coverage
Contig position (bp)
Oscillospiraceae NSJ-51 prophage
WC3-1_k141_31903 Classification: HighCovNoPattern
Match size (bp): 82000 VLP:WC ratio: 41.01
Whole-community read coverage
VLP-fraction read coverage
Contig position (bp)
Oscillospiraceae NSJ-51 prophage
WC3-2_k141_21115 Classification: Sloping
Match size (bp): 44000 Slope: -0.0019
Whole-community read coverage
VLP-fraction read coverage
Contig position (bp)
Oscillospiraceae NSJ-51 prophage
WC3-2_k141_27396 Classification: HighCovNoPattern
Match size (bp): 40000 VLP:WC ratio: Inf
Whole-community read coverage
VLP-fraction read coverage
Contig position (bp)
Oscillospiraceae NSJ-51 prophage
WC3-1_k141_2164 Classification: Prophage-like
Match size (bp): 24000 Active/highly abundant Prophage-like element
Whole-community read coverage
VLP-fraction read coverage
Contig position (bp)
Oscillospiraceae NSJ-51
WC3-1_k141_3850 Classification: Prophage-like
Match size (bp): 22000 Active/highly abundant Prophage-like element
Whole-community read coverage
VLP-fraction read coverage
Contig position (bp)
Oscillospiraceae NSJ-51
WC3-1_k141_5213 Classification: HighCovNoPattern
Match size (bp): 49000 VLP:WC ratio: 90.8356
Whole-community read coverage
VLP-fraction read coverage
Contig position (bp)
Oscillospiraceae NSJ-51
WC3-1_k141_16154 Classification: Sloping
Match size (bp): 29000 Slope: 0.001
Whole-community read coverage
VLP-fraction read coverage
Contig position (bp)
Oscillospiraceae NSJ-51

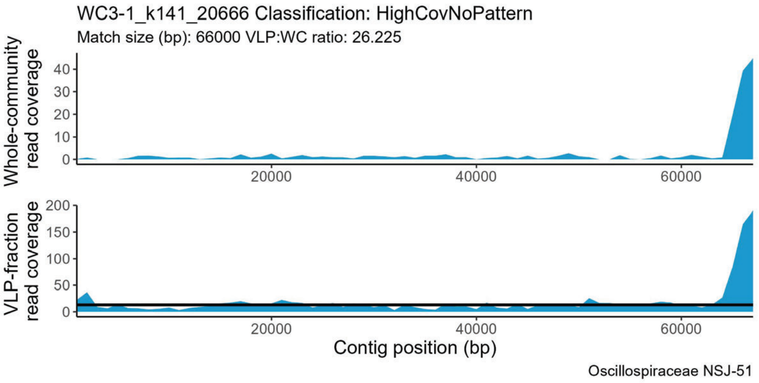
WC3-1_k141_20666 Classification: HighCovNoPattern
Match size (bp): 66000 VLP:WC ratio: 26.225
Whole-community read coverage
VLP-fraction read coverage
Contig position (bp)
Oscillospiraceae NSJ-51

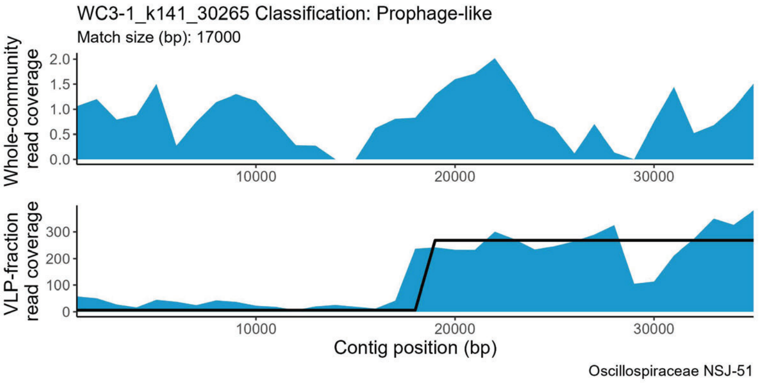
WC3-1_k141_30265 Classification: Prophage-like
Match size (bp): 17000
Whole-community read coverage
VLP-fraction read coverage
Contig position (bp)
Oscillospiraceae NSJ-51

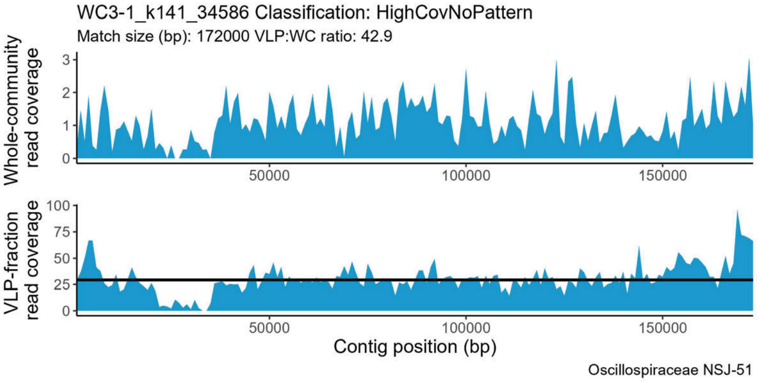
WC3-1_k141_34586 Classification: HighCovNoPattern
Match size (bp): 172000 VLP:WC ratio: 42.9
Whole-community read coverage
VLP-fraction read coverage
Contig position (bp)
Oscillospiraceae NSJ-51

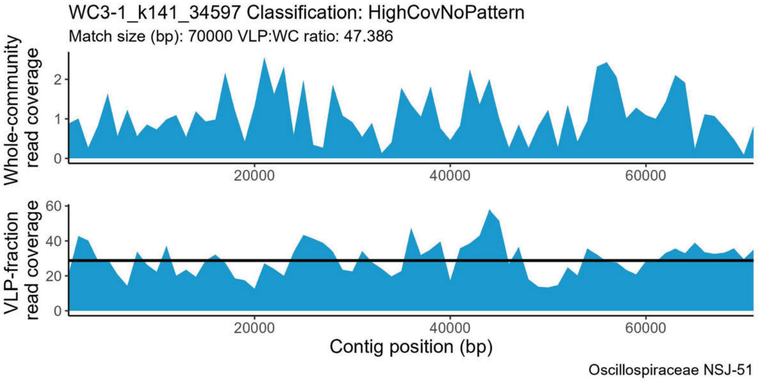
WC3-1_k141_34597 Classification: HighCovNoPattern
Match size (bp): 70000 VLP:WC ratio: 47.386
Whole-community read coverage
VLP-fraction read coverage
Contig position (bp)
Oscillospiraceae NSJ-51

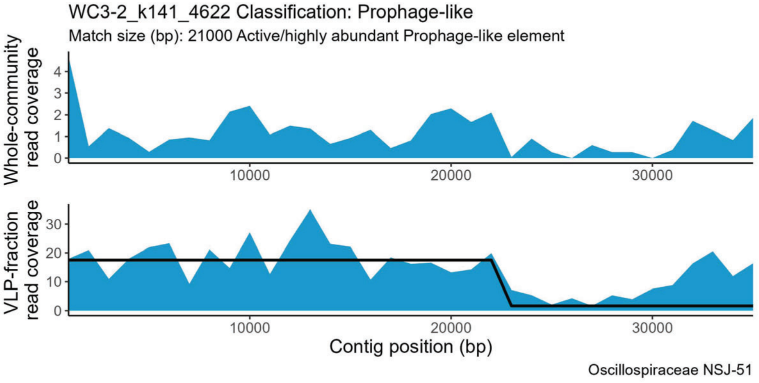
WC3-2_k141_4622 Classification: Prophage-like
Match size (bp): 21000 Active/highly abundant Prophage-like element
Whole-community read coverage
VLP-fraction read coverage
Contig position (bp)
Oscillospiraceae NSJ-51

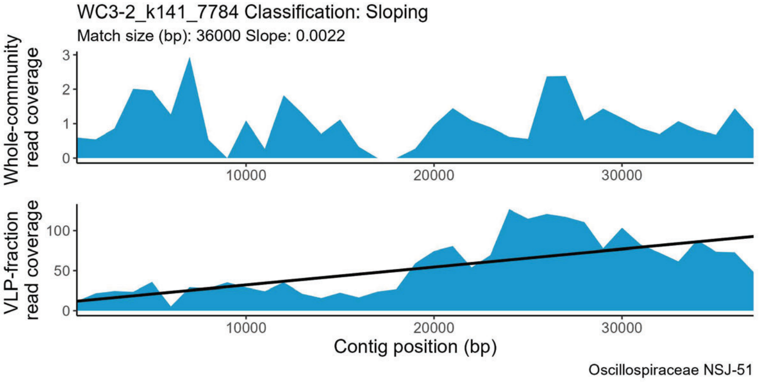
WC3-2_k141_7784 Classification: Sloping
Match size (bp): 36000 Slope: 0.0022
Whole-community read coverage
VLP-fraction read coverage
Contig position (bp)
Oscillospiraceae NSJ-51

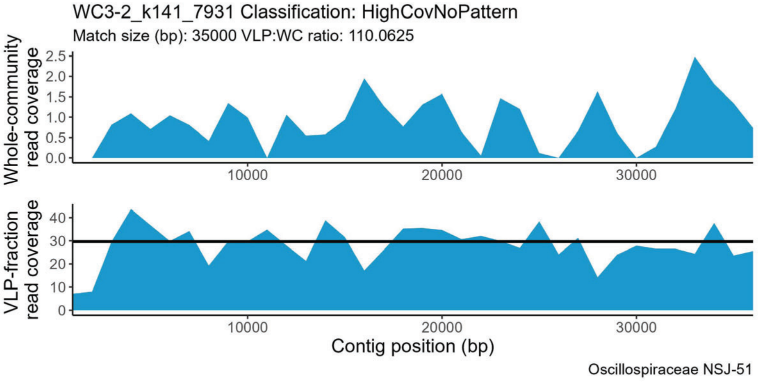
WC3-2_k141_7931 Classification: HighCovNoPattern
Match size (bp): 35000 VLP:WC ratio: 110.0625
Whole-community read coverage
VLP-fraction read coverage
Contig position (bp)
Oscillospiraceae NSJ-51

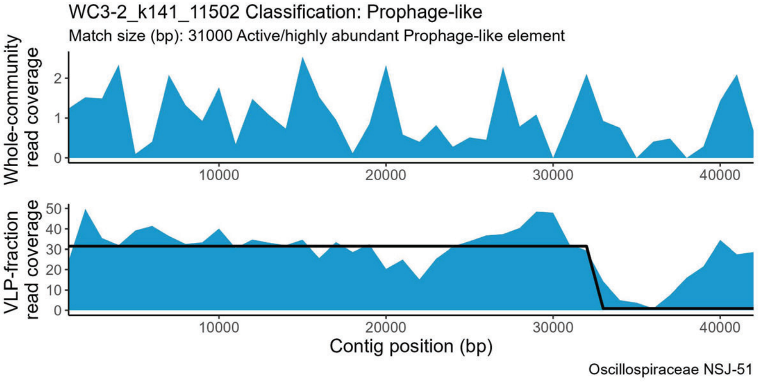
WC3-2_k141_11502 Classification: Prophage-like
Match size (bp): 31000 Active/highly abundant Prophage-like element
Whole-community read coverage
VLP-fraction read coverage
Contig position (bp)
Oscillospiraceae NSJ-51

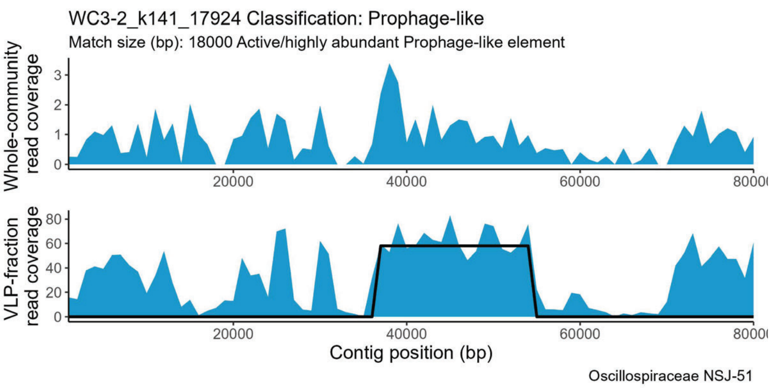
WC3-2_k141_17924 Classification: Prophage-like
Match size (bp): 18000 Active/highly abundant Prophage-like element
Whole-community read coverage
VLP-fraction read coverage
Contig position (bp)
Oscillospiraceae NSJ-51

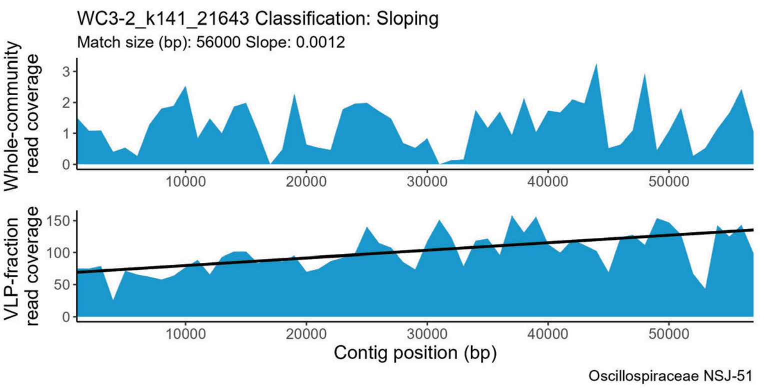
WC3-2_k141_21643 Classification: Sloping
Match size (bp): 56000 Slope: 0.0012
Whole-community read coverage
VLP-fraction read coverage
Contig position (bp)
Oscillospiraceae NSJ-51

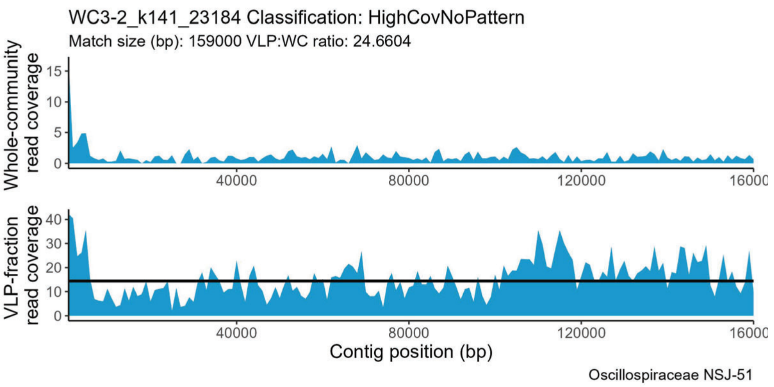
WC3-2_k141_23184 Classification: HighCovNoPattern
Match size (bp): 159000 VLP:WC ratio: 24.6604
Whole-community read coverage
VLP-fraction read coverage
Contig position (bp)
Oscillospiraceae NSJ-51

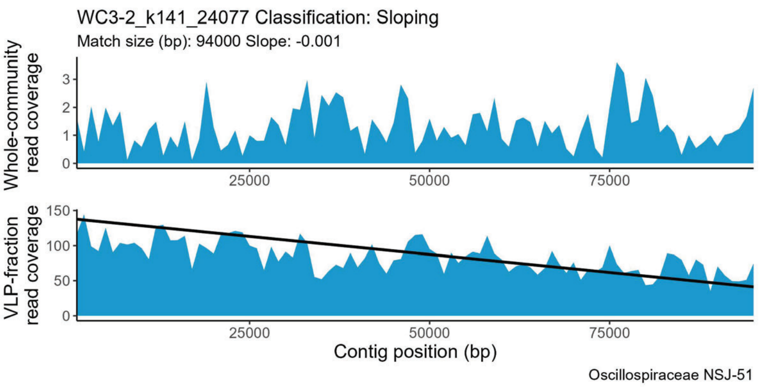
WC3-2_k141_24077 Classification: Sloping
Match size (bp): 94000 Slope: -0.001
Whole-community read coverage
VLP-fraction read coverage
Contig position (bp)
Oscillospiraceae NSJ-51

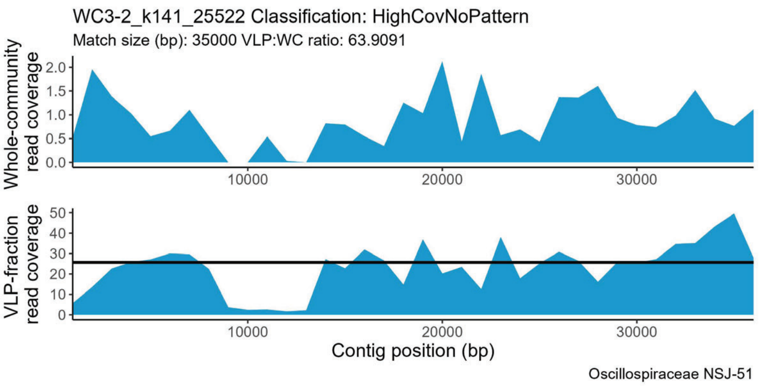
WC3-2_k141_25522 Classification: HighCovNoPattern
Match size (bp): 35000 VLP:WC ratio: 63.9091
Whole-community read coverage
VLP-fraction read coverage
Contig position (bp)
Oscillospiraceae NSJ-51

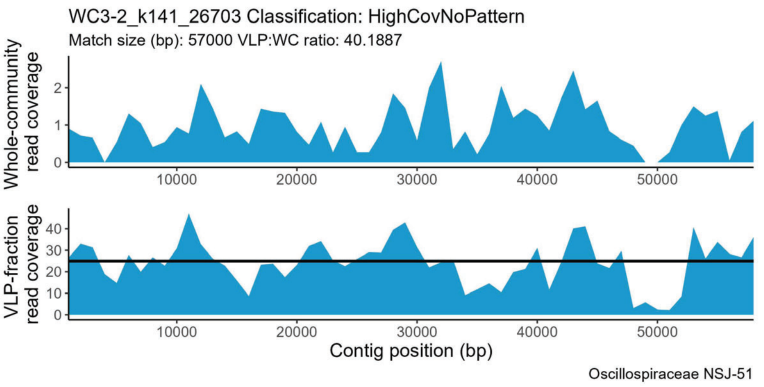
WC3-2_k141_26703 Classification: HighCovNoPattern
Match size (bp): 57000 VLP:WC ratio: 40.1887
Whole-community read coverage
VLP-fraction read coverage
Contig position (bp)
Oscillospiraceae NSJ-51

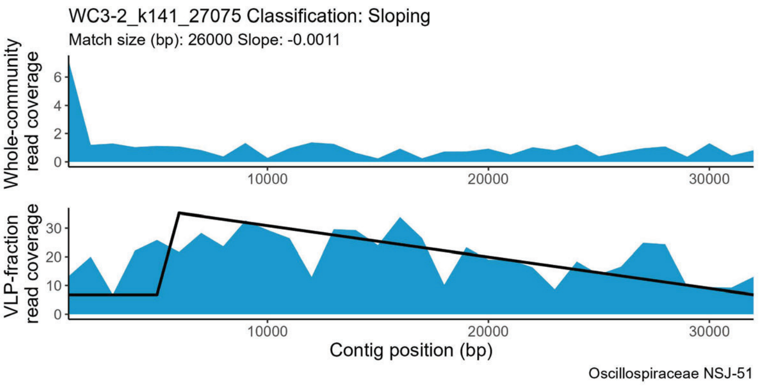
WC3-2_k141_27075 Classification: Sloping
Match size (bp): 26000 Slope: -0.0011
Whole-community read coverage
VLP-fraction read coverage
Contig position (bp)
Oscillospiraceae NSJ-51

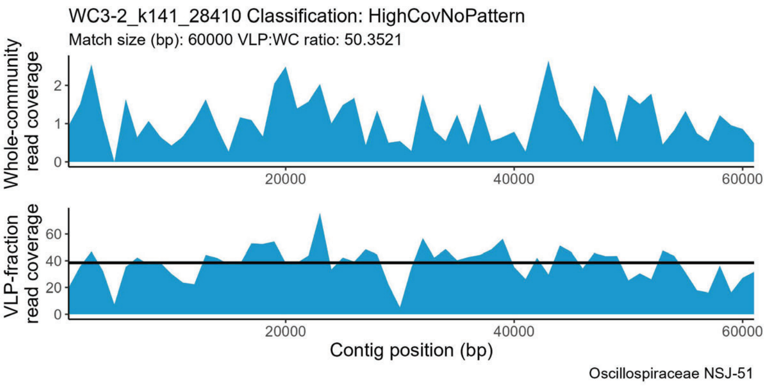
WC3-2_k141_28410 Classification: HighCovNoPattern
Match size (bp): 60000 VLP:WC ratio: 50.3521
Whole-community read coverage
VLP-fraction read coverage
Contig position (bp)
Oscillospiraceae NSJ-51

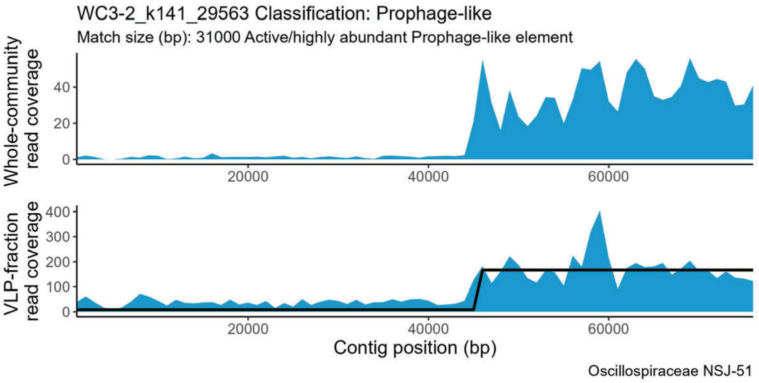
WC3-2_k141_29563 Classification: Prophage-like
Match size (bp): 31000 Active/highly abundant Prophage-like element
Whole-community read coverage
0
20
40
20000
40000
60000
VLP-fraction read coverage
0
100
200
300
400
Contig position (bp)
Oscillospiraceae NSJ-51

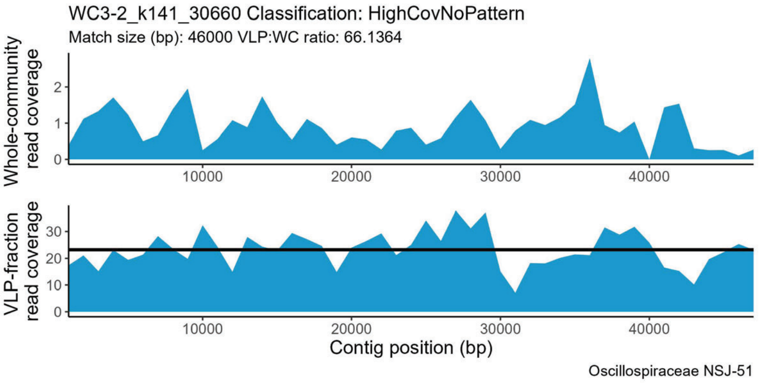
WC3-2_k141_30660 Classification: HighCovNoPattern
Match size (bp): 46000 VLP:WC ratio: 66.1364
Whole-community read coverage
0
1
2
10000
20000
30000
40000
VLP-fraction read coverage
0
10
20
30
Contig position (bp)
Oscillospiraceae NSJ-51

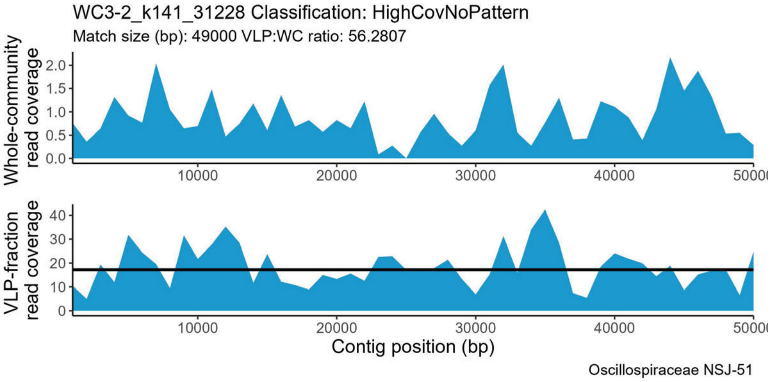
WC3-2_k141_31228 Classification: HighCovNoPattern
Match size (bp): 49000 VLP:WC ratio: 56.2807
Whole-community read coverage
0.0
0.5
1.0
1.5
2.0
10000
20000
30000
40000
VLP-fraction read coverage
0
10
20
30
40
Contig position (bp)
Oscillospiraceae NSJ-51

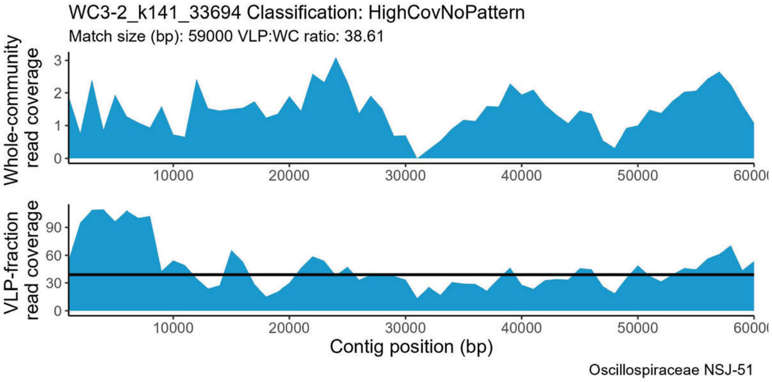
WC3-2_k141_33694 Classification: HighCovNoPattern
Match size (bp): 59000 VLP:WC ratio: 38.61
Whole-community read coverage
0
1
2
3
10000
20000
30000
40000
50000
VLP-fraction read coverage
0
30
60
90
Contig position (bp)
Oscillospiraceae NSJ-51

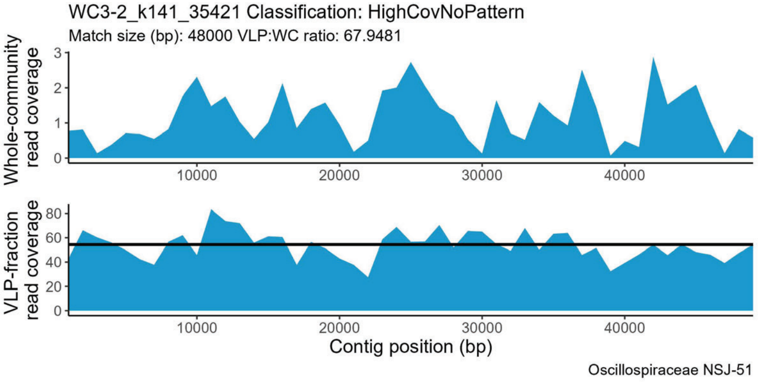
WC3-2_k141_35421 Classification: HighCovNoPattern
Match size (bp): 48000 VLP:WC ratio: 67.9481
Whole-community read coverage
0
1
2
3
10000
20000
30000
40000
VLP-fraction read coverage
0
20
40
60
80
Contig position (bp)
Oscillospiraceae NSJ-51

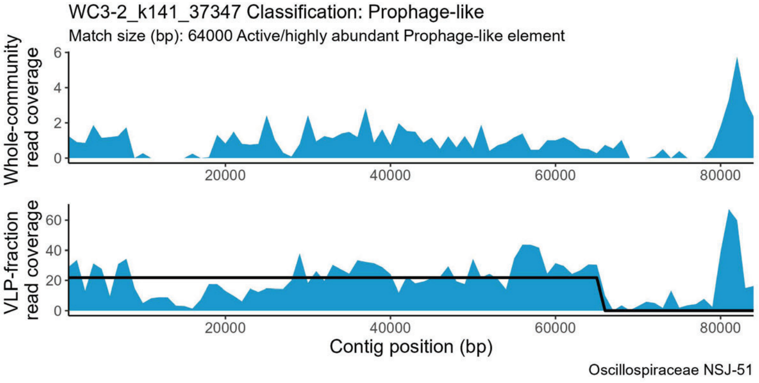
WC3-2_k141_37347 Classification: Prophage-like
Match size (bp): 64000 Active/highly abundant Prophage-like element
Whole-community read coverage
0
2
4
6
20000
40000
60000
80000
VLP-fraction read coverage
0
20
40
60
Contig position (bp)
Oscillospiraceae NSJ-51

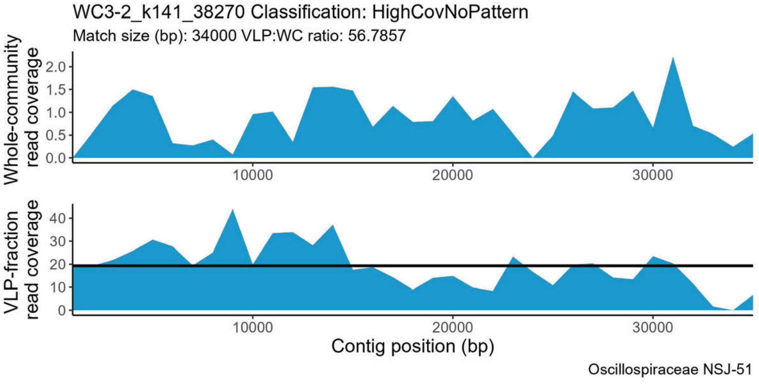
WC3-2_k141_38270 Classification: HighCovNoPattern
Match size (bp): 34000 VLP:WC ratio: 56.7857
Whole-community read coverage
0.0
0.5
1.0
1.5
2.0
10000
20000
30000
VLP-fraction read coverage
0
10
20
30
40
Contig position (bp)
Oscillospiraceae NSJ-51

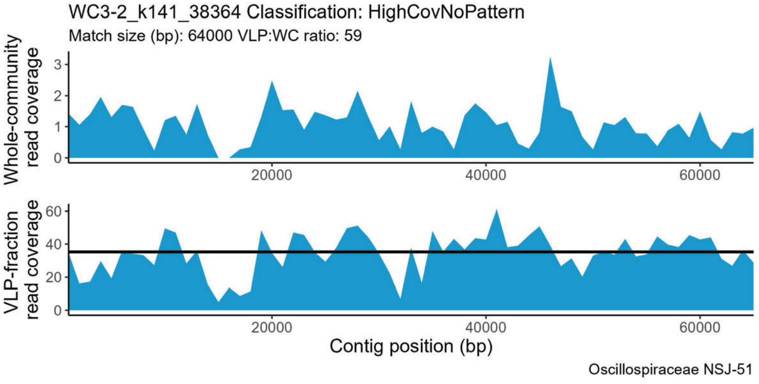
WC3-2_k141_38364 Classification: HighCovNoPattern
Match size (bp): 64000 VLP:WC ratio: 59
Whole-community read coverage
0
1
2
3
20000
40000
60000
VLP-fraction read coverage
0
20
40
60
Contig position (bp)
Oscillospiraceae NSJ-51

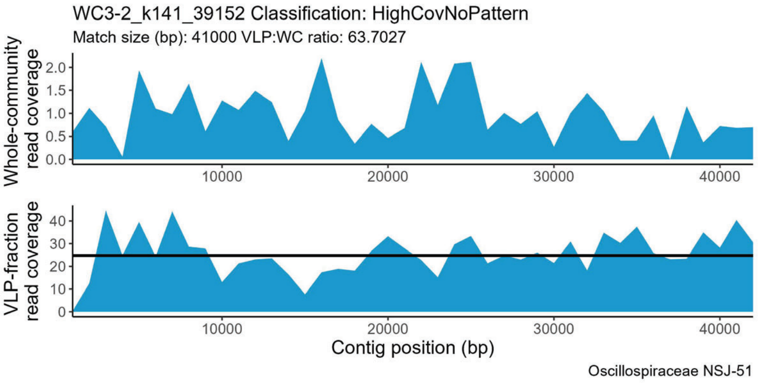
WC3-2_k141_39152 Classification: HighCovNoPattern
Match size (bp): 41000 VLP:WC ratio: 63.7027
Whole-community read coverage
VLP-fraction read coverage
Contig position (bp)
Oscillospiraceae NSJ-51

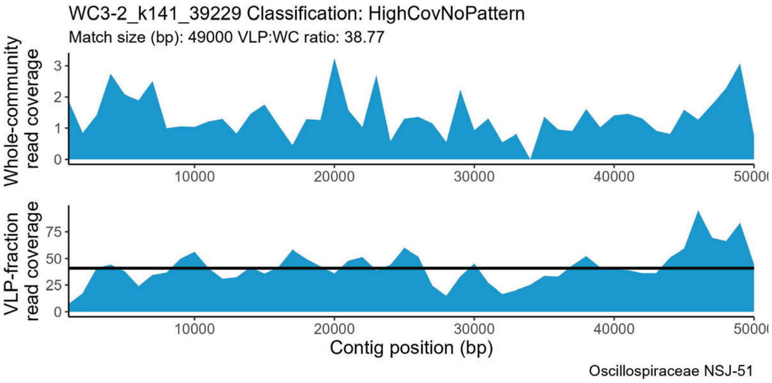
WC3-2_k141_39229 Classification: HighCovNoPattern
Match size (bp): 49000 VLP:WC ratio: 38.77
Whole-community read coverage
VLP-fraction read coverage
Contig position (bp)
Oscillospiraceae NSJ-51

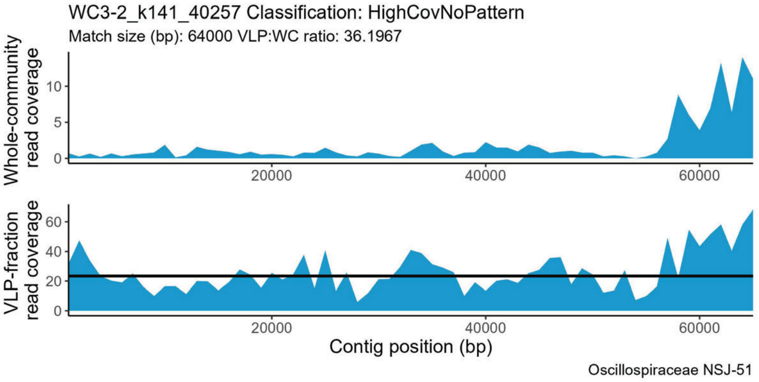
WC3-2_k141_40257 Classification: HighCovNoPattern
Match size (bp): 64000 VLP:WC ratio: 36.1967
Whole-community read coverage
VLP-fraction read coverage
Contig position (bp)
Oscillospiraceae NSJ-51

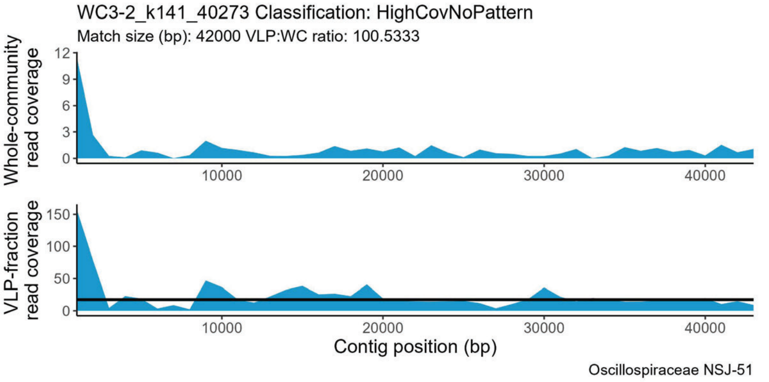
WC3-2_k141_40273 Classification: HighCovNoPattern
Match size (bp): 42000 VLP:WC ratio: 100.5333
Whole-community read coverage
VLP-fraction read coverage
Contig position (bp)
Oscillospiraceae NSJ-51

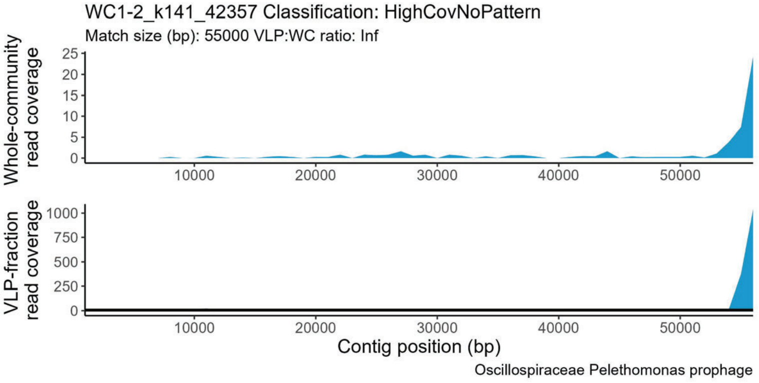
WC1-2_k141_42357 Classification: HighCovNoPattern
Match size (bp): 55000 VLP:WC ratio: Inf
Whole-community read coverage
VLP-fraction read coverage
Contig position (bp)
Oscillospiraceae Pelethomonas prophage

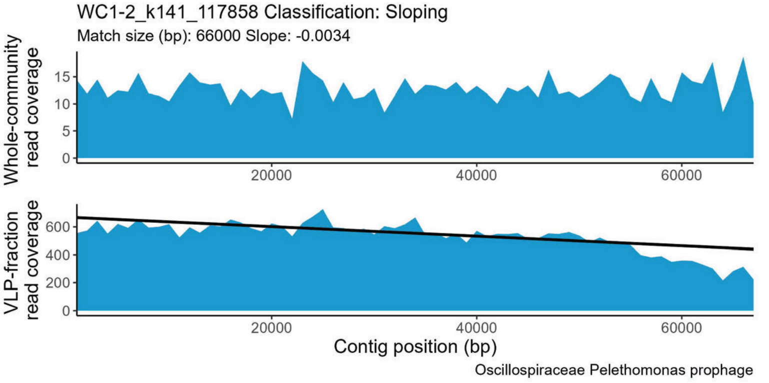
WC1-2_k141_117858 Classification: Sloping
Match size (bp): 66000 Slope: -0.0034
Whole-community read coverage
VLP-fraction read coverage
Contig position (bp)
Oscillospiraceae Pelethomonas prophage

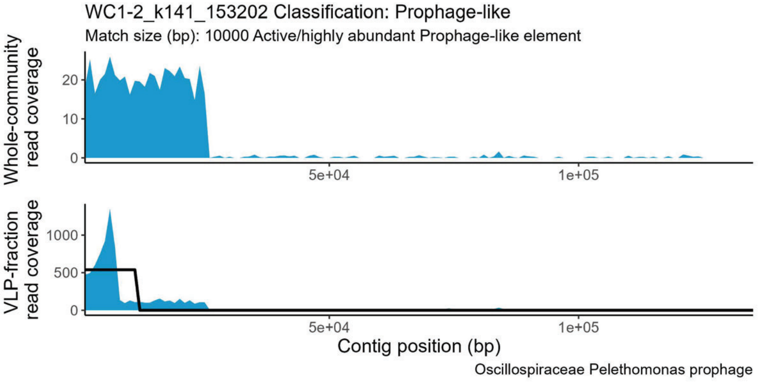
WC1-2_k141_153202 Classification: Prophage-like
Match size (bp): 10000 Active/highly abundant Prophage-like element
Whole-community read coverage
VLP-fraction read coverage
Contig position (bp)
Oscillospiraceae Pelethomonas prophage

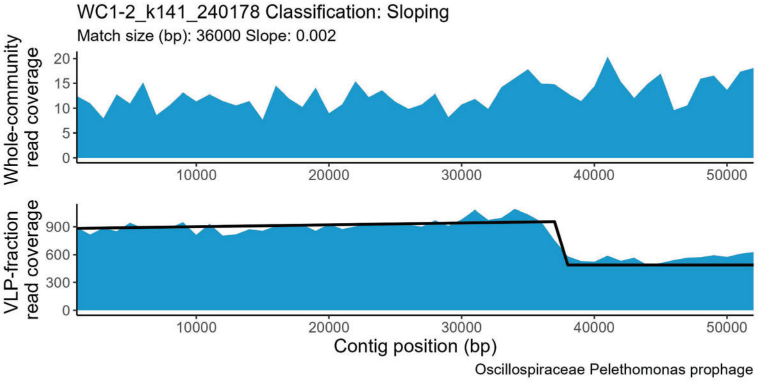
WC1-2_k141_240178 Classification: Sloping
Match size (bp): 36000 Slope: 0.002
Whole-community read coverage
VLP-fraction read coverage
Contig position (bp)
Oscillospiraceae Pelethomonas prophage

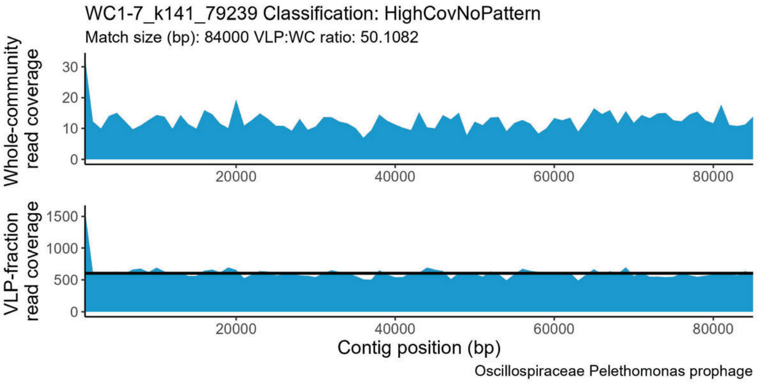
WC1-7_k141_79239 Classification: HighCovNoPattern
Match size (bp): 84000 VLP:WC ratio: 50.1082
Whole-community read coverage
VLP-fraction read coverage
Contig position (bp)
Oscillospiraceae Pelethomonas prophage

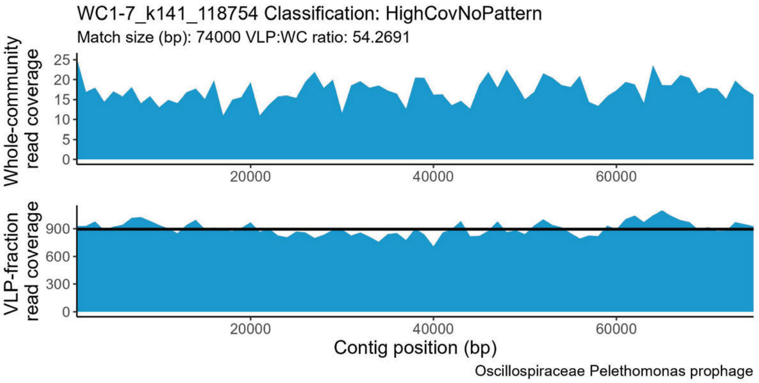
WC1-7_k141_118754 Classification: HighCovNoPattern
Match size (bp): 74000 VLP:WC ratio: 54.2691
Whole-community read coverage
VLP-fraction read coverage
Contig position (bp)
Oscillospiraceae Pelethomonas prophage

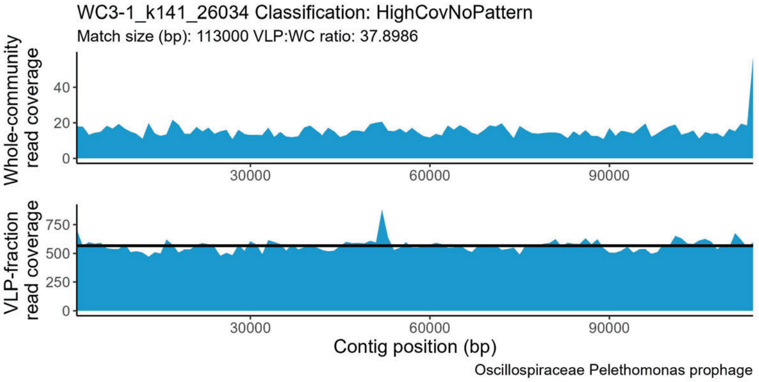
WC3-1_k141_26034 Classification: HighCovNoPattern
Match size (bp): 113000 VLP:WC ratio: 37.8986
Whole-community read coverage
VLP-fraction read coverage
Contig position (bp)
Oscillospiraceae Pelethomonas prophage

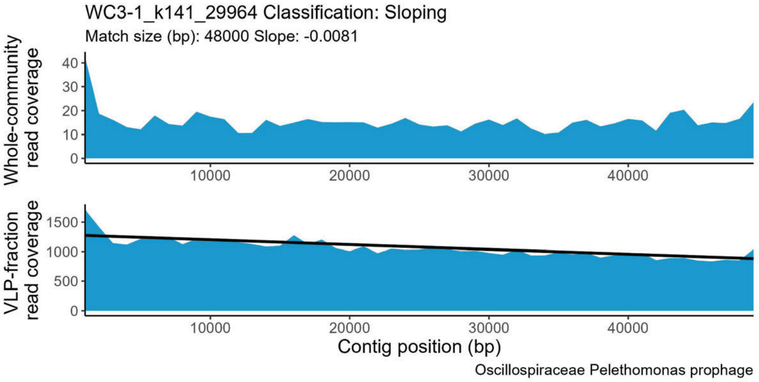
WC3-1_k141_29964 Classification: Sloping
Match size (bp): 48000 Slope: -0.0081
Whole-community read coverage
VLP-fraction read coverage
Contig position (bp)
Oscillospiraceae Pelethomonas prophage

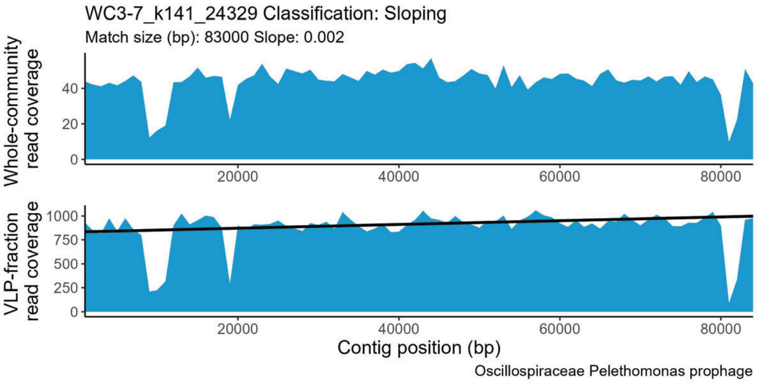
WC3-7_k141_24329 Classification: Sloping
Match size (bp): 83000 Slope: 0.002
Whole-community read coverage
VLP-fraction read coverage
Contig position (bp)
Oscillospiraceae Pelethomonas prophage

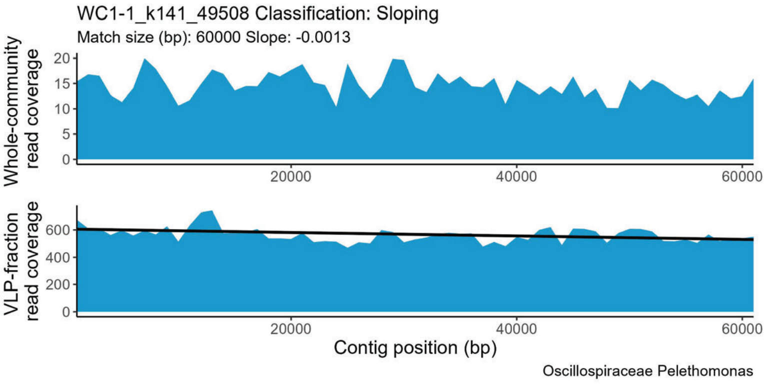
WC1-1_k141_49508 Classification: Sloping
Match size (bp): 60000 Slope: -0.0013
Whole-community read coverage
VLP-fraction read coverage
Contig position (bp)
Oscillospiraceae Pelethomonas

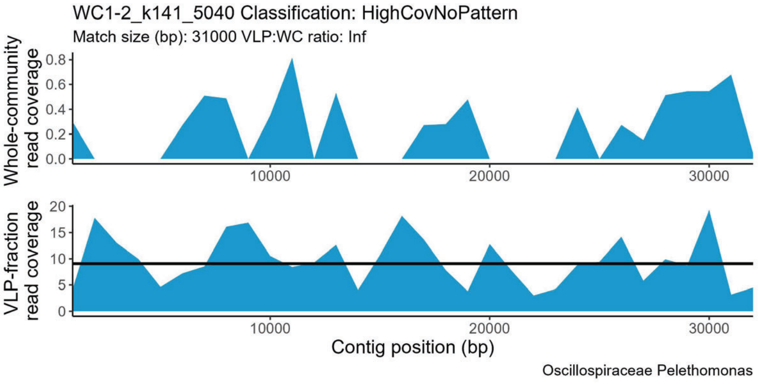
WC1-2_k141_5040 Classification: HighCovNoPattern
Match size (bp): 31000 VLP:WC ratio: Inf
Whole-community read coverage
VLP-fraction read coverage
Contig position (bp)
Oscillospiraceae Pelethomonas

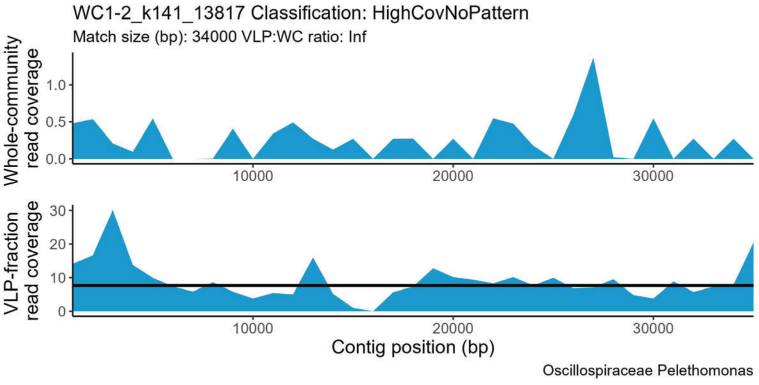
WC1-2_k141_13817 Classification: HighCovNoPattern
Match size (bp): 34000 VLP:WC ratio: Inf
Whole-community read coverage
VLP-fraction read coverage
Contig position (bp)
Oscillospiraceae Pelethomonas

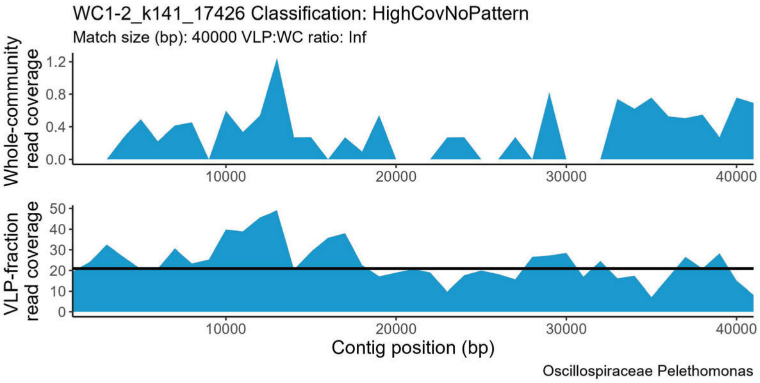
WC1-2_k141_17426 Classification: HighCovNoPattern
Match size (bp): 40000 VLP:WC ratio: Inf
Whole-community read coverage
VLP-fraction read coverage
Contig position (bp)
Oscillospiraceae Pelethomonas

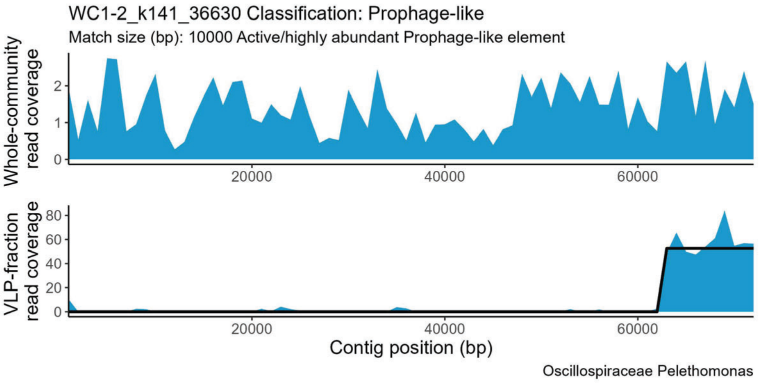
WC1-2_k141_36630 Classification: Prophage-like
Match size (bp): 10000 Active/highly abundant Prophage-like element
Whole-community read coverage
VLP-fraction read coverage
Contig position (bp)
Oscillospiraceae Pelethomonas

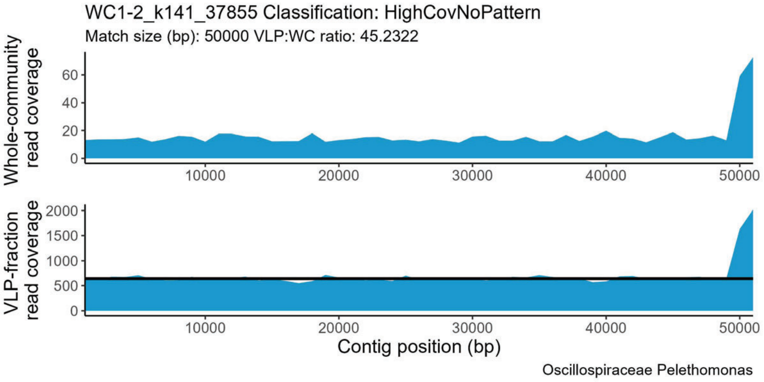
WC1-2_k141_37855 Classification: HighCovNoPattern
Match size (bp): 50000 VLP:WC ratio: 45.2322
Whole-community read coverage
VLP-fraction read coverage
Contig position (bp)
Oscillospiraceae Pelethomonas

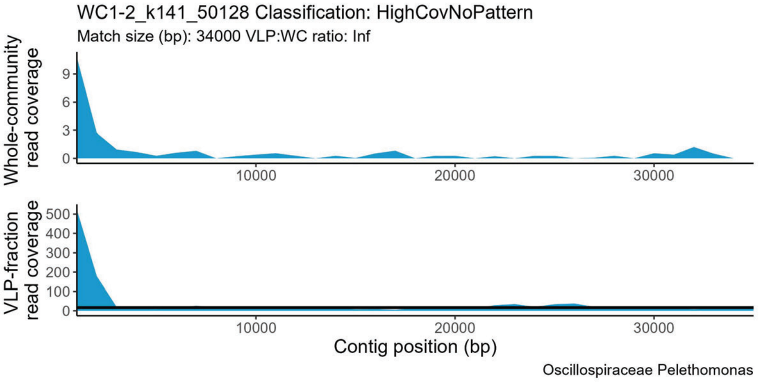
WC1-2_k141_50128 Classification: HighCovNoPattern
Match size (bp): 34000 VLP:WC ratio: Inf
Whole-community read coverage
VLP-fraction read coverage
Contig position (bp)
Oscillospiraceae Pelethomonas

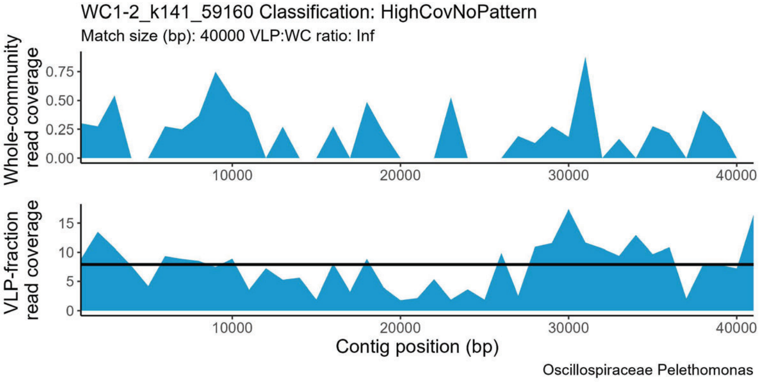
WC1-2_k141_59160 Classification: HighCovNoPattern
Match size (bp): 40000 VLP:WC ratio: Inf
Whole-community read coverage
VLP-fraction read coverage
Contig position (bp)
Oscillospiraceae Pelethomonas

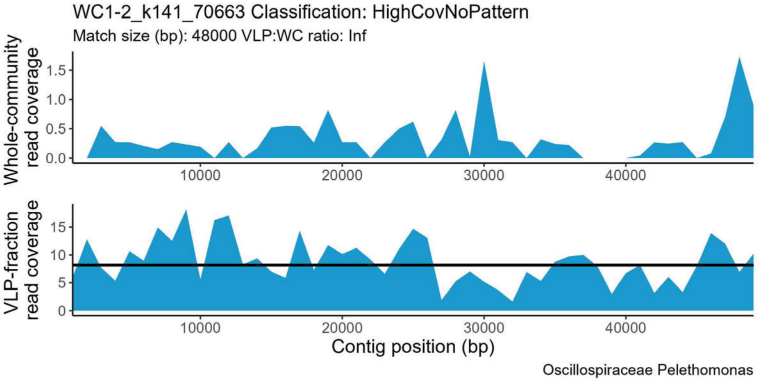
WC1-2_k141_70663 Classification: HighCovNoPattern
Match size (bp): 48000 VLP:WC ratio: Inf
Whole-community read coverage
VLP-fraction read coverage
Contig position (bp)
Oscillospiraceae Pelethomonas

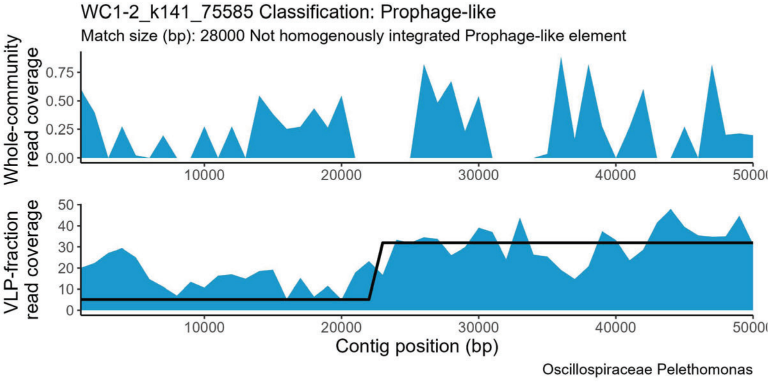
WC1-2_k141_75585 Classification: Prophage-like
Match size (bp): 28000 Not homogenously integrated Prophage-like element
Whole-community read coverage
VLP-fraction read coverage
Contig position (bp)
Oscillospiraceae Pelethomonas

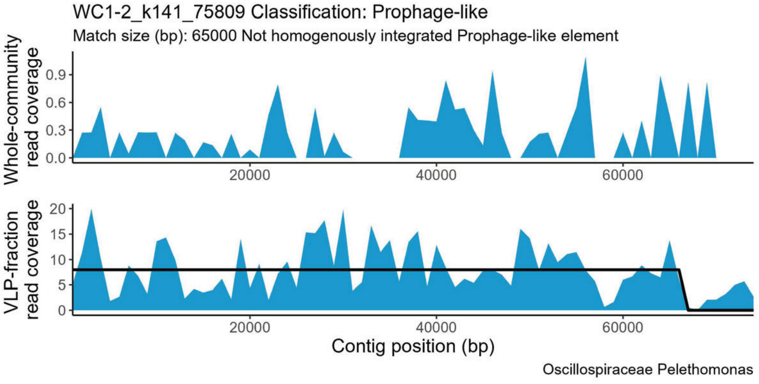
WC1-2_k141_75809 Classification: Prophage-like
Match size (bp): 65000 Not homogenously integrated Prophage-like element
Whole-community read coverage
VLP-fraction read coverage
Contig position (bp)
Oscillospiraceae Pelethomonas

WC1-2_k141_91156 Classification: HighCovNoPattern
Match size (bp): 59000 VLP:WC ratio: Inf

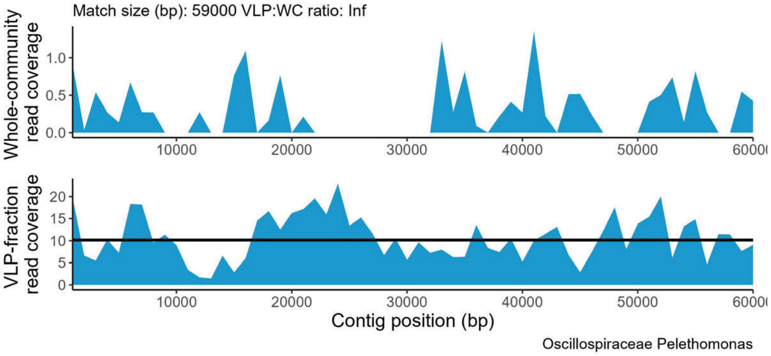


WC1-2_k141_97345 Classification: HighCovNoPattern
Match size (bp): 30000 VLP:WC ratio: Inf

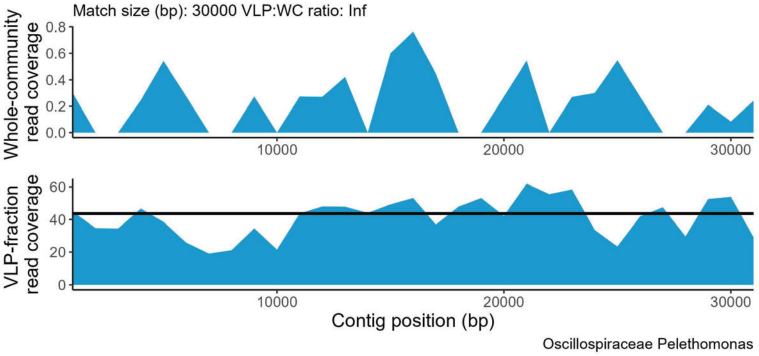


WC1-2_k141_100375 Classification: Sloping
Match size (bp): 24000 Slope: 0.0016

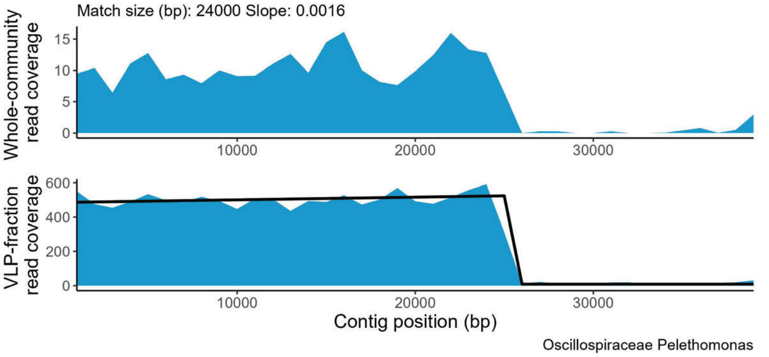


WC1-2_k141_101700 Classification: HighCovNoPattern
Match size (bp): 33000 VLP:WC ratio: Inf

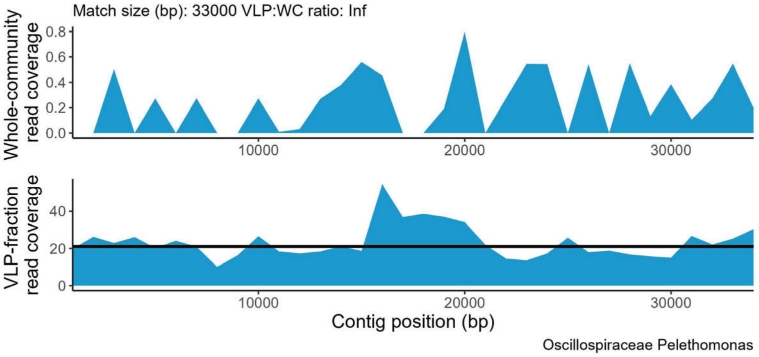


WC1-2_k141_110467 Classification: HighCovNoPattern
Match size (bp): 67000 VLP:WC ratio: Inf

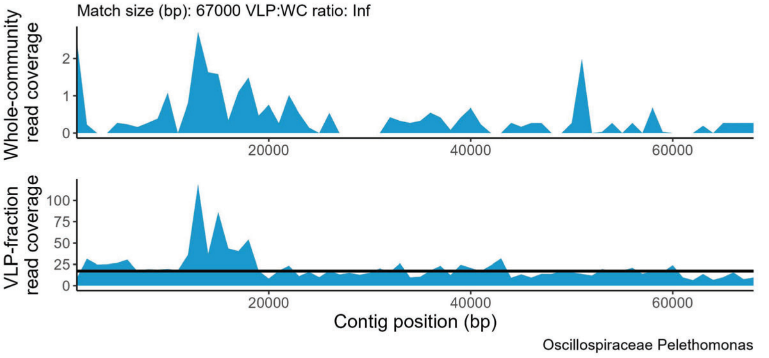


WC1-2_k141_115368 Classification: Prophage-like
Match size (bp): 64000 Not homogenously integrated Prophage-like element

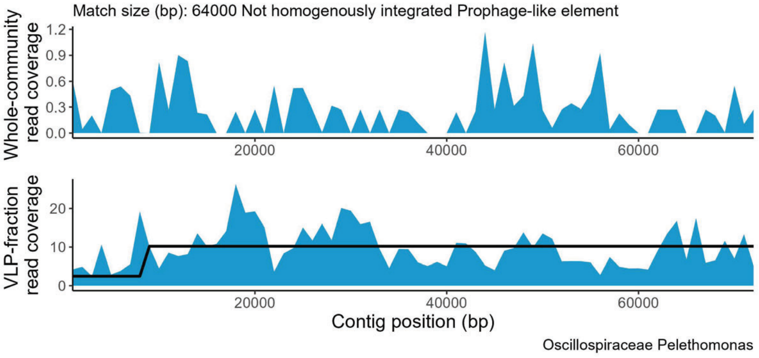


WC1-2_k141_129152 Classification: Prophage-like
Match size (bp): 29000 Not homogenously integrated Prophage-like element

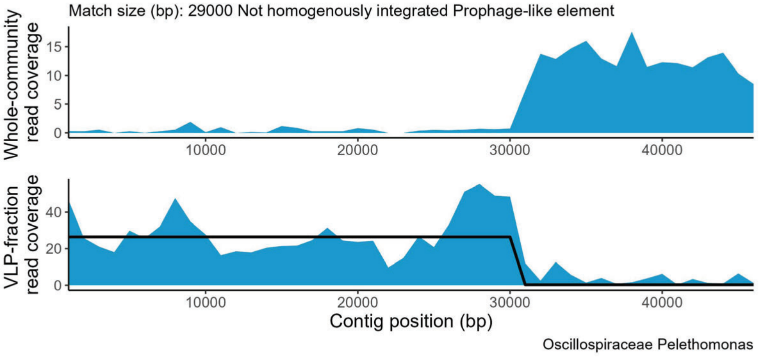


WC1-2_k141_129936 Classification: HighCovNoPattern
Match size (bp): 62000 VLP:WC ratio: Inf

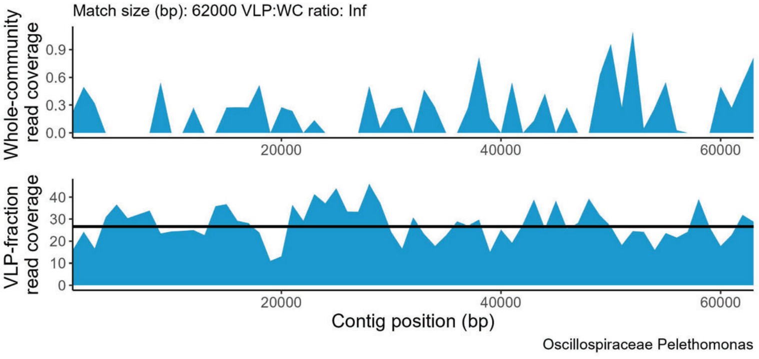

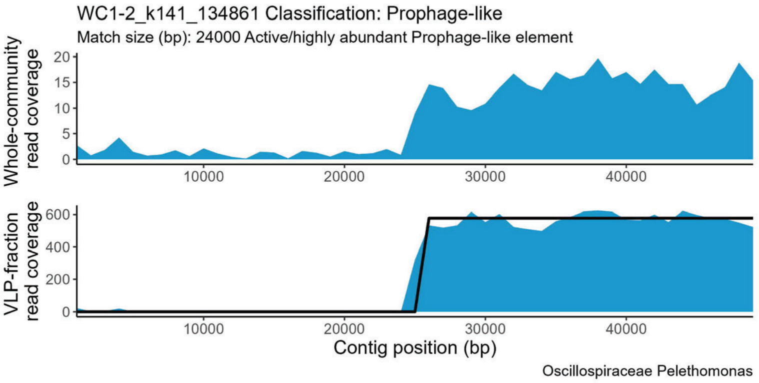
WC1-2_k141_134861 Classification: Prophage-like
Match size (bp): 24000 Active/highly abundant Prophage-like element
Whole-community read coverage
VLP-fraction read coverage
Contig position (bp)
Oscillospiraceae Pelethomonas

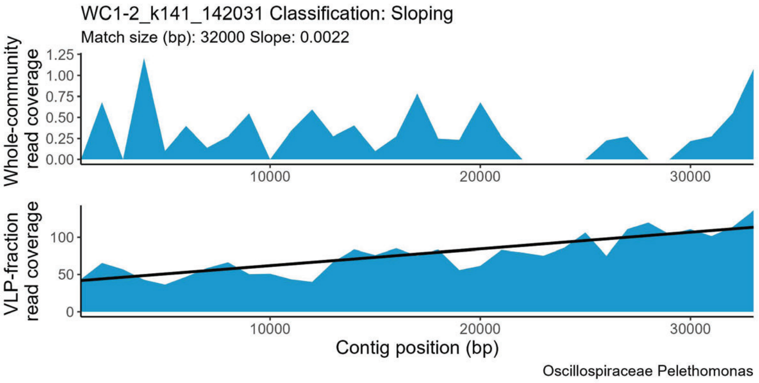
WC1-2_k141_142031 Classification: Sloping
Match size (bp): 32000 Slope: 0.0022
Whole-community read coverage
VLP-fraction read coverage
Contig position (bp)
Oscillospiraceae Pelethomonas

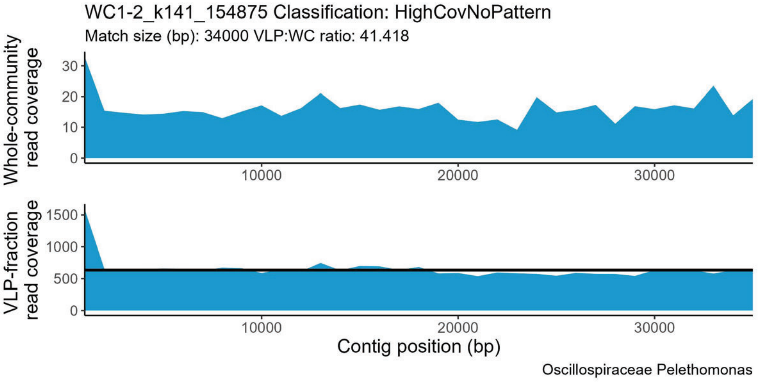
WC1-2_k141_154875 Classification: HighCovNoPattern
Match size (bp): 34000 VLP:WC ratio: 41.418
Whole-community read coverage
VLP-fraction read coverage
Contig position (bp)
Oscillospiraceae Pelethomonas

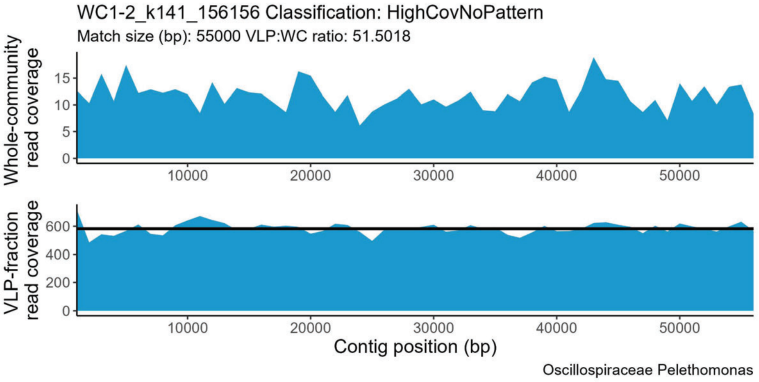
WC1-2_k141_156156 Classification: HighCovNoPattern
Match size (bp): 55000 VLP:WC ratio: 51.5018
Whole-community read coverage
VLP-fraction read coverage
Contig position (bp)
Oscillospiraceae Pelethomonas

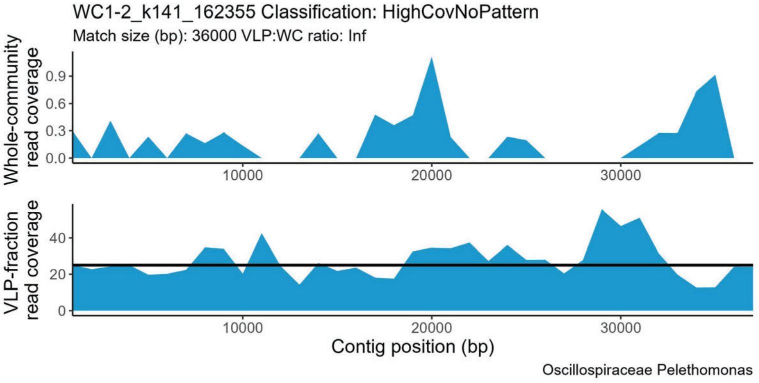
WC1-2_k141_162355 Classification: HighCovNoPattern
Match size (bp): 36000 VLP:WC ratio: Inf
Whole-community read coverage
VLP-fraction read coverage
Contig position (bp)
Oscillospiraceae Pelethomonas

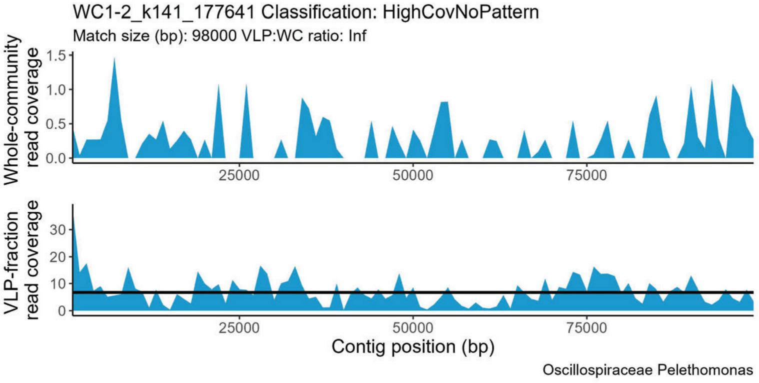
WC1-2_k141_177641 Classification: HighCovNoPattern
Match size (bp): 98000 VLP:WC ratio: Inf
Whole-community read coverage
VLP-fraction read coverage
Contig position (bp)
Oscillospiraceae Pelethomonas

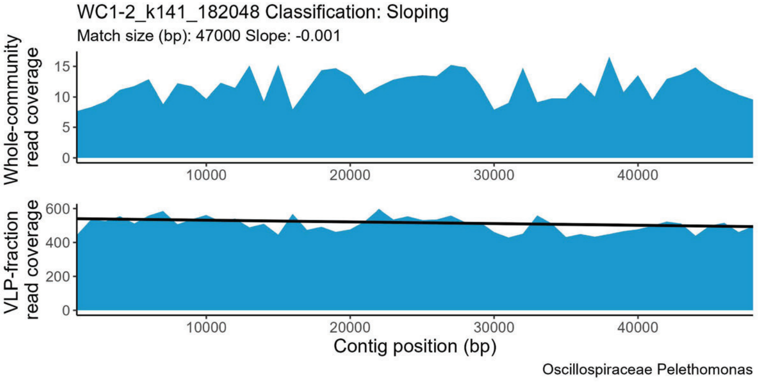
WC1-2_k141_182048 Classification: Sloping
Match size (bp): 47000 Slope: -0.001
Whole-community read coverage
VLP-fraction read coverage
Contig position (bp)
Oscillospiraceae Pelethomonas

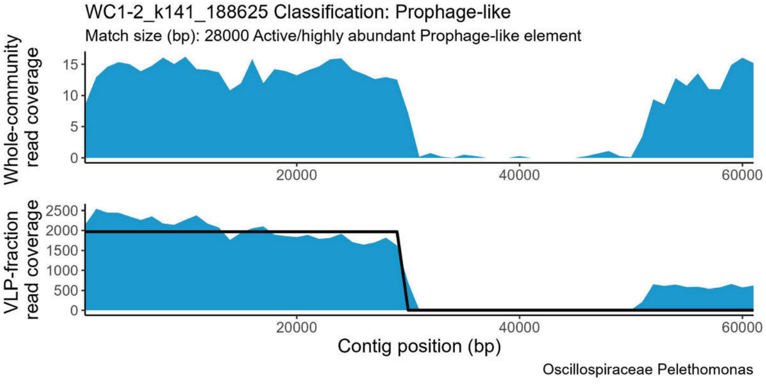
WC1-2_k141_188625 Classification: Prophage-like
Match size (bp): 28000 Active/highly abundant Prophage-like element
Whole-community read coverage
VLP-fraction read coverage
Contig position (bp)
Oscillospiraceae Pelethomonas

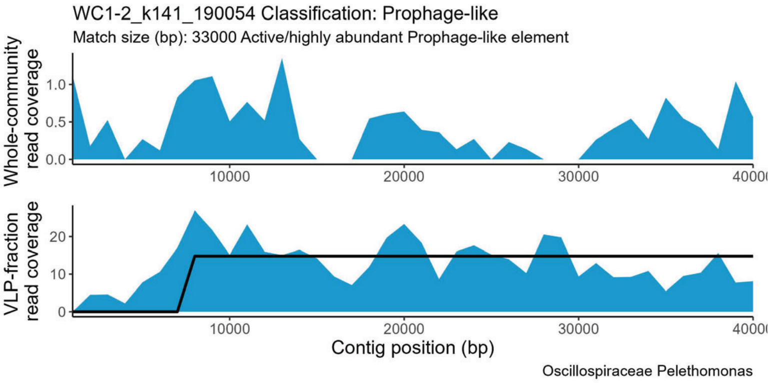
WC1-2_k141_190054 Classification: Prophage-like
Match size (bp): 33000 Active/highly abundant Prophage-like element
Whole-community read coverage
VLP-fraction read coverage
Contig position (bp)
Oscillospiraceae Pelethomonas

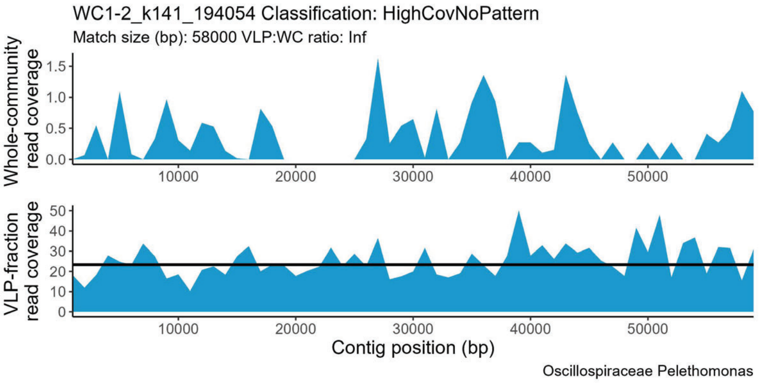
WC1-2_k141_194054 Classification: HighCovNoPattern
Match size (bp): 58000 VLP:WC ratio: Inf
Whole-community read coverage
VLP-fraction read coverage
Contig position (bp)
Oscillospiraceae Pelethomonas

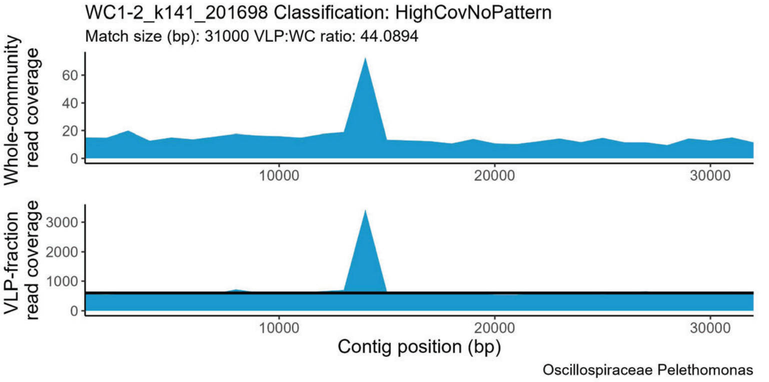
WC1-2_k141_201698 Classification: HighCovNoPattern
Match size (bp): 31000 VLP:WC ratio: 44.0894
Whole-community read coverage
VLP-fraction read coverage
Contig position (bp)
Oscillospiraceae Pelethomonas

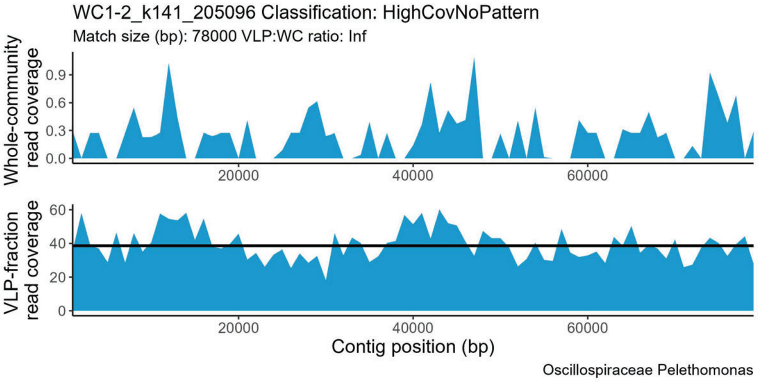
WC1-2_k141_205096 Classification: HighCovNoPattern
Match size (bp): 78000 VLP:WC ratio: Inf
Whole-community read coverage
VLP-fraction read coverage
Contig position (bp)
Oscillospiraceae Pelethomonas

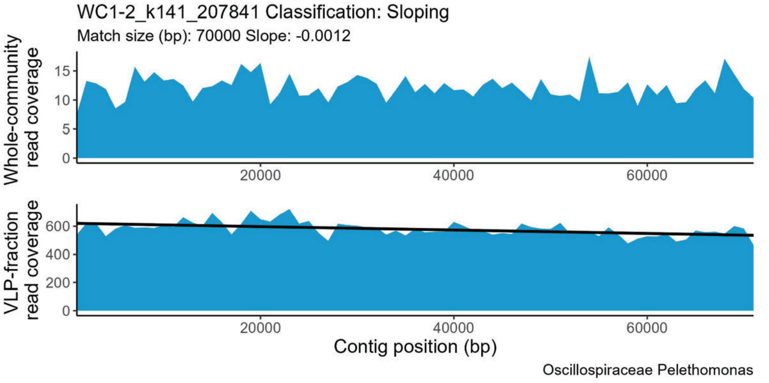
WC1-2_k141_207841 Classification: Sloping
Match size (bp): 70000 Slope: -0.0012
Whole-community read coverage
VLP-fraction read coverage
Contig position (bp)
Oscillospiraceae Pelethomonas

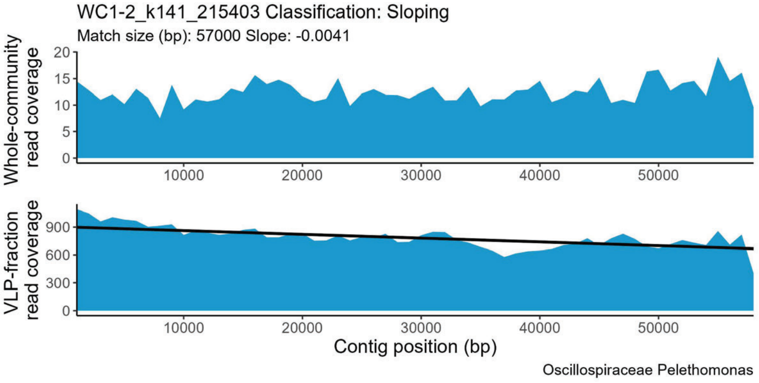
WC1-2_k141_215403 Classification: Sloping
Match size (bp): 57000 Slope: -0.0041
Whole-community read coverage
VLP-fraction read coverage
Contig position (bp)
Oscillospiraceae Pelethomonas

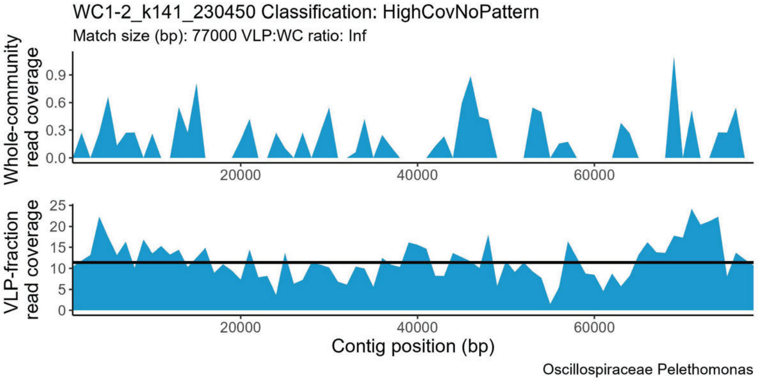
WC1-2_k141_230450 Classification: HighCovNoPattern
Match size (bp): 77000 VLP:WC ratio: Inf
Whole-community read coverage
VLP-fraction read coverage
Contig position (bp)
Oscillospiraceae Pelethomonas

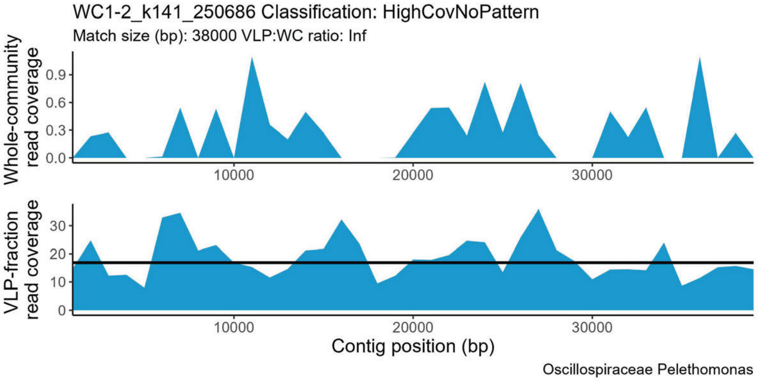
WC1-2_k141_250686 Classification: HighCovNoPattern
Match size (bp): 38000 VLP:WC ratio: Inf
Whole-community read coverage
VLP-fraction read coverage
Contig position (bp)
Oscillospiraceae Pelethomonas

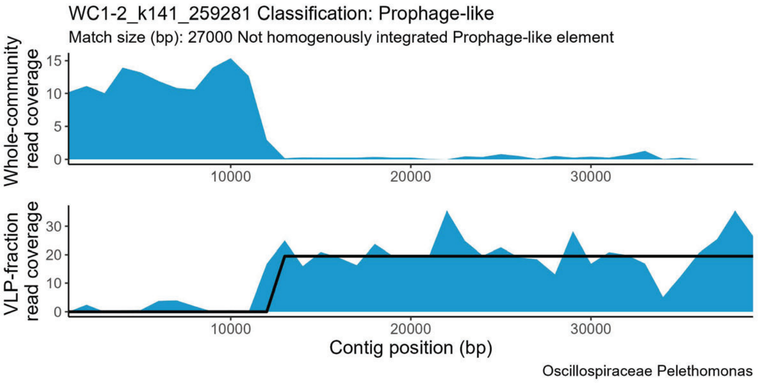
WC1-2_k141_259281 Classification: Prophage-like
Match size (bp): 27000 Not homogenously integrated Prophage-like element
Whole-community read coverage
VLP-fraction read coverage
Contig position (bp)
Oscillospiraceae Pelethomonas

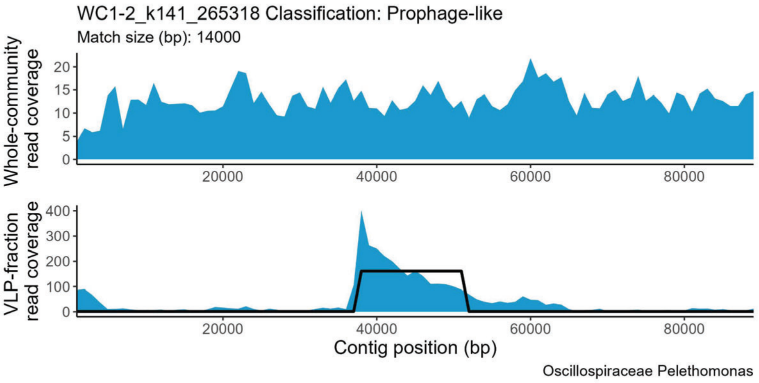
WC1-2_k141_265318 Classification: Prophage-like
Match size (bp): 14000
Whole-community read coverage
VLP-fraction read coverage
Contig position (bp)
Oscillospiraceae Pelethomonas

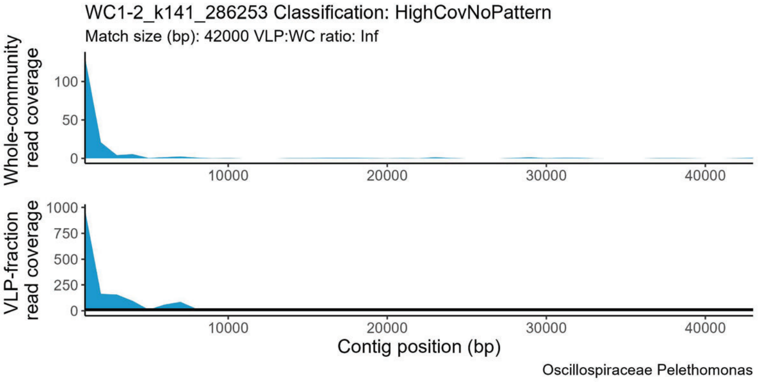
WC1-2_k141_286253 Classification: HighCovNoPattern
Match size (bp): 42000 VLP:WC ratio: Inf
Whole-community read coverage
VLP-fraction read coverage
Contig position (bp)
Oscillospiraceae Pelethomonas

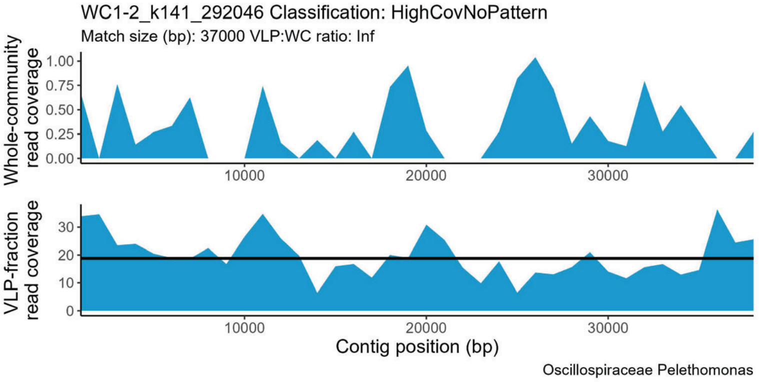
WC1-2_k141_292046 Classification: HighCovNoPattern
Match size (bp): 37000 VLP:WC ratio: Inf
Whole-community read coverage
VLP-fraction read coverage
Contig position (bp)
Oscillospiraceae Pelethomonas

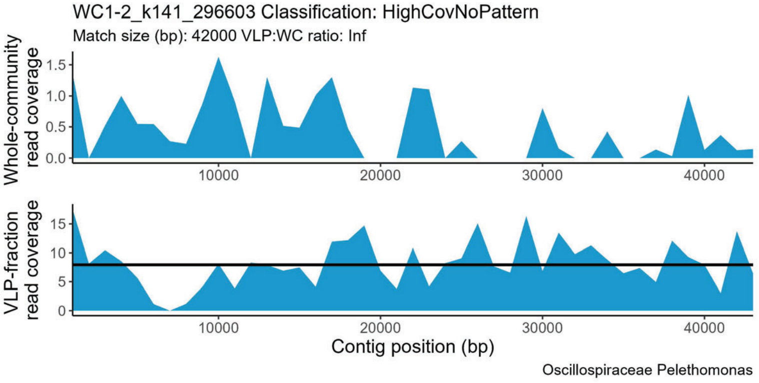
WC1-2_k141_296603 Classification: HighCovNoPattern
Match size (bp): 42000 VLP:WC ratio: Inf
Whole-community read coverage
VLP-fraction read coverage
Contig position (bp)
Oscillospiraceae Pelethomonas

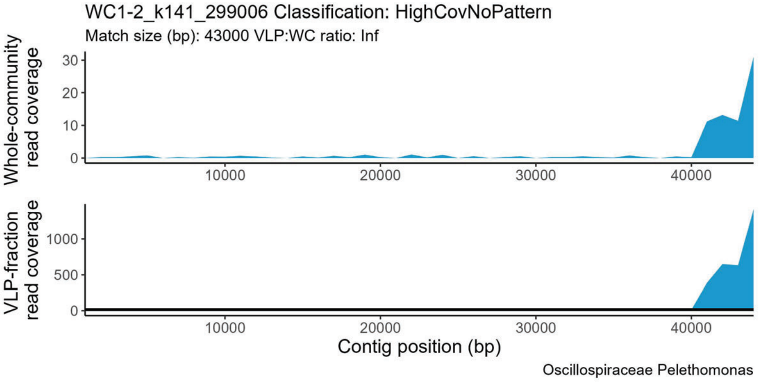
WC1-2_k141_299006 Classification: HighCovNoPattern
Match size (bp): 43000 VLP:WC ratio: Inf
Whole-community read coverage
VLP-fraction read coverage
Contig position (bp)
Oscillospiraceae Pelethomonas

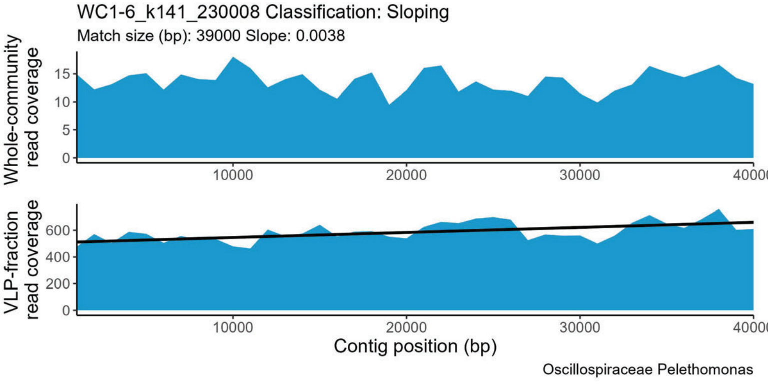
WC1-6_k141_230008 Classification: Sloping
Match size (bp): 39000 Slope: 0.0038
Whole-community read coverage
VLP-fraction read coverage
Contig position (bp)
Oscillospiraceae Pelethomonas

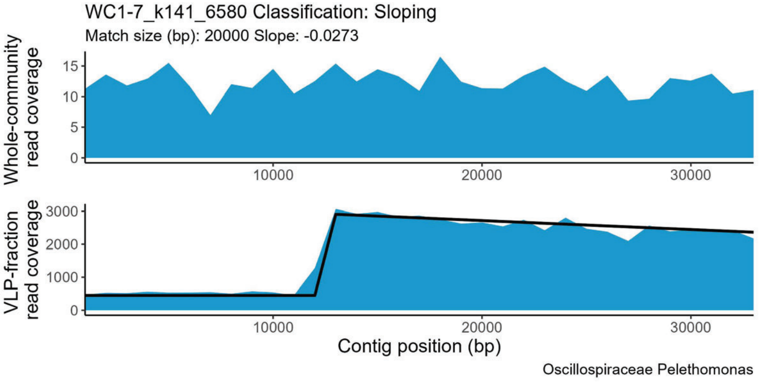
WC1-7_k141_6580 Classification: Sloping
Match size (bp): 20000 Slope: -0.0273
Whole-community read coverage
VLP-fraction read coverage
Contig position (bp)
Oscillospiraceae Pelethomonas

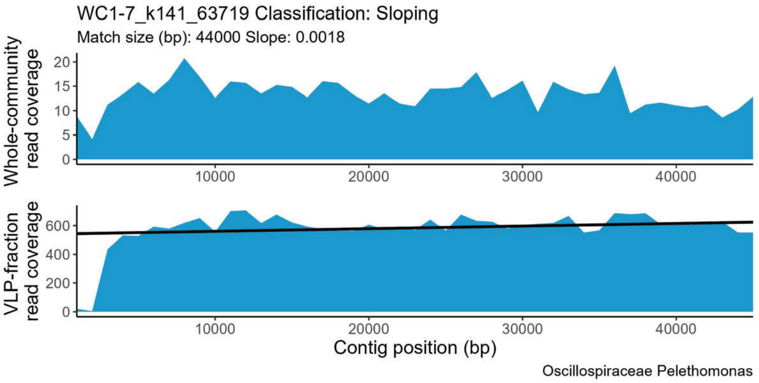
WC1-7_k141_63719 Classification: Sloping
Match size (bp): 44000 Slope: 0.0018
Whole-community read coverage
VLP-fraction read coverage
Contig position (bp)
Oscillospiraceae Pelethomonas

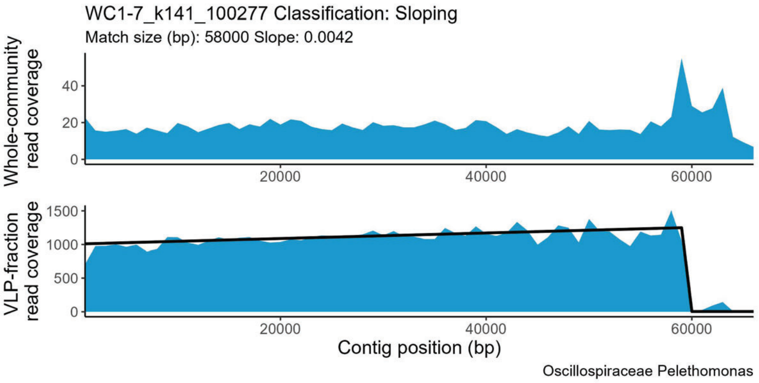
WC1-7_k141_100277 Classification: Sloping
Match size (bp): 58000 Slope: 0.0042
Whole-community read coverage
VLP-fraction read coverage
Contig position (bp)
Oscillospiraceae Pelethomonas

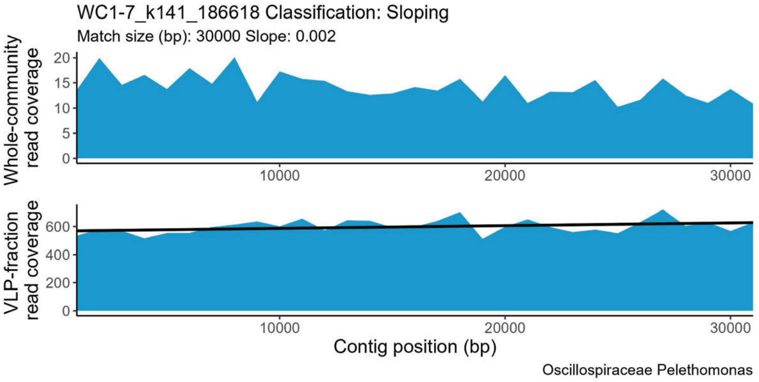
WC1-7_k141_186618 Classification: Sloping
Match size (bp): 30000 Slope: 0.002
Whole-community read coverage
VLP-fraction read coverage
Contig position (bp)
Oscillospiraceae Pelethomonas

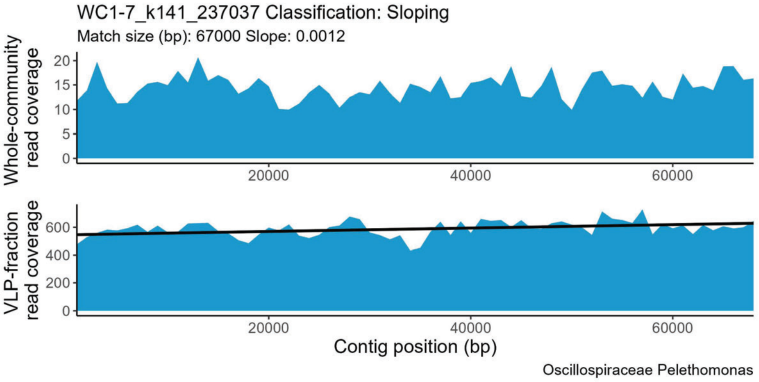
WC1-7_k141_237037 Classification: Sloping
Match size (bp): 67000 Slope: 0.0012
Whole-community read coverage
VLP-fraction read coverage
Contig position (bp)
Oscillospiraceae Pelethomonas

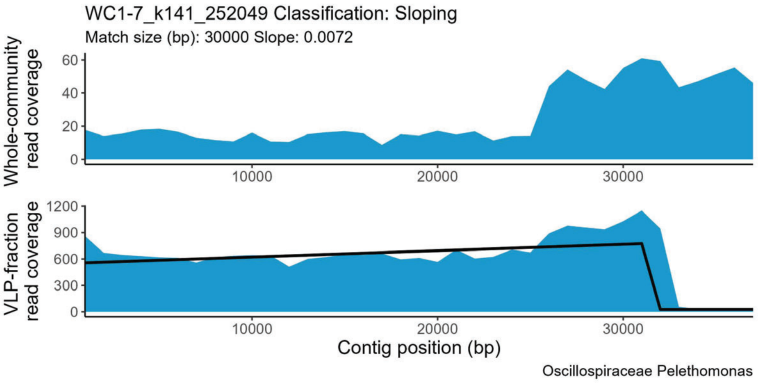
WC1-7_k141_252049 Classification: Sloping
Match size (bp): 30000 Slope: 0.0072
Whole-community read coverage
VLP-fraction read coverage
Contig position (bp)
Oscillospiraceae Pelethomonas

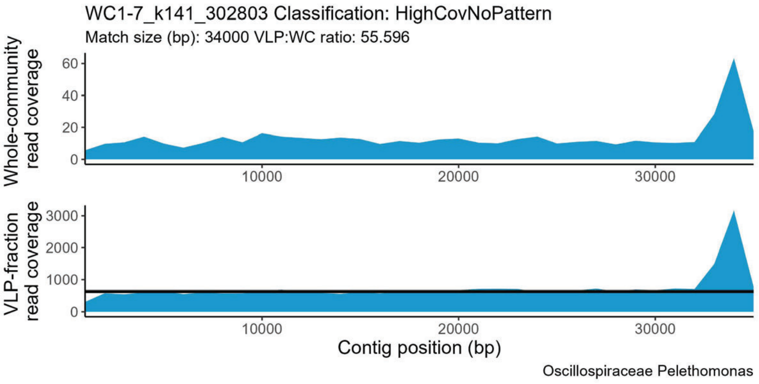
WC1-7_k141_302803 Classification: HighCovNoPattern
Match size (bp): 34000 VLP:WC ratio: 55.596
Whole-community read coverage
VLP-fraction read coverage
Contig position (bp)
Oscillospiraceae Pelethomonas

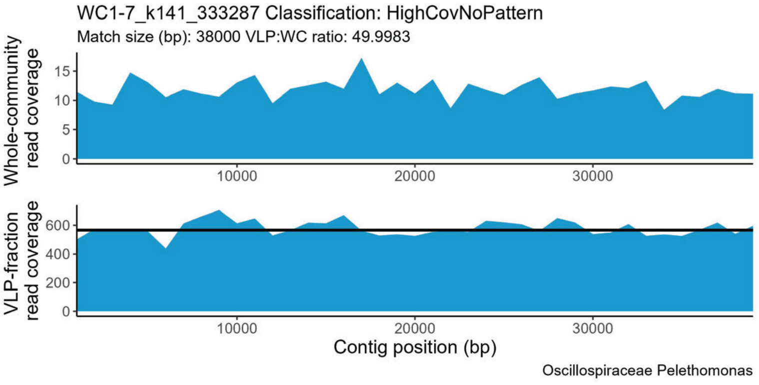
WC1-7_k141_333287 Classification: HighCovNoPattern
Match size (bp): 38000 VLP:WC ratio: 49.9983
Whole-community read coverage
VLP-fraction read coverage
Contig position (bp)
Oscillospiraceae Pelethomonas

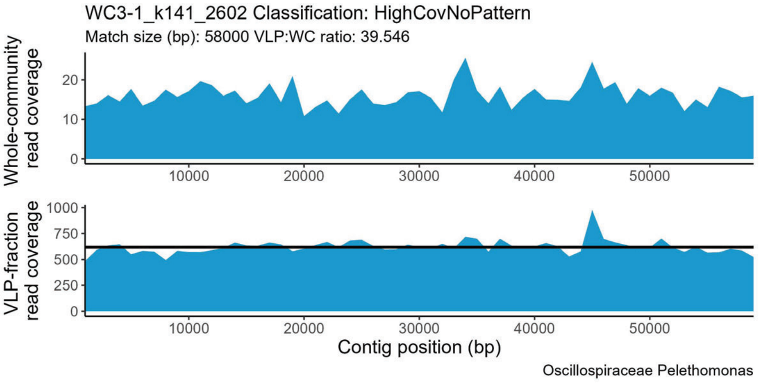
WC3-1_k141_2602 Classification: HighCovNoPattern
Match size (bp): 58000 VLP:WC ratio: 39.546
Whole-community read coverage
VLP-fraction read coverage
Contig position (bp)
Oscillospiraceae Pelethomonas

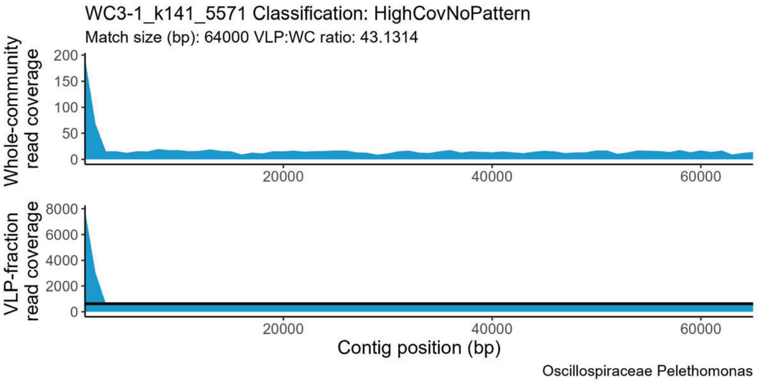
WC3-1_k141_5571 Classification: HighCovNoPattern
Match size (bp): 64000 VLP:WC ratio: 43.1314
Whole-community read coverage
VLP-fraction read coverage
Contig position (bp)
Oscillospiraceae Pelethomonas

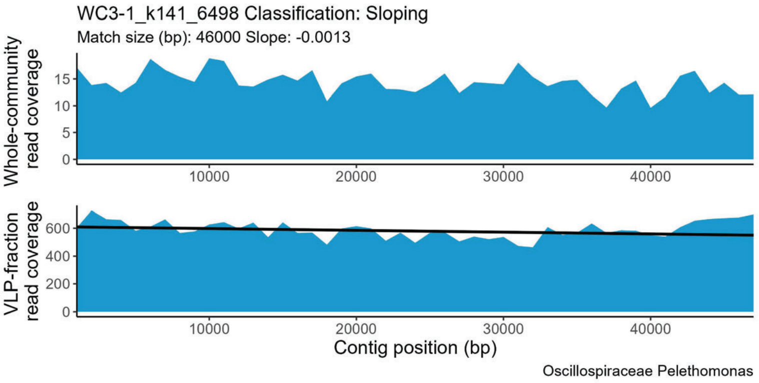
WC3-1_k141_6498 Classification: Sloping
Match size (bp): 46000 Slope: -0.0013
Whole-community read coverage
VLP-fraction read coverage
Contig position (bp)
Oscillospiraceae Pelethomonas

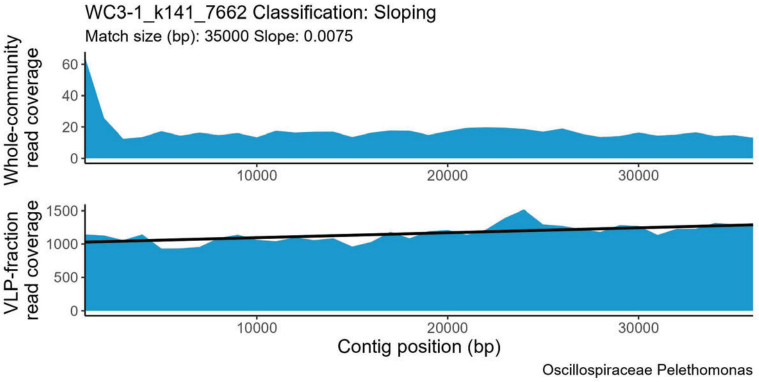
WC3-1_k141_7662 Classification: Sloping
Match size (bp): 35000 Slope: 0.0075
Whole-community read coverage
VLP-fraction read coverage
Contig position (bp)
Oscillospiraceae Pelethomonas

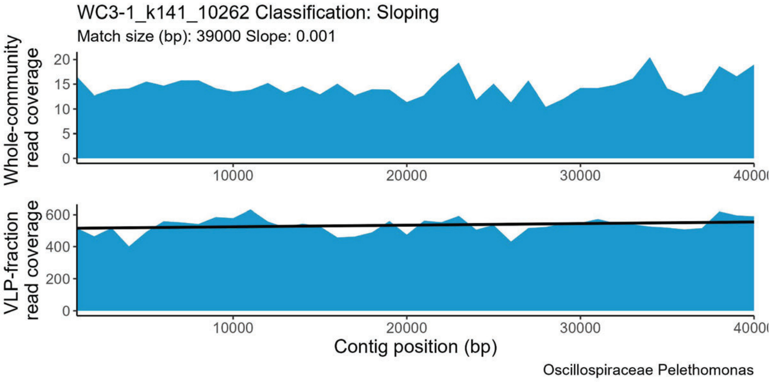
WC3-1_k141_10262 Classification: Sloping
Match size (bp): 39000 Slope: 0.001
Whole-community read coverage
VLP-fraction read coverage
Contig position (bp)
Oscillospiraceae Pelethomonas

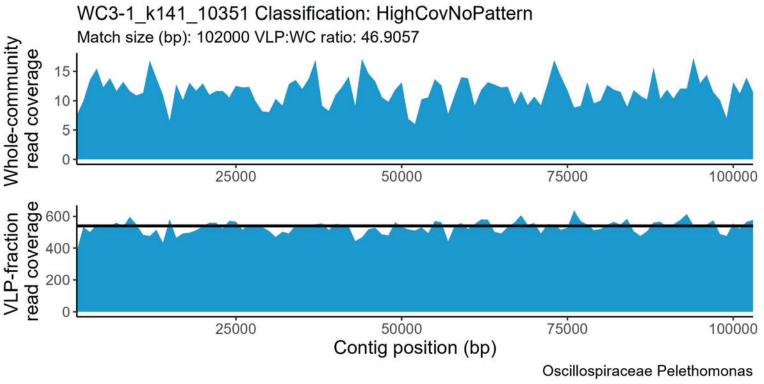
WC3-1_k141_10351 Classification: HighCovNoPattern
Match size (bp): 102000 VLP:WC ratio: 46.9057
Whole-community read coverage
VLP-fraction read coverage
Contig position (bp)
Oscillospiraceae Pelethomonas

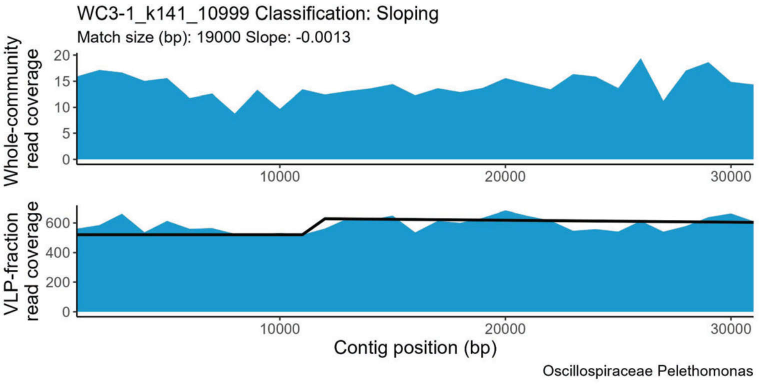
WC3-1_k141_10999 Classification: Sloping
Match size (bp): 19000 Slope: -0.0013
Whole-community read coverage
VLP-fraction read coverage
Contig position (bp)
Oscillospiraceae Pelethomonas

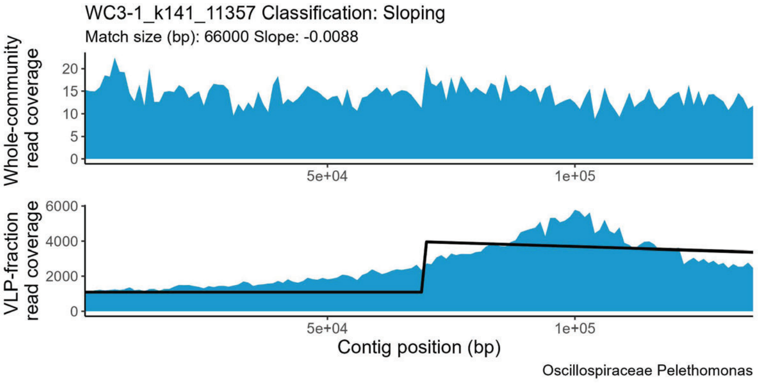
WC3-1_k141_11357 Classification: Sloping
Match size (bp): 66000 Slope: -0.0088
Whole-community read coverage
VLP-fraction read coverage
Contig position (bp)
Oscillospiraceae Pelethomonas

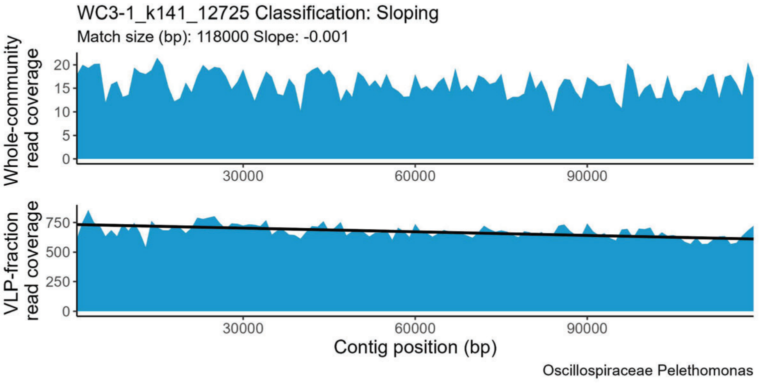
WC3-1_k141_12725 Classification: Sloping
Match size (bp): 118000 Slope: -0.001
Whole-community read coverage
VLP-fraction read coverage
Contig position (bp)
Oscillospiraceae Pelethomonas

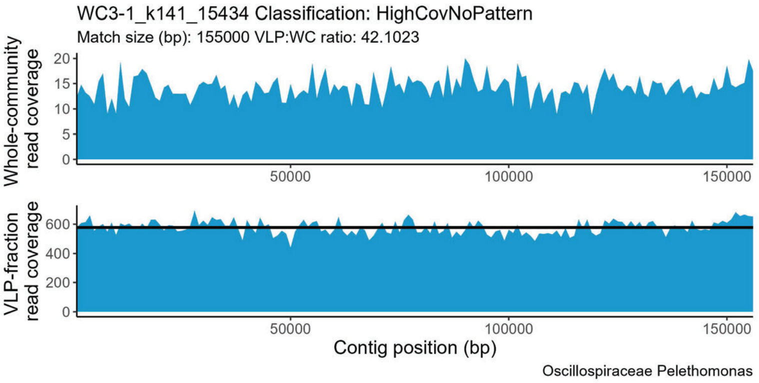
WC3-1_k141_15434 Classification: HighCovNoPattern
Match size (bp): 155000 VLP:WC ratio: 42.1023
Whole-community read coverage
0
5
10
15
20
50000
100000
150000
VLP-fraction read coverage
0
200
400
600
50000
100000
150000
Contig position (bp)
Oscillospiraceae Pelethomonas

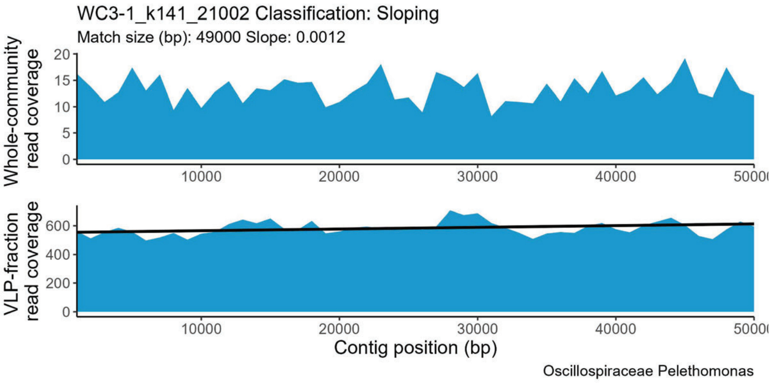
WC3-1_k141_21002 Classification: Sloping
Match size (bp): 49000 Slope: 0.0012
Whole-community read coverage
0
5
10
15
20
10000
20000
30000
40000
VLP-fraction read coverage
0
200
400
600
10000
20000
30000
40000
Contig position (bp)
Oscillospiraceae Pelethomonas

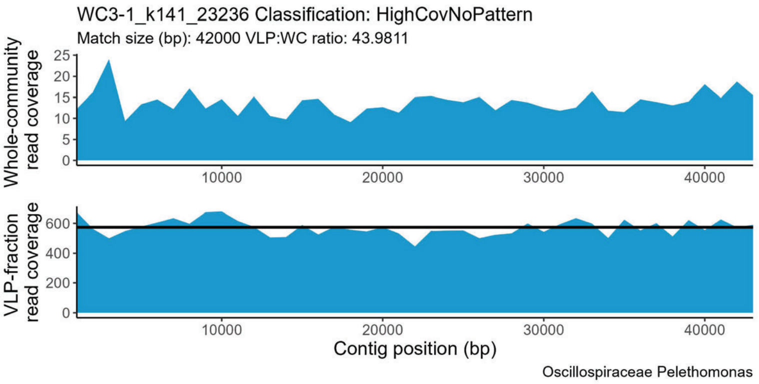
WC3-1_k141_23236 Classification: HighCovNoPattern
Match size (bp): 42000 VLP:WC ratio: 43.9811
Whole-community read coverage
0
5
10
15
20
25
10000
20000
30000
40000
VLP-fraction read coverage
0
200
400
600
10000
20000
30000
40000
Contig position (bp)
Oscillospiraceae Pelethomonas

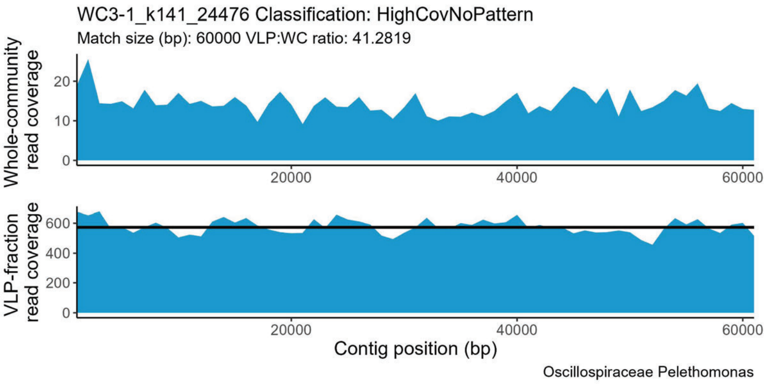
WC3-1_k141_24476 Classification: HighCovNoPattern
Match size (bp): 60000 VLP:WC ratio: 41.2819
Whole-community read coverage
0
10
20
20000
40000
60000
VLP-fraction read coverage
0
200
400
600
20000
40000
60000
Contig position (bp)
Oscillospiraceae Pelethomonas
