## Supplemental Material for "Perturbations shift the composition of bacterial DNA carried by virus-like particles in the murine gut microbiome"

**Affiliations:**
[1]Department of Plant and Microbial Biology, North Carolina State University, Raleigh, NC
[2]Population Health and Pathobiology Department, North Carolina State University, Raleigh, NC
[3]Department of Immunology and Microbiology, University of Colorado - Anschutz Medical Campus, School of Medicine, Aurora, CO

### Supplementary text:

#### Supplementary Results

Total transduction increase/decrease between conditions:

While the question as to whether or not total transduction rates increase or decrease after perturbation was of interest, issues caused by assembly fragmentation (discussed in the conclusions and Suppl. Text limitations) prevented us from drawing any conclusions. While there appears to be a large decrease in transduction events during CDI, when TrIdent is run on the individual assemblies from each replicate, there is an average of 13.7% and 13.1% positively classified contigs in Pre-ABX and CDI, respectively (Fig. S37).

NoPattern classifications:

While we removed NoPattern classifications to streamline the analysis, contigs with NoPattern classifications are not necessarily devoid of VLP-fraction coverage, they just don't have a defined pattern and the coverage is lower than the default HighCovNoPattern ratio cutoff of >2. Therefore, contigs with NoPattern classifications may still be associated with known or unknown DNA carriage mechanisms, such as being the 'tails' of generalized or lateral or GTA-mediated transduction events that have lower coverage (see Fig. S2 Kleiner et al. (2020)). These additional potential events could be detected by lowering the default minimum HighCovNoPattern ratio (`minHCNPRatio` parameter) when running TrIdentClassifier(), which however would likely also increase detection of false positives.

#### Supplementary Discussion

##### Limitations

The biggest limitation to not only the transductomics method but short-read sequencing in general is that it's impossible to know the true origin of reads that map ambiguously to two or more locations. This problem is especially pronounced in our non-redundant contig set in which, despite the efforts to reduce redundancy, there are still many contigs with shared portions. To make matters complicated, these regions are often associated with MGEs which are highly diverse and can be integrated into multiple different hosts[82]. When mapping reads for transductomics, using ambiguous=random is typically recommended to avoid artificial increases in coverage at ambiguous regions, however, we used ambiguous=all in our study. Since our non-redundant contig set originated from different assemblies, there was so much redundancy that using ambiguous=random caused dips in coverage at redundant sites. Using ambiguous=all smoothed the coverage at ambiguous sites leading to better pattern-matches. Since ambiguous=all can inflate the coverage of redundant regions, we confirmed that the coverage patterns analyzed (Figs. 3-5) persisted when reads from individual samples were mapped back to their respective assemblies with ambiguous=random (Fig. S36). Additionally, using BLASTn to search for detected MGEs in the non-filtered (redundant) contig set helps determine how much read coverage is associated with ambiguous read mapping in each sample.

Redundant regions can create read coverage artifacts regardless of the mapping method used and therefore, it may be tempting to implement strict redundancy removal methods however there are important considerations to make when removing redundancy. The goal with redundancy removal is to remove contigs that, due to sequence similarity, produce near identical read coverage patterns to other contigs in the dataset. While choosing a high minimum sequence identity is straightforward, choosing the minimum contig overlap is more difficult. The higher the minimum overlap, the more likely there are to be contigs that differ by only very small regions and produce indistinguishable coverage patterns, however the lower the minimum overlap, the more data is 'lost' or removed, some of which may represent biologically relevant SVs. We found a balance between unique coverage patterns and overall dataset size with a minoverlap=70, however we find that contigs associated with certain bacterial families, like the Butyricicoccaceae, still have a large amount of redundancy. We suspect that bacterial genera and/or species heavily involved in HGT are more likely to have redundancy due to the abundance of SVs in the population[83].

Finally, TrIdent results are affected by the number, length and quality of contigs in an assembly. A higher alpha diversity may lead to a more fragmented assembly based on the increased complexity of the community. Since transduction events that span multiple contigs will be counted as individual events by TrIdent, a more fragmented assembly will lead to inflated transduction event counts. The N50s of the assemblies used to generate the non-redundant contig set (Table S3) are lower in the Pre-ABX samples compared to CDI and therefore it's difficult to draw conclusions about the count of transduction events in each condition. In fact, when TrIdent is run on the individual WC assemblies, the percentage of positively classified contigs is similar between Pre-ABX and CDI (Fig. S37). A potential solution for the issues caused by both ambiguously mapping reads and assembly quality is long-read sequencing.

### Supplementary Tables:

| **Condition** | **Replicate** | **# of trimmed and decontaminated WC reads** |
|---|---|---|
| pre-ABX | 1-1 | 48,302,807 |
| | 1-2 | 70,752,496 |
| | 1-3 | 46,594,438 |
| | 1-4 | 76,560,783 |
| post-ABX | 2-1 | 37,477,257 |
| | 2-2 | 90,360,268 |
| | 2-3 | 128,514,825 |
| | 2-4 | 48,889,705 |
| CDI | 3-1 | 92,274,340 |
| | 3-2 | 37,117,994 |
| | 3-3 | 61,330,785 |
| | 3-4 | 45,493,445 |

**Supplementary table 1 - Number of trimmed and decontaminated reads obtained for each whole community.** ABX= antibiotics, CDI=*Clostridioides difficile* infection, WC= whole-community

| **Condition** | **Replicate** | **# of trimmed and decontaminated VLP-fraction reads** |
|---|---|---|
| Pre-ABX | 1-1 | 47,432,387 |
| Pre-ABX | 1-2 | 36,100,429 |
| Pre-ABX | 1-3 | 115,326,785 |
| Pre-ABX | 1-4 | 5,703,612 |
| Post-ABX | 2-1 | 38,189,939 |
| Post-ABX | 2-2 | 61,948,489 |
| Post-ABX | 2-3 | 50,831,719 |
| Post-ABX | 2-4 | 524,118 |
| CDI | 3-1 | 110,364,922 |
| CDI | 3-2 | 172,259,597 |
| CDI | 3-3 | 4,114,031 |
| CDI | 3-4 | 2,305,702 |

**Supplementary table 2 - Number of trimmed and decontaminated reads obtained for each VLP-fraction sample.** ABX= antibiotics, CDI=*Clostridioides difficile* infection, VLP= virus-like particle

**Table S2) MetaQuast results for whole-community assemblies.**

Assemblies were performed with MEGAHIT. The N50 values are highlighted in yellow to emphasize the difference in values between sample conditions. ABX= antibiotics, CDI= Clostridioides difficile

| Sample name in manuscript | Pre-ABX rep 1 | Pre-ABX rep 2 | Pre-ABX rep 3 | Pre-ABX rep 4 | Post-ABX rep 1 | Post-ABX rep 2 | Post-ABX rep 3 | Post-ABX rep 4 | CDI rep 1 | CDI rep 2 | CDI rep 3 | CDI rep 4 |
|---|---|---|---|---|---|---|---|---|---|---|---|---|
| Assembly | WC1-1_CP05702_L007_contigs | WC1-2_CP05702_L007_contigs | WC1-6_CP05702_L007_contigs | WC1-7_CP05570_S7_L001_contig | WC2-1_CP05702_L007_contigs | WC2-2_CP05702_L007_contigs | WC2-6_CP05702_L008_contigs | WC2-7_CP05570_S8_L001_contigs | WC3-1_CP05702_L007_contigs | WC3-2_CP05702_L007_contigs | WC3-6_CP05702_L007_contigs | WC3-7_CP05570_contigs |
| # contigs (>= 0 bp) | 285165 | 301023 | 320832 | 329946 | 9036 | 10145 | 8885 | 6656 | 33695 | 40654 | 36370 | 36116 |
| # contigs (>= 1000 bp) | 58292 | 59984 | 62900 | 67987 | 691 | 816 | 444 | 353 | 8839 | 10843 | 11926 | 11341 |
| # contigs (>= 5000 bp) | 7852 | 8932 | 8073 | 7828 | 52 | 64 | 55 | 77 | 3523 | 2500 | 4065 | 2599 |
| # contigs (>= 10000 bp) | 3666 | 4140 | 3858 | 3806 | 44 | 48 | 50 | 69 | 2486 | 1573 | 2531 | 1547 |
| # contigs (>= 25000 bp) | 1344 | 1646 | 1687 | 1637 | 31 | 31 | 32 | 46 | 1343 | 794 | 1209 | 743 |
| # contigs (>= 50000 bp) | 521 | 708 | 793 | 729 | 17 | 16 | 17 | 31 | 660 | 425 | 588 | 371 |
| Total length (>= 0 bp) | 3.54E+08 | 3.95E+08 | 4.05E+08 | 4.10E+08 | 7722316 | 8390066 | 7187710 | 8370184 | 1.43E+08 | 1.08E+08 | 1.49E+08 | 1.04E+08 |
| Total length (>= 1000 bp) | 2.40E+08 | 2.75E+08 | 2.76E+08 | 2.78E+08 | 3825239 | 4174270 | 3376520 | 5480735 | 1.31E+08 | 92621581 | 1.37E+08 | 90917420 |
| Total length (>= 5000 bp) | 1.44E+08 | 1.77E+08 | 1.72E+08 | 1.67E+08 | 2848461 | 2862893 | 2789571 | 5067977 | 1.21E+08 | 76972555 | 1.20E+08 | 74011027 |
| Total length (>= 10000 bp) | 1.15E+08 | 1.44E+08 | 1.44E+08 | 1.39E+08 | 2799201 | 2759653 | 2755720 | 5014455 | 1.13E+08 | 70452818 | 1.09E+08 | 66623693 |
| Total length (>= 25000 bp) | 80019840 | 1.07E+08 | 1.10E+08 | 1.06E+08 | 2590072 | 2450680 | 2435042 | 4631868 | 94986012 | 58092965 | 88438563 | 53953244 |
| Total length (>= 50000 bp) | 51704061 | 73573144 | 78805116 | 74071370 | 2126459 | 1947423 | 1946542 | 4119135 | 71112532 | 45014464 | 66770360 | 40744270 |
| # contigs | 149471 | 155178 | 164063 | 174837 | 3306 | 3324 | 2754 | 2186 | 18068 | 24683 | 21774 | 22637 |
| Largest contig | 561305 | 763659 | 556368 | 728877 | 353181 | 353181 | 353181 | 353181 | 714945 | 576711 | 628498 | 483925 |
| Total length | 3.02E+08 | 3.39E+08 | 3.44E+08 | 3.51E+08 | 5544372 | 5821693 | 4873103 | 6677345 | 1.38E+08 | 1.02E+08 | 1.43E+08 | 98769013 |
| GC (%) | 47.3 | 48.28 | 46.41 | 45.61 | 37.77 | 37.67 | 38.78 | 38.87 | 46.51 | 45.63 | 45.91 | 45.42 |
| N50 | 4343 | 5828 | 4999 | 4186 | 11063 | 4203 | 23820 | 79344 | 52709 | 36993 | 43149 | 32793 |
| N90 | 680 | 696 | 683 | 681 | 588 | 594 | 582 | 675 | 3148 | 1053 | 2294 | 1190 |
| auN | 31045.7 | 40759.5 | 36219.5 | 36527.9 | 66969.7 | 59854.2 | 69210.4 | 117966.9 | 89280.5 | 74221.3 | 89323.6 | 69687.8 |
| L50 | 9283 | 7497 | 8080 | 9712 | 42 | 75 | 33 | 20 | 616 | 565 | 696 | 584 |
| L90 | 97258 | 97114 | 104530 | 114217 | 2279 | 2250 | 1848 | 1020 | 4339 | 10192 | 6761 | 9484 |
| # N's per 100 kbp | 0 | 0 | 0 | 0 | 0 | 0 | 0 | 0 | 0 | 0 | 0 | 0 |

#### Supplementary figures:

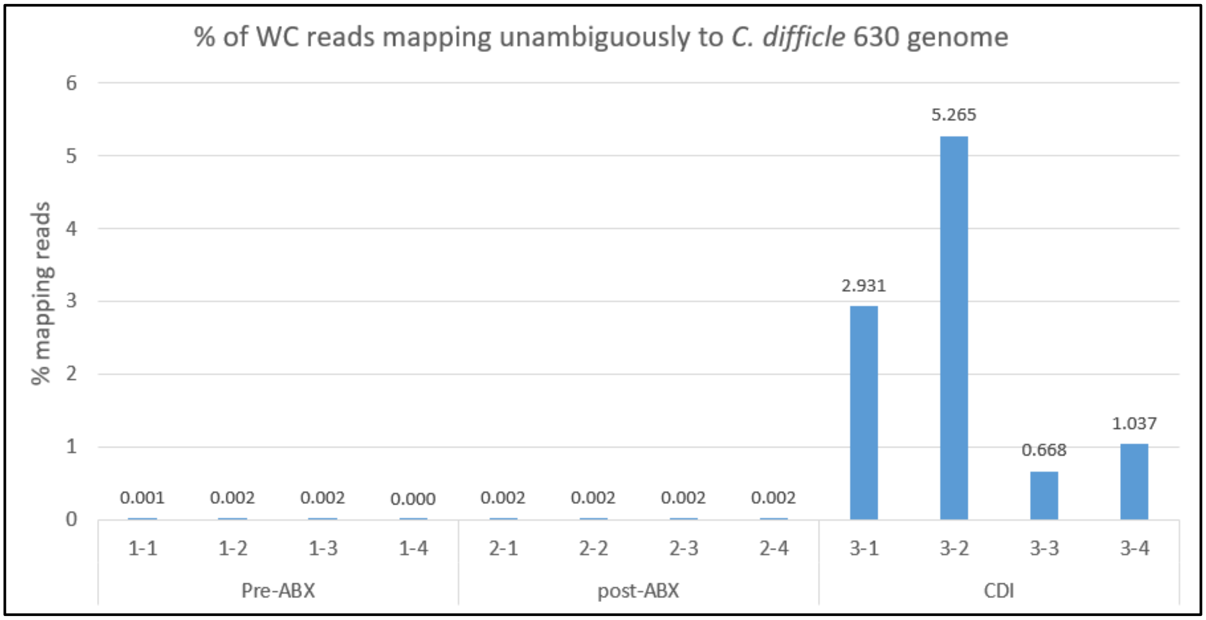


**Supplementary figure 1 - Percent of trimmed and decontaminated reads whole-community reads that mapped unambiguously to the *Clostridioides difficile* 630 reference genome in each sample condition (Pre-ABX, Post-ABX and CDI).** ABX= antibiotics, CDI=*Clostridioides difficile* infection, WC=whole-community

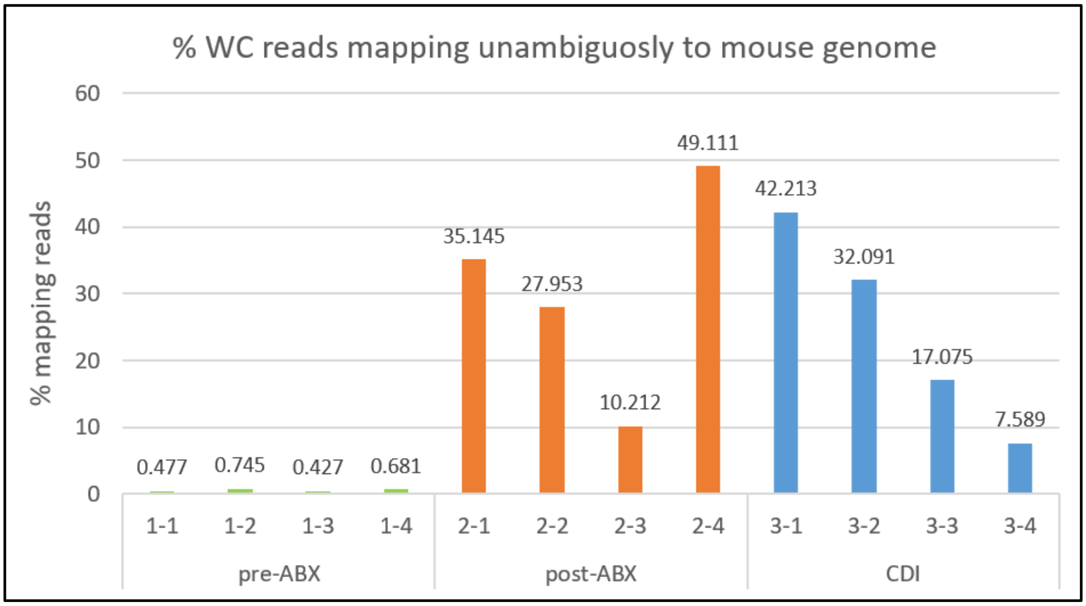


**Supplementary figure 2 - Percent of trimmed whole-community reads that map unambiguously to the mouse reference genome in each sample condition (Pre-ABX, Post-ABX and CDI).** ABX= antibiotics, CDI=*Clostridioides difficile* infection, WC=whole-community

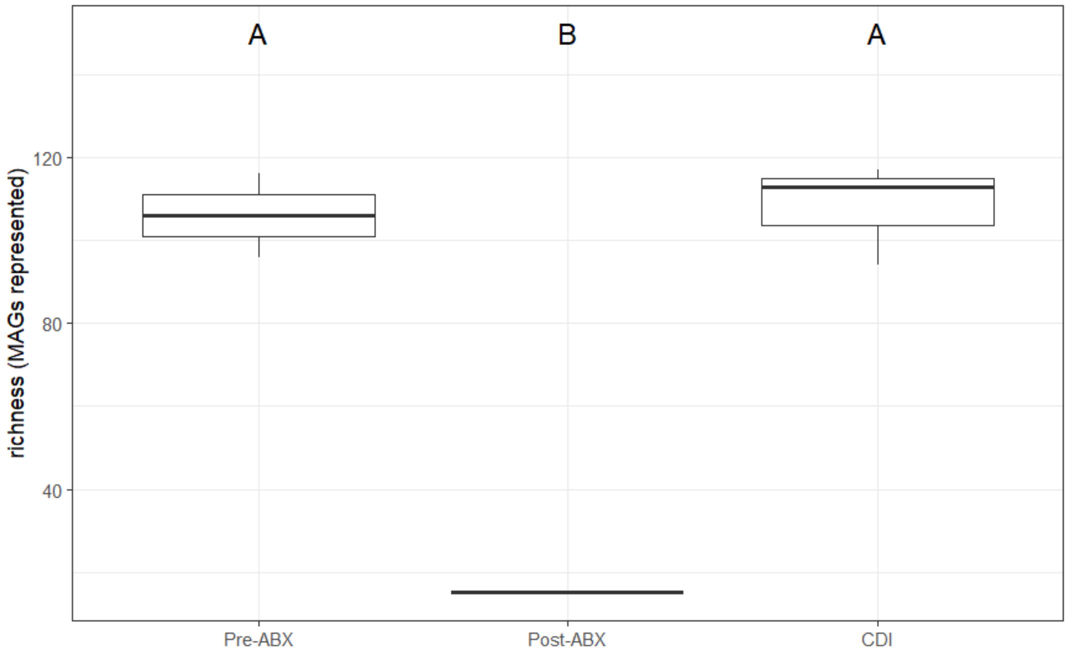


**Supplementary figure 3- Richness for whole-communities.** Determined via the number of metagenome assembled genomes (MAGs) with relative abundance >0.05% represented in the non-redundant contig set in each whole-community sample. Kruskal-Wallis test performed between samples in each condition and Dunn post-hoc test to determine significance ($p<0.05$). ABX= antibiotics, CDI=*Clostridioides difficile* infection.

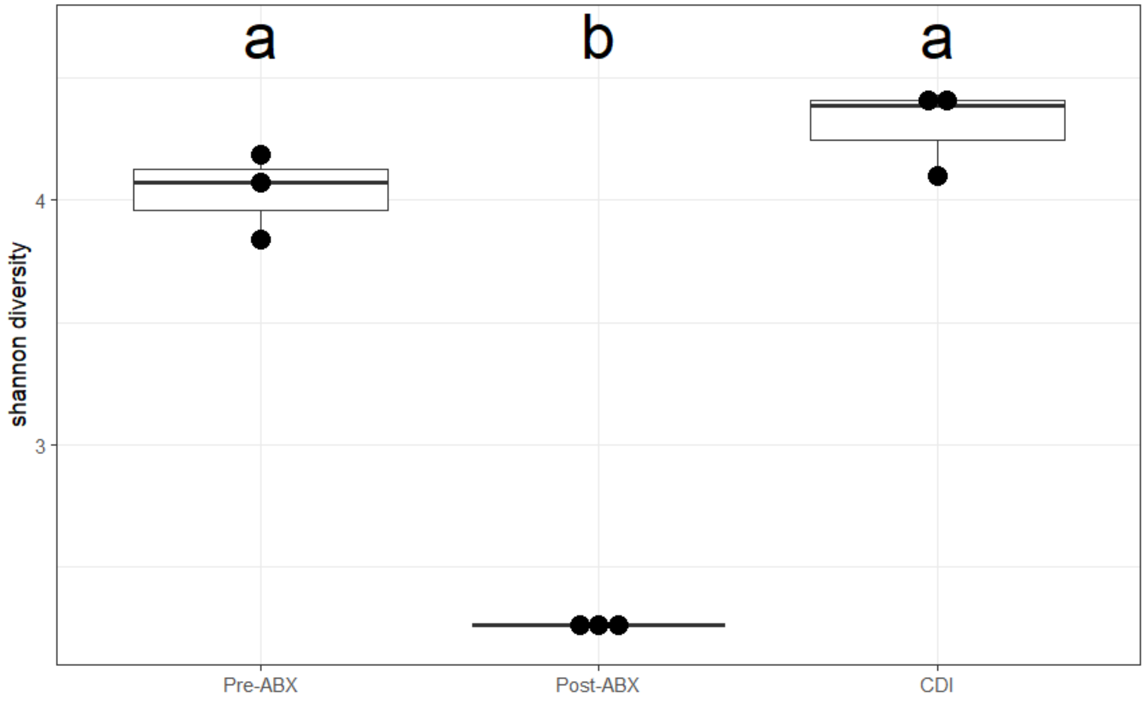

**Supplementary figure 4- Shannon diversity index for whole-communities.** Determined using the relative abundances of metagenome assembled genomes (MAGs) >0.05% represented in the non-redundant contig set in each whole-community sample. One way ANOVA performed between samples in each condition and Tukey HSD post-hoc test to determine significance ($p<0.05$). ABX= antibiotics, CDI=*Clostridioides difficile* infection.

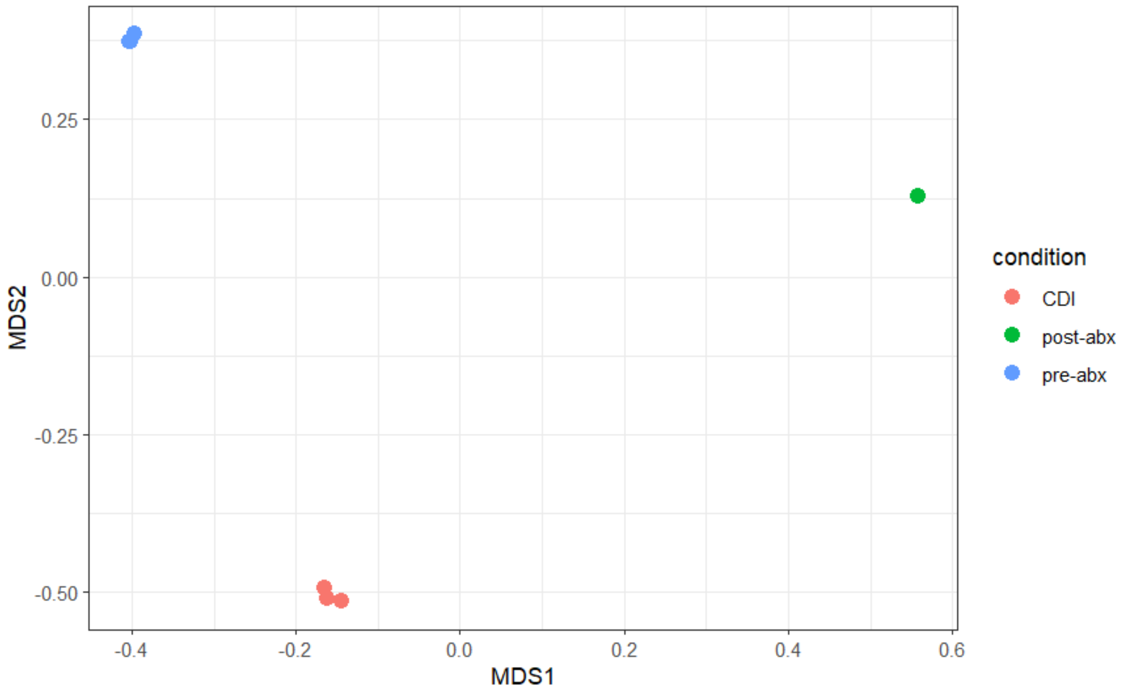


**Supplementary figure 5 - Multi-dimensional scaling visualization of the whole community (WC) taxonomic compositions across conditions based on Bray-Curtis dissimilarity.** Determined using the relative abundances of metagenome assembled genomes (MAGs) >0.05% represented in the non-redundant contig set in each whole-community sample. ABX= antibiotics, CDI=*Clostridioides difficile* infection.

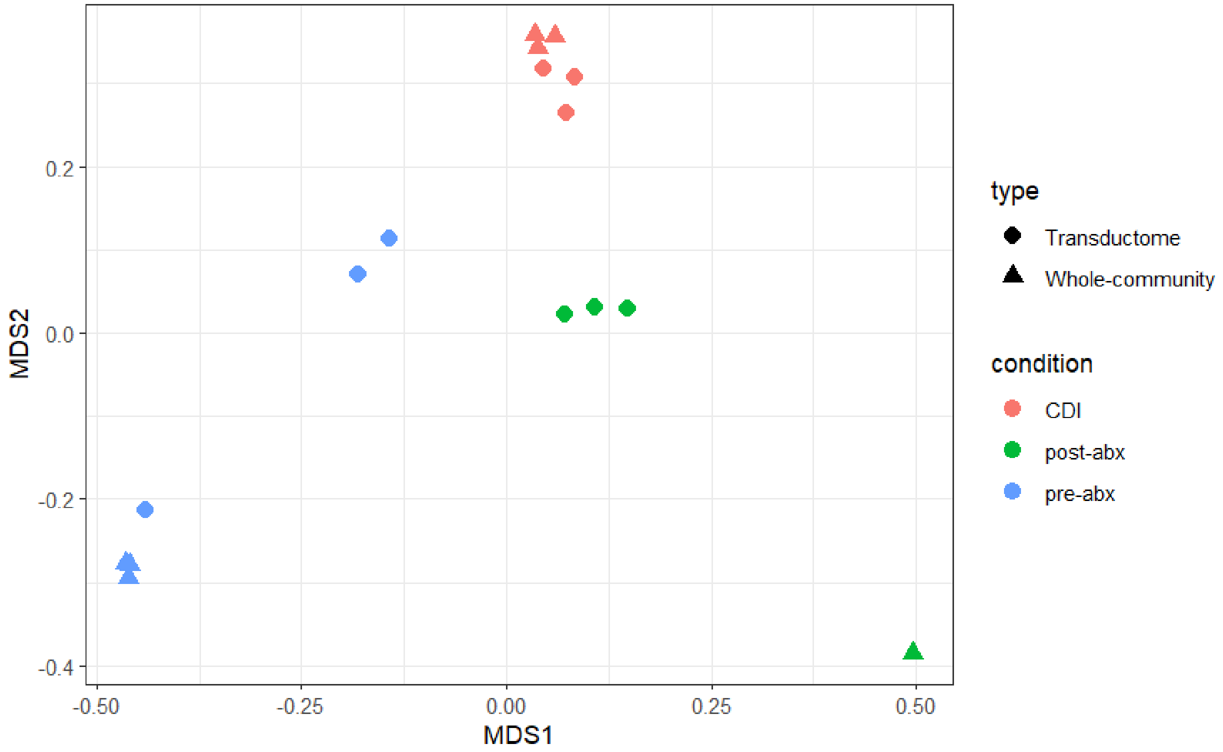


**Supplementary figure 6 - Multi-dimensional scaling visualization of the whole community (WC) and transductome taxonomic compositions across conditions based on Bray-Curtis dissimilarity.** Determined using the relative abundances of metagenome assembled genomes (MAGs) >0.05% represented in the non-redundant contig set in each whole-community and VLP-fraction sample. ABX= antibiotics, CDI=*Clostridioides difficile* infection.

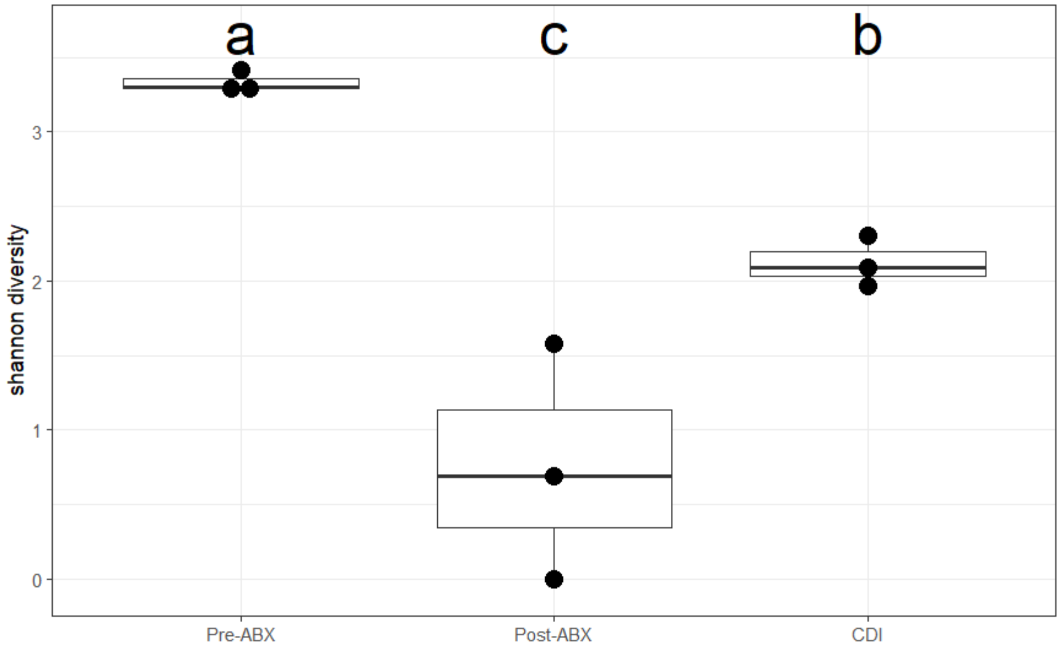


**Supplementary figure 7 - Shannon diversity index for transductomes.** One way ANOVA performed between samples in each condition and Tukey HSD post-hoc test to determine significance ($p<0.05$). Determined using the relative abundances of metagenome assembled genomes (MAGs) >0.05% represented in the non-redundant contig set in each VLP-fraction sample. ABX= antibiotics, CDI=*Clostridioides difficile* infection.

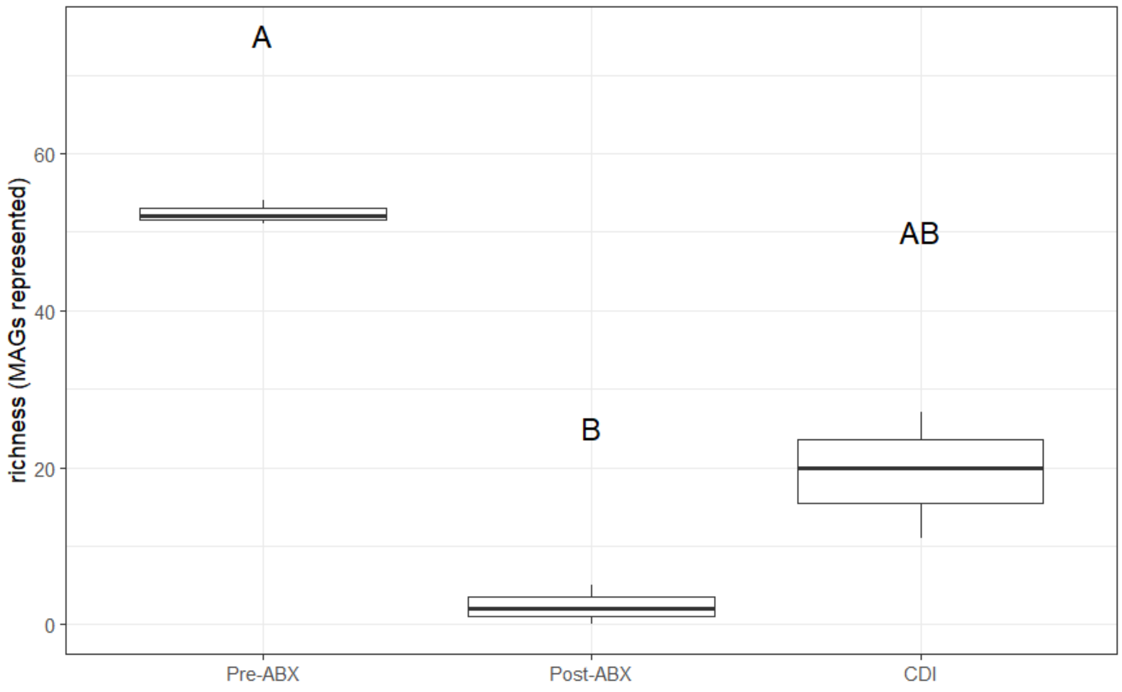

**Supplementary figure 8 - Richness for transductomes.** Determined via the number of metagenome assembled genomes (MAGs) with >0.05% relative abundance represented within the contigs present in each VLP-fraction sample. Kruskal-Wallis test performed between samples in each condition  to determine significance and Dunn test for post-hoc analysis ($p<0.05$). ABX= antibiotics, CDI=*Clostridioides difficile* infection.

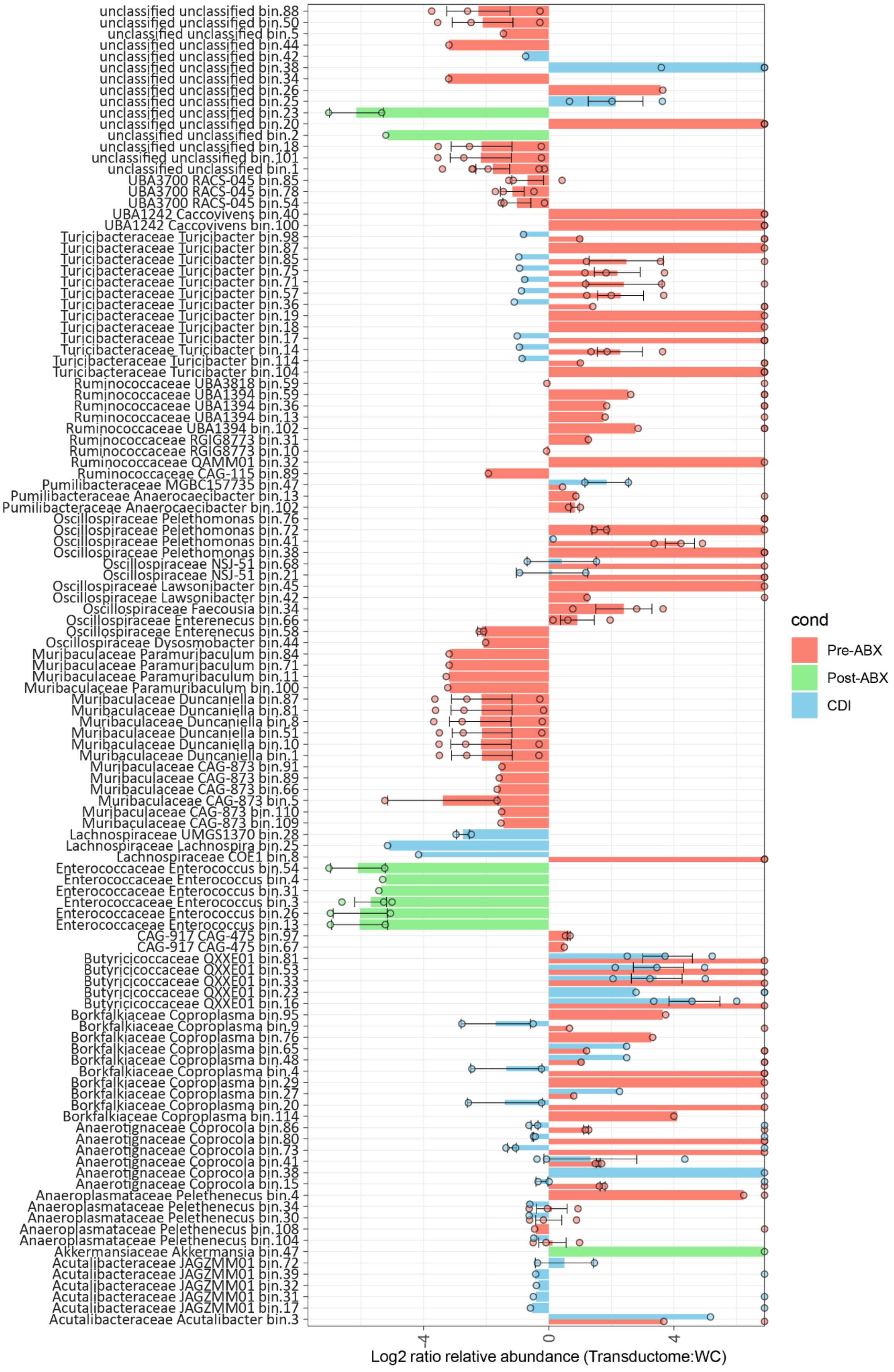

unclassified unclassified bin.88
unclassified unclassified bin.50
unclassified unclassified bin.5
unclassified unclassified bin.44
unclassified unclassified bin.42
unclassified unclassified bin.38
unclassified unclassified bin.34
unclassified unclassified bin.26
unclassified unclassified bin.25
unclassified unclassified bin.23
unclassified unclassified bin.20
unclassified unclassified bin.2
unclassified unclassified bin.18
unclassified unclassified bin.101
unclassified unclassified bin.1
UBA3700 RACS-045 bin.85
UBA3700 RACS-045 bin.78
UBA3700 RACS-045 bin.54
UBA1242 Caccovivens bin.40
UBA1242 Caccovivens bin.100
Turicibacteraceae Turicibacter bin.98
Turicibacteraceae Turicibacter bin.87
Turicibacteraceae Turicibacter bin.85
Turicibacteraceae Turicibacter bin.75
Turicibacteraceae Turicibacter bin.71
Turicibacteraceae Turicibacter bin.57
Turicibacteraceae Turicibacter bin.36
Turicibacteraceae Turicibacter bin.19
Turicibacteraceae Turicibacter bin.18
Turicibacteraceae Turicibacter bin.17
Turicibacteraceae Turicibacter bin.14
Turicibacteraceae Turicibacter bin.114
Turicibacteraceae Turicibacter bin.104
Ruminococcaceae UBA3818 bin.59
Ruminococcaceae UBA1394 bin.59
Ruminococcaceae UBA1394 bin.36
Ruminococcaceae UBA1394 bin.13
Ruminococcaceae UBA1394 bin.102
Ruminococcaceae RGIG8773 bin.31
Ruminococcaceae RGIG8773 bin.10
Ruminococcaceae QAMM01 bin.32
Ruminococcaceae CAG-115 bin.89
Pumilibacteraceae MGBC157735 bin.47
Pumilibacteraceae Anaerocaecibacter bin.13
Pumilibacteraceae Anaerocaecibacter bin.102
Oscillospiraceae Pelethomonas bin.76
Oscillospiraceae Pelethomonas bin.72
Oscillospiraceae Pelethomonas bin.41
Oscillospiraceae Pelethomonas bin.38
Oscillospiraceae NSJ-51 bin.68
Oscillospiraceae NSJ-51 bin.21
Oscillospiraceae Lawsonibacter bin.45
Oscillospiraceae Lawsonibacter bin.42
Oscillospiraceae Faecousia bin.34
Oscillospiraceae Enterenecus bin.66
Oscillospiraceae Enterenecus bin.58
Oscillospiraceae Dysosmobacter bin.44
Muribaculaceae Paramuribaculum bin.84
Muribaculaceae Paramuribaculum bin.71
Muribaculaceae Paramuribaculum bin.11
Muribaculaceae Paramuribaculum bin.100
Muribaculaceae Duncaniella bin.87
Muribaculaceae Duncaniella bin.81
Muribaculaceae Duncaniella bin.8
Muribaculaceae Duncaniella bin.51
Muribaculaceae Duncaniella bin.10
Muribaculaceae Duncaniella bin.1
Muribaculaceae CAG-873 bin.91
Muribaculaceae CAG-873 bin.89
Muribaculaceae CAG-873 bin.66
Muribaculaceae CAG-873 bin.5
Muribaculaceae CAG-873 bin.110
Muribaculaceae CAG-873 bin.109
Lachnospiraceae UMGS1370 bin.28
Lachnospiraceae Lachnospira bin.25
Lachnospiraceae COE1 bin.8
Enterococcaceae Enterococcus bin.54
Enterococcaceae Enterococcus bin.4
Enterococcaceae Enterococcus bin.31
Enterococcaceae Enterococcus bin.3
Enterococcaceae Enterococcus bin.26
Enterococcaceae Enterococcus bin.13
CAG-917 CAG-475 bin.97
CAG-917 CAG-475 bin.67
Butyricicoccaceae QXXE01 bin.81
Butyricicoccaceae QXXE01 bin.53
Butyricicoccaceae QXXE01 bin.33
Butyricicoccaceae QXXE01 bin.23
Butyricicoccaceae QXXE01 bin.16
Borkfalkiaceae Coproplasma bin.95
Borkfalkiaceae Coproplasma bin.9
Borkfalkiaceae Coproplasma bin.76
Borkfalkiaceae Coproplasma bin.65
Borkfalkiaceae Coproplasma bin.48
Borkfalkiaceae Coproplasma bin.4
Borkfalkiaceae Coproplasma bin.29
Borkfalkiaceae Coproplasma bin.27
Borkfalkiaceae Coproplasma bin.20
Borkfalkiaceae Coproplasma bin.114
Anaerotignaceae Coprocola bin.86
Anaerotignaceae Coprocola bin.80
Anaerotignaceae Coprocola bin.73
Anaerotignaceae Coprocola bin.41
Anaerotignaceae Coprocola bin.38
Anaerotignaceae Coprocola bin.15
Anaeroplasmataceae Pelethenecus bin.4
Anaeroplasmataceae Pelethenecus bin.34
Anaeroplasmataceae Pelethenecus bin.30
Anaeroplasmataceae Pelethenecus bin.108
Anaeroplasmataceae Pelethenecus bin.104
Akkermansiaceae Akkermansia bin.47
Acutalibacteraceae JAGZMM01 bin.72
Acutalibacteraceae JAGZMM01 bin.39
Acutalibacteraceae JAGZMM01 bin.32
Acutalibacteraceae JAGZMM01 bin.31
Acutalibacteraceae JAGZMM01 bin.17
Acutalibacteraceae Acutalibacter bin.3
-4
0
4
Log2 ratio relative abundance (Transductome:WC)
cond
Pre-ABX
Post-ABX
CDI

**Supplementary figure 9 - Log2 ratio of MAG-level relative abundances between transductomes and whole-communities (WCs).** Bars represent averages between replicate points. Points or bars at the edge of the plot represent infinite values. Zero values were removed.

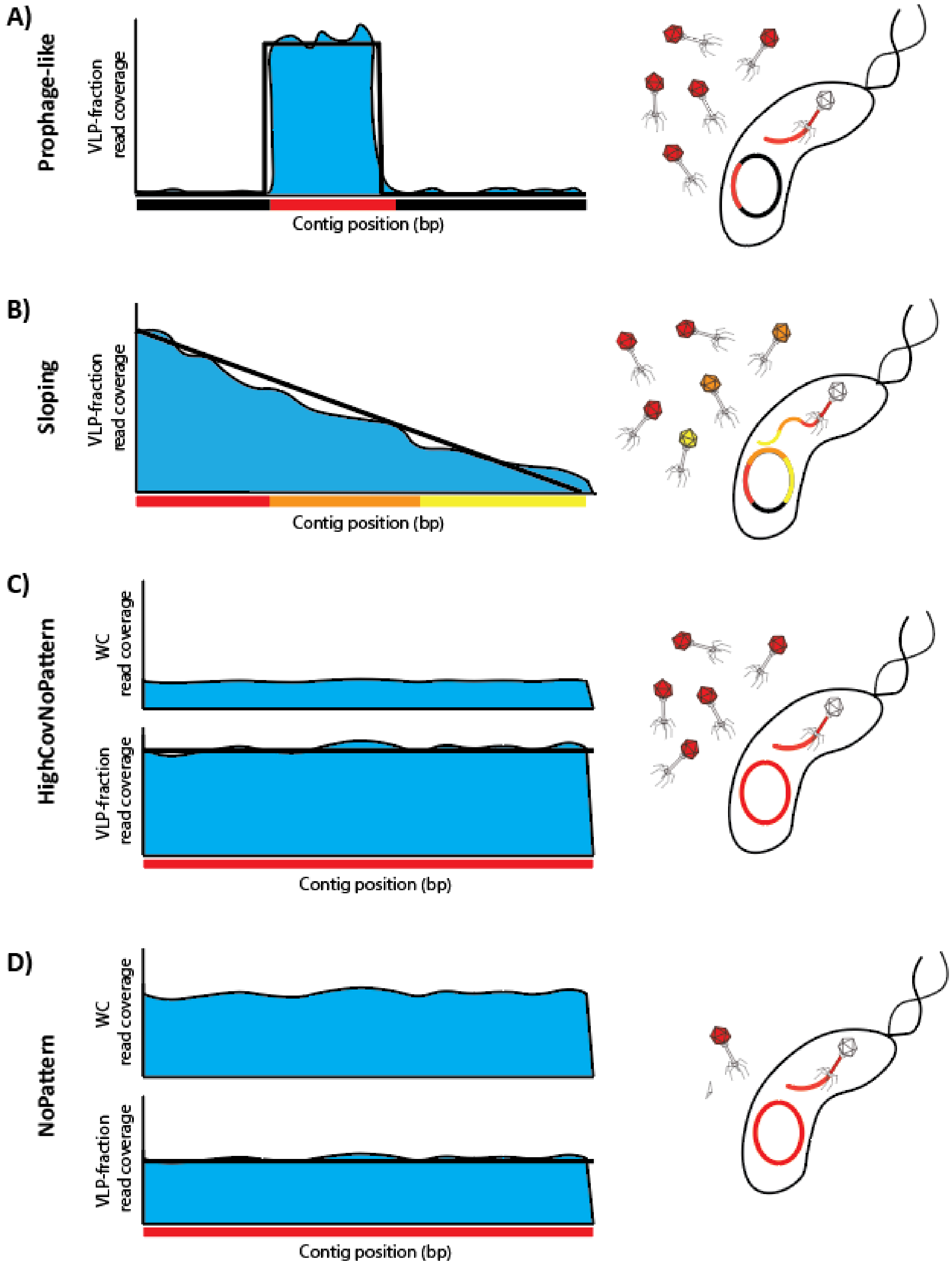

**Supplementary figure 10 - TrIdent pattern-Matching classifications and examples of their biological implications.** Virus-like particle (VLP)-fraction read coverage is represented in blue and TrIdent pattern-match in thick black line. The bacterial genome region packaged into VLPs is represented by colored bars along the contig. A) The 'Prophage-like' classification is given to contigs with a defined block of coverage in the VLP-fraction. This pattern is produced by mobile genetic elements that cleanly excise from the bacterial genome and are preferentially packaged into VLPs. B) The 'Sloping' classification is given to contigs with sloping VLP-fraction read coverage. This pattern is produced by the sequential packaging of large regions of the bacterial genome starting at a preferred packing site. The packaging frequency decreases as the distance from the packaging initiation site increases. C) The 'HighCovNoPattern' classification is given to contigs with even VLP-fraction read coverage patterns that are relatively enriched compared to the contig's whole-community (WC) coverage. This pattern may be produced by non-biased packaging of large genome regions. D) The 'NoPattern' classification is given to contigs with even VLP-fraction read coverage patterns that are not enriched relative to the WC coverage. This pattern may be associated with little to no VLP-packaged DNA or larger amounts of VLP-packaged DNA albeit not enriched.

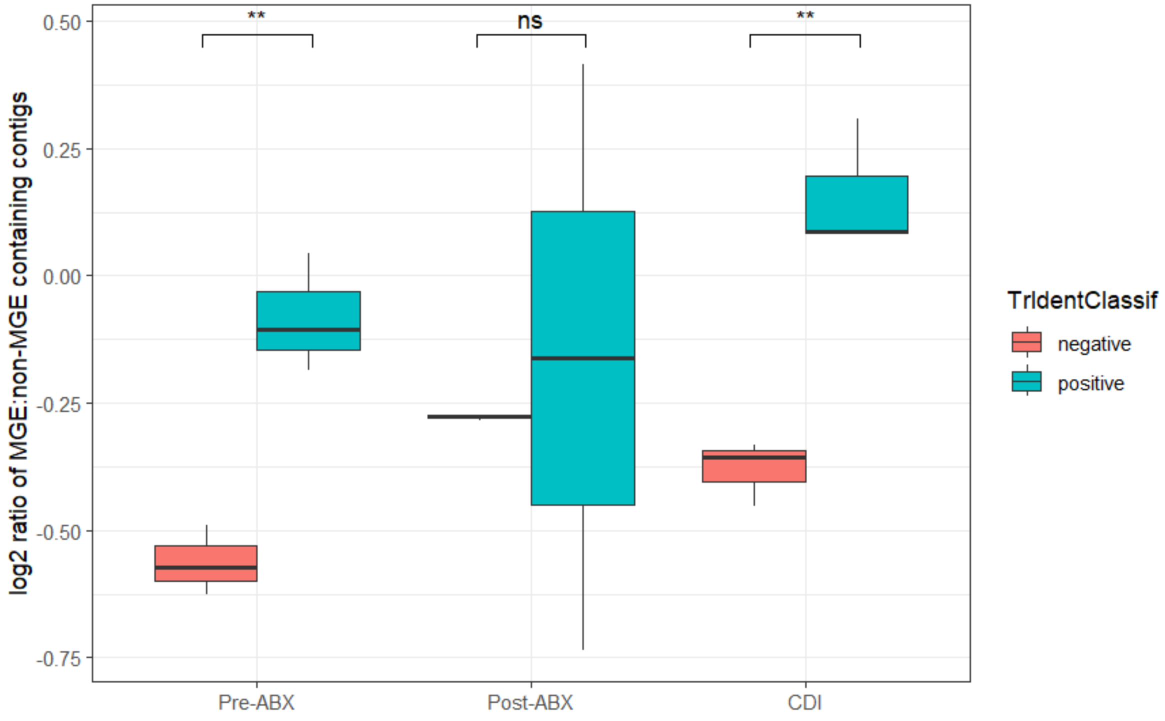


**Supplementary figure 11- Ratio of mobile genetic element (MGE)- containing contigs to non-MGE containing contigs for all positively and negatively TrIdent classified contigs.** Positive classifications include 'Prophage-like', 'Sloping' and 'HighCovNoPattern' whereas negative classifications include 'NoPattern' classifications and contigs that were filtered out for low VLP-fraction read coverage. Statistical significance between negative and positive TrIdent classification for each sample condition was determined with a t-test ($p<0.05$). ABX= antibiotics, CDI=*Clostridioides difficile* infection.

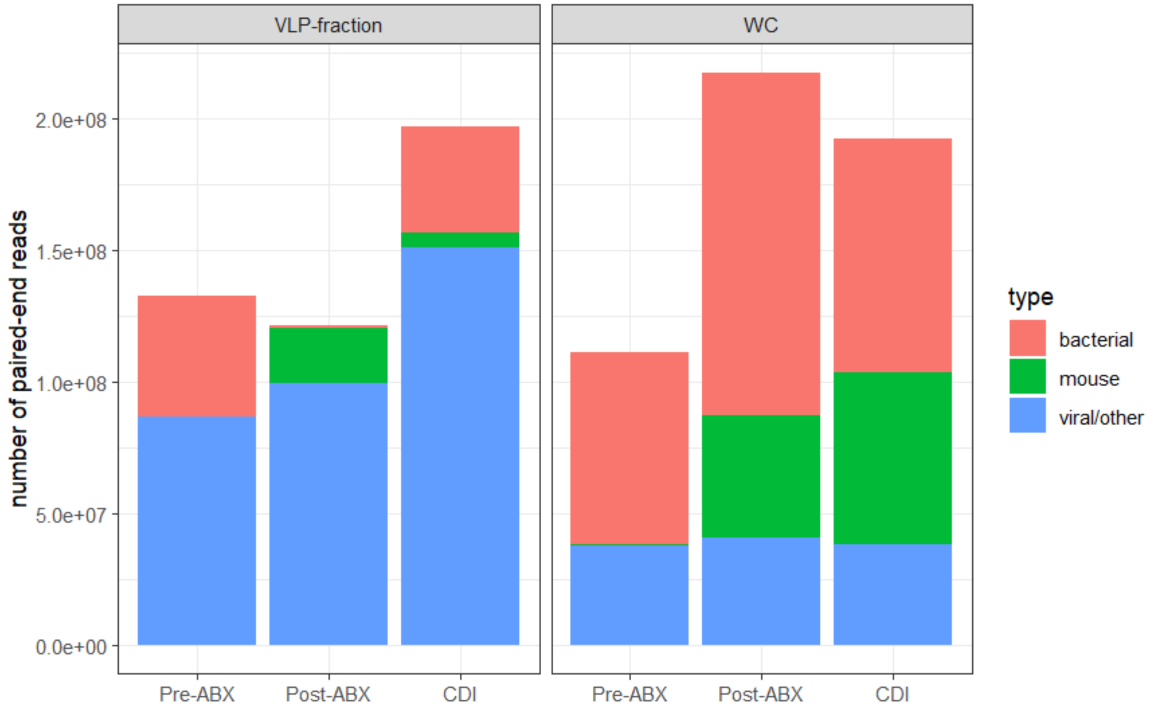


**Supplementary figure 12 - Number of trimmed whole-community (WC) and virus-like particle (VLP)-fractions sequencing reads that are bacterial, mouse, or viral/other.** Bacterial reads were mapped to the non-redundant contig set which consists only of bacterial contigs. Mouse reads were mapped to the host mouse reference genome, and viral/other reads were unmapped. Reads were averaged across replicates. ABX= antibiotics, CDI=*Clostridioides difficile* infection.

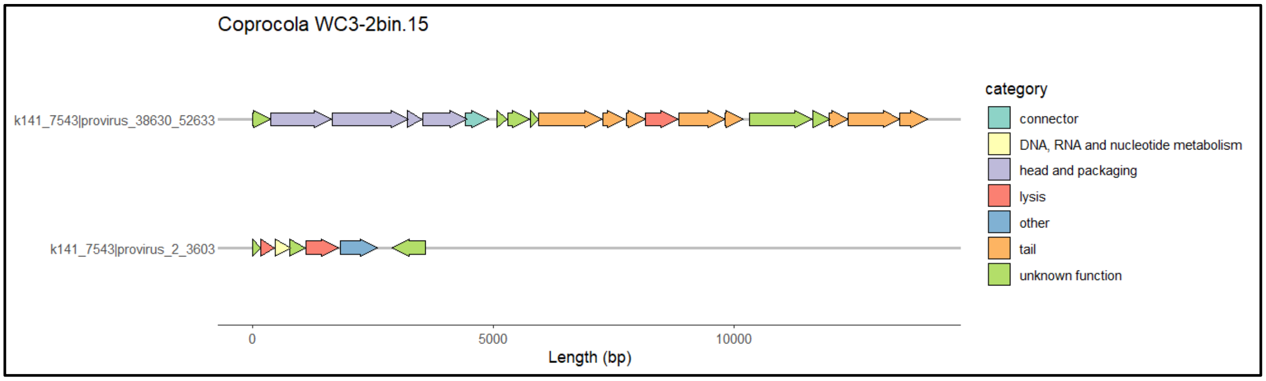

**Supplementary figure 13 - Pharokka annotation categories of geNomad predicted prophage in Anaerotignaceae *Coprocola* MAG WC3-2bin15.** A detailed view of the two predicted prophage/GTAs can be found in figure S16.

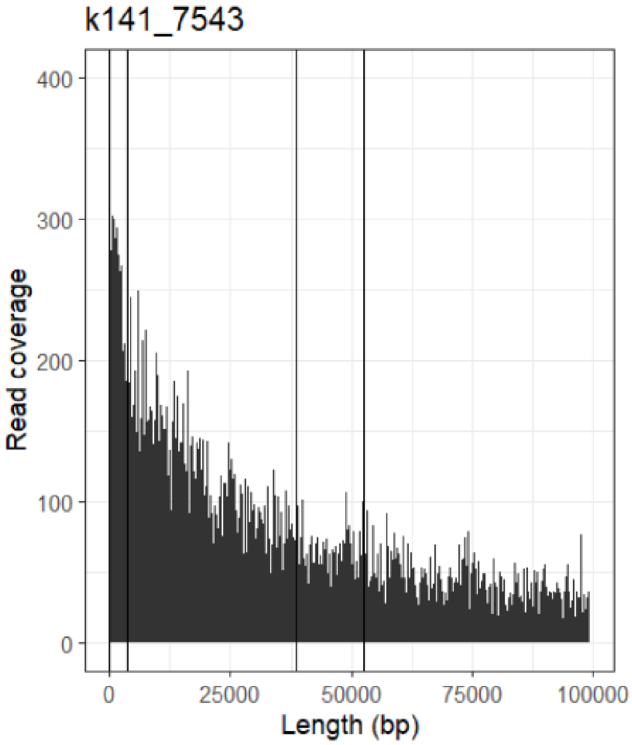


**Supplementary figure 14 - Locations of GTA candidates on contig k141_7543 with VLP-fraction read coverage in Pre-ABX replicate 1.** Start and stop locations (provided by geNomad) are marked with black vertical lines.

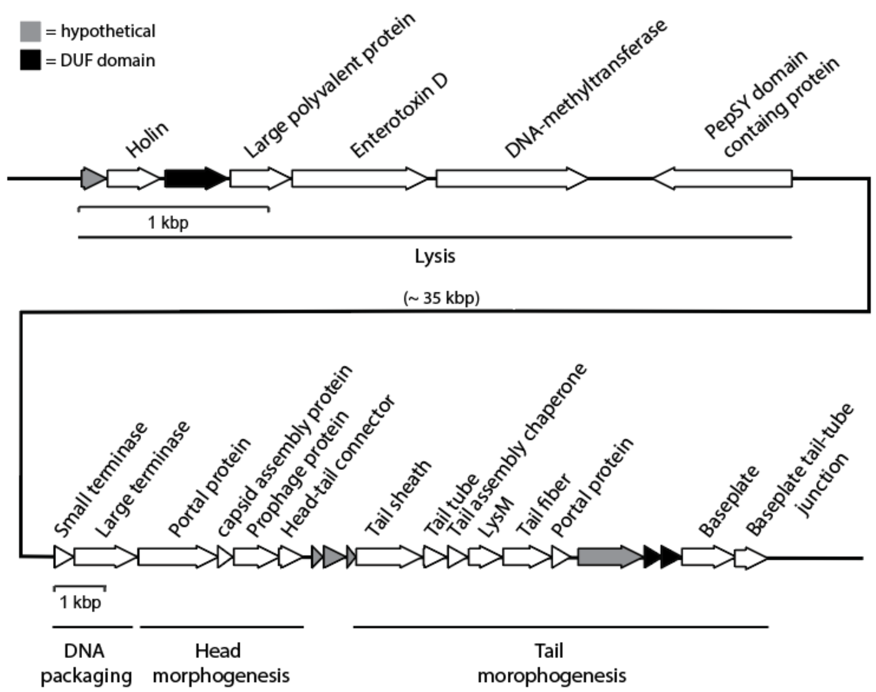


**Supplementary figure 15 - Putative GTA clusters in Anaerotignaceae *Coprocola* contig k141_7543**

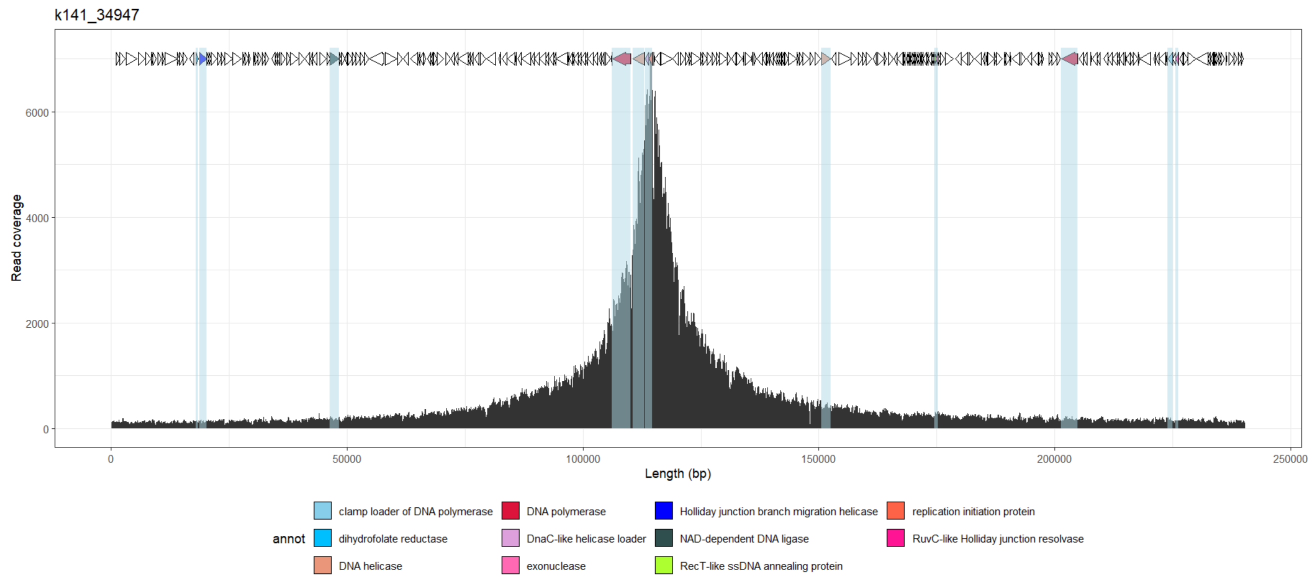


**Supplementary figure 16 - Oscillospiraceae *Pelethomonas* contig k141_34947 Pharokka gene annotation at VLP-fraction read coverage peak in Pre-ABX replicate 1.** Only genes in Pharokka's 'DNA, RNA and nucleotide metabolism' are colored by their associated Pharokka annotations and are highlighted in blue.

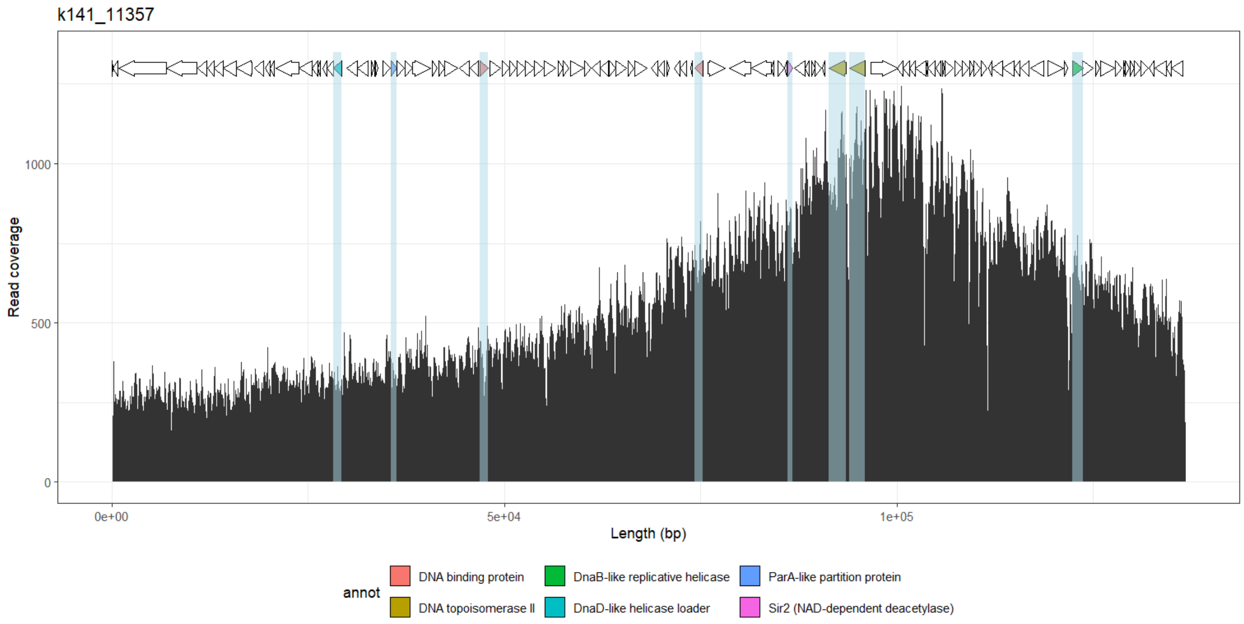


**Supplementary figure 17 - Oscillospiraceae *Pelethomonas* contig k141_11357 Pharokka gene annotation at VLP-fraction read coverage peak in Pre-ABX replicate 1.** Only genes in Pharokka's 'DNA, RNA and nucleotide metabolism' are colored by their associated Pharokka annotations and are highlighted in blue.

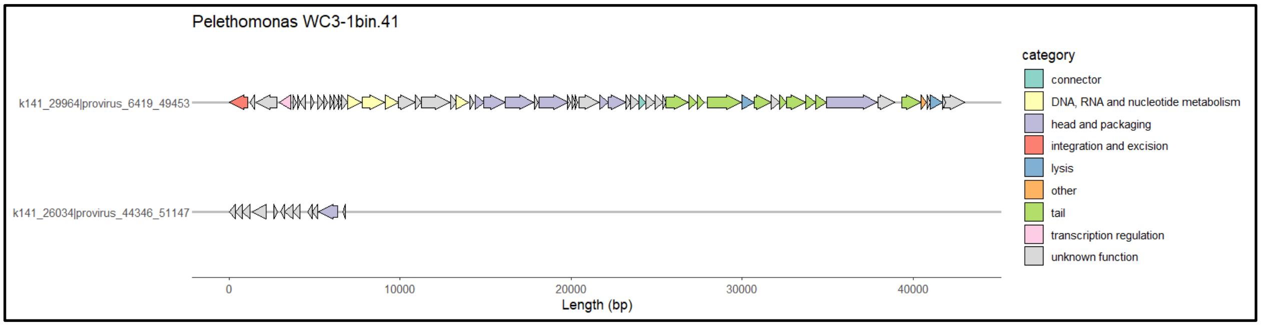


**Supplementary figure 18 - Pharokka annotation categories of geNomad predicted prophage in Oscillospiraceae *Pelethomonas* MAG WC3-1bin41.** The only *Pelethomonas* contigs containing a prophage that fit our GTA candidate criteria was contig k141_26034. The contig had full VLP-fraction coverage with no major difference in coverage at the GTA candidate site. This GTA candidate encodes a phage capsid protein and a conjugal transfer protein and is directly downstream of several tRNAs.

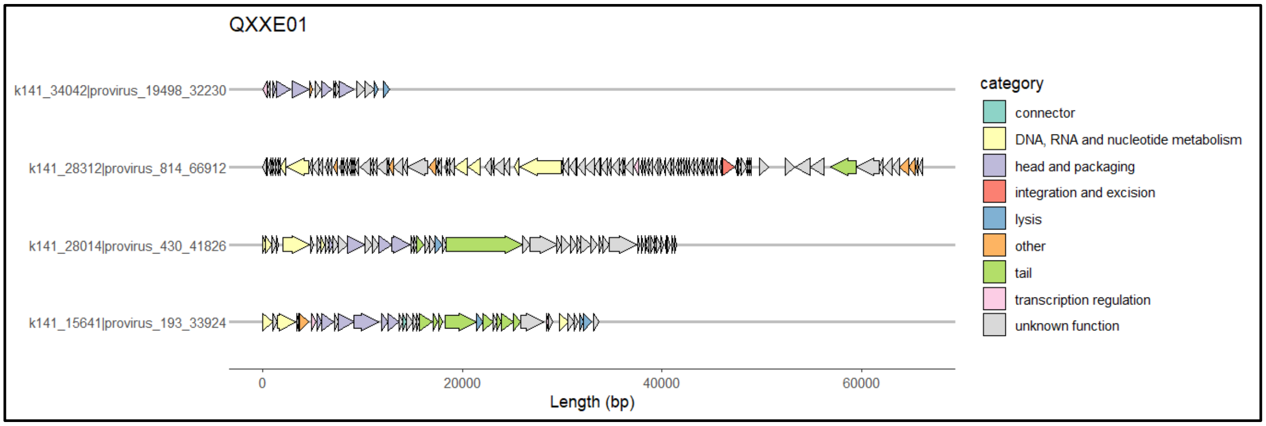


**Supplementary figure 19 -Butyricoccaceae *QXXE01* MAGs predicted prophage Pharokka annotation categories.** Contigs k141_34032, k141_28014, and k141_15641 met our GTA criteria as they did not contain gene annotations in both the 'integration and excision' and 'DNA, RNA and nucleotide metabolism' categories

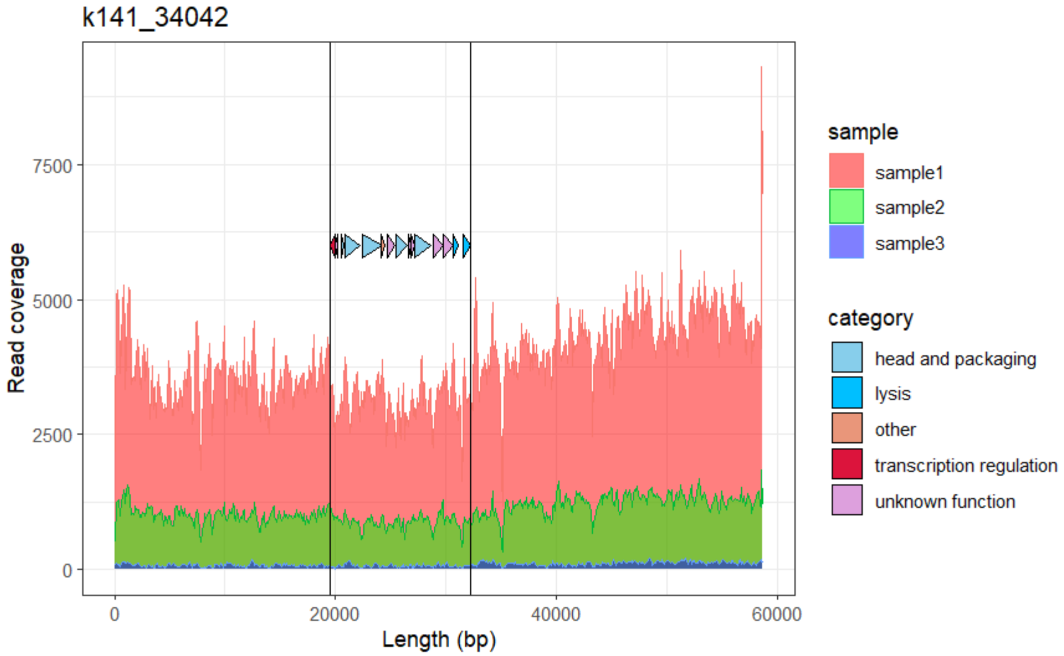


**Supplementary figure 20 - Butyricoccaceae *QXXE01* contig k141_34042 read coverage and Pharokka categories at predicted-prophage site in CDI sample.** We speculate that this is either a GTA cluster or a capsid-forming phage satellite based on the Pharokka gene categories.

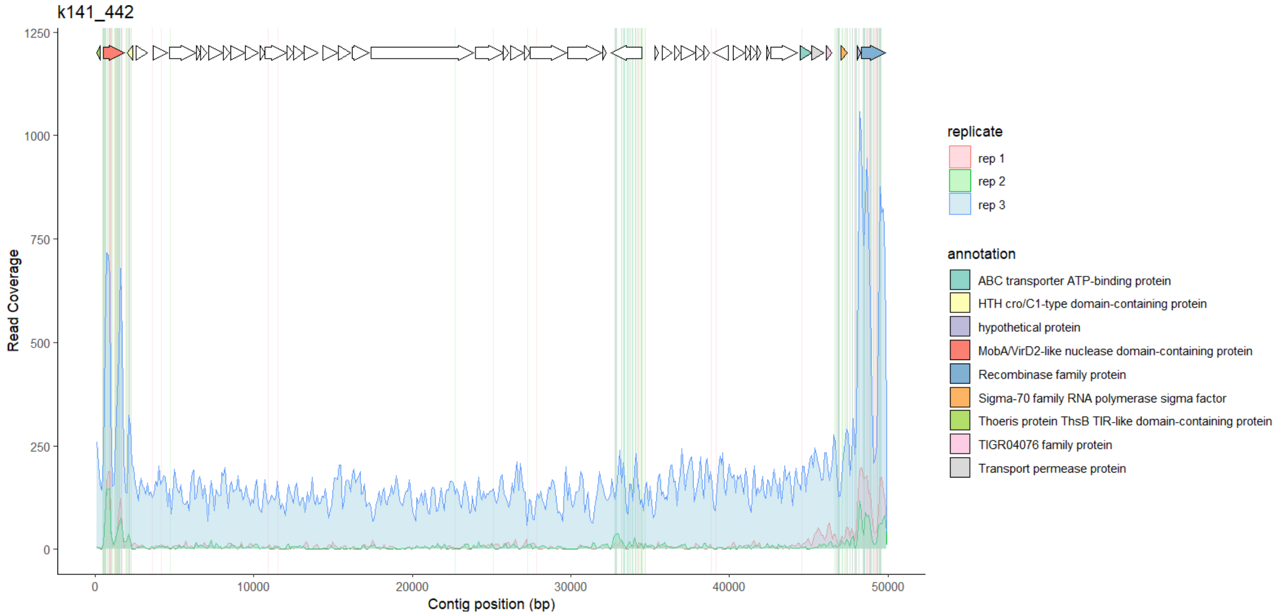


**Supplementary figure 21- Gene annotations at regions associated with elevated Pre-ABX read coverage and high density of read mapping variants.** The read coverage and read mapping variants for Pre-ABX replicates 1 (pink), 2 (blue) and 3 (green) are overlaid with the predicted open reading frames. Only the annotations for the end regions are shown.

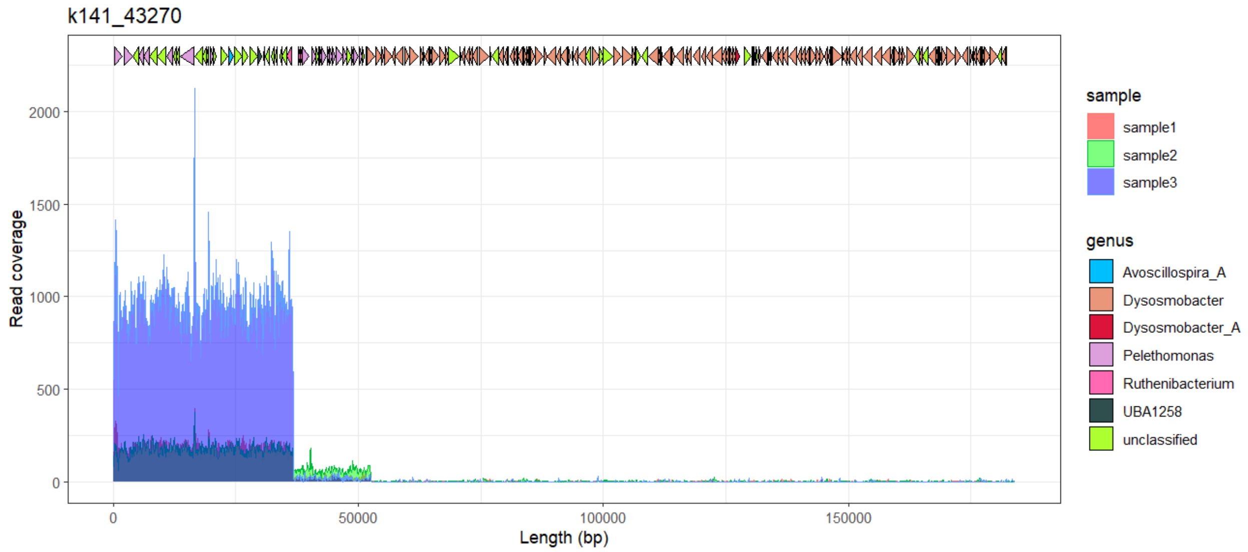


**Supplementary figure 22 - Oscillospiraceae *Dysosmobacter* contig k141_43270 gene taxonomy and read coverage in Pre-ABX replicates.** The read coverage for Pre-ABX replicates 1 (pink), 2 (blue) and 3 (green) are overlaid with the predicted open reading frames. Only the annotations for the end regions are shown.

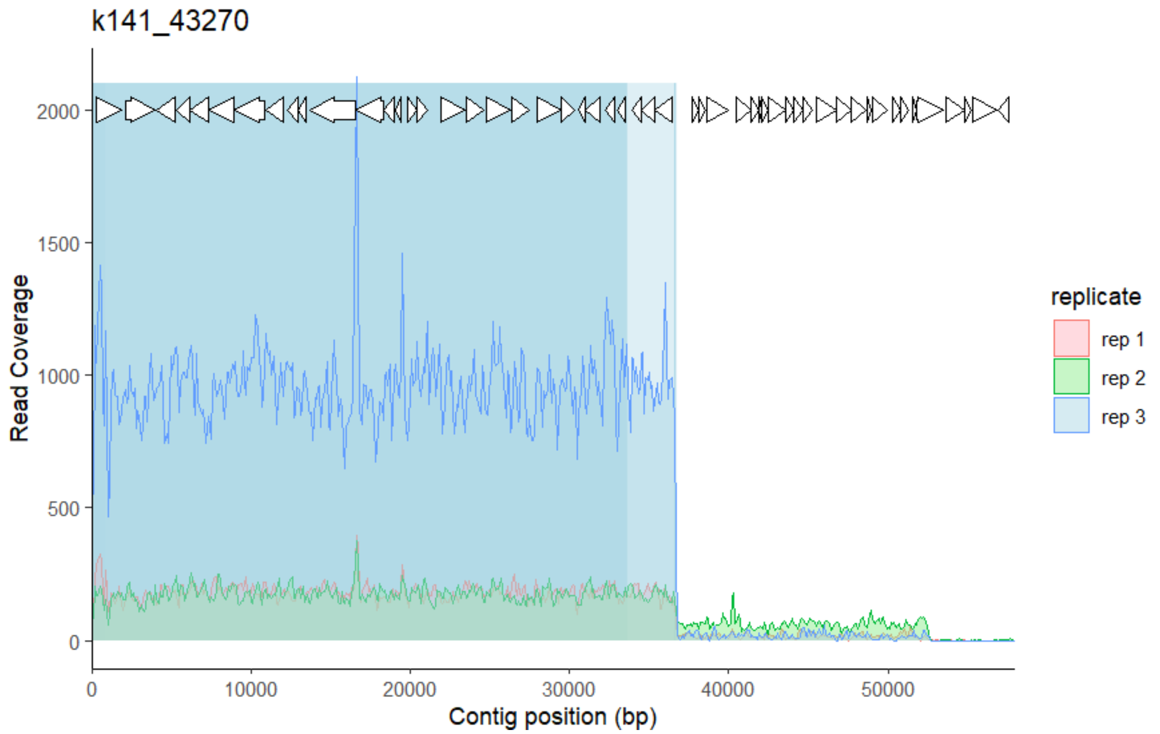


**Supplementary figure 23 - Oscillospiraceae *Dysosmobacter* contig k141_43270 subset with BLASTn hit coverage and read coverage in Pre-ABX replicates**

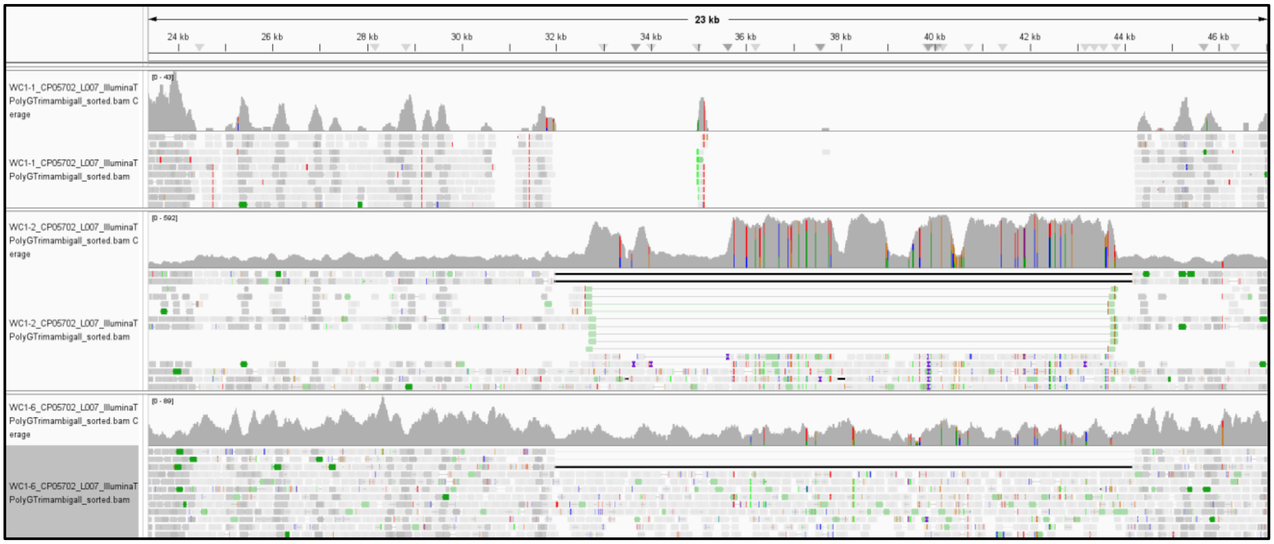


**Supplementary figure 24 - Anaeroplasmataceae *Pelethenecus* contig k141_267558 MGE large inserts in Pre-ABX WC samples.** Bam files of all three WC replicates were visualized in Integrative Genome Viewer (IGV). Light green reads and inserts indicate read pairs in the reverse-forward orientation (indicative of duplication or translocation). Black inserts indicate read pairs in the forward-reverse orientation with inferred inserts that are larger than expected (indicative of deletion).

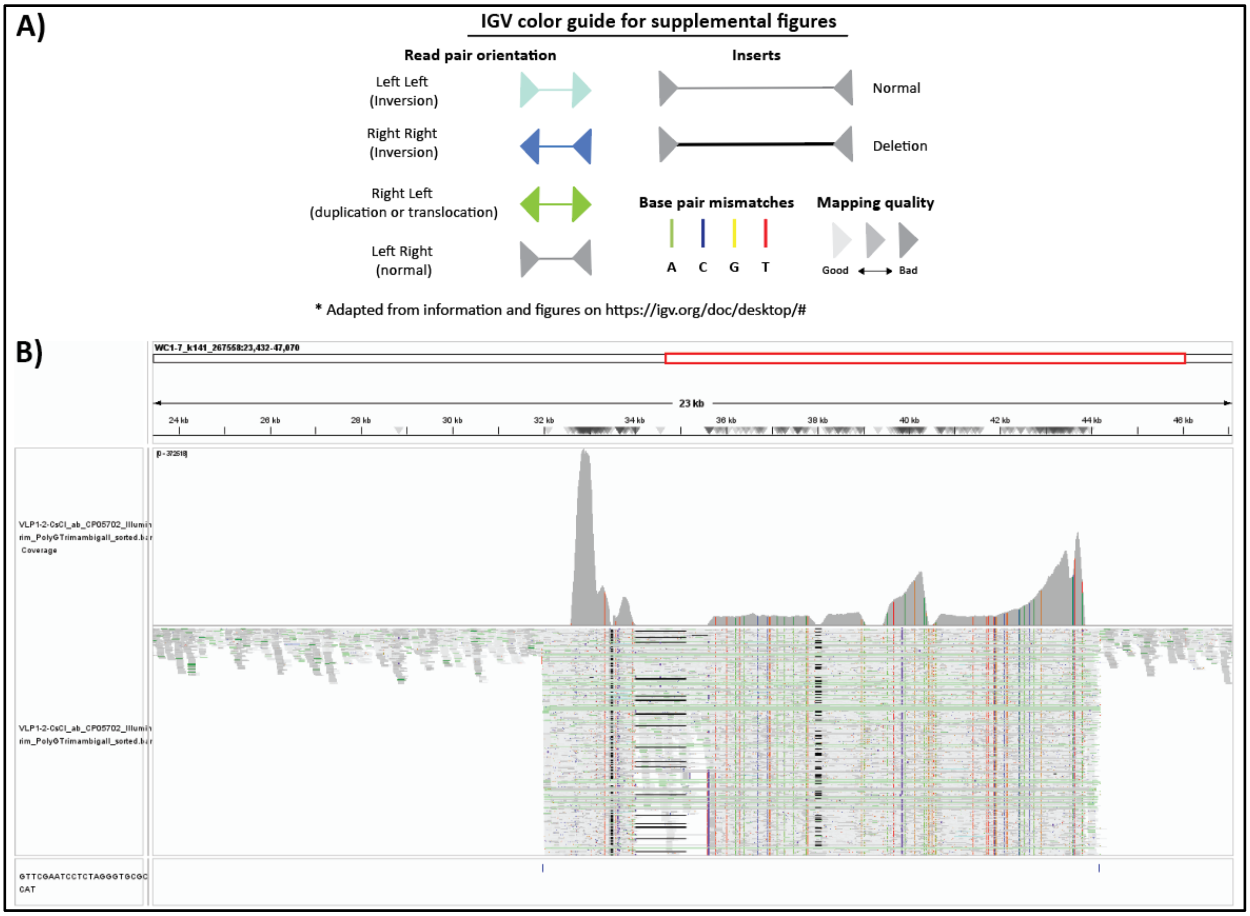


**Supplementary figure 25 - Location of direct repeats (5' GTTCGAATCCTCTAGGGTGCGCCAT 3') flanking the MGE on Anaeroplasmataceae *Pelethenecus* contig k141_267558.** A) IGV color guide for features displayed in IGV plot. B) Read coverage and read mapping visualized in IGV. Small blue vertical lines at bottom denote locations of direct repeats.

```
                                    10        20        30        40        50        60        70
                            ....|....|....|....|....|....|....|....|....|....|....|....|....|....|.
k141_267558_zot             --------MIWVLIAIALVVFYIYFYRKFAVLPDNSIMMLNGGLGSGKTSLAVGASIRDYKSRLRSWKICNVLFPW
CholeraZot_BindingDomain    GRLCVQDG--------------------------------------------------------------------

                                   110       120       130       140       150       160       170
                            ....|....|....|....|....|....|....|....|....|....|....|....|....|....|.
k141_267558_zot             ILERKVRLPYGSVVLIDEFSLVADSMSYKTINNTVLMLFIKLFRQETRGGVMYITTQSLSDCHYAVKRCVNSYLWI
CholeraZot_BindingDomain    ----------------------------------------------------------------------------

                                   210       220       230       240       250       260       270
                            ....|....|....|....|....|....|....|....|....|....|....|....|....|....|.
k141_267558_zot             INTVEDVQESRLYLLTTRCWKYYDTYAFSGLTDNLPIANKVVNGKELPDLKVSSYRDIASYIDYESEVNGNEKKKG
CholeraZot_BindingDomain    ----------------------------------------------------------------------------
```

**Supplementary figure 26 - Pairwise alignment of Anaeroplasmataceae *Pelethenecus* contig k141_267558 Zot protein sequence and Zot receptor-binding domain from *Vibrio cholera***

**Zot protein sequence.** The pairwise alignment was performed with the BioEdit software using the option 'allow sequence ends to slide'. There were 0 identities and 0 similarities.

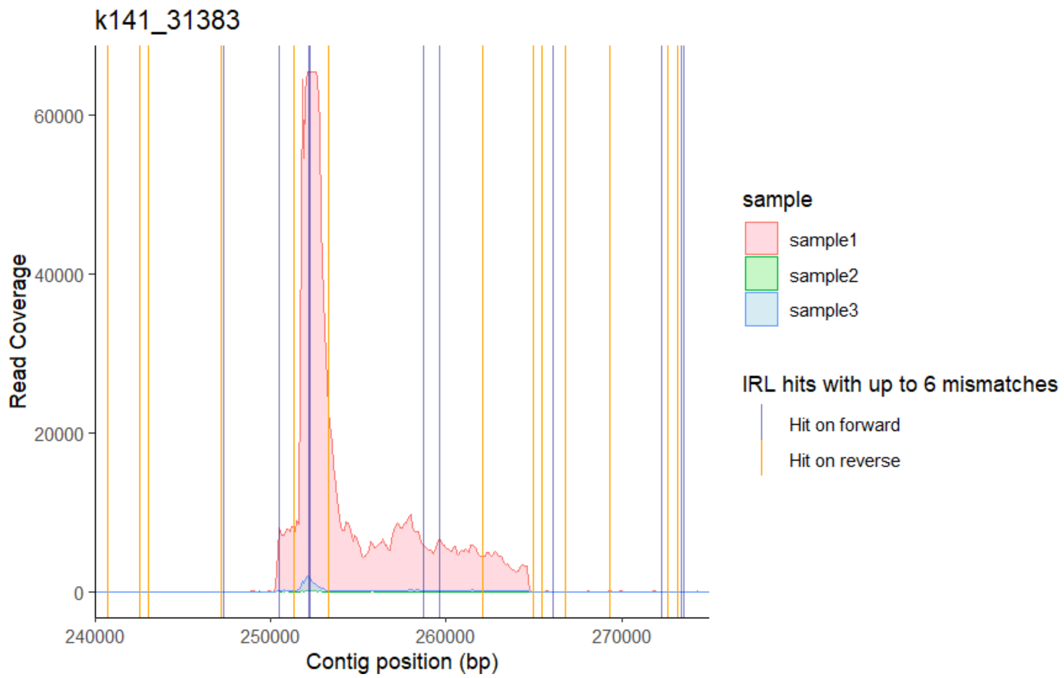


**Supplementary figure 27 - Pumilibacteraceae *MGBC157735* contig k141_31383 subset with IRL hit locations and read coverage in CDI samples**

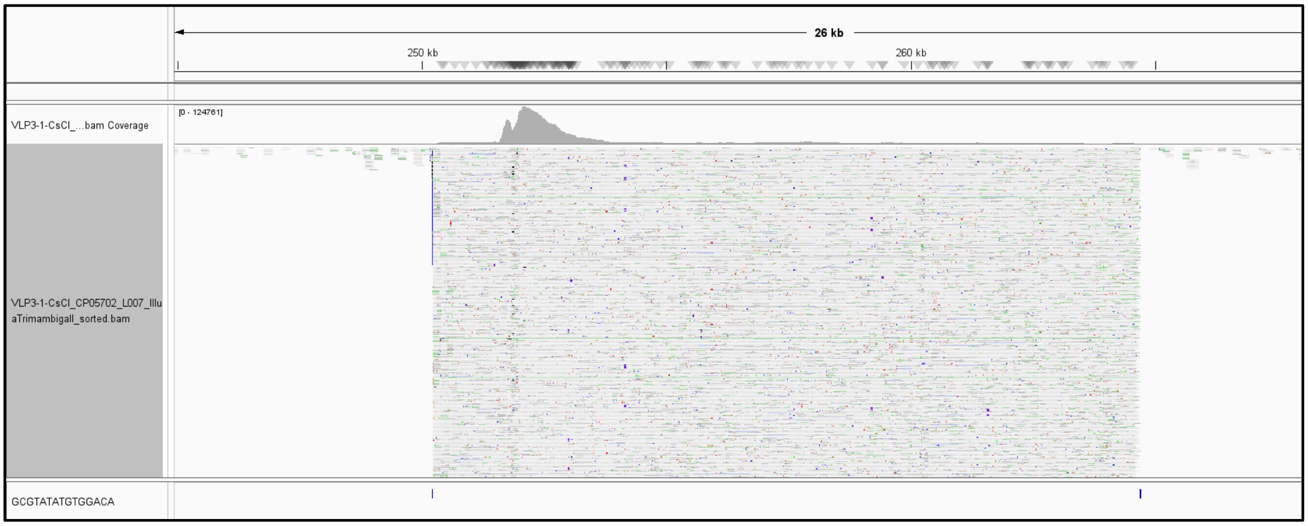


**Supplementary figure 28 - Location of direct repeats (5' GCGTATATGTGGACA 3') flanking the MGE on Pumilibacteraceae *MGBC157735* contig k141_31383.** Visualized in IGV. Small blue vertical lines at bottom denote locations of direct repeats. See Fig. S25A for IGV color guide.

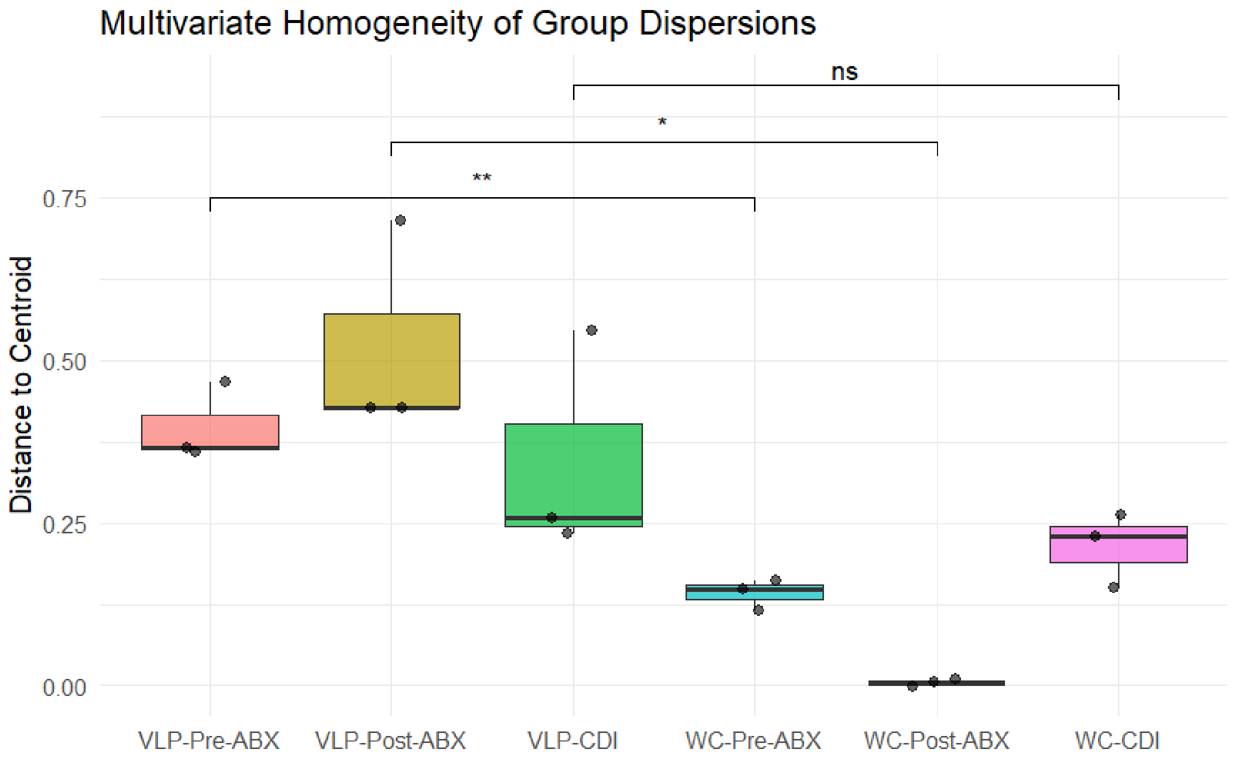


**Supplementary figure 29 - Dispersion of beta diversity values between groups.** VLP= virus-like particle, WC= whole-community. ABX=antibiotics, CDI= *Clostridioides difficile* infection. Statistical significance determined with a t-test between group means. Beta diversity values originate from Bray-Curtis dissimilarity analysis of WC and transductome compositions (Fig. S6)

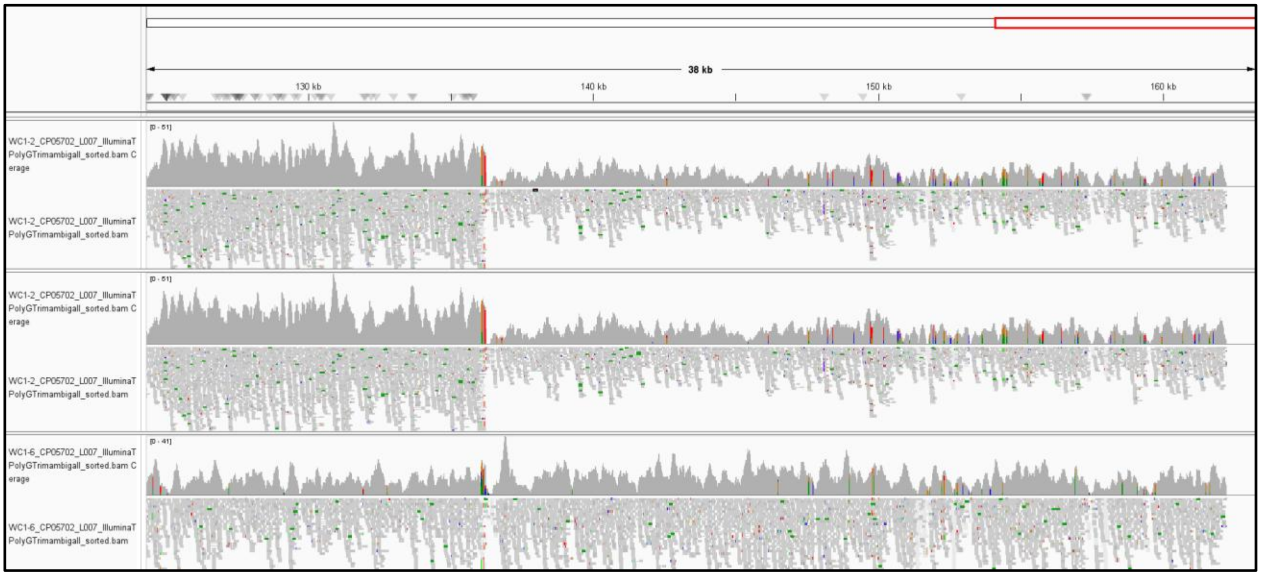


**Supplementary figure 30 - IGV read coverage of WC Pre-ABX replicates 1-3 at coverage discontinuity in contig k141_22490.** See Fig. S25A for IGV color guide.

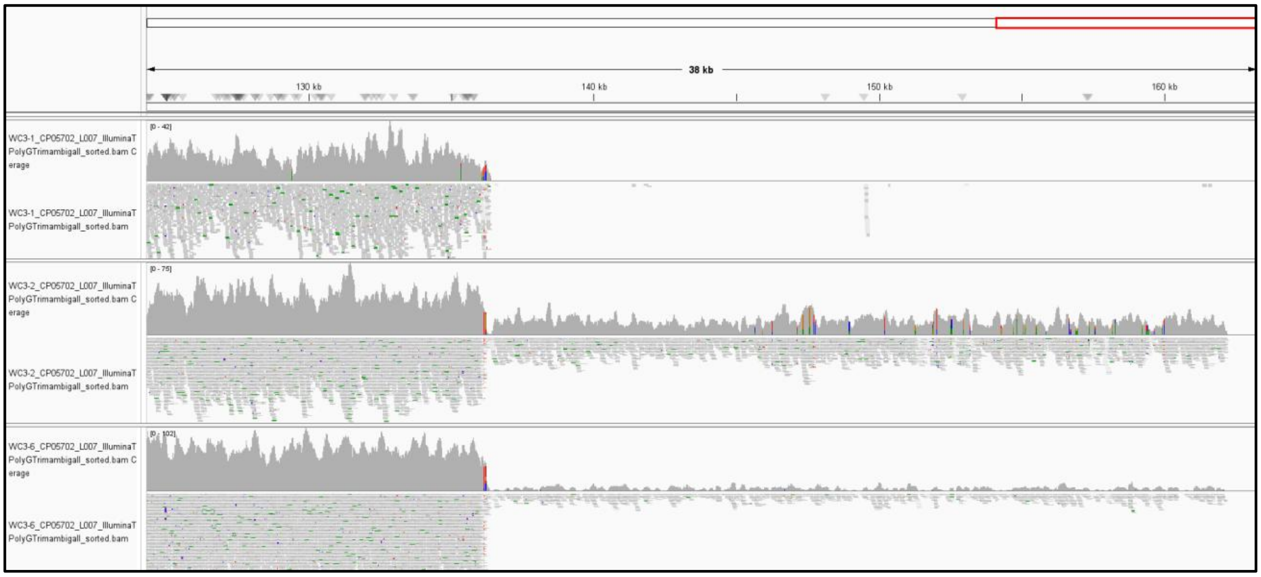


**Supplementary figure 31 - IGV read coverage of WC CDI replicates 1-3 at coverage discontinuity in contig k141_22490.** See Fig. S25A for IGV color guide.

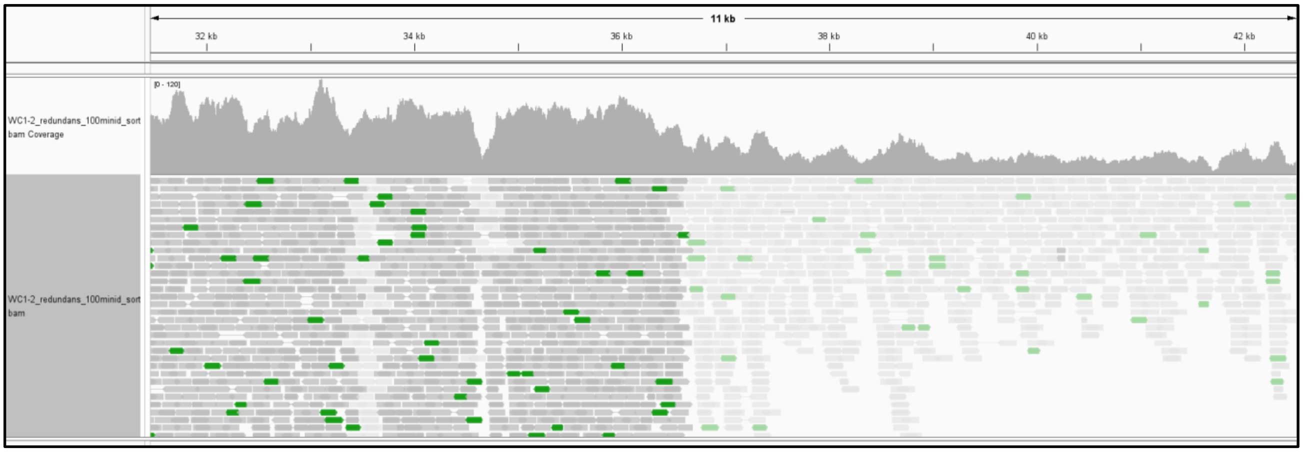


**Supplementary figure 32 - Oscillospiraceae *Dysosmobacter* contig k141_43270 read mapping at potential chimeric junction region in Pre-ABX WC replicate 2 mapped at 100% minimum sequence ID.** Contig k141_43270 originates from the Pre-ABX WC replicate 2 assembly. The read pairs that span the drop in coverage indicate that this is not a chimeric assembly but rather a genomic region that is physically present within the sample. The dark grey reads indicate ambiguous regions where reads may map to more than one location in the dataset. See Fig. S25A for IGV color guide.

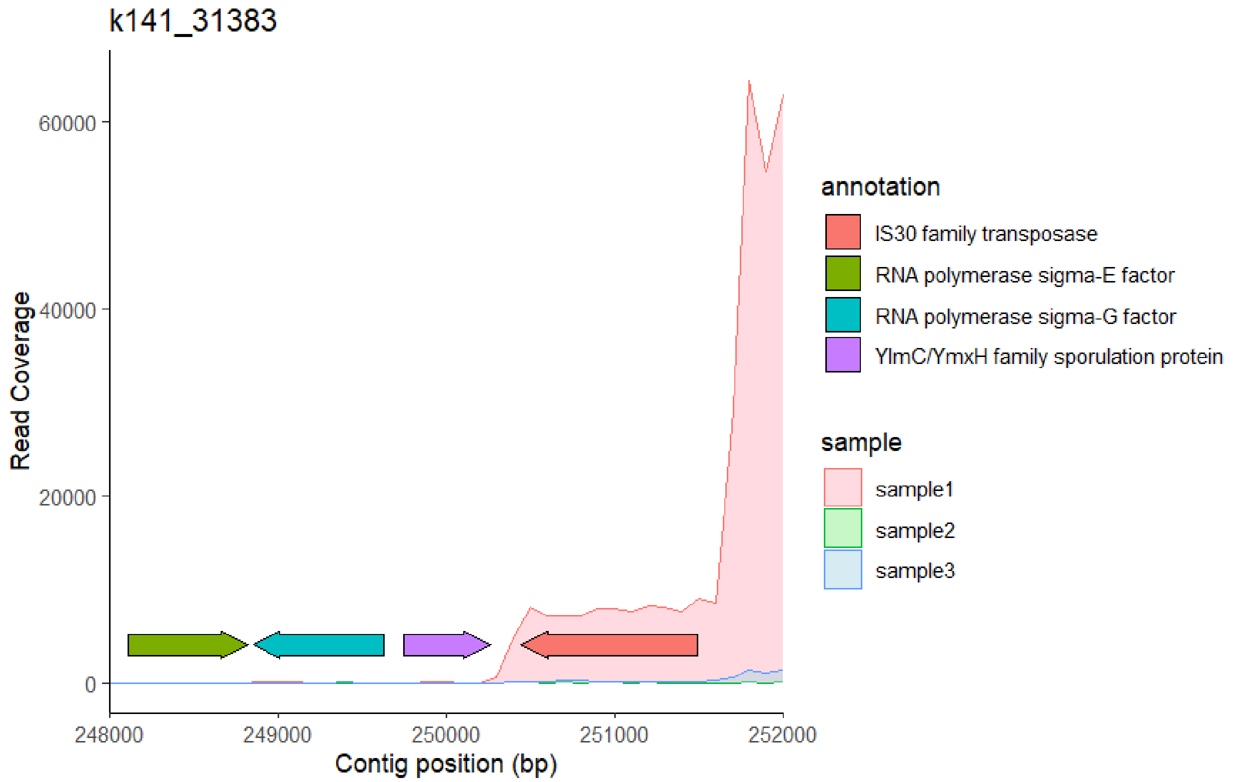


**Supplementary figure 33 - Pumilibacteraceae *MGBC157735* contig k141_31383 subset with sigma factor locations and read coverage in CDI samples**

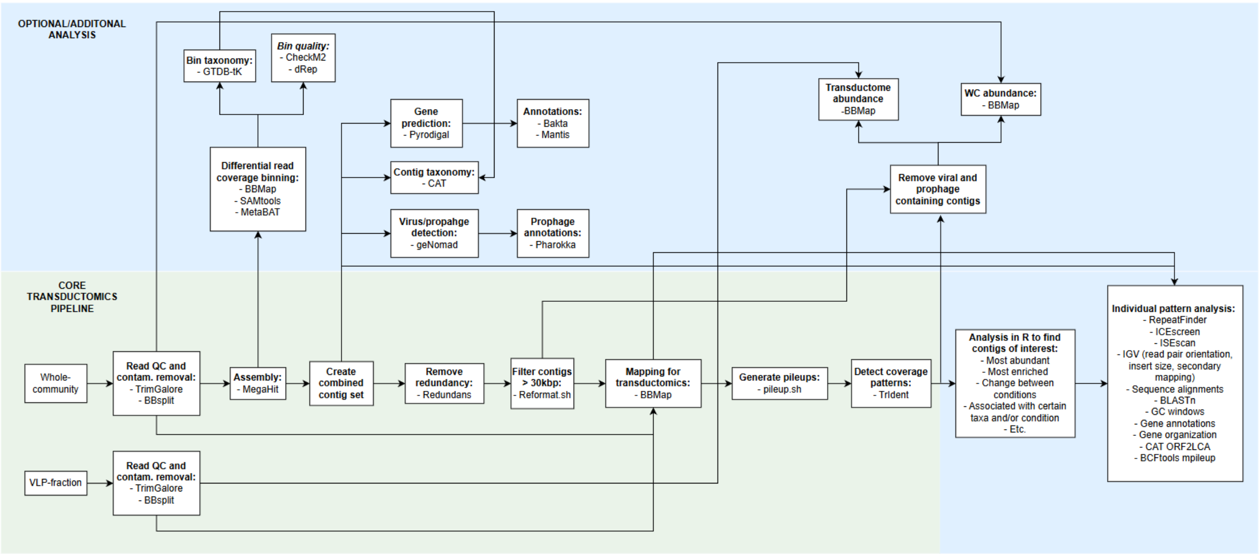


**Supplementary figure 34 - Bioinformatics pipeline used for transductomics analysis**

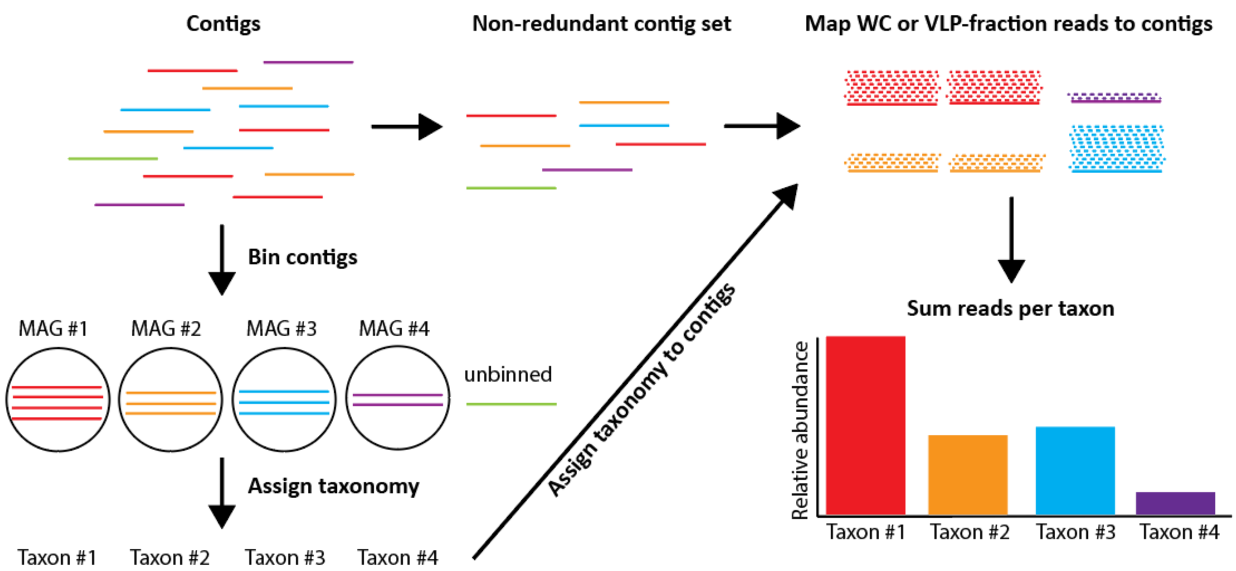


**Supplementary figure 35 - Workflow used to determine taxonomic composition of both the whole-community (WC) and transductome (VLP-fraction).** Unbinned contigs were not included when determining the taxonomic compositions, however they were included when TrIdent was run on the non-redundant contig set. If an unbinned contig was positively classified by TrIdent, its taxonomic classification was obtained with CAT. Relative abundances were calculated by dividing the summed reads by the number of trimmed and decontaminated reads in the respective WC or VLP-fraction sample.

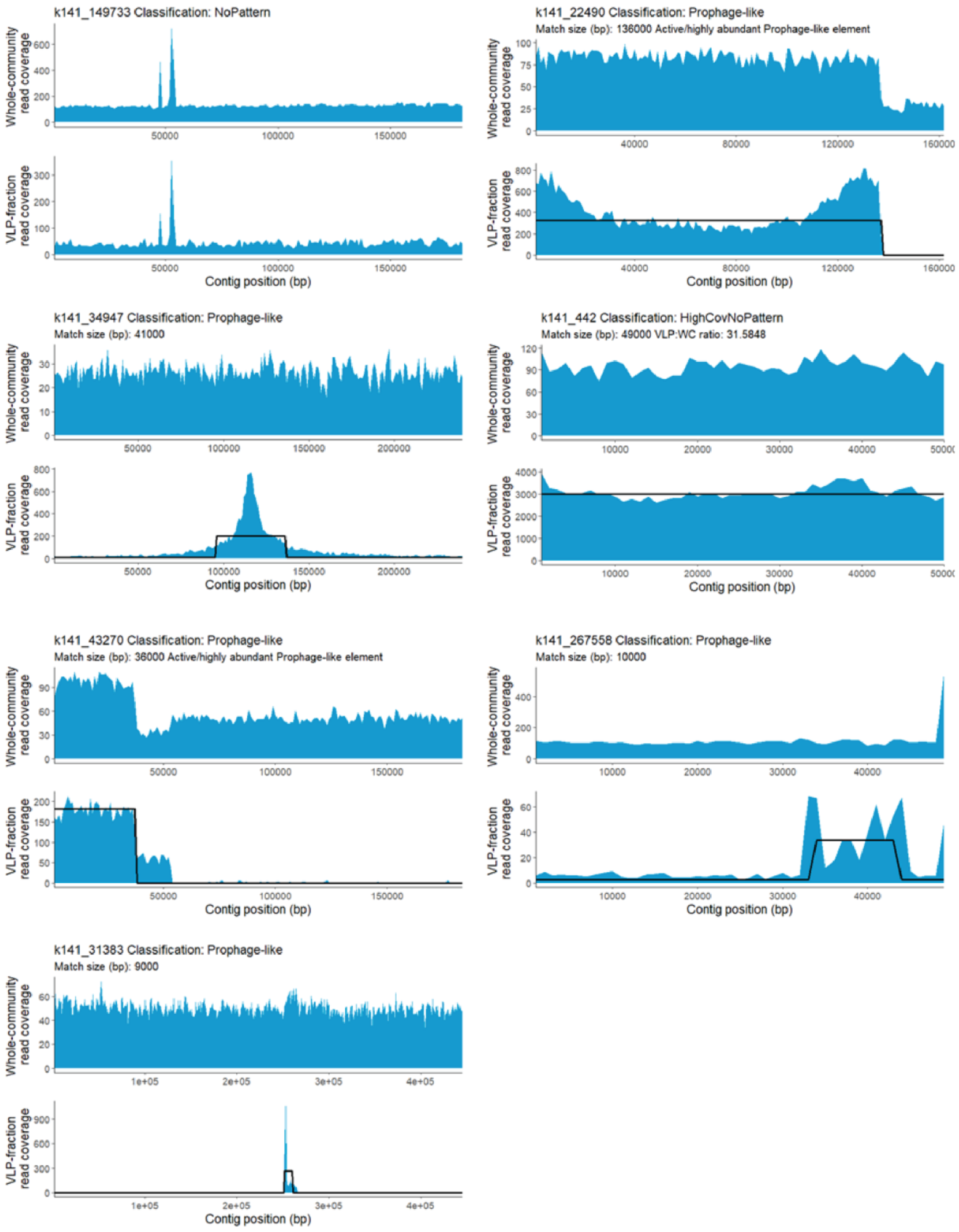


**Supplementary figure 36 - Mapping of reads from individual samples to their respective assemblies with ambiguous=random to show preservation of read coverage patterns analyzed in case study**

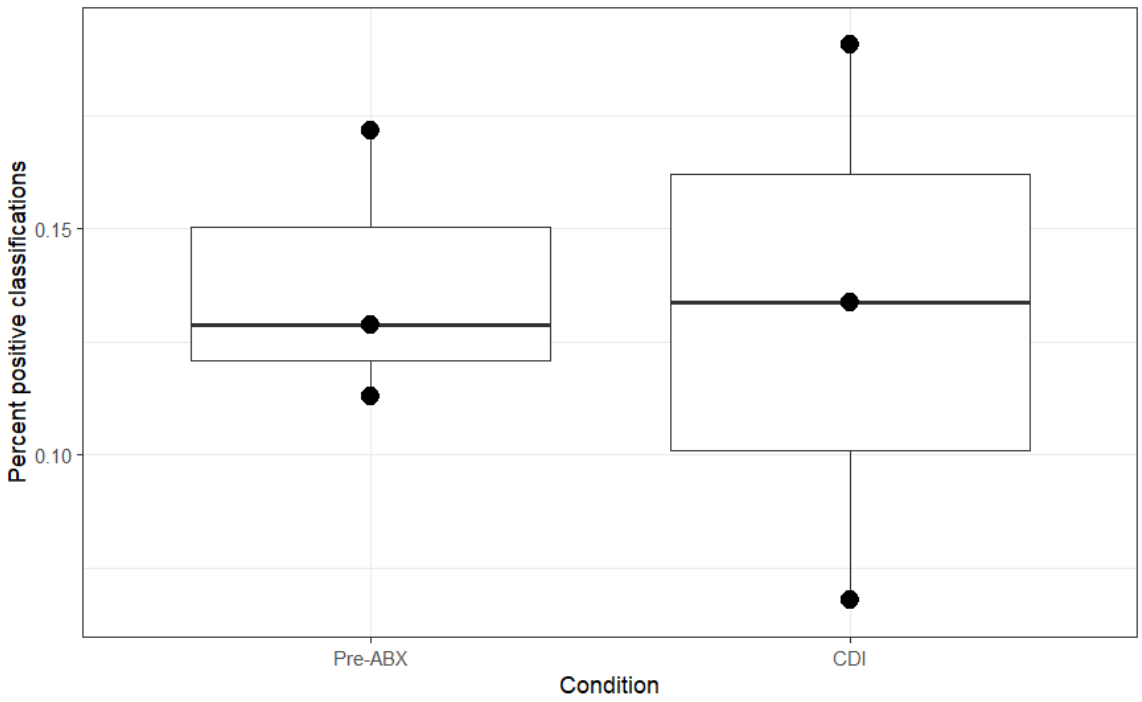


**Supplementary figure 37 - Percent of positive TrIdent classification in each replicate across Pre-ABX and CDI conditions when TrIdent is run on individual WC assemblies.** ABX= antibiotics, CDI=*Clostridioides difficile* infection.
